# A BCG toxic effector induces mitochondrial DNA compaction to refrain protective immunity

**DOI:** 10.1101/2025.11.09.687409

**Authors:** Cheng Peng, Qiu Chen, Jingping Huang, Wenyi Bu, Xuan Wu, Yongjia Duan, Shanshan Liu, Hongyu Cheng, Mingtong Ma, Yuanna Cheng, Jingxiang Wang, Hua Yang, Ruijuan Zheng, Jie Wang, Xiaochen Huang, Lin Wang, Baoxue Ge

**Author notes:** These authors contributed equally to this work (Cheng Peng, Qiu Chen, Jingping Huang). Correspondence: Baoxue Ge; Lin Wang.

## Abstract

Innate immune memory typically emphasizes nuclear chromatin remodeling that determines transcriptional memory in monocytes and macrophages. Here, we show that compaction of mitochondrial DNA (mtDNA), governed by mitochondrial transcription factor A (TFAM) abundance and mtATP-dependent remodeling, defines the transcriptional and functional state of trained macrophages. While typical pathogen-associated molecular pattern (PAMPs) triggered mtDNA accessibility, Bacillus Calmette-Guérin (BCG) effector PE18 directly interacted with and activated SLC25A5 to reduce mtATP levels, thereby inhibiting TFAM degradation by AFG3L2. Accumulated TFAM restricts mtDNA accessibility, suppressing the transcription of the mtNd1 and thereby reducing mtNd1-mediated mtROS production, which diminishes the trained capacity of macrophages against secondary infections. Notably, BCGΔ*pe18* maintains macrophage’s cytokine transcriptional capacity independent of classical nuclear histone modifications. Moreover, BCGΔ*pe18* provides robust and lifelong protective immunity against *Mycobacterium tuberculosis* (*Mtb*) infection. These findings reveal mitochondrial genome as a regulatory entity in trained immunity, and suggest a potential strategy for improving vaccine performance.

## Introduction

Innate immune memory, also known as trained immunity, has fundamentally reshaped the traditional dichotomy between innate and adaptive immunity, whereas traditionally immunological memory was solely attributed to antigen-specific lymphocytes (1–3). In response to prior stimulation or infection, the innate immune system undergoes memory-like epigenetic and metabolic reprogramming (4–7). Notably, *in vivo* epigenetic remodeling and functional tuning of hematopoietic stem cells (HSCs), the source of innate immune cells including macrophages and neutrophils, sustains long-term differentiation potential and endows their progeny innate immune cells with enhanced responsiveness to subsequent challenges (8–10). Beyond enhancing host defenses against recurrent infections, innate immune memory has been implicated in the pathogenesis of diverse inflammation-associated diseases, including cancer (11–13), cardiovascular disorders (14–16), and metabolic disorders (17–19), and neuroinflammatory pathologies (20–23).

Epigenetic reprogramming of innate immune cells is driven by metabolic rewiring, notably a metabolic shift from oxidative phosphorylation (OXPHOS) to glycolysis (4). This shift promotes the accumulation of metabolic intermediates such as lactate or fumarate. Specifically, activation of the AKT-mTOR-HIF-1α axis boosts glucose uptake and lactate production; meanwhile, the accumulation of tricarboxylic acid (TCA) cycle intermediates in mitochondria regulates the activity of epigenetic enzymes and the activation of transcription factors. For instance, Bacillus Calmette-Guérin (BCG) vaccination triggers H3K27 acetylation (H3K27ac) and H3K4 methylation (H3K4me1) at enhancer regions of pro-inflammatory genes such as Interleukin-1β (IL-1β) and tumor necrosis factor-α (TNF-α) (24, 25) in myeloid cells. More recent studies reveal H3K18 lactylation (H3K18la) as a novel epigenetic mark that integrates metabolic and epigenetic signaling. Lactate, produced via glycolysis during immune activation, is catalyzed by p300/CBP to modify histones, thereby stabilizing open chromatin states at immune response genes (26–28). Additionally, DNA demethylation at promoters or enhancers contribute to sustained gene expression, which plays a critical role in trained immunity (29–32). Collectively, these epigenetic modifications drive the transcription of nuclear-encoded inflammatory cytokines, sustaining the trained immunity phenotype (12, 33, 34).

Mitochondrial genome encodes 13 components of the electron transport chain (ETC), which are critical for OXPHOS to maintain mitochondrial energy production and calcium signaling (35–37). Structurally, mtDNA is a double-stranded, circular molecule mainly compacted by transcription factor A (TFAM), a protein with dual regulatory capacity for mtDNA transcription (38) and replication whereas TFAM deficiency causes mtDNA replication stress and cytosolic leakage, which in turn activates antiviral immunity and nuclear DNA repair through activation of cGAS-STING signaling (39–41). Emerging evidence reveals a central role of mitochondria in metabolic reprogramming during innate immune training driven by mitochondrial regulation of TCA cycle intermediates and epigenetic modifiers such as α-ketoglutarate (42), neglecting the potential of mtDNA as a genetic material. Notably, despite the fact that proinflammatory role of mtDNA is well-established (39, 43, 44), whether mtDNA dynamics, including transcriptional regulation, mtDNA accessibility or nucleoid packaging, participate in shaping innate immune memory remains largely unexplored. The century-old live attenuated BCG vaccine remains a cornerstone of tuberculosis prevention yet exhibits critical limitations: inconsistent protection against *Mycobacterium tuberculosis* (*Mtb*), particularly variable efficacy in adolescents and adults (45), and safety concerns in immunocompromised populations (46–48). BCG modification strategies such as knocking out of virulence genes like *ureC* (49, 50) or inserting immune-enhancing genes like Ag85B and ESAT-6 (51) have been shown to enhance vaccine efficacy while preserving BCG advantages. In addition, intravenous (i.v.) delivery of BCG provides protection against *Mtb* in macaques but poses safety challenges (52). A “suicidal BCG” (53) is therefore engineered to harbor conditional suicide systems by overexpression of inducible lysis genes. Although intravenous BCG confers nearly complete protection in non-human primates (NHPs) (52, 53), it only provides partial protective efficacy in mice (54, 55), suggesting that the protective efficacy of BCG may exhibit species-specific differences. Given that mice are widely used for preclinical evaluation of vaccine candidates (53, 56–58), optimizing BCG to enhance its protective potency in this model represents an important research direction. While BCG provides non-specific protection against viral and fungal pathogens through metabolic and epigenetic reprogramming of myeloid cells (24, 59), its failure to establish durable and more effective anti-*Mtb* immunity exposes a fundamental gap in our understanding of the protective and pathogenic mechanisms underlying BCG vaccination.

In this study, we identified that PE18 from BCG (encoded by BCG_1820 gene) induces mtDNA compaction and downregulated mitochondrial ROS (mtROS) production in response to secondary bacterial infection, which restricts the killing capacity of BCG-vaccinated mice derived macrophages against multiple pathogens. Mechanistically, PE18 interacts with and activates mitochondrial adenine nucleotide translocase SLC25A5 to decrease mitochondrial ATP pool and ATP-dependent protease AFG3L2-mediated TFAM degradation, leading to subsequent TFAM accumulation and mtDNA compaction. Moreover, BCGΔ*pe18* significantly upregulates gene expression of mitochondrial encoded NADH dehydrogenase subunit 1 (mtNd1) to facilitate mtROS-mediated bacterial clearance in macrophages and notably retains the transcriptional capacity of pro-inflammatory cytokines in response to secondary stimulation, independent of classical nuclear histone modifications. Lastly, BCGΔ*pe18* vaccination shows more safety and confers superior and longer protective efficacy against *Mtb* infection compared to parental BCG.

## Results

### BCG PE18 induced mtDNA compaction

Macrophages serve as the first line of defense against invading bacterial pathogens (60–62). To examine the effects of innate immune activation on mtDNA compaction, mouse bone marrow derived macrophages (BMDMs) were stimulated with different stimuli, or *Mycobacterium bovis* BCG, and mtDNA compaction was analyzed by a modified mitochondrial assay using transposase-accessible chromatin-see (mtATAC-see) probes (63) (**Figure 1A-C**). The data revealed that typical innate immune memory inducer β-glucan (64) or oxidized Low-density Lipoprotein (oxLDL) (65) markedly inhibited mtDNA compaction. Interestingly, *Mycobacterium bovis* BCG significantly enhanced mtDNA compaction in BMDMs, whereas BCG-derived pathogen-associated molecular patterns (PAMPs), such as arabinogalactan (AG) and peptidoglycan (PGN), conversely reduced compaction in BMDMs (**Figure 1A-C**). In addition, heat-killed or streptomycin-killed BCG failed to induce mtDNA compaction in BMDMs (**Supplemental Figure 1A, B**). Given that *Mycobacteria* employ various secretion systems to enable the efficient translocation of effector proteins through their hydrophobic and exceptionally impermeable cell walls directly into the cytoplasm of host cells to interfere with various cellular functions (66, 67), these results suggest that effector proteins from live BCG may induce mtDNA compaction.

**Figure 1.**
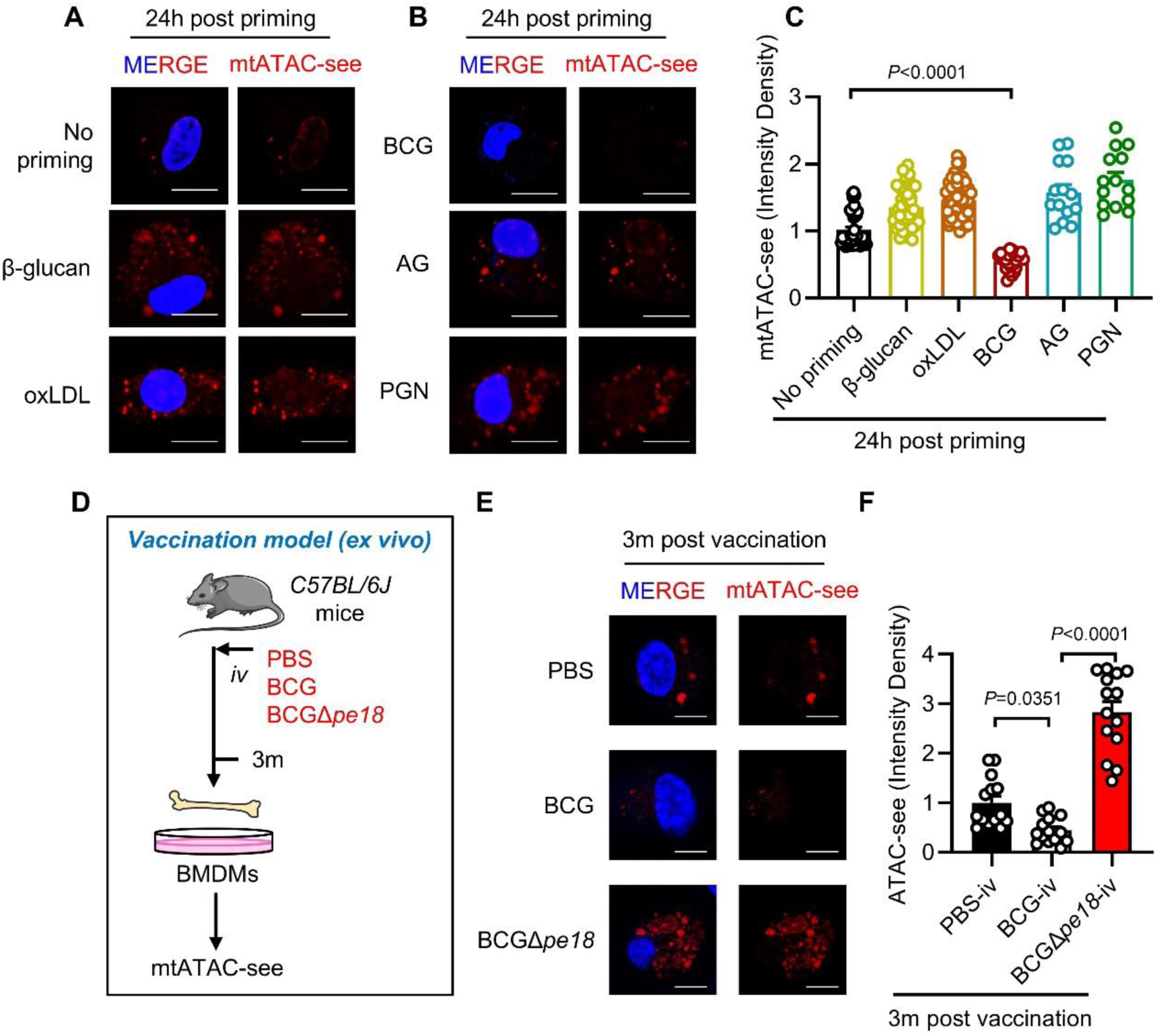
BCG PE18 induces mtDNA inaccessibility. **A-C**. Representative images (**A** and **B**) and quantification (**C**) of the mitochondrial ATAC-see (mtATAC-see) signal in BMDMs primed with β-glucan, oxidized Low-density Lipoprotein (oxLDL), BCG, arabinogalactan (AG) and peptidoglycan (PGN) for 24 h. Scale bars, 10 μm. Each single spot is representative of a single cell. (mean of n = 14 to 53). **D**. Workflow of the mtATAC-see assay in BMDMs isolated from C57BL/6J mice vaccinated with BCG or BCGΔ*pe18* for 3 months (3m) (*ex vivo* model). **E-F**. Representative images (**E**) and quantification (**F**) of the mtATAC-see signal in BMDMs derived from vaccinated mice at 3m post vaccination. Scale bars, 10 μm. Each single spot is representative of a single cell. n=14. *P* values were calculated via One-way ANOVA tests (**C, F**).

To identify BCG effectors that induce mtDNA compaction, we examined the effects of 161 mycobacterial secretory proteins on mtDNA compaction by an mtATAC-see probe in HEK293T cells (**Supplemental Figure 1C, D; Supplementary Table 1**). Among these mycobacterial proteins, PE18 (also named BCG_1820) was found to be the secretory protein that blocks the mtATAC-see signal most significantly (**Supplemental Figure 1C, D**), suggesting PE18 acts as an inducer of mtDNA compaction. Consistently, BMDMs isolated from the mice intravenously vaccinated with BCG showed more enhanced mtDNA compaction than those derived from PBS control (PBS-iv) mice (**Figure 1D-F**). Notably, deletion of *pe18* from BCG almost completely abolished the enhancement of mtDNA compaction in BMDMs isolated from BCG-iv mice (*ex vivo*) (**Figure 1E, F**). These data suggest that BCG PE18 may induce mtDNA compaction during BCG vaccination.

### PE18 interacts with SLC25A5 to induce mtDNA compaction

Given that PE18 is a secretory protein (**Supplemental Figure 2A**), we hypothesized that a host factor is likely involved in its role related to regulating mtDNA compaction. By performing coimmunoprecipitation (co-IP) combined with mass spectrometry in HEK293T cells overexpressing green fluorescent protein (GFP)-tagged PE18, we aimed to identify a PE18-binding protein in mammalian cells (**Supplemental Figure 2B**). Seventeen of the top 22 hits were mitochondrion-localized proteins (**Supplemental Figure 2C; Supplementary Table 2**). Through immunoprecipitation and immunoblotting assays, we confirmed that SLC25A5, a mitochondrial ADP/ATP translocase (68, 69), is a PE18-binding protein (**Supplemental Figure 2D**). Mitochondrial fraction and immunofluorescence further validated that the interaction of PE18 and SLC25A5 occurs within mitochondria (**Figure 2A, Supplemental Figure 2E**).

**Figure 2.**
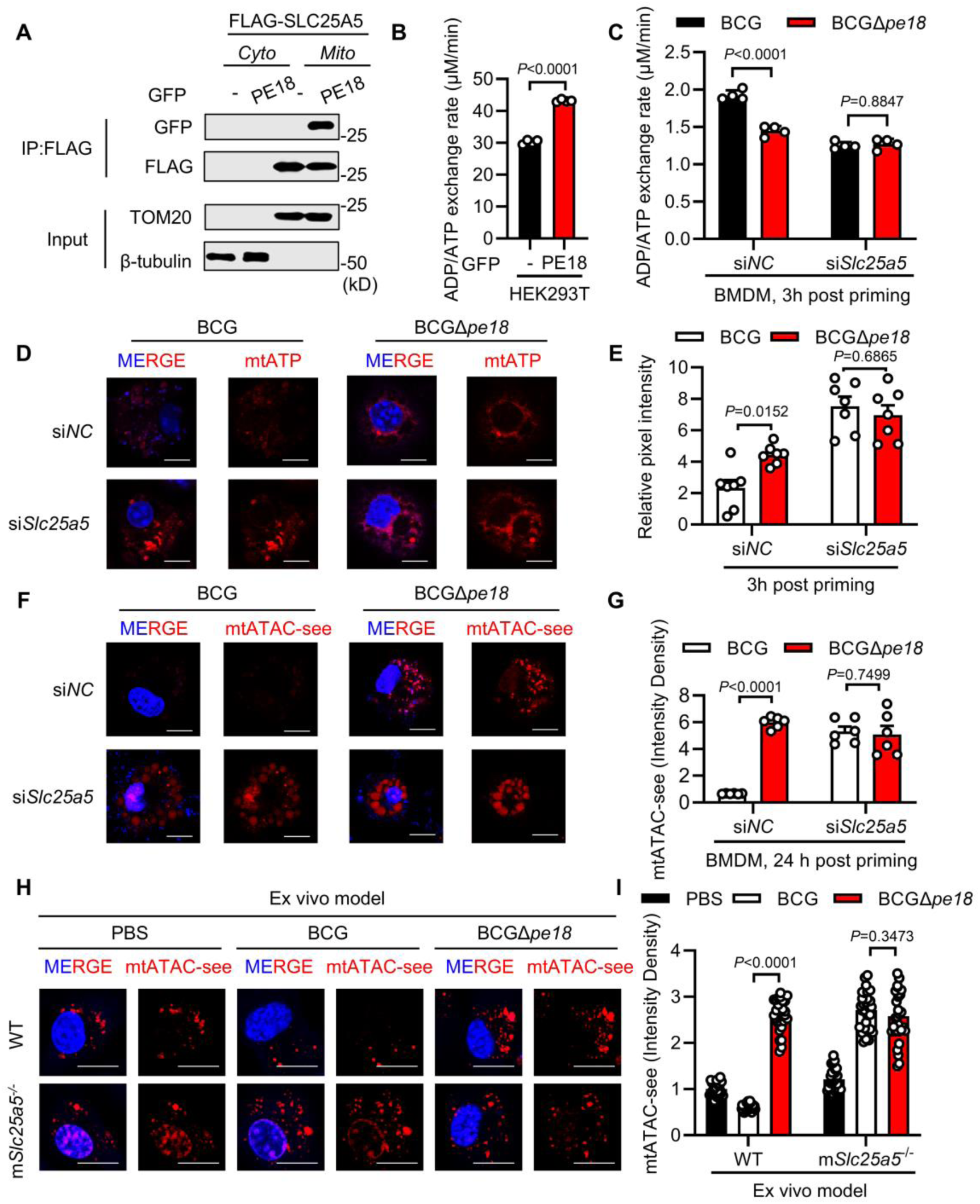
PE18 binds with SLC25A5 to decrease the mitochondrial ATP pool. **A**. Coimmunoprecipitation and immunoblot analysis of cytosolic (*Cyto*) or mitochondrial (*Mito*) fractions isolated from HEK293T cells transfected with FLAG-SLC25A5 and GFP-PE18. The GFP vector (−) was used as a control. TOMM20 and β-tubulin were used as reference proteins for lysates of either mitochondria or the cytosol. **B-C**. Mitochondrial ADP/ATP exchange assay in HEK293T cells (**B**) transfected with GFP-PE18; and BMDMs (**C**) transfected with negative control siRNA (siNC) or siRNA targeting mouse *Slc25a5* (si*Slc25a5*) for 48 h followed by priming with BCG or BCGΔ*pe18* for 3 h. **D-E**. Representative images (**D**) and quantification (**E**) of the mitochondrial ATP signal (mtATP) in BMDMs transfected with si*Slc25a5* followed by priming with BCG or BCGΔ*pe18* for 3 h. Scale bars, 10 μm. Each single spot is representative of a single cell. (mean ± s.e.m. of n = 7). **F-G**. Representative images (**F**) and quantification (**G**) of the mtATAC-see signal in BMDMs transfected with si*Slc25a5* followed by priming with BCG or BCGΔ*pe18* for 24 h. Scale bars, 10 μm. Each single spot is representative of a single cell. (mean ± s.e.m. of n = 6). **H-I**. Representative images (**H**) and quantification (**I**) of the mtATAC-see signal in BMDMs isolated from wild-type (WT) mice and macrophage-conditioned *Slc25a5* knockout mice (m*Slc25a5*^-/-^) at 1 month after intravenous vaccination with BCG or BCGΔ*pe18*. Scale bars, 10 μm. Each single spot is representative of a single cell. (mean ± s.e.m. of n = 30). *P* values were calculated via two-tailed unpaired Student’s *t* tests (**B**) or Two-way ANOVA tests (**C, E, G, I**).

SLC25A5 belongs to the adenine nucleotide transporter (ANT) family and transports ATP across the inner mitochondrial membrane into the cytosol via exchange of equal amounts of ADP into the mitochondria (68, 69). We next investigated the effect of PE18 on mitochondrial ADP/ATP exchange rates (68). Overexpression of PE18 significantly increased the ADP-triggered ATP release from mitochondria in HEK293T cells (**Figure 2B**). Similarly, BMDMs primed with BCGΔ*pe18* exhibited much lower mitochondrial ADP/ATP exchange efficiency than those primed with BCG (**Figure 2C**). Knockdown of *Slc25a5* using a specific siRNA decreased the mitochondrial ADP/ATP exchange rates and eliminated the differential ADP/ATP exchange rates between BMDMs primed with BCG or BCGΔ*pe18* (**Figure 2C**), suggesting that PE18 may promote mitochondrial ADP/ATP exchange by targeting SLC25A5. As determined by the mitochondrial ATP (mtATP) probe (70), the mtATP level in wild-type BMDMs primed with BCGΔ*pe18* was much higher than that in those primed with parental BCG, which was not observed in *Slc25a5*-knockdown BMDMs (**Figure 2D, E**). These results suggest that PE18 may decrease mtATP levels by promoting SLC25A5-mediated mitochondrial ADP/ATP exchange.

Because PE18 induces mtDNA compaction and promotes SLC25A5-mediated mitochondrial ADP/ATP exchange, we next examined whether PE18 induces mtDNA compaction via its binding target SLC25A5 in BMDMs. As shown in **Figure 2F, G**, silencing of *Slc25a5* dramatically increased the mtATAC-see signal, suggesting that SLC25A5 might regulate mtDNA compaction during BCG priming. Furthermore, compared with wild-type BCG, BCGΔ*pe18* only reduced mitochondrial DNA compaction in control BMDMs but not in *Slc25a5*-knockdown BMDMs (**Figure 2F,G**). Furthermore, we crossed *Slc25a5*^flox/flox^ (*Slc25a5*^fl/fl^) mice with LysM-Cre transgenic mice to generate *Slc25a5*^flox/flox^; LysM-Cre (m*Slc25a5*^-/-^) mice in which the *Slc25a5* gene is deleted in myeloid cells. BMDMs derived from BCG-vaccinated m*Slc25a5*^-/-^ mice presented highly reduced mtDNA compaction compared with those derived from wild-type mice (**Figure 2H, I**). Compared with parental BCG, BCGΔ*pe18* vaccination did not reduce mtDNA compaction in m*Slc25a5*^-/-^ mice as it did in wild-type mice (**Figure 2H, I**). Together, these results suggest that PE18 may induce mtDNA compaction through SLC25A5.

### PE18 suppresses mtDNA accessibility through accumulating TFAM

Mitochondrial transcription factor A (TFAM) compacts mtDNA into nucleoids (71). The TFAM:mtDNA ratio is the main determining factor for mtDNA accessibility, and a high TFAM:mtDNA ratio leads to mitochondrial DNA inaccessibility (72). The TFAM:mtDNA ratio was significantly increased in BMDMs primed with BCG but markedly decreased in BMDMs primed with BCGΔ*pe18* (**Figure 3A, B**). Immunoblotting analysis confirmed that the TFAM protein level was significantly increased in BMDMs primed with BCG but not in those primed with BCGΔ*pe18* (**Figure 3C**). Moreover, knockdown of *Tfam* abolished the mtDNA accessibility difference between BCG– and BCGΔ*pe18-*primed BMDMs (**Figure 3D-F**), indicating that BCG PE18 may suppress mtDNA accessibility by promoting TFAM protein accumulation.

**Figure 3.**
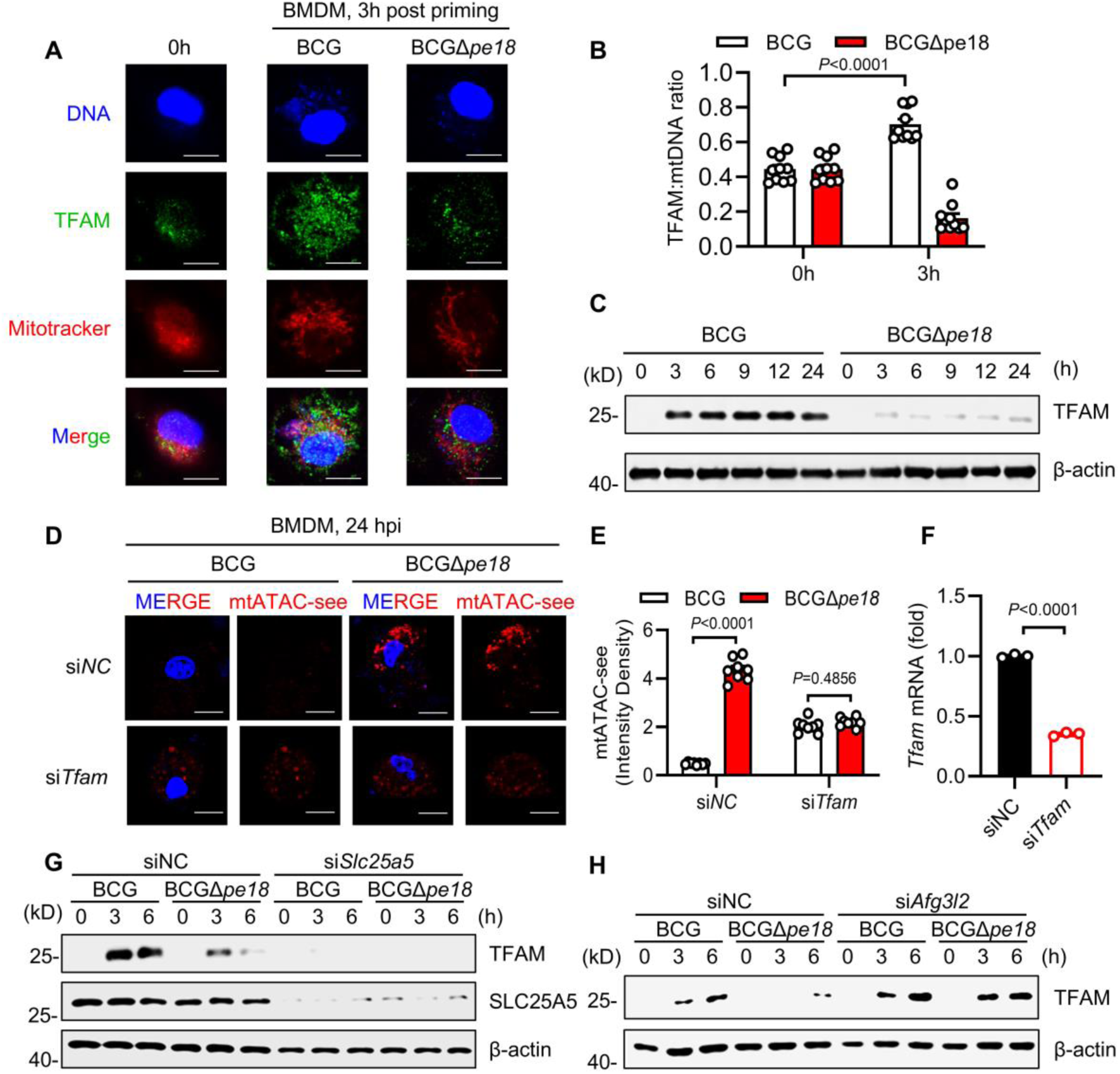
PE18 leads to TFAM accumulation-mediated mtDNA inaccessibility. **A-B**. Representative images (**A**) and quantification (**B**) of the TFAM:mitochondrial DNA (mtDNA) ratio of BMDMs stimulated with BCG or BCGΔ*pe18* for 3 h. (mean ± s.e.m. of n = 10). **C**. Immunoblot analysis of BMDMs stimulated with BCG or BCGΔ*pe18* for the indicated times. **D-E**. Representative images (**D**) and quantification (**E**) of the mtATAC-see signal in BMDMs transfected with si*Tfam* for 48 h followed by stimulation with BCG or BCGΔ*pe18* for 24 h. Scale bars, 10 μm. Each single spot is representative of a single cell. (mean ± s.e.m. of n = 8). **F**. Q-PCR analysis of *Tfam* mRNA expression in BMDMs transfected with si*Tfam* for 48 h. (mean ± s.e.m. of n=3). Representative **G**. Immunoblot analysis of BMDMs transfected with si*Slc25a5* and stimulated with BCG or BCGΔ*pe18* for 24 h. **H**. Immunoblot analysis of BMDMs transfected with si*Afg3l2* and stimulated with BCG or BCGΔ*pe18* for 24 h. *P* values were calculated via Two-way ANOVA tests (**B, E**) or two-tailed unpaired Student’s *t* tests (**F**).

We next investigated the underlying mechanism by which PE18 promotes TFAM accumulation. RT-PCR analysis revealed that deletion of *pe18* did not significantly change the *Tfam* mRNA levels in BMDMs (**Supplemental Figure 3A**). In addition, treatment of BMDMs with the lysosome inhibitor chloroquine (CQ) or the proteasome inhibitor MG132 had no significant effect on the PE18-mediated accumulation of the TFAM protein (**Supplemental Figure 3B, C**), suggesting that the lysosomal or proteasomal degradation pathways may not be involved in PE18-mediated TFAM protein accumulation. Notably, silencing of *Slc25a5* reduced TFAM protein level and eliminated the PE18-mediated accumulation of the TFAM protein (**Figure 3G**), linking mtATP depletion to TFAM accumulation (**Figure 2D, E**). Mitochondrial proteins are primarily degraded by a series of ATP-dependent proteases (73). By screening key mitochondrial proteases, including LONP1 (74, 75), AFG3L2 (76, 77), YME1L (78), Clpp and Clpx (79), we found that only *Afg3l2* knockdown markedly abolished the decreased TFAM protein level in BCGΔ*pe18-*infected BMDMs (**Figure 3H; Supplemental Figure 3D-G**). These results suggest that BCG PE18 may deplete mtATP via SLC25A5, inhibiting ATP-dependent degradation of TFAM by AFG3L2, thus leading to TFAM accumulation and mtDNA inaccessibility.

### mtDNA accessibility regulates innate immune memory

BCG promotes bone marrow monocyte/macrophage lineage nuclear DNA accessibility to increase the production of cytokines in response to secondary *Mtb* infection. However, the role of mtDNA accessibility in BCG-mediated innate immune memory remains unknown. To determine the direct impact of PE18-induced mtDNA inaccessibility on macrophages bacterial killing, we conducted an intracellular *Mtb* survival assay in macrophages primed with BCG or BCGΔ*pe18*. Deletion of *pe18* in BCG significantly enhanced *Mtb* clearance in BMDMs during secondary infection (**Figure 4A, B**), which was abolished by treatment with 2’,3’-dideoxycytidine (ddC), a nucleoside analog that specifically depletes mtDNA activity (80) (**Figure 4A, B**). To confirm this *ex vivo*, we isolated BMDMs from mice intravenously vaccinated with BCG or BCGΔ*pe18* for 3 months and assessed their intracellular *Mtb* killing activity at 48 h postinfection (**Figure 4C**). Consistently, BMDMs from BCGΔ*pe18*-*iv* mice exhibited much greater killing capacity against intracellular *Mtb* than those from BCG-iv mice (**Figure 4D**). Notably, ddC treatment did not significantly alter the differential *Mtb* killing observed between PBS-control and BCG-trained BMDMs but strikingly ablated the *Mtb* killing difference between BCG– and BCGΔ*pe18*-trained BMDMs. This effect was consistent across both *in vitro* and *ex vivo* models (**Figure 4A, B**). These results suggest that *pe18* deletion may increase BCG-induced innate immune memory through restoring mtDNA accessibility.

**Figure 4.**
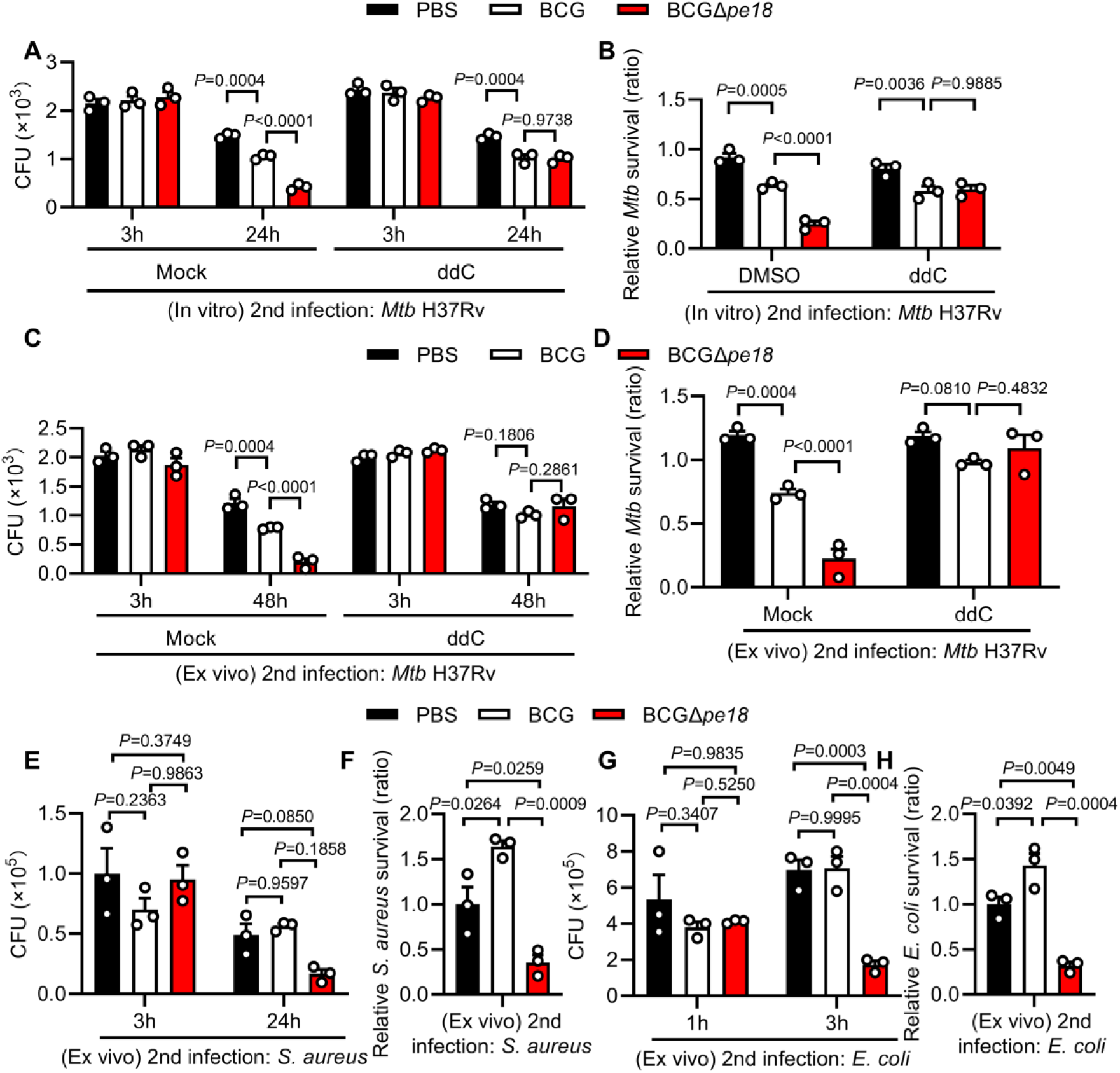
mtDNA inaccessibility impairs innate immune memory. **A-B**. CFU counts (**A**) and relative intracellular CFU ratios (**B**) of secondary *Mtb* H37Rv infection in BMDMs primed with PBS control, BCG or BCGΔ*pe18* for 5 days followed by pretreatment with ddC and infection with *Mtb* H37Rv (MOI = 2) at 3 or 24 h. (mean ± s.e.m. of n = 3). **C-D**. CFU counts (**C**) and relative intracellular CFU ratios (**D**) of secondary *Mtb* H37Rv infection after ddC pretreatment in BMDMs isolated from unvaccinated mice (PBS) or mice intravenously vaccinated with BCG or BCGΔ*pe18* for 3 months at 3 or 48 h post-secondary infection (MOI = 2). (mean ± s.e.m. of n = 6). **E‒F**. CFU counts (**E**) and relative intracellular CFU ratios (**F**) of secondary *S. aureus* infection in BMDMs isolated from mice vaccinated with BCG or BCGΔ*pe18* for 3 months at 3 or 24 h post-secondary infection (MOI = 10). (mean ± s.e.m. of n = 3). **G‒H**. CFU counts (**G**) and relative intracellular CFU ratios (**H**) of secondary *E. coli* infection in BMDMs isolated from mice vaccinated with BCG or BCGΔ*pe18* for 3 months at 1 or 3 h post-secondary infection (MOI = 10). (mean ± s.e.m. of n = 3). *P* values were calculated via One-way or Two-way ANOVA tests (**A-H**).

BCG-mediated innate immune memory provides nonspecific protection against *Staphylococcus aureus* via increased cytokines production (81, 82), but its impact on intracellular bacterial killing of unrelated pathogens such as *S. aureus* or *Escherichia coli* in the myeloid cells from BCG-vaccinated mice has not been investigated. In our *ex vivo* model, BMDMs from BCG*-iv* mice and PBS*-iv* mice showed similar capacity to clear intracellular *S. aureus* or *E. coli* (**Figure 4E-H**), but deletion of *pe18* markedly increased BCG-induced killing activity of these two unrelated pathogens in BMDMs (**Figure 4E-H**). These results suggest that PE18 may suppress the functional state of trained macrophages.

### PE18 inhibits mtNd1-mediated mtROS production

mtDNA accessibility is usually associated with the transcription of mtDNA-encoding genes (63, 83). Transcriptomic analysis revealed that, compared with BCGΔ*pe18*-primed BMDMs, BCG-primed BMDMs presented lower mitochondrial DNA accessibility and extensively suppressed the expression of nearly all the mitochondrial DNA-encoding genes in BMDMs (**Supplemental Figure 4A; Supplementary Table 3**). These genes are associated with multiple mitochondrial functions, such as those related to the electron transport chain (ETC) (84), mitochondrial fusion/fission (85), and the generation of mtROS, the main byproduct of respiratory complex I, which is critical for intracellular bacterial clearance (86, 87).

To investigate key mechanisms involved in BCGΔ*pe18*-mediated killing advantage, we treated BCG– or BCGΔ*pe18-*primed BMDMs with rotenone (88) (an inhibitor of the electron respiratory chain), Mdivi-1 (89) (an inhibitor of mitochondrial fusion/fission) or mitoTEMPO (90) (an inhibitor of mtROS) prior to secondary *Mtb* infection. Only rotenone or mitoTEMPO eliminated the differential survival of *Mtb* in BCG– and BCGΔ*pe18-*primed BMDMs (**Figure 5A, B; Supplemental Figure 4B, C**). These results suggest that mitochondrial respiratory complex I or mtROS may contribute to the PE18-enhanced survival of secondary *Mtb* infection.

**Figure 5.**
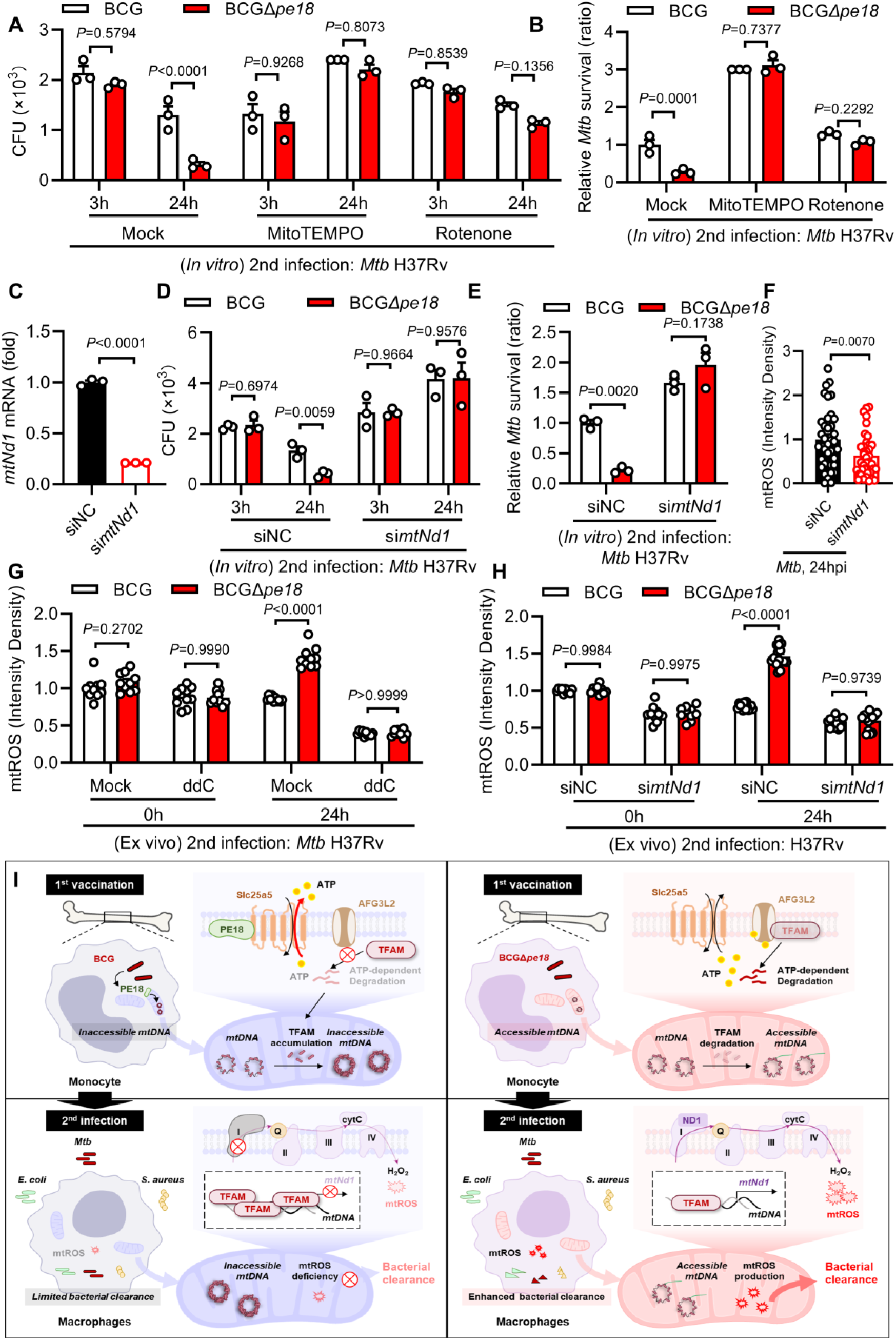
PE18 inhibits mtNd1-mediated mtROS production. **A-B**. CFU counts (**A**) and relative intracellular CFU ratios (**B**) of secondary *Mtb* H37Rv infection in BMDMs isolated from unvaccinated mice and primed with BCG or BCGΔ*pe18* for 5 days followed by pretreatment with MitoTEMPO and Rotenone before infection with *Mtb* H37Rv at 3 or 24 h after secondary infection (MOI = 2) (mean ± s.e.m. of n = 3). **C**. Q-PCR analysis of *mtNd1* mRNA expression in BMDMs transfected with si*mtNd1* for 48 h. (mean ± s.e.m. of n=3). **D-E**. CFU counts (**D**) and relative intracellular CFU ratios (**E**) of secondary *Mtb* H37Rv infection in in vitro-isolated BMDMs primed with BCG or BCGΔ*pe18* for 5 days followed by transfection with si*mtNd1* for 48 h and infection with *Mtb* H37Rv (MOI = 2) at 3 or 24 h after secondary infection (mean ± s.e.m. of n = 3). **F**. Quantification of mitochondrial ROS (mtROS) fluorescence intensity in BMDMs transfected with si*mtNd1* followed by *Mtb* infection for 24 h. (mean ± s.e.m. of n = 38). **G-H**. Quantification of the mtROS signal in BMDMs isolated from mice vaccinated with BCGΔ*pe18* for 1 month followed by pretreatment with ddC (**F**) or transfection with si*mtNd1* (**G**) and *Mtb* infection (MOI = 2) for 0 or 24 h (mean ± s.e.m. of n = 10). Quantification of mtROS signals in BMDMs isolated from mice. **I**. Diagram: BCG PE18 interacts with SLC25A5 to decrease the mitochondrial ATP pool, thus inhibiting ATP-dependent degradation of TFAM by the protease AFG3L2, leading to TFAM accumulation and blocking mtDNA accessibility to downregulate mt*Nd1* expression as well as downstream mtROS production during secondary infection, thereby impairing innate immune memory. Quantification of mtROS signals in BMDMs isolated from mice. *P* values were calculated via Two-way ANOVA tests (**A-B, D-H**) or two-tailed unpaired Student’s *t* tests (**C**).

mtNd1 is a crucial component of mitochondrial respiratory complex I encoded by mtDNA (91, 92). The silencing of *mtNd1* markedly increased the intracellular survival of *Mtb* H37Rv in BMDMs (**Figure 5C-E**) and abolished the killing advantage of BCGΔ*pe18*-primed BMDMs (**Figure 5C-E**). By using an mtROS probe, we observed that *mtNd1* knockdown markedly decreased mtROS production in *Mtb*-infected BMDMs (**Figure 5F**). *Ex vivo* model reveals that BCGΔ*pe18* significantly increased mtROS levels during secondary infection with *Mtb* H37Rv. Furthermore, depletion of mtDNA by ddC treatment or blockade of mitochondrial respiratory complex I via *mtNd1* knockdown eliminated the mtROS increase in BCGΔ*pe18*-primed BMDMs during secondary infection with *Mtb* H37Rv (**Figure 5G, H**). *mtNd1* knockdown also significantly increased the intracellular survival of *S. aureus* and *E. coli* in BMDMs, whereas inhibition of mtROS eliminated the differential effects of si*mtNd1* and siNC on the clearance of *S. aureus* and *E. coli* (**Supplemental Figure 4D, E**). These data suggest that suppression of mtDNA accessibility by PE18 may decrease the expression of the mtDNA-encoded gene *mtNd1* to reduce mtROS production, thus impairing the ability of macrophages to kill bacteria intracellularly (**Figure 5I**).

### BCGΔ*pe18* maintains cytokines transcription

Cytokine production following heterogeneous secondary stimulation is the hallmark functional readout for trained immunity (93–95). To determine the role of PE18 in regulating cytokine responses during trained immunity, we examined *Il1b*, *Il6*, and *Tnf* expression and production in BCG-primed BMDMs following secondary LPS stimulation or *Mtb* infection. Unexpectedly, at 6 hours post LPS stimulation, macrophages primed with BCG or BCGΔ*pe18* showed comparable expression and production of *Il1b, Il6* and *Tnf* (**Supplemental Figure 5A, B**). In contrast, upon secondary *Mtb* infection, deletion of *pe18* increased both mRNA and protein level of these cytokines in BCG-primed BMDMs (**Supplemental Figure 5C, D**).

Metabolic rewiring and histone modifications usually modify chromatin accessibility and lead to enhanced transcription of innate immune genes (26, 96). Trained immunity is canonically linked to metabolic reprogramming driven by AKT-mTOR-HIF-1α signaling (26, 97, 98), which is characterized by heightened aerobic glycolysis and the conversion of pyruvate to lactate (5). Notably, a recent breakthrough study revealed that BCG vaccination-induced glycolysis-mediated lactate production enhances IL-1β expression *in vivo* via long-term lactylation of histone H3 at lysine residue 18 (H3K18la) predominantly at distal regulatory regions (26). To explore whether BCGΔ*pe18* alters metabolic profiles, we performed metabolomics in BCG– or BCGΔ*pe18*-primed BMDMs. As previously reported, BCG priming increased lactate production compared to the PBS control; however, this effect was significantly attenuated in BCGΔ*pe18*-primed macrophages, which displayed notably lower intracellular lactate levels (**Supplemental Figure 5E, F; Supplementary Table 4**). Macrophages primed with BCGΔ*pe18* showed a significant decrease in H3K18la levels compared to those primed with wild-type BCG (**Supplemental Figure 5G**), which is consistent with the role of lactate in mediating H3K18la (26). BCG vaccination upregulates H3K18la at the distal regions of the *Il1b* promoter and enhancer (as indicated by H3K27ac and H3K4me3 peaks) in BMDMs (26). In contrast, these specific H3K18la enrichment peaks were significantly reduced in BMDMs isolated from BCGΔ*pe18*-vaccinated mice (**Supplemental Figure 6A**), suggesting that classical H3K18la may not be involved in BCGΔ*pe18*-mediated maintenance of cytokine transcriptional capacity.

In addition, fumarate is a key metabolite that induces chromatin-based trained immunity by promoting histone H3K4me3 and H3K27ac modification, which upregulate the accessibility and transcription of *Il6* and *Tnf* (99). Compared with BCG-primed BMDMs, a significant reduction of fumarate and subsequent histone H3K27ac/H3K4me3 was detected in BCGΔ*pe18*-primed BMDMs (**Supplemental Figure 5E-G; Supplementary Table 4**). The enrichment of histone H3K27ac and H3K4me3 at the promoter regions of the *Il6* and *Tnf* loci was reduced in BMDMs isolated from BCGΔ*pe18*-iv mice relative to BCG-iv mice (**Supplemental Figure 6B, C**), suggesting that classical H3K4me3 and H3K27ac may not be involved in BCGΔ*pe18*-mediated maintenance of cytokine transcriptional capacity.

### Deletion of *pe18* from BCG enhances protection against *Mtb*

We next examined the protection efficiency of BCGΔ*pe18* against *Mtb* infection. C57BL/6J mice were intravenously vaccinated for one month and subjected to H37Rv infection for another month (**Figure 6A**). While intravenous BCG confers nearly complete protection against *Mtb* infection in rhesus macaques (52), wild-type BCG-iv only provided partial protection in mice (54, 55, 100). Consistently, nearly ∼10-fold reduction of lung bacterial loads was observed in the lung tissues from BCG-vaccinated mice compared to PBS-treated counterparts. However, bacterial loads were notably lower in BCGΔ*pe18*-vaccinated mice than in parental BCG-iv mice after vaccination for one month (reduced >36.30-fold; ≈1.56-fold (log_10_)) or three months (reduced >9.06 × 10^3^-fold; ∼3.96-fold (log_10_)) (**Figure 6B**). Accordingly, mice immunized with BCGΔ*pe18* for one month presented dramatically reduced histopathological changes in their lung tissues, including much less immune cell infiltration and fewer inflammatory lesions, in response to *Mtb* infection (**Figure 6C**). To clarify the duration of protection provided by BCGΔ*pe18*, we extended the immunization time of the mice to 24 months, which is equivalent to ∼96 years of human age, before H37Rv aerosol infection. Surprisingly, BCGΔ*pe18* profoundly altered the protective outcome of *Mtb*-challenged mice, in which nine out of ten mice vaccinated with BCGΔ*pe18* were sterilized with *Mtb*, indicating a lifespan protection effect induced by BCGΔ*pe18*, which showed an absolute advantage over BCG in aged mice (**Figure 6D**). These results suggest that mtDNA accessibility-mediated protection may offer a paradigm-shifting strategy for enhancing TB vaccine durability through mitochondrial nucleoid manipulation.

**Figure 6.**
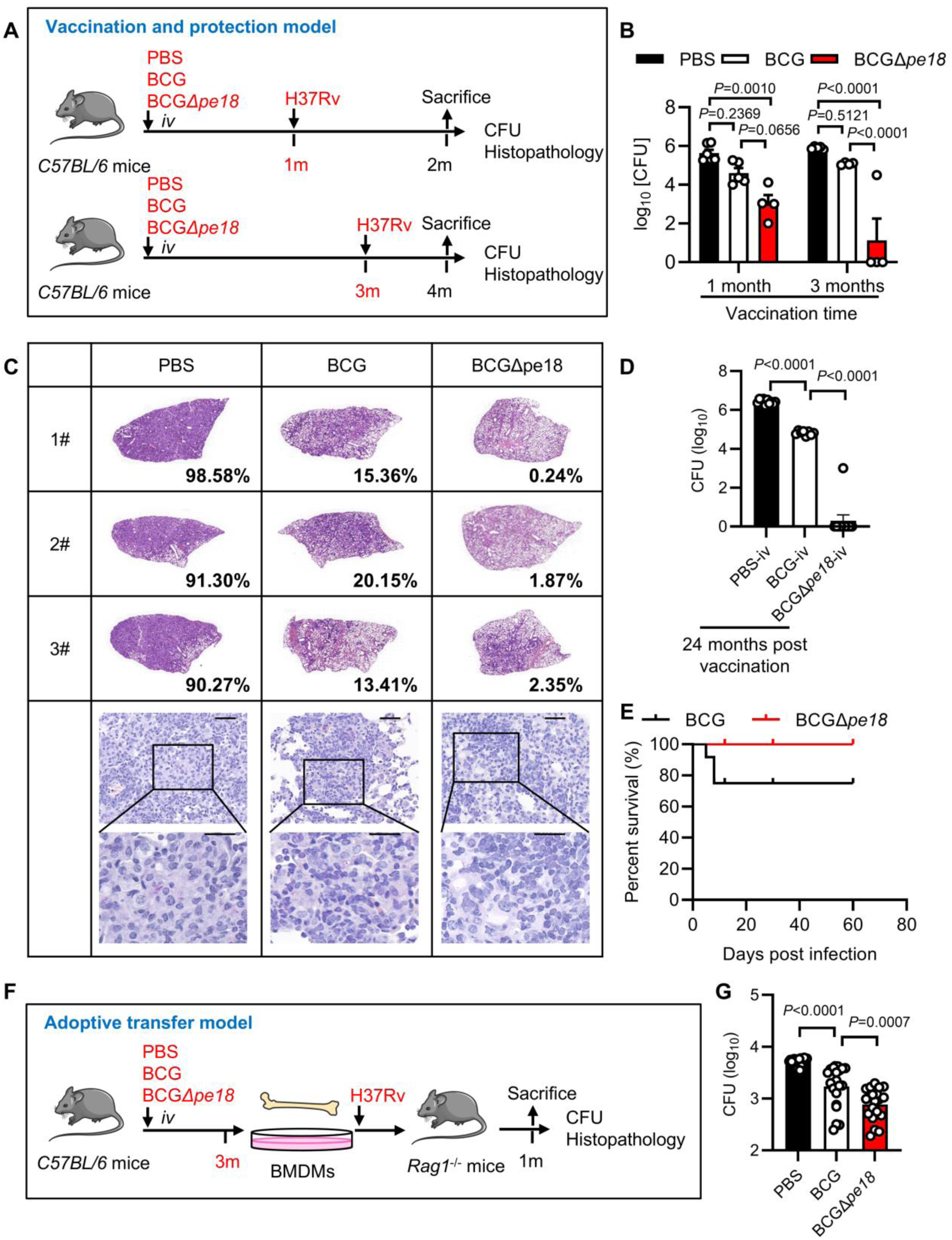
Deletion of PE18 from BCG enhances protection against tuberculosis. **A-C**. Workflow (**A**), CFU counts (**B**), histopathological assay (**C**) via acid-fast staining and hematoxylin and eosin (H & E) staining and histological scoring of the lung tissues of C57BL/6J mice intravenously vaccinated with PBS, BCG or BCGΔ*pe18* for 1 or 3 months followed by aerosol infection with *Mtb* H37Rv for 1 month and assessment of the bacterial burden in the lungs and histopathology. Scale bars, 100 μm (top; original magnification, 400×) and 20 μm (bottom; original magnification, 100×). **D**. Pulmonary CFU counts of mice vaccinated with BCG or BCGΔ*pe18* for 24 months followed by aerosol infection with *Mtb* H37Rv for 1 month. (mean ± s.e.m. of n=8 for the BCG-iv group and n = 10 for the PBS-iv or BCGΔ*pe18*-iv group). **E**. Survival curve of SCID mice vaccinated with BCG or BCGΔ*pe18* (mean ± s.e.m. of n=12). **F-G**. Workflow (**F**) and CFU counts (**G**) in the lung tissues of *Rag1*^-/-^ mice intratracheally transplanted with BMDMs derived from mice intravenously vaccinated with PBS, BCG or BCGΔ*pe18* for 3 months and infected with *Mtb* H37Rv for 1 month. #1, #2 and #3 in (**C**) represent lung tissues from 3 mice vaccinated for 1 month. *P* values were calculated via One-way or Two-way ANOVA test (**B, D, G**).

The administration of the BCG vaccine carries the potential risk of causing disseminated BCGosis, which can lead to life-threatening complications in immunocompromised individuals (101). This includes children with HIV (102) and those suffering from genetic immunodeficiency (103). We also evaluated vaccine safety in severe combined immunodeficiency (SCID) mice, which are severely deficient in functional B and T lymphocytes but retain elements of the innate immune system (104). Three of the ten BCG*-iv* mice died during our observation period, and no mice died in BCGΔ*pe18* vaccination group (**Figure 6E**). These results suggest PE18 as a toxic factor of live attenuated BCG, and that deletion of *pe18* from BCG may provide an excellent safety, accompanied by an outstanding protection effect and duration against TB.

To determine whether BCGΔ*pe18* enhances anti-TB immunity via innate immune memory, BMDMs were generated from mice after 3 months of BCG-iv or BCGΔ*pe18*-iv vaccination and infected with *Mtb in vitro* for 30 min, followed by adoptive transfer (intratracheally) into adaptive immunity-deficient *Rag1*^-/-^ mice (**Figure 6F**). After 30 days of adoptive transfer, the *Mtb* bacterial burden in the lungs of *Rag1*^-/-^ mice that received BMDMs derived from BCG-iv mice was significantly lower than that in the lungs of PBS-iv mice (**Figure 6G**), which is consistent with the findings of a previous study showing that BCG-iv vaccination trains HSCs to generate monocytes/macrophages with enhanced anti-*Mtb* activity *in vivo* (9). Moreover, BCGΔ*pe18*-iv mice-derived BMDMs presented increased anti-*Mtb* capacity in *Rag1*^-/-^ mice compared with those from BCG-iv mice (**Figure 6G**).

Given that PE18 can boost BCG-induced antigen-specific CD4^+^ T cells responses (105), we further explored whether CD4^+^ T cells contribute to BCGΔ*pe18*-mediated protection. We isolated CD4^+^CD44⁺ T cells from the spleens of BCG– or BCGΔ*pe18*-immunized mice and adoptively transferred them into *Rag1*^-/-^ mice, followed by *Mtb* challenge. Notably, the lung bacterial load in *Rag1*^-/-^ mice receiving CD4^+^CD44⁺ T cells from BCGΔ*pe18*-immunized mice was much higher than that in the parental BCG group (**Supplemental Figure 7A, B**).

## Discussion

The induction of innate immune memory has been attributed to epigenetic reprogramming of chromatin accessibility, which modulates nuclear genes encoding proinflammatory cytokines, thereby enabling enhanced transcriptional responsiveness upon subsequent stimulation (10, 106). Here we found that BCG suppresses protective immunity through mitochondrial DNA condensation, linking mitochondrial genome accessibility to innate immune memory. Our identification of PE18 as a previously uncharacterized secreted toxic effector of BCG, which directly inhibits innate immune memory by restricting mitochondrial bioenergetic output and mitochondrial DNA accessibility, introduces the concept of live attenuated vaccines that escape immunity via metabolic evasion, suggesting that pathogen-derived factors may have co-evolved to undermine the persistence of trained immunity. Additionally, our research advances the integration of mitochondrial metabolism and immune training concepts by shifting focus from mitochondria as static sources of reactive oxygen species or ATP production to a more nuanced perspective that mitochondrial genome accessibility and transcriptional activity as regulatory signals for the immune system. This framing of the mitochondrial genome as a regulatory entity in trained immunity is apparently different from the existing chromatin-centered framework. Moreover, a rationally engineered BCGΔ*pe18* strain, which confers robust and lifelong protection against *Mtb* in mice, introduces a potentially actionable strategy for improving vaccine performance through targeted manipulation of mitochondrial nucleoid biology—a paradigm previously unexplored in the context of live attenuated vaccine design. From an evolutionary perspective, cytokines such as IL-1β first appeared ∼420 million years ago around the emergence of the vertebrate subphylum (107), while mtNd1 is present in not only vertebrates but also ancestral insects (108), hinting at a potentially conserved role of mitochondrial DNA transcription in innate immune memory. Whether and how mtDNA accessibility contributes to other types of diseases in the context of different stresses or stimuli remains to be determined.

Mitochondrial genome encodes essential subunits of the oxidative phosphorylation system but is also a major damage-associated molecular pattern (DAMP) that engages innate immune sensors when released into the cytoplasm, extracellular space or circulation (39, 71). The packaging of mtDNA into nucleoids regulates the number of mtDNA molecules available for both transcription and replication (71). Depletion or haploinsufficiency (*Tfam^+/−^*) of TFAM, an abundant mtDNA-binding protein essential for transcription-primed mtDNA replication, nucleoid formation and mtDNA segregation during mitochondrial fission (109–111), leads to enlarged nucleoids, elongated mitochondria and the release of mtDNA into the cytoplasm, which activates the cGAS-STING innate immune signaling pathway (39, 40). We found that infection with PE18-deficient BCG increased ATP-dependent degradation of TFAM by the protease AFG3L2. Unlike TFAM deficiency-mediated mtDNA replication stress (39), BCGΔ*pe18* priming facilitates the decompaction of mtDNA to increase the transcription of mitochondrial-encoded genes that mediate the mitochondrial oxidative response against pathogens, which is distinct from the cGAS-STING mediate activation of type I IFN signaling by mtDNA itself (43, 112). In addition, our work demonstrates that BCG PE18-mediated mtDNA inaccessibility inhibits secondary bacterial clearance in monocytes/macrophages, demonstrating for the first time the critical importance of mtDNA accessibility in regulating innate immune memory. Overall, our research uncovers a previously undiscovered function of mitochondrial DNA in innate immune memory. It will be valuable to explore whether mtDNA accessibility is involved in adaptive immune memory (113, 114) and the protective efficacy of other vaccines (115, 116).

Innate immune memory is canonically linked to metabolic reprogramming and histone modifications, thus leading to subsequent enhancer activation of pro-inflammatory cytokine genes including IL-1β, IL-6 and TNF-α (5, 10, 26, 106). Indeed, we found that BCGΔ*pe18*-trained BMDMs exhibited higher expression of *Il1b*, *Il6* and *Tnf* following secondary *Mtb* challenge compared to BCG-trained macrophages. Innate immune memory typically drives metabolic reprogramming by AKT-mTOR-HIF-1α signaling (26, 97, 99), wherein this reprogramming is characterized by heightened aerobic glycolysis and the conversion of pyruvate to lactate (99). A very recent breakthrough study revealed that increased lactate production upon BCG vaccination enhanced IL-1β expression, a process associated with long-term histone lactylation *in vivo* (26). In contrast, we observed a marked decrease in lactate and fumarate levels in BCGΔ*pe18*-primed BMDMs which is a divergence from the canonical metabolic reprogramming where elevated levels of these metabolites drive proinflammatory trained immunity through histone modifications including H3K18la, H3K27ac, and H3K4me3. Consistently, levels of these histone modifications including H3K27ac, H3K4me3, or H3K18la showed significant reduction in BCGΔ*pe18*-trained BMDMs relative to BCG-trained macrophages (**Supplemental Figure 5g**), suggesting that BCGΔ*pe18*-mediated maintenance of cytokine transcriptional capacity may be independent of classical metabolic rewiring and histone modifications. We illustrated that the BCG PE18-mediated inhibition of the bactericidal capacity of macrophages is dependent on mtDNA-encoded respiratory complex I subunit mtNd1, which leads to the production of mitochondrial ROS and supports the bacterial killing capacity of innate immune memory. Given that mtROS activates nuclear factor-κB (NF-κB) or MAPKs signal to increase cytokines expression (117, 118), both of which are essential for the transcription of *Il1b*, *Il6* and *Tnf* in macrophages. These results suggest that BCGΔ*pe18* may elevate mtROS production upon secondary mycobacterial stimulation, thus activating NF-κB or MAPK pathways to increase cytokines expression.

As tuberculosis (TB) has again become the world’s leading cause of death from a single infectious agent, *Mtb* is a globally prioritized endemic pathogen for vaccine research and development (119). BCG remains the sole TB vaccine licensed for use in humans and provides protection against miliary and meningeal tuberculosis in children, but it is less effective for the prevention of pulmonary tuberculosis, especially in adults. In this study, intravenous immunization with BCGΔ*pe18* provided excellent protection against *Mtb* challenge, in which nine out of ten mice achieved complete *Mtb* clearance while intravenous BCG immunization did not elicit this effect. Although BCG exerts pathogenicity in immunodeficient populations (101–103), BCGΔ*pe18* shows greater safety than BCG in immunodeficient mice, which may be due to the removal of inhibitory effect of PE18 on innate immune memory. In addition, BMDMs isolated from BCGΔ*pe18*-vaccinated mice exhibited increased clearance ability against *S. aureus* and *E. coli*, suggesting its nonspecific protective effects against other pathogens. This study revealed that BCGΔ*pe18* has four advantages over BCG: (i) greater safety, (ii) stronger protective efficacy, (iii) longer protective duration, and (iv) protection against a broader spectrum of bacteria. Taken together, these data support the clinical development of BCGΔ*pe18* against TB and other diseases in adults, especially in elderly people.

A surprising finding in this study was that BCGΔ*pe18* had strong effects against *Mtb* almost two years after intravenous vaccination, which is almost lifelong protection against TB in mice. For intravenous BCG vaccination in a nonhuman primate (NHP) infection models, the longest verified protection time of BCG is 6 months (52). Based on current knowledge, innate immune memory is assumed to have a short lifespan and is thought to be difficult to sustain for such a long duration. To explain this phenomenon, we propose two hypotheses: (i) self-renewing and long-lived macrophages (120–122) could exist in the pulmonary immune system or might be induced by BCGΔ*pe18* vaccination; (ii) mtDNA could be transferred to other cells, such as long-lived memory T/B-cells, as mitochondria could be transferred via intercellular crosstalk (123–125). Nevertheless, the immune mechanism behind the profound protective effects of BCGΔ*pe18* is intriguing to be further explored.

A very recent study report that a trivalent mRNA vaccine consisting of PPE20 (Rv1387), EsxG (Rv0287), and PE18 (Rv1788) augments and exceeds BCG protection in multiple mouse models through boosting BCG-induced CD4^+^ T cells (105). Notably, our adoptive transfer experiments revealed that the lung bacterial load in the *Rag1*^-/-^ mice that receiving CD4^+^CD44⁺ T cell from BCGΔ*pe18*-iv mice was higher than that in the BCG group, which aligns with PE18’s known role in activating protective CD4^+^ T cell responses, suggesting that CD4^+^ T cells induced by BCGΔ*pe18* are less effective at mediating anti-TB protection. CD8^+^ T cells are critical for controlling intracellular pathogens such as *Mtb* through mechanisms including granule exocytosis and cytokine secretion, and previous studies have shown that BCG can induce CD8^+^ T cell responses (126, 127). However, we have not identified the role of CD8^+^ T cells in BCGΔ*pe18*-mediated protection. Future studies using CD8^+^ T cell depletion or adoptive transfer approaches as well as CD4^+^ T cell depletion experiments to clarify the necessity of CD4^+^ T cells for BCGΔ*pe18*-mediated protection will be necessary to clarify these questions.

It has been shown that BCG educated HSCs *in vivo* to generate epigenetically modified macrophages that provide significantly better protection against *Mtb* infection (9), which is proved to be sustainable *in vivo* by using chimeric mice and adoptive transfer approaches. However, BCG was unable to infect HSCs either *in vitro* or *in vivo* (9), while BCG-infected monocyte/macrophage lineage cells was observed (9). Here we demonstrated that BMDMs from BCGΔ*pe18*-vaccinated mice exhibit higher mtDNA accessibility (**Figure 1f**) and confer superior protection against *Mtb* infection in *Rag1*^-/-^ recipients (**Figure 6g**) compared with those from BCG-vaccinated mice, suggesting a sustainable mtDNA reprogramming and protection of HSC-derived monocyte/macrophage lineage in BCGΔ*pe18*-vaccinated mice. Given that BCG-mediated training stems from the primary infection of monocyte/macrophage, our observation of BCG PE18-mediated mtDNA reprogramming in *in vitro* isolated BMDMs may contribute to intracellular mechanisms triggered by BCG internalization in monocyte/macrophage lineage cells. For *in vivo* mtDNA reprogramming of HSCs, mitochondrial transfer from adjacent cells (128, 129), especially macrophages is found as emerging key players in mitochondrial transfers (130). We propose that mtDNA reprogramming in primary infected bone marrow derived monocytes/macrophages might modulate the activation and functions of HSCs via mitochondrial transfer, which may partially explain the robust and long-term protection effects of BCGΔ*pe18* vaccination. However, whether and how mitochondria transfer between BCGΔ*pe18*-trained macrophages/monocytes and HSCs in bone marrow, and whether and how mitochondrial transfer between these cells contributes to broad influence and long-term protection effects of BCGΔ*pe18* needs further exploration.

## Materials and Methods

### Mice

All mice were on the C57BL/6J background. Both males and females were used. All experiments were started at the age of 6-8 weeks unless otherwise stated. *Lyz2*^cre^ mice, *Slc25a5*^flox/flox^ mice and *Rag1*^-/-^ mice were purchased from GemPharmatech Co., Ltd. (Nanjing, China). *Tfam*^flox/flox^ mice were purchased from Shanghai Model Organisms Center, Inc. (Shanghai, China). All animals were maintained in Specific pathogen-free (SPF) animal facilities and were handled in accordance with protocols approved by Laboratory Animal Welfare and Ethics Review Working Committee of Tongji University. Mice were housed under a 12-h light/12-h dark cycle, at ambient temperature of 22 ± 2 °C and relative humidity of 45–65%. No wild animals and no field-collected samples were used in the study.

### Cell culture

Both cell lines and primary cultured cells were used in this study.

#### Cell lines

HEK293T cells (ATCC CRL-3216) were obtained from the American Type Culture Collection (ATCC) and maintained in Dulbecco’s modified Eagle’s medium (DMEM, Gibco, Cat#C11995500BT) supplemented with 10% (v/v) heat-inactivated fetal bovine serum (FBS, ThermoFisher, Cat#26010066) and 1% (v/v) penicillin-streptomycin (100 U/ml penicillin, 100 μg/ml streptomycin; Gibco, Cat#15140122).

#### Primary cultured cells

Primary bone marrow-derived macrophages (BMDMs) were isolated from 6-8-week-old C57BL/6J mice (consistent with the mouse background in previous sections) and cultured in RPMI 1640 medium (Gibco, Cat#C11875500BT) supplemented with 10% (v/v) heat-inactivated FBS, 1% (v/v) penicillin-streptomycin, and 20 ng/ml recombinant mouse M-CSF protein (ABclonal, Cat#RP01216).

All cells were routinely tested for mycoplasma contamination, and only mycoplasma-negative cells were used for experiments. Cells were incubated in a humidified atmosphere of 5% CO₂ at 37 ℃.

### Bacterial culture

*Mycobacterium tuberculosis (M. tuberculosis*, *Mtb)* H37Rv (ATCC 27294), BCG Tokyo 172 (ATCC 35737), BCGΔ*pe18* (a *pe18* gene deletion mutant of BCG) strains were cultured in either liquid or solid medium:

#### Liquid medium

Middlebrook 7H9 broth (7H9, Becton Dickson, Cat#271310) supplemented with 10% (v/v) oleic acid-albumin-dextrose-catalase (OADC, Becton Dickson, Cat#212351), 0.5% (v/v) glycerol (Sigma, Cat#G6279), and 0.05% (v/v) Tween-80 (Sigma, Cat#9005-65-6).

#### Solid medium

Middlebrook 7H10 agar (Becton Dickson, Cat#262710) supplemented with 10% (v/v) OADC.

And antibiotic supplements as required. The antibiotics used were 50 μg/mL penicillin (Sigma, Cat#P4333). All *Mtb* and BCG strains were cultured at 37 ℃ under aerobic conditions (with gentle shaking at 120 rpm for liquid cultures); *Mtb* and BCG strains were grown to mid-log phase (optical density at 600 nm (OD_600_) ∼0.6). *Staphylococcus aureus* (*S. aureus*) or *Escherichia coli* (*E. coli*) strains were grown at 37 °C with shaking in Luria-Bertani broth (LB, ThermoFisher).

### *In vitro* infection model

BMDMs were separately primed with one of the following stimuli: β-glucan (MedChemExpress, Cat#HY-W145521) (10 μg/mL), Oxidized low density lipoprotein (mouse) (oxLDL, MedChemExpress, Cat#HY-NP013) (10 μg/mL), Arabinogalactan (AG, Sigma, Cat#10830) (1 μg/mL), Peptidoglycan (PGN, Sigma, Cat#77140) (10 μg/mL), Muramyl dipeptide (MDP, MedChemExpress, Cat#HY-127090) (10 μg/mL), BCG or BCGΔ*pe18* (MOI = 2). After 24 hours of incubation at 37 ℃ with 5% CO_2_, cells were washed three times with 1× PBS, then cultured in fresh RPMI-1640 medium containing 10% FBS and 50 μg/mL isoniazid (INH, an anti-tuberculosis drug; MedChemExpress, Cat#HY-B0329) for 5 days to eliminate the pre-infected BCG/BCGΔ*pe18.* All BMDM cultures were confirmed to be BCG-free before secondary infection. BMDMs were then infected with *Mtb* H37Rv for secondary infection at indicated time points. Collect cell samples for subsequent experiments.

### *Ex vivo* infection model

C57BL/6J mice or conditional knockout (CKO) mice (*Lyz2*^cre^, *Slc25a5*^flox/flox^), 6-8 weeks old, were intravenously (iv) vaccinated with 2×10^6^ colony-forming units (CFU) of mid-log phase BCG or BCGΔ*pe18* (prepared as single-cell suspension in 100 μL sterile PBS and mice injected i.v. with 100 μL 1× PBS served as the PBS-only control group.

After 1 month or 3 months post-vaccination, BMDMs were isolated from the vaccinated/control mice and cultured in RPMI-1640 medium containing 10% (v/v) heat-inactivated FBS and 50 μg/mL INH for 5 days to eliminate residual BCG. All BMDM cultures were BCG-free at the time of secondary infection. Subsequently, these BCG-free BMDMs were subjected to secondary infection with one of the following bacteria: mid-log phase *Mtb* H37Rv (MOI = 2), mid-log phase *E. coli* (MOI = 10), or mid-log phase *S. aureus* (MOI = 10) for subsequent experiments.

### *In vivo* mice vaccination-challenge infection model

C57BL/6J mice, 6-8 weeks old, were intravenously (iv) vaccinated with 2×10^6^ single-suspended BCG or BCGΔ*pe18* in 100 μL 1× PBS, with PBS used as a control. 30 days after vaccination, mice were challenged via aerosol with 200 CFUs of *Mtb* H37Rv per mouse. Animal procedures were approved by the Animal Experiment Administration Committee of Shanghai Pulmonary Hospital (K18-033), and this study was carried out in strict accordance with the China National Research Council’s Guide for Care and Use of Laboratory Animals. All surgeries were performed under sodium pentobarbital anesthesia, and all efforts were made to minimize suffering. Pulmonary tissues were collected for bacterial burden analysis and histopathology analysis. Pathological sections were stained with hematoxylin and eosin (H&E), and bacterial load in lung sections was analyzed by acid-fast staining. Digital images were captured using the 3Dhistech Pannoramic Scan system (3DHISTECH Ltd.) and processed with the CaseViewer™ application. Histopathological scores were calculated by dividing the inflammatory infiltrated area by the total lung area.

### Adoptive Transfer infection Model

C57BL/6J mice, 6-8 weeks old were intravenously (iv) vaccinated with BCG, BCGΔ*pe18* (2×10^6^ CFU), or PBS. 3 months later, BMDMs were harvested and infected with *Mtb* H37Rv (MOI = 2) for 30 minutes at 37 ℃ and 5% CO_2_ with frequent agitation. Extracellular bacteria were then removed by washing three times with cold RPMI-1640 medium; each wash was followed by centrifugation at 1,500 rpm for 10 minutes at 4 °C. BMDMs (5×10^5^ cells) were resuspended in 40 μL 1× PBS and then intratracheally transferred into naive *Rag1*^-/-^ mice, as described previously (9). CD4^+^CD44^+^ T cells (5×10^4^ cells) isolated from spleen tissues of vaccinated mice were resuspended in 100 μL 1× PBS and intravenously injected into naive *Rag1*^-/-^ mice.

### Small interfering RNA-mediated gene interference

Three small interfering RNAs (siRNA) sequences were designed to silence one gene. Non-targeting siRNA was used as the negative control siRNA (siNC). BMDMs were transfected with a mixture of the three siRNAs using siRNA-Mate transfection reagent jetPRIME^®^ Buffer (ERPAN TECH, Cat#101000046) according to the manufacturer’s instructions. Taking 6-well plate as an example: dilute 22 pmole of siRNA in 200 μL of jetPRIME^®^ Buffer. Vortex for 10 seconds and spin down. Add 4 μL of jetPRIME^®^ reagent. Vortex for 1 second, spin down, and incubate for 10 minutes at room temperature. Add the transfection mix to the cells in serum-containing medium. At 48 hours post siRNA transfection, gene knockdown efficiency was assayed by Western blot or quantitative PCR (qPCR) analysis.

### Cell transfection

HEK293T cells were plated with a density of 2×10^6^ cells/mL in 100 mm cell culture dishes. Cells were transiently transfected using Polyethylenimine (PEI, Polysciences, Cat#23966). Two groups were set up: GFP-vector and GFP-PE18. First, add 10 μg of plasmid DNA to a 1.5 mL EP tube containing 400 μL of DMEM. Then add PEI transfection reagent to the DMEM solution at a recommended PEI/DNA ratio of 3:1. Subsequently, add the PEI solution to the plasmid solution in equal proportions. After co-incubation for 20 minutes at room temperature, slowly drop the mixture into the cell culture. After 48 hours, remove the cell culture medium and wash the cells twice with 1×PBS. For Co-IP experiments, lyse the cells in 1 mL of Western and IP cell lysis buffer (Beyotime, Cat#P0013) supplemented with 1% protease inhibitor cocktail (MedChemExpress, Cat#HY-K0011) at 4 °C. For immunofluorescence analysis, fix the cells with cold 4% paraformaldehyde (PFA, Sangon Biotech, Cat#E672002).

### Mitochondrial isolation

HEK293T cells were plated at a density of 2×10^6^ cells/mL in 100 mm cell culture dishes. Mitochondria were isolated using the Mitochondria Isolation Kit (ThermoFisher, Cat#89874.) according to the manufacturer’s instructions. Pellet 2×10^7^ cells by centrifuging the harvested cell suspension in a 2.0 mL microcentrifuge tube at ∼850 ×g for 2 minutes. Carefully remove and discard the supernatant. Add 800 µL of Mitochondria Isolation Reagent A. Vortex at medium speed for 5 seconds and incubate the tube on ice for exactly 2 minutes. Add 10 µL of Mitochondria Isolation Reagent B. Vortex at maximum speed for 5 seconds. Incubate the tube on ice for 5 minutes, vortexing at maximum speed every minute. Add 800 µL of Mitochondria Isolation Reagent C. Invert tube several times to mix. Centrifuge the tube at 700 ×g for 10 minutes at 4 °C. Transfer the supernatant to a new 2.0 mL tube and centrifuge it at 3,000 ×g for 15 minutes at 4 °C. Transfer the supernatant to a new tube. The pellet contains the isolated mitochondria. Add 500 µL Mitochondria Isolation Reagent C to the pellet, and centrifuge at 12,000 ×g for 5 minutes. Discard the supernatant and maintain the mitochondrial pellet on ice before downstream processing. Freezing and thawing may compromise mitochondrial integrity.

### Tn5 transposome assembly

The assembly of Tn5 transposome was performed as described with minor modifications (63). In brief, adaptor oligos were purchased from Integrated DNA Technologies (IDT) with HPLC purification, including:

**Reverse oligo Tn5Merev:** 5’[phos]CTGTCTCTTATACACATCT-3’

**Tn5ME-A-ATTO488:** 5’/ATTO488N/TCGTCGGCAGCGTCAGATGTGTATAAGAGACAG-3’

**Tn5ME-B-ATTO488:** 5’/ATTO488N/GTCTCGTGGGCTCGGAGATGTGTATAAGAGACAG-3’.

100 μM Tn5MErev was mixed in equal amounts either with 100 μM Tn5ME-A-ATTO488 or 100 μM Tn5ME-BATTO488. The oligo mixtures were denatured for 5 minutes at 95 °C in a thermocycler. The cycler was shut off and the oligos left inside the thermomixer until they reached room temperature. The transposome was assembled in a mixture of mixed oligos (Tn5MErev/Tn5MEA-ATTO488 and Tn5MErev/Tn5ME-B-ATTO488) (final concentration 12.5 μM for each mix), 31.75% glycerol (final concentration reached 47.9%), 0.24× Tn5 Dialysis Buffer and 5 μM Tn5 (Beyotime, Cat#D7102M). The mixture was mixed by carefully pipetting up and down and incubated at room temperature for 60 minutes. The assembled transposome was stored at –20 °C.

### Mitochondrial DNA accessibility assay

BMDMs were plated at a density of 0.5 × 10^6^ cells/mL in glass-bottom culture dishes. Then cells either transfected with plasmids or infected with *Mtb* were fixed with 4% PFA at 4 °C for 24 hours, and then washed three times with 1× PBS. After three washes with 1× PBS, cells were permeabilized with 1× PBS containing 0.25% Triton X-100 (Sangon Biotech, Cat#A417820) for 20 minutes. Cells were washed three times with 1× PBS and then incubated at 37 °C for 1 hour in blocking buffer (e.g., 5% bovine serum albumin (BSA, Sangon Biotech, Cat#A600332) in 1‰ PBST). Tn5 (100 nM) was used in tagmentation buffer (16.5 mM Tris, pH 7.8, 33 mM potassium acetate, 5.5 mM magnesium acetate and 8% N, N-dimethylformamide). For the inactive Tn5 control, the mixture included 50 mM EDTA. After mixing by pipetting, the transposase solution was centrifuged for either 10 or 20 minutes at maximum speed at 4 °C in a tabletop centrifuge to reduce aggregates in the sample. The cells were incubated with the transposase solution for 60 minutes at 37 °C in an oven in the dark. The transposase was inactivated and washed away in three washes for 15 minutes at 37 °C with prewarmed washing buffer (1× PBS, 50 mM EDTA, 0.1% Triton X-100). Afterwards, the cells were rinsed three times with 1× PBS at room temperature, incubated with 5 μg/mL DAPI (Beyotime, Cat#C1002) at room temperature for 30 minutes, and imaged using a Leica SP8 confocal microscope (Leica Microsystems). Confocal images were acquired with the Leica SP8 confocal microscope (Leica Microsystems) and analyzed using Leica Application Suite Las X software (v2.0.1.14392).

### Mitochondrial ATP/ADP exchange assay

The mitochondrial pellet was resuspended in a reaction buffer (10 mM HEPES [pH 7.4], 250 mM sucrose, and 10 mM KCl). The ADP/ATP exchange process was initiated by the addition of ADP at a final concentration of 2 mM. After 5 minutes of incubation, the amount of ATP transported from mitochondria was determined using an ATP measurement kit (Invitrogen, Cat#A22066). Luminescence of samples was detected using a BioTek reader (Synergy 2, BioTek Instruments). For each experiment, a standard curve was generated with serially diluted ATP and was used to calculate the concentration of ATP in samples to determine ADP/ATP exchange rates.

### Mitochondrial ATP assay

The mitochondrial ATP probe Rh6G-NH-PBA (70) was diluted to 10 μM with RPMI-1640 medium and incubated with cells in the dark at 37 °C for 30 minutes. Then cells were washed with 1×PBS and fixed with 4% PFA at 4 ℃ for 24 hour, and then washed three times with 1× PBS. Afterwards, the cells were incubated with 5 μg/mL DAPI at room temperature for 30 minutes and imaged using a confocal fluorescence microscopy.

### Mitochondrial ROS assay

MitoSOX Red (MedChemExpress, Cat#HY-D1055) was diluted to 5 μM with RPMI-1640 medium and incubated with cells in the dark at 37 °C for 30 minutes. Then cells were washed with 1× PBS and fixed with 4% PFA at 4 ℃ for 24 hour, and then washed three times with 1× PBS. Afterwards, the cells were incubated with 5 μg/mL DAPI at room temperature for 30 minutes, imaged using a Leica SP8 confocal microscope (Leica Microsystems), and analyzed using Leica Application Suite Las X (v2.0.1.14392) software.

### Immunofluorescence

BMDMs were primed with BCG or BCGΔ*pe18* (MOI = 2), as described previously. After 3 hours of incubation at 37 ℃ with 5% CO_2_, the supernatant was discarded. MitoTracker™ Red FM (Invitrogen, Cat#A66444) was diluted with RPMI-1640 medium (1:1000) and incubated with cells in the dark at 37 °C for 30 minutes. Then cells were fixed with cold 4% PFA for 1 hour. After three washes with 1× PBS, cells were permeabilized with 1× PBS containing 0.5% Triton X-100 for 20 minutes. Cells were washed once with 1‰ (v/v) PBST and then blocked with 3% BSA for 1 hour at room temperature, washed once with 1‰ PBST, and incubated with primary antibody (1:100) overnight at 4 ℃. The primary antibody was recovered, and cells were washed three times with 1‰ PBST (10 minutes per wash), then incubated with fluorescent secondary antibody (1:500) for 1 hour at room temperature. Cells were wash three times with 1‰ PBST (10 minutes per wash). Afterwards, the cells were incubated with 5 μg/mL DAPI at room temperature for 30 minutes and imaged using a confocal fluorescence microscopy.

### Intracellular CFUs in BMDMs

BMDMs were differentiated into mature macrophages, 30 minutes prior to priming with BCG, BCGΔ*pe18*, or PBS, the inhibitor 2’,3’-dideoxycytidine (ddC, Sigma, Cat#D5782) was added at a final concentration of 150 μM, with dimethyl sulfoxide (DMSO) used as a control. Subsequent experiments were performed as described previously. BMDMs were primed and rested as described previously. 30 minutes prior to the secondary infection with *Mtb* H37Rv, the inhibitors Mito-TEMPO (MedChemExpress, Cat#HY-112879) (10 μM), Rotenone (MedChemExpress, Cat#HY-B1756) (10 nM), Mdivi-1 (MedChemExpress, Cat#HY-15886) (10 μM) and Agaric acid (MedChemExpress, Cat#HY-N4104) (10 μM) were added, with DMSO used as the control. BMDMs were then infected with *Mtb* H37Rv for 3 hour at 37 ℃ in 5% CO_2_. Cells were washed twice with 1 × PBS and subsequently supplemented with RPMI-1640 medium containing 10% FBS. Infected cells were lysed at 3 hours and 24 hours with 1× PBS containing 1% (v/v) Triton X-100. Serial dilutions of the lysate in PBS were plated on 7H10 agar plates with 10% OADC enrichment and 50 μg/mL penicillin. Plates were incubated at 37 ℃ and CFUs were counted after 21-30 days. For *S. aureus* or *E. coli* infection, bacterial strains were cultured overnight in liquid broth and used to infect BMDMs at MOI = 10. 1 hour post infection, cells were washed twice with 1×PBS and subsequently supplemented with RPMI-1640 medium containing 10% FBS and 1% gentamicin. Infected cells were lysed at 1 hour, 3 hour, 9 hours, 12 hours and 24 hours with 1× PBS containing 1% (v/v) Triton X-100. Serial dilutions of the lysate in PBS were plated on LB agar plates and CFUs were counted after 24 hours.

30 minutes prior to the secondary infection with *Mtb* H37Rv, the inhibitor 2’,3’-dideoxycytidine was added to *Mtb*-infected BMDMs at a final concentration of 150 μM, with DMSO used as a control. Then the cells were infected with *Mtb* for 3 hours at 37 ℃ in 5% CO_2_. The cells were washed twice with 1 × PBS and subsequently supplemented with RPMI-1640 medium containing 10% FBS. Infected cells were lysed at 3 hours and 48 hours with 1× PBS containing 1% (v/v) Triton X-100. Serial dilutions in PBS were plated on 7H10 agar plates with 10% OADC enrichment and 50 μg/mL penicillin. Plates were then incubated at 37 ℃ and counted after 21-30 days. For *S. aureus* or *E. coli* infection, bacterial strains were cultured in liquid broth overnight and infected BMDMs at MOI = 10. After 1 h post infection, the cells were washed twice with PBS. Cells were subsequently supplemented with RPMI-1640 containing 10% FBS, 1% Gentamicin. Infected cells were lysed at 1 h and 9 h with 1% (v/v) Triton X-100 in PBS. Serial dilutions in PBS were plated on LB agar plates and counted after 24 h.

### Immunoprecipitation (IP) and mass spectrometry

Sample preparation: HEK293T cells (5 × 10^6^ cells) were transiently transfected using PEI. Two groups were set up: GFP-vector and GFP-PE18. After 48 hours, the cell culture medium was removed, the cells were washed twice with cold 1 × PBS, and the cells were lysed in 1 mL Western and IP cell lysis buffer supplemented with 1% protease inhibitor cocktail. The cells were placed in a 4 ℃ shaker for 30 minutes to ensure complete lysis. The cell lysates were collected into 1.5 mL EP tubes, centrifuged at 12,000 rpm at 4 ℃ for 10 minutes, and the supernatant was retained. The 100 μL of the cell lysate was mixed with 2 × loading buffer (ABclonal, Cat#RM00001) and boiled at 95 ℃ for 10 minutes; The remaining cell lysate was added to anti-GFP beads (MedChemExpress, Cat#HY-K0246) and incubated overnight at 4 ℃. The samples were centrifuged at 6,000 rpm for 3 minutes to remove the supernatant, and washed 5 times with 1 mL 1‰ (v/v) PBST (10 minutes per wash), and the supernatant was completely removed after the final wash. The immunoprecipitated samples were boiled with the SDS-loading buffer and separated on a 10% SDS-PAGE gel for subsequent mass spectrometry analysis.

### Co-immunoprecipitation (Co-IP)

Two groups were set up: GFP-PE18 and GFP-PE18+Flag-SLC25A5. After 48 hours, mitochondria were isolated using the Mitochondria Isolation Kit as described previously. The separated mitochondria were lysed in 1 mL of Western and IP cell lysis buffer supplemented with 1% protease inhibitor cocktail. After centrifugation, the supernatants were incubated with anti-FLAG Magnetic Beads (MedChemExpress. Cat#HY-K0207) overnight at 4 °C. The same operation was also performed for the cytosol fraction. Then the samples were washed three times with 1‰ (v/v) PBST (10 minutes per wash). These protein samples were boiled with the SDS-loading buffer and separated on a 10% SDS-PAGE gel for subsequent immunoblotting analysis.

### Western Blot

For western blot analysis, 20 μg of proteins was separated on a 10% or 15% SDS-polyacrylamide gel. The concentration of the separating gel should be determined according to the molecular weight of the target protein. Stacking gel electrophoresis conditions: 80 V, 30 minutes; Separating gel electrophoresis conditions: 100 V until bromophenol blue migrated out of the separating gel; Transfer conditions: 100 V, 1-2 hours (transfer time can be adjusted according to the molecular weight of the target protein). After transfer on ice, the membranes were blocked with 5% BSA for 1 hour at room temperature, washed three times with 1‰ TBST (v/v), and incubated with primary antibody overnight at 4 ℃. The primary antibody was recovered, and the membranes were washed three times with 1‰ TBST (v/v) (10 minutes per wash), then incubated with horseradish peroxidase-conjugated secondary antibody for 1 hour at room temperature. The membranes were washed three times with 1‰ (v/v) TBST (10 minutes per wash), followed by the addition of chemiluminescence developing solution and exposure. Images were acquired using an Amersham™ Imager 680 (Cytiva) with chemiluminescence reagent (Thermo Scientific, Cat#34580), and subsequent analysis was performed. Following antibodies were used: Rabbit polyclonal anti-TFAM (Affinity, Cat#AF0531); Rabbit polyclonal anti-SLC25A5 (Abclonal, Cat#A15639);

Rabbit monoclonal anti-NRF2Abclonal, Cat#A21176); Rabbit monoclonal anti-β-Actin (Abclonal, Cat#AC026); Mouse monoclonal anti-TOMM20 (Abcam, Cat#AB56783); Mouse monoclonal anti-β-Tubulin (Abclonal, Cat#AC021); Rabbit polyclonal anti-GFP, Abclonal, Cat#AE011); Mouse monoclonal anti-Flag (Sigma, Cat#F3165); Goat anti-Rabbit IgG Secondary antibody (Abclonal, Cat#AS014); Goat anti-Mouse IgG Secondary antibody (Abclonal, Cat#AS003).

### RNA-seq Analyses

BMDBs were set as three treatment groups: BCG-infected, BCGΔ*pe18*-infected, and the control (PBS-treated), with 2 biological replicates per group. After 6 hours of infection, the culture supernatant was discarded, and the cells were washed with 1× PBS. Cell samples were collected with 1mL Trizol reagent (ThermoFisher, Cat#15596026). Total RNA was extracted in accordance with the manufacturer’s instructions (ThermoFisher, Cat#15596026). High-quality total RNA was used to prepare cDNA libraries following the Illumina standard library construction protocol. Sequencing was performed on the Illumina Novaseq 6000 instrument with a read length of 2×150 bp. Raw sequencing data were trimmed using Skewer to remove low-quality reads and adaptor sequences. Clean reads were aligned to the murine genome (mm10) using STAR. Transcript expression levels were calculated as fragments per kilobase of exon model per million mapped reads (FPKM) using Perl scripts. Differentially expressed transcripts (DETs) between groups (BCG-infected vs. control, BCGΔpe18-infected vs. control, and BCG-infected vs. BCGΔpe18-infected) were identified using the MA-plot-based method with random sampling (MARS) model in the DEGseq package. DETs were subjected to Gene Ontology (GO) and Kyoto Encyclopedia of Genes and Genomes (KEGG) pathway enrichment analyses.

### Q-PCR analysis

RNA was extracted from BMDMs subjected to primary treatment with PBS, BCG, or BCGΔ*pe18* and secondary infection with *Mtb* H37Rv. Total RNA was extracted with 1 mL of Trizol reagent in accordance with the manufacturer’s instructions (ThermoFisher, Cat#15596026). Then 1 μg of total RNA was uesed for cDNA synthesis with the ABScript III RT Master Mix for qPCR (ABclonal, Cat#RK20428). Real-time quantitative PCR was performed using the 2× Universal SYBR Green Fast qPCR Mix (ABclonal, Cat#RK21203) on an LC480 thermocycler (Roche, Indianapolis, IN). The 2^-ΔΔCt^ method was adopted to analyze the relative gene expression, with the gene expression normalized to that of *β-actin* (eukaryotic reference gene). Real-time qPCR data were collected from at least three independent experiments, with three technical replicates per experiment.

### Metabolomics

BMDMs cells were plated with a density of 2 × 10^6^ cells/mL in 90 mm plates. BMDM cells were primed with BCG or BCGΔ*pe18* (MOI = 2) for 24 h. The cell pellets were taken, mixed with 1000 μL of extraction solution (MeOH:ACN:H2O, 2:2:1 (v/v)), the extraction solution contain deuterated internal standards, the mixed solution were vortexed for 30 s and incubated in liquid nitrogen for 1 min. The samples were then allowed to thaw at room temperature and vortexed for 30 s. This freeze-thaw cycle was repeated three times.Then the samples were sonicated for 10 min in 4 ℃ water bath, and incubated for 1 h at –40 ℃ to precipitate proteins. The samples were centrifuged at 12000 rpm (RCF=13800(×g),R= 8.6cm) for 15 min at 4 ℃. The supernatant was transferred to a fresh glass vial for analysis. The quality control (QC) sample was prepared by mixing an equal aliquot of the supernatant of samples.

For polar metabolites, LC-MS/MS analyses were performed using anUHPLC system (Vanquish, Thermo Fisher Scientific) with a Waters ACQUITYUPLC BEH Amide (2.1 mm × 50 mm, 1.7 μm) coupled to Orbitrap Exploris 120mass spectrometer (Orbitrap MS, Thermo). The mobile phase consisted of 25mmol/L ammonium acetate and 25 ammonia hydroxide in water (pH =9.75) (A) and acetonitrile (B). The auto-sampler temperature was 4 ℃, and theinjection volume was 2 μL. The Orbitrap Exploris 120 mass spectrometer wasused for its ability to acquire MS/MS spectra on information-dependentacquisition (IDA) mode in the control of the acquisition software (Xcalibur,Thermo). In this mode, the acquisition software continuously evaluates the fullscan MS spectrum. The ESI source conditions were set as following: sheath gasflow rate as 50 Arb, Aux gas flow rate as 15 Arb, capillary temperature 320 ℃,full MS resolution as 60000, MS/MS resolution as 15000, collision energy: SNCE 20/30/40, spray voltage as 3.8 kV (positive) or –3.4 kV (negative), respectively.

The raw data were converted to the mzXML format using ProteoWizard and processed with an in-house program. which was developed using R and based on XCMS, for feature detection, extraction, alignment, and integration. The R package and the BiotreeDB (V3.0) were applied in metabolite Identification (131).

### Chromatin immunoprecipitation followed by sequencing (ChIP-seq)

ChIP-seq analysis were conducted using a Hyperactive Universal CUT&Tag Assay Kit (#TD903, Vazyme Biotech) following the manufacturers’ protocols. Briefly, BMDMs were collected and bound to the ConA-coated beads. Cells were then resuspended in antibody buffer and sequentially incubated with primary antibodies against histone modifications, including H3K18la (PTM Bio, Cat#PTM-1406), H3K27ac (Abmart, Cat#T59439M), and H3K4me3 (Abmart, Cat#MB9009), followed by incubation with secondary antibody (Goat anti-Rabbit IgG H&L, #AB206-01-AA, Vazyme Biotech). Subsequently, samples were incubated with pA-Tn5 transposase (Hyperactive PG-TN5/PA-TN5 Transposon) to mediate transposon activation and tagmentation. Fragmented DNA was isolated, amplified by PCR, and purified using VAHTS DNA Clean Beads (N411, Vazyme Biotech). Libraries were constructed with the TruePrep Index Kit V2 for Illumina. Library quality control included assessment of fragment size distribution and effective concentration using the Quant-iTTM dsDNA HS Assay Kit (Invitrogen, MA, USA) and Agilent 2100 Bioanalyzer (Agilent Technologies). Library quantification was performed with the VAHTS Library Quantification Kit for Illumina before sequencing on an Illumina Novaseq platform with 150 bp paired-end reads.

### Library preparation, sequencing, and data processing of ChIP samples

The quality control of raw sequencing reads was measured by FastQC (v0.12.1). The cutadapt software (v2.6) is then applied to filter the raw reads by removing adapters, low-quality reads and short reads to obtain high-quality clean reads. The filtered clean reads are aligned to the mouse genome assembly mm39 using bowtie2 (v2.5.1). Based on the mapped reads that align to the reference genome, the distribution of DNA insert lengths is calculated. The MAC2 suite (v2.2.9.1) was used for peak calling. Peak annotation was carried out by Homer (v4.11.1). The deepTools (v3.5.1) webserver (GitHub) was used for generating genome coverage bigwig files. Gene sites overrepresentation analysis was performed by Enrichr. Using IGV (Integrative Genomics Viewer) (v2.1.0) software to load the bigwig files allows for the visual inspection of read enrichment across the genome in each group.

### ELISA

BMDMs were plated in 12-well plates at a density of 2 × 10⁶ cells/mL. 30 minutes prior to infection, the inhibitor 2’,3’-dideoxycytidine was added at a final concentration of 150 μM, with DMSO used as a control. BMDMs were primed with BCG or BCGΔpe18 (MOI = 2). After 5 days, the BMDMs were infected with LPS (Sigma, Cat#LPS25) (1μg/mL) for secondary stimulation. and cell supernatants were collected at 6 hours post LPS treatment. The concentration of Interleukin-1β (IL-1β) in the supernatants was measured using a mouse IL-1β ELISA kit (ABclonal, Cat#RK00006), a mouse IL-6 ELISA kit (ABclonal, Cat#RK00008) or a mouse TNF-α ELISA kit (ABclonal, Cat#RK00027) according to the manufacturer’s protocols.

For the ELISA procedure: 100 μL of the standard working solution or sample was added to the corresponding well, followed by incubation at 37 °C for 2 hours. After discarding the liquid in the wells, the plate was washed three times; immediately thereafter, 100 μL of the biotinylated antibody working solution was added and incubated at 37 °C for 60 minutes. The liquid was discarded, and the plate was washed three times. 100 μL of the horseradish peroxidase (HRP)-conjugated enzyme working solution was added to each well and incubated at 37 °C for 30 minutes. The liquid was discarded, and the plate was washed three times. 100 μL of 3,3’,5,5’-tetramethylbenzidine (TMB) substrate solution was added to each well and incubated at 37 °C for approximately 15 minutes. 50 μL of stop solution was added to each well, and the absorbance was immediately measured at 450 nm (primary wavelength) and 570 nm (reference wavelength) for data analysis.

### Fluorescence-Activated Cell Sorting (FACS)

Spleen tissues isolated from vaccinated mice were gently homogenized through a 70 μm cell strainer into a fresh 50 mL centrifuge tube to obtain a single-cell suspension. The strainer was rinsed with 5 mL 1× PBS to collect residual cells. The cell suspension was centrifuged at 400×g for 5 minutes at 4°C. The supernatant was discarded, and the cell pellet was resuspended in 10 mL ice-cold RBC lysis buffer. And the suspension was incubated at 4°C for 10 minutes to lyse RBCs. Then lysis was terminated by adding 10 mL ice-cold 1× PBS, followed by centrifugation at 400×g for 5 minutes at 4°C. The supernatant was discarded, and final cell pellet was resuspended in 1 mL 1× PBS and count the cells. Cells were adjusted to a concentration of 1×10⁸ cells/mL in 1× PBS. The suspension was transferred to a 5 mL polystyrene round-bottom tube. For every 1×10⁸ cells, 100 μL of Biotinylated Antibody Cocktail (from the kit (BioLegend, Cat# 480006)) was added. The suspension was gently mixed and incubated at 4°C for 15 minutes. And 100 μL of Streptavidin Nanobeads (from the kit) was added, the suspension was mixed gently and incubated at 4°C for 10 minutes. To minimize the loss of cell number, 2 mL 1× PBS was added in to the tube. The tube was placed in the MojoSort™ Magnet and incubated at room temperature for 3 minutes to allow magnetic separation of bead-bound (non-CD4^+^ T) cells. Repeat the procedure once. The supernatant was carefully transferred to a new 50 mL centrifuge tube. The isolated CD4^+^ T cells were centrifuged at 400×g for 5 minutes at 4°C. The supernatant was discarded. Staining for CD4^+^ T cells: Mouse CD4 FITC GK1.5 (BD Pharmingen, Cat# 557307), PE Rat Anti-Mouse CD44 (IM7) (BD Pharmingen, Cat# 553134), all biotin-conjugated were added at a 1:100 dilution and incubated with cells at 4 °C for 30 minutes. Cells were subsequently washed with 0.5% BSA. Perform sorting using a flow cytometer (BD Biosciences). Sorting was performed using a flow cytometer (BD FACSAria™ II), and CD4⁺CD44⁺ cells were collected.

## Statistical analyses

All data were analyzed in Prism 9.0 software (GraphPad). Statistically significant differences between two groups were determined using two-tailed unpaired Student’s *t* tests between two groups. For comparisons involving three or more groups, one-way or two-way analysis ANOVA test was employed, respectively. P-values for all statistical comparisons are shown in the figures. A *P*<0.05 value was deemed statistically significant. All data were taken from technical replicate averages from several independent experiments and represented by the mean ± standard error of the mean (SEM).

## Data and code availability

All data needed to evaluate the conclusions in the paper are available in the main text and supplementary materials. Additional relevant data are available from the corresponding author upon reasonable request.

## Supporting information

Supplementary Table 1. mtATAC-see signal in HEK293T cells transfected with 162 Mycobacterial scretary protein plasmids

Supplementary Table 2. co-IP combined mass spectromery analysis of GFP-PE18 in HEK293T cells

Supplementary Table 3. RNA-seq analysis of mitochondrial encoded genes in BMDMs

## Acknowledgements

We thank Professor Yicheng Sun (Chinese Academy of Medical Sciences and Peking Union Medical College) for providing CRISPR knockout plasmids, Professor Yen Wei (Tsinghua University) for providing mitochondrial ATP probe (Rh6Gsingle bond NH single bond PBA), Professor Junhao Zhu (Institute of Microbiology, Chinese Academy of Sciences) for helpful methods of fluorescence imaging. This project was supported by the National Natural Science Foundation of China (32188101, 32030038, 91842303, and 3170025 to B.G.); the National Key R&D Program of China (2023YFC2307300; 2022YFC2302900, 2021YFA1300902); China Postdoctoral Science Foundation (2024M762428 to C. P.); The Most Important Clinical Discipline in Shanghai (2017ZZ02003); the Shanghai Clinical Research Center for Infectious Disease (tuberculosis) (19MC1910800).

## Author contributions

B.G., L.W., and C.P. conceived the study and designed experiments. C.P., Q.C. and J. H. developed methodology. C.P., and Q.C. performed CFU assays and vaccination assay. C.P., and J.H., H.C., Y.C., J.W. and M.M. performed mouse experiments and histopathology analysis. J.H., C.P., Q.C., W.B. and X.W. performed western blot, and confocal experiments. B.G., L.W. and C.P. wrote the manuscript, and all authors commented on the manuscript, data, and conclusion.

---

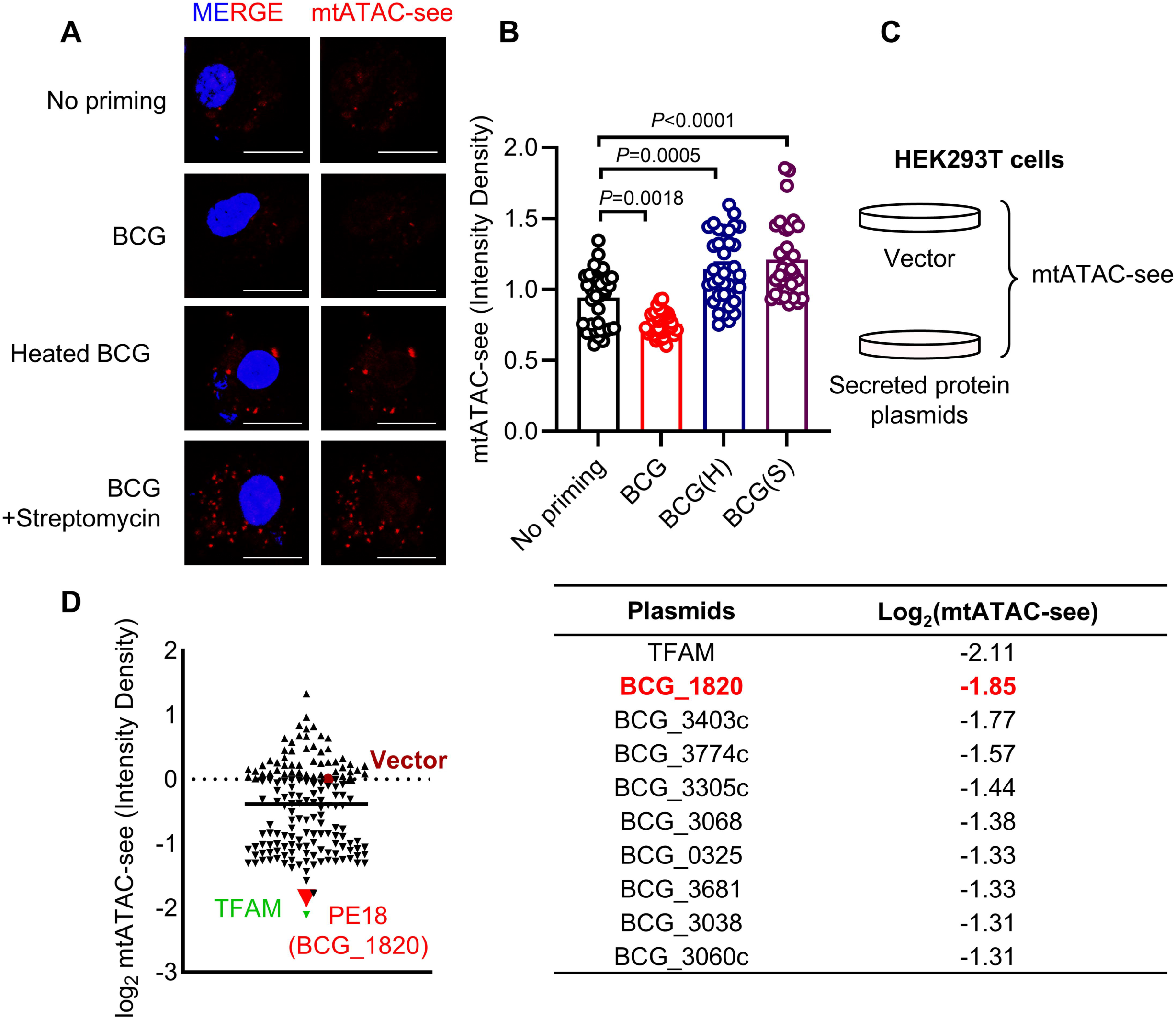

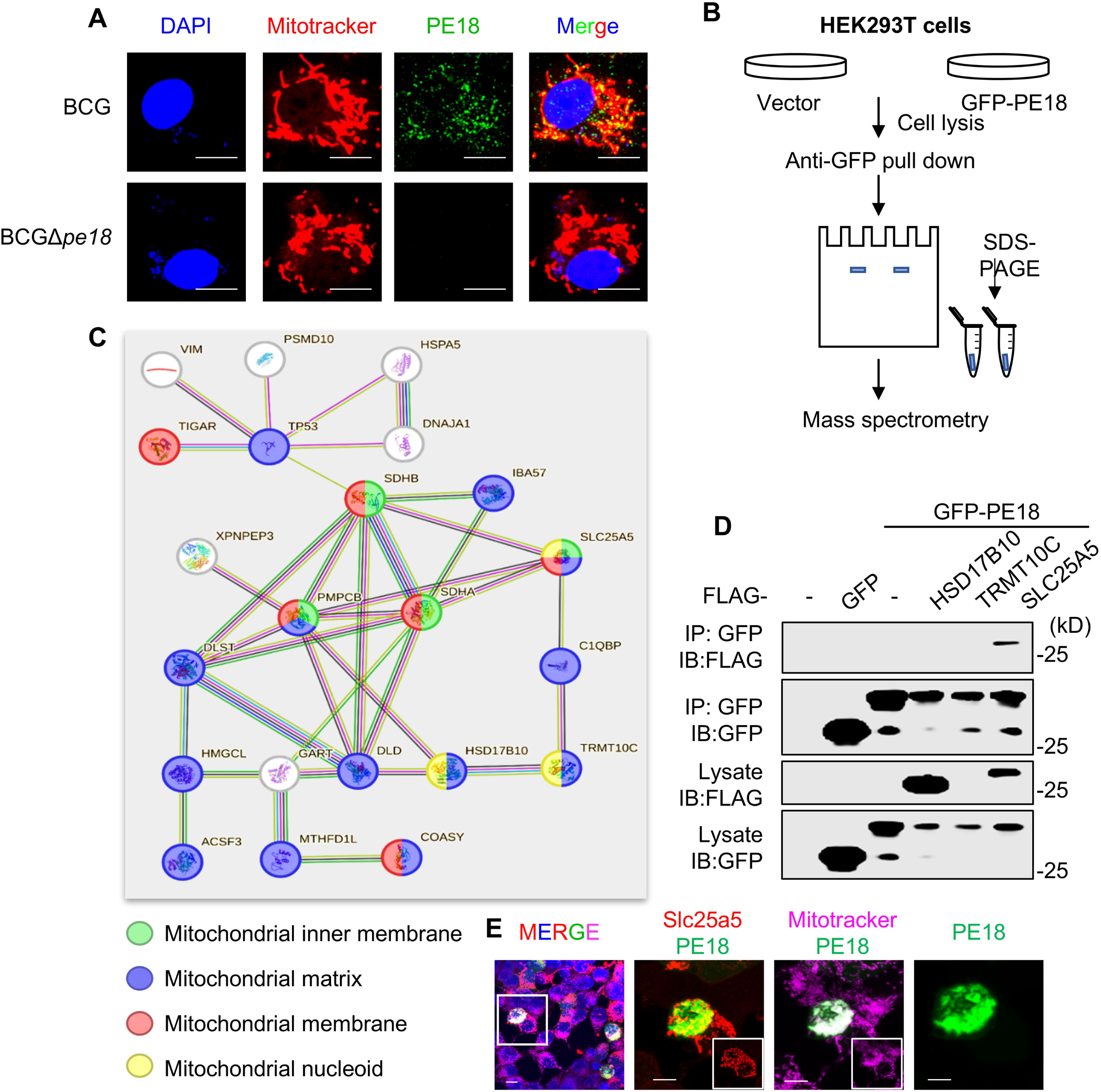

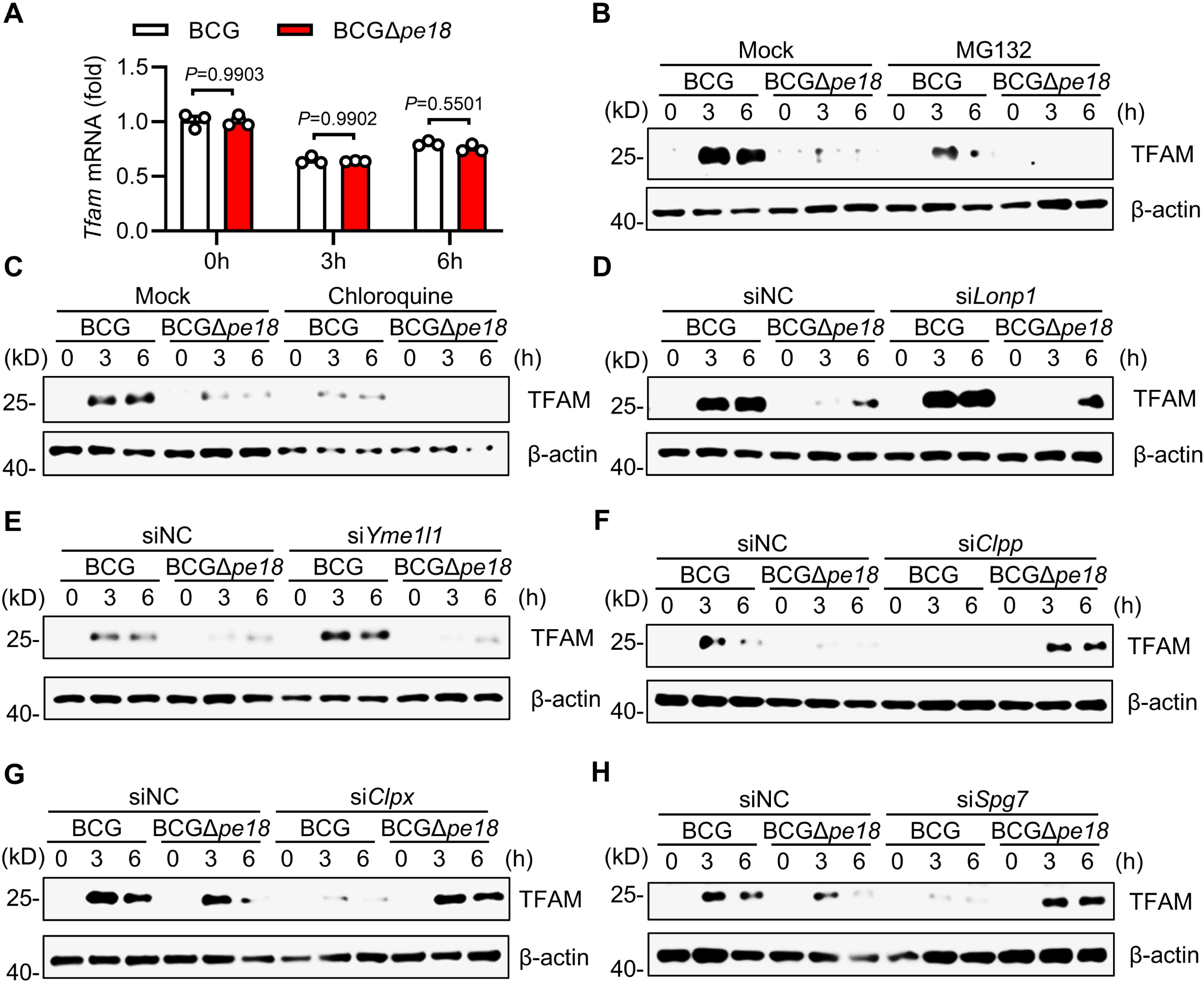

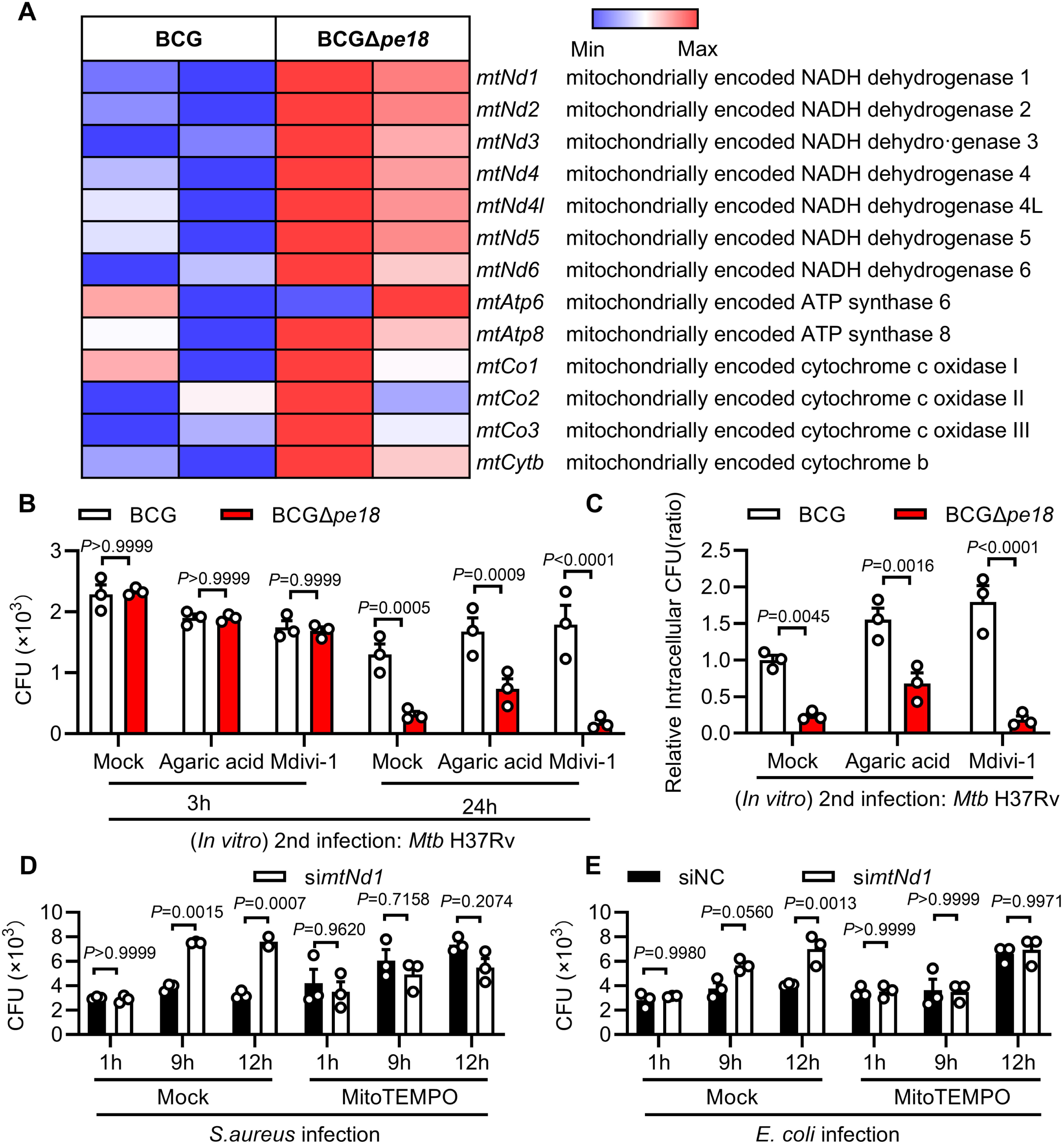

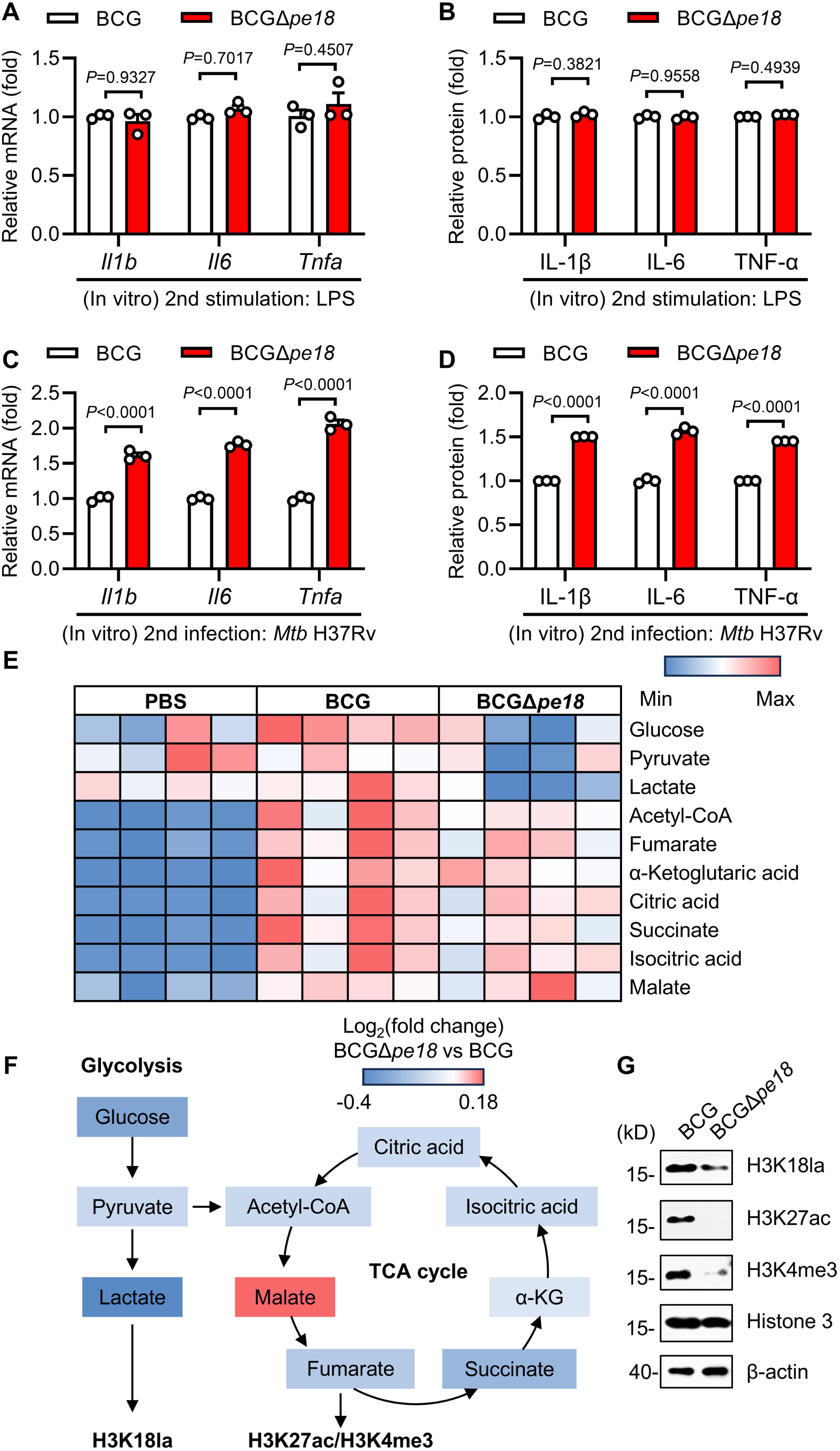

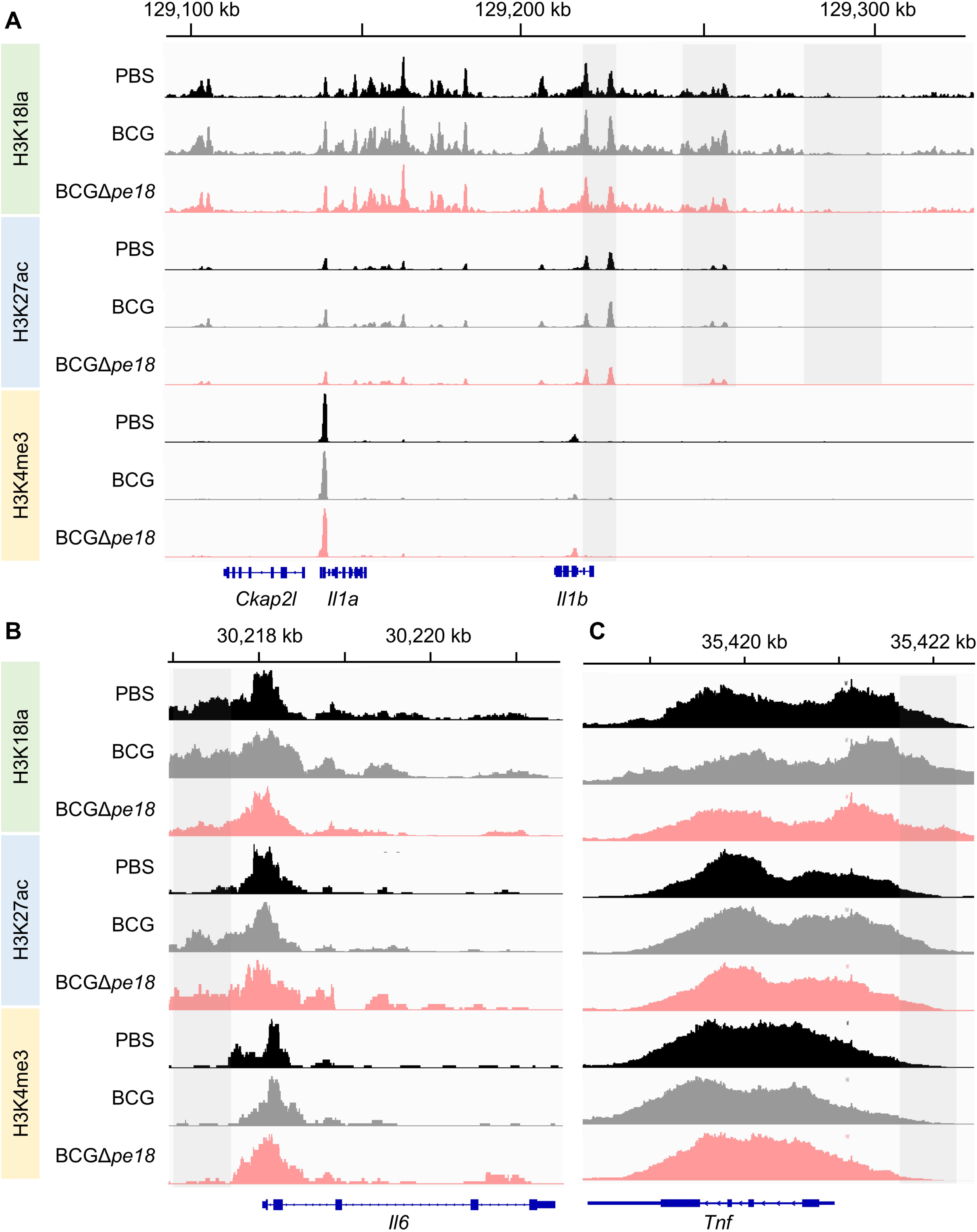

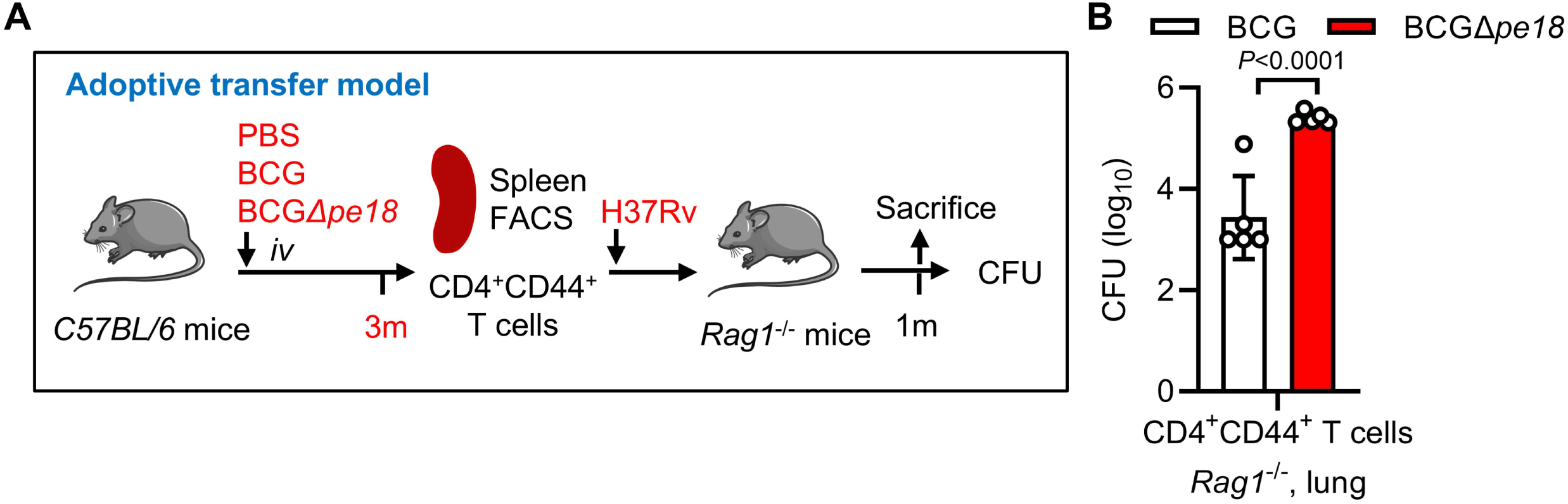

