## Supplemental Figure for "A BCG toxic effector induces mitochondrial DNA compaction to refrain protective immunity"

**Supplemental Figure 1. BCG PE18 inhibits mtDNA accessibility**

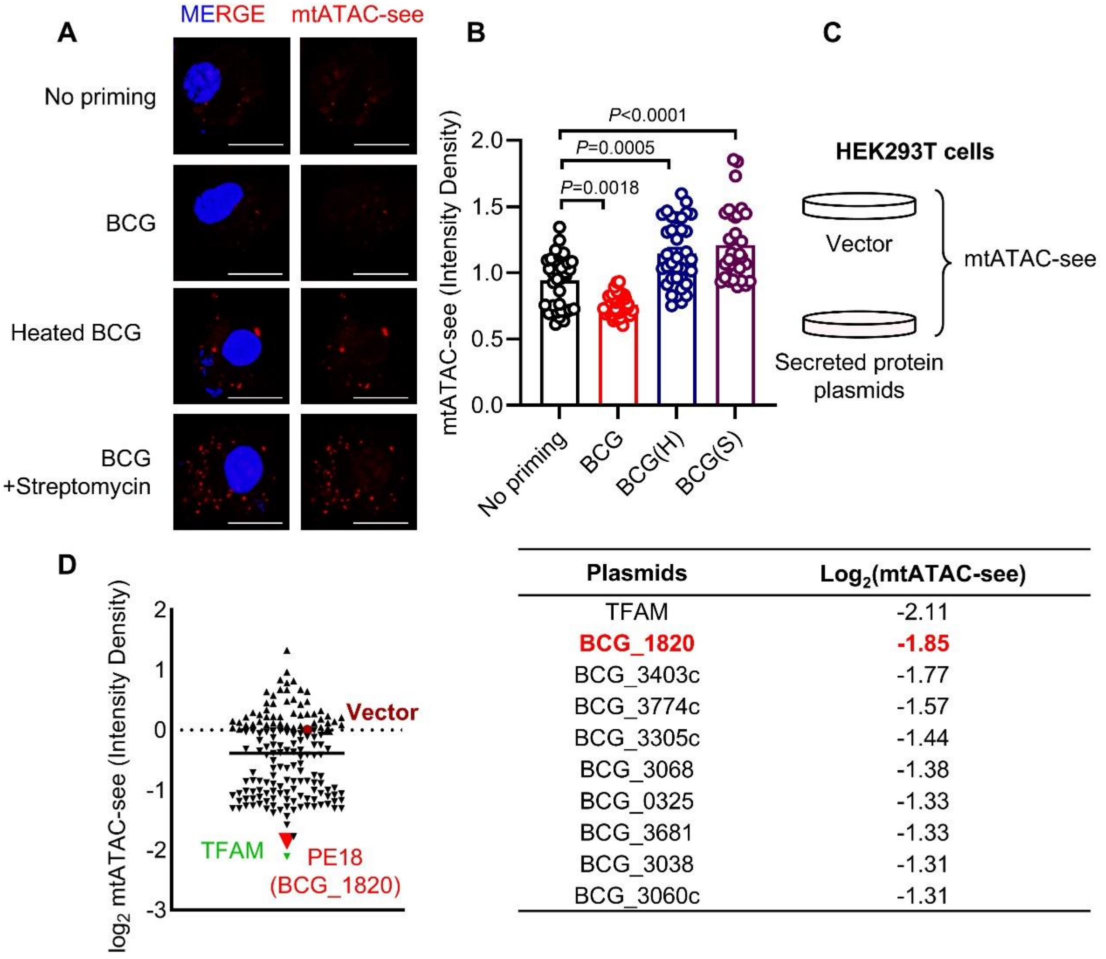


| Plasmids | $Log_2$(mtATAC-see) |
|---|---|
| TFAM | -2.11 |
| **BCG_1820** | **-1.85** |
| BCG_3403c | -1.77 |
| BCG_3774c | -1.57 |
| BCG_3305c | -1.44 |
| BCG_3068 | -1.38 |
| BCG_0325 | -1.33 |
| BCG_3681 | -1.33 |
| BCG_3038 | -1.31 |
| BCG_3060c | -1.31 |

**A-B**. Representative images (**A**) and quantification (**B**) of the mtATAC-see signal in BMDMs primed with BCG, heated BCG (H), or streptomycin-treated BCG (S) for 24 h. Scale bars, 10 μm. Each single spot is representative of a single cell. (mean ± s.e.m. of n=35). **C-D**. Workflow (**C**), mtATAC-see signal analysis and bottom 10 plasmids (**D**) of HEK293T cells transfected with empty vector or plasmids encoding 161 Mycobacterial secretory proteins independently. *P* values were calculated via One-way ANOVA test (**B**).

**Supplemental Figure 2. PE18 binds with SLC25A5 in mitochondria**

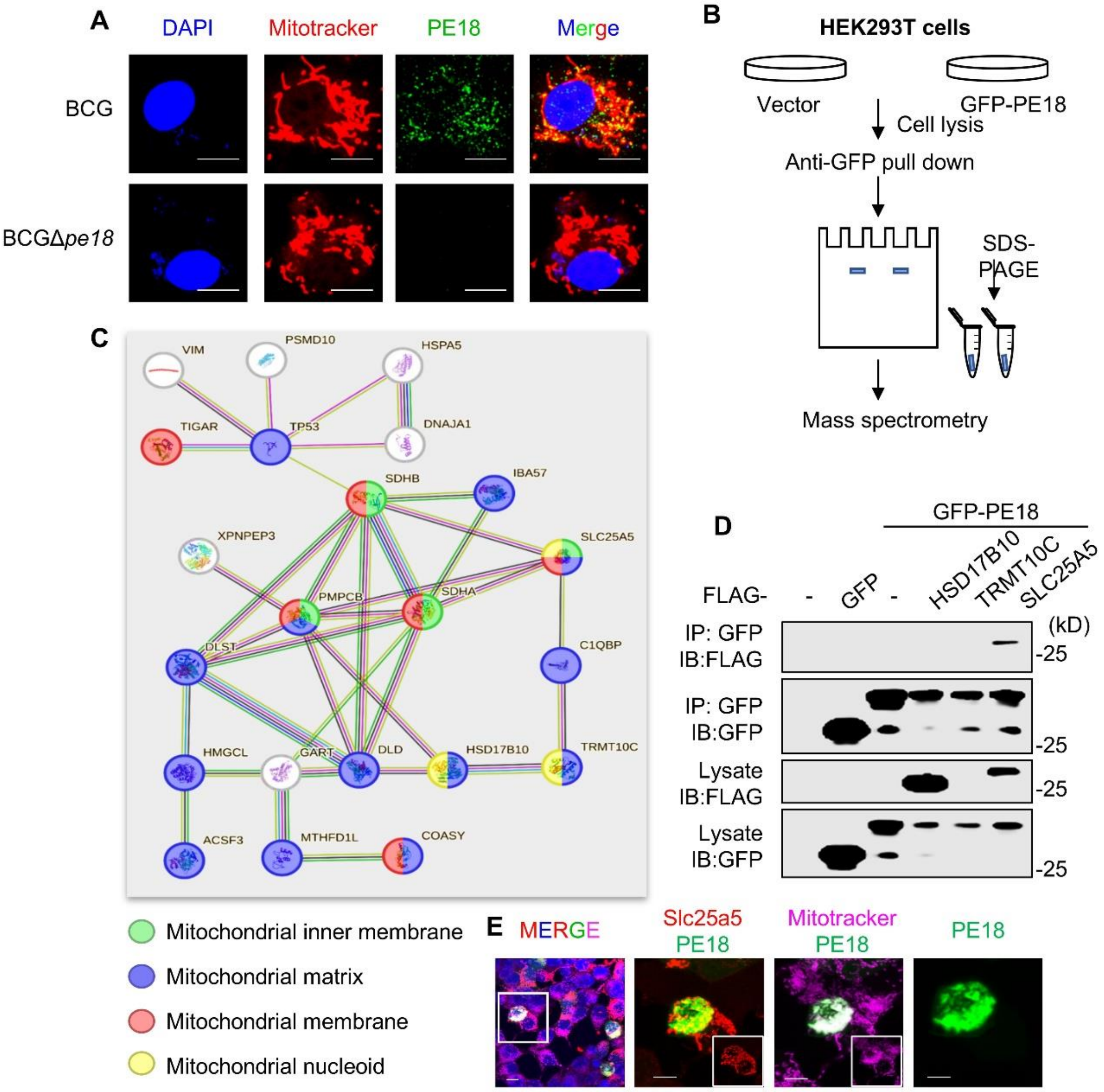


**A**. Confocal image of PE18 in BMDMs primed with BCG or BCGΔ*pe18* for 3 h. Scale bars, 10 μm. **B**. Workflow of HEK293T cells transfected with the GFP vector or GFP-PE18 followed by coimmunoprecipitation with anti-GFP beads and mass spectrometry analysis. Scale bars, 10 μm. Each single spot is representative of a single cell. (mean ± s.e.m. of n=35). **C**. STRING pathway of the top 22 proteins from the spectrometry analysis. **D**. Coimmunoprecipitation and immunoblot analysis of HE293T cells transfected with the indicated plasmids. **E**. Fluorescence colocalization analysis of HE293T cells transfected with GFP-PE18 and FLAG-SLC25A5.

**Supplemental Figure 3. PE18 increases TFAM protein level**

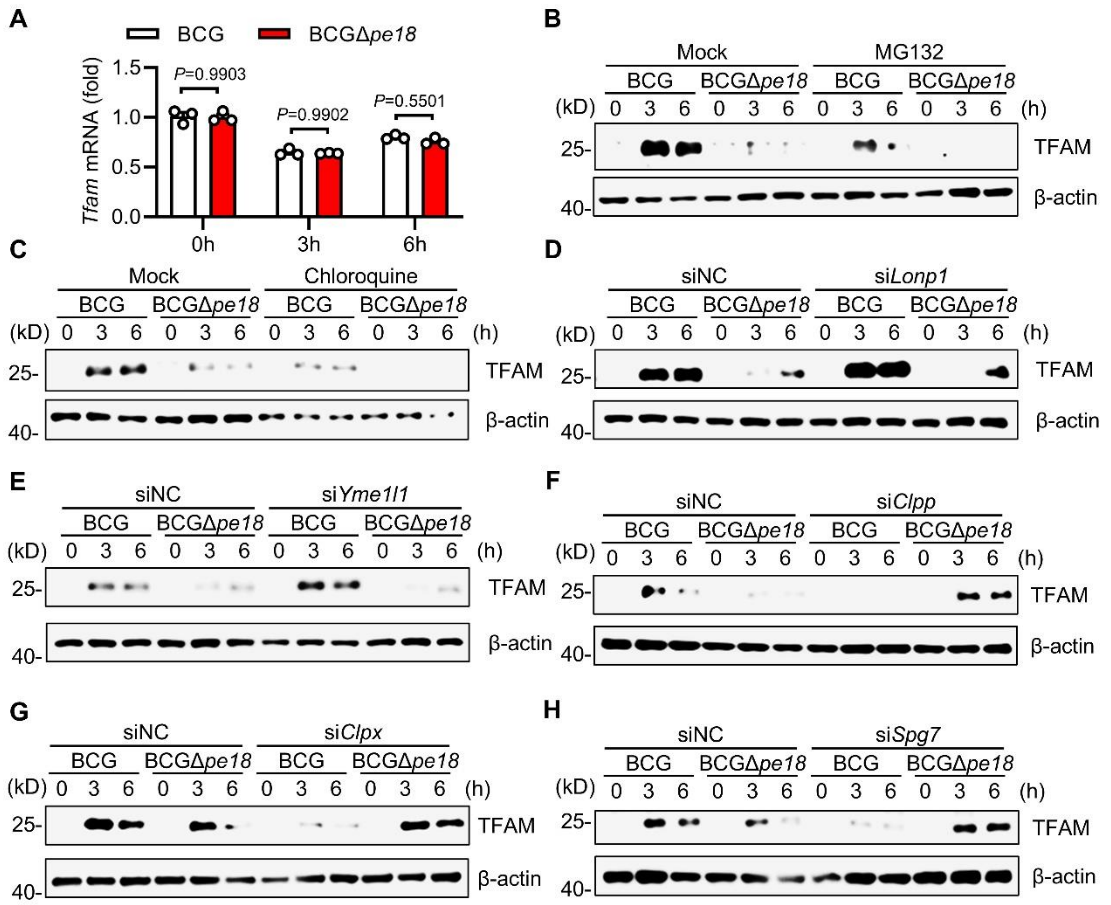


**A**. Q-PCR analysis of *Tfam* mRNA expression in BMDMs stimulated with BCG or BCGΔ*pe18* for the indicated times. (MOI=2) (mean ± s.e.m. of n=3). **B-C**. Immunoblot analysis of BMDMs pretreated with MG132 (**B**) or chloroquine (**C**) followed by stimulation with BCG or BCGΔ*pe18* for 3 or 6 h. **D-H**. Immunoblot analysis of BMDMs transfected with si*Lonp1*, si*Yme1l1*, si*Clpp*, si*Clpx* or si*Spg7* followed by stimulation with BCG or BCGΔ*pe18* for 3 or 6 h. *P* values were calculated via two-way ANOVA tests (**A**).

**Supplemental Figure 4. BCGΔ*pe18* promotes innate immune memory via mtDNA**

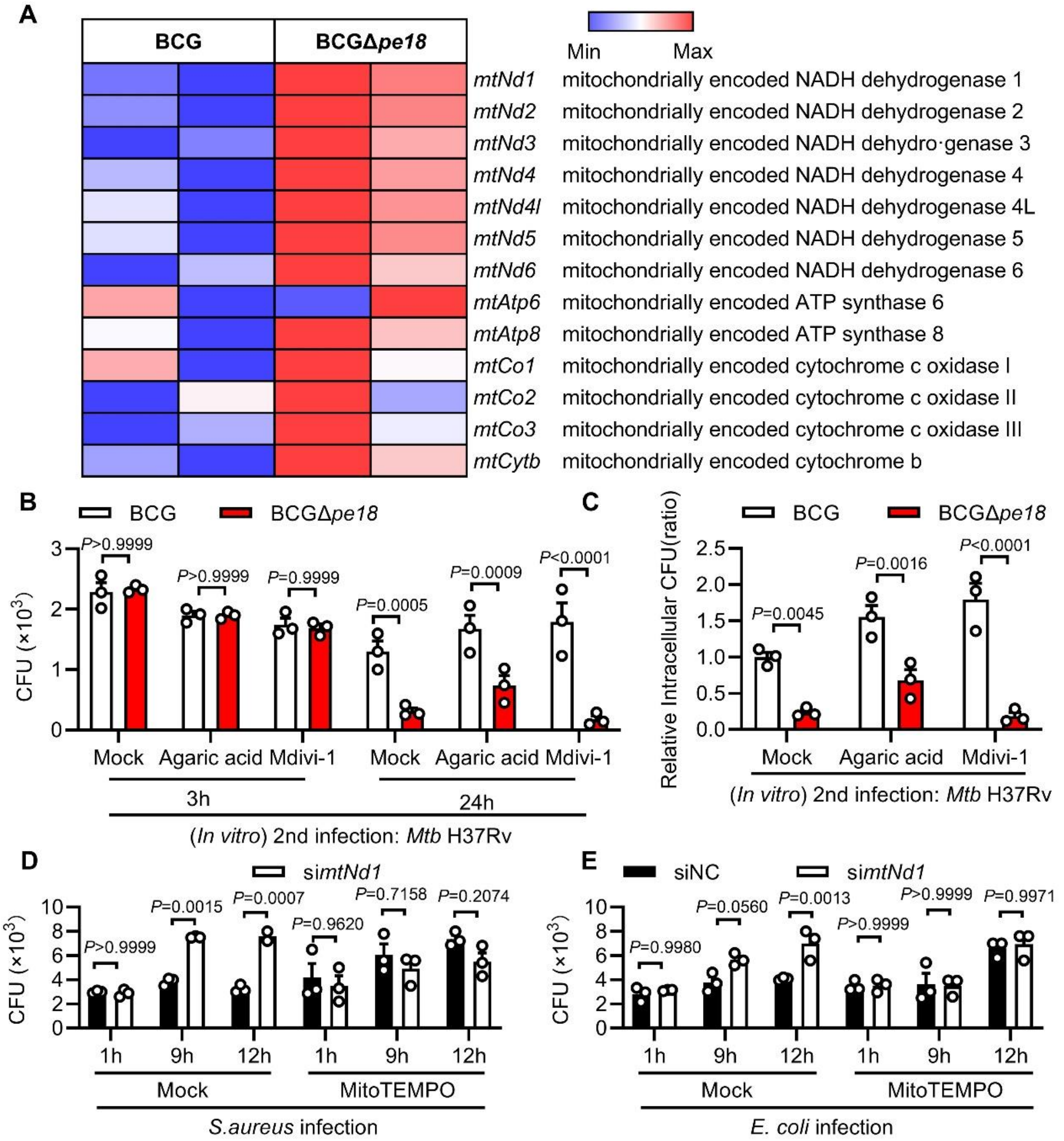


**A.** Heatmap analysis for transcription of 13 mitochondrial DNA encoded genes in BCG- or BCGΔ*pe18*-primed BMDMs infected with *Mtb* H37Rv for 6 h. **B-C**. CFU counts (**B**) and relative intracellular CFU ratios (**C**) of secondary *Mtb* H37Rv infections in BMDMs primed with BCG or BCGΔ*pe18* followed by pretreatment with agaric acid or Mdivi-1 and infection with *Mtb* H37Rv (n=3). **D-E**. CFU counts of BMDMs transfected with si*mtNd1* followed by pretreatment with MitoTEMPO and infection with *S. aureus* (**D**) or *E. coli* (**E**). (n=3). *P* values were calculated via two-way ANOVA tests (**B-E**).

**Supplemental Figure 5 Cytokine expression and metabolic reprogramming in BCG and BCGΔ*pe18*-primed macrophages**

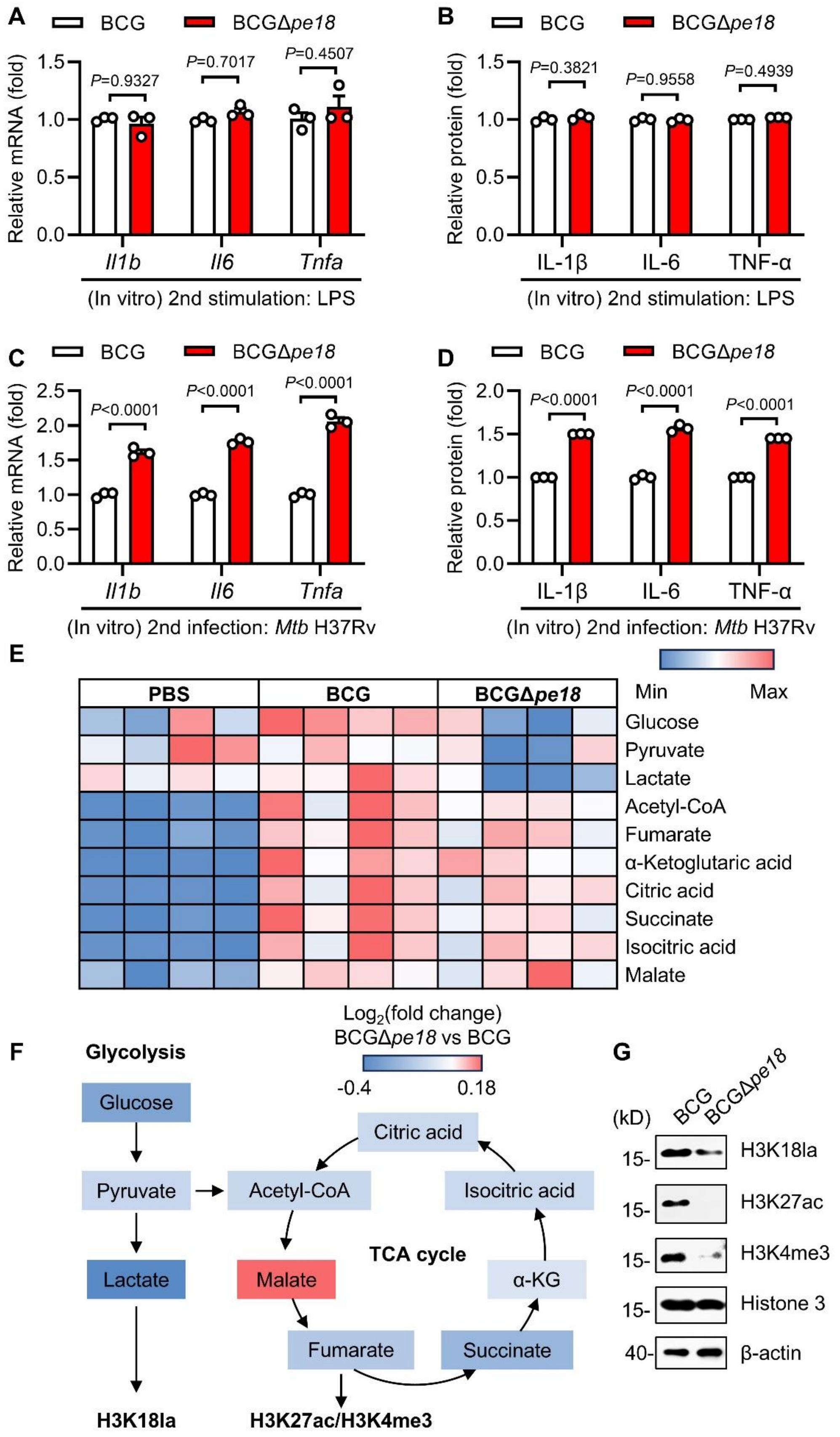

**A-D**. Q-PCR analysis and ELISA analysis of BMDMs primed with BCG or BCGΔ*pe18* followed by secondary LPS stimulation of *Mtb* H37Rv infection (n=3). **E.** Heatmap analysis for metabolites involved in glycolysis and TCA cycle in BCG- or BCGΔ*pe18*-primed BMDMs for 24 h. **F**. Metabolic pathway analysis of relative metabolites abundance in BCGΔ*pe18*-primed BMDMs compared with BCG-primed macrophages. **G**. Immunoblot analysis of si*Slc25a5*-knockdown and siNC control BMDMs primed with BCG or BCGΔ*pe18*. *P* values were calculated via two-way ANOVA tests (**A-D**).

**Supplemental Figure 6. Epigenetic reprogramming in BCG- and BCGΔ*pe18*-primed macrophages**

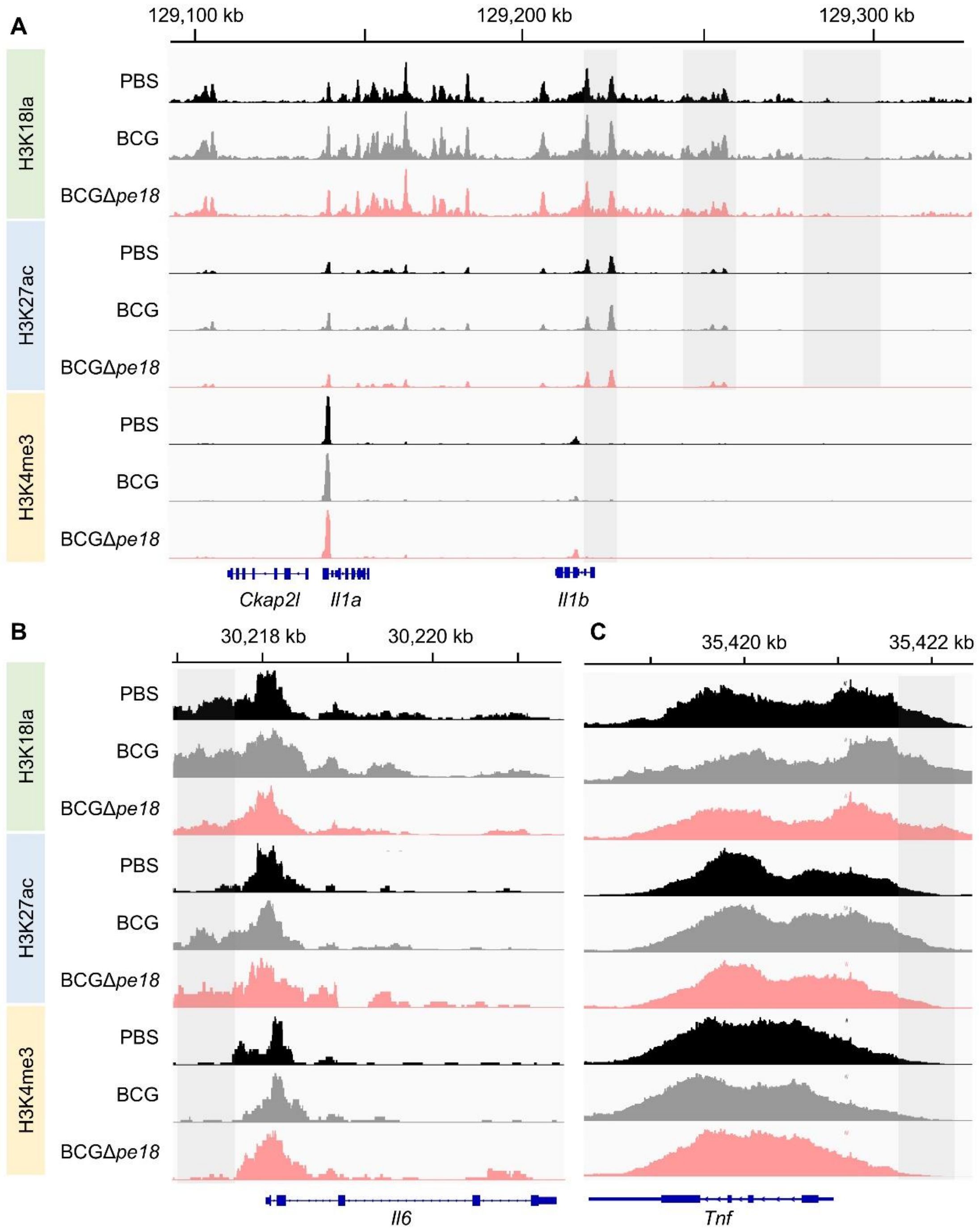


**A-C.** Tracks of H3K18la, H3K27ac and H3K4me3 modification levels in promoter and coding region of *Il1b* (**A**), *Il6* (**B**) and *Tnf* (**C**) in BMDMs from BCG-iv or BCGΔ*pe18*-iv vaccinated mice with PBS-control mice after 21 days of vaccination (gray rectangle are representative peaks).

**Supplemental Figure 7. Adoptive transfer of CD4$^+$CD44$^+$ cells in *Rag1*$^{-/-}$ mice**

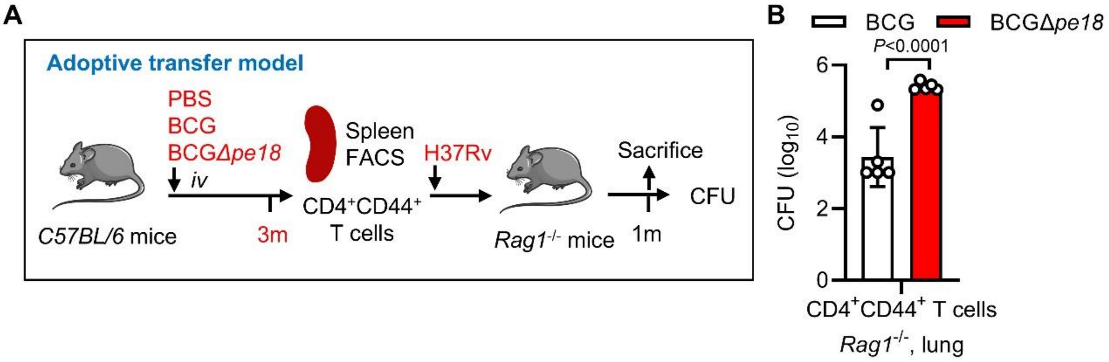


**A-B**. Workflow (**A**) and CFU counts (**B**) in the lung tissues of *Rag1*$^{-/-}$ mice intravenously transplanted with CD4$^+$CD44$^+$ T cells derived from mice intravenously vaccinated with PBS, BCG or BCGΔ*pe18* for 3 months and infected with *Mtb* H37Rv for 1 month. *P* values were calculated via two-tailed unpaired Student's *t* tests (**A**).
