## Supplementary Table 1. mtATAC-see signal in HEK293T cells transfected with 162 Mycobacterial scretary protein plasmids for "A BCG toxic effector induces mitochondrial DNA compaction to refrain protective immunity"

**Table S1. mtATAC-see signal in HEK293T cells transfected with 162 Mycobacterial scretary protein plasmids**

| Plasmid | $log_2$mtATAC-see(Intensity Density) |
|---|---|
| Vector | 1 |
| TFAM | 0.232095051 |
| BCG_1820 | 0.277434166 |
| BCG_3403c | 0.292367092 |
| BCG_3774c | 0.335561949 |
| BCG_3305c | 0.36941381 |
| BCG_3068 | 0.38522397 |
| BCG_0325 | 0.397338423 |
| BCG_3681 | 0.39742819 |
| BCG_3038 | 0.401999828 |
| BCG_3060c | 0.404662168 |
| BCG_3128c | 0.404753688 |
| BCG_2317 | 0.404813598 |
| BCG_2390c | 0.405449527 |
| BCG_2932 | 0.410309262 |
| BCG_3854 | 0.412164606 |
| BCG_2967c | 0.413674651 |
| BCG_1837c | 0.416805018 |
| BCG_3915 | 0.417309672 |
| BCG_1329c | 0.417316135 |
| BCG_3889 | 0.41814394 |
| BCG_1777 | 0.421500983 |
| BCG_2305 | 0.423815556 |
| BCG_3216c | 0.425678144 |
| BCG_3932 | 0.437951755 |
| BCG_3559c | 0.438023659 |
| BCG_3869c | 0.439064656 |
| BCG_3026 | 0.440701353 |
| BCG_1593c | 0.44440681 |
| BCG_2022c | 0.447475443 |
| BCG_1580c | 0.448934245 |
| BCG_3649 | 0.451176666 |
| BCG_0750 | 0.452728103 |
| BCG_1550 | 0.452979468 |
| BCG_1447 | 0.453649232 |
| BCG_3864c | 0.461581653 |
| BCG_3405 | 0.463769142 |
| BCG_1773c | 0.466288855 |
| BCG_2271 | 0.470238221 |

| Plasmid | $log_2$mtATAC-see(Intensity Density) |
|---|---|
| BCG_2241c | 0.472464342 |
| BCG_0689 | 0.480820918 |
| BCG_2541c | 0.480990092 |
| BCG_3079c | 0.481800771 |
| BCG_1532 | 0.48269592 |
| BCG_2672c | 0.482736595 |
| BCG_2486c | 0.485927817 |
| BCG_2723 | 0.491017134 |
| BCG_2673c | 0.492814634 |
| BCG_2287 | 0.49403332 |
| BCG_0614 | 0.498298283 |
| BCG_2050c | 0.499443683 |
| BCG_2926 | 0.500390352 |
| BCG_0112 | 0.506141919 |
| BCG_2585 | 0.513130845 |
| BCG_2886c | 0.519065935 |
| BCG_2155 | 0.538713308 |
| BCG_0188c | 0.540019244 |
| BCG_3937c | 0.544046065 |
| BCG_3865c | 0.544073201 |
| BCG_2536c | 0.553953909 |
| BCG_0449c | 0.554410895 |
| BCG_2087c | 0.554537516 |
| BCG_1480 | 0.555378526 |
| BCG_3777 | 0.556070835 |
| BCG_1098c | 0.559240888 |
| BCG_3792 | 0.560159713 |
| BCG_0890 | 0.585301448 |
| BCG_0383c | 0.590469139 |
| BCG_2686 | 0.595196976 |
| BCG_1122c | 0.611411219 |
| BCG_2897 | 0.621333236 |
| BCG_2368 | 0.644835508 |
| BCG_3832c | 0.644841092 |
| BCG_0232 | 0.651875128 |
| BCG_2472c | 0.662166617 |
| BCG_1893 | 0.678869008 |
| BCG_2060c | 0.701833391 |
| BCG_0328 | 0.735091299 |
| BCG_3268c | 0.745722338 |
| BCG_1918c | 0.74828888 |
| BCG_0349 | 0.7513055 |

| Plasmid | $log_2$mtATAC-see(Intensity Density) |
|---|---|
| BCG_0988 | 0.751937458 |
| BCG_2553c | 0.765352582 |
| BCG_2124 | 0.770210355 |
| BCG_3300c | 0.777268869 |
| BCG_0046c | 0.786675679 |
| BCG_0465c | 0.792525107 |
| BCG_1929 | 0.795177037 |
| BCG_2703c | 0.803091475 |
| BCG_2576c | 0.805200033 |
| BCG_1089 | 0.821663095 |
| BCG_1539 | 0.834427446 |
| BCG_1828 | 0.840934112 |
| BCG_3669c | 0.841775785 |
| BCG_3378c | 0.841875461 |
| BCG_0595c | 0.856018252 |
| BCG_0436c | 0.884846912 |
| BCG_1237c | 0.886655009 |
| BCG_3022 | 0.916483103 |
| BCG_0940 | 0.920176293 |
| BCG_1274c | 0.925243873 |
| BCG_0720 | 0.945603284 |
| BCG_2895 | 0.950815263 |
| BCG_0441c | 0.959431226 |
| BCG_1921c | 0.959840308 |
| BCG_3117c | 0.971694769 |
| BCG_2554c | 0.97730355 |
| BCG_2403c | 0.979767201 |
| BCG_3560c | 0.987612965 |
| BCG_2206c | 0.990541415 |
| BCG_1073c | 0.990709007 |
| BCG_0209 | 0.99659661 |
| BCG_0457 | 1.003858351 |
| BCG_3935c | 1.006547302 |
| BCG_3400 | 1.027251389 |
| BCG_0729c | 1.027959645 |
| BCG_3172 | 1.031184485 |
| BCG_0450c | 1.051342553 |
| BCG_0869c | 1.0534883 |
| BCG_0374c | 1.057939585 |
| BCG_3549c | 1.059519879 |
| BCG_2470c | 1.060378235 |
| BCG_0435c | 1.066916739 |

| Plasmid | $log_2$mtATAC-see(Intensity Density) |
|---|---|
| BCG_0518 | 1.078672279 |
| BCG_1472c | 1.080995901 |
| BCG_1487c | 1.083201204 |
| BCG_0760 | 1.085143249 |
| BCG_3483c | 1.092843392 |
| BCG_3754 | 1.101009822 |
| BCG_3693 | 1.101234472 |
| BCG_3437c | 1.108689211 |
| BCG_2863 | 1.13338238 |
| BCG_1295 | 1.153181289 |
| BCG_0917 | 1.158919413 |
| BCG_3557c | 1.163864197 |
| BCG_2493 | 1.165814934 |
| BCG_3488c | 1.168558982 |
| BCG_3302 | 1.177068695 |
| BCG_3866c | 1.187270727 |
| BCG_2802c | 1.193792225 |
| BCG_2620 | 1.200614749 |
| BCG_1312c | 1.20664816 |
| BCG_1356 | 1.214430371 |
| BCG_2994c | 1.223807569 |
| BCG_2900c | 1.249765376 |
| BCG_3822 | 1.262464107 |
| BCG_0071c | 1.346996943 |
| BCG_0782 | 1.359461707 |
| BCG_0150c | 1.365859326 |
| BCG_3115 | 1.389852603 |
| BCG_0210 | 1.395090172 |
| BCG_2783c | 1.426601821 |
| BCG_0163c | 1.490738019 |
| BCG_1406 | 1.556017213 |
| BCG_3556c | 1.563068202 |
| BCG_1462 | 1.590220676 |
| BCG_1327c | 1.594460184 |
| BCG_3766c | 1.595949312 |
| BCG_0604c | 1.722497152 |
| BCG_1430 | 1.762593175 |
| BCG_1479 | 1.788922808 |
| BCG_3742 | 1.95396988 |
| BCG_3555 | 2.513735606 |
