## Supplementary Table 2. co-IP combined mass spectromery analysis of GFP-PE18 in HEK293T cells for "A BCG toxic effector induces mitochondrial DNA compaction to refrain protective immunity"

**Table S2. co-IP combined mass spectromery analysis of GFP-PE18 in HEK293T cells**

| Protein name | Description |
|---|---|
| ACSF3 | Malonate--CoA ligase ACSF3, mitochondrial; Catalyzes the initial reaction in intramitochondrial fatty acid synthesis, by activating malonate and methylmalonate, but not acetate, into their respective CoA thioester. May have some preference toward very-long-chain substrates ; Belongs to the ATP-dependent AMP-binding enzyme family. |
| C1QBP | Complement component 1 Q subcomponent-binding protein, mitochondrial; Is believed to be a multifunctional and multicompartmental protein involved in inflammation and infection processes, ribosome biogenesis, protein synthesis in mitochondria, regulation of apoptosis, transcriptional regulation and pre-mRNA splicing. At the cell surface is thought to act as an endothelial receptor for plasma proteins of the complement and kallikrein-kinin cascades. Putative receptor for C1q; specifically binds to the globular 'heads' of C1q thus inhibiting C1; may perform the receptor function through a [...] |
| COASY | Phosphopantetheine adenylyltransferase; Bifunctional enzyme that catalyzes the fourth and fifth sequential steps of CoA biosynthetic pathway. The fourth reaction is catalyzed by the phosphopantetheine adenylyltransferase, coded by the coaD domain; the fifth reaction is catalyzed by the dephospho-CoA kinase, coded by the coaE domain. May act as a point of CoA biosynthesis regulation. |
| DLD | Dihydrolipoyl dehydrogenase, mitochondrial; Lipoamide dehydrogenase is a component of the glycine cleavage system as well as an E3 component of three alpha-ketoacid dehydrogenase complexes (pyruvate-, alpha-ketoglutarate-, and branched- chain amino acid-dehydrogenase complex). The 2-oxoglutarate dehydrogenase complex is mainly active in the mitochondrion. A fraction of the 2- oxoglutarate dehydrogenase complex also localizes in the nucleus and is required for lysine succinylation of histones: associates with KAT2A on chromatin and provides succinyl-CoA to histone succinyltransferase KA [...] |
| DLST | 2-oxoglutarate dehydrogenase E2 component (dihydrolipoamide succinyltransferase); Dihydrolipoamide succinyltransferase (E2) component of the 2- oxoglutarate dehydrogenase complex. The 2-oxoglutarate dehydrogenase complex catalyzes the overall conversion of 2-oxoglutarate to succinyl- CoA and $CO(2)$. The 2-oxoglutarate dehydrogenase complex is mainly active in the |

| Protein name | Description |
|---|---|
| | mitochondrion. A fraction of the 2-oxoglutarate dehydrogenase complex also localizes in the nucleus and is required for lysine succinylation of histones: associates with KAT2A on chromatin and provides succinyl-CoA to histone s [...] |
| DNAJA1 | DnaJ homolog subfamily A member 1; Co-chaperone for HSPA8/Hsc70. Stimulates ATP hydrolysis, but not the folding of unfolded proteins mediated by HSPA1A (in vitro). Plays a role in protein transport into mitochondria via its role as co-chaperone. Functions as co- chaperone for HSPA1B and negatively regulates the translocation of BAX from the cytosol to mitochondria in response to cellular stress, thereby protecting cells against apoptosis. Promotes apoptosis in response to cellular stress mediated by exposure to anisomycin or UV. |
| GART | Trifunctional purine biosynthetic protein adenosine-3; Phosphoribosylglycinamide formyltransferase, phosphoribosylglycinamide synthetase, phosphoribosylaminoimidazole synthetase; In the central section; belongs to the AIR synthase family. |
| HMGCL | Hydroxymethylglutaryl-CoA lyase, mitochondrial; Mitochondrial 3-hydroxymethyl-3-methylglutaryl-CoA lyase that catalyzes a cation-dependent cleavage of (S)-3-hydroxy-3- methylglutaryl-CoA into acetyl-CoA and acetoacetate, a key step in ketogenesis. Terminal step in leucine catabolism. Ketone bodies (beta-hydroxybutyrate, acetoacetate and acetone) are essential as an alternative source of energy to glucose, as lipid precursors and as regulators of metabolism. |
| HSD17B10 | 3-hydroxyacyl-CoA dehydrogenase type-2; Mitochondrial dehydrogenase that catalyzes the beta-oxidation at position 17 of androgens and estrogens and has 3-alpha- hydroxysteroid dehydrogenase activity with androsterone. Catalyzes the third step in the beta-oxidation of fatty acids. Carries out oxidative conversions of 7-alpha-OH and 7-beta-OH bile acids. Also exhibits 20-beta-OH and 21-OH dehydrogenase activities with C21 steroids. By interacting with intracellular amyloid-beta, it may contribute to the neuronal dysfunction associated with Alzheimer disease (AD). Essential for structural [...] |
| HSPA5 | Endoplasmic reticulum chaperone BiP; Endoplasmic reticulum chaperone that plays a key role in protein folding and quality control in the endoplasmic reticulum lumen. Involved in the correct folding of proteins and degradation of misfolded proteins via its interaction with DNAJC10/ERdj5, probably to facilitate the release of DNAJC10/ERdj5 from its substrate (By similarity). Acts |

| Protein name | Description |
| --- | --- |
| | as a key repressor of the ERN1/IRE1-mediated unfolded protein response (UPR). In the unstressed endoplasmic reticulum, recruited by DNAJB9/ERdj4 to the luminal region of ERN1/IRE1, leading to disrupt the dimeriz [...] |
| IBA57 | Putative transferase CAF17, mitochondrial; Involved in the maturation of mitochondrial 4Fe-4S proteins functioning late in the iron-sulfur cluster assembly pathway. |
| MTHFD1L | Monofunctional C1-tetrahydrofolate synthase, mitochondrial; May provide the missing metabolic reaction required to link the mitochondria and the cytoplasm in the mammalian model of one-carbon folate metabolism in embryonic an transformed cells complementing thus the enzymatic activities of MTHFD2; In the N-terminal section; belongs to the tetrahydrofolate dehydrogenase/cyclohydrolase family. |
| PMPCB | Mitochondrial-processing peptidase subunit beta; Catalytic subunit of the essential mitochondrial processing protease (MPP), which cleaves the mitochondrial sequence off newly imported precursors proteins (Probable). Preferentially, cleaves after an arginine at position P2 (By similarity). Required for PINK1 turnover by coupling PINK1 mitochondrial import and cleavage, which results in subsequent PINK1 proteolysis. |
| PSMD10 | 26S proteasome non-ATPase regulatory subunit 10; Acts as a chaperone during the assembly of the 26S proteasome, specifically of the PA700/19S regulatory complex (RC). In the initial step of the base subcomplex assembly is part of an intermediate PSMD10:PSMC4:PSMC5:PAAF1 module which probably assembles with a PSMD5:PSMC2:PSMC1:PSMD2 module. Independently of the proteasome, regulates EGF-induced AKT activation through inhibition of the RHOA/ROCK/PTEN pathway, leading to prolonged AKT activation. Plays an important role in RAS-induced tumorigenesis. |
| SDHA | Succinate dehydrogenase [ubiquinone] flavoprotein subunit, mitochondrial; Flavoprotein (FP) subunit of succinate dehydrogenase (SDH) that is involved in complex II of the mitochondrial electron transport chain and is responsible for transferring electrons from succinate to ubiquinone (coenzyme Q). Can act as a tumor suppressor ; Belongs to the FAD-dependent oxidoreductase 2 family. FRD/SDH subfamily. |
| SDHB | Succinate dehydrogenase [ubiquinone] iron-sulfur subunit, mitochondrial; Iron-sulfur protein (IP) subunit of succinate dehydrogenase (SDH) that is involved in complex II of the |

| Protein name | Description |
|---|---|
| | mitochondrial electron transport chain and is responsible for transferring electrons from succinate to ubiquinone (coenzyme Q). |
| SLC25A5 | ADP/ATP translocase 2, N-terminally processed; Catalyzes the exchange of cytoplasmic ADP with mitochondrial ATP across the mitochondrial inner membrane. As part of the mitotic spindle-associated MMXD complex it may play a role in chromosome segregation. |
| TIGAR | Fructose-2,6-bisphosphatase TIGAR; Fructose-bisphosphatase hydrolyzing fructose-2,6-bisphosphate as well as fructose-1,6-bisphosphate. Acts as a negative regulator of glycolysis by lowering intracellular levels of fructose-2,6-bisphosphate in a p53/TP53-dependent manner, resulting in the pentose phosphate pathway (PPP) activation and NADPH production. Contributes to the generation of reduced glutathione to cause a decrease in intracellular reactive oxygen species (ROS) content, correlating with its ability to protect cells from oxidative or metabolic stress-induced cell death. Plays a [...] |
| TP53 | Cellular tumor antigen p53; Acts as a tumor suppressor in many tumor types; induces growth arrest or apoptosis depending on the physiological circumstances and cell type. Involved in cell cycle regulation as a trans-activator that acts to negatively regulate cell division by controlling a set of genes required for this process. One of the activated genes is an inhibitor of cyclin-dependent kinases. Apoptosis induction seems to be mediated either by stimulation of BAX and FAS antigen expression, or by repression of Bcl-2 expression. Its pro-apoptotic activity is activated via its intera [...] |
| TRMT10C | tRNA methyltransferase 10 homolog C; Mitochondrial tRNA N(1)-methyltransferase involved in mitochondrial tRNA maturation. Component of mitochondrial ribonuclease P, a complex composed of TRMT10C/MRPP1, HSD17B10/MRPP2 and PRORP/MRPP3, which cleaves tRNA molecules in their 5'-ends. Together with HSD17B10/MRPP2, forms a subcomplex of the mitochondrial ribonuclease P, named MRPP1-MRPP2 subcomplex, which displays functions that are independent of the ribonuclease P activity. The MRPP1-MRPP2 subcomplex catalyzes the formation of N(1)-methylguanine and N(1)-methyladenine at position 9 (m1G9 a [...] |
| VIM | Vimentin; Vimentins are class-III intermediate filaments found in various non-epithelial cells, especially mesenchymal cells. Vimentin is attached to the nucleus, endoplasmic reticulum, and mitochondria, either laterally or terminally. |

| Protein name | Description |
|---|---|
| XPNPEP3 | Xaa-Pro aminopeptidase 3; Catalyzes the removal of a penultimate prolyl residue from the N-termini of peptides, such as Leu-Pro-Ala. Also shows low activity towards peptides with Ala or Ser at the P1 position. |
