## Supplementary Table 3. RNA-seq analysis of mitochondrial encoded genes in BMDMs for "A BCG toxic effector induces mitochondrial DNA compaction to refrain protective immunity"

**Table S3. RNA-seq analysis of mitochondrial encoded genes in BMDMs**

| Symbol | Description | BG_1 | BG_2 | PE_1 | PE_2 |
|---|---|---|---|---|---|
| mt-Nd1 | mitochondrially encoded NADH dehydrogenase 1 | 994.35 | 778.35 | 2078.77 | 1915.57 |
| mt-Nd2 | mitochondrially encoded NADH dehydrogenase 2 | 393.67 | 246.72 | 830.33 | 749.17 |
| mt-Nd3 | mitochondrially encoded NADH dehydrogenase 3 | 93.51 | 117.89 | 208.44 | 183.25 |
| mt-Nd4 | mitochondrially encoded NADH dehydrogenase 4 | 505.76 | 294.17 | 833.81 | 730.35 |
| mt-Nd4l | mitochondrially encoded NADH dehydrogenase 4L | 365.88 | 69.86 | 621.37 | 527.25 |
| mt-Nd5 | mitochondrially encoded NADH dehydrogenase 5 | 550.42 | 197.84 | 876.46 | 773.55 |
| mt-Nd6 | mitochondrially encoded NADH dehydrogenase 6 | 571.63 | 740.82 | 981.47 | 866.79 |
| mt-Atp6 | mitochondrially encoded ATP synthase 6 | 1695.82 | 335.60 | 470.58 | 2051.43 |
| mt-Atp8 | mitochondrially encoded ATP synthase 8 | 2369.23 | 860.89 | 3354.09 | 2690.20 |
| mt-Co1 | mitochondrially encoded cytochrome c oxidase I | 8130.30 | 5819.77 | 8796.44 | 7675.41 |
| mt-Co2 | mitochondrially encoded cytochrome c oxidase II | 2196.23 | 2666.19 | 2935.70 | 2444.87 |
| mt-Co3 | mitochondrially encoded cytochrome c oxidase III | 1205.38 | 1518.61 | 2069.20 | 1694.51 |
| mt-Cytb | mitochondrially encoded cytochrome b | 750.45 | 499.47 | 1300.69 | 1076.30 |
| mt-Rnr1 | mitochondrially encoded 12S rRNA | 437.89 | 683.33 | 527.84 | 589.30 |
| mt-Rnr2 | mitochondrially encoded 16S rRNA | 368.67 | 655.06 | 506.93 | 560.81 |
| mt-Ta | mitochondrially encoded tRNA alanine | 80.36 | 104.84 | 134.74 | 115.87 |
| mt-Tc | mitochondrially encoded tRNA cysteine | 77.43 | 109.38 | 123.78 | 105.30 |
| mt-Td | mitochondrially encoded tRNA aspartic acid | 1.32 | 0.98 | 0.85 | 1.90 |
| mt-Te | mitochondrially encoded tRNA glutamic acid | 105.56 | 205.38 | 167.90 | 155.42 |
| mt-Tf | mitochondrially encoded tRNA phenylalanine | 0.38 | 0.59 | 1.18 | 1.71 |
| mt-Tg | mitochondrially encoded tRNA glycine | 5.66 | 7.56 | 13.94 | 15.12 |
| mt-Th | mitochondrially encoded tRNA histidine | 1.68 | 5.28 | 16.49 | 18.16 |
| mt-Ti | mitochondrially encoded tRNA isoleucine | 6.74 | 13.48 | 6.64 | 6.06 |

| Symbol | Description | BG_1 | BG_2 | PE_1 | PE_2 |
|---|---|---|---|---|---|
| mt-Tk | mitochondrially encoded tRNA lysine | 0.59 | 2.23 | 4.51 | 5.63 |
| mt-Tl1 | mitochondrially encoded tRNA leucine 1 | 97.35 | 39.10 | 161.24 | 137.39 |
| mt-Tl2 | mitochondrially encoded tRNA leucine 2 | 1.80 | 8.53 | 20.53 | 18.52 |
| mt-Tm | mitochondrially encoded tRNA methionine | 2.58 | 5.44 | 8.04 | 8.77 |
| mt-Tn | mitochondrially encoded tRNA asparagine | 137.99 | 181.03 | 221.28 | 192.86 |
| mt-Tp | mitochondrially encoded tRNA proline | 67.78 | 52.94 | 97.29 | 87.14 |
| mt-Tq | mitochondrially encoded tRNA glutamine | 6.04 | 12.47 | 7.28 | 8.88 |
| mt-Tr | mitochondrially encoded tRNA arginine | 4.20 | 4.28 | 10.59 | 12.03 |
| mt-Ts1 | mitochondrially encoded tRNA serine 1 | 9.97 | 41.16 | 11.50 | 12.74 |
| mt-Ts2 | mitochondrially encoded tRNA serine 2 | 0.19 | 0.23 | 0.22 | 0.27 |
| mt-Tt | mitochondrially encoded tRNA threonine | 1.87 | 1.47 | 7.34 | 2.48 |
| mt-Tv | mitochondrially encoded tRNA valine | 0.66 | 0.06 | 2.17 | 1.72 |
| mt-Tw | mitochondrially encoded tRNA tryptophan | 3.22 | 4.71 | 7.27 | 6.26 |
| mt-Ty | mitochondrially encoded tRNA tyrosine | 27.93 | 29.88 | 39.73 | 33.83 |
