## Supplemental Data 1 for "A BCG toxic effector induces mitochondrial DNA compaction to refrain protective immunity"

**Supplementary Table 4. Metabolomics in BCG- and BCGΔ*pe18*-primed macrophages**

| Metabolites | PBS_1 | PBS_2 | PBS_3 | PBS_4 | BCG_1 | BCG_2 | BCG_3 | BCG_4 | BCGΔ*pe18*_1 | BCGΔ*pe18*_2 | BCGΔ*pe18*_3 | BCGΔ*pe18*_4 |
|---|---|---|---|---|---|---|---|---|---|---|---|---|
| 2-methylcitrate | 401245.4397 | 303079.3632 | 392086.7317 | 453734.6055 | 13850740.1 | 8575815.308 | 11143522.46 | 8717770.437 | 17564667.14 | 17584694.29 | 14250179.32 | 17342504.32 |
| Citrulline | 97102.05423 | 128190.2933 | 82082.56649 | 129031.6629 | 6082878.769 | 4261537.47 | 4742130.268 | 4656924.952 | 9579413.255 | 5794283.718 | 5554840.735 | 8403061.137 |
| Adenosine 5'-diphosphate (ADP) | 378323.2436 | 503121.1916 | 379543.4121 | 411862.2418 | 454716.9533 | 372587.8604 | 172244.3132 | 244547.0491 | 300716.4026 | 671351.7392 | 350092.3163 | 1560790.557 |
| Malic acid | 48260952.69 | 21016451.16 | 47164534.84 | 38567746.88 | 81087057 | 119751299 | 106123109.1 | 81487057.28 | 66042565.93 | 102806137.9 | 198809775.9 | 71916856.3 |
| Succinate | 20194428.97 | 15318342.21 | 32486123.49 | 19612287.85 | 216229740.3 | 144740857.1 | 212808570.8 | 165026018.3 | 129042382.9 | 152274495.8 | 155691916.6 | 118825196.9 |
| Lactate | 834499757.7 | 658399698.8 | 795444443.7 | 669463945.4 | 1048826437 | 714518879 | 1266963661 | 819839606.2 | 677868500.6 | 480293765.2 | 486393922.1 | 562765838 |
| Pyruvate | 16128047.32 | 14879933.82 | 25237042.33 | 22746841.57 | 16318177.68 | 20722281.28 | 16718558.01 | 16494640.14 | 17975018.23 | 11641086.34 | 12156073.25 | 19082125.79 |
| alpha-Ketoglutaric acid (alpha-KG) | 1210845.299 | 1040203.676 | 1305692.61 | 1006386.54 | 9733459.987 | 7005530.817 | 8769426.199 | 7734544.994 | 8703162.381 | 7843439.545 | 7067468.695 | 6865314.111 |
| Isocitric acid | 2891611.047 | 2851816.456 | 2901590.825 | 2661868.864 | 6782849.293 | 4939959.764 | 8134935.19 | 6290105.526 | 4624584.912 | 6610850.832 | 5623783.497 | 6019268.092 |
| Fumaric acid | 1509193.973 | 1198232.562 | 2318500.508 | 1524724.946 | 7100770.378 | 5510062.338 | 10470958.42 | 7160146.623 | 4476847.132 | 8250214.932 | 7209278.5 | 4739741.857 |
| Nicotinamide | 120782 | 937882. | 119929 | 110356 | 359677 | 210072 | 331378 | 262264 | 255536 | 315114 | 257599 | 264074 |

| Metabolites | PBS_1 | PBS_2 | PBS_3 | PBS_4 | BCG_1 | BCG_2 | BCG_3 | BCG_4 | BCGΔ*pe18*_1 | BCGΔ*pe18*_2 | BCGΔ*pe18*_3 | BCGΔ*pe18*_4 |
|---|---|---|---|---|---|---|---|---|---|---|---|---|
| adenine dinucleotide (NAD) | 1.033 | 0218 | 9.79 | 2.911 | 1.071 | 4.382 | 6.452 | 5.057 | 7.661 | 8.52 | 7.275 | 7.044 |
| Glucose | 5229170.992 | 4183664.979 | 8368353.282 | 6016796.427 | 8880878.945 | 8453295.493 | 7785932.909 | 8083357.086 | 7710053.785 | 4173089.389 | 3238113.928 | 6683050.574 |
| Citric acid | 2891611.047 | 2851816.456 | 2901590.825 | 2661868.864 | 6782849.293 | 4939959.764 | 8134935.19 | 6290105.526 | 4624584.912 | 6610850.832 | 5623783.497 | 6019268.092 |
| Acetyl-CoA | 136468.8842 | 127547.8736 | 162614.4215 | 140400.4804 | 540226.3451 | 359507.1351 | 557106.6155 | 467532.4555 | 405350.5667 | 431038.5842 | 429723.9056 | 404818.7837 |
| Glutamine | 5242310.898 | 4896585.405 | 9228495.134 | 7059804.184 | 26367063.72 | 20726371.32 | 23607861.22 | 21230388.86 | 23177556.37 | 12984323.27 | 15650037.81 | 19638355.95 |
| Itaconic acid | 36423784.15 | 24293593.32 | 59315185.46 | 37642727.23 | 728452365.5 | 453193811.1 | 737099582.8 | 551101366.1 | 426261451.5 | 577805304.2 | 612611096.8 | 368292521.3 |
| Desoxypipradrol | 1999451.365 | 20256356.17 | 43548106.8 | 31342000.55 | 2636683.311 | 954794.9096 | 3265538.299 | 583639.2978 | 3751776.112 | 5811495.286 | 2284611.859 | 156065755.2 |
| 1-(Tetrahydro-2H-thiopyran-4-yl)piperazine | 1379438.193 | 6980557.375 | 11679867.77 | 12760226.19 | 851289.5459 | 518383.2082 | 1650992.023 | 410143.3503 | 1362676.374 | 1902380.608 | 1048988.508 | 80472268.72 |
| Radicicol | 590606.7 | 314705.5444 | 819625.6186 | 457957.5826 | 1280903.231 | 1230344.193 | 1129995.509 | 1107751.73 | 200615567.7 | 1013535.014 | 1137136.476 | 1404817.356 |
| 6-Amino-1-hexanol | 1881147.971 | 19760807.02 | 38622045.58 | 22967240.05 | 3950347.615 | 1140883.251 | 6108370.819 | 1909473.882 | 5314446.079 | 9134870.766 | 1478344.333 | 149537163.8 |
| 6-Methoxymethylone | 898279.7253 | 43487535.19 | 60066043.46 | 53446927.79 | 3973312.921 | 1157191.652 | 10893466.91 | 2108514.848 | 8306499.337 | 9928947.161 | 5251630.198 | 177223901.4 |

| **Metabolites** | **PBS_1** | **PBS_2** | **PBS_3** | **PBS_4** | **BCG_1** | **BCG_2** | **BCG_3** | **BCG_4** | **BCGΔ*pe18*_1** | **BCGΔ*pe18*_2** | **BCGΔ*pe18*_3** | **BCGΔ*pe18*_4** |
|---|---|---|---|---|---|---|---|---|---|---|---|---|
| Mepanipyrim-2-hydroxypropyl | 130543.5686 | 2035872.153 | 4195896.016 | 3596870.189 | 250391.8082 | 79814.55751 | 528700.1124 | 59854.99496 | 568763.9159 | 783928.1053 | 330764.1673 | 18901409.61 |
| Benzyl-L-valine methyl ester | 4075786.772 | 54071386.32 | 70199057.07 | 79031225.73 | 8971840.215 | 2646637.8 | 11661409.7 | 2814672.668 | 10079888.66 | 13198703.25 | 7142725.1 | 175673872.3 |
| (.+/-.)-N-Propyl-3,4-methylenedioxyamphetamine | 17977135.86 | 227157032.2 | 293601910.8 | 329176804.9 | 38547286.35 | 13042695.76 | 53090065.16 | 12857607.45 | 41150039.24 | 57580541.88 | 29425725.26 | 686997034 |
| M339T450 | 588138.4371 | 2396067.49 | 3948149.681 | 2780319.863 | 551848.3159 | 330314.4334 | 689173.8368 | 246765.0785 | 607611.0089 | 904573.0192 | 640408.1979 | 19348733.79 |
| N-Nervonoyl Glutamine | 0 | 338475.6136 | 772671.0892 | 170116.6512 | 23765.74133 | 11043.37894 | 411383.5939 | 0 | 0 | 27544.42841 | 83944.79335 | 3570604.322 |
| Nadolol | 31858.2404 | 811783.3476 | 2705988.133 | 871187.1914 | 57778.54037 | 47014.80854 | 0 | 0 | 0 | 0 | 0 | 4164750.501 |
| Dianicline | 167520.116 | 849260.3844 | 1637952.561 | 1360271.327 | 274013.4619 | 249292.7355 | 255346.5754 | 303221.9796 | 346665.3252 | 133901.1546 | 255468.3912 | 13825542.99 |
| N-Methylanabasine | 1557538.744 | 19298717.52 | 44222370.09 | 34374196.91 | 1590765.569 | 1353562.579 | 6213489.687 | 1659156.032 | 5148647.436 | 2694762.596 | 1569519.158 | 93231779.46 |
| 3-Piperidin-4-ylpropanoic acid | 2394460.774 | 22868897.5 | 27994460.76 | 34744593.28 | 8895551.472 | 1063919.22 | 6199355.806 | 1981687.834 | 6808964.733 | 9790310.246 | 8566889.426 | 121097439.3 |
| 4'-Methyl-.alpha.-pyrrolidinohexanophenone | 68383.72969 | 814768.2657 | 945489.8106 | 686263.1328 | 370892.1401 | 276301.6787 | 305153.9195 | 297202.26 | 393085.193 | 469052.8882 | 325705.597 | 9023005.163 |
| Cyclohexylaminopropanesulfonic acid | 503213.6609 | 5853442.172 | 5631866.132 | 8444955.223 | 1608189.016 | 293782.2482 | 1308793.13 | 648722.8666 | 1266344.871 | 1334380.205 | 979777.4551 | 22613742.64 |

| **Metabolites** | **PBS_1** | **PBS_2** | **PBS_3** | **PBS_4** | **BCG_1** | **BCG_2** | **BCG_3** | **BCG_4** | **BCGΔ*pe18*_1** | **BCGΔ*pe18*_2** | **BCGΔ*pe18*_3** | **BCGΔ*pe18*_4** |
|---|---|---|---|---|---|---|---|---|---|---|---|---|
| PI 40:10 | 0 | 0 | 0 | 34579.09797 | 90043.96607 | 169612.0559 | 99598.66783 | 73652.78717 | 2132366.143 | 1335164.786 | 1577796.8 | 1151016.681 |
| Huperzine A | 138208.302 | 2678170.688 | 6869229.82 | 5465544.972 | 434155.4847 | 277066.2421 | 518037.578 | 0 | 456934.8021 | 1541065.3 | 0 | 5965811.915 |
| Hyaluronic_acid | 52537.75597 | 51776.55738 | 0 | 94151.72382 | 628174.361 | 650663.2539 | 639547.9537 | 568016.3407 | 3674894.026 | 3084729.432 | 3499287.175 | 10151331.4 |
| (2R)-3-(tert-Butylamino)propane-1,2-diol | 295046.3792 | 4600411.901 | 11243513.9 | 6084132.707 | 607776.0046 | 295746.3992 | 1547560.086 | 159503.3848 | 1035271.574 | 1831106.118 | 110569.4034 | 20462889.15 |
| 4-Piperidinecarboxamide, 4-amino-N-[(1S)-1-(4-chlorophenyl)-3-hydroxypropyl]-1-(7H-pyrrolo[2,3-d]pyrimidin-4-yl)- | 275111.0022 | 258556.07 | 79190.35894 | 251988.9565 | 3251435.575 | 2581468.843 | 3086900.945 | 2434157.95 | 16968526.01 | 12794599.54 | 16211148.44 | 15679558.68 |
| 4-(7-Hydroxy-6-methoxy-2,3-dimethyl-1,2,3,4-tetrahydronaphthalen-1-yl)benzene-1,2-diol | 5301351.671 | 16047004.27 | 30177665.74 | 20639631.83 | 3588378.713 | 2516019.084 | 5605059.339 | 1470044.903 | 5411796.682 | 7475385.671 | 5625454.104 | 43744256.48 |
| N-Methyl-N-[2-oxo-2-(1-pyrrolidinyl)ethyl]amine | 573468.4195 | 3742626.233 | 5730807.443 | 4061107.671 | 535006.4459 | 313597.6017 | 952380.9719 | 262109.1783 | 627457.8372 | 1056879.727 | 272203.4037 | 7754055.038 |

| Metabolites | PBS_1 | PBS_2 | PBS_3 | PBS_4 | BCG_1 | BCG_2 | BCG_3 | BCG_4 | BCGΔ*pe18*_1 | BCGΔ*pe18*_2 | BCGΔ*pe18*_3 | BCGΔ*pe18*_4 |
|---|---|---|---|---|---|---|---|---|---|---|---|---|
| Nabilone | 1368939.984 | 1234383.407 | 2189186.843 | 3347809.916 | 6125001.634 | 4067211.14 | 5294726.944 | 4415552.792 | 26235760.96 | 22809395.29 | 25511581.79 | 17315735.23 |
| 3-Benzylpiperidine | 3269356.619 | 3524507.044 | 2809725.766 | 3048561.159 | 3336487.511 | 3336742.067 | 3169633.93 | 3276832.701 | 3339436.449 | 3273709.003 | 3239491.908 | 66585177.24 |
| Decanoic acid, 3-amino- | 1101839.901 | 9508872.517 | 9515723.244 | 14579482.41 | 3323553.109 | 387444.55 | 1776337.085 | 389209.5713 | 2178871.092 | 5233739.67 | 2720076.179 | 20433930.83 |
| Lifibrol | 1905775.884 | 2029371.389 | 2793471.16 | 1277877.056 | 535339.3936 | 378711.7228 | 517353.1347 | 168482.2332 | 457363.9351 | 1445025.366 | 517956.8829 | 5784985.119 |
| 1-Ethyl-5-methyl[1,2,4]triazolo[4,3-a]quinolin-8-yl methyl ether | 533686.4948 | 2872246.043 | 10012397.77 | 9147172.743 | 347407.7186 | 146497.932 | 272756.6732 | 64229.7236 | 529507.8624 | 1144567.646 | 74148.63435 | 7513437.447 |
| 2-(4-Chlorobenzyl)-1H-benzimidazole | 1441663.702 | 6951829.545 | 13849546.45 | 9293422.833 | 1393150.434 | 541397.005 | 2098835.509 | 451344.8592 | 1773731.36 | 2748348.442 | 1885756.01 | 16333605.06 |
| DL-beta-Leucine | 8517840.513 | 58232897.49 | 77919802.93 | 82171973.7 | 11795888.38 | 3924843.026 | 17863875.72 | 4909259.647 | 19593957.53 | 28533370.81 | 22097283.68 | 110865654.5 |
| Furmecyclox | 2564685.364 | 21307580.75 | 30886824.47 | 32152481.86 | 5811453.069 | 1197576.74 | 5954724.149 | 928252.2589 | 4950484.352 | 8801519.712 | 4197119.785 | 39322045.05 |
| 2-(Piperidin-1-yl)benzo[d]oxazole | 391362.2667 | 7166420.406 | 7309563.298 | 7990279.804 | 106002.1981 | 65137.22836 | 725128.469 | 179517.6106 | 417781.7323 | 360775.7438 | 442564.2455 | 65404029.08 |

| Metabolites | PBS_1 | PBS_2 | PBS_3 | PBS_4 | BCG_1 | BCG_2 | BCG_3 | BCG_4 | BCGΔ*pe18*_1 | BCGΔ*pe18*_2 | BCGΔ*pe18*_3 | BCGΔ*pe18*_4 |
|---|---|---|---|---|---|---|---|---|---|---|---|---|
| (R)-4-((7R,8S,9S,10R,13R,14S,17R)-7-hydroxy-10,13-dimethyl-3-oxo-2,3,6,7,8,9,10,11,12,13,14,15,16,17-tetradecahydro-1H-cyclopenta[a]phenanthren-17-yl)pentanoic acid | 176791.0365 | 108337.7095 | 144957.481 | 200891.9567 | 373865.7683 | 285005.726 | 621721.5183 | 323705.8844 | 3737830.383 | 3527495.242 | 3611907.127 | 2583760.925 |
| 2-Hydroxy-3-(2-hydroxy-3-methylbut-3-enyl)-4-methoxy-6-(2-phenylethyl)benzoic acid | 2327681.662 | 9711614.258 | 17092440.97 | 10913200.16 | 2322700.716 | 736061.6857 | 2232782.337 | 347322.4199 | 1942970.079 | 3506682.951 | 2506140.976 | 26885586.04 |
| Ampkinone | 3309.203931 | 3731.277841 | 5318.568734 | 0 | 96701.42522 | 68806.10159 | 84237.22202 | 60201.15248 | 552569.999 | 411887.1727 | 541171.264 | 463788.8034 |
| 3-(4-Methyl-1-piperazinyl)propanohydrazide | 34794.78406 | 1465177.318 | 2899206.787 | 1976413.006 | 337351.0183 | 72296.19395 | 382330.9465 | 14990.63472 | 344356.0272 | 470692.3112 | 32783.02234 | 4671483.486 |
| 5-Cyclohexyl-1H-1,2,3,4-tetrazole | 949782.3964 | 1249036.582 | 2477856.521 | 1519339.251 | 733427.0172 | 726936.6605 | 272384.0763 | 71670.57236 | 263974.944 | 499326.058 | 757801.93 | 5643375.486 |
| Acetylhydrazine | 2137027.398 | 20148908.45 | 7623980.665 | 13121300.25 | 2176173.67 | 390919.1867 | 4131667.048 | 2839979.054 | 3147189.014 | 3576497.166 | 1254536.506 | 44514549.82 |
| N,N'-Diacetylchitob | 0 | 0 | 0 | 4711.10 | 117196. | 99406.4 | 228727. | 112856. | 611930. | 539431. | 107605 | 500003. |

| Metabolites | PBS_1 | PBS_2 | PBS_3 | PBS_4 | BCG_1 | BCG_2 | BCG_3 | BCG_4 | BCGΔ*pe18*_1 | BCGΔ*pe18*_2 | BCGΔ*pe18*_3 | BCGΔ*pe18*_4 |
|---|---|---|---|---|---|---|---|---|---|---|---|---|
| iose | | | | 7437 | 3167 | 6006 | 0912 | 8262 | 1505 | 5281 | 9.121 | 0281 |
| Butylphosphonic acid | 1414940.163 | 936840.1225 | 859493.8646 | 1119794.526 | 751637.0743 | 909308.2695 | 677886.6771 | 698655.165 | 853333.9576 | 1121064.888 | 818480.7821 | 16388522.8 |
| 2-(4-Chlorophenyl)-1H-benzimidazole | 7523826.491 | 37564912.51 | 77223642.12 | 50779010.6 | 6670239.596 | 2977802 | 11433841.07 | 2745683.998 | 10916853.61 | 14730707.03 | 14159593.81 | 81750317.9 |
| N,N-Dimethylhistidine | 1744089.029 | 7843095.453 | 16156661.68 | 13767240.5 | 2708016.261 | 2085881.554 | 2343807.845 | 1856204.194 | 3041347.401 | 2956967.565 | 2633955.883 | 21853944.05 |
| Ethyldiethanolamine | 959336.668 | 6942225.228 | 17864800.56 | 11382681.09 | 1407067.374 | 149284.4744 | 3171041.766 | 804754.0359 | 1816060.307 | 2614691.567 | 984513.7666 | 48197490.19 |
| Isopropylurea | 1007093.749 | 9626346.355 | 13838609.86 | 12442250.42 | 705266.3137 | 726998.4221 | 2633938.411 | 470901.3568 | 2188336.551 | 2271175.58 | 477560.2646 | 18982278.01 |
| Dicyclohexylamine | 1390096.501 | 4138597.617 | 5159846.504 | 4680174.299 | 930720.7587 | 826640.7763 | 1163376.277 | 696585.0542 | 984029.4268 | 1114615.852 | 1016510.992 | 8495495.78 |
| Tetrahydrozoline | 285416.5111 | 3039680.073 | 5267951.852 | 3992833.925 | 304907.8402 | 132150.8833 | 375814.1839 | 43734.16416 | 435698.9288 | 1092318.548 | 338134.6954 | 21557552.9 |
| PI 17:2_18:2 | 1893417.438 | 2468249.806 | 609415.5825 | 1516466.409 | 190775.1156 | 99732.81645 | 239162.3652 | 276104.6981 | 5745411.687 | 397657.3373 | 100408.3835 | 164427.0537 |
| (S)-1-(2-Chloroacetyl)pyrrolidine-2-carbonitrile | 495505.6472 | 1049543.735 | 1344348.663 | 1252464.725 | 286171.233 | 275593.8753 | 382738.4534 | 657918.0087 | 416889.1091 | 400166.0875 | 1159202.013 | 2842835.34 |
| ST 24:1;O3;T/22:5 | 1129399.227 | 658876.5004 | 813199.6429 | 1106261.433 | 398683.3126 | 610045.1225 | 354445.4227 | 207436.1381 | 2235590.968 | 619228.1456 | 1617828.989 | 1088710.14 |
| (+)-Muscarine cation | 401059.1551 | 1960803.001 | 3061297.157 | 2916095.007 | 1609242.841 | 461518.2481 | 954298.9649 | 177991.5733 | 984073.0646 | 2579106.041 | 878004.8238 | 4811209.829 |

| Metabolites | PBS_1 | PBS_2 | PBS_3 | PBS_4 | BCG_1 | BCG_2 | BCG_3 | BCG_4 | BCGΔ*pe18*_1 | BCGΔ*pe18*_2 | BCGΔ*pe18*_3 | BCGΔ*pe18*_4 |
|---|---|---|---|---|---|---|---|---|---|---|---|---|
| PC O-35:2 | 2719066.617 | 1740615.967 | 5946489.052 | 5775514.013 | 1283116.242 | 251395.2518 | 1619607.251 | 788945.5554 | 5570746.288 | 4317335.186 | 2521715.862 | 2082555.084 |
| 1H-Indole-3-carboxylic acid, 1-(cyclohexylmethyl)- | 193309.5902 | 205634.2311 | 205350.5311 | 195126.4625 | 10134590.33 | 7197104.826 | 12462171.8 | 7164379.955 | 30865281.19 | 20909153.33 | 28647244.62 | 15251561.73 |
| 4,4-Dimethyl-4,5-dihydro-1,3-oxazol-2-amine | 455166.3445 | 1048805.118 | 1518204.321 | 2562844.61 | 1051168.501 | 594823.2421 | 2368001.631 | 895859.0318 | 676324.8372 | 1798417.903 | 2244613.483 | 9901304.186 |
| Cer 12:1;2O/2:0 | 9384014.636 | 30243840.54 | 42388898.85 | 37576631.27 | 17592456.56 | 14799371.53 | 18254019.29 | 8307330.815 | 11975408.85 | 13497121.72 | 11860499.15 | 117435154.9 |
| Swainsonine | 447059.2704 | 5092251.377 | 5672982.398 | 8285198.632 | 3203956.273 | 525368.1153 | 1148151.601 | 676807.2209 | 933979.8512 | 890290.0499 | 1143375.553 | 11273876.73 |
| N-(5-Methyl-3-oxohexyl)alanine | 1383515.16 | 4271486.074 | 7961655.614 | 4154620.24 | 1330649.44 | 1339647.899 | 1801781.708 | 1330443.874 | 2001230.393 | 2133604.086 | 1319473.876 | 19891373.1 |
| 1-hexylamine | 17767180.16 | 4572290.732 | 5264272.292 | 3369466.286 | 28642.37739 | 3718248.516 | 4933420.872 | 4323027.488 | 8227248.071 | 15401734.57 | 5280813.853 | 5409107.026 |
| Bis(2,2,6,6-tetramethyl-4-piperidyl) sebacate | 652895.1048 | 1050585.282 | 1422066.282 | 682360.4278 | 731166.85 | 998104.1054 | 974445.3965 | 689685.7801 | 660247.5472 | 610385.5898 | 2951569.906 | 8958451.801 |
| PC 8:0_30:3 | 29153141.92 | 6626836.195 | 12656105.64 | 19179641.1 | 6301763.377 | 1891516.863 | 9506768.072 | 6912606.998 | 18936617.55 | 30881691.81 | 9389639.24 | 4882971.447 |
| 3',4'-Dihydroxy-.beta.-naphthoflavone | 0 | 0 | 0 | 0 | 1476887.638 | 1265191.203 | 1004605.111 | 1150296.234 | 3709926.696 | 2516619.528 | 2962822.313 | 3934732.468 |

| Metabolites | PBS_1 | PBS_2 | PBS_3 | PBS_4 | BCG_1 | BCG_2 | BCG_3 | BCG_4 | BCGΔ*pe18*_1 | BCGΔ*pe18*_2 | BCGΔ*pe18*_3 | BCGΔ*pe18*_4 |
|---|---|---|---|---|---|---|---|---|---|---|---|---|
| 6-(Ethylsulfonyl)-2H-1,4-benzoxazin-3(4H)-one | 593260.1765 | 516092.8632 | 631449.2573 | 444291.2642 | 1593729.658 | 1384480.688 | 1363661.098 | 1425607.78 | 5687143.793 | 5787864.293 | 4859577.447 | 1442477.936 |
| Cyclo(alanylleucyl) | 3172584.393 | 12418303.43 | 25521980.86 | 45491360.94 | 3642543.165 | 1082401.421 | 8436262.059 | 3235182.922 | 6501811.523 | 8985305.177 | 1347644.129 | 35739720.56 |
| Cholestan-3-One | 70492613.79 | 16350027.52 | 20124622.7 | 62710009.26 | 9487027.81 | 8346587.953 | 9667075.476 | 9672425.01 | 54093995.01 | 11351840.64 | 9083180.225 | 16760359.06 |
| N-(2-Hydroxyethyl) valine | 230576.2461 | 656523.298 | 3905204.758 | 2159188.356 | 256891.279 | 126608.3968 | 409693.3626 | 225740.208 | 219808.7681 | 312799.62 | 182113.3609 | 4966893.006 |
| 2-Aminobiphenyl | 900408.113 | 4580447.63 | 10093755.04 | 7115330.657 | 836542.0331 | 884900.0702 | 1635955.715 | 914445.1893 | 1139441 | 1194343.255 | 792022.6468 | 7869982.074 |
| 5-Pyrimidinecarboxamide, 2-[(2,4-dichlorophenyl)amino]-N-[(tetrahydro-2H-pyran-4-yl)methyl]-4-(trifluoromethyl)- | 45272.20504 | 171553.3739 | 175928.6178 | 160209.2625 | 424968.3291 | 239761.5134 | 102512.1984 | 100903.7596 | 347210.5995 | 1194909.568 | 335587.9289 | 587819.9964 |
| 2,4-Quinazolinediamine, N2,N4-bis(phenylmethyl)- | 2239797.483 | 15507639.28 | 14473978.5 | 6239079.257 | 1136399.39 | 644393.0026 | 1587243.663 | 365108.6219 | 1267283.268 | 2059206.379 | 1307912.379 | 14588828.95 |

| Metabolites | PBS_1 | PBS_2 | PBS_3 | PBS_4 | BCG_1 | BCG_2 | BCG_3 | BCG_4 | BCGΔ*pe18*_1 | BCGΔ*pe18*_2 | BCGΔ*pe18*_3 | BCGΔ*pe18*_4 |
|---|---|---|---|---|---|---|---|---|---|---|---|---|
| (6-Chloro-2-hydroxy-4-phenyl-3-quinolinyl)acetic acid | 345104.2499 | 327791.3313 | 797552.5207 | 456190.0489 | 4097301.211 | 3073705.579 | 5315898.861 | 4459260.164 | 9453848.573 | 5170528.091 | 6598863.474 | 19826024.07 |
| SL 23:3;O/26:6 | 4767474.315 | 4400805.704 | 3419336.028 | 6066878.584 | 3302819.396 | 404479.9872 | 1928158.593 | 1782206.218 | 6678100.256 | 5256907.954 | 3615514.992 | 2171585.616 |
| 5-Isopropyl-5-methylhydantoin | 2257692.391 | 6900514.279 | 18125279.57 | 19656685.99 | 1501892.374 | 2959921.171 | 6398448.482 | 2037277.903 | 4321346.723 | 5276331.087 | 1068594.517 | 23693211.34 |
| 5-Cyclohexyl-1,3,4-thiadiazol-2-amine | 1534952.216 | 531007.6921 | 3048664.931 | 664242.1253 | 4675563.466 | 11673717.07 | 20954760.05 | 20203722.32 | 38377034.04 | 20086074.18 | 34007735.11 | 38878020.24 |
| 2-Chloro-5-fluorobenzoic acid | 6914855.459 | 3726776.191 | 11141839.29 | 3476311.37 | 29078195.88 | 32774663.41 | 50234844.99 | 26166131.16 | 66163191.11 | 80126465.62 | 103255597.1 | 62959039.91 |
| Ethyl 5-(2-chloroacetyl)-2,4-dimethyl-1H-pyrrole-3-carboxylate | 458656.724 | 205399.488 | 930324.7742 | 233517.4588 | 4919180.016 | 5417272.394 | 13536262.65 | 11533221.08 | 24339663.89 | 11439343.53 | 20743336.26 | 23039622.23 |
| 2-(2-Naphthyl)-1,3-benzoxazol-5-amine | 296545.5482 | 162947.3333 | 566500.1084 | 135083.8388 | 3157831.113 | 3480102.122 | 8590485.98 | 7286782.33 | 15650740.07 | 7199465.232 | 13211864.74 | 14729242.47 |
| PE O-16:1_20:3 | 1017564.645 | 830612.9444 | 926332.5816 | 792897.0778 | 575717.3029 | 0 | 585667.2799 | 413819.3135 | 1000452.634 | 1088804.016 | 836193.9499 | 702067.3903 |
| Damnacanthal | 138860.345 | 126389.7212 | 441160.9675 | 67047.74572 | 2203088.95 | 2629993.579 | 6217813.726 | 4159911.513 | 8052532.899 | 7167971.803 | 7373427.59 | 11153369.63 |

| **Metabolites** | **PBS_1** | **PBS_2** | **PBS_3** | **PBS_4** | **BCG_1** | **BCG_2** | **BCG_3** | **BCG_4** | **BCGΔ*pe18*_1** | **BCGΔ*pe18*_2** | **BCGΔ*pe18*_3** | **BCGΔ*pe18*_4** |
|---|---|---|---|---|---|---|---|---|---|---|---|---|
| 2,8,9-Trimethyl-2,5,8,9-tetraaza-1-phosphabicyclo[3.3.3]undecane | 16753.91921 | 1232816.111 | 3055486.879 | 2042589.227 | 114664.3559 | 0 | 121680.9196 | 0 | 54009.88234 | 243741.4694 | 0 | 5817547.069 |
| PE O-26:5_13:0 | 9043811.575 | 5824500.6 | 6188843.65 | 5503108.754 | 4796480.571 | 227115.2541 | 4013781.485 | 4682063.674 | 4551640.997 | 10708534.25 | 13354241.08 | 4189355.724 |
| 3,4-Dicaffeoyl-1,5-quinolactone | 568759.8011 | 2539070.27 | 6457141.887 | 3843458.597 | 526426.0711 | 286198.1262 | 859567.2336 | 33725.48921 | 530975.2955 | 829271.1698 | 178808.567 | 7544574.103 |
| PI(18:0/20:4(5Z,8Z,11Z,14Z)) | 699483.7885 | 749367.6987 | 517359.2105 | 716831.6891 | 686352.6294 | 758363.4331 | 383454.7553 | 723640.1005 | 710420.5636 | 9345638.3 | 737200.4907 | 656782.5583 |
| Desmetryn | 466421.502 | 148064.6041 | 541925.4364 | 166899.773 | 1918657.879 | 2147316.988 | 3912589.424 | 4406893.433 | 7552205.898 | 4318804.818 | 6578719.14 | 8595512.925 |
| 9-Phenyl-1-(2,4,6-trihydroxyphenyl)nonan-1-one | 3011077.143 | 5242082.719 | 8938174.823 | 5630474.094 | 700259.234 | 623811.4283 | 956514.4765 | 335150.0009 | 960656.3709 | 863975.5116 | 1227855.742 | 11238226.9 |
| 7-Methoxy-1,2,3,4-tetrahydroacridin-9-amine | 337099.1074 | 2142588.966 | 6138116.618 | 5686275.509 | 505771.2131 | 312731.8191 | 658657.512 | 156502.0888 | 838835.0863 | 1331568.617 | 230706.4088 | 4638699.033 |
| PI 18:0_18:1 | 1946623.99 | 5264257.688 | 7962304.129 | 2632643.707 | 4964503.729 | 110729.2318 | 1802891.13 | 776555.9007 | 8093951.864 | 4321952.694 | 4295999.451 | 1084563.929 |
| PC 9:0_24:1 | 40153533.37 | 26065663.49 | 66068029.21 | 48464734.06 | 37121490.53 | 6486540.71 | 27615236.16 | 10718438.64 | 46552976.32 | 48281842.56 | 76118741.75 | 12803166.32 |
| 2-(Naphthalen-1-yl)-1H-1,3-benzodiazo | 395753.5772 | 415691.0925 | 101483.7767 | 283338.4248 | 2255706.305 | 991326.6335 | 7858104.604 | 5843078.148 | 13448375.72 | 12731504.67 | 5962311.09 | 3575649.914 |

| **Metabolites** | **PBS_1** | **PBS_2** | **PBS_3** | **PBS_4** | **BCG_1** | **BCG_2** | **BCG_3** | **BCG_4** | **BCGΔ *pe18*_1** | **BCGΔ *pe18*_2** | **BCGΔ *pe18*_3** | **BCGΔ*pe18*_4** |
|---|---|---|---|---|---|---|---|---|---|---|---|---|
| le | | | | | | | | | | | | |
| .beta.-Alanylleucine | 385540.5353 | 2323952.588 | 3066243.059 | 4468383.837 | 1650109.604 | 141946.2097 | 1029864.366 | 264728.3741 | 958641.6081 | 3010429.622 | 1602128.381 | 3939618.787 |
| D-Tyrosine tert-butyl ester | 202719.907 | 2948651.044 | 2622463.138 | 3282734.258 | 1137527.935 | 112701.7091 | 436418.1195 | 178684.8162 | 333030.5659 | 384384.3975 | 277412.674 | 4097170.035 |
| [2-Amino-1-(2-chlorophenyl)ethyl]dimethylamine | 314445.5642 | 215453.5574 | 491206.1699 | 162128.7223 | 2213255.617 | 2601404.789 | 5636336.324 | 5123566.218 | 9133677.127 | 4779965.088 | 8224244.393 | 10234334.78 |
| (3-Carboxypropyl)trimethylammonium cation | 2623555.788 | 15543148.96 | 25036529.44 | 37360867.23 | 11603443.97 | 1009148.555 | 5573411.015 | 2083879.912 | 5897207.27 | 13807827.07 | 7634919.859 | 47892293.96 |
| Sinapoylcholine | 74886.15388 | 1023425.204 | 2514074.59 | 1903855.679 | 81878.24059 | 89758.4649 | 256875.2037 | 57761.41448 | 194637.9835 | 407356.4779 | 70949.34073 | 3697749.062 |
| Triphenyl(propyl)phosphonium cation | 51206.93245 | 79649.67772 | 45821.24645 | 69064.22396 | 1202786.825 | 712505.8113 | 806902.7335 | 662918.463 | 2072367.952 | 1381805.748 | 1439171.81 | 2501485.663 |
| PC O-31:0 | 9873042.97 | 26689192.81 | 10239430.4 | 10333872.85 | 8320444.58 | 24820.90664 | 15347134.74 | 7919195.221 | 13368491.49 | 13196815.73 | 35773409.79 | 7101534.722 |
| ST 24:1;O4;T/20:1 | 1951021.169 | 1364673.199 | 2404312.37 | 4016459.214 | 1872023.667 | 164092.2095 | 1587462.515 | 1033985.508 | 1949344.304 | 5246501.629 | 1712886.542 | 801320.0538 |
| 4-(4-Amino-3-nitrophenyl)-1(2H)-phthalazinone | 869104.5956 | 253321.0515 | 1836145.254 | 291107.6225 | 11880541.67 | 11929768.31 | 27820028.89 | 23719911.93 | 46204918.53 | 22476189.09 | 37065076.25 | 40083029.16 |
| 1'-Naphthoyl-2-met | 190479. | 103069 | 207468 | 127777 | 141755. | 46764.4 | 224566. | 41198.2 | 165796. | 331319. | 184520. | 293182 |

| Metabolites | PBS_1 | PBS_2 | PBS_3 | PBS_4 | BCG_1 | BCG_2 | BCG_3 | BCG_4 | BCGΔ*pe18*_1 | BCGΔ*pe18*_2 | BCGΔ*pe18*_3 | BCGΔ*pe18*_4 |
|---|---|---|---|---|---|---|---|---|---|---|---|---|
| hylindole | 1181 | 3.11 | 7.799 | 1.593 | 1767 | 4664 | 8675 | 7636 | 1581 | 2825 | 1653 | 1.505 |
| NAGly 26:7/22:4 | 76833139.77 | 58422160.51 | 88350623.89 | 76571727.03 | 61758517.53 | 821418.7029 | 50296694.06 | 27495701.91 | 105573912.2 | 68099501.78 | 69781862.8 | 25485850.55 |
| PE-Cer 12:1;2O/12:0 | 8374.461092 | 0 | 0 | 0 | 793699.7873 | 482610.0549 | 1008731.894 | 853150.7836 | 1749735.666 | 1622501.595 | 2093840.108 | 904347.5568 |
| SM 19:2;2O/13:0 | 2112044.601 | 1792050.611 | 2414208.048 | 1956743.033 | 1585901.781 | 13381.24202 | 1342014.681 | 438281.2411 | 2212207.501 | 1689356.525 | 1824310.813 | 526021.626 |
| PE 8:0_28:1 | 21401939.22 | 9404426.326 | 25377763.47 | 17468571.94 | 13148604.18 | 2000691.753 | 10484381.35 | 5738573.752 | 19435760.31 | 14888692.48 | 13125289.42 | 10496273.24 |
| PC O-31:2 | 21505028.92 | 9023095.203 | 24698310.59 | 17355422.23 | 13157188.96 | 264950.3113 | 11057681.29 | 6656594.295 | 19344815.93 | 17925812.2 | 13404910.95 | 10299183.01 |
| SM 23:1;2O/13:0 | 30713872.73 | 17145301.88 | 36445470.76 | 37841880.38 | 25165136.38 | 814523.7437 | 15534819.13 | 9306070.678 | 37837242.06 | 24522699.93 | 16334188.2 | 25890853.22 |
| PC 9:0_28:2 | 7228393.286 | 3682352.397 | 9903102.522 | 12479506.85 | 4996415.993 | 1994099.621 | 6107888.344 | 1580704.221 | 12245436.14 | 10130818.57 | 9936735.825 | 1365756.607 |
| PC O-36:1 | 1165218.733 | 4105781.721 | 4268887.808 | 2400870.997 | 1471646.592 | 3505040.461 | 2174031.307 | 4883844.912 | 3209208.918 | 1942819.333 | 17532192.4 | 3285256.671 |
| Cymiazole | 676878.5425 | 563718.5294 | 1062718.88 | 642869.5893 | 7973303.682 | 5563565.998 | 6291638.989 | 5863870.916 | 13724502.93 | 11286391.87 | 10577386.04 | 11474026.8 |
| 2-Hydroxyatrazine | 907481.2366 | 555164.5483 | 1135458.965 | 506514.1788 | 2697922.415 | 3223985.284 | 5904276.52 | 5268840.354 | 9468361.9 | 5679378.581 | 8868246.303 | 10377068.02 |
| PC 8:0_26:0 | 30812726.24 | 10988740.8 | 66448257.24 | 26646184.69 | 22943303.07 | 741117.4188 | 26951043.35 | 31332513.66 | 50468645.6 | 50843724.04 | 20096792.57 | 36170310.52 |

| Metabolites | PBS_1 | PBS_2 | PBS_3 | PBS_4 | BCG_1 | BCG_2 | BCG_3 | BCG_4 | BCGΔ*pe18*_1 | BCGΔ*pe18*_2 | BCGΔ*pe18*_3 | BCGΔ*pe18*_4 |
|---|---|---|---|---|---|---|---|---|---|---|---|---|
| 2-Amino-4,6-dichloropyrimidine-5-carbaldehyde | 23724544.66 | 21960224.15 | 23370135.08 | 11832771.18 | 13593774.18 | 2438470.024 | 11929611.2 | 12462286.2 | 21915857.65 | 14090153.08 | 23214590.71 | 12509726.74 |
| NAGlySer 22:6/22:5 | 17303356.61 | 2656691.036 | 4804326.418 | 6961663.846 | 7573406.183 | 11195.12535 | 3992720.165 | 3873443.14 | 10897146.69 | 10834525.74 | 2802073.283 | 2858311.225 |
| Palmitoylcholine [M] | 2007692.265 | 2785471.115 | 2427779.992 | 2314210.427 | 915169.5506 | 855494.2918 | 1310774.952 | 1108839.522 | 1468063.112 | 1445091.111 | 1228925.857 | 6442100.858 |
| 4-Chloro-5-ethyl-7H-pyrrolo[2,3-d]pyrimidine | 3292665.48 | 3798014.757 | 2899547.367 | 3598850.582 | 2488092.119 | 2403192.137 | 2735439.265 | 2821374.037 | 3164062.341 | 2413627.131 | 3863934.844 | 9692496.825 |
| PE-Cer 12:2;2O/26:2;O | 1167572.672 | 1818988.45 | 1699039.221 | 2214502.952 | 632866.4034 | 0 | 980246.2838 | 1188386.463 | 2449260.332 | 976932.628 | 1048770.254 | 1476795.541 |
| 7-Keto-8-aminopelargonic acid | 27130417.72 | 22757473.9 | 25978476.18 | 155394291.1 | 4716341.09 | 34279933.4 | 180839967.3 | 4794580.541 | 139202207 | 56131526.62 | 7477156.557 | 189186944.5 |
| PC 8:0_28:1 | 38967089 | 16895474.36 | 63180038.35 | 32190292.66 | 25558051.86 | 518021.8925 | 22784341.87 | 13860708.1 | 41468403.96 | 27946172.72 | 23132177.28 | 34381491.14 |
| Piperazine-2-carbonitrile | 965204.9948 | 1278532.339 | 4465391.251 | 2064613.818 | 1012920.098 | 978635.984 | 659668.0105 | 669909.6634 | 610853.0365 | 743837.7786 | 549785.6966 | 12636890.08 |
| L-Glycine ether ester | 2599067.886 | 2588117.299 | 4067984.714 | 2557022.308 | 70351389.69 | 57271622.63 | 79023763.04 | 61553469.79 | 126836269 | 113879535.1 | 118150446.1 | 95966046.45 |
| 1-(1,2R-Dioctanoylphosphatidyl)inositol-3,4-bisphosphate | 3377.632283 | 9732.09672 | 0 | 0 | 626565.5239 | 585767.0657 | 501158.8326 | 513416.8358 | 993324.5998 | 819607.1843 | 855360.9665 | 1353925.879 |
| SM 39:5;3O | 890524 | 889977 | 511529 | 206979 | 250681 | 170982 | 225184 | 220435 | 305369 | 282072 | 627064 | 479224 |

| Metabolites | PBS_1 | PBS_2 | PBS_3 | PBS_4 | BCG_1 | BCG_2 | BCG_3 | BCG_4 | BCGΔ*pe18*_1 | BCGΔ*pe18*_2 | BCGΔ*pe18*_3 | BCGΔ*pe18*_4 |
|---|---|---|---|---|---|---|---|---|---|---|---|---|
| | 3.074 | 6.619 | 9.75 | 34.8 | 0.134 | 3.49 | 0.321 | 0.997 | 8.993 | 1.924 | 5.167 | 2.371 |
| Norharmane | 6755749.257 | 4015872.426 | 10723796.41 | 9679049.045 | 13493231.05 | 15837851.67 | 6899360.869 | 5631916.624 | 11158896.33 | 6418557.785 | 14534337.47 | 42182736.38 |
| N-Acetyl-D-lactosamine | 9707.054027 | 5321.174163 | 14385.92556 | 6906.05937 | 2754901.778 | 1767403.817 | 1786972.2 | 1512451.097 | 3597099.97 | 3019808.712 | 3370429.321 | 3266865.365 |
| 6-Hydroxy-5-nitronicotinic acid | 4588553.601 | 4139012.858 | 4710075.28 | 4468803.877 | 4452472.022 | 4481109.415 | 4449764.924 | 6084206.159 | 4535716.478 | 4892147.84 | 4772517.805 | 42009866.86 |
| Flunixine | 0 | 13352.94775 | 123436.6432 | 0 | 46637.65904 | 16764.39745 | 28492184.41 | 622453.5665 | 111767.3438 | 23769716.07 | 24983825.09 | 475370.1708 |
| aurapten | 136422092.5 | 189875679.2 | 219264949.8 | 179935745.1 | 82405120.72 | 95122945.31 | 99402307.62 | 83916296.57 | 87320290.82 | 91930906.08 | 88195824.12 | 347250929.6 |
| PE 19:2_19:2 | 89624245.96 | 49133575.43 | 77614251.53 | 69361471.4 | 66586848.15 | 1919613.585 | 65563440.45 | 41622077.84 | 95304069.24 | 85433277.47 | 62100497.97 | 61765539.81 |
| PE O-24:4_14:0 | 5625012.311 | 3096470.852 | 6299674.267 | 4722568.967 | 2726206.053 | 214888.2654 | 2252251.309 | 1979970.606 | 3666551.038 | 3465153.667 | 1990134.104 | 4498424.084 |
| 1,2-dioctadecanoyl-phosphatidyl-ethanolamine | 19759630.37 | 21808150.5 | 47079522.97 | 25443543.21 | 18524111.04 | 802598.0019 | 11973905.34 | 3824465.462 | 23045826.79 | 24043187.67 | 15497792 | 5123545.925 |
| Triptophenolide | 421576455.1 | 494057610.1 | 613959338.5 | 438428159.8 | 270587484.5 | 324433487.1 | 311458081.8 | 300962452.6 | 276225417.2 | 281869877.4 | 302649365.1 | 1114959441 |
| 1,3-Butadiene-1,1,3-tricarbonitrile, 2-amino-4-(3,4,5-trihydroxyphenyl)-, | 603080.2348 | 1749498.344 | 2517961.86 | 1898042.86 | 797464.4563 | 586322.9882 | 1231209.133 | 776517.8021 | 1015394.056 | 1042644.789 | 3870377.954 | 4208745.927 |

| Metabolites | PBS_1 | PBS_2 | PBS_3 | PBS_4 | BCG_1 | BCG_2 | BCG_3 | BCG_4 | BCGΔ *pe18*_1 | BCGΔ *pe18*_2 | BCGΔ *pe18*_3 | BCGΔ*pe18*_4 |
|---|---|---|---|---|---|---|---|---|---|---|---|---|
| (3Z)- | | | | | | | | | | | | |
| Canrenone | 1344985837 | 1483556654 | 1250483454 | 1364035550 | 862973517.6 | 946543436.7 | 824732417.4 | 1064143098 | 769462472.8 | 857089564.5 | 900964489.9 | 3504260292 |
| Methyl(oxolan-2-ylmethyl)amine | 25434377.27 | 20151182.99 | 42537013.15 | 19920.43461 | 69021437.58 | 5224199.484 | 5304029.652 | 75703162.15 | 85796712.93 | 74669565.84 | 5450844.337 | 89392832.91 |
| S-Acetyldihydrolipoamide-E | 318173.1594 | 221389.3567 | 607020.9715 | 365545.0323 | 4297299.402 | 4751772.402 | 3851678.314 | 4122983.333 | 7772558.264 | 5185401.097 | 6937576.918 | 8237591.815 |
| 2-{1-[(2-Methoxy-5-pyrimidinyl)methyl]-3-piperidinyl}-6-(2-thienyl)-4-pyrimidinol | 4398.441383 | 0 | 4106.786139 | 3312.889545 | 1739369.152 | 1176807.498 | 1187559.069 | 1007815.358 | 2407158.1 | 1976078.174 | 2219023.665 | 2188234.83 |
| phosphatidylglycerol (1,2-bis-cis-vaccenoyl) | 2571860.292 | 1237798.405 | 13142994.42 | 9219432.358 | 7801029.697 | 102975.092 | 4046961.516 | 963811.9772 | 12317096.1 | 5697690.477 | 2729100.521 | 891022.3392 |
| PG 16:0_20:1 | 3073370.208 | 4407765.905 | 3710547.308 | 3091538.258 | 3640292.712 | 246597.1157 | 2014226.09 | 1529240.55 | 8332599.984 | 3213495.41 | 3958035.96 | 3727390.529 |
| 2-[4-Amino-5-oxopentyl]guanidine | 142401.8386 | 644347.3104 | 1212890.934 | 1069928.223 | 59296.02845 | 198791.9322 | 487354.8131 | 138017.3942 | 112263.3136 | 79596.24744 | 121101.5651 | 2912898.008 |

| **Metabolites** | **PBS_1** | **PBS_2** | **PBS_3** | **PBS_4** | **BCG_1** | **BCG_2** | **BCG_3** | **BCG_4** | **BCGΔ*pe18*_1** | **BCGΔ*pe18*_2** | **BCGΔ*pe18*_3** | **BCGΔ*pe18*_4** |
|---|---|---|---|---|---|---|---|---|---|---|---|---|
| 4-(2-Chlorophenyl)-3,4-dihydro[1,3,5]triazino[1,2-a]benzimidazol-2-amine | 1049109.249 | 934838.6377 | 1595045.778 | 1070552.362 | 5475533.044 | 4906368.147 | 8266314.127 | 7614035.575 | 16332835.77 | 8459639.962 | 9816561.68 | 37809363.62 |
| N-Methyltetrahydro-2H-pyran-4-amine | 3162314.904 | 119203948.6 | 122662400.1 | 112339817.4 | 114496958.1 | 120107619.1 | 22974850.84 | 21587107.76 | 116667524.4 | 112425247.5 | 112853646.1 | 118622918.9 |
| (E)-2-Phenylethene sulfonic acid | 15034994.24 | 20243259.12 | 21973653.84 | 19389758.19 | 9422499.364 | 10232266.24 | 9422056.975 | 10206304.68 | 8944022.034 | 10243130.92 | 9889588.69 | 39177577.36 |
| 3-[(4-Amino-2-quinazolinyl)amino]phenol | 2515793.097 | 5690839.542 | 16419021.41 | 10345213.24 | 2836888.415 | 1994181.684 | 3506090.67 | 1809662.161 | 2703697.244 | 3292166.613 | 5064171.879 | 24210851.34 |
| PC 8:0_25:0 | 16029312.11 | 13620518.39 | 20593156.63 | 10880676.08 | 18018972.29 | 3817550.244 | 17147233.79 | 5934979.543 | 20175275.24 | 17637661.16 | 15373989.68 | 18812563.07 |
| PE O-22:2_16:3 | 21921577.33 | 12036392.24 | 17379755.65 | 17457757.56 | 17964411.99 | 1365930.667 | 18806203.27 | 13279203.57 | 24787837.36 | 24533656.74 | 14403630.43 | 20157867.82 |
| Pterostilbene | 2615683.197 | 2515882.651 | 4117299.991 | 3810308.183 | 304126.4148 | 180838.1248 | 206324.762 | 82963.16709 | 254441.7894 | 366120.1918 | 96805.04214 | 541980.4593 |
| 7H-[1,2,4]Triazolo[4,3-b][1,2,4]triazole-3,7-diamine | 5420574.896 | 4324076.469 | 5289901.348 | 5651302.748 | 29698284.21 | 23941483.15 | 31159613.06 | 31180627.98 | 58650960 | 40825884.63 | 50104117.26 | 34420980.25 |
| LNAPS 15:1/N-26:3 | 7202473.955 | 2106289.871 | 2393533.151 | 8405970.447 | 3115192.643 | 125016.299 | 1239894.487 | 186511.2243 | 10811279.38 | 931114.9105 | 1997511.688 | 161775.9197 |
| PG 18:1_20:4 | 110951722.1 | 97592602.6 | 55948733.87 | 71695338.03 | 67042530.7 | 40762040.25 | 123067140.6 | 48611610.43 | 101684045.6 | 172549848.9 | 112166939.3 | 54514798.62 |

| Metabolites | PBS_1 | PBS_2 | PBS_3 | PBS_4 | BCG_1 | BCG_2 | BCG_3 | BCG_4 | BCGΔ*pe18*_1 | BCGΔ*pe18*_2 | BCGΔ*pe18*_3 | BCGΔ*pe18*_4 |
|---|---|---|---|---|---|---|---|---|---|---|---|---|
| N-Cyclohexyl-N'-(4-pyridinylmethyl)urea | 1373046.102 | 2354339.702 | 5762645.959 | 2784453.043 | 2694805.146 | 2630516.905 | 2838575.009 | 1142651.629 | 3198217.767 | 3160883.193 | 2940265.47 | 9772448.044 |
| 2-Propanone, 1-(3-ethyl-5-methoxy-2(3H)-benzothiazolylidene)- | 163429.7723 | 182214.0651 | 201953.2682 | 185373.502 | 1829214.85 | 1728782.942 | 1663345.869 | 1588643.727 | 2895907.757 | 2422875.736 | 2682816.675 | 2825238.219 |
| 1-Pyrrolidineethanol, .alpha.-phenyl-.beta.-propyl- | 251634.6975 | 814646.266 | 2283457.107 | 1192513.204 | 419658.7235 | 358616.3591 | 235069.726 | 264101.9133 | 318875.2515 | 252685.8133 | 347609.6362 | 3586037.125 |
| Benzenesulfonic acid | 4729822.258 | 8517846.762 | 11262167.71 | 13233707.48 | 5239276.551 | 4671576.634 | 5712893.677 | 4720927.877 | 2934818.852 | 9162162.303 | 5803631.649 | 14563162.91 |
| 5-tert-Butyl-2,4-dihydroxybenzophenone | 33121653.73 | 34768495.89 | 34674110.2 | 142171.3642 | 156441.2665 | 148410.4818 | 36311385.68 | 34161203.34 | 35263869.53 | 34305846.15 | 33489198.95 | 32924252.37 |
| 8-(p-Chlorophenyl)theophylline | 31041.10501 | 36055.0596 | 31947.11193 | 56246.92532 | 13388152.7 | 10093708.54 | 9259660.518 | 9107942.775 | 18587365.91 | 14113282.93 | 14762081.73 | 17732745.22 |
| 1-Stearoyl-2-arachidonoyl-sn-glycero-3-phosphoserine | 4101183.956 | 6849516.641 | 7167372.782 | 9446168.37 | 8512295.587 | 1629407.145 | 5064545.008 | 17131427.31 | 20351654.53 | 15525953.49 | 9736253.905 | 6131436.952 |
| 2-Hydroxy-N-[3-(trifluoromethyl)phenyl]benzamide | 547451.0478 | 857794.5303 | 1150646.796 | 1031964.491 | 499973.7943 | 496808.9171 | 546888.4323 | 396108.7779 | 599336.4088 | 533426.2379 | 501694.1421 | 1445000.243 |
| TG 10:0_16:0_18:5 | 124279 | 103488 | 140673 | 126162 | 658428 | 360918 | 103316 | 700492 | 152268 | 133405 | 975581 | 710628 |

| Metabolites | PBS_1 | PBS_2 | PBS_3 | PBS_4 | BCG_1 | BCG_2 | BCG_3 | BCG_4 | BCGΔ *pe18*_1 | BCGΔ *pe18*_2 | BCGΔ *pe18*_3 | BCGΔ*pe18*_4 |
|---|---|---|---|---|---|---|---|---|---|---|---|---|
| | 20.7 | 64 | 58.67 | 17.29 | 4.382 | 3.885 | 85.47 | 3.225 | 04.82 | 06.03 | 7.72 | 9.612 |
| Glu-Ala | 75739.56091 | 71125.3734 | 126730.2541 | 53238.18811 | 1499286.916 | 945018.8641 | 1097925.575 | 873154.2639 | 2209718.217 | 1723432.042 | 1867950.783 | 2004874.303 |
| 2-(4-Nitrophenyl)-1,3-dioxo-5-isoindolinecarboxylic acid | 18835.23087 | 15298.81708 | 7766.162962 | 17876.66807 | 7155039.978 | 5832541.14 | 5057376.375 | 4933728.749 | 10067230.88 | 7103485.08 | 8036897.168 | 10185140.76 |
| LNAPS 22:3/N-13:0 | 1123123.913 | 1774449.975 | 466203.6053 | 929921.4751 | 1885978.762 | 114883.3283 | 1147321.621 | 1012443.647 | 1281336.355 | 3493979.001 | 1821650.309 | 1785592.33 |
| PE-Cer 14:3;2O/26:2;O | 2138615.035 | 1315700.787 | 1686874.493 | 1629081.463 | 1613481.687 | 93273.35952 | 1488996.39 | 1451861.835 | 2214653.468 | 2003109.617 | 1482205.832 | 1529064.399 |
| Methyl tetradecanoate | 20954087.51 | 18465254.89 | 15611341.43 | 13703879.07 | 6666960.401 | 8721808.674 | 7506929.13 | 6445461.909 | 8213152.129 | 19087192.14 | 9373990.389 | 8606955.243 |
| (3-Ethyl-2-imino-2,3-dihydro-1H-benzimidazol-1-yl)acetic acid | 289099.3446 | 98968.61878 | 481291.1208 | 121512.4288 | 9390616.81 | 4159786.055 | 7642724.884 | 6588957.27 | 13600704.24 | 6874425.177 | 10731045.76 | 10903670.96 |
| Betamethasone 9,11-epoxide 21-propionate | 2168120.236 | 100818.2999 | 1305222.738 | 205549.871 | 100294.7926 | 1583780.23 | 256329.2003 | 415226.7815 | 2024095.213 | 152544.2366 | 2969131.466 | 170975.6931 |
| 4-(4-Piperidinyl)morpholine | 2718545.149 | 4804141.844 | 9557265.103 | 8491585.088 | 3048088.924 | 3114000.206 | 3193848.689 | 2817107.198 | 3047820.362 | 4947240.287 | 2967957.918 | 10319878.37 |
| N-Ethylglycine | 12799163.47 | 3532427.941 | 24911881.33 | 14636764.4 | 4248571.312 | 3758704.811 | 4050202.518 | 2702763.825 | 6195145.01 | 11201503.69 | 14687459.14 | 4117930.954 |
| PC 9:0_26:2 | 138876 | 544524 | 712279 | 790145 | 591838 | 168027. | 407498 | 537663. | 786569 | 604720 | 778981 | 509202. |

| Metabolites | PBS_1 | PBS_2 | PBS_3 | PBS_4 | BCG_1 | BCG_2 | BCG_3 | BCG_4 | BCGΔ*pe18*_1 | BCGΔ*pe18*_2 | BCGΔ*pe18*_3 | BCGΔ*pe18*_4 |
|---|---|---|---|---|---|---|---|---|---|---|---|---|
| | 76.02 | 7.082 | 8.102 | 9.622 | 6.702 | 746 | 1.71 | 8744 | 5.329 | 5.02 | 6.699 | 2376 |
| Pyrazolo[1,5-a]pyrimidine-3-carbonitrile, 4,7-dihydro-5-methyl-6-(1-methylethyl)-7-oxo- | 3486560.09 | 3313579.262 | 6011077.615 | 6480249.683 | 3682489.785 | 4342120.061 | 3706214.231 | 3896683.629 | 5141201.615 | 3034628.684 | 4834469.963 | 12721327.91 |
| PC O-15:1_20:4 | 2949401.849 | 2624162.32 | 1407986.057 | 885504.8687 | 2781808.007 | 295306.271 | 1775277.32 | 3484379.866 | 2099058.494 | 3179374.784 | 2968245.335 | 4389897.549 |
| PG 22:5_22:6 | 204379461.8 | 153964496.5 | 64302577.53 | 162272266.3 | 96821318.84 | 44764323.34 | 58157399.48 | 98691784.2 | 181980024.8 | 130553562.2 | 60738119.14 | 81349703.01 |
| 2-Naphthalenesulfonic acid | 4536493.333 | 4913596.853 | 6319589.534 | 6576990.375 | 1944901.014 | 2168382.959 | 3192484.997 | 1698984.809 | 3389543.603 | 2803903.195 | 2866056.442 | 6857865.544 |
| METHYL 4-FORMYLBENZOATE | 5929831.202 | 7275548.718 | 5581345.448 | 14827253.85 | 2537937.382 | 4088630.505 | 5638930.789 | 2314908.027 | 11965949.78 | 2475524.892 | 3011243.818 | 4533513.116 |
| 1-Palmitoyl-2-(5-hydroxy-8-oxo-6-octenoyl)-sn-glycero-3-phosphocholine | 2161413.542 | 2471823.773 | 2478350.716 | 2561298.759 | 2453008.667 | 2689684.293 | 2641013.379 | 2388308.905 | 4752152.528 | 4551926.142 | 4385067.373 | 4437976.774 |
| PG 18:1_22:5 | 87241596.6 | 69082859.88 | 39069376.01 | 43662105.24 | 100608319 | 45588444.38 | 85569484.46 | 72618557.61 | 150908023.8 | 189888777.7 | 59761724.8 | 56991555.26 |
| Niflumic acid | 249904.9855 | 252002.7147 | 2830766.12 | 261678.1156 | 652843.9422 | 281556.4724 | 4297767.455 | 449983.7306 | 443941.492 | 3043484.912 | 5529502.225 | 398311.0827 |

| Metabolites | PBS_1 | PBS_2 | PBS_3 | PBS_4 | BCG_1 | BCG_2 | BCG_3 | BCG_4 | BCGΔ*pe18*_1 | BCGΔ*pe18*_2 | BCGΔ*pe18*_3 | BCGΔ*pe18*_4 |
|---|---|---|---|---|---|---|---|---|---|---|---|---|
| PI 18:1_20:4 | 19274240.44 | 15111046.29 | 6117000.001 | 4695300.687 | 12944057.5 | 10711387.87 | 24296326.69 | 32477022.73 | 11382537.92 | 62190597.86 | 19212556.15 | 28400347.13 |
| 5-VALEROLACTONE | 319969.0336 | 8205097.415 | 12098747.97 | 10882276.72 | 2699845.093 | 2828445.892 | 3021419.822 | 2953491.066 | 17472693.14 | 14330020.05 | 14741463.54 | 16884480.04 |
| PI 8:0_26:1 | 8386031.515 | 14268157.45 | 10246370.51 | 8004109.764 | 7645130.104 | 2052762.196 | 5587047.96 | 3423711.739 | 11124238.45 | 8236109.17 | 6158798.568 | 4453332.647 |
| delta-guanidovaleramide | 617642.7329 | 714038.8791 | 2456452.808 | 3806189.966 | 520796.4933 | 582217.9567 | 656040.9417 | 583507.6285 | 558398.6747 | 516686.5565 | 400276.5081 | 4100222.129 |
| 2-amino-3-({hydroxy[2-(icosanoyloxy)-3-(octadecanoyloxy)propoxy]phosphoryl}oxy)propanoic acid | 2724194.659 | 1746769.335 | 1186676.637 | 1586992.403 | 10591197.51 | 3958201.192 | 3902040.055 | 12732717.49 | 15975169.56 | 17619282.21 | 12049984.83 | 5511154.643 |
| Ethyl 2-amino-5-ethyl-3-thiophenecarboxylate | 709521.3943 | 625989.5764 | 766851.7608 | 731566.9915 | 4972184.331 | 3539670.872 | 4737730.585 | 3836344.547 | 7120036.144 | 6010971.987 | 6591382.764 | 5679265.75 |
| gamma-Glutamyl-methionine sulfoxide | 108815.8396 | 108536.2212 | 143156.5849 | 152158.5993 | 6737788.91 | 3692682.931 | 1803490.329 | 1485862.879 | 5334425.246 | 5536934.633 | 4738569.746 | 4816211.922 |
| PC O-37:6 | 9911824.79 | 9485161.358 | 14816167.48 | 11538938.53 | 11023143.34 | 804828.4237 | 9656354.219 | 7223709.389 | 10674902.02 | 11381676.92 | 9976067.57 | 10555806.05 |
| .beta.-Estradiol | 5082.65 | 0 | 0 | 0 | 206648 | 167412 | 148953 | 141700 | 280084 | 197258 | 237376 | 308058 |

| Metabolites | PBS_1 | PBS_2 | PBS_3 | PBS_4 | BCG_1 | BCG_2 | BCG_3 | BCG_4 | BCGΔ*pe18*_1 | BCGΔ*pe18*_2 | BCGΔ*pe18*_3 | BCGΔ*pe18*_4 |
|---|---|---|---|---|---|---|---|---|---|---|---|---|
| 3,17-disulfate | 3624 | | | | 8.063 | 8.584 | 1.22 | 9.282 | 3.742 | 1.79 | 4.205 | 4.039 |
| 1H-1,2,4-Triazol-3-amine | 669142.7164 | 3612981.757 | 7517220.512 | 5492577.832 | 434748.4393 | 133668.6953 | 782491.3466 | 213529.294 | 1010038.565 | 823587.0901 | 733564.5461 | 14766808.71 |
| 3(2H)-Pyridazinone, 4-chloro-5-(4-morpholinyl)- | 3244893.444 | 2892876.362 | 3587199.348 | 3466810.398 | 24446169.41 | 17101656.78 | 23471930.51 | 19169895.66 | 35523119.18 | 28054936.73 | 31403168.19 | 27352026.41 |
| PE O-28:4_8:0 | 3570969.926 | 1915659.668 | 5398386.135 | 2708677.395 | 2758376.848 | 11312.3558 | 3142358.033 | 1947774.617 | 3101869.523 | 3991459.039 | 2824838.976 | 2246799.828 |
| PC O-33:3 | 7655097.757 | 2997603.552 | 13068858.4 | 5612209.798 | 6851710.827 | 94720.96229 | 5798225.47 | 2025231.654 | 6128786.221 | 9397567.79 | 4327997.422 | 3490523.861 |
| Lormetazepam | 23696.66075 | 28129.25797 | 35909.10856 | 19875.3853 | 1412342.742 | 1210618.1 | 1006793.237 | 1036927.235 | 2084308.848 | 1403864.559 | 1642067.552 | 2070415.245 |
| PE 8:0_26:1 | 23706810.71 | 17091812.02 | 22912239.12 | 22139138.67 | 19250571.72 | 374853.5486 | 14618954.03 | 3870541.422 | 31741011.44 | 24448724.42 | 20924709.03 | 5133141.655 |
| Torin 2 | 1049364.65 | 1529479.291 | 1835513.008 | 1496838.759 | 766341.1693 | 1655733.053 | 898820.2597 | 836271.2594 | 521230.7368 | 579001.1769 | 983776.8043 | 4120343.47 |
| N-Nitrosopyrrolidine | 284013.2023 | 805307.6909 | 1078974.141 | 1411895.355 | 440058.604 | 421348.2708 | 549303.3699 | 339947.6372 | 338568.8037 | 570070.8862 | 511348.1883 | 5018162.53 |
| 1H-Pyrazolo[3,4-b]pyridine | 6529282.662 | 2006266.8 | 2373920.357 | 4091686.659 | 2664900.043 | 3935026.367 | 4627016.949 | 4973989.541 | 9626019.277 | 8326608.869 | 4620675.233 | 2780482.702 |
| SMGDG O-28:3_6:0 | 362991959.3 | 438510616 | 196733784 | 298242045.8 | 86024987.01 | 44009057.57 | 91250692.42 | 70615131.39 | 125839404.6 | 153156960.5 | 75499496.51 | 78275900.83 |
| PC O-35:5 | 558463 | 411454 | 587858 | 420894 | 458341 | 107802 | 492773 | 519835 | 565318 | 659256 | 466241 | 821618 |

| Metabolites | PBS_1 | PBS_2 | PBS_3 | PBS_4 | BCG_1 | BCG_2 | BCG_3 | BCG_4 | BCGΔ*pe18*_1 | BCGΔ*pe18*_2 | BCGΔ*pe18*_3 | BCGΔ*pe18*_4 |
|---|---|---|---|---|---|---|---|---|---|---|---|---|
| | 36.76 | 83.12 | 33.92 | 74.75 | 42.08 | 8.212 | 12.35 | 12.01 | 74.2 | 21.79 | 73.15 | 16.86 |
| SM 34:1;3O | 22772398.75 | 17760680.16 | 20277792.06 | 17022970.33 | 17998849.68 | 799653.0736 | 13660700.01 | 31840932.46 | 34633477.08 | 24177241.36 | 7800831.904 | 25645699.81 |
| Cavipetin B | 152809.5394 | 145036.1735 | 674411.205 | 166200.9559 | 287301.6163 | 181687.8729 | 573127.5151 | 410451.2451 | 214668.5288 | 460953.2913 | 910001.7056 | 1011748.473 |
| 2,4-Dimethoxy-2'-hydroxy-5'-methylchalcone | 1660648.609 | 5993237.473 | 12045675.9 | 7526087.714 | 1067173.506 | 484312.2055 | 1332486.305 | 261735.5538 | 987832.0111 | 1822701.963 | 1277646.926 | 19059003.02 |
| DGCC 9:0_28:5 | 22261173.59 | 17172235.37 | 22448354.27 | 22293835.68 | 13760586 | 4395186.618 | 15170552.85 | 22272624.58 | 33620145.51 | 22050941.43 | 17520584.87 | 13407858.78 |
| SM 42:5;3O | 6239910.202 | 5536640.841 | 5505063.822 | 6045343.405 | 6021128.546 | 833714.5164 | 6086295.573 | 13014977.75 | 14605418.76 | 5583786.136 | 3044025.551 | 16313946.05 |
| 5-Methyldodecanoylcarnitine | 27246584.36 | 23373291.32 | 18109642.56 | 26395768.03 | 26818450.67 | 29109599.43 | 31316177.9 | 24043029.72 | 78346650.22 | 29056735.71 | 31898204.19 | 22313493.75 |
| Taprostene (free acid) | 229720.1595 | 590545.596 | 285835.3896 | 26495326.06 | 212455.2162 | 890594.7999 | 39800329.96 | 179780.2908 | 13433935.22 | 2764010.11 | 193059.849 | 44589166.09 |
| LPE 15:1 | 679325.1025 | 507608.3911 | 527763.205 | 1020817.45 | 394130.9493 | 144809.0167 | 415411.4389 | 270792.4529 | 940891.1161 | 625111.0685 | 400304.9612 | 424645.655 |
| LPI 15:0 | 1370134.062 | 1587892.743 | 1288451.534 | 1453550.652 | 9341679.454 | 5089148.368 | 8061418.869 | 35459569.5 | 35128493.18 | 34879783.33 | 40743142 | 6818782.562 |
| Risperidone (Risperdal) | 566842.3698 | 615821.0133 | 446959.2039 | 2170152.17 | 571910.3857 | 763834.6498 | 892845.2376 | 377743.1301 | 1081910.905 | 1026781.498 | 599663.0998 | 1352974.65 |
| PC(20:3(5Z,8Z,11Z)/14:0) | 123913943.9 | 49094077.72 | 57346906.71 | 78671487.69 | 77316128.51 | 49757371.73 | 208076537.4 | 55203093.37 | 152941660.5 | 215810201.7 | 91902033.05 | 104609010.1 |

| Metabolites | PBS_1 | PBS_2 | PBS_3 | PBS_4 | BCG_1 | BCG_2 | BCG_3 | BCG_4 | BCGΔ*pe18*_1 | BCGΔ*pe18*_2 | BCGΔ*pe18*_3 | BCGΔ*pe18*_4 |
|---|---|---|---|---|---|---|---|---|---|---|---|---|
| PC(P-16:0/16:0) | 49954778.93 | 39383170.85 | 66612756.12 | 37564206.45 | 39414318.8 | 1765694.211 | 30423591.57 | 70725141.79 | 75630495.98 | 73096208.51 | 17048812.49 | 55703770.21 |
| Lipofectin | 9274985.85 | 42224895.25 | 13729851.07 | 27667796.96 | 13348028.59 | 8877163.395 | 9998229.146 | 12067440.54 | 16278621.17 | 39327867.81 | 20142568.09 | 8558365.726 |
| 6-(Chloromethyl)-N2-(2,4-dimethylphenyl)-1,3,5-triazine-2,4-diamine | 439883.3067 | 115478.0255 | 499188.8723 | 39047.62763 | 1181338.25 | 958302.795 | 649365.3316 | 517149.7184 | 1369898.194 | 1215299.459 | 1118236.752 | 1304663.142 |
| 8-Bromo-2,5-dihydro-3H-[1,2,4]triazino[5,6-b]indole-3-thione | 1752095.981 | 2824783.65 | 1280081.35 | 2032013.805 | 990553.2936 | 829260.7802 | 1273077.996 | 2017965.75 | 2429530.771 | 1879044.033 | 2192387.522 | 1305928.458 |
| 2-(4-Amino-1-piperidinyl)acetamide | 389705.99 | 910974.0106 | 1738960.19 | 1397951.334 | 52171.35659 | 677563.2371 | 602905.8036 | 844766.7659 | 545963.3369 | 479144.2723 | 418682.9112 | 2437002.387 |
| Ethyl 2-acetylpentanoate | 832094.2693 | 819655.8949 | 678383.0112 | 5868614.737 | 871939.7278 | 857618.8113 | 8173198.738 | 969209.6592 | 3838437.689 | 1232724.05 | 855974.5402 | 10038352.27 |
| PC O-22:6_20:3 | 1363687.822 | 839769.9699 | 1562420.401 | 1669041.969 | 1748559.924 | 1795914.315 | 1622104.393 | 1703527.888 | 3517188.594 | 2099607.708 | 2256871.701 | 1841017.879 |
| PS 18:0_22:5 | 6195262.819 | 9209029.765 | 20406837.82 | 7614301.171 | 4401792.499 | 1017308.618 | 10948767.36 | 2987606.151 | 5771170.974 | 11102057.39 | 8521028.385 | 8011391.482 |
| Phenyl-p-tolylamine | 1688377.836 | 6780713.6 | 12263154.99 | 8476288.776 | 1891859.628 | 1723468.265 | 4166367.977 | 1780780.029 | 1986796.574 | 2705156.472 | 1798308.958 | 15143556.86 |
| 5-Amino-1-methyl-2-piperidinone | 3670914.438 | 5419430.443 | 4901006.749 | 4981728.715 | 1949900.864 | 1977193.538 | 2110203.097 | 1855203.382 | 2440494.864 | 2358264.013 | 2666556.842 | 8710786.183 |

| Metabolites | PBS_1 | PBS_2 | PBS_3 | PBS_4 | BCG_1 | BCG_2 | BCG_3 | BCG_4 | BCGΔ*pe18*_1 | BCGΔ*pe18*_2 | BCGΔ*pe18*_3 | BCGΔ*pe18*_4 |
|---|---|---|---|---|---|---|---|---|---|---|---|---|
| PE O-22:5_18:1 | 5426634.956 | 3912776.826 | 4991758.342 | 4927541.248 | 4467152.203 | 604314.4403 | 4295580.801 | 3110119.759 | 6729184.117 | 5695458.536 | 4322907.708 | 5020889.709 |
| LNAPS 22:3/N-14:0 | 613128.7108 | 421020.1747 | 646559.2589 | 646520.1491 | 662244.5996 | 0 | 703413.6793 | 240637.6856 | 2270596.901 | 594943.0103 | 654036.405 | 306690.0429 |
| Triciribine | 800800.0065 | 655646.3947 | 783989.7158 | 798863.3932 | 670410.0252 | 872317.9098 | 679561.9527 | 674390.2395 | 791629.2251 | 636266.4213 | 2006759.131 | 701253.4286 |
| Promethazine sulfoxide | 3433496.502 | 3647941.983 | 4761958.801 | 12697055.43 | 9373114.711 | 3785107.741 | 5208039.741 | 8099413.403 | 4490440.829 | 15127825.31 | 14803635.04 | 4147009.309 |
| Acamprosate | 808483.9802 | 1217055.918 | 1764272.073 | 1545547.015 | 839316.5558 | 848056.0206 | 1256765.375 | 692641.7646 | 809359.1577 | 750113.9816 | 977568.5928 | 3406495.106 |
| Eplerenone | 1236977562 | 199516522.1 | 943864909.5 | 214827617.1 | 218045479.8 | 1060658062 | 462668899 | 608117349.2 | 1293372938 | 266055222.6 | 1385554561 | 291757209.7 |
| PC 18:0_16:1 | 9820119.822 | 5995592.152 | 9376358.575 | 7835575.274 | 5756135.017 | 93666.78911 | 6217180.966 | 5316732.757 | 7790986.425 | 6186130.363 | 7603815.154 | 4891585.964 |
| PE O-26:6_14:1 | 12491783.62 | 9575562.73 | 16854795.72 | 13529186.48 | 14226335.97 | 2319046.147 | 12999887.96 | 10069660.34 | 12508341.05 | 12984541.89 | 15242451.35 | 13733106.37 |
| PC 16:0_17:0 | 126466.3513 | 957940.7743 | 1249289.904 | 1016505.604 | 542918.6996 | 292263.3104 | 460498.0497 | 312002.8428 | 997106.5932 | 744723.0412 | 591481.7439 | 1008092.807 |
| Pioglitazone | 600146.3113 | 927195.8939 | 1014537.809 | 768432.7804 | 367610.7161 | 487204.8101 | 436732.2818 | 349762.5751 | 373901.7154 | 377410.5301 | 402633.5366 | 1492446.295 |
| PC 8:0_36:9 | 2352212.593 | 2502928.265 | 1135564.805 | 1502384.214 | 490078.4915 | 493645.5844 | 545778.7593 | 633366.6874 | 920430.228 | 671840.5666 | 623084.8698 | 766827.417 |
| PC(34:4) | 61273604.39 | 42943671.53 | 24903756.23 | 46150409.58 | 59153579.26 | 52344254.6 | 167732987.6 | 65012429.66 | 158162149 | 188778338.5 | 65427413.12 | 48135974.62 |

| Metabolites | PBS_1 | PBS_2 | PBS_3 | PBS_4 | BCG_1 | BCG_2 | BCG_3 | BCG_4 | BCGΔ*pe18*_1 | BCGΔ*pe18*_2 | BCGΔ*pe18*_3 | BCGΔ*pe18*_4 |
|---|---|---|---|---|---|---|---|---|---|---|---|---|
| [2-(Trifluoromethyl)-1,3-thiazol-4-yl]methanol | 36330337.96 | 22016699.81 | 33033457.99 | 31950493.34 | 29509398.54 | 30471071.88 | 29847434.26 | 35078843.55 | 75443344.17 | 30707918.41 | 29689322.01 | 33185869.72 |
| SM 35:1;3O | 6123540.33 | 5679235.164 | 10420375.39 | 10082030.5 | 5097971.056 | 347109.4179 | 4641198.564 | 4125158.742 | 7324576.531 | 6318790.66 | 6193070.122 | 2477909.912 |
| 1,2,2,6,6-Pentamethyl-4-piperidinol | 917468.7938 | 1158633.966 | 1969460.157 | 1595651.64 | 788040.0387 | 720793.91 | 983783.529 | 753990.0632 | 814551.5079 | 775072.5655 | 835778.2755 | 2734679.873 |
| N-Nitrosopiperazine (NPAZ) | 73373612.49 | 72162055.41 | 68495092.6 | 70559753.34 | 419982.0324 | 75300553.72 | 76318644.16 | 655223.9131 | 445319.1771 | 63751881.7 | 68600571.78 | 77514443.28 |
| LPC(30:0) | 8057963.667 | 6970328.433 | 7254909.025 | 4186281.618 | 6182207.97 | 5362388.179 | 6814999.335 | 5594868.537 | 15067489.07 | 7341172.195 | 5240789.701 | 7180918.064 |
| PI 16:0_19:0 | 12663660.03 | 5234451.95 | 8490898.256 | 17532124.89 | 3682118.175 | 233718.2225 | 4836175.267 | 369091.3792 | 19260407.65 | 6334157.391 | 7005568.775 | 280746.7194 |
| PE 18:0_20:3 | 496629.7649 | 231199.1989 | 992389.3183 | 564417.3924 | 632064.0849 | 0 | 411487.2849 | 66601.46971 | 690338.2472 | 689031 | 639836.3626 | 199340.5767 |
| Argininic_acid | 931661.1638 | 1092437.963 | 1007495.648 | 988414.8403 | 65847645.52 | 44364220.61 | 56946450.86 | 44975519.42 | 92357167.38 | 53010163.91 | 59113340.09 | 83850362.86 |
| PG 17:1_22:5 | 2914501.809 | 3109076.714 | 1316985.412 | 1985606.771 | 4814773.903 | 5922365.023 | 5151057.636 | 7454398.811 | 10821589.36 | 12040621.32 | 3412967.291 | 5677544.835 |
| 1-O-Hexadecyl-2-O-(5Z,8Z,11Z,14Z,17Z-eicosapentaenoyl)-sn-glyceryl-3-phosphorylcholine | 84794913.66 | 76976381.53 | 44975479.86 | 54512211.78 | 16232421.7 | 14395494.64 | 18092384.57 | 16647189.85 | 30112361.59 | 35156184.16 | 14983081.61 | 12381206.37 |

| **Metabolites** | **PBS_1** | **PBS_2** | **PBS_3** | **PBS_4** | **BCG_1** | **BCG_2** | **BCG_3** | **BCG_4** | **BCGΔ*pe18*_1** | **BCGΔ*pe18*_2** | **BCGΔ*pe18*_3** | **BCGΔ*pe18*_4** |
|---|---|---|---|---|---|---|---|---|---|---|---|---|
| 4-Hydroxynonenal alkyne | 6257997.277 | 5046819.618 | 7709344.992 | 31248557.44 | 1051265.593 | 7894491.611 | 40783557.82 | 1055767.712 | 21168467.99 | 19561717.79 | 1745115.962 | 32244195.19 |
| 1H-Indazole-3-carboxamide, N-[(1S)-1-(aminocarbonyl)-2-methylpropyl]-1-pentyl- | 3170341.932 | 2482046.45 | 2516139.859 | 2196168.704 | 3773583.171 | 2546618.09 | 3151295.777 | 1495000.768 | 3511458.816 | 6369219.352 | 3358343.258 | 2930657.349 |
| PS 10:0_26:2 | 5897249.204 | 3522870.285 | 5853321.798 | 6128498.975 | 4554440.864 | 573695.316 | 3213883.065 | 4435984.797 | 5811602.26 | 5037074.71 | 2895846.603 | 4326754.952 |
| 3-(5,6-Dihydroxyheptyl)-4-methyl-2H-furan-5-one | 401552.8101 | 1230822.12 | 4023528.895 | 2973045.998 | 220259.4925 | 244865.3076 | 630950.1405 | 275272.249 | 218580.9955 | 512829.8343 | 310685.815 | 3349764.356 |
| 3-Pyridinecarbonitrile, 1,2-dihydro-4-methyl-2-thioxo-6-(trifluoromethyl)- | 776250571.5 | 1065350727 | 998516829 | 995246441.8 | 882037119 | 938046204.3 | 1037215323 | 1201598510 | 1171207771 | 1668023658 | 1044081289 | 1464627260 |
| 4-(4-Chlorobenzoyl)-1-(2H)-phthalazinone | 2094136.923 | 15081681.34 | 26000668.41 | 17911080.53 | 2445345.506 | 248490.7667 | 4245519.782 | 811238.8511 | 4239357.352 | 7620067.225 | 5414028.313 | 26101406.01 |
| PC O-30:0 | 23561691.6 | 9951880.611 | 16139272.4 | 16569083.01 | 20748833.04 | 243758.8399 | 19985453.03 | 21812332.26 | 28667525.6 | 26144138.18 | 14189522.3 | 23266480.38 |
| PE 8:0_32:5 | 19880204.09 | 13912227.97 | 25789452.51 | 20105559.34 | 20196431.07 | 130189.8368 | 14270077.7 | 6022518.936 | 19035226.47 | 19915106.28 | 19357071.05 | 9643999.291 |

| Metabolites | PBS_1 | PBS_2 | PBS_3 | PBS_4 | BCG_1 | BCG_2 | BCG_3 | BCG_4 | BCGΔ*pe18*_1 | BCGΔ*pe18*_2 | BCGΔ*pe18*_3 | BCGΔ*pe18*_4 |
|---|---|---|---|---|---|---|---|---|---|---|---|---|
| PC 16:0_20:2 | 2977506.606 | 2591652.576 | 2530690.387 | 2661207.036 | 1548747.34 | 497552.2369 | 1763772.848 | 727592.828 | 2158856.827 | 1980021.34 | 1764352.2 | 611058.872 |
| PC O-36:6 | 5559837.004 | 6351187.252 | 2263648.599 | 3855418.186 | 2783532.215 | 1488172.984 | 2330405.893 | 964649.5875 | 3466174.078 | 2846945.245 | 2304635.662 | 2595575.252 |
| PG 18:2_20:4 | 49575258.49 | 23238274.26 | 27578770.99 | 13360774.49 | 36651487.45 | 32731220.97 | 57323425.03 | 44098403.56 | 53177562.23 | 92907917.02 | 46763041.09 | 43076637.53 |
| PS 16:0_16:1 | 2374740.754 | 651047.2592 | 1853999.046 | 1507796.025 | 1139670.252 | 20584.6355 | 1204073.908 | 266838.9236 | 1605270.946 | 1190069.023 | 2444550.872 | 162756.2505 |
| Asp-Val | 546199.6001 | 676929.5865 | 801954.1592 | 708520.319 | 3216112.038 | 2883629.26 | 2864621.285 | 2303837.269 | 3765343.191 | 4165268.305 | 3663445.261 | 3227780.427 |
| Nebracetam (hydrochloride) | 5399948.287 | 32329981.65 | 50394216.05 | 56465835.97 | 3387220.552 | 1783130.038 | 6783432.493 | 1434977.772 | 6013186.139 | 7365952.214 | 3502689.081 | 364909831.9 |
| CAR 13:1 | 1247503.638 | 1749938.08 | 2129152.588 | 1430213.782 | 1155291.654 | 1137352.884 | 1222430.626 | 1145766.169 | 1167781.262 | 1173596.004 | 1210579.963 | 2799555.885 |
| CE 22:5 | 2822086.358 | 3850438.673 | 6588618.93 | 3575214.355 | 3522589.379 | 4291504.793 | 2319910.118 | 1293140.321 | 3126616.688 | 3114759.801 | 6812672.408 | 3313127.764 |
| N-hydroxyhistamine | 2107024.388 | 1146601.739 | 1836168.566 | 1394478.757 | 4036146.308 | 2503099.367 | 3076326.3 | 1936507.937 | 6900374.802 | 3274929.031 | 4649654.568 | 4565373.162 |
| 2-(glutathion-S-yl)-2-methylbut-3-enoate | 6886.595176 | 0 | 3981.519204 | 7741.759314 | 1885142.021 | 1374468.247 | 1376927.591 | 1132735.096 | 2713402.595 | 2144518.918 | 2606343.86 | 2652701.801 |
| 4-Piperidinecarboxamide | 4894733.874 | 12667199.32 | 13195885.87 | 9094196.439 | 3596345.833 | 3832138.264 | 4836302.936 | 5263046.014 | 6865912.481 | 8263804.728 | 3274751.069 | 18381134.15 |
| NAGlySer | 164753 | 838389 | 151884 | 163654 | 180120 | 145349 | 271822 | 206747 | 260965 | 217718 | 117296 | 255261 |

| Metabolites | PBS_1 | PBS_2 | PBS_3 | PBS_4 | BCG_1 | BCG_2 | BCG_3 | BCG_4 | BCGΔ*pe18*_1 | BCGΔ*pe18*_2 | BCGΔ*pe18*_3 | BCGΔ*pe18*_4 |
|---|---|---|---|---|---|---|---|---|---|---|---|---|
| 21:2/16:2 | 202.4 | 82.72 | 151.2 | 653 | 110.7 | 3.313 | 797.1 | 576.6 | 319.1 | 092.5 | 186.2 | 103.1 |
| 3-(1-Isobutyl-1H-tetraazol-5-yl)aniline | 2214520.557 | 18160422.19 | 26201052.27 | 69767320.22 | 67042932.57 | 2857329.293 | 5229290.632 | 1582110.284 | 6045561.206 | 11641191.74 | 20465987.46 | 61353844 |
| NAOrn 26:7/17:2 | 3578683.257 | 2139359.593 | 4493285.768 | 4412066.475 | 4030336.94 | 1057730.485 | 3314390.247 | 3329927.27 | 5268735.666 | 4865195.597 | 3634673.274 | 2815885.732 |
| LNAPE 18:0/N-20:4 | 37033192.11 | 21211743.17 | 24638462.75 | 23606616.76 | 29780870.02 | 1052847.201 | 33467763.84 | 24289295.19 | 37058880.66 | 28063801.43 | 31700702.61 | 19126059.14 |
| LNAPE 2:0/N-6:0 | 1771250.509 | 1982893.434 | 1520474.734 | 1587274.318 | 3116384.868 | 2237300.137 | 2327498.721 | 2257358.521 | 3639010.161 | 2989615.028 | 2993183.273 | 3434007.339 |
| 10,13-dimethyl-17-oxo-3-sulfooxy-1,2,3,4,7,8,9,11,12,14,15,16-dodecahydrocyclopenta[a]phenanthrene | 1715929.253 | 7827116.646 | 16936940.61 | 10106945 | 1263766.853 | 508749.099 | 1666461.017 | 314499.8119 | 1503040.507 | 2255311.878 | 1848479.651 | 19993470.37 |
| PE 8:0_28:2 | 14459795.31 | 10192395.31 | 28216793.98 | 16697457.47 | 15494824.85 | 413676.9137 | 11704369.47 | 3772021.936 | 18313313.66 | 18409776.27 | 12676472.7 | 5255957.416 |
| (1S,3R)-1-Aminocyclopentane-1,3-dicarboxylic acid | 196859.524 | 146133.7168 | 262796.0087 | 20659.4655 | 989429.7287 | 1627214.087 | 938587.9328 | 830962.527 | 3729531.436 | 1157425.079 | 1249874.073 | 1133050.178 |
| PC O-33:1 | 16443167.43 | 11795291.51 | 21451643.84 | 21398981.59 | 7844314.73 | 1145842.296 | 12181270.97 | 13635971.68 | 15576314.17 | 16423180.38 | 12203587.05 | 5702600.868 |
| PI 18:0_18:0;O | 779523.3609 | 857689.3298 | 173037.7336 | 1405642.711 | 159556.6503 | 0 | 265162.5246 | 11595.5893 | 1228446.664 | 159567.6701 | 139924.2766 | 22755.2969 |

| Metabolites | PBS_1 | PBS_2 | PBS_3 | PBS_4 | BCG_1 | BCG_2 | BCG_3 | BCG_4 | BCGΔ*pe18*_1 | BCGΔ*pe18*_2 | BCGΔ*pe18*_3 | BCGΔ*pe18*_4 |
|---|---|---|---|---|---|---|---|---|---|---|---|---|
| 3-(4-Bromophenyl)-7-hydroxy-4H-chromen-4-one | 2341394.971 | 2306196.906 | 2578671.75 | 2932210.457 | 2103652.597 | 1150398.034 | 1418183.647 | 1480334.085 | 2174389.503 | 1850541.42 | 1799338.769 | 2061592.961 |
| 1,2-Dilinoleoyl-sn-glycero-3-phospho-(1'-rac-glycerol) | 5841030.293 | 11913110.84 | 2650535.705 | 4153088.185 | 21662824.47 | 18653971.92 | 27608639.74 | 21331483.79 | 40073756.8 | 36532770.48 | 32635463.33 | 16954402.77 |
| Curvulin | 4407970.052 | 5776652.807 | 2027826.505 | 2404771.002 | 5109488.057 | 5002956.23 | 5445882.158 | 4328312.698 | 5120823.188 | 5053363.463 | 5932133.705 | 9486885.608 |
| 1-Hexadecyl-2-(5Z,8Z,11Z,14Z-eicosatetraenoyl)-sn-glycero-3-phosphocholine | 669777116.5 | 782392776.4 | 303712664.8 | 425471052.2 | 128897163.3 | 107033462.4 | 114591950.2 | 147526684.1 | 162760119.2 | 222263474.9 | 140754822.3 | 106688508 |
| PE 8:0_30:3 | 4863554.774 | 2072236.419 | 5560735.81 | 4032294.543 | 4140016.443 | 243397.3402 | 3393204.494 | 1205309.527 | 5455098.131 | 5909539.483 | 5150011.86 | 1883361.238 |
| Metaproterenol | 6558893.736 | 8735188.332 | 3784371.076 | 9163097.803 | 3432646.471 | 2727592.341 | 18637906.77 | 2757679.56 | 4503535.893 | 7931953.178 | 3255864.804 | 24893611.67 |
| Monoglyceride citrate | 1083154.731 | 988783.5227 | 1074696.923 | 1021652.304 | 8378826.97 | 7747907.079 | 6113374.199 | 10211592.63 | 12781297.17 | 18139684.45 | 10302478.07 | 12960150.02 |
| PC(18:0/18:3(6Z,9Z,12Z)) | 205922585.1 | 268960540.8 | 154479599.5 | 170854775.1 | 251307866.5 | 294722549.9 | 227542489.5 | 397224073.7 | 469691019.5 | 448292015 | 374919588.5 | 194251957.5 |
| epsilon-Decalactone | 7132897.94 | 4518312.05 | 5131667.128 | 4172854.591 | 7354338.744 | 5453942.523 | 5728917.012 | 2958423.124 | 4975231.007 | 5403416.884 | 13835664.02 | 5167289.07 |
| PI(18:1/18:1) | 942182.147 | 1205177.763 | 1262524.89 | 1872300.386 | 3196023.802 | 2852636.62 | 3352874.056 | 2792185.519 | 6583732.85 | 5976680.26 | 5676711.446 | 1777174.237 |

| Metabolites | PBS_1 | PBS_2 | PBS_3 | PBS_4 | BCG_1 | BCG_2 | BCG_3 | BCG_4 | BCGΔ*pe18*_1 | BCGΔ*pe18*_2 | BCGΔ*pe18*_3 | BCGΔ*pe18*_4 |
|---|---|---|---|---|---|---|---|---|---|---|---|---|
| PG 20:4_22:6 | 453291252.1 | 613550939.4 | 238460635.2 | 437864038.6 | 79763211.11 | 44445522.94 | 87976469.48 | 96932650.33 | 168064168.2 | 165101371.8 | 65242619.33 | 67715895.12 |
| PG 16:3_22:6 | 750513.9568 | 789499.4005 | 458430.3586 | 627201.6649 | 520230.3742 | 458429.2462 | 597876.6074 | 1002493.489 | 1621127.378 | 856558.7416 | 442700.1752 | 631519.0697 |
| Pyrimidinol | 4279465.451 | 1066518.803 | 15937031.89 | 16350115.79 | 2535686.543 | 11746618.7 | 10831151.25 | 9017307.114 | 9030251.726 | 9023116.175 | 9375410.346 | 24825835.25 |
| 19-(4-Hydroxyphenyl)nonadecanoate | 8382252.455 | 20352227.87 | 18184032.45 | 9352821.346 | 15651743.39 | 16022215.15 | 16766569.57 | 5705433.603 | 15795572.4 | 19157841.29 | 16870741.17 | 16559401.99 |
| PE O-18:1_22:6 | 5672357.832 | 4877860.673 | 6389957.273 | 9020302.696 | 4391654.662 | 499369.8166 | 4107047.137 | 4366478.503 | 16112928.26 | 3970828.583 | 8496384.281 | 6289384.193 |
| PI 29:5 | 128464.6069 | 147028.8896 | 70145.06892 | 112303.0962 | 93256.62125 | 88920.88344 | 110280.8731 | 81475.24016 | 155246.7196 | 161868.9338 | 139059.2019 | 169935.4415 |
| Isopentenyladenine | 471753.5808 | 357304.9044 | 765370.7598 | 510912.5717 | 993896.5699 | 833072.5633 | 1364115.804 | 912258.1379 | 1220817.71 | 1273516.569 | 1589941.001 | 1153448.369 |
| 6-tert-Butyl-2H-1,2,4-triazine-3,5-dione | 856978.4827 | 5446393.418 | 11798031.11 | 8367111.498 | 1070827.522 | 1014966.911 | 2072517.138 | 893470.2015 | 1053592.029 | 1721076.589 | 872783.9637 | 10311650.83 |
| PE O-23:0_14:1 | 65322740.15 | 49496032.34 | 104664758.6 | 101597960.1 | 80127986.87 | 2310880.98 | 47392171.02 | 82764864.17 | 117755979.1 | 72082629.83 | 37871797.17 | 56469176.69 |
| (+\|-)-Sulfobutanedioic_acid | 19635688.94 | 20171352.75 | 15433083.6 | 18561958.11 | 7051105.242 | 16320716.18 | 18106348.01 | 7573433.533 | 17873201.39 | 19681721.06 | 9439296.956 | 16274665.32 |
| PG 18:1_20:1 | 1127044.002 | 221320.5654 | 1126281.541 | 695034.262 | 1039302.462 | 0 | 856332.4364 | 270030.5929 | 1484271.097 | 3330759.982 | 1237748.052 | 223945.2552 |
| PG 16:0_19:1 | 123120 | 113264 | 201668 | 425771 | 164110 | 143164. | 164213 | 924222. | 226853 | 292622 | 123869 | 696035. |

| Metabolites | PBS_1 | PBS_2 | PBS_3 | PBS_4 | BCG_1 | BCG_2 | BCG_3 | BCG_4 | BCGΔ*pe18*_1 | BCGΔ*pe18*_2 | BCGΔ*pe18*_3 | BCGΔ*pe18*_4 |
|---|---|---|---|---|---|---|---|---|---|---|---|---|
| | 7.502 | 6.114 | 2.462 | 9.496 | 8.287 | 0805 | 0.32 | 0123 | 8.618 | 2.05 | 5.967 | 1459 |
| apo-14'-lycopenal | 2579697.628 | 2614139.148 | 961110.4829 | 2102213.26 | 737292.0067 | 983887.8309 | 1106415.991 | 748832.5545 | 1256351.286 | 1324307.306 | 674876.7281 | 1232114.214 |
| SL 12:2;O/13:1;O | 121709307.4 | 74890415.52 | 61422929.95 | 93421770.21 | 68987382.81 | 56534107.31 | 49494153.02 | 41108300.06 | 61950667.16 | 82202084.36 | 84470091.89 | 44098431.12 |
| Triacsin C | 1758604.631 | 1845253.836 | 2576505.199 | 3413349.722 | 3685880.445 | 7556739.745 | 1751883.339 | 1007943.769 | 2490525.867 | 4865076.102 | 9125266.416 | 2587595.978 |
| LPS 22:4 | 8503385.308 | 9544458.968 | 11012588.58 | 7434522.053 | 15101049.27 | 14743982.51 | 12862334.33 | 12519191.23 | 15496092.16 | 15273709.3 | 27566636.14 | 13751245.76 |
| PMeOH 18:1_26:4 | 4588190.125 | 4008325.617 | 3349320.893 | 3468850.904 | 3869327.836 | 94888.74275 | 2906280.703 | 2495866.102 | 3340860.945 | 2895299.015 | 3549420.096 | 1843127.702 |
| 5-[[17-(5,6-dihydroxy-6-methylheptan-2-yl)-3,12-dihydroxy-4,4,10,13,14-pentamethyl-2,3,5,6,7,11,12,15,16,17-decahydro-1H-cyclopenta[a]phenanthren-2-yl]oxy]-3-hydroxy-3-methyl-5-oxopentanoic acid | 282100.7245 | 319059.0013 | 323592.0504 | 295604.4507 | 336690.5555 | 333292.0375 | 423764.8447 | 274649.9099 | 682191.0618 | 620406.4148 | 648194.5414 | 677811.3303 |
| Ser-Met | 2806064.871 | 1045282.579 | 6218173.425 | 1255449.236 | 2229792.031 | 1910061.669 | 2381010.41 | 2228781.29 | 1722016.168 | 1413595.313 | 2334159.588 | 6004444.722 |

| **Metabolites** | **PBS_1** | **PBS_2** | **PBS_3** | **PBS_4** | **BCG_1** | **BCG_2** | **BCG_3** | **BCG_4** | **BCGΔ*pe18*_1** | **BCGΔ*pe18*_2** | **BCGΔ*pe18*_3** | **BCGΔ*pe18*_4** |
|---|---|---|---|---|---|---|---|---|---|---|---|---|
| Statine | 8015475.389 | 11246283.17 | 14481977.73 | 16534584.81 | 13963264.33 | 10167801.75 | 12110386.63 | 11033705.26 | 17608366.26 | 4059032.088 | 11122441.74 | 31072002.44 |
| M272T63 | 6418151.842 | 7066048.947 | 7250127.442 | 7283695.18 | 7307004.558 | 1801001.043 | 2146478.439 | 7450022.748 | 7251863.286 | 6170724.89 | 7294254.105 | 7421380.672 |
| M398T320 | 221166.3524 | 214360.9772 | 134160.0592 | 195673.3819 | 10736794.89 | 6120114.32 | 3869476.564 | 7189323.661 | 13263475.64 | 11736699.34 | 5585076.364 | 9777155.228 |
| Daltogen | 6366971.989 | 11045294.61 | 20358291.06 | 13472332.1 | 5320842.966 | 4878734.514 | 11668429.66 | 5345766.267 | 8638699.443 | 7725803.477 | 5620643.384 | 39944127.93 |
| PC 8:0_23:0 | 35558350.8 | 18534184.58 | 36504500.56 | 42225969.75 | 29389944.25 | 9365601.598 | 33866883.88 | 22241265.95 | 46379313.02 | 35788239.94 | 27909661.61 | 13444491.52 |
| Dihydroisotanshinone I | 234249.3525 | 234455.872 | 228688.9216 | 233384.7021 | 12598448.93 | 16413554.53 | 34785619.82 | 35575078.11 | 43327280.86 | 14157285.53 | 28507103.01 | 46141520.22 |
| 2-Hydrazinyl-4,6-di(morpholin-4-yl)-1,3,5-triazine | 1567398.904 | 1739078.827 | 1580846.425 | 1867543.741 | 2484567.624 | 2376446.698 | 2222268.004 | 1941894.956 | 2681241.868 | 2773253.755 | 2622811.06 | 3589912.221 |
| PI 18:0_20:4 | 53983987.26 | 33032779.47 | 49066384.7 | 69742910.09 | 27919969.03 | 30779949.28 | 40973874.31 | 34137736.19 | 29398075.68 | 46979548.72 | 56226512.93 | 36741662.62 |
| SPB 17:1;2O | 2754006.441 | 35240199.58 | 15766017.98 | 35266419.26 | 3982712.513 | 4306601.012 | 21786165.9 | 17728193.6 | 15740404.12 | 7738949.467 | 16252970 | 21173223.86 |
| 1-Palmitoyl-2-linoleoyl-sn-glycero-3-phosphocholine | 22725953.09 | 18995693.92 | 21959844.29 | 20194194.5 | 17716656.28 | 8590532.298 | 15221026.84 | 9340494.392 | 21650066.76 | 17655039.85 | 19119468.63 | 7348871.665 |

| Metabolites | PBS_1 | PBS_2 | PBS_3 | PBS_4 | BCG_1 | BCG_2 | BCG_3 | BCG_4 | BCGΔ*pe18*_1 | BCGΔ*pe18*_2 | BCGΔ*pe18*_3 | BCGΔ*pe18*_4 |
|---|---|---|---|---|---|---|---|---|---|---|---|---|
| (4R)-4-((3R,5R,6S,7R,9S,10R,12S,13R,14S,17R)-3,6,7,12-tetrahydroxy-10,13-dimethylhexadecahydro-1H-cyclopenta[a]phenanthren-17-yl)pentanoic acid | 28418.52736 | 181037.117 | 12362.06815 | 4976192.549 | 198133.9802 | 129326.0363 | 4587185.763 | 134101.804 | 1906307.885 | 279002.8908 | 72903.23217 | 10767516.59 |
| 2-(2-Thienyl)-1H-phenanthro[9,10-d]imidazole | 8092607.467 | 7324283.662 | 8152697.701 | 7413213.747 | 11112206.18 | 15834299.01 | 30965357.78 | 30085146.53 | 35272182.47 | 11871252.05 | 24746377.44 | 36350262.94 |
| Tri(3-chloropropyl) phosphate | 9102303.44 | 12870132.34 | 10216092.68 | 10218486.78 | 6898329.705 | 7043521.88 | 6534598.224 | 6548640.798 | 6136430.43 | 7140781.209 | 8739247.699 | 15921558.29 |
| DGTS 8:0_8:0 | 9470860.101 | 9441048.714 | 8964419.638 | 12709483.54 | 9201927.313 | 6702135.594 | 9778330.664 | 9187746.918 | 13742004.66 | 14118775.36 | 13575935.67 | 8703767.022 |
| LPC O-2:0 | 2148110.822 | 2132702.951 | 2490274.941 | 1890783.654 | 1421270.623 | 1217639.904 | 1477998.127 | 1202740.298 | 1189018.844 | 1800667.828 | 2337296.84 | 1585814.271 |
| ST 24:1;O3;T/20:4 | 1350481.154 | 832552.8243 | 1100179.898 | 1316622.35 | 1137514.361 | 30518.73693 | 1672718.535 | 2379586.761 | 1326268.616 | 1522830.401 | 1285610.451 | 2901047.776 |
| Probenecid (Benemid) | 37600.97211 | 19879.68107 | 44424.8798 | 14909.65955 | 1302060.256 | 901717.2426 | 801467.1456 | 946050.5372 | 1740800.259 | 1288448.762 | 1424776.457 | 1616711.04 |
| PG 20:4_22:5 | 300582804.3 | 326571138.3 | 140221780.3 | 237448513.5 | 50703965.26 | 20390123.66 | 60456916.2 | 57453191.16 | 48931461.04 | 92742217.34 | 44073481.58 | 44200708.47 |
| Ribose-1-arsenate | 599058 | 273711 | 381237 | 229269 | 283721 | 222802 | 304364 | 223976 | 339471 | 538518 | 254507 | 575577 |

| **Metabolites** | **PBS_1** | **PBS_2** | **PBS_3** | **PBS_4** | **BCG_1** | **BCG_2** | **BCG_3** | **BCG_4** | **BCGΔ*pe18*_1** | **BCGΔ*pe18*_2** | **BCGΔ*pe18*_3** | **BCGΔ*pe18*_4** |
|---|---|---|---|---|---|---|---|---|---|---|---|---|
| | 3.515 | 6.557 | 4.836 | 6.481 | 8.191 | 5.2 | 5.051 | 3.733 | 6.334 | 7.193 | 9.082 | 9.761 |
| 2-(4-Amino-2-nitroanilino)ethanol | 773543.7459 | 482709.7544 | 767534.2315 | 552999.7322 | 65138587.88 | 46363571.32 | 58119875.88 | 44384024.88 | 86688004.4 | 51551381.23 | 57558423.64 | 76654416.99 |
| 1H-Pyrrolo[2,3-b]pyridine, 2-bromo- | 1810281.587 | 1587505.742 | 1805124.551 | 1709744.124 | 1268603.782 | 1456272.133 | 1614086.657 | 1625374.72 | 1577914.004 | 1586643.121 | 1900679.297 | 2669600.602 |
| (11.Xi.)-4,7-Dihydroxy-12,13-epoxytrichothec-9-en-8-one | 56052716.89 | 8612309.06 | 37399324.29 | 9732419.923 | 9399480.54 | 46642473.51 | 20627334.04 | 25928120.55 | 51415125.05 | 11487911.51 | 53260818.3 | 12410419.22 |
| (2R)-N-(3-Ethoxypropyl)-2,4-dihydroxy-3,3-dimethylbutanamide | 583760.8692 | 2517259.195 | 3517500.368 | 2984337.961 | 712289.8377 | 726335.9207 | 845695.5523 | 726263.2577 | 949873.7678 | 861013.5369 | 651502.5497 | 4650614.125 |
| PG 16:0_18:0 | 2844809.234 | 1589189.527 | 2728453.257 | 2077624.904 | 2065660.572 | 15980.42229 | 2057685.229 | 1546937.748 | 3246099.252 | 2073482.105 | 1367829.255 | 2261286.847 |
| 1-(alpha-linolenoyl)-sn-glycero-3-phosphoethanolamine | 2627260.909 | 14475339.34 | 25134794.54 | 11860027.37 | 26047252.78 | 14752019.53 | 26653488.52 | 16369677.78 | 30835988.11 | 36614541.38 | 34712794.09 | 5333902.388 |
| 7-Methoxy-6-(1,2,3-trihydroxy-3-methylbutyl)chromen-2-one | 24191361.42 | 3742180.94 | 15368219.69 | 4138302.12 | 3824343.494 | 20277894.76 | 8742947.092 | 10761101.76 | 20997586.15 | 4673779.325 | 22131367.2 | 5301589.492 |
| Methylprednisolone | 26404367.02 | 4222667.944 | 18045042.78 | 4907652.96 | 4571555.136 | 22969388.56 | 9887049.122 | 11838888.76 | 23689931.35 | 5184361.754 | 25386082.58 | 6253022.628 |
| MGDG | 105472 | 307836. | 139570 | 805011. | 498091. | 4717.52 | 407689. | 178321. | 663710. | 431152. | 576230. | 309194. |

| Metabolites | PBS_1 | PBS_2 | PBS_3 | PBS_4 | BCG_1 | BCG_2 | BCG_3 | BCG_4 | BCGΔ*pe18*_1 | BCGΔ*pe18*_2 | BCGΔ*pe18*_3 | BCGΔ*pe18*_4 |
|---|---|---|---|---|---|---|---|---|---|---|---|---|
| O-13:1_18:0 | 1.473 | 6214 | 1.459 | 2493 | 5899 | 613 | 56 | 8483 | 1671 | 5687 | 5518 | 3273 |
| PI 27:0_18:4 | 93285.43984 | 92361.84088 | 54833.65028 | 77157.99083 | 125487.8222 | 119951.6979 | 91858.04781 | 133583.3575 | 280876.8724 | 137313.2312 | 166826.3145 | 228075.578 |
| SM 38:3;3O | 19566001.28 | 10837111.31 | 14901730.66 | 22284796.1 | 15010542.57 | 521511.584 | 17712615.45 | 4210389.682 | 26490261.18 | 19900307.4 | 11421766.54 | 9332601.897 |
| SM 40:8;3O | 2517906.496 | 1777095.489 | 2580697.108 | 2709346.148 | 1993436.917 | 165034.1181 | 1624711.093 | 1828227.079 | 2490869.248 | 2226419.795 | 1553925.092 | 1875086.477 |
| PE O-16:4_6:0 | 10347320.79 | 16423496.97 | 20405086.67 | 14089224.56 | 12716297.88 | 16158142.09 | 17739858.48 | 17085826.68 | 13359225.84 | 24034593.46 | 24343435.78 | 16196957.93 |
| N2-[4-(Aminomethyl)phenyl]-5-fluoro-N4-phenyl-2,4-pyrimidinediamine | 249217.8046 | 3641860.187 | 10981046.51 | 7596251.824 | 326464.5511 | 145257.9665 | 984057.8241 | 256242.7399 | 828001.1498 | 1464580.238 | 0 | 14952065.67 |
| 4-Bromobenzenecarboximidamide | 13161661.84 | 10196325.13 | 17661020.56 | 12513048.49 | 12355897.81 | 9928045.537 | 11505880.68 | 11874290.97 | 10782073.4 | 14960388.32 | 11166496.37 | 17441785.21 |
| PC O-18:1 | 4189746.566 | 4691113.199 | 4548997.169 | 484139.8703 | 14443201.73 | 1101544.872 | 1712605.753 | 11290181.44 | 1165309.925 | 6634806.786 | 17376317.69 | 11395984.86 |
| PG O-24:6_16:0 | 759604.6925 | 604555.9505 | 884522.3093 | 653002.2362 | 835851.9012 | 144883.7426 | 361400.0446 | 92354.37743 | 537032.8175 | 709517.4238 | 725294.1585 | 40031.64647 |
| Ergothioneine | 611438.5097 | 134964.8095 | 1204702.108 | 245981.8622 | 6043320.151 | 5313106.163 | 13657805.52 | 12733675.02 | 23892842.3 | 11137556.86 | 18121272.72 | 26121454.46 |
| 3-(2-Chloro-5-fluorophenyl)-3-oxoprop | 5793615.55 | 10897219.92 | 5290384.709 | 6049163.376 | 5393508.257 | 5858022.279 | 5190940.696 | 6116796.391 | 4978376.747 | 4931837.196 | 5904921.875 | 10927047.75 |

| **Metabolites** | **PBS_1** | **PBS_2** | **PBS_3** | **PBS_4** | **BCG_1** | **BCG_2** | **BCG_3** | **BCG_4** | **BCGΔ*pe18*_1** | **BCGΔ*pe18*_2** | **BCGΔ*pe18*_3** | **BCGΔ*pe18*_4** |
|---|---|---|---|---|---|---|---|---|---|---|---|---|
| ionitrile | | | | | | | | | | | | |
| SHexCer 34:1;3O | 23724017.69 | 27822123.69 | 9494835.982 | 5407169.846 | 17590289.13 | 15511568.38 | 29603570.72 | 21063290.24 | 25454859.25 | 37320967.9 | 22144307.03 | 18539615.06 |
| Verapamil | 3920464.729 | 4511621.493 | 3641613.728 | 4163029.827 | 10820712.66 | 10266311.52 | 10926582.22 | 10758896.28 | 12872453.49 | 11968097.05 | 14162344.54 | 11368171.93 |
| PC O-34:3 | 3676066.175 | 2662082.252 | 3169598.814 | 3815678.427 | 4055057.95 | 3151984.91 | 4902973.261 | 3504734.838 | 8009888.048 | 7046109.199 | 3522473.519 | 3091364.876 |
| PG 18:1_16:3 | 294923.8406 | 456739.8568 | 163774.969 | 206323.533 | 1756864.96 | 1485892.933 | 2387310.05 | 1965316.675 | 3052566.361 | 3270767.473 | 1883590.729 | 2257212.565 |
| LACTULOSE | 146563.8781 | 62066.13722 | 254048.7218 | 129028.3953 | 2555472.095 | 5224649.833 | 4048897.941 | 1946906.214 | 11995451.25 | 5474510.567 | 4499384.902 | 6193876.314 |
| 6-[(1-Phenylethyl)sulfanyl]-9H-purine | 1597302.288 | 7281766.346 | 14343449.76 | 10041422.62 | 1520207.293 | 603470.3426 | 1930329.956 | 396890.9439 | 1764271.797 | 2767498.002 | 1741636.261 | 18484808.12 |
| LNAPE 10:0/N-26:6 | 18965555.01 | 15980841.82 | 16355978.11 | 17489588.67 | 6528308.44 | 10984453.97 | 7123008.812 | 9517586.26 | 10189698.92 | 8234127.429 | 10514290.48 | 11146596.41 |
| PC 9:0_28:4 | 12815225.07 | 10960601.64 | 10563682.2 | 9557372.757 | 10756724.27 | 8912959.382 | 13137057.7 | 14760093.16 | 19167816.57 | 14577341.03 | 11173905.26 | 13330234.57 |
| PC 9:0_26:3 | 1253146.499 | 1255440.676 | 1146530.974 | 963102.4904 | 4551276.422 | 5281342.548 | 4240660.086 | 5073384.87 | 5610092.376 | 9245198.205 | 4533542.399 | 4822900.542 |
| PE 9:0_28:4 | 1516433.783 | 1493771.982 | 580798.2707 | 1381007.654 | 3897123.782 | 3582310.311 | 4201166.796 | 3855743.304 | 3896207.413 | 4946325.334 | 3725192.083 | 7767586.828 |
| PG 16:1_20:5 | 994781.9914 | 1161069.907 | 473459.617 | 374310.9199 | 2528398.38 | 1750859.356 | 2549731.191 | 1560209.724 | 3369359.727 | 3376905.355 | 1980064.92 | 1543230.323 |

| Metabolites | PBS_1 | PBS_2 | PBS_3 | PBS_4 | BCG_1 | BCG_2 | BCG_3 | BCG_4 | BCGΔ*pe18*_1 | BCGΔ*pe18*_2 | BCGΔ*pe18*_3 | BCGΔ*pe18*_4 |
|---|---|---|---|---|---|---|---|---|---|---|---|---|
| PE O-16:1_20:4 | 65017541.47 | 117337405.8 | 76499166.87 | 88062966.8 | 54685457 | 43372698.54 | 68309527.24 | 60916261.01 | 66143819.2 | 89708172.85 | 60280498.89 | 59153781.79 |
| LNAPE 17:1/N-16:0 | 2408072.308 | 1134394.488 | 2027556.826 | 1531398.464 | 2073016.176 | 0 | 1265271.518 | 132461.5195 | 2320549.067 | 1829784.03 | 1591645.317 | 0 |
| PC 20:2_20:2 | 8496392.986 | 10425794.84 | 7136525.201 | 8186324.171 | 5276021.445 | 58998.37658 | 15610140.12 | 6292028.653 | 12025750.18 | 11145502.7 | 4682175.955 | 8509886.984 |
| PI 18:2_20:4 | 1313071.018 | 665350.8132 | 532890.7962 | 1215014.864 | 1711244.252 | 1617462.772 | 1926679.712 | 1706573.136 | 1819217.952 | 2356777.199 | 1944456.962 | 2261080.279 |
| 4-Aminomethylcyclohexanecarboxylic acid | 2474067.354 | 2855716.876 | 1827956.876 | 5281327.491 | 1739069.973 | 2383015.896 | 3851514.147 | 1507655.85 | 2967316.955 | 2855459.69 | 1718323.101 | 6522082.289 |
| PG 18:0_22:4 | 223079.2604 | 0 | 115708.2637 | 52153.27826 | 346252.5142 | 34397.02787 | 460321.5104 | 283829.2761 | 937618.1789 | 370592.3969 | 276244.9305 | 271045.2514 |
| LPI 18:0 | 49957191.99 | 52959111.8 | 31820076.74 | 44431788.85 | 54365309.32 | 53849348.03 | 46283636.98 | 46149576.13 | 73038823.93 | 58642335.7 | 56539521.28 | 48310797.47 |
| PC(18:0/18:0) | 21560618.9 | 13725268.32 | 57434594.34 | 34033646.61 | 14104288.34 | 638618.662 | 12353944.34 | 10649464.24 | 25059954.18 | 18351374.15 | 16334383.08 | 9801258.416 |
| 4-(3,5-Dimethyl-4H-1,2,4-triazol-4-yl)piperidine | 140600.4156 | 2208139.069 | 4447791.602 | 3038418.382 | 291026.6923 | 252170.5332 | 519335.2231 | 88984.89885 | 508040.3989 | 783935.2672 | 164737.2328 | 4947079.25 |
| PE O-22:2_19:2 | 8688436.426 | 7635484.395 | 7748611.536 | 19163408.23 | 6966964.997 | 270782.5229 | 15411463.73 | 12279174.89 | 22770482.58 | 9842013.453 | 4301225.042 | 12132611.92 |
| Glycerophospho-N-palmitoylethanolam | 16707432.32 | 19733342.23 | 15082271 | 16500433.56 | 44416627.78 | 42053715.16 | 46750560.83 | 46350576.56 | 52039664.82 | 50525425.56 | 58847514.92 | 46691484.83 |

| Metabolites | PBS_1 | PBS_2 | PBS_3 | PBS_4 | BCG_1 | BCG_2 | BCG_3 | BCG_4 | BCGΔ *pe18*_1 | BCGΔ *pe18*_2 | BCGΔ *pe18*_3 | BCGΔ*pe18*_4 |
|---|---|---|---|---|---|---|---|---|---|---|---|---|
| ine | | | | | | | | | | | | |
| SM 35:3;3O | 3468872.301 | 1435337.986 | 5733723.809 | 4324204.971 | 3002745.087 | 53735.32728 | 2598608.697 | 1600366.478 | 2566829.581 | 4254604.303 | 1797372.357 | 1573395.438 |
| 3'-Azido-3'-deoxythymidine 5'-monophosphate | 7671491.234 | 9243478.201 | 8078164.824 | 8914366.279 | 16573591.99 | 15655700.18 | 15447350.04 | 15525819.53 | 17748648.16 | 17862687.78 | 17135721.29 | 22119218.88 |
| 4-Hydroxy-3-oxo-1,3-dihydronaphtho[2,3-c]furan-5-yl .beta.-D-glucopyranoside | 2139037.97 | 2520441.248 | 3147562.965 | 3170559.06 | 518674883.9 | 398815913.5 | 376369485.8 | 380286612.3 | 550155061.6 | 474165731.7 | 400182814.4 | 512216204.4 |
| 1-(1Z-Octadecenyl)-2-(5Z,8Z,11Z,14Z-eicosatetraenoyl)-sn-glycero-3-phosphocholine | 11134705.75 | 19707391.18 | 13731756.79 | 27683708.42 | 14954515.38 | 415688.2515 | 30254433.34 | 49864644.53 | 19936673.76 | 13328571.36 | 12240994.2 | 67143490.74 |
| 8-AMINOOCTANOIC ACID | 4251361.791 | 4171600.547 | 4039844.651 | 9249885.458 | 3585882.328 | 5012457.38 | 13979767.73 | 3379700.296 | 8910592.038 | 6646342.442 | 4327203.46 | 11451980.45 |
| TG 8:0_16:4_20:5 | 155171.2532 | 216691.3686 | 250986.9763 | 249594.6316 | 5989897 | 4227193.643 | 3434409.902 | 3365009.76 | 6758950.458 | 3750852.708 | 7276617.926 | 2541288.35 |
| N-Decanoylglycine | 48985366.74 | 42639492.84 | 60559097.56 | 42029843.01 | 51312719.35 | 49248259.41 | 38873752.69 | 32506534.63 | 49135285.57 | 52424206.11 | 50836318.79 | 51331537.27 |

| **Metabolites** | **PBS_1** | **PBS_2** | **PBS_3** | **PBS_4** | **BCG_1** | **BCG_2** | **BCG_3** | **BCG_4** | **BCGΔ*pe18*_1** | **BCGΔ*pe18*_2** | **BCGΔ*pe18*_3** | **BCGΔ*pe18*_4** |
|---|---|---|---|---|---|---|---|---|---|---|---|---|
| 2-(Benzoylamino)-4,5,6,7-tetrahydro-1-benzothiophene-3-carboxylic acid | 1151049.376 | 1100511.873 | 1202752.385 | 1058835.073 | 1079653.379 | 1690430.714 | 3255792.507 | 3186353.281 | 3745806.998 | 1254060.317 | 2487595.487 | 3913887.517 |
| LNAPS 24:1/N-10:0 | 11760898.76 | 5516453.461 | 9682754.985 | 7719555.955 | 6451551.253 | 943703.8356 | 5526462.904 | 8424992.244 | 8281287.027 | 9197744.925 | 5890574.983 | 6351751.406 |
| PG 22:4_22:4 | 867327.4398 | 11254283.48 | 5145395.017 | 3930592.15 | 880555.0539 | 633372.3598 | 2221742.538 | 1179546.719 | 1700723.72 | 3254437.464 | 1525751.047 | 745515.6721 |
| SM 32:0;3O | 7927643.211 | 5561910.792 | 24378559.19 | 7783932.181 | 7293064.398 | 62851.10364 | 13938455.28 | 8373644.087 | 13893906.95 | 8547669.397 | 7229675.411 | 6699710.572 |
| 3-[2-(4-Chlorophenoxy)-2-methylpropanoyl]-1,3-thiazolidine-4-carboxylic acid | 428455.0463 | 473148.4286 | 485677.8138 | 1880329.701 | 343796.7855 | 538352.1103 | 2398518.333 | 342497.096 | 1189226.087 | 691425.1456 | 388129.2615 | 2130791.524 |
| 2-Chloro-N,N-dimethylpropan-1-amine | 643454.3124 | 437926.2374 | 1018612.674 | 1419425.698 | 802901.9036 | 739978.9991 | 248903.8591 | 177759.2459 | 687516.6463 | 193722.0614 | 794997.4602 | 2757055.155 |
| PC 16:0_18:3;O | 1277278.286 | 1309987.949 | 974519.5632 | 1024901.303 | 642367.0279 | 656894.4111 | 614496.7787 | 743931.2676 | 715379.5245 | 856351.4286 | 872102.4831 | 812578.6999 |
| 4-Amino-2-oxo-1,2-dihydropyrimidine-5-carboxylic acid | 12902812.59 | 9080106.554 | 114509611.1 | 21990012.15 | 127494174.1 | 228870002.4 | 149716746.8 | 144454297.8 | 164099046.9 | 86301782.03 | 294163966.1 | 245183583.8 |
| LPE 20:4 | 63586977.08 | 64617931.47 | 74951015.23 | 64212745.55 | 62829196.23 | 52998507.03 | 58436303.88 | 49738618.71 | 52861397.39 | 69270075.16 | 77019748.96 | 60403986.33 |

| Metabolites | PBS_1 | PBS_2 | PBS_3 | PBS_4 | BCG_1 | BCG_2 | BCG_3 | BCG_4 | BCGΔ*pe18*_1 | BCGΔ*pe18*_2 | BCGΔ*pe18*_3 | BCGΔ*pe18*_4 |
|---|---|---|---|---|---|---|---|---|---|---|---|---|
| 2-(2-oxo-8,9-dihydrofuro[2,3-h]chromen-8-yl)propan-2-yl acetate | 3128429.978 | 2249696.859 | 5054919.496 | 3270909.005 | 16901865.83 | 10666500.32 | 15888499.5 | 12538172.69 | 16714790.8 | 16697087.3 | 18526463.44 | 13280291.92 |
| PC 8:0_24:0 | 3554636.998 | 2455932.327 | 3745029.254 | 3311914.029 | 3670078.848 | 124568.5347 | 3813213.272 | 5761886.119 | 3348029.825 | 3830474.265 | 3183452.672 | 5675426.648 |
| 2-[(2-Methylpropyl)amino]acetic acid | 4604391.114 | 19451842.37 | 25726082.05 | 27116453.68 | 5791336.121 | 5482987.584 | 17953959.38 | 8541956.378 | 10702751.21 | 7416255.547 | 5872018.295 | 37858885.19 |
| Maltose | 6229.943972 | 13027.11296 | 34877.51122 | 434233.6736 | 2321743.264 | 1925094.058 | 1673306.673 | 1465547.267 | 2194829.585 | 1865984.062 | 2112761.721 | 2429479.646 |
| 3-Hydroxy-5-methoxy-3-(methoxycarbonyl)-5-oxopentanoic acid | 89815.51273 | 432529.4393 | 450662 | 828639.6697 | 1136049.727 | 612474.2591 | 797498.2658 | 757058.8207 | 1013029.127 | 937359.7829 | 992356.6293 | 907214.2335 |
| cis-4,10,13,16-Docosatetraenoic acid | 19013164.78 | 18509179.2 | 39100399.26 | 21283644.1 | 4803513.526 | 6208779.452 | 7150709.729 | 6601317.569 | 6689401.506 | 8247483.539 | 10370953.93 | 5431109.421 |
| FAHFA 16:0/3:0 | 419056.9094 | 427323.4893 | 277247.7212 | 359726.3357 | 620424.4935 | 760401.9745 | 407190.5287 | 529336.4243 | 489602.1206 | 358688.6441 | 577799.0197 | 1560500.575 |
| DGDG O-8:0_22:0 | 5725337.261 | 6721515.515 | 2939134.285 | 5413733.289 | 505753.4562 | 446596.1534 | 556248.2415 | 441240.0638 | 555336.9379 | 695226.2112 | 646775.3513 | 823984.6966 |
| PG 18:1_22:4 | 18028064.44 | 28756528.65 | 10581572.61 | 14561856.01 | 28351789.97 | 11244994.03 | 21394959.13 | 11247261.16 | 21787350.79 | 33034428.62 | 33602561.46 | 20253971.24 |
| HexCer | 439045 | 263003 | 308949 | 293718 | 374731 | 0 | 412663 | 311680 | 460215 | 347417 | 397931 | 238571 |

| Metabolites | PBS_1 | PBS_2 | PBS_3 | PBS_4 | BCG_1 | BCG_2 | BCG_3 | BCG_4 | BCGΔ*pe18*_1 | BCGΔ*pe18*_2 | BCGΔ*pe18*_3 | BCGΔ*pe18*_4 |
|---|---|---|---|---|---|---|---|---|---|---|---|---|
| 8:0;2O/32:8 | 8.237 | 8.583 | 9.273 | 3.727 | 9.531 |  | 4.398 | 3.167 | 9.589 | 8.326 | 6.467 | 8.561 |
| PC(42:8) | 20193489.26 | 24415549.67 | 13220409.29 | 14809065.21 | 8608707.183 | 9100663.619 | 11068099.38 | 12865011 | 12845470.87 | 16909666 | 9568121.935 | 8951964.558 |
| Mestranol | 38485516.3 | 36351587.05 | 30197746.45 | 38561229.31 | 25832848.91 | 28869037.47 | 30647018.08 | 30331506.48 | 30701243.27 | 33616692.89 | 28355734.09 | 44707629.29 |
| Heptadecasphinganine | 192672060.2 | 138710552.2 | 106108206.2 | 141602456.9 | 113997703.8 | 92465087.5 | 92818195.98 | 84728936.6 | 102331675.9 | 116700100.9 | 155203102.6 | 70575296.37 |
| Pentaethylene glycol | 21404134.73 | 34150940.65 | 43135429.52 | 36895811.89 | 25411553.99 | 21485614.55 | 23690019.62 | 18412857.11 | 44652859.5 | 26045825.18 | 25133089.6 | 62287785.99 |
| SL 24:0;O/16:4 | 41484918.87 | 35209416.85 | 35200572.99 | 38411664.73 | 15155416.05 | 24392251.18 | 16255601.27 | 20854639.09 | 20489933.64 | 18655395.98 | 23076738.24 | 25010418.59 |
| Ginkgolide C | 543112.1678 | 741418.2816 | 469301.5969 | 480070.1826 | 310530.4628 | 1208357.918 | 283835.8295 | 389519.1663 | 294727.7909 | 316270.2939 | 441024.506 | 2507120.374 |
| PE 38:10 | 20926398.55 | 17789219.8 | 17454482.82 | 18223868.52 | 9174479.319 | 14581969.59 | 9904613.109 | 11739858.61 | 12461959.96 | 10937238.16 | 13457830.94 | 14603055.31 |
| NAE 19:1 | 2737633.231 | 3183335.528 | 2779436.687 | 2911015.409 | 2164800.374 | 2471746.132 | 2460176.159 | 2210282.459 | 2055013.08 | 2999337.258 | 2576587.859 | 5192668.923 |
| Ethiozin | 273786.3276 | 2784467.808 | 6510212.643 | 1040549.335 | 200475.1979 | 189370.9208 | 958456.4788 | 250348.7902 | 227502.3332 | 1454373.206 | 333438.1112 | 2190283.61 |
| (2Z)-4,6-Di(1-piperidinyl)-1,3,5-triazin-2(3H)-one hydrazone | 2599886.448 | 4522407.674 | 2576178.129 | 1433042.365 | 3593760.209 | 2484857.842 | 4538971.777 | 4260883.949 | 2698639.412 | 3582997.254 | 11383416.09 | 3339339.03 |
| LPG O-17:2 | 151606 | 151234 | 156276 | 137892 | 102328 | 114689 | 128273 | 103442 | 162230 | 113253 | 165043 | 853702. |

| Metabolites | PBS_1 | PBS_2 | PBS_3 | PBS_4 | BCG_1 | BCG_2 | BCG_3 | BCG_4 | BCGΔ*pe18*_1 | BCGΔ*pe18*_2 | BCGΔ*pe18*_3 | BCGΔ*pe18*_4 |
|---|---|---|---|---|---|---|---|---|---|---|---|---|
| | 6.772 | 8.273 | 0.925 | 5.862 | 7.694 | 8.884 | 9.149 | 5.919 | 7.075 | 3.322 | 4.503 | 4998 |
| SMGDG O-28:7_10:0 | 2267834.325 | 1290409.911 | 614802.4896 | 2271787.856 | 1143220.304 | 1419395.774 | 982313.9947 | 1373738.344 | 1437405.421 | 1352654.167 | 997488.4029 | 2492319.27 |
| 15(S)-15-Methylprostaglandin F2.alpha. isopropyl ester | 1915287.556 | 2046574.44 | 1643409.146 | 2611304.504 | 1898544.476 | 2392555.323 | 2055977.324 | 2004984.554 | 2708432.513 | 2804686.225 | 2002478.155 | 2789868.016 |
| Benzonitrile, 2-[[(6-bromo-3H-imidazo[4,5-b]pyridin-2-yl)thio]methyl]- | 24758836.98 | 23522192.16 | 28400594.25 | 26696838.46 | 17022832.15 | 20121712.17 | 16412480.12 | 18906279.92 | 1946665 8.5 | 20182023.34 | 23799905.75 | 27203764.01 |
| CerP 15:0;2O/24:4 | 25627959.76 | 22454188.62 | 22447726.65 | 23587360.04 | 9853223.744 | 17126511.11 | 10207298.38 | 12974325.17 | 14143453.93 | 11161375.15 | 15249268.65 | 16401875.51 |
| 11-Dehydrocorticosterone | 2316178134 | 3088819226 | 3223295793 | 2671088591 | 1203989328 | 1303960890 | 1221777660 | 1369151432 | 1092795311 | 1165078358 | 1203074393 | 4891398570 |
| CerP 19:1;2O/2:0 | 10595466.83 | 12123604.45 | 12865887.17 | 12333178.35 | 13776979.56 | 14897842.23 | 12462902.09 | 11780489.28 | 10941422.68 | 11568494.71 | 21057121.24 | 16940823.24 |
| Tetraethylene glycol | 47172679.19 | 53703747.51 | 46430254.79 | 60031391.61 | 58522759.62 | 39517309.77 | 44292031.35 | 37206811.56 | 114787027 | 48745215.01 | 55074266.26 | 65642969.21 |
| PC 17:0_18:2 | 962241.8673 | 1272767.397 | 2017820.035 | 1617483.127 | 787439.4121 | 48314.15498 | 1963597.356 | 4219727.625 | 1716180.024 | 3162727.483 | 1535572.478 | 3126004.784 |
| 1-Stearoyl-2-docosahexaenoyl-sn-glycero-3-phosphocholine | 3485526.751 | 2130562.756 | 1066954.716 | 3331343.997 | 1476063.107 | 1638535.139 | 1405867.526 | 1612627.725 | 5881080.352 | 1945088.83 | 1462248.165 | 1256454.302 |

| Metabolites | PBS_1 | PBS_2 | PBS_3 | PBS_4 | BCG_1 | BCG_2 | BCG_3 | BCG_4 | BCGΔ*pe18*_1 | BCGΔ*pe18*_2 | BCGΔ*pe18*_3 | BCGΔ*pe18*_4 |
|---|---|---|---|---|---|---|---|---|---|---|---|---|
| 2,3-Dihydroxy-2-methylbutanoic acid | 1061281.981 | 1466665.74 | 1084315.887 | 1109273.304 | 13252237.7 | 7770403.613 | 12235700.91 | 3880763.575 | 14458397.08 | 15532459.72 | 10648680.6 | 8086943.479 |
| Ginsenoside F3 | 28075177.37 | 29406791.64 | 12049481.82 | 23798945.87 | 8977617.351 | 8968432.823 | 9084071 | 10115151.45 | 16915481.47 | 11009626.34 | 7565032.851 | 8879001.976 |
| SM 12:0;2O | 91353.48975 | 78302.18699 | 113945.3187 | 125865.7205 | 161079.7445 | 88836.07568 | 159243.7111 | 136853.168 | 145203.7608 | 226354.3941 | 160773.3417 | 196778.539 |
| Ribose | 42514152.7 | 26725502.98 | 37444730.79 | 35427989.75 | 11530695.68 | 2201706.403 | 13986506.84 | 2174199.123 | 18959147.37 | 12120750.4 | 11492388.41 | 16274654.13 |
| C8-homoserine lactone | 1040423.891 | 2214064.261 | 2235246.128 | 2101467.288 | 1092369.9 | 1023308.74 | 1349425.281 | 1004095.486 | 1110636.271 | 1204412.559 | 945233.2013 | 2664209.212 |
| Cer 26:2;2O/13:1;(2OH) | 56119780.96 | 51974296.75 | 50344012.32 | 52393395.93 | 22423934.51 | 35608053.13 | 23417520.52 | 30309638.55 | 30361828.43 | 27088676.38 | 33280752.25 | 35370914.64 |
| PI 16:0_20:4 | 1834601.044 | 2228462.229 | 1091743.553 | 2586768.115 | 2221546.57 | 1792254.597 | 2447985.088 | 2174944.3 | 2579036.796 | 2943909.554 | 2571652.166 | 2450127.174 |
| Pyridoxamine | 238577.6417 | 4311060.509 | 3197387.068 | 3367151.261 | 468917.8116 | 228113.4643 | 442402.7092 | 346553.6755 | 782265.6062 | 2258000.497 | 277177.7358 | 5309849.309 |
| PG 16:1_18:1 | 1206417.45 | 1232169.247 | 768732.0593 | 1198810.647 | 28327198.35 | 16479726.08 | 26347896.91 | 13702337.68 | 24347387.74 | 37329639.81 | 23721733.66 | 12418678.04 |
| HEPES | 3139279.212 | 3387193.251 | 41909758.81 | 3937335.762 | 5295594.149 | 3826726.045 | 60121144.69 | 5780112.354 | 3412647.483 | 35752424.88 | 40501534.85 | 3942498.708 |
| PG O-21:1_20:4 | 663656.2354 | 887842.7102 | 922161.8328 | 692727.3133 | 812163.2159 | 300486.1572 | 853109.1621 | 464611.0841 | 907699.4554 | 590134.8027 | 830057.999 | 477556.8051 |

| **Metabolites** | **PBS_1** | **PBS_2** | **PBS_3** | **PBS_4** | **BCG_1** | **BCG_2** | **BCG_3** | **BCG_4** | **BCGΔ*pe18*_1** | **BCGΔ*pe18*_2** | **BCGΔ*pe18*_3** | **BCGΔ*pe18*_4** |
|---|---|---|---|---|---|---|---|---|---|---|---|---|
| N,N-Dimethyl-2,2-diphenylethanamine | 3917177.25 | 2813452.859 | 5536848.99 | 2944995.524 | 4875473.794 | 3716234.453 | 4442789.995 | 3562696.511 | 3612062.559 | 3874764.323 | 4321400.005 | 7479796.808 |
| SHexCer 36:2;3O | 109895282.8 | 110240264.2 | 52226646.81 | 97739824.8 | 180809910.7 | 55663441.02 | 174505793.3 | 125238332.3 | 215556846.9 | 148395530 | 119511039.4 | 115358441.7 |
| Beclomethasone | 1976724.715 | 1934491.881 | 2125496.169 | 2177175.403 | 1883815.409 | 1681002.659 | 1850188.052 | 1750980.533 | 1989777.996 | 1904490.693 | 2172930.613 | 2176338.147 |
| LNAPS 20:1/N-14:1 | 1093865.116 | 861453.3486 | 1633118.302 | 1355371.048 | 1595354.483 | 789591.0995 | 2041265.273 | 1841354.903 | 2086286.527 | 1962079.622 | 2361316.675 | 1158864.128 |
| 1-Nonadecanoyl-glycero-3-phosphoserine | 1942257.29 | 2015554.71 | 2122404.925 | 2100207.666 | 2005705.69 | 2226003.117 | 2126850.339 | 2207738.66 | 3649793.233 | 3265776.986 | 3356711.779 | 3447239.171 |
| PI O-24:5_26:7 | 0 | 0 | 0 | 0 | 62336.87433 | 48929.74166 | 85885.28136 | 83751.99736 | 121005.3953 | 104520.2585 | 121954.0201 | 78605.05335 |
| LNAPE 18:2/N-18:0 | 3777387.533 | 2583908.999 | 3792912.478 | 3083806.496 | 2762543.171 | 65762.77719 | 2503231.828 | 1186881.79 | 2749243.263 | 2800620.668 | 2723569.005 | 1218605.216 |
| 3-(2-Ethylhexoxy)propan-1-amine | 12436602.43 | 18281593.17 | 8429527.276 | 10893410.83 | 7575067.074 | 19302919.87 | 14021866.57 | 9359070.497 | 14345572.52 | 7356755.806 | 10331686.93 | 23646739.62 |
| Ile-Ile | 1317051.972 | 1144363.278 | 1651946.976 | 1098799.942 | 1250518.771 | 740832.0051 | 1050860.633 | 804284.594 | 1338659.635 | 1509650.789 | 1719942.259 | 1452668.403 |
| PE O-16:1_2:0 | 2836641.8 | 3050718.704 | 2562685.64 | 1866574.134 | 4165581.475 | 3213009.461 | 2333741.025 | 2552038.368 | 3469300.271 | 5137703.113 | 4364469.412 | 2785742.56 |
| PE(22:5(4Z,7Z,10Z,13Z,16Z)/16:1(9Z)) | 9401183.996 | 12356574.15 | 15121600.64 | 11162419.44 | 26136701.88 | 20452526.84 | 22428239.54 | 25773782.41 | 30128191.49 | 39606973.79 | 30613626.96 | 22863074.07 |

| **Metabolites** | **PBS_1** | **PBS_2** | **PBS_3** | **PBS_4** | **BCG_1** | **BCG_2** | **BCG_3** | **BCG_4** | **BCGΔ*pe18*_1** | **BCGΔ*pe18*_2** | **BCGΔ*pe18*_3** | **BCGΔ*pe18*_4** |
|---|---|---|---|---|---|---|---|---|---|---|---|---|
| PS 37:1 | 0 | 0 | 0 | 42332.26453 | 2492193.547 | 2574135.926 | 2714956.263 | 2103886.18 | 2943618.666 | 3360593.858 | 2778487.947 | 2075762.985 |
| PE(18:3(9Z,12Z,15Z)/15:0) | 400749.5552 | 413804.9188 | 177824.5839 | 228115.4635 | 1222299.486 | 1407045.577 | 1967460.447 | 2193230.573 | 2109071.98 | 3982502.792 | 1261343.76 | 1108055.521 |
| 1-Hexadecyl-2-(8Z,11Z,14Z-eicosatrienoyl)-sn-glycero-3-phosphocholine | 158641115.1 | 211253222.6 | 89028406.92 | 121400975.6 | 97195580.86 | 67606076.87 | 100267228.2 | 87966052.25 | 117644708.6 | 126019883.6 | 89583220.28 | 61462528.32 |
| CYCLOSERINE | 9636367.775 | 10528603.51 | 9521518.368 | 10146800.87 | 9502298.726 | 9518692.241 | 9857156.996 | 9229202.545 | 10162130.85 | 15525279.19 | 9420735.372 | 9560647.872 |
| PG 17:1_18:1 | 608652.5003 | 532659.0028 | 484357.7147 | 634303.6029 | 10219126.71 | 6618404.343 | 13762373.75 | 5090464.304 | 10030604.63 | 15148844.04 | 11293857.57 | 3081660.192 |
| PG 20:4_22:4 | 157539895.3 | 249614089.3 | 78271448.82 | 144381500.2 | 29786680.55 | 18771521.25 | 29666264.58 | 12877470.29 | 16018708.56 | 58960680.56 | 30640808.71 | 11919061.26 |
| Phosphonic acid, P-[[4-[(1-oxotetradecyl)amino]phenyl]methyl]- | 16944413.17 | 18611646.15 | 21039402.72 | 20284933.06 | 20798333.82 | 21153059.2 | 19871983.98 | 18601007.28 | 17136474.63 | 17903553.6 | 30672723.17 | 24068871.7 |
| 2-Hydroxy-4,6-dimethylpyrimidine | 3014528.233 | 6122612.208 | 5386812.052 | 5302651.616 | 2234379.774 | 6570397.002 | 9628373.071 | 5386787.858 | 6482636.98 | 6191844.789 | 6766483.362 | 7025578.106 |
| Cytochalasin J | 110960.0116 | 52204.75184 | 66884.93457 | 57162.04787 | 667947.6581 | 408962.1609 | 554201.7388 | 521060.9194 | 698458.036 | 889867.9784 | 727569.0979 | 468317.7865 |
| PG 20:4_20:4 | 4749455.643 | 5475302.426 | 2534741.333 | 13056255.21 | 867992.6122 | 673681.827 | 1263241.659 | 1231821.327 | 1327446.324 | 1636339.36 | 973061.2967 | 878711.0044 |

| Metabolites | PBS_1 | PBS_2 | PBS_3 | PBS_4 | BCG_1 | BCG_2 | BCG_3 | BCG_4 | BCGΔ*pe18*_1 | BCGΔ*pe18*_2 | BCGΔ*pe18*_3 | BCGΔ*pe18*_4 |
|---|---|---|---|---|---|---|---|---|---|---|---|---|
| N-palmitoyl-aspartate | 3857384.456 | 4667608.699 | 1907947.719 | 3255007.548 | 4489237.634 | 3620788.966 | 5634422.214 | 3864464.014 | 5151184.315 | 5232571.93 | 4418761.339 | 4920034.129 |
| Propanedioic acid, 2-(2-methylpropyl)-, 1,3-dimethyl ester | 6701487.362 | 7039442.832 | 5769240.718 | 11238599.09 | 5145449.1 | 7114807.049 | 13689143.18 | 5085067.588 | 9759112.119 | 8143539.819 | 4949448.78 | 13626018.48 |
| PE O-16:4_4:0 | 93203874.86 | 118726053.5 | 129286397.9 | 106081889.4 | 99929264.13 | 88783238.54 | 95519752.08 | 93288095.24 | 87771538.93 | 122353195.5 | 113287797.9 | 101711638.1 |
| LPE 18:5 | 56550.36235 | 73057.40842 | 71143.15877 | 100853.2642 | 1218084.006 | 890357.1371 | 870969.5441 | 628058.0599 | 1182193.307 | 1345924.23 | 1137470.585 | 960613.7572 |
| PC O-26:6_5:0 | 18209645.17 | 15880499.18 | 15046305.42 | 16812179.81 | 5789934.947 | 10939392.86 | 7226076.251 | 8635236.017 | 8781604.659 | 6905284.548 | 9861234.039 | 11253370.45 |
| Cer 8:1;2O/4:0 | 36622241.26 | 40232100.8 | 40767199.78 | 41073037.17 | 37679476.39 | 34535181.71 | 38259519.07 | 31655976.98 | 34827246.49 | 38204416.97 | 36202130.67 | 51141907.51 |
| 4,6-Bis(4-fluorophenyl)-2-methyl-5-(4-pyridyl)pyrazolo[3,4-b]pyridine | 13003234.1 | 13717519.96 | 15168697.82 | 13259986.04 | 21896870.04 | 17340478.52 | 23949354.18 | 18690571.55 | 23338316.27 | 25543441.93 | 24214298.31 | 20849098.74 |
| 4-Hydroxy-2,2',5'-trichlorobiphenyl | 1540061.099 | 848004.7581 | 2322034.51 | 1285856.825 | 18817163.35 | 12368123.79 | 11691742.22 | 18018778.44 | 11385195.78 | 13511334.9 | 38876099.86 | 7342614.375 |
| PC 14:0_18:1 | 3953478.095 | 3049860.911 | 6146130.667 | 3288722.381 | 3637661.244 | 497790.6337 | 3332555.746 | 2082296.535 | 3500500.904 | 3850945.399 | 4224942.627 | 1518063.456 |
| LPS 24:4 | 15041983.74 | 15494462.01 | 9439484.252 | 14251513.31 | 16035835.75 | 15534794.02 | 13716006.77 | 12991882.79 | 21623803.41 | 18174432.33 | 16748352.53 | 13567535.9 |

| **Metabolites** | **PBS_1** | **PBS_2** | **PBS_3** | **PBS_4** | **BCG_1** | **BCG_2** | **BCG_3** | **BCG_4** | **BCGΔ*pe18*_1** | **BCGΔ*pe18*_2** | **BCGΔ*pe18*_3** | **BCGΔ*pe18*_4** |
|---|---|---|---|---|---|---|---|---|---|---|---|---|
| 4-Iodosuberoylanilide hydroxamic acid | 24646098.01 | 14663458.84 | 22961806.37 | 13189930.29 | 21592123.24 | 25740219.64 | 15655892.2 | 11339705.43 | 19989684.49 | 14473272.98 | 25989880.07 | 21976792.7 |
| 2',4'-Dihydroxy-2-methoxychalcone | 67199630.88 | 44922985.62 | 75407120.91 | 49083372.37 | 38695742.81 | 54893770.28 | 25569347.83 | 1922757.956 | 9671982.826 | 13240112.11 | 9604667.398 | 98954280.54 |
| Umbelliferone | 7000810.124 | 8632597.359 | 8206714.931 | 9358067.592 | 10937096.55 | 9468959.198 | 10092065.37 | 8376228.008 | 8434144.96 | 10144873.24 | 8258517.254 | 18287902.21 |
| LPG 22:5 | 5075976.772 | 5152560.013 | 34894733.21 | 5746666.271 | 2986267.825 | 2900972.744 | 6419252.265 | 3795854.968 | 2951935.662 | 5992455.833 | 5735980.268 | 3312307.854 |
| 2-Chloro-3-chloromethyl-6-fluoroquinoline | 2791661.718 | 2712914.969 | 2938953.585 | 2720060.789 | 2856127.027 | 2474192.894 | 1962006.355 | 2809167.538 | 2758733.957 | 2967105.943 | 2958631.927 | 2700720.595 |
| D-myo-Inositol-3,4,5-triphosphate | 56916905.83 | 56160112.7 | 38425652.52 | 46815566.1 | 41861921.53 | 52791144.42 | 28642515.58 | 39918045.48 | 36593109.19 | 49581243.57 | 60184394.6 | 35246472.7 |
| Dipyridamole_(Persantine) | 2544645.188 | 1036949.041 | 2607382.538 | 906955.2052 | 906131.1901 | 1202006.524 | 1019458.271 | 1136411.845 | 1306509.94 | 1823664.299 | 2737601.658 | 1607067.396 |
| Sulfaphenazole | 481894.3645 | 890759.1029 | 1881086.006 | 2123382.612 | 494883.6537 | 287986.7892 | 586809.3993 | 567112.8255 | 680250.4644 | 612663.5709 | 505860.1981 | 2045764.397 |
| 2,4,5,6-Pyrimidinetetramine | 48045412.02 | 41289499.77 | 41870482.28 | 28463875.64 | 51280610.2 | 29931792.91 | 31724683.32 | 49676146.4 | 58850826.98 | 32464085.98 | 34485692.71 | 54777967.99 |
| PMeOH 16:1_26:4 | 386154.3455 | 339688.6763 | 300033.5797 | 430095.334 | 510244.0062 | 479826.4491 | 635038.5081 | 526656.7153 | 646805.688 | 642188.987 | 606550.0565 | 617710.4702 |
| cis-3-Amino-2-pyrrolidinecarboxylic acid | 124972.5851 | 180038.0545 | 499483.1394 | 446022.3388 | 994218.5853 | 1807119.897 | 894843.0535 | 1014888.621 | 1643251.511 | 926435.0624 | 1482857.321 | 2246461.887 |

| **Metabolites** | **PBS_1** | **PBS_2** | **PBS_3** | **PBS_4** | **BCG_1** | **BCG_2** | **BCG_3** | **BCG_4** | **BCGΔ*pe18*_1** | **BCGΔ*pe18*_2** | **BCGΔ*pe18*_3** | **BCGΔ*pe18*_4** |
|---|---|---|---|---|---|---|---|---|---|---|---|---|
| Ethyl 3-(piperidin-4-yl)propanoate | 11184666.25 | 19893571.52 | 6405073.427 | 12248757.27 | 9526522.509 | 7601372.063 | 9193777.148 | 7885890.616 | 9948044.575 | 10175650.55 | 8133199.545 | 10145746.4 |
| Docebenone | 151285939.3 | 307651330.3 | 357800964.8 | 221626133.5 | 59964424.08 | 56891086.51 | 60811569.16 | 53779300.9 | 19095011.07 | 17239136.98 | 55547460.98 | 226179876.5 |
| PC O-25:2 | 1618391.236 | 1871936.209 | 1620482.879 | 1666585.843 | 1826084.644 | 1849916.887 | 1776501.03 | 1911925.96 | 2227683.537 | 2078524.47 | 1865369.565 | 1855030.81 |
| 2-[4-(2-Carboxy-5-oxopyrrolidin-1-yl)-6,6,9a-trimethyl-5-[(2E,4E,6E)-octa-2,4,6-trienoyl]oxy-1-oxo-4,5,5a,7,8,9-hexahydro-3H-benzo[e]isoindol-2-yl]pentanedioic acid | 818091.8955 | 902811.5463 | 830181.638 | 541261.9967 | 313742.5902 | 353220.239 | 163302.3176 | 397446.4295 | 321511.8064 | 383511.3235 | 446045.2858 | 429399.3786 |
| Ifenprodil | 505137.0512 | 366907.0777 | 488941.4389 | 881632.6656 | 2438604.516 | 1049351.816 | 1352481.015 | 1175652.575 | 1845546.546 | 1927080.685 | 3116353.113 | 1678916.958 |
| PG 19:2_22:6 | 2275190.751 | 1036738.993 | 761408.0476 | 1936024.963 | 2001727.016 | 2170127.725 | 2068942.04 | 1835085.899 | 2004198.629 | 2345854.252 | 2462923.239 | 2102113.784 |

| Metabolites | PBS_1 | PBS_2 | PBS_3 | PBS_4 | BCG_1 | BCG_2 | BCG_3 | BCG_4 | BCGΔ*pe18*_1 | BCGΔ*pe18*_2 | BCGΔ*pe18*_3 | BCGΔ*pe18*_4 |
|---|---|---|---|---|---|---|---|---|---|---|---|---|
| 5,7-Dihydroxy-2-(4-hydroxyphenyl)-3-[3,5,7-trihydroxy-2-(4-hydroxyphenyl)-3,4-dihydro-2H-chromen-8-yl]-2,3-dihydrochromen-4-one | 1014971.251 | 1072337.57 | 765000.2803 | 1285644.594 | 1135604.741 | 1119738.828 | 757440.7601 | 1116681.515 | 1273678.647 | 1343758.623 | 1282477.65 | 1000886.449 |
| 1-Methylpyrrolidine | 41944363.84 | 16922715.09 | 53422462.77 | 41664107.17 | 25178620.9 | 22785796.53 | 23803297.53 | 41581802.06 | 53644125.77 | 25490336.34 | 34805214.06 | 40020334.56 |
| N-(9-Oxodecyl)acetamide | 12405186.66 | 15802111.13 | 11684023.97 | 13772601.6 | 11338253.42 | 9682233.681 | 9184195.358 | 8962493.727 | 9281232.078 | 10860891.11 | 9031181.409 | 13373284.68 |
| Nitazoxanide | 1510203.104 | 1394226.033 | 1594049.652 | 1266733.481 | 2615210.252 | 2377454.037 | 3204913.922 | 3171593.748 | 3091174.71 | 3156254.33 | 2928959.651 | 3769096.76 |
| 4-(2,4-Difluorophenyl)piperidine | 442047.5549 | 697125.0512 | 218120.7576 | 3550861.216 | 330531.9083 | 545562.8129 | 1124450.475 | 75612.06445 | 1176471.011 | 765214.5819 | 150603.4278 | 2391044.118 |
| 4-(Aminomethyl)-L-phenylalanine | 734353.4518 | 842957.9894 | 900400.5089 | 1343188.265 | 950578.6539 | 784540.4589 | 916058.2657 | 911620.4861 | 1054985.437 | 864596.6717 | 924531.3279 | 1085099.73 |
| Cer 14:2;2O/15:1;(2OH) | 41181385.54 | 51477809.83 | 22893066.01 | 37590106.54 | 21351832.98 | 21302853.27 | 22855337.31 | 23603975.07 | 28095941.66 | 27848311.69 | 23768044.95 | 18213097.89 |
| SL 11:1;O/20:2;O | 9710495.539 | 6924923.752 | 6400564.221 | 8515814.725 | 6645950.947 | 7112606.15 | 6384933.33 | 4852398.81 | 5848672.782 | 7674091.044 | 9578685.996 | 4778497.202 |

| Metabolites | PBS_1 | PBS_2 | PBS_3 | PBS_4 | BCG_1 | BCG_2 | BCG_3 | BCG_4 | BCGΔ*pe18*_1 | BCGΔ*pe18*_2 | BCGΔ*pe18*_3 | BCGΔ*pe18*_4 |
|---|---|---|---|---|---|---|---|---|---|---|---|---|
| 5,8-Dihydroxy-6-methoxy-2-oxo-2H-chromen-7-yl .beta.-D-glucopyranoside | 148075.0388 | 160513.9851 | 245799.017 | 187212.3681 | 16587221.43 | 18168220.78 | 14970085.33 | 15509986.2 | 17661753.97 | 15392932.2 | 16880292.31 | 20018613.57 |
| PI 18:1_22:4 | 555736.0063 | 440884.6461 | 1576104.403 | 496642.214 | 709470.6239 | 62477.4685 | 528505.5607 | 737211.3168 | 708436.6633 | 602654.3167 | 572538.9926 | 508867.2602 |
| (1,3,5-Trimethyl-1H-pyrazol-4-yl)methanol | 71216154.11 | 75392327.2 | 198312838.8 | 297453410.9 | 362398016.5 | 100633429.3 | 186825763 | 166476288.4 | 235999582.2 | 286828205.7 | 55291819.35 | 303318973 |
| Methyprylon | 6718135.162 | 7732503.286 | 5831594.004 | 11551443.9 | 7021272.06 | 6631821.383 | 8971657.414 | 6164208.116 | 8111712.083 | 7855511.554 | 6540929.681 | 10491712.28 |
| Acetic acid N',N'-dimethylhydrazide | 413345.7953 | 623349.9013 | 1259400.112 | 852651.0349 | 465711.0337 | 467403.6766 | 133809.1325 | 386106.648 | 410998.3154 | 338312.3261 | 459896.9103 | 1387442.884 |
| Epigallocatechin | 21324790.37 | 20404359.2 | 20983605.77 | 19966056.4 | 36267110.73 | 33186451.15 | 46834512.16 | 47160877.62 | 41794209.85 | 42106026.01 | 40870147.16 | 51354781.02 |
| NAOrn 22:6/17:2 | 13517861.23 | 7105477.787 | 15201031.43 | 3485186.898 | 8295304.918 | 0 | 2691173.928 | 3029638.02 | 3422180.469 | 3878371.468 | 8733665.952 | 1936858.701 |
| 4-Methyl-3-(trifluoromethyl)benzoic acid | 17306087.95 | 11332104.76 | 11841939.35 | 13223791.45 | 9271694.137 | 15873243.6 | 12765987.76 | 9351743.043 | 11836755.67 | 11743646.62 | 11682781.04 | 15994222.08 |
| 1,7-Dihydroxyxanthone | 2508541.602 | 2464162.995 | 2210247.06 | 1385513.453 | 2429616.092 | 1607386.014 | 1282470.376 | 1237223.506 | 1966570.959 | 1526192.969 | 1669311.885 | 2115501.309 |

| Metabolites | PBS_1 | PBS_2 | PBS_3 | PBS_4 | BCG_1 | BCG_2 | BCG_3 | BCG_4 | BCGΔ*pe18*_1 | BCGΔ*pe18*_2 | BCGΔ*pe18*_3 | BCGΔ*pe18*_4 |
|---|---|---|---|---|---|---|---|---|---|---|---|---|
| N-Cyclohexyl-N'-(3,5-dimethyl-1H-pyrazol-4-yl)thiourea | 6802828.515 | 6265933.619 | 6071913.302 | 6116988.622 | 6473508.32 | 7597504.223 | 6815868.842 | 6384262.504 | 7764141.765 | 6762186.259 | 6409845.113 | 16058965.21 |
| LPE 22:4 | 5412222.335 | 5702707.192 | 11346254.75 | 6289746.772 | 8440867.981 | 8747573.855 | 10198066.66 | 8017948.287 | 6813222.12 | 13266002.13 | 11927229.18 | 10304564.16 |
| Deoxyhypusine | 7325170.11 | 8998720.124 | 7503927.128 | 8548898.934 | 8080473.357 | 8117954.481 | 9243256.179 | 6855123.697 | 8832199.833 | 9121570.656 | 7576998.303 | 9118099.298 |
| Saccharin | 4977423.002 | 8670398.429 | 14409046.54 | 8167852.167 | 4419774.868 | 4052031.907 | 5362718.361 | 3697932.021 | 5727700.82 | 4939995.671 | 4953006.791 | 12362659.53 |
| 2-Amino-N-(4-fluorophenyl)benzamide | 1336313.015 | 1478499.882 | 1286545.059 | 1364707.751 | 1365227.676 | 68490.97404 | 1385153.817 | 1361926.951 | 1352749.476 | 1269249.707 | 1368458.94 | 1516831.46 |
| DGCC 16:3_20:5 | 1260360.552 | 1878121.44 | 1686270.241 | 1493715.955 | 2257383.672 | 3000546.93 | 2295886.203 | 2456261.635 | 2436063.593 | 2657502.173 | 4224333.778 | 2025730.442 |
| PE O-20:4_19:0 | 98426969.56 | 68276464.09 | 109468335.3 | 115738372.5 | 89523303.54 | 1014092.958 | 158488033.8 | 147848177.2 | 108580884.8 | 95390626.71 | 75854284.99 | 146998450.9 |
| 5-(Ethoxymethyl)-2-methyl-4-pyrimidinamine | 4278884.161 | 3977745.143 | 7915493.453 | 4913717.045 | 342949.2647 | 247898.4732 | 1070081.607 | 251570.5838 | 895340.6693 | 1102369.746 | 4919502.601 | 8049559.614 |
| 3,3'-Dimethoxybenzidine | 149404.4365 | 165836.5836 | 151948.5097 | 171138.8125 | 134072.2538 | 113421.4824 | 137586.9115 | 129719.7842 | 121086.3939 | 148366.4696 | 115720.0731 | 197840.9419 |
| Pentadecylamine | 15840010.15 | 20983592.41 | 25167055.74 | 20926235.82 | 16040527.1 | 15617032.86 | 14811607.03 | 14925682.95 | 17058377.84 | 15759995.76 | 15050316.9 | 36244876.81 |
| 2-(Diisopropylamino)ethyl | 3369571.398 | 4034067.02 | 2702097.786 | 3827392.082 | 3919040.744 | 4375628.258 | 10320376.16 | 3516207.891 | 7033537.966 | 6882508.37 | 3846497.817 | 11074784.77 |

| **Metabolites** | **PBS_1** | **PBS_2** | **PBS_3** | **PBS_4** | **BCG_1** | **BCG_2** | **BCG_3** | **BCG_4** | **BCGΔ*pe18*_1** | **BCGΔ*pe18*_2** | **BCGΔ*pe18*_3** | **BCGΔ*pe18*_4** |
|---|---|---|---|---|---|---|---|---|---|---|---|---|
| methacrylate | | | | | | | | | | | | |
| PE O-16:1_22:5 | 6230573.957 | 3341320.884 | 5615513.105 | 4296831.4 | 4156037.439 | 25934.17594 | 4270975.166 | 3252367.82 | 6491280.411 | 4290672.856 | 3473624.866 | 2306178.786 |
| 3-(5-Amino-3-methyl-1H-pyrazol-1-yl)propanenitrile | 20644158 | 22078427.96 | 19836996.45 | 27944378.28 | 22928329.23 | 21807250 | 20696826.73 | 20283428.32 | 21314788.43 | 25929741.69 | 22057440.75 | 24477553.98 |
| PC(30:0) | 4277056.013 | 3299964.437 | 3409079.426 | 2498734.023 | 743619.1674 | 10583922.54 | 18377286.19 | 12054074.71 | 9029951.841 | 17163910.52 | 14255260.61 | 9056122.057 |
| Methanedisulfonic acid | 34792757.45 | 34617985.86 | 36072627.88 | 37326360.93 | 36308968.07 | 34930742.12 | 40855828.42 | 39634057.21 | 42448529.81 | 43211850.05 | 35770541.97 | 39459353.43 |
| M239T381 | 4587121.381 | 2774190.059 | 3260579.282 | 3093357.094 | 3897053.858 | 3863084.872 | 3513015.631 | 1976182.917 | 3966135.042 | 4457517.83 | 3132228.503 | 3901441.732 |
| HexCer 24:2;3O/15:0;(2OH) | 928964.2779 | 1003177.306 | 951683.9436 | 1723147.882 | 620029.9452 | 489306.2218 | 1015434.371 | 493157.805 | 1349737.968 | 1170979.465 | 744550.0369 | 362443.891 |
| Betaine | 33759378.99 | 34284156.92 | 41705843.46 | 32923402.88 | 113644261.8 | 90279520.22 | 106139189.9 | 90235930.45 | 146494488.7 | 126825061.5 | 124412810 | 120247200.9 |
| Icaridin | 14421106.22 | 13275582.16 | 18314726.62 | 16960322.12 | 13725920.44 | 14751905.6 | 12586934.24 | 9853480.527 | 14792707.61 | 11286911.88 | 13837987.52 | 20479219.98 |
| Uzarigenin | 23401823.99 | 24577216.01 | 22300536.49 | 22991531.77 | 22755292.96 | 24025739.77 | 23062766.53 | 23038003.13 | 29004597.05 | 24885947.79 | 23459869.58 | 21002830.43 |
| Diethanolamine | 107493283.6 | 87022393.91 | 90201081.35 | 114765471.6 | 65706723.91 | 74798902.73 | 50528812.21 | 46916857.04 | 66047648.71 | 87419238.46 | 75969152.07 | 66170892.72 |

| Metabolites | PBS_1 | PBS_2 | PBS_3 | PBS_4 | BCG_1 | BCG_2 | BCG_3 | BCG_4 | BCGΔ*pe18*_1 | BCGΔ*pe18*_2 | BCGΔ*pe18*_3 | BCGΔ*pe18*_4 |
|---|---|---|---|---|---|---|---|---|---|---|---|---|
| N-Nervonoyl-D-erythro-sphingosylphosphorylcholine | 19923047.71 | 15723334.25 | 34651241.35 | 77630843.32 | 23855307.14 | 18964737.43 | 13688918.79 | 32260503.52 | 24815026.84 | 19053182.57 | 4892835.396 | 48049218.07 |
| Mammea_E/BB | 5606778.015 | 784622.7921 | 1321276.625 | 1374891.994 | 1446431.742 | 7109126.843 | 1182434.372 | 2195456.092 | 5161607.718 | 1190911.802 | 6554854.473 | 838842.2001 |
| (Dimethoxymethyl)benzene | 7082469.406 | 5615522.38 | 5465969.427 | 48024773.22 | 1813177.518 | 8517431.946 | 72180887.12 | 1895454.09 | 31736781.62 | 15033415.59 | 2088520.303 | 51183240.61 |
| ST 24:1;O4;T/17:2 | 1519671.706 | 1569528.635 | 1406028.403 | 892895.1295 | 4769259.126 | 3807856.368 | 4322845.062 | 4885093.275 | 4374050.509 | 6195803.545 | 5805123.266 | 2929731.372 |
| 1-Piperidinepropanol, .alpha.-(4-hydroxyphenyl)-.beta.-methyl-4-(phenylmethyl)-, (.alpha.R,.beta.S)- | 4653463.667 | 6053714.478 | 2044163.641 | 4171504.778 | 2512598.362 | 3624041.577 | 2683041.445 | 3709857.882 | 2235427.932 | 3439595.612 | 3210882.524 | 5443979.322 |
| Arg-Gly-Arg | 499298.2378 | 336803.9217 | 537665.2105 | 1690282.473 | 355651.1448 | 1119245.812 | 2800343.987 | 446770.5124 | 996011.3933 | 1684001.143 | 656182.9984 | 1750945.019 |
| CAR 16:3 | 1773452.859 | 1711616.349 | 2719429.236 | 2190232.575 | 207996.3807 | 295536.4609 | 384145.8713 | 361238.5113 | 298304.3191 | 530151.4605 | 318835.7071 | 328521.7951 |
| 3,4-Dimethylbenzaldehyde | 11574571.08 | 1920973.007 | 7776919.599 | 2370645.53 | 2171162.765 | 10376688.21 | 4559625.888 | 5491804.929 | 11384592.64 | 2751035.767 | 13155051.52 | 2830660.477 |
| 2'-Deoxyuridine | 7733942.538 | 6227497.391 | 13582155.13 | 8647473.058 | 43652464.02 | 28704815.83 | 41243925.91 | 33122445.01 | 42692570.03 | 42331563.71 | 48141536 | 35496052.94 |
| Sambutoxin | 645287. | 613004. | 559838. | 548452. | 181876 | 143192 | 181533 | 161227 | 195007 | 199844 | 197301 | 154926 |

| Metabolites | PBS_1 | PBS_2 | PBS_3 | PBS_4 | BCG_1 | BCG_2 | BCG_3 | BCG_4 | BCGΔ*pe18*_1 | BCGΔ*pe18*_2 | BCGΔ*pe18*_3 | BCGΔ*pe18*_4 |
|---|---|---|---|---|---|---|---|---|---|---|---|---|
| | 107 | 6592 | 3077 | 6599 | 9.088 | 3.637 | 0.698 | 0.872 | 2.55 | 1.176 | 8.409 | 7.636 |
| Pivagabine | 26121184.39 | 28900734.45 | 29359546.9 | 127428387.7 | 21654192.4 | 33592243.63 | 166024437.8 | 20363802.53 | 73790989.17 | 44634070.22 | 22252562.48 | 154263732.7 |
| 5-(2-Hydroxyethyl)-4-methylthiazole | 270454.5728 | 60897.57366 | 336494.9252 | 164999.2515 | 234247.8095 | 428599.9048 | 128148.301 | 280973.478 | 527786.6469 | 133288.8496 | 417816.4091 | 495839.1768 |
| 1-Hexadecyl-2-(9Z-octadecenoyl)-sn-glycero-3-phosphocholine | 10275403.88 | 4006243.527 | 2129300.563 | 1717428.323 | 11680336 | 4132692.6 | 9172585.426 | 2103441.875 | 2507480.907 | 14729287.9 | 6852972.569 | 6304230.959 |
| 4-Hydroxycyclofenil | 7817958.563 | 10491072.26 | 12240960.3 | 11200476.13 | 5250958.241 | 5539215.823 | 6535009.199 | 6517940.035 | 5831329.693 | 5287901.752 | 5325517.75 | 14403523.89 |
| NAGly 22:4/15:1 | 6549879.307 | 10003501.5 | 4836928.863 | 9957007.462 | 6055261.912 | 6514321.459 | 5575220.636 | 7666587.874 | 5161641.057 | 10768668.7 | 6809597.768 | 6318776.782 |
| NAGly 24:6/22:5 | 78496615.98 | 80720343.52 | 77631834.81 | 71042049.23 | 153927056.5 | 158581185.7 | 188312934.4 | 132860826.4 | 217037383.6 | 172839527.2 | 148997807.4 | 125797053.1 |
| PG 22:6_22:6 | 283473584.7 | 326562059.7 | 91838909.19 | 216908528.7 | 173241331.1 | 147987532.9 | 134276475.4 | 205173502.3 | 288385921.2 | 175149425.5 | 96668704.74 | 137228836.2 |
| Cer 8:0;2O/26:1 | 2079637.689 | 1519620.5 | 1836483.461 | 2257004.786 | 1731597.086 | 7020.597283 | 1843930.396 | 2137766.892 | 2583234.68 | 1916102.384 | 1599528.978 | 2075430.828 |
| 3-Aminononanoic acid | 940352.1437 | 971526.147 | 1010013.714 | 1086240.909 | 4491957.921 | 15192939.86 | 82908009.95 | 2726177.522 | 50148224.46 | 18768239.09 | 5107447.654 | 72429725.68 |
| 2-Chloro-N-(4H-1,2,4-triazol-3-yl)acetamide | 11762821.59 | 18678731.91 | 17983287.74 | 18332473.05 | 9070195.618 | 13918635.36 | 18962308.71 | 3667018.774 | 18909830.15 | 19063355.63 | 11297352.29 | 17765179.11 |

| Metabolites | PBS_1 | PBS_2 | PBS_3 | PBS_4 | BCG_1 | BCG_2 | BCG_3 | BCG_4 | BCGΔ*pe18*_1 | BCGΔ*pe18*_2 | BCGΔ*pe18*_3 | BCGΔ*pe18*_4 |
|---|---|---|---|---|---|---|---|---|---|---|---|---|
| 6-[(E)-2,8-Dihydroxy-5,7-dimethylnon-5-enyl]-4-hydroxy-3-methylpyran-2-one | 122087514.7 | 112917325.3 | 88092692.38 | 115709799 | 85729417.87 | 91355416.94 | 88735131.52 | 113253180.1 | 74805214.32 | 94488520.83 | 89923230 | 144973112.1 |
| 4-Chloro-5,6,7,8-tetrahydroquinazolin-2-amine | 49123841.92 | 53182893.83 | 54449458.61 | 53056638.35 | 47822488.71 | 49687425.69 | 48715670.09 | 47638594.32 | 47625418.59 | 46534312.82 | 47971056.3 | 69241966.64 |
| Midafotel | 152649.5193 | 108276.6557 | 245848.9244 | 162000.8003 | 1766955.133 | 1022800.732 | 2011853.879 | 1278092.339 | 1413086.863 | 1688094.54 | 1980578.946 | 1426167.13 |
| LPS 20:4 | 17587399.64 | 14028642.17 | 24410470.35 | 15513538.69 | 9357091.442 | 8324897.915 | 13933304.89 | 12256395.73 | 11415157.8 | 17791449.39 | 13998678.84 | 12478725.17 |
| Levulinic acid | 2554983.296 | 2391859.067 | 2326887.71 | 2223922.932 | 1924249.51 | 2232714.109 | 2273852.271 | 2263936.287 | 2030736.062 | 2260358.728 | 5055761.225 | 2511064.828 |
| 1,2-Diarachidonoyl-sn-glycero-3-phosphoethanolamine | 4929828.11 | 5487168.944 | 6387720.779 | 4274582.082 | 16415226.44 | 11674624.75 | 18748225.85 | 17596602.74 | 16963412.41 | 18801758.48 | 19126116.28 | 13438280.41 |
| N-Cyclohexyl-N'-(4-pyridinylmethyl)thiourea | 7989915.851 | 6317726.206 | 10625993.94 | 52625460.33 | 1288189.42 | 18714735.15 | 75841063.18 | 1023074.144 | 28311843.06 | 20861194.29 | 1133222.161 | 56182066.81 |
| 3-(Aminosulfonyl)-2,6-difluorobenzoic acid | 3208069.502 | 965903.7537 | 2715479.869 | 1160623.31 | 5716057.999 | 3109427.661 | 4445647.057 | 4847808.847 | 5898300.213 | 7269074.107 | 3666403.102 | 2751988.247 |

| Metabolites | PBS_1 | PBS_2 | PBS_3 | PBS_4 | BCG_1 | BCG_2 | BCG_3 | BCG_4 | BCGΔ*pe18*_1 | BCGΔ*pe18*_2 | BCGΔ*pe18*_3 | BCGΔ*pe18*_4 |
|---|---|---|---|---|---|---|---|---|---|---|---|---|
| N1-(3-Aminopropyl)-N1-methylpropane-1,3-diamine | 6700144.102 | 7114996.205 | 8627226.668 | 15246728.79 | 9380680.502 | 6018943.968 | 5022055.731 | 9103122.852 | 5853282.693 | 8192555.399 | 6925914.813 | 14853713.3 |
| PI 17:0_15:1 | 339920.492 | 448759.2823 | 176130.7317 | 492068.2197 | 1588442.552 | 1890565.161 | 1643876.081 | 1622326.005 | 1923555.545 | 1909186.16 | 1706359.763 | 1598657.476 |
| 4-Ethyl-5-hydroxy-3,5-dimethylfuran-2-one | 3065284.608 | 3356118.157 | 2352664.949 | 3242168.074 | 3341785.329 | 1523675.89 | 1515681.072 | 1584488.742 | 3203512.09 | 3209004.327 | 1427135.485 | 1419268.461 |
| PC(40:7) | 68965786.55 | 73996302 | 36002815.36 | 41921271.52 | 73919977.38 | 60953596.86 | 77551153.02 | 71114559.51 | 82091705.62 | 107743455.3 | 60040216.52 | 61559160.1 |
| 3-Amino-5,7-dimethyl-1-adamantanol | 11465177.8 | 13377504.67 | 8746404.578 | 8671276.707 | 10684383.44 | 10618857.39 | 10269742.75 | 10224344.37 | 11159258.78 | 10878484.63 | 11063216.39 | 12225942.22 |
| cis-2-tert-Butoxycarbonylamino-2-methylcyclohexanecarboxylic acid | 757793.7459 | 819448.612 | 843319.204 | 702152.4233 | 648784.7449 | 616808.654 | 666374.7462 | 615378.6486 | 625750.0014 | 696087.0795 | 590899.8768 | 931198.0376 |
| presphingofungin | 11501743.07 | 11238727.3 | 5366260.006 | 10487573.11 | 6970953.351 | 7811134.816 | 6446679.303 | 6395820.383 | 8040015.083 | 8112847.704 | 6633490.144 | 6628406.231 |
| S-Diclofenac | 13347586.93 | 13292236.8 | 12541016.99 | 14406737.04 | 12483492.33 | 13494234.86 | 13012665.23 | 13048600.74 | 13755954.13 | 13883966.09 | 13679597.2 | 13389890.19 |
| PC(20:2(11Z,14Z)/20:5(5Z,8Z,11Z,14Z,17Z)) | 110172261.6 | 135826647.6 | 79665532.39 | 93174181.53 | 147795144.2 | 112517878.2 | 152300673.4 | 142133295.8 | 164701324.2 | 186808423.2 | 129806446.6 | 94238949.47 |
| DGCC 9:0_11:0 | 494110 | 484871 | 491593 | 471573 | 689664 | 474350 | 513215 | 535674 | 519766 | 668133 | 666690 | 449226 |

| Metabolites | PBS_1 | PBS_2 | PBS_3 | PBS_4 | BCG_1 | BCG_2 | BCG_3 | BCG_4 | BCGΔ*pe18*_1 | BCGΔ*pe18*_2 | BCGΔ*pe18*_3 | BCGΔ*pe18*_4 |
|---|---|---|---|---|---|---|---|---|---|---|---|---|
| | 24.52 | 08.33 | 46.42 | 70.77 | 97.73 | 63.32 | 03.9 | 81.69 | 96.66 | 44.5 | 84.13 | 04 |
| 1-Phenyl-1,3,8-triazaspiro[4.5]decan-4-one | 127508.9541 | 1482866.346 | 3007368.774 | 2642447.148 | 172441.6572 | 111675.7261 | 715931.8037 | 22896.68272 | 437275.3159 | 770487.46 | 51742.67848 | 1522180.755 |
| PC O-34:0 | 2688595.517 | 9684896.195 | 4501517.479 | 19678976.92 | 19430067.9 | 2571459.309 | 3668686.644 | 10786081.96 | 18088127.96 | 3281878.018 | 16813756.09 | 3595512.401 |
| PC 8:0_20:1 | 297706.4599 | 187791.6184 | 225944.533 | 244411.5466 | 4725802.975 | 7285041.996 | 5125482.521 | 8456177.563 | 10864905.68 | 11413467.35 | 6684844.175 | 6758175.377 |
| Gallopamil | 6626314.462 | 7710434.203 | 5868195.61 | 7489985.062 | 7475083.821 | 4819530.859 | 8446682.452 | 7075249.262 | 5153756.938 | 7353038.788 | 6550029.661 | 13308562.45 |
| SM 14:3;2O/8:0 | 5823345.708 | 6001147.498 | 5897942.002 | 6122817.077 | 6381848.693 | 5596319.933 | 5984867.791 | 6080181.438 | 5706721.64 | 7316656.906 | 5459479.295 | 6354425.169 |
| 4-Oxo-2-nonenal | 618318.6648 | 958470.181 | 593017.9674 | 2411514.7 | 570481.6192 | 695357.5445 | 949027.4235 | 506630.7969 | 770694.1927 | 962015.8064 | 462116.644 | 1218701.226 |
| PE O-8:0_26:3 | 7915366.982 | 3611853.778 | 8070268.479 | 7024784.551 | 6909184.581 | 0 | 5221171.078 | 3103984.52 | 6298255.584 | 7196406.962 | 5032454.095 | 3032276.402 |
| SM 23:1;2O/11:0 | 149566518.9 | 68944899.92 | 126226928 | 73176480.4 | 38557445.63 | 386508.9604 | 130595105.8 | 86962465.03 | 75505913.18 | 110531860.7 | 30979349.83 | 50585269.02 |
| Emodin-8-Beta-D-Glucoside | 68980.74236 | 55487.32233 | 89654.87546 | 72789.85927 | 3471199.763 | 4463761.876 | 3243089.285 | 3637787.257 | 3673865.228 | 3520415.548 | 3973269.309 | 4386698.179 |
| Phosphate | 211387239.8 | 254366506.5 | 163874187.3 | 198569649.7 | 227848566.5 | 184338180.7 | 224335870 | 216107477.1 | 227036354.2 | 265981936.5 | 215719638 | 247076025.1 |
| PC O-37:5 | 10337732.64 | 9498362.994 | 5383755.887 | 8878774.515 | 2471977.513 | 2397941.616 | 2573204.547 | 2654335.046 | 10948884.96 | 3087452.066 | 2378957.478 | 5995255.473 |

| Metabolites | PBS_1 | PBS_2 | PBS_3 | PBS_4 | BCG_1 | BCG_2 | BCG_3 | BCG_4 | BCGΔ*pe18*_1 | BCGΔ*pe18*_2 | BCGΔ*pe18*_3 | BCGΔ*pe18*_4 |
|---|---|---|---|---|---|---|---|---|---|---|---|---|
| PC 8:0_19:1 | 1216153.046 | 1282697.69 | 1088356.517 | 1112259.315 | 1321796.116 | 1384164.75 | 1348372.466 | 1323208.332 | 3731015.011 | 1306954.512 | 1231578.303 | 6814094.176 |
| Met-Glu | 118763.1178 | 140497.5344 | 103843.5564 | 94190.84404 | 1168518.086 | 1215778.19 | 2952175.679 | 2637394.703 | 3156439.274 | 1172901.633 | 2076900.793 | 2886280.195 |
| PE O-18:5_4:0 | 16883175.28 | 21286608.19 | 22155988.5 | 19833809.14 | 31317427.51 | 31420615.62 | 25125479.87 | 31687334.08 | 32664760.88 | 30740373.08 | 28489276.04 | 33850161.9 |
| OMEGA-HYDROXYDODECANOIC ACID | 3998945.543 | 3271473.141 | 3778750.518 | 2462426.944 | 4706155.538 | 4763765.686 | 4448865.291 | 4213224.74 | 4116752.171 | 4189117.426 | 4907823.539 | 5816936.824 |
| 1,5-dimethyl-1-vinyl-4-hexenyl acetate | 4444623.709 | 4617910.941 | 4423421.004 | 4103672.96 | 4104797.589 | 4210584.05 | 4318101.013 | 4238848.231 | 4568209.294 | 4115358.115 | 4120561.458 | 4521129.641 |
| PG 22:5_22:5 | 218285604.3 | 234568297.6 | 78003783.82 | 158876447.4 | 20609431.69 | 17506544.41 | 31214300.93 | 29422525.35 | 28980848.91 | 40827068.32 | 22892028.56 | 23839732.24 |
| 1-(Chloroacetyl)-6-fluoro-2-methyl-1,2,3,4-tetrahydroquinoline | 1261073.141 | 1311850.408 | 1247474.856 | 132237.663 | 1354404.627 | 1290653.156 | 1420134.643 | 1312517.282 | 1371886.659 | 1427085.112 | 1689723.608 | 1187829.636 |
| DGTS 17:2_22:4 | 3813059.514 | 1818409.209 | 2726305.851 | 3282469.58 | 2435672.554 | 644359.7273 | 4271075.819 | 8376607.279 | 3622095.83 | 2984426.495 | 1834975.176 | 8180423.248 |
| 1,3,4,6,7,8-Hexahydro-1-methyl-2H-pyrimido[1,2-a]pyrimidine | 3281125.173 | 4739103.652 | 7747863.717 | 6697943.12 | 4203323.763 | 3300082.513 | 5082082.66 | 2982821.249 | 4970921.551 | 3917171.126 | 2927883.853 | 4631544.082 |

| Metabolites | PBS_1 | PBS_2 | PBS_3 | PBS_4 | BCG_1 | BCG_2 | BCG_3 | BCG_4 | BCGΔ*pe18*_1 | BCGΔ*pe18*_2 | BCGΔ*pe18*_3 | BCGΔ*pe18*_4 |
|---|---|---|---|---|---|---|---|---|---|---|---|---|
| (2S,3aS,7aS)-Octahydro-1H-indole-2-carboxylic acid | 1226060.38 | 743677.5093 | 1132232.268 | 668231.8162 | 647324.0029 | 875309.7089 | 776030.4149 | 475465.7585 | 1004728.205 | 1191037.72 | 1236570.236 | 1053707.569 |
| Isomaltose | 473170.8303 | 447306.8038 | 656803.8536 | 523157.5728 | 154225535.8 | 139888358.6 | 115797351.5 | 117316471.7 | 160291042.3 | 122417727.8 | 130177395.6 | 159424984.6 |
| Erythronolactone | 123505584.7 | 111364212.5 | 126106664.9 | 128763595.1 | 145863081.4 | 239016858.8 | 107462695.6 | 77784133.73 | 104471283.6 | 98448306.63 | 178074910.9 | 201842585.8 |
| LPG(17:1) | 220105.1362 | 102717.8739 | 118809.8965 | 254463.1797 | 1578031.432 | 441134.6905 | 796665.1049 | 964145.1274 | 879662.5219 | 1270894.893 | 1903264.393 | 581369.6168 |
| LPI 19:0 | 539619.6236 | 576089.6707 | 396163.9165 | 649532.3319 | 18294820.96 | 34104136.26 | 20447616.35 | 18169927.43 | 26826639.66 | 21963142.97 | 33029734.95 | 24589256.24 |
| 1-Trifluoromethoxyphenyl-3-(1-propionylpiperidin-4-yl)urea | 130883.2523 | 130955.4574 | 180968.5143 | 125310.8822 | 27464326.14 | 21104434.72 | 17874273.96 | 17577741.62 | 25211825.99 | 19197854.73 | 18270961.53 | 24565296.9 |
| DGDG 13:1_14:1 | 140737.6729 | 253945.4431 | 136036.0198 | 171327.2684 | 657820.8408 | 960589.1117 | 601185.109 | 555797.8234 | 1132271.823 | 925289.8704 | 823301.5797 | 801229.5382 |
| SM 21:2;2O/18:0 | 19280969.21 | 16152841.3 | 11849463.08 | 18853285.59 | 4382673.228 | 8730094.847 | 7095823.249 | 5610047.163 | 7220819.369 | 9969476.783 | 6563006.698 | 5256824.905 |
| PC(20:4(8Z,11Z,14Z,17Z)/15:0) | 154832517.4 | 189630910.1 | 286178246.4 | 216594311 | 327744120.7 | 264788171.3 | 401187175 | 300736526.4 | 331815078.5 | 461666391.7 | 388480929.4 | 178486642.8 |
| 1-Stearoyl-2-docosahexaenoyl-sn-glycero-3-phospho-(1'-sn | 80122786.72 | 91284471.71 | 34005535.73 | 57410033.84 | 107524322.9 | 71820938.56 | 116490233.9 | 98636561.63 | 128443044.2 | 136554414.3 | 96704793.27 | 65027281.13 |

| Metabolites | PBS_1 | PBS_2 | PBS_3 | PBS_4 | BCG_1 | BCG_2 | BCG_3 | BCG_4 | BCGΔ*pe18*_1 | BCGΔ*pe18*_2 | BCGΔ*pe18*_3 | BCGΔ*pe18*_4 |
|---|---|---|---|---|---|---|---|---|---|---|---|---|
| -glycerol) | | | | | | | | | | | | |
| 3-Methylpyridine | 1019217.269 | 281574.6172 | 2742193.217 | 1975667.284 | 3049251.015 | 3044156.671 | 557370.6874 | 1488915.803 | 1558007.826 | 2730255.463 | 2198259.02 | 2852442.67 |
| UDP-4-dehydro-6-deoxy-glucose | 0 | 0 | 18723.56538 | 22912.89682 | 1513883.776 | 1415916.651 | 1107277.666 | 1068016.377 | 1560698.578 | 1223292.888 | 1437703.956 | 1747909.738 |
| LPI 17:0 | 1040695.893 | 866863.7641 | 607042.9888 | 1509057.636 | 3655984.929 | 3869164.362 | 2819101.591 | 2977971.431 | 3794563.985 | 5989008.123 | 3064539.637 | 2231773.042 |
| 2-Bromo-5-fluoro-N-(2-furylmethyl)benzamide | 130074149.4 | 122731503 | 136303050.4 | 130154907.8 | 127092492 | 127456072 | 116866061.2 | 133455204.4 | 120203644.8 | 190543710 | 134812442.6 | 76802000.67 |
| 1-22:0-2-18:3-phosphatidylserine | 110139999.7 | 156661372.1 | 96112051.56 | 108688056.1 | 110022837 | 92643181.25 | 112459035.2 | 100285290.3 | 110638394.8 | 128871387.4 | 112126938.9 | 78955105.48 |
| Panaxydiol | 11583034.03 | 12123976.55 | 10250593.69 | 11071893.67 | 11561817.29 | 10238862.85 | 11761423.5 | 11338978.48 | 12757619.44 | 11640864.92 | 11126762.68 | 10786636.9 |
| PC(18:1(9Z)/18:1(9Z)) | 941344703 | 1234966723 | 500747256.7 | 714596905.1 | 213720200.9 | 180616572.3 | 282220283.3 | 244272138.2 | 294029579.7 | 335947924.2 | 241886390.7 | 161995009.7 |
| Cer 8:1;2O/3:0 | 51391433.85 | 56980076.08 | 54090057.23 | 56240327.86 | 53274266.74 | 52204461.52 | 60553758.86 | 51204238.31 | 54538370.25 | 54667960.29 | 51295095.27 | 60536004.82 |
| Ammonia pyruvate | 11201922.91 | 14375281.82 | 13392279.08 | 15267637.33 | 10325385.12 | 27965347.01 | 13288223.88 | 8274224.681 | 13564759.42 | 12546476.78 | 21506058.6 | 13382659.23 |
| Lumichrome | 10022811.09 | 9077624.488 | 13726977.82 | 18654559.77 | 19469517.27 | 18904889.46 | 20375263.85 | 18091404.03 | 19988251.95 | 18606015.57 | 18455644.53 | 34209530.52 |

| **Metabolites** | **PBS_1** | **PBS_2** | **PBS_3** | **PBS_4** | **BCG_1** | **BCG_2** | **BCG_3** | **BCG_4** | **BCGΔ*pe18*_1** | **BCGΔ*pe18*_2** | **BCGΔ*pe18*_3** | **BCGΔ*pe18*_4** |
|---|---|---|---|---|---|---|---|---|---|---|---|---|
| N-(4-Chlorophenyl)-N'-(5-methyl-2-pyridinyl)urea | 7432.820253 | 69697.49556 | 21002.70371 | 11505.94349 | 674208.9177 | 990433.0338 | 1957020.932 | 1851839.136 | 2153761.291 | 699671.0259 | 1423263.712 | 2102684.214 |
| 2-Methylpropanamine | 92627934.23 | 82844323.83 | 85213448.83 | 84445620.75 | 82909308.75 | 86407474.98 | 57767794.31 | 90211025.04 | 83055396.4 | 85142243.5 | 83724401.67 | 81237647.02 |
| 4,4'-Sulfonylbisphenol | 23176563.28 | 23275319.03 | 21954754.1 | 22990204.25 | 3281920.972 | 23373790.91 | 22887289.94 | 21484632.45 | 2974392.42 | 23017906.24 | 23451751.77 | 22847422.85 |
| LPE 18:3 | 785536.4651 | 568171.5255 | 468201.9284 | 897066.2374 | 448119.2835 | 233453.6203 | 536122.3619 | 461837.0547 | 917044.1317 | 772408.9754 | 496948.1964 | 505921.3019 |
| 1-Hexadecanoyl-2-(9Z-octadecenoyl)-sn-glycero-3-phospho-1'-myo-inositol | 978512.6272 | 883145.9769 | 1093638.878 | 1024376.869 | 1193630.216 | 112262.1814 | 869559.1113 | 883639.73 | 809143.8422 | 726779.6675 | 1124446.837 | 1117534.005 |
| SM 38:4;3O | 3943692.816 | 5998041.851 | 3379093.874 | 12443391.99 | 5250297.487 | 564214.1986 | 9610190.135 | 49049815.21 | 18676202.65 | 5881192.365 | 3028145.732 | 43210873.15 |
| n-Octadecylamine | 49621728.09 | 67771259.05 | 35922778.47 | 42279377.8 | 13234075.46 | 11271447.37 | 16850902.66 | 17031620.77 | 14699731.1 | 19080581.25 | 15186765.76 | 10142379.93 |
| DGCC 16:4_20:5 | 313743151.1 | 338611207.6 | 275819038.8 | 297008067.6 | 310532189.8 | 286338801.2 | 314424727.7 | 277588421.3 | 297849411.8 | 314422365.3 | 315526374.1 | 279724742.5 |
| Leu-Leu | 654679.901 | 570747.0039 | 752705.4549 | 537415.9655 | 584405.7111 | 392442.771 | 544427.277 | 401921.2885 | 642002.5541 | 731994.4778 | 859688.6794 | 451306.4991 |
| Azimsulfuron | 0 | 0 | 4243.228444 | 0 | 2459671.828 | 2206227.283 | 1940424.74 | 1898747.937 | 2374525.151 | 2142443.595 | 2066010.035 | 2353455.045 |
| Curcumenol | 130739 | 123276 | 376914 | 172973 | 151872 | 148090 | 106344 | 126218 | 142268 | 118908 | 148200 | 129226 |

| Metabolites | PBS_1 | PBS_2 | PBS_3 | PBS_4 | BCG_1 | BCG_2 | BCG_3 | BCG_4 | BCGΔ*pe18*_1 | BCGΔ*pe18*_2 | BCGΔ*pe18*_3 | BCGΔ*pe18*_4 |
|---|---|---|---|---|---|---|---|---|---|---|---|---|
| | 4.843 | 9.965 | 0.056 | 5.591 | 836.6 | 037.2 | 452.2 | 880.3 | 239.7 | 736.7 | 196.9 | 353.7 |
| Cys-Cys | 126978558.4 | 126451325.4 | 129100163.6 | 127208757.8 | 123332635.2 | 109002907.2 | 67219450.21 | 33975943.55 | 132414704 | 31117822.67 | 81515734.09 | 115499649.7 |
| PE O-13:1_5:0 | 11884471.82 | 13072411.5 | 15617782.58 | 11104434.93 | 41278700.94 | 41667399.33 | 40765683.31 | 35937723.76 | 37264398.58 | 42178574.83 | 59028769.29 | 35124305.9 |
| Galactinol | 473170.8303 | 447306.8038 | 656803.8536 | 523157.5728 | 154225535.8 | 139888358.6 | 115797351.5 | 117316471.7 | 160291042.3 | 122417727.8 | 130177395.6 | 159424984.6 |
| Genistein | 4055347.606 | 4029023.857 | 4529539.836 | 4764293.551 | 2136364.327 | 2519116.061 | 1898186.169 | 2399000.991 | 2434398.355 | 1784012.321 | 2107931.062 | 2925102.574 |
| M261T452_2 | 121617052.7 | 126221336.7 | 92450701.87 | 112213310.6 | 140990697.3 | 164149153.8 | 90276874.18 | 146370077.7 | 139329017.3 | 152931184.1 | 141080318.2 | 117385708.2 |
| 2-Arachidonoyl-1-stearoyl-sn-glycero-3-phosphoethanolamine | 54244235.33 | 55336773.91 | 72952579.39 | 61266078.77 | 86535450.78 | 75826621.89 | 128681399.4 | 95390093.76 | 88085544.57 | 146382196.7 | 109143346.9 | 55154681.84 |
| Dodeca-2(E),4(E)-dienoic acid | 52106532.26 | 41172648.83 | 44991265.92 | 35284742.49 | 50668153.92 | 40881490.14 | 47298850.8 | 35365251.29 | 46496580.64 | 43342958.39 | 45821405.71 | 43008622.47 |
| FA 15:1 | 7644929.162 | 8573943.328 | 5681783.975 | 6286871.967 | 5961397.16 | 7836075.442 | 7079084.989 | 7108025.719 | 6571032.394 | 8410443.63 | 8889317.23 | 7064655.293 |
| 1-Palmitoyl-2-docosahexaenoyl-sn-glycero-3-phospho-(1'-rac-glycerol) | 48545564.27 | 57515365.92 | 26840104.87 | 39985586.35 | 41806395.57 | 36644922.45 | 60213187.38 | 53571029.39 | 76345361.66 | 80209529.36 | 53346843 | 38643950.64 |

| Metabolites | PBS_1 | PBS_2 | PBS_3 | PBS_4 | BCG_1 | BCG_2 | BCG_3 | BCG_4 | BCGΔ*pe18*_1 | BCGΔ*pe18*_2 | BCGΔ*pe18*_3 | BCGΔ*pe18*_4 |
|---|---|---|---|---|---|---|---|---|---|---|---|---|
| 2,4-Dichloro-6-fluorobenzoic acid | 13579605.81 | 12011272.52 | 3934742.634 | 16651301.92 | 15134591.72 | 5653954.742 | 7291632.314 | 2301296.327 | 11313426.9 | 2946426.648 | 13680556.33 | 7223230.944 |
| Sulfadimethoxine | 1663349.334 | 1231965.678 | 2732746.024 | 1780325.254 | 8936176.323 | 5557302.075 | 7961316.043 | 6616103.525 | 8528545.429 | 7593392.138 | 9287401.532 | 7330143.25 |
| gamma-Glutamylmethionine | 56544.27256 | 49149.14592 | 56260.10222 | 52026.3233 | 8478970.772 | 13106927.53 | 24899347.52 | 24805651.03 | 30051150.25 | 11250201.62 | 19654506.53 | 31556106.95 |
| PC O-16:1 | 17961283.55 | 15914339.68 | 25114072.49 | 15986555.26 | 71394341.38 | 55213410.2 | 58954558.2 | 51773605.03 | 62018081.96 | 59286880.9 | 65584107.17 | 53163080.49 |
| 4-[3-(4-Butoxyphenoxy)propyl]morpholine | 75666868.74 | 80717156.46 | 71387670.88 | 78514910.78 | 76794505.35 | 76561467.96 | 77195364.73 | 75256484.91 | 81477330.96 | 79440161.73 | 76712189.95 | 73847672.42 |
| PC O-32:2 | 3670539.287 | 3403856.96 | 4885536.855 | 3281621.627 | 3757229.722 | 2668337.406 | 3834468.02 | 3097131.365 | 4229047.805 | 8046873.06 | 3239690.499 | 2991182.013 |
| 3-(Hepta-1,3-dienyl)hexanedioic acid | 68279451.28 | 51271313.19 | 53309874.3 | 43332066.9 | 65105295.36 | 51502207.34 | 60922259.92 | 45959747.07 | 59254720.14 | 54783986.35 | 56677379.11 | 54272623.54 |
| Dinor-12-oxophytodienoic acid | 6985520.132 | 7248051.551 | 6742746.597 | 7055867.566 | 6903997.841 | 6891614.6 | 8205662.494 | 6773014.822 | 7343725.161 | 6798314.444 | 6944832.96 | 8005671.015 |
| 2-[(4-Morpholinylcarbothioyl)sulfanyl]propanoic acid | 1276984.232 | 1266755.833 | 1258183.99 | 1135919.915 | 1184118.747 | 1207957.347 | 1155938.384 | 1129024.57 | 1213283.733 | 1102685.663 | 1236540.988 | 1215674.039 |
| 3-(1-Pyrrolidinylcarbonyl)piperidine | 46859740.96 | 54175811.65 | 32750264.96 | 47325684.03 | 50112032.83 | 39932040.56 | 49808440.35 | 50326341.41 | 54262867.65 | 48209435.78 | 47130152.99 | 43101566.34 |
| 1,9-Dideoxyforskol | 730788 | 732387 | 736237 | 717772 | 702633 | 739926 | 704237 | 708513 | 744998 | 728703 | 753210 | 665945 |

| Metabolites | PBS_1 | PBS_2 | PBS_3 | PBS_4 | BCG_1 | BCG_2 | BCG_3 | BCG_4 | BCGΔ*pe18*_1 | BCGΔ*pe18*_2 | BCGΔ*pe18*_3 | BCGΔ*pe18*_4 |
|---|---|---|---|---|---|---|---|---|---|---|---|---|
| in | 07.26 | 33.28 | 88.05 | 05.45 | 32.65 | 25.42 | 58.46 | 34.35 | 49.22 | 04.98 | 12.12 | 38.1 |
| 3-(2-Fluorophenoxy)-7-hydroxy-4H-chromen-4-one | 25738.12085 | 49858.56554 | 55231.21211 | 31716.0478 | 644879.0914 | 357276.749 | 706767.3982 | 476474.1647 | 509889.9383 | 693172.4153 | 788611.3497 | 461365.9939 |
| Prolinamide | 695743.8234 | 3436775.865 | 1973988.753 | 1939018.616 | 2066292.088 | 3047611.318 | 1301732.894 | 664609.1518 | 4239712.134 | 1381043.023 | 1002510.227 | 1423541.978 |
| LDGCC 12:0 | 2082131.991 | 2186650.488 | 1920542.053 | 2290641.753 | 2242225.696 | 2148393.431 | 2251276.166 | 2400791.649 | 2340633.575 | 2729773.501 | 2056919.119 | 1997801.137 |
| PC(22:2(13Z,16Z)/P-18:1(9Z)) | 22972152.53 | 30509914.26 | 23790894.27 | 23784431.79 | 7806746.021 | 7919241.78 | 12809791.39 | 9012970.624 | 12151608.61 | 16030817.59 | 10397790.27 | 4982616.55 |
| 7-(2-Aminophenyl) heptanoic acid | 294296.5729 | 1017034.498 | 1161328.632 | 1267026.416 | 362370.1031 | 310334.4169 | 429990.7743 | 446820.7514 | 462610.4566 | 447231.1931 | 369952.5521 | 24270206.36 |
| MGDG O-10:0_18:1 | 12798308.25 | 13259605.39 | 10775029.98 | 11837600.77 | 12481922.6 | 12189238.84 | 11641984.25 | 11965033.35 | 13381001.95 | 11992160.18 | 11600719.73 | 11471018.79 |
| PS 15:1_22:6 | 494519.9662 | 430365.9879 | 282259.7235 | 485078.6959 | 341445.9795 | 637157.197 | 507669.7954 | 549098.8227 | 812152.0977 | 622637.8049 | 461230.6373 | 482590.3147 |
| LDGCC 17:1 | 808345004.7 | 935365607.5 | 766482469.5 | 810269862 | 825860129.6 | 822693572.4 | 815379485.6 | 824480279.5 | 827512214.5 | 825242646.4 | 795089312.5 | 850810245.8 |
| PC(38:5) | 15303265.37 | 22286524.65 | 5745819.005 | 10688160.47 | 5331737.689 | 5259339.216 | 5086751.654 | 5801125.631 | 7230335.165 | 7491528.915 | 3511025.468 | 3459599.545 |
| dTDP-4-acetamido-4,6-dideoxy-D-galactose | 718433.1961 | 570561.0671 | 860763.9703 | 589000.9995 | 2727495.967 | 1859008.513 | 3794174.525 | 2998087.84 | 2507819.056 | 2485292.791 | 2992489.454 | 3712113.526 |
| PC O-22:4 | 417321 | 161836 | 998235 | 154385 | 217663 | 126290 | 152867 | 139667 | 245786 | 310470 | 348503 | 129544 |

| Metabolites | PBS_1 | PBS_2 | PBS_3 | PBS_4 | BCG_1 | BCG_2 | BCG_3 | BCG_4 | BCGΔ *pe18*_1 | BCGΔ *pe18*_2 | BCGΔ *pe18*_3 | BCGΔ*pe18*_4 |
|---|---|---|---|---|---|---|---|---|---|---|---|---|
| | 4.126 | 0.829 | 9.921 | 7.164 | 0.901 | 7.766 | 6.239 | 2.518 | 5.879 | 5.838 | 9.256 | 5.259 |
| 1-(7Z-Hexadecenoyl)-2-(4Z,7Z,10Z,13Z,16Z,19Z-docosahexaenoyl)-sn-glycero-3-phosphocholine | 1635223.729 | 1588807.02 | 1335847.663 | 1600976.775 | 1872110.274 | 1547979.877 | 2391923.056 | 1725974.399 | 2102464.108 | 2117592.818 | 2480494.186 | 1339927.216 |
| 1H-Indazole-3-carboxamide, N-[1-(aminocarbonyl)-2,2-dimethylpropyl]-1-pentyl- | 8975308.635 | 9711494.611 | 10435815.03 | 10589316.12 | 8587388.352 | 9201625.616 | 9300634.39 | 9056189.822 | 9220753.62 | 8813139.898 | 9282280.697 | 12048958.35 |
| PC O-16:0_20:4 | 2646526.442 | 8181611.682 | 2317334.638 | 6433525.436 | 2988253.293 | 31022.7295 | 12600448.19 | 14762879.63 | 13603700.92 | 8050011.747 | 2344577.295 | 28883142.83 |
| N-Dodecanoyl-N-methylglycine | 70382952.09 | 77517764.69 | 71473258.79 | 71768779.93 | 71111413.31 | 73032114.52 | 75812091.08 | 72580644.86 | 76629384.43 | 75374282.58 | 72227952.65 | 71315740.85 |
| N-Acetylputrescine | 663564.1549 | 775395.0211 | 1335167.312 | 864200.7445 | 10713991.61 | 7559264.434 | 12346229.02 | 9387035.76 | 9998944.433 | 13706353.78 | 13401317.37 | 9862621.321 |
| Isocaproic acid | 11200798.14 | 11753945.74 | 10540995.71 | 22623580.32 | 12857259.35 | 12596840.34 | 30523389.54 | 10663412.18 | 18247375.38 | 13504266.64 | 11089852.81 | 30914008.55 |
| 2-(2-Hydroxypropan-2-yl)-2,3-dihydro-7H-furo[3,2-g]chromen-7-one | 7932183.04 | 6890851.373 | 8203457.22 | 6327343.22 | 33325069.74 | 24116777.56 | 32285873.88 | 25068274.08 | 23251994.65 | 25243929.07 | 40925025.7 | 25637541.2 |

| Metabolites | PBS_1 | PBS_2 | PBS_3 | PBS_4 | BCG_1 | BCG_2 | BCG_3 | BCG_4 | BCGΔ *pe18*_1 | BCGΔ *pe18*_2 | BCGΔ *pe18*_3 | BCGΔ*pe18*_4 |
|---|---|---|---|---|---|---|---|---|---|---|---|---|
| 2-Arachidonoyl-1-palmitoyl-sn-glycero-3-phosphoethanolamine | 29283738.73 | 35938298.01 | 34368323.99 | 30248360.35 | 81263530.05 | 61757605.11 | 92763821.99 | 73324984.37 | 78246910.68 | 99506849.96 | 87167355.05 | 45889079 |
| LPI 18:1 | 41733053.83 | 39112577.14 | 35237031.16 | 32742130.17 | 89178295.49 | 80030138.82 | 78135176.3 | 70599959.61 | 75030749.96 | 83301974.9 | 97449529.78 | 64480190.5 |
| Trimethylaminoacetone | 15164599.07 | 30734460.32 | 31609039.59 | 15155579.17 | 17531564.27 | 26259340.72 | 27594088.05 | 24704029.04 | 21067586.12 | 21153042.16 | 33003003.52 | 27982951.52 |
| PG(16:1(9Z)/18:0) | 27404963.76 | 27534925.88 | 22613316.68 | 29490337.94 | 38331836.7 | 32773734.35 | 44280719.06 | 25375127.04 | 34192731.53 | 46014587.28 | 37214748.31 | 31442326.19 |
| 2-(Hydroxymethyl)-2-(octylamino)-1,3-propanediol | 258352942.3 | 270360167.8 | 249827687.2 | 240999147.2 | 289219057 | 269177653.7 | 256791148.8 | 254647114.3 | 317845477.3 | 268862090.7 | 243285677.4 | 249810604.9 |
| PC(14:1(9Z)/16:1(9Z)) | 1310460.572 | 1349572.233 | 1600447.474 | 1129609.593 | 16695854.38 | 12286024.22 | 18461167.79 | 14852911.83 | 18641880.42 | 19497977.51 | 14823264.64 | 9284987.905 |
| DGGA 12:0_14:0 | 745788774.2 | 778904499.9 | 637363012.4 | 685008141.6 | 759429913.4 | 715664474.1 | 697698085.1 | 668978247.2 | 762854978.6 | 742504044.1 | 706169294 | 658210172 |
| FA 22:4 | 32659818.06 | 32810013.08 | 62570903.57 | 34391723.55 | 10874735.34 | 12797235.22 | 16175826.49 | 12753642.39 | 11766833.99 | 15224536.99 | 17131891.07 | 12845583.17 |
| PE 42:0_26:5 | 1842657.483 | 1747573.937 | 1662738.65 | 1766368.506 | 1628685.935 | 1732408.209 | 1652652.457 | 1767355.428 | 1789550.575 | 1718406.711 | 1734648.512 | 1826135.447 |
| LNAPE 20:4/N-24:3 | 1447732.772 | 686583.656 | 751943.648 | 1674578.349 | 754060.7155 | 26137.99931 | 514637.0211 | 337846.2951 | 1593529.571 | 1959175.274 | 897892.8994 | 408820.7839 |
| N-Methylcyclohexa | 129278 | 147374 | 160902 | 156190 | 142967 | 129806 | 137108 | 140717 | 126991 | 135807 | 134087 | 169658 |

| Metabolites | PBS_1 | PBS_2 | PBS_3 | PBS_4 | BCG_1 | BCG_2 | BCG_3 | BCG_4 | BCGΔ*pe18*_1 | BCGΔ*pe18*_2 | BCGΔ*pe18*_3 | BCGΔ*pe18*_4 |
|---|---|---|---|---|---|---|---|---|---|---|---|---|
| namine | 13.47 | 70.35 | 70.35 | 19.58 | 37.21 | 41.69 | 88.49 | 60.1 | 52 | 79.94 | 00.72 | 62.13 |
| PC(18:4(6Z,9Z,12Z,15Z)/20:4(5Z,8Z,11Z,14Z)) | 11240134.13 | 13947794.58 | 6273440.302 | 9740371.702 | 17424662.01 | 14789712.48 | 18083367.42 | 18220381.02 | 21224902.26 | 21176461.21 | 15962841.94 | 14952223.81 |
| PG(16:0/20:3(5Z,8Z,11Z)) | 23304786.29 | 25873779.89 | 19877318.63 | 20985126.2 | 24914397.51 | 19736393.79 | 24910144.95 | 13941641.42 | 25052657.08 | 26184653.9 | 22628334.42 | 15822684.03 |
| Allopurinol-1-ribonucleoside | 2840127.608 | 2096157.34 | 4909354.043 | 3028713.914 | 36873297.93 | 22205570.17 | 39857064.5 | 26789214.7 | 29461613.59 | 36040009.97 | 42510478.42 | 24865145.63 |
| Frangulin B | 40401364.03 | 38711412.93 | 36872160.82 | 38733844.39 | 39233946.2 | 39116796.46 | 38826889.54 | 37637341.1 | 38000101.99 | 38197405.32 | 38918799.79 | 39340343.85 |
| DGTS 18:5_22:6 | 862194.4452 | 85524.84212 | 813086.0952 | 770345.622 | 39980298.51 | 35338398.58 | 41499974.3 | 32707693.9 | 37734878.55 | 36917781.89 | 43008175.59 | 32556981.56 |
| LPE 36:5 | 4238164.882 | 3019270.119 | 4537285.596 | 1990994.235 | 3687066.86 | 0 | 3757649.739 | 5100080.579 | 2514190.037 | 4635481.733 | 4625159.471 | 3391527.833 |
| 3,3-pentamethylene-4-butyrolactam | 1287029.425 | 1607570.53 | 1004820.037 | 1121062.608 | 1135267.1 | 937582.1668 | 1067806.101 | 910525.0242 | 1150679.775 | 897698.763 | 1387216.31 | 1439460.121 |
| PE(34:2) | 4853203.098 | 4964565.536 | 5644898.454 | 4149828.804 | 36075739.43 | 28313434.77 | 38332110.08 | 31580697.36 | 39401562.17 | 40136526.68 | 33644327.59 | 20207621.66 |
| SM 19:2;2O/15:0 | 14018808.49 | 27714835.84 | 15040366.43 | 12643419.37 | 22318549.32 | 43105566.13 | 22061584.45 | 25885315.9 | 26659070.4 | 38725300.98 | 24212496.78 | 24095516.4 |
| SL 23:3;O/26:7 | 10137578.49 | 18789271.82 | 8229037.313 | 12929403 | 13368535.86 | 8913959.978 | 11404932.14 | 9307159.514 | 11120984.07 | 13956787.05 | 10771947.99 | 7298730.784 |
| PG(16:0/18:1) | 81555994.32 | 89229373.62 | 58026558.87 | 59820300.19 | 137595920.4 | 92800933.17 | 146066910.7 | 118510570.9 | 137194149.4 | 167139909.9 | 129379221.1 | 77276162.94 |

| Metabolites | PBS_1 | PBS_2 | PBS_3 | PBS_4 | BCG_1 | BCG_2 | BCG_3 | BCG_4 | BCGΔ*pe18*_1 | BCGΔ*pe18*_2 | BCGΔ*pe18*_3 | BCGΔ*pe18*_4 |
|---|---|---|---|---|---|---|---|---|---|---|---|---|
| PS 17:0 | 1042431.26 | 1222703.997 | 699129.938 | 1043179.352 | 880147.1721 | 888420.7668 | 635356.8723 | 872166.192 | 915865.6839 | 952424.5235 | 781285.6631 | 815494.9172 |
| PS 18:0_22:4 | 6094834.809 | 3310671.34 | 5250040.585 | 7114014.864 | 5030626.163 | 2288524.024 | 5492627.499 | 5188997.518 | 7968181.261 | 6697625.835 | 4045817.392 | 7294359.928 |
| N-Methyltrimethylacetamide | 3713344.989 | 13084972.93 | 19385773.35 | 15378836.92 | 25957242.94 | 3972374.274 | 12767839.74 | 4056587.526 | 6660006.075 | 6577149.018 | 8204700.526 | 30305293.33 |
| DGGA 12:0_26:2 | 12951056.67 | 15524803.78 | 10407380.39 | 11331038.08 | 2134571.774 | 1924272.773 | 2461342.578 | 1608083.472 | 2518618.844 | 3224372.18 | 2406173.818 | 1550128.89 |
| sphingofungin B1 | 3569191.798 | 2945283.026 | 2878229.764 | 3106034.845 | 3626948.77 | 2668668.685 | 3344218.497 | 2772799.651 | 3534442.586 | 3200021.862 | 3028330.943 | 3128288.431 |
| Cer 8:0;2O/16:3 | 3976516.74 | 4396704.708 | 3216080.499 | 3465918.632 | 3381188.329 | 3798093.744 | 3864396.201 | 3928831.476 | 3604228.801 | 4416590.996 | 3861809.217 | 3382679.12 |
| PE O-16:3_2:0 | 1559646.019 | 1947015.046 | 1536730.839 | 1606216.219 | 4452065.674 | 4707885.427 | 4599660.165 | 4688781.497 | 4834490.418 | 5158146.759 | 6057646.933 | 4177395.967 |
| 1-Heptanamine | 1029363292 | 1044771926 | 1047655203 | 986637666.7 | 977639526.5 | 990010249.7 | 1015498027 | 472176989.5 | 436229447 | 958908430.1 | 1008186136 | 1018878178 |
| PC O-38:4 | 23221806.78 | 14568655.2 | 22418694.59 | 20434984.81 | 18118723.27 | 2006946.986 | 32879268.37 | 61151128.3 | 30359152.87 | 21468544.53 | 36523620.09 | 60212338.07 |
| PMeOH 20:4_26:4 | 40444.7261 | 26772.86873 | 73581.64814 | 41641.96501 | 274790.8727 | 272003.982 | 262160.8918 | 330376.1683 | 413241.8109 | 270798.8264 | 355243.8143 | 259581.7327 |
| MGDG O-8:0_26:6 | 1235415.199 | 846490.7315 | 1676251.887 | 1825530.473 | 1258250.586 | 2569232.605 | 3568715.588 | 1597970.807 | 2827815.223 | 3179083.85 | 2613204.831 | 2242398.944 |

| Metabolites | PBS_1 | PBS_2 | PBS_3 | PBS_4 | BCG_1 | BCG_2 | BCG_3 | BCG_4 | BCGΔ*pe18*_1 | BCGΔ*pe18*_2 | BCGΔ*pe18*_3 | BCGΔ*pe18*_4 |
|---|---|---|---|---|---|---|---|---|---|---|---|---|
| 2-(Dipentylamino)-2-(hydroxymethyl)-1,3-propanediol | 2498804.902 | 1506840.392 | 2241226.316 | 1466166.033 | 1509044.296 | 1495378.917 | 1463090.439 | 1493133.123 | 2140237.254 | 1447388.149 | 1458236.284 | 1445129.965 |
| PI 18:1_22:6 | 1685924.425 | 2389214.719 | 956801.6643 | 1755984.312 | 2017086.348 | 2011073.677 | 1683018.707 | 1550996.013 | 2357895.029 | 2134466.923 | 1877505.817 | 1603554.511 |
| PC(20:2(11Z,14Z)/18:3(9Z,12Z,15Z)) | 287727388.4 | 298223876.7 | 103296485.1 | 204735435 | 329398823.4 | 183912839.7 | 378918085.3 | 342410195.2 | 341068292.9 | 482197095.4 | 298305122.8 | 234520256.9 |
| Cer 10:1;2O/8:0 | 4515728.656 | 4845603.591 | 4142726.182 | 4201855.383 | 4797402.628 | 4025507.101 | 4403453.102 | 4801291.252 | 4493748.789 | 7986949.301 | 4429815.882 | 4252151.06 |
| 2-(3-Isobutoxyphenoxy)-N-(4-methoxyphenyl)acetamide | 2906463.016 | 3109910.824 | 2681894.195 | 2850308.447 | 3037613.964 | 2889622.176 | 3059489.564 | 3017700.361 | 3112624.83 | 3051731.102 | 2983086.663 | 2826691.629 |
| Cholesteryl sulfate | 70939608.02 | 78698221.51 | 47226593.77 | 55063950.38 | 94754238.97 | 74403494.85 | 106163594.1 | 100662749 | 106390035.9 | 120957860 | 94141773.05 | 64471590.8 |
| 3-Isobutylpentanedioic acid | 2042386.022 | 2437546.181 | 2914767.038 | 2467452.053 | 2290615.233 | 3736205.382 | 2293037.064 | 2051435.714 | 2234742.727 | 2249258.184 | 2301418.618 | 3593078.453 |
| 5-(2-Chlorophenyl)-4-pyrimidinamine | 16612486.95 | 16885755.7 | 17107323.29 | 15923982.52 | 15087779.99 | 15325880.21 | 14852657.98 | 15924403.94 | 15421149.96 | 14419095.39 | 15249889.8 | 23895379.78 |
| PE O-16:2_2:0 | 5010570.033 | 4746300.839 | 6325223.671 | 4125261.767 | 12250951.62 | 9865891.528 | 11122569.12 | 11508168.42 | 12390906.28 | 10992531.56 | 12650885.25 | 11543627.68 |
| PC(14:0/18:3(9Z,12Z,15Z)) | 22897345.45 | 14381013.39 | 7020509.391 | 18474650.04 | 62046989.23 | 30247369.92 | 29507924.25 | 22114075.52 | 68175230.94 | 74392503.69 | 19849095.74 | 21361938.18 |
| PC(33:2) | 19712349.75 | 22689775.97 | 38607549.85 | 26034114.66 | 164414089.8 | 81322663.28 | 110712584.8 | 90645391.7 | 157062400.5 | 122778578.9 | 115604327.9 | 58556160.58 |

| **Metabolites** | **PBS_1** | **PBS_2** | **PBS_3** | **PBS_4** | **BCG_1** | **BCG_2** | **BCG_3** | **BCG_4** | **BCGΔ*pe18*_1** | **BCGΔ*pe18*_2** | **BCGΔ*pe18*_3** | **BCGΔ*pe18*_4** |
|---|---|---|---|---|---|---|---|---|---|---|---|---|
| 5-Norbornene-2,3-dicarboximide, N-(2-ethylhexyl)- | 46091951.71 | 47519323.31 | 41394174.71 | 44166410.42 | 44686160.86 | 44220177.01 | 45151106.71 | 43728840.43 | 47728657.9 | 43160149.52 | 42865621.58 | 42110010.94 |
| 2-(4,8,12-Trimethylcyclotetradeca-3,7,11-trien-1-yl)propan-2-ol | 166862139.9 | 172651520.5 | 158784239.5 | 158823440.5 | 160181288.5 | 157660503.5 | 166825400.1 | 165797085.1 | 163006919.5 | 166142895.8 | 162268459.6 | 155710636.4 |
| 2-Aminopyridyl-5-thioamide | 8755805.273 | 8549215.305 | 7972899.189 | 8108257.891 | 8071031.779 | 8271210.06 | 8298733.786 | 8453392.93 | 8451487.852 | 7847423.263 | 8264993.827 | 8035259.319 |
| PC(42:6) | 13367513.61 | 18724392.27 | 8145502.775 | 11151563.88 | 3982483.455 | 3306816.104 | 4275696.93 | 4037197.505 | 5277074.506 | 5190620.093 | 3665195.301 | 2945666.065 |
| 3-Ethyl-2,8-dimethyl-4-quinolinol | 19523452.23 | 19894852.38 | 17178533.67 | 18258238.55 | 16006859.87 | 15635877.07 | 16764518.76 | 14669391.51 | 15931885.08 | 15567227.74 | 15831882.77 | 17193195.92 |
| 2-Chloro-L-phenylalanine | 2583608.455 | 3054160.329 | 4155181.669 | 2710492.111 | 3720135.206 | 3450461.439 | 3668784.735 | 3623985.389 | 4451446.038 | 3477429.709 | 3203541.503 | 3865548.106 |
| LNAPS 3:0/N-4:0 | 7366271.437 | 7397081.651 | 7381407.297 | 7415698.055 | 7405060.933 | 7354090.36 | 7333917.446 | 7474696.136 | 7244361.049 | 7344438.637 | 7414603.331 | 7465815.631 |
| Prop-2-yn-1-ylglycine | 2848627.917 | 2654610.901 | 2281683.536 | 2644773.688 | 3121986.915 | 2870423.048 | 2496567.362 | 3011322.188 | 3187727.748 | 3246410.552 | 2241860.829 | 2757537.928 |
| (2-aminoethoxy)[2-[icosa-5.8.11-trienoyloxy]-3-(octadecanoyloxy)propoxy]phosphinic acid | 24224477.2 | 30054559.69 | 42969212.24 | 33935621.9 | 58031986.77 | 43014504.02 | 64409973.73 | 45927977.17 | 49538142.86 | 67503067.57 | 63766860.44 | 27286601.63 |

| Metabolites | PBS_1 | PBS_2 | PBS_3 | PBS_4 | BCG_1 | BCG_2 | BCG_3 | BCG_4 | BCGΔ*pe18*_1 | BCGΔ*pe18*_2 | BCGΔ*pe18*_3 | BCGΔ*pe18*_4 |
|---|---|---|---|---|---|---|---|---|---|---|---|---|
| 15HpETrE | 54088739.55 | 30598165.78 | 63284874.81 | 79497792.32 | 47479306.78 | 54305989.37 | 50328225.97 | 58678242.86 | 65983548.54 | 25641723.23 | 52497493.18 | 66615995.76 |
| PA 2:0_21:2 | 1781609.55 | 1779421.05 | 1747467.354 | 1797780.154 | 1832594.159 | 1790082.554 | 1757498.572 | 1703959.351 | 1765213.48 | 1793338.962 | 1875346.232 | 1763344.873 |
| 1,2,4,5-Benzenetetracarboxamide | 2125296.747 | 1663205.982 | 3857372.829 | 2361733.67 | 27126057.99 | 16807854.13 | 30128993.2 | 20407047.39 | 21852272.68 | 26715966.77 | 30083249.93 | 17696962.84 |
| PC(35:4) | 44963722.28 | 55376188.3 | 92917554.32 | 70356867.32 | 135033421.8 | 105929389.3 | 206750928.9 | 142949764.2 | 104806656.4 | 217348960.8 | 167707715.4 | 87663178.24 |
| Ethyl 4-hydroxy-2-methyl-6-quinolinecarboxylate | 746013.3098 | 737515.1347 | 349859.6524 | 481783.126 | 3121949.863 | 2876576.385 | 3449736.888 | 3483398.021 | 3699141.279 | 3387658.089 | 3021827.785 | 3377460.38 |
| SM 8:1;2O/32:9 | 7139099.465 | 7850771.054 | 6137553.757 | 6950112.291 | 7149365.033 | 6674584.544 | 7245966.869 | 6212597.484 | 6960856.899 | 7407931.884 | 7513558.625 | 6540556.922 |
| N-Benzyl-N,N-dimethyl-1-hexadecanaminium cation | 9136529.165 | 12014593.09 | 7840608.496 | 8237229.331 | 7370351.345 | 11198077.22 | 8629996.689 | 11733626.41 | 12047966.39 | 7877041.498 | 11559249.61 | 7788591.153 |
| Pentalenic acid | 3779065.399 | 2774668.244 | 2969810.781 | 3300740.637 | 5658683.247 | 3651275.41 | 32060.06913 | 4080848.962 | 3956456.459 | 5060171.249 | 3917708.301 | 3508602.252 |
| LPC 40:11-SN2 | 153011342.7 | 179734663.8 | 155262989.2 | 155381128.9 | 169528866.7 | 156301819.6 | 175878666.9 | 158730245.3 | 163035407.5 | 168833271.9 | 161573052.5 | 152340883.9 |
| 4-Chloro-N'-(2-chloroacetyl)benzohydrazide | 44881558.98 | 45628157.05 | 17088823.9 | 35007803.18 | 40083983.34 | 31564670.78 | 39809165.39 | 39719550.24 | 37789338.05 | 46474909.28 | 33081579.68 | 35616910.05 |

| Metabolites | PBS_1 | PBS_2 | PBS_3 | PBS_4 | BCG_1 | BCG_2 | BCG_3 | BCG_4 | BCGΔ*pe18*_1 | BCGΔ*pe18*_2 | BCGΔ*pe18*_3 | BCGΔ*pe18*_4 |
|---|---|---|---|---|---|---|---|---|---|---|---|---|
| LNAPS 24:2/N-11:0 | 506254.3005 | 495281.2155 | 382221.4175 | 822227.9927 | 1334407.284 | 701593.5079 | 1375539.159 | 1075284.693 | 1630966.378 | 2008484.785 | 1165760.657 | 890055.9366 |
| 5-Carbethoxycytosine | 8683741.516 | 6755975.621 | 5544074.148 | 4593772.762 | 28696845.93 | 19363959.05 | 27524732.48 | 24226477.81 | 24402199.72 | 29386003.05 | 27961706.77 | 15546875.12 |
| PE 18:0_22:5;2O | 2584303.328 | 3273332.144 | 1514171.35 | 2436113.564 | 2809610.047 | 1696721.219 | 2860799.989 | 2762431.858 | 3475166.491 | 2573575.08 | 2956048.593 | 1648666.546 |
| PI 19:0_15:1 | 1786777.296 | 1367761.168 | 2171683.501 | 1616556.815 | 3029788.879 | 1947687.686 | 2928599.905 | 2709142.773 | 2748350.61 | 3518961.756 | 3676833.495 | 1742556.366 |
| SL 10:1;O/26:6 | 351214533.1 | 339320089.9 | 294640216.4 | 325285578 | 371846420.5 | 351465798.6 | 334970512.3 | 310088789.7 | 342154247.5 | 344415849.7 | 335240407 | 312986764 |
| MG 17:0 | 9545231.185 | 8659957.61 | 7696489.874 | 8174974.061 | 9135085.976 | 2236372.427 | 8031613.179 | 7805312.76 | 7012864.2 | 6847803.095 | 6716769.984 | 6304757.569 |
| 2,2-Dimethylpiperazine-1,4-diol | 4585019.303 | 4210625.591 | 6388447.823 | 4508371.57 | 10379523.07 | 7695821.77 | 2315096.334 | 6742839.986 | 7093157.57 | 5833201.719 | 7206423.137 | 6546507.227 |
| 16-Feruloyloxypalmitate | 1079120.281 | 1078874.277 | 2225464.183 | 960552.0697 | 962062.5753 | 1093108.074 | 917087.2309 | 913705.1679 | 1102055.721 | 864706.1995 | 1064382.289 | 852084.783 |
| 2,5-Dichlorothiophene-3-carboxylic acid | 12065074.52 | 10363492.88 | 13158000.74 | 11787660.01 | 9821007.379 | 12590800.77 | 10054399.07 | 13288088.76 | 10630672.18 | 10725121.95 | 12470859.94 | 11749732.47 |
| Methyl gallate | 2130467.848 | 2383172.725 | 1735653.076 | 2418053.951 | 1267990.207 | 1208833.587 | 1297290.764 | 2240593.053 | 939631.4109 | 2193898.535 | 1770643.02 | 1452373.558 |
| Trifolirhizin | 15115806.23 | 15266027.21 | 15263353.74 | 15083600.7 | 15145901.49 | 15468932.84 | 15558244.26 | 15614503.98 | 15054033.16 | 15115933.72 | 15493884.18 | 15148972.21 |
| PI(18:1(9Z)/16:0) | 180702 | 201969 | 123463 | 155761 | 470707 | 502705 | 832965 | 616562 | 833930 | 741981 | 577777 | 524450 |

| **Metabolites** | **PBS_1** | **PBS_2** | **PBS_3** | **PBS_4** | **BCG_1** | **BCG_2** | **BCG_3** | **BCG_4** | **BCGΔ*pe18*_1** | **BCGΔ*pe18*_2** | **BCGΔ*pe18*_3** | **BCGΔ*pe18*_4** |
|---|---|---|---|---|---|---|---|---|---|---|---|---|
| | 80.22 | 04.55 | 05.11 | 61.95 | 2.786 | 7.941 | 4.819 | 1.741 | 1.627 | 5.556 | 6.467 | 9.274 |
| MGDG 3:0_21:1 | 956389236.7 | 929038168 | 801773175.5 | 884256947.6 | 999041157.2 | 952936374 | 909590349.9 | 838650243 | 941286727.5 | 928148334.7 | 897190731.6 | 846583851.9 |
| [1,1':4',1''-Terphenyl]-3,4'',5-triol | 166225.385 | 153705.6497 | 221963.7447 | 165444.5488 | 36104693.91 | 53681090.26 | 102627707.6 | 102929014.2 | 115726866.4 | 41464518.49 | 78378644.78 | 113849978.4 |
| Dibutylamine | 29716898.76 | 31676386.1 | 25420815.92 | 30701503.98 | 24797323.65 | 21704409.97 | 24623460.8 | 28034526.3 | 23184592.34 | 18960879.65 | 20355035.97 | 34705756.75 |
| 1-Monopalmitin | 4530518.018 | 5214029.733 | 4838390.692 | 4562730.955 | 4816981.139 | 4459903.146 | 4198955.645 | 4654101.978 | 4343577.188 | 4541563.233 | 4656767.448 | 4976661.364 |
| Trehalose-6-phosphate | 0 | 0 | 0 | 4658.744758 | 6010696.927 | 4844554.971 | 3645017.958 | 4092474.099 | 6196300.119 | 6254939.147 | 5510992.735 | 5654076.259 |
| LPE 19:0 | 1323110.026 | 1206983.552 | 874607.1971 | 854760.0714 | 1000199.916 | 858388.1239 | 1487533.269 | 1147234.307 | 1923654.834 | 5333741.566 | 1658509.324 | 913910.5979 |
| M162T198 | 4002098.513 | 875872.5418 | 2454102.393 | 1082476.92 | 10164971.2 | 1084212.841 | 9462547.774 | 3203832.376 | 7857130.787 | 456517.8724 | 5738022.597 | 9830604.503 |
| 1,3,4,5-Tetrahydrothiopyrano[4,3-b]indole | 685843.3744 | 580928.2004 | 878375.7171 | 580193.5418 | 4145776.984 | 2763313.945 | 3417993.833 | 2837031.41 | 3238156.839 | 3333039.198 | 3475513.707 | 2896423.082 |
| 4-(4-Methyl-1-piperazinyl)benzaldehyde | 26484997.36 | 30641920.01 | 18620964.64 | 25950976.44 | 26843449.92 | 21767379.84 | 27336279.84 | 27786767.04 | 30275241.94 | 26478511.19 | 26105170.95 | 23529025.21 |
| 1-Methylnicotinamide | 1320762.664 | 3426151.909 | 3433134.593 | 3394914.449 | 3308022.021 | 2236064.068 | 5988596.917 | 2878874.02 | 3447725.74 | 3221281.703 | 2533591.461 | 11536456.72 |
| Adenosine | 895852 | 806967 | 122056 | 107653 | 200552 | 182653 | 136941 | 148816 | 171029 | 136501 | 154539 | 214412 |

| Metabolites | PBS_1 | PBS_2 | PBS_3 | PBS_4 | BCG_1 | BCG_2 | BCG_3 | BCG_4 | BCGΔ*pe18*_1 | BCGΔ*pe18*_2 | BCGΔ*pe18*_3 | BCGΔ*pe18*_4 |
|---|---|---|---|---|---|---|---|---|---|---|---|---|
| | 19.83 | 75.45 | 78.38 | 526.4 | 040.1 | 072.2 | 788.7 | 124.5 | 263.3 | 645.8 | 137.2 | 420.1 |
| TG 16:4_16:4_16:4 | 3386485.541 | 3421267.792 | 1710135.187 | 3513896.783 | 2776202.289 | 5002487.995 | 2600817.338 | 4713761.243 | 5117110.638 | 5795322.643 | 3375700.647 | 3559174.331 |
| 4-(4-Methyl-1-piperazinyl)butanoic acid | 1588675.493 | 2263672.84 | 1957231.864 | 2678684.325 | 1872191.71 | 1466897.751 | 2607251.611 | 2121877.596 | 2073954.237 | 2103431.423 | 1809285.21 | 3354987.994 |
| DGTS 13:0_18:5 | 8876260.902 | 14790360.08 | 7981037.216 | 12115639.58 | 9883257.707 | 8458228.506 | 9304636.147 | 12132136.71 | 8610076.92 | 10836667.26 | 9558453.989 | 11775193.9 |
| Cer(d18:1/16:0) | 47703506.76 | 53002477.84 | 66692669.63 | 49097130.66 | 45406142.02 | 37115484.53 | 56771180.39 | 49937969.98 | 49685398.48 | 61788666.98 | 54897453.82 | 26000589.45 |
| SPI_298.2741_15.1 | 10449859.17 | 12093063.67 | 7735735.786 | 8106438.364 | 10103222.31 | 9720832.65 | 10746279.27 | 9980721.249 | 10882387.3 | 11050473.95 | 11165549.27 | 9347241.485 |
| SL 10:0;O/18:5 | 789973.6536 | 1115779.278 | 372053.3209 | 615426.0924 | 636739.0988 | 476694.2276 | 428277.1515 | 395385.8631 | 537311.8558 | 561725.3606 | 447212.655 | 407356.143 |
| PE(P-18:0/20:4(5Z,8Z,11Z,14Z)) | 225751290 | 290191490.6 | 341158397.2 | 291461703.7 | 236378328.3 | 179174301.5 | 315780745.3 | 210665118.8 | 217957122.1 | 312099433.5 | 271298385.1 | 111797296 |
| 4-Dodecylphenol | 57070667.89 | 43704558.15 | 38335037.6 | 47486373.63 | 53945403.03 | 52011658.13 | 63880426.99 | 50656946.78 | 50882936.25 | 56692469.18 | 65179632.28 | 42374549.46 |
| SL 12:2;O/18:5 | 110378799.9 | 117072881.4 | 102612452.6 | 105853251 | 116280969.5 | 124997275.6 | 114173397.1 | 106126901.1 | 105100135.8 | 120125801.9 | 108960906.9 | 114098678.3 |
| 9H-Carbazole, 2-[(2E)-2-(1-azabicyclo[2.2.2]oct-3-ylidene)-2-fluoroethox | 5051769.278 | 4624854.97 | 6566018.522 | 6245386.971 | 4993028.16 | 4279292.929 | 6415265.551 | 1621164.046 | 5200835.982 | 6075861.589 | 5414788.402 | 4338993.384 |

| Metabolites | PBS_1 | PBS_2 | PBS_3 | PBS_4 | BCG_1 | BCG_2 | BCG_3 | BCG_4 | BCGΔ *pe18*_1 | BCGΔ *pe18*_2 | BCGΔ *pe18*_3 | BCGΔ*pe18*_4 |
|---|---|---|---|---|---|---|---|---|---|---|---|---|
| y]- | | | | | | | | | | | | |
| N-(2'-Oxobi(cyclohex)yl)acetamide | 29461131.52 | 31558980.42 | 30604755.67 | 30282179.75 | 35863228.91 | 31577331.05 | 31691026.33 | 31124409.7 | 29546973.96 | 33069177.1 | 31977220.14 | 32455264.92 |
| PC(40:4) | 161894885.8 | 210197950.7 | 74159853.08 | 125869290.2 | 53755590.6 | 41147835.61 | 48235212.13 | 56803867.51 | 70194292.73 | 66506830.79 | 41997849.31 | 33914378.87 |
| Retreversine | 507169.3998 | 496230.6231 | 586473.0431 | 525726.4441 | 426479.5975 | 470903.8513 | 442834.5034 | 411089.6102 | 451772.1999 | 420344.7587 | 458527.8873 | 450739.7987 |
| SMGDG O-9:0_16:2 | 1899038.287 | 1928215.527 | 1854288.661 | 1829860.721 | 1820066.011 | 1834961.292 | 1796567.104 | 1798916.717 | 1839323.564 | 1737095.842 | 1946453.29 | 1845748.578 |
| N,N-Dimethyl-N-(2-phenoxyethyl)-1-dodecanaminium cation | 47389853.14 | 35890351.45 | 30016575.65 | 23990453.81 | 17180771.02 | 28906408.77 | 23031020.95 | 20136316.84 | 21383981.33 | 22280239.42 | 24438368.83 | 20544040.87 |
| [8-[2-(3-methylbutanoyloxy)propan-2-yl]-2-oxo-8,9-dihydrofuro[2,3-h]chromen-9-yl] (Z)-2-methylbut-2-enoate | 2601262.223 | 2722310.799 | 2451073.286 | 2738898.915 | 2737960.341 | 2852787.858 | 2810752.127 | 2681972.804 | 2757143.978 | 2884991.651 | 2692671.764 | 2604554.523 |
| 2-Ethylbutan-1-amine | 2536683647 | 2486431142 | 1994431853 | 3069457892 | 3098746982 | 2910322355 | 2507816172 | 2381249155 | 2527248338 | 2335199145 | 2922530345 | 2886442730 |

| Metabolites | PBS_1 | PBS_2 | PBS_3 | PBS_4 | BCG_1 | BCG_2 | BCG_3 | BCG_4 | BCGΔ*pe18*_1 | BCGΔ*pe18*_2 | BCGΔ*pe18*_3 | BCGΔ*pe18*_4 |
|---|---|---|---|---|---|---|---|---|---|---|---|---|
| 4-tert-Butyl-N-(3-quinolinyl)benzamide | 269313609.7 | 279427835.5 | 275337517.2 | 297148558.9 | 260176332.9 | 272640471.1 | 303228598.6 | 303856500.3 | 275092535.1 | 266830305.8 | 302636529.3 | 258050774.9 |
| Phenylalanine betaine | 4269704.807 | 4764758.011 | 3204798.139 | 3712904.349 | 3877835.804 | 3244465.358 | 4684560.569 | 3945756.639 | 4766520.369 | 4070211.77 | 4061045.711 | 3969314.892 |
| Fenoxaprop-P-ethyl | 2841325.546 | 2563153.045 | 2542342.994 | 2854517.54 | 4789428.147 | 6226345.167 | 6802163.444 | 5252063.613 | 4668172.651 | 4168864.891 | 8837550.202 | 5082984.262 |
| Methyl 6-(acetyloxy)-1,3,7,11-tetrahydroxyabieta-8,11,13-trien-18-oate | 99310152.28 | 96360116.09 | 89144531.63 | 95638738.83 | 98148732.12 | 101169680.4 | 99905543.89 | 97209999.37 | 98726218.91 | 95321267.57 | 94893661.8 | 95326023.42 |
| Rhodinyl Acetate | 42988247.45 | 46628910.01 | 46877243.67 | 44984912.81 | 54301077.41 | 47615202.15 | 48256902.73 | 46241749.29 | 44217106.29 | 49301053.99 | 47366709.86 | 48743867.95 |
| DGCC 9:0_10:0 | 1031217728 | 1127406465 | 937317590.4 | 958793597.7 | 1074684275 | 1128712541 | 1031100569 | 998073257.1 | 961194491 | 1086940047 | 1009145487 | 1025937332 |
| 2-(2-Thienyl)-1H-benzimidazole | 2868592.601 | 3017690.791 | 2935277.723 | 3024164.607 | 2839397.081 | 2807487.61 | 2944015.971 | 2952226.876 | 3063540.775 | 2737477.277 | 2809465.208 | 2787471.779 |
| MGDG O-20:2_3:0 | 55505227.64 | 58277753.69 | 56261104.27 | 56957603.46 | 50832411.17 | 57414797.21 | 55802002.03 | 63560947.61 | 57287235.6 | 60426775.13 | 51832624.24 | 52762095.04 |
| Cer 8:1;2O/7:0 | 110769207.8 | 113437557.7 | 97658773.43 | 112654698.6 | 106808369.1 | 105443216.4 | 113287381.6 | 113240235.1 | 105891678.5 | 109928496.9 | 110069077.3 | 105548101.5 |
| 16-Hydroxypalmitic acid | 5418747.5 | 5474283.146 | 9957641.59 | 6289064.226 | 6651421.582 | 5746813.062 | 8032245.145 | 5536076.467 | 5664795.733 | 6260828.948 | 7975095.729 | 9616192.856 |

| Metabolites | PBS_1 | PBS_2 | PBS_3 | PBS_4 | BCG_1 | BCG_2 | BCG_3 | BCG_4 | BCGΔ*pe18*_1 | BCGΔ*pe18*_2 | BCGΔ*pe18*_3 | BCGΔ*pe18*_4 |
|---|---|---|---|---|---|---|---|---|---|---|---|---|
| PEtOH 20:3_22:5 | 2446199.892 | 1986079.988 | 2086699.533 | 1944990.82 | 1798162.346 | 0 | 1754565.681 | 729851.9466 | 1524154.839 | 1701395.772 | 2331177.396 | 1216337.899 |
| PS 18:0_20:3 | 958346.715 | 759324.5683 | 1440003.237 | 784801.1568 | 1324727.188 | 317705.3725 | 2929903.566 | 1177702.477 | 1684658.133 | 947673.3148 | 1235965.235 | 2972669.543 |
| PE(18:1(9Z)/18:3(6Z,9Z,12Z)) | 15499736.79 | 18370200.28 | 13399319.51 | 14907956.61 | 57994797.66 | 38833598.5 | 43077733.5 | 42433411.38 | 50926519.7 | 70672324.28 | 43766873.22 | 40251593.23 |
| LNAPS 24:4/N-16:2 | 10399728.05 | 7101403.217 | 11231337.07 | 8681104.136 | 10301721.05 | 3999836.246 | 9302471.868 | 12192330.84 | 10846155.65 | 9642753.7 | 8632843.015 | 10363396.63 |
| PC O-30:7 | 15168257.89 | 15690367.37 | 13565076.37 | 14248070.48 | 16240713.4 | 15270588.12 | 15427074.63 | 15367957.67 | 14720756.27 | 15517622.91 | 15366705.55 | 14414330.76 |
| MG 11:0 | 46466277.8 | 48773705.14 | 41481027.48 | 47102962.87 | 48948201.13 | 46380330.55 | 47905277.49 | 49011552.17 | 48652264.16 | 48349308.64 | 45343703.3 | 43308676.22 |
| Cer 14:0;2O/4:0 | 8919363.023 | 10236672.25 | 9279954.981 | 9425901.78 | 8377114.234 | 8645784.632 | 8300278.905 | 8465162.131 | 8559394.778 | 8954417.574 | 8918953.189 | 8843874.831 |
| 7-Methoxy-3-(4-methoxyphenyl)-4-phenylcoumarin | 5610652.489 | 3611154.723 | 5474538.258 | 3160791.991 | 6823340.597 | 5081981.371 | 8356896.455 | 6746716.013 | 6780574.551 | 7403053.247 | 7399763.54 | 5184043.003 |
| N-(1,1-Dimethyl-2-hydroxyethyl)-3-amino-2-hydroxypropanesulfonic acid | 4759077.603 | 2246262.776 | 2383198.484 | 1994749.962 | 7279463.209 | 2164662.51 | 2444642.238 | 3261576.089 | 2378763.775 | 5773228.23 | 4753656.744 | 2236806.992 |
| Moxisylyte | 6947964.544 | 7411016.494 | 7779983.029 | 7256226.691 | 7178336.147 | 6526011.341 | 6177998.064 | 6163771.035 | 6541969.323 | 5368948.302 | 6705842.809 | 7203846.166 |
| PC(16:0/20:4) | 998694 | 119882 | 631050 | 743730 | 793957 | 607043 | 104678 | 812168 | 894692 | 123089 | 899273 | 522381 |

| Metabolites | PBS_1 | PBS_2 | PBS_3 | PBS_4 | BCG_1 | BCG_2 | BCG_3 | BCG_4 | BCGΔ *pe18*_1 | BCGΔ *pe18*_2 | BCGΔ *pe18*_3 | BCGΔ*pe18*_4 |
|---|---|---|---|---|---|---|---|---|---|---|---|---|
| | 913.4 | 9982 | 574.8 | 960.2 | 916.4 | 228.4 | 1351 | 576.7 | 745.5 | 8334 | 097.9 | 297.2 |
| (3.beta.,9.Xi.)-3-(.beta.-D-Glucopyranosyloxy)-14-hydroxycarda-5,20(22)-dienolide | 25702.01805 | 54294.73932 | 38019.14797 | 30003.12468 | 286553.0618 | 187308.9725 | 314238.0255 | 229268.1034 | 368428.981 | 364670.5516 | 454941.974 | 286195.2427 |
| Cer 18:0;2O/4:0 | 1676283.731 | 1584734.507 | 1135327.581 | 1218664.751 | 1021978.509 | 1384691.261 | 4198097.649 | 1052742.073 | 2109710.102 | 1177742.826 | 1228751.485 | 3485018.941 |
| ISOPALMITIC ACID | 29034671.57 | 25793416.99 | 28944144.24 | 2675500 6.1 | 30593470.05 | 31215712.58 | 28817000.69 | 29246337.21 | 28563111.38 | 30161010.18 | 31822269.13 | 29355394.24 |
| LysoPE(20:4(5Z,8Z,11Z,14Z)/0:0) | 6454165.685 | 6840416.204 | 8274859.505 | 6763984.756 | 6784883.114 | 5642576.165 | 7092099.504 | 6310721.465 | 5903647.955 | 8502687.727 | 8092781.062 | 6705616.99 |
| 1,3-Benzenediol, 5-(10-nonadecenyl)-, (Z)- | 3582627.049 | 3385291.108 | 5260080.955 | 12686254.64 | 995379.1711 | 5231623.473 | 17453064.48 | 380696.6825 | 13816408.73 | 13947414.92 | 958558.9775 | 13460261.89 |
| phosphatidylglycerol (1-palmitoyl, 2-cis-9,10-methylene hexadecanoyl) | 2659646.583 | 2532654.185 | 2575344.864 | 2537486.47 | 2987757.887 | 2754396.518 | 3111705.66 | 3186673.651 | 4685961.883 | 4643547.722 | 3153760.795 | 1903615.104 |
| SL 14:3;O/10:0 | 1564931.603 | 1716197.059 | 1480171.469 | 1572346.857 | 1466129.12 | 1589471.748 | 1531803.992 | 1598460.441 | 1546059.429 | 1491500.784 | 1499682.396 | 1524598.687 |
| 4-(Hydroxymethyl)benzenesulfonic acid | 8538787.484 | 212410.4156 | 4772264.649 | 24756513.02 | 1956255.063 | 241737.3937 | 842053.1384 | 170231.4092 | 2265188.084 | 2805807.208 | 3056994.503 | 1072856.449 |

| Metabolites | PBS_1 | PBS_2 | PBS_3 | PBS_4 | BCG_1 | BCG_2 | BCG_3 | BCG_4 | BCGΔ *pe18*_1 | BCGΔ *pe18*_2 | BCGΔ *pe18*_3 | BCGΔ*p e18*_4 |
|---|---|---|---|---|---|---|---|---|---|---|---|---|
| MGDG O-19:2_3:0 | 102707 8916 | 106372 7646 | 890459 001 | 942968 824.4 | 110363 3856 | 108024 8811 | 104805 0438 | 976288 681.4 | 990396 765.3 | 106833 6819 | 993305 080.5 | 990960 165.7 |
| Sodium (4-methoxybenzoyl oxy)acetate | 767488 11.91 | 721701 26.11 | 740356 04.02 | 927795 22.24 | 729510 87.1 | 919464 79.91 | 747487 78.47 | 744088 27.49 | 746200 34.72 | 734481 08.73 | 754331 74.94 | 781773 14.12 |
| 2-amino-3-({[3-(he xadecanoyloxy)-2-( octadecanoyloxy)pr opoxy](hydroxy)ph osphoryl}oxy)propa noic acid | 340593 35.06 | 410555 58.68 | 289923 95.49 | 313424 07.39 | 426201 32.88 | 341992 45.22 | 509687 56.45 | 437975 25.11 | 448400 71.01 | 632797 48.21 | 509441 77.03 | 217749 42.94 |
| Tri-2-ethylhexyl trimellitate | 336722 803.4 | 351671 631.8 | 303187 867.2 | 315797 317.8 | 350550 904.6 | 368843 197 | 333357 074.7 | 323030 885.7 | 312138 427.9 | 353494 366.9 | 330090 534.3 | 333252 631.3 |
| N-(1-Azabicyclo[2. 2.2]oct-3-yl)-1-met hyl-1H-indole-6-car boxamide | 627676. 5347 | 674022. 6018 | 779207. 3938 | 765335. 5775 | 584488. 206 | 620668. 8861 | 643106. 1374 | 574640. 8117 | 688139. 0614 | 660039. 6861 | 632313. 5258 | 970735. 1419 |
| Nona-2,4,7-trienoyl carnitine | 202495 3.282 | 236131 4.216 | 259888 3.634 | 209694 6.569 | 183284 5.423 | 200029 1.64 | 191684 9.272 | 171398 1.596 | 206972 8.769 | 198816 3.404 | 185586 7.748 | 283978 7.599 |
| 1H-Indole-3-carbox amide, N-1-naphthalenyl-1 -pentyl- | 337749 9.538 | 336952 1.649 | 286308 5.052 | 338195 1.815 | 322461 7.345 | 348673 8.025 | 326563 2.429 | 326306 1.88 | 332460 2.558 | 325831 9.976 | 321411 8.946 | 308769 3.09 |
| 11-Keto-.beta.-bos | 738319. | 759700. | 757273. | 739891. | 741593. | 760897. | 717865. | 733749. | 750653. | 734767. | 732313. | 733543. |

| Metabolites | PBS_1 | PBS_2 | PBS_3 | PBS_4 | BCG_1 | BCG_2 | BCG_3 | BCG_4 | BCGΔ*pe18*_1 | BCGΔ*pe18*_2 | BCGΔ*pe18*_3 | BCGΔ*pe18*_4 |
|---|---|---|---|---|---|---|---|---|---|---|---|---|
| wellic acid | 3243 | 6699 | 3666 | 1747 | 1221 | 8651 | 2353 | 2363 | 4611 | 325 | 5386 | 9646 |
| PA 5:0_18:5 | 18905052.55 | 20505395.65 | 14299368.19 | 17352092.2 | 17789782.12 | 18571041.26 | 18928086.98 | 17042893.18 | 17797475.01 | 18038502.47 | 17057272 | 17347483.77 |
| 8Z,11Z,14Z-Eicosatrienoic acid, ethyl ester | 81773708.37 | 89785438.22 | 73282224 | 80164199.83 | 87081986.34 | 80503388.85 | 87266492.27 | 86169154.61 | 83989748.62 | 83977201.04 | 83130832.24 | 75996730.59 |
| PC(20:1(11Z)/20:3(5Z,8Z,11Z)) | 32542856.8 | 41086445.09 | 41869016.7 | 32230073.82 | 33387450.38 | 27814513.18 | 47590471.39 | 34472932.88 | 40887622.13 | 52121548.77 | 40519883.86 | 20288983.51 |
| 2-Mercaptomethylbenzimidazole | 210367921.2 | 229662652.3 | 254472391.6 | 324549353.8 | 320395125.6 | 295427476.9 | 205838310.4 | 185719453.1 | 214386501.4 | 213115805.1 | 281178501 | 265663312.5 |
| Isoleucine | 72265927.07 | 60343723.63 | 99592569.84 | 78615060.69 | 52770114.91 | 101281583.1 | 96035389.02 | 45890716.95 | 109327845.5 | 67114693.6 | 36086366.7 | 88021012.79 |
| 1-(Cyclohexylmethyl)proline | 4752433.434 | 6774737.49 | 8324341.275 | 8135185.268 | 5334229.95 | 6680272.831 | 9305934.984 | 4076377.805 | 5879542.061 | 6150695.127 | 5759171.554 | 10501966.26 |
| NAGly 16:3/16:3 | 21678219.76 | 27993722.26 | 18988189.21 | 22325256.09 | 12823030.7 | 13924720.59 | 15928624.34 | 11573780.99 | 11335771.81 | 13995742.08 | 18204320.07 | 9933630.618 |
| Dodecyl gallate | 58267508.44 | 68672619.53 | 31386760.44 | 73667472.08 | 34328589.96 | 42579574.28 | 36400800.55 | 50483831.81 | 31370119.37 | 49286148.39 | 38462529.99 | 78563683.11 |
| 1-palmitoleoyl-sn-glycero-3-phosphoethanolamine | 483512.6125 | 389140.7376 | 414092.9824 | 408532.1038 | 409550.9447 | 366179.7376 | 350953.3717 | 409245.6143 | 804563.7116 | 737470.0334 | 704692.1051 | 790068.2414 |
| MMV689480 | 13453911.33 | 14117002.33 | 12496130.92 | 13875453.89 | 14253880.65 | 14116993.98 | 14532389.92 | 14487883.22 | 14458782.43 | 14648063.79 | 13518892.46 | 12710698.8 |

| Metabolites | PBS_1 | PBS_2 | PBS_3 | PBS_4 | BCG_1 | BCG_2 | BCG_3 | BCG_4 | BCGΔ*pe18*_1 | BCGΔ*pe18*_2 | BCGΔ*pe18*_3 | BCGΔ*pe18*_4 |
|---|---|---|---|---|---|---|---|---|---|---|---|---|
| Cer 9:0;2O/26:7 | 8397112.17 | 13259323.81 | 7544922.551 | 18206358.73 | 8266794.157 | 7916642.302 | 6269819.841 | 10498406.06 | 7358746.541 | 10458819.05 | 9685480.288 | 8453650.866 |
| 2-Amino-4,5-dimethyl-3-thiophenecarbonitrile | 163247946 | 156320306.4 | 148018263 | 177424925.1 | 152147213.8 | 157895009.2 | 198676799.1 | 186513481.9 | 152101903 | 173957983.4 | 199990496.8 | 156813735.3 |
| Rimantadine | 136874228.9 | 128723845.6 | 118873713 | 130079220.7 | 132261143.2 | 135115198.8 | 139866619.2 | 143763948 | 141535449.1 | 138237748.5 | 128147106.5 | 124521457.3 |
| 11-(3-Methoxybenzyl)-4-piperidinylamine | 10836943.12 | 14275139.91 | 9474456.416 | 10044786.77 | 10800821.39 | 8014832.589 | 15400266.61 | 14122613.09 | 13886360.75 | 13361251.87 | 11322764.44 | 11332200.83 |
| 5,7-Dibromo-2,1-benzisothiazol-3-amine | 36386738.4 | 38347494.85 | 36458641.88 | 36270777.97 | 35981164.25 | 38860458.36 | 36617628.88 | 36672677.41 | 35254920.68 | 34840477.07 | 36776793.13 | 37029007.24 |
| (R)-Aminocarnitine | 2155263.234 | 1384249.418 | 4500922.722 | 1856994.213 | 22417774.94 | 27084490.23 | 22982072.96 | 23362138.77 | 25277900.87 | 20683498.31 | 24347442.77 | 26289908.12 |
| Hydroheptelidic acid | 2597668.868 | 2383210.641 | 1907742.595 | 2834620.191 | 2205558.939 | 3039440.173 | 2451225.142 | 2372265.346 | 2366303.143 | 2039128.472 | 7486986.303 | 2968464.713 |
| PC(18:1(9Z)/18:3(9Z,12Z,15Z)) | 15597289 | 15486833.16 | 14808086.79 | 10779180.27 | 15644760.97 | 10908393.67 | 18706956.9 | 8889197.625 | 16178050.59 | 21658707.14 | 12326224.18 | 13805945.95 |
| 1-Dodecanesulfonic acid | 637098.5737 | 1103945.236 | 2655617.092 | 1374058.774 | 757182.078 | 1467326.624 | 816785.8476 | 718824.4425 | 1486011.107 | 763528.7471 | 672702.3145 | 4561196.67 |
| Cyphenothrin | 1746143.292 | 1713109.596 | 1540831.12 | 1620523.192 | 1697242.153 | 1696624.63 | 1646307.835 | 1588817.487 | 1636635.412 | 1723688.21 | 1710517.602 | 1484673.883 |

| Metabolites | PBS_1 | PBS_2 | PBS_3 | PBS_4 | BCG_1 | BCG_2 | BCG_3 | BCG_4 | BCGΔ*pe18*_1 | BCGΔ*pe18*_2 | BCGΔ*pe18*_3 | BCGΔ*pe18*_4 |
|---|---|---|---|---|---|---|---|---|---|---|---|---|
| 2-Methoxy-5-methyl-3-(2-methylbut-3-en-2-yl)chromen-4-one | 1912763.449 | 2020925.932 | 1829326.283 | 1815915.633 | 1658856.218 | 1716605.968 | 1830472.602 | 1741649.168 | 1880942.292 | 1703879.747 | 1792019.453 | 1822021.985 |
| SM(d18:1/14:0) | 10570090.22 | 11767549.58 | 6488319.543 | 7514452.801 | 17076563.91 | 12979539 | 18262140.14 | 16261150.26 | 19409696.79 | 19961872.96 | 14736591.81 | 10674274.54 |
| Mevalonic Acid Lactone | 1713908.43 | 2174601.78 | 2614196.334 | 2248618.179 | 1568875.144 | 1583645.274 | 2296173.532 | 1576346.628 | 2375332.665 | 2220883.323 | 1246338.205 | 1415306.702 |
| N-Acetylaminooctanoic acid | 3115606.427 | 3972261.727 | 3577341.511 | 5540616.567 | 3054037.628 | 3021329.552 | 5067452.956 | 2542685.206 | 3302461.763 | 3643251.194 | 2982093.541 | 6526661.903 |
| Trehalose | 473170.8303 | 447306.8038 | 656803.8536 | 523157.5728 | 154225535.8 | 139888358.6 | 115797351.5 | 117316471.7 | 160291042.3 | 122417727.8 | 130177395.6 | 159424984.6 |
| PG 3:0_13:1 | 2795901.011 | 2952647.472 | 2721351.421 | 2682998.992 | 2795343.019 | 2562060.365 | 2869658.099 | 2778018.857 | 2949590.007 | 2712112.369 | 2605674.47 | 2676945.317 |
| DGTS 16:4_18:5 | 567167094.4 | 565848416.8 | 441788865.3 | 475473629.7 | 481744528 | 471991843.7 | 518882827.2 | 471479346.4 | 503614893.7 | 506016390.2 | 507541345.1 | 438163863.1 |
| MGDG O-18:4_18:1 | 6404880.342 | 8070240.551 | 6053539.29 | 6349139.547 | 7188706.846 | 191920.9924 | 6320805.585 | 5110692.956 | 8297304.453 | 6849739.147 | 7880886.752 | 3830031.348 |
| PC(22:4(7Z,10Z,13Z,16Z)/P-18:0) | 79658877.52 | 108237658.7 | 54020498.8 | 68928021.13 | 25608309.53 | 20154676.06 | 29798627.13 | 26483005.69 | 34863602.18 | 41305418.88 | 27031433.9 | 16352422.65 |
| M314T99 | 4747619.649 | 4398354.903 | 4150511.489 | 4203101.83 | 3464773.887 | 4098037.784 | 3248221.229 | 3777745.239 | 4016518.707 | 3665015.566 | 4199864.958 | 4007416.92 |
| LPI 20:4 | 136794832.8 | 142154745.6 | 168646998.3 | 154152849.7 | 59077586.67 | 43972229.31 | 44769721.5 | 42226340.68 | 49713414.51 | 51257737.26 | 59318547.71 | 42311643.38 |

| Metabolites | PBS_1 | PBS_2 | PBS_3 | PBS_4 | BCG_1 | BCG_2 | BCG_3 | BCG_4 | BCGΔ*pe18*_1 | BCGΔ*pe18*_2 | BCGΔ*pe18*_3 | BCGΔ*pe18*_4 |
|---|---|---|---|---|---|---|---|---|---|---|---|---|
| Gossypol | 5896037.12 | 759166.7614 | 3173136.021 | 720782.1159 | 724342.4741 | 4300355.854 | 1853481.746 | 2258647.04 | 4938767.509 | 962715.8211 | 5210000.393 | 985178.3578 |
| Methyl (2E,8E)-9-[3a-hydroxy-6,7-dimethyl-1-(2-methylpropyl)-3-oxo-2,4,7,7a-tetrahydro-1H-isoindol-4-yl]-4,5-dihydroxy-8-methylnona-2,8-dienoate | 37660520.84 | 41071361.94 | 33935417.49 | 37171916.66 | 37882718.35 | 35979053.97 | 37783350.2 | 42719021.79 | 36850965.3 | 39363023.54 | 38590464.31 | 33962728.62 |
| Adenosine monophosphate (AMP) | 6926531.746 | 7734686.061 | 8084103.07 | 8897266.363 | 16821627.04 | 17953996.68 | 15960013.3 | 16415541.14 | 17813181.27 | 15659450.17 | 18704840.82 | 23342325.51 |
| Deserpidine | 517387.8594 | 453890.4842 | 766854.5539 | 702841.3659 | 1184887.133 | 1348250.33 | 766820.3785 | 1213060.988 | 1133557.498 | 604241.131 | 1025606.617 | 1721727.596 |
| NAE 16:2 | 135388823.4 | 115979038.4 | 75645193.01 | 129537536.8 | 114244625.4 | 111301200.3 | 94474467.5 | 120030587.2 | 112197150.6 | 119575548.1 | 110777244.5 | 83374933.84 |
| Kahweol acetate | 5312816.791 | 5281597.327 | 4627909.574 | 5338706.465 | 5952030.817 | 5143818.482 | 5775787.451 | 5584677.087 | 5719775.499 | 5586691.864 | 5387886.132 | 4893215.18 |
| LPS 22:1 | 3919742.29 | 3027753.943 | 6451996.433 | 4889358.341 | 9925313.39 | 9994118.383 | 9895264.315 | 9799473.302 | 14630114.27 | 8868665.721 | 10801882.51 | 9097683.582 |
| SM(d18:0/16:0) | 58403081.55 | 39868494.93 | 55886349.02 | 67600800.45 | 43141994.38 | 696787.7196 | 50123516.62 | 22466743.17 | 60198228.22 | 48706903.38 | 54736386.95 | 16261587.44 |

| Metabolites | PBS_1 | PBS_2 | PBS_3 | PBS_4 | BCG_1 | BCG_2 | BCG_3 | BCG_4 | BCGΔ*pe18*_1 | BCGΔ*pe18*_2 | BCGΔ*pe18*_3 | BCGΔ*pe18*_4 |
|---|---|---|---|---|---|---|---|---|---|---|---|---|
| FA 14:1 | 11768997.84 | 10640846.46 | 10384991.1 | 8832640.675 | 8991180.845 | 10732045.41 | 9923251.606 | 10409804.86 | 9033070.777 | 12793014.85 | 11159293.42 | 10101144.05 |
| L-Thyronine | 10237593.8 | 10578958.09 | 8939626.591 | 10304038.31 | 10073537.01 | 9666216.928 | 10066082.08 | 10395563.29 | 9811314.5 | 10092978.01 | 10046191.23 | 9758391.735 |
| TG 12:0_14:1_18:5 | 5638026.67 | 7836104.544 | 7266881.212 | 5645128.68 | 7258653.15 | 5912480.426 | 7620446.252 | 6269599.985 | 7725393.228 | 7757058.853 | 7911529.272 | 6240102.062 |
| 2,2,4,7-Tetramethyl-1,2-dihydroquinoline | 147207350.8 | 178829710.1 | 143190773 | 81779194.42 | 132217358.8 | 115481049.5 | 123622204.7 | 112732892.1 | 124199668 | 117291178.1 | 64414254.68 | 181635181.3 |
| M419T416 | 1937481.28 | 2140977.812 | 1831310.195 | 1931265.965 | 2000463.048 | 1913522.56 | 2146777.996 | 1996639.458 | 2005104.263 | 2066665.127 | 1972392.378 | 1948416.495 |
| PC O-34:2 | 11958203.18 | 10891211.48 | 18790468.42 | 14648713.61 | 15626981.87 | 367498.8998 | 9664387.213 | 10169285.15 | 11173832.4 | 12002698.44 | 13380301.58 | 7310263.924 |
| (2-aminoethoxy)[2-[octadec-9-enoyloxy]-3-(octadecanoyloxy)propoxy]phosphinic acid | 20995865.1 | 25065490.46 | 45440078.64 | 30034030.42 | 75028578.29 | 54311498.45 | 77763051.82 | 57927165.81 | 61909931.73 | 80390496.67 | 75805024.78 | 35444144.27 |
| Bistris | 1916806.714 | 4080005.757 | 3833875.192 | 7479054.979 | 1251520.104 | 5111282.238 | 2679631.141 | 3781330.977 | 3955899.867 | 4583682.084 | 2353342.987 | 2407067.907 |
| Octaethylene glycol | 2612801.517 | 2496453.43 | 2868568.971 | 2944859.567 | 2616137.523 | 2383284.034 | 2914501.315 | 2389516.573 | 2909854.537 | 2901106.172 | 3022027.129 | 3033879.861 |
| SM 40:6;3O | 22718893.45 | 30417861.67 | 13138477.68 | 20199557.46 | 5149023.017 | 5235008.705 | 5067420.884 | 6253867.708 | 6907197.078 | 6286603.49 | 5194114.195 | 5782705.604 |

| **Metabolites** | **PBS_1** | **PBS_2** | **PBS_3** | **PBS_4** | **BCG_1** | **BCG_2** | **BCG_3** | **BCG_4** | **BCGΔ*pe18*_1** | **BCGΔ*pe18*_2** | **BCGΔ*pe18*_3** | **BCGΔ*pe18*_4** |
|---|---|---|---|---|---|---|---|---|---|---|---|---|
| VAE 21:0 | 23063729.41 | 19890017.16 | 10713935.41 | 12379593.87 | 20426028.17 | 20126696.99 | 16703636.87 | 19044382.78 | 21744714.05 | 21487074.32 | 22150080.74 | 13131886.6 |
| 3-Isopropylmalic acid | 661238.9901 | 605361.1204 | 546571.8097 | 531895.0449 | 15002523.28 | 7546254.865 | 9712871.644 | 7272613.69 | 9345675.286 | 11280232 | 12064901.5 | 9194907.26 |
| MGDG O-10:0_26:7 | 5063638.066 | 7237363.288 | 4214888.449 | 6797358.034 | 4778863.023 | 4496739.936 | 5079563.48 | 5731490.01 | 4666656.202 | 6147062.834 | 4927805.755 | 6119796.417 |
| SM 8:0;2O/15:1 | 9513710.483 | 9923904.946 | 8611283.788 | 8646255.119 | 10007803.11 | 10177919.45 | 9797008.182 | 9574168.671 | 8789263.379 | 10455393.69 | 9012407.068 | 9679907.283 |
| PG 16:0_16:1 | 631526.8013 | 592553.4034 | 419725.8431 | 416577.2512 | 2341980.368 | 1989774.739 | 2043590.745 | 5081116.487 | 5749208.118 | 7248629.533 | 6031996.048 | 1877437.292 |
| SQDG 24:2_18:3 | 9323759.553 | 10635809.98 | 8817408.704 | 9334243.806 | 10367302.22 | 10558981.01 | 9365735.404 | 10273505.49 | 9370761.454 | 10456942.37 | 10396033.21 | 9371845.381 |
| SM 25:1;2O/22:0 | 779097.4337 | 452638.4505 | 501022.3659 | 518139.1132 | 697822.6374 | 0 | 462257.0124 | 500179.5495 | 556007.3714 | 604528.9185 | 718547.8354 | 443195.0879 |
| (2-{[2-(2,6-Dimethylphenoxy)ethyl]sulfanyl}-1H-benzimidazol-1-yl)acetic acid | 2207070.115 | 1739555.09 | 5730111.84 | 2083047.04 | 14878860.44 | 3235961.585 | 19065309.69 | 9510971.642 | 10027711.87 | 16272866.29 | 14563560.62 | 9776857.47 |
| SM(d34:1) | 440080891.8 | 451085815.7 | 453385216.4 | 312291818.4 | 505180535 | 421341776 | 714628756.6 | 531022789.4 | 561127493.3 | 667650491.3 | 547875317.7 | 296416676.5 |
| (ent-16alpha)-16-Kauranol | 298844017 | 320568272.1 | 305734579.8 | 297900547.9 | 316687487.7 | 309810982.6 | 310877079.8 | 306288870.3 | 308106816.8 | 296494859 | 290058365.7 | 283869439.7 |
| M274T141_2 | 101669 | 105641 | 925924 | 958488 | 964122 | 939497 | 932620 | 942479 | 927969 | 931841 | 947063 | 846226 |

| Metabolites | PBS_1 | PBS_2 | PBS_3 | PBS_4 | BCG_1 | BCG_2 | BCG_3 | BCG_4 | BCGΔ*pe18*_1 | BCGΔ*pe18*_2 | BCGΔ*pe18*_3 | BCGΔ*pe18*_4 |
|---|---|---|---|---|---|---|---|---|---|---|---|---|
| | 699.3 | 196 | 04.88 | 48.83 | 75.42 | 56.42 | 13.15 | 47.06 | 00.32 | 16.93 | 11.95 | 74.13 |
| 3,8-dihydroxy-methylanthraquinone carboxylate | 22422422.94 | 13349795.43 | 13068252.33 | 16001865.61 | 15341004.19 | 17832756.75 | 21089326.23 | 16693633.08 | 19288897.82 | 22006474.33 | 11888769.36 | 17103494.24 |
| PE O-26:6_12:0 | 40780953.82 | 54696172 | 45469824.29 | 50203473.2 | 40366072.57 | 34285522.93 | 33952163.5 | 34007791.27 | 38730078.55 | 47829947.91 | 35769364.98 | 24716646.24 |
| Pipericine | 198749433.9 | 191686511.9 | 157495948.3 | 171396856.9 | 228238031.4 | 212813909.9 | 258847759 | 167440751.5 | 178227792 | 258582138.1 | 207098346.5 | 179885174 |
| PG 16:0_20:4 | 3350451.84 | 12324390.27 | 2067831.867 | 2542628.758 | 16275519.23 | 12931127.02 | 33838146.83 | 18854239.37 | 27412147.78 | 25128877.07 | 19876254.91 | 14185722.31 |
| Glu-Phe | 21714.84092 | 27217.88955 | 36927.19616 | 40183.9979 | 1134122.552 | 844407.1678 | 856410.8146 | 664652.9679 | 865925.005 | 804967.0587 | 963691.7277 | 807131.0282 |
| 4-Chloro-6-(1H-imidazol-1-yl)pyrimidine | 199873403.5 | 579169558.1 | 400360199 | 307560294.7 | 371279438.5 | 429080945.3 | 340444668.3 | 322728507.3 | 197696777.6 | 338884256.9 | 423085891.3 | 430162328.1 |
| Lucidenic_acid_M | 219663793.3 | 204986959 | 163807802.4 | 192987062.2 | 214809059.8 | 214364631.8 | 228701332.9 | 195526107 | 212211441.7 | 218462722 | 232437797.5 | 152434615.4 |
| Dimethylethanolamine | 2853377.353 | 4816723.635 | 7604199.604 | 5699006.908 | 5514343.013 | 5333546.229 | 7529544.623 | 6073466.473 | 4994533.819 | 3880503.299 | 8459052.13 | 8554442.735 |
| Cer 12:0;2O/4:0 | 21669192.55 | 21050111.11 | 20813279.92 | 19603288.1 | 21607608.43 | 20875524.47 | 20570019.78 | 21031434.75 | 20353193.64 | 20070494.92 | 20729932.84 | 19349419.53 |
| Daminozide | 530955.6704 | 1591319.645 | 9673538.837 | 2414969.495 | 2319362.773 | 1500900.401 | 2108312.451 | 1164502.995 | 1501857.246 | 1922434.474 | 1802726.815 | 5806217.267 |
| 4-Hydroxy-threonin | 148749 | 118272 | 199259 | 155509 | 125767 | 149142 | 114251 | 119961 | 140654 | 104712 | 123102 | 193210 |

| Metabolites | PBS_1 | PBS_2 | PBS_3 | PBS_4 | BCG_1 | BCG_2 | BCG_3 | BCG_4 | BCGΔ*pe18*_1 | BCGΔ*pe18*_2 | BCGΔ*pe18*_3 | BCGΔ*pe18*_4 |
|---|---|---|---|---|---|---|---|---|---|---|---|---|
| e | 2.419 | 5.659 | 0.843 | 5.391 | 2.124 | 2.884 | 6.218 | 1.919 | 4.112 | 9.488 | 5.874 | 1.55 |
| PE 20:3_20:3 | 7384129.603 | 8816885.872 | 11980152.54 | 10017095.22 | 19909827.8 | 17108050.9 | 17481351.38 | 18930328 | 19398462.04 | 19952267.9 | 16797632.2 | 13293353.07 |
| N-Tetradecanoyl-L-homoserine lactone | 22580737.43 | 22287465.66 | 19431629.7 | 20707055.84 | 19547839.08 | 20597317.88 | 21385992.68 | 20656784.88 | 21070820.2 | 21188327.38 | 20352545.59 | 19090351.3 |
| LDGTS 17:1 | 5758488.129 | 6808551.111 | 6017969.309 | 6288679.578 | 6167528.179 | 6142846.331 | 6150469.867 | 6284044.531 | 6341142.808 | 6226104.689 | 6539358.298 | 6311815.437 |
| SM 9:0;2O/24:5 | 1493556.397 | 1069761.42 | 981819.3728 | 1467382.299 | 1052715.268 | 1326605.799 | 1047237.568 | 1147253.108 | 888342.662 | 885798.1821 | 1182368.314 | 1533473.031 |
| 2-Oxopentanoic acid | 7756239.07 | 5976131.287 | 10382710.82 | 9805191.404 | 11476771.64 | 10872531.72 | 8200368.556 | 9359322.29 | 10634252.75 | 8347824.524 | 10873931.44 | 10778967.4 |
| Ethanesulfonic acid | 39791080.33 | 45238472.21 | 41158979.4 | 43558575.3 | 45114959.8 | 47516591.12 | 86532960.53 | 49962291.55 | 48832317.85 | 42071145.01 | 85271329.67 | 39984832.46 |
| LysoPC(16:1(9Z)) | 1962628.481 | 1554567.505 | 2742492.871 | 1873256.696 | 7523719.456 | 6391796.087 | 2152870.284 | 6484418.948 | 8426288.595 | 6613084.488 | 10235184.98 | 5761260.243 |
| MGDG O-14:0_6:0 | 4920343.388 | 5463580.475 | 4768778.198 | 5165712.732 | 6048753.533 | 5074335.51 | 6024182.421 | 6153282.459 | 5871249.443 | 5541947.569 | 5434066.271 | 5159316.3 |
| 3-Hydroxy-2-iodo-4-methoxybenzaldehyde | 26662064.14 | 28235998.21 | 26479114.63 | 26918292.71 | 28501286.34 | 27374319.78 | 26029394.5 | 26508314.57 | 27990101.33 | 26931300.09 | 26465741.94 | 26826064.38 |
| SPB 16:0;2O | 114943821.6 | 117341960.5 | 105423967.3 | 113073470.7 | 108080612.3 | 116402946.2 | 118337354.7 | 117244398.1 | 110748112.2 | 117808869.7 | 110748317.3 | 104665049.9 |
| Tegaserod | 6026906.129 | 6244585.558 | 6608049.597 | 8027143.045 | 6509636.353 | 7859164.738 | 7136289.785 | 5515167.812 | 6699813.027 | 6555382.165 | 7888380.763 | 5889777.202 |

| **Metabolites** | **PBS_1** | **PBS_2** | **PBS_3** | **PBS_4** | **BCG_1** | **BCG_2** | **BCG_3** | **BCG_4** | **BCGΔ*pe18*_1** | **BCGΔ*pe18*_2** | **BCGΔ*pe18*_3** | **BCGΔ*pe18*_4** |
|---|---|---|---|---|---|---|---|---|---|---|---|---|
| Nicotinate | 3869974.369 | 7174557.91 | 6743822.849 | 8511258.185 | 6627930.801 | 4058247.505 | 2627977.764 | 4788994.375 | 7658888.17 | 1404747.68 | 8889052.335 | 7646665.261 |
| t-Butyldimethylsilyl-Corey lactone aldehyde | 5921013.8 | 6888953.786 | 10905712.97 | 11084466.23 | 10979006.19 | 10963958.59 | 7300482.35 | 6894238.622 | 11174696.36 | 7389382.503 | 10710225.4 | 7324652.642 |
| Kinoprene | 6656203.693 | 8263890.381 | 7070443.682 | 7089148.617 | 7686469.861 | 5054452.632 | 4206425.782 | 5789585.929 | 7336939.279 | 7138493.735 | 5532540.977 | 8575301.673 |
| Cer 12:1;2O/4:0 | 9851651.009 | 10743911.92 | 9856521.006 | 10534802.94 | 9544502.442 | 10040878.55 | 10656692.19 | 10613534.32 | 9982227.021 | 9978130.795 | 10643264.65 | 10047350.38 |
| Salicylic acid | 14860120.86 | 16186832.2 | 14526950.27 | 14175524.59 | 17508748.62 | 17459309.97 | 17832047.98 | 19669297.1 | 17792245.72 | 17793777.36 | 19913052.06 | 17191381.74 |
| PC(20:4(8Z,11Z,14Z,17Z)/14:1(9Z)) | 7173728.098 | 7989759.458 | 4264565.037 | 5706975.501 | 27301136.56 | 21817387.25 | 27585119.43 | 25261495.44 | 29508480.88 | 29683991 | 23278272.94 | 18895438.99 |
| 1H-Indazole-3-carboxamide, N-[1-(aminocarbonyl)-2-methylpropyl]-1-(4-hydroxypentyl)- | 1135629.238 | 1094579.599 | 1570068.746 | 1052026.691 | 847966.3766 | 1153769.558 | 948021.1383 | 844652.1015 | 986333.6449 | 944033.0882 | 1693928.853 | 1036003.482 |
| MGDG O-14:1_22:5 | 6992553.277 | 4621149.099 | 4799352.21 | 4127290.787 | 4973422.657 | 4897959.54 | 4361749.76 | 4897270.914 | 4169013.75 | 4583646.921 | 4643903.221 | 4671894.905 |
| PE-Cer 15:3;2O/26:2;O | 3046176.393 | 2353503.221 | 3235701.621 | 1759058.993 | 2003892.415 | 46252.47303 | 5860293.379 | 1677885.356 | 1397043.088 | 3005831.486 | 4702142.044 | 2503580.06 |
| Panaxynol | 811948 | 806816 | 808357 | 749628 | 875952 | 930876 | 947627 | 861917 | 851519 | 860240 | 811734 | 891304 |

| **Metabolites** | **PBS_1** | **PBS_2** | **PBS_3** | **PBS_4** | **BCG_1** | **BCG_2** | **BCG_3** | **BCG_4** | **BCGΔ*pe18*_1** | **BCGΔ*pe18*_2** | **BCGΔ*pe18*_3** | **BCGΔ*pe18*_4** |
|---|---|---|---|---|---|---|---|---|---|---|---|---|
| | 11.32 | 50.52 | 51.01 | 15.71 | 03.9 | 18.02 | 07.48 | 46.8 | 92.31 | 67 | 10.62 | 73.06 |
| MGDG O-18:5_22:5 | 487475.1199 | 520192.6185 | 560639.2089 | 483715.9892 | 535444.1811 | 559031.9067 | 489812.3318 | 487133.1233 | 532205.0673 | 501791.185 | 539493.3577 | 490312.6059 |
| (2R)-Amino(4-hydroxyphenyl)ethanoic acid | 10912649.76 | 12149446.48 | 10650216.6 | 11026672.96 | 10527869.94 | 10770228.32 | 12367393.02 | 10782900.33 | 10715343.71 | 11147405 | 10748841.61 | 11633647.08 |
| [2-(1-Adamantyl)-1,1-dimethylethyl]amine | 1425228.725 | 1574825.991 | 1320725.755 | 1289631.62 | 1079799.495 | 943233.3221 | 1396065.139 | 869862.8634 | 991915.9181 | 2534330.971 | 1003496.066 | 1555115.122 |
| Hexahydro-6,7-dihydroxy-5-(hydroxymethyl)-3-(2-hydroxyphenyl)-2H-pyrano[2,3-d]oxazol-2-one | 95010820.41 | 76114207.48 | 110887910.4 | 92815887.34 | 577955809.9 | 599055538.3 | 523888556.6 | 529064768.8 | 574964574.5 | 442399024.6 | 529949947.6 | 545030940.8 |
| 4,4,4-Trifluorobutyric acid | 374194251.3 | 364346098.2 | 353321856.1 | 364346504 | 370910913.5 | 359081080.5 | 422901991.4 | 368788045.8 | 370416653.7 | 359409502 | 372150561.6 | 367694802.4 |
| PE(20:3(8Z,11Z,14Z)/18:2(9Z,12Z)) | 220867045.1 | 203088604.1 | 196947508.3 | 206141425.3 | 240215045 | 207722935.2 | 210977027.9 | 235264586.5 | 221931548.2 | 222222287.6 | 211819512.9 | 191432263.7 |
| Zoxazolamine | 937020.255 | 2103797.491 | 2384710.334 | 2239041.849 | 1211219.643 | 925070.6441 | 1260318.36 | 804438.5769 | 993095.0479 | 1706953.379 | 2131448.235 | 3372034.343 |
| PE(22:6(4Z,7Z,10Z,13Z,16Z,19Z)/20:4(8Z,11Z,14Z,17Z)) | 3999348.536 | 3406252.459 | 5638087.672 | 4651838.57 | 8636973.29 | 7801549.216 | 9538862.189 | 9493167.655 | 8483464.853 | 9629692.196 | 8643454.024 | 6887569.975 |

| Metabolites | PBS_1 | PBS_2 | PBS_3 | PBS_4 | BCG_1 | BCG_2 | BCG_3 | BCG_4 | BCGΔ*pe18*_1 | BCGΔ*pe18*_2 | BCGΔ*pe18*_3 | BCGΔ*pe18*_4 |
|---|---|---|---|---|---|---|---|---|---|---|---|---|
| PS(16:0/15:0) | 397250500.7 | 376558505.5 | 335068540.4 | 360648517.2 | 421851678.2 | 380097461.2 | 379683724.9 | 418839710.2 | 375425270.6 | 397830731 | 382590925.4 | 347822501.2 |
| 5-Methylisoxazol-3-amine | 12504533.31 | 13128338.58 | 10744940.01 | 12664748.92 | 12494247.54 | 14416985.28 | 11085156.91 | 12388643.22 | 11893541.39 | 11253150.48 | 12722937.89 | 12356663.01 |
| 6-Aminohexanamide | 1388537.244 | 1488176.586 | 6939867.883 | 1346564.351 | 2650058.973 | 1746835.095 | 9497492.432 | 1775182.778 | 1739330.055 | 4829009.533 | 6667947.878 | 1624331.851 |
| N-Tetradecanoyl-DL-homoserine lactone | 23014989.57 | 20773242.88 | 17370535.78 | 30515491.67 | 20677881.37 | 22451290.9 | 31880594.16 | 19358775.21 | 26114118.37 | 23118197.81 | 19505247.29 | 27561768.71 |
| 3-Heptanone, 1,7-bis(3,4-dihydroxyphenyl)-6-methoxy- | 1148837.981 | 1316361.333 | 1104697.298 | 1215690.289 | 1272116.81 | 1188865.676 | 1147598.492 | 1159987.467 | 1079271.895 | 1261950.14 | 1183482.184 | 1153034.948 |
| MGDG 4:0_22:6 | 18516480.69 | 19238677.43 | 18335160.66 | 18738482.53 | 17554944.03 | 18163356.08 | 18300827.38 | 18067804.08 | 17992448.36 | 19077768.34 | 18160800.22 | 19159326.25 |
| Suberic acid | 2483322.41 | 2871814.09 | 3038729.713 | 2945117.552 | 2467989.223 | 2243646.436 | 3046438.779 | 2458489.194 | 2513330.18 | 2303894.665 | 2450723.663 | 3841676.308 |
| 2-Ethylbutyric acid | 11200798.14 | 11753945.74 | 10540995.71 | 22623580.32 | 12857259.35 | 12596840.34 | 30523389.54 | 10663412.18 | 18247375.38 | 13504266.64 | 11089852.81 | 30914008.55 |
| [2-[docosa-4.7.10.13.16.19-hexaenoyloxy]-3-(octadecanoyloxy)propoxy]({[2.3.4.5.6-pentahydroxy | 546745.1757 | 762839.8257 | 383581.7616 | 549273.5908 | 1599490.67 | 1156061.644 | 1239774.257 | 1491980.772 | 1753971.529 | 1627736.588 | 1159077.5 | 1126450.494 |

| Metabolites | PBS_1 | PBS_2 | PBS_3 | PBS_4 | BCG_1 | BCG_2 | BCG_3 | BCG_4 | BCGΔ*pe18*_1 | BCGΔ*pe18*_2 | BCGΔ*pe18*_3 | BCGΔ*pe18*_4 |
|---|---|---|---|---|---|---|---|---|---|---|---|---|
| cyclohexyl]oxy})phosphinic acid | | | | | | | | | | | | |
| 5'-Methylthioadenosine | 7464657.693 | 7919148.817 | 11046591.15 | 701294.4512 | 54628605.51 | 49851870.56 | 48403178.09 | 47435709.25 | 61242859.98 | 40837368.05 | 48607496.92 | 43116561.36 |
| Nelfinavir | 1096082.003 | 1196122.962 | 1013767.265 | 1193049.057 | 1215069.078 | 997283.8966 | 1067302.15 | 1185716.582 | 1120468.703 | 1137800.841 | 1110331.861 | 1076330.096 |
| PE O-22:2_17:1 | 5966779.777 | 3019979.456 | 5646075.05 | 4721584.203 | 4865856.516 | 1182699.557 | 3454271.134 | 5825413.685 | 4968587.8 | 4233937.409 | 3889486.842 | 3973594.92 |
| 5-Bromo-4-fluoro-2-hydroxybenzoic acid | 27351218.71 | 30541652.64 | 41240935.69 | 35414025.93 | 35093069.25 | 36167521.84 | 39741202.72 | 35806971.67 | 34323328.75 | 33801169.2 | 35942830.06 | 34151193.36 |
| PC(44:10) | 7949753.285 | 10307427.46 | 3821185.672 | 4352808.817 | 5105110.332 | 2782787.735 | 5063947.752 | 5458460.323 | 6360559.84 | 7392768.451 | 4045634.508 | 2459600.287 |
| 7-Bromo-1H-pyrrolo[3,2-d]pyrimidin-4(5H)-one | 64105419.64 | 72404653.96 | 104044578.1 | 62170955.47 | 65287898.34 | 65307438.28 | 55212355.16 | 60382361.53 | 60911892.41 | 65652585.81 | 62427984.78 | 75682971.7 |
| PC(37:4) | 11176947.69 | 14805624.76 | 65297119.07 | 11583606.33 | 128011905 | 109583539.4 | 169041205.4 | 124621082.5 | 132450822.1 | 144665563.2 | 151806506.8 | 68687334.99 |

| Metabolites | PBS_1 | PBS_2 | PBS_3 | PBS_4 | BCG_1 | BCG_2 | BCG_3 | BCG_4 | BCGΔ*pe18*_1 | BCGΔ*pe18*_2 | BCGΔ*pe18*_3 | BCGΔ*pe18*_4 |
|---|---|---|---|---|---|---|---|---|---|---|---|---|
| 6-[(6,8-Dihydroxy-7-methoxy-3-methyl-1-oxo-3,4-dihydroisochromen-4-yl)oxy]-4,8-dihydroxy-7-methoxy-3-methyl-3,4-dihydroisochromen-1-one | 13290423.64 | 12629194.54 | 18983269.44 | 14640030.83 | 14127634.33 | 15220626.38 | 16302545.2 | 13168970.24 | 15869369.78 | 13496117.55 | 13490779.35 | 13181237.17 |
| M258T85 | 43872633.75 | 52647286.41 | 42606156.7 | 50975640.11 | 50480901.27 | 48253245.75 | 46793501.05 | 47766592.98 | 46722983.46 | 44143071.09 | 41912132.05 | 48405634.07 |
| Oligomycin A | 15981230.95 | 19126344.25 | 26403795.62 | 20513201.28 | 38734771.58 | 31216840.58 | 41391563.43 | 37781073.34 | 37764845.96 | 44567231.38 | 40436801.49 | 24006405.73 |
| 3-Methyl-4-nitrophenol | 9627812.08 | 10080542.43 | 8887984.962 | 9383165.456 | 8868439.669 | 8369666.131 | 9093259.935 | 8888131.003 | 8477177.552 | 8637667.189 | 8513562.022 | 9055112.059 |
| Palmitoyl sphingomyelin | 440080891.8 | 451085815.7 | 453385216.4 | 312291818.4 | 505180535 | 421341776 | 714628756.6 | 531022789.4 | 561127493.3 | 667650491.3 | 547875317.7 | 296416676.5 |
| 4-deoxy-threo-hex-4-enopyranosiduronate-alpha-rhamnopyranose 3-sulfate | 3971196.616 | 5436772.325 | 7949137.28 | 3497542.682 | 3588922.749 | 4535570.248 | 2511013.467 | 2528798.38 | 4886000.622 | 3975376.449 | 5537593.983 | 9383479.651 |
| PG(16:0/16:0) | 157152574.9 | 154747123.3 | 137136044.2 | 145683222.7 | 172035585.3 | 149190202.3 | 157536035.8 | 168748298.5 | 159566121 | 157741463.7 | 156881884.5 | 141053870.5 |

| Metabolites | PBS_1 | PBS_2 | PBS_3 | PBS_4 | BCG_1 | BCG_2 | BCG_3 | BCG_4 | BCGΔ*pe18*_1 | BCGΔ*pe18*_2 | BCGΔ*pe18*_3 | BCGΔ*pe18*_4 |
|---|---|---|---|---|---|---|---|---|---|---|---|---|
| N-(4-Aminophenyl)-N-methyl-2-(4-methylpiperazin-1-yl)acetamide | 1679255.841 | 2056226.625 | 1560899.994 | 1728885.631 | 1615930.997 | 1644108.444 | 1536199.139 | 1531261.218 | 1502119.093 | 1682131.822 | 2134046.078 | 1710222.609 |
| SPB 17:0;2O | 23994585.83 | 22803260.29 | 21243681.08 | 22921024.69 | 22019839.72 | 28076074.62 | 20283049.93 | 22788181.72 | 20942147.57 | 25818650.54 | 22423955.36 | 19727318.84 |
| 1-oleoyl-2-myristoyl-sn-glycero-3-phosphocholine | 271060261.3 | 294761398.8 | 198638238.7 | 194362158.1 | 911682519.5 | 655316453.7 | 963868084.5 | 772648636.9 | 913910368.8 | 1032127814 | 804488033 | 449771319.9 |
| trans-Hydroxyperhexiline | 2478041.759 | 3081165.628 | 1135753.252 | 1731030.187 | 1633802.939 | 2128675.9 | 1546907.357 | 2035901.346 | 2129695.389 | 1875832.823 | 1892076.169 | 1261798.434 |
| PC(22:6(4Z,7Z,10Z,13Z,16Z,19Z)/P-18:1(11Z)) | 81408529.47 | 90190227.11 | 32703434.39 | 54247131.17 | 14244817.38 | 11311128.78 | 14670277.51 | 19673484.16 | 18906776.33 | 23655322.46 | 12881522.86 | 15265141.84 |
| 7-ethenyl-1,4a,7-trimethyl-3,4,6,8,8a,9,10,10a-octahydro-2H-phenanthrene-1-carboxylic acid | 208328801.9 | 167873052.5 | 136265694.3 | 159561119.9 | 201638927.2 | 185864452.5 | 180988777.2 | 208917293 | 198800882.8 | 211474939.4 | 214661638.3 | 125361571 |
| 3-O-Benzyl-16-O-tert-butoxycarbonyl-16.alpha.-hydroxyestrone | 26498215.48 | 354431.3917 | 26306639.4 | 9087385.154 | 7763109.41 | 23923404.49 | 12351360.74 | 14289465.85 | 30208364.86 | 9742495.539 | 31431558.77 | 13346225.17 |
| SM 8:0;2O/26:0 | 123834 | 942559. | 969401. | 158673 | 896714. | 8086.68 | 820706. | 490764. | 860252. | 100526 | 830187. | 361471. |

| Metabolites | PBS_1 | PBS_2 | PBS_3 | PBS_4 | BCG_1 | BCG_2 | BCG_3 | BCG_4 | BCGΔ *pe18*_1 | BCGΔ *pe18*_2 | BCGΔ *pe18*_3 | BCGΔ*p e18*_4 |
|---|---|---|---|---|---|---|---|---|---|---|---|---|
| | 7.249 | 3442 | 0539 | 9.329 | 9699 | 1108 | 2217 | 0055 | 2615 | 0.687 | 5354 | 8511 |
| LPC 34:6-SN2 | 293704 41.92 | 296844 26.73 | 269518 15.01 | 290996 55.3 | 240057 80.3 | 306094 45.8 | 303495 46.95 | 216344 66.02 | 223127 41.08 | 308982 41.01 | 229588 19.17 | 261250 55.69 |
| 2,4,6-Trimethylpyri dine | 470156 2.953 | 434200 3.875 | 448387 8.598 | 418372 8.829 | 409994 3.837 | 439090 1.46 | 454488 5.77 | 421443 8.429 | 421969 8.619 | 477751 7.048 | 409382 6.097 | 393628 8.083 |
| MGDG O-16:2_5:0 | 720451 2.106 | 860424 4.819 | 903934 2.091 | 917966 2.594 | 853800 9.374 | 108746 23.13 | 811725 2.954 | 880837 4.103 | 726889 7.321 | 908721 3.355 | 978673 1.459 | 107313 37.23 |
| N-(3-Hydroxyphen yl)-2-(4-isobutylphe nyl)propanamide | 618081 9.704 | 648168 3.581 | 558085 2.586 | 588864 6.128 | 583275 4.844 | 587732 3.398 | 616752 6.806 | 591761 1.9 | 637232 9.215 | 591767 5.733 | 570010 3.415 | 537842 1.066 |
| Dimethylglycine | 241814 47.3 | 237620 78.33 | 234786 36.85 | 232051 57.2 | 253918 26.02 | 237182 45.62 | 260652 54.79 | 239925 51.83 | 250970 55.24 | 228183 49.53 | 259855 13.64 | 251139 94.93 |
| DGDG O-9:0_18:5 | 523847 5.428 | 616416 3.899 | 543376 2.272 | 518417 9.428 | 564430 3.621 | 538780 4.65 | 576262 9.633 | 515024 3.276 | 537288 9.909 | 584210 8.802 | 527661 0.292 | 535231 3.481 |
| PC O-37:2 | 172943 9.335 | 184305 3.989 | 179914 3.333 | 175763 1.516 | 180639 5.057 | 176389 9.897 | 176275 0.045 | 180862 7.363 | 179007 8.756 | 172985 6.881 | 178622 1.346 | 178565 3.373 |
| PC(35:2) | 164229 69.64 | 181608 22.15 | 172496 22.4 | 132603 53.41 | 937465 85.54 | 679778 81.02 | 986331 21.25 | 789817 92.19 | 921792 24.51 | 103811 566 | 738184 57.8 | 498037 46.23 |
| PC O-18:2 | 181810 86.12 | 190451 18.04 | 320195 75.93 | 194266 56.29 | 729295 50.63 | 484631 89.2 | 493531 88.16 | 502170 68.91 | 487428 95.52 | 465946 28.66 | 749045 30.37 | 547172 33.71 |
| 2-Hydroxy-6-methy l-4-(trifluoromethyl )nicotinonitrile | 258322 484.5 | 362424 342 | 338202 691.6 | 252844 705.4 | 367248 759.6 | 344101 252.6 | 354043 075 | 343964 056.1 | 361698 534 | 267921 042.7 | 346788 226.1 | 360023 779.3 |
| Flurandrenolide | 198328 | 225622 | 225069 | 210379 | 213931 | 181194 | 207056 | 200053 | 206386 | 201358 | 207405 | 222749 |

| Metabolites | PBS_1 | PBS_2 | PBS_3 | PBS_4 | BCG_1 | BCG_2 | BCG_3 | BCG_4 | BCGΔ*pe18*_1 | BCGΔ*pe18*_2 | BCGΔ*pe18*_3 | BCGΔ*pe18*_4 |
|---|---|---|---|---|---|---|---|---|---|---|---|---|
| | 66.62 | 10.55 | 22.25 | 63.23 | 56 | 23.73 | 21.75 | 76.55 | 14.72 | 55.86 | 78.45 | 15.72 |
| M545T512 | 4898087.461 | 4036125.436 | 6907503.111 | 4484016.354 | 2620072.058 | 2200559.184 | 3899138.964 | 2721126.393 | 3218814.574 | 3401225.861 | 3939429.491 | 3331287.493 |
| Eicosanedioic acid | 425987.7679 | 320488.6708 | 265561.3525 | 348135.8929 | 237726.8156 | 332695.4659 | 259946.9387 | 351426.7422 | 172220.1011 | 232619.2078 | 201007.7807 | 850129.6724 |
| Terbinafine | 1898225.637 | 1914296.157 | 1705396.818 | 1786770.439 | 1905951.26 | 1679148.504 | 2014263.314 | 1872029.899 | 1739719.351 | 1816306.065 | 1706760.171 | 1742815.444 |
| 9S,12S,13S-trihydroxy-10E-octadecenoic_acid | 4434767.706 | 3734863.683 | 5425387.726 | 4457126.847 | 4920454.899 | 5958621.531 | 4908850.965 | 4846852.559 | 4853004.509 | 4542167.051 | 5370399.696 | 4529724.627 |
| OPHTHALMIC ACID | 2445809.624 | 2019374.559 | 3941809.072 | 2123211.395 | 2808902.388 | 1788761.719 | 4600070.206 | 2471421.949 | 1822539.058 | 3476161.305 | 3696318.904 | 1922050.242 |
| Benzo[1,2-b:4,5-b']difuran-4-ethanamine, 8-bromo-2,3,6,7-tetrahydro- | 2236766.216 | 1525281.544 | 2356940.394 | 1976200.448 | 12570783.01 | 5058839.727 | 10143588.54 | 7987294.57 | 8657185.609 | 9325631.235 | 9936060.966 | 5897871.205 |
| [3-Hydroxy-2-[[tetracosa-5,8,11,14-tetraenoyl]amino]octadecyl] 2-(trimethylazaniumyl)ethyl phosphate | 3119245.537 | 3493602.728 | 3613376.06 | 2859221.758 | 3456295.491 | 2817510.506 | 4353786.884 | 3807382.454 | 3964918.553 | 4360091.88 | 3465024.618 | 1760731.032 |

| Metabolites | PBS_1 | PBS_2 | PBS_3 | PBS_4 | BCG_1 | BCG_2 | BCG_3 | BCG_4 | BCGΔ*pe18*_1 | BCGΔ*pe18*_2 | BCGΔ*pe18*_3 | BCGΔ*pe18*_4 |
|---|---|---|---|---|---|---|---|---|---|---|---|---|
| 2-(3-Hydroxy-5-oxotetrahydrofuran-3-yl)propyl 6-O-((2E)-3-(3,4-dihydroxyphenyl)prop-2-enoyl)-.beta.-D-glucopyranoside | 1240594.606 | 1624534.604 | 1630349.222 | 2407253.325 | 481344.1183 | 338379.5351 | 729693.9645 | 440009.9855 | 1031923.506 | 480550.6598 | 662548.7114 | 3924476.246 |
| LPS 18:0 | 72219758.11 | 72622467.26 | 41560991.61 | 52203355.55 | 26201525.23 | 29085379.75 | 30681052.63 | 24261665.23 | 32261399.16 | 31696503.55 | 32838308.57 | 19085815.88 |
| LPG 20:4 | 40105112.69 | 55709725.86 | 48955192.26 | 14886971.64 | 6376291.17 | 6046391.841 | 11089907.09 | 8732368.248 | 7211666.731 | 13154083.18 | 11807943.17 | 8335580.609 |
| PC(14:0/18:4(6Z,9Z,12Z,15Z)) | 1941510.932 | 2045098.478 | 1543463.543 | 2636087.056 | 16341039.95 | 11750876.99 | 17448697.86 | 14680787.29 | 17496876.49 | 17632287.11 | 13815364.3 | 9483355.448 |
| 4-Chlorophenylalanine | 5380542.476 | 5776538.454 | 6317799.603 | 5490293.597 | 7854313.338 | 7567321.772 | 7098979.201 | 7041305.472 | 7491168.125 | 6911081.854 | 7361792.796 | 7469588.427 |
| NAE 15:1 | 3838941.773 | 4646434.908 | 3181308.383 | 3123409.227 | 3405798.556 | 3383645.977 | 3740740.767 | 3824344.207 | 4104326.591 | 4179069.907 | 3618776.979 | 3361058.768 |
| (12Z)-9,10,11-Trihydroxyoctadec-12-enoic acid | 16376377.9 | 15843012.92 | 19129710.48 | 15594223.55 | 16538802.68 | 19255434.08 | 17807420.27 | 16382713.42 | 16612731.98 | 15572484.16 | 17771140.52 | 16020253.34 |
| 2-Hydroxystearic acid | 49072864.57 | 49992545.52 | 47968819.31 | 52079233.27 | 53954887.04 | 58120844.39 | 49498400.69 | 52837085.08 | 53738533.09 | 50792946.9 | 52904767.16 | 52676388.03 |
| 8-Deoxy-11,13-dihydroxygrosheimin | 32133358.83 | 5468306.288 | 25017283.46 | 5321738.817 | 6014582.826 | 29129317.79 | 12059602.03 | 16016600 | 34990210.36 | 7240752.803 | 39111739.18 | 8479163.961 |

| **Metabolites** | **PBS_1** | **PBS_2** | **PBS_3** | **PBS_4** | **BCG_1** | **BCG_2** | **BCG_3** | **BCG_4** | **BCGΔ*pe18*_1** | **BCGΔ*pe18*_2** | **BCGΔ*pe18*_3** | **BCGΔ*pe18*_4** |
|---|---|---|---|---|---|---|---|---|---|---|---|---|
| PS(20:1(11Z)/20:2(11Z,14Z)) | 19644668.77 | 25870188.55 | 8794105.657 | 15414098.7 | 5450077.087 | 4154521.463 | 4230173.534 | 6143285.517 | 7692139.728 | 6583161.79 | 4287327.219 | 4455298.432 |
| Butyl_3-hydroxy-2-methylidenebutanoate | 1563972.488 | 1562471.306 | 3275208.472 | 4601044.037 | 696436.8911 | 3867392.464 | 12988440.49 | 432237.0106 | 7282759.522 | 3489704.818 | 903966.5994 | 8653842.71 |
| 2,2,6,6-Tetramethyl-4-piperidinyl 2-methylacrylate | 785012289 | 753067899.8 | 752914142.9 | 751509641.9 | 823606240.8 | 899495541.1 | 870373159.1 | 839324865.1 | 815420964.5 | 853139599.1 | 799804892.5 | 787122990.8 |
| 4'-Hydroxyacetophenone | 5009499.178 | 4556363.001 | 5363823.602 | 5104252.86 | 4426366.153 | 5289304.239 | 4599329.201 | 4056302.014 | 4577051.366 | 4923855.535 | 4570113.843 | 5078414.036 |
| (2-aminoethoxy)[2-[octadec-9-enoyloxy]-3-[octadeca-1.11-dien-1-yloxy]propoxy]phosphinic acid | 8333965.101 | 11415152.07 | 13618236.85 | 10941887.22 | 25768310.73 | 16402305.96 | 24459074.29 | 18043413.15 | 21070888.31 | 24369888.02 | 23768999.76 | 11474127.79 |
| Ectoine | 2754273.089 | 2536485.449 | 2715957.751 | 2658951.1 | 2718062.675 | 2867480.201 | 2788360.955 | 2599919.853 | 2578360.862 | 2775342.608 | 2743721.114 | 2594961.106 |
| PS 10:0_24:1 | 4954043.575 | 2336870.779 | 4465810.67 | 3375375.685 | 3525525.658 | 1044291.53 | 2544404.182 | 2225805.616 | 5099253.768 | 4272144.82 | 1947645.258 | 1915759.979 |
| LNAPE 23:0/N-10:0 | 1375108.09 | 1395891.417 | 920669.0451 | 517158.4598 | 64307.98884 | 226814.6893 | 56449.42551 | 140892.4942 | 179827.1504 | 119122.2904 | 265502.0603 | 288479.6866 |
| Austdiol | 1597593.327 | 1617346.827 | 1497026.205 | 1578549.907 | 1516933.643 | 1590298.492 | 1497634.727 | 1580360.064 | 1593850.702 | 1516401.244 | 3088939.398 | 1558311.946 |

| Metabolites | PBS_1 | PBS_2 | PBS_3 | PBS_4 | BCG_1 | BCG_2 | BCG_3 | BCG_4 | BCGΔ*pe18*_1 | BCGΔ*pe18*_2 | BCGΔ*pe18*_3 | BCGΔ*pe18*_4 |
|---|---|---|---|---|---|---|---|---|---|---|---|---|
| PE-Cer 14:3;2O/13:1 | 25260840.45 | 22154378.51 | 26744558.57 | 21842111.42 | 37929094.25 | 33856938.22 | 37511608.34 | 32881224.3 | 35810570.78 | 35273669.71 | 38658772.59 | 21000556.78 |
| alpha-(1,2-Dihydroxyethyl)-1,2,3,4-tetrahydro-7-hydroxy-9-methoxy-3,4-dioxocyclopenta[c][1]benzopyran-6-acetaldehyde | 161047.9779 | 95755.82797 | 483053.28 | 136603.9205 | 426831.9091 | 2589063.442 | 2501657.363 | 3155955.619 | 4760927.771 | 403307.6579 | 7142019.589 | 12716197.14 |
| 4-(4-Iodophenyl)-1,3-thiazol-2-ylamine | 1797736.04 | 1059365.178 | 2783787.187 | 1794843.75 | 18987126.27 | 16880604.81 | 19789889.38 | 18031231.97 | 15365423.77 | 16870897.68 | 20607064.62 | 14811744.59 |
| 7-[3,5-Dihydroxy-2-[3-hydroxyocta-1,5-dien-1-yl]cyclopentyl]hept-5-enoylcarnitine | 1411044.668 | 1452541.943 | 1265014.803 | 1405448.65 | 1577152.461 | 1346719.459 | 1565247.371 | 1431697.96 | 1423113.751 | 1648202.254 | 1444541.315 | 1323481.466 |
| UDP-alpha-galactofuranose | 3345080.66 | 3391471.213 | 3324581.918 | 3312645.847 | 3226709.432 | 3312624.522 | 3433044.919 | 3532142.012 | 3379622.112 | 3228639.057 | 2922090.455 | 3371209.017 |
| Diphenylcyclopropenone | 5258592.559 | 5344602.818 | 4798553.408 | 5366009.88 | 4782070.448 | 4995890.506 | 4935576.422 | 5013437.017 | 5116834.208 | 4783403.503 | 4745879.572 | 4797452.258 |
| Di-N-propylamine | 78450152.76 | 146723552 | 72133817.99 | 111545823.6 | 72590493.55 | 121667839.1 | 73583138.23 | 106675147 | 90700787.83 | 74646421.47 | 135702541.2 | 46041170.76 |
| Mildronate | 17344353.95 | 17948846.41 | 25802879.02 | 15801047.44 | 34299275.95 | 30181019.25 | 38463319.68 | 32330012.18 | 28431703.52 | 30779032.03 | 36046493.67 | 34109218.21 |

| Metabolites | PBS_1 | PBS_2 | PBS_3 | PBS_4 | BCG_1 | BCG_2 | BCG_3 | BCG_4 | BCGΔ*pe18*_1 | BCGΔ*pe18*_2 | BCGΔ*pe18*_3 | BCGΔ*pe18*_4 |
|---|---|---|---|---|---|---|---|---|---|---|---|---|
| THYROTROPIN RELEASING HORMONE | 13631457.85 | 14193627.52 | 12908945.82 | 13631079.92 | 13321878.26 | 12538397.85 | 13680242.41 | 13493050.19 | 12573855.86 | 13020565.64 | 12764176.04 | 12467186.61 |
| PEtOH 19:0_22:4 | 1626791.574 | 1138122.089 | 1746683.448 | 1828610.593 | 1730478.198 | 4781.004994 | 1850458.345 | 2761208.385 | 1628544.173 | 1868999.441 | 1303996.223 | 1642285.905 |
| 2-Isopropyl-5-methylfuran | 3944319.203 | 3574962.572 | 3335809.378 | 24159337.19 | 1373037.367 | 4710159.384 | 36744122.94 | 1047825.301 | 16516218.09 | 7642262.647 | 1386278.318 | 26015525.2 |
| 4-Hydroxyphenylacetaldehyde | 5009499.178 | 4556363.001 | 5363823.602 | 5104252.86 | 4426366.153 | 5289304.239 | 4599329.201 | 4056302.014 | 4577051.366 | 4923855.535 | 4570113.843 | 5078414.036 |
| PC O-14:0 | 12503213.38 | 11317408.12 | 17840202.38 | 12950700.02 | 43106745.31 | 32034135.4 | 45220202.45 | 36386864.69 | 36639342.19 | 37901705.73 | 40727533.78 | 32619979.7 |
| Ethoxyquin | 68445824.69 | 72645366.14 | 58937247.79 | 60302438.7 | 64129422.53 | 78881739.25 | 67300515.71 | 71789299.17 | 72401152.54 | 63907236.4 | 69191863.36 | 63938267.36 |
| Phosphatidylinositols,soya | 17016845.08 | 21058159.97 | 10717620.56 | 15288989.3 | 29584167.79 | 24195725.97 | 24320560.3 | 26662039.02 | 31231359.55 | 26488852.35 | 22764010.8 | 20697919.33 |
| SL 11:1;O/16:4 | 325029.218 | 1740042.017 | 152122.7051 | 2122906.347 | 1298248.127 | 671332.9047 | 356118.3829 | 1751546.032 | 488671.4926 | 525438.838 | 367631.7127 | 2770722.251 |
| NAGlySer 22:4/13:0 | 94326688.74 | 93294024.03 | 138580848.6 | 78084350.81 | 165595540.1 | 144069730.5 | 199677979.8 | 142997980.9 | 153270787.6 | 166084334.7 | 184350479.1 | 94832236.34 |
| 1-(1-Adamantylcarbonyl)-4-ethylpiperazine | 1606224.331 | 1948312.831 | 1319984.043 | 1365874.813 | 1695630.486 | 1432168.515 | 1880334.536 | 1973840.82 | 2021658.904 | 1851644.103 | 1786928.806 | 1530900.422 |
| CAR 15:1 | 3334934.11 | 3128141.003 | 2822335.918 | 3110309.568 | 3571727.157 | 3329362.427 | 3818714.388 | 3236825.836 | 3358260.756 | 3771376.492 | 3458830.835 | 3262779.205 |

| Metabolites | PBS_1 | PBS_2 | PBS_3 | PBS_4 | BCG_1 | BCG_2 | BCG_3 | BCG_4 | BCGΔ*pe18*_1 | BCGΔ*pe18*_2 | BCGΔ*pe18*_3 | BCGΔ*pe18*_4 |
|---|---|---|---|---|---|---|---|---|---|---|---|---|
| 3-Piperidinecarboxamide | 17610232.73 | 72276734.97 | 13891594.77 | 94403905.34 | 209222055.2 | 160578533.1 | 5308335.229 | 8782664.737 | 73057963.73 | 151531727.7 | 123339156.4 | 6978095.693 |
| MG 19:0 | 18490735.42 | 17949121.41 | 14437393.55 | 16723128.7 | 15861560.94 | 17094864.25 | 16771018.8 | 16917899.37 | 17172684.72 | 15084321.73 | 16815049.86 | 14596127.98 |
| PE(20:3(8Z,11Z,14Z)/22:0) | 1939524.91 | 2448366.854 | 1647209.216 | 1954192.78 | 8504351.994 | 7395573.224 | 6907826.287 | 8135508.204 | 11274939.43 | 8249995.753 | 9461255.068 | 5410441.384 |
| (2-aminoethoxy)[2-[docosa-4.7.10.13.16-pentaenoyloxy]-3-(octadecanoyloxy)propoxy]phosphinic acid | 9180487.53 | 8587547.589 | 12419223.64 | 10468614.61 | 24985193.2 | 20836458.76 | 26563941.7 | 24311232.45 | 22200784.92 | 27253838.22 | 24688964.94 | 16288413.38 |
| Platanic acid | 212795512.3 | 7705090.857 | 205895939.8 | 201589836.3 | 224890625.1 | 216793429.4 | 224222421 | 220566409.6 | 204289148.8 | 224177007.8 | 221338117.8 | 195641103.4 |
| 1-Phenyl-2-decanoylamino-3-morpholino-1-propanol | 9835691.041 | 9180400.382 | 8381674.82 | 9081745.568 | 10070531.63 | 9664265.67 | 9983572.797 | 9195642.586 | 9309336.709 | 9601034.74 | 9459543.747 | 9024876.852 |
| Methanone, 1-naphthalenyl(1-pentyl-1H-benzimidazol-2-yl)- | 1140672.36 | 1133473.954 | 1112156.206 | 1079596.029 | 1149471.388 | 1124349.562 | 1127973.123 | 1133141.317 | 1112686.627 | 1140034.122 | 1128394.206 | 1019171.462 |
| licoflavanone | 2684823.725 | 2899178.164 | 2676117.999 | 2869262.802 | 2234657.251 | 2717126.123 | 2489474.499 | 2725772.961 | 2748661.169 | 2534668.853 | 2868548.372 | 2771393.351 |

| Metabolites | PBS_1 | PBS_2 | PBS_3 | PBS_4 | BCG_1 | BCG_2 | BCG_3 | BCG_4 | BCGΔ*pe18*_1 | BCGΔ*pe18*_2 | BCGΔ*pe18*_3 | BCGΔ*pe18*_4 |
|---|---|---|---|---|---|---|---|---|---|---|---|---|
| N,N'-DIACETYL-1,6-DIAMINOHEXANE | 25967144.71 | 32868913.01 | 16431554.98 | 27279598.07 | 30227655.54 | 19557056.68 | 35833081.86 | 32419251.82 | 30653497.29 | 28342112.35 | 25220472.64 | 23565893.92 |
| 4,4'-Oxydibenzohydrazide | 244768.7008 | 227066.4515 | 320010.4681 | 302902.0639 | 795538.8104 | 901471.7229 | 694919.2644 | 672445.5336 | 619748.7942 | 621353.4227 | 626698.9265 | 1015803.937 |
| 5-Propyl-1H-1,2,3,4-tetrazole | 601652.662 | 6714999.371 | 14360596.35 | 11878282.19 | 676066.799 | 238984.0271 | 2537129.704 | 852911.7281 | 1423930.138 | 1404238.48 | 685286.9948 | 9588461.88 |
| Ganoderic acid H | 1431058.244 | 1440873.764 | 1402917.068 | 1419334.878 | 3218325.875 | 5174938.194 | 1881300.842 | 2263680.811 | 2865749.699 | 2353575.258 | 4904553.787 | 3609916.656 |
| DGDG O-18:5_3:0 | 498629.4385 | 568561.6238 | 304372.474 | 510892.4777 | 450788.4471 | 464830.3569 | 430342.516 | 414162.3124 | 486326.0384 | 543222.8734 | 478964.0717 | 380720.5751 |
| 1,3-Dipalmitoyl-sn-glycero-2-phosphoethanolamine | 1531905.385 | 1431722.334 | 2123708.722 | 1536128.366 | 6181608.127 | 5404288.355 | 5176345.476 | 5162734.007 | 6481139.397 | 6880430.261 | 6674976.163 | 4444772.286 |
| 1H-Imidazo[4,5-c]quinoline-1-ethanol,4-amino-2-(ethoxymethyl)-.alpha.,.alpha.-dimethyl- | 635271.446 | 754567.4597 | 734921.1976 | 738397.4695 | 660761.786 | 961368.2773 | 765229.0787 | 764994.4102 | 867260.6143 | 831962.5903 | 589896.7799 | 986035.2509 |
| DGGA 12:0_13:0 | 8806427.299 | 12726715.96 | 7604076.325 | 9635374.346 | 8695518.161 | 11789908.02 | 8841388.468 | 8904144.147 | 12234444.33 | 9247693.378 | 9193869.916 | 11161783.84 |
| LPE O-16:1 | 92720459.09 | 120729122.7 | 63190591.07 | 86853723.43 | 52457132.29 | 48921061.71 | 54677223.12 | 53808868.41 | 64030495 | 58385153.28 | 53172358.37 | 35546894.16 |

| Metabolites | PBS_1 | PBS_2 | PBS_3 | PBS_4 | BCG_1 | BCG_2 | BCG_3 | BCG_4 | BCGΔ*pe18*_1 | BCGΔ*pe18*_2 | BCGΔ*pe18*_3 | BCGΔ*pe18*_4 |
|---|---|---|---|---|---|---|---|---|---|---|---|---|
| 3-Chloro-4-fluoro-5-trifluoromethylaniline | 2620249.035 | 2217919.979 | 130028.7714 | 2005840.003 | 11808242.22 | 9933408.141 | 10946981.08 | 12327452.32 | 11479796.78 | 10111057.02 | 10819026.38 | 10745038.45 |
| PI O-18:1_20:3 | 611357.8455 | 364071.4939 | 531381.5573 | 600486.576 | 385872.4708 | 10545.41923 | 498660.606 | 628460.7465 | 350995.0728 | 468499.1951 | 258076.8417 | 871970.842 |
| Promethazin | 3636720.965 | 3929325.309 | 3151794.249 | 3416565.764 | 3395720.35 | 3292275.071 | 3009929.346 | 3335019.223 | 3219234.963 | 3112612.673 | 3164157.133 | 3324023.034 |
| PC 8:0_16:1 | 14268798.31 | 14832275.12 | 12164629.84 | 11139306.78 | 13863674.37 | 10630817.7 | 10923525.65 | 11596396.27 | 13667994.42 | 10739982.96 | 11166137.45 | 10091746.04 |
| METHYL GALACTOSIDE | 1789317.004 | 2114616.424 | 1900778.965 | 2453578.196 | 1848760.415 | 1685399.685 | 2334380.063 | 1980944.718 | 1460004.016 | 2096236.514 | 2397671.29 | 1728491.477 |
| 2,5-Di-tert-butylhydroquinone | 12950504.02 | 10712642.33 | 12702798.5 | 15365917.65 | 10457822.57 | 16836487.55 | 12686452.65 | 15635680.85 | 14271749.24 | 17136491.93 | 12211331.46 | 13965702.43 |
| PC 9:0_30:4 | 2921659.347 | 1099075.661 | 1465743.804 | 1579502.802 | 812613.0779 | 1657544.837 | 1231480.281 | 1272027.879 | 4146792.394 | 1490582.474 | 1360387.88 | 202092.2799 |
| PE O-19:0_28:0 | 70486.2618 | 38025.88685 | 49624.21536 | 53381.09489 | 246277.5489 | 235514.9136 | 195726.0555 | 180781.9471 | 164278.9928 | 152765.5878 | 301923.0042 | 342019.222 |
| Sulfatide(d18:1/17:0) | 515038.6368 | 520882.1455 | 401332.2426 | 594594.0186 | 1310426.837 | 1283097.316 | 654399.2683 | 1049258.148 | 1260858.569 | 1542505.382 | 1568227.946 | 1303343.87 |
| 1-Allylpiperazine | 51142.3998 | 10694897.54 | 5233177.052 | 10019941.84 | 12764763.69 | 3257320.306 | 5744592.569 | 9196551.777 | 11047623.55 | 2571714.555 | 4855160.854 | 12243120.54 |
| Cer 8:0;3O/16:2;(2OH) | 945657.0549 | 1026648.381 | 817800.4566 | 856078.6326 | 1035834.359 | 881795.4951 | 926249.0334 | 808793.1561 | 803602.2561 | 927998.7122 | 926985.746 | 854609.941 |
| PA 2:0_21:0 | 188397 | 180948 | 174517 | 167489 | 178734 | 220133 | 208803 | 176280 | 203394 | 174820 | 197750 | 161940 |

| Metabolites | PBS_1 | PBS_2 | PBS_3 | PBS_4 | BCG_1 | BCG_2 | BCG_3 | BCG_4 | BCGΔ*pe18*_1 | BCGΔ*pe18*_2 | BCGΔ*pe18*_3 | BCGΔ*pe18*_4 |
|---|---|---|---|---|---|---|---|---|---|---|---|---|
| | 3.802 | 7.021 | 6.984 | 2.363 | 4.642 | 0.609 | 0.076 | 2.075 | 0.82 | 4.987 | 3.402 | 5.144 |
| Cer 9:1;2O/5:0 | 10542691.47 | 12016445.83 | 10766802.07 | 11062043.25 | 11075370.31 | 10699338.38 | 10797948.93 | 10603200.31 | 10892875.48 | 10756122.97 | 11015231.22 | 11066349.68 |
| Capric acid | 33763475.48 | 33357399.83 | 31076570.45 | 30625766.98 | 32308188.94 | 36940314.09 | 33219947.31 | 31996444.91 | 34437937.69 | 33858627.67 | 32861329.54 | 36359862.5 |
| CAI-1 | 6622437.836 | 9627196.366 | 6347255.455 | 5581397.332 | 6761165.857 | 6223839.176 | 6006841.736 | 5463171.771 | 5975884.071 | 6694887.367 | 5798586.695 | 5981320.076 |
| LPI 22:6 | 8835585.68 | 8155224.783 | 7460720.901 | 8341809.995 | 5974627.065 | 5180551.824 | 4562351.75 | 5010585.372 | 7566548.841 | 5341692.569 | 5400622.915 | 5174668.502 |
| SM 40:5;3O | 2115384.604 | 46393886.73 | 983881.3363 | 7584366.351 | 1168400.862 | 335428.4486 | 1182903.533 | 1420765.004 | 3626590.998 | 3646903.093 | 525781.5379 | 2182359.806 |
| 1-Hexadecylpyridinium cation | 14700363.88 | 15352593.14 | 15064313.28 | 14437877.21 | 15298772.72 | 13925819.59 | 17677469.71 | 15032815.15 | 14754334.47 | 16294774.8 | 16257429.6 | 20384941.69 |
| Ethylphosphonic acid | 62956401.62 | 65156293.59 | 66117933.12 | 66315500.62 | 66018684.59 | 64193870.18 | 70089447.85 | 59212815.23 | 58545534.5 | 67583168.82 | 65816687.95 | 62571597.03 |
| Vanillic acid diethylamide | 1782263.763 | 1566849.408 | 2211440.966 | 2024285.165 | 2587861.546 | 2624117.872 | 2263777.072 | 2130749.97 | 2250638.758 | 1897666.796 | 2377638.791 | 2826163.772 |
| 5-ketocaproate | 3737502.271 | 3281309.418 | 3921972.428 | 3890210.469 | 3307583.242 | 3026924.566 | 3016867.142 | 1865337.854 | 2928508.25 | 3300237.114 | 2555864.44 | 2821489.742 |
| LPE O-18:2 | 30456146.54 | 30520663.65 | 28733021.62 | 28866668.73 | 27788922.21 | 35531581.3 | 30974056.51 | 25336943.33 | 26323530.42 | 28146135.92 | 37810049.11 | 19268486.92 |

| Metabolites | PBS_1 | PBS_2 | PBS_3 | PBS_4 | BCG_1 | BCG_2 | BCG_3 | BCG_4 | BCGΔ*pe18*_1 | BCGΔ*pe18*_2 | BCGΔ*pe18*_3 | BCGΔ*pe18*_4 |
|---|---|---|---|---|---|---|---|---|---|---|---|---|
| 7-[2,6-dimethyl-8-(2-methylbutanoyloxy)-1,2,6,7,8,8a-hexahydronaphthalen-1-yl]-3,5-dihydroxyheptanoic acid | 41629521.94 | 37591369.11 | 36834746.88 | 39215347.37 | 38134594.26 | 38733341.79 | 38587173.52 | 38430592.93 | 37916089.65 | 37232473.25 | 38603699.23 | 37844541.88 |
| 5-(5-Methoxycarbonyl-5,8a-dimethyl-2-methylidene-3,4,4a,6,7,8-hexahydro-1H-naphthalen-1-yl)-3-methylpentanoic acid | 918055.2803 | 1041577.609 | 748942.051 | 992647.983 | 609501.565 | 689424.769 | 1264560.909 | 885327.4193 | 986713.7718 | 1195682.598 | 1043050.911 | 1612626.145 |
| 3-(2,6-Dichlorophenyl)-5-(propan-2-yl)-1,2-oxazole-4-carboxylic acid | 12521311.06 | 13133671.11 | 12836424.87 | 16236177.89 | 12477331.34 | 14317624.45 | 15461362.89 | 11909832.09 | 14569069.43 | 13739178.9 | 11108505.64 | 13549524.82 |
| 17-Acetoxygrindelic acid | 17850742.74 | 18410135.24 | 17168001.97 | 18523430.81 | 18060379.86 | 18487742.23 | 18435453.25 | 18495903.31 | 18316948 | 17826980.42 | 17966135.76 | 18001872.16 |
| apo-13-zeta-carotenone | 4396032.187 | 3990067.158 | 2748065.213 | 4546530.17 | 4453798.397 | 4061402.429 | 4923984.018 | 5044445.45 | 4185121.448 | 5229733.501 | 4464062.559 | 3855489.758 |
| PG 19:1_22:6 | 6086937.401 | 6644108.57 | 2352543.667 | 4581894.664 | 4308513.341 | 2142153.132 | 2810813.409 | 3246283.39 | 4366426.295 | 3581718.629 | 2846737.476 | 1654228.266 |

| Metabolites | PBS_1 | PBS_2 | PBS_3 | PBS_4 | BCG_1 | BCG_2 | BCG_3 | BCG_4 | BCGΔ*pe18*_1 | BCGΔ*pe18*_2 | BCGΔ*pe18*_3 | BCGΔ*pe18*_4 |
|---|---|---|---|---|---|---|---|---|---|---|---|---|
| 9S,15S-Dihydroxy-5Z,13E-prostadienoic acid | 6223754.435 | 5461556.586 | 5274306.112 | 7152603.176 | 5326018.485 | 8362239.492 | 8753513.015 | 6521594.409 | 6663070.291 | 6359725.265 | 5022175.528 | 8176263.999 |
| 18-(3-Ethylcycloprop-1-en-1-yl)-14-hydroxyoctadeca-4,7,10,12,16-pentaenoylcarnitine | 1353188.014 | 2858577.091 | 2622058.922 | 846652.5834 | 2967378.734 | 962643.4752 | 2738501.787 | 3036009.873 | 3021754.157 | 1300704.845 | 3173764.235 | 3426547.932 |
| 2-Linoleoyl-1-palmitoyl-sn-glycero-3-phosphoethanolamine | 3307343.161 | 3895914.091 | 4753941.377 | 3468006.163 | 22544189.37 | 16383671.56 | 22868586.55 | 19443408.31 | 22785996.79 | 26067197.29 | 24386757.49 | 13123082.87 |
| PC(39:6) | 5830850.305 | 6415826.904 | 5838481.738 | 5551954.404 | 8127325.786 | 5429156.22 | 8464077.255 | 6211016.02 | 7010391.574 | 7509465.718 | 6615701.526 | 4575134.377 |
| DGDG O-10:0_22:0 | 685805.6542 | 889900.8162 | 527316.8378 | 720101.2449 | 749011.8197 | 958522.9865 | 978757.3508 | 912080.6002 | 742063.9602 | 954231.8609 | 787152.4822 | 803436.559 |
| PG 16:1_20:4 | 3470212.38 | 1957589.419 | 1925062.139 | 1114952.371 | 10367051.68 | 6664513.339 | 13482268.9 | 7965349.044 | 15166218.73 | 13538998.37 | 11898776.69 | 4760808.31 |
| 2-Oxooctadecanoic acid | 9341651.001 | 6772462.871 | 14424987.25 | 8816281.215 | 9661931.034 | 11149750.31 | 7136530.37 | 12038580.53 | 11125451.12 | 5211505.808 | 10238677.31 | 14581381.92 |
| PC(40:6) | 90473729.66 | 105006241.5 | 59919370.28 | 72425867.43 | 139632104.9 | 103436851 | 125480457.4 | 128095204.8 | 150020189.5 | 157213102 | 108875264.2 | 80066688.95 |
| N-(4-Methylphenyl)-N'-(6-methyl-2-py | 546414.1212 | 434350.1592 | 682305.5737 | 543685.1602 | 1761657.512 | 1272642.519 | 1409307.766 | 1266070.728 | 2566826.34 | 1261034.978 | 1675116.155 | 2774989.536 |

| Metabolites | PBS_1 | PBS_2 | PBS_3 | PBS_4 | BCG_1 | BCG_2 | BCG_3 | BCG_4 | BCGΔ *pe18*_1 | BCGΔ *pe18*_2 | BCGΔ *pe18*_3 | BCGΔ*pe18*_4 |
|---|---|---|---|---|---|---|---|---|---|---|---|---|
| ridinyl)thiourea | | | | | | | | | | | | |
| Pehanorm | 11110687.23 | 4880527.288 | 10115462.19 | 9211813.127 | 7301634.93 | 9551264.138 | 4743370.322 | 8595950.184 | 8370560.578 | 8825141.095 | 4034663.001 | 7833857.683 |
| PC O-20:5 | 1207594.72 | 896960.4328 | 1020180.344 | 1021685.85 | 2012384.537 | 1995915.528 | 1959060.291 | 2263572.554 | 1695237.168 | 1343367.999 | 2303133.471 | 2057382.362 |
| SHexCer 12:1;2O/18:0;O | 5069639.827 | 5401865.878 | 4830454.179 | 5412906.979 | 5333112.742 | 5329602.516 | 6083623.942 | 5862443.786 | 5992287.446 | 5164633.153 | 5225757.453 | 5440867.966 |
| Myristic acid | 160372505.1 | 131148594.7 | 92195255.14 | 108107327.1 | 109844256.1 | 151540767.3 | 107300967.2 | 126756146 | 110850148.6 | 99552336.99 | 128953929.1 | 119343003.2 |
| MGDG O-10:0_28:7 | 1735743.514 | 2634365.657 | 1145860.921 | 2545873.713 | 1442206.454 | 2168785.663 | 1806361.264 | 1828857.016 | 1409886.93 | 2146604.361 | 1706144.118 | 2421940.196 |
| DGDG O-8:0_20:5 | 1473189.838 | 1263041.067 | 350038.3904 | 227059.1441 | 2075081.679 | 2005848.54 | 1496544.548 | 1979975.844 | 3145738.421 | 1725302.158 | 1428862.158 | 1105463.545 |
| 2-Isopropyl-N-methylaniline | 548077.1816 | 1685239.198 | 2413883.389 | 2238014.216 | 577821.5979 | 454100.7011 | 717259.2767 | 499119.9005 | 614529.2915 | 454567.9268 | 708821.8918 | 4764507.521 |
| 1-Hexadecylamine | 69873837.99 | 75500505.5 | 43961169.22 | 52663525.25 | 23594808.15 | 20742844.48 | 25753751.38 | 24074867.73 | 25202015.94 | 28551564.25 | 24664042.78 | 17322480.93 |
| PE O-24:5_12:0 | 4931251.158 | 1964202.141 | 3450318.042 | 2130836.353 | 3002907.364 | 163215.7188 | 4102995.018 | 4844389.49 | 3201435.111 | 3587930.697 | 2435798.342 | 2009176.44 |
| DGCC 8:0_10:0 | 16334098.4 | 12347232.21 | 11719196.31 | 12327566.87 | 11604856.07 | 12696001.35 | 17789818.51 | 17906794.19 | 13050791.55 | 12966659.17 | 17766833.81 | 16751262.08 |
| Piperazine-2-carboxamide | 43956259.7 | 12307305.49 | 26278013.99 | 50142175.73 | 49620669.9 | 26244277.48 | 24720779.59 | 46033031.27 | 50875056.44 | 4944864.893 | 21097529.57 | 54606043.26 |

| Metabolites | PBS_1 | PBS_2 | PBS_3 | PBS_4 | BCG_1 | BCG_2 | BCG_3 | BCG_4 | BCGΔ*pe18*_1 | BCGΔ*pe18*_2 | BCGΔ*pe18*_3 | BCGΔ*pe18*_4 |
|---|---|---|---|---|---|---|---|---|---|---|---|---|
| PC(16:1(9Z)/22:6(4Z,7Z,10Z,13Z,16Z,19Z)) | 59288754.75 | 118524770.7 | 41653616.11 | 62894907.07 | 87760792.7 | 71827120.2 | 110655916.1 | 97752719.03 | 107496463.6 | 154115704.2 | 78938108.12 | 65288871.85 |
| CE 24:6 | 34998826.26 | 25863768.74 | 55287732.78 | 53109658.33 | 42278789.13 | 629836.6673 | 24904820.25 | 40494814.57 | 61517682.8 | 38052003.51 | 20066421.87 | 28720481.47 |
| LPE 20:3 | 1303369.834 | 1857948.815 | 1618023.381 | 1329691.909 | 3004500.143 | 3594386.074 | 3023572.937 | 2435935.601 | 2805974.347 | 3651158.77 | 3712873.692 | 2514276.201 |
| (2-aminoethoxy)[3-[hexadec-1-en-1-yloxy]-2-[icosa-5.8.11.14-tetraenoyloxy]propoxy]phosphinic acid | 398721308.9 | 468610714.1 | 412869538.1 | 379211308.7 | 331810121.2 | 243440426.5 | 389756954.2 | 300397697.2 | 295014482.9 | 394899083 | 324363074.3 | 160292242.7 |
| LysoPE(P-16:0/0:0) | 16076870.42 | 15115821.3 | 10350620.22 | 13020345.64 | 8642602.485 | 6263620.627 | 8900911.122 | 8546338.369 | 9171551.688 | 8410907.646 | 8571709.931 | 5315182.446 |
| PG(18:0/18:0) | 25833515.86 | 33158314.94 | 33221052.64 | 31884460.09 | 42417544.03 | 32990078.39 | 44914414.98 | 37828560.8 | 41203180.64 | 45671864.69 | 39785643.22 | 20470117.35 |
| 1-(1Z-Octadecenyl)-2-(4Z,7Z,10Z,13Z,16Z,19Z-docosahexaenoyl)-sn-glycero-3-phosphocholine | 58552467.16 | 54870159.69 | 26477030.83 | 45333322.15 | 15281553.53 | 17813200.52 | 12826099.33 | 16257039.98 | 19413796.92 | 15108589.24 | 13038108.88 | 14417959.78 |
| LPI 20:3 | 6145689.72 | 6267279.883 | 6870700.068 | 8354305.736 | 9114851.337 | 4204662.466 | 12198629.58 | 4874457.55 | 6022065.561 | 13945236.25 | 14674464.32 | 6971075.472 |

| Metabolites | PBS_1 | PBS_2 | PBS_3 | PBS_4 | BCG_1 | BCG_2 | BCG_3 | BCG_4 | BCGΔ*pe18*_1 | BCGΔ*pe18*_2 | BCGΔ*pe18*_3 | BCGΔ*pe18*_4 |
|---|---|---|---|---|---|---|---|---|---|---|---|---|
| NAGlySer 10:0/16:2 | 7517301.594 | 10376049.26 | 5716043.977 | 7277118.726 | 4076907.01 | 5296940.434 | 5754783.484 | 4499950.306 | 4986728.967 | 6235630.086 | 5884807.76 | 2786494.046 |
| TOBRAMYCIN | 702876923.3 | 706707431.2 | 496104785.9 | 473225675.3 | 608748942.3 | 591187610.6 | 586657412.2 | 590748548.5 | 483642836.5 | 619055353 | 581123119.8 | 441931318.2 |
| CerP 19:0;2O/2:0 | 100913828.7 | 114361337.9 | 71164584.12 | 84875702.93 | 57913633.41 | 74215115.78 | 77935866.75 | 58541344.96 | 59761102.91 | 63186105.93 | 73121794.28 | 44474793 |
| 2-[(2-Fluorophenyl)amino]acetohydrazide | 1153325.121 | 1184113.328 | 1128931.754 | 1173155.547 | 1067362.278 | 1148013.379 | 1160501.621 | 1205264.172 | 1157901.5 | 1112844.395 | 1105910.244 | 1099576.975 |
| PC 9:0_24:2 | 3658844.971 | 2502718.644 | 2142218.955 | 1586991.371 | 18092226.45 | 11750446.69 | 10322674.35 | 17276193.34 | 12887370.34 | 11142485.65 | 16648753.33 | 14536523.49 |
| PC(16:0/16:0) | 347850136.8 | 173751485.9 | 235876111.2 | 391682467 | 314807886.7 | 268293272.9 | 254117241.4 | 136266291.6 | 433021301.5 | 135370214.2 | 280659052.8 | 54405183.64 |
| PI(18:1(9Z)/20:4(5Z,8Z,11Z,14Z)) | 4200901.363 | 6136322.647 | 5237258.171 | 4817489.399 | 1510212.647 | 1748032.579 | 2303244.359 | 1775115.234 | 2326228.502 | 3893181.233 | 2560697.291 | 1229196.371 |
| piperettine | 2154983.196 | 2366084.2 | 2105730.062 | 2148659.955 | 2261819.659 | 2338372 | 2186682.548 | 2259712.791 | 2291563.991 | 2417409.832 | 2258332.197 | 2092601.656 |
| PC(37:1) | 3211651.128 | 563657.4086 | 1118210.261 | 918017.3931 | 15592468.52 | 12072332.66 | 15124400.51 | 13354575.5 | 14577999.68 | 16572973.36 | 13009066.06 | 7251187.747 |
| PC 9:0_28:5 | 8230430.725 | 8600943.441 | 3943786.14 | 5569739.323 | 9929572.316 | 11616939.96 | 11047668.97 | 11306990.79 | 11501254.28 | 13191689.19 | 11071296.18 | 8542572.573 |
| 4-Amino-6-chloropyrimidine-5-carbaldehyde | 139534749.1 | 123266223.7 | 147099513.3 | 151719126.3 | 175725225.6 | 126735195.2 | 139800902.2 | 138683003.7 | 147855149.8 | 131667103.7 | 137887067.5 | 111663266.6 |

| **Metabolites** | **PBS_1** | **PBS_2** | **PBS_3** | **PBS_4** | **BCG_1** | **BCG_2** | **BCG_3** | **BCG_4** | **BCGΔ*pe18*_1** | **BCGΔ*pe18*_2** | **BCGΔ*pe18*_3** | **BCGΔ*pe18*_4** |
|---|---|---|---|---|---|---|---|---|---|---|---|---|
| Succinamide | 7948214.061 | 6479073.542 | 8703420.413 | 7344724.187 | 34366030.9 | 28666861.53 | 33640665.62 | 31910723.71 | 31935368.86 | 32773383.33 | 29796033.21 | 27218428.04 |
| LPC O-25:0 | 4891941.638 | 4036794.065 | 2578509.627 | 9131917.423 | 1086371.489 | 4746078.08 | 12354764.42 | 1113983.117 | 9985470.596 | 6939748.5 | 881696.1014 | 9443846.865 |
| Stearic acid | 134587862.8 | 132421529.9 | 139441990.2 | 136147609.8 | 129889005.4 | 180965459.1 | 139757276.8 | 134536168.6 | 120488008.3 | 119071525.8 | 153095240.1 | 151220927 |
| Xanthoxylin | 3010544.167 | 2961774.722 | 3159897.597 | 2934802.952 | 2889312.78 | 2978772.528 | 2884575.356 | 2923885.603 | 2927212.782 | 3068553.413 | 3003445.803 | 2980574.587 |
| MGDG O-16:2_7:0 | 4528924.943 | 5181073.162 | 4389991.771 | 4498576.739 | 4471120.561 | 5051535.038 | 4647844.919 | 5049986.207 | 4706811.608 | 4525096.827 | 4845848.823 | 4486195.474 |
| 1-Amino-1-cyclobutanecarboxylic acid | 32045663.43 | 42685595.35 | 40597409.77 | 45728523.07 | 41480989.94 | 36444852.9 | 34711046.23 | 44559451.83 | 43738711.98 | 47681130.07 | 35286404.09 | 49568487.54 |
| 4-Amino-5-(2-chlorophenyl)-4H-1,2,4-triazole-3-thiol | 3634891.188 | 3787416.898 | 2379025.193 | 3949583.28 | 1925309.865 | 3474087.256 | 3615718.513 | 1756838.17 | 963345.818 | 1719887.373 | 4301786.762 | 3779153.35 |
| 4-(1-Phenylcyclohexyl)phenol | 3145425.244 | 3189888.189 | 3769421.136 | 3365710.169 | 2477111.455 | 3039627.329 | 3197791.452 | 2443473.022 | 2551990.957 | 3031405.317 | 2840455.36 | 3436441.612 |
| 12,13-Epoxy-9,15-octadecadienoic_acid | 5779042.748 | 5611915.827 | 8552991.171 | 6611871.091 | 5572613.114 | 6941679.522 | 5300293.592 | 5934989.721 | 5065790.499 | 4469679.138 | 5988588.344 | 6863463.516 |
| 9-Oxooctadecanoic acid | 21814186.26 | 19338157.91 | 14295680.62 | 16900976.27 | 19546276.73 | 15127058.05 | 17308560.5 | 17442449.13 | 20122943.47 | 16930871.47 | 15776641.01 | 14170033.4 |
| Paprazine | 779303.5851 | 760933.4434 | 776649.8159 | 997117.9886 | 839543.6473 | 859197.3164 | 788116.1963 | 744810.9692 | 745912.2904 | 818760.3052 | 784733.2426 | 640605.6428 |

| Metabolites | PBS_1 | PBS_2 | PBS_3 | PBS_4 | BCG_1 | BCG_2 | BCG_3 | BCG_4 | BCGΔ*pe18*_1 | BCGΔ*pe18*_2 | BCGΔ*pe18*_3 | BCGΔ*pe18*_4 |
|---|---|---|---|---|---|---|---|---|---|---|---|---|
| N-Cyclohexyl-N'-[2-(1H-imidazol-4-yl)ethyl]urea | 10005542.39 | 6378035.397 | 7618983.24 | 7034941.001 | 10485621.5 | 7232221.247 | 7913350.894 | 4883916.4 | 7489653.689 | 8396983.466 | 7641581.203 | 7376203.823 |
| PE 20:2_20:2 | 16042264.86 | 5917750.149 | 13141390.19 | 11863710.1 | 10783947.07 | 875842.8028 | 9328247.819 | 3698328.713 | 18061740.46 | 15936858.33 | 10912757.57 | 9076916.285 |
| 15-Ketoprostaglandin F2.alpha. | 1969050.934 | 1999858.215 | 2604724.426 | 2093094.611 | 2278058.28 | 2534251.55 | 2245298.412 | 2219933.554 | 2150522.033 | 2017358.922 | 2468840.346 | 2276800.266 |
| Palmitoyleicosapentaenoyl phosphatidylcholine | 53366755.31 | 57796910.57 | 28416952.11 | 40912861.16 | 82575100.33 | 90918038.53 | 87789760.77 | 71047149.06 | 90017737.04 | 98087568.17 | 81290990.23 | 43692249.63 |
| 2,4-bis(Ethylamino)-6-diethylamino-1,3,5-triazine | 1301873.471 | 1385991.152 | 1531535.65 | 1311184.93 | 1024658.487 | 1115511.141 | 1276098.347 | 1183933.835 | 1229625.837 | 1253123.496 | 1085105.415 | 1215328.574 |
| 4-Cyanobenzene-1-sulfonamide | 63055825.71 | 71822392.92 | 74339121.13 | 65648919.71 | 66815033.37 | 67566181.54 | 65261146.65 | 69056366.81 | 70407041.15 | 71686698.16 | 68100873.85 | 69767725.85 |
| 1,3,4,6,7,8-Hexahydro-2H-pyrimido[1,2-a]pyrimidine | 11146652.07 | 18637728.88 | 31995067.24 | 27162306.51 | 11523370.62 | 11123090.69 | 13861100.86 | 10503878.45 | 12754282.21 | 13594798.75 | 11766263.16 | 21525759.1 |
| Benztropine (mesylate) | 2719182.77 | 2928088.527 | 2674167.243 | 2581488.137 | 2764298.912 | 3018073.005 | 2603002.289 | 2890571.172 | 3206168.841 | 2664868.392 | 2619141.789 | 2834865.654 |
| Ethyl 3-(N-butylacetamido)propionate | 14187271.08 | 11482327.85 | 11358386.73 | 13651206.73 | 11312405.7 | 14021018.66 | 16285556.74 | 13335092.92 | 13203029.64 | 15053529.95 | 15001624.96 | 19112751.83 |
| PE 18:1_22:6 | 269756 | 288538 | 243829 | 256100 | 712335 | 790590 | 685088 | 830882 | 727604 | 804019 | 771561 | 536847 |

| Metabolites | PBS_1 | PBS_2 | PBS_3 | PBS_4 | BCG_1 | BCG_2 | BCG_3 | BCG_4 | BCGΔ*pe18*_1 | BCGΔ*pe18*_2 | BCGΔ*pe18*_3 | BCGΔ*pe18*_4 |
|---|---|---|---|---|---|---|---|---|---|---|---|---|
| | 2.026 | 6.413 | 4.548 | 2.118 | 7.884 | 7.16 | 5.429 | 2.253 | 7.991 | 0.558 | 9.311 | 6 |
| PE 8:0_24:1 | 771715.6899 | 243393.3351 | 597441.249 | 602240.1865 | 2154459.598 | 3114681.714 | 3533621.231 | 1796636.214 | 3449195.619 | 3278617.516 | 3458327.405 | 2422982.603 |
| 17-Phenyltrinor-13,14-dihydroprostaglandin A2 | 3124506.59 | 3481322.429 | 3241476.043 | 3212042.81 | 3370025.43 | 3420951.395 | 3429051.401 | 3246006.776 | 3457921.798 | 3408124.778 | 3249664.818 | 3380372.849 |
| (2-aminoethoxy)[2-[icosa-5.8.11-trienoyloxy]-3-[octadec-1-en-1-yloxy]propoxy]phosphinic acid | 37873915.6 | 48896421.33 | 52515772.46 | 39030827.83 | 46165407.22 | 32584640.67 | 55454489.26 | 37579917.22 | 39883921.08 | 54052883.2 | 47494718.23 | 12618914.14 |
| Tetraglyme | 9583951.839 | 5490378.879 | 7172643.047 | 5750809.602 | 10532244.47 | 7899053.151 | 8344949.385 | 4650976.953 | 7679503.143 | 8156881.297 | 7667316.278 | 6628437.047 |
| 1-Methylhistamine | 9322040.155 | 8539284.725 | 3080429.506 | 6235627.567 | 5505351.417 | 4189692.733 | 6082232.171 | 6671142.015 | 7894197.634 | 6310922.973 | 4936769.826 | 6238778.604 |
| LPC 16:1 | 3830636.016 | 3611388.596 | 5782893.1 | 4133935.632 | 14238997.49 | 11540707.28 | 15726560.15 | 11371707.91 | 12118192.69 | 12593502.8 | 14830220.92 | 10205598.33 |
| LPC 16:3-SN2 | 2184661.903 | 2215351.375 | 3163204.694 | 2307758.882 | 7778980.963 | 6162498.593 | 7731515.137 | 6941412.559 | 6698193.497 | 6951420.973 | 7746552.799 | 5846529.338 |
| NAGly 10:0/10:0 | 15515447.1 | 17932488.42 | 12633265.46 | 13946917.68 | 12722111.9 | 20254713.83 | 15592759.84 | 13659492.46 | 12278937.47 | 15317521.6 | 15524289.28 | 14450125.96 |
| Calcimycin | 15455520.72 | 18999568.52 | 20550933.34 | 18328942.65 | 17549995.97 | 16678527.46 | 16552475.93 | 15035692.89 | 13651815.1 | 21576077.83 | 20052119.14 | 17503760.32 |

| Metabolites | PBS_1 | PBS_2 | PBS_3 | PBS_4 | BCG_1 | BCG_2 | BCG_3 | BCG_4 | BCGΔ*pe18*_1 | BCGΔ*pe18*_2 | BCGΔ*pe18*_3 | BCGΔ*pe18*_4 |
|---|---|---|---|---|---|---|---|---|---|---|---|---|
| 4,4,6a,9,12b-Pentamethyl-1,11-dioxo-1,3,4,4a,5,6,6a,10,11,12,12a,12b-dodecahydro-2H,9H-benzo[f]pyrano[2,3-b]chromen-3-yl acetate | 5253252.943 | 4323476.04 | 4203488.997 | 4157445.956 | 4612467.857 | 5618489.908 | 5167382.426 | 4906483.169 | 4997302.672 | 5094689.659 | 6437727.892 | 4204278.075 |
| Tapentadol O-sulfate | 45455796.84 | 56198715.95 | 46819533.2 | 53193705.62 | 59515780.91 | 56755171.33 | 57966457.81 | 60455439.98 | 54152101.57 | 52533645.26 | 48717447.3 | 65930400.66 |
| Solasodine | 1098500.843 | 930054.9628 | 1091451.157 | 866392.1084 | 1521251.776 | 875957.4499 | 1416501.121 | 1215969.317 | 1172598.063 | 1201494.661 | 1313072.42 | 806776.6902 |
| PS 9:0_26:2 | 0 | 0 | 0 | 0 | 2422418.107 | 2944618.231 | 2360863.536 | 2022694.937 | 2101164.862 | 2775809.376 | 2553155.296 | 1826018.339 |
| Ambrettolide | 3868102.163 | 2312963.478 | 1530637.412 | 1512595.417 | 1792041.79 | 1699466.984 | 2148393.591 | 1991145.394 | 2237528.782 | 2044755.65 | 2013930.553 | 1963680.177 |
| 3'-N-Acetyl-4'-O-(9-octadecenoyl)fusarochromanone | 3311451.693 | 3910580.114 | 1422421.922 | 2881461.773 | 2705258.183 | 1713813.76 | 1996110.422 | 2213808.032 | 3157079.589 | 3068341.627 | 2040553.306 | 1580154.06 |
| 3,4'-Isopropylidenediphenol | 16764861.03 | 18456883.36 | 16381890.56 | 17652718.61 | 16902291.84 | 16461127.84 | 18305933.94 | 15978210.98 | 16853638.75 | 17577919.56 | 15972459.31 | 15021872.28 |
| Rhizonin_A | 4781826.099 | 5228111.567 | 4097002.478 | 3906863.963 | 9956311.458 | 7223651.908 | 9415197.509 | 8885170.892 | 8566488.482 | 10151872.69 | 9353288.051 | 7232937.547 |
| Drostanolone propionate | 19288211.55 | 23113719.26 | 13969629.4 | 18045651.31 | 3520337.885 | 3577078.134 | 6381383.667 | 4520973.557 | 5740080.974 | 8862016.041 | 6131766.392 | 3959733.108 |

| **Metabolites** | **PBS_1** | **PBS_2** | **PBS_3** | **PBS_4** | **BCG_1** | **BCG_2** | **BCG_3** | **BCG_4** | **BCGΔ*pe18*_1** | **BCGΔ*pe18*_2** | **BCGΔ*pe18*_3** | **BCGΔ*pe18*_4** |
|---|---|---|---|---|---|---|---|---|---|---|---|---|
| Valine betaine | 2652777.35 | 3258491.43 | 4739078.481 | 4264116.803 | 4007608.693 | 2345013.068 | 5381513.993 | 3335360.448 | 2705524.209 | 4695762.675 | 4517228.966 | 5035275.79 |
| PI 8:0_24:1 | 6717818.829 | 7112844.564 | 6376137.427 | 6986227.785 | 7287966.795 | 7518587.912 | 7622044.421 | 6714323.66 | 6955886.662 | 7243240.003 | 7597461.59 | 6273357.845 |
| PC(18:1(9Z)/P-18:0) | 20728057 | 24269161.41 | 15154788.51 | 17509272.73 | 22957794.22 | 24068343.38 | 20246955.76 | 10723252.32 | 14148198.75 | 22163057.28 | 34964478.69 | 24136009.88 |
| .beta.-Glutamic acid | 25833518.42 | 21568315.17 | 34308816.01 | 24867909.33 | 291879393.2 | 212064421.4 | 277265839.3 | 233819224.4 | 242598393 | 210397748.4 | 218839824.7 | 220422568.5 |
| Diphenylamine | 1641149.92 | 2836468.063 | 3875394.689 | 2320921.028 | 2149904.051 | 2280837.658 | 2503640.207 | 3591197.919 | 1154175.931 | 1206125.243 | 3313871.749 | 3656706.995 |
| (2-aminoethoxy)[2-[hexadec-9-enoyloxy]-3-[octadeca-1.11-dien-1-yloxy]propoxy]phosphinic acid | 7308328.859 | 8415946.488 | 8615772.26 | 7742270.672 | 19945609.83 | 12180349.27 | 19171233.4 | 14617185.88 | 18516299.69 | 17954917.56 | 16582228.56 | 8874895.539 |
| 2-Hydroxycinnamic acid | 3020025.158 | 1901113.13 | 3072897.948 | 2809204.703 | 2352956.531 | 2272980.563 | 2313933.566 | 1983547.09 | 2144999.24 | 1759846.436 | 2651770.808 | 3333727.07 |
| .alpha.-Hydroxymetoprolol | 8129586.928 | 8762453.575 | 7414409.864 | 8482367.647 | 8810832.474 | 8703402.007 | 8530803.922 | 8842301.132 | 8716530.001 | 8894319.632 | 8436786.534 | 8054603.629 |
| Adenosine 3'-monophosphate | 4864458.172 | 5909957.639 | 5411059.995 | 5453164.442 | 12818238.29 | 11023349.15 | 11909430.58 | 11662407.13 | 14357871.79 | 12143921.99 | 14366023.66 | 18371797.46 |

| Metabolites | PBS_1 | PBS_2 | PBS_3 | PBS_4 | BCG_1 | BCG_2 | BCG_3 | BCG_4 | BCGΔ*pe18*_1 | BCGΔ*pe18*_2 | BCGΔ*pe18*_3 | BCGΔ*pe18*_4 |
|---|---|---|---|---|---|---|---|---|---|---|---|---|
| 1-(4-(7H-Purin-6-yl)morpholin-2-yl)methanamine | 2029562.515 | 1278133.069 | 1109726.698 | 1198746.681 | 1304558.48 | 1173498.322 | 1203157.298 | 1188761.476 | 1152490.375 | 1105937.489 | 1190404.811 | 1348170.515 |
| N-(4-Hydroxy-2-methyl-6-quinolinyl)acetamide | 10247864.18 | 10400069.67 | 13573669.72 | 6090613.204 | 11387620.07 | 13437976.72 | 9245893.069 | 9107595.441 | 4603996.944 | 9281811.445 | 12668711.3 | 12848907.59 |
| 10-Methylicosanoylcarnitine | 1485091.517 | 1110031.707 | 1038252.692 | 1121903.855 | 1165922.924 | 1812835.448 | 1413088.274 | 1211079.557 | 1381917.48 | 1541339.118 | 1465858.359 | 1090105.651 |
| 4-Hydroxyisophthalic acid | 2775513.156 | 1164220.382 | 3048294.703 | 2955551.082 | 1221839.117 | 2852523.309 | 2736407.497 | 2783435.492 | 1536368.346 | 2683053.705 | 1373497.49 | 3087215.796 |
| Leucyl-Alanine | 8557978.643 | 8735514.921 | 5769423.172 | 7086461.2 | 6496878.825 | 7141646.611 | 11197491.5 | 8577370.045 | 8222378.681 | 7013983.425 | 7630417.605 | 8445459.675 |
| Cer 14:2;2O/13:1;(2OH) | 3421725.428 | 3491307.945 | 2511065.501 | 2990412.06 | 3203906.86 | 2801199.109 | 3189515.205 | 2853609.819 | 3002732.689 | 3098820.085 | 3117197.998 | 2403263.563 |
| LPE 34:5 | 6520035.974 | 5742106.33 | 7415614.918 | 6835461.193 | 5855121.71 | 6676033.889 | 5987581.443 | 6752093.714 | 5620201.805 | 5116775.041 | 6448964.603 | 5614170.779 |
| Doxepin | 1448398.215 | 1612907.81 | 1353194.887 | 1409098.014 | 1519862.818 | 1549764.23 | 1588616.125 | 1623910.493 | 1574438.814 | 1487967.768 | 1444775.955 | 1373007.752 |
| PC(37:6) | 47251217.23 | 53192699.33 | 70913531.82 | 61498821.53 | 155481210.1 | 110060788.3 | 127790093.6 | 122659333.8 | 126847815.8 | 134732768.4 | 119138525.9 | 73948971.65 |
| HEPTANOIC ACID | 7489939.94 | 7816063.107 | 6555259.061 | 7709560.139 | 7905142.551 | 7566869.086 | 8165521.969 | 7116978.386 | 9285089.808 | 6839318.633 | 7619915.405 | 7959035.508 |

| Metabolites | PBS_1 | PBS_2 | PBS_3 | PBS_4 | BCG_1 | BCG_2 | BCG_3 | BCG_4 | BCGΔ*pe18*_1 | BCGΔ*pe18*_2 | BCGΔ*pe18*_3 | BCGΔ*pe18*_4 |
|---|---|---|---|---|---|---|---|---|---|---|---|---|
| N-[5-(2,5-Dioxopyrrolidin-1-yl)pentyl]-N-hydroxyacetamide | 92333.35989 | 83886.59377 | 56566.26175 | 78103.24563 | 17330571.28 | 15403301.58 | 74561.98443 | 47261.69083 | 55374.97847 | 15375508.09 | 67604.30319 | 14879228.54 |
| PI 18:0_18:2 | 9224965.319 | 8277453.259 | 12744579.36 | 9029004.905 | 6251346.596 | 103203.1427 | 6676260.651 | 1575319.198 | 18329557.72 | 6650687.249 | 11773407.8 | 2229109.692 |
| Peonidin_3-sambubioside_5-glucoside | 15077083.96 | 6671438.402 | 17671277.17 | 20569302.27 | 19472785.92 | 15577306.43 | 18153141.83 | 15614126.02 | 14873703.07 | 17199122.08 | 19070674.1 | 9870044.606 |
| PG 20:1_22:6 | 7116355.794 | 3352452.658 | 1408847.378 | 1287766.243 | 1658783.238 | 784709.6738 | 2991214.42 | 1046897.593 | 2487547.135 | 4150888.66 | 1215096.845 | 1244869.447 |
| DGDG 2:0_2:0 | 377175.4269 | 422448.4405 | 383944.6752 | 399743.2639 | 436656.3279 | 393374.4854 | 378979.4859 | 424777.859 | 431365.4325 | 442962.2615 | 408636.7565 | 383416.9827 |
| 2-Isobutyl-1,3-dioxo-2,3-dihydro-1H-Isoindole-5-carboxylic acid | 4924602.922 | 5903849.964 | 6656273.129 | 8496458.111 | 4710210.018 | 5258396.429 | 5490984.449 | 4760477.935 | 5472378.167 | 3099690.476 | 3470711.741 | 7247092.917 |
| Cer 10:1;2O/6:0 | 1372553.84 | 2396968.924 | 2763144.764 | 2759595.292 | 1537654.662 | 1204354.478 | 1556036.063 | 1197771.781 | 1503842.773 | 1386723.334 | 1341507.025 | 9269467.79 |
| GKK 1032B | 1297726.735 | 2033925.09 | 2397628.354 | 1987704.39 | 2183918.208 | 1634741.21 | 2068985.588 | 1465116.423 | 1871978.184 | 2514111.553 | 2630578.01 | 1602262.441 |
| LPA 19:0 | 599010.2269 | 502477.3972 | 750996.2428 | 748982.8964 | 2942539.452 | 2039551.979 | 1812332.506 | 2519418.106 | 3337979.29 | 2666081.91 | 2264365.958 | 1637727.417 |
| PG 20:5_22:6 | 22760962.35 | 22168095.75 | 9235444.993 | 19037374.93 | 9086818.461 | 10389846.15 | 7430637.251 | 11598353 | 19135489.13 | 9532876.775 | 5926277.17 | 11741164.9 |

| Metabolites | PBS_1 | PBS_2 | PBS_3 | PBS_4 | BCG_1 | BCG_2 | BCG_3 | BCG_4 | BCGΔ*pe18*_1 | BCGΔ*pe18*_2 | BCGΔ*pe18*_3 | BCGΔ*pe18*_4 |
|---|---|---|---|---|---|---|---|---|---|---|---|---|
| 9-Methyluric acid | 372561.9955 | 379872.8553 | 898854.5605 | 772233.5161 | 1033037.587 | 750818.6218 | 1002169.225 | 1130502.093 | 694088.6696 | 710536.9394 | 921263.6233 | 1286945.085 |
| 3,3-Dimethyl-2-oxobutanoic acid | 61611634.21 | 62414563.68 | 84336808.65 | 64267082.48 | 34453677.78 | 42090198.06 | 118283889.3 | 37197914.08 | 40921145.46 | 33353558.71 | 28098394.27 | 116480882.5 |
| 2,5-Di-tert-butylaniline | 2236404.23 | 2317994.647 | 2255406.054 | 2332661.275 | 2318488.707 | 2201449.583 | 2650517.29 | 2438658.309 | 2376897.967 | 1850245.527 | 2103882.22 | 2118748.707 |
| 2-[3-[3-Hydroxy-4-(4-hydroxyphenyl)-2,5-dimethoxyphenyl]-5-oxo-2H-furan-2-yl]acetic acid | 4265267.588 | 4677130.5 | 4289021.799 | 4831091.453 | 3501711.66 | 4770168.194 | 3162987.968 | 2878816.414 | 3498442.225 | 2964594.451 | 3447116.252 | 3390317.749 |
| Cer 20:3;2O/42:1;(3OH) | 1157593.532 | 1329502.935 | 1247324.252 | 1172452.308 | 1190363.249 | 1178508.059 | 916498.3498 | 1161495.25 | 1257706.563 | 1032517.645 | 1211077.849 | 821486.4949 |
| PHOSPHONOACETATE | 5233271.213 | 5534933.9 | 5615497.135 | 5356793.579 | 5435356.329 | 5570035.263 | 5036880.38 | 5470646.361 | 5342052.002 | 4822149.006 | 5442257.461 | 5665574.967 |
| Reserpine | 5975800.665 | 7107560.84 | 5460650.695 | 6078449.688 | 6231865.867 | 6059286.463 | 6030744.067 | 6092651.535 | 6033593.559 | 6368803.168 | 5893500.382 | 6120725.678 |
| SM 42:6;3O | 1903276.304 | 1770506.885 | 785610.6367 | 1476272.757 | 368268.7745 | 401191.8398 | 223695.0022 | 340784.4701 | 439576.49 | 347380.0569 | 330501.4107 | 356849.2721 |
| 5-Ethyl-5-(propan-2-yl)imidazolidine-2,4-dione | 2065264.845 | 3909817.982 | 9358359.865 | 11372612.26 | 1415926.543 | 2491348.219 | 4967113.356 | 1758645.871 | 2539369.739 | 4442924.794 | 958834.5572 | 3577568.713 |

| Metabolites | PBS_1 | PBS_2 | PBS_3 | PBS_4 | BCG_1 | BCG_2 | BCG_3 | BCG_4 | BCGΔ*pe18*_1 | BCGΔ*pe18*_2 | BCGΔ*pe18*_3 | BCGΔ*pe18*_4 |
|---|---|---|---|---|---|---|---|---|---|---|---|---|
| PE(20:4(8Z,11Z,14Z,17Z)/20:5(5Z,8Z,11Z,14Z,17Z)) | 3480583.999 | 3687715.501 | 3620958.799 | 3240536.605 | 3517762.847 | 2726400.96 | 3788852.359 | 3367561.06 | 3226978.769 | 5116652.56 | 3773278.34 | 1797969.249 |
| Dihydrocodeine | 7264360.242 | 7351814.689 | 7080605.836 | 6910365.141 | 7076871.329 | 7124695.878 | 7317624.993 | 7401383.908 | 7205849.697 | 7440161.653 | 7137748.854 | 7247324.922 |
| 4-Aminobenzoic acid hydrazide | 4041313.044 | 4212631.557 | 4189547.656 | 4107985.664 | 3991439.451 | 3427807.972 | 715672.1686 | 1215231.086 | 2199504.352 | 2600277.823 | 2298327.367 | 1259739.479 |
| Laurylsulfuric acid | 1491654680 | 1403260756 | 1375999343 | 1592338471 | 1351129394 | 1625477862 | 1299492127 | 1389204244 | 1170507359 | 1224848456 | 1317232070 | 1320294413 |
| N-oleoyl-tryptophan | 2079981.785 | 2105457.596 | 2155353.249 | 1990552.31 | 1990654.991 | 2176725.828 | 1878309.146 | 1924533.486 | 2042302.135 | 2088770.379 | 2069184.086 | 1877047.683 |
| 1-22:2-2-18:3-phosphatidylcholine | 10877018.68 | 17825095.19 | 5237113.902 | 6227293.667 | 7883542.722 | 10566505.51 | 7891606.74 | 5826436.02 | 8395939.53 | 9320825.675 | 9367286.157 | 4247186.346 |
| Ceftriaxone | 7534846.009 | 14992832.67 | 11302615.15 | 43131746.35 | 15379668.53 | 16440879.15 | 9732434.623 | 8809847.534 | 16808760.82 | 11187687.04 | 11039363.89 | 6524121.215 |
| N-Acetylserine | 9071972.888 | 7224853.29 | 9135102.076 | 6931480.586 | 45707559.07 | 33800164.19 | 40867328.62 | 36651189.95 | 38144170.59 | 38097202.7 | 34518953.19 | 33415571.55 |
| Val-Leu | 758736.7932 | 1318381.642 | 906294.4056 | 682375.6509 | 906603.3577 | 584854.1452 | 684160.7563 | 659005.3416 | 796376.7697 | 1142364.472 | 1300824.174 | 658561.7467 |
| 4-Phenylbutanoic acid | 10724728.34 | 12959874.63 | 10312479.28 | 10411009.99 | 11935211.19 | 12694274.71 | 12337900.45 | 12297777.56 | 11455727.33 | 11325005.58 | 11725846.84 | 10193325.45 |
| 9-Methylxanthine | 2956424.132 | 2154728.87 | 4177694.665 | 2954253.022 | 3235422.734 | 3409066.89 | 4860711.009 | 3401118.132 | 4131224.159 | 2905012.882 | 2857548.656 | 3275142.888 |
| PC 15:0_15:0 | 361193. | 143789. | 204148. | 240228. | 297484. | 0 | 247455. | 134552. | 210829. | 394956. | 307385. | 44486.3 |

| **Metabolites** | **PBS_1** | **PBS_2** | **PBS_3** | **PBS_4** | **BCG_1** | **BCG_2** | **BCG_3** | **BCG_4** | **BCGΔ<br>*pe18*_1** | **BCGΔ<br>*pe18*_2** | **BCGΔ<br>*pe18*_3** | **BCGΔ*p*<br>*e18*_4** |
|---|---|---|---|---|---|---|---|---|---|---|---|---|
| | 0755 | 6786 | 5973 | 8509 | 7761 | | 2617 | 4977 | 7896 | 3376 | 062 | 9018 |
| Petunidin_3-arabin<br>oside | 71229.9<br>4544 | 39518.8<br>9613 | 102296.<br>7603 | 46589.0<br>0539 | 557076.<br>0111 | 485914.<br>2748 | 622706.<br>5998 | 483108.<br>9889 | 681680.<br>7774 | 590160.<br>5907 | 600898.<br>8437 | 510728.<br>3169 |
| 13-Oxo-8(14)-podo<br>carpen-18-oic acid | 512102<br>5.202 | 354438<br>9.031 | 293318<br>2.85 | 369542<br>7.795 | 285040<br>2.677 | 316001<br>2.416 | 312539<br>4.61 | 390361<br>0.07 | 292425<br>2.429 | 275928<br>7.765 | 340070<br>3.171 | 335549<br>6.264 |
| LPS 22:0 | 147774<br>19.29 | 152448<br>33.24 | 231839<br>16.58 | 147754<br>59.87 | 384573<br>71.04 | 346708<br>41.7 | 431383<br>69.15 | 337605<br>89.68 | 305952<br>04.71 | 340155<br>34.3 | 386402<br>28.48 | 272917<br>96.47 |
| Hexamethylenetetra<br>mine | 633497<br>7.017 | 603971<br>2.166 | 725947<br>9.498 | 827358<br>1.079 | 642503<br>3.392 | 629953<br>4.87 | 728538<br>2.073 | 690434<br>9.887 | 669124<br>3.738 | 694030<br>6.455 | 687128<br>5.248 | 651638<br>7.384 |
| 9-Benzyl-5-[(furan-<br>2-ylmethyl)amino]-<br>7,8,9,10-tetrahydro[<br>1,2,4]triazolo[3,4-a]<br>[2,7]naphthyridine-<br>6-carbonitrile | 569252<br>8.062 | 536517<br>8.854 | 438104<br>9.662 | 473591<br>3.29 | 435833<br>8.294 | 425621<br>3.752 | 447649<br>6.462 | 553418<br>4.309 | 450609<br>4.074 | 498907<br>2.496 | 442124<br>9.148 | 799736<br>5.545 |
| 3-Chloro-2-hydroxy<br>-5-phenylbenzoic<br>acid | 131417<br>0.454 | 895719.<br>2658 | 136861<br>6.229 | 117217<br>9.452 | 304671<br>7.928 | 281523<br>4.156 | 313869<br>1.43 | 301546<br>3.76 | 287508<br>4.041 | 244545<br>4.701 | 254674<br>8.691 | 306239<br>0.976 |
| PE(18:0/18:2) | 149909<br>33.55 | 174837<br>28.48 | 288287<br>37.42 | 196502<br>34.04 | 809854<br>10.22 | 573287<br>22.72 | 817527<br>30.51 | 629908<br>54.61 | 721749<br>41.46 | 869490<br>06.11 | 815300<br>58.62 | 418075<br>06.9 |
| Pyridoxal (Vitamin<br>B6) | 434567<br>4.21 | 441552<br>9.631 | 540540<br>0.085 | 491695<br>3.896 | 596981<br>5.181 | 539437<br>8.273 | 438265<br>0.662 | 453561<br>8.079 | 634360<br>5.055 | 491802<br>2.983 | 428090<br>0.99 | 830121<br>1.257 |
| Rhein | 173873<br>27.68 | 147075<br>56 | 369325<br>43.93 | 179745<br>92.1 | 130077<br>76.34 | 156686<br>68.02 | 770542<br>9.338 | 163257<br>20.18 | 144653<br>74.53 | 115684<br>14.69 | 106541<br>91.72 | 970859<br>5.456 |

| Metabolites | PBS_1 | PBS_2 | PBS_3 | PBS_4 | BCG_1 | BCG_2 | BCG_3 | BCG_4 | BCGΔ*pe18*_1 | BCGΔ*pe18*_2 | BCGΔ*pe18*_3 | BCGΔ*pe18*_4 |
|---|---|---|---|---|---|---|---|---|---|---|---|---|
| Chlorothalonil-4-hydroxy | 10196769.71 | 7734821.174 | 7259842.866 | 8168776.243 | 6274005.009 | 7024979.854 | 12021676.46 | 9209090.502 | 10872730.05 | 6942415.386 | 5961660.59 | 8182496.488 |
| Caprylic acid | 76924997.5 | 79054326.54 | 74556549.09 | 78961041.58 | 77172446.79 | 75854224.77 | 76616867.53 | 110190302.2 | 77181871.53 | 76897030.24 | 75785682.16 | 81791063.03 |
| N-Dodecanoyl-L-homoserine lactone | 1989559.825 | 2147676.875 | 1707822.868 | 1914772.068 | 2080000.923 | 1985576.904 | 2027008.259 | 1894770.489 | 1988444.546 | 2020352.082 | 1917042.031 | 1956100.345 |
| Caftaric acid | 350040.6578 | 254572.4244 | 293957.5967 | 432521.7999 | 2543327.671 | 2404307.145 | 1803993.059 | 2208920.713 | 4447900.968 | 3687760.159 | 3343239.42 | 4602474.813 |
| 1-(2-Hydroxyethyl)-2,2,6,6-tetramethyl-4-piperidinol | 59471598.02 | 35272206.47 | 35017654.66 | 85686627.61 | 34900659.38 | 30261386.88 | 27552405.11 | 21889426.23 | 26461526.17 | 28592490.42 | 46687434.83 | 16137025.63 |
| PEtOH 16:1_20:5 | 14514916.82 | 17634085.2 | 14190520.14 | 15101873.12 | 15114958.19 | 15400461.71 | 15773840.99 | 14786876.31 | 14481317.47 | 14907399.38 | 16265528.62 | 13770331.32 |
| (4-Oxo-1,2,3-benzotriazin-3(4H)-yl)acetic acid | 610799.6915 | 516719.8375 | 603952.3945 | 686543.6548 | 3036164.49 | 1719484.251 | 2931630.251 | 1790101.118 | 5525031.844 | 2226161.182 | 1973303.298 | 1192774.772 |
| 1-(5-Methyl-3-pyridinyl)-1-decanone | 1332275.973 | 1596206.952 | 2833277.261 | 1498048.554 | 1227203.355 | 1499069.32 | 196333.22 | 1222897.319 | 1250146.535 | 480568.7313 | 1524825.694 | 13801268.5 |
| Spisulosine | 7522692.195 | 9050130.241 | 5124920.033 | 7239099.313 | 5891177.291 | 5867338.657 | 5835432.99 | 6313277.517 | 7014512.958 | 6743457.118 | 5809723.135 | 5256222.749 |
| M345T426 | 21026017.73 | 20892582.43 | 22518958.03 | 22081072.34 | 20418392.06 | 19935045.35 | 17286064.11 | 18478322.52 | 17682644.73 | 16563982.8 | 23895863.2 | 10448169.13 |
| LPE 22:5 | 9249083.095 | 9172690.297 | 12870465.92 | 11393042.18 | 18919739.19 | 17932108.84 | 13228007.67 | 23506765.05 | 16230771.69 | 16732930.5 | 15825356.16 | 19555308.22 |

| Metabolites | PBS_1 | PBS_2 | PBS_3 | PBS_4 | BCG_1 | BCG_2 | BCG_3 | BCG_4 | BCGΔ*pe18*_1 | BCGΔ*pe18*_2 | BCGΔ*pe18*_3 | BCGΔ*pe18*_4 |
|---|---|---|---|---|---|---|---|---|---|---|---|---|
| PC(34:2) | 7021822.595 | 9697394.488 | 6476992.761 | 5372097.904 | 10787964.13 | 39099218.43 | 47354737.29 | 41990433.81 | 45049977.41 | 47125802.57 | 7430493.747 | 27033329.88 |
| PC(16:0/16:1(9Z)) | 2825407.594 | 6455794.278 | 5295273.208 | 4327772.989 | 15905591 | 16425999.51 | 21132978.87 | 10676397.25 | 9834453.146 | 37869499.2 | 15687292.99 | 4766774.204 |
| 1-Hexylpiperazine | 1009186.843 | 1576031.822 | 1785267.941 | 1574876.678 | 1351367.283 | 1370127.069 | 1276336.759 | 2017034.502 | 1532140.406 | 1737022.052 | 1477816.827 | 1468995.127 |
| N-Cyclohexyl-N-propylamine | 784724.2298 | 3171383.728 | 4865344.66 | 4943135 | 953934.5375 | 589469.5967 | 1100236.647 | 587511.2991 | 1199074.104 | 1733969.511 | 881353.6994 | 10299793.99 |
| Ecgonine ethyl ester | 8113968.666 | 7666618.441 | 6655629.483 | 6896041.574 | 7554104.525 | 6775306.429 | 7921257.804 | 8206302.056 | 7608185.593 | 7595909.185 | 6838469.658 | 6442237.651 |
| 3-Hydroxy-9,10-Dimethoxypterocarpan | 337718.2916 | 287092.1392 | 483266.4702 | 254356.6675 | 2542389.584 | 2793506.405 | 2507007.038 | 2604068.943 | 2502011.959 | 1984997.668 | 2449669.917 | 2574140.789 |
| PC 9:0_30:5 | 2908181.654 | 2825656.88 | 2479252.81 | 2706323.283 | 3945204.967 | 2758025.399 | 2612444.681 | 2230212.855 | 3023335.319 | 2954417.002 | 2477238.443 | 3234441.133 |
| PE 18:0_22:4 | 1618098.32 | 491832.474 | 1600023.068 | 1090756.614 | 1404427.678 | 5803.957527 | 1292322.2 | 696702.2085 | 1917197.528 | 1864663.488 | 1440167.503 | 1421065.871 |
| Squalene | 9126790.013 | 14532251.01 | 12119902.65 | 15720712.67 | 9743514.949 | 10444258.15 | 47675568.19 | 9158824.424 | 21819933.35 | 16841228.72 | 8093542.205 | 22684372.05 |
| NAGly 22:6/22:5 | 6668998.068 | 7001526.905 | 4081820.052 | 4728763.548 | 37192221.27 | 32220022.05 | 33916392.98 | 35877218.92 | 29358439.33 | 42398574.77 | 29635935.19 | 24523088.42 |
| Cer 17:0;2O/17:1 | 12229724.39 | 8270130.85 | 9237206.97 | 11004499.99 | 10997796.9 | 508608.2527 | 11228584.42 | 14615110.48 | 13069739.49 | 11665387.23 | 10141507.65 | 13426394.61 |
| Phe-Ile | 791385 | 867883 | 665109 | 706710 | 231926. | 753550 | 738081 | 777068 | 774638 | 774884 | 764707 | 641044 |

| **Metabolites** | **PBS_1** | **PBS_2** | **PBS_3** | **PBS_4** | **BCG_1** | **BCG_2** | **BCG_3** | **BCG_4** | **BCGΔ *pe18*_1** | **BCGΔ *pe18*_2** | **BCGΔ *pe18*_3** | **BCGΔ*pe18*_4** |
|---|---|---|---|---|---|---|---|---|---|---|---|---|
| | 0.35 | 3.827 | 6.109 | 1.388 | 8714 | 1.491 | 7.047 | 8.485 | 4.528 | 3.722 | 2.982 | 0.73 |
| 2,2',2'',2'''-(1,2-Phenylenebis(azanetriyl))tetraethanol | 27681287.15 | 30350676.55 | 27179184.1 | 28572754.58 | 28163542.49 | 26784363.26 | 30150231.62 | 28811789.33 | 28203521.92 | 27321554.73 | 27273177.72 | 26958516.69 |
| PE(P-18:1(11Z)/22:4(7Z,10Z,13Z,16Z)) | 27610690.83 | 27960994.38 | 24434189.63 | 24193714.64 | 30121650.54 | 25339052.53 | 40349952.85 | 29365336.5 | 28065814.47 | 34469751.59 | 28212004.1 | 19423982.94 |
| Cuminaldehyde | 2653650.931 | 2722416.951 | 2469336.903 | 1679266.485 | 2534676.747 | 3015932.919 | 2546758.125 | 2704704.938 | 1964715.668 | 2796804.366 | 2299430.727 | 2297296.598 |
| PC(18:4(6Z,9Z,12Z,15Z)/P-16:0) | 11720604.44 | 12179245.93 | 11821753.66 | 12598708.93 | 7765284.995 | 8110383.262 | 11227702.23 | 7730019.058 | 6835134.075 | 9978207.365 | 7807257.475 | 5952785.72 |
| PG 21:2_22:6 | 1774252.317 | 1042405.947 | 400243.2951 | 757462.6728 | 430944.4231 | 208500.7861 | 522980.9347 | 397644.8751 | 519681.9989 | 684019.4144 | 336271.9871 | 286197.5215 |
| 10E,12Z-octadecadienoic acid | 18516113.07 | 15374121.85 | 21246909.28 | 27933988.61 | 15882859.82 | 17595765.91 | 10778798.75 | 30332486.06 | 12070313.02 | 14195407.42 | 16048292.07 | 25637875.77 |
| Cyanofenphos | 126884.9022 | 113436.4588 | 140168.7462 | 124312.9137 | 855986.0801 | 741297.6586 | 1086988.705 | 1241140.458 | 659699.0601 | 1469849.24 | 901990.5011 | 649481.3709 |
| Benzyldimethyltetradecylammonium cation | 19518155.99 | 19746731.24 | 11223186.53 | 17474634.85 | 19012815.7 | 16623250.32 | 17714887.08 | 17197105.99 | 15496862.29 | 17882019.04 | 17144786.79 | 17519628.12 |
| 4-Guanidinobutanamide | 4820727.012 | 2829041.888 | 2431851.917 | 5186777.664 | 4762221.141 | 5048489.329 | 3688059.033 | 5306537.619 | 5722009.183 | 3200551.486 | 2823574.71 | 5704214.978 |

| Metabolites | PBS_1 | PBS_2 | PBS_3 | PBS_4 | BCG_1 | BCG_2 | BCG_3 | BCG_4 | BCGΔ*pe18*_1 | BCGΔ*pe18*_2 | BCGΔ*pe18*_3 | BCGΔ*pe18*_4 |
|---|---|---|---|---|---|---|---|---|---|---|---|---|
| N-(3-PHENYLPROPIONYL)GLYCINE | 1700956.278 | 1971247.767 | 1609478.856 | 1547348.693 | 1724105.654 | 1660284.032 | 1576234.025 | 1702500.632 | 1613813.349 | 1609407.06 | 1616754.896 | 1435302.684 |
| Carbamic acid, [(3,4-dichlorophenyl)methyl]-, (1S)-2,3-dihydro-1H-inden-1-yl ester | 82218346.17 | 71332561.29 | 109985416.4 | 77400646.57 | 93389693.31 | 87588073.87 | 71061602.69 | 97981271.89 | 67375276.62 | 63976696.83 | 78303185.47 | 91861901.96 |
| PA 21:0_26:4 | 344955.3024 | 327182.7174 | 323905.328 | 333812.3803 | 343051.1666 | 328156.923 | 348526.0977 | 335233.9644 | 312400.35 | 340571.511 | 339040.7059 | 312560.6712 |
| Anabasine Hydrochloride | 26805912.64 | 27767174.2 | 26319250.27 | 21405091.36 | 26624428.93 | 29607433.01 | 27529761.08 | 26199430.01 | 23532189.41 | 27073465.81 | 26600043.87 | 20293679.73 |
| beta-Solamarine | 4351020.126 | 4540054.132 | 2852222.101 | 3901345.357 | 1844954.476 | 1690468.634 | 2232367.773 | 2496293.335 | 2533367.657 | 2231765.114 | 1498269.873 | 1657096.512 |
| Citral_propylene_glycol_acetal | 10603982.68 | 11890520.63 | 11067916.07 | 12395561.98 | 12232194.22 | 10119964.14 | 10555690.39 | 9650643.195 | 9845718.204 | 10923350.73 | 10291878.17 | 10750706.78 |
| Sch 210972 | 6552917.534 | 2031181.67 | 5591492.072 | 2268876.356 | 2218704.544 | 6048179.475 | 3907680.243 | 4577071.827 | 6566653.687 | 2589610.973 | 6879247.684 | 2903511.731 |
| PS 37:0 | 24294.58072 | 0 | 0 | 0 | 13818870.57 | 11914298.7 | 13621850.59 | 11340436.95 | 10784476.65 | 13089699.2 | 14574021.57 | 10084737.97 |
| Ritalinic acid | 6460306.085 | 6272060.903 | 5654329.2 | 6675177.409 | 6417142.836 | 6325743.583 | 6325364.005 | 6672987.006 | 6160645.571 | 6664524.459 | 6857118.943 | 6432898.328 |

| **Metabolites** | **PBS_1** | **PBS_2** | **PBS_3** | **PBS_4** | **BCG_1** | **BCG_2** | **BCG_3** | **BCG_4** | **BCGΔ*pe18*_1** | **BCGΔ*pe18*_2** | **BCGΔ*pe18*_3** | **BCGΔ*pe18*_4** |
|---|---|---|---|---|---|---|---|---|---|---|---|---|
| 4-Amino-N-(2,4-difluorophenyl)benzenesulfonamide | 85072499.62 | 54598159.36 | 45059097.91 | 85421427.32 | 76664131.15 | 76255962.45 | 73085448.28 | 70972524.85 | 74083769.25 | 69419118.1 | 75004468.43 | 82515594.75 |
| Methanone, (4-methyl-1-naphthalenyl)[1-[2-(4-morpholinyl)ethyl]-1H-indol-3-yl]- | 236945.8937 | 202562.3112 | 316400.3961 | 222068.3576 | 570326.0717 | 391371.2399 | 628921.446 | 501783.0375 | 675858.6987 | 764760.8202 | 761851.8839 | 508787.4247 |
| PC 8:0_32:9 | 3325721.876 | 3560996.869 | 1202201.516 | 2567347.784 | 1357647.585 | 1599737.401 | 1441296.127 | 1982450.894 | 1965135.552 | 1891812.769 | 1167034.526 | 1128349.84 |
| Cer 12:1;2O/12:0 | 5762595.385 | 7784050.742 | 3393732.756 | 4453315.308 | 4247355.64 | 5739966.71 | 5828176.026 | 8473954.493 | 7104427.766 | 5072271.678 | 4223642.916 | 5319591.056 |
| 5-(2-Furyl)-4H-1,2,4-triazol-3-amine | 4990168.729 | 5815076.39 | 6075804.019 | 5706705.87 | 7528678.976 | 7283775.488 | 7444407.03 | 7210710.726 | 9364670.974 | 6582148.275 | 6075609.332 | 6907050.073 |
| 2,3-Diethylpyrazine | 50299609.94 | 7945599.294 | 53660640.43 | 21498546.21 | 46852881.42 | 50345402.47 | 51563191.17 | 49644578.51 | 20318635.09 | 49873650.13 | 50153155.37 | 49666978.15 |
| PC(14:0/14:0) | 3436495.252 | 3587394.347 | 2696597.194 | 2708513.7 | 38641863.95 | 25385802.78 | 42640162.29 | 32896122.84 | 34699289.78 | 41813635.14 | 33278357.71 | 17069385.86 |
| 1-Phenyl-3-(trifluoromethyl)-1H-pyrazol-5-ol | 1502549.589 | 1550831.134 | 1368872.65 | 1526536.121 | 1313655.938 | 1403291.682 | 1360300.213 | 1452145.26 | 1376120.081 | 1379866.965 | 1377690.052 | 1311499.767 |
| PS(18:0/18:2) | 11875397.28 | 6672292.543 | 12458700.34 | 13354760.05 | 9061007.155 | 1409228.482 | 7788422.694 | 13755603.42 | 12296254.51 | 10119624.66 | 8921168.709 | 9709867.347 |
| 2-Furyl methyl | 119846 | 113104 | 139190 | 113231 | 108749 | 121012 | 118149 | 119822 | 112125 | 115430 | 119527 | 117068 |

| Metabolites | PBS_1 | PBS_2 | PBS_3 | PBS_4 | BCG_1 | BCG_2 | BCG_3 | BCG_4 | BCGΔ*pe18*_1 | BCGΔ*pe18*_2 | BCGΔ*pe18*_3 | BCGΔ*pe18*_4 |
|---|---|---|---|---|---|---|---|---|---|---|---|---|
| ketone | 6.597 | 3.35 | 0.119 | 7.277 | 6.531 | 6.28 | 9.62 | 2.829 | 3.542 | 6.345 | 3.469 | 4.032 |
| 4-(2,4-Dichlorophenyl)-1,3-thiazole-2-carboxylic acid | 3865484.859 | 6562336.728 | 6170680.885 | 6676525.859 | 5530661.451 | 3553901.991 | 3374541.733 | 3273822.489 | 3458591.147 | 1258768.447 | 5644294.258 | 3395460.533 |
| MG 13:0 | 6287698.478 | 6876418.215 | 5880711.115 | 6157332.702 | 6893340.919 | 6605093.606 | 6747528.112 | 6889659.375 | 6586191.333 | 6846073.272 | 6678917.992 | 6117836.024 |
| 10-(Acetyloxy)decyl acetate | 217470.2902 | 819822.6711 | 1286424.387 | 1054691.39 | 2565172.595 | 355869.333 | 2543080.555 | 243388.205 | 315706.3203 | 293563.4134 | 1614823.895 | 3310249.324 |
| PE(22:6(4Z,7Z,10Z,13Z,16Z,19Z)/20:3(8Z,11Z,14Z)) | 7315001.278 | 8196750.63 | 10687931.1 | 8940586.602 | 20243094.71 | 16331394.09 | 17806372.75 | 18306706.45 | 19218170.26 | 19881353.47 | 17990956.71 | 14477728.13 |
| Betazole | 5151495.501 | 6264050.227 | 10364735.89 | 7808811.566 | 5896672.279 | 4084379.053 | 7113708.278 | 6644518.708 | 6171170.678 | 6191910.053 | 3803073.869 | 14291113.64 |
| 3-Methylbenzoic acid | 4279321.074 | 4132279.539 | 4066495.789 | 4236505.462 | 4183719.433 | 4280401.266 | 4059330.917 | 4014509.357 | 3949747.523 | 4322890.347 | 3899046.431 | 4347539.039 |
| 3,4-Dimethyl-2,6-dinitrophenol | 7022212.806 | 7366367.353 | 6404820.75 | 7275920.976 | 8056285.595 | 7669512.133 | 7835502.31 | 7770106.703 | 6999148.851 | 7404991.822 | 7041523.324 | 5494150.288 |
| Dioctylphosphinic acid | 22265022.88 | 27960262.02 | 18326321.01 | 19705398.89 | 23498248.64 | 20673266.49 | 26076614.41 | 30922127.55 | 27889931 | 24205391.87 | 23036350.62 | 21558787.77 |
| Hexacyclinic acid | 522360.3805 | 853109.7813 | 555351.2314 | 422621.7934 | 450514.6734 | 737605.6507 | 472309.3215 | 499831.9952 | 1806345.287 | 476595.9222 | 721009.8187 | 780190.2571 |
| SL 22:2;O/14:0;O | 15969124.08 | 20118634.6 | 15866798.18 | 19772565.67 | 16688998.7 | 18193841.49 | 17020551.47 | 14894083.73 | 15802066.23 | 14999455.54 | 15152380.4 | 15785633.54 |

| Metabolites | PBS_1 | PBS_2 | PBS_3 | PBS_4 | BCG_1 | BCG_2 | BCG_3 | BCG_4 | BCGΔ*pe18*_1 | BCGΔ*pe18*_2 | BCGΔ*pe18*_3 | BCGΔ*pe18*_4 |
|---|---|---|---|---|---|---|---|---|---|---|---|---|
| 3,6-Ditert-butyl-1,2-benzenediol | 11457462.94 | 10859937.22 | 10467350.24 | 14613165.46 | 9700965.91 | 12701361.15 | 10966604.33 | 14421683.26 | 12097540.26 | 16292977.59 | 9687129.592 | 11996850.62 |
| SM 18:3;2O | 14729848.66 | 15370529.28 | 16524756.69 | 14061419.88 | 18072819.41 | 20329022.65 | 15263991.98 | 14026945.51 | 17094271.55 | 13632386.58 | 17136579.9 | 11125702.67 |
| Mitragynine | 9052850.598 | 6765223.752 | 8357574.801 | 7067854.844 | 9972771.042 | 7031145.238 | 10591731.2 | 9734258.822 | 7105504.668 | 9443943.279 | 7423737.213 | 8088416.554 |
| [2.3-dihydroxypropoxy][2-[docosa-4.7.10.13.16.19-hexaenoyloxy]-3-[octadec-9-enoyloxy]propoxy]phosphinic acid | 1666296.611 | 2695391.748 | 669170.4718 | 1484913.873 | 1963772.273 | 2411212.153 | 2074458.687 | 2293241.351 | 2100813.44 | 2207898.943 | 2404016.784 | 1782029.985 |
| Hexaethylene glycol | 24442898.9 | 27068212.6 | 33555625.09 | 33289775.29 | 24316499.2 | 25846369.93 | 24897053.6 | 22297615 | 27607941.58 | 23867052.72 | 25707452.76 | 42596224.46 |
| UHQ (mixed MS2 C11:db:UHQ) | 4143504.221 | 9020745.295 | 3116348.037 | 3186584.057 | 4729928.988 | 4669704.912 | 2308524.304 | 6298728.209 | 3178280.212 | 4852781.426 | 5304712.062 | 2282438.308 |
| 4-(Isopentyloxy)benzoic acid | 6915680.633 | 7225105.095 | 6193746.601 | 6778306.509 | 7216846.92 | 7195135.356 | 7122062.065 | 7267683.67 | 7144259.796 | 7037916.11 | 6816707.803 | 6653286.485 |
| PI 18:1_18:2 | 2284532.782 | 2379234.614 | 2057398.185 | 4503880.643 | 8016023.344 | 6606912.509 | 9207859.119 | 13930653.98 | 16948511.09 | 17363225.92 | 9322308.102 | 6306614.097 |
| Trospium | 1608625.98 | 1635656.312 | 1175057.48 | 1349986.923 | 1678546.038 | 1578554.685 | 1687818.432 | 1793733.336 | 1645362.274 | 1748217.165 | 1524042.052 | 1678030.245 |
| 1,1,3,3-Tetramethylguanidine | 656186.694 | 5009416.171 | 9314655.617 | 5398659.046 | 540720.1734 | 238899.1223 | 1412326.792 | 435842.5819 | 1216707.407 | 1637408.06 | 7119216.781 | 16172965.32 |

| Metabolites | PBS_1 | PBS_2 | PBS_3 | PBS_4 | BCG_1 | BCG_2 | BCG_3 | BCG_4 | BCGΔ*pe18*_1 | BCGΔ*pe18*_2 | BCGΔ*pe18*_3 | BCGΔ*pe18*_4 |
|---|---|---|---|---|---|---|---|---|---|---|---|---|
| Piperolein_B | 57857370.99 | 62781651.99 | 70637480.32 | 71929465.24 | 60555433.4 | 61154170.26 | 61059440.24 | 58226317.41 | 65014386.07 | 60546148.03 | 62896931.36 | 78560017.94 |
| 2.4-Dichlorobenzoic acid | 2436322.546 | 1565284.792 | 2567619.61 | 4543856.723 | 472387.5153 | 2577829.823 | 948486.364 | 7212873.052 | 340505.4345 | 7150500.649 | 2177882.654 | 4443508.394 |
| 3-Chloro-9H-carbazole | 10016429.3 | 10161441.21 | 10095738.17 | 10204447.59 | 9660829.243 | 9882100.82 | 10190982.78 | 10169772.81 | 10106750.27 | 9601535.152 | 9762187.713 | 9533947.459 |
| Cloperidone | 280956.6655 | 778364.2109 | 1342405.524 | 932687.7198 | 236758.0029 | 141719.3877 | 1032429.012 | 1192939.321 | 229263.2207 | 418234.1186 | 253094.5289 | 2731812.076 |
| TG 16:3_16:4_18:5 | 1184636.791 | 1192957.055 | 698011.1484 | 994921.9515 | 491898.1069 | 352383.1666 | 492411.2111 | 456794.3729 | 692350.9895 | 707990.4363 | 512412.0535 | 524139.8866 |
| Pelargonic acid | 72668385.85 | 54967171.09 | 56019601.95 | 72430693.77 | 61879481.13 | 71574933.89 | 84383778.97 | 66717094.82 | 69712945.85 | 77060018.01 | 56453506.88 | 83802929.94 |
| PC 15:1_15:1 | 979461.696 | 990531.0123 | 592571.1641 | 860598.6779 | 7872816.317 | 12730858.77 | 7468442.182 | 8122667.35 | 8723254.201 | 15787265.02 | 5762689.917 | 6788403.921 |
| TG O-8:0_16:4_18:3 | 5335679.221 | 5080175.845 | 4385915.691 | 4262035.993 | 4619928.487 | 4847245.972 | 5741772.394 | 5658860.447 | 5583678.027 | 5482754.413 | 4595510.642 | 3883937.537 |
| Methyl 1-piperazinecarboxylate | 66393.73784 | 803462.9639 | 1447775.864 | 1087993.437 | 88605.24656 | 109455.2801 | 247571.7582 | 161641.1665 | 308938.6725 | 159104.807 | 77150.10491 | 3249212.553 |
| 5-(2-methoxyethylamino)-3H-1,3,4-thiadiazole-2-thione | 2640547.754 | 2098985.334 | 3568906.143 | 2561852.654 | 12279222.43 | 10781421.72 | 13282662.34 | 12080567.39 | 10957495.62 | 9554136.185 | 11145181.97 | 10223916.28 |
| CoQ1 | 2390644.676 | 2353706.09 | 2384404.062 | 2346930.148 | 2455271.95 | 2335218.098 | 2429273.679 | 2630323.892 | 2248181.313 | 2578612.077 | 2783305.432 | 2451409.701 |

| **Metabolites** | **PBS_1** | **PBS_2** | **PBS_3** | **PBS_4** | **BCG_1** | **BCG_2** | **BCG_3** | **BCG_4** | **BCGΔ*pe18*_1** | **BCGΔ*pe18*_2** | **BCGΔ*pe18*_3** | **BCGΔ*pe18*_4** |
|---|---|---|---|---|---|---|---|---|---|---|---|---|
| Fluphenazine | 3762020.47 | 3870743.698 | 2700055.007 | 3364348.829 | 4169269.878 | 3601075.241 | 5511880.085 | 4163945.104 | 4088148.223 | 4600693.193 | 4155680.165 | 4628494.905 |
| LNAPS 3:0/N-6:0 | 852134.3245 | 902158.2963 | 820600.9831 | 936700.9059 | 806319.7842 | 788477.5379 | 915143.5963 | 899169.5093 | 861716.3005 | 872335.5786 | 841903.2479 | 885264.4853 |
| 1-Palmitoyl-2-thiopalmitoyl phosphatidylcholine | 948819.5906 | 3411676.284 | 3520470.296 | 581854.8518 | 2659154.09 | 5535167.935 | 3614660.304 | 3238695.945 | 2116353.874 | 5170021.506 | 6109571.331 | 3001809.61 |
| PG 18:1_18:2 | 6804315 | 4961162.1 | 7546042.38 | 4281785.813 | 56933629.36 | 34851773.9 | 101933874.3 | 64193346.53 | 90565255.13 | 92821944.27 | 59207601.85 | 27438547.38 |
| Deoxyribose triphosphate | 262248878.4 | 392544428.5 | 393536332.7 | 389815080.3 | 407968137 | 384246200.2 | 378612683.4 | 387410925.8 | 270347469.3 | 286865060.9 | 377581659.9 | 387082193.3 |
| 2-Hydroxy-6-methylquinoline-3-carbaldehyde | 40141144.26 | 34445153.03 | 35260320.4 | 42532409.83 | 48832056.76 | 41908193.1 | 76953126.36 | 49066151.4 | 50126817.23 | 48529550.4 | 37340560.16 | 51620116.29 |
| N-(3-((3-((3-Hydroxy-6-methylheptanoyl)oxy)-6-methylheptanoyl)oxy)-8-methylnonanoyl)serine | 9745633.666 | 6131156.09 | 5709907.052 | 6926213.999 | 6359882.618 | 6262191.946 | 5227402.433 | 4740842.476 | 5940888.389 | 7643402 | 7726568.897 | 3468098.029 |
| 1-(3,4-Dihydroxyphenyl)-6,7-dihydroxy-1,2-dihydronaphthalene-2,3-dicarboxylic acid | 7811137.735 | 7888370.287 | 7981288.442 | 7568437.145 | 7700938.215 | 7834513.607 | 7991134.699 | 8729148.636 | 7360579.882 | 7392541.143 | 7774118.418 | 7389523.191 |

| Metabolites | PBS_1 | PBS_2 | PBS_3 | PBS_4 | BCG_1 | BCG_2 | BCG_3 | BCG_4 | BCGΔ*pe18*_1 | BCGΔ*pe18*_2 | BCGΔ*pe18*_3 | BCGΔ*pe18*_4 |
|---|---|---|---|---|---|---|---|---|---|---|---|---|
| PI(16:0/18:1) | 823118.1846 | 893718.2019 | 908613.3154 | 754550.6625 | 4353631.151 | 1589440.402 | 3615620.104 | 2517784.982 | 3314178.121 | 3211347.374 | 2775919.855 | 1578270.592 |
| LysoPE(P-18:1(9Z)/0:0) | 24301349.74 | 22607435.72 | 19550167.89 | 18630688.23 | 2088415 7.3 | 19845116.42 | 22559443.27 | 19337355.63 | 19259796.55 | 22612056.52 | 22715718.27 | 13977874.07 |
| PS(15:0/22:4(7Z,10Z,13Z,16Z)) | 15352657.6 | 18522147.68 | 16694711.25 | 17645995.19 | 25726378.6 | 19722842.44 | 22802962.67 | 22472389.48 | 24428539.19 | 22650477.41 | 21326479 | 14027994.41 |
| xi-2,2,6-Trimethyl-1,4-cyclohexanedione | 4533958.181 | 8253795.978 | 3690563.073 | 27048481.22 | 2291598.256 | 6665586.309 | 51889698.46 | 1794562.913 | 15114210.78 | 10394218.5 | 1948359.527 | 32426882.56 |
| 4'-Methoxy-.alpha.-pyrrolidinopropiophenone | 1082954.121 | 6603492.107 | 6767060.948 | 5096584.843 | 5368669.158 | 6727690.529 | 5340717.257 | 6536188.771 | 5049907.559 | 6286207.296 | 5204497.93 | 5237181.671 |
| Maleic acid | 1683535.914 | 1631914.935 | 2559095.56 | 1813733.152 | 7600857.602 | 5593661.464 | 10906267.63 | 7802191.549 | 6201043.121 | 9570146.104 | 8687435.217 | 4705368.687 |
| N-(3-Chlorophenyl)-N'-(2-pyridinyl)thiourea | 11032699.67 | 7075810.681 | 13055830.16 | 8073794.12 | 14013586.62 | 10482825.21 | 11858148.93 | 13305375.93 | 12399501.68 | 13048213.5 | 10272797.03 | 8187209.811 |
| 1-(2,4-Dihydroxyphenyl)-2-(4-fluorophenyl)ethanone | 4542535.849 | 3157339.277 | 6086827.688 | 3441168.674 | 9896328.448 | 11487714.98 | 12529216.58 | 9925410.457 | 9298553.146 | 9385346.485 | 9855615.224 | 8638255.339 |
| Telon Blue RR 01 | 0 | 0 | 0 | 4123.412867 | 347063.679 | 105122.5571 | 382238.3273 | 217108.6437 | 216283.7089 | 372874.9639 | 242904.1277 | 184739.6534 |
| DGGA 13:0_14:0 | 34923071.38 | 38016605.56 | 32077715.16 | 35660416.69 | 45969466.81 | 44137769.47 | 36004161.21 | 36346780.88 | 34626901.62 | 46828451.02 | 44080130.27 | 37696934.6 |

| Metabolites | PBS_1 | PBS_2 | PBS_3 | PBS_4 | BCG_1 | BCG_2 | BCG_3 | BCG_4 | BCGΔ *pe18*_1 | BCGΔ *pe18*_2 | BCGΔ *pe18*_3 | BCGΔ*pe18*_4 |
|---|---|---|---|---|---|---|---|---|---|---|---|---|
| 1-[(4-Nitrophenyl)sulfonyl]piperazine | 0 | 797002.3389 | 1934792.174 | 1268493.151 | 0 | 0 | 295650.7371 | 25503.55413 | 144403.3963 | 57955.59316 | 0 | 8114602.42 |
| Pellitorine | 14738515.58 | 17638139.11 | 11659403.57 | 12261955.21 | 14609726 | 17910551.2 | 14929146.48 | 15184597.01 | 14641769.62 | 15104782.91 | 14006392.74 | 16318816.13 |
| LPS(18:1) | 1301955.473 | 1074136.169 | 1382818.02 | 1218639.382 | 2005058.891 | 1551417.92 | 1930976.706 | 1591757.997 | 2062201.081 | 2350882.964 | 2644304.117 | 1681560.193 |
| Azelaic acid | 8893622.618 | 9921428.343 | 11965754.65 | 11880810.7 | 9014364.449 | 8561551.754 | 18931068.52 | 8021929.294 | 10108643.7 | 9196914.307 | 9598334.365 | 18305415.52 |
| PE(18:3(9Z,12Z,15Z)/P-18:1(9Z)) | 4050819.163 | 4917432.207 | 4255490.404 | 4044635.385 | 3628988.657 | 2441719.094 | 4483948.457 | 3691644.056 | 3461399.95 | 4773077.39 | 3593917.335 | 2109014.737 |
| LNAPE 7:0/N-26:4 | 955125.8236 | 857564.586 | 1035188.952 | 1159901.851 | 1114087.601 | 791708.0833 | 1599174.897 | 1223960.946 | 1299006.952 | 1220673.456 | 943351.6319 | 1097953.938 |
| SM 23:1;2O/9:0 | 8803572.538 | 8925519.071 | 2625575.662 | 6444451.695 | 5443052.797 | 9773775.919 | 9754100.495 | 10420201.02 | 12439803.48 | 11520284.74 | 5481708.802 | 7735504.221 |
| DGDG O-20:1_22:6 | 1180463.189 | 1002146.299 | 1212735.362 | 1010036.506 | 2321589.756 | 2114491.381 | 241402.1738 | 1930766.831 | 1786628.675 | 1787561.301 | 2747350.058 | 916575.3604 |
| 3,7,8-Trihydroxy-3-methyl-1,4-dihydropyrano[4,3-b]chromen-10-one | 1093419.522 | 1053549.528 | 1092804.759 | 978987.6626 | 4208462.775 | 3818364.712 | 4047805.438 | 3618380.394 | 3842539.574 | 3836465.399 | 3657530.908 | 3633128.212 |
| PC(18:0/14:0) | 871523737.7 | 931223409.1 | 1120497661 | 742983594.8 | 672064812 | 555662963.5 | 929118013.1 | 720967246.9 | 698055314.8 | 805792479 | 699757036.8 | 295160198.5 |
| Maltobionate | 329493 | 275009 | 358337 | 385950 | 930307 | 904785 | 769026 | 848366 | 857550 | 643394 | 744826 | 785398 |

| **Metabolites** | **PBS_1** | **PBS_2** | **PBS_3** | **PBS_4** | **BCG_1** | **BCG_2** | **BCG_3** | **BCG_4** | **BCGΔ*pe18*_1** | **BCGΔ*pe18*_2** | **BCGΔ*pe18*_3** | **BCGΔ*pe18*_4** |
|---|---|---|---|---|---|---|---|---|---|---|---|---|
| | 1.248 | 4.642 | 6.329 | 5.315 | 5.352 | 7.234 | 0.591 | 5.572 | 9.786 | 6.189 | 6.458 | 6.718 |
| LDGTS 17:2 | 9930175.538 | 13058833.47 | 6906124.896 | 8435514.453 | 8452598.29 | 7252805.299 | 7457707.414 | 7283503.886 | 7409624.055 | 5356666.841 | 6883834.25 | 6715954.204 |
| 4,5-Dinitro-1H-imidazole | 8745893.648 | 3301387.418 | 4799037.333 | 2426954.899 | 7290736.101 | 4048946.216 | 3865258.375 | 3586798.357 | 7624195.745 | 7155562.023 | 3609614.727 | 4464498.643 |
| BMP 3:0_13:0 | 1214728.869 | 1353888.693 | 1483225.623 | 1280514.243 | 11313119.79 | 1213401.585 | 1183986.047 | 1211026.268 | 1371488.527 | 1200596.802 | 1447039.931 | 1322034.955 |
| LDGCC 18:5 | 43780973.72 | 51589182.07 | 42567641.41 | 44904701.65 | 46348553.11 | 47122092.75 | 45875302.55 | 45865757.09 | 47071559.12 | 46354253.67 | 45368629.58 | 50684599.02 |
| Guanfacine | 18007899.96 | 15907661.48 | 19937574.95 | 17205568.22 | 25532655.06 | 20484902.41 | 25558097.23 | 22634055.6 | 20394213.52 | 23003191.54 | 23638999.96 | 17942783.2 |
| 2,6-Di-tert-butylbenzoquinone | 1321787.24 | 1392455.683 | 1159091.924 | 1193293.222 | 1014199.272 | 2076371.041 | 942953.5028 | 890902.839 | 977118.4378 | 986615.994 | 976066.1521 | 1366359.911 |
| NAE 22:5 | 16271646.44 | 15625799.94 | 7721726.84 | 10257807.88 | 10950336.79 | 16036155.01 | 11655836.9 | 8804636.946 | 10966702.91 | 12728668.99 | 11753552.72 | 11423889 |
| PC(15:0/P-18:1(11Z)) | 10427772.48 | 10429774 | 15942309.83 | 12676700.45 | 26869036.43 | 17201674.86 | 24873662.59 | 17668839.86 | 19247749.37 | 22763534.72 | 24425094.03 | 9650203.916 |
| Glutamate | 14381374.64 | 11600628.32 | 20492313.17 | 13701544.67 | 159990118.8 | 109345225.6 | 142193227.6 | 133291203.3 | 133798181.3 | 108723103.7 | 113407352.2 | 113958562.2 |
| Methyl dihydrojasmonate, mixture of cis and trans | 5278443.011 | 5830522.843 | 5258161.079 | 5360161.623 | 5300743.889 | 5119028.15 | 5216569.344 | 4982713.74 | 5216457.5 | 5416691.161 | 5537552.786 | 5418651.595 |
| Cer 23:1;2O/38:0 | 945172. | 791402. | 940216. | 607184. | 524846. | 758928. | 564891. | 441189. | 441471. | 651566. | 675019. | 744770. |

| Metabolites | PBS_1 | PBS_2 | PBS_3 | PBS_4 | BCG_1 | BCG_2 | BCG_3 | BCG_4 | BCGΔ*pe18*_1 | BCGΔ*pe18*_2 | BCGΔ*pe18*_3 | BCGΔ*pe18*_4 |
|---|---|---|---|---|---|---|---|---|---|---|---|---|
| | 4055 | 5508 | 6744 | 8638 | 6412 | 6499 | 3806 | 5146 | 8922 | 9904 | 8987 | 1357 |
| SPB 16:1;2O | 23374218.22 | 25339994.7 | 23514186.84 | 26479741.7 | 22747870.11 | 25856104.28 | 27412674.4 | 26213198.04 | 20465756.79 | 23962852.17 | 20106692.95 | 21419061.38 |
| 3-Amino-3-(4-hydroxyphenyl)propanoic acid | 35480114.22 | 29166515.09 | 44357788.75 | 35775673.24 | 59875756.12 | 48961009.59 | 59004586.36 | 54753674.98 | 59563947.49 | 41798664.58 | 41491418.21 | 50744577.78 |
| Flubendazole | 212957.8534 | 132918.6536 | 185969.8165 | 165417.841 | 1230755.568 | 1196701.854 | 1086899.821 | 1093045.856 | 1142097.485 | 1058941.317 | 1142718.58 | 829704.5292 |
| Nafamostat | 3645537.983 | 3981923.81 | 3491283.418 | 3501773.058 | 3607278.12 | 3905293.429 | 3508805.126 | 3441179.473 | 3729059.99 | 3900490.391 | 3554668.017 | 3421290.741 |
| 1,7-Dioxa-4,10-diazacyclododecane | 16862661.78 | 15852700.42 | 24687564.88 | 17544930.37 | 18743157.45 | 16507349.27 | 27763747.42 | 19502888.78 | 16981729.03 | 19320270.37 | 22863411.22 | 15243869.05 |
| PC(31:0) | 55301264.11 | 59418233.13 | 59044622.46 | 45438178.03 | 102593735.8 | 81943512.59 | 122502414.4 | 100004692.7 | 103849138.7 | 115187096.8 | 93567207.42 | 44997419.05 |
| 3-(4-Hydroxyanilino)-5,5-dimethyl-2-cyclohexen-1-one | 789077.8392 | 847916.2279 | 742412.5111 | 802398.894 | 790025.2567 | 816898.1015 | 860125.2925 | 877837.7732 | 808288.3328 | 827507.7853 | 776684.9486 | 789950.4914 |
| Tris(2,4-di-tert-butylphenyl)phosphate | 251612955.9 | 281835899.8 | 158447324.3 | 209534774.1 | 214253123.3 | 178236758.9 | 241959676.7 | 210659792.2 | 206834123 | 245224092.7 | 183325727.3 | 213740703 |
| PC(39:4) | 3898903.362 | 4399530.799 | 4914063.402 | 5029640.333 | 7660045.107 | 5507345.403 | 7604308.139 | 6190969.358 | 7916803.073 | 7906885.745 | 7368842.611 | 3154978.518 |
| PE O-18:3_2:0 | 181446.8114 | 204108.7135 | 196455.8136 | 195786.191 | 193237.7096 | 184341.2015 | 207950.2861 | 166516.4137 | 180735.9268 | 151369.308 | 190389.24 | 206800.0822 |
| PE O-9:0_9:0 | 272124 | 251793 | 179018 | 164711 | 223994 | 246870 | 243490 | 193298 | 194668 | 229491 | 225868 | 149335 |

| **Metabolites** | **PBS_1** | **PBS_2** | **PBS_3** | **PBS_4** | **BCG_1** | **BCG_2** | **BCG_3** | **BCG_4** | **BCGΔ*pe18*_1** | **BCGΔ*pe18*_2** | **BCGΔ*pe18*_3** | **BCGΔ*pe18*_4** |
|---|---|---|---|---|---|---|---|---|---|---|---|---|
| | 7.752 | 5.879 | 5.104 | 8.041 | 9.593 | 4.932 | 7.632 | 5.274 | 5.882 | 2.164 | 1.871 | 9.738 |
| Ethyl nitroacetate | 2847691.947 | 1636093.342 | 2808371.481 | 1727518.406 | 9132825.548 | 4421164.053 | 7668445.887 | 6398408.229 | 7015511.377 | 6271569.092 | 4652302.835 | 5733367.681 |
| Ipratropium cation | 6564485.548 | 6663212.225 | 2662801.849 | 5313784.875 | 2804339.361 | 2663811.338 | 1719001.091 | 2613050.117 | 3602586.273 | 3298371.754 | 2014312.158 | 3137237.583 |
| 2-Iodo-3-(trifluoromethyl)benzoic acid | 1141591.419 | 603712.3029 | 1763416.986 | 1160509.377 | 7354299.176 | 7943682.825 | 8504402.056 | 7876319.644 | 4401936.818 | 7534560.909 | 7860469.867 | 9676409.427 |
| 4-Bromo-2,6-di-tert-butylphenol | 6689367.648 | 6232506.599 | 7640031.751 | 6340464.245 | 7514491.002 | 8419139.082 | 8263203.674 | 11003927.8 | 7474366.622 | 7138616.168 | 7054891.639 | 8811744.123 |
| PE O-16:1_22:4 | 4227849.29 | 2632479.636 | 3390414.625 | 3635191.379 | 3273760.831 | 160234.9567 | 3652576.933 | 3045074.898 | 4556412.539 | 4037146.495 | 3307493.778 | 3245880.061 |
| 5-Chloro-N,2-dihydroxybenzamide | 34325661.49 | 28164144.62 | 42712428.55 | 36096096.82 | 33634868.96 | 38163986.92 | 36195224.2 | 35520360.92 | 36174353.37 | 26025031.4 | 28548445.72 | 33150069.11 |
| PG(20:4(5Z,8Z,11Z,14Z)/22:6(4Z,7Z,10Z,13Z,16Z,19Z)) | 8323673.164 | 8533007.809 | 5202581.714 | 6596272.414 | 8345214.789 | 7332781.032 | 7716958.384 | 8647486.932 | 8638592.259 | 9086255.462 | 7564603.523 | 7154894.445 |
| SM(d18:1/22:0) | 18139639.48 | 18130702.01 | 23240803.62 | 18091385.25 | 17009253 | 15164747.06 | 22380689.47 | 17383317.53 | 17094808.88 | 20716348.87 | 19382024.69 | 7660931.784 |
| 3-(1,3-Benzothiazol-2-yl)-7-hydroxy-2H-chromen-2-one | 6635778.808 | 6392199.621 | 6273474.511 | 6181023.346 | 7002883.679 | 6367510.085 | 5947212.699 | 6198524.332 | 6105663.947 | 6060172.585 | 6045823.586 | 5604613.524 |
| Laurocapram | 2350763485 | 2343550037 | 1876869598 | 1549712778 | 1427315351 | 1215490753 | 1392781813 | 1860156447 | 1443901106 | 1382369511 | 1374124335 | 831038899.2 |

| **Metabolites** | **PBS_1** | **PBS_2** | **PBS_3** | **PBS_4** | **BCG_1** | **BCG_2** | **BCG_3** | **BCG_4** | **BCGΔ*pe18*_1** | **BCGΔ*pe18*_2** | **BCGΔ*pe18*_3** | **BCGΔ*pe18*_4** |
|---|---|---|---|---|---|---|---|---|---|---|---|---|
| 4-(Hydroxymethyl)benzenediazonium(1+) | 2017109.85 | 5458856.956 | 1204692.932 | 2469500.301 | 15344852.5 | 5631746.355 | 8666027.425 | 9244574.875 | 4395084.142 | 16355914.82 | 10714330.4 | 5596492.741 |
| Polysorbate 60 | 1882235.182 | 154378.5271 | 1715827.517 | 1547355.398 | 805084.2641 | 1355531.825 | 855723.4176 | 1055756.75 | 1803539.498 | 495259.2934 | 454961.2125 | 783105.2898 |
| PC(32:1) | 81817209.95 | 72523479.8 | 131340557.3 | 82574807.41 | 552748179.6 | 259390519 | 257218997.3 | 556003474.5 | 258408674.2 | 226614722.1 | 551595112.7 | 346813140 |
| PS(20:4(5Z,8Z,11Z,14Z)/15:0) | 26190775.78 | 27493641.93 | 21661375.23 | 22263336.72 | 35120981.7 | 26873841.43 | 31748110.53 | 32884700.14 | 33525758.25 | 30949548.52 | 26375320.2 | 21062216.74 |
| Ethyl 2-(4-fluorophenyl)-3-(trifluoromethyl)pyrazole-4-carboxylate | 118227.5285 | 124087.91 | 140864.1598 | 124713.1225 | 389982.9035 | 611383.8669 | 1245993.37 | 1180293.088 | 1354665.93 | 462333.7917 | 885022.6166 | 1703875.405 |
| Tetrapropylammonium cation | 4957922.685 | 7536336 | 4755058.177 | 4847626.667 | 4273940.027 | 5148596.988 | 4950423.614 | 5204083.269 | 5621705.27 | 4123566.943 | 4362750.237 | 5161530.373 |
| MeAIB | 19730511.71 | 16077165.34 | 32512009.39 | 22865807.49 | 57679622.72 | 63635514.51 | 59525607.22 | 51149234.73 | 58808485.22 | 41963322.33 | 49058031.23 | 51458190.61 |
| Methionine sulfoxide | 369824.7861 | 466489.368 | 270113.502 | 373846.947 | 3054007.853 | 1933532.74 | 1425626.896 | 1167677.007 | 1783913.922 | 1825356.471 | 1689826.14 | 1554954.17 |
| 2-(1-Hydroxycyclohexyl)butanoic acid | 2120266.458 | 2109920.917 | 2075299.722 | 2077158.235 | 2070966.007 | 2158024.293 | 2039707.507 | 1950167.602 | 2137846.352 | 2278163.536 | 2039284.343 | 2313313.845 |

| Metabolites | PBS_1 | PBS_2 | PBS_3 | PBS_4 | BCG_1 | BCG_2 | BCG_3 | BCG_4 | BCGΔ*pe18*_1 | BCGΔ*pe18*_2 | BCGΔ*pe18*_3 | BCGΔ*pe18*_4 |
|---|---|---|---|---|---|---|---|---|---|---|---|---|
| Methanone, [6-bromo-2-methyl-1-[2-(4-morpholinyl)ethyl]-1H-indol-3-yl](4-methoxyphenyl)- | 77349.74951 | 100996.9963 | 175825.1804 | 152310.4073 | 1043701.613 | 859194.7483 | 885223.276 | 791659.2447 | 800441.9059 | 757743.5733 | 860835.5599 | 1058778.627 |
| 4-Allyl-5-(2-methylphenyl)-4H-1,2,4-triazol-3-yl hydrosulfide | 1357302.194 | 1458287.389 | 1487336.982 | 6166748.374 | 1010251.314 | 1638370.721 | 7707839.93 | 1085587.316 | 3672546.126 | 2006630.576 | 1111438.425 | 6732161.18 |
| PC 8:0_26:3 | 11578138.66 | 25255604.84 | 12431450.94 | 13280586.08 | 40873188.55 | 73225546.16 | 60315447 | 79990908.45 | 84910117.66 | 52489259.39 | 53772799.02 | 59894905.33 |
| Carnitine | 16449773.4 | 17361393.14 | 19813881.85 | 21902417.65 | 21673281.27 | 12575248.46 | 17253919.02 | 14640921.71 | 12011582.2 | 14131707.56 | 13026834.78 | 27654051.98 |
| DGTS 3:0_16:3 | 23224915.41 | 23763542.93 | 21007980.48 | 22849090.14 | 22199084.45 | 24661061.33 | 24586728.12 | 24113286.89 | 22148490.16 | 24366887.38 | 23464810.02 | 21317292.21 |
| 8-Oxononanoic acid | 2940207.717 | 2719367.301 | 2538875.656 | 2760898.918 | 2302053.734 | 6215348.123 | 27210262.06 | 2918026.949 | 7945719.684 | 3487373.819 | 2046202.702 | 24185478.69 |
| 1-Cyclopentyl-3-methyl-1H-pyrazol-5-amine | 17721675.68 | 9245041.616 | 19621705.01 | 19527375.8 | 15507327.58 | 15209413.79 | 17247660.72 | 14338932.42 | 14298946.72 | 17886007.86 | 21543402.71 | 21129550.38 |
| Monobutyl phthalate | 1356785.911 | 2302691.548 | 1530582.985 | 1721511.521 | 2278589.218 | 3006415.746 | 1691811.811 | 1869735.353 | 1851068.534 | 2342733.698 | 2622696.69 | 2282344.385 |
| Mianserin N-oxide | 427469 | 542048 | 433517 | 526200 | 628633 | 613064 | 627141 | 619524 | 578307 | 593750 | 510616 | 714358 |

| Metabolites | PBS_1 | PBS_2 | PBS_3 | PBS_4 | BCG_1 | BCG_2 | BCG_3 | BCG_4 | BCGΔ*pe18*_1 | BCGΔ*pe18*_2 | BCGΔ*pe18*_3 | BCGΔ*pe18*_4 |
|---|---|---|---|---|---|---|---|---|---|---|---|---|
| | 6.688 | 1.611 | 8.329 | 1.112 | 0.666 | 3.364 | 1.223 | 2.806 | 9.523 | 3.286 | 2.319 | 3.553 |
| Glucose chloride | 51669906.45 | 39564142.86 | 61385711.11 | 51372087.31 | 33079622.35 | 44689951.04 | 36957207.68 | 24675503.87 | 39338321.02 | 23981667.15 | 23979805.27 | 39630731.95 |
| Ethyl 6-chloro[1,2,4]triazolo[4,3-b]pyridazine-3-carboxylate | 462293.9823 | 395055.6558 | 576764.5121 | 427157.6268 | 2916214.149 | 2227036.386 | 2926995.023 | 2632772.82 | 2588018.359 | 2375096.521 | 2356055.337 | 2250803.833 |
| PC O-11:0_16:0 | 1813554.831 | 1616868.276 | 1871905.79 | 1960433.114 | 1863351.334 | 2516689.118 | 2205371.021 | 2103318.407 | 1836389.845 | 1636511.325 | 2377071.514 | 1678795.949 |
| 1-Propyl-1H-tetraazol-5-ylamine | 197420.9932 | 2242509.679 | 4551651.427 | 3048273.561 | 275975.8513 | 1884990.842 | 956033.6213 | 212134.9225 | 808263.2884 | 1172185.846 | 1859365.773 | 2011126.988 |
| Benzoic acid | 68022799.92 | 53951336.24 | 46804104.78 | 51543782.85 | 53595831.79 | 56786380.98 | 56843838.8 | 36514278.36 | 52001816.65 | 51730132.88 | 47984383.4 | 61606854.03 |
| Myrsinone | 24406455.87 | 12877424.59 | 11537334.8 | 16247566.48 | 19607981.43 | 34729235.91 | 17283912.85 | 32355840.81 | 21027548.45 | 28843878.51 | 22842481.09 | 16720874.24 |
| .alpha.-Methyl-L-tyrosine | 2860052.208 | 3109395.011 | 2759203.416 | 3758975.461 | 3729022.909 | 2893828.054 | 3146332.936 | 2945153.734 | 2849509.829 | 3941581.577 | 2818831.926 | 2904582.29 |
| Euscaphic acid | 5760504.175 | 6068215.244 | 6021343.486 | 5636281.281 | 5491537.479 | 6051650.327 | 5466073.433 | 5346128.226 | 5806990.756 | 5972636.166 | 5890183.795 | 5561265.019 |
| DG 26:4 | 7202099.513 | 8234613.086 | 6276730.272 | 8095039.562 | 8387269.221 | 8709979.062 | 9904048.64 | 8089710.865 | 7679645.033 | 7633279.168 | 7350393.418 | 7762058.327 |
| 5-Chloro-2-(trifluoromethyl)-3H-imidazo[4,5-b]pyridine | 37839518.06 | 47560529.17 | 48311285.8 | 34593743.19 | 20556236.51 | 46263184.21 | 31436476.54 | 30901701.34 | 29835347.59 | 34465937.85 | 32045577.21 | 28975196.25 |

| **Metabolites** | **PBS_1** | **PBS_2** | **PBS_3** | **PBS_4** | **BCG_1** | **BCG_2** | **BCG_3** | **BCG_4** | **BCGΔ*pe18*_1** | **BCGΔ*pe18*_2** | **BCGΔ*pe18*_3** | **BCGΔ*pe18*_4** |
|---|---|---|---|---|---|---|---|---|---|---|---|---|
| LPC O-18:5 | 38172377.85 | 35275900.31 | 27812135.91 | 25328555.47 | 32986933.31 | 37722343.61 | 33099619.94 | 31334635.29 | 29711654.74 | 36082831.39 | 34646248.04 | 26727325.85 |
| Plastoquinone-9 | 7545932.762 | 8465214.03 | 7312187.466 | 7077272.825 | 5134322.007 | 3863399.751 | 6447356.818 | 4770815.924 | 4825616.992 | 5559418.71 | 4766383.424 | 1675608.926 |
| PE O-13:1_6:0 | 361168.1565 | 1584356.492 | 213249.0233 | 1703610.645 | 395400.8452 | 246565.1188 | 212709.3859 | 142404.4789 | 358823.9891 | 428502.1499 | 227813.838 | 182328.5113 |
| 5-(4-Hydroxyphenyl)-5-phenylhydantoin | 658239.5107 | 457323.212 | 554973.5813 | 599145.8698 | 64018.74397 | 834294.1968 | 1098180.345 | 966682.4634 | 1223678.599 | 704225.88 | 1120054.581 | 851131.5201 |
| 4-(dimethylamino)butanoate | 19228390.47 | 17559386 | 27664863.81 | 21523278.61 | 27948823.78 | 25753887.04 | 27926919.99 | 25750362.27 | 27992538.68 | 19739541.63 | 17808707.96 | 24815507.65 |
| PC(22:5(7Z,10Z,13Z,16Z,19Z)/20:5(5Z,8Z,11Z,14Z,17Z)) | 27341443.18 | 37064782.07 | 12098681.27 | 23042659.14 | 19700272.47 | 23376079.27 | 21415115.58 | 20995494.77 | 18918794.35 | 25249019.29 | 25079833.91 | 10252692.87 |
| Candoxatril | 16523319.38 | 6311613.02 | 16493487.83 | 8329780.126 | 8319875.356 | 26408621.89 | 14383819.15 | 16199257.56 | 20908104.29 | 10442319.7 | 16557698.82 | 6198183.157 |
| PI 17:1_18:1 | 639218.3232 | 795938.423 | 341407.9693 | 498193.9941 | 852227.0468 | 203696.6978 | 939527.1428 | 842951.7997 | 847691.8894 | 1046993.327 | 790522.4522 | 326969.7363 |
| 1-(3,5-Dihydroxyphenyl)-12-hydroxytridecan-2-one | 11955575.75 | 13130391.44 | 7339464.193 | 12030333.42 | 7838627.647 | 8775278.479 | 7155886.117 | 8752608.139 | 7018709.008 | 9312774.953 | 7676989.149 | 22570068.27 |

| Metabolites | PBS_1 | PBS_2 | PBS_3 | PBS_4 | BCG_1 | BCG_2 | BCG_3 | BCG_4 | BCGΔ*pe18*_1 | BCGΔ*pe18*_2 | BCGΔ*pe18*_3 | BCGΔ*pe18*_4 |
|---|---|---|---|---|---|---|---|---|---|---|---|---|
| 1,3-Dimethyl-6-(propylamino)-2,4(1H,3H)-pyrimidinedione | 3153827.124 | 5488627.026 | 11950203.53 | 7422710.425 | 3037619.396 | 3425019.971 | 3683747.491 | 3309855.19 | 3097915.52 | 3136722.036 | 3495611.671 | 9280931.976 |
| Isopeonidin_3-rutinoside | 52839838.22 | 48372608.52 | 49098083.19 | 61520802.33 | 53510874.28 | 50007364.99 | 63233841.67 | 55551447.56 | 33559357.44 | 52990565.84 | 51483927.66 | 50268614.69 |
| 3,8-Dibromo-6-chloroimidazo[1,2-a]pyridine | 3114900.369 | 2897196.027 | 3065769.419 | 3071224.132 | 2794888.028 | 3172208.43 | 3069307.986 | 2790177.075 | 2976645.78 | 2825774.426 | 3206642.599 | 3004214.547 |
| SMGDG O-24:5_18:2 | 7202.950937 | 0 | 0 | 0 | 536011.8891 | 711316.1596 | 355681.6252 | 478461.2255 | 707623.48 | 539881.3524 | 672644.6721 | 513640.1834 |
| LPE(22:4) | 824869.7275 | 1264085.784 | 1987985.685 | 1279103.941 | 747317.3038 | 1482623.381 | 1994029.492 | 1475709.165 | 1785821.536 | 2814687.884 | 2504896.295 | 1808347.095 |
| DGDG 3:0_26:0 | 4459669.158 | 4409856.254 | 4025985.242 | 4223506.669 | 5523086.467 | 3762757.325 | 4166083.887 | 4603215.58 | 4137609.215 | 4290103.119 | 4550396.965 | 3482388.465 |
| Cer 23:1;3O/21:0;(2OH) | 1353501.113 | 1331799.388 | 1144979.373 | 1325385.959 | 1282934.493 | 1477681.019 | 1195491 | 1244197.959 | 1137427.727 | 1028812.58 | 1329214.88 | 1201160.866 |
| Neotame | 1157047.063 | 1628071.209 | 1053623 | 138890.3534 | 1258337.01 | 2088762.55 | 1621776.547 | 2175804.863 | 2072958.52 | 1568402.866 | 527580.5022 | 2399854.247 |
| (3beta)-3-hydroxyolean-12-en-28-oic acid | 3067824.992 | 3418520.476 | 3124790.806 | 3406931.733 | 4037447.717 | 2079437.535 | 3550799.403 | 3255679.265 | 3431425.589 | 3491273.932 | 2419344.786 | 2748495.3 |
| SL 23:2;O/11:0 | 161535 | 221728 | 143209 | 200827 | 182431 | 159088 | 161273 | 198536 | 145553 | 183678 | 158909 | 182117 |

| Metabolites | PBS_1 | PBS_2 | PBS_3 | PBS_4 | BCG_1 | BCG_2 | BCG_3 | BCG_4 | BCGΔ*pe18*_1 | BCGΔ*pe18*_2 | BCGΔ*pe18*_3 | BCGΔ*pe18*_4 |
|---|---|---|---|---|---|---|---|---|---|---|---|---|
| | 93.95 | 26.49 | 72.81 | 49.09 | 36.22 | 41.13 | 70.84 | 77.69 | 64.77 | 96.91 | 18.33 | 50.55 |
| Cer 10:1;2O/9:0 | 2982326.828 | 3491770.734 | 2815855.165 | 3013345.783 | 2952225.658 | 3060239.664 | 3164654.887 | 3485550.203 | 3187389.202 | 3190972.094 | 2795719.819 | 2967551.467 |
| Ethyl 4-aminobutanoate | 291107913.6 | 239052067.5 | 393682381.7 | 315526929.7 | 433981480.3 | 393153773.3 | 437114228.3 | 414743760.1 | 421491456.3 | 288439274.1 | 294593854.2 | 380869315 |
| 2-oleoyl-sn-glycero-3-phosphoglycerol | 279424.2994 | 205314.6088 | 186514.8089 | 443954.5622 | 931672.6365 | 971563.9283 | 1333634.984 | 893354.7543 | 810304.5231 | 650575.6296 | 1198451.404 | 2263422.427 |
| PC 18:1_18:2 | 2168269.394 | 2425853.875 | 2555060.796 | 871530.5793 | 2054362.641 | 37490.64447 | 1719198.715 | 1210241.189 | 2158160.979 | 2107450.603 | 2225616.768 | 950169.6176 |
| trans-11-Methyl-2-dodecenoic acid | 201688.285 | 156997.1461 | 203694.4303 | 116452.2938 | 5370563.816 | 5387878.627 | 3728189.97 | 4289982.861 | 4966207.122 | 4344474.154 | 5236399.523 | 4561125.847 |
| DGDG 9:0_18:2 | 233754.1884 | 268346.7655 | 79144.19689 | 228739.2778 | 159473.4505 | 115765.9061 | 105968.2918 | 213581.6666 | 306921.6689 | 123647.5082 | 83080.78863 | 82394.04615 |
| Puerariaglycoside 3 | 12027276.54 | 11764748.27 | 11590757.58 | 11805304.86 | 11363815.32 | 11986164.8 | 11755528 | 11658435.38 | 11783529.25 | 11572616.24 | 11575315.98 | 11429847.79 |
| PC(33:1) | 1723958.814 | 5605127.397 | 4554596.836 | 4689695.61 | 6718160.711 | 5566317.217 | 9129616.666 | 6406701.627 | 7600647.021 | 7278809.59 | 5936887.607 | 3962809.111 |
| 3,3-Dimethylpyrrolidin-2-one | 14492336.77 | 12836370.7 | 9074823.879 | 8726062.673 | 8498363.695 | 10391585.77 | 8067492.764 | 8146171.575 | 8474702.868 | 8094361.028 | 8879380.719 | 9911230.907 |
| Vernakalant | 2913780.523 | 2914513.46 | 2280225.33 | 2508544.004 | 4549798.637 | 3274050.178 | 2816929.747 | 1998785.151 | 2650688.78 | 2540833.264 | 5279262.529 | 2452011.967 |
| 3-[[5-(2-Cyanoethylthio)-2-thioxo-1,3-dithiol-4-yl]thio]prop | 339287.1557 | 303821.7293 | 547240.6601 | 426919.0692 | 3607777.68 | 2471972.54 | 3741852.165 | 2918336.221 | 2212764.676 | 3568749.641 | 2982292.868 | 2333408.063 |

| Metabolites | PBS_1 | PBS_2 | PBS_3 | PBS_4 | BCG_1 | BCG_2 | BCG_3 | BCG_4 | BCGΔ*pe18*_1 | BCGΔ*pe18*_2 | BCGΔ*pe18*_3 | BCGΔ*pe18*_4 |
|---|---|---|---|---|---|---|---|---|---|---|---|---|
| ionitrile | | | | | | | | | | | | |
| Isovaleramide | 51259022.76 | 66509570.73 | 42989590.94 | 205808942 | 196162245.2 | 31709773.29 | 217575263.8 | 170216855.3 | 198150124.5 | 45444953.06 | 181210069.9 | 164799942.3 |
| 20-Deoxynarasin | 62563168.04 | 71852143.83 | 54057037.44 | 59908765.5 | 96668251.18 | 73181147.92 | 88204296.03 | 87660937.81 | 90099918.77 | 82200321.88 | 70763120.86 | 48494522.11 |
| SM 38:5;3O | 9675359.315 | 11948614.89 | 5753270.35 | 7996376.623 | 2371526.232 | 2206012.287 | 3210930.208 | 3142282.213 | 3171102.313 | 4598059.89 | 2981777.679 | 2189269.659 |
| Tetraethylammonium cation | 58170191.71 | 39715187.89 | 48277011.54 | 54937392.2 | 48243727.17 | 46966333.19 | 49016611.97 | 38840764.08 | 32490674.26 | 39073065.98 | 42236581.03 | 50703463.84 |
| Hygric acid | 440740.8101 | 1445393.795 | 871437.3605 | 682521.1711 | 1451960.762 | 1117660.815 | 1325728.389 | 1156766.075 | 1137878.626 | 1000753.362 | 3547999.645 | 1131518.544 |
| 9(10)-Epoxy-12Z-octadecenoic acid | 3046203.324 | 2606395.567 | 2462015.788 | 2730599.864 | 2511184.123 | 3541566.755 | 2432196.013 | 2710689.702 | 2104001.631 | 2904550.148 | 2664077.773 | 3378286.744 |
| N-Stearoyltaurine | 15816676.6 | 18078146.71 | 10876014.57 | 15885209.38 | 28698190.74 | 32921280.48 | 28046259.99 | 21913729.4 | 31043753.49 | 26746745.19 | 32441552 | 23052179.66 |
| LPE 22:6 | 34572068.14 | 38043188.02 | 37477426.63 | 34152739 | 75818855.1 | 76695904.92 | 55131059.93 | 64863813.42 | 61886743.76 | 67752915 | 64479910.51 | 65383541.24 |
| Bicalutamide | 606090.5309 | 621305.7792 | 929418.4789 | 509191.0435 | 2494334.468 | 936319.253 | 2755025.194 | 1536497.613 | 1399690.265 | 2026655.715 | 2045636.601 | 1184835.04 |
| PC(22:5(7Z,10Z,13Z,16Z,19Z)/22:6(4Z,7Z,10Z,13Z,16Z, | 4790966.93 | 5159479.781 | 2257132.611 | 3131756.35 | 3848731.498 | 3884259.86 | 3610966.485 | 5322296.745 | 4577739.229 | 4496281.561 | 3394601.738 | 3880771.564 |

| **Metabolites** | **PBS_1** | **PBS_2** | **PBS_3** | **PBS_4** | **BCG_1** | **BCG_2** | **BCG_3** | **BCG_4** | **BCGΔ*pe18*_1** | **BCGΔ*pe18*_2** | **BCGΔ*pe18*_3** | **BCGΔ*pe18*_4** |
|---|---|---|---|---|---|---|---|---|---|---|---|---|
| 19Z)) | | | | | | | | | | | | |
| Phthalic anhydride | 11036707.47 | 12608242.3 | 9100711.903 | 8301413.716 | 19268069.95 | 21857940.62 | 18916057.86 | 20166644.85 | 17214867.55 | 16844981.25 | 16486052.53 | 15488017.69 |
| 3'-Hydroxyprostanozol | 829416.2457 | 581980.0848 | 671442.3657 | 1198376.909 | 879496.5108 | 714292.1028 | 891987.64 | 947240.7307 | 687070.0625 | 826623.1368 | 1015704.825 | 716572.2886 |
| Triphenylphosphine oxide | 6303622.265 | 7701043.043 | 8055083.016 | 7463921.474 | 6372920.255 | 6188206.034 | 6518019.015 | 5897074.68 | 6315716.395 | 6481929.927 | 6180783.958 | 9526005.706 |
| Cer 14:1;2O/32:3;2O | 818295.5639 | 861973.5448 | 822245.0231 | 938135.9065 | 2583776.607 | 918083.2866 | 908493.1723 | 339259.3933 | 1654332.209 | 950119.2821 | 964220.6396 | 809417.5439 |
| Phenylalanine | 138614097.3 | 85182268.13 | 187547721.4 | 121238293 | 163673565.1 | 180176961.1 | 134811544.7 | 139646174.3 | 116227091.4 | 149979958 | 125844515.8 | 138561620.6 |
| 1-Deoxy-D-glucitol | 9586713.207 | 8828104.614 | 12922625.25 | 10812259.08 | 19372050.7 | 17205005.15 | 19199150.36 | 16303612.75 | 16288952.88 | 14069311.71 | 15213277.45 | 15150128.98 |
| N-Butyl-N'-phenylthiourea | 28810518.69 | 27165141.71 | 24490258.26 | 44940218.75 | 25675978.67 | 26752176.99 | 33819844.78 | 31654452.46 | 25392393.83 | 30586816.06 | 32679284.89 | 27718697.82 |
| Roseoflavin | 437055.462 | 470061.2546 | 655143.0671 | 390988.6943 | 716995.8984 | 462866.2089 | 786582.3532 | 556897.3879 | 448495.8636 | 569930.849 | 673191.9583 | 424484.6008 |
| Moguisteine | 15333.86252 | 23878.60015 | 40207.19187 | 30415.48891 | 3145722.073 | 3685351.004 | 2898180.125 | 3106735.407 | 3950776.955 | 3175730.703 | 2777115.836 | 3105470.729 |
| Methanone, (1-pentyl-1H-indol-3-yl)(2,2,3,3-tetram | 4303676.453 | 4468666.806 | 4155361.305 | 4036178.281 | 4043074.205 | 4528486.105 | 4618445.163 | 4454818.952 | 4274717.602 | 4038598.633 | 5088873.128 | 4119955.023 |

| Metabolites | PBS_1 | PBS_2 | PBS_3 | PBS_4 | BCG_1 | BCG_2 | BCG_3 | BCG_4 | BCGΔ *pe18*_1 | BCGΔ *pe18*_2 | BCGΔ *pe18*_3 | BCGΔ*pe18*_4 |
|---|---|---|---|---|---|---|---|---|---|---|---|---|
| ethylcyclopropyl)- | | | | | | | | | | | | |
| PG(18:1(9Z)/20:3(8Z,11Z,14Z)) | 7584226.614 | 9181116.906 | 8048415.164 | 8453115.205 | 12682387.22 | 9934389.716 | 10713549.69 | 10984857.13 | 11869622.13 | 11103801.55 | 10223590.1 | 6605820.69 |
| Vomitoxin | 3104418.896 | 3394932.43 | 2652462.348 | 3254757.33 | 3074593.313 | 3238100.703 | 2954869.242 | 2861943.096 | 3155497.196 | 3021804.415 | 3109314.007 | 2588973.914 |
| ADENOSINE DIPHOSPHATE RIBOSE | 10769.94815 | 34474.4209 | 25627.66374 | 36141.83689 | 171028.3334 | 163360.3437 | 142668.6789 | 135729.5033 | 218510.092 | 271375.113 | 213964.1991 | 269133.307 |
| Oxymatrine | 2310148.68 | 3179433.02 | 3389335.365 | 2906914.598 | 2878544.353 | 2317926.318 | 2703947.066 | 2399096.157 | 2750416.418 | 3004075.613 | 2573856.887 | 4311604.975 |
| Cer 12:1;2O/3:0 | 1839106.678 | 2439995.565 | 2571086.126 | 2817787.697 | 1894593.606 | 2043561.427 | 2176767.454 | 2074394.122 | 2069201.11 | 2080731.376 | 1991375.512 | 4991395.439 |
| PC O-40:8 | 9842388.472 | 11648053.8 | 3134093.702 | 7822550.329 | 2235509.28 | 2418533.339 | 2197921.151 | 2796552.246 | 3754403.328 | 2560922.78 | 2079535.39 | 2585511.094 |
| 2-(HYDROXYMETHYL)BUTANOIC ACID | 1375535.945 | 1386164.888 | 2022172.244 | 1628898.165 | 1807991.135 | 1881810.684 | 1809441.558 | 1862280.909 | 1919659.373 | 1627642.128 | 1942885.777 | 1713587.805 |
| PC O-16:4_22:4 | 750009.2768 | 300044.1849 | 833304.0409 | 724836.0354 | 655851.4513 | 388870.3152 | 868627.2163 | 786746.7526 | 789028.5348 | 795115.862 | 517991.9286 | 403356.2331 |
| 3-Dodecyloxypropylamine | 41516760.48 | 25575563.5 | 19566279.08 | 26169288.17 | 20428877.26 | 16979299.88 | 15299560.52 | 22682667.27 | 16992111.96 | 17754031.46 | 25785918.04 | 12941226.43 |
| Difloxacin | 116562. | 64639.4 | 97312.0 | 92256.5 | 116002 | 733125. | 921697. | 473265. | 875215. | 821819. | 877806. | 811323. |

| Metabolites | PBS_1 | PBS_2 | PBS_3 | PBS_4 | BCG_1 | BCG_2 | BCG_3 | BCG_4 | BCGΔ *pe18*_1 | BCGΔ *pe18*_2 | BCGΔ *pe18*_3 | BCGΔ*pe18*_4 |
|---|---|---|---|---|---|---|---|---|---|---|---|---|
| | 2997 | 6942 | 1294 | 1952 | 9.703 | 3364 | 8246 | 1371 | 1604 | 6693 | 2543 | 5305 |
| FR-33289 | 4735567.689 | 3806076.177 | 5189858.071 | 3909018.062 | 24981356.45 | 14490003.12 | 24608302.52 | 19119825.55 | 19573855.75 | 19378309.69 | 17862938.47 | 11170192.8 |
| Isovalerylcarnitine | 306194.7722 | 399093.608 | 355458.7175 | 560240.5052 | 289700.5068 | 251044.6085 | 499447.8043 | 233377.0207 | 267371.6862 | 279118.9823 | 248527.9647 | 510827.4309 |
| PS 10:0_26:1 | 4157437.044 | 4861572.925 | 5181599.884 | 6120617.687 | 3341050.965 | 4016077.043 | 3548940.732 | 4444217.038 | 5130559.11 | 3982957.827 | 733332.5668 | 5752784.307 |
| Ethyl 6-chloro-4-hydroxyquinoline-3-carboxylate | 1635863.965 | 1294091.599 | 2236129.064 | 1534255.262 | 8562840.063 | 7188366.615 | 8913100.047 | 7894543.206 | 8009693.769 | 6569991.994 | 7639704.373 | 7773874.262 |
| PS(18:0/20:2(11Z,14Z)) | 2304658.135 | 2027569.698 | 1443162.074 | 1654736.839 | 4033905.72 | 2498079.081 | 4859343.135 | 3218776.488 | 3582563.262 | 5741495.137 | 5094768.875 | 2402916.096 |
| Trilostane | 379681.8571 | 460868.2807 | 415803.524 | 420895.7819 | 513578.9136 | 481688.6675 | 368280.8572 | 483930.0824 | 390717.618 | 442760.3625 | 515370.9747 | 393139.1835 |
| 2,3-Dimethylbenzene-1-sulfonic acid | 3184773.916 | 3300923.539 | 3219529.26 | 3189301.827 | 3166334.967 | 3197715.256 | 3477743.364 | 3180404.757 | 3207500.793 | 3074119.567 | 3152897.651 | 3311523.494 |
| (17Z)-16-(Acetyloxy)-6-hydroxy-4,8,14-trimethyl-3,7-dioxo-18-norcholesta-17,24-dien-21-oic acid | 2262794.113 | 2278487.318 | 3257914.352 | 2846781.037 | 4722657.2 | 3868974.286 | 3567074.128 | 5920665.323 | 3855801.814 | 4088334.194 | 4099672.263 | 4596450.234 |

| Metabolites | PBS_1 | PBS_2 | PBS_3 | PBS_4 | BCG_1 | BCG_2 | BCG_3 | BCG_4 | BCGΔ*pe18*_1 | BCGΔ*pe18*_2 | BCGΔ*pe18*_3 | BCGΔ*pe18*_4 |
|---|---|---|---|---|---|---|---|---|---|---|---|---|
| 1-Stearoyl-2-arachidonoyl-sn-glycero-3-phospho-(1'-myo-inositol) | 16462447.8 | 23530189.28 | 17808550.56 | 18506354.61 | 17033008.6 | 13112534.05 | 19986823.13 | 16456112.07 | 17975024.45 | 24367710.94 | 18041212.73 | 11870178.19 |
| 2-Methylbenzoic acid | 5009499.178 | 4556363.001 | 5363823.602 | 5104252.86 | 4426366.153 | 5289304.239 | 4599329.201 | 4056302.014 | 4577051.366 | 4923855.535 | 4570113.843 | 5078414.036 |
| Methyl_nigakinone | 42434286.95 | 27423019.74 | 53800069.35 | 32131156.42 | 123749308.7 | 82421186.91 | 108508154.5 | 99206223.81 | 93151669.84 | 81333825.77 | 89744014.02 | 76949961.67 |
| SL 22:0;O/16:2;O | 5564552.936 | 7908714.365 | 3824012.184 | 7490796.469 | 4526313.635 | 5169973.149 | 4872901.923 | 6667958.49 | 4510638.221 | 7174283.689 | 5487634.32 | 4919059.874 |
| Rhamnose | 2627918.891 | 2354748.98 | 2221437.961 | 2075213.509 | 2533295.806 | 2864647.27 | 3964119.076 | 2307136.928 | 2523437.491 | 2254014.5 | 2862986.932 | 2849226.625 |
| Sethoxydim | 334639.0129 | 527975.8773 | 661329.3672 | 640204.2703 | 328391.1389 | 287239.9026 | 441162.3832 | 362081.768 | 387111.4046 | 414284.5598 | 332911.1294 | 739419.8008 |
| (9Z,12E)-15,16-Dihydroxyoctadeca-9,12-dienoic acid | 6799874.814 | 6628416.29 | 9347879.476 | 7843109.67 | 5884339.01 | 8035523.072 | 6431061.19 | 11378208.42 | 6150689.931 | 5376555.486 | 6324397.54 | 8355853.478 |
| 1-[3,4-Dihydroxy-5-(hydroxymethyl)oxolan-2-yl]-4-hydroxyhydropyridin-2-one | 5682320.769 | 6342556.766 | 5842317.3 | 6171175.283 | 5301989.899 | 5668730.83 | 5345104.53 | 5535422.698 | 5281853.571 | 5309623.326 | 5538314.893 | 8133610.185 |
| M274T76 | 103636525.6 | 122498887.5 | 124106316.5 | 115773721.2 | 143128658.6 | 115885208.4 | 137710223.5 | 145461031.1 | 119260943.4 | 117055017.9 | 101094357.5 | 108276429.1 |

| Metabolites | PBS_1 | PBS_2 | PBS_3 | PBS_4 | BCG_1 | BCG_2 | BCG_3 | BCG_4 | BCGΔ*pe18*_1 | BCGΔ*pe18*_2 | BCGΔ*pe18*_3 | BCGΔ*pe18*_4 |
|---|---|---|---|---|---|---|---|---|---|---|---|---|
| PS 10:0_26:4 | 1698573.85 | 1158275.876 | 1898908.711 | 1768794.955 | 1711606.905 | 990602.4249 | 1715540.222 | 1860961.348 | 2334831.549 | 2642965.522 | 993014.8815 | 1532575.158 |
| LPE(O-18:2) | 5505481.225 | 4408860.777 | 4567534.071 | 4339718.735 | 5003590.391 | 3656489.052 | 5311387.729 | 4450856.656 | 4934594.497 | 5482300.564 | 5372799.473 | 3405622.757 |
| 13-Oxo-9,11-tridecadienoic acid | 34968569.36 | 2782873.106 | 2423554.829 | 31352485.69 | 1931590.639 | 2050418.496 | 1990907.036 | 2480897.851 | 2079938.051 | 1522269.403 | 2669614.107 | 1829458.243 |
| LPG 22:4 | 20366527.33 | 19593379.29 | 18775450.18 | 16190705.08 | 7115276.415 | 7824021.497 | 12380475.5 | 9689144.119 | 8334637.479 | 11469502.87 | 12292052.15 | 7560764.94 |
| PE O-16:0_22:5 | 1878896.594 | 1408030.746 | 1208341.055 | 1711719.741 | 1457917.02 | 109296.4298 | 1619198.805 | 1298105.379 | 3381001.962 | 1387744.018 | 1596683.589 | 1929861.935 |
| Deoxyadenosine | 7475705.384 | 8755795.751 | 342544.7164 | 7545135.332 | 8284309.279 | 6700079.309 | 4387238.841 | 4890008.777 | 8002116.805 | 4944583.601 | 2906766.459 | 8233514.289 |
| Piperazine-N,N'-bis(2-hydroxypropanesulfonic acid) | 304898.3796 | 207251.1687 | 257085.5507 | 198912.8537 | 15717253.8 | 17928996.66 | 13838817.3 | 15074248.01 | 20363251.8 | 15556623.81 | 12354272.81 | 14369715.42 |
| MGDG 11:0_20:4 | 1936266.839 | 1467232.973 | 1340111.399 | 1137268.383 | 1414035.671 | 1495377.624 | 1148865.511 | 1482266.133 | 1212424.268 | 1626881.084 | 1354055.187 | 1070145.211 |
| LNAPS 16:3/N-26:6 | 381304.1975 | 284683.7404 | 182221.8694 | 346846.9847 | 641362.7074 | 759002.1973 | 939322.9539 | 577207.8971 | 635281.1917 | 678652.2407 | 554182.5297 | 773252.1903 |
| DMPO | 10078656.73 | 5151926.892 | 12935368.13 | 18096898.97 | 9861110.998 | 8874055.662 | 7682195.049 | 11594636.62 | 8697339.135 | 8577926.334 | 9433326.104 | 9538891.907 |
| LPC O-18:1 | 9436523.68 | 12957233.8 | 10199890.48 | 11483600.92 | 6842448.494 | 7489205.647 | 6758857.682 | 6618285.135 | 6426356.928 | 6682197.38 | 6866635.751 | 5558987.878 |
| PC(33:0) | 267914 | 286535 | 350651 | 299518 | 746766 | 561722 | 898264 | 675886 | 695388 | 709998 | 604508 | 317620 |

| Metabolites | PBS_1 | PBS_2 | PBS_3 | PBS_4 | BCG_1 | BCG_2 | BCG_3 | BCG_4 | BCGΔ *pe18*_1 | BCGΔ *pe18*_2 | BCGΔ *pe18*_3 | BCGΔ*pe18*_4 |
|---|---|---|---|---|---|---|---|---|---|---|---|---|
| | 56.87 | 45.42 | 02.94 | 97.91 | 98.9 | 39.92 | 24.95 | 64.05 | 88.1 | 78.12 | 46.88 | 72.57 |
| Arginine | 3795566.708 | 2718470.109 | 1557506.268 | 3614096.458 | 3899003.865 | 3435211.847 | 1024460.493 | 3732936.193 | 4267582.151 | 1834366.898 | 1625609.504 | 3888531.995 |
| Di(3-methyl-4-octyl) phthalate | 2749055.59 | 4907317.019 | 3244642.392 | 3603322.487 | 3589399.385 | 3388012.768 | 2588614.7 | 3422737.852 | 2837546.501 | 3479723.586 | 2587753.984 | 3490573.05 |
| 5,6,7,8-Tetrahydro-1H-spiro[benzo[4,5]thieno[2,3-d]pyrimidine-2,1'-cyclohexane]-4(3H)-thione | 3348231.985 | 3129846.4 | 3193418.588 | 2872370.161 | 4111586.024 | 3798807.257 | 4003309.051 | 3919703.131 | 4034356.552 | 3480260.763 | 3397733.66 | 3391921.051 |
| 4-Chloro-N-(5-fluoro-2-methylphenyl)benzamide | 6818543.379 | 4883597.846 | 8003492.75 | 5420592.14 | 12192074.15 | 7591007.373 | 9238084.698 | 9127366.277 | 8582745.556 | 7846811.615 | 8263884.12 | 6291312.99 |
| PC(14:0/P-18:1(9Z)) | 11347288.84 | 13238315.12 | 7405939.652 | 7779148.315 | 14642929.21 | 13953948.82 | 19626709.24 | 13986335.57 | 18205158.44 | 21970347.8 | 16934632.02 | 11112051.08 |
| N-Allyl-N-(2-bromo-2-propenyl)amine | 895451.988 | 735779.8521 | 1760598.784 | 991783.234 | 3446882.505 | 2851386.936 | 3123183.946 | 2941138.373 | 3453769.293 | 2348282.372 | 3416286.341 | 3365060.214 |
| NAE 22:4 | 7108168.189 | 5328079.808 | 4768386.162 | 5248594.031 | 4001269.206 | 6979502.416 | 4829508.83 | 4958306.818 | 5089043.556 | 5214545.191 | 7327467.622 | 4709801.961 |
| 3-{4-[(Phenylcarbonyl)amino]phenyl} propanoic acid | 3791795.72 | 4003980.759 | 3408724.835 | 3602906.23 | 3976236.464 | 3582722.822 | 3704538.116 | 3609525.624 | 3973091.541 | 3714052.355 | 3728339.216 | 3372146.237 |
| 2,4(1H,3H)-Quinazolinedithione | 395776.2052 | 289805.8009 | 589177.7943 | 362915.3117 | 1267058.112 | 748312.6991 | 959535.8525 | 844672.7248 | 958534.9938 | 709129.4905 | 786526.0459 | 679306.3854 |

| Metabolites | PBS_1 | PBS_2 | PBS_3 | PBS_4 | BCG_1 | BCG_2 | BCG_3 | BCG_4 | BCGΔ*pe18*_1 | BCGΔ*pe18*_2 | BCGΔ*pe18*_3 | BCGΔ*pe18*_4 |
|---|---|---|---|---|---|---|---|---|---|---|---|---|
| Benazepril | 5781813.926 | 7020741.175 | 5403090.846 | 5225340.122 | 4907510.368 | 5362480.05 | 4518793.197 | 4707936.243 | 4363742.761 | 4697746.639 | 4603588.651 | 7203285.377 |
| 4-Nitrophenol | 14454889.1 | 16009102.4 | 15017848.95 | 15001927.68 | 13628374.17 | 13634574.16 | 13777098.16 | 13033761.34 | 13264445.98 | 13224245.44 | 13281000.11 | 16931852.46 |
| LPC O-17:0 | 3493518.545 | 5735523.17 | 3252928.869 | 3716969.588 | 3941695.618 | 3340621.191 | 3249500.878 | 2017323.173 | 3355290.095 | 2946300.671 | 3515998.707 | 1700820.43 |
| C20 sphinganine | 3676326.509 | 3683222.197 | 2728822.884 | 3625174.843 | 3244633.245 | 3372289.559 | 3362196.776 | 3534407.464 | 3324251.283 | 3261629.409 | 3367945.773 | 2773056.593 |
| Piericidin C1 | 5727207.784 | 5532990.535 | 7012565.493 | 6776411.303 | 6802741.888 | 6363929.402 | 7008183.569 | 6925338.801 | 6974837.795 | 7302752.787 | 7166642.526 | 6849575.364 |
| Benzoic acid, 3-[2-[2-chloro-4-[[3-(2,6-dichlorophenyl)-5-(1-methylethyl)-4-isoxazolyl]methoxy]phenyl]ethenyl]- | 2400094.555 | 2194772.283 | 2456130.088 | 1963370.804 | 11306201.4 | 7154704.012 | 10953032.3 | 8648954.788 | 7760057.562 | 7624494.691 | 8936150.097 | 8897817.929 |
| 2-Chloro-4,5-dimethoxybenzoic acid | 16154554.36 | 10564466.73 | 16271348.62 | 16301776.74 | 9933274.133 | 12303252.77 | 9330339.965 | 11576232.99 | 10680056.37 | 5997688.98 | 6500534.663 | 11713432.82 |
| Lithocholate 3-O-glucuronide | 47902.52667 | 66647.89984 | 114074.3841 | 63619.95461 | 144693.1787 | 145331.203 | 134158.1587 | 149590.6408 | 169428.8419 | 171651.5822 | 147094.5597 | 169578.4837 |
| SPB 20:0;2O | 3699356.275 | 3460557.625 | 2108609.645 | 2754468.759 | 2725793.909 | 2693516.181 | 2610102.702 | 2908802.465 | 2530984.577 | 2350854.18 | 2571504.639 | 2515625.703 |
| DL-.beta.-Hydroxy palmitic acid | 1985022.278 | 2759796.185 | 2121753.226 | 2442344.718 | 3148279.789 | 3090853.514 | 2268406.623 | 2610172.697 | 2019533.04 | 2313669.488 | 2233616.474 | 4509568.31 |

| Metabolites | PBS_1 | PBS_2 | PBS_3 | PBS_4 | BCG_1 | BCG_2 | BCG_3 | BCG_4 | BCGΔ*pe18*_1 | BCGΔ*pe18*_2 | BCGΔ*pe18*_3 | BCGΔ*pe18*_4 |
|---|---|---|---|---|---|---|---|---|---|---|---|---|
| PG(18:1(9Z)/18:3(9Z,12Z,15Z)) | 11747846.24 | 13001535.63 | 9836900.795 | 10889883.7 | 16730491.65 | 12977656.13 | 14690965.95 | 15808641.25 | 15619196.88 | 13631168.37 | 12097678.66 | 9892857.905 |
| PE O-18:1_2:0 | 347260.8927 | 355459.46 | 535290.142 | 316487.2819 | 1157331.912 | 842000.5572 | 1040943.554 | 932345.4613 | 707874.8221 | 908894.6949 | 1183020.947 | 646199.0083 |
| Cer 18:1;2O/10:0 | 1658181.425 | 1693959.147 | 695859.2594 | 872777.1383 | 408555.9301 | 413862.0715 | 375059.4127 | 556180.2208 | 501054.7746 | 298350.2758 | 473698.1743 | 383833.4116 |
| Ornithine | 378575.9234 | 158428.7849 | 1465157.951 | 332292.5098 | 1790703.685 | 1324471.053 | 2714483.927 | 1806329.668 | 1589981.345 | 2044809.374 | 2060018.22 | 1722754.075 |
| LPI 22:5 | 11576050.71 | 11028063.74 | 10378685.48 | 10135217.48 | 5100766.481 | 4293227.407 | 3283376.001 | 6083423.123 | 4807129.794 | 3946486.042 | 4667858.324 | 4374421.942 |
| LPC 20:2 | 991801.804 | 1015358.442 | 857375.6266 | 677425.4989 | 1076386.344 | 1287714.344 | 1043242.051 | 911635.6765 | 932733.1758 | 1006003.125 | 1326202.527 | 836282.4848 |
| SHexCer 27:0;2O | 246859.0324 | 303904.311 | 173957.2883 | 259740.3551 | 293060.7284 | 274687.3621 | 341998.2633 | 278654.8059 | 317833.6744 | 275573.2004 | 233490.2146 | 242064.8642 |
| Oleic acid methyl ester | 2149888.284 | 2720249.954 | 1543036.442 | 2554815.49 | 1965842.316 | 2172561.383 | 2043697.351 | 2265721.46 | 2001484.646 | 2357787.116 | 2099684.337 | 2138902.751 |
| PE(P-16:0/20:5(5Z,8Z,11Z,14Z,17Z)) | 13421766.21 | 14202389.85 | 8858276.501 | 11002875.06 | 27773252.04 | 18659001.1 | 20304934.8 | 21904869.61 | 22703957.58 | 21579458.51 | 17218205.78 | 12926117.88 |
| 5-(4-Hydroxybenzylidene)-1,3-dimethyl-2,4,6(1H,3H,5H)-pyrimidinetrione | 2230264.256 | 2335003.673 | 42153211 | 2387156.388 | 3386397.415 | 2365249.625 | 71092059.66 | 4031573.864 | 2259125.497 | 41076514.75 | 41584068.66 | 2383474.132 |

| Metabolites | PBS_1 | PBS_2 | PBS_3 | PBS_4 | BCG_1 | BCG_2 | BCG_3 | BCG_4 | BCGΔ*pe18*_1 | BCGΔ*pe18*_2 | BCGΔ*pe18*_3 | BCGΔ*pe18*_4 |
|---|---|---|---|---|---|---|---|---|---|---|---|---|
| (4S,5Z,6S)-4-(2-methoxy-2-oxoethyl)-5-[2-[(E)-3-phenylprop-2-enoyl]oxyethylidene]-6-[(2S,3R,4S,5S,6R)-3,4,5-trihydroxy-6-(hydroxymethyl)oxan-2-yl]oxy-4H-pyran-3-carboxylic acid | 123111930.1 | 66476735.75 | 121037794.5 | 121874574.5 | 85422027.36 | 145193513.9 | 115068385.6 | 129892928.3 | 128318567.2 | 68690125.52 | 103649999.9 | 80732496.97 |
| SHexCer 36:4;3O | 19373509.82 | 22379751.05 | 8147670.838 | 9256268.908 | 3720990.387 | 4528350.043 | 5589869.823 | 4290024.505 | 7577913.732 | 8063061.001 | 4532438.533 | 5203005.746 |
| DGCC 9:0_13:1 | 1556027.7 | 2311227.617 | 1448722.172 | 1472995.68 | 1431927.451 | 1667402.632 | 1126755.748 | 1623904.459 | 1164328.419 | 1416727.229 | 1497359.389 | 1461107.863 |
| 4-Fluoro-2-[(methylsulfonyl)amino]benzoic acid | 1142502.704 | 956357.8581 | 1574941.909 | 1121107.373 | 6239620.149 | 3910431.123 | 6923790.01 | 4621172.012 | 4105846.431 | 4920839.744 | 4881370.318 | 3861308.508 |
| Fudosteine | 1304473.628 | 4019135.306 | 1298230.634 | 3140974.996 | 12937893.99 | 6712875.85 | 2755641.219 | 2239056.049 | 4290005.912 | 8889407.303 | 6097934.012 | 5997910.101 |
| N-(Piperidin-3-yl)methanesulfonamide | 3174786.787 | 3220298.469 | 2758343.738 | 2817529.641 | 2028674.07 | 2271691.123 | 1875590.081 | 2112441.366 | 2322643.201 | 2115032.238 | 2344007.218 | 2135223.041 |
| PG(18:2(9Z,12Z)/16:0) | 11921833.32 | 11437802.31 | 12110499.86 | 10682927.08 | 11231087.04 | 8742461.055 | 12256952.6 | 10917103.3 | 9000718.665 | 13354780.19 | 10904648.24 | 7775947.546 |
| skullcapflavone II | 198278 | 217043 | 236225 | 201864 | 228045 | 238177 | 227745 | 233240 | 229747 | 240334 | 250301 | 211978 |

| Metabolites | PBS_1 | PBS_2 | PBS_3 | PBS_4 | BCG_1 | BCG_2 | BCG_3 | BCG_4 | BCGΔ*pe18*_1 | BCGΔ*pe18*_2 | BCGΔ*pe18*_3 | BCGΔ*pe18*_4 |
|---|---|---|---|---|---|---|---|---|---|---|---|---|
| | 6.511 | 8.13 | 3.579 | 6.953 | 8.878 | 1.542 | 1.528 | 1.118 | 8.208 | 5.679 | 0.023 | 8.475 |
| 4-Hydroxy-1-methyl-7,8,9,10-tetrahydro-6H-[1,3]thiazocino[2,3-f]purin-2(1H)-one | 1516254.067 | 4747075.869 | 1936498.989 | 1437706.92 | 1690063.297 | 1987775.522 | 1792383.701 | 1874689.928 | 2152832.69 | 1332615.532 | 6336429.892 | 1766243.308 |
| Indoxyl sulfate | 4962529.583 | 2101121.869 | 8880060.28 | 8406367.133 | 10441770.74 | 8517332.323 | 8676881.946 | 11446190.64 | 8645080.148 | 9246120.64 | 11791673.86 | 5667419.411 |
| 1,2-dihydroxyheptadec-16-yn-4-yl acetate | 3828988.575 | 3204503.949 | 5982484.483 | 5217120.771 | 20795024.23 | 12109558.38 | 20235900.97 | 14450774.15 | 10781944.75 | 17129263.07 | 17279903.22 | 8861920.202 |
| 3-Chloro-6-ethyl-1-benzothiophene-2-carboxylic acid | 10938313.19 | 8827400.133 | 18010728.03 | 13180121.86 | 16982866.38 | 16094172.98 | 17330159.93 | 16562933.2 | 14311192.1 | 11421995.69 | 17266242.01 | 14329029.15 |
| PE O-24:5_14:1 | 8845058.704 | 7278548.316 | 9312657.381 | 8604963.559 | 8699759.157 | 10081338.76 | 8970634.455 | 9324272.55 | 6914664.592 | 8785559.066 | 9688544.825 | 7433756.247 |
| LPG 22:6 | 32495470.08 | 31428918.91 | 43052464.64 | 41329603.55 | 80652782.26 | 81589793.54 | 50329982.27 | 97518815.32 | 88395840.53 | 45970675.71 | 45395618.93 | 70748874.03 |
| SMGDG O-12:0_28:5 | 224313.7239 | 268871.1016 | 264413.2045 | 232058.8965 | 309050.9852 | 300417.9863 | 282498.6668 | 234146.3584 | 267337.6805 | 264009.9821 | 244694.4466 | 286111.7081 |
| PG 18:1_20:3 | 10600207.25 | 12015698.49 | 4359742.259 | 8196242.88 | 13467397.42 | 3330866.96 | 32524512.27 | 11112126.31 | 23769737.23 | 52340293.41 | 19118523.36 | 10248511.2 |
| PI O-14:1_16:1 | 10737301.24 | 11185060.02 | 10152709.7 | 10872098.87 | 10768679.15 | 12130526.75 | 12266723.23 | 10797638.95 | 11156454.45 | 11270962.83 | 12174676.32 | 10615406.96 |

| Metabolites | PBS_1 | PBS_2 | PBS_3 | PBS_4 | BCG_1 | BCG_2 | BCG_3 | BCG_4 | BCGΔ*pe18*_1 | BCGΔ*pe18*_2 | BCGΔ*pe18*_3 | BCGΔ*pe18*_4 |
|---|---|---|---|---|---|---|---|---|---|---|---|---|
| 5-(Diphenylphosphinyl)pentanoic acid | 28807433.31 | 24452874.34 | 50722416.53 | 33228690.78 | 37796151.06 | 33556516.09 | 39949494.36 | 34701358.25 | 33444527.83 | 25713941.41 | 31373928.43 | 29863872.97 |
| LPC 18:5 | 535369.7079 | 4414505.334 | 219940.124 | 4075402.311 | 6424115.335 | 5333754.3 | 792066.4181 | 1073150.618 | 5493443.479 | 4880725.856 | 6845718.16 | 985448.1298 |
| Ethyl sulfate | 3744443.875 | 14204035.61 | 30342341.24 | 20091683.86 | 3236075.594 | 2508270.715 | 8667298.451 | 2565099.492 | 4773756.107 | 3948583.364 | 3245276.575 | 68529266.18 |
| 7-Hydroxy-4,8-dimethyl-2H-chromen-2-one | 1922438.25 | 1484942.575 | 1786650.126 | 1492047.245 | 3174046.062 | 4294431.616 | 3114800.79 | 2785697.635 | 2955828.077 | 2148885.3 | 2857952.81 | 2779983.221 |
| Moxonidine | 31128397.35 | 23393592.78 | 34608135.88 | 25259762.59 | 49479580.02 | 30930704.9 | 37608703.59 | 37357149.29 | 35224157.63 | 33270012.24 | 32247278.73 | 24883740.15 |
| Palmatine cation | 1284588.36 | 1258074.881 | 1210232.509 | 1321619.424 | 1272142.726 | 1334565.403 | 1262022.831 | 1281584.661 | 1323310.635 | 1284337.205 | 1266431.515 | 1185582.894 |
| N-[3-(Trifluoromethyl)phenyl]thiourea | 1749745.199 | 1105372.135 | 1614105.979 | 1286182.002 | 1545145.237 | 1886261.049 | 1787336.63 | 1820332.719 | 1720561.44 | 1285014.129 | 1366200.752 | 1386167.137 |
| 1-(4-Hydroxyphenyl)-2-methylaminoethanone | 19194149.14 | 24640997.3 | 24834974.35 | 27207203.05 | 24069439.51 | 23365977.05 | 31327036.66 | 26922945.57 | 25897100.99 | 14873493.12 | 16175659.93 | 28200872.63 |
| Isovaleric acid | 4610531.432 | 5014207.947 | 5469556.202 | 6150762.824 | 7775688.678 | 7468101.793 | 5678493.762 | 5217739.064 | 6479145.63 | 4776850.898 | 4866906.478 | 5711832.664 |
| DGGA 12:0_12:0 | 2574948.545 | 2621979.686 | 1918813.105 | 2784029.865 | 2651415.353 | 2391201.647 | 2329845.404 | 2141714.833 | 2758665.611 | 2832334.382 | 1918247.694 | 1754423.301 |
| Caprolactam | 100786 | 515192 | 129353 | 180968 | 986111 | 887405 | 768219 | 115946 | 869733 | 857792 | 943332 | 953889 |

| Metabolites | PBS_1 | PBS_2 | PBS_3 | PBS_4 | BCG_1 | BCG_2 | BCG_3 | BCG_4 | BCGΔ*pe18*_1 | BCGΔ*pe18*_2 | BCGΔ*pe18*_3 | BCGΔ*pe18*_4 |
|---|---|---|---|---|---|---|---|---|---|---|---|---|
| | 56.73 | 6.892 | 68.13 | 98.97 | 0.998 | 5.662 | 5.049 | 36.62 | 9.135 | 6.334 | 6.104 | 1.907 |
| Fenoterol | 1586635.581 | 1385193.938 | 1936729.821 | 1562008.708 | 2474633.574 | 2076639.513 | 2611293.581 | 2087861.121 | 1922062.417 | 2064978.601 | 2150055.414 | 1756000.801 |
| Thymidine | 89583967.07 | 76726582.97 | 158037930.4 | 101553973.5 | 117403808.8 | 105033474.4 | 125080380 | 109031453.9 | 104993277.3 | 79851632.81 | 97475094 | 93839673.87 |
| 6-Chloro-7-fluoroisatin | 942577.2146 | 585384.0028 | 724957.1385 | 773214.9569 | 2469562.922 | 1362441.268 | 2176714.292 | 2091789.014 | 1833309.681 | 1558758.488 | 1607888.768 | 1783634.439 |
| Methylprednisolone succinate | 5045595.642 | 7080598.51 | 3582459.675 | 7693141.287 | 2319994.797 | 5643422.332 | 2807093.878 | 5729230.943 | 5589138.502 | 2280349.768 | 5135518.945 | 1916427.356 |
| 1-(4-Chlorophenyl)-2-[(5-methyl-4H-1,2,4-triazol-3-yl)sulfanyl]ethanone | 4841663.471 | 4347547.077 | 6041671.567 | 4689534.655 | 7607378.669 | 6047572.434 | 7936804.908 | 7098088.951 | 5735209.195 | 6646335.649 | 6932374.015 | 5081509.554 |
| PC(31:1) | 9591612.004 | 9556379.919 | 10364104.1 | 9436108.177 | 25089938 | 16352627.58 | 23934855.27 | 19234319.24 | 22436296.18 | 22630115.71 | 22024389.21 | 11587953.86 |
| Mannitol | 54690520.57 | 59095992.69 | 83746485.28 | 58618125.82 | 167571737.2 | 106694960.3 | 154652380.9 | 124644203.9 | 153673786.3 | 199505098.8 | 113063096.1 | 116150809.1 |
| N1-(alpha-D-ribosyl)-5,6-dimethyl-benzimidazole | 504065.5154 | 354580.5088 | 553486.0633 | 526476.8581 | 3141117.462 | 2962125.104 | 2643426.698 | 2564729.083 | 3498986.63 | 2690767.261 | 3439167.974 | 3212417.174 |
| 3-Amino-1,1,1-trifluoro-2-butanol | 2856971.712 | 2376823.327 | 4081289.055 | 2698639.146 | 9291491.535 | 6824825.339 | 9839107.164 | 8455393.657 | 6968855.519 | 6808371.75 | 7419450.284 | 6152916.494 |
| PG 16:1_18:2 | 889514.4989 | 509574.3965 | 298002.4412 | 79048.03902 | 10636657.71 | 4686019.354 | 12853206.09 | 8180503.2 | 7116625.968 | 7731397.41 | 7537913.533 | 6491931.908 |

| **Metabolites** | **PBS_1** | **PBS_2** | **PBS_3** | **PBS_4** | **BCG_1** | **BCG_2** | **BCG_3** | **BCG_4** | **BCGΔ*pe18*_1** | **BCGΔ*pe18*_2** | **BCGΔ*pe18*_3** | **BCGΔ*pe18*_4** |
|---|---|---|---|---|---|---|---|---|---|---|---|---|
| Clobetasol_propionate | 4719824.5 | 5161664.722 | 4572032.608 | 5391891.261 | 4866716.435 | 4761928.461 | 4450552.181 | 4544784.331 | 4728247.829 | 4173757.761 | 4568485.284 | 3697502.192 |
| Turanose | 473170.8303 | 447306.8038 | 656803.8536 | 523157.5728 | 154225535.8 | 139888358.6 | 115797351.5 | 117316471.7 | 160291042.3 | 122417727.8 | 130177395.6 | 159424984.6 |
| Sophocarpine | 6671221.783 | 7031139.16 | 7181384.796 | 7095544.29 | 6772706.289 | 6695493.213 | 6703195.93 | 6333080.66 | 6503483.848 | 6479530.747 | 6762556.853 | 6877140.543 |
| Docosapentaenoic acid (DPA) | 137985653.2 | 124219070.2 | 167425842.4 | 142192600.4 | 52952528.24 | 78989291.16 | 52212151.22 | 74787015.3 | 63503745.18 | 38250515.73 | 54761039.18 | 57244668.21 |
| PA 2:0_15:1 | 1738454.515 | 1888022.985 | 1644177.676 | 1694694.045 | 1845872.463 | 1708759.7 | 1914576.216 | 1786147.689 | 1809117.819 | 1755978.324 | 1798446.721 | 1737876.169 |
| DL-threo-1-Phenyl-2-palmitoylamino-3-morpholino-1-propanol | 619986.0702 | 388648.7176 | 450741.2606 | 557874.2741 | 487937.1955 | 550328.1216 | 480979.0632 | 264642.7834 | 424662.9505 | 507274.1973 | 505295.6682 | 542463.1057 |
| N6-(1-Iminoethyl)-L-lysine | 5879310.758 | 7512336.266 | 4781339.341 | 6202942.88 | 5861112.788 | 5374602.752 | 5131883.445 | 1506501.27 | 5101443.503 | 1476715.154 | 1441995.05 | 7563631.543 |
| Bursin | 2334179.751 | 2444386.462 | 2388991.058 | 2422813.325 | 2275618.287 | 2429783.572 | 2459893.172 | 2311588.098 | 2304478.917 | 2091560.721 | 2349548.491 | 2445314.227 |
| PI 2:0_8:0 | 632585.1448 | 27272.01884 | 48906.59205 | 57714.5004 | 37935.91161 | 663045.4362 | 17959.34801 | 34811.47061 | 399466.5887 | 36276.19457 | 432119.5515 | 14589.02215 |
| Nordihydroguaiaretic acid | 3678134.708 | 4149667.079 | 3755695.189 | 3744245.984 | 3895499.136 | 3847412.637 | 3891455.36 | 3811638.582 | 3936145.858 | 4040439.277 | 3770131.724 | 3625372.58 |
| LPG 18:2 | 5146208.915 | 5231459.14 | 10373787.42 | 6454339.69 | 15751083.9 | 17400882.99 | 18549768.67 | 14544576.12 | 15559617.89 | 17179929.9 | 23051548.87 | 18448718.39 |

| Metabolites | PBS_1 | PBS_2 | PBS_3 | PBS_4 | BCG_1 | BCG_2 | BCG_3 | BCG_4 | BCGΔ*pe18*_1 | BCGΔ*pe18*_2 | BCGΔ*pe18*_3 | BCGΔ*pe18*_4 |
|---|---|---|---|---|---|---|---|---|---|---|---|---|
| N-Lauroylglycine | 5299451.699 | 2137835.582 | 1525684.45 | 2552603.677 | 1422836.816 | 2022967.433 | 1858691.217 | 2729198.139 | 1490851.337 | 1156290.7 | 1614967.512 | 2465825.769 |
| L-.beta.-Homoglutamine | 10433019.87 | 9052543.431 | 15404000.32 | 10059781.1 | 36657616.72 | 27618435.32 | 38504221.84 | 33233701.72 | 27790374.12 | 27298005.79 | 29463731.5 | 23409294.81 |
| Eremopetasinorol | 30096608.7 | 25936298.39 | 25420643.51 | 31700598.4 | 13300189.16 | 18740215.44 | 19080067.45 | 17620222.34 | 18495329.16 | 15076768.15 | 15020248.51 | 11026141.6 |
| (8.Xi.,9.Xi.,14.Xi.)-3-((6-Deoxy-.alpha.-L-mannopyranosyl)oxy)-1,5,14-trihydroxycard-20(22)-enolide | 1194620.832 | 1199657.395 | 1057741.219 | 1183253.047 | 1226734.793 | 1129500.671 | 1217754.567 | 1126193.19 | 1167227.15 | 1163190.916 | 1094727.7 | 1076064.653 |
| Nordentatin | 1063387.73 | 3608005.539 | 7407934.799 | 4337605.807 | 560774.2622 | 305859.1768 | 687135.7901 | 188232.0794 | 520578.1943 | 746880.0401 | 677374.231 | 13600752.45 |
| Methyl 9H-purine-6-carboxylate | 33130579.98 | 24761251.15 | 40613075.34 | 34553344.14 | 17737977.63 | 23670488.74 | 17396112.07 | 22198648.76 | 20353655.27 | 11611989.76 | 13534779.68 | 19937223.95 |
| Phenethyl decanoate | 12920276.64 | 9302995.253 | 7191053.711 | 8058896.833 | 13456381.14 | 15214912.6 | 10279666.94 | 8501011.859 | 8226313.483 | 8619517.895 | 8312919.609 | 16419979.25 |
| Daunomycinone | 21213.25707 | 69745.67276 | 0 | 57074.08905 | 5615903.507 | 4030912.966 | 3530430.48 | 3152798.395 | 5778412.565 | 2938785.384 | 6268534.731 | 6104523.526 |
| Rosmaricine | 4709167.737 | 4569894.041 | 4622679.153 | 4480871.936 | 4977885.542 | 5142708.651 | 4955335.815 | 5180113.784 | 4886230.447 | 4777360.394 | 4945160.114 | 4542247.441 |

| Metabolites | PBS_1 | PBS_2 | PBS_3 | PBS_4 | BCG_1 | BCG_2 | BCG_3 | BCG_4 | BCGΔ*pe18*_1 | BCGΔ*pe18*_2 | BCGΔ*pe18*_3 | BCGΔ*pe18*_4 |
|---|---|---|---|---|---|---|---|---|---|---|---|---|
| 5-[(4-Fluorophenoxy)methyl]-4-(prop-2-en-1-yl)-4H-1,2,4-triazole-3-thiol | 7268821.619 | 4631383.139 | 8225575.777 | 5238133.812 | 10166704.15 | 8072125.054 | 9499209.687 | 9092760.643 | 8505238.432 | 7948979.102 | 8363458.723 | 6417541.991 |
| Deoxyinosine | 4222347.735 | 4533390.401 | 6518497.822 | 4551827.379 | 5911642.454 | 3874676.054 | 6244268.08 | 4662693.813 | 3700338.903 | 4409063.78 | 5302739.746 | 3018754.815 |
| Tazarotenic acid | 14105396.49 | 12734010.13 | 24228301.42 | 16983839.53 | 17637033.52 | 16846255.61 | 17737798.4 | 16737912.91 | 16065547.96 | 12286700 | 14831713.51 | 14302105.31 |
| MGDG O-19:2_20:4 | 2385184.096 | 2265015.642 | 2146174.366 | 2283294.796 | 1478448.944 | 855534.4038 | 1776542.693 | 732621.1684 | 1840230.51 | 1707203.265 | 1732521.882 | 694690.2711 |
| Ropinirole | 2429788.852 | 2257289.92 | 1888604.882 | 2321697.569 | 2361015.189 | 2172844.636 | 3039679.641 | 2261817.595 | 2280970.297 | 2463682.07 | 2085404.625 | 2157189.819 |
| Persicachrome | 14327940.6 | 15938605.92 | 7963086.524 | 13058710.74 | 14092315.18 | 11400909.88 | 12435057.02 | 15709055.63 | 17322896.03 | 15506900.19 | 11692273.9 | 10769422.52 |
| Narasin | 21892895.52 | 25557830.48 | 27037161.87 | 24987343.02 | 20715476.31 | 17078418.85 | 25345480.67 | 26936532.45 | 17749243.34 | 26516608.81 | 21401240.32 | 14587515.65 |
| Trp-Leu | 702722.7309 | 772226.1181 | 831624.0389 | 682163.9269 | 817914.9989 | 596944.3265 | 909195.4429 | 712622.4182 | 950979.4898 | 1079092.781 | 1042658.201 | 558693.9351 |
| PI 18:0_20:3 | 9494050.063 | 1797198.976 | 7264039.9 | 4072430.553 | 2562042.347 | 66260.47026 | 2435902.796 | 462334.117 | 6906165.034 | 3959307.868 | 3205027.511 | 1747489.593 |
| trans-2-Ethoxy-5-(1-propenyl)phenol | 3958535.774 | 3584931.526 | 3404177.852 | 2803803.308 | 2565839.664 | 2698949.339 | 2595334.171 | 2445433.274 | 2680857.416 | 3088156.862 | 2880121.857 | 4807951.476 |
| Dihydroxyoctadecanoic acid | 7336628.369 | 8845039.124 | 10680230.76 | 8801982.961 | 8226459.171 | 10642540.31 | 8314431.585 | 9713074.166 | 7721563.519 | 6941117.199 | 9852136.162 | 8321023.676 |

| Metabolites | PBS_1 | PBS_2 | PBS_3 | PBS_4 | BCG_1 | BCG_2 | BCG_3 | BCG_4 | BCGΔ*pe18*_1 | BCGΔ*pe18*_2 | BCGΔ*pe18*_3 | BCGΔ*pe18*_4 |
|---|---|---|---|---|---|---|---|---|---|---|---|---|
| 2-Chloro-N-hexylacetamide | 1383727.645 | 1204211.124 | 982545.5703 | 1537850.348 | 935408.7646 | 1097820.42 | 2153231.814 | 889611.2396 | 1152394.572 | 1543314.536 | 940775.937 | 2944078.205 |
| N-Acetyl-arginine | 4357778.926 | 6013920.504 | 3277016.958 | 5892689.242 | 26751479.33 | 14304033.87 | 17853361.8 | 14910098.05 | 17607052.84 | 15088930.89 | 16071328.68 | 14472280.95 |
| 13-Keto-9Z,11E-octadecadienoic acid | 23383782.98 | 28543765.28 | 28746590.07 | 29758252.61 | 21468449.8 | 42231795.42 | 21269472.04 | 27511703.77 | 20775770.99 | 19806201.64 | 46293523.99 | 19638893.68 |
| M274T156 | 6069617.284 | 6353358.261 | 5215244.917 | 5530083.042 | 6286917.49 | 6114406.441 | 6156666.664 | 4791761.091 | 4686186.064 | 5209776.717 | 5295616.986 | 4421241.08 |
| 6-Nitro-2,3,4,9-tetrahydro-1H-carbazole | 1030149.258 | 600661.5344 | 1631711.787 | 1340425.068 | 1448146.159 | 1868068.84 | 787319.272 | 613755.3429 | 1427113.528 | 1047419.36 | 1748111.815 | 6654433.048 |
| PS(22:4(7Z,10Z,13Z,16Z)/20:3(8Z,11Z,14Z)) | 38742777.79 | 42024073.29 | 21852944.95 | 31270395.19 | 6961587.019 | 6522784.956 | 9553136.961 | 9017725.431 | 11056041.69 | 11901049.52 | 8214600.835 | 6626549.839 |
| Enniatin B1 | 3468052.191 | 2122419.286 | 2081424.553 | 2465009.893 | 2333303.647 | 2243820.225 | 2030927.16 | 1716613.736 | 2034137.98 | 2552188.965 | 2759807.637 | 1100377.32 |
| Pro-Trp | 7601439.106 | 7395963.785 | 7348446.859 | 7680022.64 | 7406797.722 | 7209052.108 | 7328437.612 | 7645114.098 | 7576438.504 | 7353994.068 | 7574906.091 | 7569138.417 |
| PC 8:0_30:7 | 5686082.625 | 5859200.317 | 3103774.179 | 4792004.204 | 17486198.61 | 17655645.86 | 16229343.76 | 16561540.67 | 20804971.57 | 20576805.73 | 14803973.82 | 15416976.14 |
| taspine | 744475.9893 | 798301.8146 | 681932.9868 | 563380.3985 | 642989.2866 | 528957.179 | 532623.6302 | 415792.6587 | 498243.9161 | 714412.9792 | 540153.0074 | 542198.5555 |
| 4-Methoxy-.alpha.-pyrrolidinopentioph | 10692222.44 | 12207126.02 | 9500174.071 | 10975214.03 | 10218421.75 | 10023992.49 | 9710961.272 | 9836882.756 | 9723588.231 | 9769716.805 | 9620174.024 | 10474478.52 |

| Metabolites | PBS_1 | PBS_2 | PBS_3 | PBS_4 | BCG_1 | BCG_2 | BCG_3 | BCG_4 | BCGΔ*pe18*_1 | BCGΔ*pe18*_2 | BCGΔ*pe18*_3 | BCGΔ*pe18*_4 |
|---|---|---|---|---|---|---|---|---|---|---|---|---|
| enone | | | | | | | | | | | | |
| LDGCC 18:1 | 41794204.79 | 42358220.19 | 37186670.33 | 51046998.87 | 46507023.23 | 43064401.94 | 42961582.36 | 41129052.29 | 39021222.07 | 36745550.98 | 43246975.98 | 34166666.34 |
| CROTONIC ACID | 951308.0973 | 892822.1362 | 1273304.82 | 1111735.61 | 1734523.953 | 1394348.016 | 1837707.324 | 1456136.519 | 1505726.146 | 1694015.18 | 1701451.423 | 1389791.48 |
| PA 3:0_13:1 | 870549.5384 | 1011033.783 | 829897.9977 | 645688.0131 | 521955.214 | 1281829.539 | 658423.7487 | 548769.5031 | 452843.8947 | 498958.5792 | 690903.9599 | 2422279.095 |
| 1-(2-Bromophenylsulfonyl)-4-methyl-1H-imidazole | 1797287.477 | 1875582.11 | 953531.071 | 809357.2982 | 1375468.002 | 1118595.945 | 960845.205 | 553768.0789 | 687063.5446 | 1824723.252 | 1094494.801 | 614981.0732 |
| Cer 29:2;3O/14:0;(2OH) | 2152773.588 | 1854130.327 | 1820436.077 | 1732468.767 | 773534.5101 | 1120856.64 | 1547919.329 | 1322233.21 | 1140636.077 | 1249743.103 | 950385.6816 | 630225.3589 |
| N-Benzyl-L-isoleucine methyl ester | 9434435.916 | 10022275.68 | 10289019.91 | 8416704.882 | 10914403.25 | 11244089.91 | 9028625.698 | 9659055.853 | 8172904.029 | 9617081.786 | 10129270.01 | 8138781.761 |
| Etilefrine | 4035409.621 | 4339074.879 | 3632562.36 | 4115083.547 | 3831241.913 | 3957000.848 | 5232096.079 | 3895150.06 | 4110566.576 | 3982353.142 | 3883099.001 | 4044862.988 |
| Melibiose | 9323.637144 | 25329.23171 | 4599.533498 | 4659.405172 | 1384845.161 | 1257705.463 | 1229426.906 | 901894.675 | 2700593.234 | 1722700.875 | 418484.0459 | 1211222.305 |
| Galeterone | 1157058.705 | 2618126.329 | 1826342.249 | 2040037.201 | 1846132.178 | 1915471.261 | 1652884.503 | 1813922.568 | 931776.2845 | 1789390.296 | 1780050.796 | 1725999.186 |
| LPI(18:0) | 12587.63302 | 315804.2904 | 308758.3186 | 283552.4399 | 30283.62757 | 447935.1545 | 710738.0781 | 25505.24455 | 137351.817 | 28456.55003 | 315370.0579 | 603831.1697 |

| Metabolites | PBS_1 | PBS_2 | PBS_3 | PBS_4 | BCG_1 | BCG_2 | BCG_3 | BCG_4 | BCGΔ*pe18*_1 | BCGΔ*pe18*_2 | BCGΔ*pe18*_3 | BCGΔ*pe18*_4 |
|---|---|---|---|---|---|---|---|---|---|---|---|---|
| PE 20:4_22:5 | 1330653.935 | 926950.6664 | 1189759.833 | 1260849.72 | 1125310.505 | 844232.4334 | 1151112.033 | 1331433.348 | 1556009.599 | 1258139.107 | 465336.3727 | 1749206.013 |
| N-Cyclohexylformamide | 3138331.554 | 2329095.794 | 2549261.756 | 2205935.496 | 3464666.338 | 2018666.381 | 2216828.685 | 1794856.696 | 2516047.113 | 2644287.708 | 2935386.573 | 5369829.982 |
| FGIN 1-27 | 9259236.612 | 4738741.288 | 7821745.202 | 5843168.503 | 10746888.5 | 10101939.22 | 11210219.13 | 11313798.15 | 8183847.836 | 8733225.131 | 13278093.43 | 3961351.737 |
| 3-Methylcyclohexanethiol | 415511.179 | 502661.6298 | 716309.897 | 490831.066 | 3145510.984 | 2323734.714 | 3235405.52 | 2745009.181 | 2166261.965 | 2415031.659 | 2463073.079 | 2022842.193 |
| N-Methyl-L-proline | 8826780.616 | 8464861.89 | 13647962.34 | 9466729.437 | 52562715.44 | 39029377.27 | 50458325.9 | 42094627.38 | 35124634.17 | 37349656.18 | 42047715.25 | 35052453.96 |
| CAR 15:0 | 7713673.976 | 7060094.786 | 9642953.526 | 7560589.826 | 26374027.68 | 16372334.28 | 24093081.57 | 22486203.48 | 15113702.91 | 21111460.16 | 22139995.4 | 12704487.11 |
| Corypalline | 10552252.41 | 12193993.62 | 9040362.926 | 10951759.17 | 11006562.96 | 9517476.829 | 11014341.01 | 10050817.44 | 10379546.32 | 10720436.08 | 10000651.35 | 8515831.785 |
| 1-(linoleoyl)-sn-glycero-3-phosphoethanolamine | 257374.615 | 246669.4926 | 371960.5944 | 214403.52 | 768485.534 | 629312.5145 | 717511.3435 | 750912.5294 | 762327.1758 | 780043.4784 | 773361.4744 | 718176.0693 |
| Docosahexaenoic acid (DHA) | 98333635.27 | 101041478.1 | 111785686.3 | 98828074.91 | 53076075.05 | 77505121.68 | 37929881.46 | 65789183.08 | 52927848.07 | 37028227.55 | 44535274.58 | 57014361.09 |
| 7,16,17-Trihydroxydocosa-4,8,10,12,14,19-hexaenoylcarnitine | 36248719.08 | 37415075.94 | 33820883.68 | 35540745.54 | 33662937.89 | 33004771.21 | 37320684.15 | 32337037.17 | 32812969.73 | 31301824.09 | 33344673 | 33380452.14 |
| LPA 11:0 | 211713. | 145928. | 282145. | 254631. | 348327 | 265342 | 311232 | 288606 | 274189 | 344357 | 272682 | 218620 |

| Metabolites | PBS_1 | PBS_2 | PBS_3 | PBS_4 | BCG_1 | BCG_2 | BCG_3 | BCG_4 | BCGΔ*pe18*_1 | BCGΔ*pe18*_2 | BCGΔ*pe18*_3 | BCGΔ*pe18*_4 |
|---|---|---|---|---|---|---|---|---|---|---|---|---|
| | 5913 | 9382 | 7313 | 9939 | 8.617 | 3.752 | 9.549 | 7.869 | 9.704 | 0.649 | 6.857 | 0.112 |
| Cer 9:0;2O/3:0 | 3251695.058 | 9432545.921 | 1200857.759 | 4785354.712 | 7483004.122 | 3703061.9 | 1354177.688 | 2114274.238 | 2432295.566 | 3619260.27 | 3304021.154 | 2955039.117 |
| 2,2-Dimethyl-N-[3-(trifluoromethyl)phenyl]propanamide | 386106.4549 | 311689.4069 | 382877.845 | 379582.8492 | 2910825.54 | 1541161.374 | 1891025.994 | 1919368.181 | 1934764.166 | 1523793.955 | 1305394.838 | 2719843.723 |
| DGCC 8:0_9:0 | 1045220.084 | 897113.0507 | 840602.0499 | 797617.0252 | 898645.7949 | 748144.1397 | 865055.7336 | 928310.5561 | 882293.8773 | 898864.2615 | 842886.9914 | 710400.2748 |
| (4-Isobutylphenyl)acetic acid | 1331848.727 | 831434.4344 | 950559.7418 | 9538355.538 | 312512.5536 | 3043279.953 | 13509198.82 | 167025.6207 | 4991965.399 | 4787711.242 | 344030.7096 | 8904715.457 |
| MGDG 3:0_22:1 | 53893938.27 | 69095638.34 | 48053821.66 | 52374912.27 | 56408146.69 | 50718319.44 | 53427033.56 | 63056747.82 | 50810487.46 | 55339116.25 | 56320357.76 | 57150952.48 |
| CerP 15:0;2O/10:0 | 2127921.565 | 3003374.984 | 1158800.042 | 1433042.722 | 855283.2906 | 1372825.687 | 1035843.416 | 904858.88 | 811532.3114 | 1018028.874 | 970961.5406 | 575141.4441 |
| Glycine | 270632.7768 | 844776.7663 | 543543.8479 | 292874.9573 | 404451.9661 | 324398.0951 | 326642.2121 | 296833.2189 | 469116.5753 | 332957.5901 | 342394.3745 | 223490.1331 |
| diazoxide | 1206110.516 | 1460935.916 | 1863926.26 | 1751227.781 | 1755370.068 | 2194677.516 | 1409833.778 | 1593416.848 | 1638251.908 | 1569160.123 | 2338852.659 | 1805013.736 |
| SPB 14:0;2O | 11405326.42 | 11730318.86 | 10399919.54 | 11684047.02 | 11434020.8 | 11021006.14 | 11053376.63 | 11293721.36 | 11425207.32 | 11364276.14 | 11355166.25 | 10339145.96 |
| IS6 | 329654618.9 | 314734114.7 | 294763754.4 | 355081192 | 300449968 | 307212598.9 | 388431420.7 | 364276387.8 | 304768277.4 | 351118878.3 | 385210058.7 | 317185563 |
| PI 12:0_24:2 | 3475330.348 | 2032228.358 | 3183352.957 | 2882760.143 | 2756999.794 | 15022.85721 | 2216766.529 | 1010775.483 | 3343095.58 | 2630405.095 | 2292428.02 | 864553.655 |

| Metabolites | PBS_1 | PBS_2 | PBS_3 | PBS_4 | BCG_1 | BCG_2 | BCG_3 | BCG_4 | BCGΔ*pe18*_1 | BCGΔ*pe18*_2 | BCGΔ*pe18*_3 | BCGΔ*pe18*_4 |
|---|---|---|---|---|---|---|---|---|---|---|---|---|
| Tafluprost ethylamide | 4043024.494 | 4735790.737 | 4389077.269 | 3826975.837 | 3849537.859 | 3873442.888 | 3961842.719 | 4073066.033 | 3857936.751 | 3862722.882 | 3821882.799 | 4519225.102 |
| N-[2-Chloro-5-(trifluoromethyl)phenyl]-3,4,5-trimethoxybenzamide | 556451.0819 | 420423.3689 | 424405.5315 | 363724.0802 | 648227.8937 | 614941.4722 | 921293.1647 | 868214.1236 | 750372.7536 | 749531.3437 | 652967.6942 | 653783.9616 |
| NAGly 26:7/22:5 | 1393006.562 | 1330032.271 | 212573.2917 | 1041797.132 | 831878.9315 | 514868.2346 | 823268.1221 | 726306.4313 | 896929.1996 | 947468.3272 | 925322.4392 | 597978.3757 |
| DGDG O-9:0_15:0 | 17335.75477 | 0 | 0 | 16925.82209 | 4734277.868 | 4362951.277 | 4816185.134 | 3608501.167 | 3885687.778 | 3854874.313 | 4857052.186 | 3398238.031 |
| MG 9:0 | 669945.6006 | 772961.1997 | 726586.3637 | 684252.7627 | 909074.6051 | 789754.4682 | 991988.348 | 827467.0552 | 816121.5809 | 769253.7145 | 836514.4503 | 782171.5021 |
| 5-[2-(4-Chlorophenyl)hydrazin-1-ylidene]-2,2-dimethyl-1,3-dioxane-4,6-dione | 55137150.37 | 35351081.8 | 71371554.9 | 44668837.08 | 39116928.48 | 41068022.36 | 34884378.95 | 37802565.61 | 35397464.64 | 7733917.305 | 39629627.4 | 36975153.63 |
| S-Adenosylmethionine | 1088455.044 | 857545.8313 | 1092456.436 | 880663.7713 | 12756045.78 | 10127393.75 | 12544691.46 | 10429474.06 | 10643516.07 | 9811131.972 | 11578010.37 | 11817101.83 |
| 4-Hydroxy-3-[(2E,6E)-4-hydroxy-3,7,11-trimethyldodeca-2,6,10-trienyl]benzoic acid | 3629438.992 | 4362849.365 | 971643.0014 | 3853175.495 | 3469583.405 | 3588463.434 | 3556599.778 | 3604931.473 | 3954760.141 | 3147948.651 | 3430569.643 | 5080961.647 |

| Metabolites | PBS_1 | PBS_2 | PBS_3 | PBS_4 | BCG_1 | BCG_2 | BCG_3 | BCG_4 | BCGΔ*pe18*_1 | BCGΔ*pe18*_2 | BCGΔ*pe18*_3 | BCGΔ*pe18*_4 |
|---|---|---|---|---|---|---|---|---|---|---|---|---|
| N-(1,3-Thiazol-2-yl)benzenesulfonamide | 12642917.32 | 12851134.72 | 15688116.81 | 12950917.43 | 17501538.44 | 13186599.38 | 15537776.82 | 15272813.12 | 13311109.62 | 14157196.21 | 13888889.68 | 12943035.33 |
| FA 28:7 | 666361.1673 | 724231.6037 | 339029.2425 | 645406.9342 | 527459.7453 | 423611.1485 | 465358.3051 | 501045.89 | 469849.8719 | 659481.2609 | 531501.1899 | 290963.2182 |
| SM 39:4;3O | 1747247.131 | 1255162.682 | 528530.83 | 573294.1336 | 508687.9281 | 84832.53158 | 711591.7213 | 2112028.48 | 568727.766 | 715961.5168 | 502337.6437 | 1317645.499 |
| PC O-19:0 | 2154743.008 | 2112375.858 | 1080622.984 | 1688763.512 | 7324902.508 | 12645454.69 | 7111158.161 | 7140833.189 | 7791558.425 | 10777761.23 | 7760338.399 | 8075028.256 |
| 4-Hydroxycinnamic acid | 3020025.158 | 1901113.13 | 3072897.948 | 2809204.703 | 2352956.531 | 2272980.563 | 2313933.566 | 1983547.09 | 2144999.24 | 1759846.436 | 2651770.808 | 3333727.07 |
| PS 14:1_24:4 | 2320664.998 | 2567025.2 | 2523325.335 | 2772824.215 | 2318652.332 | 1474579.669 | 2356485.927 | 2491755.482 | 4351790.586 | 2958500.541 | 2090479.099 | 2879686.837 |
| tert-Butyl 4-(2-aminoethyl)-1-piperazinecarboxylate | 24199320.91 | 20304280.98 | 26858542.52 | 24144274.45 | 54132762.78 | 41078906.05 | 52278912.71 | 44540088.63 | 35443066.32 | 35651437.47 | 45534298.21 | 31821524.78 |
| 4,4,6-Trimethyl-1-(2-naphthyl)-3,4-dihydro-2(1H)-pyrimidinethione | 856855.9551 | 739985.1287 | 1191630.35 | 832734.7219 | 3912954.742 | 3496397.587 | 3374928.951 | 3046118.13 | 2450411.02 | 1954611.763 | 2180920.814 | 6402680.895 |
| Dropropizine | 5532568.969 | 7893224.524 | 5264433.07 | 7336798.119 | 5501625.906 | 5571871.928 | 7560587.584 | 5773776.896 | 5573930.176 | 5723611.68 | 5865703.091 | 5132891.922 |
| 3-Hydroxy-4-amino | 541978 | 445599 | 601650 | 690062 | 393458 | 561139 | 482822 | 624514 | 487338 | 359598 | 448844 | 523762 |

| **Metabolites** | **PBS_1** | **PBS_2** | **PBS_3** | **PBS_4** | **BCG_1** | **BCG_2** | **BCG_3** | **BCG_4** | **BCGΔ*pe18*_1** | **BCGΔ*pe18*_2** | **BCGΔ*pe18*_3** | **BCGΔ*pe18*_4** |
|---|---|---|---|---|---|---|---|---|---|---|---|---|
| pyridine | 0.948 | 4.21 | 4.438 | 6.797 | 4.583 | 8.704 | 6.16 | 8.539 | 1.826 | 6.029 | 8.021 | 3.399 |
| PC O-20:2 | 3268250.07 | 533756.2021 | 4791380.554 | 3263535.127 | 5781330.023 | 6292763.326 | 5789380.791 | 4783628.612 | 5230211.092 | 5884174.214 | 8263722.127 | 5216374.037 |
| Lys-Leu | 154586389.9 | 133471571.4 | 176124893.3 | 155775225.3 | 177117813.9 | 119754840.1 | 188540767.7 | 148991038.3 | 118615773.5 | 125205803.3 | 137537682.6 | 107445125 |
| 2'-Deoxyguanosine 5'-monophosphate (dGMP) | 1115721.737 | 1301832.009 | 1328538.498 | 1193063.56 | 2249605.421 | 2022044.223 | 1733065.354 | 6149874.528 | 2337698.62 | 1988248.408 | 2586439.257 | 2911919.118 |
| 4-Bromo-1-methyl-1H-indazole | 2845132.32 | 2232233.144 | 4445575.862 | 2474593.504 | 8927588.201 | 6538230.941 | 8209581.398 | 7112427.756 | 6924183.525 | 6285633.923 | 5331982.454 | 5695617.582 |
| PC O-20:3 | 4200895.774 | 4128526.889 | 6318336.006 | 3592594.412 | 9490084.933 | 8467900.005 | 8335722.278 | 8340105.492 | 7300712.427 | 7675244.279 | 9568129.819 | 6942114.855 |
| 1,2-Diarachidonoyl-sn-glycero-3-phosphocholine | 39797547.31 | 40823056.78 | 22124469.42 | 30333602.65 | 17575062.04 | 12336191.57 | 16883194.23 | 16525805.42 | 21313695.12 | 22361761.83 | 15957548.34 | 15565632.03 |
| Butyl levulinate | 17919726.03 | 2474526.446 | 9528021.785 | 18918413.81 | 2194254.426 | 2589355.084 | 2861801.626 | 2675653.841 | 2368317.014 | 5326451.389 | 2419925.944 | 3844239.34 |
| 3-Methylthiopropionic acid | 6180133.54 | 7306544.106 | 5370723.957 | 6990415.689 | 7257644.767 | 9812431.684 | 7008348.642 | 7523060.657 | 5853373.777 | 6860521.904 | 7836297.467 | 8082209.495 |
| Arachidonic acid (AA) | 242531305.7 | 224285813.5 | 385687118.2 | 208242919.7 | 60110875.17 | 94382525.72 | 83883256.31 | 80230334.44 | 69234691.41 | 55539894.68 | 72873708.02 | 62561595.27 |
| CAR 12:1 | 2872724.365 | 3134517.268 | 3554108.58 | 3241674.139 | 3079924.605 | 2979204.597 | 3097655.596 | 3012230.688 | 3029528.313 | 2970832.81 | 3031606.936 | 3880918.741 |

| Metabolites | PBS_1 | PBS_2 | PBS_3 | PBS_4 | BCG_1 | BCG_2 | BCG_3 | BCG_4 | BCGΔ*pe18*_1 | BCGΔ*pe18*_2 | BCGΔ*pe18*_3 | BCGΔ*pe18*_4 |
|---|---|---|---|---|---|---|---|---|---|---|---|---|
| 1-Palmitoyl-2-docosahexaenoyl-sn-glycero-3-phosphocholine | 95983240.97 | 108231486 | 51835996.8 | 100212200.2 | 101406023.1 | 116424591.7 | 96373716.61 | 124010588.7 | 110056538.5 | 107759570.8 | 99335125.48 | 84081648.87 |
| Ethyl 3-chloro-1H-indole-2-carboxylate | 4546785.119 | 3367609.017 | 4159469.773 | 3105095.719 | 2503485.979 | 4013058.136 | 1999807.325 | 2229477.563 | 2519488.144 | 1537105.727 | 1991809.636 | 2594832.906 |
| Methyl hydrogen azelate | 4137878.534 | 4930402.678 | 5222309.372 | 5354135.891 | 4371105.44 | 7226228.45 | 5489413.19 | 4370555.08 | 4664403.503 | 4207537.4 | 4786468.604 | 6441749.989 |
| LPE 24:5 | 235503.1924 | 258943.897 | 826301.8878 | 673463.5406 | 655631.3167 | 708686.6034 | 600783.8588 | 615979.694 | 755819.0708 | 677369.7283 | 646106.2214 | 533262.4763 |
| SM 25:1;2O/30:5 | 226460.3237 | 99728.3812 | 268346.9796 | 252600.118 | 0 | 0 | 0 | 39769.21461 | 83257.16505 | 92456.58522 | 40879.10959 | 131768.2943 |
| LPA 16:1 | 589552.7056 | 729446.5502 | 568366.6079 | 654879.9959 | 1630449.609 | 1519816.525 | 1673576.927 | 1348586.056 | 1113601.245 | 1104523.124 | 1463527.42 | 1150591.817 |
| 1-Ethyl-3-methylimidazolium cation | 10866197.25 | 9895715.364 | 20256640.87 | 13312017.7 | 22752036.43 | 13743115.05 | 5554230.818 | 22070865.22 | 20628150.59 | 14668344.18 | 17500587.28 | 22706788.71 |
| Mepirizole | 1641973.132 | 1625891.427 | 1930949.777 | 1713273.434 | 4763007.422 | 3131889.437 | 5212747.948 | 4604119.034 | 4962462.408 | 5274031.998 | 4736784.462 | 3300254.205 |
| 2'-Hydroxy-4'-methylacetophenone | 2237953.471 | 2071438.942 | 2308548.613 | 1880807.729 | 2538795.677 | 2579975.376 | 2396801.463 | 2750267.283 | 2410171.615 | 2081644.404 | 2695592.331 | 2479502.749 |
| dehydrohomospermidine | 9845443.088 | 13955075.38 | 10381216.47 | 10302567.45 | 61969876.64 | 12918095.5 | 47023099.43 | 54667563.22 | 58132114.03 | 60623501.35 | 8780316.297 | 10652025.85 |
| 2-Acetylthiazole | 199500 | 148564 | 239933 | 177926 | 377225 | 261953 | 418248 | 325812 | 236937 | 300432 | 322815 | 203892 |

| Metabolites | PBS_1 | PBS_2 | PBS_3 | PBS_4 | BCG_1 | BCG_2 | BCG_3 | BCG_4 | BCGΔ*pe18*_1 | BCGΔ*pe18*_2 | BCGΔ*pe18*_3 | BCGΔ*pe18*_4 |
|---|---|---|---|---|---|---|---|---|---|---|---|---|
| | 13.02 | 42.74 | 74.41 | 27.78 | 97.1 | 92.23 | 01.35 | 17.31 | 58.94 | 98.16 | 58.8 | 48.79 |
| Miglustat | 292028.6588 | 3512389.372 | 6883481.189 | 4469622.96 | 421621.6801 | 417106.4528 | 662709.2248 | 289818.2161 | 446524.6937 | 687583.4326 | 396951.9376 | 10601389.79 |
| SL 13:2;O/18:5 | 5445696.794 | 5371935.775 | 6913121.406 | 4680119.483 | 7770452.978 | 5468728.758 | 5271906.703 | 5577657.934 | 5071665.751 | 5463067.502 | 5336358.381 | 4367731.547 |
| 11.beta.,16.alpha.,17.alpha.,21-Tetrahydroxypregna-1,4-diene-3,20-dione | 4390614.403 | 4581754 | 4049354.943 | 4417764.812 | 4389826.582 | 4397401.36 | 4238493.333 | 4408941.131 | 4203306.949 | 4251354.78 | 4419536.075 | 4039097.379 |
| PE(P-18:0/22:6(4Z,7Z,10Z,13Z,16Z,19Z)) | 6828771.277 | 7339860.408 | 6346638.647 | 7515775.951 | 6449672.673 | 5828301.952 | 6041749.887 | 5884445.121 | 7242999.198 | 6421222.193 | 5830907.547 | 5130870.698 |
| SM 34:1;2O | 1939600.026 | 2557197.298 | 3573335.163 | 2278247.535 | 3022028.8 | 27778.74388 | 3781856.586 | 2559785.331 | 2123192.462 | 2927015.879 | 3324777.27 | 1788217.979 |
| 1H-3-Benzazepine-7,8-diol, 6-chloro-2,3,4,5-tetrahydro-1-phenyl- | 3233667.433 | 2752609.717 | 2308471.677 | 2735761.569 | 6343555.763 | 3502163.822 | 5609119.893 | 4805383.26 | 5791944.318 | 4476293.146 | 2732513.408 | 3792403.366 |
| PC O-15:0 | 10993344.53 | 10326842.07 | 12055519.81 | 11245535.46 | 24600945.86 | 20371568.65 | 20570006.39 | 20587711.8 | 18247121.69 | 22360428.66 | 21348251.69 | 18409110.93 |
| PS(14:0/22:0) | 896858.5277 | 1129554.666 | 903167.0545 | 1186785.699 | 13119310.55 | 7508517.404 | 13789193.04 | 8323283.646 | 8093799.237 | 13975888.3 | 12551367.88 | 5634605.104 |
| LPA 16:3 | 1322242.381 | 2318483.745 | 1041688.244 | 775106.5214 | 339579.1696 | 1182499.219 | 478063.6161 | 682018.7143 | 511244.3382 | 455242.8744 | 844942.9244 | 4735331.78 |

| Metabolites | PBS_1 | PBS_2 | PBS_3 | PBS_4 | BCG_1 | BCG_2 | BCG_3 | BCG_4 | BCGΔ*pe18*_1 | BCGΔ*pe18*_2 | BCGΔ*pe18*_3 | BCGΔ*pe18*_4 |
|---|---|---|---|---|---|---|---|---|---|---|---|---|
| Cyclic adenosine diphosphate-ribose (cADPR) | 991391.4554 | 861925.399 | 979590.2721 | 823911.5953 | 2744034.804 | 1932065.358 | 2532831.875 | 2439902.02 | 2364751.501 | 2115667.681 | 2241710.579 | 2429015.49 |
| SM 23:1;2O/28:1 | 44415.6353 | 74717.47663 | 53735.05976 | 39125.1262 | 555510.9192 | 25967.03445 | 596217.253 | 710427.381 | 293455.7599 | 215215.4311 | 602435.6123 | 435468.3182 |
| 2-Amino-3-methoxybenzoic acid | 5176851.656 | 4648738.599 | 6557563.778 | 5616309.029 | 6024639.165 | 6097639.891 | 5111813.842 | 5074695.593 | 5809652.524 | 4491018.627 | 4184200.499 | 8652173.889 |
| NAE 20:1 | 43756785.96 | 44930007.41 | 29969342.64 | 37696898.8 | 31188357.37 | 45437290.5 | 37229222.56 | 35420221.5 | 37482259.15 | 35453083.45 | 35223458.9 | 33699882.78 |
| Tetradecylamine | 7725586.252 | 10647359.32 | 8106818.118 | 11228396.88 | 11190744.83 | 10620927.53 | 9217551.418 | 9641128.06 | 9684040.696 | 8689486.922 | 8141511.929 | 10435139.59 |
| 4-(4-Nitrophenyl)-1H-imidazole | 4291184.55 | 3604432.532 | 3888926.627 | 4200754.482 | 3765135.043 | 3921872.517 | 3110772.482 | 3831173.222 | 3520333.712 | 2293611.146 | 3796107.581 | 3964478.631 |
| PEtOH 19:2_22:4 | 6587155.221 | 4929605.367 | 8583936.173 | 7758608.91 | 5837029.716 | 4934698.055 | 4399133.093 | 7572319.394 | 6163724.134 | 6636546.314 | 1149285.899 | 7526690.455 |
| Lactosylceramide_(d18:1/12:0) | 929450.6261 | 973424.8703 | 428584.6672 | 743258.9256 | 1153882.097 | 835357.6234 | 1218089.057 | 860767.9069 | 837514.0181 | 774151.1074 | 1060110.087 | 668599.4759 |
| N1,N2-Diisobutylethanediamide | 51559497.83 | 56779892.76 | 32453420.62 | 50620715.07 | 58513161.75 | 41230930.55 | 71908287.69 | 66632341.04 | 57791990.84 | 54626640.54 | 50652383.8 | 47406484.84 |
| 4,6-Quinazolinediamine, N4-[2-(4-phenoxyphenyl)ethyl]- | 1779587.179 | 1926615.741 | 1853286.511 | 1797858.276 | 1889557.893 | 1923502.156 | 1841783.012 | 1971250.834 | 1952733.513 | 1934875.481 | 1948524.195 | 1852241.224 |

| Metabolites | PBS_1 | PBS_2 | PBS_3 | PBS_4 | BCG_1 | BCG_2 | BCG_3 | BCG_4 | BCGΔ*pe18*_1 | BCGΔ*pe18*_2 | BCGΔ*pe18*_3 | BCGΔ*pe18*_4 |
|---|---|---|---|---|---|---|---|---|---|---|---|---|
| 9Z,11E,13E-Octadecatrienoic acid | 64643909.82 | 56710535.3 | 35849313.05 | 59830993.86 | 50531062.27 | 47977378.26 | 40620793.15 | 48192712.03 | 44463664.98 | 52441148.18 | 41802355.79 | 35972709.32 |
| 3,5-Di-tert-butyl-4-hydroxybenzoic acid | 3100296.144 | 3218610.655 | 3520251.345 | 3463482.245 | 4110776.183 | 3121366.452 | 3111619.296 | 3266833.966 | 3632293.349 | 3838212.995 | 3447658.142 | 3612426.946 |
| Oligomycin C | 12922234.17 | 12003388.07 | 12311074.77 | 12972807.51 | 15563952.25 | 19258039.33 | 15870369.98 | 15895090 | 14511934.86 | 17072923.74 | 14277148.94 | 15514170.53 |
| Acridine orange 10-nonyl cation | 2621147.987 | 2927671.469 | 1882550.074 | 2553531.871 | 2239701.405 | 3514147.551 | 2739214.965 | 2583919.046 | 2186211.463 | 2411677.845 | 2841990.544 | 2337430.208 |
| LDGTS 10:0 | 2506891.645 | 2640494.015 | 2373817.459 | 2553021.516 | 2567026.063 | 2486915.135 | 2652200.737 | 2457329.152 | 2727122.985 | 2678536.909 | 2528714.596 | 2478457.937 |
| 6-Benzylaminouracil | 1138235.169 | 995694.0046 | 4440335.537 | 1404124.661 | 1611290.746 | 3290982.445 | 1382534.892 | 1784595.672 | 1595514.928 | 1131110.917 | 1497913.207 | 2083657.782 |
| MGDG O-16:2_6:0 | 91439787.63 | 95306117.28 | 77591944.42 | 83701537.87 | 95502570.62 | 98212123.11 | 93616859.25 | 84119553.28 | 88569282.26 | 94522760.77 | 87295274.18 | 91588565.37 |
| PI 25:5 | 47643.55336 | 47810.74539 | 31126.27895 | 66717.07553 | 56365.33803 | 38722.2398 | 18260.03016 | 18660.16152 | 51238.09253 | 39411.74594 | 36208.43672 | 90823.62983 |
| PC O-35:7 | 30529258.02 | 32443862.64 | 30174753.63 | 25490755.15 | 36373176.92 | 35743226.88 | 29982922.2 | 38612110.91 | 31289663.64 | 26544374.89 | 29580221.11 | 27182472.49 |
| Fucose 1-phosphate | 889156.2069 | 723189.738 | 1512172.573 | 977605.1085 | 3092276.844 | 1762065.565 | 2090948.776 | 2084914.053 | 2300598.899 | 2020145.227 | 3184863.248 | 3292517.213 |
| 1,3-Oxazinan-2-imine | 763729.8628 | 1558816.311 | 2047027.51 | 1818202.621 | 994796.751 | 859766.5783 | 1147408.113 | 1263620.834 | 1125924.693 | 11971764.27 | 1125563.067 | 6107003.882 |
| .beta.-Peltatin | 283830 | 288478 | 310050 | 394169 | 273832 | 391125 | 321453 | 393156 | 307699 | 447536 | 261890 | 356515 |

| Metabolites | PBS_1 | PBS_2 | PBS_3 | PBS_4 | BCG_1 | BCG_2 | BCG_3 | BCG_4 | BCGΔ*pe18*_1 | BCGΔ*pe18*_2 | BCGΔ*pe18*_3 | BCGΔ*pe18*_4 |
|---|---|---|---|---|---|---|---|---|---|---|---|---|
| | 2.566 | 3.675 | 7.99 | 8.13 | 9.303 | 0.42 | 1.63 | 7.349 | 6.753 | 6.025 | 4.777 | 1.138 |
| 2-(2,6-Dimethylmorpholin-4-yl)ethanol | 12277325.58 | 10415335.94 | 18002714.03 | 11770249.61 | 14969252.83 | 11148827.28 | 22671378.97 | 14235015.19 | 10381771.49 | 15622786.81 | 16957860.08 | 9601694.671 |
| Prostaglandin G2 2-glyceryl Ester | 5440043.884 | 5782033.329 | 5730160.659 | 5297226.15 | 4273498.128 | 5077898.91 | 4842516.643 | 5249939.044 | 3228986.617 | 4937687.146 | 4512846.804 | 6420735.471 |
| 2-PHENYLACETAMIDE | 2539792.623 | 2246149.115 | 3347351.249 | 2669814.639 | 4432633.196 | 3719678.14 | 4258838.554 | 4349383.16 | 4220555.287 | 3262046.851 | 3243260.899 | 3760329.277 |
| 2-(4-Amino-1-piperidinyl)ethanol | 2753990.619 | 2798332.864 | 2622162.861 | 2569311.019 | 2632132.127 | 2807644.717 | 3014338.353 | 2683630.385 | 2383036.808 | 3013630.173 | 3040280.044 | 2073973.109 |
| PC(18:1(9Z)/P-16:0) | 14803236.94 | 57593569.34 | 12723571.22 | 10398501.37 | 86711414.83 | 33246671.19 | 87741869.24 | 56844128.05 | 38952093.06 | 69104228.32 | 69941956.2 | 22199936.98 |
| 5-{2-[2-(1H-Tetraazol-5-yl)ethoxy]ethyl}-1H-tetraazole | 773423.456 | 741689.585 | 535275.1803 | 605086.425 | 667343.8373 | 867015.7009 | 709158.1589 | 956721.2268 | 662962.2388 | 799877.678 | 589006.8159 | 589162.8619 |
| Oxamniquine | 3561443.65 | 3171879.803 | 3610911.748 | 2751092.185 | 5706890.353 | 7429705.579 | 5766958.195 | 5330154.602 | 5330342.953 | 4019421.187 | 4883331.109 | 5022419.01 |
| DGGA 13:0_20:1 | 194546.4231 | 121535.7593 | 236096.7775 | 206396.4013 | 1626300.851 | 2591857.153 | 1782785.702 | 1388679.391 | 1618967.265 | 1559666.961 | 1729829.458 | 1168183.484 |
| Symmetric dimethylarginine | 329109.0587 | 396230.4385 | 343038.6291 | 559289.9953 | 296669.9807 | 295018.2956 | 511556.9431 | 263359.5623 | 329286.7537 | 391296.4301 | 302401.4527 | 657682.256 |
| 3-(4-tert-Butylphenyl)propanoic acid | 6318793.708 | 6308162.552 | 6320342.617 | 6598049.089 | 6204665.488 | 6087011.22 | 6166006.377 | 6729325.054 | 6103882.58 | 6031778.343 | 6223855.22 | 6299687.977 |
| Glu-His | 281629.7618 | 171066.3094 | 582384.8119 | 215571.7405 | 2759151.655 | 1913451.403 | 2398170.102 | 1956764.974 | 2177096.3 | 2177967.34 | 1651395.523 | 1920713.697 |

| **Metabolites** | **PBS_1** | **PBS_2** | **PBS_3** | **PBS_4** | **BCG_1** | **BCG_2** | **BCG_3** | **BCG_4** | **BCGΔ*pe18*_1** | **BCGΔ*pe18*_2** | **BCGΔ*pe18*_3** | **BCGΔ*pe18*_4** |
|---|---|---|---|---|---|---|---|---|---|---|---|---|
| 2-Chloro-N-(1-cyanocyclopentyl)acetamide | 1771880.717 | 1732886.531 | 1391295.387 | 1289039.13 | 1262884.906 | 1363879.761 | 1260067.429 | 1198188.733 | 1240924.901 | 1305665.18 | 1328698.954 | 1633826.903 |
| 3-Hydroxy-2-((9Z,12Z)-octadeca-9,12-dienoyloxy)propyl 2-(trimethylazaniumyl)ethyl phosphate | 1897405.939 | 1134948.564 | 2778015.662 | 2029188.368 | 6962257.925 | 6419880.879 | 2596379.079 | 6575361.523 | 6942784.53 | 5893134.739 | 7329662.42 | 5511840.482 |
| (1Z)-1-(3-Ethyl-5-hydroxy-1,3-benzothiazol-2-ylidene)acetone | 1357051.186 | 1077849.537 | 2164937.965 | 1479154.811 | 2647571.465 | 2320403.391 | 2542971.565 | 2501713.891 | 2426032.143 | 1566081.254 | 1602790.871 | 2210582.063 |
| 8-Methyl-2-(trifluoromethyl)quinolin-4-ol | 1082261.047 | 790001.0534 | 1246086.789 | 814914.3258 | 1593755.189 | 983665.5426 | 1171098.002 | 1190464.294 | 1123012.419 | 972457.775 | 1040366.547 | 788859.5849 |
| SM(d18:1/18:0) | 11732540.09 | 11810557.7 | 18114292.71 | 10390237.29 | 11920102.36 | 10506568.76 | 18295371.23 | 12930688.95 | 13454014.7 | 17525592.99 | 15331203.53 | 6749206.982 |
| PS 2:0_16:2 | 586549.0954 | 637767.5003 | 353931.2128 | 326941.4971 | 410572.7711 | 293610.5684 | 568016.2251 | 479549.1887 | 631335.0697 | 366051.2469 | 317182.5393 | 445653.1233 |
| Uridine diphosphate glucose (UDP-glucose) | 376869.1525 | 418769.3176 | 523284.4217 | 410241.5669 | 1999079.733 | 1273707.858 | 1998778.231 | 1622996.324 | 1668910.138 | 2072308.57 | 1832264.393 | 2021528.689 |

| Metabolites | PBS_1 | PBS_2 | PBS_3 | PBS_4 | BCG_1 | BCG_2 | BCG_3 | BCG_4 | BCGΔ *pe18*_1 | BCGΔ *pe18*_2 | BCGΔ *pe18*_3 | BCGΔ*pe18*_4 |
|---|---|---|---|---|---|---|---|---|---|---|---|---|
| PG 20:2_22:4 | 2593320.847 | 3109461.576 | 1667116.278 | 2020069.415 | 1948159.781 | 1174897.65 | 1394412.086 | 1542288.662 | 2126700.46 | 3236121.511 | 1984703.694 | 988816.3502 |
| Geneticin | 21616689.83 | 20679032.9 | 19146295.93 | 19315954.01 | 22797564.72 | 20066049.9 | 19648784.62 | 21195407.61 | 20862503.33 | 21744786.88 | 20231936.58 | 15193942.34 |
| N-Desmethyltapentadol | 26728750.17 | 25846450.44 | 27514348.76 | 29512753.11 | 25083581.09 | 11629849.31 | 21235609.25 | 10653169.39 | 21983935.11 | 9920689.137 | 9375059.747 | 10421756.01 |
| FA 17:1 | 10602477.09 | 9899072.662 | 8700062.481 | 8698093.512 | 16085201.83 | 19796340.2 | 16345534.05 | 17459591.94 | 11811609.46 | 11868619.25 | 16153781.78 | 15776665.32 |
| Ethyl (3-thioxo-1,2,4-triazaspiro[4.7]dodec-2-yl)acetate | 1098536.522 | 1259144.921 | 1410070.004 | 1501315.079 | 1158082.21 | 1470971.814 | 4401273.703 | 1523300.075 | 1528408.448 | 4524678.785 | 1102333.036 | 1445875.339 |
| Imazapyr | 559617.9861 | 533829.8823 | 687053.1121 | 526432.0637 | 502685.3737 | 382039.5959 | 580086.1796 | 511745.4066 | 492846.7999 | 520906.4126 | 578993.99 | 356957.8454 |
| Pyroglutamic acid | 152754472.8 | 114002886.5 | 224070379 | 165433355.1 | 246482053.9 | 234275957.8 | 263989489.5 | 219834365.2 | 214426290.8 | 153662170.2 | 175721986.2 | 203915295.6 |
| PC(14:0/16:0) | 4277056.013 | 3299964.437 | 3409079.426 | 2498734.023 | 743619.1674 | 10583922.54 | 18377286.19 | 12054074.71 | 9029951.841 | 17163910.52 | 14255260.61 | 9056122.057 |
| SM 36:0;3O | 4993217.329 | 2252751.087 | 4375639.637 | 5665486.838 | 2215180.637 | 1930289.682 | 5120220.458 | 1794791.295 | 3140009.313 | 2052125.964 | 2488948.513 | 1744953.625 |
| 9-Deoxy-9-methylene-16,16-dimethylprostaglandin E2 | 432326.3809 | 547203.1571 | 506729.7726 | 367632.2457 | 294639.5467 | 289753.776 | 589766.8221 | 386483.132 | 351400.1215 | 618021.649 | 482814.4626 | 245661.5449 |
| Nitrofurazone | 105736 | 801110 | 186318 | 426011 | 125412 | 138424 | 173378 | 130895 | 124069 | 960065 | 104605 | 135345 |

| Metabolites | PBS_1 | PBS_2 | PBS_3 | PBS_4 | BCG_1 | BCG_2 | BCG_3 | BCG_4 | BCGΔ *pe18*_1 | BCGΔ *pe18*_2 | BCGΔ *pe18*_3 | BCGΔ*pe18*_4 |
|---|---|---|---|---|---|---|---|---|---|---|---|---|
| | 67.6 | 8.197 | 44.31 | 6.871 | 45.58 | 95.22 | 50.8 | 67.62 | 07.08 | 8.255 | 12.68 | 67.77 |
| 6-(Chloromethyl)-N2-(2-naphthyl)-1,3,5-triazine-2,4-diamine | 978283.24 | 828164.8765 | 1174711.058 | 860225.5208 | 4520728.803 | 2347332.612 | 3957072.113 | 2918133.385 | 3120708.224 | 3429790.396 | 2899267.415 | 2080988.978 |
| 6-({[6-tert-Butyl-3-(methoxycarbonyl)-4,5,6,7-tetrahydro-1-benzothien-2-yl]amino}carbonyl)-3-cyclohexene-1-carboxylic acid | 91207.16543 | 50273.60314 | 160947.6883 | 114607.0365 | 159064.6539 | 134241.7176 | 313601.7552 | 138767.4058 | 160223.3764 | 155287.1028 | 195365.1045 | 197106.8311 |
| SM 12:1;2O/38:2 | 434892.957 | 357126.7769 | 458500.2708 | 408991.1921 | 3930179.561 | 0 | 3125750.497 | 3362565.201 | 2719931.609 | 1814862.718 | 3659981.755 | 1923050.757 |
| 7-Methoxy-4-methylcoumarin | 5423074.136 | 7145819.989 | 5565131.981 | 7011450.587 | 7387300.089 | 6762698.284 | 7243489.252 | 7626300.483 | 7275324.349 | 6881486.142 | 6374397.539 | 8318624.853 |
| PA(18:4(6Z,9Z,12Z,15Z)/14:1(9Z)) | 71367889.9 | 56492666.72 | 26301163.64 | 50323813.51 | 74755671.93 | 66081298.07 | 62097161.05 | 77715957.93 | 46897355.52 | 66649019.41 | 81299619.95 | 24830130.51 |
| PI 18:0_20:4;O | 0 | 25121.07887 | 0 | 0 | 1289363.403 | 1122847.533 | 861750.3698 | 1042354.098 | 1223674.892 | 1149107.234 | 754540.7424 | 814870.9314 |
| Bis(2-ethylhexyl) adipate | 27340826.4 | 28574003.75 | 22186705.61 | 18362852.91 | 24819175.89 | 32425211.19 | 27346802.4 | 28233761.8 | 17874000.9 | 22505735.09 | 25003164.37 | 22451381.27 |
| Volkenin | 20099994.07 | 5708568.712 | 6295288.377 | 11337587.5 | 18560194.86 | 21128161.64 | 27038306.45 | 22013843.28 | 20400334.28 | 4517323.569 | 20770757.53 | 21611897.2 |

| Metabolites | PBS_1 | PBS_2 | PBS_3 | PBS_4 | BCG_1 | BCG_2 | BCG_3 | BCG_4 | BCGΔ*pe18*_1 | BCGΔ*pe18*_2 | BCGΔ*pe18*_3 | BCGΔ*pe18*_4 |
|---|---|---|---|---|---|---|---|---|---|---|---|---|
| Phenylacetic acid | 4369117.575 | 4447660.94 | 4033120.028 | 4185256.447 | 3831018.848 | 4052172.489 | 4060204.056 | 4128527.295 | 4057207.805 | 4022759.004 | 3993075.046 | 4027391.705 |
| Dacarbazine | 873935.6319 | 963725.2994 | 929154.5255 | 955812.4603 | 922118.177 | 923985.7406 | 926557.1637 | 849546.8223 | 874517.7688 | 909572.1165 | 894350.6052 | 1052215.925 |
| 3-Hydroxypropionic acid (beta-lactic acid) | 34642312.28 | 41226506.08 | 48458651.29 | 39888494.07 | 54540001.9 | 39963602.55 | 39967297.35 | 38725766.53 | 48021368.81 | 27221231.27 | 23602624.84 | 37768902.87 |
| PC(18:3(6Z,9Z,12Z)/P-16:0) | 30348857.21 | 36719517.92 | 16705384.26 | 22763837.22 | 11250317.66 | 9180514.836 | 15351751.37 | 12750006.36 | 15651937.36 | 19608999.61 | 13464051.19 | 8727608.585 |
| PC(16:0/18:0) | 208830843.2 | 233506071.8 | 294860075 | 198650546 | 252504868.3 | 190656303.9 | 307315361.5 | 224138121.9 | 238099897.2 | 218968930.3 | 236489026.8 | 116895037 |
| 2-(3-Hydroxycyclohexyl)-5-(2-methyl-2-octanyl)phenol | 865897.1248 | 1101506.179 | 766677.4962 | 844491.9091 | 862009.4061 | 1032694.831 | 834476.4042 | 834716.6056 | 867304.3191 | 769875.6594 | 815856.8472 | 809039.8342 |
| Dimethyl sulfoxide | 11149197.91 | 10575392.15 | 3854946.862 | 10435936.49 | 10713422.1 | 11297500.6 | 6869230.814 | 5328791.433 | 4309633.418 | 10238275.96 | 12779644.34 | 9440232.917 |
| Fomepizole | 5457679.191 | 5293826.763 | 6279509.728 | 6086944.33 | 8656094.143 | 8243701.536 | 10994429.05 | 7898250.925 | 9313690.643 | 10007075.65 | 10829025.27 | 7624756.542 |
| 2-Phosphonoethylphosphonic acid | 6609874.954 | 7311507.46 | 6066773.469 | 6963776.982 | 7790579.653 | 6529895.737 | 6627334.537 | 6449388.907 | 6509075.767 | 6978308.211 | 6374615.867 | 6038623.241 |
| 3-(3-Methoxybenzyl)piperidine | 3220828.836 | 3286717.852 | 3019896.898 | 3035261.2 | 3332602.373 | 3339240.407 | 3214659.507 | 3244830.549 | 3220421.875 | 3178659.141 | 2903239.796 | 2859335.489 |
| 5-Hexyl-2,4-dihydroxybenzaldehyde | 2586409.446 | 2729851.457 | 2431765.811 | 2300708.392 | 2654411.719 | 2656656.003 | 2513376.257 | 2711649.869 | 2447597.595 | 2533283.688 | 2617780.215 | 2295576.705 |

| Metabolites | PBS_1 | PBS_2 | PBS_3 | PBS_4 | BCG_1 | BCG_2 | BCG_3 | BCG_4 | BCGΔ*pe18*_1 | BCGΔ*pe18*_2 | BCGΔ*pe18*_3 | BCGΔ*pe18*_4 |
|---|---|---|---|---|---|---|---|---|---|---|---|---|
| Palmitamide | 7550878.359 | 6262593.381 | 4118601.733 | 4133014.794 | 6504395.805 | 5999408.176 | 4479365.329 | 5159850.455 | 6307514.56 | 4250871.874 | 4809178.875 | 3232107.388 |
| 5-{8(Z),11(Z)-pentadecadienyl}resorcinol | 2246157.175 | 3846064.577 | 1626451.721 | 5717468.078 | 1722420.963 | 2519470.482 | 2827112.572 | 2104975.043 | 2943737.496 | 1914506.272 | 1583666.845 | 2220708.312 |
| Ethyl 4-methyl-5-[(propan-2-yl)oxy]-9H-pyrido[3,4-b]indole-3-carboxylate | 10428768.9 | 9911606.122 | 11773173.51 | 10667099.64 | 7184954.764 | 8913026.456 | 6004877.819 | 7967736.44 | 7465466.495 | 5379242.791 | 7093513.541 | 8686951.856 |
| Telon Yellow FG01 | 973711.4762 | 864422.9424 | 1036306.565 | 920357.796 | 1165087.105 | 1075407.274 | 1155137.991 | 1069168.11 | 1139933.269 | 1167926.192 | 1066896.649 | 1002671.084 |
| SMGDG O-18:5_3:0 | 3383446.255 | 3705795.081 | 2814433.634 | 3479183.054 | 3235300.47 | 3486768.807 | 3668434.569 | 3353461.036 | 3398995.694 | 3107022.4 | 3204062.453 | 3097711.919 |
| .beta.-Damascone | 5591528.577 | 6315479.256 | 4667113.904 | 4681337.182 | 5636241.19 | 7249249.218 | 5341627.161 | 5969369.522 | 5277459.908 | 4782261.486 | 5227891.124 | 5339490.09 |
| 3-Pyridol | 1250533.231 | 1473387.228 | 1448939.095 | 1306007.258 | 1155805.012 | 1158994.787 | 1130555.656 | 1306086.859 | 1258252.246 | 1257035.067 | 1430089.913 | 1212939.58 |
| N-Cycloheptyl-N'-cyclohexylurea | 3906291.641 | 3850268.944 | 3388817.787 | 3853784.05 | 4206802.029 | 3725928.276 | 5084944.196 | 3719231.721 | 3818050.327 | 4312962.31 | 3525158.041 | 3672393.551 |
| Enalaprilat butyl tert-butyl diester | 2540600.506 | 1887617.823 | 2858254.5 | 3334498.743 | 2774410.885 | 1657223.481 | 1939600.834 | 1822110.208 | 1314486.794 | 2210042.289 | 1885031.1 | 715529.9335 |
| Norleucine | 16375892.26 | 13243922.04 | 14555978.35 | 17865563.53 | 13671128.08 | 13679648.4 | 19263540.74 | 24609234.54 | 15785040.42 | 15091386.41 | 12825853.21 | 14514744.8 |

| **Metabolites** | **PBS_1** | **PBS_2** | **PBS_3** | **PBS_4** | **BCG_1** | **BCG_2** | **BCG_3** | **BCG_4** | **BCGΔ*pe18*_1** | **BCGΔ*pe18*_2** | **BCGΔ*pe18*_3** | **BCGΔ*pe18*_4** |
|---|---|---|---|---|---|---|---|---|---|---|---|---|
| PI 16:0_22:6 | 1395412.586 | 2535423.581 | 450415.06 | 1534500.203 | 2101680.298 | 1980292.169 | 2374223.891 | 6231635.538 | 7549296.804 | 2538479.053 | 3714798.152 | 1812585.359 |
| 4-[4-(Trifluoromethoxy)phenoxy]piperidine | 1000850.957 | 631286.6765 | 991504.8694 | 674414.1845 | 3860743.889 | 3047494.928 | 2941056.751 | 3274933.926 | 3267272.47 | 2921503.05 | 2651577.574 | 3453210.237 |
| Oxalic acid | 13498891.85 | 13372299.71 | 14047512.45 | 13182399.49 | 12899215.29 | 13174426.26 | 14211220.19 | 12875175.34 | 14848598.3 | 4432279.056 | 13500304.91 | 14000495.83 |
| Nicotinamide | 34609049.32 | 30024510.93 | 73412540.3 | 41296680.97 | 268878321.2 | 221478906.5 | 386437939.1 | 283898276.5 | 186456110.8 | 196649482.4 | 280881349.7 | 220710529.6 |
| Cer 8:1;2O/17:0 | 524458.0166 | 304788.8221 | 515131.9655 | 224129.4283 | 640515.7832 | 943694.0669 | 954799.3646 | 816305.2916 | 648598.8137 | 442014.8122 | 634336.543 | 925137.1889 |
| (2R)-2-(2,5-Difluorophenyl)pyrrolidine | 7283521.985 | 5514501.2 | 6065256.45 | 5734894.484 | 14430037.46 | 5598636.295 | 10206171.02 | 9249458.591 | 7098746.646 | 9255215.324 | 7896319.764 | 5003032.958 |
| 2-(1-Hydroxy-1-methoxy-3-methylbutyl)-6-methyl-5H-[1,3]dioxolo[4,5-c]pyridin-4-one | 1631460.27 | 1772252.251 | 1843087.675 | 7618270.24 | 1364955.587 | 2120488.48 | 9451795.777 | 1301363.068 | 4438176.834 | 2665121.158 | 1387549.079 | 8691195.718 |
| TG 8:0_16:4_24:5 | 2942580.209 | 4805082.53 | 3274468.177 | 3209661.966 | 3104302.02 | 3569730.69 | 2752435.72 | 3326001.175 | 4398251.154 | 2931548.653 | 4308401.094 | 3276920.706 |
| MG 22:2 | 61595495.74 | 62693189.48 | 44479010.8 | 51327821.24 | 49799989.79 | 30181274.45 | 53809531.14 | 46329212.32 | 43767703.25 | 35925003.32 | 32150265.97 | 23331666.91 |
| MG 18:2 | 31346246.99 | 24978637.08 | 42825140.49 | 31016091.94 | 108924711.7 | 68046624.56 | 109462644.4 | 83030719.09 | 58060715.25 | 83931122.15 | 86783352.78 | 46578009.99 |

| Metabolites | PBS_1 | PBS_2 | PBS_3 | PBS_4 | BCG_1 | BCG_2 | BCG_3 | BCG_4 | BCGΔ*pe18*_1 | BCGΔ*pe18*_2 | BCGΔ*pe18*_3 | BCGΔ*pe18*_4 |
|---|---|---|---|---|---|---|---|---|---|---|---|---|
| 2-(2-Thienyl)-2,3-dihydro-1H-perimidine | 1696380.412 | 1664976.229 | 2083365.31 | 2069968.248 | 6349993.522 | 4076347.862 | 4806962.62 | 5117880.555 | 3782973.044 | 3826841.631 | 3968592.935 | 3890883.32 |
| SM(d18:1/17:0) | 5939540.653 | 6183868.615 | 7004738.846 | 4266452.711 | 7272810.202 | 6845468.782 | 10665648.75 | 8311635.877 | 8030791.936 | 9490524.541 | 7840425.282 | 4099405.314 |
| (5-Bromo-3-fluoro-2-pyridinyl)methanol | 1169719.646 | 977916.1144 | 1555515.734 | 1261842.746 | 7356917.32 | 4809162.102 | 6292874.34 | 5937806.042 | 5436606.6 | 4738469.095 | 4789003.036 | 3703361.588 |
| PI(18:1(11Z)/18:1(11Z)) | 18595119.82 | 20410677.22 | 10194613.77 | 15026906.62 | 2992801.086 | 3425143.688 | 4610220.031 | 4379329.037 | 5215202.13 | 5429385.72 | 3073251.232 | 3331289.336 |
| N-(3-Fluorophenyl)-5-nitro-2-furamide | 328982.5016 | 1523161.133 | 2969381.108 | 2350905.749 | 588805.0971 | 193906.4229 | 715076.0506 | 123069.2053 | 553053.1907 | 1110188.471 | 630171.546 | 2184380.489 |
| PC 16:2_16:2 | 462457.1354 | 543200.0818 | 451851.3475 | 696750.4223 | 3436817.377 | 3462925.268 | 4099285.643 | 3968825.511 | 4387851.325 | 5080217.708 | 3831010.176 | 2423116.839 |
| p-Hydroxynonanophenone | 1800937.329 | 2081259.298 | 1874152.055 | 1870329.783 | 1611508.849 | 2112958.366 | 1577655.954 | 1786019.469 | 2011561.514 | 1781187.382 | 1757917.348 | 2125782.512 |
| 9-Riburonosylhypoxanthine | 5430029.313 | 4346041.307 | 8095419.088 | 5406240.704 | 38282414.86 | 21922058.97 | 36718410.52 | 26930152.36 | 25646948.86 | 27637007.93 | 29512218.92 | 18463007.58 |
| Phenylsulfate | 70364185.75 | 54282038.04 | 88583818.92 | 61865024.23 | 78911080.47 | 56059425.81 | 108282682.2 | 70372652.07 | 43913172.76 | 74463081.87 | 84448187.47 | 42969904.85 |
| 2-(2-Methoxyethoxy)ethanamine | 83084382.83 | 36921893.41 | 40618579.34 | 39731217.16 | 81948502.89 | 63158694.97 | 54246413.47 | 61265467.35 | 56700375.77 | 53454549.45 | 35442041.34 | 57678635.34 |
| N-Acetylphenylalanine | 2484587.782 | 863358.506 | 1651166.673 | 1164205.141 | 1236512.51 | 2466853.469 | 1093616.073 | 2545478.271 | 996891.8143 | 1611485.192 | 2431527.008 | 1366408.961 |

| **Metabolites** | **PBS_1** | **PBS_2** | **PBS_3** | **PBS_4** | **BCG_1** | **BCG_2** | **BCG_3** | **BCG_4** | **BCGΔ*pe18*_1** | **BCGΔ*pe18*_2** | **BCGΔ*pe18*_3** | **BCGΔ*pe18*_4** |
|---|---|---|---|---|---|---|---|---|---|---|---|---|
| TG 12:0_14:0_16:4 | 3115642.222 | 4159390.742 | 2905000.745 | 5283236.893 | 2983693.072 | 131937.6235 | 5730086.273 | 4288132.225 | 5490160.391 | 3887648.91 | 2639916.651 | 4412091.011 |
| 2-Adamantylamine | 8179462.219 | 5012659.157 | 7173586.15 | 8147987.753 | 5100293.729 | 8155758.977 | 5736052.193 | 9413411.952 | 10350546.84 | 9717167.103 | 10824024.69 | 4355109.093 |
| SM 34:2;3O | 91842.23944 | 160273.5966 | 243021.2929 | 213761.3451 | 1855969.579 | 1972898.448 | 1552498.703 | 1612163.222 | 3147113.508 | 1926365.42 | 2196320.203 | 2395287.038 |
| N-Nitroso-N-methyl-3-aminopropionic acid, methyl ester | 23385123.77 | 20860882 | 26469935.67 | 24298690.53 | 104778455.5 | 87276763.99 | 89558429.68 | 66620928.14 | 92969065.14 | 52527756.85 | 55957901.53 | 56752156.79 |
| Spermidine | 16174522.04 | 12390507.48 | 21948488.5 | 18918320.43 | 20755862.88 | 22386591.03 | 25022238.41 | 23718662.77 | 20731629.15 | 17527496.87 | 23371947.62 | 22756887.19 |
| 2-Naphthalenecarboxamide, 1-hydroxy-N-[2-(5-hydroxy-1H-indol-3-yl)ethyl]- | 145664.5478 | 1718756.008 | 2712332.304 | 2002957.558 | 150133.0759 | 57343.58655 | 493439.663 | 60475.68405 | 371905.3616 | 272254.4164 | 373568.2819 | 7040859.125 |
| 3-{2-[2-(2-Cyanoethoxy)ethoxy]ethoxy}propanenitrile | 6075889.694 | 6295107.543 | 5589577.842 | 5755432.515 | 5543268.634 | 5474689.216 | 5397981.352 | 5343818.815 | 5610860.056 | 5375614.68 | 5398518.151 | 5434825.003 |
| 3'-Hydroxydiclofenac | 1442999.432 | 1233124.296 | 2366487.927 | 1637443.671 | 3260342.749 | 3322235.939 | 3664066.082 | 3054355.553 | 2832665.216 | 2654174.097 | 2645153.507 | 3119495.73 |
| Propyl 2,4-decadienoate | 15767594.22 | 15207501.85 | 13984455.52 | 15775887.75 | 14215104.3 | 11887655.61 | 12701503.56 | 11921686.52 | 12554963.83 | 12964162.41 | 12373379.01 | 12935348.35 |
| Piperidine-1-carbox | 263964. | 972217. | 129521 | 124968 | 281306 | 215340. | 273909. | 275989 | 391204. | 551145. | 245922. | 401087 |

| Metabolites | PBS_1 | PBS_2 | PBS_3 | PBS_4 | BCG_1 | BCG_2 | BCG_3 | BCG_4 | BCGΔ *pe18*_1 | BCGΔ *pe18*_2 | BCGΔ *pe18*_3 | BCGΔ*pe18*_4 |
|---|---|---|---|---|---|---|---|---|---|---|---|---|
| imidamide | 9791 | 3102 | 4.08 | 1.553 | 6.226 | 8133 | 5068 | 6.275 | 4658 | 1645 | 0773 | 2.826 |
| LNAPS 22:3/N-16:1 | 6751654.289 | 817335.8162 | 2096435.693 | 1189586.362 | 2190731.945 | 2034941.051 | 2050086.293 | 1896103.68 | 3290943.099 | 1856121.38 | 533837.8435 | 1856949.866 |
| PI O-16:2_14:0 | 647290.9394 | 1267695.439 | 599349.4907 | 723342.6953 | 213439.9722 | 562358.1715 | 253297.271 | 295218.7613 | 252196.2387 | 301897.7361 | 327193.1445 | 857713.4251 |
| Aspochalasin D | 1755567.572 | 1801646.981 | 1618480.126 | 1744194.998 | 1632410.741 | 1677321.351 | 1718868.618 | 1638676.577 | 1848082.845 | 1795595.92 | 1760476.627 | 1739890.409 |
| LPS 20:3 | 2021927.599 | 2549795.852 | 2592533.056 | 2021545.063 | 3984524.305 | 2579255.983 | 2049105.244 | 1821479.442 | 2890745.335 | 3105396.18 | 2928882.489 | 3138269.53 |
| 3,6-Dichloro-N-(pyridin-2-ylmethyl)pyridazine-4-carboxamide | 2883969.089 | 3107984.365 | 9700847.05 | 3039230.335 | 1770909.114 | 1731936.714 | 1615639.635 | 1830516.898 | 2107452.8 | 2841739.137 | 3889636.348 | 3600231.677 |
| 1,10a-Dihydroxy-4,4,7,11b-tetramethyl-1,2,3,4a,5,6,6a,7,11,11a-decahydronaphtho[2,1-f][1]benzofuran-9-one | 1477971.591 | 169823.6892 | 1228049.289 | 173954.8902 | 2033349.546 | 2857068.841 | 1256187.453 | 2391210.426 | 2925906.09 | 783533.0914 | 1029621.388 | 3652970.419 |
| columbianetin | 4623457.68 | 4377606.37 | 4989558.221 | 4519947.621 | 4082461.263 | 4977516.69 | 4295772.923 | 4868946.8 | 4157278.184 | 3818283.553 | 4515600.322 | 4461500.261 |
| PI 17:0_20:4 | 1889119.249 | 1799333.9 | 987456.7041 | 2071483.199 | 1382005.081 | 739699.4189 | 967810.9688 | 1708986.37 | 2307897.556 | 2810724.421 | 776516.2533 | 909593.6615 |
| DGDG O-9:0_18:1 | 112096 | 874034 | 103027 | 802196 | 118002 | 115050 | 873522 | 101391 | 778588 | 114660 | 113003 | 903095 |

| Metabolites | PBS_1 | PBS_2 | PBS_3 | PBS_4 | BCG_1 | BCG_2 | BCG_3 | BCG_4 | BCGΔ*pe18*_1 | BCGΔ*pe18*_2 | BCGΔ*pe18*_3 | BCGΔ*pe18*_4 |
|---|---|---|---|---|---|---|---|---|---|---|---|---|
| | 95.32 | 5.733 | 91.79 | 9.917 | 89.78 | 62.61 | 5.29 | 91.33 | 6.015 | 15.38 | 44.3 | 9.665 |
| 19-Noretiocholanolone | 4566097.73 | 4703541.588 | 4025159.582 | 4699237.871 | 1800669.711 | 4010148.352 | 5293238.05 | 3989430.918 | 6084868.644 | 2779707.836 | 5426129.424 | 4635883.607 |
| NAGly 11:0/16:3 | 9915.491322 | 272401.6828 | 923195.5591 | 97646.78603 | 45281.33613 | 9439.658173 | 118961.9602 | 0 | 0 | 20055.48678 | 64438.66334 | 2391151.188 |
| 3-O-Acetyl-2-O-((2E)-3-(4-hydroxyphenyl)prop-2-enoyl)hexopyranose | 26676341.94 | 26663368.63 | 15474973.27 | 33987619.21 | 9091544.548 | 38230789.7 | 18694846.82 | 17548266.5 | 14250394.64 | 6277401.475 | 21913409.27 | 27139389.23 |
| 2-(Methylamino)-1-(morpholin-4-yl)ethan-1-one | 2777100.772 | 2968215.815 | 2794834.594 | 2709066.409 | 2763745.666 | 2765666.405 | 2794695.775 | 3834449.931 | 2765705.765 | 2840075.431 | 2653750.127 | 2719078.362 |
| (2E,4E,9Z,12Z)-1-Piperidin-1-yloctadeca-2,4,9,12-tetraen-1-one | 637141.9988 | 931810.5835 | 599654.9167 | 530321.769 | 2206563.103 | 2181341.184 | 1901242.054 | 1594370.385 | 1820753.636 | 1898449.593 | 1166267.708 | 941845.1642 |
| MGDG O-18:2_3:0 | 14545239.42 | 18013228.38 | 11431855.22 | 11879911.65 | 11265724.43 | 12431122.62 | 16621282.37 | 12411071.87 | 12339662.52 | 14007247.56 | 17214327.01 | 11865486.79 |
| 1'-Naphthoylindole | 577367.3034 | 589868.6901 | 844226.3528 | 590881.118 | 1690097.439 | 1297672.724 | 1723526.62 | 1466703.723 | 1369233.108 | 1059202.861 | 1182963.034 | 1099556.553 |
| Acitretin | 3299852.203 | 3176490.766 | 8340303.062 | 4911043.886 | 7220453.619 | 3602247.027 | 7853074.497 | 8946256.89 | 9164163.607 | 2509939.962 | 2468598.965 | 8249905.014 |
| Neostigmine cation | 14568987.18 | 18598985.83 | 9308352.651 | 16115796.93 | 16345521.78 | 11211165.55 | 21303759.7 | 18545997.11 | 17674894.67 | 15669437.92 | 13905549.33 | 13337310.76 |

| Metabolites | PBS_1 | PBS_2 | PBS_3 | PBS_4 | BCG_1 | BCG_2 | BCG_3 | BCG_4 | BCGΔ*pe18*_1 | BCGΔ*pe18*_2 | BCGΔ*pe18*_3 | BCGΔ*pe18*_4 |
|---|---|---|---|---|---|---|---|---|---|---|---|---|
| 2-[Bis(5-methyl-2-furyl)methyl]-6-bromo-4-nitrophenol | 20468722.96 | 17121978.39 | 6792745.215 | 17882073.09 | 11123463.21 | 4263223.604 | 16278431.01 | 10042352.53 | 16980085.03 | 16548025.06 | 8642805.145 | 7243469.364 |
| Allose | 158677464.2 | 121019993.9 | 163578314.7 | 148581193.5 | 122842412.9 | 145645511.8 | 121670957.5 | 128739498.5 | 128503914 | 74437631.99 | 66910977.38 | 124800632.7 |
| C20 phytosphingosine | 2906182.66 | 3246140.773 | 2244269.696 | 2966658.546 | 2699531.141 | 2285737.649 | 2444261.499 | 2602855.482 | 2760532.915 | 2128802.658 | 2835903.967 | 8609838.228 |
| PC(20:5(5Z,8Z,11Z,14Z,17Z)/20:4(5Z,8Z,11Z,14Z)) | 30383251.99 | 33443664.85 | 15107332.2 | 22651292.4 | 39861907.68 | 30730286.16 | 35511793.89 | 39617713.93 | 45639005.33 | 39124605.67 | 30614062.38 | 30597814.84 |
| Fulvestrant 9-sulfone | 391818.4229 | 236460.7812 | 269973.7497 | 400455.0928 | 508512.3217 | 80741.40283 | 404847.4602 | 410680.4501 | 347957.9393 | 297898.2107 | 311092.7691 | 329487.4432 |
| Nicergoline | 254523.9984 | 252317.8768 | 266850.6009 | 233013.1537 | 457197.2815 | 427855.4497 | 578093.5854 | 580467.9317 | 517910.5332 | 545150.2971 | 540240.6236 | 655301.1778 |
| SM 8:0;2O/10:0 | 6198372.207 | 7996626.594 | 4457567.671 | 5698407.949 | 6770016.826 | 6912527.049 | 6232433.584 | 5398004.338 | 5353743.201 | 5548444.873 | 6821240.15 | 4679452.51 |
| Cer 14:3;2O/5:0 | 674574.7487 | 867691.1834 | 391250.161 | 75819.27619 | 525456.4574 | 464317.6915 | 563692.576 | 438126.1057 | 442037.8212 | 491305.5953 | 586629.1071 | 358758.8941 |
| Meperidine | 3087894.014 | 3131417.966 | 2765460.917 | 2960783.904 | 3005368.277 | 2972318.303 | 3202089.32 | 3373135.502 | 3008948.984 | 3182300.95 | 2947692.815 | 2759163.639 |
| LPC 18:3 | 21902.40844 | 64706.91763 | 67563.66998 | 45969.54052 | 271108.9686 | 332272.223 | 474391.2613 | 360042.6612 | 451851.9651 | 260148.9057 | 372889.3711 | 299012.0824 |
| LPG 20:3 | 5601495.356 | 2633221.123 | 5145960.513 | 2893204.257 | 5325800.029 | 5615144.704 | 5989541.683 | 6063601.857 | 3857814.538 | 6707981.914 | 5146588.262 | 4292064.013 |

| Metabolites | PBS_1 | PBS_2 | PBS_3 | PBS_4 | BCG_1 | BCG_2 | BCG_3 | BCG_4 | BCGΔ*pe18*_1 | BCGΔ*pe18*_2 | BCGΔ*pe18*_3 | BCGΔ*pe18*_4 |
|---|---|---|---|---|---|---|---|---|---|---|---|---|
| Phenol, 5-[(1Z)-2-(3,5-dimethoxyphenyl)ethenyl]-2-methoxy- | 2249195.762 | 2493438.436 | 2232137.787 | 2520127.858 | 2386770.401 | 2524464.058 | 2387980.213 | 2440079.641 | 2332826.439 | 2477460.689 | 2521146.202 | 2262149.578 |
| Arachidonoyl-N-methylamide | 872938.3365 | 918515.7928 | 742579.3942 | 831353.4769 | 790424.3889 | 783350.9674 | 838191.4288 | 822294.5189 | 989453.7863 | 847590.6774 | 770598.2288 | 767458.8226 |
| o-Tyrosine | 10761983.29 | 9096234.18 | 14444038.27 | 11318398.37 | 22093684.58 | 15963815.57 | 19915821.77 | 18184252.52 | 19243070.94 | 12997049.09 | 14623883.45 | 17887845.21 |
| alanyl-isoleucine | 1210810.155 | 1291967.233 | 812135.2463 | 955348.7526 | 819055.5639 | 953213.3586 | 1510331.531 | 1230053.605 | 1068170.236 | 961084.8139 | 1043846.22 | 1117775.167 |
| Oxeladin | 3925109.227 | 4438088.591 | 2630687.863 | 2319828.328 | 2914216.812 | 3721480.706 | 5786112.385 | 5643061.728 | 3153977.498 | 3524512.281 | 4439235.619 | 3086802.571 |
| 2-Amino-4-[carbamimidoyl(methyl)amino]butanoic acid | 15050461.28 | 8879407.302 | 22749941.33 | 7758218.534 | 17453839.95 | 3209182.15 | 7159047.837 | 11519888.38 | 4407104.668 | 9031895.726 | 8345561.05 | 11107933.75 |
| 2-Amino-5-bromo-3-chloropyrazine | 2238231.937 | 2539440.836 | 2166292.108 | 2722357.8 | 2285092.336 | 2194462.361 | 2294260.297 | 2565749.037 | 2355417.581 | 2304723.187 | 2169296.859 | 2096884.415 |
| Methyl 4-formyl-3-hydroxy-2-naphthoate | 2394522.533 | 3477036.734 | 2943738.208 | 2642557.093 | 3506655.429 | 3036456.171 | 3429742.672 | 3241066.77 | 3333662.351 | 2614729.704 | 3356246.412 | 3672439.898 |
| 9,10-Epoxy-18-hydroxystearate | 1413217.72 | 1791259.334 | 1318263.339 | 1385894.13 | 4671289.093 | 4254739.512 | 3604100.884 | 3264933.16 | 2481108.496 | 2042731.731 | 3578311.773 | 5120360.074 |
| N-Cyclooctyl-4-methylbenzenesulfona | 2225873.214 | 2167997.648 | 2257033.065 | 2221576.675 | 3164387.6 | 3662491.926 | 3363586.675 | 3221612.428 | 2757240.353 | 1921190.297 | 2093184.76 | 3071236.53 |

| **Metabolites** | **PBS_1** | **PBS_2** | **PBS_3** | **PBS_4** | **BCG_1** | **BCG_2** | **BCG_3** | **BCG_4** | **BCGΔ*pe18*_1** | **BCGΔ*pe18*_2** | **BCGΔ*pe18*_3** | **BCGΔ*pe18*_4** |
|---|---|---|---|---|---|---|---|---|---|---|---|---|
| mide | | | | | | | | | | | | |
| LPS 16:0 | 17805728.76 | 23061693.78 | 16937629.03 | 15029352.43 | 17011527.46 | 15549823.08 | 15307701.14 | 15391486.44 | 16923497.5 | 17851607.55 | 20569908.25 | 12433537.76 |
| beta-Alanine | 3394823.365 | 2857013.822 | 4838427.035 | 3196684.619 | 16706744.12 | 9755094.368 | 18417068.59 | 11705299.09 | 11193832.57 | 11843630.53 | 13191529.6 | 7337559.256 |
| Gly-Ala | 1681742.824 | 1688220.9 | 1981922.724 | 1739981.68 | 6489290.496 | 5483850.853 | 5875438.602 | 5765094.048 | 5931315.09 | 4731312.104 | 3871408.726 | 3999472.212 |
| PS 3:0_21:2 | 271638.0805 | 277153.8925 | 273730.3069 | 236763.182 | 307830.252 | 320420.2831 | 333527.4435 | 305830.3109 | 269877.5132 | 287812.2815 | 257916.2343 | 182378.0443 |
| Valproic acid | 76924997.5 | 79054326.54 | 74556549.09 | 78961041.58 | 77172446.79 | 75854224.77 | 76616867.53 | 110190302.2 | 77181871.53 | 76897030.24 | 75785682.16 | 81791063.03 |
| PE O-10:0_7:0 | 379250.6106 | 1835254.96 | 1119416.493 | 1366330.962 | 2622229.947 | 5890349.94 | 3113850.612 | 3013755.198 | 3149955.069 | 2541296.71 | 3345021.261 | 1912582.934 |
| DLCL 16:1_18:1 | 340100.6571 | 369417.2795 | 39608.68405 | 179623.3181 | 313731.3005 | 323133.7341 | 378583.3106 | 308301.2263 | 547200.9906 | 231386.1883 | 291604.1932 | 388101.1755 |

| Metabolites | PBS_1 | PBS_2 | PBS_3 | PBS_4 | BCG_1 | BCG_2 | BCG_3 | BCG_4 | BCGΔ *pe18*_1 | BCGΔ *pe18*_2 | BCGΔ *pe18*_3 | BCGΔ*pe18*_4 |
|---|---|---|---|---|---|---|---|---|---|---|---|---|
| (2S,3R)-3-((2'R,3S,4b'R,7'S,10a'R)-7'-(2-O-(6-Deoxy-.alpha.-L-mannopyranosyl)-.beta.-D-glucopyranosyl)oxy)-4b',8',8',10a'-tetramethyl-5-oxotetradecahydro-2'H-spiro[furan-3,1'-phenanthren]-2'-yl)-1-((2R)-4-methyl-5-oxo-2,5-dihydrofuran-2-yl)butan-2-yl acetate | 771810.6518 | 745398.4045 | 613860.5809 | 743655.9179 | 747799.1522 | 775631.4606 | 777472.9168 | 878493.5236 | 697068.022 | 903711.5015 | 769866.9743 | 739536.3019 |
| Mesaconic acid | 36423784.15 | 24293593.32 | 59315185.46 | 37642727.23 | 728452365.5 | 453193811.1 | 737099582.8 | 551101366.1 | 426261451.5 | 577805304.2 | 612611096.8 | 368292521.3 |
| 5-(2-Chlorophenyl)-4-phenyl-4H-1,2,4-triazol-3-yl hydrosulfide | 4573747.622 | 3364395.621 | 6141464.285 | 4817027.216 | 6420973.043 | 6618754.769 | 6957285.646 | 5722568.554 | 5532143.562 | 4302894.893 | 4432715.031 | 5390439.819 |
| SL 13:2;O/12:1;O | 6083413.301 | 3853746.048 | 3699327.391 | 4431787.964 | 3735612.075 | 4509303.649 | 3210396.278 | 2466964.035 | 2999476.416 | 3851647.125 | 5950579.756 | 2570127.624 |
| 11-nitro-1-undecene | 24834455.65 | 25441651.07 | 21188003.34 | 30334101.84 | 19620656.49 | 20189874.37 | 29788407.96 | 19121141.12 | 23282558.68 | 22982476.28 | 21226256.22 | 30696912.22 |

| Metabolites | PBS_1 | PBS_2 | PBS_3 | PBS_4 | BCG_1 | BCG_2 | BCG_3 | BCG_4 | BCGΔ*pe18*_1 | BCGΔ*pe18*_2 | BCGΔ*pe18*_3 | BCGΔ*pe18*_4 |
|---|---|---|---|---|---|---|---|---|---|---|---|---|
| 4-(Trifluoromethyl)piperidine | 6850191.854 | 5335366.358 | 9081844.065 | 6746310.666 | 8744036.985 | 9136814.002 | 8555405.508 | 8503322.554 | 9643015.557 | 5831588.783 | 5986764.606 | 7293087.976 |
| Zeatin | 453344.3902 | 688406.7126 | 827761.9762 | 677730.2029 | 958813.4414 | 1436386.892 | 1100593.245 | 919793.5255 | 980905.5704 | 686654.1835 | 747696.5609 | 893278.7828 |
| Chlorthalidone | 4847509.817 | 4778097.178 | 4484787.031 | 6213341.112 | 5105941.66 | 5380582.121 | 4201904.88 | 5679572.321 | 5083484.975 | 5412625.528 | 6580154.233 | 4284435.136 |
| MGDG O-22:6_6:0 | 4173193.385 | 2642370.991 | 4196000.668 | 2971268.334 | 2393573.192 | 2843570.593 | 2461905.317 | 2360361.798 | 1662826.416 | 2129207.925 | 2535871.759 | 2393786.534 |
| (4-Oxo-1,3-thiazolidin-3-yl)acetic acid | 135325.8512 | 59210.50377 | 258430.0979 | 149057.5254 | 1621402.358 | 1604330.738 | 2146601.49 | 1515503.346 | 1353159.345 | 1549132.231 | 1794724.178 | 657441.2157 |
| PE O-16:3_4:0 | 2438629.566 | 1547747.086 | 2484152.347 | 2000615.044 | 4639250.83 | 5574755.495 | 3935556.031 | 3625106.134 | 3166235.358 | 4547295.176 | 4764396.009 | 4910955.922 |
| Sorbose | 158677464.2 | 121019993.9 | 163578314.7 | 148581193.5 | 122842412.9 | 145645511.8 | 121670957.5 | 128739498.5 | 128503914 | 74437631.99 | 66910977.38 | 124800632.7 |
| NAE 16:3 | 6989254.095 | 7106842.735 | 6078757.921 | 6267489.913 | 6703951.39 | 6502629.513 | 6866310.634 | 6523826.554 | 6563203.368 | 7038560.88 | 6242060.287 | 6659138.644 |
| PS(20:5(5Z,8Z,11Z,14Z,17Z)/15:0) | 11474801.24 | 11617294.12 | 11476894.3 | 10515050.83 | 8026732.789 | 6353014.573 | 9305187.988 | 7722513.142 | 7059886.175 | 8586474.21 | 7786337.634 | 5541173.893 |
| N-(4-Phenoxyphenyl)-2-pyrazinecarboxamide | 594250.2416 | 594911.4588 | 954303.4314 | 648619.2257 | 1810736.74 | 1443783.638 | 1867309.533 | 1602695.494 | 1582472.716 | 1254270.302 | 1225055.735 | 1224745.379 |
| 5-Amino-1-pentanol | 1755216.949 | 1825730.763 | 1502696.06 | 1716222.431 | 2005839.168 | 1767348.377 | 2341507.765 | 1473841.984 | 1764831.664 | 1436462.576 | 1926115.789 | 1690423.001 |

| Metabolites | PBS_1 | PBS_2 | PBS_3 | PBS_4 | BCG_1 | BCG_2 | BCG_3 | BCG_4 | BCGΔ*pe18*_1 | BCGΔ*pe18*_2 | BCGΔ*pe18*_3 | BCGΔ*pe18*_4 |
|---|---|---|---|---|---|---|---|---|---|---|---|---|
| 5,6-Dihydroxy-8Z,11Z,14Z-eicosatrienoic acid | 1759718.07 | 2163910.092 | 3069769.206 | 2241548.752 | 2293942.284 | 2693492.122 | 2113638.778 | 2405337.146 | 740007.8535 | 1813549.185 | 2727489.094 | 2352395.488 |
| 4-Chloro-2-methyl-5-(4-morpholinyl)-3(2H)-pyridazinone | 1826228.331 | 1407922.378 | 2934592.955 | 1936397.116 | 3735684.051 | 3728493.13 | 3723150.336 | 3331887.067 | 3477293.978 | 2011116.563 | 2220134.725 | 3051876.988 |
| N-Methyl-O-methyl-L-tyrosine | 1057979.285 | 1137598.027 | 1029392.296 | 1084067.462 | 1741065.747 | 1090472.291 | 1155026.184 | 1107137.912 | 1134402.514 | 1047081.293 | 1746263.535 | 1014427.042 |
| N-Methyl-N-(methylsulfonyl)glycine | 2004612.278 | 3540513.21 | 2590579.059 | 3510160.261 | 11999810.03 | 5311239.859 | 8439245.334 | 6456723.455 | 5921805.676 | 7557502.61 | 7249142.15 | 4905230.124 |
| N-(2-Propynyl)cyclohexanamine | 3014094.894 | 3512443.064 | 2111398.296 | 3073450.169 | 3244357.348 | 2596427.941 | 3216692.12 | 3166292.729 | 3429469.35 | 3048713.093 | 2935066.526 | 2707521.695 |
| PC(18:3(9Z,12Z,15Z)/P-16:0) | 24999449.67 | 11239029.06 | 4013445.704 | 8501502.101 | 14149359.04 | 3325084.868 | 4323434.985 | 5936393.753 | 15548106.68 | 4906981.113 | 12837338.47 | 10304503.79 |
| Methyl 1-aminocyclopropanecarboxylate | 5434780.548 | 1819956.148 | 3602284.367 | 2518494.604 | 5516594.607 | 5574029.725 | 5231423.466 | 4834779.704 | 4213236.393 | 4264307.751 | 4833443.784 | 4158750.668 |
| PG 3:0_20:5 | 13830088.96 | 12729711.71 | 10645425.42 | 12499688.22 | 14184366.35 | 12668274.88 | 12168081.19 | 13322296.58 | 13263878.01 | 13633702.82 | 12720565.51 | 13630994.84 |
| 15-Ketofluprostenol | 334864.6218 | 2022489.09 | 1729221.001 | 1841838.754 | 2136134.097 | 2093177.592 | 2098178.671 | 2310546.332 | 1837981.814 | 1813841.522 | 2059912.99 | 1483488.998 |
| 4-methyloct-3-enoate | 599427.1271 | 424800.0539 | 511638.2097 | 3534651.83 | 84313.23434 | 817322.4218 | 4102012.11 | 62179.50321 | 1848340.994 | 1078103.06 | 224619.4676 | 2911215.635 |
| 7,8-Dihydro-.alpha. | 440443. | 59067.2 | 430573. | 349878. | 759655 | 781446 | 443702 | 592561 | 622586 | 572035 | 790391 | 615085 |

| Metabolites | PBS_1 | PBS_2 | PBS_3 | PBS_4 | BCG_1 | BCG_2 | BCG_3 | BCG_4 | BCGΔ*pe18*_1 | BCGΔ*pe18*_2 | BCGΔ*pe18*_3 | BCGΔ*pe18*_4 |
|---|---|---|---|---|---|---|---|---|---|---|---|---|
| -ionone | 1189 | 5981 | 9708 | 9971 | 9.424 | 6.006 | 0.563 | 4.779 | 7.843 | 2.415 | 1.414 | 2.081 |
| 3-(2,6-Dimethyl-4-morpholinyl)-N-phenylpropanamide | 2960746.209 | 2537140.99 | 2154394.204 | 2432289.738 | 2568960.503 | 2422510.686 | 2939186.062 | 3350361.74 | 2995260.192 | 2660875.097 | 5240147.178 | 2281451.626 |
| N,1-Dimethyl-4-nitro-1H-imidazol-5-amine | 3385515.828 | 3912354.32 | 3168845.259 | 3319585.52 | 3779731.177 | 3401250.353 | 3448588.429 | 3506683.459 | 4148353.88 | 4241201.656 | 3200497.961 | 3648095.533 |
| PE(P-36:4) | 505619757.1 | 593934696.4 | 520160150.7 | 489893195.6 | 414037865.8 | 315015526.7 | 495585006 | 384034434.4 | 372129030.9 | 490374527.9 | 449229609.3 | 209236901 |
| N-Cyclohexyl-N'-[3-(1H-imidazol-1-yl)propyl]thiourea | 4354125.343 | 5156584.008 | 2243725.652 | 3396688.773 | 3010709.345 | 5266480.885 | 2991002.73 | 3478165.607 | 3475831.063 | 2445224.486 | 3105235.514 | 4326306.674 |
| PS 3:0_19:2 | 167494.0034 | 129171.105 | 184472.9488 | 163244.5351 | 173658.3721 | 227038.7797 | 175561.1567 | 181426.836 | 154932.0407 | 164609.4467 | 150283.2363 | 91543.47902 |
| PI 8:0_24:0 | 1941548.207 | 1576975.077 | 857840.9208 | 1291035.962 | 1928769.474 | 1469297.851 | 1736772.311 | 2230188.212 | 1987322.527 | 1989677.87 | 2097928.116 | 1932566.681 |
| Crotamiton | 10826178.98 | 10802057.97 | 9689848.393 | 10183902.64 | 10301866.9 | 10098552.07 | 10254396.58 | 10407539.67 | 10303991.27 | 9918064.073 | 9871568.094 | 9126626.738 |
| N-(3-Hydroxy-2-methylbenzoyl)-L-valyl-N-[(1S,2E)-4-ethoxy-1-(2-methylpropyl)-4-oxo-2-butenyl]-L-serinamide | 582495.3081 | 599513.4959 | 756014.6264 | 712526.665 | 828597.743 | 976710.1371 | 1007388.232 | 812938.3964 | 714867.084 | 688273.7411 | 789582.8862 | 607830.7908 |

| **Metabolites** | **PBS_1** | **PBS_2** | **PBS_3** | **PBS_4** | **BCG_1** | **BCG_2** | **BCG_3** | **BCG_4** | **BCGΔ*pe18*_1** | **BCGΔ*pe18*_2** | **BCGΔ*pe18*_3** | **BCGΔ*pe18*_4** |
|---|---|---|---|---|---|---|---|---|---|---|---|---|
| Lutein 5,6-epoxide | 2668097.948 | 3151298.483 | 1922782.464 | 3424649.531 | 1530394.895 | 1401504.838 | 2001255.311 | 1692151.777 | 1856652.669 | 0 | 2117983.46 | 940383.1133 |
| Menthyl isovalerate | 675882.9373 | 665519.549 | 699858.5938 | 608890.1381 | 604168.2441 | 582280.4619 | 634635.9355 | 648391.9561 | 596966.0739 | 629621.3882 | 660408.5929 | 554093.0422 |
| (2R)-2,4-Dihydroxy-3,3-dimethylbutanoic acid | 700896.3388 | 667228.5756 | 435870.7919 | 507738.5645 | 7611961.492 | 4741156.166 | 6310991.401 | 5472569.818 | 4972057.733 | 5981152.113 | 4879347.984 | 3906989.877 |
| Benzimidazo[2,1-b]quinazolin-12(5H)-one | 1375994.776 | 4878640.387 | 7730782.245 | 2589809.973 | 790662.553 | 764529.7546 | 3773934.5 | 1630770.687 | 3634096.809 | 2704672.811 | 2139953.11 | 12678704.71 |
| Cer 37:1;2O/40:7 | 2851108.584 | 2783357.545 | 3080093.79 | 2942382.74 | 2611609.417 | 2652625.817 | 2609285.794 | 2756400.076 | 2744366.882 | 2700053.604 | 2647725.544 | 2644796.767 |
| Glu-Leu | 481830.354 | 451271.148 | 551809.6051 | 484159.7734 | 578401.261 | 430353.5004 | 648964.0258 | 488051.8855 | 657380.1591 | 772713.8448 | 775437.5781 | 666583.2311 |
| Mono(2-ethylhexyl) phthalate | 9679141.836 | 12762729.37 | 14589908.74 | 10769428.87 | 18025017.22 | 20670732.11 | 14489301.48 | 17576198.45 | 16096234.73 | 12884920.83 | 15642578.44 | 13962998.71 |
| CMP-2-aminoethyl phosphonate | 812135.0032 | 581757.8774 | 1401776.28 | 707987.6566 | 3266667.352 | 1465521.272 | 3137515.888 | 2009018.785 | 1675072.252 | 2148068.391 | 2772173.421 | 1618004.798 |
| 5-Acetoacetamido-2-benzimidazolone | 2486167.247 | 2278596.44 | 3487793.122 | 2229806.84 | 7802892.737 | 6000330.062 | 8052319.665 | 6352734.837 | 6416389.214 | 6666494.149 | 7071447.968 | 6260721.118 |
| (1,3-Benzothiazol-2-ylsulfonyl)acetic acid | 1684474.971 | 1560159.271 | 2243653.699 | 1935623.597 | 3486697.934 | 2632699.214 | 3320339.063 | 2962775.639 | 2593129.391 | 2499208.345 | 2539775.568 | 1989657.295 |

| **Metabolites** | **PBS_1** | **PBS_2** | **PBS_3** | **PBS_4** | **BCG_1** | **BCG_2** | **BCG_3** | **BCG_4** | **BCGΔ*pe18*_1** | **BCGΔ*pe18*_2** | **BCGΔ*pe18*_3** | **BCGΔ*pe18*_4** |
|---|---|---|---|---|---|---|---|---|---|---|---|---|
| 1-O-Hexadecyl-sn-glycero-3-phosphocholine (LPC(O-16:0/0:0)) | 23242488.68 | 24492505.08 | 15324023.24 | 21006356.06 | 15916800.24 | 12219072.24 | 15822386.1 | 16121172.71 | 14997455.38 | 14896131.92 | 14165023.01 | 10120341.6 |
| 2,3 Dihydroamentoflavone | 4741847.304 | 4787869.477 | 4991154.811 | 5493905.351 | 4547296.781 | 4166813.139 | 5237585.612 | 4651685.139 | 5053469.32 | 4863069.953 | 4534431.3 | 5339799.182 |
| MGDG O-24:2_11:0 | 310947.5693 | 79267.38227 | 304092.9308 | 342197.6709 | 2268931.427 | 2216821.807 | 2017924.235 | 1604946.517 | 1562830.601 | 2021499.628 | 3845933.531 | 973555.9538 |
| (22E,24R)-Stigmasta-4,22-diene-3,6-dione | 1594851.877 | 1162541.829 | 2354445.538 | 1518359.943 | 1589050.846 | 1419013.26 | 1975655.554 | 951932.8698 | 1017246.29 | 1285529.564 | 1236923.262 | 711507.2095 |
| PC O-32:1 | 10844280.96 | 12900268.08 | 5010695.904 | 11716310.43 | 9560920.277 | 262412.4139 | 8596570.806 | 11581935.18 | 28430066.8 | 5448887.572 | 8521330.514 | 7743066.789 |
| PG O-22:5_20:4 | 2244094.208 | 1677877.43 | 1959987.404 | 1943160.844 | 1723763.722 | 20897.38895 | 1965884.025 | 1152001.851 | 2268395.941 | 2031918.764 | 1875492.2 | 1680426.317 |
| DGDG O-10:0_16:3 | 443575.7961 | 224738.8089 | 682543.8416 | 275296.8908 | 645070.3981 | 577091.2251 | 936392.3614 | 826495.0342 | 1466961.37 | 1664084.304 | 739566.9452 | 1016687.421 |
| 4-Amino-2,8-bis(trifluoromethyl)quinoline | 356380.7264 | 176250.2898 | 498785.8594 | 419956.2477 | 407074.4778 | 400982.7403 | 508219.3293 | 329233.0886 | 364003.5866 | 374925.1746 | 435482.324 | 529371.0849 |
| (E)-9-(1,3-Benzodioxol-5-yl)non-8-enoic acid | 100922.9065 | 143067.3199 | 171142.7629 | 163426.2 | 3288961.498 | 2915462.27 | 2552924.01 | 2362620.698 | 1893615.312 | 2080183.553 | 1947282.328 | 2173289.498 |

| Metabolites | PBS_1 | PBS_2 | PBS_3 | PBS_4 | BCG_1 | BCG_2 | BCG_3 | BCG_4 | BCGΔ*pe18*_1 | BCGΔ*pe18*_2 | BCGΔ*pe18*_3 | BCGΔ*pe18*_4 |
|---|---|---|---|---|---|---|---|---|---|---|---|---|
| Prunetin | 6527120.652 | 2744788.564 | 2827861.395 | 6947230.075 | 3212122.448 | 2776268.168 | 2564699.28 | 6504413.463 | 3299021.769 | 3764579.627 | 4599272.683 | 4468462.991 |
| 8-Methylquinolin-2(1H)-one | 7983488.682 | 6562862.329 | 6016866.261 | 5492727.115 | 13422647.19 | 10070609.09 | 17502534.5 | 16778631.6 | 14306354.86 | 9267069.124 | 11865153.1 | 7445462.376 |
| Heptadecanoic acid | 29014712.13 | 24761710.35 | 16421579.19 | 26643330.82 | 23897126.62 | 38232779.56 | 25584083.65 | 24244094.31 | 17134123.09 | 15601927.31 | 31700380.91 | 20902492.83 |
| alpha-Naphthoflavone | 357831.3653 | 264068.9757 | 377623.8767 | 340965.4438 | 1024765.772 | 798107.7319 | 1036973.818 | 910629.0943 | 872848.0837 | 708608.5047 | 743585.24 | 577597.8473 |
| 3-Piperidone | 9528731.209 | 7402561.901 | 7200607.219 | 6848821.838 | 1885827.934 | 7923165.069 | 14900093.47 | 7022882.024 | 6970240.726 | 6813564.538 | 5338582.954 | 7605537.156 |
| 3-Hydroxy-carbofuran | 2003814.846 | 2017397.892 | 2748359.219 | 2322961.58 | 2012621.66 | 2153482.058 | 1978535.977 | 1946746.483 | 2011423.039 | 1813902.942 | 2156645.022 | 2313868.184 |
| 2-amino-4-phosphonobutanoate | 2160262.589 | 1939400.679 | 2259273.385 | 1887770.612 | 3442813.408 | 2062099.723 | 2825831.839 | 2687697.665 | 2321696.193 | 2567344.877 | 2494179.837 | 1446721.796 |
| SL 22:2;O/10:0 | 1206535.302 | 1223666.873 | 1178652.629 | 1231455.74 | 1188829.29 | 1143635.521 | 1436296.642 | 1220474.895 | 1225672.454 | 1234984.329 | 1122198.358 | 1405229.529 |
| 1,3,5-Trimethyl-2,6-diphenyl-4-piperidinol | 13979314.55 | 15113062.96 | 13222781.98 | 13959929.64 | 13231733.11 | 14389911.14 | 14904546.96 | 14076031.94 | 13770488.22 | 13892825.96 | 12932708.14 | 13185014.42 |
| PC 16:0_20:6;O | 1655251.089 | 1785807.523 | 1660859.26 | 1650979.4 | 1744881.361 | 1721830.711 | 1683103.889 | 1680022.517 | 1649393.26 | 1685567.452 | 1760749.745 | 1595066.276 |
| Cer 18:1;2O/24:2 | 4014979.609 | 6454987.74 | 6202693.438 | 6614535.778 | 5781463.656 | 6203028.042 | 4013179.191 | 4667345.77 | 4509221.863 | 3621834.377 | 11219.027 | 6522909.794 |
| Quinoclamine | 449906 | 217125 | 170371 | 719547 | 130370 | 182376 | 138659 | 138606 | 110985 | 316911 | 125775 | 155070 |

| Metabolites | PBS_1 | PBS_2 | PBS_3 | PBS_4 | BCG_1 | BCG_2 | BCG_3 | BCG_4 | BCGΔ*pe18*_1 | BCGΔ*pe18*_2 | BCGΔ*pe18*_3 | BCGΔ*pe18*_4 |
|---|---|---|---|---|---|---|---|---|---|---|---|---|
| | 0.059 | 6.402 | 33.74 | 5.128 | 48.15 | 08.67 | 72.4 | 81.99 | 95.14 | 4.47 | 20.04 | 04.02 |
| (3-Bromophenyl)(3-fluorophenyl)methanone | 895936.7354 | 466862.1566 | 1487442.633 | 860148.1694 | 12984473.97 | 10958504.27 | 13594576.55 | 11501160.98 | 8894982.524 | 9427300.245 | 13658641.08 | 9240978.454 |
| SHexCer 38:4;3O | 184890895.7 | 210486567.9 | 87685846.33 | 151979862.5 | 14285726.23 | 14264246.02 | 36181104.72 | 35158255.22 | 56427792.05 | 29566343.85 | 26469666.67 | 27579944.82 |
| Phosphocreatine | 3255893.14 | 2555524.17 | 4254388.534 | 2984600.107 | 19647333.81 | 11740325.88 | 19795160.17 | 14188302.37 | 11498125.52 | 15199963.1 | 14764189.95 | 10658961.36 |
| sclareol | 2478313.23 | 2790402.345 | 3086542.516 | 2756191.335 | 12101189.05 | 14297802.26 | 11551299.94 | 10358959.54 | 7945334.026 | 7834766.17 | 11632139.32 | 6740287.581 |
| PC 17:2_17:2 | 19472352.34 | 24410338.61 | 13134164.08 | 17022313.04 | 41198412.63 | 50327144.08 | 50925653.48 | 46774126.6 | 43372967.71 | 51991965.13 | 42871578.03 | 29846789.37 |
| 1,2-Didocosahexaenoyl-sn-glycero-3-phosphocholine | 1157740.21 | 1664571.87 | 366473.0751 | 921651.8387 | 2821899.575 | 2951741.755 | 1585181.42 | 2736990.531 | 2580665.916 | 2299648.555 | 1713926.806 | 1667007.154 |
| Cyclovirobuxin D | 849952.4215 | 1150830.385 | 548610.7286 | 686467.3109 | 958884.2356 | 732129.7107 | 1112184.645 | 1107353.981 | 1424539.541 | 1025940.932 | 909781.445 | 596971.4084 |
| 1-(3,5-Dimethyl-1H-pyrazol-4-yl)ethanamine | 3556025.889 | 3176655.176 | 1965086.178 | 3414039.284 | 3414317.839 | 2835159.764 | 3261531.175 | 3040352.851 | 2444743.039 | 2819517.24 | 1654115.027 | 3463229.855 |
| Cysteine-glutathione disulfide | 2054889.387 | 1671378.252 | 4960742.919 | 2253429.698 | 7014149.836 | 12470849.21 | 1916168.391 | 1766836.757 | 5682301.873 | 5466493.175 | 5308208.894 | 5446154.713 |
| Isopyrazam | 261899.4947 | 282188.237 | 336553.8896 | 200313.0138 | 880008.9876 | 580229.5293 | 1080935.03 | 820762.8663 | 884269.5695 | 1060938.876 | 1115282.892 | 777213.9119 |

| Metabolites | PBS_1 | PBS_2 | PBS_3 | PBS_4 | BCG_1 | BCG_2 | BCG_3 | BCG_4 | BCGΔ*pe18*_1 | BCGΔ*pe18*_2 | BCGΔ*pe18*_3 | BCGΔ*pe18*_4 |
|---|---|---|---|---|---|---|---|---|---|---|---|---|
| PE 8:0_15:1 | 455301.792 | 325755.774 | 399852.9139 | 343169.8711 | 4244556.974 | 1704504.646 | 3159695.63 | 2062979.083 | 2217275.225 | 2673974.751 | 3535515.41 | 1710226.321 |
| PMeOH 16:3_20:4 | 10146688.26 | 10748568.03 | 8130869.177 | 8828529.57 | 9611641.849 | 9334849.239 | 9424511.853 | 9941715.804 | 9012217.616 | 9072519.026 | 8995412.515 | 8402438.735 |
| Glycerophosphocholine | 21643068.83 | 14427660.59 | 20037410.26 | 13110271.95 | 54130503.34 | 29575820.99 | 41078859.06 | 40306251.63 | 37860214.73 | 34467706.7 | 30083305.25 | 22031426.22 |
| Purpurin | 541001.3616 | 427505.2898 | 876918.1052 | 557604.3787 | 456448.591 | 685532.4953 | 595783.7332 | 642254.5848 | 794725.1375 | 659647.1837 | 818607.4363 | 722568.9745 |
| PC O-22:5 | 1244498.911 | 1623574.149 | 2896343.858 | 1172220.158 | 1343054.332 | 1378533.957 | 2082124.665 | 1523675.033 | 1188847.689 | 1605561.204 | 2176581.913 | 1058802.956 |
| cis-11-Eicosenoic acid | 4258147.981 | 5239092.69 | 5234650.067 | 6353618.579 | 78164829.44 | 98354984.84 | 70152328.85 | 96228349.24 | 65891293.53 | 37449552.88 | 76542747.08 | 77017560.53 |
| LPC 22:5 | 3260742.168 | 2706445.164 | 3752543.615 | 2890484.239 | 2609137.607 | 3137521.082 | 3354259.693 | 3041379.142 | 3825454.335 | 2165650.194 | 2585889.967 | 2699272.845 |
| 5'-Dehydroadenosine | 10378214.06 | 11700783.98 | 10104188.7 | 9850856.775 | 8543685.763 | 8512950.38 | 8241015.054 | 8866610.434 | 9234957.308 | 6968458.358 | 8343668.818 | 9532875.249 |
| 2-Propylphenol | 733404.6185 | 628096.5642 | 503537.7124 | 2523362.559 | 106489.6682 | 1037775.732 | 4110776.617 | 524676.7459 | 2468132.426 | 2838176.204 | 211627.107 | 2803584.4 |
| 3-Oxocyclobutanecarboxylic acid | 623971.2741 | 598873.1665 | 633515.8558 | 612739.8777 | 1279939.271 | 943098.0064 | 1162499.73 | 1089429.005 | 862562.4499 | 853462.5824 | 791590.3714 | 900041.5439 |
| PE(22:6(4Z,7Z,10Z,13Z,16Z,19Z)/20:5(5Z,8Z,11Z,14Z,17Z)) | 1189350.372 | 1714103.249 | 1181882.173 | 1260043.889 | 2114840.209 | 1663963.335 | 1745321.826 | 2631813.833 | 2239265.393 | 2466358.898 | 1861381.08 | 1167191.501 |

| **Metabolites** | **PBS_1** | **PBS_2** | **PBS_3** | **PBS_4** | **BCG_1** | **BCG_2** | **BCG_3** | **BCG_4** | **BCGΔ*pe18*_1** | **BCGΔ*pe18*_2** | **BCGΔ*pe18*_3** | **BCGΔ*pe18*_4** |
|---|---|---|---|---|---|---|---|---|---|---|---|---|
| Phencyclidine | 1141458.517 | 1083126.437 | 997024.3903 | 1040367.13 | 1214296.096 | 1146404.646 | 1112911.614 | 1100670.364 | 1101841.079 | 1085497.638 | 1106939.719 | 1115378.746 |
| 1-(4-Bromo-2-fluorophenyl)cyclopropanecarboxylic acid | 1808640.834 | 1797264.652 | 1940452.726 | 1094861.598 | 2048217.165 | 2136132.109 | 1084040.489 | 1192957.419 | 1130502.586 | 1915010.609 | 1827571.92 | 1791187.632 |
| LPE 38:8 | 1553738.902 | 1083478.708 | 1448461.202 | 1376506.172 | 1694581.262 | 0 | 1582625.633 | 2282525.35 | 1537186.022 | 1550895.5 | 1647737.152 | 1446064.318 |
| 5-Fluoro-1H-pyrazolo[3,4-b]pyridin-3-amine | 9641267.725 | 10843066.12 | 9214924.034 | 9259608.339 | 9926311.537 | 9646308.39 | 9723215.824 | 9876728.801 | 9464142.331 | 9886250.193 | 10231343.01 | 13526354.42 |
| LNAPE 4:0/N-19:2 | 1213758.569 | 1340515.972 | 960203.8823 | 1240011.736 | 1351892.525 | 1821885.138 | 1373522.068 | 1355139.429 | 1106733.758 | 1389561.397 | 1246037.762 | 827390.1742 |
| PS 24:5 | 269641.5047 | 217220.1963 | 231139.967 | 254403.982 | 263737.1509 | 238860.2093 | 255449.5185 | 242746.314 | 263553.2432 | 288548.5267 | 325486.0353 | 223126.9832 |
| 1-O-Octadecyl-sn-glyceryl-3-phosphorylcholine | 47905954 | 52482281.06 | 24642341.7 | 36750392.74 | 22515846.96 | 27549002.33 | 25079055.7 | 22190770.98 | 21372497.46 | 22571544.39 | 22268246 | 13846769.66 |
| Cer 13:0;2O/4:0 | 5034447.226 | 5299249.815 | 5120623.643 | 5069953.905 | 4973151.843 | 5210837.601 | 5037484.92 | 5226705.666 | 5057565.187 | 5262567.48 | 4975273.989 | 5182110.851 |
| PC(36:5) | 77552455.24 | 80660697.8 | 42143262.16 | 57589425.16 | 172810413.3 | 128912549.4 | 179832447.4 | 169310664.7 | 189237802.1 | 69250221.03 | 141312089.2 | 131987430 |
| Butylate | 4860217.195 | 1755562.75 | 2639637.7 | 2240072.14 | 207761.6985 | 6905953.057 | 5855440.674 | 6508719.48 | 326030.2857 | 878217.9632 | 454159.6386 | 23437992.03 |
| Trifluoroacetic acid | 444961 | 587143 | 538290 | 542987 | 600091 | 600771 | 717732 | 614689 | 582768 | 583438 | 650296 | 565738 |

| **Metabolites** | **PBS_1** | **PBS_2** | **PBS_3** | **PBS_4** | **BCG_1** | **BCG_2** | **BCG_3** | **BCG_4** | **BCGΔ*pe18*_1** | **BCGΔ*pe18*_2** | **BCGΔ*pe18*_3** | **BCGΔ*pe18*_4** |
|---|---|---|---|---|---|---|---|---|---|---|---|---|
| | 9.892 | 1.691 | 3.51 | 6.041 | 6.112 | 9.645 | 2.827 | 4.59 | 1.063 | 2.843 | 1.378 | 5.557 |
| Neomangiferin | 115181.2787 | 124373.7246 | 192178.8469 | 151539.0874 | 1519195.075 | 1237522.501 | 1277780.393 | 1117606.195 | 1045949.305 | 1070257.815 | 817602.304 | 1411619.128 |
| 2-(3-Carboxypropionylamino)-4-methylsulfanylbutyric acid | 110511.5574 | 95530.1526 | 48745.83168 | 100929.0975 | 3056276.36 | 4362800.418 | 2881076.424 | 1443271.954 | 3713639.025 | 5429222.044 | 4737278.468 | 3037379.005 |
| risedronate | 25667713.13 | 22023010.48 | 31149936.78 | 24175338.07 | 39275750.97 | 31673519.51 | 41262454.92 | 34022890.75 | 30319825.7 | 33940092.44 | 36039029.03 | 27813947.28 |
| Oxopentadecanoylcarnitine (Car(15:1-O)) | 3914108.949 | 4017227.527 | 3335687.649 | 3718693.027 | 4114066.046 | 3209075.671 | 3709127.514 | 3998822.309 | 3792802.232 | 3816142.385 | 3797281.937 | 3179649.282 |
| gamma-Glutamylglutamine | 325758.6685 | 445602.9545 | 482036.6619 | 335923.8394 | 10566203.57 | 7924314.309 | 7252666.455 | 6635293.311 | 8669310.551 | 6968547.792 | 7442095.972 | 9053453.515 |
| PE O-16:2_22:6 | 6566084.689 | 3966451.212 | 9539929.167 | 4133811.411 | 4632394.63 | 5775740.431 | 6418260.966 | 6185562.669 | 2709408.025 | 4748564.008 | 5795341.191 | 3778482.144 |
| LPE O-22:5 | 1762901.515 | 1938057.427 | 1163887.452 | 1640840.854 | 7283805.027 | 8494752.444 | 10944162.86 | 7476685.28 | 5035469.254 | 4332898.65 | 9574484.476 | 5435977.725 |
| PC 8:0_22:1 | 12917650.45 | 14809860.15 | 10528344.9 | 11278148.14 | 66354202.27 | 59361657.11 | 69932103.3 | 55050826.06 | 65695712.33 | 70940047.66 | 93488057.54 | 46369035.52 |
| N,N-Dimethylformamide | 9649849.276 | 14280806.25 | 15562355.01 | 11829561.49 | 1583040.923 | 15671062.57 | 15422285 | 13265468.21 | 16201706.76 | 7526274.499 | 2272487.831 | 14817830.12 |
| Cer 14:1;2O/9:0 | 9557451.778 | 10584521.57 | 8443455.161 | 8706212.464 | 9226638.099 | 9551328.762 | 9507285.092 | 10155060.39 | 8997193.906 | 9317657.329 | 9544928.494 | 8766656.091 |
| 6-methyloctanoate | 273586 | 285902 | 254941 | 671991 | 204620 | 307654 | 713871 | 485242 | 345314 | 265661 | 182148 | 658689 |

| **Metabolites** | **PBS_1** | **PBS_2** | **PBS_3** | **PBS_4** | **BCG_1** | **BCG_2** | **BCG_3** | **BCG_4** | **BCGΔ*pe18*_1** | **BCGΔ*pe18*_2** | **BCGΔ*pe18*_3** | **BCGΔ*pe18*_4** |
|---|---|---|---|---|---|---|---|---|---|---|---|---|
| | 6.019 | 4.357 | 7.091 | 0.227 | 7.907 | 3.816 | 2.766 | 4.786 | 2.065 | 6.946 | 5.78 | 2.542 |
| Cer 8:1;2O/18:5 | 154050.3394 | 153422.12 | 148081.2726 | 147572.7677 | 144461.4217 | 155295.7505 | 184864.8628 | 150093.1785 | 152279.4014 | 152490.2747 | 152353.7552 | 191297.0843 |
| Pentylenetetrazole | 1201599.203 | 1303699.77 | 1728510.014 | 1673422.768 | 4216725.589 | 2911727.741 | 4579622.253 | 3234667.231 | 3050427.811 | 3919469.925 | 3026418.397 | 1348684.138 |
| 1-Stearoyl-2-linoleoyl-sn-glycero-3-phosphocholine | 3683447.27 | 1504686.843 | 1159580.79 | 1543108.02 | 5609964.358 | 6722104.866 | 4763398.844 | 3660160.604 | 6019438.321 | 5998443.192 | 2353758.814 | 4158726.235 |
| Embelin | 120540278.4 | 116053776.3 | 111001177.2 | 159833472.5 | 106476184.6 | 145326256.4 | 114535146.5 | 155941197.2 | 131542620.8 | 171732409.6 | 105158114.2 | 128056904.4 |
| PC 19:0_18:3 | 4702391.646 | 1767383.638 | 2909069.565 | 2487239.608 | 5204805.897 | 5927852.65 | 2870682.261 | 4331370.764 | 4071710.338 | 3646534.684 | 3445751.499 | 2133261.905 |
| PC 9:0_28:3 | 3776641.497 | 2340273.383 | 4118653.688 | 2791935.258 | 6177663.139 | 6275452.575 | 4805210.81 | 4377904.505 | 6202089.331 | 4803303.258 | 2693041.874 | 2971642.163 |
| SHexCer 35:0;2O | 2790267.472 | 2912573.619 | 2259418.062 | 2692896.775 | 2538807.099 | 25586.3152 | 2074316.744 | 1908863.711 | 3162712.145 | 2372691.196 | 1878816.759 | 1674844.208 |
| Serotonin | 3040484.556 | 2397755.889 | 2278300.676 | 2043454.341 | 5319071.553 | 3789349.765 | 6872960.706 | 6451000.827 | 5578177.579 | 3493867.664 | 4422598.653 | 2759976.968 |
| PG 18:0_22:6 | 7348612.513 | 948871.5505 | 875857.2769 | 8860931.989 | 7621167.037 | 2396378.276 | 23765592.38 | 5824892.202 | 9944080.026 | 13589468.3 | 7162027.151 | 4230472.937 |
| Oroxyloside | 5412694.287 | 5082201.343 | 5751884.589 | 5363750.264 | 5469596.017 | 5292175.937 | 5306271.895 | 5317400.583 | 5140275.969 | 5042070.881 | 5109300.93 | 5376256.883 |
| Diallyl Sulfide | 6283768.39 | 3177395.771 | 8612539.488 | 6277418.483 | 65408582.85 | 61550833.81 | 157608189 | 52972301.78 | 57465969.59 | 78780546.31 | 71109910.74 | 31455117.88 |

| Metabolites | PBS_1 | PBS_2 | PBS_3 | PBS_4 | BCG_1 | BCG_2 | BCG_3 | BCG_4 | BCGΔ*pe18*_1 | BCGΔ*pe18*_2 | BCGΔ*pe18*_3 | BCGΔ*pe18*_4 |
|---|---|---|---|---|---|---|---|---|---|---|---|---|
| 2,2'-[(Phosphonomethyl)imino]diacetic acid | 661804.7031 | 455247.5596 | 910779.6278 | 697892.3713 | 2812541.373 | 1730310.549 | 2487833.079 | 2277863.188 | 1948665.095 | 1697486.019 | 1761910.647 | 1489495.546 |
| N-Acetylaspartic acid | 11114293.87 | 7111107.296 | 10218059.37 | 12019754.29 | 10299830.9 | 11005063.49 | 9390426.401 | 9239025.807 | 6477703.817 | 10855362.16 | 7212486.717 | 4659532.618 |
| gamma-Glutamyl-propargylglycine | 2636516.441 | 2414316.481 | 4383500.611 | 2920324.32 | 3244758.278 | 2760103.927 | 3554727.437 | 3092540.072 | 2891429.329 | 2334322.98 | 2522195.033 | 3059452.651 |
| SM(d17:1/24:1(15Z)) | 4684119.805 | 5198315.744 | 6038989.645 | 4702106.705 | 5745355.801 | 4789066.122 | 6888844.451 | 5979110.147 | 5643336.12 | 7111248.273 | 5490999.15 | 2643581.805 |
| Pinacidil | 3668854.743 | 2618052.668 | 4663690.977 | 3417097.83 | 21271223.97 | 12075829.03 | 17805055.47 | 15764437.81 | 13279909.57 | 12491179.64 | 14346358.31 | 10464915.51 |
| LNAPS 24:4/N-14:1 | 3376158.93 | 3377776.211 | 3226507.315 | 3397457.751 | 3392338.207 | 3406823.818 | 3603074.604 | 3184844.986 | 3452521.628 | 3405617.927 | 3421928.02 | 2992081.364 |
| Caproic acid | 11200798.14 | 11753945.74 | 10540995.71 | 22623580.32 | 12857259.35 | 12596840.34 | 30523389.54 | 10663412.18 | 18247375.38 | 13504266.64 | 11089852.81 | 30914008.55 |
| LPC(20:4) | 4918089.323 | 3146550.197 | 4043058.486 | 2392061.552 | 4575244.105 | 1876657.394 | 3155239.916 | 5569975.637 | 2110504.112 | 3909579.541 | 3087900.855 | 1675629.788 |
| PC O-14:0_2:0 | 49056644.95 | 53384667.49 | 62245576.85 | 133005286.4 | 160211785.3 | 126697847.5 | 136785087.4 | 113263982.3 | 55905578.07 | 138658716.7 | 77814149.71 | 102854749.5 |
| Triethylamine | 27805021.91 | 38840905.68 | 41450153.84 | 37555026.9 | 212222283.3 | 37213990.45 | 34045237.78 | 90639942.93 | 103975127 | 29724941.14 | 30858573.72 | 94722001.76 |
| 2-oxobut-3-enanoate | 2299008.495 | 2728870.786 | 1963991.21 | 3201245.119 | 4453403.975 | 2853565.145 | 10930736.46 | 4529398.87 | 4807268.464 | 4296275.055 | 6655191.582 | 4042781.921 |
| 2-hydroxytridecano | 116423 | 103565 | 248741 | 930563. | 326133 | 613847 | 274725 | 406001 | 240952 | 102980 | 428795 | 427995 |

| Metabolites | PBS_1 | PBS_2 | PBS_3 | PBS_4 | BCG_1 | BCG_2 | BCG_3 | BCG_4 | BCGΔ*pe18*_1 | BCGΔ*pe18*_2 | BCGΔ*pe18*_3 | BCGΔ*pe18*_4 |
|---|---|---|---|---|---|---|---|---|---|---|---|---|
| ate | 1.566 | 4.033 | 7.992 | 3183 | 75.3 | 17.81 | 30.72 | 25.72 | 53.22 | 35.5 | 42.12 | 88.72 |
| 6a_Methylprednisolone | 841389.8381 | 841565.9474 | 598610.4283 | 567880.5498 | 2491697.451 | 3255747.221 | 1822784.052 | 2729300.825 | 3503275.076 | 1125439.317 | 1594428.601 | 4040356.573 |
| 2-Piperidone | 9549017.035 | 10265450.68 | 10802598.51 | 11008992.47 | 11030403.62 | 10547050.92 | 9410114.132 | 9620081.981 | 9865030.808 | 10427566.57 | 10770635.83 | 10456074.65 |
| PE(P-18:0/22:4(7Z,10Z,13Z,16Z)) | 31836754.11 | 48312533.22 | 59314201.31 | 49675253.61 | 46209545.85 | 37360002.79 | 64664993.56 | 46761072.8 | 47244757.56 | 65246866.54 | 60731260.14 | 22849298.04 |
| (5S)-1,2,3,4,5,6,7,8,9,10-Decahydro-5,9-methanobenzo[8]annulen-11-one | 15965655.29 | 13280042.12 | 13956504.81 | 17253931.03 | 8449784.484 | 10260858.94 | 9477446.092 | 9641283.297 | 9168395.512 | 9253737.102 | 7613305.898 | 5806398.795 |
| Cuscohygrine | 13828623.6 | 16516615.28 | 16139152.04 | 18050120.71 | 16748233.41 | 15127942.25 | 17100823.08 | 16479173.83 | 17177671.32 | 11852788.57 | 15804739.9 | 9018854.215 |
| PI O-22:3_26:6 | 0 | 0 | 0 | 0 | 1102468.042 | 883811.349 | 1153215.961 | 1336072.741 | 1654440.139 | 1448832.504 | 1685092.149 | 1469309.918 |
| Methylglycinate | 286241.5306 | 1419918.107 | 744309.5893 | 1391237.015 | 1379865.449 | 661180.8537 | 1315579.13 | 1227774.921 | 1422553.426 | 1473449.797 | 1441426.666 | 1379092.515 |
| O-Methyltyramine | 578845.4258 | 660176.6827 | 617982.796 | 2264398.304 | 528591.8367 | 665326.6868 | 3126352.667 | 503365.1033 | 1470439.636 | 943239.5624 | 582827.7435 | 2674813.087 |
| 2-Acetoxy-3-geranylgeranyl-1,4-dihydroxybenzene | 73532.08711 | 77221.63201 | 94160.58984 | 49852.90334 | 71991.40621 | 78436.8812 | 83214.18677 | 99490.04582 | 74296.18796 | 52856.79114 | 76548.36897 | 62964.07 |
| LPG 16:0 | 9998172.631 | 8052524.225 | 6838993.836 | 6809669.288 | 36551524.88 | 35379855.61 | 53778123.4 | 36713789.31 | 30543705.13 | 27980978.6 | 35096217.61 | 26545252.15 |

| Metabolites | PBS_1 | PBS_2 | PBS_3 | PBS_4 | BCG_1 | BCG_2 | BCG_3 | BCG_4 | BCGΔ*pe18*_1 | BCGΔ*pe18*_2 | BCGΔ*pe18*_3 | BCGΔ*pe18*_4 |
|---|---|---|---|---|---|---|---|---|---|---|---|---|
| N-palmitoyl-threonine | 20128887.69 | 20242667.27 | 13501847.35 | 14538203.06 | 18100906.94 | 17185325.41 | 13809401.09 | 17918614.92 | 18011044.35 | 19653151.27 | 19925179.32 | 12852024.91 |
| 2-(Cyclobutylamino)-N-[(2S)-3-(3,4-dihydro-1H-isoquinolin-2-yl)-2-hydroxypropyl]isonicotinamide | 425397.1902 | 415874.5132 | 397958.8532 | 1595438.126 | 1288442.95 | 900965.4358 | 1471081.487 | 777516.2381 | 1069295.413 | 710120.3747 | 369420.6336 | 2135913.535 |
| LPC(O-18:1) | 11490575.94 | 12370823.31 | 11502414.24 | 12214240.39 | 8488639.664 | 6871477.791 | 7988574.395 | 7455321.816 | 7826677.134 | 7242577.567 | 8234311.508 | 5846418.955 |
| 6-Oxocativic acid | 1247415.616 | 1582383.993 | 1783256.508 | 1784017.718 | 3350307.056 | 4762853.628 | 2870668.297 | 3656416.864 | 4096837.235 | 2144971.081 | 3385474.628 | 4322022.07 |
| S-Formylglutathione | 2567704.915 | 1599805.61 | 3140526.977 | 1960522.445 | 6788873.237 | 4362620.526 | 5287188.567 | 5498690.196 | 5687400.987 | 4646693.36 | 4713244.785 | 4227905.486 |
| Montecristin | 386572.6604 | 318919.1937 | 36046.24985 | 136033.7535 | 1475663.931 | 1357234.066 | 1878134.464 | 1752850.092 | 2445346.845 | 2231138.525 | 1396568.722 | 1248820.4 |
| Irgarol | 1826331.509 | 2013771.112 | 1619888.283 | 1668134.924 | 1764434.297 | 1791467.21 | 1562965.743 | 1713110.547 | 1690248.985 | 1676853.765 | 1650573.612 | 1638864.719 |
| PS O-22:5_17:1 | 700189.677 | 840302.7773 | 837301.9214 | 737831.1547 | 343755.9533 | 498855.5803 | 244487.9539 | 240941.5718 | 257884.6071 | 292635.6956 | 332096.9039 | 1518229.595 |
| CAR 18:1 | 63867670.52 | 54021632.73 | 95265718.86 | 63480895.54 | 127717233.2 | 88238510.38 | 135678443.2 | 118723037.8 | 74480291.98 | 95309418.49 | 94782635.41 | 59303272.06 |
| Cer 21:2;2O/10:0 | 327290.0243 | 278220.2897 | 197502.2136 | 135026.9737 | 510414.5146 | 981637.4577 | 515911.7902 | 647134.6254 | 548768.1542 | 414120.854 | 584927.1023 | 331382.5155 |

| **Metabolites** | **PBS_1** | **PBS_2** | **PBS_3** | **PBS_4** | **BCG_1** | **BCG_2** | **BCG_3** | **BCG_4** | **BCGΔ*pe18*_1** | **BCGΔ*pe18*_2** | **BCGΔ*pe18*_3** | **BCGΔ*pe18*_4** |
|---|---|---|---|---|---|---|---|---|---|---|---|---|
| 1-palmitoyl-2-linoleoyl-sn-glycero-3-phospho-(1'-rac-glycerol) | 5525809.806 | 7712520.907 | 2622184.658 | 3741762.511 | 49270109.64 | 50709781.11 | 23612998.79 | 20934865.76 | 27052545.18 | 31327374.55 | 27871710.51 | 15500446.58 |
| Ile-Glu | 1775974.049 | 1396047.207 | 2561763.658 | 1699289 | 37144348.84 | 33124334.44 | 26667065.22 | 27107504.66 | 23554756.62 | 18504620.79 | 18627534.23 | 24684663.98 |
| Radiclonic acid | 41935385.12 | 44539184.42 | 39269134.98 | 43193056.57 | 55080292.84 | 49584148.15 | 47638620.69 | 49894689.48 | 45462700.26 | 257184.7732 | 48822675.15 | 45742566.63 |
| 5-Hydroxydopamine | 15911226.1 | 12704663.27 | 23121152.81 | 18147374.86 | 17236011.37 | 19391181.69 | 17000426.76 | 16434887.16 | 17762636.49 | 8871611.376 | 7911308.646 | 17258106.48 |
| 3-Quinolinecarboxamide, 1,4-dihydro-8-methoxy-4-oxo-1-pentyl-N-tricyclo[3.3.1.13,7]dec-1-yl- | 108369.7941 | 65875.74213 | 112782.1177 | 41463.31031 | 183407.3152 | 251266.0839 | 498873.7712 | 168744.033 | 197809.9214 | 496807.5192 | 564892.7905 | 411279.908 |
| Lysine | 1240802.306 | 1497332.319 | 925045.499 | 1493839.663 | 4840600.222 | 3538754.079 | 4163300.16 | 3938325.155 | 4659470.454 | 2870445.917 | 3839594.79 | 4031866.266 |
| M337T522_1 | 4602057.266 | 3713217.605 | 4674194.47 | 5041424.479 | 3337907.533 | 5041712.615 | 3044659.677 | 4629119.85 | 3612288.304 | 2890320.024 | 3043378.173 | 13588814.6 |
| SM 9:1;2O/24:6 | 1583914.375 | 1910912.873 | 1529723.848 | 1669531.206 | 1625827.815 | 1703797.925 | 1772599.63 | 1652802.732 | 1618172.688 | 1552284.198 | 1789650.2 | 1477518.077 |
| M209T374_1 | 10539517.05 | 3069969.939 | 2970263.238 | 3328740.634 | 2865093.081 | 7068715.303 | 4961126.52 | 4666164.227 | 4848504.46 | 4183941.112 | 2968157.958 | 2096852.934 |

| Metabolites | PBS_1 | PBS_2 | PBS_3 | PBS_4 | BCG_1 | BCG_2 | BCG_3 | BCG_4 | BCGΔ*pe18*_1 | BCGΔ*pe18*_2 | BCGΔ*pe18*_3 | BCGΔ*pe18*_4 |
|---|---|---|---|---|---|---|---|---|---|---|---|---|
| Valine | 8379850.643 | 6473834.689 | 12519295.45 | 8379231.709 | 20119956.27 | 17589817.62 | 20992754.69 | 16885845.96 | 18389902.59 | 12327477.6 | 14416469.8 | 15318332.59 |
| Dioctylphthalate | 7341314.152 | 1807267.21 | 1990665.116 | 1797292.021 | 4782491.419 | 4299888.803 | 18357223.21 | 15392783.84 | 2725304.657 | 10793757.56 | 17533407.78 | 2593452.63 |
| 2-((1-(4-Nitrobenzoyl)prolyl)amino)benzoic acid | 6927.454553 | 20525.827 | 77245.89037 | 42857.90539 | 60954.84035 | 264630.5197 | 83834.36079 | 243605.0825 | 203997.5264 | 375487.4012 | 34263.10332 | 250378.0357 |
| L-Tyrosine, O-(1,1-dimethylethyl)- | 1369563.573 | 1516127.404 | 1326124.97 | 1449590.662 | 1286517.4 | 1268828.068 | 1524011.244 | 1251645.522 | 1289204.882 | 1379251.407 | 1217054.811 | 1371331.842 |
| SL 13:0;O/21:2 | 24045660.5 | 27540407.48 | 19596893.13 | 26631828.56 | 22685192.39 | 25636896.28 | 21539543.72 | 20952023.43 | 21371089.12 | 19881545.96 | 20871204.78 | 20352782.32 |
| Cytochalasin H | 2231626.034 | 2051070.864 | 1854929.896 | 2016347.82 | 2211836.526 | 1998802.374 | 1888650.338 | 2010185.38 | 2126303.527 | 2000651.226 | 2009688.885 | 1763602.983 |
| LPG 19:0 | 6110101.594 | 8654591.06 | 130547.9687 | 122454.3542 | 17831182.08 | 20843277.03 | 19481015.18 | 17216472.4 | 15974059.8 | 10079956.32 | 17279432.37 | 8657673.332 |
| 2-Aminopyridine | 2013121.804 | 263757.3904 | 1978192.26 | 1959451.706 | 19964704.96 | 22100674.38 | 23952906.08 | 30559267.38 | 24686054.13 | 14348812.56 | 16573198.23 | 20605688.54 |
| LNAPS 24:1/N-8:0 | 726842.9945 | 504021.2004 | 862265.7318 | 907320.9826 | 1535780.624 | 636284.7743 | 1096558.257 | 887371.6376 | 1490983.203 | 1326020.208 | 1717782.707 | 654876.0109 |
| 5-(4-Hydroxy-3-methoxybenzylidene)-1,3-dimethyl-2,4,6(1H,3H,5H)-pyrimid | 3085986.555 | 3045679.993 | 4767683.066 | 3249044.38 | 8764525.263 | 7136440.088 | 9399334.002 | 7879783.777 | 7787481.286 | 6064647.806 | 6161135.215 | 6203910.149 |

| Metabolites | PBS_1 | PBS_2 | PBS_3 | PBS_4 | BCG_1 | BCG_2 | BCG_3 | BCG_4 | BCGΔ *pe18*_1 | BCGΔ *pe18*_2 | BCGΔ *pe18*_3 | BCGΔ*pe18*_4 |
|---|---|---|---|---|---|---|---|---|---|---|---|---|
| inetrione | | | | | | | | | | | | |
| Lyso-sphingomyelin | 951402.8931 | 787457.7415 | 661144.9675 | 821033.0423 | 1132638.848 | 855795.5423 | 1104324.893 | 1233903.14 | 1174500.145 | 1094577.814 | 1170466.336 | 845030.8104 |
| MGDG O-13:1_20:4 | 2191993.579 | 457630.8305 | 895208.4811 | 788233.5591 | 662397.5008 | 545938.6731 | 694943.6495 | 1787470.224 | 794040.8514 | 1049100.775 | 673932.8366 | 758852.8103 |
| PG 17:1_18:2 | 171715.1774 | 79772.50068 | 129739.8267 | 155502.5547 | 3006485.493 | 2035440.233 | 3003392.181 | 1897270.226 | 3581197.201 | 4157335.239 | 2487247.854 | 1451751.841 |
| L-Fucitol | 1080660.516 | 1119592.031 | 1357208.257 | 1293800.874 | 2128916.934 | 1930242.667 | 2079613.098 | 2106521.534 | 1872580.363 | 1703211.128 | 1790811.736 | 1597229.669 |
| Fenazaflor | 1478830.306 | 1377298.188 | 2031232.566 | 1575057.381 | 2756150.005 | 2125372.876 | 2765894.279 | 2804748.573 | 2422582.222 | 2878043.501 | 2210916.632 | 2383293.241 |
| Chlozolinate | 912567.6226 | 704320.3571 | 1037577.091 | 766270.396 | 1501573.666 | 1145873.147 | 1580764.751 | 1289258.114 | 1031615.02 | 1288366.73 | 1402833.878 | 797078.6173 |
| PC(20:3(5Z,8Z,11Z)/P-18:0) | 184347243.4 | 179624563.2 | 91868873.21 | 125233512.9 | 38751893.61 | 24186559.83 | 38716951.59 | 37559301.77 | 48763331.91 | 57258801.76 | 43598433.71 | 29733920.16 |
| Tyrosine | 10761983.29 | 9096234.18 | 14444038.27 | 11318398.37 | 22093684.58 | 15963815.57 | 19915821.77 | 18184252.52 | 19243070.94 | 12997049.09 | 14623883.45 | 17887845.21 |
| 1-Octadecanoyl-2-(15S-hydroxy-5Z,8Z,11Z,13E-eicosatetraenoyl)-sn-glycero- | 1549335.432 | 1733634.852 | 2738687.923 | 2195917.384 | 12043233.43 | 3150816.198 | 3660327.393 | 8051781.061 | 3213651.035 | 3284211.955 | 9903484.661 | 2356256.793 |

| Metabolites | PBS_1 | PBS_2 | PBS_3 | PBS_4 | BCG_1 | BCG_2 | BCG_3 | BCG_4 | BCGΔ*pe18*_1 | BCGΔ*pe18*_2 | BCGΔ*pe18*_3 | BCGΔ*pe18*_4 |
|---|---|---|---|---|---|---|---|---|---|---|---|---|
| 3-phosphoethanolamine | | | | | | | | | | | | |
| N-Cyclohexyl-N'-(tetrahydro-2-furanylmethyl)urea | 741329.7864 | 970235.2681 | 1354408.61 | 2039731.418 | 534560.275 | 1187519.807 | 899359.0124 | 661191.5064 | 682974.2499 | 632723.7287 | 636868.0941 | 981454.9486 |
| SL 10:0;O/18:4 | 2141385.082 | 2414313.723 | 360585.575 | 1487487.308 | 340028.1068 | 231214.3681 | 350750.3789 | 431557.5208 | 492385.3751 | 547454.7287 | 303006.0037 | 335215.7924 |
| SM 36:4;3O | 5819456.169 | 6471250.035 | 4925014.907 | 4524299.11 | 1896762.074 | 1949979.231 | 2532506.901 | 2481492.976 | 2547350.393 | 3049666.812 | 2482362.544 | 2025136.286 |
| DGGA 12:0_24:2 | 10285463.7 | 9308061.078 | 3832408.545 | 7881881.171 | 2345750.763 | 1917949.184 | 1608539.315 | 2716291.133 | 2866154.81 | 1909067.528 | 1510651.093 | 2246458.55 |
| PE 8:0_30:5 | 6856733.786 | 6524059.693 | 5889357.693 | 4880035.769 | 13414391.89 | 13353891.11 | 14486934.85 | 14186683.77 | 13678003.36 | 16317236.79 | 16696872.36 | 14171788.41 |
| 5-(Allylamino)-1,3,4-thiadiazole-2(3H)-thione | 12229608.34 | 9471384.137 | 16354690.48 | 13195159.66 | 12364821.85 | 14716716.59 | 13461209.84 | 13166318.11 | 12836910.1 | 9245657.218 | 10975897.35 | 12313442.68 |
| 2,4,6-Tri-tert-butylaniline | 6377256.615 | 7364618.486 | 5387292.953 | 5464686.278 | 6637022.676 | 6510055.591 | 5869687.382 | 6637933.365 | 6455294.627 | 6257569.027 | 6170110.926 | 7091573.438 |
| Ouabain | 5284200.937 | 4996508.777 | 5969840.857 | 5266659.614 | 6239953.256 | 5174555.508 | 6056812.013 | 5576949.937 | 5566608.557 | 5817635.77 | 5720670.333 | 5572596.675 |
| Heneicosapentaenoic acid | 1073507.55 | 856015.9515 | 772322.6211 | 852997.7967 | 1140147.833 | 871867.9276 | 1309152.28 | 765428.6297 | 890169.5292 | 1106037.193 | 898637.864 | 833277.0632 |

| Metabolites | PBS_1 | PBS_2 | PBS_3 | PBS_4 | BCG_1 | BCG_2 | BCG_3 | BCG_4 | BCGΔ*pe18*_1 | BCGΔ*pe18*_2 | BCGΔ*pe18*_3 | BCGΔ*pe18*_4 |
|---|---|---|---|---|---|---|---|---|---|---|---|---|
| PE(P-18:0/18:1(9Z)) | 7034042.505 | 9209889.235 | 13570703.81 | 10895154.94 | 20866280.35 | 11798611.05 | 18362822.39 | 12602986.05 | 14527258.42 | 17051129.8 | 18144934.12 | 7568065.631 |
| PC O-18:5_18:1 | 621118.8802 | 575842.7504 | 1141324.435 | 569224.0957 | 663373.0446 | 76249.11638 | 632064.6054 | 349401.8685 | 707736.9964 | 328957.3701 | 799406.171 | 182576.5645 |
| Timosaponin A1 | 118214.2773 | 175489.1377 | 193813.2467 | 289805.645 | 236822.7128 | 248108.7618 | 170966.1219 | 163383.4301 | 149576.2607 | 190005.2274 | 149705.7127 | 242266.4854 |
| N.alpha.-Benzoyl-DL-arginine-4-nitroanilide | 8685159.824 | 6691157.85 | 7345914.639 | 7000932.024 | 7045257.38 | 7692312.125 | 6769125.153 | 7795121.599 | 6909467.318 | 7874491.623 | 7488803.142 | 6170141.555 |
| Undecanedioic acid | 1515252.983 | 1698284.723 | 2095890.929 | 1992064.705 | 1627474.81 | 1577869.82 | 3962885.431 | 1398007.246 | 1617154.428 | 1525186.849 | 1650133.03 | 2768929.084 |
| Undecanoic acid | 12399302.49 | 11900935.92 | 10602411.46 | 10046075.58 | 28626352.67 | 11153775.35 | 11070563.19 | 9796416.575 | 11839570.81 | 11576177.65 | 11499111.5 | 11107264.78 |
| .delta.-Decalactone | 5306289.67 | 3089979.723 | 3785076.357 | 1736039.702 | 5408324.257 | 4241079.119 | 3850324.326 | 7738269.871 | 2117695.903 | 6309898.293 | 4746229.182 | 3464487.618 |
| 5-Chloro-3-iodo-2-pyridone | 4773730.24 | 2079883.913 | 5888286.215 | 3479387.128 | 53034811.03 | 53096178.22 | 13124866.24 | 52558188.12 | 43808110 | 21708239.94 | 29438939.97 | 22015244.12 |
| Dimethirimol | 62624.34921 | 461897.9082 | 1254416.288 | 1401922.625 | 138912.5159 | 30429.31285 | 135362.5884 | 35523.98955 | 121656.7614 | 115446.5677 | 28952.03439 | 1299662.55 |
| HexCer 17:0;2O/17:1 | 5034670.471 | 2833721.296 | 8684026.088 | 3794000.469 | 4596720.358 | 6605008.484 | 6133971.262 | 5888187.294 | 4218494.647 | 4212595.91 | 6037875.869 | 3559101.643 |
| 3-(2-Aminoethylthio)-2-methylpropionic acid | 2406054.444 | 1204502.652 | 2784548.009 | 1447611.717 | 5931705.919 | 4546548.1 | 6589183.961 | 7125245.439 | 3931965.853 | 3841154.023 | 5436422.015 | 3842091.362 |

| Metabolites | PBS_1 | PBS_2 | PBS_3 | PBS_4 | BCG_1 | BCG_2 | BCG_3 | BCG_4 | BCGΔ*pe18*_1 | BCGΔ*pe18*_2 | BCGΔ*pe18*_3 | BCGΔ*pe18*_4 |
|---|---|---|---|---|---|---|---|---|---|---|---|---|
| Valeric acid | 2667419.678 | 2603687.625 | 3227771.812 | 3407182.379 | 3543685.929 | 3676122.834 | 2774402.756 | 2749659.765 | 3105805.824 | 2082018.866 | 2299049.942 | 3091169.054 |
| Milameline | 4983721.484 | 4396762.279 | 7954825.899 | 4569778.678 | 5350281.664 | 5742042.408 | 5113016.666 | 4677825.502 | 4052425.195 | 2558855.242 | 5769103.998 | 5520207.907 |
| PE-Cer 12:0;2O/20:5 | 428176.205 | 447338.5227 | 816885.0374 | 525605.2578 | 1747324.502 | 2070480.496 | 2053554.996 | 1378944.005 | 1616834.525 | 1481267.31 | 1927169.296 | 723489.879 |
| PC 9:0_36:6 | 1514517.274 | 1275429.647 | 1062867.322 | 1565641.807 | 1037692.626 | 1213774.593 | 1014849.901 | 1079389.371 | 1315560.704 | 1027794.98 | 1040911.292 | 1320638.225 |
| Pyrrolidine | 14980685.86 | 12208566.33 | 19325993.88 | 14995234.64 | 32547840.1 | 26855499.86 | 35956189.16 | 31010489.85 | 27493320.18 | 24913501.3 | 26005917.25 | 23486246.04 |
| 5-[5-(Acetyloxymethyl)-1,2,4a-trimethyl-7-oxo-3,4,8,8a-tetrahydro-2H-naphthalen-1-yl]-3-methylpentanoic acid | 3110067.568 | 1757814.426 | 2461371.056 | 1888965.138 | 65470191.46 | 96960383.98 | 60284675.75 | 73007502.5 | 49083716.3 | 26237981.97 | 65608525.73 | 73483519.43 |
| Dodecylbenzenesulfonic acid | 2408996.302 | 2878298.123 | 2703198.923 | 2364529.412 | 2385665.856 | 2461257.533 | 2600072.535 | 2713319.038 | 2314956.333 | 5061768.661 | 2315326.68 | 2527282.226 |
| L-Erythrulose | 8064638.164 | 7667452.216 | 10742180.56 | 7069458.53 | 9945849.837 | 7998574.863 | 11294842.57 | 8857007.386 | 10863838.74 | 5971627.584 | 3864766.92 | 7053605.084 |
| (.+/-.)-11-Hydroxy-.DELTA.9-tetrahydrocannabinol | 3226350.787 | 3258887.139 | 3359343.547 | 2997827.226 | 3024565.294 | 3171840.725 | 4347911.614 | 4269099.285 | 3315449.814 | 3278806.629 | 3055262.528 | 3223087.17 |
| Taurolithocholic | 472027 | 491100 | 837748 | 561558 | 761902 | 790278 | 101895 | 782527 | 658855 | 610836 | 605798 | 578616 |

| Metabolites | PBS_1 | PBS_2 | PBS_3 | PBS_4 | BCG_1 | BCG_2 | BCG_3 | BCG_4 | BCGΔ*pe18*_1 | BCGΔ*pe18*_2 | BCGΔ*pe18*_3 | BCGΔ*pe18*_4 |
|---|---|---|---|---|---|---|---|---|---|---|---|---|
| acid | 2.015 | 2.486 | 7.032 | 0.07 | 6.243 | 3.349 | 96.87 | 4.214 | 9.983 | 3.246 | 2.031 | 9.447 |
| DGCC 8:0_28:7 | 2863979.923 | 2207517.515 | 1804375.829 | 1800593.98 | 1727549.402 | 1421410.511 | 1949201.043 | 1413991.074 | 1925391.049 | 2125900.417 | 2030686.495 | 1370631.691 |
| DL-Isocitric acid lactone | 2053855.864 | 1881208.244 | 2020671.518 | 1724188.838 | 6952770.372 | 4171372.277 | 8628934.566 | 6377002.936 | 4221668.263 | 6104134.221 | 5196458.754 | 3908149.004 |
| PC(18:2(9Z,12Z)/P-16:0) | 9545457.266 | 14914202.48 | 9647453.217 | 7067491.365 | 11247243.39 | 9152373.691 | 8790227.802 | 3883083.311 | 5237835.115 | 7522831.294 | 3934734.729 | 7623620.34 |
| 3-(Morpholinomethyl)-1H-indole-6-carboxylic acid | 2006875.808 | 2035111.557 | 4065870.061 | 2431673.294 | 2907406.53 | 3591034.852 | 3833990.12 | 2829363.164 | 2946489.429 | 1752209.347 | 2196076.85 | 2696216.713 |
| PC(20:2(11Z,14Z)/18:1(9Z)) | 78164438.11 | 98325119.11 | 44460577.16 | 72981401.77 | 94180471.12 | 64959353.98 | 109088672.7 | 79412975.75 | 95463459.78 | 119050220.8 | 93408151.65 | 49077348.96 |
| 3-Amino-4-hydroxy-N-methylbenzenesulfonamide | 3135109.211 | 3009274.943 | 2891115.286 | 2924050.442 | 2893280.934 | 3037916.201 | 3041341.869 | 3034206.181 | 3057006.929 | 2922482.158 | 2978697.09 | 2869846.331 |
| Isoginkgetin | 624167.9031 | 637264.1189 | 605603.6931 | 646646.5086 | 726758.5616 | 790997.1476 | 690894.9256 | 654568.869 | 708271.6675 | 725462.3843 | 712486.5073 | 723296.5188 |
| Tropinone | 1020635.419 | 927531.7788 | 1025309.59 | 1025928.601 | 1060824.249 | 382345.8861 | 969973.8598 | 356837.909 | 960174.5556 | 1002111.003 | 949816.1948 | 1105615.657 |
| Carbadox | 3614554.734 | 2432543.3 | 4045463.157 | 2571942.676 | 15545778.42 | 7350299.798 | 13055295.16 | 11158381.33 | 9583897.656 | 10526504.33 | 9521509.575 | 5733412.973 |
| 1H-Pyrazolo[3,4-d]pyrimidine-3,4-diamine, | 2081303.154 | 1427566.2 | 2865255.955 | 2070101.771 | 1985496.015 | 2458431.686 | 2369862.777 | 2385011.433 | 2223085.208 | 1011996.575 | 1162979.058 | 2069884.585 |

| **Metabolites** | **PBS_1** | **PBS_2** | **PBS_3** | **PBS_4** | **BCG_1** | **BCG_2** | **BCG_3** | **BCG_4** | **BCGΔ*pe18*_1** | **BCGΔ*pe18*_2** | **BCGΔ*pe18*_3** | **BCGΔ*pe18*_4** |
|---|---|---|---|---|---|---|---|---|---|---|---|---|
| N3-(4-fluorophenyl)- | | | | | | | | | | | | |
| Prolylhydroxyproline | 1418104.843 | 460955.612 | 966009.8527 | 563834.0327 | 1281421.574 | 1071418.217 | 1397257.492 | 1124382.294 | 936083.7373 | 799845.1275 | 904274.5143 | 873961.7393 |
| (Diethylamino)(oxo)acetic acid | 540151.441 | 742034.2472 | 521961.1975 | 640059.4704 | 865798.1153 | 593374.1726 | 825607.839 | 748574.976 | 676790.0427 | 671950.6887 | 708398.8345 | 564060.3458 |
| LysoPC(18:1(11Z)) | 7167752.368 | 1423830.582 | 1847133.364 | 1409440.858 | 2697407.083 | 3342410.087 | 15546984.02 | 12623585.94 | 12964525 | 13880033.63 | 3673733.968 | 2633615.676 |
| Ethyl 4-(4-methylpiperazin-1-yl)butanoate | 12300610.37 | 11210966.52 | 11287078.35 | 11493731.53 | 17629461.22 | 22271407.72 | 16215894.8 | 19456723.91 | 16390918.87 | 12563829.28 | 18788940.62 | 11895160.37 |
| Arachidonyl trifluoromethyl ketone | 22432827.24 | 25559664.42 | 23286675.09 | 22304908.83 | 27635089.54 | 26897305.24 | 22691107.71 | 20785215.64 | 23284265.7 | 24875102.47 | 24519895.25 | 22357887.99 |
| LPC(22:6) | 592217.9154 | 1874345.167 | 2587704.882 | 1880243.12 | 3936590.453 | 3953546.308 | 3504092.351 | 3477003.376 | 3906347.355 | 19341.69149 | 3255367.588 | 3621004.709 |
| Dodemorph | 258001911.5 | 287444796.7 | 150965110.9 | 213953293.2 | 57346201.27 | 51048869.21 | 62655572.6 | 66682125.32 | 47562750.59 | 44312592.55 | 42847115.65 | 24929064.25 |
| 2-Formyl-3-thienyl boronic acid | 3231856.31 | 3112784.838 | 1538868.194 | 1737943.163 | 3007545.112 | 1516027.725 | 1939694.477 | 1493538.688 | 2961187.001 | 3148338.398 | 1666653.619 | 2837185.906 |
| Pirimiphos-methyl-N-desethyl | 857331.1835 | 1130921.45 | 2254809.92 | 1270803.114 | 16123317.44 | 8557711.126 | 14487227.71 | 12501414.62 | 9764493.363 | 8408828.008 | 10609026.5 | 6198773.543 |

| Metabolites | PBS_1 | PBS_2 | PBS_3 | PBS_4 | BCG_1 | BCG_2 | BCG_3 | BCG_4 | BCGΔ*pe18*_1 | BCGΔ*pe18*_2 | BCGΔ*pe18*_3 | BCGΔ*pe18*_4 |
|---|---|---|---|---|---|---|---|---|---|---|---|---|
| Tetracosa-6,9,12,15,18,21-hexaenoylcarnitine | 3296101.971 | 3111455.901 | 2701610.023 | 2871895.378 | 4197766.877 | 4820554.425 | 5852139.625 | 5785855.905 | 4301746.599 | 3987628.328 | 3711098.696 | 2570039.172 |
| N,N-Diethylcathinone | 1716061.301 | 2304233.859 | 1450472.54 | 1810995.87 | 1836580.57 | 1366597.038 | 2061194.714 | 2181429.208 | 1994601.52 | 1965759.114 | 1674454.805 | 1645205.708 |
| NAGly 18:4/22:5 | 703345.9054 | 719760.8849 | 713394.4322 | 708919.3427 | 726851.4661 | 709314.8902 | 727132.8667 | 719123.3399 | 714262.7516 | 725410.0432 | 739548.9095 | 684966.8181 |
| NAE 18:4 | 306332.7043 | 1784712.19 | 1146451.607 | 1301317.897 | 1270810.009 | 1137012.701 | 1404333.255 | 887516.2138 | 1319833.442 | 714219.6785 | 1163973.237 | 945520.2994 |
| Altamisic acid | 5949767.519 | 8491588.147 | 12450305.16 | 11084869.01 | 6677158.189 | 5529386.169 | 6224875.021 | 5992120.828 | 5753106.509 | 5949044.342 | 6162015.595 | 17712533.67 |
| 3-Hydroxyheptadecanoylcarnitine | 1797728.318 | 2082560.947 | 1289339.504 | 2239618.628 | 1979760.856 | 2367397.179 | 1941226.888 | 1557269.351 | 1014082.543 | 1973374.982 | 1344898.911 | 1704469.899 |
| 5-(4-Aminophenyl)-4-phenyl-2,4-dihydro-3H-1,2,4-triazole-3-thione | 0 | 0 | 0 | 0 | 28154.94805 | 25810.09566 | 22841.58284 | 28672.38132 | 124908.845 | 95621.50271 | 90898.71907 | 87229.59625 |
| cis-5-Dodecenoic acid | 2562113.905 | 1963888.387 | 1775474.526 | 1615751.317 | 2103105.582 | 2232611.295 | 2115353.629 | 1893889.477 | 2050125.142 | 3005601.539 | 2255289.375 | 2073971.726 |
| Nonaethylene glycol | 3671438.628 | 3573701.426 | 4241688.491 | 3566388.932 | 3742516.788 | 3722787.862 | 3500054.107 | 3567512.672 | 3611048.063 | 3676349.665 | 3684610.703 | 3601809.996 |
| 2-Acetyl-3-ethylpyrazine | 306625.0334 | 3784384.392 | 3681022.558 | 4145269.633 | 764894.9899 | 192813.0274 | 762286.0019 | 860145.5379 | 1114945.67 | 2068885.954 | 223477.5237 | 6423286.711 |

| **Metabolites** | **PBS_1** | **PBS_2** | **PBS_3** | **PBS_4** | **BCG_1** | **BCG_2** | **BCG_3** | **BCG_4** | **BCGΔ*pe18*_1** | **BCGΔ*pe18*_2** | **BCGΔ*pe18*_3** | **BCGΔ*pe18*_4** |
|---|---|---|---|---|---|---|---|---|---|---|---|---|
| Dodecyltriphenylphosphonium cation | 102521.709 | 59670.96702 | 69862.83556 | 68240.86713 | 91450.63134 | 74066.56098 | 58175.97242 | 77575.34183 | 84514.2566 | 81054.88484 | 75452.99533 | 61347.61528 |
| LNAPE 22:6/N-18:0 | 2299285.717 | 1623497.048 | 3964879.564 | 2564178.309 | 4701845.388 | 4945234.667 | 4247353.611 | 4743010.627 | 2957882.225 | 3430723.779 | 4174886.947 | 2219204.681 |
| SL 15:3;O/20:2;O | 71500.92026 | 198246.0624 | 26560.50784 | 62312.79425 | 2208377.506 | 1423836.284 | 1986934.554 | 1260196.501 | 1042081.94 | 1460842.493 | 1891712.205 | 940487.571 |
| 3-Sulfamoylbenzoic acid | 6092284.41 | 8892104.909 | 10360066.53 | 9659026.942 | 6176214.267 | 5906115.695 | 6618309.169 | 4729352.553 | 6450794.452 | 6445498.154 | 6505131.87 | 13661529.46 |
| SPB 22:1;2O | 10459473.4 | 15280913.33 | 5769414.878 | 11476622.49 | 6492428.256 | 8593293.543 | 7160247.712 | 10703933.46 | 9515469.471 | 8420621.972 | 7076862.421 | 6051380.192 |
| N-Oleoyl-D-erythro-sphingosylphosphorylcholine | 1524303.029 | 1312862.068 | 1123111.195 | 1217182.154 | 912708.2889 | 679550.7542 | 945224.3962 | 1322286.839 | 989527.8623 | 1053077.929 | 1099724.986 | 1281159.358 |
| 13,15-Dihydroxy-9-methyl-10-oxabicyclo[10.4.0]hexadeca-1(12),13,15-trien-11-one | 1482777.494 | 3194391.911 | 5460725.366 | 3205264.759 | 1190720.78 | 1021454.497 | 2377265.022 | 1018991.967 | 1008372.174 | 1493681.032 | 1519103.882 | 8790831.872 |
| 2-Amino-5,5,5-trifluoropentanoic acid | 725444.777 | 506610.2254 | 1373581.51 | 610527.5095 | 4437150.424 | 2514090.599 | 4948182.595 | 3105941.971 | 2320815.616 | 2827657.399 | 3716019.997 | 1911439.256 |
| L-tyrosine-methyl-ester | 1764063.31 | 1538377.633 | 923881.6637 | 2103649.856 | 9798588.27 | 10381806.92 | 5479018.938 | 8771749.803 | 4469502.067 | 6047710.71 | 9538375.162 | 7514365.6 |
| Allocryptopine | 2760094.009 | 2822206.257 | 2997282.661 | 2736112.098 | 2652083.359 | 2505593.298 | 2684211.017 | 2824567.108 | 2883717.639 | 2511665.326 | 2702501.478 | 2566250.394 |

| Metabolites | PBS_1 | PBS_2 | PBS_3 | PBS_4 | BCG_1 | BCG_2 | BCG_3 | BCG_4 | BCGΔ*pe18*_1 | BCGΔ*pe18*_2 | BCGΔ*pe18*_3 | BCGΔ*pe18*_4 |
|---|---|---|---|---|---|---|---|---|---|---|---|---|
| Animicin A | 7766.209651 | 0 | 7019.184385 | 14740.77133 | 455464.8924 | 390181.8615 | 416320.0932 | 375731.1249 | 334327.7672 | 238462.111 | 268354.3431 | 305318.6456 |
| PC(P-32:0) | 27005499.99 | 41558036.44 | 15617409.49 | 33585118.71 | 34635641.69 | 40077598.1 | 36584524.55 | 32461175.44 | 35549856.78 | 31064745.94 | 31669033.35 | 30588207.44 |
| N-Linoleoyl Methionine | 7679318.945 | 6566954.558 | 9556379.727 | 10011250.05 | 15970892.07 | 15774544.41 | 14797744.46 | 12306088.45 | 9583740.957 | 9052315.535 | 15696668.9 | 10521886.01 |
| Pro-Pro | 103961.7909 | 97198.74117 | 415758.8934 | 511018.8743 | 531289.1417 | 406252.9752 | 364546.5352 | 376479.9704 | 469353.3784 | 484514.2263 | 595293.5727 | 471282.047 |
| N-Phenylbenzo[cd]indol-2-amine | 145836.4369 | 626543.9184 | 874338.8722 | 1035398.984 | 94513.27773 | 85381.24719 | 121925.5425 | 172124.3941 | 125588.613 | 123546.2886 | 58256.83319 | 1587671.895 |
| N-cis-Hexadec-9-enoyl-L-homoserine lactone | 8923395.2 | 6278278.724 | 6399022.334 | 6989875.872 | 7295447.574 | 7276254.479 | 7788677.946 | 6382417.503 | 7450816.612 | 7960656.88 | 8278081.725 | 8721958.218 |
| PI O-10:0_22:4 | 303127.8201 | 313774.5143 | 47657.69817 | 330684.2939 | 12496860 | 11217586.97 | 12125732.31 | 10041297.43 | 10369541.1 | 4966273.066 | 8690109.337 | 7338963.573 |
| 2-Bromo-5-hydroxybenzaldehyde | 10602364.08 | 3540382.779 | 10628592.41 | 10595387.11 | 10676498.65 | 1736708.899 | 2799663.145 | 6100862.488 | 10040603.54 | 8316928.372 | 1395671.258 | 2257196.684 |
| MGDG 3:0_17:2 | 234100.5297 | 202742.3024 | 187930.371 | 207188.5129 | 400086.3209 | 116238.087 | 507019.3894 | 368798.9921 | 269078.7867 | 449583.3522 | 222954.7411 | 165478.5438 |
| Palmitoylserinol | 4274585.166 | 4493729.069 | 4510978.628 | 6623442.445 | 3954071.731 | 4619731.073 | 5896694.129 | 4517033.888 | 3987182.161 | 4594265.723 | 5003814.591 | 3913272.2 |
| N-Tetracosenoyl-4-sphingenine | 674790.0508 | 9523263.662 | 18618014.76 | 10600912.04 | 7363658.89 | 13332660.36 | 7891093.432 | 5953204.153 | 8068971.319 | 6549339.536 | 9643795.489 | 15049346.73 |
| LysoPC(18:3(6Z,9 | 907437. | 929644. | 914031. | 105111 | 138501 | 112640 | 187746 | 139349 | 135752 | 144042 | 159757 | 134617 |

| Metabolites | PBS_1 | PBS_2 | PBS_3 | PBS_4 | BCG_1 | BCG_2 | BCG_3 | BCG_4 | BCGΔ *pe18*_1 | BCGΔ *pe18*_2 | BCGΔ *pe18*_3 | BCGΔ*pe18*_4 |
|---|---|---|---|---|---|---|---|---|---|---|---|---|
| Z,12Z)) | 1559 | 063 | 6455 | 4.197 | 7.021 | 1.308 | 6.902 | 9.005 | 7.847 | 0.731 | 7.017 | 3.548 |
| 5,5,7,12,12,14-Hexamethyl-1,4,8,11-tetraazacyclotetradecane | 10086761.42 | 11384270.04 | 6875814.57 | 8718641.366 | 3997782.011 | 6028415.153 | 4335805.426 | 3864997.94 | 3491235.529 | 5632851.165 | 4253743.047 | 3318129.928 |
| 5-oxo-prolyl-glutamine | 61352.61853 | 40062.11169 | 121753.835 | 98939.75634 | 1307810.07 | 940296.8743 | 1079739.928 | 785484.0167 | 792724.3419 | 899857.7761 | 840640.2305 | 645222.7073 |
| DGTS 2:0_17:2 | 10904094.42 | 10972520.14 | 8299275.312 | 10872383.31 | 10630227.54 | 17522383.2 | 13245228.36 | 11760828.49 | 10767977.71 | 8567076.837 | 9996653.438 | 7132379.106 |
| 5-Bromo-1H-indazole | 14145973.61 | 14493421.23 | 18953888.63 | 13029123.18 | 16192425.92 | 17135155.13 | 12689138.7 | 14038501.24 | 13059009.83 | 13334355.19 | 17423423.04 | 12093211.81 |
| 2,5-Dimethyl-3,4-diphenyl-2,4-cyclopentadien-1-one | 372985.3885 | 330585.4388 | 198331.1668 | 405980.6457 | 410019.9608 | 356951.7129 | 271363.7801 | 313548.0685 | 408461.9874 | 271462.3596 | 248879.2009 | 453126.1989 |
| 1-Stearoyl-2-arachidonyl-sn-glycero-3-phosphocholine | 4224857.032 | 7068558.507 | 5728875.819 | 5419123.825 | 3551982.849 | 3773997.001 | 4545882.869 | 6060577.285 | 3531791.265 | 9516077.267 | 6512146.926 | 16871827.82 |
| Iraldeine | 16531429.28 | 18449384.85 | 18173399.5 | 18805160.48 | 17454306.93 | 15865431.34 | 17954407.89 | 17360157.97 | 15240884.91 | 16991434.5 | 14478604.84 | 16740546.98 |
| (1S,2R)-2-(Diethylamino)-1-phenyl-1-propanol | 30870131.88 | 42678006.44 | 39349146.32 | 39336629.68 | 29333252.29 | 38021684.86 | 40317632.06 | 39334553.06 | 36799576.79 | 40433651.55 | 38755207.88 | 35471920.06 |
| Diethyl 3-hydroxyphenylph | 1789517.862 | 1979916.088 | 1639451.956 | 1879561.23 | 1880230.206 | 1905458.912 | 1881565.071 | 1805301.759 | 1915408.673 | 1931104.445 | 1836422.462 | 1719065.942 |

| Metabolites | PBS_1 | PBS_2 | PBS_3 | PBS_4 | BCG_1 | BCG_2 | BCG_3 | BCG_4 | BCGΔ *pe18*_1 | BCGΔ *pe18*_2 | BCGΔ *pe18*_3 | BCGΔ*pe18*_4 |
|---|---|---|---|---|---|---|---|---|---|---|---|---|
| osphonate | | | | | | | | | | | | |
| (E)-5-((1S,4R)-3,3-Dimethylbicyclo[2.2.1]heptan-2-yl)pent-3-en-2-one | 5391822.712 | 6571479.495 | 4410416.776 | 4465330.581 | 6036640.132 | 6909171.236 | 5445912.638 | 5637147.718 | 5409327.539 | 5952095.603 | 6117748.644 | 5697990.062 |
| DG 18:5_18:5 | 760512.3958 | 818445.5413 | 1116978.064 | 687156.4327 | 757619.0388 | 858847.5298 | 853979.7164 | 833429.1818 | 755832.7164 | 791526.6675 | 895655.4901 | 480755.446 |
| PC 9:0_32:6 | 804685.2134 | 1356182.432 | 625924.5 | 1252994.118 | 810770.5008 | 913015.7806 | 677188.2893 | 868395.3806 | 798133.2379 | 1167463.544 | 887405.5313 | 855998.1468 |
| Thymine | 1399862.482 | 2617109.382 | 4773891.287 | 2145323.73 | 7396519.29 | 5934842.081 | 7042811.785 | 5707309.318 | 5469432.686 | 3938271.429 | 5764631.627 | 4880573.39 |
| DG 38:9 | 1055462.332 | 1167445.696 | 883495.6087 | 818120.1268 | 1111661.325 | 1105867.089 | 578930.1314 | 689566.1901 | 1119688.905 | 1002805.581 | 1096315.242 | 904833.0205 |
| Butyrylcarnitine (Car(4:0)) | 1498770.229 | 1415680.123 | 2266337.753 | 1842132.578 | 6270345.198 | 4135892.423 | 5959990.308 | 4654622.898 | 4491243.977 | 4185253.664 | 5537269.119 | 4838675.842 |
| BMP 20:4_20:4 | 3994181.362 | 5113455.055 | 3021977.242 | 4237965.535 | 1543872.762 | 2320329.774 | 2359563.469 | 2279128.68 | 3018714.909 | 2040573.641 | 2036792.183 | 2596274.886 |
| Ethyl 6-methyl-4-(4-methylphenyl)-2-oxo-1,2,3,4-tetrahydro-5-pyrimidinecarboxylate | 1344485.505 | 1679340.124 | 1612481.559 | 1458452.098 | 1970202.511 | 1943617.592 | 1325097.459 | 1633882.19 | 1663262.848 | 1210179.609 | 1241296.863 | 2353163.258 |
| LysoPC(14:1(9Z)) | 192595. | 76588.6 | 105795. | 169263. | 200617. | 169693. | 189996. | 240744. | 175743. | 5067.72 | 250939. | 135015. |

| **Metabolites** | **PBS_1** | **PBS_2** | **PBS_3** | **PBS_4** | **BCG_1** | **BCG_2** | **BCG_3** | **BCG_4** | **BCGΔ*pe18*_1** | **BCGΔ*pe18*_2** | **BCGΔ*pe18*_3** | **BCGΔ*pe18*_4** |
|---|---|---|---|---|---|---|---|---|---|---|---|---|
| | 5793 | 1801 | 7576 | 2329 | 2065 | 6199 | 4003 | 4196 | 3077 | 066 | 3637 | 1833 |
| PC 9:0_26:4 | 12478216.02 | 13756443.47 | 7253850.649 | 8874570.412 | 9143671.894 | 10841801.86 | 12842242.13 | 14887776.07 | 11643579.47 | 14190625.48 | 11619544.1 | 8544980.043 |
| 4-(Dimethylamino)-3-methylphenol | 1266748.248 | 1228410.534 | 1164557.881 | 1243216.114 | 1422224.368 | 1283138.644 | 1201036.879 | 1431643.943 | 1112698.096 | 1089024.844 | 1194011.702 | 744256.0805 |
| Stearidonic acid | 1245461.052 | 1386247.086 | 1110402.652 | 1394738.765 | 1685105.571 | 1421104.68 | 1551584.937 | 1144143.344 | 879155.0364 | 1071754.276 | 1563980.45 | 2372145.909 |
| Antibiotic FR 901512 | 22653853.06 | 9139831.861 | 8370170.342 | 8403696.61 | 8637903.53 | 10094135.45 | 9206412.588 | 9772164.163 | 9146470.435 | 9284167.853 | 9108548.193 | 8249226.6 |
| 2-Ethylbutanedioic acid | 2832383.991 | 3797607.025 | 4381643.897 | 6040507.281 | 7429366.196 | 7497073.555 | 3156327.92 | 2069161.336 | 5221917.522 | 4402644.719 | 3586794.318 | 6864003.991 |
| 1-Stearoyl-2-oleoyl-sn-glycero-3-phospho-(1'-sn-glycerol) | 4009589.916 | 5126776.143 | 3830176.951 | 4309734.657 | 93676633.74 | 60864587.24 | 32765092.24 | 19356702.99 | 21372437.79 | 32182256.84 | 86899508.55 | 11847836.95 |
| alpha-C-Mannosyltryptophan | 5760537.347 | 4976799.268 | 7302217.654 | 5235540.942 | 6574041.084 | 4135171.153 | 7665762.227 | 5424051.166 | 4587276.615 | 5408776.096 | 5730556.974 | 4290190.856 |
| PC(37:5) | 3032873.571 | 975960753.4 | 18186651.74 | 7236164.269 | 63660508.33 | 47425092.59 | 63372068.33 | 65562264.16 | 57706674.11 | 79080643.91 | 54238722.25 | 41859532.11 |
| PC(22:2) | 900529.8647 | 845522.6213 | 1941328.703 | 847322.2988 | 525644.7309 | 565526.3707 | 333872.6851 | 691042.4778 | 598848.5285 | 0 | 738331.8169 | 494007.4605 |
| 2-Phenylpropionic acid | 2237953.471 | 2071438.942 | 2308548.613 | 1880807.729 | 2538795.677 | 2579975.376 | 2396801.463 | 2750267.283 | 2410171.615 | 2081644.404 | 2695592.331 | 2479502.749 |
| N-Demethylsambutoxin | 560280.8646 | 574108.4597 | 623981.64 | 469351.5285 | 493119.8376 | 508207.6519 | 806924.6267 | 498720.908 | 516185.5718 | 710405.674 | 875814.4629 | 296872.5725 |

| Metabolites | PBS_1 | PBS_2 | PBS_3 | PBS_4 | BCG_1 | BCG_2 | BCG_3 | BCG_4 | BCGΔ*pe18*_1 | BCGΔ*pe18*_2 | BCGΔ*pe18*_3 | BCGΔ*pe18*_4 |
|---|---|---|---|---|---|---|---|---|---|---|---|---|
| Cer 17:2;2O/2:0 | 633637.8333 | 652829.3985 | 1580002.633 | 434578.8706 | 767031.6164 | 1412951.629 | 750544.1171 | 563599.4086 | 681747.7334 | 640063.4763 | 644897.7904 | 619509.1678 |
| N,N-Dimethyl-L-valine | 137835782.3 | 117051014.6 | 200302285.1 | 135351757.4 | 139594760.4 | 85494688.53 | 171286708.9 | 108928014.8 | 76655833.9 | 108499553.3 | 118673368.6 | 66215072.55 |
| PS(22:6(4Z,7Z,10Z,13Z,16Z,19Z)/22:4(7Z,10Z,13Z,16Z)) | 24911605.6 | 26131194.1 | 9938541.922 | 19467549.5 | 22093289.28 | 19841059.34 | 16754044.21 | 28175858.57 | 33010290.17 | 18959793.02 | 13966466.13 | 18977302.09 |
| LPC O-16:1 | 3128028.058 | 3447135.117 | 2554768.773 | 2670431.003 | 3218389.957 | 3291348.963 | 2660736.795 | 2528952.866 | 2268309.539 | 2201192.291 | 2982504.261 | 1901036.326 |
| PEtOH 18:3_22:6 | 1914555.181 | 1997530.722 | 1658183.758 | 1861260.372 | 2007298.855 | 1855872.74 | 2046370.696 | 2058141.194 | 1821133.713 | 1918580.71 | 1827419.03 | 1815129.376 |
| 3-Aminosalicylic acid | 8286103.245 | 7537939.119 | 5109249.824 | 5580614.551 | 8401624.549 | 9944771.996 | 8007085.594 | 8993085.918 | 8648740.475 | 7578760.194 | 8675553.133 | 4468908.64 |
| 1-O-Hexadecyl-2-O-acetyl-sn-glyceryl-3-phosphorylcholine | 192721140.5 | 203418626.3 | 146187128.4 | 197182017.2 | 245014371.7 | 312188553.3 | 228625964.7 | 210353735.8 | 177026357.8 | 168894851.7 | 181075900.4 | 127232608.2 |
| 3-Methylcrotonylglycine | 3885387.738 | 4257668.891 | 3377109.529 | 4295765.67 | 3135113.468 | 3862165.341 | 4197716.261 | 4260814.094 | 4019333.383 | 2567175.841 | 4698242.768 | 3892568.884 |
| SPB 22:0;2O | 163724168.1 | 80699673.94 | 58212602.83 | 74209074.47 | 39547996.08 | 52356807.42 | 49104791.7 | 47845257.39 | 40207093.13 | 43800769.34 | 41937069.2 | 35341562.92 |
| Flavin mononucleotide (FMN) | 257057.7853 | 270384.2905 | 426575.9917 | 295679.9411 | 2172595.384 | 1954386.1 | 1667184.948 | 1638815.076 | 1727975.892 | 1666389.6 | 1942296.308 | 2201300.16 |

| **Metabolites** | **PBS_1** | **PBS_2** | **PBS_3** | **PBS_4** | **BCG_1** | **BCG_2** | **BCG_3** | **BCG_4** | **BCGΔ*pe18*_1** | **BCGΔ*pe18*_2** | **BCGΔ*pe18*_3** | **BCGΔ*pe18*_4** |
|---|---|---|---|---|---|---|---|---|---|---|---|---|
| 3-Carboxy-6,8-difluoro-7-hydroxycoumarin succinimidyl ester | 301151.5065 | 299870.2679 | 337161.8912 | 292733.3839 | 1418628.214 | 1050132.258 | 1125775.444 | 1204942.308 | 1493413.14 | 1375082.611 | 1216191.103 | 1090770.19 |
| p-(Isoamyloxy)aniline | 7043065.62 | 7867883.745 | 7198403.88 | 8279606.577 | 8890700.623 | 6928053.174 | 7291051.452 | 7248830.826 | 7225814.781 | 7364103.033 | 6964594.37 | 6835950.863 |
| 5-(3-Chlorophenyl)-5-methylimidazolidine-2,4-dione | 152775081.3 | 102978614.5 | 202130296 | 93403545.33 | 177634652.3 | 158944532.8 | 134728273.2 | 110080940.9 | 131068095.7 | 100895824.3 | 99786283.59 | 63297431.01 |
| Tramadol | 5653896.711 | 4516304.335 | 3535760.58 | 3967356.913 | 3585997.039 | 4384019.905 | 4018810.505 | 10109221.25 | 3867413.533 | 3951530.438 | 3547338.83 | 3748116.897 |
| Prostaglandin A2-biotin | 21103.96443 | 22970.38075 | 23179.04803 | 20487.26637 | 19811.14815 | 25137.58924 | 7695.04367 | 14953.0853 | 103525.0499 | 81936.12896 | 143082.4138 | 88360.79263 |
| CAR 16:1 | 10393161.19 | 8061804.803 | 15222867.59 | 11173016.62 | 41649381.25 | 29571323.66 | 48631127.18 | 39254383.91 | 23038835.05 | 30480467.69 | 31993080.74 | 19011212.62 |
| Lopinavir | 633474.6374 | 292741.3967 | 260756.0306 | 571712.9537 | 657714.7578 | 703273.0162 | 318295.1849 | 662482.7839 | 677107.1122 | 443362.2819 | 264471.8593 | 608052.5003 |
| 3-Hydroxyisobutyric acid | 2752904.267 | 1129923.182 | 2140696.638 | 2820984.262 | 2929405.815 | 1769671.562 | 1923798.13 | 4033504.525 | 2325692.257 | 3103243.586 | 1420908.079 | 1532185.252 |
| PC O-24:5 | 789708.3375 | 820194.4667 | 1341901.892 | 695194.5876 | 2361413.653 | 2216604.755 | 2276349.812 | 2373231.442 | 2170172.467 | 889239.9396 | 1687578.256 | 1631770.766 |
| PS 16:0_22:5 | 6307220.868 | 5258721.408 | 2009498.082 | 3598616.303 | 4820783.088 | 3844627.852 | 4017721.974 | 3925242.681 | 5405808.881 | 5808005.894 | 4843888.912 | 6506916.722 |
| Neoeriocitrin | 380210 | 357830 | 270800 | 398023 | 218719 | 240429 | 354800 | 378494 | 206145 | 157842 | 383205 | 405941 |

| Metabolites | PBS_1 | PBS_2 | PBS_3 | PBS_4 | BCG_1 | BCG_2 | BCG_3 | BCG_4 | BCGΔ<br>*pe18*_1 | BCGΔ<br>*pe18*_2 | BCGΔ<br>*pe18*_3 | BCGΔ*p*<br>*e18*_4 |
|---|---|---|---|---|---|---|---|---|---|---|---|---|
| | 54.9 | 55.47 | 45.16 | 12.45 | 74.04 | 53.53 | 34.58 | 05.82 | 15.04 | 42.29 | 30.22 | 52.72 |
| PE(P-16:0/18:1(9Z)<br>) | 202189<br>02.11 | 207556<br>99.87 | 315960<br>64.47 | 256215<br>44.14 | 614782<br>34.42 | 376766<br>19.07 | 552833<br>34.7 | 403985<br>31.33 | 434997<br>32.9 | 510386<br>27.97 | 525165<br>91.16 | 231384<br>44.22 |
| Phloretin_xylosyl-g<br>alactoside | 4339.81<br>9673 | 3520.94<br>4681 | 7393.97<br>3619 | 0 | 939934.<br>48 | 461257.<br>5105 | 114305<br>8.509 | 640040.<br>0293 | 829417.<br>3079 | 776626.<br>243 | 857143.<br>3642 | 525860.<br>3729 |
| 6-Methoxyguanine | 148687.<br>8387 | 76821.2<br>2619 | 341816.<br>4931 | 122787.<br>2676 | 322738<br>2.666 | 221716<br>2.574 | 418658<br>4.295 | 305167<br>2.519 | 260619<br>3.583 | 269012<br>9.497 | 196701<br>8.228 | 267338<br>9.58 |
| MGDG 2:0_10:0 | 158974.<br>7519 | 116629.<br>8436 | 358995.<br>1243 | 374288.<br>4431 | 823963.<br>2542 | 620410.<br>9778 | 867374.<br>2091 | 916611.<br>9392 | 571676.<br>4355 | 476479.<br>2285 | 812860.<br>9997 | 594220.<br>2408 |
| N,N-Dimethyl-4-(6<br>-methylbenzo[d]thi<br>azol-2-yl)aniline | 710296.<br>0965 | 527184.<br>2563 | 931128.<br>5342 | 680615.<br>9358 | 238524<br>7.703 | 200496<br>1.987 | 209120<br>5.925 | 177549<br>1.999 | 135400<br>1.894 | 127452<br>8.174 | 141496<br>4.418 | 142513<br>9.166 |
| PC(36:3) | 109196<br>75.02 | 219743<br>20.53 | 160183<br>30.44 | 144273<br>37.14 | 305715<br>38.04 | 184279<br>42.35 | 242842<br>20.41 | 129834<br>01.37 | 183330<br>55.51 | 826762<br>5.956 | 278564<br>50.89 | 134128<br>55.25 |
| LPC 20:3 | 827399.<br>982 | 830649.<br>2512 | 124809<br>3.698 | 110565<br>3.769 | 166493<br>1.425 | 158387<br>8.671 | 180505<br>7.307 | 157365<br>7.541 | 165932<br>0.05 | 178692<br>2.969 | 171594<br>2.103 | 159290<br>3.76 |
| Kuwanon C | 405426<br>6.231 | 415824<br>6.567 | 364545<br>5.379 | 368315<br>2.624 | 465399<br>8.935 | 485486<br>3.016 | 423394<br>4.92 | 423294<br>5.239 | 420662<br>9.199 | 419700<br>2.731 | 541061<br>2.763 | 363203<br>6.958 |
| CAR 11:1 | 270495<br>6.957 | 280621<br>6.584 | 221993<br>3.74 | 246015<br>8.637 | 243844<br>7.414 | 256348<br>4.521 | 254475<br>7.408 | 240596<br>1.025 | 257163<br>1.672 | 242508<br>9.765 | 235403<br>0.39 | 231973<br>7.962 |
| 5-Bromo-2,4-dimet<br>hoxypyrimidine | 882288<br>5.923 | 839644<br>1.424 | 832825<br>2.079 | 840921<br>8.665 | 857751<br>6.839 | 850842<br>3.393 | 818381<br>9.546 | 814563<br>8.056 | 806875<br>9.097 | 835402<br>0.287 | 811936<br>5.62 | 817363<br>4.51 |
| Tetradecylphospho<br>nate | 244701<br>5.052 | 233259<br>2.62 | 213017<br>5.134 | 225150<br>1.525 | 185301<br>5.127 | 186710<br>1.71 | 203540<br>6.929 | 205547<br>1.04 | 191378<br>5.276 | 193928<br>4.249 | 172843<br>2.982 | 176580<br>1.169 |

| **Metabolites** | **PBS_1** | **PBS_2** | **PBS_3** | **PBS_4** | **BCG_1** | **BCG_2** | **BCG_3** | **BCG_4** | **BCGΔ *pe18*_1** | **BCGΔ *pe18*_2** | **BCGΔ *pe18*_3** | **BCGΔ*pe18*_4** |
|---|---|---|---|---|---|---|---|---|---|---|---|---|
| 3-(4-Bromoanilino)-5,5-dimethyl-2-cyclohexen-1-one | 4447753.533 | 3171511.374 | 3298780.182 | 2648931.947 | 4315216.526 | 3743136.682 | 3576342.758 | 4116012.39 | 3335738.778 | 3371391.532 | 3114357.86 | 1746350.704 |
| PS 18:1_22:6 | 766366.5646 | 922125.9981 | 692379.0235 | 555526.3528 | 1009127.851 | 787929.0662 | 835540.1341 | 1586456.85 | 1007878.545 | 1100373.659 | 978448.2167 | 1045508.238 |
| feruloylserotonin | 860803.8797 | 866978.3205 | 836422.6728 | 846451.3708 | 853197.8345 | 866330.2873 | 862672.596 | 842473.1603 | 826705.2134 | 834112.2338 | 838476.5851 | 822498.064 |
| 1-Butyl-3-methyl-1H-imidazol-3-ium cation | 6414330.904 | 7736660.923 | 2075724.16 | 5271324.116 | 6809720.046 | 6160804.714 | 9373322.146 | 5521107.9 | 6237940.253 | 8932642.966 | 5416727.292 | 4508377.837 |
| Hydroxyacetone | 2367664.343 | 1965313.675 | 3911996.874 | 2351510.915 | 30657228.19 | 19063455.83 | 34051881 | 23549901.41 | 16634331.86 | 24645657.07 | 23895663.12 | 16507286.77 |
| PG 18:0_18:1;O | 858100.3613 | 686959.715 | 1535587.854 | 812719.8224 | 1936834.79 | 2390119.128 | 1879099.964 | 1918217.432 | 1118180.702 | 1443950.361 | 1824677.113 | 981313.9836 |
| (3Z,5E)-3,5-Bis(4-methoxybenzylidene)tetrahydro-4H-thiopyran-4-one | 1181854.97 | 2165969.047 | 825109.0804 | 2461037.171 | 6350582.942 | 2276732.882 | 2379627.136 | 1128172.171 | 2645703.409 | 5881482.483 | 2559994.479 | 1576673.448 |
| Caplamin | 19228390.47 | 17559386 | 27664863.81 | 21523278.61 | 27948823.78 | 25753887.04 | 27926919.99 | 25750362.27 | 27992538.68 | 19739541.63 | 17808707.96 | 24815507.65 |
| PE-Cer 12:2;2O/16:1;O | 10076738.03 | 10882834.41 | 9812632.703 | 11161568.64 | 10693040.56 | 10812607.06 | 10353999.8 | 9879961.475 | 9922088.757 | 10093587.91 | 9939990.521 | 8950205.033 |
| FAHFA 2:0/18:0 | 303503.0505 | 456994.495 | 289450.9569 | 195098.9792 | 505472.8588 | 251171.9693 | 246401.3525 | 399549.2028 | 326422.3935 | 679852.4052 | 348318.7737 | 661475.2642 |

| Metabolites | PBS_1 | PBS_2 | PBS_3 | PBS_4 | BCG_1 | BCG_2 | BCG_3 | BCG_4 | BCGΔ*pe18*_1 | BCGΔ*pe18*_2 | BCGΔ*pe18*_3 | BCGΔ*pe18*_4 |
|---|---|---|---|---|---|---|---|---|---|---|---|---|
| 1-palmitoyl-2-vernoloyl-phosphatidylcholine | 166027.4757 | 185888.0004 | 129291.4943 | 182849.5789 | 3694054.794 | 2315277.316 | 3726667.765 | 2593796.799 | 2819866.424 | 3314634.638 | 3407631.233 | 1488084.104 |
| Cer 8:1;2O/9:0 | 1220575.031 | 1333692.693 | 1184246.943 | 1174362.292 | 1296699.551 | 1315410.637 | 1349442.158 | 1312112.461 | 1346526.333 | 1254246.997 | 1225907.372 | 1254218.91 |
| (2R)-7-Methoxy-3-oxo-3,4-dihydro-2H-1,4-benzoxazin-2-yl .beta.-D-glucopyranoside | 5409330.877 | 4923834.956 | 6047027.82 | 5407595.362 | 54913959.16 | 47811590.17 | 47290857.11 | 45929564.68 | 50206143.39 | 42837447.99 | 48618725.13 | 43716623.78 |
| Acetic acid, 2-[4-[(5,6-diphenyl-2-pyrazinyl)(1-methylethyl)amino]butoxy]- | 2757047.534 | 3882382.681 | 2046999.802 | 1963134.731 | 1066447.207 | 2201336.677 | 1293562.573 | 1421055.878 | 1049320.897 | 875140.156 | 1641987.915 | 6076558.163 |
| 10-Hydroxydecanoic acid | 4201385.048 | 3577645.123 | 3585596.051 | 3140968.291 | 3621251.137 | 4458876.747 | 3944254.46 | 2756969.87 | 2959485.635 | 3664676.18 | 3226333.106 | 3890517.245 |
| Palmitoylcarnitine (Car(16:0)) | 109201692.4 | 93467889.13 | 120405461.5 | 97730120.46 | 241901444.5 | 143868290.2 | 249844872.5 | 192397659.8 | 127108874.5 | 169380553.2 | 179959249.1 | 89405021.49 |
| Testosterone cypionate | 960085.6372 | 646544.6365 | 694131.9297 | 707443.5338 | 8618005.293 | 11180874.5 | 5458199.354 | 7629737.391 | 7762612.697 | 3550888.669 | 4617715.093 | 6440086.367 |
| PC O-42:6 | 1101419.203 | 1016725.517 | 1226004.359 | 1783900.319 | 755337.2158 | 0 | 1369229.976 | 1917212.635 | 1365865.836 | 2760459.737 | 568191.1752 | 3400630.538 |
| Coumestrol | 416820 | 331016 | 595645 | 538052 | 547248 | 581051 | 566379 | 524580 | 507545 | 393785 | 428489 | 467818 |

| Metabolites | PBS_1 | PBS_2 | PBS_3 | PBS_4 | BCG_1 | BCG_2 | BCG_3 | BCG_4 | BCGΔ*pe18*_1 | BCGΔ*pe18*_2 | BCGΔ*pe18*_3 | BCGΔ*pe18*_4 |
|---|---|---|---|---|---|---|---|---|---|---|---|---|
| | 0.685 | 9.281 | 0.943 | 5.602 | 4.396 | 3.903 | 9.031 | 4.919 | 0.926 | 7.767 | 8.404 | 9.265 |
| gamma-Glutamylleucine | 4383889.723 | 3226888.892 | 5759628.632 | 3896520.465 | 29447550.56 | 25532273.72 | 23642309.23 | 23672470.68 | 18150093.14 | 15910252.86 | 17200722.13 | 18632660.91 |
| FAHFA 7:0/20:2 | 87794.73672 | 589355.3505 | 522834.8271 | 533472.1385 | 1330617.083 | 599082.279 | 513357.1292 | 795606.7026 | 666267.4037 | 822904.2776 | 527824.5545 | 801327.411 |
| LPE O-20:5 | 2254133.13 | 2448876.102 | 2192372.798 | 2030323.183 | 2102602.538 | 3900166.626 | 2416457.222 | 2041137.571 | 1940702.252 | 2262831.006 | 2811807.654 | 1669524.474 |
| LPG O-22:5 | 365596.866 | 264436.0831 | 326558.2973 | 323136.1992 | 718764.9901 | 657126.5909 | 725828.0273 | 599231.4868 | 704705.4396 | 566080.0739 | 748242.8063 | 487820.3036 |
| PC O-37:4 | 1605469.618 | 913804.7306 | 1343136.424 | 1889936.888 | 1762283.003 | 239868.9365 | 3297867.801 | 4647882.945 | 2811235.473 | 1856087.577 | 1667817.376 | 3117277.059 |
| SL 10:1;O/16:3 | 1928525.409 | 1976506.404 | 3121969.666 | 2159916.252 | 3088089.197 | 3233526.427 | 4150216.571 | 3023696.193 | 2657724.388 | 2261675.166 | 2262002.33 | 2390269.99 |
| PE O-18:0_20:4 | 618850.7133 | 369428.5129 | 863928.6125 | 614691.7389 | 899921.8054 | 6470.494325 | 878111.1892 | 870499.7453 | 876554.4821 | 1690813.64 | 776020.5722 | 1218536.972 |
| [2-[docosa-7.10.13.16-tetraenoyloxy]-3-(octadecanoyloxy)propoxy]({[2.3.4.5.6-pentahydroxycyclohexyl]oxy})phosphinic acid | 462982.4471 | 599737.2366 | 405922.634 | 475326.2766 | 544440.7109 | 40830.20774 | 1036096.616 | 532158.3296 | 580699.1594 | 548215.5653 | 258600.7552 | 417369.3745 |
| (1,5-Dimethyl-1H-pyrazol-3-yl)methyla | 2442591.414 | 2588638.491 | 1907643.538 | 3118774.246 | 2870977.252 | 5157300.994 | 2813191.019 | 5003662.327 | 3381368.658 | 2306789.635 | 2427905.985 | 2204678.362 |

| Metabolites | PBS_1 | PBS_2 | PBS_3 | PBS_4 | BCG_1 | BCG_2 | BCG_3 | BCG_4 | BCGΔ*pe18*_1 | BCGΔ*pe18*_2 | BCGΔ*pe18*_3 | BCGΔ*pe18*_4 |
|---|---|---|---|---|---|---|---|---|---|---|---|---|
| mine | | | | | | | | | | | | |
| N-Cyclohexylcyclohexanecarboxamide | 16242501.6 | 18433912.87 | 12854269.28 | 12896278.21 | 14958810.65 | 18975396.7 | 16777079.27 | 16658011.12 | 16134154.52 | 14938980.34 | 15160168.68 | 17974792.3 |
| Piperazine-2-carboxylic acid | 596458.8039 | 4037253.343 | 3425649.121 | 3346716.899 | 7104755.329 | 1796779.064 | 1385787.891 | 1150121.103 | 2407398.883 | 6208902.483 | 2481023.171 | 2086994.113 |
| UDP-N-acetylglucosamine | 1556144 | 1517204.241 | 1776758.213 | 1541326.704 | 6073912.678 | 3485381.035 | 6198793.944 | 4376482.058 | 4356450.29 | 4481503.54 | 4672777.144 | 4148729.162 |
| Linoleic acid | 80539996.79 | 71323316.76 | 105907536.1 | 93882371.18 | 1053268719 | 1280782831 | 988615374.1 | 1091616553 | 769056221 | 454834722.3 | 970687816.4 | 889755476.9 |
| 2-(4-Chloro-2-methylphenoxy)-N-(2-hydroxyethyl)acetamide | 668196.0437 | 835317.3469 | 1361062.706 | 783987.7962 | 772506.3309 | 1117910.699 | 534573.7 | 848893.9794 | 637188.4722 | 504256.6137 | 831964.0047 | 811869.0459 |
| 5-[2-Chloro-5-(trifluoromethyl)anilino]-5-oxopentanoic acid | 11528971.68 | 4446364.641 | 4007014.42 | 6509284.166 | 9488187.649 | 11382384.34 | 12631068.78 | 9282570.589 | 10296399.1 | 6772114.207 | 9051174.211 | 4410325.834 |
| SMGDG O-9:0_8:0 | 350119.5066 | 539147.1205 | 497779.6783 | 457608.0631 | 1630522.853 | 1832296.403 | 229594.5425 | 192794.7319 | 275826.2022 | 2089243.15 | 2166354.258 | 859904.3063 |
| PC 8:0_28:4 | 73452652.77 | 87341118.73 | 75862290.52 | 90942425.96 | 40599774.86 | 45007728.91 | 51727582.75 | 39783796.92 | 44713780.17 | 50915400.74 | 52046448.29 | 13458830.48 |
| Zopiclone | 5041759.515 | 4241574.615 | 6582502.721 | 5275902.228 | 5257289.771 | 6361807.761 | 5694120.2 | 5137755.097 | 5790186.577 | 5020310.45 | 5384811.456 | 5179302.11 |

| **Metabolites** | **PBS_1** | **PBS_2** | **PBS_3** | **PBS_4** | **BCG_1** | **BCG_2** | **BCG_3** | **BCG_4** | **BCGΔ*pe18*_1** | **BCGΔ*pe18*_2** | **BCGΔ*pe18*_3** | **BCGΔ*pe18*_4** |
|---|---|---|---|---|---|---|---|---|---|---|---|---|
| 5-Bromo-3-(2,2,2-trifluoroethoxy)pyrazin-2-amine | 3794211.321 | 3399292.422 | 3073526.778 | 3528922.587 | 11085630.92 | 10160923.1 | 10801574.89 | 10207849.45 | 10706478.83 | 9517759.261 | 9418738.472 | 8235986.42 |
| 4-(4-Chlorophenyl)-4,5,6,7-tetrahydro-3H-imidazo[4,5-c]pyridine | 6559.740238 | 0 | 0 | 0 | 2095686.032 | 2424706.206 | 4079114.585 | 4242268.185 | 4577421.334 | 1680045.202 | 2919375.304 | 5625506.837 |
| 6.beta.-Hydroxyeplerenone | 2338871.507 | 269726.2192 | 346904.9787 | 523814.222 | 364609.121 | 3499337.398 | 88070.47563 | 491133.9009 | 2545631.803 | 367560.7969 | 1767927.446 | 200131.258 |
| 9Z-Octadecenoic acid, (2S)-3-[(hydroxymercaptophosphinyl)oxy]-2-methoxypropyl ester | 379147.7197 | 397945.0485 | 533169.9729 | 445884.8387 | 611654.6377 | 694313.6117 | 611949.5536 | 643861.1832 | 533659.4258 | 398821.3802 | 450371.9222 | 462514.8177 |
| PS 22:5_22:5 | 945100.9322 | 871345.9891 | 889253.3511 | 1846894.512 | 2914478.702 | 1367943.123 | 1678853.658 | 4364181.321 | 5231625.733 | 5446631.963 | 1635669.633 | 1422831.671 |
| PC(14:0/18:1(9Z)) | 11092789.77 | 12578761.85 | 9814804.281 | 14451390.82 | 12247670.69 | 13244343.12 | 13450721.52 | 12708308.18 | 9717510.642 | 11065764.59 | 9627417.702 | 11213466.69 |
| 4(1H)-Pyridinone, 5-acetyl-2,3-dihydro-2,6-dimethyl- | 969063.9068 | 1056852.183 | 888855.1035 | 1311500.936 | 962381.1545 | 935485.3546 | 999036.5356 | 920033.1916 | 941913.7008 | 945772.7235 | 918306.7793 | 974847.0881 |
| Ceanothic acid | 674472.193 | 714667.5468 | 696785.8111 | 771754.4664 | 1105795.707 | 887249.0817 | 1253755.236 | 746751.726 | 756914.0408 | 756414.8734 | 759191.928 | 1044712.084 |

| Metabolites | PBS_1 | PBS_2 | PBS_3 | PBS_4 | BCG_1 | BCG_2 | BCG_3 | BCG_4 | BCGΔ*pe18*_1 | BCGΔ*pe18*_2 | BCGΔ*pe18*_3 | BCGΔ*pe18*_4 |
|---|---|---|---|---|---|---|---|---|---|---|---|---|
| 2-Amino-3-(1H-pyrazol-1-yl)propanoic acid | 7868384.739 | 6662452.716 | 47607932.25 | 35433378.45 | 12627539.32 | 5744125.502 | 30337695.44 | 7320142.395 | 12570887.49 | 7133380.486 | 32911839.29 | 12556837.94 |
| Glu-Val | 1883764.563 | 1418714.704 | 2414188.229 | 1635178.984 | 6224683.265 | 5004145.794 | 5302358.616 | 4843896.872 | 3558662.175 | 3497609.908 | 3652681.471 | 3490494.582 |
| FA 16:2 | 2139570.396 | 3339533.158 | 2314354.683 | 2014040.502 | 2506266.949 | 2630842.101 | 2441535.83 | 2322002.487 | 1874341.805 | 2233635.628 | 2404576.704 | 2309793.87 |
| Levonorgestrel butyrate | 1741165.338 | 2746863.743 | 1054277.605 | 1763252.518 | 1237855.593 | 1728288.735 | 1313623.841 | 1486880.824 | 1077134.551 | 1000040.05 | 949717.2503 | 1149469.121 |
| PS 9:0_26:1 | 1298865.017 | 1284567.631 | 900785.3056 | 691617.7134 | 1236953.26 | 1124979.047 | 944295.8369 | 1213169.286 | 1154841.387 | 941363.9918 | 1024896.355 | 1012052.491 |
| 1-Dodecylpyridinium cation | 3955528.209 | 4231143.563 | 3888329.702 | 3830487.418 | 3933671.828 | 4582611.578 | 4403813.054 | 4174282.365 | 4099647.887 | 4025843.467 | 4651353.423 | 3765274.215 |
| Pristanic acid | 1787303.981 | 1309296.41 | 645124.5159 | 1031230.664 | 5254245.92 | 7186545.201 | 5219022.585 | 6710372.989 | 6715164.199 | 4916376.128 | 6194490.035 | 2711212.478 |
| PC 9:0_22:2 | 1041209.383 | 791276.2545 | 256426.2458 | 269854.9262 | 6730374.312 | 5551860.905 | 6540397.497 | 5311676.126 | 5840244.627 | 9315171.032 | 4658502.548 | 3542975.37 |
| 5.alpha.-Hydroxy-6-ketocholesterol | 220819.3739 | 203402.6432 | 141021.7728 | 202206.1104 | 295290.3532 | 304632.6468 | 219488.5492 | 232481.5709 | 225493.8806 | 211234.9113 | 233345.4626 | 202687.1687 |
| 1-Hexadecanoyl-2-(9Z-octadecenoyl)-sn-glycero-3-phosphocholine [M] | 32593292.15 | 24743065.58 | 131916388.3 | 25995051.7 | 38086367.83 | 1910061.041 | 17754752.5 | 14918706.79 | 15676159.25 | 22955335.16 | 24672875.88 | 8553122.662 |

| Metabolites | PBS_1 | PBS_2 | PBS_3 | PBS_4 | BCG_1 | BCG_2 | BCG_3 | BCG_4 | BCGΔ*pe18*_1 | BCGΔ*pe18*_2 | BCGΔ*pe18*_3 | BCGΔ*pe18*_4 |
|---|---|---|---|---|---|---|---|---|---|---|---|---|
| 5-Heptenoic acid, 7-[(1S,2S,3S,5R)-3-[[(4-iodophenyl)sulfonyl]amino]-6,6-dimethylbicyclo[3.1.1]hept-2-yl]-, (5Z)- | 4538911.259 | 2841006.694 | 3761049.758 | 3138958.729 | 3622451.832 | 3537004.677 | 3596075.359 | 2661241.389 | 3533562.094 | 1781896.624 | 1730053.366 | 2528402.997 |
| 5-Nitro-N-(3-pyridinylmethyl)-2-furamide | 81148000.11 | 54218600.78 | 92671107.99 | 55791149.29 | 113527110 | 91411980.5 | 100756905.9 | 97544517.69 | 93446128.26 | 90376458.55 | 97473613.69 | 72773922.39 |
| Propanoic acid, 2-[[4-[2-[[(cyclohexylamino)carbonyl](4-cyclohexylbutyl)amino]ethyl]phenyl]thio]-2-methyl- | 0 | 54991.13573 | 0 | 27723.23121 | 10701853.32 | 11555054.26 | 9900884.45 | 12356361.36 | 9961291.504 | 7113478.755 | 10954404.03 | 7304400.887 |
| Pro-Glu | 551270.7556 | 454222.355 | 625794.5423 | 473103.4427 | 2211620.442 | 1472746.658 | 2268356.374 | 1673780.672 | 2321365.567 | 2165213.742 | 2453158.035 | 2123815.394 |
| (3beta,23E)-3-Hydroxy-27-norcycloart-23-en-25-one | 8638323.112 | 6373313.781 | 11078852.58 | 7808453.714 | 17119031.04 | 11897786.86 | 18529545.06 | 15670323.06 | 9054924.195 | 12377294.65 | 12261880.33 | 6918433.743 |
| PA 2:0_16:3 | 397463.4126 | 323081.5691 | 399219.2119 | 575978.6189 | 259821.0644 | 403659.9338 | 212908.4978 | 185299.6216 | 179979.9131 | 288713.7016 | 247436.6749 | 210718.8248 |
| 4-Methylcyclohexaneacetic acid | 3602973.993 | 2592971.937 | 3141066.038 | 3745140.021 | 2858893.81 | 3242489.513 | 4758964.581 | 2394120.771 | 3482063.735 | 3498404.454 | 2853533.209 | 3939383.167 |

| Metabolites | PBS_1 | PBS_2 | PBS_3 | PBS_4 | BCG_1 | BCG_2 | BCG_3 | BCG_4 | BCGΔ*pe18*_1 | BCGΔ*pe18*_2 | BCGΔ*pe18*_3 | BCGΔ*pe18*_4 |
|---|---|---|---|---|---|---|---|---|---|---|---|---|
| 2,4-Dihydroxy-6-(8-pentadecenyl)benzoic acid | 1514114.836 | 750753.9486 | 2101901.013 | 1166859.57 | 14588260.33 | 15277584.16 | 24108620.55 | 17238811.93 | 12034452.09 | 7951124.326 | 15039011.99 | 12824103.89 |
| Kaur-16-en-18-al | 10958880.38 | 11456850.83 | 9752538.139 | 9871712.239 | 4232005.619 | 6577130.215 | 4161466.963 | 5577989.37 | 5624548.14 | 5067018.051 | 4957112.687 | 5693671.367 |
| Masitinib (AB1010) | 1493857.973 | 916964.8825 | 1630074.567 | 1033935.744 | 1397649.561 | 1694782.551 | 1399508.712 | 1404311.997 | 1032680.957 | 1445328.47 | 1642419.315 | 990355.0401 |
| 5,6,7,8-Tetrahydro-2(1H)-quinolinone | 617519.3766 | 691869.1396 | 580808.0455 | 673587.6022 | 596998.5054 | 587018.0377 | 591109.1731 | 549014.1545 | 577162.1556 | 553550.0382 | 545393.1253 | 725563.7961 |
| Hex2Cer 16:2;2O/14:0 | 3190593.829 | 2311179.967 | 2307447.193 | 2984595.881 | 18365439.86 | 23970082.66 | 17029290.73 | 18673455.23 | 15125492.16 | 7482631.493 | 14367787.08 | 12728845.06 |
| Carpropamid | 3437538.496 | 2664112.382 | 5586789.703 | 3679263.863 | 6037297.359 | 4798094.469 | 6153605.058 | 5628105.08 | 5462229.591 | 3817387.399 | 3973041.259 | 4557422.425 |
| SQDG 13:1_18:5 | 617167.3615 | 677852.8375 | 301801.3381 | 493786.5826 | 778266.6833 | 767141.9005 | 707920.8809 | 984954.5521 | 1218508.417 | 915828.8992 | 910143.9526 | 678057.268 |
| Aviprin | 1346429.045 | 2872660.29 | 3967668.673 | 3333092.391 | 1079072.249 | 1065967.916 | 1370233.418 | 1016505.068 | 1476350.791 | 1207044.2 | 1220620.597 | 7825123.625 |
| Protoporphyrin IX | 309121.7115 | 310492.7846 | 499383.2778 | 375223.635 | 1366848.604 | 2037890.359 | 1847727.994 | 1254370.065 | 5520717.448 | 827529.5569 | 1077468.812 | 1506588.481 |
| 2-(3,4,5-Trimethoxyphenyl)-1H-benzimidazole | 5442651.263 | 5789200.565 | 5504582.942 | 4879904.802 | 5665551.826 | 6262296.139 | 5722674.902 | 6304687.575 | 5895898.519 | 5221040.54 | 6070869.7 | 4892963.222 |
| O-Acetylserine | 5943135.322 | 4369407.873 | 9930433.663 | 5913451.819 | 17283168.11 | 9338702.859 | 18536796.29 | 12043296.05 | 9154812.451 | 11429635.46 | 11437205.88 | 7455358.399 |

| **Metabolites** | **PBS_1** | **PBS_2** | **PBS_3** | **PBS_4** | **BCG_1** | **BCG_2** | **BCG_3** | **BCG_4** | **BCGΔ*pe18*_1** | **BCGΔ*pe18*_2** | **BCGΔ*pe18*_3** | **BCGΔ*pe18*_4** |
|---|---|---|---|---|---|---|---|---|---|---|---|---|
| Benzenebutanamide, N-(4-oxo-4H-3,1-benzothiazin-2-yl)- | 9534916.642 | 7986072.732 | 17570625.43 | 11292731.9 | 11471191.19 | 10693638.82 | 11420793.84 | 10194073.86 | 9947507.81 | 6679646.658 | 9608695.921 | 9863249.6 |
| N-Acetyl mesalazine | 72623.65198 | 65819.64924 | 144235.1711 | 156063.5863 | 3365220.162 | 2455953.649 | 3828400.762 | 3085948.006 | 1849736.991 | 2315884.67 | 2878542.098 | 1657204.809 |
| 2'-O-Methylguanosine | 35617.39858 | 62236.73052 | 84747.8086 | 65231.19976 | 1149747.939 | 623994.8793 | 1082223.505 | 806154.1849 | 909545.7401 | 907166.9842 | 1106006.168 | 734863.341 |
| 4-[2-(Trifluoromethyl)phenyl]-4H-1,2,4-triazole-3-thiol | 413233.8302 | 351135.9168 | 562190.0815 | 456911.9186 | 3362644.179 | 2522731.168 | 3268863.468 | 3270327.479 | 3162416.576 | 2644313.373 | 2635546.191 | 2538510.854 |
| Cohibin_A | 0 | 0 | 0 | 0 | 1233186.788 | 890843.2199 | 1213532.111 | 828015.8107 | 1164406.358 | 1427445.697 | 1202084.621 | 713224.2239 |
| DEOXYCARNITINE | 14230878.58 | 14668293.69 | 20817017.06 | 19172770.8 | 12695832.88 | 7486553.263 | 14714883.03 | 10036480.56 | 7536877.171 | 12739255.85 | 10764491.39 | 17687544.94 |
| PC(36:6) | 2331311.766 | 4362142.143 | 2316387.459 | 2676910.082 | 12630111.17 | 7001776.192 | 13512984.56 | 6883125.763 | 5874736.646 | 1512192.398 | 13201766.57 | 6359561.836 |
| Resolvin_D2 | 342066.9728 | 207219.6941 | 3980022.318 | 8086138.77 | 8852202.811 | 12078138.98 | 9439869.332 | 11168271.81 | 7123143.499 | 3738854.246 | 9097943.255 | 10111009.97 |
| Phe-Glu | 632000.6873 | 518773.3691 | 835778.926 | 449255.5198 | 7161674.842 | 9915606.326 | 8202975.196 | 8184961.512 | 6438301.311 | 4165334.397 | 4442809.934 | 6330035.804 |
| Butorphanol | 2549391.003 | 2621669.908 | 3748388.915 | 2432665.518 | 2357769.533 | 2487811.124 | 2542724.375 | 3573568.264 | 2376616.654 | 2352351.833 | 1989569.38 | 2129697.337 |
| Cinidon-ethyl | 129606 | 117243 | 222616 | 153159 | 237254 | 230690 | 249984 | 221297 | 234764 | 148684 | 157143 | 193592 |

| **Metabolites** | **PBS_1** | **PBS_2** | **PBS_3** | **PBS_4** | **BCG_1** | **BCG_2** | **BCG_3** | **BCG_4** | **BCGΔ*pe18*_1** | **BCGΔ*pe18*_2** | **BCGΔ*pe18*_3** | **BCGΔ*pe18*_4** |
|---|---|---|---|---|---|---|---|---|---|---|---|---|
| | 5.57 | 3.155 | 5.326 | 8.088 | 6.703 | 8.509 | 4.871 | 5.274 | 1.965 | 5.438 | 8.78 | 0.532 |
| 2-Oxo-6-phenyl-4-(trifluoromethyl)-1,2-dihydropyridine-3-carbonitrile | 899140.282 | 1023383.517 | 1792245.119 | 1019152.036 | 5900105.338 | 3348652.473 | 5712919.928 | 4014982.985 | 6165541.355 | 6181630.764 | 5845279.774 | 5073906.327 |
| CAR 16:2 | 841000.6561 | 387933.77 | 842910.4686 | 973348.3134 | 756815.1602 | 745989.0794 | 1133931.263 | 850223.3552 | 548981.1466 | 686270.4859 | 602753.8283 | 535862.4983 |
| 5-(4-Aminophenyl)-4-methyl-2,4-dihydro-3H-1,2,4-triazole-3-thione | 2545423.67 | 2365017.92 | 2117612.99 | 2388277.845 | 2389604.248 | 2078751.216 | 2160100.098 | 2299974.636 | 1968265.614 | 2175688.196 | 2373028.644 | 2294774.461 |
| Ricinoleic acid | 1542112.664 | 932925.462 | 3173085.832 | 2271013.22 | 1442432.103 | 1973002.314 | 2689046.252 | 2585761.571 | 2775408.59 | 1096126.344 | 2383978.784 | 2056199.039 |
| 3-Ureidopropionic acid | 437330.27 | 342806.3179 | 463023.4898 | 257896.7347 | 1507851.344 | 614432.0744 | 807477.9705 | 1045173.871 | 988776.2317 | 1250058.25 | 650863.0446 | 746852.3368 |
| DGCC 8:0_30:7 | 1956477.464 | 1229046.843 | 1763927.14 | 1699642.572 | 4477643.385 | 4820362.751 | 3389046.457 | 3672051.253 | 3600324.709 | 2704461.5 | 2753031.733 | 2482450.557 |
| Fructose | 158677464.2 | 121019993.9 | 163578314.7 | 148581193.5 | 122842412.9 | 145645511.8 | 121670957.5 | 128739498.5 | 128503914 | 74437631.99 | 66910977.38 | 124800632.7 |
| SL 21:2;O/14:0 | 11639355.58 | 13287165.81 | 8666318.472 | 14237076.07 | 27602876.04 | 35527828.59 | 14319797.63 | 23141139.65 | 29866604.25 | 8437921.11 | 7826691.49 | 18526069.84 |
| Ganoderic Acid G | 2020179.171 | 2381912.436 | 1839639.952 | 2150062.048 | 5736459.262 | 9204477.118 | 4280660.071 | 4893808.219 | 6647375.594 | 5121129.461 | 9038102.71 | 5410270.596 |

| Metabolites | PBS_1 | PBS_2 | PBS_3 | PBS_4 | BCG_1 | BCG_2 | BCG_3 | BCG_4 | BCGΔ*pe18*_1 | BCGΔ*pe18*_2 | BCGΔ*pe18*_3 | BCGΔ*pe18*_4 |
|---|---|---|---|---|---|---|---|---|---|---|---|---|
| Anomalin A | 646478.1539 | 604649.6888 | 1184275.008 | 688317.0065 | 1641668.224 | 1326804.56 | 1749418.143 | 1521292.021 | 1352534.912 | 1021138.782 | 1103338.457 | 1229412.738 |
| Antibiotic F 1839A | 259714.3483 | 369476.3276 | 295426.4209 | 290691.6857 | 344033.5485 | 393310.4757 | 329841.2368 | 347196.3279 | 313899.5948 | 344488.5452 | 348082.8717 | 281820.0905 |
| Cytosine | 3375278.156 | 2734241.967 | 4203952.682 | 3154101.383 | 6116536.472 | 4020507.048 | 4784645.029 | 4640644.112 | 3836833.005 | 3189070.827 | 3909107.861 | 2905792.016 |
| 6-Methyluracil | 2283071.174 | 4658153.679 | 4455820.253 | 2977935.623 | 3457180.66 | 3145516.486 | 3492190.948 | 2929402.127 | 3138483.218 | 2274327.61 | 2862304.327 | 2886945.854 |
| PC 8:0_24:3 | 5742773.322 | 6431491.333 | 3473067.588 | 4889256.884 | 12085245.74 | 11952329.63 | 12792567.47 | 12488015.12 | 20350870.43 | 14029981.79 | 10769099.92 | 14101681.62 |
| 1-O-(9Z-Octadecenyl)-sn-glycero-2,3-cyclic-phosphate | 715921.9789 | 1183055.042 | 508566.8286 | 863761.1322 | 2477597.344 | 2598968.242 | 2411256.381 | 1975156.737 | 1573141.692 | 1132957.558 | 2092224.521 | 1797962.838 |
| PS 10:0_26:0 | 1081249.265 | 585295.2506 | 731934.5178 | 714576.6802 | 634231.3819 | 566457.6786 | 709997.5476 | 1037500.807 | 577262.1117 | 674192.8254 | 928848.9181 | 905444.0597 |
| DGDG O-17:0_22:3 | 1748304.347 | 1021798.618 | 1290386.446 | 1558551.206 | 4307434.204 | 2981606.765 | 4166704.175 | 3368016.625 | 2552090.784 | 2901679.597 | 4137852.752 | 1351172.559 |
| MGDG O-16:4_20:4 | 428433.4891 | 541253.6943 | 200104.2868 | 188218.0969 | 549516.0931 | 979494.5462 | 751066.9268 | 618150.0672 | 481142.5736 | 596694.607 | 700334.6458 | 545732.6434 |
| beta-arginine | 1053536.35 | 1346788.573 | 5666588.447 | 2176615.942 | 9016831.959 | 4992290.528 | 11661432.69 | 6990722.295 | 4020470.491 | 5268731.712 | 6190947.342 | 8800516.22 |
| ACRYLAMIDE | 2489483.655 | 1188959.525 | 803401.0671 | 3832405.694 | 1049752.838 | 999806.594 | 1013223.944 | 1293346.187 | 996380.6782 | 897231.4747 | 1043327.851 | 1181868.638 |
| PC O-16:4_22:6 | 289361. | 307800. | 250183. | 238938. | 331113. | 449294. | 366626. | 307341. | 265615. | 303239. | 338888. | 344469. |

| **Metabolites** | **PBS_1** | **PBS_2** | **PBS_3** | **PBS_4** | **BCG_1** | **BCG_2** | **BCG_3** | **BCG_4** | **BCGΔ*pe18*_1** | **BCGΔ*pe18*_2** | **BCGΔ*pe18*_3** | **BCGΔ*pe18*_4** |
|---|---|---|---|---|---|---|---|---|---|---|---|---|
| | 9986 | 213 | 5814 | 1069 | 8502 | 8886 | 7202 | 4732 | 2963 | 9545 | 3159 | 4102 |
| Bensulfuron-methyl | 3971551.361 | 3501983.291 | 5751179.123 | 3666232.581 | 4686975.492 | 4288146.625 | 3872292.791 | 3776488.679 | 4245780.766 | 3590789.907 | 4236245.56 | 3015795.48 |
| 9-Oxo-10(E),12(E)-octadecadienoic acid | 11897721.46 | 7659639.407 | 13705272.04 | 11809958.4 | 6916785.446 | 8521065.084 | 6537416.972 | 7433629.22 | 7263813.224 | 6561388.124 | 8316009.909 | 6523464.681 |
| Celastrol | 494725.4475 | 414288.8426 | 872413.6079 | 1101989.362 | 11050791.97 | 9512968.192 | 12557496.82 | 10200998.12 | 6864871.674 | 6383439.303 | 8887070.036 | 5192678.811 |
| Cer 15:0;2O/2:0 | 3726767.684 | 8714861.983 | 3487493.972 | 3650784.105 | 8827361.858 | 3619112.445 | 3923063.887 | 3449397.868 | 3354865.489 | 3315713.055 | 3598974.418 | 3237858.737 |
| 3-hydroxybenzaldehyde | 4827781.538 | 4546814.131 | 4376810.209 | 5521992.087 | 6802046.662 | 6911741.092 | 4234523.184 | 6761522.72 | 4336655.422 | 4380442.328 | 4307332.475 | 4419341.884 |
| Pentazocine | 4295249.744 | 4506498.741 | 3890984.96 | 4206626.306 | 4412887.804 | 4250035.546 | 4482685.987 | 4324949.582 | 4265676.48 | 4125850.37 | 3862160.659 | 4120969.976 |
| MGDG O-13:1_22:3 | 585822.0325 | 129214.3626 | 628252.6077 | 342457.7304 | 547592.4769 | 7023.37584 | 509234.505 | 187905.5728 | 810310.2606 | 761826.7834 | 542338.4379 | 482772.8628 |
| PE O-26:4_18:4 | 895887.0701 | 790074.7671 | 902840.576 | 903392.8153 | 819669.39 | 1001348.808 | 846695.3179 | 868371.8661 | 797496.7373 | 835701.9767 | 918832.8696 | 969097.4736 |
| 2-oxindole-3-acetate | 577009.1924 | 444918.3002 | 542949.5218 | 540813.9465 | 744606.5409 | 677989.9439 | 662778.4342 | 782480.8615 | 646893.4556 | 342973.5424 | 388606.9536 | 601473.4175 |
| N,N-Dimethylarginine (ADMA) | 44527991.81 | 37597247.04 | 61740108.82 | 43134045.77 | 71165942.84 | 48446983.98 | 72306790.62 | 36819340.52 | 54487217.3 | 50164742.72 | 56233690.58 | 48652255.69 |
| PC O-12:0 | 59546.59684 | 89275.41698 | 96460.19559 | 90699.2578 | 660651.5068 | 552977.7083 | 770289.6969 | 594041.8357 | 661256.8695 | 705791.3425 | 751096.1728 | 523624.4515 |

| Metabolites | PBS_1 | PBS_2 | PBS_3 | PBS_4 | BCG_1 | BCG_2 | BCG_3 | BCG_4 | BCGΔ*pe18*_1 | BCGΔ*pe18*_2 | BCGΔ*pe18*_3 | BCGΔ*pe18*_4 |
|---|---|---|---|---|---|---|---|---|---|---|---|---|
| 5-[(2-Chlorophenoxy)methyl]-1H-tetrazole | 1245676.861 | 956761.4805 | 2252856.256 | 1196576.761 | 6479496.383 | 4230441.43 | 6200379.128 | 4666577.537 | 4307834.895 | 3319930.487 | 2873503.421 | 3126742.494 |
| 9,12,13-TriHOME | 26100359.01 | 24047356.28 | 30984621.46 | 27229400.74 | 25712777.24 | 29393989.73 | 27521421.09 | 27461016.77 | 25481743.93 | 28877686.97 | 26469018.26 | 25235389.65 |
| Ethoxyacetic acid | 4399360.742 | 4016482.749 | 3873866.109 | 2485475.343 | 5727629.546 | 5211978.097 | 6156902.136 | 4961756.454 | 3606665.57 | 3259820.611 | 3616525.336 | 3458315.675 |
| N-Palmitoyltaurine | 6925775.792 | 8064446.509 | 6285695.536 | 8143599.395 | 18707399.04 | 11866620.01 | 15766700.23 | 8615064.148 | 14749441 | 12502925.43 | 7185126.338 | 8337894.406 |
| 5,10-Dihydro-10-phenophosphazinol 10-oxide | 635993.3056 | 1319707.096 | 1118593.714 | 591380.7697 | 3438222.069 | 2541228.165 | 2679378.98 | 2861420.907 | 2787555.669 | 1754840.618 | 2065031.258 | 1918625.101 |
| Quinapril | 0 | 0 | 0 | 0 | 400195.6783 | 158431.2679 | 481569.573 | 351917.8497 | 492489.416 | 669006.7964 | 611828.2622 | 324313.6895 |
| lesquerolate | 3616372.084 | 3902355.938 | 4659458.512 | 4509560.572 | 7444073.207 | 5622002.651 | 9396682.941 | 6306697.368 | 4966555.805 | 6943900.04 | 7057620.253 | 3653483.437 |
| Butacaine | 3109655.743 | 3529006.995 | 1391778.431 | 2659461.021 | 4414488.735 | 2153918.514 | 5556501.403 | 4131831.421 | 3072120.464 | 5018904.825 | 2798560.676 | 2394529.717 |
| N6-Methyllysine | 1063090.78 | 919168.8594 | 1549232.433 | 1127528.285 | 1912968.294 | 1341157.993 | 2133233.72 | 1316189.263 | 16302503.05 | 1092026.285 | 1722608.185 | 1306836.054 |
| 3-Methylglutaric acid | 6059716.555 | 5622443.16 | 7470447.804 | 7074297.591 | 15401839.23 | 11852446.76 | 13392913.3 | 13713585.69 | 9329621.416 | 7294356.507 | 10060825.76 | 9384663.594 |
| alpha-CEHC (alpha-Carboxyethy | 24476299.69 | 22530157.47 | 31844382.88 | 26265732.88 | 44539316.71 | 40860314.28 | 38855818.23 | 37556452.65 | 32160558.42 | 24120798.78 | 29532793.39 | 38640895.95 |

| Metabolites | PBS_1 | PBS_2 | PBS_3 | PBS_4 | BCG_1 | BCG_2 | BCG_3 | BCG_4 | BCGΔ *pe18*_1 | BCGΔ *pe18*_2 | BCGΔ *pe18*_3 | BCGΔ*pe18*_4 |
|---|---|---|---|---|---|---|---|---|---|---|---|---|
| l hydrochroman) | | | | | | | | | | | | |
| Dodecanedioic acid | 1159006.965 | 1342094.957 | 1593992.921 | 1459525.645 | 1221781.159 | 1156502.897 | 1356850.889 | 1055400.249 | 1209478.898 | 1380992.547 | 1302013.187 | 2155096.014 |
| 2-{4-[2-(Trifluoromethyl)benzoyl]-1-piperazinyl}-1,3-thiazole-5-carboxamide | 1849840.045 | 1798657.3 | 1942110.376 | 1828136.007 | 1822065.47 | 1848800.34 | 1841064.542 | 1727317.517 | 1841072.796 | 1839011.082 | 1830446.961 | 1866936.976 |
| 7,7-Dimethyl-(5Z,8Z)-eicosadienoic acid | 669205.7371 | 802114.7428 | 961212.5693 | 953761.2522 | 10473584.24 | 15745523.34 | 10306693.19 | 12800091.75 | 8273818.783 | 7655039.159 | 10060097.63 | 7230386.123 |
| Hydroxyphenyllactic acid | 1032744.871 | 887459.9951 | 1457493.588 | 1151745.929 | 2548380.486 | 1465274.139 | 1985694.984 | 2409755.049 | 1879409.836 | 2025218.037 | 2371922.585 | 1705937.707 |
| Oxprenolol | 4964860.849 | 2367511.807 | 2250607.534 | 2351629.682 | 2369950.323 | 2364084.835 | 3875015.462 | 2366018.42 | 2377130.56 | 2376749.595 | 2343826.21 | 2603508.853 |
| Methylsuccinic acid | 11174977.38 | 8577077.431 | 8631044.876 | 11232337.04 | 68555567.43 | 41996145.02 | 56586076.01 | 53208723.67 | 40721465.99 | 31478998.08 | 43123974.9 | 32728210.57 |
| PC(20:3(8Z,11Z,14Z)/15:0) | 12127863.8 | 14034252.26 | 16000832.99 | 11242957.11 | 48088197.69 | 36863912.09 | 50402363.92 | 46152976.67 | 53989603.71 | 57435431.01 | 44840536.4 | 28339190.04 |
| 2-amino-4-pentenoate | 8848228.013 | 7959511.791 | 840931.111 | 7992129.586 | 1550075.67 | 10975847.28 | 11739890.37 | 11439010.46 | 5337330.613 | 11242946.65 | 4705708.938 | 1186784.11 |
| Oleic acid | 69070676.43 | 65400494.57 | 77988791.82 | 67750970.46 | 930195874.1 | 1626449473 | 992781726 | 1056128272 | 661239512.6 | 437449743.1 | 992267020.8 | 829030185.7 |
| 3,8'-Biapigenin | 0 | 0 | 20406.8 | 0 | 399617 | 392123 | 404095 | 394678 | 393847 | 345371 | 378463 | 385449 |

| **Metabolites** | **PBS_1** | **PBS_2** | **PBS_3** | **PBS_4** | **BCG_1** | **BCG_2** | **BCG_3** | **BCG_4** | **BCGΔ*pe18*_1** | **BCGΔ*pe18*_2** | **BCGΔ*pe18*_3** | **BCGΔ*pe18*_4** |
|---|---|---|---|---|---|---|---|---|---|---|---|---|
| | | | 0013 | | 0.202 | 9.535 | 3.386 | 4.932 | 8.587 | 7.421 | 6.309 | 5.514 |
| Methyl 5-hydroxypyridine-2-carboxylate | 8286103.245 | 7537939.119 | 5109249.824 | 5580614.551 | 8401624.549 | 9944771.996 | 8007085.594 | 8993085.918 | 8648740.475 | 7578760.194 | 8675553.133 | 4468908.64 |
| Karanjin | 2606208.41 | 2081443.698 | 4369891.862 | 2893770.948 | 5031726.758 | 4534570.493 | 4890205.779 | 4624701.736 | 4769634.779 | 2897172.536 | 3059573.691 | 4169497.686 |
| N,2-Dimethyl-N-phenylbutanamide | 665831.8058 | 678688.7141 | 477787.2144 | 5634935.289 | 121255.0329 | 795544.9415 | 9211131.666 | 119705.9085 | 3501524.523 | 2975379.096 | 199121.1227 | 6791561.939 |
| FAHFA 22:6/5:0 | 269149.0362 | 290598.8265 | 305763.5096 | 280198.7314 | 1004201.171 | 919908.1772 | 843963.627 | 1001846.985 | 980046.2453 | 655624.4524 | 678648.5457 | 831454.7792 |
| Erythraddison II | 401954.4414 | 475532.6894 | 578909.1336 | 485999.3626 | 948902.5713 | 968160.2394 | 703604.6585 | 833911.6785 | 888594.0509 | 625355.581 | 531010.1063 | 816301.1308 |
| Uracil | 24644788.54 | 18945647.06 | 39763836.72 | 24954535.51 | 48506060.23 | 45863990.69 | 57505348.96 | 49452137.03 | 36538807.04 | 28675633.95 | 30975931 | 31771614.05 |
| Jasmonic acid | 1134088.504 | 1176655.054 | 1013463.269 | 1192019.087 | 1059733.065 | 1144968.533 | 1203813.006 | 1134023.563 | 1108238.79 | 1152904.403 | 1085769.232 | 1039983.498 |
| Tolmetin | 28917035.63 | 8594644.533 | 14709880.41 | 8205771.365 | 42449671.32 | 20531932.77 | 31707955.81 | 27664044.87 | 27166149.41 | 23663456.32 | 25925955.44 | 18824424.53 |
| 1,4'-Bipiperidine | 4665702.916 | 1320454.089 | 1335690.808 | 4746899.032 | 3696881.596 | 5497698.333 | 2818931.733 | 1665677.298 | 5109633.693 | 4792301.969 | 3262675.503 | 3542568.267 |
| 2-Propyl-1-indanone | 860376.7032 | 703920.8076 | 707642.6577 | 6787826.614 | 240216.5384 | 1270629.113 | 10687801.6 | 229643.3497 | 3814989.833 | 2064930.028 | 421681.9234 | 8221922.323 |
| Heptadecanoyl_carnitine | 2782634.581 | 2211489.828 | 2560297.104 | 2538332.112 | 8665395.686 | 6023101.889 | 8576252.18 | 7924723.27 | 5017380.57 | 5729484.279 | 5479222.129 | 3211480.804 |

| **Metabolites** | **PBS_1** | **PBS_2** | **PBS_3** | **PBS_4** | **BCG_1** | **BCG_2** | **BCG_3** | **BCG_4** | **BCGΔ *pe18*_1** | **BCGΔ *pe18*_2** | **BCGΔ *pe18*_3** | **BCGΔ*pe18*_4** |
|---|---|---|---|---|---|---|---|---|---|---|---|---|
| N-{5-[(2-Amino-1,3-thiazol-4-yl)methyl]-1,3,4-thiadiazol-2-yl}-N-methylamine | 472095.6072 | 354480.0284 | 486782.6917 | 307220.8351 | 1984294.644 | 995298.9314 | 1851040.628 | 1493649.616 | 1264876.303 | 1421234.073 | 1748876.171 | 874456.8993 |
| trans-Vaccenic acid | 69070676.43 | 65400494.57 | 77988791.82 | 67750970.46 | 930195874.1 | 1626449473 | 992781726 | 1056128272 | 661239512.6 | 437449743.1 | 992267020.8 | 829030185.7 |
| PE 8:0_26:0 | 24195726.48 | 12436124.6 | 32051625.49 | 25711894.19 | 21636128.59 | 1535202.751 | 26828847.2 | 4219228.929 | 1871109.596 | 18912032.95 | 16852889.81 | 1752872.724 |
| Chaetoglobosin A | 198894.3568 | 228388.5125 | 302154.1453 | 141373.0414 | 6769330.679 | 5711774.576 | 6227841.08 | 7463446.311 | 5619235.648 | 4533640.702 | 7231747.721 | 5150419.776 |
| DGGA 16:4_24:4 | 1247882.52 | 1036972.009 | 434579.8066 | 888305.1581 | 350517.6239 | 53160.60033 | 179235.5716 | 77808.4093 | 404280.7531 | 192703.1615 | 164987.0838 | 239260.8122 |
| Cellobiose | 0 | 0 | 0 | 0 | 1866199.671 | 1306014.031 | 1055738.505 | 1086160.176 | 1597887.449 | 1170675.953 | 1212277.35 | 1581540.231 |
| LPE 17:1 | 140063.8265 | 116746.2464 | 125189.8912 | 181255.138 | 1020682.1 | 1026515.502 | 960279.7581 | 840417.4608 | 944333.5142 | 887160.1636 | 979170.0372 | 823524.9501 |
| His-Arg | 1154950.29 | 1193129.079 | 1134851.624 | 1145164.49 | 1148567.86 | 1175598.12 | 1121430.107 | 1279738.769 | 1137216.752 | 1181050.636 | 1214328.288 | 1221465.768 |
| PS 16:0_22:6 | 1190654.122 | 1039275.568 | 918906.2321 | 465341.5414 | 2010476.855 | 3017672.874 | 2520771.556 | 2200299.566 | 2132452.422 | 2028258.637 | 2301393.198 | 2125556.23 |
| Serine | 988876.2655 | 1109781.917 | 878764.4873 | 1274106.636 | 3944859.397 | 1692836.649 | 2904180.67 | 2452780.423 | 2871234.898 | 2161910.452 | 2104975.4 | 2272678.236 |
| .beta.-Hydroxyphen | 373240 | 369882 | 472056 | 387210 | 633652 | 505696 | 627074 | 537785 | 601445 | 442386 | 422199 | 529033 |

| Metabolites | PBS_1 | PBS_2 | PBS_3 | PBS_4 | BCG_1 | BCG_2 | BCG_3 | BCG_4 | BCGΔ*pe18*_1 | BCGΔ*pe18*_2 | BCGΔ*pe18*_3 | BCGΔ*pe18*_4 |
|---|---|---|---|---|---|---|---|---|---|---|---|---|
| ylalanine | 8.694 | 2.802 | 4.533 | 1.93 | 8.812 | 1.012 | 7.21 | 0.931 | 7.246 | 6.833 | 0.119 | 8.784 |
| Parabanic acid | 2378927.373 | 8743410.588 | 8161724.808 | 8493885.209 | 1867073.377 | 1398496.838 | 2571859.145 | 1458756.881 | 2095652.295 | 1985661.527 | 2171657.168 | 6957282.009 |
| Tetradecanedioic acid | 1365639.548 | 1899429.328 | 1928078.72 | 1899883.84 | 3480237.712 | 5863425.964 | 3903400.216 | 3677533.016 | 2377082.803 | 1737166.82 | 2801185.01 | 5039324.783 |
| 4-Amino-2-(1,3-benzoxazol-2-yl)phenol | 524184.0726 | 432097.5351 | 637967.9618 | 447533.8609 | 706994.8053 | 528027.3139 | 518029.5275 | 581434.6318 | 533038.8379 | 496368.0554 | 466604.2231 | 391991.2494 |
| Ouabagenin | 1308556.529 | 1233939.468 | 1129857.811 | 1047358.523 | 1700277.839 | 1152162.879 | 1203768.369 | 1911826.859 | 1204842.288 | 1185193.537 | 1100685.118 | 1094725.775 |
| IS4 | 27570884.45 | 28019006.82 | 33090963.48 | 28258783.1 | 40602437.23 | 42090779.32 | 39772575.01 | 37902416.49 | 37411127.85 | 33971783.47 | 36309557.33 | 39505231.13 |
| PE O-26:6_15:0 | 708351.1467 | 370489.0737 | 777167.2961 | 723700.6867 | 115474.2663 | 266912.3967 | 746257.2906 | 2228515.281 | 349854.2164 | 40416.81744 | 489795.2023 | 1356345.008 |
| LPC 17:1 | 506672.9355 | 418316.5509 | 709441.644 | 554745.4249 | 2071400.411 | 1918682.942 | 2018895.553 | 1841103.858 | 1789467.241 | 1814277.357 | 2004864.947 | 1538892.548 |
| ST 24:1;O4;T/14:1 | 8029691.517 | 10955903.49 | 8374199.165 | 8695671.73 | 8723151.937 | 9685612.861 | 6716092.972 | 5937854.437 | 8370349.226 | 8836728.402 | 6322229.569 | 7800545.071 |
| LNAPE 3:0/N-22:3 | 5190958.54 | 7574842.756 | 1468601.341 | 3985976.636 | 797955.4428 | 1034541.944 | 899481.8254 | 984940.2047 | 870918.7215 | 991555.4224 | 1128819.558 | 772491.8015 |
| Terephthalic-Acid | 23576126.15 | 11819149.18 | 12904513.25 | 12028889.27 | 14698960.97 | 11251662.81 | 26462272.64 | 15641109.25 | 11634731.06 | 8874172.185 | 14620900.95 | 9039246.11 |
| 4-Guanidinobutanal | 10851319.29 | 8152971.522 | 10415503.86 | 14973246.06 | 38590292.94 | 35290301.66 | 17879245.17 | 29263185.54 | 30246466.46 | 8667986.041 | 12604648.52 | 33295029.28 |

| **Metabolites** | **PBS_1** | **PBS_2** | **PBS_3** | **PBS_4** | **BCG_1** | **BCG_2** | **BCG_3** | **BCG_4** | **BCGΔ*pe18*_1** | **BCGΔ*pe18*_2** | **BCGΔ*pe18*_3** | **BCGΔ*pe18*_4** |
|---|---|---|---|---|---|---|---|---|---|---|---|---|
| cis-4-Hydroxycyclohexanecarboxylic acid | 3268597.794 | 2655046.859 | 3059060.194 | 5885550.569 | 2454214.551 | 5283153.724 | 1956635.895 | 3006988.436 | 5009750.446 | 3647965.747 | 2621282.394 | 2340325.987 |
| Dihydroflavopereirine | 2268710.118 | 2262082.753 | 1968374.208 | 2079605.126 | 2171646.949 | 2119863.385 | 2192304.246 | 2287007.356 | 2263836.952 | 2145953.895 | 2172670.071 | 2072604.737 |
| PC 13:0_13:0 | 22709.1645 | 10624.01798 | 0 | 24727.06327 | 3240267.504 | 2415617.255 | 3494279.115 | 2616078.033 | 3515774.518 | 4030937.15 | 3018032.396 | 1862196.737 |
| PE-Cer 12:1;2O/17:2 | 508063.7538 | 370528.8517 | 524843.8033 | 508158.7895 | 319908.2236 | 584051.7907 | 415940.5368 | 508351.0847 | 320423.9465 | 225091.7821 | 124796.2969 | 983896.3259 |
| Trabectedin | 2055207.411 | 755952.5397 | 5030763.681 | 1068551.804 | 21919774.08 | 8284368.503 | 13527655.58 | 13891305.84 | 11963438.23 | 8272307.621 | 10588039.46 | 6087590.339 |
| Oleoylcarnitine (Car(18:1)) | 32015359.63 | 23606256.08 | 41490628.74 | 29645334.02 | 64271694.4 | 43276879.24 | 66974154.12 | 58065387.34 | 33548167.78 | 45575893.48 | 45189903.09 | 24762215.47 |
| CerP 14:1;2O/10:0 | 489956.548 | 513524.9543 | 418508.5038 | 480973.1621 | 506910.2265 | 491603.1492 | 579324.9722 | 546477.3221 | 559666.8096 | 553818.9505 | 522448.238 | 460828.5871 |
| Methyl 2-oxocycloheptane-1-carboxylate | 2326076.528 | 2469663.788 | 2436812.75 | 5940540.971 | 1654629.974 | 3088569.254 | 7061747.572 | 1830546.194 | 5404985.854 | 3984791.215 | 1704316.955 | 7663723.112 |
| LPE O-20:3 | 1344637.587 | 1577687.074 | 1178915.341 | 1137321.153 | 918880.6997 | 1353052.32 | 933289.3711 | 1050856.479 | 1186408.213 | 981584.3506 | 1318771.439 | 634755.9002 |
| 4-(9H-Fluoren-2-yl)-1,3-thiazol-2-amine | 3913941.638 | 3170987.371 | 7271167.333 | 4438188.487 | 5970897.365 | 5518878.957 | 6567613.423 | 5495888.09 | 5189667.67 | 4349599.433 | 4829306.128 | 4499451.738 |
| L-Serine methyl | 187452 | 147692 | 268316 | 168746 | 136802 | 157352 | 135703 | 150717 | 192021 | 121733 | 167612 | 149469 |

| Metabolites | PBS_1 | PBS_2 | PBS_3 | PBS_4 | BCG_1 | BCG_2 | BCG_3 | BCG_4 | BCGΔ*pe18*_1 | BCGΔ*pe18*_2 | BCGΔ*pe18*_3 | BCGΔ*pe18*_4 |
|---|---|---|---|---|---|---|---|---|---|---|---|---|
| ester | 2.704 | 1.23 | 7.171 | 4.217 | 9.103 | 8.966 | 7.764 | 8.797 | 3.917 | 8.257 | 4.733 | 5.586 |
| Gentisaldehyde | 1083399.382 | 1004728.392 | 742798.9651 | 768438.1867 | 1132445.958 | 1059289.28 | 984675.9136 | 1139198.492 | 1146877.544 | 980785.3089 | 986963.1195 | 1122554.927 |
| Dephospho-CoA | 28185.17973 | 0 | 29578.50872 | 8816.955686 | 126845.5226 | 240813.5011 | 385537.9712 | 421691.1036 | 440269.1264 | 395261.9489 | 423065.8762 | 5245.988692 |
| 3-Succinoylpyridine | 8694211.055 | 7015990.241 | 12491346.19 | 10337166.31 | 12215982 | 12227509.25 | 12065679.5 | 11615946.26 | 10683229 | 6334765.789 | 7500149.006 | 9991033.155 |
| SL 10:0;O/16:0 | 22868.51844 | 80724.0965 | 49074.8218 | 86780.09895 | 48746.69566 | 24665.34828 | 80369.85188 | 39068.94301 | 64203.38366 | 39805.03694 | 48598.4463 | 28478.06806 |
| 7,8-Dihydrobiopterin | 118574.9906 | 118278.9929 | 197721.3485 | 88712.7876 | 2930268.169 | 1626527.512 | 4677795.891 | 3284019.925 | 2238379.885 | 1911784.229 | 3054562.478 | 1592387.918 |
| Cer 8:1;2O/5:0 | 21450811.37 | 14609208.81 | 13206510.02 | 20139513.48 | 8655657.579 | 11584520.71 | 12577845.79 | 8347919.664 | 10352576.06 | 9617263.452 | 8912284.964 | 6732744.543 |
| 8-Hydroxy-3-(7-hydroxyheptyl)-6-oxo-3,4-dihydroisochromene-7-carboxylic acid | 3779239.188 | 2968750.12 | 3103282.954 | 2454582.647 | 3675496.427 | 2867217.093 | 3321702.777 | 2563588.492 | 3388730.201 | 3024016.782 | 3265485.222 | 3025347.747 |
| Psychotrine | 1761040.653 | 1174324.781 | 3375922.914 | 3434235.537 | 1886522.637 | 1252503.793 | 1984045.926 | 1846229.643 | 3244411.907 | 2784330.375 | 1064938.365 | 2706104.183 |
| PE O-16:2_5:0 | 1201309.256 | 1266580.532 | 1208829.63 | 1191871.236 | 1217100.918 | 1199322.154 | 1212257.563 | 866863.1323 | 1328363.468 | 1149893.543 | 1265733.063 | 1181346.734 |
| 4-Amino-2-isopropyl-5-methylphenol | 1747442.127 | 2112138.911 | 1261547.241 | 1858489.751 | 1944529.849 | 1569913.052 | 2000700.008 | 1944857.499 | 2049946.557 | 1815851.077 | 1863802.693 | 1725930.422 |

| **Metabolites** | **PBS_1** | **PBS_2** | **PBS_3** | **PBS_4** | **BCG_1** | **BCG_2** | **BCG_3** | **BCG_4** | **BCGΔ *pe18*_1** | **BCGΔ *pe18*_2** | **BCGΔ *pe18*_3** | **BCGΔ*pe18*_4** |
|---|---|---|---|---|---|---|---|---|---|---|---|---|
| 1-alpha-linolenoyl-sn-glycero-3-phospho-(1'-sn-glycerol) | 8893.13308 | 15306.57195 | 21491.38048 | 0 | 523033.8091 | 398975.7803 | 585709.7502 | 521907.6146 | 520467.7479 | 448952.8991 | 460243.4811 | 492346.7944 |
| Diethylcarbamazine | 2080010.179 | 7377347.605 | 1437371.458 | 4056139.064 | 16051508.37 | 8065719.582 | 3000596.745 | 1276616.971 | 2990952.771 | 4659577.496 | 8682023.746 | 5807439.612 |
| Cer 12:2;2O/7:0 | 8052313.189 | 6797019.72 | 4347271.334 | 6288424.481 | 4356487.21 | 4704826.547 | 5170928.361 | 4066368.96 | 6156595.124 | 5375080.168 | 4520299.443 | 4908275.267 |
| 2,3,4-Pentanetrione, 3-[(4-nitrophenyl)hydrazone] | 735203.5864 | 850269.9378 | 1210084.979 | 852151.2987 | 1539153.21 | 1164751.08 | 1695451.671 | 1527418.591 | 1258752.939 | 1002071.002 | 1171185.717 | 1167677.821 |
| LNAPS 3:0/N-21:0 | 705333.3415 | 992799.8849 | 607984.0666 | 634036.3352 | 661359.1143 | 960760.6207 | 658049.7216 | 524462.7551 | 476200.6034 | 503411.3254 | 710304.3474 | 368655.3839 |
| Citraconic acid | 11303185.32 | 6695110.517 | 32516990.67 | 9566902.76 | 5948506.895 | 3672718.865 | 7890060.771 | 9305293.487 | 5547407.488 | 7771549.209 | 5502483.325 | 8487360.545 |
| 3-((Ethoxy(hydroxy)phosphoryl)oxy)-2-hydroxypropyl (9Z,12Z)-octadeca-9,12-dienoate | 1466739.238 | 3840716.897 | 1896238.805 | 1691457.5 | 650773.2023 | 414098.1339 | 705300.6737 | 484657.4829 | 606768.885 | 713840.7295 | 823727.2484 | 296189.1026 |
| trans-2,5-Bis-(3,4,5-trimethoxyphenyl)-1,3-dioxolane | 529322.7134 | 639791.8932 | 662296.1372 | 904982.291 | 618366.5746 | 912160.0317 | 640250.279 | 836287.4145 | 758832.1839 | 979903.9772 | 559716.5148 | 835035.0302 |
| 2-(4-Chlorophenyl)-3H-imidazo[4,5-c] | 6320620.613 | 6512940.687 | 6737723.309 | 5974623.271 | 6905547.174 | 6100389.69 | 9183763.543 | 7264105.452 | 6102236.46 | 3586482.821 | 12051693.14 | 4373610.505 |

| Metabolites | PBS_1 | PBS_2 | PBS_3 | PBS_4 | BCG_1 | BCG_2 | BCG_3 | BCG_4 | BCGΔ *pe18*_1 | BCGΔ *pe18*_2 | BCGΔ *pe18*_3 | BCGΔ*pe18*_4 |
|---|---|---|---|---|---|---|---|---|---|---|---|---|
| pyridine | | | | | | | | | | | | |
| PE-Cer 13:1;2O/22:6 | 808023.8472 | 1138867.87 | 542686.6604 | 855837.5956 | 1216085.87 | 1213750.974 | 1104509.63 | 1437472.617 | 1347365.612 | 1432084.022 | 992076.5811 | 769318.7 |
| Methyl 2-{[(4-methoxyphenyl)carbonyl]amino}-5,6-dihydro-4H-cyclopenta[b]thiophene-3-carboxylate | 496445.0404 | 230698.1234 | 401108.9882 | 366545.345 | 2207169.772 | 1571118.639 | 1092198.185 | 1121685.806 | 1175725.279 | 1214340.407 | 2451812.245 | 987188.5723 |
| CAR 14:1 | 2231173.285 | 1833258.454 | 4122622.797 | 3512805.806 | 6554318.328 | 4121648.828 | 7513371.588 | 6182842.563 | 3153865.846 | 4831624.855 | 4531462.596 | 2788940.258 |
| FA 19:1 | 2198204.975 | 2061025.344 | 2167758.464 | 1883853.259 | 5401604.898 | 9737276.881 | 5138922.809 | 6663371.576 | 3794169.666 | 2805840.101 | 5870350.173 | 4168781.465 |
| 5-Hydroxy-2-(hydroxymethyl)pyridine | 1269562.783 | 1329308.469 | 1421773.199 | 1342171.635 | 1356473.579 | 1243742.769 | 1395203.75 | 1372558.961 | 1195977.293 | 1153813.052 | 1229437.956 | 1636307.417 |
| trans-2-Dodecenoylcarnitine | 8618143.645 | 11365789.98 | 3786617.283 | 6277087.16 | 14452075.03 | 2155650.874 | 16484450.83 | 7675720.078 | 8148410.405 | 19851832.03 | 3922808.535 | 4132551.143 |
| Butylamine | 38266098.55 | 61030287.52 | 35620583.09 | 67091516.28 | 64094703.67 | 37082188.18 | 37974449.13 | 37437691.29 | 38767115.22 | 37849526.2 | 35946576.08 | 35912256.12 |
| 2-Amino-6-ethyl-4(3H)-pyrimidinone | 1914443.171 | 1385604.198 | 4892991.222 | 1908431.554 | 3271471.965 | 2956821.487 | 7114446.853 | 3162290.298 | 2898564.6 | 5764611.599 | 5947688.135 | 2987433.127 |
| PC O-28:2 | 1998626.129 | 2025736.382 | 1865453.505 | 1980641.86 | 1993287.422 | 2088340.113 | 1988182.37 | 1876933.523 | 1874825.093 | 1955462.124 | 1875783.252 | 1994189.01 |

| Metabolites | PBS_1 | PBS_2 | PBS_3 | PBS_4 | BCG_1 | BCG_2 | BCG_3 | BCG_4 | BCGΔ*pe18*_1 | BCGΔ*pe18*_2 | BCGΔ*pe18*_3 | BCGΔ*pe18*_4 |
|---|---|---|---|---|---|---|---|---|---|---|---|---|
| 6-(Acetylamino)hexanoic acid | 1005292.378 | 1116930.877 | 1021346.56 | 1557230.185 | 761774.8519 | 831435.4938 | 1529187.749 | 744856.894 | 981445.9008 | 1255378.359 | 902242.9141 | 2046864.774 |
| Erucic acid | 693089.8365 | 499510.7587 | 1246798.912 | 691441.9963 | 80047309.68 | 118271944.9 | 79262533.32 | 91284263.98 | 50671990.36 | 37640863.11 | 82666189.39 | 71782870.02 |
| Methanone, [1-(4-penten-1-yl)-1H-indol-3-yl](2,2,3,3-tetramethylcyclopropyl)- | 29272570.01 | 41374598.65 | 26405069.76 | 29980273.34 | 34226920.92 | 30050765.76 | 29618640.78 | 32543984.83 | 31490774.51 | 36076943.07 | 29064886.51 | 26019450.85 |
| MGDG O-20:2_18:0 | 10610983.03 | 7736663.632 | 7202598.963 | 10219700.48 | 12451320.61 | 10156314.3 | 7184037.352 | 19579535.78 | 11722223.11 | 9301105.046 | 1990756.137 | 20506380.44 |
| CDP-Ethanolamine | 217308.0943 | 273941.1354 | 385878.087 | 284714.2823 | 787220.1209 | 392089.2578 | 765801.6211 | 659698.4231 | 623471.6784 | 720493.3256 | 619235.0078 | 610526.3917 |
| Imatinib | 625944.5369 | 663767.7227 | 560570.9228 | 640476.9758 | 672907.9127 | 610916.4229 | 699120.1263 | 580775.9063 | 572280.6328 | 653655.6791 | 533768.3369 | 549315.6169 |
| HexCer 14:3;2O/5:0 | 1287753.585 | 1725604.195 | 1219114.647 | 1896190.636 | 1747984.408 | 1434265.681 | 1855967.63 | 1702364.078 | 1310621.318 | 1749766.166 | 1222966.673 | 1748107.255 |
| DL-Tryptophan, methyl ester | 910825.0656 | 780698.6236 | 1149717.499 | 882870.9239 | 2945832.376 | 2107644.341 | 3227169.66 | 2584083.965 | 2256033.764 | 2425974.912 | 2439912.993 | 1085209.131 |
| PS 18:1_20:4 | 2608408.375 | 1787470.627 | 1916881.553 | 2590494.449 | 2316045.785 | 106884.5525 | 2176069.44 | 1436679.66 | 2257416.274 | 2138352.29 | 2486146.055 | 1108587.144 |
| Monomethyl fumarate | 11303185.32 | 6695110.517 | 32516990.67 | 9566902.76 | 5948506.895 | 3672718.865 | 7890060.771 | 9305293.487 | 5547407.488 | 7771549.209 | 5502483.325 | 8487360.545 |
| DGTS 14:1_14:1 | 426325. | 130717 | 955798. | 111593 | 161114 | 118177 | 176692 | 655234. | 481042. | 793934. | 162146 | 119160 |

| Metabolites | PBS_1 | PBS_2 | PBS_3 | PBS_4 | BCG_1 | BCG_2 | BCG_3 | BCG_4 | BCGΔ*pe18*_1 | BCGΔ*pe18*_2 | BCGΔ*pe18*_3 | BCGΔ*pe18*_4 |
|---|---|---|---|---|---|---|---|---|---|---|---|---|
| | 2871 | 7.746 | 6742 | 3.148 | 8.992 | 8.164 | 7.045 | 8321 | 1261 | 3826 | 8.029 | 9.816 |
| Mifepristone | 864434.5942 | 985371.1793 | 736941.6069 | 795136.9132 | 876374.4263 | 812188.4743 | 751937.5653 | 1088100.895 | 586696.1758 | 912503.611 | 746965.7684 | 706477.4691 |
| 2-Methoxy-N-(1H-tetraazol-5-yl)benzamide | 208051.9494 | 150626.5403 | 503641.9625 | 199550.0836 | 2828883.278 | 1489279.845 | 3351564.072 | 1758443.581 | 2257563.32 | 2506874.455 | 2948258.398 | 1453571.076 |
| 1-(1Z-Octadecenyl)-sn-glycero-3-phosphocholine | 7392346.951 | 8705198.666 | 4188207.977 | 7303303.323 | 7054008.483 | 8171037.529 | 6006629.677 | 6077335.129 | 5748459.37 | 5007944.634 | 5130072.667 | 3699211.8 |
| MPL-dm | 4053910.607 | 3941779.536 | 3304057.456 | 4972966.79 | 3180938.168 | 3255938.278 | 4570051.722 | 3011636.871 | 3929560.724 | 3484648.232 | 2785140.639 | 4387367.751 |
| 3-O-Sulfogalactosylceramide (d18:1/22:0) | 1367142.372 | 2007761.01 | 1351218.918 | 1469993.05 | 1580148.745 | 1412027.775 | 1459966.178 | 1393662.073 | 1270099.705 | 1496015.066 | 1403845.995 | 1559977.604 |
| SMGDG O-9:0_26:6 | 498679.8464 | 496893.7832 | 448893.6166 | 471226.9405 | 514503.982 | 489067.8593 | 453506.9001 | 486848.5449 | 491707.2974 | 442535.7019 | 503586.2313 | 434905.9446 |
| N-3,4-Tridhydroxybenzamide | 944075.8617 | 911698.1833 | 726074.2532 | 943757.6761 | 922382.2934 | 887962.9871 | 894648.1134 | 626371.5129 | 943272.1894 | 923233.8109 | 914537.6044 | 922423.6467 |
| 6,7-Diethoxy-4-methylcoumarin | 2252853.52 | 2302490.71 | 2262577.905 | 2390735.965 | 2329765.102 | 2080923.034 | 2413262.62 | 2245802.672 | 1942080.671 | 2414117.885 | 1996832.556 | 2072603.15 |
| Nordihydrocapsaicin | 1656283.917 | 1839078.963 | 1480983.58 | 1705285.549 | 1661337.639 | 1630881.423 | 1806342.428 | 1754486.431 | 1647852.072 | 1646213.675 | 1674454.761 | 1591906.592 |
| Warfarin | 1081684.45 | 1530995.295 | 1356687.366 | 1578465.489 | 1072691.57 | 1917580.01 | 1357099.351 | 916357.1124 | 1363100.167 | 901580.4879 | 1565524.525 | 1258944.719 |

| **Metabolites** | **PBS_1** | **PBS_2** | **PBS_3** | **PBS_4** | **BCG_1** | **BCG_2** | **BCG_3** | **BCG_4** | **BCGΔ*pe18*_1** | **BCGΔ*pe18*_2** | **BCGΔ*pe18*_3** | **BCGΔ*pe18*_4** |
|---|---|---|---|---|---|---|---|---|---|---|---|---|
| Pachyrrhizin | 2730636.659 | 2279383.98 | 3892869.719 | 2668265.471 | 2688200.257 | 2580175.078 | 3719036.102 | 2700743.342 | 3223027.173 | 2452582.541 | 2407057.083 | 2375496.482 |
| LDGTS 12:0 | 1179690.163 | 1215114.131 | 1130707.512 | 1119232.452 | 1130816.025 | 1078279.716 | 1244460.514 | 1132659.246 | 1231509.791 | 1220235.438 | 1201874.027 | 1064044.473 |
| Choline | 331582112.1 | 307182214 | 369097734.1 | 317783441.9 | 326583143 | 235149079.4 | 343580237.4 | 288105445.1 | 237456001.5 | 270719488.5 | 219443654.1 | 233159391.8 |
| N-arachidonoyl-phenylalanine | 14714609.18 | 15510761.5 | 5561135.641 | 7508728.045 | 5955454.295 | 5259394.53 | 5191628.727 | 5645957.224 | 5936826.269 | 3764230.664 | 3402267.273 | 2001254.736 |
| Taurine | 17878203.58 | 14545873.45 | 19763209.1 | 15124615.16 | 44497451.67 | 24352744.42 | 49269250.36 | 32880692.8 | 21995663.05 | 36302874.26 | 30048839.81 | 16690295.02 |
| 5-(9-Anthrylmethylene)-2,4,6(1H,3H,5H)-pyrimidinetrione | 251253.4528 | 229255.9961 | 440755.6518 | 317448.7746 | 5705181.765 | 3142051.124 | 5351324.226 | 4293633.655 | 4530319.757 | 2885711.123 | 3575769.542 | 3695263.43 |
| tetradecasphinganine | 8442479.962 | 9083202.046 | 9454517.182 | 8902836.386 | 8841344.037 | 9135964.685 | 8138745.243 | 8899344.798 | 8412641.745 | 7706295.384 | 8675663.075 | 9121641.018 |
| (2R)-3-Hydroxyisovaleroylcarnitine | 1485635.568 | 1157878.859 | 2345712.507 | 1746860.024 | 7862764.149 | 5137105.536 | 8842518.651 | 6499794.242 | 4333944.427 | 4147325.782 | 4852495.649 | 3689635.066 |
| 1H-Indazole-3-carboxamide, N-[1-(aminocarbonyl)-2,2-dimethylpropyl]-1-[(4-fluorophenyl)methyl]- | 200894.8242 | 176156.1904 | 137321.5678 | 115086.448 | 8964710.241 | 14775867.9 | 11415147.23 | 9925750.135 | 7031520.084 | 2881862.493 | 9762465.892 | 8748167.835 |
| 2-Methylvaleric | 112007 | 117539 | 105409 | 226235 | 128572 | 125968 | 305233 | 106634 | 182473 | 135042 | 110898 | 309140 |

| Metabolites | PBS_1 | PBS_2 | PBS_3 | PBS_4 | BCG_1 | BCG_2 | BCG_3 | BCG_4 | BCGΔ *pe18*_1 | BCGΔ *pe18*_2 | BCGΔ *pe18*_3 | BCGΔ*pe18*_4 |
|---|---|---|---|---|---|---|---|---|---|---|---|---|
| acid | 98.14 | 45.74 | 95.71 | 80.32 | 59.35 | 40.34 | 89.54 | 12.18 | 75.38 | 66.64 | 52.81 | 08.55 |
| LPS 16:1 | 1888777.585 | 1377437.883 | 2526583.736 | 1830575.754 | 4452361.808 | 3561694.718 | 4489718.754 | 3622762.056 | 4448499.19 | 4163329.938 | 5350336.608 | 3708428.669 |
| N-alpha-methylhistamine | 19274902.58 | 19567973.15 | 20950919.43 | 19718805.28 | 19201202.18 | 19776166.8 | 19354566 | 18992442.53 | 19252403.1 | 18310753.73 | 19575694.99 | 18869858.43 |
| 5-Aminolevulinic acid hexyl ester | 4810196.721 | 4977101.186 | 4160460.273 | 5953532.902 | 2533840.717 | 4037273.731 | 4651216.921 | 3291795.175 | 4278373.784 | 3703172.404 | 3149767.388 | 3422014.897 |
| 3-Hydroxylauric acid | 3512170.269 | 2948882.468 | 3288217.157 | 2938567.705 | 3749900.625 | 2504992.088 | 2405407.758 | 2449885.611 | 2847385.06 | 3199369.277 | 4355722.098 | 2860752.929 |
| LPE O-18:1 | 24585696.09 | 30430438.29 | 14079638.35 | 17237444.8 | 8286103.451 | 10450410.09 | 10817779.64 | 8991986.523 | 6928120.127 | 8169344.479 | 9009888.001 | 4835062.666 |
| 6-Bromo-2-phenyl-4-quinolinecarboxylic acid | 2020731.882 | 1544103.875 | 3546753.888 | 2431354.001 | 5650563.833 | 5794001.682 | 5123900.015 | 4992146.697 | 4939601.827 | 3913006.784 | 4473158.326 | 4529397.923 |
| Isoxsuprine hydrochloride | 1526749.016 | 1444038.061 | 1386065.833 | 1292499.421 | 1204929.25 | 1492175.07 | 1555918.572 | 1539097.184 | 1282145.975 | 1668369.815 | 1290250.984 | 1453954.485 |
| cis-3-Hexenyl tiglate | 1200674.116 | 1212618.582 | 1003524.125 | 1407000.768 | 990668.1963 | 1046986.714 | 1439304.805 | 1121484.761 | 1112555.21 | 1074047.293 | 1147670.049 | 1484534.238 |
| 9-Octadecenoic acid , (2-hydroxy-2-oxido-1,2-oxaphospholan-4-yl)methyl ester | 91391.96809 | 227589.9844 | 125206.3815 | 330826.9173 | 1230306.245 | 892857.0846 | 334050.3774 | 429687.7519 | 569475.8457 | 803305.3767 | 957159.9318 | 138937.1993 |
| Ethephon | 106066 | 103486 | 151441 | 106983 | 129011 | 129180 | 115269 | 117177 | 110849 | 121210 | 108911 | 119825 |

| Metabolites | PBS_1 | PBS_2 | PBS_3 | PBS_4 | BCG_1 | BCG_2 | BCG_3 | BCG_4 | BCGΔ*pe18*_1 | BCGΔ*pe18*_2 | BCGΔ*pe18*_3 | BCGΔ*pe18*_4 |
|---|---|---|---|---|---|---|---|---|---|---|---|---|
| | 3.042 | 2.336 | 2.151 | 5.062 | 2.529 | 6.793 | 7.861 | 3.994 | 0.418 | 7.045 | 9.949 | 8.962 |
| Cer 9:0;3O/16:2;(2OH) | 0 | 9030.061562 | 0 | 0 | 184245.2682 | 180371.3308 | 132474.6457 | 112964.4493 | 345974.2265 | 384850.5794 | 350996.0265 | 278685.6857 |
| Heptanedioic acid, 1-(2-cyclopentylidenehydrazide) | 2628427.935 | 2811492.142 | 3933232.606 | 2589843.138 | 2597705.379 | 1717638.928 | 2795137.166 | 2102302.511 | 1544360.244 | 2019986.588 | 2046257.521 | 1093217.561 |
| PG 17:1_22:6 | 2136021.817 | 2444276.508 | 646513.3 | 1644085.585 | 8258996.561 | 5647725.348 | 7042818.918 | 7408546.53 | 11020593.27 | 8702510.726 | 4992484.171 | 6274805.239 |
| PC(35:5) | 44772920.09 | 47079402.57 | 36889781.54 | 40651606.67 | 63807590.55 | 52020768.32 | 58006432.45 | 73596367.83 | 58038493.29 | 62886515.68 | 47663401.86 | 47679568.16 |
| Leucine methyl ester | 2937473.864 | 2251712.422 | 4079977.049 | 2867053.521 | 12411545.31 | 10135135.41 | 11051132.02 | 9606303.959 | 8935507.058 | 8875742.649 | 11573693.73 | 10700562.03 |
| Pseudouridine | 469222.3014 | 369566.3695 | 685032.9555 | 444421.4569 | 2212430.869 | 1331381.341 | 2041874.48 | 1616085.177 | 1567032.166 | 1464811.043 | 1432719.655 | 1247977.76 |
| PC(14:0/14:1(9Z)) | 251759.9191 | 168426.0973 | 36980.53523 | 165772.0846 | 9060207.635 | 5946297.533 | 9383885.792 | 7512046.278 | 8855645.473 | 9225095.452 | 7310768.912 | 4102497.157 |
| prolyl-isoleucine | 23736906.49 | 18575836.2 | 30965448.34 | 21948924.82 | 21776003.88 | 13821872.31 | 24314148.1 | 17158497.67 | 11864345.75 | 15401865.79 | 16300022.46 | 9903216.889 |
| Sarcosine | 921091.933 | 1726621.24 | 1850578.28 | 849914.3759 | 9985984.966 | 3622395.064 | 9873332.789 | 5661336.361 | 4246504.242 | 5472696.098 | 5088721.626 | 4503867.145 |
| 1-(4-Bromophenyl)-2-(4-morpholinyl)ethanol | 623047.758 | 422694.5617 | 760977.2613 | 473300.8921 | 918454.2542 | 649013.1888 | 658054.7727 | 708707.3733 | 655240.7061 | 539616.4397 | 592846.1662 | 450426.6514 |
| PC O-13:0_2:0 | 268763 | 206081 | 218248 | 216839 | 443914 | 426386 | 419542 | 354279 | 374466 | 373979 | 449986 | 315660 |

| **Metabolites** | **PBS_1** | **PBS_2** | **PBS_3** | **PBS_4** | **BCG_1** | **BCG_2** | **BCG_3** | **BCG_4** | **BCGΔ*pe18*_1** | **BCGΔ*pe18*_2** | **BCGΔ*pe18*_3** | **BCGΔ*pe18*_4** |
|---|---|---|---|---|---|---|---|---|---|---|---|---|
| | 0.011 | 8.912 | 3.121 | 9.574 | 4.976 | 4.756 | 5.31 | 4.589 | 2.144 | 3.71 | 0.116 | 6.048 |
| Phosphorylcholine | 149414274 | 120661081.1 | 179311113.3 | 142045331.1 | 763727400.5 | 355152687.1 | 662902244.2 | 482282379.8 | 532542279.8 | 579158781.2 | 601673849 | 365791846.6 |
| Dihydroethidium | 2317107.99 | 2419295.364 | 2252841.647 | 2344889.442 | 2322865.144 | 2308295.498 | 2322763.4 | 2323182.368 | 2327123.68 | 2776508.255 | 2384639.541 | 2284982.129 |
| 5-Hydroxyindole | 1288436.886 | 1227815.568 | 1152160.565 | 1286948.175 | 1223788.37 | 1417690.338 | 1014977.753 | 1264329.629 | 1818423.845 | 1168767.943 | 1373194.844 | 1298716.779 |
| Dimethyloxallylglycine | 3281394.646 | 5816444.98 | 3301979.399 | 3857381.394 | 5401699.369 | 3776875.202 | 4425114.657 | 4092555.458 | 3472024.392 | 3799196.278 | 3810589.245 | 2208142.242 |
| CAR 17:1 | 2275040.537 | 1325086.345 | 2350295.755 | 1633256.644 | 7571230.13 | 5555999.732 | 6784949.063 | 6317057.99 | 4145673.785 | 5647107.578 | 5371065.624 | 3694841.982 |
| PE(20:5(5Z,8Z,11Z,14Z,17Z)/20:2(11Z,14Z)) | 33535569.86 | 39943657.88 | 53311350.53 | 42922710.01 | 80052264.31 | 62822457.64 | 83601430.45 | 76634663.12 | 78128062.4 | 91997578.76 | 81861927.16 | 50201206.74 |
| Proline | 22786500.93 | 19500516.83 | 33049398.75 | 23577897.93 | 61410086.98 | 70473960.06 | 54931364.44 | 58822022.09 | 80956887.5 | 64209924.34 | 72016860.97 | 72840591.65 |
| 5Z,8Z,11Z,14Z-Eicosatetraenoic acid, 3-theinylmethyl ester | 551446.7811 | 650004.584 | 1676904.481 | 763397.5031 | 1272931.856 | 961390.4528 | 1607665.055 | 1253616.019 | 834526.8833 | 833240.5627 | 864470.1124 | 585497.005 |
| LPC O-22:5 | 71500.81068 | 38575.43626 | 92910.71025 | 69480.19838 | 585989.7323 | 501128.9306 | 647792.8577 | 471769.8722 | 301051.6615 | 372616.0037 | 562226.185 | 277784.7641 |
| Stauprimide | 126126.1718 | 261410.8477 | 67776.42653 | 142545.5031 | 794568.2358 | 2224007.137 | 813774.0459 | 596503.4804 | 553803.7432 | 491219.798 | 746907.0175 | 825908.325 |

| **Metabolites** | **PBS_1** | **PBS_2** | **PBS_3** | **PBS_4** | **BCG_1** | **BCG_2** | **BCG_3** | **BCG_4** | **BCGΔ*pe18*_1** | **BCGΔ*pe18*_2** | **BCGΔ*pe18*_3** | **BCGΔ*pe18*_4** |
|---|---|---|---|---|---|---|---|---|---|---|---|---|
| Valerylcarnitine | 4601123.421 | 2871245.15 | 7445127.121 | 4206500.272 | 26880874.23 | 16784801.38 | 22623128.6 | 21246584.51 | 17897543.35 | 12476106.55 | 15774395.99 | 13415659.7 |
| LPG O-24:5 | 409740.105 | 490749.7927 | 444402.0947 | 1293876.504 | 849379.1377 | 1060188.77 | 648652.5759 | 694199.3883 | 788037.1128 | 678706.908 | 971182.626 | 6315636.136 |
| HexCer 8:0;2O/6:0 | 1059064.104 | 946171.9398 | 510716.7754 | 899619.3881 | 1717276.594 | 1433141.339 | 1121687.788 | 1837243.824 | 2148042.539 | 1398002.132 | 728536.2048 | 817842.6282 |
| Beta-Caryophyllene Alcohol | 634007.9622 | 653728.0725 | 177861.071 | 769146.7066 | 175772.9846 | 744763.226 | 682205.1688 | 892290.643 | 448564.8314 | 425069.4148 | 353432.3471 | 482695.4763 |
| Palmitoleylcarnitine (Car(16:1)) | 5756467.437 | 4682388.598 | 8275897.722 | 5625551.029 | 24771707.21 | 16275734.55 | 25551791.47 | 20499819.16 | 12373049.82 | 16092772.25 | 16088119.56 | 9483408.993 |
| Corynanthin | 34012.54954 | 20172.55479 | 57192.54482 | 33801.94943 | 194666.6137 | 152984.7541 | 263269.3794 | 193277.2502 | 159938.1326 | 144390.1715 | 214231.4338 | 140370.9564 |
| Quizalofop-ethyl | 2943071.881 | 2654900.376 | 4319213.569 | 2780279.713 | 4265095.586 | 3730224.018 | 2942755.064 | 2683575.665 | 2945780.636 | 2431564.126 | 2773996.427 | 2282550.619 |
| 3-ketosphinganine | 627168.9606 | 1257966.215 | 1648770.295 | 902666.1332 | 1127750.261 | 334000.284 | 397262.0546 | 311779.4202 | 987604.3852 | 265555.956 | 332696.2231 | 11826466.03 |
| 1-(1-Hydroxynaphthalen-2-yl)-3-phenylpropane-1,3-dione | 1705871.929 | 1786376.716 | 3026648.073 | 1638321.255 | 1596910.425 | 1541581.873 | 1978004.278 | 1544462.102 | 2253178.492 | 1401169.162 | 1585167.96 | 2239167.719 |
| CAR 17:0 | 1205717.383 | 1019971.28 | 1244382.742 | 1098519.725 | 7688658.295 | 6326134.981 | 7175009.67 | 4877484.473 | 3600188.028 | 4391773.249 | 5338804.454 | 2976952.226 |
| ACARBOSE | 47044.26022 | 0 | 79644.94839 | 67071.86725 | 648287.0582 | 408414.7722 | 573872.2794 | 569452.5399 | 355499.0813 | 281274.7975 | 472788.8906 | 420001.5594 |
| Palmitic acid | 458369 | 424801 | 381378 | 416998 | 310350 | 335383 | 311898 | 292007 | 306562 | 339794 | 335143 | 333011 |

| Metabolites | PBS_1 | PBS_2 | PBS_3 | PBS_4 | BCG_1 | BCG_2 | BCG_3 | BCG_4 | BCGΔ*pe18*_1 | BCGΔ*pe18*_2 | BCGΔ*pe18*_3 | BCGΔ*pe18*_4 |
|---|---|---|---|---|---|---|---|---|---|---|---|---|
| | 945.9 | 878 | 306.9 | 288.1 | 507.4 | 212 | 523.9 | 956.8 | 421 | 867.3 | 668.5 | 936.5 |
| Methylmalonic acid | 20194428.97 | 15318342.21 | 32486123.49 | 19612287.85 | 216229740.3 | 144740857.1 | 212808570.8 | 165026018.3 | 129042382.9 | 152274495.8 | 155691916.6 | 118825196.9 |
| N-(2,14-Eicosadienoyl)piperidine | 110491808 | 106286703.8 | 74673000.8 | 94974962.65 | 83198793.35 | 84233304.22 | 82071913.55 | 89307809.38 | 98047203.63 | 97366272.24 | 89123333.42 | 68392996.23 |
| Ile-Asp | 2064697.88 | 1665727.384 | 2678086.537 | 1917922.212 | 6155956.916 | 5256619.84 | 5426684.229 | 5158177.924 | 3758111.671 | 3755536.852 | 4015365.423 | 4122451.558 |
| LPG 20:1 | 794314.532 | 609387.8332 | 667689.8271 | 611445.5358 | 4753149.897 | 4504711.093 | 6083029.275 | 3658684.965 | 2721447.559 | 2432150.921 | 5461175.603 | 2783133.38 |
| Lauroylcarnitine | 177769.3371 | 1239300.685 | 2547070.671 | 2181176.571 | 9043104.429 | 5411081.454 | 8292676.648 | 6790900.927 | 3060713.739 | 7137264.743 | 6986018.435 | 3569756.655 |
| 1-(3-Methylbut-2-en-1-yl)guanidine | 2611113.703 | 584237.512 | 2697649.663 | 2904755.824 | 2844654.677 | 3092350.157 | 5971048.498 | 3166619.094 | 2908820.5 | 3312418.2 | 2786989.871 | 2755310.65 |
| N2-Methyl-L-arginine | 2606321.197 | 3008232.089 | 1275692.604 | 2157681.92 | 7472639.663 | 1399791.681 | 5737838.649 | 5923394.536 | 4953476.821 | 6778913.254 | 3174574.701 | 1138490.621 |
| antimycin A1a | 796456.4092 | 814066.253 | 1181562.922 | 1008824.176 | 1163060.488 | 1378416.615 | 1268479.755 | 1114919.625 | 1101070.54 | 661369.2444 | 750371.9569 | 1186784.372 |
| Dioctyltin_isooctylthioglycolate | 1953568.666 | 2062594.371 | 1762782.23 | 1930872.034 | 1806146.111 | 2087138.632 | 1667206.034 | 2167024.398 | 2012393.371 | 1748410.325 | 1843833.073 | 1635500.3 |
| IS2 | 14141032.03 | 13912681.68 | 17051093.15 | 14935113.02 | 21061791.82 | 18905563.1 | 21769652.5 | 20704754.95 | 20375151.76 | 18684484.95 | 18695277.1 | 18803517.47 |
| PE O-16:0_20:4 | 3300537.652 | 3450454.538 | 2441898.43 | 3780675.945 | 2146586.514 | 0 | 2594016.345 | 2854885.6 | 4426679.167 | 2509072.82 | 1517441.9 | 3546252.219 |

| Metabolites | PBS_1 | PBS_2 | PBS_3 | PBS_4 | BCG_1 | BCG_2 | BCG_3 | BCG_4 | BCGΔ*pe18*_1 | BCGΔ*pe18*_2 | BCGΔ*pe18*_3 | BCGΔ*pe18*_4 |
|---|---|---|---|---|---|---|---|---|---|---|---|---|
| 1-(3-Phenyl-6,7-dihydro-2H-pyrazolo[4,3-c]pyridin-5(4H)-yl)propan-1-one | 90088.8709 | 741803.2939 | 2084558.016 | 1102780.536 | 34150.03018 | 31266.14496 | 47781.6241 | 31373.63817 | 99814.85164 | 159293.9203 | 20308.7599 | 2712179.522 |
| 1,3-Bis(tris(hydroxymethyl)methylamino)propane | 3067601.596 | 3281806.502 | 3117023.756 | 3890926.085 | 2565808.794 | 4017791.623 | 2749319.598 | 2869082.861 | 2836088.122 | 2564326.699 | 2817433.989 | 3279905.299 |
| Linoleoylcarnitine (Car(18:2)) | 6689959.035 | 5078784.097 | 9075094.298 | 5448120.007 | 5871960.362 | 4895350.79 | 7632252.923 | 6709539.792 | 4194748.903 | 4883423.277 | 4672640.807 | 2979565.614 |
| 7-Oxo-7H-benzo[e]perimidine-4-carboxylic acid | 884315.4501 | 736243.9059 | 54813688.81 | 1511033.579 | 2665829.104 | 2514705.598 | 80279796.84 | 2988081.85 | 1643247.228 | 27425309.47 | 50198913.37 | 2174794.061 |
| 1,24-Dihydroxyvitamin D3 | 123829.9946 | 3863257.644 | 3724624.216 | 3894071.02 | 16500296.13 | 10565778.22 | 12407794.73 | 12449672.77 | 8337254.672 | 7578979.654 | 8962775.673 | 7847646.892 |
| LNAPE 17:1/N-26:6 | 1662633.515 | 1407095.244 | 2125990.886 | 1733643.607 | 1852845.361 | 43227.67778 | 1739400.788 | 7998544.329 | 1893705.192 | 1918223.668 | 1627095.244 | 1982981.449 |
| 3-HYDROXYGLUTARIC ACID | 8803937.994 | 6859326.05 | 6998271.552 | 8097818.159 | 72818776.79 | 34385993.94 | 76612874.27 | 54669493.91 | 34153634.49 | 39286464.19 | 43491275.64 | 29620771.66 |
| Casticin | 1562967.064 | 1073861.748 | 2532265.977 | 1571763.87 | 2426984.096 | 2619759.776 | 2880522.061 | 2381240.22 | 2132753.548 | 2535791.051 | 2172813.298 | 2441243.338 |
| 1-arachidonyl-sn-glycero-3-phospho-(1'-sn-glycerol) | 7179262.358 | 7878237.491 | 10468689.96 | 12138518.68 | 3098047.693 | 4594163.374 | 4426429.483 | 4670802.405 | 4251602.463 | 5648916.718 | 5308046.053 | 3428026.053 |
| Carnosine | 882050. | 573178. | 127919 | 699954. | 210562 | 145164 | 261587 | 186906 | 142929 | 144901 | 154234 | 131058 |

| Metabolites | PBS_1 | PBS_2 | PBS_3 | PBS_4 | BCG_1 | BCG_2 | BCG_3 | BCG_4 | BCGΔ*pe18*_1 | BCGΔ*pe18*_2 | BCGΔ*pe18*_3 | BCGΔ*pe18*_4 |
|---|---|---|---|---|---|---|---|---|---|---|---|---|
| | 5326 | 195 | 3.819 | 7976 | 9.221 | 2.619 | 6.338 | 9.785 | 3.917 | 7.913 | 0.267 | 2.047 |
| dibutyladipate | 1097293.188 | 1159589.564 | 879986.0686 | 729261.3818 | 1014832.16 | 1304849.145 | 1080408.34 | 1013356.719 | 656475.4277 | 858689.4848 | 1040392.722 | 940972.8651 |
| Sulprofos | 12728956.26 | 5301941.42 | 5417574.67 | 12852635.29 | 11927268.78 | 5459425.331 | 4986106.824 | 8458348.637 | 6187977.032 | 7063625.883 | 6145372.174 | 5441938.882 |
| DGDG O-22:6_6:0 | 97112.16177 | 102191.8433 | 75962.66147 | 66746.34858 | 85978.25723 | 94938.63852 | 105079.4554 | 79828.52266 | 130605.1123 | 99556.68435 | 105092.769 | 123275.2358 |
| PE O-14:1_2:0 | 945090.699 | 1399319.588 | 1178497.164 | 786876.2482 | 7302641.312 | 6122356.786 | 7866154.726 | 5457440.812 | 8073341.163 | 6720705.541 | 7941612.462 | 6930959.551 |
| 1-palmitoyl-2-linoleoyl-phosphatidylcholine | 19744761.4 | 9034810.741 | 9566367.465 | 6682176.042 | 21338876.95 | 467112.0264 | 19021877.28 | 6152741.469 | 10477248.98 | 12164711.98 | 10662478.28 | 3679614.128 |
| PA 2:0_26:4 | 2902993.652 | 3085200.495 | 4566670.962 | 3436532.844 | 7616393.589 | 7495092.066 | 8760591.439 | 6757638.625 | 5961192.212 | 6430559.288 | 7913744.822 | 5339011.881 |
| Nolatrexed | 1973008.516 | 10401931.91 | 23143827.01 | 13527141.98 | 1596247.932 | 11831653.58 | 2964049.478 | 349525.2386 | 2276217.111 | 3734717.826 | 2834363.65 | 22777392.38 |
| PI 21:0 | 159255.7811 | 175460.0625 | 169317.3507 | 164551.4191 | 165805.2752 | 166060.4079 | 172616.0563 | 162222.7368 | 156501.3117 | 163819.9226 | 164418.3607 | 165100.8659 |
| 1-Methyl-1H-pyrazolo[3,4-d]pyrimidin-4-ol | 35390554.8 | 36020122.23 | 39397051.41 | 33418187.51 | 17292396.64 | 37093178.65 | 18993536.2 | 31226485.26 | 21561726.91 | 12267884.45 | 12028850.05 | 18345123.98 |
| Piperonyl butoxide | 336775.1589 | 622407.3235 | 512906.3837 | 416638.0981 | 478584.1195 | 538888.959 | 527211.5348 | 525666.0216 | 406200.056 | 434848.6635 | 637607.6485 | 2134178.618 |

| Metabolites | PBS_1 | PBS_2 | PBS_3 | PBS_4 | BCG_1 | BCG_2 | BCG_3 | BCG_4 | BCGΔ*pe18*_1 | BCGΔ*pe18*_2 | BCGΔ*pe18*_3 | BCGΔ*pe18*_4 |
|---|---|---|---|---|---|---|---|---|---|---|---|---|
| 6-Ethoxy-3-(4-methoxyphenyl)-4-methylcoumarin | 7108847.105 | 6244188.711 | 6570131.887 | 6486790.396 | 7150288.348 | 6662349.897 | 6807099.105 | 7172049.721 | 7128297.684 | 6970363.69 | 6975740.32 | 4690443.153 |
| 17-Epimethandienone | 3213750.939 | 3940867.732 | 3160386.78 | 3741505.619 | 3361941.213 | 3525647.949 | 3596132.711 | 3090946.762 | 2913604.456 | 3313197.677 | 2811110.562 | 3543137.99 |
| coniferin | 1085496.095 | 651446.5119 | 940398.1885 | 853005.6785 | 415408.5475 | 720888.4963 | 455581.548 | 621690.0252 | 556291.2067 | 314109.1898 | 351041.2884 | 512255.3146 |
| 4-Penten-1-one, 3,3,4-trimethyl-1-(1-pentyl-1H-indol-3-yl)- | 7591379.746 | 3850093.367 | 1352025.828 | 1138973.578 | 1886595.764 | 2180076.426 | 1044100.495 | 2680219.742 | 3668995.123 | 8565673.642 | 4614849.799 | 1539835.96 |
| 5-(3,4'-Dihydroxy-4,4,7,8a-tetramethyl-6'-oxospiro[2,3,4a,5,6,7-hexahydro-1H-naphthalene-8,2'-3,8-dihydrofuro[2,3-e]isoindole]-7'-yl)pentanoic acid | 0 | 0 | 0 | 32089.93168 | 1267709.418 | 1212537.526 | 1317842.4 | 1532225.097 | 821784.7019 | 877177.8818 | 1015766.184 | 645159.7114 |
| gamma-Linolenic acid | 15175518.26 | 14979331.09 | 13851128.64 | 16748138.91 | 74509521.25 | 94759332.82 | 60990732.02 | 84838105.48 | 56379905.57 | 30252020.15 | 61463034.22 | 55974840.94 |
| PC 17:0 | 316916.0778 | 349279.3883 | 347116.4618 | 389882.1236 | 541165.7302 | 413093.7667 | 463353.5027 | 425665.4365 | 370095.0275 | 444045.7705 | 399508.8543 | 428370.3062 |
| sesamin | 120488. | 87382.2 | 189607. | 143512. | 120856 | 191718 | 205000 | 118108 | 896370. | 111164 | 196243 | 835331. |

| Metabolites | PBS_1 | PBS_2 | PBS_3 | PBS_4 | BCG_1 | BCG_2 | BCG_3 | BCG_4 | BCGΔ*pe18*_1 | BCGΔ*pe18*_2 | BCGΔ*pe18*_3 | BCGΔ*pe18*_4 |
|---|---|---|---|---|---|---|---|---|---|---|---|---|
| | 2173 | 3673 | 5784 | 0259 | 9.251 | 9.695 | 7.42 | 6.099 | 472 | 0.546 | 3.014 | 0649 |
| Ethylmalonic acid | 11174977.38 | 8577077.431 | 8631044.876 | 11232337.04 | 68555567.43 | 41996145.02 | 56586076.01 | 53208723.67 | 40721465.99 | 31478998.08 | 43123974.9 | 32728210.57 |
| SMGDG O-28:5_8:0 | 6810599.194 | 4947318.098 | 2758730.244 | 13350017.65 | 600568.6742 | 480887.0873 | 1013540.593 | 1092000.607 | 1304960.445 | 1148094.492 | 1789000.599 | 716059.9828 |
| LPE O-15:1 | 984690.6122 | 779468.6471 | 630057.4591 | 643213.7418 | 966798.2265 | 829602.8109 | 946768.5189 | 838818.9757 | 1016931.681 | 847726.5378 | 875990.2682 | 145231.864 |
| DGDG O-14:1_26:7 | 5603732.289 | 3863588.589 | 4371676.646 | 5336374.566 | 13769149.89 | 9925801.404 | 13443355.24 | 10873426.33 | 7990319.678 | 9958560.802 | 13685730.21 | 4569440.672 |
| 1-(11Z-eicosenoyl)-glycero-3-phosphate | 531538.9207 | 1027313.392 | 302186.7903 | 471975.1388 | 761774.8383 | 671266.045 | 646174.1767 | 635060.6034 | 605012.6816 | 705365.5665 | 629137.7152 | 508197.999 |
| 2-[5-(2-Hydroxypropyl)oxolan-2-yl]propanoic acid | 1122142.499 | 1142569.98 | 1018865.839 | 1044547.955 | 1118703.743 | 1102493.923 | 1114898.934 | 1072421.822 | 1097356.848 | 1117132.132 | 1086681.78 | 1097274.277 |
| Val-Phe | 419269.6723 | 399999.99 | 390498.6079 | 364491.2132 | 430373.8837 | 278948.9356 | 394136.1801 | 300296.6861 | 407368.6708 | 402042.8178 | 421247.8598 | 329410.2574 |
| PEtOH 16:0_18:4 | 2750196.235 | 2561787.019 | 1756059.224 | 1891955.465 | 1171936.444 | 1157524.326 | 1910366.677 | 1727173.261 | 2046664.441 | 1660061.913 | 1247543.634 | 1147002.634 |
| 2-Hydroxy-4-methoxy-5-(2-methylbut-3-en-2-yl)benzaldehyde | 2067528.163 | 2007102.811 | 750545.3809 | 2535694.185 | 1820840.952 | 3105367.14 | 1323944.126 | 2209862.157 | 1610131.535 | 1844968.388 | 1702499.796 | 1516233.105 |
| PC O-16:0 | 373435. | 263445. | 410635. | 392120. | 569212. | 498228. | 521905. | 575128. | 536696. | 649038. | 556068. | 514501. |

| Metabolites | PBS_1 | PBS_2 | PBS_3 | PBS_4 | BCG_1 | BCG_2 | BCG_3 | BCG_4 | BCGΔ *pe18*_1 | BCGΔ *pe18*_2 | BCGΔ *pe18*_3 | BCGΔ*pe18*_4 |
|---|---|---|---|---|---|---|---|---|---|---|---|---|
| | 07 | 9384 | 9068 | 9183 | 7763 | 9949 | 3993 | 3742 | 0892 | 1442 | 4339 | 9244 |
| LPA 20:0 | 127367.0678 | 180166.0195 | 90810.08123 | 165893.3804 | 328670.095 | 242505.9683 | 392407.825 | 384095.1387 | 365519.6673 | 1161336.723 | 242954.0565 | 529575.9464 |
| N-Cyclohexyl-N'-cyclooctylurea | 1825973.531 | 1836435.96 | 1718817.597 | 1797529.39 | 1529423.155 | 2037370.983 | 1620405.567 | 1557405.474 | 1482075.841 | 1617939.704 | 1361621.216 | 1498947.089 |
| N-(Cyclohexylmethyl)cyclohexanamine | 1754012.854 | 1999464.316 | 1954549.459 | 1799782.031 | 1356005.008 | 1572452.084 | 1261072.622 | 1285162.941 | 1397722.503 | 1009895.054 | 1247073.755 | 1865876.676 |
| 1-Butylimidazole | 3513377.686 | 2848010.36 | 3177472.321 | 3374978.425 | 2713797.256 | 3288929.203 | 2588014.142 | 749084.478 | 1205835.907 | 2969576.722 | 2838961.469 | 3353376.573 |
| 1,4-D-Xylobiose | 1506926.267 | 1072418.872 | 1221803.51 | 1270683.277 | 4568368.22 | 4552591.058 | 3985088.558 | 4322497.035 | 4363640.721 | 2058288.635 | 1976476.784 | 2177117.592 |
| LPS(18:0) | 699152.9149 | 716494.6866 | 290885.6457 | 556187.4047 | 502798.7179 | 315885.8221 | 594123.0667 | 522722.7613 | 614527.1525 | 598417.5834 | 652065.3874 | 328947.2652 |
| rac-trans-4-Cotinine carboxylic acid methyl ester | 2531341.343 | 1761984.196 | 1797282.207 | 1721284.522 | 2299996.78 | 2686491.38 | 2800594.556 | 3001016.805 | 2193284.009 | 1323148.36 | 2169747.517 | 919781.669 |
| PC O-17:0 | 16336348.45 | 16299257.35 | 21004252.51 | 15001901.19 | 49171307.45 | 35801458.16 | 27971229.08 | 25483768.78 | 20515653.56 | 17089945.25 | 23358971.48 | 18931416.18 |
| 14-(hydroxymethyl)-5,9-dimethyltetracyclo[11.2.1.0.0]hexadecane-5-carbaldehyde | 420700.9343 | 236670.0096 | 251714.8094 | 14293607.47 | 4949235.939 | 10762804.17 | 3903569.396 | 5244668.707 | 3498865.048 | 2513613.496 | 4396726.331 | 4524265.449 |

| Metabolites | PBS_1 | PBS_2 | PBS_3 | PBS_4 | BCG_1 | BCG_2 | BCG_3 | BCG_4 | BCGΔ*pe18*_1 | BCGΔ*pe18*_2 | BCGΔ*pe18*_3 | BCGΔ*pe18*_4 |
|---|---|---|---|---|---|---|---|---|---|---|---|---|
| LPC(18:0/0:0) | 16137154.38 | 6811937.892 | 15268787.42 | 5737462.316 | 17485043.61 | 11661997.07 | 6671254.792 | 21118131.26 | 14504715.4 | 12822811.6 | 11799902.18 | 2892185.953 |
| Benzyltrimethylammonium cation | 2039029.254 | 2057785.445 | 1955575.854 | 1811701.093 | 1833198.47 | 1910272.673 | 3318490.053 | 517217.7462 | 1853343.642 | 1963897.721 | 1836234.071 | 2033906.565 |
| LPS 17:0 | 1422166.669 | 1460305.524 | 872137.1713 | 1040933.958 | 1365508.332 | 1617347.346 | 1313737.549 | 1001993.559 | 1427169.134 | 1187602.962 | 1824963.392 | 714868.502 |
| Nicotinamide adenine dinucleotide phosphate (NADP) | 1472100.534 | 1074039.541 | 1205084.079 | 1207648.893 | 1486215.329 | 1286597.585 | 2045004.955 | 1406042.598 | 1721064.856 | 1612855.052 | 1329408.178 | 2490100.152 |
| p-Toluquinone | 68022799.92 | 53951336.24 | 46804104.78 | 51543782.85 | 53595831.79 | 56786380.98 | 56843838.8 | 36514278.36 | 52001816.65 | 51730132.88 | 47984383.4 | 61606854.03 |
| Dye X-6012-14H | 989767.2339 | 779576.8852 | 1165034.875 | 875532.2274 | 1591285.118 | 926287.4723 | 1682475.529 | 1161605.137 | 870734.1875 | 1097845.621 | 1269067.05 | 628370.803 |
| PI 18:0_22:6 | 2378122.044 | 1531511.083 | 2179167.542 | 4219043.602 | 3938741.38 | 3927978.361 | 4024759.846 | 3504724.426 | 10172018.87 | 4667430.866 | 7145769.433 | 7218885.804 |
| Sophoraflavonoloside | 1252385.882 | 939700.2732 | 2368718.748 | 946816.1997 | 1241929.725 | 1363418.486 | 1270394.586 | 1266948.557 | 1112471.939 | 1032476.044 | 957977.6516 | 1019211.114 |
| LNT | 0 | 0 | 3698.271998 | 0 | 168983.3295 | 56516.18492 | 161452.7431 | 81151.54409 | 156868.6087 | 109239.8765 | 140226.7121 | 112920.831 |
| CYCLOPENTANONE | 7532740.805 | 30095.94331 | 6020326.229 | 6927813.433 | 7222190.31 | 7728517.221 | 5782043.537 | 8167191.255 | 7670053.358 | 9058533.515 | 7675246.965 | 7719543.526 |
| 1-Carboxyethylisoleucine | 510888.5899 | 862906.1458 | 1111240.25 | 5935266.746 | 8521007.344 | 259366.2939 | 725294.2123 | 229626.0272 | 586470.1733 | 2889883.969 | 2772902.254 | 3324392.51 |

| Metabolites | PBS_1 | PBS_2 | PBS_3 | PBS_4 | BCG_1 | BCG_2 | BCG_3 | BCG_4 | BCGΔ*pe18*_1 | BCGΔ*pe18*_2 | BCGΔ*pe18*_3 | BCGΔ*pe18*_4 |
|---|---|---|---|---|---|---|---|---|---|---|---|---|
| 5-Hydroxyhexanoic acid | 2607442.12 | 2516791.118 | 2499580.208 | 2657883.819 | 2681318.29 | 2444323.768 | 2591915.114 | 2601494.849 | 2521355.901 | 2602310.8 | 2572239.424 | 2576690.995 |
| N-(3-Piperidinyl)propanamide | 1178191.203 | 1233741.619 | 1171931.481 | 1241871.126 | 1227257.159 | 1213525.1 | 1206417.1 | 1162630.158 | 1140825.999 | 1175815.484 | 1166122.702 | 1153964.973 |
| Theaflavin | 3424185.986 | 2829729.997 | 3997382.77 | 3291925.283 | 11671726.89 | 8860055.02 | 17033745.84 | 14158980.72 | 11437275.48 | 11433822.59 | 13149422.2 | 16100779.69 |
| Thiomorpholine 3-carboxylate | 3079706.689 | 1209699.031 | 3615602.916 | 2057879.319 | 4337021.565 | 5730629.069 | 12180684.68 | 11396393.76 | 9109485.727 | 3826419.187 | 5884746.725 | 6607867.387 |
| TG 8:0_16:4_24:6 | 1634233.206 | 1703318.59 | 1046092.51 | 1373592.001 | 1619450.33 | 1497347.622 | 1595528.614 | 1772901.605 | 1834348.972 | 1849605.089 | 1703526.211 | 1335028.328 |
| 4-Hexylresorcinol | 253250.3348 | 1933170.116 | 2473920.31 | 1995768.643 | 5325618.373 | 1881959.284 | 2340157.456 | 2412399.175 | 2339825.58 | 1681260.901 | 2680837.97 | 2417883.314 |
| Dihydroxyacetone_Phosphate_Acyl_Ester | 4103332.454 | 2922243.12 | 4817582.062 | 3697747.208 | 2948275.266 | 5116886.483 | 3280859.62 | 4772915.589 | 2920960.445 | 3057060.361 | 5040882.583 | 3071084.588 |
| LNAPS 24:2/N-12:0 | 5592406.816 | 1557772.3 | 2234396.474 | 1667384.022 | 2066153.931 | 1070593.04 | 1639856.913 | 1362349.598 | 1862804.655 | 2085949.5 | 135536.907 | 1022490.666 |
| SHexCer 32:0;3O | 1402526.837 | 749819.2997 | 498861.1582 | 448540.4304 | 2886673.37 | 2974650.425 | 5932443.165 | 3514801.911 | 3750379.632 | 6063057.529 | 5314042.114 | 2048836.261 |
| 2-Amino-7-phenyl-7,8-dihydro-5(6H)-quinazolinone | 1275742.839 | 894181.5041 | 1503647.858 | 908529.299 | 1845364.479 | 1276522.312 | 1533440.785 | 1480868.174 | 1463532.169 | 1334857.167 | 1387382.998 | 968530.7515 |
| (.+/-.)-Cromakalim | 2101389.891 | 2299325.57 | 2029418.757 | 2094644.463 | 2259518.362 | 2183982.338 | 2335579.833 | 2138559.154 | 2150338.35 | 2270343.663 | 2133710.416 | 2103967.277 |

| Metabolites | PBS_1 | PBS_2 | PBS_3 | PBS_4 | BCG_1 | BCG_2 | BCG_3 | BCG_4 | BCGΔ*pe18*_1 | BCGΔ*pe18*_2 | BCGΔ*pe18*_3 | BCGΔ*pe18*_4 |
|---|---|---|---|---|---|---|---|---|---|---|---|---|
| Sebacic acid | 3363773.833 | 3224847.884 | 3890129.222 | 4719410.58 | 3241908.052 | 3152926.493 | 3741442.335 | 2991739.872 | 3030067.384 | 2944930.176 | 3002652.83 | 4146880.431 |
| LPI 19:1 | 545212.3683 | 407674.1126 | 366511.4472 | 396479.2687 | 2975955.098 | 2871284.83 | 2349227.395 | 2192804.24 | 2157510.819 | 1872527.666 | 2719298.975 | 1731212.39 |
| Termitomycamide B | 498775.9791 | 524624.7097 | 380940.6242 | 501221.7563 | 789915.8676 | 669699.4758 | 535942.8979 | 492132.0444 | 490969.478 | 466443.535 | 657480.2867 | 528137.2089 |
| 4-Hydroxy-6-methoxycoumarin | 1115237.263 | 976318.3266 | 877065.2442 | 965231.0817 | 1044996.325 | 777657.5256 | 600137.4121 | 789695.7858 | 1061070.353 | 813848.6767 | 1609462.128 | 732398.5521 |
| LPA 19:2 | 611134.6699 | 648451.7723 | 1181030.903 | 1022955.079 | 1706858.254 | 1512732.367 | 2183839.92 | 1672723.855 | 1289618.006 | 1781360.136 | 2202035.21 | 1271537.759 |
| Prostaglandin_D2-1-glyceryl_ester_ | 933504.7947 | 1052380.652 | 1347564.211 | 1092868.705 | 1497482.389 | 1534966.44 | 1505743.736 | 1597131.727 | 1495422.648 | 1101859.928 | 1355478.809 | 1423027.684 |
| ST 24:1;O4;S | 73949.44941 | 70945.24409 | 83019.88182 | 92812.04612 | 84819.32179 | 89654.33025 | 69197.00035 | 82322.56534 | 166775.1811 | 125951.8126 | 131669.6635 | 157298.8869 |
| 6-Methyl-2,4-pyrimidinedithiol | 3676842.418 | 4383019.652 | 3367359.707 | 5967584.832 | 3738350.516 | 3042461.664 | 3928568.585 | 1498457.647 | 4068623.872 | 4631628.631 | 5533616.406 | 3346709.248 |
| Cer 8:0;2O/26:7 | 18258059.84 | 17931304.57 | 12786187.72 | 21492657.86 | 52611114.61 | 42293816.73 | 24801403.98 | 43457076.97 | 31981019.5 | 14883602.51 | 13768834.86 | 35614429.62 |
| Urobilin | 437599.6298 | 331242.1443 | 455971.8012 | 409250.3261 | 485228.6065 | 631065.6317 | 446513.9523 | 508129.1294 | 431138.8349 | 414414.2706 | 454850.1211 | 409840.8739 |
| PS 20:4_20:4 | 348673.3403 | 841407.9869 | 281434.0017 | 272254.4776 | 828902.4613 | 514957.0007 | 614260.2006 | 897507.1316 | 938045.7306 | 1082871.164 | 662212.9558 | 586454.5876 |
| FAHFA 2:0/20:4 | 759684.2093 | 710802.5316 | 750505.0845 | 737654.368 | 3180472.601 | 5629125.879 | 2864834.402 | 3216955.048 | 2332201.605 | 2002971.056 | 3044554.992 | 2622582.65 |

| Metabolites | PBS_1 | PBS_2 | PBS_3 | PBS_4 | BCG_1 | BCG_2 | BCG_3 | BCG_4 | BCGΔ*pe18*_1 | BCGΔ*pe18*_2 | BCGΔ*pe18*_3 | BCGΔ*pe18*_4 |
|---|---|---|---|---|---|---|---|---|---|---|---|---|
| 2-Amino-6-phosphonohexanoic acid | 2416009.183 | 1783411.151 | 2884686.082 | 1959955.015 | 4442966.693 | 2761356.641 | 3514318.385 | 3495170.909 | 3391712.922 | 2860532.713 | 2818211.748 | 2403987.53 |
| PE-Cer 19:3;2O/26:7 | 642834.4707 | 304336.035 | 762614.9722 | 569086.9154 | 1042578.434 | 489134.199 | 1049227.988 | 1147752.939 | 814797.5373 | 1466328.533 | 1072463.929 | 681962.7767 |
| 2-Oleoyl-1-stearoyl-sn-glycero-3-phosphoserine | 3380309.616 | 4315300.952 | 3241595.588 | 4678349.961 | 1937222.782 | 12710118.92 | 4581836.134 | 3197402.19 | 6124161.478 | 2695557.798 | 1356066.304 | 3246050.829 |
| 4-O-(beta-Arabinofuranosyl)-4-hydroxyproline | 6554171.24 | 5250441.095 | 15986740.96 | 5727002.406 | 4394618.882 | 3462875.473 | 5851308.942 | 4021266.766 | 2061157.37 | 2271946.261 | 4321799.49 | 2284728.202 |
| Capsi-amide | 52807385.01 | 51840761.3 | 55749519.73 | 39981338.06 | 76312914.39 | 47142137.38 | 54417985.49 | 62527312.29 | 62551172.75 | 71396306.84 | 61047859.93 | 32587207.83 |
| 13(S)-HODE | 5121272.172 | 5225397.573 | 6120695.296 | 5453166.026 | 51513838.67 | 53705779.39 | 61899333.47 | 42525806.54 | 36387025.84 | 16015271.42 | 39660987.14 | 29742425.4 |
| 6-Bromo-2-methyl-4(1H)-quinolinone | 17202655.38 | 2456179.183 | 8015245.451 | 8703753.614 | 6603408.072 | 7668067.323 | 7626823.908 | 11342767.89 | 8489949.072 | 1792005.288 | 8070611.843 | 11439052.06 |
| Dihydroxyacetone | 834499757.7 | 658399698.8 | 795444443.7 | 669463945.4 | 1048826437 | 714518879 | 1266963661 | 819839606.2 | 677868500.6 | 480293765.2 | 486393922.1 | 562765838 |
| PEtOH 19:0_22:6 | 541533.9563 | 554540.0245 | 1103927.523 | 991184.0913 | 869439.6433 | 1067658.803 | 1240005.436 | 1045193.241 | 1012684.05 | 1632640.961 | 275701.1303 | 808146.1656 |
| 2-oxopent-4-enoate | 75807.56852 | 9570357.542 | 11488779.97 | 19497379.67 | 10709779 | 15277039.88 | 9991731.963 | 13604790.01 | 7116834.42 | 7516026.495 | 8270724.284 | 12535887.73 |
| PC 9:0_22:1 | 5547714.632 | 4807686.023 | 2871818.653 | 2310374.329 | 24408271.36 | 15509704.01 | 22929957.54 | 23450660.68 | 23017806.01 | 14129382.07 | 14781411.57 | 18033662.59 |

| Metabolites | PBS_1 | PBS_2 | PBS_3 | PBS_4 | BCG_1 | BCG_2 | BCG_3 | BCG_4 | BCGΔ*pe18*_1 | BCGΔ*pe18*_2 | BCGΔ*pe18*_3 | BCGΔ*pe18*_4 |
|---|---|---|---|---|---|---|---|---|---|---|---|---|
| 1-(Cyclopropylcarbonyl)-3-piperidinamine | 1408910.403 | 1379351.673 | 1089526.496 | 1294992.617 | 1315025.46 | 1103543.66 | 1559219.509 | 1266711.722 | 1565024.789 | 1596470.562 | 1314274.676 | 1311595.475 |
| LPC 22:4 | 1586217.387 | 1490533.827 | 3026758.1 | 1317453.279 | 786123.5812 | 948713.6612 | 1775789.206 | 1038045.799 | 1103665.376 | 1400066.632 | 1350027.379 | 734319.3395 |
| PS 39:1 | 63974529.3 | 76859176.31 | 62717939.23 | 67822105.38 | 74653397.49 | 69451229.75 | 68435834.7 | 68239366.85 | 72363989.94 | 76489628.4 | 69005689.32 | 66661314.46 |
| N-(2-Phenylethyl)-1-pentanamine | 1956745.285 | 2215714.768 | 1889818.772 | 2056560.925 | 1949145.87 | 1898454.441 | 1993815.908 | 2068175.382 | 2041356.044 | 2025669.513 | 2031088.783 | 1852598.553 |
| 2-Hydroxylauroylcarnitine | 75554.90851 | 277451.5549 | 613612.935 | 540682.7204 | 612357.9249 | 1251781.412 | 2282305.451 | 403410.1943 | 1042472.446 | 665662.3372 | 1323862.702 | 656929.8494 |
| LPE(P-16:0) | 22231909.15 | 20079921.16 | 13990462.82 | 16882320.61 | 11878352.54 | 8821888.31 | 12035872.7 | 11079477.19 | 12736606.79 | 11307666.36 | 11679375.2 | 6992898.204 |
| Benoxacor | 2627292.264 | 1542403.625 | 3840851.943 | 2010710.676 | 10674210.49 | 4846165.778 | 9917007.373 | 6699759.706 | 6071269.965 | 5577367.933 | 7010868.382 | 4885826.95 |
| Thr-Tyr | 563384.3224 | 511525.0928 | 830786.2769 | 568537.3997 | 4154702.281 | 3562232.685 | 3361557.023 | 3205388.437 | 2613618.397 | 2020896.347 | 2256058.939 | 2565473.501 |
| Glyceraldehyde | 834499757.7 | 658399698.8 | 795444443.7 | 669463945.4 | 1048826437 | 714518879 | 1266963661 | 819839606.2 | 677868500.6 | 480293765.2 | 486393922.1 | 562765838 |
| Pro-Asp | 562023.5662 | 737607.0968 | 1153606.756 | 824491.9775 | 2422201.21 | 2135333.455 | 1334448.714 | 688781.6129 | 2690543.013 | 2680615.503 | 2635715.28 | 2152008.512 |
| 1,2-Dipalmitoyl-sn-glycero-3-phosphoethanolamine | 2752953.088 | 2388074.804 | 2474643.295 | 2663382.831 | 2069141.775 | 53608.47967 | 2170213.805 | 895811.5121 | 2906803.142 | 2319557.043 | 2535418.631 | 816633.6901 |

| Metabolites | PBS_1 | PBS_2 | PBS_3 | PBS_4 | BCG_1 | BCG_2 | BCG_3 | BCG_4 | BCGΔ*pe18*_1 | BCGΔ*pe18*_2 | BCGΔ*pe18*_3 | BCGΔ*pe18*_4 |
|---|---|---|---|---|---|---|---|---|---|---|---|---|
| Enniatin B | 903021.1846 | 664982.6904 | 655748.4509 | 689013.8247 | 708236.1472 | 624539.9638 | 588564.6531 | 508111.9406 | 588923.2599 | 762694.3356 | 830787.3064 | 374124.2813 |
| MGDG O-10:0_6:0 | 1552169.183 | 1392741.061 | 1412865.013 | 1366613.23 | 2043049.492 | 1489540.063 | 1616968.22 | 1747789.183 | 1493325.341 | 1558380.104 | 1425098.811 | 1351290.54 |
| SM 9:0;2O/22:4 | 1274392.458 | 888208.4776 | 573311.3264 | 43875547.24 | 0 | 3089360.467 | 104105998.7 | 0 | 12222520.35 | 6505677.948 | 0 | 45762546.18 |
| MGDG 2:0_8:0 | 0 | 9467.09318 | 15571.10663 | 6293.180728 | 355812.8852 | 303070.5853 | 289170.7702 | 264852.2271 | 182533.3841 | 195088.3089 | 225337.9064 | 156596.7883 |
| PC 8:0_34:9 | 16197043.42 | 18115493.06 | 7711060.971 | 11837603.01 | 7568200.22 | 4431261.312 | 5722420.926 | 6059121.026 | 8112224.865 | 7837105.939 | 5254462.643 | 5929492.162 |
| DGDG 17:1_22:4 | 229602.4509 | 177002.9809 | 197009.3426 | 141150.4365 | 381271.9636 | 407277.1767 | 400444.3956 | 376235.385 | 273867.0138 | 331324.7573 | 484067.4578 | 133311.8548 |
| Haplamine | 3075338.852 | 2264773.189 | 4063059.909 | 3610910.828 | 15190041.66 | 5962789.275 | 12007248.97 | 9494307.837 | 8380862.207 | 7699887.715 | 7864479.342 | 5380072.156 |
| Heptadecasphing-4-enine | 10600601.54 | 12244375.96 | 5257075.332 | 9343390.89 | 2813038.486 | 2914883.243 | 3220220.756 | 3651191.369 | 2536628.784 | 1865773.316 | 1928979.892 | 1265338.903 |
| 4,4'-[(4-Imino-2,5-cyclohexadien-1-ylidene)methylene]bis[benzenamine] | 34995310.6 | 37223349.05 | 36598315.74 | 38459719.6 | 34591494.27 | 33754415.56 | 35406738.11 | 35734800.37 | 35131604.43 | 35186836.09 | 37832171.39 | 33606521.37 |
| Succinic semialdehyde | 1603732.83 | 1187813.442 | 1806289.215 | 1474462.535 | 8099657.845 | 5654437.696 | 8019942.812 | 6784861.885 | 5477006.153 | 8021634.831 | 6979161.128 | 4691350.805 |
| LPA 26:4 | 820701.6869 | 714630.4177 | 1213155.675 | 770113.0318 | 1460399.619 | 1536378.099 | 15230011.53 | 1215909.071 | 1335559.585 | 1270305.274 | 1600174.753 | 1093877.3 |

| Metabolites | PBS_1 | PBS_2 | PBS_3 | PBS_4 | BCG_1 | BCG_2 | BCG_3 | BCG_4 | BCGΔ*pe18*_1 | BCGΔ*pe18*_2 | BCGΔ*pe18*_3 | BCGΔ*pe18*_4 |
|---|---|---|---|---|---|---|---|---|---|---|---|---|
| Ethyl 3-amino-4-(methylsulfonyl)-5-(methylthio)thiophene-2-carboxylate | 1004428.96 | 837617.2249 | 1286579.833 | 920408.9524 | 1516999.535 | 1601854.06 | 2268638.087 | 1531526.981 | 1609223.187 | 1564889.14 | 1635417.555 | 1502015.446 |
| 4-Amino-2-(methylsulfanyl)-5-pyrimidinecarboxylic acid | 3985533.684 | 5062666.412 | 4072597.836 | 3284832.543 | 7690127.537 | 2778474.427 | 7539062.174 | 8054044.487 | 8098075.598 | 2759530.116 | 2562593.844 | 1619802.117 |
| N,N-Diethyl 3-bromobenzenesulfonamide | 20879873.01 | 20145329.53 | 15937799.77 | 21366920.93 | 12328657.65 | 17977325.87 | 18666493.52 | 12746570.52 | 14786733.43 | 9221847.669 | 5685578.925 | 5346346.654 |
| 5-Hydroxyisophthalic acid | 1137287.362 | 1990572.214 | 2432343.052 | 1102105.161 | 1671879.918 | 1396400.547 | 1658014.507 | 1175797.948 | 1550023.205 | 1417733.387 | 1381908.656 | 1637706.01 |
| 3-{[4-(Trifluoromethyl)phenyl]amino}cyclohex-2-en-1-one | 1822954.443 | 1307330.895 | 2490397.451 | 1545328.285 | 7539421.217 | 3404873.561 | 5640340.957 | 4824189.972 | 5037688.781 | 4003040.935 | 4101126.196 | 3064250.364 |
| 1-Palmitoyl-2-lauroyl-sn-glycero-3-phosphorylcholine | 1013355.948 | 349526.4142 | 561799.8378 | 382340.8677 | 32820113.57 | 8811079.434 | 17818576.37 | 13794567.35 | 6130105.75 | 27571859.18 | 5931590.044 | 4713731.467 |
| LPI 18:2 | 9358558.95 | 9456602.729 | 14733366.13 | 9832711.699 | 17626941.14 | 13486298.63 | 9696943.904 | 9389170.117 | 10449683.23 | 9849981.979 | 16457041.51 | 8883798.557 |
| DGDG 2:0_4:0 | 5162732.519 | 6152830.594 | 3732584.203 | 4935493.278 | 4995261.714 | 13231292.94 | 5079755.027 | 4651079.398 | 3824265.367 | 4305462.649 | 4806358.973 | 4311431.652 |
| LysoPC(0:0/18:0) | 654450 | 725378 | 496454 | 900657 | 655307 | 115975 | 791202 | 744373 | 641200 | 539952 | 649524 | 350302 |

| Metabolites | PBS_1 | PBS_2 | PBS_3 | PBS_4 | BCG_1 | BCG_2 | BCG_3 | BCG_4 | BCGΔ*pe18*_1 | BCGΔ*pe18*_2 | BCGΔ*pe18*_3 | BCGΔ*pe18*_4 |
|---|---|---|---|---|---|---|---|---|---|---|---|---|
| | 2.783 | 4.134 | 8.225 | 7.014 | 5.563 | 90.45 | 9.425 | 2.141 | 5.217 | 1.499 | 8.556 | 6.626 |
| Methyl 3-[(1E,3E)-3,5-dimethylhepta-1,3-dienyl]-5,6-dihydroxy-6-methyl-7-oxocyclopenta[c]pyridine-5-carboxylate | 551684.1775 | 538574.4366 | 457361.2848 | 499397.6666 | 584530.4386 | 635557.2919 | 581975.7879 | 575901.0057 | 602927.208 | 592563.0884 | 638850.9159 | 568102.7705 |
| Tridec-3-enoylcarnitine | 1275637.843 | 1260904.838 | 582354.514 | 1413823.033 | 1320678.767 | 1050020.135 | 1113931.681 | 1224764.692 | 1138896.397 | 1655042.903 | 1179202.437 | 873314.335 |
| N-Acetyl-glutamate 5-semialdehyde | 298460.5101 | 369240.1928 | 274709.6658 | 356186.8698 | 4285476.59 | 2812758.952 | 5646229.228 | 3292133.352 | 2719124.318 | 2419234.597 | 2842204.669 | 1932050.389 |
| SL 14:3;O/18:5 | 3516400.033 | 3407025.293 | 1769814.312 | 5287277.237 | 3838481.475 | 8719750.368 | 5166098.291 | 5332881.683 | 2514219.637 | 4523495.133 | 5476590.365 | 3737096.964 |
| Tridemorph | 2830077.34 | 3430140.61 | 2642612.346 | 3301240.559 | 2250150.886 | 6440539.928 | 3344997.668 | 4918812.219 | 3220740.356 | 2487015.564 | 4189580.153 | 3137117.987 |
| 3-Methylpiperidin-2-one | 86272.82053 | 9576596.337 | 16099835.8 | 9364177.677 | 9204516.296 | 9970795.399 | 9534338.539 | 9162886.543 | 9643370.742 | 8666386.789 | 9261503.178 | 9098028.796 |
| (5E,9E)-Farnesylacetone | 4047805.357 | 5570097.041 | 1657264.374 | 2525973.141 | 2340766.548 | 2370139.937 | 2431279.78 | 2335788.015 | 3892912.656 | 4740790.659 | 3706283.592 | 1707021.413 |
| SM 12:2;2O/5:0 | 147884.0893 | 190596.8896 | 280336.5471 | 256479.4591 | 233018.5011 | 234439.8452 | 242650.1479 | 247001.5226 | 208791.7023 | 187906.1214 | 221961.9306 | 223704.6901 |
| PC O-34:4 | 13834914.73 | 13777838.47 | 9022394.112 | 9609432.832 | 3947520.664 | 4370338.845 | 5365265.588 | 5280690.703 | 5502610.786 | 6440218.843 | 5465030.511 | 4060236.074 |

| Metabolites | PBS_1 | PBS_2 | PBS_3 | PBS_4 | BCG_1 | BCG_2 | BCG_3 | BCG_4 | BCGΔ*pe18*_1 | BCGΔ*pe18*_2 | BCGΔ*pe18*_3 | BCGΔ*pe18*_4 |
|---|---|---|---|---|---|---|---|---|---|---|---|---|
| Lupanine | 1811853.398 | 2091895.355 | 1264167.192 | 1542740.124 | 1707885.35 | 1387509.563 | 2174912.861 | 1938873.703 | 2019523.198 | 2172181.331 | 1537261.312 | 1670457.233 |
| DEHP | 104995972.4 | 96058370.21 | 69428142.79 | 79540784.74 | 283128953.6 | 273881722.1 | 279897290.4 | 300551422.1 | 279036206.6 | 177880767.8 | 201414337.9 | 128016046.3 |
| Norphenylephrine | 3184263.56 | 2938319.478 | 2916416.449 | 1401125.716 | 2633925.817 | 3239016.829 | 2956010.425 | 2923354.48 | 2923639.937 | 2699511.146 | 2516268.917 | 845514.1502 |
| 12alpha-Hydroxy-13,18-dehydroparain | 212847.987 | 643639.4207 | 399072.3799 | 480103.8887 | 519385.4359 | 350963.8775 | 390085.1108 | 407309.3807 | 321062.4445 | 742094.9712 | 409642.5204 | 261732.089 |
| 4-Hydroxybenzoic acid | 6098039.887 | 6768398.521 | 5931187.11 | 5967154.819 | 6450340.457 | 6483876.317 | 6730905.511 | 6701565.666 | 7282675.944 | 6812535.089 | 7637531.198 | 7404943.903 |
| 1-[(4-tert-Butylphenyl)sulfonyl]piperazine | 1926235.208 | 2047981.71 | 1747661.028 | 1921841.453 | 1912247.489 | 2033255.622 | 2097650.434 | 1957841.735 | 2051059.181 | 2098435.014 | 1834424.27 | 1893496.856 |
| PE O-23:0_22:6 | 333817.192 | 445085.2294 | 441952.625 | 550545.9215 | 160079.5489 | 298675.3222 | 194776.3676 | 224983.0264 | 226343.895 | 197648.2361 | 29703.68729 | 332099.3084 |
| NAGlySer 24:6/21:1 | 96079.52595 | 77353.25152 | 63530.89921 | 66033.5222 | 203967.7112 | 206792.2839 | 129094.8338 | 152246.3624 | 126472.1023 | 141957.9326 | 109864.1948 | 73933.81962 |
| PG 16:1_22:6 | 5097686.865 | 2502947.199 | 1461377.606 | 3189644.012 | 24921809.32 | 7958557.9 | 18619660 | 10132780.28 | 18597781.32 | 27939573.33 | 15927478.31 | 9018248.661 |
| MGDG O-13:0_24:5 | 626582.0483 | 537489.6163 | 2085388.811 | 573182.5927 | 1602568.234 | 213071.6499 | 223368.0373 | 380537.6805 | 676128.1359 | 282313.9187 | 1006823.865 | 406956.4648 |
| SM 35:5;3O | 9134506.255 | 2580494.333 | 10429276.08 | 1226876.142 | 10083153.11 | 1607551.474 | 7351175.874 | 10453574.68 | 1748108.396 | 4182846.224 | 6535840.657 | 4801657.588 |

| Metabolites | PBS_1 | PBS_2 | PBS_3 | PBS_4 | BCG_1 | BCG_2 | BCG_3 | BCG_4 | BCGΔ *pe18*_1 | BCGΔ *pe18*_2 | BCGΔ *pe18*_3 | BCGΔ*pe18*_4 |
|---|---|---|---|---|---|---|---|---|---|---|---|---|
| 2,4,6-Tri-tert-butyl-N-methylaniline | 5267530.537 | 5574004.721 | 5016242.704 | 5169480.003 | 5277145.491 | 5480477.475 | 5715299.62 | 4883136.754 | 5594265.302 | 5646000.872 | 5229704.487 | 4643465.039 |
| Cer 14:3;2O/2:0 | 1310268.883 | 1299555.659 | 2250414.808 | 1538987.492 | 1015694.115 | 1125237.43 | 1470917.795 | 982417.4942 | 1182055.357 | 1151604.491 | 955849.5074 | 1516908.781 |
| PE O-16:3_22:5 | 6100374.328 | 860410.8061 | 19865348.97 | 1121343.388 | 8737537.127 | 42053.80713 | 9563874.983 | 17979649.79 | 2414560.678 | 11102645.97 | 5895311.961 | 2216288.924 |
| DGDG 5:0_22:6 | 87393.73318 | 43250.00765 | 27044.47517 | 63820.93588 | 344878.3905 | 250992.3717 | 432547.9034 | 379662.7737 | 329492.8167 | 260531.4353 | 387276.7781 | 215299.6596 |
| 1-[6-Chloro-5-(trifluoromethoxy)-1H-benzimidazol-2-yl]-1H-pyrazole-4-carboxylic acid | 2262610.855 | 2380417.461 | 2229725.946 | 2333696.316 | 2867040.712 | 2329884.03 | 2020333.473 | 2139188.513 | 1930427.657 | 2270079.732 | 2144535.744 | 2254141.437 |
| Asticolorin_B | 99352638.63 | 39919276.75 | 178109259.1 | 56546292.8 | 724619299.5 | 348843351.5 | 545235814.8 | 498092374.3 | 475242227.5 | 349639290.5 | 402025305 | 287333567.6 |
| DGDG O-13:1_28:4 | 325327.2608 | 313529.4275 | 494026.7073 | 345136.4299 | 2491959.728 | 1695964.783 | 1898371.61 | 1656937.672 | 1603276.192 | 1844849.081 | 2451656.32 | 863470.0923 |
| Ganoderic_acid_V | 6005332.559 | 5730468.301 | 4428639.651 | 5399690.571 | 6398152.293 | 5430865.248 | 5448458.513 | 5366693.983 | 5553770.535 | 5342824.145 | 6226645.852 | 3363345.76 |
| Formoterol | 347670.7074 | 446200.8092 | 659328.1347 | 467875.0164 | 837917.7982 | 645104.4638 | 712861.9681 | 760891.5014 | 900365.6039 | 687477.9274 | 733528.962 | 1122054.369 |
| 5-(4-Nitro-1H-pyrazol-1-yl)-1H-tetraazole | 6826079.003 | 1423494.313 | 6605823.704 | 1495774.841 | 7638728.706 | 5928202.738 | 18067147.83 | 18635248.33 | 7426108.949 | 8025490.099 | 15759815.75 | 6143045.789 |

| Metabolites | PBS_1 | PBS_2 | PBS_3 | PBS_4 | BCG_1 | BCG_2 | BCG_3 | BCG_4 | BCGΔ*pe18*_1 | BCGΔ*pe18*_2 | BCGΔ*pe18*_3 | BCGΔ*pe18*_4 |
|---|---|---|---|---|---|---|---|---|---|---|---|---|
| Chicoric acid | 590759.8131 | 380391.9412 | 831980.2908 | 578199.4521 | 2637093.193 | 1601994.166 | 2163428.151 | 1756670.197 | 2445238.969 | 1954412.693 | 2688726.559 | 2081426.307 |
| TG O-16:4_16:4_16:4 | 504161.3757 | 862598.478 | 605210.7383 | 853482.8117 | 724209.2106 | 1720185.713 | 947469.7557 | 1359322.148 | 480141.659 | 872942.5193 | 504397.1591 | 1000810.835 |
| 2-Ketocaproic acid | 49166107.3 | 32297609.34 | 81815478.44 | 14211203.85 | 15056853.27 | 18693184.18 | 68481034.26 | 67389042 | 27211585.63 | 19030985.36 | 17293606.46 | 37293643.1 |
| PS(22:4(7Z,10Z,13Z,16Z)/22:2(13Z,16Z)) | 2233734.591 | 2931585.811 | 1356373.097 | 2137294.383 | 142264.9968 | 134580.832 | 492606.7797 | 195818.9953 | 216422.0043 | 512160.02 | 294412.6855 | 172344.7548 |
| Val-Ala | 385417.2511 | 1190302.815 | 2938829.749 | 1508170.636 | 10168657.12 | 258513.4682 | 1633860.768 | 1536511.437 | 911995.1542 | 345404.7843 | 411401.5513 | 9333761.882 |
| Methyl dodecadienoate | 1981151.39 | 2589437.39 | 1711122.017 | 2451182.941 | 1928913.687 | 2184926.535 | 1924094.129 | 2049751.356 | 1976234.954 | 2077959.38 | 2688919.113 | 1927288.416 |
| 5-(4-Methoxyphenyl)pentanoic acid | 1872219.342 | 1976818.142 | 2102084.638 | 2116150.922 | 255662.8732 | 287922.5416 | 2224649.957 | 1936886.437 | 1705958.209 | 1674328.776 | 1536619.963 | 1794946.825 |
| 4-Piperidinamine, 1-(9H-purin-6-yl)- | 1422270.956 | 1447096.323 | 1375837.281 | 1338634.442 | 1307130.625 | 1301765.355 | 1322537.908 | 1301365.963 | 1305152.52 | 1253206.461 | 1330376.229 | 1479808.066 |
| N-Acetyldihydrosphingosine | 4042567.418 | 4070137.279 | 2905600.456 | 3415467.998 | 3150380.862 | 4550156.852 | 3186039.516 | 2970944.588 | 3210132.925 | 3291636.771 | 3967035.672 | 3265302.126 |
| .alpha.-Piperidinobutiophenone | 2585433.052 | 2411853.095 | 1876069.183 | 2243797.096 | 2494676.496 | 2389810.929 | 2481933.539 | 2394188.144 | 2494806.144 | 442591.581 | 2346112.788 | 1833759.655 |
| Cyclocytidine | 132455136.3 | 92260073.86 | 158807594.7 | 100713361.4 | 252577559.9 | 163661995.2 | 204168208.4 | 200279998.7 | 195358778.8 | 173223997 | 168842266.2 | 133035697.9 |

| Metabolites | PBS_1 | PBS_2 | PBS_3 | PBS_4 | BCG_1 | BCG_2 | BCG_3 | BCG_4 | BCGΔ*pe18*_1 | BCGΔ*pe18*_2 | BCGΔ*pe18*_3 | BCGΔ*pe18*_4 |
|---|---|---|---|---|---|---|---|---|---|---|---|---|
| PC(P-18:1(9Z)/0:0) | 15740.41569 | 592043.9626 | 578342.4974 | 561681.7136 | 0 | 428222.0781 | 357861.4698 | 0 | 0 | 475790.2308 | 339403.9293 | 300847.1574 |
| Lysine conjugated deoxycholic acid putative | 1055413.904 | 1185648.187 | 923919.2993 | 978108.4179 | 1130195.544 | 1003505.306 | 1079597.66 | 1164479.268 | 1096393.071 | 1025918.743 | 976834.3936 | 997112.3018 |
| PE O-17:1_20:4 | 1132515.069 | 972178.707 | 1137217.071 | 951811.3804 | 1233855.156 | 4900.895508 | 1547540.412 | 1934045.626 | 1716805.066 | 1115805.349 | 730318.3838 | 2441672.439 |
| MGDG O-26:5_3:0 | 1412953.062 | 1459560.97 | 1483869.767 | 1492551.502 | 1843442.291 | 1684566.826 | 1481177.099 | 1273968.915 | 1626634.37 | 1596207.488 | 1425677.044 | 1335634.715 |
| 5-Methyl-2-(4-methylphenyl)-1,3-benzoxazole | 2952175.865 | 2991881.057 | 2556544.544 | 2693929.482 | 2385011.201 | 2322321.202 | 2459197.633 | 2266278.355 | 2399965.609 | 2344448.135 | 2371841.385 | 2514282.917 |
| LPE 20:5 | 828103.129 | 685615.3736 | 655994.847 | 734015.2688 | 2516109.732 | 1977233.162 | 1405141.938 | 1717818.252 | 1871174.717 | 1721202.122 | 1566196.964 | 2283435.717 |
| 2-PHENYLBUTYRIC ACID | 8854971.619 | 9667985.675 | 9145412.512 | 9028810.615 | 9966893.283 | 10526999.43 | 9821815.59 | 9370363.909 | 9451790.508 | 9477423.179 | 9167672.308 | 9541057.759 |
| Ferrioxamine | 865168.6282 | 841977.6758 | 859696.2463 | 706825.4884 | 751337.4307 | 901759.644 | 712563.0711 | 864158.7912 | 760005.0802 | 759082.2721 | 932589.6039 | 614684.6587 |
| Pyrazophos | 572021.1807 | 539143.4978 | 950551.7295 | 634106.5286 | 534766.1075 | 892316.4409 | 601242.9174 | 574204.5419 | 638044.5567 | 524711.6558 | 571381.6378 | 503306.7481 |
| Suberosin | 3326875.262 | 3295722.179 | 3225048.484 | 3456610.673 | 3313079.979 | 3410208.867 | 3517720.367 | 3477662.405 | 2987446.586 | 2925552.906 | 3356064.465 | 2596636.388 |
| Myristoylcarnitine | 14771904.78 | 9478415.366 | 16266365.76 | 12448803.85 | 51908147.35 | 31272263.32 | 48936870.52 | 10404847.76 | 23816436.23 | 201697.9842 | 41489685.18 | 19467191.77 |

| **Metabolites** | **PBS_1** | **PBS_2** | **PBS_3** | **PBS_4** | **BCG_1** | **BCG_2** | **BCG_3** | **BCG_4** | **BCGΔ*pe18*_1** | **BCGΔ*pe18*_2** | **BCGΔ*pe18*_3** | **BCGΔ*pe18*_4** |
|---|---|---|---|---|---|---|---|---|---|---|---|---|
| 3,4'-Dimethoxy-2'-hydroxychalcone | 37858382.91 | 43272319.05 | 85791034.77 | 56668843.67 | 7865384.722 | 35575669.19 | 10687251.35 | 2522748.493 | 34959674.18 | 33141764.72 | 34465253.82 | 89747641.78 |
| His-Tyr | 824154.3438 | 1153908.093 | 1454024.641 | 1114576.711 | 2464888.292 | 1730151.104 | 1684865.916 | 1812250.304 | 1776270.558 | 1367474.925 | 873596.4184 | 1886251.369 |
| 2-Methylbutyrylglycine | 1565397.746 | 4441054.225 | 1127410.999 | 5529055.988 | 2647262.965 | 2638687.816 | 1310122.306 | 456514.8497 | 1326041.471 | 1483623.059 | 1368397.591 | 1571834.745 |
| GLYCEROL-MYRISTATE | 2286309.158 | 2713317.42 | 2004737.721 | 3451249.941 | 3778970.124 | 2385490.857 | 1682421.546 | 3702203.641 | 1995703.779 | 1561852.801 | 2229274.746 | 2414207.463 |
| 1-Hydroxy-2-naphthoic acid | 2052006.539 | 1612351.558 | 8649187.65 | 8002309.339 | 28105222.91 | 135598254.7 | 115322613.6 | 46472700.19 | 24497596.73 | 8328590.671 | 93207051.45 | 76354426.97 |
| 4,6'-Anhydrooxysporidinone | 151527.5447 | 174405.9957 | 134449.2271 | 148379.0794 | 160652.4876 | 149444.8676 | 139425.9744 | 176182.4753 | 180768.0815 | 171643.6411 | 167894.5432 | 176550.5074 |
| Flavoglaucin | 16720467.77 | 11440225.03 | 4898190.898 | 6188336.115 | 7430611.14 | 7180767.411 | 5508626.96 | 10139317.52 | 5152288.289 | 4106870.29 | 5664924.004 | 6716283.57 |
| PS 18:0_18:0 | 882419.3214 | 411191.0424 | 646770.3997 | 366800.7585 | 1594552.722 | 1823636.433 | 1509054.208 | 1717868.317 | 905120.8966 | 868499.6996 | 1485303.355 | 837437.4082 |
| Tryptophan | 14237250.07 | 13613747.32 | 31771377.08 | 26830454.66 | 27846978.85 | 37341323.71 | 28631860.75 | 27253257.22 | 26032265.88 | 16672103.31 | 20083565.81 | 20773545.28 |
| PI O-12:0_26:4 | 1804133.513 | 1578838.301 | 1418469.241 | 1359919.769 | 1188237.488 | 0 | 1724395.149 | 3387700.505 | 1622690.886 | 1474276.598 | 1601585.73 | 3710332.569 |
| M260T421 | 1241919.526 | 1029563.396 | 1687414.296 | 982267.4783 | 980555.3774 | 573381.0795 | 990764.6708 | 727404.1234 | 564913.8487 | 800145.2039 | 857733.4634 | 616026.7052 |
| LISINOPRIL | 45154.15121 | 76990.4712 | 76966.74772 | 37037.89208 | 308086.2329 | 302070.8422 | 238369.9056 | 210826.907 | 178123.2895 | 112544.2862 | 350657.9443 | 346690.4045 |

| Metabolites | PBS_1 | PBS_2 | PBS_3 | PBS_4 | BCG_1 | BCG_2 | BCG_3 | BCG_4 | BCGΔ*pe18*_1 | BCGΔ*pe18*_2 | BCGΔ*pe18*_3 | BCGΔ*pe18*_4 |
|---|---|---|---|---|---|---|---|---|---|---|---|---|
| 5-Nitro-2-(2-pyridinyl)-1H-benzimidazole | 2504129.091 | 1934170.078 | 2614763.199 | 5405438.201 | 4856462.252 | 2264367.055 | 3055416.554 | 2372719.925 | 2905912.602 | 4510532.951 | 5505585.983 | 2070883.386 |
| 4-Nitroaniline | 752880.5974 | 643402.0512 | 516024.1394 | 634189.2476 | 474333.1226 | 613008.219 | 461750.9559 | 476016.5028 | 489499.3624 | 550220.47 | 507073.0324 | 746506.4752 |
| N-(2-Hydroxy-3-methylbutanoyl)valine | 1113116.901 | 835146.0937 | 1167445.122 | 1217574.476 | 773612.9012 | 783551.5728 | 1280796.896 | 700657.2038 | 867841.3575 | 928294.4774 | 811759.3289 | 1130454.434 |
| Glycochenodeoxycholic acid | 188605.1032 | 192399.9266 | 442060.881 | 246555.078 | 366464.2011 | 389201.018 | 463467.1173 | 343613.5413 | 300145.94 | 258527.0588 | 214514.3566 | 300549.0886 |
| gamma-Tocotrienol | 218463.2628 | 146729.5159 | 110945.0556 | 170337.1549 | 521544.8402 | 479124.0874 | 293280.8402 | 421169.9568 | 343466.8284 | 238028.9027 | 230351.1878 | 331915.1158 |
| Naloxone N-oxide | 1483302.524 | 1519654.018 | 1239360.812 | 1393126.388 | 1492554.149 | 1428830.39 | 1480980.332 | 1439149.584 | 1479213.363 | 1428516.354 | 1345608.201 | 1321591.34 |
| DGCC 9:0_26:7 | 2839421.83 | 3328595.153 | 1760492.981 | 2762274.224 | 5460913.519 | 6190618.094 | 6369718.86 | 5935223.758 | 5245837.57 | 6642458.296 | 5540384.999 | 3667615.714 |
| 1-O-Octadecyl-2-O-methyl-sn-glycerol | 4145996.014 | 4213952.079 | 2602095.043 | 3428222.866 | 3106185.533 | 3091049.365 | 3245274.531 | 3372907.049 | 3566982.062 | 3633242.927 | 3390759.14 | 2401328.362 |
| PS 2:0_18:5 | 3007664.726 | 3343589.38 | 2771569.339 | 2860999.528 | 2912406.768 | 2951794.354 | 2941084.166 | 2878062.005 | 2996424.37 | 2882168.102 | 2990868.613 | 2921497.029 |
| FAHFA 2:0/20:1 | 1845879.12 | 950139.1252 | 430863.7768 | 819948.5771 | 1716214.273 | 2720209.939 | 1993102.512 | 2055770.357 | 998021.7348 | 576329.2237 | 2340295.125 | 1527004.619 |
| DGCC 9:0_28:6 | 319231.9779 | 8846108.027 | 10274440.91 | 8917805.721 | 13345124.06 | 14676062.45 | 13973415.42 | 19730332.52 | 28705846.75 | 18313849.76 | 14084110.98 | 14027525.78 |

| Metabolites | PBS_1 | PBS_2 | PBS_3 | PBS_4 | BCG_1 | BCG_2 | BCG_3 | BCG_4 | BCGΔ*pe18*_1 | BCGΔ*pe18*_2 | BCGΔ*pe18*_3 | BCGΔ*pe18*_4 |
|---|---|---|---|---|---|---|---|---|---|---|---|---|
| SM 18:2;2O/16:1 | 1908147.731 | 1033938.556 | 1746422.44 | 1386064.923 | 1214357.914 | 308210.8133 | 1081158.091 | 349741.6616 | 1506968.371 | 1307416.429 | 1445724.155 | 574686.4552 |
| LPC O-14:1 | 405281.2649 | 227142.0841 | 398100.7031 | 359931.1248 | 1785662.743 | 1415446.973 | 812588.1536 | 1475196.252 | 1145079.128 | 609030.1753 | 726917.1738 | 515724.1829 |
| (asparaginyl)adenylate | 1296423.523 | 1257369.066 | 1220146.603 | 1312625.9 | 1244243.745 | 1358246.466 | 1263618.944 | 1290110.587 | 1301779.181 | 1405886.85 | 1232319.019 | 1323435.128 |
| PC O-18:5_22:6 | 402093.5749 | 366710.192 | 301541.0529 | 381150.83 | 605777.1258 | 1119408.342 | 658270.8592 | 542077.0357 | 567719.8314 | 587935.2388 | 847842.5388 | 665438.1858 |
| Dextromethorphan | 844665.7362 | 963622.0925 | 759028.6663 | 1033282.754 | 813958.2936 | 943482.482 | 831743.4543 | 847552.2668 | 881950.7754 | 833238.0949 | 743585.1394 | 817034.2913 |
| Plumbagin | 5902303.839 | 5419953.411 | 3721497.869 | 9894306.771 | 18951242.32 | 107059884.8 | 143964690.2 | 49444604.46 | 22949965.34 | 11939253.36 | 105651769.9 | 31855745.02 |
| Tauroursodeoxycholic acid | 8986806.664 | 10806401.67 | 15316999.42 | 8398648.039 | 14979562.37 | 18023580.83 | 22019389.86 | 13529448.37 | 13769025.83 | 8399065.542 | 7828063.372 | 11557974.39 |
| 15-Cyclohexylpentanorprostaglandin F2.alpha. | 888033.7625 | 1442726.742 | 520202.0309 | 1395922.02 | 621765.3244 | 746543.55 | 464784.1565 | 686604.969 | 638849.8744 | 897165.5588 | 526793.3002 | 4421324.204 |
| Glu-Ile | 2760173.407 | 2254404.54 | 3329277.864 | 2273886.726 | 4872583.048 | 6066078.873 | 4894768.056 | 3988443.662 | 4578087.705 | 3048358.839 | 4201636.016 | 3901203.085 |
| cis-Parinaric acid methyl ester | 481667.0092 | 379169.9109 | 497197.7975 | 388782.547 | 627865.4326 | 281119.9222 | 572622.6938 | 502173.0663 | 452759.2861 | 518179.0854 | 434027.1087 | 326716.937 |
| PA(18:3(6Z,9Z,12Z)/15:0) | 4356234.441 | 3650794.796 | 2827955.936 | 3214535.345 | 3686428.386 | 3714162.689 | 13052120.19 | 4224556.313 | 4478730.724 | 3300528.013 | 4260991.208 | 2160350.863 |
| Tenofovir | 108841 | 764675. | 106421 | 652564. | 238133 | 221144 | 265694 | 202362 | 232747 | 239525 | 247413 | 216638 |

| Metabolites | PBS_1 | PBS_2 | PBS_3 | PBS_4 | BCG_1 | BCG_2 | BCG_3 | BCG_4 | BCGΔ*pe18*_1 | BCGΔ*pe18*_2 | BCGΔ*pe18*_3 | BCGΔ*pe18*_4 |
|---|---|---|---|---|---|---|---|---|---|---|---|---|
| | 1.004 | 4923 | 7.714 | 0064 | 1.561 | 9.033 | 6.539 | 0.541 | 3.165 | 4.813 | 0.014 | 7.922 |
| SM 9:1;2O/36:9 | 0 | 0 | 34098.31164 | 17604.14198 | 1479050.75 | 795223.0139 | 646922.4056 | 577760.5976 | 510979.82 | 478080.0361 | 399639.3445 | 694220.8984 |
| Pantethine | 941192.1565 | 290969.5198 | 166565.7862 | 708916.1056 | 1127464.413 | 520051.7854 | 1012798.127 | 287690.7146 | 815760.8515 | 1495693.114 | 740121.5196 | 642841.5292 |
| 1H-Benzotriazole | 630973.3798 | 545653.9982 | 634169.7529 | 583821.2506 | 691073.1903 | 614573.8001 | 540445.7487 | 772908.3647 | 561588.2935 | 585296.5183 | 591020.273 | 614785.5184 |
| 2-(Dicyclohexylphosphino)-2'-methylbiphenyl | 2198380.074 | 11163045.96 | 6279856.676 | 2671181.458 | 4307506.48 | 2998862.437 | 26276315.14 | 5066601.001 | 12555672.7 | 2893166.386 | 12843432.23 | 13909486.43 |
| Folic acid | 1363318.063 | 1185772.739 | 1949789.459 | 1667244.165 | 1435472.011 | 1524526.329 | 1422811.002 | 1393112.816 | 1505906.285 | 755997.6925 | 913961.8818 | 1704937.251 |
| Oxooctanoylcarnitine (Car(8:1-O)) | 3595627.558 | 3937858.575 | 3937053.859 | 4168444.887 | 3607782.561 | 3882391.062 | 4099181.993 | 3871583.532 | 3988055.153 | 3087097.411 | 4956917.199 | 4311372.974 |
| Acetylproline | 5453146.835 | 1490813.309 | 2424383.973 | 2143779.503 | 1985434.917 | 2404296.741 | 2242738.852 | 1760434.672 | 2703646.436 | 1557427.846 | 2836155.222 | 3480092.822 |
| Glabrin_C | 738622.0454 | 1035641.277 | 670276.9539 | 846154.4274 | 1317657.509 | 1102845.493 | 2164062.134 | 1654601.582 | 1399367.536 | 1744469.32 | 1225794.313 | 840884.3808 |
| Ethyl (E,Z)-decadienoate | 976384.9197 | 918938.1734 | 1461077.382 | 1035406.701 | 865201.9781 | 866070.4581 | 944032.6373 | 889248.7396 | 912620.0507 | 930748.6222 | 943472.4541 | 1000061.756 |
| PI O-14:1_26:7 | 313437.5756 | 494387.7173 | 283087.7452 | 275823.8111 | 257218.6325 | 520847.035 | 276189.2493 | 298559.972 | 233778.9783 | 261590.1337 | 277379.1677 | 290698.8991 |

| Metabolites | PBS_1 | PBS_2 | PBS_3 | PBS_4 | BCG_1 | BCG_2 | BCG_3 | BCG_4 | BCGΔ*pe18*_1 | BCGΔ*pe18*_2 | BCGΔ*pe18*_3 | BCGΔ*pe18*_4 |
|---|---|---|---|---|---|---|---|---|---|---|---|---|
| 6-(6-Aminohexanamido)hexanoic acid | 1378170.709 | 1486917.03 | 1343720.288 | 1493131.029 | 1527681.628 | 1473049.641 | 1450947.044 | 1349501.791 | 1415052.248 | 1391560.871 | 1351368.963 | 1739000.39 |
| 1-Amino(cyclohexyl)acetic acid ethyl ester | 2646853.982 | 3068450.705 | 2094061.66 | 6571021.536 | 1845514.256 | 2437612.706 | 5436394.324 | 1492122.73 | 3456499.089 | 3158098.906 | 1633218.823 | 6223467.835 |
| PG 17:2_22:6 | 327749.0972 | 345566.4498 | 92158.14372 | 268503.7486 | 439120.5363 | 235034.4453 | 453880.9032 | 400293.1587 | 692287.0682 | 838671.2789 | 410589.1366 | 302187.1683 |
| DGDG O-9:0_2:0 | 14376.95979 | 11180.00206 | 8435.041602 | 22875.08682 | 242528.1856 | 141732.7457 | 310320.9419 | 186174.6724 | 193357.8598 | 167615.889 | 205603.7287 | 94683.40646 |
| SMGDG O-13:1_28:7 | 0 | 0 | 0 | 0 | 224809.0776 | 121122.9948 | 143781.4446 | 156335.3959 | 216504.4887 | 241828.7478 | 186129.5512 | 101081.6415 |
| Aurantiamine | 10484946.63 | 8832469.434 | 6425827.709 | 11268057.07 | 11582450.79 | 14179678.05 | 10451599.11 | 10010898.69 | 10623600.83 | 9363349.969 | 9100894.629 | 9239206.669 |
| Benzenepropanoic acid, 4-[2-(2-methylphenyl)ethynyl]- | 1224040.207 | 922025.7418 | 7926824.202 | 8047007.299 | 1652600.161 | 18511875.57 | 7980938.205 | 7865367.834 | 11672612.58 | 1465798.673 | 7636768.589 | 11565776.29 |
| 6-Chloro-3-(trifluoromethyl)-2-pyridinecarbonitrile | 1518646.711 | 1314698.602 | 1630717.99 | 1525590.385 | 2174593.143 | 1800342.303 | 2229022.802 | 1932837.805 | 1806491.901 | 1927422.834 | 1971190.932 | 1604292.324 |
| Dodecylamine | 2631831.586 | 2813902.225 | 2607845.09 | 2847304.142 | 2808847.772 | 2817723.962 | 2736887.192 | 3017633.332 | 2898518.139 | 4692515.889 | 2862787.249 | 3018999.905 |
| MeOSuc-Ala-Ala-P | 477014. | 532514. | 337619. | 405728. | 377652. | 361750. | 432069. | 414258. | 369315. | 430678. | 395763. | 380354. |

| Metabolites | PBS_1 | PBS_2 | PBS_3 | PBS_4 | BCG_1 | BCG_2 | BCG_3 | BCG_4 | BCGΔ *pe18*_1 | BCGΔ *pe18*_2 | BCGΔ *pe18*_3 | BCGΔ*pe18*_4 |
|---|---|---|---|---|---|---|---|---|---|---|---|---|
| ro-Val-AMC | 7774 | 0068 | 6045 | 9737 | 588 | 6653 | 4177 | 8524 | 0679 | 8717 | 0663 | 7756 |
| HexCer 16:3;2O/14:0 | 1231762.824 | 1322062.513 | 976503.9275 | 1380094.518 | 1670503.095 | 812862.339 | 1085988.345 | 1035647.742 | 1602608.272 | 1377983.531 | 1023467.624 | 1059768.377 |
| 6-Hydroxy-1-((2-O-((2E)-3-(4-hydroxyphenyl)prop-2-enoyl)hexopyranosyl)oxy)-7-methylidene-1,4a,5,6,7,7a-hexahydrocyclopenta[c]pyran-4-carboxylic acid | 2237004.644 | 3683949.108 | 6158498.756 | 4614284.421 | 2022882.143 | 2474876.85 | 3767128.729 | 1649249.252 | 2763565.194 | 1853704.193 | 2544360.519 | 7378986.409 |
| 4-Benzyl-7-methoxy-3-phenylcoumarin | 2648381.223 | 4218651.779 | 4013305.16 | 4272833.949 | 1794792.973 | 2293151.51 | 1768728.824 | 1644688.666 | 1559373.864 | 1602884.251 | 1874237.308 | 5481626.649 |
| DGDG O-26:7_3:0 | 703862.2943 | 401806.8689 | 631664.2762 | 563325.7585 | 664519.8687 | 18174.499 | 690624.834 | 574444.8899 | 680052.7988 | 658606.3447 | 659067.9946 | 521965.0387 |
| (2-Acetyloxy-3-hydroxypropyl) octadec-9-enoate | 264019.1574 | 203271.3954 | 333736.7103 | 291115.2226 | 691307.1793 | 511028.8691 | 845030.5921 | 626579.6619 | 360274.2229 | 388134.0585 | 676054.9095 | 332986.4726 |
| M497T426 | 681049.4194 | 409318.8479 | 499595.2074 | 662398.4825 | 380307.1658 | 835437.794 | 434034.2367 | 440434.2909 | 719586.7157 | 352811.4229 | 730853.348 | 362409.9524 |
| 1-Methylpyrrolinium [M] | 3554943.226 | 3753782.388 | 4076699.373 | 4374542.812 | 4027284.66 | 4034114.087 | 3405566.312 | 8655981.996 | 3828924.049 | 3047355.609 | 3780685.245 | 4102632.626 |

| Metabolites | PBS_1 | PBS_2 | PBS_3 | PBS_4 | BCG_1 | BCG_2 | BCG_3 | BCG_4 | BCGΔ*pe18*_1 | BCGΔ*pe18*_2 | BCGΔ*pe18*_3 | BCGΔ*pe18*_4 |
|---|---|---|---|---|---|---|---|---|---|---|---|---|
| Oxo(propan-2-ylamino)acetic acid | 1657611.949 | 1456305.287 | 1834854.203 | 1465556.642 | 6463107.683 | 4394650.862 | 5434530.977 | 5715564.895 | 5178233.928 | 4724793.403 | 5010541.265 | 4115500.609 |
| ST 24:1;O4;T/22:6 | 329648.2443 | 292707.4384 | 157932.8824 | 245114.6923 | 1048621.426 | 839584.895 | 1134949.926 | 955356.4953 | 1159806.06 | 1128739.708 | 849243.3473 | 630277.0672 |
| Citrinin | 1073837.637 | 1117972.78 | 1141517.96 | 1087201.618 | 1286446.961 | 1225311.851 | 1024473.352 | 1108529.165 | 1149452.036 | 1140482.428 | 1256845.216 | 986906.6486 |
| PFOH-perfluoroalkyl_alcohol | 427211.1349 | 384542.0601 | 422854.5268 | 407066.2453 | 699351.0464 | 279410.6527 | 678882.9884 | 268919.7405 | 381706.4608 | 609576.0472 | 375573.3263 | 488269.4013 |
| (1S,2S,3R,4R)-4-Amino-1,2,3-cyclopentanetriol | 1684614.519 | 2072344.125 | 3071651.714 | 3168945.228 | 3612481.868 | 3692919.891 | 2011262.875 | 2425082.59 | 2184318.743 | 1358123.116 | 2101910.292 | 2474812.791 |
| 3,6-Diamino-9-(2-carboxyphenyl)xanthylium cation | 0 | 0 | 0 | 0 | 1593822.415 | 1342707.653 | 1454638.867 | 1113962.57 | 1344984.991 | 974729.4127 | 1595969.399 | 1043376.784 |
| Cer 18:1;2O/8:0 | 2929180.894 | 4007118.138 | 1420547.972 | 2066339.018 | 1110847.016 | 3331249.364 | 1191289.34 | 1501282.861 | 1233833.399 | 1017992.403 | 1027815.891 | 932209.0422 |
| 1-Formylpyrrolidine-2-carboxylic acid | 15964574.48 | 7359443.859 | 12096276.81 | 12379974.68 | 8308201.005 | 10423710.09 | 10979160.2 | 13946144.76 | 12654301.59 | 11510915.47 | 9081527.251 | 10033970.62 |
| MG 15:0 | 8326420.3 | 8517939.692 | 8116978.387 | 8451888.357 | 8312246.574 | 8709457.308 | 9385819.931 | 8590771.564 | 8678549.507 | 8860931.8 | 8055211.586 | 7849041.571 |

| Metabolites | PBS_1 | PBS_2 | PBS_3 | PBS_4 | BCG_1 | BCG_2 | BCG_3 | BCG_4 | BCGΔ *pe18*_1 | BCGΔ *pe18*_2 | BCGΔ *pe18*_3 | BCGΔ*pe18*_4 |
|---|---|---|---|---|---|---|---|---|---|---|---|---|
| (1aS,1bS,2S,5aR,6S,6aS)-6-((6-Deoxy-3-O-((2E)-3-(4-hydroxy-3-methoxyphenyl)prop-2-enoyl)-.alpha.-L-mannopyranosyl)oxy)-1a-(hydroxymethyl)-1a,1b,2,5a,6,6a-hexahydrooxireno[4,5]cyclopenta[1,2-c]pyran-2-yl .beta.-D-glucopyranoside | 0 | 0 | 0 | 0 | 13019328.19 | 10951967.09 | 7650339.334 | 7917787.743 | 13629126.52 | 8970651.15 | 9471640.998 | 13743295.58 |
| N-ALPHA-ACETYL-L-ASPARAGINE | 206222.2347 | 241434.1057 | 2322935.46 | 308796.2203 | 850247.3517 | 541360.7851 | 4139584.824 | 730743.3045 | 615732.9541 | 2654865.593 | 2822539.356 | 997555.2216 |
| Diethyl azelate | 5304116.776 | 6127311.414 | 5203055.689 | 5844182.822 | 5355422.288 | 5352111.174 | 5379979.683 | 5354415.647 | 5335704.527 | 5391985.97 | 5573113.428 | 5302393.569 |
| 3-tert-Butyl-1-(4-methylphenyl)-1H-pyrazol-5-ylamine | 77552.1832 | 944399.9368 | 1002587.549 | 1580404.167 | 84865.91906 | 47338.81998 | 87546.86291 | 43123.23361 | 80924.27205 | 55669.92137 | 43594.88862 | 3771462.591 |
| 4-(Thiophen-2-yl)-6-(trifluoromethyl)pyrimidin-2-ol | 1191751.078 | 806051.7691 | 1161232.965 | 933986.8058 | 2417999.761 | 2055538.828 | 3939722.947 | 2558104.589 | 2157893.697 | 1588517.557 | 1714471.657 | 831930.3808 |

| Metabolites | PBS_1 | PBS_2 | PBS_3 | PBS_4 | BCG_1 | BCG_2 | BCG_3 | BCG_4 | BCGΔ*pe18*_1 | BCGΔ*pe18*_2 | BCGΔ*pe18*_3 | BCGΔ*pe18*_4 |
|---|---|---|---|---|---|---|---|---|---|---|---|---|
| M649T444_2 | 1192548.231 | 1200602.273 | 908592.1935 | 1146726.709 | 1272694.802 | 1155713.482 | 1157365.764 | 1195272.949 | 1223330.506 | 1174083.51 | 1055730.555 | 43122.51215 |
| 11-(Dansylamino)undecanoic acid | 4335331.198 | 5153538.649 | 4000230.243 | 4335001.443 | 5452725.312 | 5013400.641 | 5311686.597 | 4775760.796 | 4610327.849 | 4967920.52 | 4536080.018 | 4350174.815 |
| Trideca-3,5-dienoyl carnitine | 1501964.916 | 1763579.146 | 1094127.237 | 1643213.022 | 1734078.799 | 1513453.078 | 1651122.822 | 1846798.174 | 1752814.269 | 1819629.944 | 1577536.059 | 1610851.814 |
| Cyproterone | 213722.9929 | 1678656.737 | 2605931.471 | 1274482.247 | 128037.2703 | 117910.8675 | 227490.7756 | 92770.19898 | 165151.7252 | 131226.1388 | 296943.5694 | 10691327.22 |
| PC(18:1(6Z)/18:1(6Z)) | 941344703 | 1234966723 | 500747256.7 | 714596905.1 | 213720200.9 | 180616572.3 | 282220283.3 | 244272138.2 | 294029579.7 | 335947924.2 | 241886390.7 | 161995009.7 |
| DG 19:0 | 29486786 | 31520999.68 | 6268735.023 | 7262389.434 | 28572783.14 | 28924605.69 | 29205782.78 | 28982047.44 | 7019981.993 | 29220095.32 | 28330719.15 | 6162279.534 |
| SHexCer 40:5;3O | 76479192.59 | 59650535.83 | 27411124.18 | 54783457.95 | 14635679.6 | 9364595.463 | 12077463.39 | 12965632.88 | 8794342.424 | 15827985.64 | 11130830.01 | 5635323.022 |
| PE O-20:4_5:0 | 706339.5929 | 922057.6615 | 673829.1993 | 553873.975 | 370301.2614 | 846970.6567 | 549230.866 | 393724.7113 | 384602.732 | 524579.2742 | 422617.7387 | 319247.7198 |
| 4-(1-Pyrrolidinyl)piperidine | 960507.2093 | 1378875.747 | 1034281.975 | 872606.539 | 960316.0234 | 1297129.606 | 943840.3789 | 1172940.689 | 940941.533 | 979029.592 | 1036524.249 | 1087419.653 |
| CAR 23:6 | 85845.69655 | 87748.31398 | 87039.25408 | 101268.9854 | 96687.14613 | 106669.6581 | 85703.68046 | 97028.02204 | 86729.30219 | 85012.22061 | 84442.60067 | 100689.9201 |
| NAE(20:2(11Z,14Z)) | 2376652.137 | 2353838.787 | 969956.2471 | 1424421.898 | 1528625.729 | 1829606.603 | 1572392.976 | 1389305.322 | 1456437.604 | 2166309.901 | 1758951.626 | 1446395.244 |
| Methyl 4-hydroxy-3,5-dinit | 654414.8471 | 709612.1197 | 531622.5904 | 555992.3219 | 2074040.366 | 1355856.985 | 2016639.323 | 1752877.321 | 1856840.536 | 2187040.623 | 1713826.621 | 1490556.683 |

| Metabolites | PBS_1 | PBS_2 | PBS_3 | PBS_4 | BCG_1 | BCG_2 | BCG_3 | BCG_4 | BCGΔ*pe18*_1 | BCGΔ*pe18*_2 | BCGΔ*pe18*_3 | BCGΔ*pe18*_4 |
|---|---|---|---|---|---|---|---|---|---|---|---|---|
| robenzoate | | | | | | | | | | | | |
| PE-Cer 12:2;2O/18:5 | 650567.7496 | 575130.4837 | 543650.2588 | 630431.9439 | 1435193.842 | 1377682.181 | 871340.2415 | 1279863.142 | 976311.4419 | 561204.5902 | 640287.819 | 1075865.55 |
| PS 34:5 | 68786.31999 | 52681.77399 | 54369.24465 | 41086.64572 | 124686.7713 | 77211.76851 | 137164.3802 | 84808.30774 | 67515.81248 | 123864.5336 | 125623.7266 | 55181.77128 |
| LPE O-18:4 | 6618562.528 | 8017348.671 | 5406417.178 | 5479405.514 | 3981530.235 | 6766462.742 | 5089330.094 | 3965032.07 | 3933582.705 | 3892271.007 | 5273752.009 | 2844240.014 |
| STIGMASTA-4,22-DIEN-3-ONE | 185954.6885 | 95810.4046 | 30095.93952 | 141716.5815 | 2436685.164 | 3435521.636 | 1239347.711 | 2196722.371 | 1394216.643 | 468141.1523 | 2224189.499 | 2741427.314 |
| Sph(d18:2) | 19195931.53 | 19848834.13 | 13708343.29 | 17571898.82 | 19751468.83 | 19846114.66 | 17810264.79 | 22347358.77 | 17868663.81 | 13966827.21 | 14229591.65 | 11269188.29 |
| 3-Hydroxybutyrylcarnitine (Car(4:0-O)) | 10536594.91 | 9437293.066 | 13769857.54 | 10808409.21 | 23907861.45 | 17409321.71 | 30757248.73 | 20653922.13 | 15810575.44 | 19415721.87 | 20517344.85 | 12953988.31 |
| [3-Me-His2] TRH | 1046666.001 | 953962.7655 | 949130.9131 | 882322.254 | 854267.7642 | 968891.732 | 909735.149 | 780239.4958 | 916614.1999 | 1055958.553 | 1063923.776 | 861793.4055 |
| trans-11-Eicosenoic acid | 931764.035 | 855715.9845 | 1294641.536 | 852424.7832 | 18031905.24 | 28823027.49 | 18234106.85 | 20245306.15 | 11365287.44 | 7544858.832 | 16659843.18 | 14978034.38 |
| Quinmerac | 3406615.652 | 3172494.881 | 2913865.085 | 3179764.918 | 2891611.033 | 2960146.089 | 2968649.385 | 3157145.444 | 3035674.672 | 2809790.246 | 2781813.349 | 3021579.759 |
| CerP 14:1;2O/24:1 | 1662405.56 | 1484532.058 | 1754333.776 | 1334357.363 | 1792505.473 | 1803919.227 | 2207335.266 | 1818089.369 | 1837945.961 | 1663166.04 | 1863015.582 | 2364729.238 |
| pseudoionone | 964306. | 125166 | 128491 | 105380 | 833342. | 872655. | 108185 | 254111 | 730514. | 108012 | 722014. | 107168 |

| Metabolites | PBS_1 | PBS_2 | PBS_3 | PBS_4 | BCG_1 | BCG_2 | BCG_3 | BCG_4 | BCGΔ *pe18*_1 | BCGΔ *pe18*_2 | BCGΔ *pe18*_3 | BCGΔ*pe18*_4 |
|---|---|---|---|---|---|---|---|---|---|---|---|---|
| | 0296 | 9.006 | 8.681 | 4.476 | 925 | 7757 | 3.368 | 7.929 | 8613 | 0.362 | 1058 | 0.903 |
| Alogliptin | 1866066.768 | 2269701.862 | 1894538.257 | 2081664.943 | 3661645.728 | 1903698.201 | 1977299.145 | 2124052.682 | 1950904.464 | 1841475.942 | 2223269.041 | 1507761.45 |
| LPG 16:1 | 437930.2336 | 259192.8546 | 839748.7606 | 541163.4259 | 5269904.52 | 5013084.323 | 9338698.778 | 7974794.723 | 3020634.549 | 3488550.308 | 4341583.354 | 4050182.667 |
| Methyl 5(S),6(R)-7-trihydroxyheptanoate | 548841.0703 | 474922.7759 | 386397.9712 | 5203018.826 | 131923.5271 | 695206.5506 | 3306457.678 | 97975.79069 | 1778001.088 | 1010348.505 | 121745.7899 | 2519164.84 |
| Oxybutynine | 306892.9445 | 667084.8299 | 1015546.487 | 676958.063 | 311563.2549 | 272936.7787 | 386242.4244 | 240700.0234 | 381124.1955 | 548211.0765 | 432118.3387 | 1687786.278 |
| Ribitol | 854335.5302 | 704609.2875 | 1220004.884 | 866572.5925 | 2879974.494 | 1727137.009 | 3082775.737 | 2243171.408 | 1885184.776 | 2201999.733 | 1883693.2 | 1614610.997 |
| Benzeneoctanol, 3-hydroxy-4-[(1R,3S)-3-hydroxycyclohexyl]-.theta.,.theta.-dimethyl-, rel- | 141339.1006 | 203590.8664 | 293320.3965 | 234186.6233 | 980979.5731 | 1628616.833 | 1094171.966 | 1063055.034 | 1508931.867 | 842841.5383 | 1225085.689 | 1499921.043 |
| Fenarimol | 202216.8042 | 269551.575 | 298921.2193 | 308768.3057 | 316357.2241 | 314701.9239 | 300249.1168 | 293226.7544 | 312882.4757 | 322426.5486 | 317686.1293 | 329453.6931 |
| CMP-N-trimethyl-2-aminoethylphosphonate [M] | 999183.8684 | 807983.1426 | 1201431.082 | 884219.0392 | 2537622.341 | 976560.1213 | 2466524.326 | 1599679.798 | 1349529.618 | 2444266.269 | 2504141.353 | 1214770.736 |
| Pipecolic acid | 440740.8101 | 1445393.795 | 871437.3605 | 682521.1711 | 1451960.762 | 1117660.815 | 1325728.389 | 1156766.075 | 1137878.626 | 1000753.362 | 3547999.645 | 1131518.544 |

| Metabolites | PBS_1 | PBS_2 | PBS_3 | PBS_4 | BCG_1 | BCG_2 | BCG_3 | BCG_4 | BCGΔ *pe18*_1 | BCGΔ *pe18*_2 | BCGΔ *pe18*_3 | BCGΔ*pe18*_4 |
|---|---|---|---|---|---|---|---|---|---|---|---|---|
| Thiamine pyrophosphate (TPP) | 181125.0024 | 166716.715 | 251372.1878 | 158124.3509 | 737450.6144 | 423743.4802 | 733538.4174 | 397322.4273 | 827152.73 | 568718.1485 | 604356.1127 | 572091.0492 |
| 4-Chloro-3-(6,7,8,9-tetrahydro-5H-[1,2,4]triazolo[4,3-a]azepin-3-yl)aniline | 483921.6382 | 570866.9844 | 1244711.111 | 771288.5986 | 654360.0663 | 828721.4954 | 597458.2257 | 649803.8973 | 461816.9097 | 263034.8885 | 521396.3929 | 696931.0861 |
| Di(2-nonyl) phthalate | 4621609.007 | 9733946.465 | 3830527.468 | 6218020.677 | 5808799.736 | 16141694.06 | 4630710.79 | 5122394.019 | 3946450.664 | 5470307.898 | 4150819.354 | 5722252.89 |
| Lactupicrin | 817999.8027 | 1422746.355 | 1279100.812 | 1090130.264 | 575140.8185 | 677592.6822 | 613523.6395 | 527699.9372 | 584913.0785 | 513788.7164 | 652797.5779 | 1975213.084 |
| PC O-13:0_26:2 | 534682.935 | 433830.4527 | 674127.6375 | 731397.6862 | 387868.5059 | 492237.6816 | 442637.6952 | 440155.7492 | 530477.918 | 426877.3011 | 12871.56855 | 709614.4525 |
| PC(18:3(9Z,12Z,15Z)/14:0) | 1803796.934 | 5646608.414 | 7494568.305 | 1130485.104 | 24584417.7 | 4875472.573 | 30824242.58 | 23804236.11 | 5434295.881 | 27411017.75 | 5049722.091 | 16555521.55 |
| 2-[(2R,3S,7R,8R,8aS)-2,3,4'-Trihydroxy-4,4,7,8a-tetramethyl-6'-oxospiro[2,3,4a,5,6,7-hexahydro-1H-naphthalene-8,2'-3,8-dihydrofuro[2,3-e]isoindole]-7'-yl]-3-methylpentanoic | 5438573.528 | 6194727.372 | 8738946.037 | 6164776.765 | 6733860.021 | 6638845.252 | 5956723.763 | 6140050.911 | 6984600.327 | 6649628.683 | 7098701.727 | 6714011.044 |

| Metabolites | PBS_1 | PBS_2 | PBS_3 | PBS_4 | BCG_1 | BCG_2 | BCG_3 | BCG_4 | BCGΔ*pe18*_1 | BCGΔ*pe18*_2 | BCGΔ*pe18*_3 | BCGΔ*pe18*_4 |
|---|---|---|---|---|---|---|---|---|---|---|---|---|
| acid | | | | | | | | | | | | |
| Glycidyl oleate | 4343794.168 | 5355825.981 | 3157684.847 | 3722064.486 | 19986639.58 | 12882449.57 | 21741830.22 | 14958120.78 | 16940480.9 | 26353729.26 | 21684271.89 | 11529046.79 |
| 2-Bromohypoxanthine | 692415.1359 | 860678.5664 | 523142.7789 | 681628.0042 | 627192.3024 | 601521.7196 | 506019.4334 | 472075.237 | 587211.7202 | 697597.547 | 516439.9172 | 567625.7392 |
| 3-Hydroxypyruvic acid | 3541349.399 | 3694496.594 | 5727361.725 | 6917198.996 | 5463140.83 | 5403920.347 | 12384927.23 | 10617475.91 | 4776787.976 | 5320126.518 | 10639237.32 | 5339423.973 |
| MGDG O-20:5_18:1 | 344589.2128 | 394159.2527 | 359200.5745 | 386447.0039 | 67163.05115 | 117128.164 | 297388.0418 | 211671.8342 | 410994.9943 | 82999.04398 | 657817.0588 | 471510.1287 |
| 1-Methyl-4-(p-tolylsulfonyl)piperazine | 1650905.974 | 1826019.901 | 1862548.176 | 1786226.373 | 1617872.006 | 1950273.326 | 1721374.111 | 1689298.287 | 1737859.482 | 1590421.791 | 1799230.948 | 2044330.315 |
| 1-[4-Hydroxy-3-(3-methylbut-2-enyl)phenyl]ethanone | 1294521.064 | 1341626.397 | 1157263.078 | 1171570.367 | 1317371.527 | 1272276.36 | 1132955.767 | 1272486.646 | 1217319.647 | 1226639.546 | 1255770.68 | 1113022.532 |
| Allopurinol | 482299.6475 | 447196.4185 | 2862490.309 | 484254.0864 | 3311597.08 | 2651104.075 | 5166625.286 | 3479507.701 | 2126999.696 | 1655408.711 | 3272932 | 1354667.327 |
| N-(4-Nitrobenzoyl)phenylalanine | 3781519.834 | 3295231.668 | 5401668.789 | 2950001.271 | 3318525.55 | 3304781.163 | 5529248.71 | 3815560.881 | 3436370.26 | 4525757.332 | 3026932.34 | 3252781.634 |

| Metabolites | PBS_1 | PBS_2 | PBS_3 | PBS_4 | BCG_1 | BCG_2 | BCG_3 | BCG_4 | BCGΔ*pe18*_1 | BCGΔ*pe18*_2 | BCGΔ*pe18*_3 | BCGΔ*pe18*_4 |
|---|---|---|---|---|---|---|---|---|---|---|---|---|
| 3-Methyl-2-oxovaleric acid | 49166107.3 | 32297609.34 | 81815478.44 | 14211203.85 | 15056853.27 | 18693184.18 | 68481034.26 | 67389042 | 27211585.63 | 19030985.36 | 17293606.46 | 37293643.1 |
| Adenosine 3',5'-cyclic phosphate (cAMP) | 133032.3017 | 126172.9382 | 215425.5102 | 282517.8788 | 2451188.681 | 2180034.986 | 2497620.802 | 2483095.469 | 1098384.36 | 1270865.957 | 1247375.109 | 1451103.579 |
| L-3-Aminobutyric acid | 653165.6996 | 621348.5185 | 755684.591 | 608777.5012 | 4800438.133 | 3225445.945 | 4542151.7 | 3873767.325 | 4183680.943 | 3544407.742 | 3602633.771 | 3644172.971 |
| 1-Palmitoyllysophosphatidate | 153054.0495 | 167981.1302 | 90834.47657 | 144151.546 | 609851.1385 | 558264.7644 | 494493.301 | 537680.5261 | 469407.4697 | 500337.9693 | 565544.2688 | 327553.1057 |
| Futalosine | 0 | 0 | 0 | 0 | 283420.6268 | 171932.9471 | 201719.4554 | 164959.7127 | 281236.9653 | 360551.9629 | 320092.6459 | 260814.8795 |
| 1,6-Anhydro-.beta.-D-glucose | 1905746.368 | 1513533.153 | 1914306.202 | 1765727.94 | 1769503.143 | 2738164.676 | 2133364.897 | 1268531.185 | 1976004.716 | 1335726.715 | 1827051.818 | 1886793.768 |
| Ethyl 4,6-dichloro-1H-indole-2-carboxylate | 1263559.056 | 1276661.121 | 1400104.35 | 1356520.136 | 1157361.414 | 1289575.89 | 1210508.271 | 1257205.365 | 1263303.287 | 1122296.527 | 1266966.485 | 1269557.428 |
| Guanidinosuccinic acid | 7052858.948 | 5386841.645 | 9105501.298 | 6937765.934 | 6833891.096 | 7831663.849 | 6520242.466 | 7044323.954 | 7886984.015 | 4759240.784 | 5499243.174 | 7105803.985 |
| 2-Methylthio-adenosine-5'-diphosphate | 11288597.85 | 2591255.721 | 7123932.155 | 5178772.652 | 4243603.114 | 2992298.999 | 2329356.781 | 5307446.691 | 4128978.817 | 2573331.183 | 2796973.831 | 2336257.076 |
| SMGDG O-24:4_13:0 | 1779549.62 | 2077124.508 | 1383662.63 | 1396740.007 | 2238689.286 | 1978480.554 | 2919796.209 | 2073333.738 | 2839907.168 | 3510458.941 | 1727499.856 | 2029269.637 |
| Lys-Pro | 2257007.554 | 15686751.14 | 833500.2879 | 15179236.77 | 21003708.59 | 11756937.96 | 639313.1601 | 801780.8956 | 3194306.741 | 10889213.69 | 6330144.354 | 1983596.162 |

| **Metabolites** | **PBS_1** | **PBS_2** | **PBS_3** | **PBS_4** | **BCG_1** | **BCG_2** | **BCG_3** | **BCG_4** | **BCGΔ*pe18*_1** | **BCGΔ*pe18*_2** | **BCGΔ*pe18*_3** | **BCGΔ*pe18*_4** |
|---|---|---|---|---|---|---|---|---|---|---|---|---|
| LPC(13:0) | 1422855.599 | 1255442.239 | 1094616.118 | 1210385.157 | 4550876.332 | 3484747.356 | 4431586.623 | 4188037.345 | 6322666.744 | 5977159.767 | 5485174.808 | 3887141.924 |
| PC(20:5(5Z,8Z,11Z,14Z,17Z)/22:6(4Z,7Z,10Z,13Z,16Z,19Z)) | 9433157.265 | 12326400.78 | 5420061.028 | 7791192.483 | 6189678.339 | 4812080.494 | 5892966.223 | 6802096.131 | 7258875.526 | 7198239.545 | 4786310.785 | 4476405.405 |
| N-(1-Deoxy-1-fructosyl)histidine | 35821187.13 | 26283073.08 | 48831925.2 | 30627849.09 | 147190196.4 | 72648598 | 107978864 | 94840124.19 | 93647248.02 | 74465218.4 | 78067693.56 | 63380273.96 |
| NAGly 19:2/17:2 | 662750.5425 | 773115.9917 | 503241.2165 | 612619.4543 | 1982203.813 | 1628450.501 | 789357.2461 | 1479332.219 | 1037798.616 | 426973.5547 | 476495.3491 | 1554379.669 |
| Methadone | 2427073.798 | 2514237.693 | 2045351.765 | 2202441.601 | 2300364.777 | 1849149.18 | 4103559.516 | 1977673.086 | 2477767.24 | 2904339.165 | 2070508.418 | 2061139.495 |
| Timonacic | 6652835.535 | 1386246.945 | 9562393.999 | 2637930.41 | 12928412.85 | 17103217.43 | 62506255.81 | 55216252.21 | 25392616.3 | 10913609.36 | 25433732.45 | 17637865.96 |
| N'-(2-Oxo-1,2-dihydro-3H-indol-3-ylidene)isonicotinohydrazide | 770019.4966 | 208692.1083 | 2543077.601 | 688484.173 | 3343799.072 | 12484401.11 | 14056677.13 | 14594210.02 | 28056689.68 | 7407264.638 | 35565592.08 | 61058222.62 |
| PE(22:6(4Z,7Z,10Z,13Z,16Z,19Z)/16:1(9Z)) | 4650867.152 | 6091631.663 | 6170285.275 | 5900551.903 | 16808251.78 | 12903457.37 | 20888213.75 | 15626925.54 | 15888188.34 | 18614776.89 | 17427226.24 | 11307225.68 |
| M463T523 | 6213902.177 | 7271655.474 | 13221601.96 | 9683024.59 | 15476958 | 14412254.45 | 7821788.552 | 12326693.83 | 7936053.544 | 4644003.419 | 5377389.363 | 8127734.673 |
| SL 10:0;O/18:3 | 430837 | 582416 | 278930 | 377806 | 104494 | 115189 | 139253 | 117173 | 131342 | 141728 | 133783 | 745976. |

| Metabolites | PBS_1 | PBS_2 | PBS_3 | PBS_4 | BCG_1 | BCG_2 | BCG_3 | BCG_4 | BCGΔ *pe18*_1 | BCGΔ *pe18*_2 | BCGΔ *pe18*_3 | BCGΔ*pe18*_4 |
|---|---|---|---|---|---|---|---|---|---|---|---|---|
| | 8.565 | 2.522 | 7 | 4.428 | 3.06 | 6.8 | 7.671 | 2.705 | 2.724 | 5.279 | 0.629 | 9231 |
| TG 8:0_8:0_22:6 | 742327.0509 | 706540.5521 | 209633.3829 | 778284.6449 | 454897.206 | 490438.7664 | 561736.1167 | 459428.9648 | 384014.255 | 805003.8195 | 487245.6066 | 332681.1529 |
| Decamethylcyclopentasiloxane | 642295.9167 | 866021.7717 | 967215.8746 | 1031734.671 | 751398.1909 | 876039.5874 | 719035.8234 | 725811.3647 | 1287988.955 | 501604.6587 | 729849.6411 | 1120135.453 |
| PC(14:1(9Z)/22:0) | 2006570.137 | 2635900.099 | 3579155.888 | 2071713.816 | 8300813.591 | 6927235.123 | 11309770.29 | 8264105.128 | 5851109.206 | 5406610.07 | 3819331.999 | 3053960.978 |
| 4-(1,3-Dioxo-1,3-dihydro-2H-isoindol-2-yl)-2-hydroxybenzoic acid | 1427674.019 | 1258948.788 | 2253659.553 | 1697565.567 | 3494926.602 | 3394630.991 | 7992357.618 | 3482613.307 | 2346921.638 | 2259050.583 | 2748316.179 | 2307277.595 |
| Isovalerylcarnitine (Car(5:0)) | 4601123.421 | 2871245.15 | 7445127.121 | 4206500.272 | 26880874.23 | 16784801.38 | 22623128.6 | 21246584.51 | 17897543.35 | 12476106.55 | 15774395.99 | 13415659.7 |
| 5-(4-Fluorobenzyl)-1,3,4-thiadiazol-2-amine | 7607188.611 | 1953044.034 | 12136801.84 | 4263990.094 | 3146004.065 | 4386899.113 | 4601162.373 | 4860611.435 | 5699563.018 | 2245469.477 | 1179365.305 | 2204343.495 |
| N-(5-Chloropyridin-2-yl)-5-propylthiophene-3-carboxamide | 45640326.13 | 40108225.51 | 64691576.6 | 51482614.02 | 122775043.3 | 104854080.6 | 248636920.1 | 108233338.2 | 74255186.71 | 69614495.3 | 85974248.41 | 70139644.82 |
| Veratridine | 6606.306349 | 0 | 25305.51289 | 6962.128784 | 162902.5002 | 163570.8117 | 137580.0493 | 142477.3104 | 85856.20958 | 101114.0935 | 157649.9495 | 75262.32046 |
| 2,3-Dinorfluprostenol | 4185894.93 | 3641390.202 | 3892546.11 | 4662058.146 | 4236562.163 | 3857186.007 | 4042273.271 | 4244311.943 | 3910022.371 | 3896706.7 | 4563901.765 | 3319869.254 |

| Metabolites | PBS_1 | PBS_2 | PBS_3 | PBS_4 | BCG_1 | BCG_2 | BCG_3 | BCG_4 | BCGΔ*pe18*_1 | BCGΔ*pe18*_2 | BCGΔ*pe18*_3 | BCGΔ*pe18*_4 |
|---|---|---|---|---|---|---|---|---|---|---|---|---|
| [(4,6-Dimethyl-2-pyrimidinyl)amino]acetic acid | 4731039.154 | 4826365.97 | 7281269.402 | 5847033.825 | 22068942.19 | 15554552.7 | 14888660.64 | 12884035.85 | 12131692.41 | 13594017.57 | 17257001.1 | 12610008.1 |
| Cer 10:1;2O/4:0 | 6737196.219 | 7180540.609 | 6443830.042 | 7148370.083 | 6918619.656 | 6760825.919 | 7575954.941 | 6691978.917 | 6818809.935 | 6994425.412 | 6583471.763 | 7798268.792 |
| Acetohexamide | 1367096.962 | 1498346.562 | 1570022.367 | 1716340.859 | 1186750.427 | 1520607.605 | 1212663.376 | 1050054.605 | 1194251.661 | 1036161.333 | 1197130.982 | 1106631.997 |
| LPA 20:4 | 1233301.468 | 1698923.205 | 2191638.993 | 1645921.269 | 881724.4858 | 957924.592 | 1199850.475 | 981143.4201 | 600412.3705 | 688761.3495 | 880636.8842 | 763093.419 |
| 3-Cyclohexyl-6-{4-[3-(trifluoromethyl)phenyl]-1-piperazinyl}-2,4(1H,3H)-pyrimidinedione | 290330.0441 | 491660.4501 | 576873.3292 | 1453483.293 | 285992.4414 | 82978.19962 | 270742.2578 | 182510.708 | 404187.6449 | 412652.2563 | 509973.5827 | 883927.9208 |
| 2-Chloro-4-methyl-7-(methylthio)quinoline | 401976.292 | 306007.6432 | 362967.4898 | 440878.4233 | 405002.2922 | 468854.2904 | 304566.2941 | 445234.6373 | 355945.9167 | 134975.5044 | 163036.0756 | 362660.731 |
| Amidosulfonic_acid | 3441189.586 | 11133048.47 | 4409896.724 | 2940745.742 | 5343909.398 | 4490214.001 | 4688942.543 | 3167352.972 | 1865269.612 | 2288792.422 | 4915320.21 | 4651577.157 |
| N-Cyclohexyl-3-methylbutanamide | 19299728.55 | 21794745.89 | 14933258.17 | 15117762.38 | 19612715.68 | 22861420.92 | 18674311.15 | 18801248.66 | 18331949.41 | 17165172.06 | 17291886.34 | 19875652.8 |
| LPI 20:2 | 1596593.572 | 1379978.666 | 1241537.712 | 1186824.952 | 3345641.867 | 2931060.732 | 3122301.827 | 2679022.37 | 2533137.821 | 2539078.382 | 3890880.013 | 1972600.915 |

| Metabolites | PBS_1 | PBS_2 | PBS_3 | PBS_4 | BCG_1 | BCG_2 | BCG_3 | BCG_4 | BCGΔ *pe18*_1 | BCGΔ *pe18*_2 | BCGΔ *pe18*_3 | BCGΔ*pe18*_4 |
|---|---|---|---|---|---|---|---|---|---|---|---|---|
| [3,4,5-trihydroxy-6-(3,4,5-trihydroxybenzoyl)oxyoxan-2-yl]methyl 3,4,5-trihydroxybenzoate | 176282.2805 | 186908.9044 | 175706.6259 | 181495.444 | 161093.4574 | 204572.9184 | 145960.5747 | 360858.8801 | 204434.5648 | 170827.8579 | 130281.4963 | 104713.44 |
| 1,2-Dilinolenoyl-sn-glycero-3-phosphoethanolamine | 4711208.731 | 5977372.382 | 2469825.636 | 4265238.115 | 5621318.694 | 14566277.99 | 8755147.12 | 13469545.55 | 19491039.36 | 19949779.86 | 18396679.76 | 6840178.55 |
| PE 8:0_28:5 | 735639.2049 | 660462.6633 | 357999.9459 | 581712.6171 | 3145731.24 | 2892849.977 | 3629094.386 | 3363307.656 | 3458484.136 | 3026993.749 | 3102531.12 | 2102986.104 |
| TG 16:4_18:5_18:5 | 483377.997 | 537431.7949 | 248325.5486 | 372030.7066 | 262478.5128 | 273423.4489 | 332604.9436 | 304906.2202 | 391948.0451 | 387266.5301 | 281410.018 | 314060.3423 |
| PI O-10:0_24:2 | 541078.8456 | 224503.257 | 558750.0544 | 492169.2166 | 554067.4864 | 0 | 366356.7233 | 285893.0497 | 664839.4455 | 692350.9145 | 383982.3036 | 210792.6773 |
| Luvangetin | 1535892.824 | 2437641.197 | 1818514.345 | 2221326.383 | 2140960.787 | 2227687.996 | 1996549.766 | 2220007.461 | 1996367.663 | 1876173.538 | 1825237.363 | 2638957.412 |
| 1-O-Cinnamoyl-6-O-p-coumaroyl-.beta.-D-glucopyranose | 871262.0689 | 842128.067 | 771201.9554 | 769660.4801 | 805530.0833 | 860910.6148 | 880193.1307 | 797972.3627 | 803193.5824 | 828028.4758 | 773223.3763 | 797326.108 |
| 11(12)-Epoxy-5Z,8Z,14Z-eicosatrienoic acid | 0 | 884626.9297 | 762231.0804 | 910562.176 | 4052080.773 | 4893477.896 | 3533715.887 | 4235663.071 | 3038277.296 | 2146112.422 | 4040937.853 | 4294401.03 |

| Metabolites | PBS_1 | PBS_2 | PBS_3 | PBS_4 | BCG_1 | BCG_2 | BCG_3 | BCG_4 | BCGΔ*pe18*_1 | BCGΔ*pe18*_2 | BCGΔ*pe18*_3 | BCGΔ*pe18*_4 |
|---|---|---|---|---|---|---|---|---|---|---|---|---|
| 5,5'-Methylenebis(2-aminobenzoic acid) | 2016889.57 | 1876561.167 | 4549114.408 | 3431390.49 | 3746393.921 | 5192362.811 | 3770799.149 | 3688075.162 | 3566058.125 | 2273556.081 | 2659480.422 | 2884310.537 |
| Cyanazine | 4897212.488 | 4047239.35 | 4948079.713 | 3738076.9 | 12403311.03 | 6371457.766 | 9438457.116 | 8768252.654 | 8162035.743 | 9227804.125 | 7468883.657 | 4845571.736 |
| SM 22:0;2O/19:2 | 4578678.822 | 8892759.275 | 4234058.578 | 10435590.79 | 5901119.26 | 2298143.071 | 4526089.642 | 3441943.586 | 5423720.333 | 3596611.665 | 287008.6014 | 5452976.474 |
| Pisiferal | 1092719.085 | 1024660.724 | 1384909.187 | 1338289.041 | 878037.6748 | 2096280.904 | 1113159.369 | 1609427.28 | 881144.7414 | 1070212.925 | 1399512.82 | 1319984.332 |
| NAE 13:0 | 22552765.63 | 26804897.58 | 24585876.49 | 20622558.08 | 28828486.92 | 29744297.61 | 20807000.4 | 26195297.32 | 23039868.97 | 26483996.66 | 24772776.64 | 23011531.45 |
| Linoleamide | 1315189943 | 1364747255 | 857341394.7 | 1079266497 | 1304435382 | 1096019844 | 1221829493 | 1447876992 | 1144715079 | 1435563509 | 1375633701 | 990885823.1 |
| SPB 19:0;2O | 3605283.101 | 2935148.216 | 3647794.993 | 8094054.592 | 3273636.658 | 5340460.59 | 8493053.892 | 8901351.345 | 3786592.794 | 7941556.733 | 2385961.981 | 2824578.393 |
| (3.beta.,5.Xi.,9.Xi.)-23-Hydroxy-3-((.beta.-D-xylopyranosyl-(1->3)-6-deoxy-.alpha.-L-mannopyranosyl-(1->2)-.alpha.-L-arabinopyranosyl)oxy)olean-12-en-28-oic acid | 512711.803 | 734335.3841 | 636693.5475 | 1160800.814 | 2663745.209 | 2473268.85 | 1596472.405 | 1501140.637 | 3032415.956 | 1807515.987 | 2201340.452 | 1650968.703 |

| Metabolites | PBS_1 | PBS_2 | PBS_3 | PBS_4 | BCG_1 | BCG_2 | BCG_3 | BCG_4 | BCGΔ*pe18*_1 | BCGΔ*pe18*_2 | BCGΔ*pe18*_3 | BCGΔ*pe18*_4 |
|---|---|---|---|---|---|---|---|---|---|---|---|---|
| 2-((3r,5r,7r)-Adamantan-1-yl)piperidine | 4464755.324 | 4456649.737 | 4390897.804 | 4295622.699 | 4639779.807 | 4531710.009 | 4421414.057 | 4272054.309 | 4763508.409 | 4779633.008 | 4269814.453 | 4180078.749 |
| N-Pentadecanoyl-L-homoserine lactone | 3977389.972 | 4455069.298 | 3815436.996 | 3487395.688 | 4520523.167 | 4149423.419 | 3209629.422 | 4367122.611 | 3200599.971 | 3711115.621 | 4423175.489 | 3470844.368 |
| N-Ethyl-2-methyl-5-nitro-N-phenylbenzenesulfonamide | 1978692.282 | 1934993.117 | 1901262.261 | 1906026.591 | 2443774.798 | 2294105.083 | 2256695.381 | 2014655.791 | 2060979.994 | 2252136.666 | 2302550.102 | 1977582.769 |
| PA(20:4(5Z,8Z,11Z,14Z)/0:0) | 1119368.077 | 1332364.99 | 1192564.868 | 1071979.222 | 1131664.057 | 1301763.729 | 1043454.162 | 1231159.532 | 953235.8785 | 812986.6666 | 1135526.64 | 1099890.023 |
| 4-Hydroxymethylpyrazole | 6342539.926 | 9874231.772 | 7145001.58 | 6990759.542 | 7221056.697 | 7450455.225 | 6905020.825 | 7622578.79 | 7416476.982 | 6775649.575 | 7037099.894 | 7439900.991 |
| 5-Aminopentanoic acid | 10552145.54 | 12426999.89 | 9134270.572 | 20260246.39 | 12426757.77 | 12653333.59 | 10994232.34 | 11346681.39 | 10277915.39 | 12998079.9 | 11566075.44 | 9513537.125 |
| 2-(3-Bromophenyl)-1H-benzimidazole | 1321093.826 | 1030181.958 | 1534790.41 | 1176080.724 | 1997335.006 | 1679120.618 | 2410592.925 | 1895317.858 | 1495317.283 | 2013731.421 | 1941568.822 | 1344603.485 |
| Asn-Arg | 2199737.04 | 1325600.598 | 1976845.827 | 859787.8431 | 1192617.084 | 1429356.418 | 1248415.793 | 705927.272 | 1725278.191 | 2046855.01 | 2603245.663 | 1561103.626 |
| 2-Docosahexaenoyl-1-stearoyl-sn-glycero-3-phosphoethanolamine | 42822201.14 | 52360574.46 | 62507516.85 | 55744440.31 | 112453285.2 | 91529329.72 | 109884933.7 | 100032886.7 | 112336906.7 | 113480868.3 | 100216325.2 | 61022136.67 |
| PS(22:4(7Z,10Z,13Z,16Z)/20:2(11Z,14 | 18667027.07 | 22155564.2 | 11420279.33 | 14739715.15 | 3927085.008 | 2477103.627 | 5378286.914 | 4546522.313 | 4492371.347 | 6173181.841 | 4727431.212 | 3217299.141 |

| Metabolites | PBS_1 | PBS_2 | PBS_3 | PBS_4 | BCG_1 | BCG_2 | BCG_3 | BCG_4 | BCGΔ *pe18*_1 | BCGΔ *pe18*_2 | BCGΔ *pe18*_3 | BCGΔ*pe18*_4 |
|---|---|---|---|---|---|---|---|---|---|---|---|---|
| Z)) | | | | | | | | | | | | |
| Imidazol-2-one | 962569.2798 | 1214019.702 | 1619392.455 | 1740308.31 | 8988954.007 | 5040906.775 | 7084543.343 | 3057306.39 | 4122501.169 | 4548397.398 | 2524676.664 | 2536775.582 |
| Enoxolone | 433784.4309 | 394812.4546 | 400328.9842 | 429524.1391 | 420352.883 | 448328.0557 | 424259.0784 | 447876.6148 | 414648.208 | 420273.115 | 398063.0804 | 401460.5954 |
| N-(1-Deoxy-1-fructosyl)threonine | 481122.4235 | 1174653.725 | 458480.2002 | 1091539.15 | 1716828.957 | 543234.1826 | 504652.8985 | 288633.4121 | 409751.5992 | 1080814.985 | 585648.345 | 341481.9257 |
| Acetylcarnitine (Car(2:0)) | 130469984.1 | 103764766.3 | 115253789.7 | 94822781.63 | 211547066.8 | 140639737 | 183990621 | 176116588.6 | 137939296.5 | 131607826 | 136403563 | 77318776.27 |
| N-3-Oxohexadec-11Z-enoyl-L-homoserine lactone | 3759112.377 | 3331885.389 | 3034512.997 | 2889447.757 | 3612863.999 | 3466128.884 | 5634945.29 | 3526506.409 | 3705970.349 | 3730521.233 | 4249410.577 | 3592956.954 |
| 4-Methoxy-2(5H)-furanone | 601948.6368 | 1378891.689 | 736495.2395 | 1277451.314 | 2703960.145 | 2543925.945 | 2533295.73 | 5284850.182 | 3035262.46 | 2198405.897 | 3234126.328 | 6350933.145 |
| Phe-Met-Arg-Phe-amide | 528576.5809 | 416652.7863 | 754864.0261 | 598000.6077 | 3310231.517 | 1954858.267 | 2728514.74 | 2155444.566 | 2344019.588 | 2085094.275 | 2573568.854 | 2102362.207 |
| Val-Gly-Val | 1443241.431 | 1629166.347 | 4982669.942 | 1855527.908 | 3656509.214 | 2508018.64 | 3580833.878 | 2889851.336 | 1893226.423 | 2616269.416 | 2429922.342 | 1523335.211 |
| PS 39:3 | 0 | 0 | 0 | 0 | 1712172.922 | 2138793.957 | 1998406.285 | 1579943.656 | 1824988.898 | 2606977.979 | 2448529.874 | 1809540.728 |
| Adenine | 3376186.606 | 1441135.2 | 2536541.14 | 1510836.882 | 3366873.197 | 3041130.619 | 5772137.852 | 3226362.601 | 2887028.465 | 2568360.88 | 3328038.748 | 2431704.92 |
| Nonadecanoic acid | 561569. | 684107. | 369925. | 579536. | 439036 | 693820 | 372271 | 457492 | 260793 | 283687 | 468757 | 243868 |

| Metabolites | PBS_1 | PBS_2 | PBS_3 | PBS_4 | BCG_1 | BCG_2 | BCG_3 | BCG_4 | BCGΔ*pe18*_1 | BCGΔ*pe18*_2 | BCGΔ*pe18*_3 | BCGΔ*pe18*_4 |
|---|---|---|---|---|---|---|---|---|---|---|---|---|
| | 3122 | 7356 | 161 | 0416 | 5.093 | 2.995 | 8.029 | 5.106 | 1.421 | 7.501 | 2.624 | 7.739 |
| M447T60 | 182799.26 | 359496.7464 | 197919.4884 | 377763.8529 | 153747.567 | 514697.2486 | 667888.1536 | 199644.7538 | 198463.5671 | 570925.9182 | 784395.0581 | 295973.1081 |
| 3-Ketocholesterol | 1928492.977 | 2661492.699 | 1111455.837 | 1490658.129 | 96976535.43 | 102428634.5 | 80924920.7 | 93400803.03 | 45725519.48 | 44986416.06 | 54966334.64 | 41583474.35 |
| Cer(d18:1/17:0) | 463447.7431 | 342116.6071 | 493960.0876 | 527915.5224 | 467716.1751 | 7677.241292 | 349758.1439 | 416766.871 | 384666.2811 | 458488.7941 | 340666.1866 | 536608.3147 |
| 5-Methyl DL-glutamate | 539520.6849 | 428825.2216 | 671707.1403 | 469837.7863 | 5968752.342 | 4307097.965 | 7266032.838 | 6036266.314 | 4031855.072 | 4296796.06 | 4497936.547 | 4708631.743 |
| PA 2:0_22:3 | 441716.3394 | 397507.3268 | 628920.5827 | 416838.7437 | 1434119.519 | 1177471.042 | 1339995.187 | 1097564.192 | 1112568.47 | 1254672.842 | 1459755.38 | 1058550.489 |
| SM(d17:1/24:0) | 3106124.364 | 3236482 | 3606089.648 | 3389997.749 | 3395916.147 | 2761263.443 | 3458452.393 | 3076741.038 | 3262386.257 | 3332654.105 | 3239669.641 | 1362923.562 |
| Prostaglandin E2 (PGE2) | 104524.1426 | 97639.58858 | 147916.1577 | 127112.4868 | 3978598.809 | 5580463.293 | 2745266.979 | 4475237.352 | 6172162.222 | 1837598.459 | 2234567.183 | 6709843.609 |
| PS 18:0_24:1 | 313361.2724 | 871721.3632 | 663531.5621 | 329378.0038 | 296829.7676 | 1587048.994 | 626250.6798 | 569187.2569 | 940984.4337 | 283920.6815 | 523448.8614 | 1642433.184 |
| LDGCC 42:12 | 535085.5159 | 568595.9511 | 71144.34811 | 398328.7457 | 3088395.308 | 2227669.448 | 2850568.309 | 2712754.266 | 3387609.492 | 4074257.546 | 2842909.219 | 1902406.513 |
| 4-Methoxy-4'-(methylthio)chalcone | 599780.8482 | 410065.825 | 1680477.266 | 1429306.077 | 8269328.505 | 6382335.91 | 7863204.184 | 8294582.118 | 1949639.542 | 6167141.884 | 2175862.356 | 5326734.396 |
| O-t-Butyl-L-threonine methyl ester | 1134724.134 | 12735545.01 | 14336281.16 | 16454109.37 | 2846999.477 | 870774.115 | 1133935.658 | 842948.5278 | 3449637.929 | 4496385.452 | 1623779.872 | 31424359.6 |
| LPG 19:1 | 363799. | 389896. | 322813. | 275561. | 349813 | 297590 | 389040 | 295228 | 152632 | 174798 | 214487 | 131738 |

| Metabolites | PBS_1 | PBS_2 | PBS_3 | PBS_4 | BCG_1 | BCG_2 | BCG_3 | BCG_4 | BCGΔ *pe18*_1 | BCGΔ *pe18*_2 | BCGΔ *pe18*_3 | BCGΔ*pe18*_4 |
|---|---|---|---|---|---|---|---|---|---|---|---|---|
| | 6257 | 4121 | 5622 | 3196 | 5.811 | 5.994 | 6.802 | 4.168 | 0.505 | 5.964 | 6.559 | 1.409 |
| Prostaglandin H2 (PGH2) | 104524.1426 | 97639.58858 | 147916.1577 | 127112.4868 | 3978598.809 | 5580463.293 | 2745266.979 | 4475237.352 | 6172162.222 | 1837598.459 | 2234567.183 | 6709843.609 |
| Bis(glycerophospho)-glycerol | 22033.91387 | 19458.92823 | 42586.51185 | 26902.40483 | 1020668.947 | 767623.8698 | 759855.1186 | 777876.1821 | 469105.8471 | 427869.7148 | 475560.434 | 381198.182 |
| LPC 36:4 | 1275782.764 | 1693423.137 | 1687433.449 | 2413924.598 | 1620996.878 | 2141832.649 | 3114627.941 | 9227081.563 | 2150772.019 | 2087515.155 | 572962.2081 | 4438861.138 |
| Diuron-desdimethyl | 34952380.08 | 30760951.78 | 37568580.77 | 32624417.99 | 46264729.54 | 39401039.37 | 47291408.97 | 43805783.09 | 38690775.45 | 43603237.09 | 43655595.49 | 34298839.45 |
| (12Z)-9,10,11,14,15,16,17,18,19,20,21,22-Dodecahydro-5,8-ethenobenzo[20]annulene-2,4,6,24-tetrol | 3657376.05 | 3145256.443 | 4969577.524 | 3624636.817 | 7839572.709 | 9564108.824 | 8179138.77 | 6898473.026 | 5288527.265 | 5494499.067 | 6794431.858 | 7002267.902 |
| LNAPS 22:1/N-16:3 | 1314774.981 | 1451308.362 | 1280271.72 | 1488228.796 | 1607727.545 | 1872150.266 | 1834654.004 | 1301788.425 | 1753047.205 | 1280590.623 | 345578.66 | 940894.2968 |
| SL 10:0;O/11:0 | 2772395.632 | 3896581.138 | 2259102.7 | 3979653.112 | 2985823.911 | 5549558.376 | 3041429.503 | 2949892.971 | 2282784.356 | 2588660.418 | 3237156.387 | 1857132.441 |
| (3R)-3-Hydroxy-L-proline | 758595.9595 | 631078.8877 | 763552.0847 | 744097.2434 | 1688364.391 | 1298332.47 | 1386287.292 | 1276221.514 | 1052721.48 | 1292317.087 | 1017502.315 | 2689148.534 |
| PC 15:0_16:0 | 1131480.923 | 2060481.8 | 1241761.404 | 1131078.074 | 815590.1556 | 73153.79535 | 2604348.583 | 851690.0984 | 1141079.676 | 1088276.425 | 2271491.365 | 638882.6786 |
| PC O-24:4 | 124084 | 159002 | 267755 | 140476 | 102094 | 161514 | 163021 | 147758 | 154609 | 151216 | 213047 | 113021 |

| Metabolites | PBS_1 | PBS_2 | PBS_3 | PBS_4 | BCG_1 | BCG_2 | BCG_3 | BCG_4 | BCGΔ *pe18*_1 | BCGΔ *pe18*_2 | BCGΔ *pe18*_3 | BCGΔ*pe18*_4 |
|---|---|---|---|---|---|---|---|---|---|---|---|---|
| | 4.758 | 6.75 | 8.291 | 7.681 | 5.777 | 7.59 | 5.542 | 7.728 | 0.966 | 5.993 | 1.892 | 3.439 |
| Cer 8:1;2O/18:1 | 3007939.976 | 3296442.066 | 1374428.685 | 1843370.546 | 1643523.612 | 1569548.396 | 2436173.822 | 2184674.074 | 2210508.215 | 2040834.615 | 1808014.196 | 1351413.716 |
| Fluticasone_propionate | 217626.2995 | 353124.6334 | 215430.2516 | 152612.2424 | 195343.4225 | 194771.5321 | 243774.2967 | 195110.652 | 200874.1778 | 247176.3713 | 412359.7543 | 181806.3759 |
| TG 8:0_16:2_22:6 | 6735120.323 | 8266860.928 | 6175819.467 | 5626146.695 | 4241518.558 | 4246498.259 | 5701212.017 | 4709222.411 | 5583224.869 | 5373392.18 | 5934797.458 | 3151583.024 |
| Butanoic acid, 4-[(1,2-dioxohexadecyl)amino]-, ethyl ester | 3937810.833 | 5903426.248 | 3583424.689 | 3424068.055 | 3876087.654 | 3995485.462 | 3519884.159 | 5900243.519 | 3783339.677 | 3872505.426 | 3916639.767 | 3730144.545 |
| 3-hydroxytridecanoyl carnitine | 1405429.905 | 1213558.554 | 1014863.316 | 1756555.323 | 1269888.907 | 1358079.39 | 1870964.878 | 1755831.017 | 1801459.632 | 2209061.696 | 1563090.947 | 1277014.59 |
| PC O-41:7 | 1594621.981 | 1687804.098 | 1439297.512 | 1804554.924 | 1438054.383 | 1653855.951 | 1477266.608 | 1772634.386 | 1323540.892 | 1856943.909 | 1530254.639 | 2101313.289 |
| Cer 14:0;2O/16:5;(3OH)(FA 20:5) | 972699.9611 | 427364.8924 | 922017.5507 | 514175.6062 | 1226105.019 | 27489.31423 | 387136.614 | 464495.6093 | 483242.3583 | 645228.9257 | 771645.0421 | 339464.6129 |
| Formylpyruvate | 822554.6158 | 761621.6489 | 923828.9276 | 945538.3973 | 3746825.409 | 3323721.637 | 4580744.817 | 3656141.767 | 2554737.909 | 3311490.916 | 3709073.457 | 2874829.256 |
| epiyangambin | 43915.53224 | 24843.94588 | 59935.69946 | 59742.20382 | 227170.7873 | 200191.4106 | 246842.7429 | 198069.842 | 168004.9781 | 128112.9957 | 261060.5042 | 162788.1083 |
| 2-Oxoadipic acid | 1186482.273 | 1375115.625 | 946382.4124 | 1278304.512 | 2056643.733 | 1974448.784 | 1854876.758 | 1793404.925 | 1669625.743 | 1231831.773 | 2247461.932 | 1144904.893 |

| Metabolites | PBS_1 | PBS_2 | PBS_3 | PBS_4 | BCG_1 | BCG_2 | BCG_3 | BCG_4 | BCGΔ*pe18*_1 | BCGΔ*pe18*_2 | BCGΔ*pe18*_3 | BCGΔ*pe18*_4 |
|---|---|---|---|---|---|---|---|---|---|---|---|---|
| 9,10-Dihydrojasmonic acid | 790302.8079 | 608859.8983 | 779801.1346 | 822774.0478 | 554301.665 | 1206096.168 | 5128460.732 | 676117.4194 | 1784935.991 | 1269845.929 | 658285.0331 | 3308033.746 |
| PG 2:0_4:0 | 183265.4347 | 144321.1859 | 273718.2861 | 195746.602 | 900703.8483 | 694417.3997 | 679803.4426 | 700973.2158 | 468591.6617 | 381638.657 | 434615.3232 | 516488.3858 |
| 1-(5-Nitro-1H-indol-3-yl)ethan-1-one | 526650.6834 | 704010.9351 | 577478.9907 | 276855.77 | 1217412.495 | 842448.1109 | 1147225.687 | 919246.3056 | 918923.4041 | 852743.5688 | 1287512.484 | 735128.9536 |
| 6(5H)-Pteridinone, 2-[(3,5-difluoro-4-hydroxyphenyl)amino]-7,8-dihydro-5,7-dimethyl-8-(3-methylbutyl)- | 2367313.695 | 2403975.426 | 2409459.551 | 2684187.196 | 11038214.45 | 9559395.093 | 7457256.123 | 10796218.57 | 9854494.943 | 4932978.338 | 5977789.853 | 7828264.303 |
| Dihomo-gamma-linolenoylcholine [M] | 273506.4235 | 407485.5022 | 435698.1649 | 435924.7172 | 289146.129 | 425230.4567 | 515122.0629 | 463918.4 | 460536.7387 | 456895.1336 | 501083.1952 | 480167.8404 |
| 3',4,5',6,8'-Pentahydroxy-2',8'-bis(4-hydroxyphenyl)spiro[1-benzofuran-2,9'-3,4-dihydro-2H-furo[2,3-h]chromene]-3-one | 2450222.795 | 1070621.902 | 3641024.313 | 1196839.628 | 1991064.158 | 1997762.626 | 1235985.496 | 898328.8046 | 1082311.942 | 816762.2136 | 1141778.506 | 973094.1497 |
| Dihydroxyoctadecenoic acid | 536650.5046 | 619870.28 | 1000686.206 | 714867.7355 | 3499283.353 | 2911709.832 | 2540025.771 | 2457100.469 | 1606842.493 | 1052658.355 | 2969592.729 | 2009432.797 |
| MGDG 5:0_19:2 | 190000 | 169637 | 187561 | 198675 | 229924 | 188958 | 196293 | 175707 | 197252 | 189988 | 188077 | 183682 |

| Metabolites | PBS_1 | PBS_2 | PBS_3 | PBS_4 | BCG_1 | BCG_2 | BCG_3 | BCG_4 | BCGΔ*pe18*_1 | BCGΔ*pe18*_2 | BCGΔ*pe18*_3 | BCGΔ*pe18*_4 |
|---|---|---|---|---|---|---|---|---|---|---|---|---|
| | 1.56 | 9.38 | 1.677 | 2.813 | 2.139 | 5.803 | 3.98 | 1.494 | 3.991 | 6.085 | 3.918 | 3.442 |
| Tetrabutylammonium cation | 1676171.546 | 2373677.843 | 1451372.464 | 1972311.548 | 1577477.491 | 1454540.231 | 1267911.031 | 1531814.481 | 1617273.713 | 1607098.252 | 1482443.364 | 2267039.242 |
| TG 16:4_18:5_28:7 | 139875.7406 | 229038.4347 | 354932.3981 | 122829.5587 | 202071.7018 | 268412.27 | 318372.8912 | 296847.4859 | 165460.1854 | 103239.1058 | 346338.1423 | 1502035.323 |
| 1-(4-Hydroxyphenyl)-1-hexanone | 1150015.907 | 1128684.269 | 992201.7235 | 1184470.696 | 1133514.546 | 1140718.318 | 1117788.221 | 1067005.567 | 1097644.27 | 1137923.241 | 1127688.049 | 1158689.264 |
| PE O-22:1_16:3 | 25982644.79 | 24821308.87 | 17707566.74 | 19918739.02 | 3024359.082 | 3118080.486 | 6180166.19 | 2910945.662 | 3926744.553 | 4423068.441 | 5839560.726 | 3801212.078 |
| (4E,10E)-13,21-Dihydroxy-8,14,19-trimethoxy-4,10,12,16-tetramethyl-3,20,22-trioxo-2-azabicyclo[16.3.1]docosa-1(21),4,10,18-tetraen-9-yl carbamate | 1810000.945 | 1699032.089 | 2181747.104 | 1584582.388 | 1544708.203 | 1650411.855 | 2172994.477 | 2033165.878 | 1588670.254 | 1077223.547 | 2099449.618 | 2220170.294 |
| 2-aminohex-4-ynoate | 868258.5937 | 859651.3561 | 867225.3859 | 1061949.245 | 1446692.287 | 1280331.869 | 1215452.952 | 1306788.901 | 1585811.929 | 1170660.775 | 1181649.615 | 1530187.439 |
| 3-(2,6-Dichlorophenyl)-N-(3-methoxyphenyl)-5-methyl-1,2-oxazole-4-carboxamide | 4011847.286 | 3167643.692 | 5530853.181 | 4459842.947 | 4774085.576 | 5436419.588 | 4889767.39 | 5330312.112 | 4750482.336 | 2228564.323 | 2417915.313 | 4021272.57 |

| **Metabolites** | **PBS_1** | **PBS_2** | **PBS_3** | **PBS_4** | **BCG_1** | **BCG_2** | **BCG_3** | **BCG_4** | **BCGΔ*pe18*_1** | **BCGΔ*pe18*_2** | **BCGΔ*pe18*_3** | **BCGΔ*pe18*_4** |
|---|---|---|---|---|---|---|---|---|---|---|---|---|
| PI 18:1_20:3 | 905297.1368 | 464734.2272 | 995875.3046 | 1016802.682 | 790262.7238 | 26666.36255 | 726867.1354 | 830875.5294 | 1177238.484 | 791914.9256 | 731269.6883 | 896905.953 |
| (6aS,13aR,13bR)-4a-(2-Hydroxypropan-2-yl)-12-methoxy-6a,8,13b-trimethyl-1,2,4a,5,6,6a,9,13,13a,13b-decahydrofuro[3,4-i]pyrano[3,2-a]xanthene-3,11-dione | 27519.5735 | 22378.67255 | 47639.11457 | 32992.73915 | 413433.3836 | 202860.8893 | 196609.2106 | 165436.6591 | 136339.5832 | 171976.8348 | 324901.2458 | 329081.1941 |
| Methyl 3,4-dihydroxybenzoate | 450622.7277 | 450482.8339 | 571921.9885 | 560050.6009 | 1039311.678 | 804648.0281 | 926127.7008 | 938626.8659 | 634012.2151 | 527536.0677 | 676592.1487 | 637312.6396 |
| MGDG O-18:5_6:0 | 10112490.59 | 9545933.007 | 8750889.724 | 9768208.763 | 9587205.329 | 10076218.34 | 10287460.53 | 9954845.889 | 9729549.863 | 9916382.151 | 9697124.899 | 8839487.513 |
| 6-Amino-2,4-dichloro-3-ethylphenol | 8443575.008 | 7192105.441 | 7932239.901 | 7572706.363 | 7333860.055 | 7213818.257 | 4342967.426 | 4737368.869 | 7513067.445 | 5256601.39 | 978474.2685 | 7667081.01 |
| NAE 16:4 | 2418189.235 | 5008178.336 | 2018389.698 | 6690564.364 | 3345929.614 | 2112840.905 | 10071888.77 | 8868412.981 | 2469618.345 | 3119301.02 | 4580723.978 | 2819596.467 |
| LPG 17:0 | 558157.0863 | 532550.5178 | 96445.46073 | 135034.6989 | 1271808.001 | 955176.9231 | 1267708.979 | 1036347.302 | 3256324.255 | 745796.8794 | 954449.7597 | 2395356.809 |
| PEtOH 18:1_24:4 | 522457.4752 | 643817.5633 | 586425.626 | 635974.6137 | 576108.4622 | 6081.896058 | 521640.682 | 508053.101 | 391004.7002 | 342942.2425 | 422415.1773 | 396562.8486 |

| Metabolites | PBS_1 | PBS_2 | PBS_3 | PBS_4 | BCG_1 | BCG_2 | BCG_3 | BCG_4 | BCGΔ*pe18*_1 | BCGΔ*pe18*_2 | BCGΔ*pe18*_3 | BCGΔ*pe18*_4 |
|---|---|---|---|---|---|---|---|---|---|---|---|---|
| PC O-39:5 | 677382.6067 | 614781.9851 | 755022.2152 | 821674.6177 | 605378.0728 | 197052.5098 | 2288734.987 | 3161489.722 | 1256051.108 | 524110.8914 | 756312.8148 | 3708851.823 |
| PS 19:0_22:5 | 680007.3311 | 823288.5813 | 1033420.736 | 398157.2803 | 599060.43 | 349983.6724 | 698586.1415 | 476690.4041 | 612102.213 | 1008964.292 | 646849.9327 | 322975.5543 |
| N,N'-Hexane-1,6-diylbis(N'-cyanoguanidine) | 37184.1363 | 1054376.203 | 947006.7859 | 971579.2149 | 787688.9266 | 763942.9597 | 783078.2383 | 721139.2543 | 21897.36172 | 978513.0217 | 774618.3773 | 1390489.992 |
| alpha-Linolenic acid | 15175518.26 | 14979331.09 | 13851128.64 | 16748138.91 | 74509521.25 | 94759332.82 | 60990732.02 | 84838105.48 | 56379905.57 | 30252020.15 | 61463034.22 | 55974840.94 |
| N-Eicosapentaenoyl Arginine | 1712962.947 | 2246959.63 | 1062965.244 | 1588444.383 | 960277.6705 | 995145.4209 | 957745.589 | 1088512.484 | 1214559.197 | 1090387.567 | 1064893.987 | 820739.4476 |
| Daidzin | 301946.0408 | 350303.1351 | 280000.5755 | 367604.5941 | 889011.349 | 790775.1985 | 774038.2269 | 1373340.175 | 1636750.677 | 828556.0981 | 1374616.581 | 1538401.129 |
| PEtOH 19:2_22:6 | 1789642.974 | 2287071.979 | 2015415.426 | 2591948.705 | 1324780.905 | 1143432.236 | 1620712.162 | 1327027.657 | 1687235.677 | 1710814.24 | 1488005.183 | 1231194.016 |
| Cer(d18:1/4:0) | 1652397.064 | 2417350.798 | 1256017.441 | 1957978.136 | 1867569.683 | 1846677.607 | 1868709.849 | 1943746.701 | 1949542.273 | 1777535.546 | 1607708.368 | 1793598.513 |
| LPI 24:4 | 649484.5649 | 706085.8875 | 653277.5856 | 585496.1166 | 384850.0115 | 305475.2561 | 555266.5878 | 263124.8737 | 390857.5756 | 499136.04 | 485656.6197 | 302328.552 |
| 1-Pyrenesulfonic acid | 1057052.168 | 1054804.996 | 1008518.669 | 441876.6035 | 135699867.8 | 72170930.66 | 79192858.32 | 75372710.82 | 7371957.972 | 81407076.58 | 72595588.24 | 28952988.09 |
| 2,6-Dimethyl-4-(4-piperidinyl)morpholine | 549620.9873 | 981175.0503 | 932501.78 | 5769848.158 | 540588.4871 | 1758622.18 | 1012375.34 | 900825.5662 | 343281.8733 | 979097.6832 | 970547.1326 | 969044.3242 |

| **Metabolites** | **PBS_1** | **PBS_2** | **PBS_3** | **PBS_4** | **BCG_1** | **BCG_2** | **BCG_3** | **BCG_4** | **BCGΔ*pe18*_1** | **BCGΔ*pe18*_2** | **BCGΔ*pe18*_3** | **BCGΔ*pe18*_4** |
|---|---|---|---|---|---|---|---|---|---|---|---|---|
| DGDG O-9:0_18:0 | 684822.2575 | 1004009.189 | 409397.4025 | 734685.1919 | 5101160.758 | 2901617.754 | 1474275.065 | 3000953.023 | 3716586.194 | 3299421.552 | 1711251.747 | 1204187.606 |
| 1,3-Cyclohexanedicarboxylic acid | 445059.0176 | 678055.2425 | 833395.5156 | 731504.8312 | 604734.5936 | 396593.6953 | 450518.1783 | 513239.726 | 636766.0712 | 451069.4766 | 445586.9679 | 1551896.71 |
| MGDG O-10:0_17:1 | 617967.3113 | 563910.4043 | 621700.6829 | 800565.6635 | 606390.0538 | 611759.1545 | 608618.2208 | 686552.7948 | 603959.6286 | 416234.6575 | 698722.3675 | 713704.6079 |
| 7H-Pyrrolo[3,2-f]quinazoline-1,3-diamine, N3-cyclopropyl-7-[[4-(1-methylethyl)phenyl]methyl]- | 121570.8377 | 898286.2227 | 780469.9171 | 1226403.936 | 932704.9518 | 949022.4445 | 877944.4463 | 888121.2667 | 928415.317 | 888138.9472 | 868858.9122 | 859457.1276 |
| FA 22:2 | 151361.8948 | 158654.727 | 298459.6636 | 151030.1013 | 2943590.939 | 6352467.57 | 3142919.717 | 3606168.975 | 1606126.514 | 1267606.228 | 3507070.41 | 2233395.443 |
| IS5 | 101094969.7 | 101593001.5 | 94898102.59 | 95221360.58 | 95947940.36 | 97397575.87 | 98835950.57 | 101026977.2 | 98738841 | 94043245.33 | 95482743.14 | 94439063.85 |
| Simvastatin | 1559131.23 | 1962983.115 | 1539545.92 | 1430150.446 | 1802933.387 | 1777559.685 | 1673616.928 | 1630666.839 | 1374639.875 | 1748036.895 | 1856289.74 | 1716927.931 |
| 3-(3-Hydroxy-5-methylphenoxy)-5-methylbenzene-1,2-diol | 1119201.694 | 1197127.939 | 1181066.44 | 1107321.606 | 1271676.056 | 1346214.492 | 1243030.523 | 26163.18168 | 1144525.045 | 1231821.338 | 1189773.1 | 1175262.444 |
| Pregabalin | 2984152.326 | 2635127.888 | 990479.2182 | 6063192.021 | 2082616.619 | 3445232.03 | 7348281.974 | 2283983.292 | 5140015.094 | 4273788.546 | 2819667.484 | 6068217.311 |

| Metabolites | PBS_1 | PBS_2 | PBS_3 | PBS_4 | BCG_1 | BCG_2 | BCG_3 | BCG_4 | BCGΔ*pe18*_1 | BCGΔ*pe18*_2 | BCGΔ*pe18*_3 | BCGΔ*pe18*_4 |
|---|---|---|---|---|---|---|---|---|---|---|---|---|
| OSELTAMIVIR PHOSPHATE | 1415014.055 | 1339517.757 | 1415510.766 | 1357072.573 | 1303968.986 | 3115203.982 | 1268047.74 | 1351283.835 | 1521565.855 | 2729413.74 | 1363795.431 | 2954816.568 |
| 3-[(1E,3E)-Hepta-1,3-dienyl]pentanedioic acid | 1227883.306 | 1056676.093 | 1013978.145 | 861484.3701 | 1353941.651 | 1004462.086 | 1173149.943 | 780394.2263 | 1161481.291 | 924933.3734 | 1180317.951 | 1276441.948 |
| Carbamic acid, [3-methyl-1-[[[1-[oxo[(2-pyridinylmethyl)amino]acetyl]butyl] amino]carbonyl]butyl]-, phenylmethyl ester | 1631388.529 | 1605152.576 | 1069811.129 | 2710280.868 | 1268870.84 | 2083597.809 | 1763321.959 | 1698135.866 | 1983310.923 | 882537.9895 | 1526084.394 | 1113564.72 |
| (2E,6E,11E)-18-(2,6-Dioxopiperidin-4-yl)-9,13-dihydroxy-8-methoxy-10,12,14-trimethyl-15-oxooctadeca-2,6,11-trienoic acid | 140493.3698 | 137752.3581 | 113762.2941 | 136940.6956 | 133987.6639 | 129200.343 | 125526.9658 | 121493.4879 | 142938.0987 | 127984.6324 | 144290.5099 | 95909.96178 |
| 6-(4-Methylpiperazin-1-yl)-9H-purin-2-amine | 1123243.177 | 1147213.927 | 882096.15 | 1370230.649 | 1139270.253 | 1022772.064 | 1660870.793 | 1003862.158 | 1108821.019 | 1247873.936 | 926200.2811 | 1098548.158 |

| Metabolites | PBS_1 | PBS_2 | PBS_3 | PBS_4 | BCG_1 | BCG_2 | BCG_3 | BCG_4 | BCGΔ*pe18*_1 | BCGΔ*pe18*_2 | BCGΔ*pe18*_3 | BCGΔ*pe18*_4 |
|---|---|---|---|---|---|---|---|---|---|---|---|---|
| N6-Carbamimidoyl-N2-((4E,6E,12E,14E)-3-hydroxy-2-(hydroxymethyl)-8,10,16-trimethyloctadeca-4,6,12,14-tetraenoyl)lysine | 2478168.463 | 2007146.028 | 1622165.274 | 2010919.426 | 2102539.204 | 2108091.197 | 2252210.601 | 1759030.542 | 2237201.186 | 2070666.731 | 1632148.278 | 1808639.583 |
| Pentadecanoic acid | 62465393.76 | 48587584.34 | 36096511.97 | 39518522.99 | 39869137.86 | 52062745.6 | 42705193.27 | 42802563.97 | 32180217.59 | 35869380.37 | 43254569.27 | 48793125.93 |
| LPC 38:5 | 1108106.559 | 3033455.862 | 1668458.214 | 2435197.657 | 2043256.193 | 0 | 3310445.889 | 12987328.8 | 2471732.892 | 2178546.939 | 877210.2461 | 4000813.353 |
| Methionine | 1912335.866 | 192944.0119 | 1201383.213 | 371355.4848 | 2357909.495 | 1324577.658 | 3755809.649 | 3290924.348 | 1770744.431 | 747964.5494 | 1628127.798 | 1695976.725 |
| 2-methylhexanoate | 307855.0004 | 388656.5261 | 914347.8205 | 958782.6266 | 351370.3917 | 300859.3133 | 431244.9255 | 337340.3008 | 342989.375 | 281023.5735 | 319933.8194 | 419178.3745 |
| PS 14:1_24:2 | 1699159.958 | 1576721.667 | 2306096.278 | 1844878.214 | 1321219.272 | 0 | 1101193.428 | 1301337.777 | 3077273.816 | 2048846.267 | 1242362.834 | 2767154.223 |
| 1-(1Z-Octadecenyl)-2-(4Z,7Z,10Z,13Z,16Z,19Z-docosahexaenoyl)-sn-glycero-3-phosphoethanolamine | 75333720.07 | 91845869.72 | 84933332.64 | 89202171.49 | 133560116.3 | 99164967.62 | 119859417.2 | 115334771 | 123598428.6 | 122843071.8 | 112067502.3 | 66364641.14 |

| Metabolites | PBS_1 | PBS_2 | PBS_3 | PBS_4 | BCG_1 | BCG_2 | BCG_3 | BCG_4 | BCGΔ*pe18*_1 | BCGΔ*pe18*_2 | BCGΔ*pe18*_3 | BCGΔ*pe18*_4 |
|---|---|---|---|---|---|---|---|---|---|---|---|---|
| 2-[2-(19-Acetamido-16,18-dihydroxy-5,9-dimethyl-6-oxoicosan-7-yl)oxy-2-oxoethyl]butanedioic acid | 2030972.066 | 1247506.558 | 1137313.535 | 1469337.208 | 1579847.488 | 1453817.767 | 1261381.45 | 1099852.01 | 1401478.701 | 1671218.153 | 1782620.423 | 552917.7169 |
| N-oleoyl-glutamine | 21157.02926 | 0 | 36410.80632 | 0 | 5255115.174 | 6820040.279 | 2809549.72 | 4007510.04 | 2379363.775 | 871482.7722 | 3026472.412 | 2950881.646 |
| N-Formylglycine | 1648587.89 | 1136770.354 | 2032729.174 | 1513034.203 | 1226646.79 | 2134331.194 | 1318585.02 | 1760902.992 | 1386657.807 | 969629.0748 | 1558656.896 | 1819676.735 |
| Thiacloprid | 2249403.51 | 1390844.651 | 3015649.045 | 1797427.521 | 6259289.99 | 3468514.229 | 7633560.004 | 4845208.245 | 2940253.822 | 4381525.298 | 4918177.056 | 2237218.231 |
| PEtOH 20:4_26:4 | 1642699.36 | 3112897.327 | 573414.6225 | 1663606.822 | 3963894.966 | 0 | 5532115.236 | 6854843.294 | 2049650.686 | 746340.6031 | 1899692.699 | 4790309.162 |
| Uridine | 13516869.54 | 11595532.49 | 18409242.5 | 15267275.21 | 35144283.42 | 29776850.91 | 74079633.87 | 30831259.45 | 21501977.33 | 20361469.54 | 24299664.26 | 18980016.23 |
| DGDG O-16:4_12:0 | 449316.2753 | 429713.0282 | 429593.2224 | 329953.7341 | 433621.6992 | 513581.5704 | 617038.3408 | 427503.5006 | 381126.0855 | 502297.3274 | 403383.5544 | 419221.7866 |
| Lys-Val | 33000440.07 | 23589304.96 | 37387833.93 | 27889172.49 | 23636693.02 | 13623473.51 | 26036381.21 | 18414002.02 | 13661068.77 | 16590531.67 | 18978618.6 | 12012105.55 |
| 3-Cyclopentene-1-octanoic acid, 2-(3-hydroxy-1-penten-1-yl)-5-oxo- | 695557.0705 | 970304.5398 | 619204.54 | 662507.9196 | 680676.7835 | 1002169.666 | 1008427.088 | 948190.5898 | 820936.5239 | 746093.8784 | 801921.3199 | 961933.2544 |

| Metabolites | PBS_1 | PBS_2 | PBS_3 | PBS_4 | BCG_1 | BCG_2 | BCG_3 | BCG_4 | BCGΔ*pe18*_1 | BCGΔ*pe18*_2 | BCGΔ*pe18*_3 | BCGΔ*pe18*_4 |
|---|---|---|---|---|---|---|---|---|---|---|---|---|
| 22-Angeloylbarringtogenol_C | 835708.0978 | 882799.8317 | 819826.7046 | 778743.5776 | 839182.1642 | 914121.251 | 601824.2367 | 830394.0087 | 839475.1063 | 780151.9089 | 748131.1958 | 620857.3778 |
| 4-Guanidinobutyric acid | 5706852.007 | 6169566.76 | 5004161.398 | 5255081.237 | 14024457.88 | 8403369.985 | 11097000.94 | 9547094.385 | 7636839.559 | 10783454.39 | 7294961.2 | 5735149.667 |
| 2-Hydroxyglutaric acid | 1645017.958 | 1291464.843 | 26335062.1 | 29374129.58 | 26473117.36 | 14597570.4 | 26549059.1 | 20691291.99 | 12241059.57 | 13621823.74 | 17085950.28 | 5792344.333 |
| PC O-20:4 | 99165446.11 | 100308647.6 | 113463019.4 | 68768027.82 | 49187687.12 | 50987920.48 | 57701247.07 | 50809458.18 | 43737448.67 | 57083439.04 | 56021541.49 | 49316724.1 |
| Eicosapentaenoic acid | 9393103.572 | 10964399.4 | 31118060.45 | 17739345.58 | 7040265.49 | 9578225.822 | 5026203.851 | 8949569.253 | 6900274.501 | 3671062.425 | 4616829.434 | 9246754.053 |
| SL 17:0;O/16:0;O | 540987.1402 | 626580.4544 | 600348.778 | 613367.9394 | 523088.3146 | 337978.4981 | 628321.7577 | 529486.519 | 535091.0248 | 604161.6326 | 440190.7833 | 210235.4353 |
| SL 10:0;O/14:0 | 740063.0742 | 992814.8428 | 274950.0686 | 393575.7125 | 494814.161 | 250786.9854 | 482110.6322 | 472782.7755 | 573041.4659 | 638153.1312 | 370128.1871 | 265724.37 |
| PIPOBROMAN | 457652.9703 | 398008.5096 | 374314.6714 | 388613.3042 | 2384257.124 | 1777708.408 | 2563926.888 | 2054074.958 | 1771744.946 | 2083035.656 | 2577590.22 | 1576867.783 |
| LNAPE 2:0/N-17:1 | 961884.7919 | 982420.6415 | 214193.2007 | 1031601.512 | 1354021.492 | 578865.4522 | 1989183.338 | 1668923.418 | 600867.7589 | 971810.2814 | 1777654.724 | 272382.771 |
| PI 17:1_20:4 | 5691485.548 | 9511752.28 | 834204.671 | 2318014.693 | 876608.2575 | 698102.2421 | 992767.0129 | 892663.6023 | 1228634.35 | 1178985.47 | 740894.5246 | 583209.5509 |

| **Metabolites** | **PBS_1** | **PBS_2** | **PBS_3** | **PBS_4** | **BCG_1** | **BCG_2** | **BCG_3** | **BCG_4** | **BCGΔ*pe18*_1** | **BCGΔ*pe18*_2** | **BCGΔ*pe18*_3** | **BCGΔ*pe18*_4** |
|---|---|---|---|---|---|---|---|---|---|---|---|---|
| 1H-2-Benzopyran-7-butanoic acid, 6,8-dihydroxy-.alpha.,.gamma.-dimethoxy-3-methyl-1-oxo-, methyl ester | 758260.2076 | 1705032.654 | 383847.4532 | 1473692.907 | 3678468.049 | 1227004.71 | 1258347.523 | 659423.8015 | 1133415.185 | 2875613.503 | 1183128.171 | 787257.1024 |
| PC 8:0_30:6 | 750309.3345 | 291601.8077 | 666143.3237 | 390969.1865 | 907769.1225 | 854062.0841 | 1140238.974 | 1038035.052 | 844002.7049 | 564626.3474 | 611628.4406 | 354541.4641 |
| LPG O-20:3 | 362052.2098 | 304804.7843 | 292345.085 | 177355.9491 | 2463707.296 | 2574202.92 | 1743165.484 | 1742273.856 | 1373071.588 | 1270426.126 | 1395993.913 | 1165983.466 |
| 2-Amino-5-[2-[[2,3-dihydroxy-2-(1-hydroxyethyl)butanoyl]oxymethyl]-4-hydroxyanilino]-5-oxopentanoic acid | 2146815.728 | 2572526.075 | 2132925.757 | 2122327.806 | 2148186.389 | 1963287.013 | 2517749.781 | 1717044.337 | 2268438.349 | 2331985.969 | 2138448.462 | 2167835.389 |
| p-phenetidine | 3235426.124 | 3105492.352 | 3195806.46 | 3165905.277 | 3263292.286 | 3672897.83 | 2999095.849 | 2977036.194 | 3234923.144 | 3028033.196 | 2954258.833 | 3328664.762 |
| (3S,6R,12S)-6-Benzyl-3-butan-2-yl-9-(6,7-dihydroxyoctyl)-1,4,7,10-tetrazabicyclo[10.4.0]hexadecane-2,5,8,11-tetron | 1483394.333 | 1764098.623 | 2372301.151 | 1107194.664 | 3325066.86 | 2961085.071 | 1003635.282 | 856741.9457 | 2720709.097 | 728325.6492 | 700908.8706 | 540729.7909 |

| Metabolites | PBS_1 | PBS_2 | PBS_3 | PBS_4 | BCG_1 | BCG_2 | BCG_3 | BCG_4 | BCGΔ *pe18*_1 | BCGΔ *pe18*_2 | BCGΔ *pe18*_3 | BCGΔ*pe18*_4 |
|---|---|---|---|---|---|---|---|---|---|---|---|---|
| e | | | | | | | | | | | | |
| 2-Furoic acid | 704553.7494 | 400846.475 | 436113.2498 | 590411.0056 | 1734139.353 | 963113.5256 | 1494484.351 | 1257774.007 | 892619.7216 | 1380282.459 | 1695549.612 | 1081845.8 |
| PC 35:7 | 2309227.942 | 2322782.733 | 2016661.775 | 2163535.414 | 2155709.908 | 2103653.377 | 2216259.256 | 2241924.895 | 2670497.135 | 2396363.4 | 2399573.118 | 2475108.681 |
| Bisindolylmaleimide I | 2160664.698 | 1452837.591 | 1359209.102 | 1496841.59 | 1489924.68 | 1812374.561 | 1489725.171 | 1472638.667 | 1858664.716 | 1592734.015 | 1710950.554 | 1419294.463 |
| 1-Cyclohexylpiperazine | 3323805.11 | 1278524.182 | 7961216.906 | 7549280.114 | 15359967.07 | 7547334.682 | 1549271.157 | 15257282.05 | 8986113.955 | 8371986.369 | 3509504.608 | 1377921.763 |
| 5-((1E,3E)-Hepta-1,3-dien-1-yl)-1,2,3-trihydroxycyclopentanecarboxylic acid | 1010680.883 | 950983.4124 | 974999.3365 | 949599.3715 | 958782.5888 | 962190.326 | 920189.7434 | 1054686.626 | 983673.8945 | 907981.3141 | 917726.6909 | 960671.9239 |
| cis- 9, 10- Epoxystearic acid | 21814186.26 | 19338157.91 | 14295680.62 | 16900976.27 | 19546276.73 | 15127058.05 | 17308560.5 | 17442449.13 | 20122943.47 | 16930871.47 | 15776641.01 | 14170033.4 |
| (2E)-2-(Propan-2-yl)but-2-enedioic acid | 856889.5395 | 5872206.436 | 1016127.332 | 6553746.86 | 5462725.188 | 4727974.976 | 4300751.8 | 4470915.62 | 4590152.177 | 4631899.388 | 5206606.79 | 4989899.582 |
| LPS 18:2 | 5336178.146 | 5230434.265 | 7705039.035 | 5247680.808 | 9447391.011 | 8921736.116 | 11409815.3 | 8451738.71 | 10756575.36 | 10121679.34 | 11446826.56 | 12088788.85 |
| Norepinephrine | 3438003.626 | 3750745.522 | 3278526.591 | 3284863.507 | 3594129.502 | 3850420.534 | 3047275.102 | 3804703.496 | 3592081.076 | 3965350.544 | 3464301.597 | 3285895.667 |

| Metabolites | PBS_1 | PBS_2 | PBS_3 | PBS_4 | BCG_1 | BCG_2 | BCG_3 | BCG_4 | BCGΔ*pe18*_1 | BCGΔ*pe18*_2 | BCGΔ*pe18*_3 | BCGΔ*pe18*_4 |
|---|---|---|---|---|---|---|---|---|---|---|---|---|
| PC(17:0/22:6(4Z,7Z,10Z,13Z,16Z,19Z)) | 909896.6717 | 937691.6569 | 1331240.304 | 1408415.208 | 1957167.543 | 1793725.788 | 2193997.907 | 1776973.646 | 2551090.961 | 2632374.106 | 2423788.259 | 1537222.746 |
| N-[4-(Dimethylamino)phenyl]-N'-(4-methoxyphenyl)urea | 11654434.45 | 11819158.44 | 11750438.04 | 8956297.117 | 11486594.71 | 11643370.71 | 11748064.68 | 11603713.95 | 11766284.86 | 11743719.33 | 2036039.386 | 11816963.97 |
| Gly-Val | 362064.055 | 1181682.451 | 396037.9052 | 2038506.232 | 21710069.42 | 14545470.35 | 30174702.24 | 14814888.24 | 13059778.94 | 13192396.26 | 12278187.13 | 11861181.25 |
| Arginine ethyl ester | 1934550.08 | 1806555.042 | 3078796.593 | 2131506.663 | 3870861.111 | 1835729.402 | 2016618.814 | 2022943.407 | 1995754.436 | 1934039.147 | 2382116.455 | 1865777.306 |
| Enterodiol | 1253017.627 | 1217900.257 | 1566701.931 | 1386842.131 | 764565.7352 | 876007.2805 | 432191.6536 | 467892.6164 | 418927.4892 | 477295.4846 | 882738.4817 | 349295.3028 |
| Telmisartan | 3118384.446 | 3460886.119 | 3723559.278 | 3819877.844 | 4030733.331 | 3453665.35 | 3755171.179 | 3545196.181 | 3821731.882 | 3863721.603 | 4034075.678 | 3793456.57 |
| Oleamide | 2350763485 | 2343550037 | 1876869598 | 1549712778 | 1427315351 | 1215490753 | 1392781813 | 1860156447 | 1443901106 | 1382369511 | 1374124335 | 831038899.2 |
| 4-Methylbenzenesulfonic acid | 27124037.05 | 8445099.238 | 8403725.195 | 13812610.58 | 26511673.85 | 10103426.77 | 9348194.866 | 8827684.774 | 23988265.62 | 8148720.005 | 8345641.764 | 18001978.36 |
| SHexCer 39:4;3O | 580053.0894 | 774798.1205 | 599426.9063 | 559854.8177 | 1008944.428 | 706285.894 | 1129935.272 | 1188575.502 | 1497049.021 | 1488982.376 | 777146.1795 | 789437.7975 |
| N-(6-Amino-1-benzyl-2,4-dioxo-1,2,3,4-tetrahydropyrimidin-5-yl)-N-methylbe | 12389706.27 | 12056039.36 | 10256831.15 | 10805117.01 | 11090896.09 | 11353112.29 | 9232024.263 | 11580535.29 | 11828196.43 | 9894639.566 | 10474892.23 | 11070259.05 |

| **Metabolites** | **PBS_1** | **PBS_2** | **PBS_3** | **PBS_4** | **BCG_1** | **BCG_2** | **BCG_3** | **BCG_4** | **BCGΔ *pe18*_1** | **BCGΔ *pe18*_2** | **BCGΔ *pe18*_3** | **BCGΔ*pe18*_4** |
|---|---|---|---|---|---|---|---|---|---|---|---|---|
| nzamide | | | | | | | | | | | | |
| PEtOH 16:0_26:4 | 685218.2534 | 276875.4734 | 1084449.473 | 1084834.6 | 551988.7296 | 4932.132456 | 466646.3647 | 479365.6924 | 664362.5686 | 594024.485 | 379409.0252 | 579923.0536 |
| Kickxioside | 328552.3793 | 372894.66 | 296890.916 | 366959.8466 | 365674.8128 | 341568.5823 | 367039.8891 | 323611.1798 | 388546.4001 | 338178.1821 | 344221.4638 | 339359.7552 |
| Brimonidine | 207998.3261 | 0 | 476825.8093 | 70848.14435 | 821199.5203 | 1066090.564 | 4221471.236 | 3616256.208 | 1671298.959 | 617142.8479 | 1467704.463 | 882833.8388 |
| 1-(1,2-Dioctanoylphosphatidyl)inositol-5-phosphate | 206702.6083 | 378596.7419 | 421742.4035 | 162152.9036 | 481059.3228 | 152546.8417 | 309717.5452 | 557299.234 | 485107.0855 | 282675.7154 | 524596.219 | 259477.6612 |
| Ricinoleic acid methyl ester | 931715.3126 | 2141151.25 | 1066317.006 | 1452262.797 | 4962101.814 | 8269643.599 | 5000814.045 | 5392491.778 | 4896421.431 | 4079265.485 | 9995325.9 | 6545252.267 |
| Regorafenib (BAY 73-4506) | 1122397.871 | 1078426.723 | 1014376.232 | 1134638.154 | 1116135.058 | 1199110.86 | 1164108.387 | 1079596.747 | 1129310.637 | 1255748.439 | 1111705.467 | 1014321.45 |
| MGDG O-9:0_18:3 | 7484932.563 | 488710.7442 | 1278155.679 | 853671.698 | 1495962.714 | 1510377.585 | 1971533.131 | 4080715.781 | 1133245.244 | 1638931.744 | 1946469.43 | 877438.3508 |
| Docosanamide | 135588829.9 | 92980125.11 | 87025969.79 | 79693406.72 | 29972484.34 | 60360098.42 | 31696937.27 | 39209281.16 | 44880342.72 | 53028901.58 | 68872691.2 | 61104885.06 |
| Aspartate | 3673965.307 | 3212074.122 | 3638799.618 | 3561429.452 | 14320016.74 | 7282834.301 | 11236244.48 | 10609175.48 | 7301313.052 | 6503418.852 | 7317706.197 | 6546002.032 |
| 5-Undecyl-1,3,4-thi | 169224 | 190037 | 154028 | 183865 | 169186 | 166656 | 171536 | 170644 | 203842 | 166889 | 157948 | 144390 |

| Metabolites | PBS_1 | PBS_2 | PBS_3 | PBS_4 | BCG_1 | BCG_2 | BCG_3 | BCG_4 | BCGΔ*pe18*_1 | BCGΔ*pe18*_2 | BCGΔ*pe18*_3 | BCGΔ*pe18*_4 |
|---|---|---|---|---|---|---|---|---|---|---|---|---|
| adiazol-2-ylamine | 8.144 | 1.098 | 1.242 | 9.672 | 4.845 | 1.419 | 8.914 | 7.791 | 4.58 | 8.281 | 8.178 | 1.72 |
| PG 16:1_17:1 | 55200.4771 | 38698.34553 | 73205.37796 | 64166.23489 | 1385343.709 | 562118.0687 | 1065242.531 | 841288.7002 | 1566657.36 | 1845993.755 | 1163722.272 | 638142.3246 |
| Bazedoxifene | 1676500.346 | 1271697.877 | 2074596.102 | 1862428.592 | 1576628.128 | 1971431.405 | 1721639.146 | 1831723.907 | 1500202.927 | 1419935.436 | 1885064.853 | 1918299.168 |
| 4-(Propan-2-yl)piperidine | 5490370.009 | 5244057.7 | 4779708.527 | 5411232.132 | 5751750.588 | 5445210.408 | 5225147.578 | 5975085.43 | 16092508.73 | 4871539.869 | 5509569.422 | 5917332.42 |
| Ethanone, 1-(3,4-dihydroxyphenyl)-2-(2-ethyl-1H-imidazol-1-yl)- | 1339988.115 | 1430119.872 | 1176826.654 | 1311402.756 | 1298393.683 | 1333239.549 | 1174670.694 | 1328689.939 | 1295846.501 | 1289834.817 | 1270951.622 | 1040017.085 |
| PI 16:0_18:2 | 1645016.154 | 1702340.381 | 1969468.792 | 2586291.597 | 5839551.062 | 4546961.959 | 6487478.85 | 5345689.739 | 5969059.051 | 7313613.978 | 10006392.9 | 4785784.004 |
| 3-Acetamidopropanal | 4428501.977 | 5329330.235 | 4076894.15 | 3672912.801 | 5324526.392 | 4760782.714 | 3727224.506 | 3813247.772 | 3742081.164 | 3455173.863 | 4808667.017 | 2665123.621 |
| NAE 12:0 | 1827416.545 | 1914922.452 | 1961182.846 | 1901969.124 | 1875950.682 | 1872874.66 | 1908368.11 | 1897889.881 | 1797283.905 | 1848589.389 | 1799382.547 | 1938915.732 |
| [4-Anilino-2-(4-fluorophenyl)-5-oxo-1-phenyl-2,5-dihydro-1H-pyrrol-3-yl]acetic acid | 97547.5686 | 76163.52406 | 170829.4727 | 80623.58212 | 326580.2475 | 357141.86 | 343471.7485 | 341838.0426 | 286459.6789 | 335358.8677 | 401081.1202 | 304712.8266 |
| Isopropalin | 301105 | 266855 | 560544 | 284348 | 877482 | 574052 | 874280 | 430624 | 767007 | 824240 | 756386 | 935124 |

| **Metabolites** | **PBS_1** | **PBS_2** | **PBS_3** | **PBS_4** | **BCG_1** | **BCG_2** | **BCG_3** | **BCG_4** | **BCGΔ*pe18*_1** | **BCGΔ*pe18*_2** | **BCGΔ*pe18*_3** | **BCGΔ*pe18*_4** |
|---|---|---|---|---|---|---|---|---|---|---|---|---|
| | 3.691 | 0.996 | 4.428 | 1.079 | 8.666 | 7.067 | 7.847 | 5.857 | 4.503 | 5.517 | 5.229 | 5.974 |
| Tris(2-ethylhexl)phosphate | 627296.5003 | 606434.917 | 583771.2526 | 564855.5345 | 565945.3347 | 706028.3838 | 808201.9282 | 578601.0404 | 583497.8813 | 755578.5775 | 849665.7222 | 421554.2602 |
| HexCer 13:0;2O/3:0 | 2262974.76 | 2542107.07 | 1886711.71 | 2012184.337 | 2013976.362 | 2406077.85 | 2152309.989 | 2217175.218 | 2511768.189 | 2297598.337 | 2229907.304 | 1976196.558 |
| Triethyl phosphate | 244425.2818 | 242651.3082 | 280532.668 | 276001.6844 | 332716.786 | 295853.6129 | 284803.1687 | 309358.1898 | 269841.1767 | 329419.9748 | 334977.4457 | 770064.1695 |
| Pterine | 232899.1274 | 111457.9803 | 496450.5414 | 242711.6324 | 10937536.84 | 7007999.929 | 17013232.91 | 14449622.78 | 8115233.116 | 9547492.792 | 6225332.973 | 5942941.116 |
| epicatechin gallate | 210444.466 | 212841.6786 | 212488.5211 | 214660.0977 | 212205.162 | 211366.9579 | 218266.4187 | 214123.0661 | 209948.3952 | 205562.7349 | 216155.0007 | 224223.7644 |
| FAHFA 18:1/18:0 | 429409.1218 | 460682.1634 | 608356.7092 | 411979.1998 | 89658144.53 | 178657732.9 | 95644516.64 | 108858393.2 | 54704530.61 | 16190828.23 | 84474786.61 | 71608560.49 |
| Evocarpine | 2052404.345 | 2587821.699 | 3140966.061 | 5151627.542 | 1330990.332 | 3614094.919 | 2813692.52 | 1694190.222 | 1487791.942 | 1403130.298 | 1417894.288 | 1881732.443 |
| Ketoleucine | 49166107.3 | 32297609.34 | 81815478.44 | 14211203.85 | 15056853.27 | 18693184.18 | 68481034.26 | 67389042 | 27211585.63 | 19030985.36 | 17293606.46 | 37293643.1 |
| Isobutyric_acid | 1166984.345 | 1147680.985 | 1257584.42 | 1175493.973 | 1363865.888 | 1379522.841 | 1400198.808 | 1346578.136 | 1207653.634 | 1369870.562 | 1382759.236 | 1697952.057 |
| Noscapine | 1154224.293 | 1020598.329 | 1680028.405 | 941712.1038 | 1084246.111 | 1886418.035 | 1768592.185 | 1870051.538 | 2045430.138 | 2119566.383 | 2351865.879 | 1831370.442 |
| PS O-24:6_17:1 | 803977.2851 | 1361509.419 | 966732.3011 | 1061346.8 | 343243.5179 | 826001.9844 | 373290.6326 | 433722.0599 | 261588.06 | 375319.5253 | 476189.9912 | 1275685.85 |
| PA 2:0_13:1 | 420639. | 188662 | 577754. | 453367. | 101596 | 664370. | 863825. | 694220. | 602166. | 191597. | 226866. | 106257 |

| Metabolites | PBS_1 | PBS_2 | PBS_3 | PBS_4 | BCG_1 | BCG_2 | BCG_3 | BCG_4 | BCGΔ*pe18*_1 | BCGΔ*pe18*_2 | BCGΔ*pe18*_3 | BCGΔ*pe18*_4 |
|---|---|---|---|---|---|---|---|---|---|---|---|---|
| | 9146 | 4.872 | 6182 | 5176 | 6.607 | 0711 | 2892 | 1966 | 4538 | 0279 | 1082 | 1.626 |
| 2-Chloro-5-nitrophenol | 483438.4741 | 521125.507 | 466833.3126 | 484319.062 | 516123.4738 | 485008.3602 | 513233.0027 | 428282.1032 | 541534.0028 | 461488.6717 | 495850.1411 | 484792.678 |
| (4aS,7aS)-6-Methyloctahydro-1H-pyrrolo[3,4-b]pyridine | 4378103.735 | 4177069.478 | 4109468.494 | 3570120.265 | 559854.3765 | 524524.2756 | 470350.1866 | 535686.9733 | 383629.5454 | 322106.1943 | 622835.9883 | 467479.3326 |
| PE O-18:4_28:7 | 889764.7863 | 876801.1791 | 1009712.44 | 870321.1437 | 786066.8994 | 958541.1885 | 938902.5407 | 962333.9138 | 830858.084 | 814659.1887 | 741255.7573 | 841771.892 |
| Pluripotin | 0 | 0 | 14091.74706 | 12038.15761 | 614339.833 | 752174.6004 | 567718.229 | 405249.8442 | 345904.2692 | 319464.3897 | 417914.3691 | 255741.9475 |
| D-Saccharic acid 1,4-lactone | 665756.4767 | 754631.8122 | 651902.2271 | 699108.9524 | 1001780.76 | 599898.0379 | 872878.0348 | 787141.9562 | 812779.2366 | 868438.3762 | 777199.5297 | 756392.1986 |
| 5'-Fluoro-2'-hydroxy-4-methylchalcone | 5373100.666 | 5183105.192 | 8953971.267 | 7070901.107 | 5981974.616 | 6944997.335 | 6586019.608 | 6277650.424 | 6023439.81 | 4688564.048 | 7290349.358 | 6816055.318 |
| 11,12,15-Trihydroxyicosa-5,8,13-trienoylcarnitine | 907385.0952 | 473438.1924 | 715701.4213 | 472538.9679 | 1066399.343 | 934990.5144 | 994639.1168 | 1478358.135 | 803589.0212 | 726031.5192 | 601484.3015 | 742551.4704 |
| Isoleucylvaline | 1886417.618 | 2280538.003 | 2029378.833 | 2067941.138 | 1592053.132 | 1751771.277 | 2389200.049 | 1786090.977 | 1777014.856 | 1593869.414 | 1750383.963 | 2937439.167 |
| LPC 22:1-SN2 | 421020.2579 | 117912.5642 | 341292.7154 | 358165.6864 | 1583096.099 | 1750889.966 | 1795141.728 | 1293528.472 | 1342597.627 | 1069324.66 | 1228609.741 | 1018542.686 |
| 1-(2-Methoxyethyl)piperazine | 3720199.07 | 4505904.296 | 2100616.917 | 4326570.075 | 4831153.768 | 3248098.346 | 3230787.262 | 2822340.368 | 2860841.542 | 1645817.102 | 2516923.287 | 5492249.279 |
| Dehydrolithocholic | 205789 | 147167 | 136299 | 258369 | 235659 | 188957 | 258078 | 199200 | 105696 | 788091. | 137839 | 218020 |

| Metabolites | PBS_1 | PBS_2 | PBS_3 | PBS_4 | BCG_1 | BCG_2 | BCG_3 | BCG_4 | BCGΔ *pe18*_1 | BCGΔ *pe18*_2 | BCGΔ *pe18*_3 | BCGΔ*pe18*_4 |
|---|---|---|---|---|---|---|---|---|---|---|---|---|
| acid | 6.027 | 0.429 | 1.302 | 2.029 | 0.125 | 1.781 | 2.02 | 6.955 | 6.822 | 9642 | 2.522 | 5.409 |
| N'-[(1E)-(5-Chloro-2-hydroxyphenyl)methylene]-2-morpholin-4-ylacetohydrazide | 3658535.32 | 3095476.302 | 3637870.471 | 3431396.338 | 33555925.96 | 29265877.65 | 28987848.22 | 27902812.47 | 30278718.69 | 25818115.69 | 29777262.96 | 26904911.34 |
| 2,4,14-Eicosatrienoic acid isobutylamide | 1714516.829 | 1536639.327 | 1513114.126 | 1551824.243 | 1487114.808 | 1529686.493 | 1602640.617 | 1535392.919 | 1592480.728 | 1547172.689 | 1485503.619 | 1569294.552 |
| HexCer 18:1;2O/24:1 | 1896847.196 | 2525451.386 | 3030909.752 | 3111194.101 | 1757317.364 | 4687417.252 | 1940569.04 | 2232960.605 | 1954339.162 | 1404188.837 | 666940.2268 | 3140882.671 |
| 22,23-Dihydro-alpha-spinasterone | 156709.4085 | 0 | 136886.9724 | 0 | 179296.0922 | 123340.587 | 384816.9837 | 323213.019 | 189605.1707 | 145941.9577 | 88449.63344 | 110181.7261 |
| 1-Hexadecyl lysophosphatidic acid | 475543.599 | 357439.3695 | 608116.9884 | 481307.4106 | 702914.4355 | 555257.2822 | 702518.0452 | 607839.0566 | 372635.737 | 462868.1845 | 600653.5906 | 372166.0966 |
| Everzol Orange BN | 38676.05889 | 57957.34212 | 62104.17625 | 22057.23521 | 377607.0624 | 336138.0173 | 299193.4374 | 302244.9356 | 234242.6233 | 189812.0658 | 185814.1788 | 220639.7752 |
| 7-[(E)-5-(3,6-Dihydroxy-2,2,6-trimethylcyclohexyl)-3-methylpent-2-enoxy]chromen-2-one | 401596.1647 | 430032.3107 | 316729.2089 | 397512.8289 | 478504.407 | 420838.6352 | 414051.4281 | 461907.3805 | 402049.6291 | 515355.649 | 420196.3622 | 392766.0794 |
| PE O-18:2_20:4 | 154480 | 104534 | 167212 | 125368 | 156515 | 40388.3 | 168656 | 164523 | 138915 | 293964 | 145076 | 103586 |

| Metabolites | PBS_1 | PBS_2 | PBS_3 | PBS_4 | BCG_1 | BCG_2 | BCG_3 | BCG_4 | BCGΔ *pe18*_1 | BCGΔ *pe18*_2 | BCGΔ *pe18*_3 | BCGΔ*pe18*_4 |
|---|---|---|---|---|---|---|---|---|---|---|---|---|
| | 4.19 | 8.458 | 6.602 | 7.28 | 3.252 | 8532 | 5.752 | 4.085 | 9.706 | 4.918 | 7.816 | 6.617 |
| PE(18:4(6Z,9Z,12Z,15Z)/P-18:0) | 28483262.57 | 33770743.42 | 33164479.59 | 27084520.28 | 38243303.72 | 21297278.8 | 32450535.31 | 24161836.83 | 23206007.64 | 36657964.38 | 42677217.84 | 14251390.19 |
| ST 24:1;O3;G/22:5 | 2430819.394 | 864078.0497 | 3104141.316 | 1695355.424 | 1443828.768 | 5049.660116 | 1252987.223 | 660882.5009 | 1743018.314 | 1404590.689 | 1594053.752 | 870030.816 |
| 2,2,5,7,8-Pentamethyl-6-hydroxychroman | 787995.221 | 875150.9503 | 794203.9196 | 844056.1971 | 756264.626 | 659248.7936 | 754920.5879 | 665982.5758 | 674911.8143 | 674934.8024 | 675710.8874 | 846640.6015 |
| 3.beta.,7.alpha.-Dihydroxy-5-cholestenoic acid | 489315.4567 | 510389.8138 | 586164.4611 | 635294.3596 | 975828.5734 | 895209.4559 | 922349.6314 | 936238.2762 | 626914.9612 | 371867.8358 | 400467.2164 | 528713.0615 |
| Threonate | 78586.40967 | 96166.95243 | 91198.74903 | 251010.9477 | 2577789.841 | 470951.6018 | 3020286.622 | 2054892.87 | 1170060.309 | 1622973.927 | 1596137.026 | 1261239.087 |
| 7-[3-[(E,3R)-3-Hydroxyoct-1-enyl]-6-thiabicyclo[3.1.1]heptan-4-yl]hept-5-enoic acid | 2857220.88 | 1465590.162 | 1293276.997 | 1377195.694 | 1001076.23 | 1287152.66 | 626742.4999 | 1682123.797 | 969862.1878 | 1105061.74 | 1187865.962 | 2776256.726 |
| Ethyl 7-chloro-1-(2,4-difluorophenyl)-6-fluoro-4-oxo-1,4-dihydro[1,8]naphthyridine-3-carboxylate | 181305.093 | 141798.4818 | 329359.1377 | 183151.7258 | 2227308.519 | 1705497.162 | 2409988.814 | 2130719.825 | 2016551.966 | 1620538.812 | 1826196.686 | 1597385.188 |

| Metabolites | PBS_1 | PBS_2 | PBS_3 | PBS_4 | BCG_1 | BCG_2 | BCG_3 | BCG_4 | BCGΔ*pe18*_1 | BCGΔ*pe18*_2 | BCGΔ*pe18*_3 | BCGΔ*pe18*_4 |
|---|---|---|---|---|---|---|---|---|---|---|---|---|
| [(7-Oxo-7H-benzo[de]anthracen-3-yl)sulfanyl]acetic acid | 79304.17185 | 95170.88134 | 357028.1259 | 277534.6278 | 3166792.553 | 1356581.221 | 2479276.516 | 1783238.612 | 2377794.495 | 1930015.144 | 2662028.929 | 1457110.026 |
| 2',6'-Dihydroxyacetophenone | 915165.9202 | 740263.8129 | 782693.1935 | 932905.0078 | 944553.2178 | 785908.9465 | 923158.334 | 943421.5404 | 854376.671 | 933475.4828 | 783359.2076 | 832788.861 |
| M406T265 | 570343.6468 | 365687.4882 | 375452.0553 | 336953.425 | 346685.0804 | 306743.7075 | 372302.3008 | 346763.055 | 366424.3026 | 347709.4711 | 400417.5437 | 371715.5293 |
| 2-(4-Piperidin-4-ylcarbonyl)-1,2,3,4-tetrahydroisoquinoline | 2090806.14 | 2348986.352 | 1454107.001 | 2073361.106 | 2207717.193 | 1583896.352 | 2018773.799 | 2392809.489 | 2258420.441 | 2144583.708 | 1932406.308 | 1659000.729 |
| FAHFA 2:0/16:2 | 1144592.097 | 1164574.734 | 815119.6611 | 900701.7998 | 929802.4279 | 1793379.856 | 894614.9028 | 979497.3762 | 783591.5256 | 765417.9156 | 1093678.62 | 915009.9117 |
| 3-Oxododecanoic acid | 3985136.364 | 3906452.914 | 4306440.485 | 4862786.748 | 4795493.826 | 4621869.204 | 3988764.717 | 4199229.262 | 4651947.443 | 5086300.934 | 4325958.618 | 4233021.708 |
| MGDG O-9:0_24:1 | 59092.35593 | 106981.5793 | 57305.32587 | 85577.0195 | 2467324.519 | 6319382.558 | 2259958.219 | 2552469.724 | 4153916.552 | 2411156.503 | 2100326.522 | 1236238.874 |
| 3,5-Dihydroxyacetophenone | 915165.9202 | 740263.8129 | 782693.1935 | 932905.0078 | 944553.2178 | 785908.9465 | 923158.334 | 943421.5404 | 854376.671 | 933475.4828 | 783359.2076 | 832788.861 |
| 3-(3-Hydroxyphenyl)-2-phenyl-4(3H)-quinazolinone | 1351794.34 | 1363229.57 | 1363523.774 | 1370281.869 | 1311261.396 | 1366786.125 | 1350311.83 | 1390075.576 | 1369777.203 | 1483586.151 | 1520945.601 | 1265455.585 |
| Isocycloheximide | 2129823.756 | 2556061.099 | 2062898.651 | 2264976.703 | 1799865.01 | 1887265.244 | 1839568.05 | 1771949.881 | 1900169.209 | 1759152.139 | 1712379.141 | 2111064.003 |
| 2,2-Dimethylglutari | 511740. | 278729. | 294972. | 496645. | 230399 | 199766 | 220890 | 301417 | 190095 | 590375. | 105584 | 101525 |

| **Metabolites** | **PBS_1** | **PBS_2** | **PBS_3** | **PBS_4** | **BCG_1** | **BCG_2** | **BCG_3** | **BCG_4** | **BCGΔ*pe18*_1** | **BCGΔ*pe18*_2** | **BCGΔ*pe18*_3** | **BCGΔ*pe18*_4** |
|---|---|---|---|---|---|---|---|---|---|---|---|---|
| c acid | 4522 | 1608 | 842 | 1361 | 6.716 | 5.083 | 5.129 | 1.584 | 5.527 | 2716 | 8.384 | 0.681 |
| gamma-Glutaminyl-4-hydroxybenzene | 8910013.879 | 9599517.598 | 128278815.7 | 10022424.46 | 13781512.18 | 10155935.66 | 195163281.4 | 16198746.37 | 9528207.986 | 119369139.6 | 126748671.2 | 9920007.18 |
| CE 20:2 | 3678229.679 | 3693729.186 | 2416728.06 | 1625347.653 | 682456.7098 | 791284.3563 | 1164918.596 | 1275462.492 | 1391976.124 | 764909.6623 | 666434.4749 | 622013.7268 |
| Isophorone | 318369.4483 | 308816.0148 | 319980.6169 | 258660.9471 | 540100.9728 | 1601348.06 | 8855815.075 | 411294.4571 | 5167812.629 | 1927224.238 | 572448.1745 | 7580758.797 |
| Thamnosin | 363424.5424 | 161539.8748 | 215168.1029 | 345201.4535 | 774657.1271 | 765619.2529 | 591584.9595 | 583935.7784 | 577116.2462 | 500788.5763 | 450663.2884 | 425379.325 |
| 4-(N-Methylacetamido)benzoic acid | 435005.1751 | 1148727.199 | 524429.5856 | 1474930.623 | 773408.5443 | 771334.319 | 757957.8563 | 279828.633 | 500532.7381 | 515553.6731 | 463444.3134 | 456833.4594 |
| 4-Hydrazino-5-phenylthieno[2,3-d]pyrimidine | 50670926.32 | 49681584.53 | 49170820.66 | 46771188.9 | 45544424.82 | 38779822.44 | 43986987.72 | 45763580.1 | 45143500.97 | 42835018.64 | 44557441.99 | 44527835.25 |
| Ginsenoside F2 | 1160374.134 | 489812.2372 | 224107.8419 | 486383.0416 | 1620835.767 | 556023.7366 | 1106179.193 | 1456688.894 | 1748843.623 | 2031385.908 | 945210.398 | 1080169.926 |
| PE(20:5(5Z,8Z,11Z,14Z,17Z)/22:5(7Z,10Z,13Z,16Z,19Z)) | 1426038.653 | 1664986 | 1207446.143 | 1373631.499 | 2220203.925 | 1748117.771 | 1422359.124 | 2063106.72 | 2238565.258 | 2069891.785 | 1737607.363 | 1016926.068 |
| Lotaustralin | 772069.5132 | 579551.7121 | 876663.0021 | 690953.4407 | 2115404.192 | 1053126.575 | 1983275.355 | 1238096.97 | 975863.9833 | 1205007.437 | 1252805.729 | 934446.488 |
| Robinin | 71955.94454 | 66270.9383 | 42686.45141 | 56524.60096 | 29488.19465 | 53137.10682 | 41781.39927 | 65002.95348 | 45919.45578 | 44748.25599 | 50359.45205 | 32618.77064 |

| **Metabolites** | **PBS_1** | **PBS_2** | **PBS_3** | **PBS_4** | **BCG_1** | **BCG_2** | **BCG_3** | **BCG_4** | **BCGΔ*pe18*_1** | **BCGΔ*pe18*_2** | **BCGΔ*pe18*_3** | **BCGΔ*pe18*_4** |
|---|---|---|---|---|---|---|---|---|---|---|---|---|
| Indolylmethyl-desulfoglucosinolate | 35570148.24 | 22045450.6 | 31303880.8 | 23982818.12 | 14053199.93 | 9986498.428 | 24667622.47 | 19793976.35 | 7423356.514 | 8757191.029 | 9405293.817 | 5529172.339 |
| Hex2Cer 24:4;2O | 142797.3563 | 129688.564 | 95094.19625 | 127485.6222 | 125478.354 | 78589.2108 | 111176.0647 | 101851.8957 | 104288.0043 | 133664.5618 | 132356.3232 | 104665.843 |
| Laudanosine | 675416.6953 | 667742.8243 | 567568.2741 | 586070.9742 | 695875.0648 | 708354.7544 | 651951.9455 | 688631.3907 | 639159.0665 | 698946.2511 | 660922.3368 | 627124.691 |
| 1-Tetrahydro-3-furanyl-4-piperidinamine | 3530538.661 | 5679405.948 | 4702937.613 | 4354956.772 | 4618107.628 | 4382935.547 | 4453606.013 | 5256973.581 | 4368772.887 | 4294214.613 | 4334080.852 | 4710747.838 |
| Lofemizole | 5362516.851 | 5231059.3 | 4679166.371 | 5070791.698 | 5808928.363 | 5588293.653 | 5477887.353 | 5503822.411 | 5199167.596 | 5403300.543 | 5493354.599 | 4855191.504 |
| Cer 17:0;2O/16:0 | 369067.8174 | 301566.9367 | 337459.8793 | 431663.7409 | 498700.5482 | 246734.1542 | 423645.4697 | 409756.5994 | 330008.8009 | 314528.0859 | 352781.5958 | 765230.1473 |
| Apabetalone | 9277971.602 | 9785779.876 | 9248605.605 | 8051465.21 | 6024503.657 | 5737089.489 | 6155486.643 | 6912378.122 | 6216185.895 | 6675861.185 | 5826356.076 | 10106505.72 |
| TG 8:0_16:4_26:7 | 748611.4337 | 977569.1126 | 397727.6228 | 673562.1988 | 963174.8048 | 1223742.492 | 1150698.2 | 1087832.258 | 1288465.168 | 1232164.084 | 1133123.316 | 965965.9023 |
| 2-Amino-5-(4-bromophenyl)-3-furonitrile | 7087120.411 | 6912275.319 | 7413771.606 | 6688936.181 | 6182145.725 | 6898033.091 | 6282879.016 | 4773748.129 | 4653182.218 | 3701472.513 | 4856808.19 | 5417655.759 |
| SL 11:1;O/18:2 | 527738.5974 | 502744.7741 | 461220.0089 | 530761.975 | 463149.7336 | 524563.2424 | 520244.6178 | 537678.5269 | 495244.9429 | 549307.3476 | 485307.5054 | 530477.1818 |
| PC(44:12) | 2849328.706 | 3790609.05 | 1263967.463 | 2456379.167 | 4608368.874 | 3369738.837 | 3903372.181 | 4827836.386 | 5246468.125 | 4282710.959 | 3378004.283 | 2861885.438 |

| Metabolites | PBS_1 | PBS_2 | PBS_3 | PBS_4 | BCG_1 | BCG_2 | BCG_3 | BCG_4 | BCGΔ*pe18*_1 | BCGΔ*pe18*_2 | BCGΔ*pe18*_3 | BCGΔ*pe18*_4 |
|---|---|---|---|---|---|---|---|---|---|---|---|---|
| 5,6-Epoxy-5,6-dihydro-10'-apo-b,y-carotene-3,10'-diol | 15882700.41 | 13949953.77 | 22629354.1 | 18199231.33 | 21561841.94 | 17865485.97 | 22958504.6 | 12946930.93 | 16427035.92 | 10989728.15 | 15293986.59 | 13135749.95 |
| Hydroxybutorphanol | 1689358.153 | 2113659.575 | 1708472.206 | 1723569.757 | 1730512.518 | 1840841.956 | 1759891.456 | 2163962.512 | 1728963.597 | 1738199.446 | 1963961.252 | 2217714.914 |
| PS 21:0_16:1 | 323426.9112 | 236813.3149 | 283264.7996 | 265551.9592 | 398836.4063 | 0 | 645970.9329 | 4986679.4 | 1545610.536 | 425984.9067 | 277958.1618 | 1052194.422 |
| (1(10)E,4E,6a,9b)-9-(2-Methylpropanoyloxy)-1(10),4,11(13)-germacratrien-12,6-olide | 3154114 | 2534878.273 | 1863866.615 | 2081500.125 | 2171904.714 | 2736431.203 | 2374523.783 | 2880581.464 | 1752266.026 | 2490605.593 | 2010280.029 | 2234269.812 |
| 2-(3-Bromo-5-ethoxy-4-hydroxybenzylidene)malononitrile | 3451076.62 | 3029691.422 | 5032632.158 | 3287519.892 | 7644726.915 | 6224057.324 | 6744446.954 | 7469833.656 | 7166878.519 | 6992196.322 | 5211855.074 | 5614522.859 |
| FAHFA 14:0/22:3 | 267713.2575 | 155206.7693 | 356438.6725 | 298006.9679 | 25322515.83 | 36496632.24 | 22773708.46 | 29578034.13 | 15276732.49 | 6038437.017 | 23056626.61 | 19136984.77 |
| LPC(O-14:0) | 699093.9915 | 2241494.139 | 2405625.209 | 1922179.204 | 742588.4068 | 381560.6017 | 9285616.498 | 565691.587 | 383814.195 | 110542.9943 | 476926.6482 | 6225754.632 |
| NAGly 20:5/22:5 | 5475684.635 | 4163454.507 | 4395136.199 | 4771543.77 | 27179724.45 | 33670084.69 | 28358847.62 | 22778201.7 | 26921296.68 | 28406390.85 | 38167877.04 | 18807350.51 |

| Metabolites | PBS_1 | PBS_2 | PBS_3 | PBS_4 | BCG_1 | BCG_2 | BCG_3 | BCG_4 | BCGΔ*pe18*_1 | BCGΔ*pe18*_2 | BCGΔ*pe18*_3 | BCGΔ*pe18*_4 |
|---|---|---|---|---|---|---|---|---|---|---|---|---|
| 2-[(3-Fluoro-4-methylphenyl)amino]-4-methylquinolin-5-ol | 5226135.043 | 4901167.911 | 5121884.316 | 5285993.862 | 3876859.343 | 3226070.367 | 2704252.183 | 2893859.504 | 1019376.776 | 1091639.861 | 1922749.315 | 2668544.409 |
| Tetradonium cation | 13575442.2 | 14198653.16 | 8449291.146 | 12715436.74 | 15047420.49 | 12942555.36 | 9822072.28 | 15190881.87 | 16813049.68 | 9567796.61 | 9674313.744 | 15024271.37 |
| Adenosine 2',3'-cyclic phosphate | 133032.3017 | 126172.9382 | 215425.5102 | 282517.8788 | 2451188.681 | 2180034.986 | 2497620.802 | 2483095.469 | 1098384.36 | 1270865.957 | 1247375.109 | 1451103.579 |
| PG(16:1(9Z)/22:5(4Z,7Z,10Z,13Z,16Z)) | 3231303.005 | 3524531.647 | 1764845.413 | 3090246.896 | 2345936.642 | 2493646.261 | 2924394.671 | 2877713.424 | 3143393.762 | 3365546.781 | 3084811.34 | 2554717.357 |
| N-Methyldioctylamine | 4294604.258 | 5150948.375 | 2477329.842 | 3479051.423 | 1754755.143 | 1470116.651 | 2386953.449 | 1669406.224 | 2061590.872 | 2049497.944 | 1840278.272 | 1209839.987 |
| Tanshinone IIA | 95143.87441 | 72463.98675 | 38939.81752 | 103440.3927 | 1231260.089 | 1125873.833 | 939118.8997 | 908573.6376 | 755848.1764 | 509154.313 | 510479.3349 | 740264.3243 |
| FA 22:3 | 1238489.849 | 1161621.514 | 1773647.109 | 1201478.977 | 2646343.383 | 4690019.585 | 2940493.853 | 3026126.682 | 1872730.656 | 1480398.09 | 3247367.32 | 2248764.83 |
| N6-Methyladenosine | 175991.9279 | 180561.3307 | 197858.411 | 201163.2437 | 2489396.6 | 520537.3961 | 1074740.227 | 1930453.265 | 1622157.884 | 674971.531 | 1933680.218 | 573267.9048 |
| PC 16:0_34:3 | 19329.06171 | 15369.86845 | 33302.52191 | 0 | 262413.4696 | 59891.44713 | 229213.7096 | 36554.2822 | 251326.1526 | 124714.3296 | 329307.2298 | 0 |
| 5,6-Dinitroacenaphthoquinone | 1228235.706 | 884286.7548 | 1662716.836 | 1056056.911 | 2727354.448 | 1559980.773 | 2955212.645 | 2180900.163 | 1432176.985 | 2009980.141 | 2150445.997 | 1168417.569 |

| Metabolites | PBS_1 | PBS_2 | PBS_3 | PBS_4 | BCG_1 | BCG_2 | BCG_3 | BCG_4 | BCGΔ*pe18*_1 | BCGΔ*pe18*_2 | BCGΔ*pe18*_3 | BCGΔ*pe18*_4 |
|---|---|---|---|---|---|---|---|---|---|---|---|---|
| Paxilline | 21660670.38 | 1124584.816 | 28893420.01 | 33631706.06 | 71388809.75 | 31589521.24 | 50945948.77 | 45722036.72 | 37080977.8 | 30374557.71 | 31608529.43 | 1196867.924 |
| PC O-32:6 | 7071926.16 | 8660901.334 | 6609694.277 | 9463081.219 | 7201755.111 | 6648729.904 | 9005643.497 | 8487216.268 | 9246080.223 | 8497742.893 | 9507834.659 | 7582347.648 |
| Glycerophosphoethanolamine | 8267440.786 | 5169257.511 | 11365558.32 | 6606816.037 | 39635226.55 | 16562762.83 | 35131126.49 | 23586852.72 | 22328641.93 | 21338456.06 | 23236234.25 | 16444823.91 |
| phosphatidylglycerol (dioctadecanoyl, n-C18:0) | 2739135.999 | 1968252.605 | 2454244.86 | 2815978.439 | 2351549.598 | 78939.09561 | 2095697.62 | 1463561.097 | 3265357.175 | 2804770.312 | 2097871.234 | 2458963.198 |
| PC ae (16:0/22:6) | 1221479.668 | 55584809.79 | 25136650.5 | 38631985.43 | 9712886.868 | 10192987.12 | 9313767.855 | 12184174.33 | 13302277.36 | 12157523.89 | 10113874.81 | 10730440.28 |
| M256T374 | 1599316.017 | 1238725.798 | 1715561.986 | 1275505.503 | 9002328.438 | 3302603.611 | 6982230.447 | 5500064.445 | 5409353.24 | 4815503.995 | 4523499.386 | 2139328.11 |
| Benserazide | 3436636.814 | 12540745.22 | 8880140.351 | 1708736.175 | 11263464.67 | 80282213.5 | 2491110.101 | 2926544.87 | 6197150.503 | 13176047.6 | 2565651.286 | 21538066.88 |
| Cer 12:0;3O/20:2;(2OH) | 1644027.677 | 1957120.965 | 980019.8095 | 1423906.685 | 1507241.788 | 1812204.857 | 1386329.216 | 1505219.132 | 1376931.643 | 1362931.462 | 1580724.274 | 984441.6731 |
| SMGDG O-13:1_7:0 | 133497.0932 | 147219.4964 | 152740.8934 | 123420.4498 | 140244.6644 | 118805.0584 | 117735.5497 | 136841.6967 | 126219.5521 | 111802.0455 | 156125.1155 | 157821.3956 |
| Methyl arachidonyl fluorophosphonate | 3886396.569 | 3815374.872 | 3648755.431 | 3674518.581 | 3672718.114 | 3628881.655 | 3517826.375 | 3908542.068 | 3839621.812 | 3715667.83 | 3647758.511 | 3261558.029 |
| Biphenylindanone A | 576818.9333 | 780197.771 | 470081.0648 | 758726.5664 | 715266.374 | 799690.2516 | 887730.4347 | 677611.779 | 911438.9713 | 640553.5309 | 719220.7266 | 668473.1245 |

| Metabolites | PBS_1 | PBS_2 | PBS_3 | PBS_4 | BCG_1 | BCG_2 | BCG_3 | BCG_4 | BCGΔ*pe18*_1 | BCGΔ*pe18*_2 | BCGΔ*pe18*_3 | BCGΔ*pe18*_4 |
|---|---|---|---|---|---|---|---|---|---|---|---|---|
| SM 34:3;3O | 0 | 0 | 0 | 0 | 1826009.85 | 2959022.749 | 1881490.222 | 2119567.594 | 1187617.346 | 1087293.431 | 944858.7551 | 792151.8073 |
| Sebacic acid monomethyl ester | 986150.2017 | 1117890.019 | 1155765.518 | 1160811.218 | 946210.1249 | 1080359.119 | 1024804.44 | 976528.6853 | 1043197.53 | 996309.2398 | 1011230.841 | 1295632.467 |
| N-Lauroyl Serine | 1657302.936 | 1589430.235 | 1641827.827 | 1745567.708 | 1731826.223 | 1459276.534 | 1514776.158 | 1359790.149 | 1354572.446 | 1516925.326 | 1556807.543 | 1463909.22 |
| PS 22:4 | 0 | 0 | 0 | 0 | 137778.7019 | 69795.66382 | 124283.1155 | 89721.17273 | 104589.2795 | 92725.66908 | 139243.5327 | 114727.1781 |
| N,N-Diethyl-2-aminoethanol | 687691.6821 | 1485823.886 | 3318482.095 | 2429523.515 | 2093744.183 | 902793.8025 | 764669.0457 | 890150.1781 | 931686.3134 | 988334.58 | 633009.6449 | 4247092.664 |
| Glutaric acid | 11583159.92 | 8384527.356 | 9487275.246 | 11480896.95 | 94593857.98 | 46266627.98 | 73122940.06 | 64257037.67 | 40316647.55 | 40379479.2 | 51073174.07 | 34777838.44 |
| Dodecenoylcarnitine (Car(12:1)) | 620007.1303 | 573040.4455 | 727500.2309 | 760134.6566 | 1181621.419 | 1284450.896 | 1260267.26 | 1092679.551 | 906590.5396 | 479854.7563 | 1107594.783 | 1029263.402 |
| (E,E)-farnesylacetone | 4566418.246 | 5073653.444 | 4395930.313 | 4642174.485 | 4902115.133 | 4881237.207 | 4928054.693 | 4701627.718 | 4662656.594 | 4917688.006 | 4804689.543 | 4479426.043 |
| Cyclohexanamine | 2169258.777 | 4024794.576 | 3598951.826 | 3649586.313 | 3934967.284 | 3832031.087 | 4208198.936 | 2293956.488 | 3990975.642 | 4452492.382 | 957910.3041 | 633296.1399 |
| PE-Cer 12:0;2O/24:1 | 878745.4991 | 633814.2983 | 867302.4664 | 849586.7338 | 1084821.469 | 0 | 731608.8568 | 673005.2339 | 999551.9267 | 797523.0319 | 1020740.789 | 408209.629 |
| MGDG O-8:0_12:0 | 543991.5667 | 636711.1369 | 735220.363 | 1229573.309 | 341809.3813 | 842860.23 | 842407.0398 | 437613.024 | 453435.122 | 401064.2299 | 442420.5097 | 612365.0391 |
| SL 12:2;O/18:2;O | 8070661.641 | 8813032.454 | 7365291.152 | 7620131.37 | 10219054.37 | 8670213.992 | 8887361.799 | 8370565.357 | 7968881.503 | 8422684.214 | 8165080.156 | 8782125.638 |

| Metabolites | PBS_1 | PBS_2 | PBS_3 | PBS_4 | BCG_1 | BCG_2 | BCG_3 | BCG_4 | **BCGΔ*pe18*_1** | **BCGΔ*pe18*_2** | **BCGΔ*pe18*_3** | **BCGΔ*pe18*_4** |
|---|---|---|---|---|---|---|---|---|---|---|---|---|
| N-(5-Chloro-2-hydroxyphenyl)piperidine-4-carboxamide | 2022899.573 | 1901015.272 | 1883022.45 | 2060694.191 | 1846769.649 | 1858684.815 | 1855646.935 | 1856892.433 | 1834867.371 | 1900790.13 | 1829378.439 | 1851888.677 |
| PS(22:4(7Z,10Z,13Z,16Z)/20:4(5Z,8Z,11Z,14Z)) | 53618702.45 | 58525640.83 | 29407129.52 | 45389382.19 | 10595249.81 | 10614587.64 | 13841983.89 | 15093466.68 | 18198223.96 | 17528510.3 | 12956982.92 | 11620272.97 |
| 2-Heptylquinolin-4(1H)-one | 546535.9301 | 1170660.667 | 1037431.383 | 812702.5964 | 742086.3042 | 1182489.935 | 919010.983 | 903025.6097 | 537570.8582 | 602627.9299 | 713916.7828 | 783490.809 |
| Coptisine cation | 1651674.579 | 1140169.184 | 1701812.601 | 1235701.442 | 3424069.634 | 2833283.553 | 6760644.574 | 6640256.51 | 2973567.251 | 2479800.431 | 4414804.721 | 1102551.042 |
| PC 16:3_26:7 | 14742506.57 | 16277856.26 | 7439449.304 | 10976330.35 | 9277952.269 | 8343146.184 | 10360385.37 | 9636151.456 | 11650908.98 | 12281218.87 | 7955023.19 | 8100421.572 |
| 2-Hydroxy-3-isopropyl-6-methylbenzoic acid | 1261870.145 | 1347494.046 | 1423626.106 | 1180628.353 | 379097.3162 | 1544788.714 | 449943.3311 | 1043275.252 | 1383380.327 | 1131691.785 | 941058.5217 | 1438103.541 |
| PG 18:0_18:1 | 4981338.62 | 2463327.593 | 9685935.87 | 5244921.056 | 4795077.422 | 125307.7458 | 2895851.991 | 962384.6112 | 8459170.272 | 4858343.582 | 5580427.187 | 869761.7103 |
| Cer(d18:0/16:0) | 5056557.012 | 5908317.31 | 3985322.624 | 3937735.488 | 4041168.681 | 2586101.355 | 3986262.108 | 4193048.939 | 6935412.678 | 7914691.391 | 2696284.13 | 2702216.69 |
| N-Methylglutamic acid | 224112.8297 | 216616.8301 | 271197.5213 | 219482.6833 | 920037.0218 | 645100.4851 | 858388.3674 | 699315.5283 | 1203892.765 | 1058429.639 | 1377301.055 | 1110594.251 |
| LPE 17:2 | 1192756.322 | 1553146.422 | 2075051.384 | 1464934.118 | 2380070.359 | 2099424.189 | 2737747.518 | 2291964.956 | 1994086.172 | 1228392.474 | 907968.9645 | 1924346.378 |
| Linoleyl-carnitine | 117247 | 100738 | 169836 | 113774 | 955287 | 946632 | 127267 | 121425 | 821766 | 876522 | 799394 | 584033 |

| Metabolites | PBS_1 | PBS_2 | PBS_3 | PBS_4 | BCG_1 | BCG_2 | BCG_3 | BCG_4 | BCGΔ*pe18*_1 | BCGΔ*pe18*_2 | BCGΔ*pe18*_3 | BCGΔ*pe18*_4 |
|---|---|---|---|---|---|---|---|---|---|---|---|---|
| | 63.65 | 39.44 | 73.33 | 34.74 | 1.354 | 2.296 | 56.63 | 59.18 | 7.212 | 9.741 | 0.016 | 6.373 |
| N-Methyl-L-asparagine | 5242310.898 | 4896585.405 | 9228495.134 | 7059804.184 | 26367063.72 | 20726371.32 | 23607861.22 | 21230388.86 | 23177556.37 | 12984323.27 | 15650037.81 | 19638355.95 |
| 1-24:1-2-18:3-phosphatidylserine | 60702053.97 | 72148866.94 | 42651918.55 | 52822988.09 | 75485010.63 | 61959060.24 | 78071689.21 | 68427520.66 | 83540008.15 | 93143757.94 | 72300264.98 | 55712444.13 |
| PC O-18:5_20:4 | 1002302.804 | 736056.2721 | 926184.3127 | 950848.9888 | 805409.8727 | 21152.40189 | 822521.2213 | 445814.0364 | 1028430.315 | 928522.8406 | 894673.5712 | 656977.2902 |
| 1,2-Diethylhydrazine | 696998.8022 | 611523.0505 | 948501.6854 | 680159.8464 | 2777939.658 | 1308714.479 | 2002791.891 | 1961811.867 | 1497334.859 | 3233346.159 | 2028364.813 | 1326804.677 |
| Methyl 1-pentyl-1H-indole-3-carboxylate | 1465276.477 | 416292.7681 | 1890627.226 | 1261003.043 | 9563009.087 | 9648579.419 | 50365887.55 | 18386157.68 | 15047610.23 | 8942450.635 | 9145501.023 | 9067760.259 |
| 4-(4-(1H-Indol-3-yl)-3,6-dihydro-1(2H)-pyridinyl)-1H-pyrazolo[3,4-d]pyrimidine | 3139757.279 | 3021054.269 | 2518299.335 | 1981848.181 | 4034067.88 | 3072084.923 | 2723268.371 | 2521131.096 | 3189741.143 | 3403242.973 | 2763018.764 | 3188889.124 |
| N2-Acetylornithine | 0 | 60418.03672 | 22761.64034 | 70766.93594 | 7070871.002 | 4804845.15 | 9290138.131 | 4356984.021 | 3767017.113 | 3397317.577 | 3545986.613 | 3443857.132 |
| Benzoic acid, 2-[[(2Z)-3-(3,4-dimethoxyphenyl)-1-oxo-2-propen-1-yl]amino]-3-hydroxy- | 4808316.398 | 4245588.945 | 4035625.073 | 4071142.798 | 4092875.273 | 3893339.487 | 4050237.01 | 3731147.424 | 4028137.478 | 4141150.038 | 4110148.708 | 3890257.824 |

| Metabolites | PBS_1 | PBS_2 | PBS_3 | PBS_4 | BCG_1 | BCG_2 | BCG_3 | BCG_4 | BCGΔ*pe18*_1 | BCGΔ*pe18*_2 | BCGΔ*pe18*_3 | BCGΔ*pe18*_4 |
|---|---|---|---|---|---|---|---|---|---|---|---|---|
| Eurostoside | 2558958.051 | 2410893.68 | 2224521.096 | 2170629.614 | 2244531.953 | 2309387.49 | 1941581.53 | 2272661.596 | 2040214.389 | 2242195.355 | 2267919.257 | 2205414.57 |
| 6-Hydroxyeurycomalactone | 1097415.797 | 1158427.472 | 1510033.387 | 1536190.189 | 1334560.565 | 1067654.019 | 1049608.381 | 1125869.374 | 1359010.546 | 1205477.309 | 1310925.977 | 1685918.819 |
| M511T534 | 2173085.394 | 1713460.006 | 1476543.444 | 2373311.879 | 655472.5837 | 3724819.359 | 2774325.437 | 888823.352 | 1449712.107 | 3093815.639 | 1718950.635 | 3545518.215 |
| PC(18:0/16:0) | 208830843.2 | 233506071.8 | 294860075 | 198650546 | 252504868.3 | 190656303.9 | 307315361.5 | 224138121.9 | 238099897.2 | 218968930.3 | 236489026.8 | 116895037 |
| PC 18:0 | 1260462.623 | 1181078.793 | 1390689.148 | 1311726.507 | 1179230.468 | 1164123.059 | 1407874.732 | 1243092.312 | 1329576.258 | 1128323.804 | 1337035.825 | 1286642.746 |
| 2-[(Phenylsulfonyl)amino]-5-propylbenzoic acid | 826629.0082 | 589739.1663 | 895698.7248 | 637046.1869 | 1679923.012 | 1536788.046 | 4617978.768 | 4523847.905 | 1607502.98 | 1202024.46 | 2322541.723 | 597706.7816 |
| Pergolide | 2597349.372 | 2893896.695 | 2592404.42 | 2520439.378 | 2745931.13 | 2668802.518 | 2911441.45 | 2696176.259 | 2763205.337 | 2750838.215 | 2844655.586 | 2612694.864 |
| SM 12:2;2O/2:0 | 175325.4237 | 188240.7196 | 162446.3902 | 169114.6451 | 212128.6784 | 174807.8407 | 190023.3212 | 153994.0722 | 185648.3973 | 167664.5658 | 218935.0039 | 157436.1013 |
| 1-O-Hexadecyl-2-O-methyl-sn-glyceryl-3-phosphorylcholine | 1008900.61 | 969346.8838 | 551949.9971 | 1108668.061 | 3651288.992 | 2923795.231 | 1516278.203 | 2851060.832 | 2102933.197 | 472520.6743 | 842901.3101 | 1815791.239 |
| M523T148 | 57068.36669 | 54898.51366 | 57306.7095 | 50541.08785 | 1120184.137 | 963668.1702 | 1063019.249 | 1469446.861 | 543537.1622 | 425141.4441 | 778215.7532 | 291294.3583 |
| PC(20:1) | 159444 | 163434 | 173713 | 179079 | 340816 | 215275 | 357234 | 233590 | 277100 | 310964 | 244560 | 216256 |

| Metabolites | PBS_1 | PBS_2 | PBS_3 | PBS_4 | BCG_1 | BCG_2 | BCG_3 | BCG_4 | BCGΔ*pe18*_1 | BCGΔ*pe18*_2 | BCGΔ*pe18*_3 | BCGΔ*pe18*_4 |
|---|---|---|---|---|---|---|---|---|---|---|---|---|
| | 5.863 | 6.979 | 3.818 | 3.065 | 6.611 | 3.352 | 4.012 | 2.108 | 7.967 | 9.486 | 6.636 | 0.738 |
| SMGDG O-9:0_10:0 | 10125.1651 | 0 | 0 | 0 | 103106.755 | 172664.7123 | 101289.4528 | 77256.03635 | 115826.9649 | 101946.7241 | 149315.0283 | 61438.71528 |
| LNAPE 2:0/N-16:1 | 319807.0202 | 356007.5513 | 285593.2989 | 320877.6752 | 337080.1395 | 302252.1636 | 313250.5949 | 316164.5836 | 323261.2612 | 330751.5073 | 308213.7252 | 318500.6344 |
| .beta.-Homoproline | 3052293.125 | 4095538.973 | 4918991.247 | 5197386.732 | 10909366.4 | 9037076.087 | 14475422.19 | 8748469.232 | 7742610.471 | 7876013.939 | 8934953.222 | 10987666.39 |
| Platyphylloside | 0 | 0 | 0 | 14138.26944 | 2683237.795 | 2656432.276 | 1926662.861 | 1846591.711 | 2338264.987 | 1966792.447 | 2331666.79 | 2116751.714 |
| 3-methylhex-2-enoate | 632865.9142 | 623324.2696 | 588911.135 | 631641.1534 | 677578.4722 | 1010806.659 | 620113.4665 | 595196.4054 | 648623.2127 | 582240.3718 | 972054.7683 | 661214.5801 |
| 1-Pentanone, 1-(3,4-dihydroxyphenyl)-2-(1-pyrrolidinyl)- | 5204531.713 | 6885352.079 | 6067891.854 | 6437110.781 | 3630839.998 | 3389940.857 | 5355447.176 | 4123540.64 | 3462682.629 | 4100130.839 | 4192872.439 | 4748575.37 |
| M300T369 | 773499.3504 | 760738.1148 | 702519.1854 | 787473.8286 | 740474.7753 | 669999.5035 | 724631.3321 | 646580.4576 | 655171.3301 | 704587.0975 | 690995.3891 | 889521.2353 |
| PG O-17:2_22:5 | 454284.298 | 266313.0012 | 369071.2529 | 362815.6114 | 1150284.744 | 1631461.22 | 1356943.086 | 933288.0737 | 1029746.449 | 1466853.237 | 1189937.725 | 1279749.281 |
| PI O-24:1_16:4 | 178534.5109 | 204596.8477 | 222261.2456 | 171752.3886 | 274465.0694 | 0 | 144483.2046 | 226314.0567 | 207908.9984 | 190203.7573 | 141674.737 | 362050.6042 |
| L-.beta.-Homolysine | 69211714.96 | 48788412 | 87958161.28 | 58934533.23 | 96758729.75 | 66776655.3 | 114519667.2 | 91584561.22 | 63805366.57 | 78581295.71 | 85589075.86 | 51824442.61 |
| ADGGA | 181609. | 109576 | 112014. | 453548. | 277878. | 156569. | 309821. | 254520. | 48420.0 | 632654. | 143507. | 358123. |

| Metabolites | PBS_1 | PBS_2 | PBS_3 | PBS_4 | BCG_1 | BCG_2 | BCG_3 | BCG_4 | BCGΔ*pe18*_1 | BCGΔ*pe18*_2 | BCGΔ*pe18*_3 | BCGΔ*pe18*_4 |
|---|---|---|---|---|---|---|---|---|---|---|---|---|
| 22:6_20:3_22:6 | 3474 | 5.377 | 3058 | 8249 | 8058 | 9698 | 3906 | 8707 | 9238 | 9488 | 8527 | 3235 |
| 2,4-Diphenyl-3-azabicyclo[3.3.1]nonan-9-ol | 2884611.109 | 2936756.139 | 2618721.591 | 2526143.136 | 2709047.788 | 2779890.234 | 2762508.756 | 2785649.246 | 2740976.718 | 2656178.147 | 2534451.462 | 2411278.453 |
| Propionic acid | 1307507.238 | 1323524.126 | 1232800.631 | 874325.4249 | 1977943.239 | 2053683.22 | 2257209.836 | 2195832.352 | 2095039.167 | 4346146.572 | 2086168.902 | 1675096.449 |
| 6-(3-Chloro-4-methylanilino)-2,4(1H,3H)-pyrimidinedione | 361196.5379 | 265261.1061 | 423791.0985 | 246697.8677 | 4935129.141 | 3139526.329 | 7946442.982 | 6167493.113 | 6244030.826 | 6415829.715 | 6517936.212 | 5234306.049 |
| (5alpha,17beta)-3-Oxoandrostan-17-yl undecanoate | 1253482.827 | 1395773.648 | 1289410.897 | 1365337.741 | 1431874.046 | 1532953.086 | 1336716.275 | 1304372.645 | 1379388.48 | 1409556.094 | 1369841.356 | 1425681.81 |
| 2,3,4,5-tetranor-thromboxane B1 | 1131841.495 | 1177198.417 | 1213849.261 | 1127926.14 | 1100109.681 | 1034801.821 | 1408695.369 | 1050585.284 | 1055116.379 | 1132178.787 | 1091588.797 | 1762485.28 |
| N4-Acetylaminobutanal | 614533.9243 | 553610.9485 | 611281.0378 | 632477.0334 | 6970164.015 | 2607761.723 | 8361334.574 | 2574261 | 2778356.752 | 1970377.8 | 2547962.34 | 2284080.496 |
| PMeOH 26:2_20:4 | 1458114.144 | 1313231.106 | 1446357.856 | 1358674.541 | 816355.0961 | 821545.7641 | 800818.5277 | 339427.8535 | 1179926.979 | 1023502.417 | 820124.6693 | 181238.7276 |
| alpha-Methylene-gamma-butyrolactone | 1751547.942 | 1702553.051 | 1705056.854 | 1606184.098 | 1754409.169 | 1746286.523 | 1913839.677 | 1904078.744 | 1767603.761 | 1745021.302 | 1776258.72 | 1649386.572 |
| 5Z,8Z,11Z-Eicosatrienoic acid | 6793137.142 | 5592952.771 | 11145103.15 | 7259646.606 | 6826189.394 | 5206025.869 | 2955615.526 | 4617555.158 | 4220957.502 | 3390521.639 | 4439786.871 | 4543765.471 |
| 2-Amino-4,6-dinitrotoluene | 841728.5187 | 506898.1242 | 1143409.097 | 724783.3086 | 4165269.846 | 1739209.399 | 3601567.654 | 2467492.724 | 2351782.807 | 2203295.361 | 2424099.44 | 1763268.863 |

| **Metabolites** | **PBS_1** | **PBS_2** | **PBS_3** | **PBS_4** | **BCG_1** | **BCG_2** | **BCG_3** | **BCG_4** | **BCGΔ*pe18*_1** | **BCGΔ*pe18*_2** | **BCGΔ*pe18*_3** | **BCGΔ*pe18*_4** |
|---|---|---|---|---|---|---|---|---|---|---|---|---|
| PC(32:2) | 8916647.657 | 3798246.064 | 2257260.361 | 2096147.047 | 43351821.82 | 23701278.58 | 77250699.7 | 71379339.34 | 37412661.71 | 30226228.92 | 25055855.65 | 9424983.836 |
| Leukotriene_F4 | 8970505.203 | 8828019.5 | 7705616.401 | 9299340.556 | 9301189.583 | 10587945.62 | 9305868.888 | 9998862.536 | 9865466.483 | 8312795.323 | 8802879.257 | 8117734.468 |
| 4-Methoxy-2,4-dioxobutanoic acid | 250095.1599 | 199138.2836 | 293766.8539 | 223328.7947 | 2297163.066 | 1560805.878 | 2323083.133 | 1880180.875 | 1539143.351 | 2262505.101 | 2101282.146 | 1140079.652 |
| PE O-26:5_11:0 | 4144082.584 | 2645493.526 | 4319211.886 | 3516041.653 | 4041475.798 | 45655.44573 | 4591384.309 | 5821069.707 | 7891206.709 | 4425271.22 | 3928835.238 | 6804635.123 |
| (6R,12aS,12bS,13R)-8,9-Dimethoxy-5,6-dihydro-3H,4H-6,12b,12a-(propane[1,1,3]triyl)-3a,10b-diazabenzo[7,8]cycloocta[1,2,3-cd]inden-11(12H)-one | 4209216.181 | 894681.4829 | 5381025.757 | 1338518.512 | 6745129.321 | 8796863.59 | 30645702.84 | 25665338.51 | 10967955.4 | 6620047.653 | 11847398.62 | 3591125.424 |
| Tropine | 3416433.777 | 6568794.063 | 10620613.65 | 7673122.916 | 11074225.59 | 10954553.53 | 10371129.05 | 5445906.98 | 5335795.478 | 10253309.36 | 4441592.837 | 5163134.25 |
| DGDG O-9:0_16:3 | 21145.42551 | 15319.01156 | 33105.82306 | 34865.32365 | 193255.0894 | 163867.7557 | 194519.5036 | 146533.6123 | 123617.5607 | 115253.5269 | 185378.7862 | 93225.7725 |
| PC O-19:3 | 1143282.006 | 203699.8351 | 1083397.822 | 883367.8991 | 2513396.751 | 3217978.173 | 2520348.809 | 1053360.66 | 1584452.251 | 1643584.992 | 1481279.539 | 1394970.915 |
| cis-Pinonic acid | 483949.921 | 514965.38 | 493965.954 | 535823.6168 | 494110.2335 | 530506.9874 | 534448.9451 | 537307.2755 | 519093.5417 | 578212.24 | 517863.1527 | 606288.4738 |

| Metabolites | PBS_1 | PBS_2 | PBS_3 | PBS_4 | BCG_1 | BCG_2 | BCG_3 | BCG_4 | BCGΔ*pe18*_1 | BCGΔ*pe18*_2 | BCGΔ*pe18*_3 | BCGΔ*pe18*_4 |
|---|---|---|---|---|---|---|---|---|---|---|---|---|
| N-(4-Piperidinyl)acetamide | 4889182.943 | 1133670.959 | 1655356.062 | 1770021.609 | 1924180.322 | 1547195.071 | 1720233.345 | 2038863.137 | 1820904.68 | 1037837.671 | 1941539.12 | 958001.7674 |
| (1-Acetyloxy-3-hydroxy-6,8a-dimethyl-7-oxo-3-propan-2-yl-2,3a,4,8-tetrahydro-1H-azulen-4-yl) 4-hydroxybenzoate | 2653990.259 | 4507679.288 | 1042352.983 | 2409449.653 | 1753026.349 | 4869107.527 | 1929103.648 | 1491738.375 | 1618229.714 | 1344037.616 | 1895654.719 | 1214151.428 |
| FAHFA 9:0/25:0 | 1205360.577 | 1186053.916 | 1217646.01 | 1274492.682 | 6501591.399 | 11441784.78 | 7417804.071 | 6557787.193 | 5173867.083 | 3980661.145 | 6726727.143 | 5551541.672 |
| Piperundecalidine | 2304006.241 | 1574513.982 | 1531095.715 | 1592919.126 | 2237989.721 | 1630341.421 | 1879929.966 | 2196015.517 | 1729850.362 | 1651001.247 | 1606017.246 | 1580636.935 |
| 13,14-Dihydro-15-ketoprostaglandin J2 | 1982959.595 | 1850268.385 | 1837842.785 | 1781386.048 | 1810107.866 | 1782140.708 | 1769986.868 | 1727594.777 | 1777991.078 | 2013686.927 | 1767138.739 | 1802430.556 |
| Nabumetone | 8419606.486 | 3857310.674 | 3210340.171 | 4917092.628 | 7377445.242 | 7541564.592 | 5707420.865 | 7622363.263 | 6219777.603 | 8278905.696 | 7300871.035 | 3229752.123 |
| Val-Ile | 469623.0197 | 493328.4911 | 469500.7772 | 349399.356 | 415618.4998 | 251868.8787 | 349398.5556 | 312471.822 | 424218.2854 | 533094.5722 | 499549.307 | 233442.1776 |
| LNAPS 26:4/N-10:0 | 1876742.58 | 697761.9254 | 841900.9732 | 1000590.72 | 1487192.323 | 48369.82153 | 818304.3643 | 484449.3945 | 1308700.409 | 1591769.857 | 798926.495 | 309322.9818 |
| PE O-9:0_26:5 | 1636395.717 | 1668767.746 | 1342348.946 | 1020424.158 | 1185643.03 | 1056631.163 | 1222294.09 | 1159220.144 | 1027427.694 | 1023964.102 | 1206801.597 | 864032.1464 |
| PGF3alpha | 26310769.15 | 23054395.61 | 25073556.75 | 27151055.22 | 24277462.35 | 28413355.51 | 19915892.12 | 22667653.64 | 21109395.75 | 19957157.86 | 24773381.08 | 27532940.47 |

| Metabolites | PBS_1 | PBS_2 | PBS_3 | PBS_4 | BCG_1 | BCG_2 | BCG_3 | BCG_4 | BCGΔ*pe18*_1 | BCGΔ*pe18*_2 | BCGΔ*pe18*_3 | BCGΔ*pe18*_4 |
|---|---|---|---|---|---|---|---|---|---|---|---|---|
| Icariside II | 303664.2933 | 318325.0298 | 401265.4372 | 281333.996 | 329917.6686 | 444402.3088 | 542392.6124 | 349327.1598 | 463790.7434 | 0 | 136349.2556 | 494638.0542 |
| 6,7,8,9,10,11-Hexahydro-5H-cycloocta[b]indole | 1682497.75 | 1801790.008 | 1727101.006 | 1517108.138 | 1684313.724 | 1624403.113 | 1472366.859 | 2048358.289 | 1831168.981 | 1604429.582 | 1922675.451 | 1621211.02 |
| 2-Naphthalenecarboxamide, N-[(1S)-2-[4-(5-chloro-2,3-dihydro-2-oxo-1H-benzimidazol-1-yl)-1-piperidinyl]-1-methylethyl]- | 0 | 0 | 0 | 0 | 625765.612 | 499839.1959 | 391706.9511 | 370134.9562 | 544880.4991 | 520255.4312 | 629474.5184 | 555518.3395 |
| 8-Geranyl-7-hydroxycoumarin | 10167783.61 | 1188879.942 | 2563990.623 | 2638916.846 | 1587099.534 | 2862606.779 | 1797540.228 | 1699193.572 | 1677504.522 | 913275.7204 | 2451781.925 | 1362578.571 |
| LPG 18:0 | 6574130.797 | 7479633.436 | 3807052.235 | 6556493.8 | 27419401.23 | 33104130.4 | 41765717.7 | 27932734.86 | 19689082.75 | 17047822.65 | 35212508.74 | 21465673.1 |
| (6E,8E)-3-Hydroxy-10-methoxy-4,9-dimethyl-10-oxodeca-6,8-dienoic acid | 303684.9631 | 263469.907 | 315365.81 | 319813.912 | 272062.775 | 370699.9675 | 728123.8315 | 351662.9633 | 406040.4442 | 1261084.654 | 320405.9956 | 816246.9453 |
| Cyromazine | 388000.9717 | 405398.3175 | 484787.3723 | 487889.7606 | 437484.9936 | 556934.7528 | 456641.9485 | 577270.7157 | 471697.2604 | 493525.9053 | 1249587.024 | 458051.1144 |
| Coumarin 334 | 5184295.646 | 4822344.583 | 4682762.855 | 4722551.865 | 3786025.759 | 4231059.88 | 3730995.769 | 4287700.157 | 4251859.345 | 3349197.068 | 3981883.148 | 3871686.614 |

| Metabolites | PBS_1 | PBS_2 | PBS_3 | PBS_4 | BCG_1 | BCG_2 | BCG_3 | BCG_4 | BCGΔ*pe18*_1 | BCGΔ*pe18*_2 | BCGΔ*pe18*_3 | BCGΔ*pe18*_4 |
|---|---|---|---|---|---|---|---|---|---|---|---|---|
| PG 20:3_22:4 | 25344702.61 | 33361462.68 | 3897619.983 | 17963449.88 | 5702660.581 | 2691732.843 | 8300992.55 | 5456126.181 | 12757197.43 | 11718166.39 | 5056855.17 | 3480658.861 |
| N-Arachidonylmaleimide | 993282.2279 | 171564.8772 | 631905.8431 | 205802.0206 | 1339958.67 | 1706395.607 | 3195338.943 | 2652536.679 | 1942436.464 | 1988772.477 | 2459608.298 | 397892.676 |
| 1-O-beta-D-Glucopyranosyl-2,3-di-O-(8-hexadecenoyl)glycerol | 13168957.8 | 14422223.47 | 14847417.11 | 13404554.05 | 14645569.98 | 9984529.447 | 17106034.61 | 11176218.84 | 13187349.77 | 17574352.87 | 13016021.6 | 7347578.009 |
| SL 15:2;O/26:7 | 241909.4943 | 258430.5319 | 263242.1271 | 257765.848 | 279622.8955 | 280074.8873 | 318973.4675 | 286142.6456 | 247966.9776 | 278123.6666 | 293390.2222 | 246296.6268 |
| 1-24:0-2-18:1-phosphatidylserine | 742156.8017 | 1123594.836 | 1271294.622 | 1010241.345 | 3469742.814 | 1946943.21 | 2711481.239 | 1989046.083 | 2466769.048 | 2692843.014 | 2706972.41 | 1617536.848 |
| Levonorgestrel acetate | 3560295.417 | 9573822.747 | 6133383.413 | 4811556.62 | 888719.2725 | 912074.7771 | 692885.7488 | 900531.433 | 879116.3298 | 790746.1159 | 608670.9985 | 4030504.44 |
| TG 8:0_16:4_22:6 | 830247.4693 | 281253.528 | 777140.3042 | 1002996.496 | 6704827.959 | 4781670.072 | 5171690.448 | 3671166.874 | 3272223.199 | 4696636.789 | 4549187.594 | 2376661.393 |
| Norfenfluramine | 678506.08 | 718424.3353 | 644553.8138 | 709579.204 | 664649.5803 | 680848.0465 | 685225.4163 | 672472.0808 | 693192.1774 | 681819.1863 | 620286.034 | 1017324.581 |
| DGDG O-16:4_2:0 | 469351.6406 | 513045.2891 | 389299.9198 | 453210.9464 | 454485.7945 | 453804.7245 | 442015.1762 | 492571.5253 | 442168.7226 | 408823.3872 | 424341.1002 | 353293.2453 |
| 14-Hydroxymyristic acid | 1265814.591 | 1238723.009 | 1596654.418 | 921009.5152 | 1249221.908 | 1204353.027 | 1103119.866 | 909588.1904 | 1377164.504 | 1210046.501 | 1525673.069 | 1199607.563 |

| Metabolites | PBS_1 | PBS_2 | PBS_3 | PBS_4 | BCG_1 | BCG_2 | BCG_3 | BCG_4 | BCGΔ*pe18*_1 | BCGΔ*pe18*_2 | BCGΔ*pe18*_3 | BCGΔ*pe18*_4 |
|---|---|---|---|---|---|---|---|---|---|---|---|---|
| 5-(1-Naphthylmethylene)-2,4,6(1H,3H,5H)-pyrimidinetrione | 721583.5837 | 472268.6158 | 1005899.572 | 694946.7827 | 668012.1982 | 826721.6847 | 852749.7642 | 1058408.001 | 750194.7823 | 329001.8538 | 409295.3866 | 661883.9444 |
| MGDG O-9:0_28:7 | 938426.1987 | 1339242.153 | 656096.223 | 1212510.527 | 2855677.557 | 2510437.302 | 1103107.194 | 1807120.701 | 1707075.547 | 455827.641 | 324679.0466 | 2732987.194 |
| PG 22:4_22:5 | 55463794.27 | 28811523.78 | 4431985.012 | 22840281.57 | 1480033.201 | 719267.5106 | 2620854.784 | 1294923.831 | 2092335.851 | 3465882 | 1482330.333 | 829174.961 |
| D-myo-Inositol-1,4-diphosphate | 57109.58959 | 26019.34751 | 67726.3489 | 65706.4463 | 2829451.385 | 2062674.721 | 3191511.276 | 2461193.646 | 1988834.87 | 2468831.075 | 3192185.842 | 1594169.427 |
| SM 42:8;3O | 5124636.11 | 6016174.979 | 1701391.783 | 3952408.572 | 1186618.979 | 1545063.226 | 1156313.008 | 1435167.389 | 1997266.41 | 1286312.868 | 1176574.037 | 1474003.717 |
| PC O-12:0_2:0 | 255651.601 | 253353.7448 | 2306594.706 | 1468309.57 | 5167922.883 | 4245912.847 | 1086979.432 | 3822576.309 | 4210441.462 | 4532465.065 | 5301382.326 | 678056.3152 |
| Cervonyl carnitine | 926509.4282 | 910936.8258 | 769508.0155 | 865721.0428 | 997108.1589 | 1089823.377 | 1616193.028 | 1093384.266 | 1057418.931 | 1110232.413 | 997416.6005 | 901292.4852 |
| 2-[(6-Ethoxy-1,3-benzothiazol-2-yl)thio]-1-phenylethanone | 79310.20071 | 91445.13637 | 130426.8996 | 192921.2176 | 1628300.252 | 1481473.721 | 1671139.361 | 1744555.689 | 742315.8206 | 836156.5656 | 784241.2082 | 978081.3182 |
| (6E)-8-Methyl-6-nonenoic acid | 654700.8404 | 851472.8814 | 1726630.548 | 600230.7724 | 648384.1461 | 613077.3611 | 737753.1282 | 537864.3897 | 637919.5785 | 644662.0451 | 1643282.36 | 611258.2478 |
| Desogestrel | 2923850.799 | 2858382.902 | 1736748.191 | 2254881.401 | 2524988.346 | 2215523.645 | 2752677.767 | 2468052.57 | 3232109.643 | 2653769.773 | 2389615.33 | 2237504.107 |
| LPI 11:0 | 926644. | 104466 | 123559 | 120588 | 156958 | 158424 | 124627 | 125716 | 133517 | 378212. | 465460. | 139643 |

| Metabolites | PBS_1 | PBS_2 | PBS_3 | PBS_4 | BCG_1 | BCG_2 | BCG_3 | BCG_4 | BCGΔ*pe18*_1 | BCGΔ*pe18*_2 | BCGΔ*pe18*_3 | BCGΔ*pe18*_4 |
|---|---|---|---|---|---|---|---|---|---|---|---|---|
| | 2069 | 6.871 | 2.547 | 3.936 | 5.82 | 5.591 | 4.293 | 8.206 | 5.188 | 9421 | 8679 | 3.851 |
| Methyl angolensate | 335012.521 | 367952.3818 | 248625.7124 | 264046.4167 | 382603.5664 | 192858.9876 | 343580.686 | 197671.3821 | 168550.387 | 251239.7646 | 339919.9415 | 318556.3753 |
| 3-[5-(2-Methylpropyl)-3,6-dioxopiperazin-2-yl]propanoic acid | 287415.9663 | 294162.3735 | 379477.051 | 364756.6582 | 1150081.26 | 511923.2752 | 672640.0988 | 555484.8597 | 651052.862 | 627074.4334 | 664896.7217 | 743593.5065 |
| Cer 10:1;2O/10:0 | 3327499.451 | 4184255.393 | 2421565.004 | 3026034.014 | 3532642.529 | 2944201.861 | 3580480.015 | 4033450.917 | 4427236.743 | 4763646.104 | 4208950.803 | 3268584.856 |
| 1-Docosahexaenoyl-2-stearoyl-sn-glycero-3-phosphocholine | 19485311.71 | 17812092.42 | 20713745.38 | 19457343.88 | 22641896.11 | 19022148.7 | 20478045.27 | 30651176.91 | 21841342.71 | 21747634.09 | 17769229.85 | 14397489.06 |
| PC(17:0/17:0) | 208830843.2 | 233506071.8 | 294860075 | 198650546 | 252504868.3 | 190656303.9 | 307315361.5 | 224138121.9 | 238099897.2 | 218968930.3 | 236489026.8 | 116895037 |
| SM 21:1;2O/19:1 | 696127.9556 | 2013985.556 | 2517306.134 | 2188398.364 | 825880.9553 | 4108545.306 | 1553224.761 | 3834657.284 | 1644227.763 | 4031053.409 | 0 | 1821194.679 |
| 4-Chloro-N-(4-methyl-1,3-thiazol-2-yl)benzenesulfonamide | 8510590.97 | 6383850.475 | 9223244.885 | 7301824.902 | 12580685.08 | 10138552.13 | 13713428.66 | 11775434.76 | 9597579.754 | 11389051.56 | 11895892.7 | 7889924.168 |
| Flavone base + 2O, 2MeO, C-Hex | 164450.1914 | 88093.01588 | 130007.9026 | 117957.5747 | 50869.47399 | 99739.32273 | 278295.2222 | 294793.2291 | 130900.0958 | 49659.27533 | 150794.8423 | 29400.52836 |
| PA 13:1_13:1 | 774927 | 879741 | 638795 | 770252 | 915418 | 134194 | 983090 | 973193 | 753351 | 104832 | 897726 | 507182 |

| Metabolites | PBS_1 | PBS_2 | PBS_3 | PBS_4 | BCG_1 | BCG_2 | BCG_3 | BCG_4 | BCGΔ*pe18*_1 | BCGΔ*pe18*_2 | BCGΔ*pe18*_3 | BCGΔ*pe18*_4 |
|---|---|---|---|---|---|---|---|---|---|---|---|---|
| | 2.615 | 8.889 | 2.33 | 5.489 | 4.197 | 82.22 | 6.082 | 2.152 | 5.828 | 78.85 | 6.315 | 4.594 |
| 1-Hydroxy Vitamin D2 | 17797.33653 | 138284.3628 | 320871.6139 | 195512.8144 | 982724.2546 | 716220.2042 | 1049544.732 | 940300.1745 | 597276.18 | 669086.3926 | 719676.352 | 374488.0555 |
| Tridecenoylcarnitine (Car(13:1)) | 24717287.32 | 23113776.23 | 19481970.05 | 20440329.79 | 23689910.66 | 18985545.91 | 21706142.36 | 21996942.34 | 22179205.11 | 22436511.09 | 21609372.6 | 18456519.55 |
| Anastrozole | 341339.245 | 1070685.497 | 1147149.54 | 1707323.65 | 594367.4403 | 512097.8595 | 641605.2131 | 220616.5823 | 578996.5433 | 516172.1273 | 540405.469 | 2213079.688 |
| ST 24:1;O4;T/17:1 | 927244.9981 | 626932.5311 | 875656.2133 | 659016.0725 | 1440022.214 | 798808.7978 | 1562740.1 | 1266458.584 | 1401836.642 | 2582968.13 | 1461530.009 | 848166.6396 |
| Monoisobutyl phthalate | 1886431.818 | 1889622.622 | 2666897.878 | 2281102.02 | 2553172.53 | 2343080.943 | 2669942.691 | 1969640.63 | 1861059.625 | 1850136.02 | 2082164.874 | 2434838.355 |
| Bergapten | 3325075.949 | 4047462.52 | 3833276.277 | 4542080.42 | 4997453.18 | 4162842.194 | 4551018.783 | 3901446.994 | 3928616.106 | 4389896.676 | 3583474.739 | 8105592.9 |
| Simazine | 419749.2798 | 300521.699 | 446119.5578 | 390935.8303 | 601174.221 | 593669.6656 | 187262.0564 | 505955.2695 | 618876.6047 | 257290.5993 | 133475.9498 | 944953.655 |
| 1,2-Di-O-(8-hexadecenoyl)-3-O-(6-sulfoquinovopyranosyl) glycerol | 2637761.553 | 3365390.326 | 3993686.903 | 3881498.111 | 1991252.764 | 1430358.873 | 2866724.888 | 2057920.761 | 1592294.192 | 2527497.744 | 2436432.012 | 2159576.097 |
| Cystine-glutathione | 668286.1442 | 520331.1806 | 903435.4628 | 729428.9134 | 3103169.951 | 4213276.141 | 1488115.203 | 787380.1844 | 2049929.491 | 6917922.059 | 3535897.915 | 3449753.939 |
| 4-Hydroxybenzoyl-adenylate | 113151.1891 | 136265.774 | 125557.5032 | 93927.31347 | 391599.3299 | 324313.2527 | 428531.0968 | 355929.1108 | 369924.1379 | 433743.7913 | 375229.6359 | 464060.2415 |

| Metabolites | PBS_1 | PBS_2 | PBS_3 | PBS_4 | BCG_1 | BCG_2 | BCG_3 | BCG_4 | BCGΔ*pe18*_1 | BCGΔ*pe18*_2 | BCGΔ*pe18*_3 | BCGΔ*pe18*_4 |
|---|---|---|---|---|---|---|---|---|---|---|---|---|
| Cytidine 2',3'-cyclic phosphate | 329618.9793 | 418015.896 | 537763.4065 | 463665.6386 | 384264.0047 | 546299.0875 | 298448.0919 | 344609.6401 | 308706.9073 | 261454.2374 | 440077.1714 | 718064.4743 |
| LPI 14:0 | 308888.8498 | 314073.7878 | 291828.1088 | 299769.8601 | 405077.9542 | 712090.4281 | 785458.7757 | 639808.7452 | 739780.0226 | 728706.4868 | 455630.41 | 718494.2957 |
| PI 19:1_20:4 | 261833.8523 | 369998.8134 | 294664.2963 | 338215.2863 | 1064327.183 | 983558.6563 | 1317860.893 | 1733694.076 | 822906.6425 | 1366268.866 | 1041679.208 | 511386.9067 |
| PC(P-19:0) | 350089.4705 | 220351.9596 | 321280.076 | 250718.4124 | 722719.1729 | 609436.9533 | 633863.1857 | 669167.9011 | 276825.6629 | 589991.0411 | 672200.4239 | 473772.4163 |
| N-[1-(Phenylmethyl)-4(1H)-quinolinylidene]-1-pentanamine | 1282191.033 | 1584746.857 | 1332643.115 | 1097266.522 | 1192559.571 | 1460382.732 | 1373713.249 | 1359493.023 | 1207139.63 | 1385086.99 | 1352044.862 | 1090862.698 |
| 4-Allylcatechol | 469357.3227 | 411192.0641 | 540137.8528 | 464280.5025 | 565449.2124 | 521610.515 | 447724.3909 | 518253.8203 | 518079.5871 | 312234.0585 | 393581.8531 | 486326.1621 |
| Hex2Cer 12:0;2O | 311133.1348 | 362586.1731 | 278117.305 | 338792.2814 | 283421.3495 | 328674.8311 | 308596.4675 | 321638.0624 | 359661.9555 | 328937.0132 | 286673.2911 | 259089.2705 |
| 4,7,8-Trimethoxy-2-oxo-1,2-dihydro-3-quinolinecarbaldehyde | 519024.6304 | 499566.8665 | 733062.357 | 620689.4378 | 933757.2286 | 852349.5492 | 955496.9133 | 916330.3369 | 929798.1565 | 639595.7931 | 734758.2292 | 813350.0607 |
| DG 34:7 | 1660889.397 | 1755401.203 | 1271222.824 | 1761630.791 | 1468544.828 | 1940694.347 | 1566244.674 | 1340065.071 | 1518161.035 | 1397562.596 | 1409404.726 | 1340648.045 |
| N2,N4-Bis(2-methoxyethyl)-2,4-pyridi | 13802121.16 | 12946638.9 | 13909878.54 | 13008805.58 | 15758391.53 | 14108948.63 | 15269156.83 | 14854161.6 | 15877815.26 | 15560471.39 | 13597364.83 | 13803026.89 |

| **Metabolites** | **PBS_1** | **PBS_2** | **PBS_3** | **PBS_4** | **BCG_1** | **BCG_2** | **BCG_3** | **BCG_4** | **BCGΔ*pe18*_1** | **BCGΔ*pe18*_2** | **BCGΔ*pe18*_3** | **BCGΔ*pe18*_4** |
|---|---|---|---|---|---|---|---|---|---|---|---|---|
| nedicarboxamide | | | | | | | | | | | | |
| 4-(4-Cyano-2-fluorophenyl)-2-morpholino-5-(1H-1,2,4-triazol-5-yl)thiophene-3-carbonitrile | 357305.7542 | 311577.9873 | 482321.1466 | 272482.7789 | 2193743.868 | 1980651.389 | 1831903.783 | 1759353.723 | 1251237.04 | 1284392.766 | 1211008.248 | 1204572.682 |
| 5-(2-Fluorophenyl)-2-furaldehyde | 308855.6737 | 194322.8111 | 438715.2833 | 255379.3033 | 614001.1427 | 785093.0583 | 1863242.045 | 1947514.38 | 909223.3128 | 496955.4732 | 1061859.072 | 563043.7923 |
| PEtOH 16:0_24:4 | 526164.3892 | 533628.924 | 606790.4884 | 755699.7789 | 386358.7628 | 13024.56747 | 435907.1391 | 623317.5345 | 580241.6752 | 380937.7435 | 549422.9114 | 406075.3512 |
| 1-(Cyclohexylcarbonyl)-4-piperidinylamine | 2141059.107 | 3036544.776 | 1986293.241 | 2337669.445 | 2812253.41 | 2078628.883 | 2360139.979 | 2327918.956 | 2512646.286 | 2385344.976 | 2692458.024 | 2594036.568 |
| 5-Amino-2-methoxyphenol | 576387.6406 | 634810.1376 | 401666.9232 | 555347.2266 | 321093.9308 | 589421.4715 | 390728.7631 | 335098.2756 | 485622.9432 | 543390.6782 | 426148.6577 | 265428.6452 |
| 9-HODE | 708654.367 | 1076319.925 | 1405679.105 | 516541.9476 | 1468032.28 | 1311935.721 | 1189903.61 | 1268430.249 | 909264.7496 | 886857.0172 | 1225043.673 | 1144491.463 |
| Homostachydrine | 3395996.367 | 4010796.419 | 5498179.432 | 10330762.42 | 6814677.55 | 6215301.081 | 10067489.67 | 4571429.895 | 5134360.642 | 3617287.345 | 5069368.229 | 6152751.595 |
| PS 24:2 | 143788.1676 | 141972.2499 | 118494.4651 | 138962.0022 | 121112.4112 | 114265.1223 | 135913.9933 | 79363.53777 | 100413.6609 | 66153.90149 | 114404.631 | 130077.1216 |
| His-Trp | 7055676.139 | 6956979.714 | 6311267.963 | 6904406.304 | 7157335.024 | 6721943.074 | 6820815.852 | 7364185.136 | 7240389.448 | 7202998.755 | 6390685.067 | 6088317.747 |

| **Metabolites** | **PBS_1** | **PBS_2** | **PBS_3** | **PBS_4** | **BCG_1** | **BCG_2** | **BCG_3** | **BCG_4** | **BCGΔ*pe18*_1** | **BCGΔ*pe18*_2** | **BCGΔ*pe18*_3** | **BCGΔ*pe18*_4** |
|---|---|---|---|---|---|---|---|---|---|---|---|---|
| PC 9:0_9:0 | 66073.23945 | 105844.7102 | 292558.8688 | 0 | 8610793.33 | 3574864.248 | 6293298.507 | 4081573.925 | 4378573.572 | 4495321.759 | 4938810.135 | 3055747.326 |
| 2-Ketobutyric acid | 12893256.64 | 21557577.24 | 13960355.3 | 13182762 | 9653804.634 | 19094182.65 | 19713529.08 | 14078375.07 | 16437283.06 | 4953738.224 | 4197795.114 | 10493261.05 |
| SHexCer 30:3;2O | 864955.8984 | 3970481.846 | 683667.8554 | 2643923.623 | 2749129.124 | 1111766.146 | 4567923.885 | 3614852.301 | 936848.3109 | 3624847.282 | 920800.5439 | 2508274.858 |
| Xylulose | 4808909.485 | 3669935.588 | 7374570.04 | 4524665.782 | 7206998.893 | 4367759.759 | 2491310.777 | 3153536.392 | 3116316.12 | 3104219.989 | 3793164.995 | 5366180.928 |
| (4E,8E,12E,16E)-2,3,7,11,15-Pentahydroxy-4,6,8,10,12,14,16-heptamethyloctadeca-4,8,12,16-tetraenoic acid | 38563.2591 | 73710.17342 | 81523.83186 | 105754.2908 | 77556.71645 | 94223.37296 | 129528.153 | 67454.96038 | 58838.2344 | 104532.6028 | 33620.12384 | 1138472.621 |
| Myriocin | 111647.6571 | 89111.07547 | 163658.8188 | 129722.153 | 518495.876 | 335562.5245 | 533337.4203 | 416973.2491 | 302557.6898 | 504227.6492 | 551555.4599 | 224888.1793 |
| a-Santal-10-en-12-oic acid | 1614348.088 | 1402656.823 | 997182.1043 | 1838421.712 | 1491596.116 | 1685532.913 | 1387589.257 | 1514019.199 | 1528951.812 | 1363351.508 | 1604025.906 | 1410777.357 |
| 3'-O-methylcytidine | 350981.1227 | 353055.7777 | 719480.0649 | 563822.4959 | 1887707.508 | 1188184.591 | 1795630.215 | 1324007.797 | 1263115.321 | 1179729.943 | 1252865.079 | 925475.4044 |
| 3-(4-Ethoxyphenyl)-6-fluoro-4-methylcoumarin | 897509.9584 | 993115.5069 | 878340.9562 | 881085.9631 | 899312.3274 | 891303.1739 | 964712.0481 | 887748.8544 | 848771.5851 | 873481.6735 | 845692.176 | 899005.8392 |
| LPE O-16:2 | 344332. | 292365. | 485504. | 370104. | 530726. | 523829. | 746501. | 456654. | 503570. | 524838. | 508505. | 417235. |

| **Metabolites** | **PBS_1** | **PBS_2** | **PBS_3** | **PBS_4** | **BCG_1** | **BCG_2** | **BCG_3** | **BCG_4** | **BCGΔ*pe18*_1** | **BCGΔ*pe18*_2** | **BCGΔ*pe18*_3** | **BCGΔ*pe18*_4** |
|---|---|---|---|---|---|---|---|---|---|---|---|---|
| | 9347 | 6844 | 7858 | 8594 | 2994 | 182 | 4105 | 6606 | 8854 | 5542 | 1118 | 9934 |
| 4-Oxo-4-((3-oxodecan-2-yl)amino)butanoic acid | 1047508.665 | 1027373.184 | 991237.2651 | 1098690.523 | 920714.6078 | 941432.2387 | 921165.9498 | 930177.9739 | 981400.5437 | 959647.7213 | 952893.6135 | 977594.1018 |
| Cer 9:0;3O/12:1;(2OH) | 1548095.736 | 1393773.646 | 2287341.339 | 1671145.934 | 4915192.492 | 3608173.177 | 7422983.357 | 4704125.483 | 3208844.398 | 4081931.385 | 3922972.233 | 2316558.922 |
| Penbutolol | 35375575.89 | 34128306.13 | 19967634.76 | 30819955.13 | 32084189.7 | 27736644.32 | 30329727.77 | 30459100.08 | 30488725.49 | 33289163.82 | 26825347.63 | 27983473.48 |
| ST 24:1;O4;T/18:0 | 474929.4418 | 384319.7058 | 738143.9158 | 477211.927 | 803974.6432 | 974316.0664 | 1157091.114 | 897911.7047 | 591494.3249 | 818319.3887 | 988566.6937 | 663868.2796 |
| 6,9,12-Pentadecatrien-2-one | 5844772.051 | 5070513.398 | 4639948.279 | 5153582.932 | 5533047.912 | 4116332.283 | 3830381.423 | 6166953.169 | 6865537.255 | 6756829.959 | 5553884.759 | 5310021.892 |
| N1,N12-Diethylspermine | 5061647.503 | 6856392.409 | 3206074.049 | 4960359.583 | 2546137.636 | 2495805.645 | 2800082.575 | 3461055.108 | 3613149.658 | 3856254.545 | 2798845.787 | 2542496.933 |
| LPG O-14:0 | 377492.235 | 405422.4529 | 798592.5836 | 671540.6898 | 607851.8889 | 974618.8328 | 509622.77 | 804412.5223 | 497732.6574 | 218167.5667 | 352463.316 | 447127.8211 |
| Apo-10'-violaxanthal | 1265152.251 | 630771.1555 | 2149077.129 | 1375076.745 | 3439676.013 | 2744569.198 | 3364911.346 | 2639192.965 | 2343643.101 | 2554379.054 | 3375362.005 | 2478432.976 |
| PC(20:5(5Z,8Z,11Z,14Z,17Z)/P-16:0) | 7980050.041 | 6839011.28 | 3188463.68 | 6699603.548 | 4169651.371 | 3024278.4 | 3537557.158 | 4021024.377 | 5521086.596 | 13434803.9 | 3360068.563 | 3325813.093 |
| PC 9:0_26:1 | 40078341.89 | 13867361.72 | 37521958.11 | 24219770.05 | 22673415.91 | 289421.5202 | 23220440.03 | 8707627.547 | 46339214.49 | 27633036.36 | 20885577.75 | 3989425.126 |

| Metabolites | PBS_1 | PBS_2 | PBS_3 | PBS_4 | BCG_1 | BCG_2 | BCG_3 | BCG_4 | BCGΔ*pe18*_1 | BCGΔ*pe18*_2 | BCGΔ*pe18*_3 | BCGΔ*pe18*_4 |
|---|---|---|---|---|---|---|---|---|---|---|---|---|
| 1-Aminocyclopentanecarboxylic acid | 1474012.904 | 1022495.764 | 794192.8724 | 914417.5509 | 829824.4307 | 912349.4842 | 859520.9279 | 904420.8008 | 1049566.06 | 861505.907 | 942654.7297 | 951866.2358 |
| M463T516 | 428706.4847 | 290609.5609 | 1087406.404 | 599384.9648 | 695288.1943 | 1162347.704 | 711486.6586 | 1142463.646 | 656440.0033 | 440082.6043 | 482723 | 687416.1454 |
| Prostaglandin F2.alpha. dimethylamine | 2738638.565 | 2275840.294 | 1669854.354 | 2208491.421 | 684336.3467 | 1783026.362 | 1124569.309 | 803872.2923 | 1082683.484 | 805237.4791 | 824442.693 | 826752.3813 |
| gamma-Glutamylphenylalanine | 513991.6433 | 462739.6945 | 621726.5271 | 1902811 | 5442189.889 | 7638130.245 | 3786466.65 | 5833643.261 | 4408368.181 | 2914216.01 | 3080932.95 | 4671720.051 |
| 2,2'-Methylenebis(4-methyl-6-tert-butylphenol) | 581808.3271 | 1141175.16 | 684626.3595 | 829446.7695 | 1354257.596 | 1570039.925 | 2579335.723 | 1703060.892 | 1153530.014 | 1253288.308 | 1147295.795 | 1616997.418 |
| LPG O-19:1 | 74613.09512 | 129085.4726 | 53523.90243 | 85579.59167 | 176734.2153 | 158136.9131 | 129556.3658 | 132597.2157 | 207254.1266 | 261514.4603 | 160488.6618 | 112435.7379 |
| N-((5-Bromopyridin-3-yl)carbonyl)-.beta.-alanine | 919573.196 | 1059283.36 | 884032.89 | 1017583.707 | 1021336.293 | 956468.1423 | 1200550.693 | 876573.1448 | 852720.0385 | 991813.6253 | 854289.7065 | 764571.6826 |
| SM 40:4;3O | 293178.2522 | 818302.6366 | 502828.5761 | 1117977.037 | 1213677.562 | 0 | 2663664.962 | 4656605.365 | 1689505.289 | 418861.4589 | 707349.7152 | 2430504.43 |
| M491T64 | 495001.2898 | 566846.2498 | 421926.2568 | 453409.2653 | 538933.4018 | 437457.4014 | 576314.3311 | 420779.7932 | 495879.0514 | 383397.1734 | 535479.7296 | 139470.2397 |
| 4-(1-Pyrazolyl)benzaldehyde | 1017564.986 | 753872.6262 | 1046768.665 | 831089.6803 | 43472296.24 | 26115347.96 | 52324088.25 | 3836239.207 | 33011075.88 | 1879984.598 | 26037006.77 | 1004823.095 |

| **Metabolites** | **PBS_1** | **PBS_2** | **PBS_3** | **PBS_4** | **BCG_1** | **BCG_2** | **BCG_3** | **BCG_4** | **BCGΔ*pe18*_1** | **BCGΔ*pe18*_2** | **BCGΔ*pe18*_3** | **BCGΔ*pe18*_4** |
|---|---|---|---|---|---|---|---|---|---|---|---|---|
| Dibutyl phthalate | 9679141.836 | 12762729.37 | 14589908.74 | 10769428.87 | 18025017.22 | 20670732.11 | 14489301.48 | 17576198.45 | 16096234.73 | 12884920.83 | 15642578.44 | 13962998.71 |
| Heptylparaben | 510943.7621 | 453579.7939 | 342618.7821 | 472033.5665 | 477461.1619 | 356533.3219 | 372281.6363 | 342595.124 | 394578.0082 | 403363.8311 | 387982.2565 | 426764.1807 |
| PC 16:0_22:4 | 1085294.446 | 801445.1301 | 863248.9447 | 616996.3238 | 653756.1178 | 217757.1325 | 980322.1375 | 1054733.964 | 1021329.225 | 523100.3278 | 932097.4974 | 620600.5954 |
| Malonic acid | 3541349.399 | 3694496.594 | 5727361.725 | 6917198.996 | 5463140.83 | 5403920.347 | 12384927.23 | 10617475.91 | 4776787.976 | 5320126.518 | 10639237.32 | 5339423.973 |
| Tuberostemonine | 5167449.648 | 5097960.642 | 5036996.261 | 5170417.255 | 5112816.541 | 5180913.338 | 5163092.563 | 5186087.092 | 5186965.205 | 5183695.312 | 5300921.416 | 4710495.226 |
| Quinine di-N-oxide | 1216334.407 | 1212684.067 | 1156039.183 | 1156552.441 | 1298427.968 | 1327655.613 | 1389561.194 | 1306610.07 | 1290585.098 | 1299565.396 | 1297804.152 | 1208800.605 |
| Cytidine 5'-diphosphocholine (CDP-choline) | 244771.1256 | 223652.8193 | 213889.6935 | 279490.972 | 582563.54 | 287477.0673 | 583824.1502 | 414039.2623 | 387761.6322 | 583032.6696 | 529945.1581 | 342534.6209 |
| Leukotriene F4 | 951729.5206 | 1106439.97 | 1072537.103 | 1006918.156 | 1079912.591 | 1069113.097 | 1009842.172 | 1091537.078 | 1036587.5 | 999013.9528 | 1059710.379 | 1013294.422 |
| (2E,4E)-3,5,7-Trimethylocta-2,4-dienedioic acid | 67031.98171 | 124059.8131 | 93864.5564 | 62574.42028 | 9890015.62 | 7178977.093 | 6415036.063 | 5746989.323 | 3072202.293 | 3008740.869 | 3615828.123 | 2952923.834 |
| Glyzarin | 846981.2856 | 1307709.617 | 1110275.631 | 720634.6257 | 38091682.05 | 12925822.2 | 4488507.06 | 3951965.121 | 14565281.3 | 18123512.81 | 16608083.7 | 13232549.51 |
| Lovastatin | 1687116.271 | 1972208.109 | 619421.8973 | 2001233.576 | 1860330.138 | 1652473.86 | 1864714.981 | 1764966.857 | 2138353.108 | 3113786.888 | 1794536.226 | 1792552.169 |

| Metabolites | PBS_1 | PBS_2 | PBS_3 | PBS_4 | BCG_1 | BCG_2 | BCG_3 | BCG_4 | BCGΔ*pe18*_1 | BCGΔ*pe18*_2 | BCGΔ*pe18*_3 | BCGΔ*pe18*_4 |
|---|---|---|---|---|---|---|---|---|---|---|---|---|
| LPA 14:0 | 323182.9988 | 257115.9797 | 319563.9496 | 295203.0878 | 1228525.207 | 1382864.261 | 1004905.369 | 1100219.495 | 844568.6265 | 1047493.595 | 1075096.607 | 924314.1156 |
| Dienogest | 3021438.076 | 3239710.471 | 2447921.76 | 2710628.943 | 2253626.738 | 2252312.132 | 2299656.354 | 1883336.565 | 2246998.059 | 2487957.757 | 2009236.224 | 2670663.937 |
| 1-Pyrrolidinecarboximidamide | 1777914.645 | 2002750.97 | 1537166.731 | 1765120.511 | 1848605.8 | 1689582.435 | 1743277.198 | 2177159.973 | 1677262.844 | 1698798.713 | 1631136.027 | 2244077.368 |
| Ligustilide | 1002012.742 | 1215108.221 | 1471030.99 | 1159983.979 | 898020.2433 | 1167638.913 | 1146298.49 | 857151.5579 | 937910.7827 | 876040.3561 | 1069479.111 | 1054754.83 |
| PE-Cer 13:1;2O/18:4 | 417025.2098 | 363183.8249 | 550264.2547 | 445655.3316 | 169567.1494 | 131604.6872 | 223174.5285 | 151677.3693 | 98238.90193 | 91842.78664 | 132076.5809 | 74244.71936 |
| Ethanol, 2-[[3-(2,3-dichlorophenoxy)propyl]amino]- | 840718.3472 | 659947.5076 | 1526491.918 | 1103894.97 | 1860234.323 | 1681346.736 | 1970927.43 | 1654355.31 | 1825493.298 | 952387.4443 | 1127345.12 | 1398192.108 |
| PS 14:0_26:4 | 3439423.41 | 1748064.322 | 3077742.421 | 2788314.11 | 1846504.238 | 651324.8286 | 1852922.507 | 2312846.236 | 3480966.74 | 2716919.472 | 1120727.906 | 3153911.99 |
| 3-[8-(2-Hydroxypropan-2-yl)-3-methoxycarbonyl-3,4a,7,10a-tetramethyl-5-methylidene-1,4-dioxo-6,6a,8,9,10,10b-hexahydrobenzo[h]isochromen-7-yl]propan | 827871.1733 | 945561.4044 | 839784.8842 | 859550.5929 | 886975.5871 | 1000685.769 | 922562.0638 | 915665.3735 | 890777.1717 | 903582.3301 | 953180.2888 | 897096.8544 |

| Metabolites | PBS_1 | PBS_2 | PBS_3 | PBS_4 | BCG_1 | BCG_2 | BCG_3 | BCG_4 | BCGΔ *pe18*_1 | BCGΔ *pe18*_2 | BCGΔ *pe18*_3 | BCGΔ*pe18*_4 |
|---|---|---|---|---|---|---|---|---|---|---|---|---|
| oic acid | | | | | | | | | | | | |
| gamma-Dodecalactone | 2004314.738 | 1342434.068 | 1718758.267 | 1038078.078 | 1996520.46 | 1493203.943 | 1620247.043 | 909828.7928 | 1780556.197 | 1655298.7 | 2156124.063 | 1615060.599 |
| 2-(4-Pyridinyl)ethanol | 19259167.46 | 9715069.47 | 12211073.44 | 9111118.266 | 9744565.64 | 10092074.88 | 8473589.953 | 9025700.439 | 9793698.75 | 9399442.541 | 8819563.713 | 7709119.119 |
| DG 36:7 | 313970.0115 | 585784.7803 | 115833.0862 | 611554.9964 | 302024.6271 | 412648.5139 | 335218.4814 | 537132.5427 | 311836.9768 | 622668.0329 | 295234.911 | 396874.6434 |
| PC(38:2) | 2430326.001 | 5284799.441 | 8095792.062 | 5443234.475 | 19847924.03 | 9667989.565 | 22445121.97 | 4788101.884 | 16408547.74 | 23503206.3 | 17295116.14 | 9581412.13 |
| Albendazole | 2483612.522 | 2521067.071 | 1939301.479 | 1278714.863 | 4848669.447 | 2271153.518 | 3391561.483 | 3571058.565 | 2485058.69 | 2852543.448 | 2333628.12 | 2034533.316 |
| N1-Ethyl-N2-(4-nitrophenyl)-1,2-ethanediamine | 541073.8882 | 556752.4392 | 470343.4811 | 454683.5817 | 487159.49 | 482098.6085 | 470119.8116 | 534415.3029 | 537352.7538 | 462234.7583 | 487320.1626 | 472293.9746 |
| N-(3-Chloro-2-methylphenyl)-1-cyclohexyl-5-oxopyrrolidine-3-carboxamide | 1195177.587 | 1161448.047 | 1060499.617 | 1142017.216 | 898481.7558 | 2079661.634 | 841918.3861 | 802483.3133 | 929667.2834 | 960084.7434 | 856880.0064 | 771720.1897 |
| PC 8:0_28:3 | 116728 | 122983 | 110416 | 131029 | 107934 | 86235.7 | 106819 | 824672 | 104114 | 109331 | 106024 | 122201 |

| Metabolites | PBS_1 | PBS_2 | PBS_3 | PBS_4 | BCG_1 | BCG_2 | BCG_3 | BCG_4 | BCGΔ *pe18*_1 | BCGΔ *pe18*_2 | BCGΔ *pe18*_3 | BCGΔ*pe18*_4 |
|---|---|---|---|---|---|---|---|---|---|---|---|---|
|  | 04.3 | 65.75 | 61.6 | 23.7 | 60.07 | 5365 | 84.07 | 0.56 | 11.64 | 60.03 | 04.35 | 25.87 |
| PE-Cer 14:3;2O/12:1;O | 1300303.507 | 1574314.166 | 709221.3195 | 1394382.306 | 835144.8358 | 1064705.211 | 967769.7248 | 1057856.193 | 750516.2369 | 1233712.228 | 923317.1827 | 1183785.203 |
| Dihydrojasmonic Acid | 1425712.835 | 1352947.877 | 1287516.562 | 1528817.263 | 1385770.785 | 1383468.497 | 1450240.337 | 1339491.475 | 1341786.795 | 1390326.851 | 1406880.009 | 1443562.838 |
| PS 18:1_18:2 | 3144881.914 | 684052.6773 | 1542751.33 | 1572726.192 | 4667020.551 | 193644.2876 | 795361.6064 | 495007.0182 | 1240674.372 | 1158752.025 | 1234264.146 | 280451.7999 |
| PE(P-38:4) | 94530148.45 | 109719855.1 | 133065652.3 | 112676202.7 | 93635996.7 | 94457003.48 | 126096990.2 | 115946729.9 | 80140383.85 | 126205893.4 | 98776116.52 | 66495555.01 |
| PC O-24:0 | 858668.2609 | 723065.9181 | 1112734.402 | 733708.0502 | 419783.7715 | 1078169.773 | 684665.7603 | 771769.0699 | 789486.8791 | 534388.0813 | 671649.5828 | 863129.7898 |
| Monomethyl glutaric acid | 2627918.891 | 2354748.98 | 2221437.961 | 2075213.509 | 2533295.806 | 2864647.27 | 3964119.076 | 2307136.928 | 2523437.491 | 2254014.5 | 2862986.932 | 2849226.625 |
| 2-Aminoadipic acid | 224112.8297 | 216616.8301 | 271197.5213 | 219482.6833 | 920037.0218 | 645100.4851 | 858388.3674 | 699315.5283 | 1203892.765 | 1058429.639 | 1377301.055 | 1110594.251 |
| Desferrioxamine B | 2342561.51 | 2911882.254 | 1948789.917 | 2793036.872 | 2481875.886 | 2792979.929 | 2087279.192 | 1939485.795 | 2043821.684 | 2112516.634 | 2073425.142 | 2160545.58 |
| Ethanone, 2-(4-methylphenyl)-1-(1-pentyl-1H-indol-3-yl)- | 537925.373 | 557150.1884 | 445409.8717 | 491869.8346 | 222584.4372 | 498210.4133 | 510534.7527 | 476568.9862 | 501140.2259 | 478172.635 | 232882.7976 | 245302.1407 |
| FAHFA 18:2/20:4 | 102896.9378 | 103371.3223 | 81780.57661 | 93391.55703 | 2656523.28 | 7091642.26 | 2206485.642 | 2864643.344 | 1667642.038 | 848549.9971 | 2560104.559 | 1974500.695 |
| Karbutilate | 437373 | 521684 | 405229 | 507392 | 632696 | 603951 | 634152 | 576691 | 553055 | 569103 | 512000 | 755186 |

| Metabolites | PBS_1 | PBS_2 | PBS_3 | PBS_4 | BCG_1 | BCG_2 | BCG_3 | BCG_4 | BCGΔ *pe18*_1 | BCGΔ *pe18*_2 | BCGΔ *pe18*_3 | BCGΔ*pe18*_4 |
|---|---|---|---|---|---|---|---|---|---|---|---|---|
| | 19.73 | 23.41 | 84.21 | 45.93 | 13.53 | 04.3 | 33.94 | 38.77 | 08.54 | 93.72 | 45.11 | 82.4 |
| PI 2:0_13:1 | 579791.8918 | 611127.3091 | 843665.876 | 792201.4769 | 759257.9939 | 968506.6903 | 904424.1963 | 749463.3563 | 774987.5717 | 487355.0698 | 488711.2002 | 782696.5416 |
| PC O-40:9 | 473509.3037 | 621796.908 | 537725.2915 | 639349.8367 | 656197.4293 | 547101.881 | 633192.6538 | 653389.0764 | 660263.3227 | 614771.7482 | 510910.6349 | 653384.7264 |
| Pelletierine | 15292064.06 | 5071767.516 | 7249707.16 | 6642336.298 | 4440721.014 | 32957319.48 | 7466187.76 | 17995728.13 | 4210794.486 | 6574130.714 | 9971726.102 | 7733025.136 |
| Hexadecadienoylcarnitine (Car(16:2)) | 593511.3169 | 477696.5798 | 1126939.469 | 600950.3365 | 873607.8072 | 736930.9153 | 1068571.276 | 978575.6298 | 518572.9702 | 0 | 587773.5166 | 426041.159 |
| Methyl (2R)-((4S,4aS,7R,7aR,8R,11aS,12aS)-4-(furan-3-yl)-4a,7a,9,9-tetramethyl-13-methylidene-2,10-dioxododecahydro-4H-7,12a-methanopyrano[4,3-b][1]benzoxocin-8-yl)(hydroxy)acetate | 694179.2034 | 700617.1482 | 715163.9791 | 681643.7171 | 722971.0685 | 672195.7977 | 733430.162 | 735114.518 | 699087.9154 | 727453.8003 | 659122.4187 | 567443.6774 |
| 1-(1-Piperidinyl)-3-(1,2,3,4-tetrahydro-9H-carbazol-9-yl)-2-propanol | 978286.6703 | 1170360.23 | 840987.1971 | 769380.4796 | 996009.7034 | 780203.6294 | 1014403.866 | 1092955.464 | 1116975.075 | 931259.5406 | 855025.9799 | 718611.2215 |

| Metabolites | PBS_1 | PBS_2 | PBS_3 | PBS_4 | BCG_1 | BCG_2 | BCG_3 | BCG_4 | BCGΔ *pe18*_1 | BCGΔ *pe18*_2 | BCGΔ *pe18*_3 | BCGΔ*pe18*_4 |
|---|---|---|---|---|---|---|---|---|---|---|---|---|
| SHexCer 22:1;2O/26:0;O | 398465.4394 | 398442.6954 | 336952.5924 | 547792.6717 | 313931.3892 | 283586.8986 | 398784.8172 | 434832.0822 | 386360.7519 | 329406.3224 | 349464.9961 | 464174.0034 |
| Glycoursodeoxycholic acid | 977378.0212 | 823916.2479 | 1443940.316 | 958467.455 | 1636621.286 | 1607943.031 | 2323764.907 | 1508212.308 | 1375287.552 | 960682.4198 | 988933.4887 | 1083126.185 |
| (2-{[2-[hexadec-9-enoyloxy]-3-(tetradecanoyloxy)propyl phosphonato]oxy}ethyl)trimethylazanium | 1343124.187 | 1507505.748 | 1958475.125 | 1366600.954 | 11890971.75 | 8402583.076 | 12061233.31 | 9353439.685 | 10209944.61 | 12671319.9 | 11106806.34 | 5920003.879 |
| 4,5-Epoxy-7Z,10Z,13Z,16Z,19Z-docosapentaenoic acid, methyl ester | 1788932.2 | 1724835.802 | 1172421.151 | 6294616.007 | 1806924.774 | 1919096.38 | 6596789.525 | 2904241.689 | 5101564.157 | 2405050.473 | 1929593.798 | 7849891.639 |
| SM 37:5;3O | 596684.0381 | 377405.4068 | 517339.7982 | 623986.3331 | 508550.2965 | 27066.10762 | 639901.9686 | 863584.7052 | 89630.24678 | 359635.1743 | 630380.7858 | 872408.9407 |
| LPE(22:6) | 2646808.81 | 2698847.479 | 3311966.418 | 2604563.314 | 5284504.6 | 5346316.554 | 4131838.092 | 5278303.171 | 5037673.128 | 4966491.8 | 4828797.488 | 5241858.785 |
| Leucine | 16375892.26 | 13243922.04 | 14555978.35 | 17865563.53 | 13671128.08 | 13679648.4 | 19263540.74 | 24609234.54 | 15785040.42 | 15091386.41 | 12825853.21 | 14514744.8 |
| Atrazine | 392436.3005 | 384722.2459 | 340007.3841 | 350275.7506 | 325846.6217 | 322855.9427 | 332225.0353 | 328182.1238 | 323846.0234 | 319643.1249 | 317569.0384 | 337211.2625 |
| 4,6-Dichloro-2-cyclopropyl-5-methylpy | 1099092.582 | 813886.8976 | 1626504.142 | 1058808.249 | 2190391.772 | 1667148.894 | 2425187.756 | 1922008.866 | 1602047.303 | 1344663.519 | 1469564.912 | 1591466.662 |

| **Metabolites** | **PBS_1** | **PBS_2** | **PBS_3** | **PBS_4** | **BCG_1** | **BCG_2** | **BCG_3** | **BCG_4** | **BCGΔ*pe18*_1** | **BCGΔ*pe18*_2** | **BCGΔ*pe18*_3** | **BCGΔ*pe18*_4** |
|---|---|---|---|---|---|---|---|---|---|---|---|---|
| rimidine | | | | | | | | | | | | |
| N-Undecanoylglycine | 1730889.1 | 1292365.91 | 1525169.618 | 1785157.473 | 1433206.9 | 1223819.605 | 1616726.124 | 1557123.796 | 2650681.909 | 1194465.757 | 2871276.899 | 1198467.654 |
| dimethyl(tetradecyl)amine | 1554124.829 | 1506527.467 | 1380047.131 | 1508594.501 | 1552600.348 | 1492086.101 | 1572239.146 | 1522366.475 | 1513592.317 | 1465583.315 | 1527038.145 | 1422129.385 |
| Himbacine | 1160420.41 | 2020655.153 | 1020587.527 | 1089280.819 | 1295807.378 | 1266168.951 | 1171437.789 | 1196183.654 | 1971553.949 | 1182336.749 | 1144542.269 | 1134558.16 |
| PI 8:0_26:0 | 1333560.721 | 1215806.461 | 1419283.21 | 1338075.712 | 1473948.421 | 215993.4844 | 1350639.631 | 1925556.058 | 1772704.45 | 1042060.339 | 1115698.515 | 2862704.465 |
| Tri-Boc-hydrazinoacetic acid | 35543.02963 | 32906.80125 | 70638.00357 | 5692.628683 | 416127.6944 | 283533.7663 | 574203.4238 | 224163.4658 | 345614.3547 | 459546.9438 | 785096.0734 | 173840.8044 |
| Ginkgolic acid I | 652043.552 | 395860.601 | 425023.5591 | 561859.3712 | 3090642.758 | 4114509.652 | 1510802.229 | 2807664.069 | 4146810.541 | 1360041.905 | 1518614.221 | 3955458.112 |
| 2-Isopropylmalic acid | 661238.9901 | 605361.1204 | 546571.8097 | 531895.0449 | 15002523.28 | 7546254.865 | 9712871.644 | 7272613.69 | 9345675.286 | 11280232 | 12064901.5 | 9194907.26 |
| Dikegulac acid | 1461925.1 | 1454623.567 | 1434497.939 | 1456508.457 | 1434899.179 | 1426385.407 | 1539059.948 | 1494131.27 | 1470878.61 | 1490635.022 | 1455486.458 | 1469131.555 |
| Kaempferol 3-.alpha.-L-arabinoside | 732591.6068 | 631875.4487 | 635843.9599 | 748410.4142 | 630021.6531 | 582717.1838 | 638358.3505 | 638529.8275 | 597082.8065 | 600905.8216 | 750958.7697 | 746362.9339 |
| (-)-erythro-Anethole_glycol_2-glucoside | 199346.6761 | 653491.6299 | 88330.944 | 946492.6277 | 4657262.181 | 2898049.163 | 77423.20979 | 115513.6805 | 679754.5265 | 1869491.513 | 1925665.94 | 74860.24954 |

| Metabolites | PBS_1 | PBS_2 | PBS_3 | PBS_4 | BCG_1 | BCG_2 | BCG_3 | BCG_4 | BCGΔ*pe18*_1 | BCGΔ*pe18*_2 | BCGΔ*pe18*_3 | BCGΔ*pe18*_4 |
|---|---|---|---|---|---|---|---|---|---|---|---|---|
| 2-(4-Fluorophenyl)-2,3-dihydro-1H-perimidine | 1760515.065 | 1252272.272 | 2149841.75 | 633698.0214 | 1829375.716 | 1436091.874 | 4052855.711 | 1113824.877 | 1159186.718 | 628851.1562 | 1371286.4 | 1985046.263 |
| SM 8:1;2O/3:0 | 3984.230125 | 0 | 0 | 0 | 198593.8318 | 133465.3882 | 237114.8397 | 193264.1602 | 170602.8018 | 299453.0441 | 204229.899 | 168294.4494 |
| SL 11:1;O/16:2 | 2478916.272 | 2872408.655 | 2100135.398 | 3315856.522 | 3207531.118 | 3437638.309 | 2712445.098 | 2612206.118 | 2492485.596 | 2227102.16 | 1951555.942 | 2345328.761 |
| 1,2-Dimethylimidazole | 3928386.528 | 3886151.829 | 4014064.798 | 3969464.442 | 3761455.275 | 4027969.178 | 3679723.826 | 3622081.373 | 3673888.891 | 3680291.176 | 3814778.509 | 3928952.12 |
| Cer 8:0;2O/8:0 | 529982.7265 | 514031.4936 | 476900.4515 | 580325.4172 | 516674.5016 | 589087.3851 | 545126.358 | 555150.5997 | 576039.1889 | 625147.8091 | 571184.232 | 532597.0262 |
| LNAPS 3:0/N-18:1 | 1256014.495 | 1215849.849 | 3096537.411 | 1444778.822 | 1031938.899 | 885269.9775 | 1264064.333 | 1201516.689 | 772638.4391 | 1446474.431 | 992912.244 | 766797.3806 |
| N,N-Diethyl-4-piperidinamine | 1153565.415 | 943131.9597 | 766927.7279 | 904862.6268 | 662917.8901 | 746659.3691 | 471455.1822 | 306865.1084 | 524259.8923 | 632365.3232 | 642184.2618 | 648162.9984 |
| 2-Pyrrolidinone, 5-[(3S)-3-hydroxy-4-phenyl-1-buten-1-yl]-1-[6-(2H-tetrazol-5-yl)hexyl]-, (5R)- | 582027.2879 | 649243.9506 | 560821.9321 | 588515.9204 | 609568.7831 | 605922.6375 | 625257.4289 | 643373.5865 | 566285.0175 | 609724.6206 | 586849.3148 | 670096.5122 |
| CerP(d18:1/22:0) | 2094365.551 | 1713336.666 | 2384572.031 | 1485652.406 | 4018670.125 | 3701097.19 | 4926160.276 | 4423845.774 | 5030056.251 | 5924215.824 | 4798787.471 | 2427399.243 |
| 1-Hexadecyl-1H-indole-2,3-dione | 2199508.025 | 2452331.354 | 2370024.206 | 2260206.756 | 2409038.942 | 1991165.19 | 2258074.098 | 2510437.615 | 1617634.633 | 1334292.435 | 2061147.859 | 1202618.022 |

| Metabolites | PBS_1 | PBS_2 | PBS_3 | PBS_4 | BCG_1 | BCG_2 | BCG_3 | BCG_4 | BCGΔ*pe18*_1 | BCGΔ*pe18*_2 | BCGΔ*pe18*_3 | BCGΔ*pe18*_4 |
|---|---|---|---|---|---|---|---|---|---|---|---|---|
| Arachidonoyl-2'-fluoroethylamide | 7346987.084 | 7016734.27 | 6496647.435 | 6686869.183 | 7518585.934 | 7770124.113 | 7278256.154 | 7071109.132 | 7097337.84 | 6798341.534 | 7464784.143 | 6267042.85 |
| Geniposide (Not validated) | 459636.5442 | 470576.772 | 860458.5293 | 567218.7768 | 1668078.902 | 1096304.169 | 1288530.769 | 1307410.882 | 1779363.268 | 1261755.34 | 1026803.552 | 1211477.166 |
| PI O-16:0_22:5 | 13720748.26 | 10431247.44 | 6106191.077 | 13864453.87 | 2188014.554 | 1873304.846 | 2651559.365 | 3556220.071 | 3952299.889 | 2970875.934 | 2581583.681 | 1356140.983 |
| SL 10:1;O/22:4 | 1521167.911 | 1906993.133 | 1649310.502 | 1596957.228 | 777082.8932 | 849654.6281 | 798293.4893 | 719113.4148 | 729291.9392 | 855850.3801 | 1202606.923 | 590625.5517 |
| PS 13:1_22:4 | 768778.9274 | 888585.3824 | 668912.8104 | 892573.3297 | 824684.8845 | 863256.9123 | 995006.2544 | 866207.841 | 878386.235 | 825815.1082 | 852055.3923 | 768033.8223 |
| 1,2-Dipalmitoleoyl-sn-glycero-3-phosphocholine | 24968891.21 | 14907221.35 | 9451188.926 | 11319634.91 | 86417957.7 | 75227778.12 | 78363003.15 | 83397111.02 | 67348214.23 | 98428588.49 | 69283094.81 | 59446624.92 |
| PC O-29:2 | 593321.4997 | 534687.3632 | 552364.3688 | 485948.914 | 483384.3653 | 483190.0337 | 417908.2254 | 579819.8732 | 508373.7243 | 526542.5161 | 514810.0301 | 473999.4335 |
| Cucurbitin | 73105.58028 | 583643.7959 | 650832.7389 | 555393.6177 | 725045.0319 | 388426.6169 | 1418492.898 | 201719.4445 | 590543.8809 | 549042.5018 | 328812.4436 | 3404093.982 |
| ST 24:1;O4;T/15:0 | 132686.6908 | 147665.0903 | 204927.2881 | 150693.4084 | 132189.5907 | 104679.7645 | 160132.0452 | 138316.5476 | 92615.6861 | 86351.16715 | 83668.56361 | 65629.39229 |
| N-.alpha.-(tert-Butoxycarbonyl)-L-proline | 1161641.981 | 368373.7466 | 440471.0273 | 1082676.252 | 4109938.893 | 8509885.81 | 8162001.831 | 7668679.274 | 6214497.019 | 4812996.825 | 4866760.495 | 6157066.366 |
| N-Naphthalen-2-yl-5-oxoprolinamide | 3662723.174 | 4358875.764 | 2434963.247 | 2713349.433 | 3496778.499 | 3037527.647 | 3334767.029 | 3168706.692 | 2475686.371 | 2434191.692 | 3348027.775 | 2411316.41 |

| Metabolites | PBS_1 | PBS_2 | PBS_3 | PBS_4 | BCG_1 | BCG_2 | BCG_3 | BCG_4 | BCGΔ*pe18*_1 | BCGΔ*pe18*_2 | BCGΔ*pe18*_3 | BCGΔ*pe18*_4 |
|---|---|---|---|---|---|---|---|---|---|---|---|---|
| Dihydroterrein | 639676.0042 | 505143.0593 | 470742.9703 | 487786.2613 | 524342.6879 | 515466.0361 | 485036.7322 | 471342.1598 | 485815.0419 | 507065.6225 | 503239.0047 | 510246.3823 |
| LPG 22:3 | 318486.0075 | 299438.1 | 362293.186 | 342901.0546 | 441726.6919 | 404121.6188 | 624681.9708 | 468344.0516 | 401833.9243 | 559120.381 | 565633.1794 | 417992.7133 |
| N-(3-Methoxypropyl)-9H-purin-6-amine | 897855.0425 | 1077098.3 | 865745.5015 | 976587.069 | 1202024.395 | 1197405.705 | 1302926.524 | 1122111.044 | 1115224.865 | 1106031.583 | 1177870.445 | 1142705.919 |
| 4-Oxohexanoic acid | 3737502.271 | 3281309.418 | 3921972.428 | 3890210.469 | 3307583.242 | 3026924.566 | 3016867.142 | 1865337.854 | 2928508.25 | 3300237.114 | 2555864.44 | 2821489.742 |
| Pronethalol | 4093693.662 | 3150092.09 | 5251848.657 | 2731492.617 | 3472357.675 | 3601294.802 | 3493032.546 | 4072848.457 | 3731644.567 | 3564051.431 | 3519203.988 | 8143718.474 |
| 5-(Furan-2-yl)thiophene-2-carboxylic acid | 1931975.62 | 1761108.251 | 3003995.249 | 1702498.389 | 4267276.897 | 3848056.657 | 4787477.902 | 3453758.988 | 3503689.811 | 3632192.624 | 3324180.746 | 3162892.443 |
| 1-Deoxymannojirimycin (hydrochloride) | 3852286.672 | 7078851.14 | 7441767.208 | 11409839.44 | 4917978.885 | 5560222.51 | 2268178.667 | 2193834.832 | 2907040.949 | 3287479.731 | 6233894.435 | 6524766.826 |
| PG 4:0_13:0 | 193261.7691 | 1440974.968 | 1122450.073 | 1132991.701 | 584742.2239 | 608062.2449 | 392628.8041 | 494709.018 | 529202.28 | 351166.4402 | 333358.1335 | 342080.223 |
| N,N-Dimethyldodecylamine | 3719051.976 | 5230940.987 | 4219357.905 | 5772593.313 | 4977468.927 | 5142722.425 | 4590450.315 | 4908489.021 | 4473028.408 | 4187618.467 | 4013762.66 | 5213520.38 |
| Royal jelly acid | 1746530.198 | 1741333.641 | 1916262.72 | 1705234.212 | 1390265.939 | 1620479.474 | 1657782.564 | 1480068.054 | 1518191.726 | 2355155.245 | 1688529.572 | 2432846.301 |
| SM 9:0;2O/24:1 | 135695 | 760798. | 140520 | 153441 | 582991. | 0 | 184945 | 163155 | 144710 | 645124. | 526554. | 939268. |

| Metabolites | PBS_1 | PBS_2 | PBS_3 | PBS_4 | BCG_1 | BCG_2 | BCG_3 | BCG_4 | BCGΔ*pe18*_1 | BCGΔ*pe18*_2 | BCGΔ*pe18*_3 | BCGΔ*pe18*_4 |
|---|---|---|---|---|---|---|---|---|---|---|---|---|
| | 9.762 | 887 | 7.033 | 6.383 | 0944 | | 0.93 | 3.359 | 6.212 | 8444 | 9334 | 3378 |
| S-Adenosylhomocysteine | 191324.2834 | 95819.57816 | 286940.5027 | 148126.1806 | 1401165.072 | 1297695.173 | 1628559.908 | 1502909.019 | 1754081.881 | 1335744.328 | 1813959.494 | 1942137.548 |
| PG(18:1(9Z)/20:4(5Z,8Z,11Z,14Z)) | 2538462.593 | 2057610.209 | 4342662.074 | 4729678.179 | 16052009.16 | 4095642.022 | 6858573.264 | 5113450.553 | 4457738.212 | 8020604.865 | 6006903.499 | 8265213.714 |
| LPC 16:4-SN2 | 3499.51746 | 42175.75671 | 4152.440346 | 53311.70299 | 142393.0809 | 92284.04856 | 143246.8966 | 132763.8202 | 87644.82686 | 126649.425 | 128883.1446 | 59991.24861 |
| 1,3-Dicyclohexylurea | 24261524.18 | 41274560.28 | 37464887.98 | 37493332.12 | 26514881.9 | 27233120.18 | 40207743.38 | 28101377.53 | 38687927 | 29758608.5 | 29830544.22 | 35614739.44 |
| Hexamethylquercetagetin | 146083.6904 | 97622.46303 | 215708.5213 | 142022.2983 | 1345355.83 | 1733898.318 | 738350.1434 | 1413107.073 | 1343027.379 | 1145779.339 | 904304.7257 | 2786191.639 |
| 4,6-Dihydroxy-3-(1-hydroxyethyl)-5-methoxy-3H-2-benzofuran-1-one | 151211505.8 | 93733993.69 | 199286918.7 | 81423382.52 | 169794913 | 155936557.6 | 124403290.2 | 101146619.6 | 118214872.8 | 92802766.08 | 90789775.02 | 54386466.08 |
| PC O-27:2 | 440618.9262 | 379659.9774 | 423620.7587 | 385927.8763 | 316233.6504 | 499875.3275 | 572625.6537 | 553874.164 | 323863.0251 | 379895.7506 | 482284.2497 | 278357.8914 |
| N-Stearoyl Threonine | 281984.905 | 239723.8045 | 312336.675 | 279418.1096 | 1559439.016 | 1422388.345 | 1529409.595 | 1337956.839 | 1065807.144 | 1152780.613 | 1665638.301 | 619038.0308 |
| UDP-2-acetamido-2-deoxy-alpha-ribo-hex-3-uluronate | 153074.2627 | 120514.9652 | 190964.9154 | 106385.0343 | 159426.5051 | 213732.6763 | 168412.8585 | 164269.485 | 154739.5481 | 162260.1769 | 175196.1826 | 163499.7732 |
| 4-Aminohex-5-ynoic acid | 3818327.155 | 3437167.447 | 5132533.248 | 3687851 | 4218217.459 | 6176440.046 | 4179083.268 | 7429700.15 | 5416257.886 | 6184108.692 | 5738888.656 | 4001140.748 |

| Metabolites | PBS_1 | PBS_2 | PBS_3 | PBS_4 | BCG_1 | BCG_2 | BCG_3 | BCG_4 | BCGΔ*pe18*_1 | BCGΔ*pe18*_2 | BCGΔ*pe18*_3 | BCGΔ*pe18*_4 |
|---|---|---|---|---|---|---|---|---|---|---|---|---|
| PC 8:0_32:7 | 21344659.49 | 25270042.71 | 11845447.34 | 27537204.35 | 36034898.05 | 38549783.91 | 32709318.01 | 37914919.51 | 35233245.05 | 55225769.25 | 38020855.94 | 42224071.15 |
| Lariciresinol | 1092269.6 | 1004850.914 | 1212881.8 | 1211616.016 | 2006994.828 | 1880529.554 | 1765135.401 | 1755233.314 | 1674503.472 | 1127472.621 | 1416806.999 | 1521293.928 |
| DGCC 9:0_30:7 | 2107688.719 | 5954654.368 | 2033488.792 | 1825429.555 | 4082784.332 | 2550657.658 | 3695727.562 | 2610155.158 | 3283292.923 | 4253742.14 | 3237642.368 | 2209448.652 |
| PC O-10:0_4:0 | 211141.8585 | 186797.3876 | 314672.7885 | 161713.1245 | 1635210.756 | 1169499.173 | 1868073.254 | 1351726.214 | 1150276.707 | 960185.8642 | 1757953.218 | 858590.4927 |
| LPG 21:1 | 389116.0878 | 435878.9512 | 377164.062 | 397650.4149 | 370946.9436 | 455291.1152 | 401797.5427 | 371934.5303 | 439143.7177 | 440914.3966 | 417830.6782 | 407839.9152 |
| (3-Oxo-2-piperazinyl)acetic acid | 10244974.53 | 6859163.374 | 5032436.427 | 1843384.4 | 2549153.234 | 6414431.146 | 2160565.805 | 5999867.701 | 2404218.518 | 3102256.316 | 1719031.606 | 2095846.697 |
| 2'-Hydroxy-5'-methylacetophenone | 1354788.059 | 1340383.755 | 1655224.445 | 1488469.605 | 1533787.596 | 1690916.787 | 1580695.619 | 1318905.915 | 1728622.346 | 1359242.775 | 1324397.399 | 1883944.536 |
| N-cis-Tetradec-9-enoyl-L-homoserine lactone | 2919481.63 | 2761940.327 | 1797639.2 | 2616987.696 | 3190740.658 | 2448811.857 | 1965010.885 | 2234794.062 | 2128239.224 | 2326736.854 | 2274910.093 | 1972379.577 |
| Fenamiphos | 3747330.349 | 2800264.758 | 4629833.237 | 3156335.89 | 7268034.284 | 4545923.319 | 5526840.237 | 5697202.605 | 5408516.253 | 4764526.35 | 4576079.589 | 3600648.981 |
| LNAPS 24:6/N-16:0 | 1746986.6 | 1284176.369 | 1549751.352 | 1454365.914 | 1584353.132 | 691006.2219 | 1371770.445 | 1081854.884 | 1057041.526 | 1149505.301 | 899163.7883 | 1276989.902 |
| Serine-Cholic Acid | 1264926.944 | 1377586.191 | 1102833.409 | 1259476.526 | 1315733.72 | 1192618.882 | 1331582.984 | 1222487.282 | 1384301.627 | 1272772.055 | 1162180.128 | 1153727.97 |

| **Metabolites** | **PBS_1** | **PBS_2** | **PBS_3** | **PBS_4** | **BCG_1** | **BCG_2** | **BCG_3** | **BCG_4** | **BCGΔ*pe18*_1** | **BCGΔ*pe18*_2** | **BCGΔ*pe18*_3** | **BCGΔ*pe18*_4** |
|---|---|---|---|---|---|---|---|---|---|---|---|---|
| N-Nonanoyl-L-homoserine lactone | 8198068.148 | 10923224.32 | 9978250.402 | 10456953.92 | 7434172.103 | 6468424.802 | 8568025.964 | 7533671.574 | 6406379.941 | 6737159.884 | 6353731.863 | 9483588.401 |
| 1,2-Dithiolane-3-pentanamide, N,N'-1,3-propanediylbis- | 527308.8844 | 86434.0064 | 676904.819 | 164707.477 | 795111.965 | 943070.0697 | 3726708.993 | 3068950.005 | 1309435.86 | 827518.4212 | 1341992.009 | 467663.3422 |
| 5a,6-Dihydroxy-13-methoxy-5,5,7a,9,14b-pentamethyl-1,2,5a,6,7,7a,10,14,14a,14b-decahydro-5H-furo[3,4-i]oxepino[4,3-a]xanthene-3,12-dione | 3372402.462 | 383561.5393 | 2106043.266 | 345237.9903 | 379262.05 | 2809917.865 | 970977.6439 | 1284841.713 | 3173609.821 | 497290.6497 | 3508121.725 | 466080.3058 |
| 10-Methylbenzo[g]pteridine-2,4(3H,10H)-dione | 67082.26592 | 135858.0433 | 165885.9888 | 144756.979 | 981398.1796 | 517817.2175 | 960343.7528 | 678012.6581 | 417781.9362 | 861428.6299 | 713052.5179 | 476835.6049 |
| 1-Oleoyl-2-acetyl-sn-glycerol | 7073130.174 | 6267231.49 | 3718976.399 | 6569982.785 | 19415278.63 | 16182054.73 | 7339283.245 | 18242324.81 | 12513363 | 4349638.088 | 5055325.24 | 11722355.48 |
| Adenylyl sulfate | 168447.5625 | 229762.8244 | 204665.9029 | 372036.8865 | 145621.833 | 157562.718 | 521885.2849 | 158148.3677 | 165062.643 | 168214.51 | 187740.3507 | 329067.9349 |

| Metabolites | PBS_1 | PBS_2 | PBS_3 | PBS_4 | BCG_1 | BCG_2 | BCG_3 | BCG_4 | BCGΔ*pe18*_1 | BCGΔ*pe18*_2 | BCGΔ*pe18*_3 | BCGΔ*pe18*_4 |
|---|---|---|---|---|---|---|---|---|---|---|---|---|
| 3-Hydroxy-3-methyl-5-oxo-5-(((2R,3S,4S,5R,6R)-3,4,5-trihydroxy-6-(octan-3-yloxy)tetrahydro-2H-pyran-2-yl)methoxy)pentanoic acid | 10576642.96 | 5054922.705 | 2056827.11 | 4144608.322 | 8625918.294 | 8140003.687 | 6305588.059 | 9621037.05 | 8838142.093 | 667370.7461 | 3846787.631 | 4113088.127 |
| PI O-20:3_20:4 | 464145.238 | 353906.7589 | 299804.5093 | 268793.8917 | 500619.9658 | 647052.0145 | 444512.8767 | 433889.2954 | 433612.9221 | 524352.1907 | 449753.6132 | 593099.296 |
| 3-Oxopentanoic acid | 1257560.24 | 1282732.497 | 933712.3518 | 671381.3007 | 2897331.539 | 1495936.095 | 2821417.994 | 2752757.217 | 1004090.13 | 1288633.735 | 1375847.776 | 880136.1769 |
| SM 35:2;3O | 44751.69316 | 61602.44886 | 180571.7616 | 201593.7461 | 3003007.16 | 2466435.57 | 2381476.086 | 2978605.536 | 1125028.034 | 1001550.535 | 1169409.752 | 989950.0406 |
| Abiraterone acetate | 690669.0534 | 692804.0611 | 699543.3598 | 793139.709 | 813772.9026 | 983109.8423 | 834419.4993 | 869276.6928 | 681251.4041 | 661227.5924 | 1114502.662 | 896081.4218 |
| 17.beta.-Nandrolone decanoate | 0 | 0 | 0 | 0 | 1146982.795 | 1239482.93 | 996736.1265 | 1128560.114 | 345666.6632 | 489912.2589 | 483396.3314 | 988041.9253 |
| LPI 16:1 | 170074.7672 | 133724.9223 | 222237.6944 | 178153.7316 | 316713.164 | 197310.2782 | 344336.1205 | 266307.8942 | 183994.6798 | 211750.6371 | 181645.33 | 137931.0018 |
| Arg-Ala | 1107761.781 | 737469.1724 | 1455746.933 | 652748.7355 | 1774171.337 | 1273209.269 | 2065788.656 | 1423445.032 | 1764343.576 | 1837599.842 | 2166191.238 | 1375619.854 |
| N-(4-Fluorobenzyl)-7H-purin-6-amine | 240798.8938 | 566980.7427 | 878201.7017 | 597971.3078 | 165385.8293 | 142835.7551 | 191832.9971 | 142754.1976 | 191269.1123 | 196381.2523 | 171139.5244 | 2438642.437 |

| Metabolites | PBS_1 | PBS_2 | PBS_3 | PBS_4 | BCG_1 | BCG_2 | BCG_3 | BCG_4 | BCGΔ*pe18*_1 | BCGΔ*pe18*_2 | BCGΔ*pe18*_3 | BCGΔ*pe18*_4 |
|---|---|---|---|---|---|---|---|---|---|---|---|---|
| Linolenoylcarnitine (Car(18:3)) | 372973.9088 | 319850.4712 | 534781.0017 | 378573.6613 | 656664.9392 | 568925.7712 | 803706.0709 | 883111.1209 | 441498.6144 | 544331.6321 | 519476.7456 | 347107.0142 |
| Xanthohumol | 830830.7007 | 978542.3079 | 897239.0218 | 817745.0835 | 1625860.214 | 1433554.23 | 1325995.752 | 1333401.583 | 1391865.615 | 1164745.4 | 1494779.175 | 1818777.171 |
| 4-sulfophenyl acetate | 2720954.894 | 2770888.524 | 3259986.572 | 3292701.82 | 2232511.175 | 1367772.299 | 1750702.626 | 2541216.376 | 2957752.678 | 2393530.856 | 2570632.491 | 2712109.18 |
| PC O-22:2_16:2 | 173214.6034 | 158136.1972 | 204070.5906 | 203847.7995 | 741493.1427 | 1417606.336 | 1232344.772 | 1489052.79 | 487337.7971 | 426741.482 | 459901.7008 | 482313.1444 |
| LPC 14:1-SN2 | 417043.0293 | 518920.7467 | 802412.9825 | 538987.5354 | 756299.8579 | 797336.9142 | 1087559.999 | 780170.1002 | 612412.9818 | 657361.4349 | 633943.3398 | 581520.1757 |
| (2E,4E,10Z)-N-(2-Methylpropyl)hexadeca-2,4,10-trienamide | 4028993.401 | 4218593.437 | 1470015.328 | 2789646.957 | 2447056.063 | 2201196.947 | 2978465.702 | 2803689.925 | 3137599.398 | 3147659.091 | 3210811.638 | 2386088.041 |
| Pseudopelletierine | 7518984.944 | 9642593.83 | 8677408.902 | 9251131.38 | 9067909.031 | 9427278.348 | 9975388.369 | 7198635.196 | 10248533.22 | 9235612.511 | 9028415.564 | 9320147.272 |
| (+\|-)-Sulfinpyrazone | 70658.47464 | 41599.1517 | 132726.3531 | 55831.37955 | 2941157.481 | 1017546.339 | 2836965.609 | 1564039.681 | 1043407.319 | 1734334.812 | 1668408.012 | 772942.7213 |
| pomiferin | 0 | 0 | 0 | 0 | 526448.4292 | 467290.6218 | 306275.375 | 240294.3678 | 767903.1216 | 508715.0322 | 977822.2695 | 638691.9625 |
| Indoline | 31368772.62 | 18783718.85 | 42086502.4 | 26900291.72 | 36899463.08 | 40872350.67 | 30170419.46 | 31268395.69 | 25818561.42 | 32499633.39 | 28184275.58 | 30602566.84 |
| 3-Hydroxyacetophenone | 5009499.178 | 4556363.001 | 5363823.602 | 5104252.86 | 4426366.153 | 5289304.239 | 4599329.201 | 4056302.014 | 4577051.366 | 4923855.535 | 4570113.843 | 5078414.036 |

| Metabolites | PBS_1 | PBS_2 | PBS_3 | PBS_4 | BCG_1 | BCG_2 | BCG_3 | BCG_4 | BCGΔ*pe18*_1 | BCGΔ*pe18*_2 | BCGΔ*pe18*_3 | BCGΔ*pe18*_4 |
|---|---|---|---|---|---|---|---|---|---|---|---|---|
| 5-Oxooctanoic acid | 1657475.987 | 1520095.202 | 1508759.477 | 1541869.846 | 1348873.262 | 1535256.615 | 1377292.03 | 1436101.587 | 1428023.54 | 1632210.198 | 1385775.279 | 1665610.534 |
| LPE 20:2 | 557458.2171 | 639243.1281 | 648278.9295 | 548758.4137 | 1758918.915 | 1240431.629 | 1625751.687 | 1159886.85 | 1266020.495 | 1589009.953 | 1591307.723 | 1082171.784 |
| N5-acetyl-l-ornithine | 0 | 60418.03672 | 22761.64034 | 70766.93594 | 7070871.002 | 4804845.15 | 9290138.131 | 4356984.021 | 3767017.113 | 3397317.577 | 3545986.613 | 3443857.132 |
| 4-Hydroxyphenylmaraviroc | 137120.8843 | 127154.4067 | 213408.769 | 148768.6024 | 494742.7403 | 396274.2324 | 432619.0625 | 361144.2034 | 362762.3851 | 455201.7632 | 505242.4999 | 356625.2555 |
| Theanine | 362064.055 | 1181682.451 | 396037.9052 | 2038506.232 | 21710069.42 | 14545470.35 | 30174702.24 | 14814888.24 | 13059778.94 | 13192396.26 | 12278187.13 | 11861181.25 |
| PG O-17:2_22:4 | 1514012.238 | 1035093.526 | 983559.0278 | 1381906.204 | 3800834.431 | 2538130.57 | 2592033.744 | 2796303.939 | 1874075.548 | 2008736.557 | 3184272.738 | 2256833.201 |
| 2-(Pentylamino)acetamide | 289083.9811 | 1221491.377 | 1874159.992 | 1359323.421 | 419825.6649 | 254177.0845 | 479287.5102 | 327870.2585 | 530544.71 | 813281.9085 | 266716.4308 | 3652979.598 |
| 4-({[(4-Fluorophenyl)sulfonyl]amino}methyl)-N1-(3-pyridylmethyl)benzamide | 79339.57682 | 55360.12764 | 37383.46602 | 63426.21892 | 132066.0517 | 132820.2864 | 115791.4406 | 113027.5931 | 115745.4134 | 130929.788 | 191490.3431 | 152767.1145 |
| N-Benzyl-N-cyclohexylamine | 2076985.585 | 2162666.295 | 1796831.721 | 1850661.269 | 2134199.098 | 2032202.293 | 2081974.19 | 2149355.545 | 1969912.707 | 2023233.612 | 1965123.681 | 1764470.826 |
| 2-Methylglutaric acid | 6059716.555 | 5622443.16 | 7470447.804 | 7074297.591 | 15401839.23 | 11852446.76 | 13392913.3 | 13713585.69 | 9329621.416 | 7294356.507 | 10060825.76 | 9384663.594 |
| Tetracosanoic acid | 524269 | 394550 | 290614 | 391238 | 235318 | 365140 | 274325 | 395103 | 178148 | 208660 | 119236 | 169405 |

| Metabolites | PBS_1 | PBS_2 | PBS_3 | PBS_4 | BCG_1 | BCG_2 | BCG_3 | BCG_4 | BCGΔ *pe18*_1 | BCGΔ *pe18*_2 | BCGΔ *pe18*_3 | BCGΔ*pe18*_4 |
|---|---|---|---|---|---|---|---|---|---|---|---|---|
| | 3.567 | 0.638 | 6.68 | 3.568 | 6.316 | 4.937 | 8.755 | 4.492 | 9.331 | 3.649 | 9.459 | 8.84 |
| LPC 20:5 | 290953.8637 | 291791.2581 | 286533.1205 | 260798.2713 | 511579.2982 | 626948.5393 | 389800.4474 | 661955.2195 | 478784.5615 | 336816.4395 | 365078.4756 | 423122.6355 |
| 3-(2-Methyl-3-oxo-2,5,6,7,8,8a-hexahydro-1H-naphthalen-1-yl)propanoic acid | 16231380.5 | 14064257.69 | 12191473.21 | 13539107.18 | 14457814.97 | 15672702.08 | 16455697.21 | 14511374.56 | 14830756.96 | 13799031.71 | 13707983.43 | 12063090.98 |
| [3,5-Dihydroxy-2-[3-hydroxy-2-[[2-hydroxytetracosanoyl]amino]octadec-4-enoxy]-6-(hydroxymethyl)oxan-4-yl] hydrogen sulfate | 622571.383 | 667472.409 | 431013.6188 | 603726.3669 | 621473.7147 | 610381.9397 | 561868.8318 | 648922.5052 | 485677.8584 | 680425.1734 | 537854.1072 | 599498.7913 |
| 3-Nitro-N,N-di(propan-2-yl)benzamide | 1703982.446 | 1575861.571 | 3559791.653 | 2082869.529 | 1586940.636 | 1293425.387 | 1631584.773 | 1432371.184 | 721221.3528 | 786029.6387 | 989624.4697 | 661991.7482 |
| Pesticide3_Pirimicarb_C11H18N4O2_2-(Dimethylamino)-5,6-dimethyl-4-pyrimidinyl dimethylcarbamate | 4415632.286 | 6245147.742 | 8110635.243 | 7240551.866 | 4309343.685 | 3909654.136 | 4414377.546 | 3696494.898 | 7664653.298 | 4634413.671 | 4702238.575 | 10876006.69 |
| PA 6:0_22:3 | 7150078.017 | 8379617.18 | 5589783.203 | 7115110.064 | 8574558.906 | 12992136.37 | 8798331.138 | 8786214.243 | 6875259.042 | 9057810.005 | 8114671.485 | 6051652.805 |

| Metabolites | PBS_1 | PBS_2 | PBS_3 | PBS_4 | BCG_1 | BCG_2 | BCG_3 | BCG_4 | BCGΔ *pe18*_1 | BCGΔ *pe18*_2 | BCGΔ *pe18*_3 | BCGΔ*pe18*_4 |
|---|---|---|---|---|---|---|---|---|---|---|---|---|
| 6-(Difluoromethyl)-4-methyl-1H-pyrazolo[3,4-b]pyridin-3-ylamine | 1657453.598 | 1752738.518 | 1181760.208 | 1360190.553 | 1226591.206 | 1397349.022 | 1281195.854 | 13818477.04 | 1409329.515 | 1296767.351 | 1293457.283 | 937626.8923 |
| Allantoin | 28510186.69 | 23223421.24 | 40196944 | 29570073.21 | 55859968.19 | 43486940.87 | 61190893.93 | 49129469.15 | 40006355.39 | 34715994.24 | 38098423.12 | 39984684.72 |
| {[3-Ethoxy-2-(ethoxycarbonyl)-3-oxo-1-propenyl]amino}acetic acid | 2337960.085 | 1657067.416 | 3893925.679 | 2140412.692 | 2425928.815 | 12881188.76 | 4390683.456 | 2091549.718 | 828884.4229 | 17358999.57 | 1891422.136 | 1348009.778 |
| Saluamine | 2241177.412 | 2403729.368 | 2262533.268 | 2209741.219 | 943250.2605 | 2655639.664 | 27454646.31 | 862551.5184 | 2239491.196 | 1858074.718 | 28836933.2 | 2557948.182 |
| 6b-Hydroxy-2,2-dimethyl-14a-(2-methylbut-3-en-2-yl)-2,6b,7,7a,10,11,12,12a,14a,15-decahydro-8H,13H-pyrano[3,2-g]pyrrolo[1",2":4',5']pyrazino[1',2':1,5]pyrrolo[2,3-b]indole-8,13-dione | 124994.3447 | 129256.6104 | 110525.5091 | 129811.8131 | 139039.0968 | 152586.2433 | 149996.6151 | 134106.8878 | 147167.4006 | 129072.1814 | 141118.5783 | 168737.4985 |
| PC 9:0_34:6 | 731810.8766 | 575512.998 | 652295.977 | 970289.6016 | 569022.3436 | 376671.5248 | 408417.0807 | 715380.7496 | 503062.7779 | 775056.0138 | 718334.9816 | 1104907.657 |

| Metabolites | PBS_1 | PBS_2 | PBS_3 | PBS_4 | BCG_1 | BCG_2 | BCG_3 | BCG_4 | BCGΔ*pe18*_1 | BCGΔ*pe18*_2 | BCGΔ*pe18*_3 | BCGΔ*pe18*_4 |
|---|---|---|---|---|---|---|---|---|---|---|---|---|
| 2-(S-Glutathionyl)acetyl_chloride | 1309455.78 | 1083673.454 | 1161937.117 | 1235620.724 | 2321224.371 | 3321428.463 | 3371448.453 | 2638286.15 | 2294727.314 | 1806843.18 | 5424307.483 | 2847993.162 |
| Putrescine | 513893.156 | 1011331.811 | 1000843.107 | 937939.4433 | 3346828.854 | 2389238.01 | 5155346.65 | 3510312.292 | 2777601.397 | 3825591.688 | 3844852.144 | 2967293.187 |
| TG O-18:5_10:0_14:1 | 0 | 0 | 0 | 0 | 659050.7488 | 559061.7848 | 437731.7969 | 626322.3646 | 869377.4102 | 655828.7555 | 651087.5824 | 641787.6542 |
| 2(1H)-Pyrimidinone, 5-[3-[(1S,2S,4R)-bicyclo[2.2.1]hept-2-yloxy]-4-methoxyphenyl]tetrahydro- | 799650.5882 | 219662.8893 | 928329.125 | 302739.2348 | 1236965.075 | 1315701.231 | 4259061.733 | 3305143.61 | 1709468.187 | 1164313.01 | 1734312.801 | 684394.6362 |
| SM 17:0;2O | 1503097.688 | 1482579.541 | 845017.9398 | 1317939.568 | 1675470.679 | 1837107.946 | 1005926.793 | 1009693.221 | 1003683.486 | 1180970.808 | 1036145.61 | 552646.2154 |
| Leu-Ile | 654679.901 | 570747.0039 | 752705.4549 | 537415.9655 | 584405.7111 | 392442.771 | 544427.277 | 401921.2885 | 642002.5541 | 731994.4778 | 859688.6794 | 451306.4991 |
| PC O-15:1_22:5 | 3009051.989 | 3865724.023 | 2758051.322 | 2793234.49 | 2407913.977 | 141806.0241 | 3074163.524 | 1912130.78 | 4061272.984 | 3035619.227 | 3762417.165 | 1978628.677 |
| M547T525 | 531132.154 | 790186.4286 | 758216.71 | 594531.8869 | 1163060.707 | 628503.9408 | 739930.0367 | 664321.2783 | 847546.8382 | 757235.0392 | 723397.2245 | 920083.5832 |
| LNAPE 21:2/N-26:7 | 602144.4528 | 624409.9383 | 538097.3156 | 581184.292 | 562667.9123 | 812636.8165 | 566446.3507 | 506462.922 | 585398.8408 | 508115.9662 | 585049.362 | 614643.4282 |
| Naphthalene_epoxide | 1932615.945 | 1513591.722 | 2669232.759 | 1887100.183 | 3860694.163 | 2742255.014 | 4424172.719 | 3314568.807 | 2399714.879 | 3104625.535 | 3416727.004 | 2120210.13 |

| **Metabolites** | **PBS_1** | **PBS_2** | **PBS_3** | **PBS_4** | **BCG_1** | **BCG_2** | **BCG_3** | **BCG_4** | **BCGΔ*pe18*_1** | **BCGΔ*pe18*_2** | **BCGΔ*pe18*_3** | **BCGΔ*pe18*_4** |
|---|---|---|---|---|---|---|---|---|---|---|---|---|
| 16,16-Dimethyl-6-ketoprostaglandin E1 | 1862414.734 | 2558458.073 | 1647425.253 | 1927783.616 | 1953145.239 | 2152298.075 | 3071926.249 | 2286097.162 | 1756968.863 | 2194014.155 | 1854357.345 | 2150979.004 |
| (3E)-4-Hydroxy-3-[hydroxy-[2-[(1E,3E)-6-hydroxyhepta-1,3-dienyl]-1,3,6-trimethyl-4a,5,6,7,8,8a-hexahydro-2H-naphthalen-1-yl]methylidene]-5-(hydroxymethyl)-1-methylpyrrol-2-one | 263285.5751 | 275104.4303 | 279888.9315 | 289652.9844 | 197731.3047 | 274722.7894 | 253060.8553 | 96499.50851 | 216703.3832 | 35047.31642 | 270305.6963 | 202718.1285 |
| 17-Desacetylnorgestimate | 1623145.628 | 1767256.528 | 1380773.244 | 1463683.473 | 1613329.371 | 1401112.823 | 1664007.979 | 1454697.114 | 1647160.596 | 1495977.562 | 1602500.473 | 1499746.855 |
| 2,6-Bis(4-hydroxy-3-methoxybenzylidene)cyclohexanone | 2654416.643 | 2085894.484 | 3609126.413 | 2421675.522 | 2671245.914 | 1709703.24 | 2998279.337 | 2188416.675 | 1825817.942 | 2123478.51 | 2199856.719 | 1616671.758 |
| naringenin | 240965.6337 | 204655.8905 | 462900.1756 | 310452.3439 | 630523.8214 | 604509.8473 | 506578.0599 | 526938.3349 | 614657.285 | 655240.3217 | 531481.3708 | 635701.5802 |
| 18.beta.-Glycyrrhetic acid methyl ester | 340897.0243 | 340645.1703 | 339302.8875 | 363350.6153 | 356810.6687 | 377428.6171 | 387311.4396 | 370558.8163 | 332936.7445 | 348289.4985 | 377969.2076 | 364255.8371 |
| N-Methylbutane-1,4-diamine | 36240934.97 | 2476295.988 | 2355226.028 | 33767618.05 | 1510221.379 | 39225352.98 | 37484283.31 | 1206087.369 | 38323750.83 | 1986900.844 | 10560844.57 | 3005904.376 |
| N-acetyldopamine | 176406 | 153837 | 923881. | 210364 | 979858 | 103818 | 547901 | 877174 | 446950 | 604771 | 953837 | 751436 |

| Metabolites | PBS_1 | PBS_2 | PBS_3 | PBS_4 | BCG_1 | BCG_2 | BCG_3 | BCG_4 | BCGΔ*pe18*_1 | BCGΔ*pe18*_2 | BCGΔ*pe18*_3 | BCGΔ*pe18*_4 |
|---|---|---|---|---|---|---|---|---|---|---|---|---|
| | 3.31 | 7.633 | 6637 | 9.856 | 8.27 | 06.92 | 8.938 | 9.803 | 2.067 | 0.71 | 5.162 | 5.6 |
| BUTYLATED HYDROXYTOLUENE | 4964155.627 | 4605267.678 | 4566523.506 | 4736279.365 | 9039579.829 | 7754563.764 | 6819541.114 | 9183322.764 | 7807478.132 | 6233210.65 | 5728924.922 | 6682961.425 |
| Pyridoxine | 1064418.428 | 714481.4208 | 1198170.388 | 990059.2348 | 639470.2242 | 1191673.534 | 936459.4279 | 3686085.704 | 874248.6748 | 450418.5367 | 686485.1446 | 927372.2286 |
| SM 42:3;3O | 932099.6248 | 958189.335 | 1144025.283 | 1144161.083 | 619332.1058 | 1088203.863 | 699917.3514 | 760382.6591 | 914131.6493 | 695067.6945 | 77036.08958 | 1185055.682 |
| THYMIDINE-5'-DIPHOSPHO-ALPHA-GLUCOSE | 782217.0787 | 457176.9207 | 511957.6587 | 501655.6247 | 1452734.837 | 863097.8685 | 1104648.918 | 1200761.375 | 898004.273 | 1029380.022 | 1179937.287 | 979041.9784 |
| 2-octyl-3-hydroxydodecanoate | 1259211.157 | 1747194.883 | 1505025.779 | 1358452.77 | 1709556.133 | 1513835.001 | 1721580.121 | 1847283.661 | 923145.9712 | 1259763.353 | 1619985.453 | 677616.9678 |
| Dihexadecyldimethylammonium cation | 1943416.41 | 1964379.683 | 2133235.482 | 2150504.242 | 1974676.661 | 181238.897 | 5151692.101 | 17437980.04 | 2729297.615 | 2016241.87 | 1688007.849 | 24983925.86 |
| 2-Hydroxyoctanoic acid | 716486.2646 | 574860.1635 | 712320.1999 | 758454.5445 | 705392.0914 | 753633.2435 | 846331.5239 | 754515.1778 | 690746.3245 | 943313.4448 | 741314.9074 | 810168.7605 |
| 3,4-Dimethoxy-N-methylamphetamine | 505004.7331 | 827133.9416 | 1227087.865 | 3105880.617 | 1071064.974 | 761230.4126 | 5337995.47 | 2348447.712 | 1689310.939 | 1365913.104 | 1391739.393 | 3396533.252 |
| Methyl 3-benzoyloxy-8-(3-hydroxypropyl)-8-azabicyclo[3.2.1]oct | 3390796.359 | 3086231.146 | 3278316.943 | 3231681.619 | 3225706.73 | 3390207.042 | 3342075.913 | 3411900.422 | 3447023.282 | 3413980.609 | 3605105.751 | 3301350.742 |

| Metabolites | PBS_1 | PBS_2 | PBS_3 | PBS_4 | BCG_1 | BCG_2 | BCG_3 | BCG_4 | BCGΔ *pe18*_1 | BCGΔ *pe18*_2 | BCGΔ *pe18*_3 | BCGΔ*pe18*_4 |
|---|---|---|---|---|---|---|---|---|---|---|---|---|
| ane-2-carboxylate | | | | | | | | | | | | |
| Esmolol | 120385.2881 | 433565.3455 | 1401127.554 | 477215.985 | 90632.05415 | 104277.3765 | 104301.4979 | 69823.7401 | 50373.58182 | 183789.5061 | 240855.4475 | 2047507.778 |
| Phenylgalactoside | 1554839.503 | 2189496.996 | 2997664.636 | 2432807.16 | 1184092.976 | 1605506.314 | 2447947.876 | 1418410.492 | 1703425.199 | 1423995.683 | 1290121.771 | 5251153.414 |
| Vilazodone | 17742.10329 | 35798.03474 | 68136.77442 | 669572.5079 | 38837.74183 | 8700.916423 | 92434.49034 | 0 | 49627.43025 | 69173.98858 | 817532.1881 | 514376.7394 |
| Borrelidin | 1842223.615 | 1711736.691 | 1565360.092 | 1541313.472 | 1434008.269 | 1632176.922 | 1545415.557 | 1505857.178 | 1494640.473 | 1493320.093 | 1555758.042 | 1496304.521 |
| Norbuprenorphine | 140122.1281 | 131443.0029 | 580634.4191 | 116776.7468 | 415106.0762 | 399806.3619 | 556877.0074 | 425136.0944 | 448020.4837 | 544261.7479 | 572636.7521 | 325588.0201 |
| SL 11:1;O/14:1 | 1349706.987 | 1470416.769 | 1247871.725 | 1334849.702 | 1422446.146 | 1456460.165 | 1506344.669 | 1471970.474 | 1377537.379 | 1582765.007 | 1310860.584 | 1262461.071 |
| PS(16:0/16:0) | 14141610.86 | 14294515.26 | 12396646.23 | 13782688.04 | 14254036.31 | 32853006.48 | 14698662.46 | 14451402.67 | 22072094.73 | 14163233.88 | 14129011.26 | 11152796.48 |
| LNAPE 20:4/N-22:4 | 468568.4816 | 193086.4117 | 451968.0575 | 350507.2741 | 625497.0557 | 64702.38031 | 578726.3936 | 549663.4877 | 637586.8854 | 599221.0279 | 457305.531 | 747187.0542 |
| DGTS 2:0_17:0 | 4746590.231 | 6242780.309 | 4730010.192 | 4968771.137 | 4991856.8 | 4849495.41 | 4466606.384 | 4899319.397 | 5058979.008 | 4932503.775 | 4586698.704 | 6549384.946 |
| 2,4-Dodecadienoic acid isobutylamide | 1882527.961 | 2931562.901 | 1649780.541 | 2022431.871 | 2353916.415 | 1963823.545 | 3501386.343 | 2388951.493 | 2029094.608 | 2514871.793 | 4456474.781 | 2071373.124 |

| **Metabolites** | **PBS_1** | **PBS_2** | **PBS_3** | **PBS_4** | **BCG_1** | **BCG_2** | **BCG_3** | **BCG_4** | **BCGΔ*pe18*_1** | **BCGΔ*pe18*_2** | **BCGΔ*pe18*_3** | **BCGΔ*pe18*_4** |
|---|---|---|---|---|---|---|---|---|---|---|---|---|
| Polyporusterone_F | 3125534.239 | 4569710.624 | 2590981.677 | 4137768.351 | 2923480.863 | 3122042.632 | 3669978.399 | 3757156.943 | 2977103.653 | 3043782.081 | 3040883.14 | 2919822.386 |
| Methylergonovine | 1738917.835 | 1072977.851 | 1530174.788 | 1084378.452 | 225136.4695 | 128011.6075 | 304020.9317 | 206096.4723 | 160791.1796 | 183105.6529 | 223468.0572 | 128339.9363 |
| PC O-40:4 | 2046398.489 | 951024.4021 | 2040602.898 | 3234816.268 | 1216181.764 | 0 | 1448878.687 | 2410308.323 | 2212233.725 | 1706753.633 | 1017848.796 | 2396498.107 |
| 12-Hydroxy-12-octadecanoylcarnitine | 315281.2992 | 208769.7372 | 310953.133 | 211958.0012 | 543795.4221 | 361128.9975 | 431816.8526 | 327197.8739 | 382843.5654 | 0 | 512103.2667 | 205352.8185 |
| N-(5-Amino-1H-1,2,4-triazol-3-yl)acetamide | 195803.3787 | 3238599.973 | 3680924.569 | 4121689.173 | 5467821.082 | 248309.7719 | 5655688.111 | 1222603.975 | 533477.9852 | 347950.9749 | 5797690.302 | 3492957.912 |
| 1-Methylhexahydroazepin-4-one | 398076.448 | 1227208.103 | 544481.3718 | 478823.445 | 1401575.277 | 629814.9326 | 1085805.68 | 722269.4687 | 549557.1498 | 535385.5055 | 759808.7643 | 807524.0044 |
| Populoside | 3701155.255 | 1809850.576 | 1822760.473 | 1346166.569 | 748472.9117 | 949747.2933 | 702347.8538 | 578698.0812 | 729856.4307 | 1153824.588 | 936310.2608 | 665686.2337 |
| PC(20:1(11Z)/15:0) | 10093149.79 | 10117690.25 | 7298839.659 | 7838396.756 | 23616561.94 | 18107052.71 | 65856032.81 | 54175340.12 | 24643842.61 | 17066381.92 | 41761877.45 | 10922161.66 |
| CE 22:6 | 4859536.616 | 6320662.422 | 4757422.029 | 5612414.026 | 4542220.519 | 17806.07255 | 3760178.518 | 5272752.504 | 4487228.971 | 4981523.506 | 4081551.292 | 3559614.461 |
| 2-Pyridinemethanamine, N-[[4-(1,4,8,11-tetraazacyclotetradec-1-ylmethyl)phenyl]m | 988653.9731 | 1046346.38 | 1052191.761 | 1153684.509 | 1094003.648 | 975190.2031 | 1092171.257 | 1112042.314 | 937597.1489 | 1135220.902 | 977167.9118 | 922857.9912 |

| Metabolites | PBS_1 | PBS_2 | PBS_3 | PBS_4 | BCG_1 | BCG_2 | BCG_3 | BCG_4 | BCGΔ*pe18*_1 | BCGΔ*pe18*_2 | BCGΔ*pe18*_3 | BCGΔ*pe18*_4 |
|---|---|---|---|---|---|---|---|---|---|---|---|---|
| ethyl]- | | | | | | | | | | | | |
| PI 38:4 | 1111490.693 | 1663867.061 | 1599635.471 | 1645022.318 | 1082486.05 | 1396455.141 | 1179794.533 | 1403669.326 | 1429191.103 | 1289812.149 | 1221503.295 | 1341868.586 |
| Galactose | 1065467.306 | 605434.2083 | 1070021.434 | 1167649.793 | 2762647.26 | 2837624.473 | 2481604.312 | 2961109.744 | 3471254.831 | 3005389.489 | 3107806.41 | 2844716.777 |
| PI 20:1_20:4 | 3308894.134 | 1456956.452 | 1793790.666 | 1978413.88 | 1195273.171 | 0 | 1852563.554 | 1767586.849 | 2430287.502 | 1138774.416 | 1380434.978 | 946860.8316 |
| O,O-Dimethyl dithiophosphate | 16425764.12 | 32927767.28 | 32235572.74 | 32584085.54 | 33115468.25 | 32537907.06 | 33945220.52 | 34036195.91 | 33334798.67 | 33943022.38 | 34131103.62 | 31348898.18 |
| Gln-Arg | 864911.5656 | 145260.8885 | 815478.7188 | 281890.1499 | 912356.1124 | 1293966.15 | 7149132.15 | 3645812.393 | 2075854.832 | 1141994.249 | 1746936.17 | 696760.1414 |
| 3-Amino-4-methylpentanoic acid | 22959012.55 | 18683299.48 | 29326695.59 | 24659164.46 | 47678927.79 | 42141703.57 | 46535823.37 | 42751333.82 | 46186025.61 | 27749270.63 | 32662727.31 | 39229284.43 |
| 7(14)-Bisabolene-2,3,10,11-tetrol | 1573160.82 | 1550200.417 | 1458356.753 | 1292535.189 | 1059055.593 | 1401300.576 | 1222169.022 | 1236776.384 | 1301115.22 | 1221240.367 | 1216159.075 | 1094640.122 |
| 4-(3,4'-Dihydroxy-4,4,7,8a-tetramethyl-6'-oxospiro[2,3,4a,5,6,7-hexahydro-1H-naphthalene-8,2'-3,8-dihydrofuro[2,3-e | 117643.233 | 161984.8056 | 339785.9114 | 202956.3509 | 961304.6774 | 911388.8408 | 698169.6912 | 827523.6143 | 627663.5802 | 527012.293 | 678245.8487 | 703632.8802 |

| **Metabolites** | **PBS_1** | **PBS_2** | **PBS_3** | **PBS_4** | **BCG_1** | **BCG_2** | **BCG_3** | **BCG_4** | **BCGΔ *pe18*_1** | **BCGΔ *pe18*_2** | **BCGΔ *pe18*_3** | **BCGΔ*pe18*_4** |
|---|---|---|---|---|---|---|---|---|---|---|---|---|
| ]isoindole]-7'-yl)butanoic acid | | | | | | | | | | | | |
| SM 24:0;2O/18:1 | 9107016.578 | 28119428.38 | 20156919.45 | 28522486.19 | 9440798.868 | 37501833.6 | 6688146.365 | 2567775.314 | 7177893.91 | 7127379.744 | 6710389.563 | 9688971.685 |
| HexCer 14:0;2O/16:4 | 1583692.998 | 1629502.576 | 1536891.223 | 1605229.315 | 1666081.947 | 1609113.627 | 1581403.265 | 1553172.268 | 1593371.395 | 1461510.703 | 1630001.728 | 1603979.897 |
| 3-Methoxy-4-hydroxyphenylglycol sulfate | 2592278.318 | 1897805.441 | 4967812.383 | 3108876.616 | 3597732.547 | 3120940.943 | 4513569.956 | 3093452.142 | 2917787.368 | 2004983.713 | 2109535.376 | 2740040.626 |
| Terpestacin | 1506695.122 | 1651614.04 | 4108667.614 | 2628782.855 | 2850184.438 | 4208794.922 | 2651380.69 | 3318777.122 | 2146689.852 | 1244508.013 | 2180951.123 | 2549063.191 |
| Acetoacetic acid | 933262.8206 | 938641.8347 | 920354.1953 | 1018688.882 | 1633855.137 | 1189837.154 | 1483809.755 | 1273350.11 | 1322142.91 | 1281513.005 | 1323707.789 | 1168435.283 |
| 13,14-Dihydro-16,16-difluoroprostaglandin F2.alpha. | 2210120.922 | 2384455.495 | 1380299.436 | 2415724.663 | 6153005.026 | 3789864.801 | 3595411.756 | 2916946.625 | 3020194.336 | 2956104.804 | 3544374.905 | 2772414.473 |
| 5-Tetradecynoic acid | 586611.2989 | 522435.1078 | 1330852.77 | 684915.4488 | 563566.7164 | 630934.3105 | 560891.9601 | 626155.1503 | 519057.5019 | 528266.5234 | 577393.9439 | 907158.397 |
| Cer 8:0;2O/14:0 | 2768652.506 | 2660825.649 | 2297710.584 | 2614552.669 | 2244433.658 | 2453998.152 | 2327057.206 | 2516058.611 | 2404827.507 | 2521368.175 | 2440048.743 | 2430727.117 |

| Metabolites | PBS_1 | PBS_2 | PBS_3 | PBS_4 | BCG_1 | BCG_2 | BCG_3 | BCG_4 | BCGΔ*pe18*_1 | BCGΔ*pe18*_2 | BCGΔ*pe18*_3 | BCGΔ*pe18*_4 |
|---|---|---|---|---|---|---|---|---|---|---|---|---|
| SM 22:1;2O/20:1 | 0 | 646736.0534 | 482765.0413 | 758444.7093 | 516134.6844 | 406572.6474 | 785659.3715 | 578321.0077 | 458721.6547 | 789389.9846 | 536275.0053 | 613919.182 |
| Blepharin | 7404271.996 | 2575107.951 | 9056606.301 | 4465303.441 | 860707.443 | 2490344.007 | 1673566.804 | 2978390.959 | 1624484.366 | 790566.0009 | 1570752.612 | 2254636.846 |
| SMGDG O-16:2_4:0 | 749304.6681 | 555194.2452 | 674193.1304 | 682017.4199 | 495807.7506 | 317648.178 | 381589.9734 | 309966.3411 | 298734.2885 | 343802.9416 | 395473.3974 | 521316.6319 |
| Quercetin-3,7,3',4'-tetramethyl ether | 1379447.741 | 1022064.264 | 1495896.515 | 1481811.009 | 1462916.404 | 1836327.13 | 1155120.706 | 1439055.468 | 1444188.271 | 504958.4832 | 593013.4869 | 1557652.73 |
| 1,2,4-Triazin-5-amine, N-([1,1'-biphenyl]-4-ylmethyl)-6-phenyl-3-(2-pyridinyl)- | 216988.9178 | 30001.67773 | 281520.4876 | 60401.53539 | 336077.0264 | 439301.8059 | 1637101.189 | 1295157.957 | 572199.9338 | 345895.6711 | 568084.1036 | 203565.3573 |
| M286T103 | 5481163.716 | 3451324.161 | 6080849.916 | 3281773.534 | 1015107.762 | 1260666.721 | 2158797.322 | 2060592.323 | 711967.9945 | 1100374.329 | 915723.0034 | 622860.0438 |
| 1-heptadecanoyl-glycero-3-phosphate | 617037.5318 | 694935.5913 | 1181870.59 | 2086754.51 | 9472787.951 | 8223968.742 | 9289062.564 | 8316764.648 | 8597025.699 | 6673353.727 | 10173262.9 | 6142281.876 |
| PG O-19:2_18:2 | 624729.7066 | 382645.4299 | 602948.6456 | 368699.1338 | 614428.4002 | 14495.16268 | 779162.7192 | 1127507.341 | 572808.149 | 644713.4032 | 494658.8612 | 1200928.967 |
| 2-Docosahexaenoyl-1-stearoyl-sn-glycero-3-phosphoserine | 7512587.718 | 3542345.083 | 5767019.84 | 5847851.277 | 6296317.194 | 727417.5827 | 4610384.193 | 5063579.369 | 4355883.584 | 3737245.345 | 2463473.215 | 3234344.044 |
| LPC(32:0) | 92690041.86 | 45991840.47 | 42015812.81 | 112407010.7 | 30165882.28 | 13905688.41 | 51974739.84 | 22980191.8 | 20562150.01 | 35939769.73 | 35042240.1 | 26388047.1 |

| **Metabolites** | **PBS_1** | **PBS_2** | **PBS_3** | **PBS_4** | **BCG_1** | **BCG_2** | **BCG_3** | **BCG_4** | **BCGΔ*pe18*_1** | **BCGΔ*pe18*_2** | **BCGΔ*pe18*_3** | **BCGΔ*pe18*_4** |
|---|---|---|---|---|---|---|---|---|---|---|---|---|
| 3-Cyclohexyl-2-hydroxypropanoic acid | 4232548.973 | 3346342.953 | 3927455.993 | 4594452.221 | 3138969.403 | 4327045.267 | 4183407.749 | 3652006.022 | 3642703.272 | 6064072.31 | 3533811.944 | 4559909.876 |
| 3'-O-Methylinosine | 1824510.017 | 1550000.131 | 3015458.776 | 2193639.599 | 21818446.13 | 12755697.06 | 19929345.93 | 15267571.9 | 17954209.92 | 17495190.41 | 20816321.64 | 15520543.8 |
| LPS 14:0 | 321334.0269 | 287469.4345 | 316185.6766 | 327080.0548 | 862948.0031 | 398424.2413 | 538223.8704 | 398769.3868 | 454249.2485 | 465963.7585 | 487949.0112 | 470836.049 |
| Propyl gallate | 635639.6668 | 492132.3423 | 1851083.804 | 2357651.629 | 1585789.95 | 672498.7431 | 295742.1036 | 421265.157 | 490923.1323 | 300618.1842 | 485084.8048 | 1053401.482 |
| Cer 15:2;3O/14:1 | 1594731.248 | 297735.7244 | 835470.1491 | 1386095.069 | 728849.7801 | 862241.4493 | 7739532.596 | 926112.7464 | 951931.9006 | 1037979.67 | 786954.827 | 1383222.069 |
| Myrtucommulone B | 394490.5736 | 549791.4448 | 380204.4145 | 577638.0509 | 549675.1098 | 559198.7607 | 729568.6476 | 630641.6536 | 483334.5656 | 624642.9189 | 441694.2526 | 449956.3389 |
| PG 18:1_18:1;3O | 514315.6758 | 352899.1371 | 478219.3196 | 530020.2971 | 351263.1834 | 338094.009 | 501355.5977 | 368286.3068 | 397268.8025 | 394732.4286 | 335101.1435 | 365838.1461 |
| 16-Phenoxytetranor prostaglandin F2.alpha. methylamide | 2779572.631 | 2937450.005 | 2534940.34 | 2752024.903 | 2841451.18 | 2833981.07 | 2892589.987 | 2813708.044 | 2972948.977 | 2697183.005 | 2898886.725 | 2573025.552 |
| 2-Ethyl-2-hydroxybutyric acid | 2607442.12 | 2516791.118 | 2499580.208 | 2657883.819 | 2681318.29 | 2444323.768 | 2591915.114 | 2601494.849 | 2521355.901 | 2602310.8 | 2572239.424 | 2576690.995 |
| L-Homoserine lactone | 1030686.061 | 1202535.347 | 1394712.299 | 1122633.804 | 2385471.463 | 1749075.465 | 2233677.233 | 1931440.475 | 1842118.035 | 1572941.61 | 1956324.929 | 1644877.683 |
| ADGGA 16:3_14:0_18:5 | 0 | 0 | 0 | 0 | 117152.4351 | 159029.8764 | 102395.4912 | 112238.1953 | 249000.6377 | 175393.9032 | 201406.8801 | 191101.2629 |

| **Metabolites** | **PBS_1** | **PBS_2** | **PBS_3** | **PBS_4** | **BCG_1** | **BCG_2** | **BCG_3** | **BCG_4** | **BCGΔ*pe18*_1** | **BCGΔ*pe18*_2** | **BCGΔ*pe18*_3** | **BCGΔ*pe18*_4** |
|---|---|---|---|---|---|---|---|---|---|---|---|---|
| SM 9:0;2O/32:8 | 135237.8395 | 61712.37143 | 32452.53446 | 217131.3272 | 1388709.817 | 1704953.719 | 1370348.436 | 2042648.109 | 458843.9385 | 471587.0097 | 811347.6141 | 705726.3034 |
| Nordeprenyl | 1001978.264 | 1887981.859 | 864667.2687 | 993877.1853 | 795901.0826 | 1001814.784 | 941688.6199 | 947778.8161 | 865634.8941 | 1003487.245 | 982532.0366 | 1087698.927 |
| Resolvin D2 | 118847.1707 | 118514.778 | 132270.6696 | 104343.0697 | 545378.1874 | 1260064.431 | 738556.6908 | 761371.4578 | 571560.4622 | 376410.6272 | 805731.4616 | 654706.3674 |
| PC 18:1_20:3 | 377519.3737 | 745020.6082 | 222036.1136 | 409904.3192 | 178961.3744 | 188705.1761 | 291131.1857 | 202818.6928 | 336616.6434 | 591862.8126 | 272202.0471 | 365967.7081 |
| 3-Hydroxytrideca-4,6-dienoylcarnitine | 1096823.332 | 1279039.423 | 860113.3328 | 990461.3532 | 1150439.922 | 1154553.6 | 1508634.289 | 2212448.39 | 1302805.842 | 1229909.672 | 1165482.54 | 916976.7796 |
| 4-(4-tert-Butylphenyl)-1,3-thiazol-2-ylamine | 768929.3178 | 994785.7354 | 747457.4912 | 980433.9795 | 882092.6435 | 808816.4711 | 1133614.51 | 758466.2881 | 545363.0258 | 886508.5014 | 985534.0622 | 709327.5443 |
| SHexCer 23:2;3O | 0 | 5030.334234 | 0 | 0 | 156148.658 | 178182.5086 | 117994.6407 | 110768.1049 | 273631.4289 | 228419.9435 | 279995.3022 | 261065.5983 |
| Indole-3-carboxaldehyde | 593462.3806 | 664647.1763 | 618594.3464 | 811777.6198 | 814019.2171 | 698534.3739 | 714468.0983 | 670900.9009 | 673356.8779 | 600546.7604 | 826962.0827 | 707078.6883 |
| ent-15-Hydroxy-8(17)-labden-2-one | 1061831.572 | 1150382.168 | 961986.2318 | 1007007.404 | 1147065.77 | 1072694.706 | 1159361.442 | 1135631.609 | 1111702.504 | 1154681.219 | 1048465.436 | 1063100.653 |
| 4H-1,2,4-Triazole-3-thiol, 4-phenyl-5-(4-pyridyl)- | 2170862.265 | 2197107.644 | 2485016.179 | 2102418.832 | 2817759.09 | 2910426.971 | 2718297.952 | 2737439.567 | 2578961.364 | 2652895.445 | 2516360.491 | 2407631.379 |
| MGDG | 101577 | 126692 | 803315. | 116967 | 962728. | 191486 | 101457 | 930875. | 816526. | 993639. | 102015 | 985310. |

| Metabolites | PBS_1 | PBS_2 | PBS_3 | PBS_4 | BCG_1 | BCG_2 | BCG_3 | BCG_4 | BCGΔ *pe18*_1 | BCGΔ *pe18*_2 | BCGΔ *pe18*_3 | BCGΔ*pe18*_4 |
|---|---|---|---|---|---|---|---|---|---|---|---|---|
| O-18:3_22:6 | 9.934 | 7.646 | 8616 | 1.42 | 8036 | 2.139 | 0.176 | 6825 | 2743 | 7412 | 1.182 | 499 |
| 7.alpha.-Hydroxy-4-cholesten-3-one | 351424.77 | 241320.3253 | 380995.243 | 421594.4284 | 21736072.31 | 19007856.06 | 22108937.29 | 18123684.83 | 5781166.587 | 7267087.927 | 10465833.33 | 4587140.085 |
| Glutathione | 6721009.93 | 2636884.079 | 5215635.921 | 2590566.347 | 1450900.12 | 2043074.788 | 14816308.8 | 15804592.28 | 2962554.117 | 1794257.487 | 5078857.956 | 2378524.421 |
| 4-(2,3-Dihydroimidazo[2,1-b][1,3]thiazol-6-yl)phenol | 748554.2934 | 737678.7758 | 1219582.487 | 2399099.875 | 557485.2348 | 909898.9514 | 1077164.261 | 3234891.237 | 1143357.055 | 521796.9821 | 483793.4707 | 1110609.12 |
| 3-Hydroxy-2-[(Z)-oct-2-enyl]pentanedioic acid | 19856290.04 | 12712310.54 | 15676764.05 | 14271085.25 | 19827328.95 | 14548444.63 | 16171916.95 | 9681156.703 | 10963288.83 | 16566205.38 | 14752572.21 | 14583689.84 |
| (2-aminoethoxy)[2-[docosa-4.7.10.13.16.19-hexaenoyloxy]-3-[hexadec-1-en-1-yloxy]propoxy]phosphinic acid | 646835.3774 | 681720.2338 | 513495.7702 | 742945.986 | 1403244.721 | 1041172.946 | 1186024.588 | 1165486.303 | 1380274.027 | 1445068.31 | 1161384.116 | 916039.0114 |
| Cer 8:0;3O/12:0;(2OH) | 4056248.173 | 3890332.157 | 3776290.79 | 3804207.45 | 4036562.725 | 3652962.977 | 4373902.681 | 4171810.692 | 3705237.434 | 3961076.723 | 3914272.7 | 3802041.83 |
| Octamethylcyclotetrasiloxane | 453380.8588 | 248699.7523 | 476761.2995 | 298519.597 | 435743.4528 | 853991.5959 | 570670.4174 | 625150.4715 | 391620.4247 | 356581.7916 | 620489.0198 | 647262.5792 |
| MGDG O-16:3_18:0 | 345458.9572 | 339927.0862 | 340719.3646 | 358305.5667 | 196055.1094 | 164616.944 | 171877.7704 | 231789.8688 | 216875.4717 | 321665.1651 | 168748.4569 | 84725.63815 |
| Jasmine lactone | 653140. | 827340. | 554306. | 562129. | 612581. | 569999. | 512924. | 412929. | 575461. | 558305. | 607584. | 626600. |

| **Metabolites** | **PBS_1** | **PBS_2** | **PBS_3** | **PBS_4** | **BCG_1** | **BCG_2** | **BCG_3** | **BCG_4** | **BCGΔ*pe18*_1** | **BCGΔ*pe18*_2** | **BCGΔ*pe18*_3** | **BCGΔ*pe18*_4** |
|---|---|---|---|---|---|---|---|---|---|---|---|---|
| | 8265 | 4574 | 7033 | 6186 | 8164 | 5713 | 1018 | 5737 | 2696 | 4517 | 15 | 3843 |
| Diltiazem | 207744.3983 | 237671.3435 | 203999.6485 | 219280.8369 | 260650.0477 | 223406.5133 | 230892.1667 | 232111.7465 | 221516.841 | 233429.2629 | 240837.3249 | 243970.4651 |
| DGDG O-16:3_3:0 | 70760.1201 | 42139.34298 | 46844.39538 | 56512.46612 | 67046.07195 | 61086.31058 | 40329.04557 | 61691.38813 | 60837.99606 | 67278.10773 | 65599.38335 | 27532.03762 |
| PC 8:0_21:0 | 2214277.628 | 2194734.744 | 1837278.696 | 1267552.222 | 12284562.2 | 7383099.288 | 8367339.115 | 9546491.598 | 10753272.6 | 7852445.819 | 7686916.857 | 5469909.782 |
| LPG 20:2 | 992011.0823 | 1172769.265 | 2782452.195 | 1200370.392 | 2910508.812 | 3381118.524 | 4055919.263 | 2671083.697 | 2535927.703 | 2540671.327 | 2925919.307 | 987309.6813 |
| Taurodeoxycholic acid | 3455937.586 | 3194197.459 | 3754366.847 | 3263979.532 | 5894826.896 | 5201696.435 | 6431847.391 | 3372338.399 | 6596763.581 | 4795432.523 | 2891005.96 | 2630891.293 |
| Goshonoside_F1 | 5106878.887 | 4867510.005 | 4207621.629 | 4804668.062 | 5560321.501 | 9587413.235 | 7753289.184 | 5524689.177 | 8086814.564 | 5524047.756 | 5075467.233 | 3989655.941 |
| 2,5-Dihydroxyacetophenone | 915165.9202 | 740263.8129 | 782693.1935 | 932905.0078 | 944553.2178 | 785908.9465 | 923158.334 | 943421.5404 | 854376.671 | 933475.4828 | 783359.2076 | 832788.861 |
| 1-(4-Piperidinyl)-1,3-dihydro-2H-indol-2-one | 18380.46963 | 345050.635 | 553052.8591 | 435583.61 | 13416.73029 | 437642.4115 | 274017.61 | 274480.229 | 51332.87296 | 36936.01872 | 33888.45366 | 2002269.373 |
| 4'-Methoxy-6-methylflavone | 323639.5915 | 329481.8807 | 249414.7757 | 1511829.215 | 236554.9723 | 337533.401 | 2048766.307 | 204124.7679 | 902822.1214 | 535380.1664 | 224588.0373 | 1663237.828 |
| 5-Amino-1-(2-pyridinyl)-1H-pyrazole-4-carbonitrile | 1470724.193 | 1324701.654 | 1632129.144 | 1307043.392 | 4991373.779 | 2973613.608 | 5087067.204 | 3891930.053 | 3609540.224 | 4009155.662 | 3761899.661 | 3527416.029 |

| **Metabolites** | **PBS_1** | **PBS_2** | **PBS_3** | **PBS_4** | **BCG_1** | **BCG_2** | **BCG_3** | **BCG_4** | **BCGΔ*pe18*_1** | **BCGΔ*pe18*_2** | **BCGΔ*pe18*_3** | **BCGΔ*pe18*_4** |
|---|---|---|---|---|---|---|---|---|---|---|---|---|
| 4,5-Diphenyl-2-methylimidazole | 1462683.925 | 1938264.222 | 1573004.799 | 1586845.913 | 1664104.188 | 1785848.453 | 1502619.476 | 1871643.882 | 1767174.489 | 1789733.391 | 1646143.349 | 1674141.896 |
| (2-aminoethoxy)[2-[docosa-4.7.10.13.16.19-hexaenoyloxy]-3-[octadeca-1.9-dien-1-yloxy]propoxy]phosphinic acid | 10024836.62 | 11257262.33 | 9203626.105 | 7441498.134 | 15238843.71 | 10710766.62 | 14553773.31 | 14316938.63 | 15797377.22 | 2213985.296 | 7116921.834 | 8089589.75 |
| 6-Aminonicotinic acid | 6000016.02 | 1745411.803 | 560245.0746 | 2286759.666 | 714255.1935 | 1031201.394 | 823393.255 | 1717251.059 | 765025.1714 | 809863.0451 | 878894.602 | 2131247.448 |
| NAE 20:4 | 2075862.498 | 2390652.525 | 2982153.082 | 2297774.994 | 2365502.404 | 2826939.68 | 2969259.301 | 4710390.883 | 1714253.815 | 2304368.93 | 2573044.247 | 1814350.354 |
| N-Acetylmuramic acid | 5222.43041 | 13741.37532 | 13981.42178 | 0 | 1505039.982 | 1151175.398 | 1056676.288 | 983154.25 | 1148429.471 | 1130356.139 | 1328098.173 | 1097260.902 |
| 13[(2R,3S)-3-Pentyloxiranyl]-8Z-tridecenoic acid | 253900.8458 | 241981.0913 | 324621.6198 | 270976.9984 | 764018.3073 | 522256.4152 | 953377.0204 | 607910.4937 | 405500.3563 | 500740.7211 | 653513.9655 | 356349.6481 |
| Imidocarb | 1274316.291 | 1167319.636 | 1273052.006 | 1137702.831 | 1106840.628 | 5740099.782 | 1083811.411 | 1226159.966 | 1195846.922 | 1297897.042 | 1088619.207 | 976106.9468 |
| Oleic acid 2,6-diisopropylanilide | 50943.16397 | 63311.36819 | 101294.1296 | 58091.31468 | 0 | 0 | 0 | 22319.42459 | 14158.95201 | 18137.68519 | 11705.93236 | 1885799.68 |
| FAHFA 16:0/2:0 | 637617.3486 | 767289.2389 | 507565.4237 | 696089.035 | 751795.4988 | 657646.2689 | 536399.3801 | 770356.0213 | 630020.091 | 700748.2768 | 521524.3556 | 2126672.142 |

| Metabolites | PBS_1 | PBS_2 | PBS_3 | PBS_4 | BCG_1 | BCG_2 | BCG_3 | BCG_4 | BCGΔ*pe18*_1 | BCGΔ*pe18*_2 | BCGΔ*pe18*_3 | BCGΔ*pe18*_4 |
|---|---|---|---|---|---|---|---|---|---|---|---|---|
| Chloroxylenol | 322882.6166 | 429873.624 | 308206.883 | 311962.6973 | 692294.9129 | 366664.8179 | 329419.6584 | 364898.2423 | 347174.0473 | 624329.025 | 622453.8548 | 433099.636 |
| 6-{[4-(1,3-Dioxo-1,3-dihydro-2H-isoindol-2-yl)anilino]carbonyl}-3,4-dimethyl-3-cyclohexene-1-carboxylic acid | 391864.4331 | 535322.488 | 195781.8074 | 254351.2526 | 258827.7809 | 899122.0726 | 413986.4427 | 286603.9904 | 350433.745 | 292621.1392 | 143190.3508 | 600852.4782 |
| SM 8:1;2O/24:6 | 2920990.364 | 3450014.515 | 3228643.57 | 3406533.564 | 3813093.426 | 3629237.456 | 2526958.36 | 3798313.59 | 2909203.166 | 2716749.911 | 3297026.774 | 3768546.26 |
| 3-(5-Chloro-1,3-benzothiazol-2-yl)propanoic acid | 987458.72 | 861458.3212 | 1002085.783 | 1004608.047 | 1323679.492 | 936801.1746 | 21630.4911 | 1237690.009 | 740843.7331 | 738772.1185 | 943410.5285 | 681678.9239 |
| Ala-Ile | 904726.4813 | 1415941.56 | 2303770.738 | 2727634.821 | 2805734.722 | 1371529.742 | 2937282.749 | 1872087.467 | 1824677.55 | 2593365.514 | 2259731.45 | 3139003.902 |
| PC(42:4) | 4373461.502 | 4870580.627 | 2738741.757 | 4691916.991 | 4411737.841 | 3537807.872 | 4674219.507 | 4243863.429 | 6462442.266 | 5820090.322 | 4300034.938 | 2517722.412 |
| 2,7-Diamino-9H-xanthen-9-one | 4906047.601 | 2850550.709 | 4375986.482 | 5159455.75 | 5885903.911 | 5204280.826 | 9094514.201 | 6254697.633 | 6162119.642 | 5620234.028 | 4290414.671 | 6159364.176 |
| (24E)-3-Oxodammara-1,12,24-trien-26-oic acid | 621495.3542 | 2262484.894 | 268455.5328 | 1670887.877 | 1458429.627 | 384054.784 | 992955.7187 | 855342.2138 | 505119.6403 | 957131.9506 | 1166376.432 | 570926.0422 |
| DGTS 15:0_22:6 | 224865.1729 | 140130.7161 | 88638.24722 | 187654.3078 | 365845.0024 | 296869.5456 | 202559.33 | 378097.8116 | 288284.3256 | 209163.749 | 126403.1628 | 242279.4649 |

| **Metabolites** | **PBS_1** | **PBS_2** | **PBS_3** | **PBS_4** | **BCG_1** | **BCG_2** | **BCG_3** | **BCG_4** | **BCGΔ*pe18*_1** | **BCGΔ*pe18*_2** | **BCGΔ*pe18*_3** | **BCGΔ*pe18*_4** |
|---|---|---|---|---|---|---|---|---|---|---|---|---|
| LNAPE 18:0/N-2:0 | 696423.0626 | 1973582.546 | 268897.8946 | 1865215.062 | 1654225.791 | 971124.2317 | 805721.8524 | 525828.8222 | 795130.1765 | 675969.014 | 1013983.006 | 868678.3376 |
| LDGCC 16:4 | 1753608.648 | 2611552.561 | 1101781.239 | 1370218.027 | 1121418.374 | 2245422.998 | 1615405.708 | 1344940.039 | 1144217.354 | 1578499.813 | 1298619.466 | 924610.4 |
| (2-Oxo-1-pyrrolidinyl)acetic acid | 6658327.938 | 4264018.591 | 4963166.354 | 4431962.107 | 4445780.31 | 6330096.207 | 5188363.005 | 5179670.711 | 3807299.805 | 3608008.59 | 8828335.543 | 5391190.497 |
| PE-Cer 14:3;2O/18:5 | 551942.231 | 944519.9406 | 560264.9845 | 749983.5071 | 305057.4172 | 472853.9292 | 425190.8814 | 361743.265 | 314238.9016 | 379330.7154 | 466556.6645 | 257112.8236 |
| TG 8:0_16:4_22:5 | 1980207.419 | 2196421.845 | 1440150.232 | 1892353.667 | 2528191.685 | 2275394.832 | 2675720.363 | 2735981.295 | 2858171.567 | 3032777.855 | 2593464.321 | 2166491.244 |
| Naltrexone | 296007.2376 | 275433.9437 | 455915.8691 | 286809.9352 | 505761.3529 | 305767.8562 | 568933.2825 | 409909.4074 | 532328.2162 | 574193.8817 | 679071.6603 | 422569.4974 |
| Sphingosine | 35368600.88 | 33404325.56 | 20001771.08 | 24163950.66 | 6525780.484 | 6396919.708 | 6595695.367 | 6223495.51 | 6005374.9 | 6038519.034 | 5960822.039 | 5629720.489 |
| PG 17:1_20:3 | 268952.8218 | 383645.9925 | 67426.33294 | 269119.4322 | 2651384.394 | 1223889.172 | 2779665.316 | 1678050.81 | 1731669.799 | 5815005.639 | 990279.4398 | 562698.8291 |
| cis-Jasmone | 574927.2339 | 620161.1456 | 503458.9327 | 1137548.702 | 448207.8643 | 521039.3259 | 1538287.459 | 457112.1102 | 849216.0056 | 644599.616 | 544105.1525 | 1170328.91 |
| Uridine 5'-monophosphate (UMP) | 505753.2147 | 446537.8707 | 587360.5187 | 518774.21 | 5909378.153 | 3145134.288 | 2745542.995 | 5583460.284 | 3091207.323 | 3002367.978 | 3271066.42 | 3292892.377 |
| (E)-5-Hydroxyundec-2-enoic acid | 628883.1762 | 1283192.81 | 1442844.956 | 2246201.378 | 1641136.259 | 1701398.741 | 1783658.878 | 1563371.65 | 1548104.911 | 1504882.088 | 1504245.226 | 1902268.607 |

| Metabolites | PBS_1 | PBS_2 | PBS_3 | PBS_4 | BCG_1 | BCG_2 | BCG_3 | BCG_4 | BCGΔ*pe18*_1 | BCGΔ*pe18*_2 | BCGΔ*pe18*_3 | BCGΔ*pe18*_4 |
|---|---|---|---|---|---|---|---|---|---|---|---|---|
| 2-Phenyl-1H-benzimidazole-5-sulfonic acid | 1118965.705 | 5229965.649 | 4281184.275 | 19168078.08 | 26494178.28 | 553996.2066 | 1251008.437 | 869654.6988 | 541744.1922 | 1586378.204 | 850712.2529 | 13048349.3 |
| Ursolic Acid | 1398120.484 | 1314882.395 | 1250268.678 | 1441350.139 | 1397108.567 | 1344981.879 | 1440963.813 | 1414519.945 | 1366545.802 | 1414979.563 | 1401679.317 | 1341844.647 |
| Senkirkin | 500921.7496 | 577015.0333 | 480628.9494 | 521014.8615 | 553924.502 | 567766.5072 | 520581.6542 | 549396.0151 | 560025.0793 | 577086.578 | 497786.7562 | 521577.063 |
| Hippuric acid | 8560348.508 | 6584418.082 | 8474364.15 | 9291902.713 | 9816277.791 | 10570807.02 | 6849780.696 | 9443550.293 | 8138237.18 | 3121648.798 | 3608498.461 | 8000776.549 |
| PC(15:0/P-18:0) | 14248356.03 | 16655980.51 | 6871516.945 | 9251894.032 | 11076478.4 | 7985667.534 | 10023099.65 | 9502869.894 | 10611130.32 | 9731224.603 | 10243633.2 | 6154799.111 |
| N-Docosanoyltaurine | 560991.0829 | 1675343.152 | 905328.8919 | 479449.377 | 631976.5613 | 586449.2014 | 730448.2358 | 875376.2608 | 1176748.05 | 890099.0387 | 940783.3949 | 359410.6376 |
| PC(37:3) | 44523580.92 | 50391920.6 | 20860437.44 | 33056696.85 | 10822328.04 | 9790227.207 | 12929740.73 | 109148882 | 14929089.5 | 17952128.65 | 11477261.17 | 7913266.978 |
| Cer 8:1;2O/13:1 | 110861.3658 | 86218.65549 | 34581.31429 | 0 | 4061295.505 | 4326980.582 | 4417904.916 | 4537489.716 | 2412270.063 | 810371.4288 | 2620617.89 | 1004582.616 |
| S-Lactoylglutathione | 91533.10224 | 98009.56694 | 40558.3263 | 81900.4487 | 678408.9485 | 657201.1989 | 1382271.698 | 1006987.014 | 737241.2151 | 523649.2479 | 697605.4176 | 692876.5296 |
| PE O-9:0_8:0 | 601754.9347 | 518672.1567 | 320297.0579 | 550679.0552 | 846828.9839 | 792133.31 | 797142.9679 | 819637.8253 | 894232.8354 | 462831.0148 | 454996.9867 | 808574.6741 |
| Luteolin 7-methyl ether | 1660549.216 | 1500747.52 | 1490107.656 | 1678646.651 | 1431462.741 | 1728717.288 | 1717436.574 | 1429550.798 | 1625500.037 | 1576466.541 | 1446014.241 | 1547228.365 |

| Metabolites | PBS_1 | PBS_2 | PBS_3 | PBS_4 | BCG_1 | BCG_2 | BCG_3 | BCG_4 | BCGΔ*pe18*_1 | BCGΔ*pe18*_2 | BCGΔ*pe18*_3 | BCGΔ*pe18*_4 |
|---|---|---|---|---|---|---|---|---|---|---|---|---|
| 1-(2,5-Dimethylphenoxy)-3-(4-morpholinyl)-2-propanol | 708136.7256 | 1048097.941 | 3513692.551 | 972673.7499 | 682839.8785 | 649063.467 | 720744.4665 | 696809.6577 | 690751.5804 | 734527.0732 | 665183.5626 | 5926448.366 |
| Methyl 2-[(3aS,4R,5S,6S)-4-hydroxy-3-(hydroxymethyl)-6-[2-(4-hydroxyphenyl)acetyl]oxy-8-methyl-1-oxo-4,5,6,7-tetrahydro-3aH-azulen-5-yl]prop-2-enoate | 444451.6769 | 44307.96017 | 46207.91002 | 77431.25671 | 59937.51976 | 616055.7338 | 47397.99433 | 56878.39793 | 398374.0992 | 83417.48777 | 951363.5 | 53064.35292 |
| 3-(3,4-Dimethoxyphenyl)-1H-pyrazole-4-carboxylic acid | 2410166.86 | 3589878.247 | 3204681.821 | 4654840.071 | 569286.8026 | 870272.7437 | 946115.4592 | 1191402.301 | 2356703.942 | 849296.1747 | 1503950.489 | 10241947.44 |
| Arg-Trp | 562506.1417 | 803294.5855 | 494176.7031 | 430736.4797 | 519650.9868 | 776006.7387 | 431216.6822 | 542869.5869 | 595693.6256 | 544954.5038 | 572371.2076 | 684869.1026 |
| PC O-40:5 | 5861136.71 | 2889145.67 | 6856665.758 | 13417990.32 | 10556780.03 | 158744.5228 | 12694630.12 | 26309180.68 | 9539700.441 | 8494264.541 | 4432743.287 | 6584747.721 |
| Cer 12:1;2O/19:0 | 406216.255 | 43046.288 | 270044.1709 | 364935.653 | 681320.2573 | 1086307.006 | 614580.8247 | 931850.8355 | 540774.9154 | 337721.7264 | 809175.4447 | 1011552.827 |
| 2,2'-Iminodiacetic acid | 2629847.637 | 2454504.343 | 2390789.672 | 2215114.422 | 5316588.361 | 3338505.483 | 4917273.25 | 7654607.718 | 3912520.964 | 3873074.183 | 3237789.544 | 3504330.47 |
| 1-(2-Thienyl)-1-hep | 534807 | 390191 | 154653 | 435905 | 354494 | 377968 | 863531. | 222643 | 512649 | 313504 | 403700 | 305340 |

| **Metabolites** | **PBS_1** | **PBS_2** | **PBS_3** | **PBS_4** | **BCG_1** | **BCG_2** | **BCG_3** | **BCG_4** | **BCGΔ*pe18*_1** | **BCGΔ*pe18*_2** | **BCGΔ*pe18*_3** | **BCGΔ*pe18*_4** |
|---|---|---|---|---|---|---|---|---|---|---|---|---|
| tanone | 9.081 | 8.225 | 0.803 | 8.103 | 9.642 | 7.295 | 0047 | 5.327 | 4.277 | 7.573 | 0.682 | 9.609 |
| Aminocyclopyrachlor | 31185.41533 | 53214.76468 | 109927.4049 | 43584.87108 | 7862697.562 | 4151120.886 | 7282975.744 | 5936832.069 | 5508092.094 | 4969336.582 | 5012680.04 | 3537105.334 |
| LNAPE 18:1/N-17:1 | 1198042.269 | 416761.5102 | 962733.5266 | 815448.0392 | 725829.3036 | 0 | 643347.0599 | 95799.92381 | 1414981.289 | 875438.7814 | 690510.2605 | 0 |
| PC 8:0_26:5 | 786476.6416 | 1061286.757 | 501366.7978 | 1098872.267 | 2925791.606 | 4094195.914 | 3048853.881 | 3578255.27 | 3827616.55 | 3808425.037 | 2709148.629 | 2236697.102 |
| CerP 16:1;2O/18:0 | 1262975.334 | 1062974.809 | 555899.3353 | 273010.4285 | 554645.3777 | 0 | 732865.1407 | 423305.6776 | 567841.1103 | 606587.6233 | 508278.8051 | 308702.4377 |
| 2-Dodecylbenzenesulfonic_acid | 16549935.77 | 18978041.63 | 14814594.74 | 14183021.45 | 18651520.06 | 20575621.85 | 16554197.78 | 20957123.62 | 15931629.85 | 16212181.07 | 19069424.53 | 15276333.55 |
| PA 2:0_16:1 | 134223.0032 | 205071.8011 | 141486.3195 | 170292.2648 | 141636.062 | 154701.7956 | 132310.0373 | 141589.5631 | 126405.0754 | 104730.9742 | 116963.9513 | 89545.07553 |
| Ilicicolin F | 0 | 0 | 0 | 0 | 1080965.436 | 822640.0556 | 743437.928 | 640357.7873 | 945483.1534 | 987768.1291 | 1044756.433 | 893666.7028 |
| Tridodecylamine | 10588546.07 | 14008712.05 | 40140678.77 | 37978869.35 | 2978665.602 | 2904113.593 | 3593355.863 | 3729273.538 | 3576096.335 | 4884402.377 | 3201998.15 | 1831018.906 |
| MGDG O-8:0_24:5 | 867102.45 | 1002685.058 | 1280722.749 | 850173.5893 | 1358899.453 | 1546073.008 | 1000408.348 | 884134.6918 | 1025489.93 | 993520.1182 | 878951.9992 | 825702.5477 |
| PE O-13:1_7:0 | 310014.3313 | 369137.3786 | 150476.2925 | 298483.7501 | 213145.9998 | 38603.33335 | 133374.135 | 64915.07594 | 121445.5802 | 113482.6293 | 131608.2161 | 99169.02576 |
| PE O-18:1_22:4 | 943712.2989 | 204216.0587 | 592655.7695 | 714455.0968 | 373132.2694 | 0 | 339350.3475 | 152881.6719 | 524482.398 | 564495.6795 | 317624.9101 | 527639.3956 |
| NAGly 14:1/24:2 | 309373 | 396420 | 255702 | 298025 | 237330 | 288940 | 274470 | 325160 | 272701 | 345730 | 221060 | 311934 |

| **Metabolites** | **PBS_1** | **PBS_2** | **PBS_3** | **PBS_4** | **BCG_1** | **BCG_2** | **BCG_3** | **BCG_4** | **BCGΔ*pe18*_1** | **BCGΔ*pe18*_2** | **BCGΔ*pe18*_3** | **BCGΔ*pe18*_4** |
|---|---|---|---|---|---|---|---|---|---|---|---|---|
| | 7.096 | 0.46 | 7.802 | 1.987 | 4.553 | 5.426 | 4.052 | 6.699 | 9.356 | 1.839 | 6.126 | 1.4 |
| Phosphonic acid, P-[(3R)-3-amino-4-[(3-hexylphenyl)amino]-4-oxobutyl]- | 1693829.639 | 1702921.932 | 1635981.625 | 1953050.397 | 1740375.857 | 1499050.758 | 1719748.94 | 1516892.132 | 1653649.656 | 1811933.33 | 1642034.427 | 1977896.519 |
| MGDG O-16:3_6:0 | 148712.2554 | 191760.8178 | 219869.5958 | 388873.2854 | 104850.582 | 259814.9525 | 263105.0371 | 123978.0343 | 157240.3189 | 142942.1373 | 132432.3753 | 191212.7596 |
| Kaempferol 3-O-xyloside | 504110.9087 | 497541.1435 | 603597.314 | 524896.4477 | 518597.2243 | 577126.5727 | 492853.5018 | 493225.1713 | 633038.7998 | 508929.3715 | 661606.5103 | 549064.8832 |
| Propanedinitrile, 2-[[3-bromo-5-(1,1-dimethylethyl)-4-hydroxyphenyl]methylene]- | 98807.78555 | 95537.30786 | 98109.50558 | 52964.24155 | 13817604.23 | 5959752.874 | 14055471.73 | 8526130.274 | 3245946.096 | 8629974.269 | 7341599.224 | 3160671.861 |
| .beta.-Tocotrienol | 194915.5541 | 243169.8243 | 260778.7415 | 225713.2645 | 775137.3621 | 1142256.65 | 422418.8479 | 577813.3262 | 334524.1812 | 235744.1257 | 629504.6494 | 603232.0363 |
| 4-Cresol sulfate | 49946097.74 | 41062227.34 | 60486158.03 | 53302068.83 | 66794930.32 | 70267441.33 | 50144096.74 | 62374516.85 | 64651152.44 | 27917088.22 | 35063055.97 | 59119149.96 |
| 4-Ethylpiperidine | 7188939.307 | 7056389.615 | 7316185.512 | 7181560.456 | 7097764.165 | 7213424.18 | 7382444.98 | 6989240.583 | 7102396.225 | 6933410.561 | 6849881.462 | 7187534.648 |
| Desacetylcinobufotalin | 252784.2804 | 266798.8655 | 224166.1584 | 242786.5729 | 264520.1866 | 264487.4762 | 264289.8604 | 238426.136 | 294370.6063 | 284558.9304 | 400479.7935 | 221736.6603 |
| PE O-9:0_4:0 | 2243604.719 | 2566083.029 | 3355245.536 | 2090159.633 | 2731979.055 | 2259599.138 | 3869714.53 | 2712263.069 | 2247762.178 | 2939508.199 | 3471437.452 | 1983670.298 |

| Metabolites | PBS_1 | PBS_2 | PBS_3 | PBS_4 | BCG_1 | BCG_2 | BCG_3 | BCG_4 | BCGΔ*pe18*_1 | BCGΔ*pe18*_2 | BCGΔ*pe18*_3 | BCGΔ*pe18*_4 |
|---|---|---|---|---|---|---|---|---|---|---|---|---|
| LPA 20:1 | 63517.57837 | 40465.87448 | 198279.6352 | 187737.9305 | 438060.7508 | 433951.3422 | 437181.5765 | 397478.3392 | 88285.21265 | 131126.3099 | 168469.0598 | 299833.932 |
| Arg-Gly | 2523544.493 | 1514349.673 | 3247004.568 | 1323551.354 | 3373190.17 | 1146241.22 | 2019816.977 | 1307956.649 | 1417651.727 | 1608084.542 | 1584282.092 | 1039357.36 |
| 5-(2-Thienylmethylene)-2,4,6(1H,3H,5H)-pyrimidinetrione | 169382.4857 | 147387.7705 | 321207.6841 | 182348.2211 | 1844656.365 | 1433647.962 | 1997998.158 | 2018444.084 | 1095059.015 | 1932418.646 | 1676346.99 | 1219344.227 |
| Daphylloside | 441991.1521 | 998399.5126 | 610467.9906 | 41102.5077 | 237570.5417 | 1550169.639 | 311379.9734 | 351811.7798 | 321869.9437 | 285897.8703 | 428822.445 | 1837116.377 |
| 9-Hydroxyrisperidone | 7216.975509 | 16800.67001 | 22644.25485 | 10936.64051 | 315956.7012 | 210774.2419 | 220116.4561 | 241120.5885 | 280675.8779 | 282576.091 | 309717.2094 | 252878.228 |
| LDGCC 15:0 | 613712.2299 | 710746.779 | 539651.4177 | 605881.042 | 654304.1049 | 606920.0282 | 631218.381 | 566408.4639 | 631299.5554 | 595467.1526 | 664546.2936 | 538125.5042 |
| PA 2:0_18:0 | 934846.6668 | 331212.549 | 247473.7064 | 432767.6277 | 162316.2248 | 86642.18596 | 154319.6897 | 169520.172 | 178124.0482 | 218460.1166 | 192020.6866 | 116463.6218 |
| CE(5-iso PGF2VI) | 2369513.292 | 2538156.932 | 2215077.715 | 2343057.603 | 2096607.546 | 2587432.926 | 2796827.948 | 2470014.574 | 2418700.525 | 2531308.391 | 2359005.127 | 2191027.149 |
| betulinic acid | 911216.1835 | 945923.3348 | 884851.8294 | 1079782.346 | 1008911.706 | 998297.8435 | 1181132.348 | 938609.29 | 996685.5812 | 1135871.78 | 1019773.957 | 999798.3705 |

| Metabolites | PBS_1 | PBS_2 | PBS_3 | PBS_4 | BCG_1 | BCG_2 | BCG_3 | BCG_4 | BCGΔ *pe18*_1 | BCGΔ *pe18*_2 | BCGΔ *pe18*_3 | BCGΔ*pe18*_4 |
|---|---|---|---|---|---|---|---|---|---|---|---|---|
| 6b-Hydroxy-3,14b,14c-trimethyl-5-oxo-2,3,4,5,6b,7,8,8a,9,14,14b,14c,15,16-tetradecahydro-4,16a-epoxy[1]benzoxocino[7',8':6,7]indeno[1,2-b]indol-3-yl acetate | 24181.1 1135 | 14775.9 0406 | 24354.8 9488 | 17437.3 9757 | 77548.0 3325 | 50917.5 0178 | 114971. 7051 | 69054.9 6227 | 66265.7 4784 | 71690.9 1228 | 72868.5 5405 | 47346.9 233 |
| M534T246 | 463331. 2549 | 480693. 1757 | 564041. 3089 | 543386. 5452 | 462549. 7635 | 486946. 8425 | 554428. 0874 | 469028. 5496 | 664093. 3043 | 661558. 8223 | 715441. 6289 | 877172. 5023 |
| Eicoseneoylcarnitine (Car(20:1)) | 697101. 4321 | 578498. 2735 | 621870. 8803 | 613040. 6782 | 820109. 8873 | 641110. 2228 | 909482. 2574 | 919619. 6567 | 512162. 4609 | 645332. 3306 | 555522. 3404 | 325082. 1942 |
| Ferutinin | 194500 5.811 | 215279 1.893 | 169813 6.766 | 185421 3.378 | 212150 7.926 | 206549 7.773 | 211206 3.517 | 190387 4.238 | 207714 1.26 | 215195 3.729 | 201383 9.205 | 182744 2.377 |
| (3R,5aS,10bR,11aS)-3-Benzyl-10b-(2-methylbut-3-en-2-yl)-6,10b,11,11a-tetrahydro-2H-pyrazino[1',2':1,5]pyrrolo[2,3-b]indole-1,4(3H,5aH)-dione | 535504. 8974 | 521595. 9063 | 414093. 7094 | 447320. 9764 | 540540. 6372 | 510802. 4542 | 561846. 5358 | 544544. 3631 | 539697. 7006 | 479216. 5818 | 502149. 8205 | 492145. 9688 |
| 6-Ketotestosterone | 52125.5 | 50184.2 | 24786.9 | 435023. | 199895. | 217974. | 466055. | 277627. | 227234. | 108608. | 302173. | 104605 |

| **Metabolites** | **PBS_1** | **PBS_2** | **PBS_3** | **PBS_4** | **BCG_1** | **BCG_2** | **BCG_3** | **BCG_4** | **BCGΔ*pe18*_1** | **BCGΔ*pe18*_2** | **BCGΔ*pe18*_3** | **BCGΔ*pe18*_4** |
|---|---|---|---|---|---|---|---|---|---|---|---|---|
| enanthate | 4709 | 8305 | 6107 | 2167 | 7843 | 6959 | 5564 | 147 | 9302 | 8871 | 9601 | 6.279 |
| 2,5-Dimethoxy-4-ethylamphetamine | 404939.9162 | 331888.9451 | 442531.5527 | 320804.3155 | 235235.2516 | 392815.7829 | 295695.8147 | 443471.0177 | 291983.2939 | 392682.6831 | 506813.0146 | 661418.4331 |
| 2-Oxooctanoic acid | 535553.1884 | 519071.7456 | 284404.9827 | 341203.0073 | 333658.9801 | 476450.7436 | 634746.364 | 437874.8385 | 268687.9229 | 743238.9571 | 333792.5206 | 492468.5775 |
| Glutathione disulfide | 816842.8663 | 1006533.629 | 1154596.146 | 918517.767 | 1195747.177 | 728611.3147 | 1818696.834 | 1458591.562 | 1133056.972 | 1181649.65 | 1335835.162 | 1927887.66 |
| PC 8:0_30:5 | 25314375.25 | 26009226.28 | 25066673.21 | 52552373.59 | 18404149.88 | 19157328.45 | 21685666 | 18000720.44 | 21105890.13 | 22285058.8 | 20799135.08 | 17170321.3 |
| LPE O-20:4 | 2284302.074 | 2422219.366 | 1338395.368 | 1331550.262 | 609434.9934 | 1658393.946 | 815385.5791 | 660735.7767 | 471116.1034 | 587303.0335 | 878996.4557 | 423725.4394 |
| LNAPE 16:1/N-18:0 | 5997516.682 | 3978410.824 | 5908146.5 | 4511177.612 | 3712254.62 | 92991.24016 | 3127047.921 | 1056685.212 | 5199671.543 | 3796257.454 | 4403281.564 | 1330520.102 |
| Adipic acid | 979268.6711 | 1070730.182 | 1229466.806 | 1055918.896 | 1686390.971 | 1342332.777 | 1558061.187 | 1550919.376 | 1126194.905 | 1017713.658 | 1201960.373 | 1320947.329 |
| 2-[4-(1-Methyl-1-phenylethyl)phenoxy]propanoic acid | 1150859.792 | 1184068.639 | 1076629.494 | 1140884.198 | 1115132.504 | 1103428.066 | 1118083.954 | 1088020.921 | 1095090.115 | 1099187.141 | 1090809.781 | 1145188.359 |
| DG 24:3 | 1007505.045 | 1548499.909 | 680495.8183 | 611179.2152 | 1101303.077 | 2125097.047 | 1256859.921 | 1692433.782 | 567210.206 | 771357.6614 | 1688470.744 | 1346865.777 |
| LPC O-17:1 | 1435424.329 | 788729.476 | 1023749.757 | 1271970.104 | 1811233.066 | 2389637.41 | 1948277.089 | 2340263.974 | 1703179.071 | 1829791.11 | 1956652.918 | 1943312.695 |

| **Metabolites** | **PBS_1** | **PBS_2** | **PBS_3** | **PBS_4** | **BCG_1** | **BCG_2** | **BCG_3** | **BCG_4** | **BCGΔ*pe18*_1** | **BCGΔ*pe18*_2** | **BCGΔ*pe18*_3** | **BCGΔ*pe18*_4** |
|---|---|---|---|---|---|---|---|---|---|---|---|---|
| 2-Hexaprenyl-3-methyl-5-hydroxy-6-methoxy-1,4-benzoquinol | 755267.7841 | 882434.7193 | 771680.1557 | 702665.0111 | 949869.5907 | 749321.8041 | 788392.7402 | 848048.9581 | 926095.154 | 901748.2984 | 815170.6421 | 663220.6628 |
| SQDG 12:0_16:3 | 105262.7953 | 97421.03789 | 183055.126 | 103469.2413 | 126588.2309 | 136922.3082 | 160940.5422 | 121140.304 | 87855.9018 | 104863.6674 | 104081.3709 | 66617.3675 |
| 1-Palmitoyl-2-azelaoylphosphatidylcholine | 0 | 107059.7533 | 259828.6245 | 540981.3278 | 959169.8312 | 556509.3226 | 1159399.903 | 214457.43 | 757415.8092 | 840573.0576 | 716181.7933 | 450280.0767 |
| 1,2,5-Trimethyl-1H-pyrrole | 2351028.666 | 2435497.719 | 2080326.842 | 2834898.386 | 2122031.79 | 2343514.809 | 2080530.014 | 2490235.453 | 2644121.877 | 2101888.288 | 2793667.299 | 1977743.969 |
| Chlorophene | 1050455.442 | 1043614.475 | 1702338.198 | 3144034.934 | 741237.9296 | 1207771.036 | 1505975.998 | 4373631.401 | 1456047.278 | 667933.1382 | 643352.108 | 1529134.881 |
| ST 24:1;O4/12:0 | 588542.7392 | 524902.4101 | 697447.9612 | 1159971.847 | 46662.65371 | 427370.6494 | 783666.2953 | 602268.7379 | 709942.2316 | 742428.7929 | 433685.2011 | 362654.0655 |
| DGDG O-26:6_16:0 | 203631.2915 | 178825.7981 | 235377.5407 | 135407.8382 | 442930.1464 | 378185.6599 | 448299.5888 | 316727.2446 | 346226.1246 | 333000.1112 | 549141.3328 | 164807.5209 |
| PC 35:0_18:2 | 118944.4595 | 87408.33074 | 110189.2921 | 83445.2603 | 113436.8021 | 0 | 98235.47456 | 50218.55164 | 71168.97501 | 93668.00808 | 122132.1419 | 53418.20241 |
| LNAPE 4:0/N-18:4 | 530602.3262 | 471509.2942 | 863877.615 | 542773.9581 | 718990.4614 | 555200.146 | 549340.0583 | 660344.6184 | 882580.5335 | 660561.7516 | 740790.0539 | 804497.2412 |
| 2-16:1-lysophosphatidylglycerol | 0 | 0 | 0 | 0 | 1975897.09 | 1875679.487 | 1782848.042 | 1640744.222 | 1674151.158 | 1614852.612 | 1804595.878 | 872040.8828 |
| 51333-22-3 | 434917. | 471246. | 403647. | 455963. | 476724. | 424098. | 737755. | 502672. | 447217. | 501557. | 469153. | 414965. |

| Metabolites | PBS_1 | PBS_2 | PBS_3 | PBS_4 | BCG_1 | BCG_2 | BCG_3 | BCG_4 | BCGΔ*pe18*_1 | BCGΔ*pe18*_2 | BCGΔ*pe18*_3 | BCGΔ*pe18*_4 |
|---|---|---|---|---|---|---|---|---|---|---|---|---|
| | 1193 | 7462 | 2956 | 6636 | 8924 | 8641 | 2347 | 0877 | 2822 | 3173 | 8898 | 9691 |
| 1-Methyl-2-imidazolidinethione | 2854147.301 | 1008212.986 | 4763769.858 | 2020173.373 | 923434.1326 | 2457920.11 | 3892003.207 | 4860782.815 | 2814922.424 | 1265089.057 | 1738355.717 | 2997062.611 |
| 8-Isobutanoylneosolaniol | 2663707.633 | 2564605.859 | 2968868.513 | 2611684.345 | 3007098.083 | 2361029.831 | 3143262.416 | 2880548.474 | 2986207.792 | 2949058.075 | 2962691.95 | 2933361.792 |
| 1H-Imidazo[4,5-c]pyridine-6-carboxylic acid, 1-[[4-(dimethylamino)-3-methylphenyl]methyl]-5-(2,2-diphenylacetyl)-4,5,6,7-tetrahydro-, (6S)- | 252718.2655 | 298931.2309 | 234683.5613 | 302316.6848 | 393477.9807 | 366441.0842 | 250961.0037 | 369388.69 | 414620.3821 | 126205.4909 | 143563.1575 | 308434.8543 |
| LNAPE 3:0/N-18:4 | 1283451.375 | 1390970.645 | 1455228.974 | 2076448.131 | 1941572.213 | 1480392.073 | 880061.0689 | 755699.1676 | 1334117.523 | 1731214.86 | 1545475.023 | 1411002.722 |
| Carbamic acid, N-[4-(5-methoxy-2-oxo-1,3,4-oxadiazol-3(2H)-yl)-2-methylphenyl]-, phenylmethyl ester | 450662.7389 | 510215.3534 | 440034.393 | 459003.1792 | 446255.4496 | 502844.8762 | 457481.1315 | 434733.6632 | 437929.6875 | 483888.9411 | 465456.3265 | 438421.241 |

| Metabolites | PBS_1 | PBS_2 | PBS_3 | PBS_4 | BCG_1 | BCG_2 | BCG_3 | BCG_4 | BCGΔ*pe18*_1 | BCGΔ*pe18*_2 | BCGΔ*pe18*_3 | BCGΔ*pe18*_4 |
|---|---|---|---|---|---|---|---|---|---|---|---|---|
| Benzoic acid, 4-[(4,6-dihydroxy-5-methoxy-2,5-dimethyl-3-oxo-1-cyclohexen-1-yl)oxy]-2-hydroxy-3,6-dimethyl-, methyl ester | 514140.196 | 443110.3512 | 952215.6408 | 634978.6688 | 679675.0502 | 775922.5972 | 775410.0657 | 656386.0512 | 616527.5812 | 320584.0982 | 347270.524 | 679059.4831 |
| 5-(2-Acetoxy-12-hydroxytridecyl)-1,3-benzenediol | 1890083.192 | 2618265.096 | 3040948.666 | 2098909.04 | 1259283.711 | 1793710.968 | 1117796.898 | 1422434.254 | 1277583.358 | 1540315.803 | 1092080.123 | 6437715.232 |
| Cer 14:3;2O/4:0 | 15530290.85 | 15851097.47 | 5484052.735 | 12220879.01 | 4654931.922 | 6000678.312 | 4323261.018 | 4160762.93 | 8009703.223 | 7291780.173 | 4251234.571 | 7411989.02 |
| PG O-17:2_20:4 | 425703.1181 | 262960.0906 | 515321.4093 | 300183.784 | 880039.72 | 872393.5884 | 1467675.609 | 1085084.169 | 929656.1166 | 1883468.55 | 994439.3828 | 772779.8756 |
| PC 9:0_30:7 | 799273.9413 | 887953.4919 | 477336.95 | 700993.3254 | 3926940.349 | 4407834.682 | 2856273.624 | 3761847.713 | 3623393.623 | 3333896.819 | 2702917.175 | 2571838.758 |
| (-)-Caryophyllene oxide | 1373935.89 | 1450677.087 | 1276659.798 | 1343597.698 | 1018340.634 | 1065923.973 | 1222992.181 | 1164089.311 | 1051395.171 | 1093734.41 | 958067.7427 | 1221913.526 |
| 1-pentadecanoyl-glycero-3-phosphate | 178737.6866 | 186514.5142 | 301683.3871 | 310952.221 | 627138.5957 | 508305.941 | 402875.0907 | 441228.0263 | 397953.3135 | 467121.8141 | 478888.9319 | 41905.76948 |
| N-[4-(4-Fluorophenyl)-1,3-thiazol-2-yl]nicotinamide | 390350.6296 | 141863.7792 | 450253.8662 | 212673.7311 | 317406.2279 | 340430.3686 | 319374.2618 | 274934.257 | 238121.5257 | 275613.2584 | 155342.7182 | 256478.1084 |

| **Metabolites** | **PBS_1** | **PBS_2** | **PBS_3** | **PBS_4** | **BCG_1** | **BCG_2** | **BCG_3** | **BCG_4** | **BCGΔ *pe18*_1** | **BCGΔ *pe18*_2** | **BCGΔ *pe18*_3** | **BCGΔ*pe18*_4** |
|---|---|---|---|---|---|---|---|---|---|---|---|---|
| Benzeneethanamine, 2,5-dimethoxy-N-[(2-methoxyphenyl)methyl]-4-nitro- | 877354.1872 | 965296.7323 | 841025.0131 | 1013224.906 | 1022493.66 | 1074847.047 | 943073.4121 | 906282.3339 | 938817.9339 | 1029673.328 | 913122.1833 | 1023253.849 |
| Curvularin | 55545.44619 | 0 | 375897.3851 | 0 | 6302550.492 | 10756554.54 | 9651787.188 | 4863686.263 | 2950514.23 | 2397305.129 | 4551946.812 | 1095302.956 |
| 1,3,7-Trimethyluric acid | 7158829.223 | 1256490.039 | 6518058.086 | 7285084.263 | 5134712.003 | 7690840.768 | 2410524.182 | 6545671.486 | 2860270.383 | 697650.196 | 1705264.929 | 5038524.164 |
| N-Arachidonoyltaurine | 775233.8241 | 753917.7382 | 768199.5149 | 905481.6613 | 722848.3692 | 774105.0802 | 848890.5587 | 785917.7122 | 1232911.703 | 739290.7114 | 1023035.367 | 628730.2439 |
| Glycyrrhizic Acid, Ammonium Salt | 98514.84415 | 137099.6286 | 110584.9249 | 119336.0739 | 131920.8022 | 123689.4904 | 130181.2259 | 132082.8021 | 124834.442 | 150431.9787 | 135586.9361 | 135366.0682 |
| Biotin | 1543438.944 | 1111913.429 | 2279439.589 | 1625322.433 | 1587678.416 | 1937266.63 | 1878797.755 | 1802425.393 | 1677788.148 | 733232.4751 | 884262.2656 | 1684759.399 |
| 3-Hydroxyhexadecanoylcarnitine | 143644.4438 | 500065.3693 | 805908.3465 | 577672.7856 | 2594853.486 | 1742398.189 | 3278419.252 | 2044648.045 | 1549111.143 | 2554610.373 | 1810772.939 | 1050089.735 |
| Purothionin_AII | 1683199.178 | 1724953.676 | 2515233.117 | 2424964.007 | 1785301.501 | 2074202.366 | 1831075.389 | 1766952.893 | 2305010.276 | 3881167.638 | 2318502.377 | 2226768.132 |
| Hydroxyisocaproic acid | 979892.2292 | 924889.0293 | 1031719.892 | 1038714.407 | 1060642.122 | 1062600.852 | 1213939.334 | 1179399.942 | 1005065.938 | 1117498.934 | 966558.4951 | 1037359.701 |
| 6,6'-Methylenebis(2,4-di-tert-butylphen | 384369.9921 | 414097.4452 | 331816.6751 | 398276.8707 | 367591.8079 | 369518.9999 | 412183.3111 | 363770.7326 | 401866.2186 | 381337.0255 | 341098.7361 | 337955.2079 |

| Metabolites | PBS_1 | PBS_2 | PBS_3 | PBS_4 | BCG_1 | BCG_2 | BCG_3 | BCG_4 | BCGΔ*pe18*_1 | BCGΔ*pe18*_2 | BCGΔ*pe18*_3 | BCGΔ*pe18*_4 |
|---|---|---|---|---|---|---|---|---|---|---|---|---|
| ol) | | | | | | | | | | | | |
| Tetradecanamide, N-[(1R,2R)-2-hydroxy-1-(hydroxymethyl)-2-(4-nitrophenyl)ethyl]- | 898999.1314 | 653368.4129 | 1134866.577 | 817203.0987 | 2691174.214 | 1609653.754 | 1848996.156 | 1675050.65 | 1393394.312 | 1596062.27 | 2084087.273 | 1071914.448 |
| 2-Methyl-N-(4-methylphenyl)benzenesulfonamide | 1787764.941 | 1804137.168 | 2473786.896 | 1177613.374 | 1235244.322 | 763273.1447 | 1661356.585 | 1266078.654 | 535765.5488 | 668512.0392 | 888676.1407 | 771476.0158 |
| N-n-Butylpropionamide | 3934249.325 | 6065236.711 | 5117928.805 | 9108009.628 | 9608480.688 | 5189383.776 | 2986326.301 | 2969049.231 | 4921978.875 | 4804817.341 | 5009288.856 | 3609782.874 |
| 1-Heptanone, 1-(4-methoxyphenyl)-2-(1-pyrrolidinyl)- | 1257865.679 | 3323055.119 | 1132884.935 | 921570.8079 | 3143352.297 | 1231154.863 | 1418955.43 | 2998577.968 | 1308367.713 | 1272536.991 | 1365255.116 | 1282511.725 |
| Ala-His | 339611.3783 | 243628.9206 | 527983.7044 | 320054.6395 | 935056.9851 | 578726.6673 | 991077.3286 | 748297.0888 | 594338.5402 | 534488.5861 | 622772.0912 | 553923.8649 |
| 8-Acetamido-2-methyl-7-oxononanoic acid | 423709.3706 | 549101.617 | 463852.013 | 763560.0253 | 372704.3331 | 349232.0738 | 563786.7111 | 264015.89 | 402149.8817 | 425725.6756 | 339787.2592 | 811122.7414 |
| NAOrn 18:5/16:3 | 13878933.19 | 13617258.96 | 11841621.09 | 12490178.64 | 14610720.09 | 14310117.4 | 13485424.86 | 12250010.8 | 14251584.29 | 13427012.49 | 13437502.02 | 12424742.7 |

| **Metabolites** | **PBS_1** | **PBS_2** | **PBS_3** | **PBS_4** | **BCG_1** | **BCG_2** | **BCG_3** | **BCG_4** | **BCGΔ*pe18*_1** | **BCGΔ*pe18*_2** | **BCGΔ*pe18*_3** | **BCGΔ*pe18*_4** |
|---|---|---|---|---|---|---|---|---|---|---|---|---|
| 4-Methyloxazole-5-carboxylic acid | 876000.676 | 667627.9841 | 1383944.207 | 850033.0523 | 1695783.297 | 1851908.339 | 1665166.099 | 3425896.034 | 1467410.721 | 1041934.453 | 1308329.212 | 1938315.623 |
| 3,5-Dicaffeoylquinic acid | 186469.2944 | 18239.61876 | 380597.7533 | 54321.08217 | 5616401.001 | 995835.6948 | 4192411.676 | 1934021.709 | 1880259.184 | 1551375.115 | 2045378.248 | 803994.1184 |
| SM 23:3;2O/28:1 | 71206.21072 | 0 | 70424.90267 | 78149.6025 | 309324.8632 | 0 | 257577.3365 | 274438.9867 | 187894.8453 | 144805.5873 | 268174.6695 | 166975.5202 |
| 3'-O-Methylguanosine | 64575.77081 | 177874.9584 | 139493.3196 | 86340.12003 | 1593250.586 | 833919.8575 | 1524779.932 | 1197615.83 | 1110466.426 | 1153387.192 | 1395610.358 | 964356.3225 |
| sphingofungin B | 3881675.775 | 3491998.919 | 3123794.9 | 3068407.886 | 3426800.861 | 3696125.675 | 5719529.861 | 3709991.961 | 3849370.197 | 3742603.934 | 4909029.598 | 3852898.942 |
| PE-Cer 22:3;2O/26:6 | 22534.39512 | 79907.35556 | 39635.90861 | 87049.91794 | 2542996.821 | 2422681.863 | 2144580.556 | 2329617.097 | 1127970.565 | 770605.9861 | 923287.7533 | 770257.042 |
| 11-Deoxyprostaglandin E2 | 665876.3887 | 770463.2304 | 554024.4693 | 730217.9784 | 747018.3939 | 743677.9634 | 809511.9231 | 649190.1194 | 879651.7889 | 942027.1071 | 709031.6967 | 801771.0175 |
| Glu-Val-Phe | 229148.117 | 259092.8233 | 306986.02 | 248473.2206 | 437305.0306 | 347473.8195 | 505685.6434 | 398286.3921 | 475300.3209 | 690505.8725 | 703376.2716 | 414387.8563 |
| 2,2'-(2-Hydroxyethylazanediyl)diacetic acid | 14929290.6 | 10330647.9 | 17433119.49 | 14363564.9 | 8552563.138 | 2701184.921 | 8447482 | 11888551.5 | 9838075.344 | 4951592.517 | 6284129.046 | 2128445.922 |
| PG 18:1_19:1 | 1598700.431 | 611448.6762 | 2123482.831 | 988464.1405 | 1161533.379 | 436545.3709 | 1080330.161 | 650521.3678 | 1808415.716 | 1577242.218 | 888950.1072 | 565033.1434 |
| PS 18:1_22:6;4O | 6353.196667 | 5831.619265 | 25081.59343 | 11286.08271 | 113934.7227 | 109457.3381 | 93931.89831 | 92382.81822 | 45292.93494 | 63664.57657 | 115601.9457 | 49161.25881 |

| Metabolites | PBS_1 | PBS_2 | PBS_3 | PBS_4 | BCG_1 | BCG_2 | BCG_3 | BCG_4 | BCGΔ*pe18*_1 | BCGΔ*pe18*_2 | BCGΔ*pe18*_3 | BCGΔ*pe18*_4 |
|---|---|---|---|---|---|---|---|---|---|---|---|---|
| 2-[(1E)-2-(4-Methoxyphenyl)ethenyl]-4(3H)-quinazolinone | 6508293.641 | 5348076.576 | 9165124.856 | 7434349.677 | 5420068.261 | 3968284.801 | 5382461.467 | 4055028.837 | 2982384.673 | 3577259.408 | 4396459.22 | 2853318.48 |
| 4-Picoline | 1467921.696 | 1035560.097 | 784033.9759 | 1374661.448 | 1274861.239 | 887402.9126 | 1630440.74 | 2059416.216 | 505281.1476 | 4065050.935 | 5927181.249 | 950704.1717 |
| SM(d18:1/24:0) | 17962803.66 | 19890538.98 | 24953543.28 | 22458547.21 | 21386457.01 | 16567279.23 | 25586052.32 | 19568013.78 | 19250327.64 | 23540866.9 | 20266257.67 | 8290658.502 |
| Glycohyodeoxycholic acid | 977378.0212 | 823916.2479 | 1443940.316 | 958467.455 | 1636621.286 | 1607943.031 | 2323764.907 | 1508212.308 | 1375287.552 | 960682.4198 | 988933.4887 | 1083126.185 |
| M415T517 | 1219973.929 | 1874619.506 | 779918.1688 | 875531.2379 | 2290121.132 | 2566750.539 | 2350725.035 | 1817614.745 | 1020472.378 | 1231669.327 | 1246409.362 | 1675847.622 |
| PA 3:0_22:6 | 239038.3563 | 232421.0744 | 260098.3114 | 177740.7988 | 765943.0474 | 861084.3164 | 671990.1596 | 668749.402 | 858606.0089 | 616077.151 | 969246.6825 | 660450.3141 |
| LPC O-10:0 | 278559.4665 | 309923.7606 | 280290.7748 | 278141.6914 | 314140.9998 | 302193.017 | 332502.8691 | 277933.1764 | 292159.9197 | 327852.3314 | 299791.7119 | 277253.0608 |
| LPC 44:11-SN2 | 537397.7546 | 642707.5335 | 680777.1167 | 641865.0227 | 641448.9582 | 686639.6068 | 712904.5246 | 615053.2314 | 655129.5159 | 577996.0937 | 631772.7725 | 665930.6628 |
| Stachydrine | 4089958.067 | 5620248.962 | 4272811.62 | 5318271.191 | 4934121.128 | 3004285.408 | 3651862.192 | 3455145.079 | 2969867.658 | 3723486.537 | 2882295.359 | 5462897.059 |
| Didodecyldimethyl ammonium cation | 2044983.163 | 2202459.328 | 833995.1617 | 1785302.301 | 1125732.841 | 947060.5436 | 965443.5388 | 1115395.584 | 1237965.264 | 1199355.711 | 922775.8982 | 771756.0002 |
| BVT 2733 | 847129.54 | 479954.781 | 1275876.772 | 677903.4661 | 925323.6025 | 684498.6228 | 892725.5851 | 687359.461 | 658238.6643 | 577523.3036 | 527978.0182 | 469710.2948 |

| Metabolites | PBS_1 | PBS_2 | PBS_3 | PBS_4 | BCG_1 | BCG_2 | BCG_3 | BCG_4 | BCGΔ*pe18*_1 | BCGΔ*pe18*_2 | BCGΔ*pe18*_3 | BCGΔ*pe18*_4 |
|---|---|---|---|---|---|---|---|---|---|---|---|---|
| 5,7-Dichloro-1-benzofuran-2-carboxylic acid | 107561.6191 | 43100.25424 | 111282.2029 | 57106.33386 | 1758434.314 | 952781.0227 | 989062.7361 | 704478.7527 | 528228.1562 | 1198598.762 | 2703841.362 | 692022.2813 |
| Geranyllinalool | 2393960.975 | 1335857.204 | 1193893.745 | 1493747.634 | 1351174.449 | 1242778.401 | 1331136.565 | 1152137.969 | 1513425.5 | 792819.7661 | 1356752.744 | 751028.4853 |
| Pregna-1,4-diene-3,20-dione, 6,9-difluoro-11,21-dihydroxy-16,17-[(1-methylethylidene)bis(oxy)]-, (6.alpha.,11.beta.,16.alpha.)- | 98948.98335 | 82948.69543 | 113974.1968 | 93092.39514 | 134752.7842 | 99870.90795 | 95168.54216 | 106029.5695 | 107366.1034 | 139116.9149 | 120601.718 | 115566.1911 |
| LPA 22:6 | 802187.7861 | 915230.4329 | 1060800.942 | 898564.4327 | 1038046.366 | 1121540.798 | 838549.9201 | 1014735.201 | 677300.6201 | 588616.4286 | 718570.1572 | 767069.9758 |
| Ethyl dodecanoate | 1122852.308 | 892448.0056 | 1355779.371 | 1060161.192 | 3464991.237 | 2281998.127 | 3664541.177 | 2489395.935 | 1827221.74 | 2674978.749 | 2674388.358 | 1604332.531 |
| Imidazol-1-yl-acetic acid | 2910665.489 | 2739087.512 | 5128429.137 | 3427391.075 | 4672573.984 | 3771007.085 | 4957898.99 | 4119575.247 | 3883151.038 | 3335414.505 | 3703100.226 | 3488007.791 |
| PE O-28:3_6:0 | 483700.3965 | 256571.8237 | 416588.7962 | 594575.6375 | 191333.7732 | 97566.76576 | 117794.9574 | 22905.67846 | 696335.5111 | 254727.7803 | 0 | 0 |
| PC O-37:7 | 361793.7804 | 353865.6566 | 356217.7891 | 387590.1482 | 333955.2157 | 349422.5655 | 349256.6313 | 311994.9537 | 266942.5331 | 387198.651 | 365008.2623 | 557927.9667 |
| Cer 9:0;2O/16:4 | 269451 | 348766 | 220673 | 270992 | 247692 | 260107 | 303834 | 282529 | 263925 | 284771 | 265704 | 271942 |

| Metabolites | PBS_1 | PBS_2 | PBS_3 | PBS_4 | BCG_1 | BCG_2 | BCG_3 | BCG_4 | BCGΔ*pe18*_1 | BCGΔ*pe18*_2 | BCGΔ*pe18*_3 | BCGΔ*pe18*_4 |
|---|---|---|---|---|---|---|---|---|---|---|---|---|
| | 7.699 | 5.519 | 7.077 | 2.654 | 0.616 | 2.369 | 4.363 | 1.363 | 2.693 | 1.596 | 3.898 | 7.997 |
| Berberrubine | 5154068.008 | 6611429.803 | 5770331.151 | 5929878.261 | 68105127.23 | 6337041.882 | 6005564.167 | 7305990.757 | 7904872.155 | 7053488.183 | 5977500.096 | 6566389.312 |
| 4-(2,3-Dihydro-1H-perimidin-2-yl)phenol | 691407.6506 | 535473.7912 | 569578.6297 | 581186.9289 | 643915.3572 | 574875.1026 | 629361.3302 | 698591.3365 | 586543.8687 | 524020.8174 | 543889.0068 | 574202.6405 |
| 2-Anilinopropanoic acid | 4171042.859 | 3374072.932 | 9086169.705 | 4630737.77 | 9303262.995 | 8509550.65 | 10266110.23 | 8807130.444 | 8955257.986 | 5305733.733 | 6340036.953 | 9450928.338 |
| Buxifoliadine A | 565005.2535 | 642731.7754 | 404330.9841 | 540595.9833 | 511396.5103 | 526794.3766 | 551046.5347 | 380308.2986 | 434956.5236 | 425580.4627 | 562749.4929 | 544538.3559 |
| 1-Palmitoyl-2-hydroxy-sn-glycero-3-phospho-(1'-rac-glycerol) | 495327.2183 | 479560.8029 | 462483.0821 | 473077.5409 | 4856215.364 | 2981036.622 | 4272272.966 | 3168768.343 | 3436567.793 | 3337943.029 | 5370057.354 | 3173974.678 |
| 2-Bromo-1-(3,4-dimethoxyphenyl)ethanone | 6315943.017 | 2374724.563 | 6142033.527 | 7665814.773 | 2760950.63 | 6454285.859 | 4405584.078 | 7141822.358 | 7427348.999 | 8669395.436 | 6705449.592 | 5667270.211 |
| PG 16:1_16:1 | 19505.32648 | 0 | 37517.18616 | 33737.40288 | 1771785.445 | 2190849.161 | 2127608.557 | 1189075.71 | 1460968.269 | 2286365.114 | 1583323.597 | 876828.9386 |
| Dodecanoic acid | 139025071.9 | 137080165.5 | 142892748.5 | 163342358.7 | 123653050.7 | 162890147.1 | 103512596.1 | 129887322 | 122884855 | 117257766.7 | 129304274.5 | 132789612.3 |
| Trillin | 225844.2293 | 324598.3547 | 521272.2925 | 420508.7415 | 249402.1456 | 317183.1783 | 313685.273 | 286848.48 | 374516.3908 | 53967.83647 | 242749.8792 | 370195.5786 |
| SM(d16:1/24:1(15Z | 315329 | 361177 | 479221 | 306928 | 487814 | 391181 | 669448 | 471368 | 563644 | 717002 | 581135 | 250753 |

| Metabolites | PBS_1 | PBS_2 | PBS_3 | PBS_4 | BCG_1 | BCG_2 | BCG_3 | BCG_4 | BCGΔ *pe18*_1 | BCGΔ *pe18*_2 | BCGΔ *pe18*_3 | BCGΔ*pe18*_4 |
|---|---|---|---|---|---|---|---|---|---|---|---|---|
| )) | 9.761 | 2.42 | 3.399 | 1.53 | 2.569 | 0.774 | 2.242 | 8.923 | 4.031 | 4.545 | 2.401 | 0.712 |
| M547T251 | 708148.1627 | 739668.1975 | 677183.0263 | 722256.1067 | 722600.1842 | 804140.7566 | 687850.4937 | 781266.3366 | 740500.8642 | 692523.8614 | 748901.94 | 716631.9323 |
| MG 21:0 | 3176081.418 | 3812909.808 | 1947414.728 | 3029717.594 | 2403945.994 | 2206863.366 | 2555756.093 | 2789308.799 | 2728697.692 | 2394155.651 | 2211431.763 | 1795985.149 |
| Pantothenic acid | 1900129.666 | 1698670.612 | 3126476.194 | 2688265.085 | 3260085.861 | 3207020.471 | 3783958.734 | 3402379.55 | 3686148.026 | 2130034.738 | 2431638.938 | 2586394.567 |
| 5-(3-Phenylpropyl)-1H-1,2,3,4-tetrazole | 897741.5686 | 753395.7767 | 531064.2225 | 836491.4172 | 824249.9986 | 1687360.448 | 7610924.489 | 403529.5944 | 2051827.831 | 844488.1095 | 508542.9883 | 7463840.662 |
| 8,11-Eicosadiynoic acid | 1484148.276 | 749201.5275 | 1255135.345 | 1419827.835 | 1195985.02 | 1231148.653 | 966149.4006 | 1255601.865 | 921528.3465 | 926334.3034 | 1386812.168 | 762105.3842 |
| Repaglinide | 100161.4658 | 94111.39824 | 241550.3249 | 104593.1866 | 85719.39382 | 63834.70828 | 93111.16969 | 78083.76029 | 53132.95003 | 48756.74378 | 65245.74352 | 53639.91083 |
| Maclurin | 2665137.413 | 1622198.764 | 2703106.815 | 1398882.563 | 3207869.879 | 2537028.625 | 2146992.326 | 1855052.851 | 2664010.674 | 1858578.215 | 1298203.708 | 2456253.569 |
| 4-[3-(Trifluoromethyl)benzyl]piperidine | 906402.4017 | 959927.7275 | 855335.0519 | 920683.964 | 860832.4278 | 887284.4696 | 917926.2367 | 887632.3222 | 946468.3183 | 882053.4655 | 881563.4048 | 858912.2069 |
| PI(18:0/16:0) | 7646609.027 | 8435738.426 | 4854237.937 | 6034444.99 | 16592703.08 | 12266883.71 | 15474248.77 | 15269946.44 | 16957545.13 | 17901021.92 | 13886595.12 | 11061544.38 |
| PE O-18:2_16:0 | 1702117.961 | 582932.1537 | 2391990.942 | 1362111.919 | 1242896.106 | 4673.732637 | 1101678.456 | 751711.9596 | 1486789.939 | 1432757.905 | 1373914.071 | 1038319.117 |
| HexCer 8:0;2O/4:0 | 427627.9916 | 489180.7999 | 253700.628 | 481088.1827 | 1208507.684 | 731049.6517 | 590099.8515 | 536109.9594 | 593872.4835 | 613226.712 | 591227.6187 | 480071.9777 |

| Metabolites | PBS_1 | PBS_2 | PBS_3 | PBS_4 | BCG_1 | BCG_2 | BCG_3 | BCG_4 | BCGΔ *pe18*_1 | BCGΔ *pe18*_2 | BCGΔ *pe18*_3 | BCGΔ*pe18*_4 |
|---|---|---|---|---|---|---|---|---|---|---|---|---|
| 5-(6-Methyl-7-oxooctyl)furan-2(5H)-one | 870591.3406 | 1124330.781 | 5853276.578 | 937294.0164 | 837101.0626 | 747071.0061 | 688528.3387 | 5907499.479 | 738352.2516 | 749811.1855 | 5812579.342 | 5819186.262 |
| Leflunomide | 9078.874944 | 9047.312531 | 0 | 0 | 1230056.914 | 505711.3709 | 1590898.716 | 1064378.432 | 688543.1213 | 391391.6104 | 560057.0997 | 322759.6083 |
| 1-Myristoyl-sn-glycero-3-phosphocholine (LPC(14:0/0:0)) | 1801650.348 | 1885717.973 | 1937358.782 | 1849930.109 | 4801540.306 | 4729512.964 | 1094749.08 | 4981046.39 | 4154570.055 | 5255831.173 | 4759925.074 | 3174778.811 |
| Cer 25:1;2O/4:0 | 338765.1126 | 341502.7732 | 331704.3945 | 288838.4794 | 940527.4958 | 2062407.846 | 713124.6689 | 1403679.123 | 705490.5918 | 356179.5133 | 1119543.004 | 1038466.68 |
| 3-Deoxy-lyxo-heptulosaric acid | 3054082.94 | 3170074.389 | 2530732.791 | 1798535.492 | 921130.9712 | 1063000.295 | 1724580.697 | 2317511.262 | 1117078.671 | 1291387.992 | 1060774.529 | 1312809.624 |
| 3'- O- Acetylhamaudol | 170696.3308 | 82021.44149 | 217162.125 | 567898.9546 | 3104048.009 | 253495.8221 | 443525.8629 | 377260.2555 | 481829.8457 | 503615.458 | 821919.3825 | 406640.0423 |
| Cer 12:1;2O/18:0 | 30200.09797 | 51360.70883 | 59922.56308 | 13715.60658 | 543948.516 | 804254.1641 | 473602.2333 | 516933.3647 | 341238.2524 | 222676.029 | 487114.7218 | 628541.5083 |
| Latanoprost ethylamide | 224439.474 | 169742.0889 | 163759.3295 | 193611.359 | 938062.9361 | 1023252.86 | 1041030.604 | 625619.6641 | 737672.5043 | 781313.6763 | 1160107.131 | 762803.1149 |
| Cholest-4,6-Dien-3-One | 98315.84638 | 109188.7189 | 68063.61426 | 65860.74244 | 10423967.57 | 9879227.046 | 7820666.438 | 7613395.739 | 2491549.652 | 2622684.734 | 3237380.614 | 2439243.804 |
| 4-Bromo-1-ethyl-3-methyl-1H-pyrazole-5-carboxylic acid | 2688870.563 | 2621767.359 | 2576233.84 | 2613476.106 | 2744114.971 | 2746353.73 | 2436636.794 | 2635168.898 | 2681939.283 | 2629798.124 | 2538289.564 | 2614421.182 |

| Metabolites | PBS_1 | PBS_2 | PBS_3 | PBS_4 | BCG_1 | BCG_2 | BCG_3 | BCG_4 | BCGΔ*pe18*_1 | BCGΔ*pe18*_2 | BCGΔ*pe18*_3 | BCGΔ*pe18*_4 |
|---|---|---|---|---|---|---|---|---|---|---|---|---|
| N,N-Dimethyl-N-(4-piperidinyl)amine | 1211496.856 | 1035815.583 | 1794116.957 | 1691482.862 | 1471067.41 | 2245977.69 | 1709171.309 | 1861781.819 | 1474951.954 | 1291992.345 | 1996056.439 | 1740087.122 |
| Decuroside IV | 651403.9106 | 360243.7246 | 592703.289 | 761615.5863 | 330895.3819 | 589341.9758 | 413907.6735 | 792708.4663 | 530399.4543 | 770562.5809 | 511734.3368 | 656381.5724 |
| Ponatinib | 45870.02434 | 42972.91592 | 68090.08156 | 262878.93 | 125497.1908 | 160293.6087 | 1410739.665 | 95699.16916 | 344646.3797 | 214615.466 | 128271.2688 | 434059.0208 |
| Arabinono-1,4-lactone | 24873715.36 | 12526076.11 | 19187422.91 | 22540141.9 | 11033732.57 | 13721124.76 | 11536669.16 | 12717554.4 | 12732009.82 | 8777507.943 | 8192290.253 | 14930087.03 |
| Gabapentin related compound D | 442897.7989 | 431533.3464 | 413586.531 | 410055.9114 | 436137.804 | 449490.1135 | 433709.6025 | 449555.4608 | 425891.8012 | 407504.3482 | 441859.8772 | 433333.7534 |
| 16-Phenoxytetranor prostaglandin F2.alpha. cyclopropyl methyl amide | 317604.6681 | 340798.8172 | 347647.7067 | 282455.7193 | 342586.2909 | 317898.941 | 398457.8641 | 205103.5156 | 257007.2559 | 0 | 337844.2371 | 225420.7656 |
| 9-Hydroxy-5-(4-hydroxy-3,5-dimethoxyphenyl)-5,8,8a,9-tetrahydrofuro[3',4':6,7]naphtho[2,3-d][1,3]dioxol-6(5aH)-one | 252785.9259 | 186533.3447 | 166375.9867 | 149872.2332 | 19461814.78 | 29273422.13 | 8146824.937 | 24105665.99 | 33124739.19 | 28766554.53 | 28099300.97 | 33685749.64 |
| N-Acetylhistamine | 7686874.473 | 8117874.472 | 7505140.203 | 7642660.84 | 3366534.886 | 3536247.224 | 3213175.88 | 4235451.433 | 3660069.188 | 2452629.774 | 2632675.139 | 3221303.947 |

| Metabolites | PBS_1 | PBS_2 | PBS_3 | PBS_4 | BCG_1 | BCG_2 | BCG_3 | BCG_4 | BCGΔ*pe18*_1 | BCGΔ*pe18*_2 | BCGΔ*pe18*_3 | BCGΔ*pe18*_4 |
|---|---|---|---|---|---|---|---|---|---|---|---|---|
| N-(2,4-Dibromophenyl)-4-fluorobenzamide | 1555004.954 | 1685065.108 | 1727909.236 | 1740272.751 | 1482653.604 | 1983378.781 | 1343446.412 | 1858567.198 | 1539013.215 | 1824992.995 | 1542685.367 | 1848602.392 |
| PI 16:1_18:2 | 479099.2679 | 584928.079 | 64585.71477 | 31373.88878 | 540590.1552 | 331033.1908 | 497813.2625 | 522026.5952 | 581615.6699 | 393491.7433 | 268667.6064 | 508724.7815 |
| Cer 8:1;2O/8:0 | 2431359.845 | 1483506.391 | 1834500.479 | 1687951.364 | 1368603.357 | 1372622.21 | 1835773.253 | 1423602.928 | 885803.2203 | 995829.7918 | 1276795.83 | 1164886.356 |
| N-Acetylmethionine | 292784.5594 | 111757.7857 | 432635.7036 | 137702.127 | 671138.9122 | 1008585.867 | 2394630.839 | 1928362.01 | 1115034.228 | 570478.0421 | 1226466.881 | 1067193.854 |
| 6.beta.-Hydroxytriamcinolone acetonide | 122008.1094 | 117691.7763 | 141876.7857 | 140639.4048 | 194357.4238 | 141986.2106 | 172262.4117 | 153469.3583 | 239897.7393 | 187571.9814 | 225084.6288 | 300590.9887 |
| 1-(Butylamino)-3-phenoxy-2-propanol | 281177.2795 | 285239.2126 | 314561.3737 | 190992.7025 | 305284.2065 | 254523.1423 | 198768.1456 | 306957.3343 | 295996.6631 | 343518.2142 | 372588.4965 | 251601.7868 |
| Tributylphosphine oxide | 749749.0756 | 679687.6838 | 716263.168 | 269965.1182 | 682609.8521 | 731336.5839 | 460508.9635 | 550961.9719 | 172539.7113 | 275082.0155 | 195684.479 | 227329.3288 |
| N-Methyl-1-deoxynojirimycin | 1645848.375 | 2252265.573 | 1656772.146 | 1718614.408 | 1545838.44 | 1410144.778 | 1599982.677 | 1520913.655 | 1769274.273 | 1369761.434 | 1557731.072 | 1323552.287 |
| Mulberrofuran_M | 169994.3923 | 117950.1232 | 241286.5948 | 145845.2107 | 650970.526 | 477292.5392 | 940453.0003 | 712239.9159 | 612032.4159 | 573558.6966 | 668039.9005 | 888404.2692 |
| 1H-1,2,3-Triazole-1-octanamide, N-[1,1'-biphenyl]-2-yl-4-(3-pyridinyl)- | 6601642.968 | 7095362.24 | 5035501.672 | 5908877.086 | 7512773.961 | 7043080.515 | 7066490.576 | 6828588.068 | 6759052.162 | 8187769.5 | 6482234.957 | 4442708.903 |

| Metabolites | PBS_1 | PBS_2 | PBS_3 | PBS_4 | BCG_1 | BCG_2 | BCG_3 | BCG_4 | BCGΔ*pe18*_1 | BCGΔ*pe18*_2 | BCGΔ*pe18*_3 | BCGΔ*pe18*_4 |
|---|---|---|---|---|---|---|---|---|---|---|---|---|
| Dolasetron | 318779.018 | 50155.13585 | 304795.0997 | 102105.5654 | 321065.9476 | 486300.5255 | 2714359.61 | 1369529.005 | 769875.9201 | 438844.3654 | 689734.3193 | 219387.108 |
| Curan | 424183.4004 | 401984.1232 | 312673.9262 | 330915.4775 | 424571.881 | 350531.5425 | 403793.9917 | 449242.2009 | 417818.0465 | 401993.1968 | 401097.3406 | 260638.8502 |
| monolinolein | 3606384.246 | 2788765.721 | 2449988.091 | 3813041.516 | 17966591.43 | 14703715.42 | 8861940.146 | 18101804.28 | 14603413.25 | 5688679.967 | 6166961.886 | 9062471.149 |
| SHexCer 20:0;3O | 446906.6803 | 375810.5387 | 370133.6372 | 296476.8957 | 1060093.233 | 877021.8878 | 806018.5041 | 630999.2309 | 900587.8443 | 1020396.036 | 1053814.951 | 648300.8529 |
| (E)-3-[2-[(2S,3R,4S,5S,6R)-3,4,5-trihydroxy-6-(hydroxymethyl)oxan-2-yl]oxyphenyl]prop-2-enoic acid | 133982.655 | 182736.4102 | 230898.6158 | 336003.8525 | 196651.7478 | 153277.0687 | 229485.0573 | 258111.1451 | 235323.4268 | 144083.2989 | 106207.6289 | 171812.1433 |
| 1,8-Diazabicyclo[5.4.0]undec-7-ene | 1301033.607 | 921066.6498 | 1268269.77 | 1411835.046 | 1030215.726 | 946450.7298 | 939890.3235 | 1096996.609 | 1637216.32 | 1228762.434 | 1571428.758 | 843887.6698 |
| FA 24:3 | 76793.70169 | 62819.28014 | 158902.1067 | 95754.76006 | 139439.7498 | 221224.9639 | 126163.3069 | 125341.1442 | 76177.8726 | 88315.39716 | 133787.6356 | 71053.00394 |
| Bonactin | 139907.7654 | 166297.1845 | 444796.9941 | 204328.9403 | 260682.1433 | 235151.0058 | 321803.2819 | 278889.878 | 161372.7746 | 176615.6689 | 199824.6605 | 125059.9235 |
| 5-(2-Hydroxyethyl)-6-methyl-2-phenyl-4(3H)-pyrimidinon | 188259.616 | 130778.7826 | 58636.09366 | 55210.40104 | 160835.711 | 169352.9875 | 163136.7132 | 158352.5532 | 124165.628 | 176805.9869 | 181756.8313 | 99987.28509 |

| **Metabolites** | **PBS_1** | **PBS_2** | **PBS_3** | **PBS_4** | **BCG_1** | **BCG_2** | **BCG_3** | **BCG_4** | **BCGΔ*pe18*_1** | **BCGΔ*pe18*_2** | **BCGΔ*pe18*_3** | **BCGΔ*pe18*_4** |
|---|---|---|---|---|---|---|---|---|---|---|---|---|
| e | | | | | | | | | | | | |
| PC O-18:0 | 905370.2553 | 826493.6528 | 600471.1964 | 829420.1992 | 932950.6959 | 873722.1796 | 976270.2191 | 1091930.826 | 842970.2118 | 730250.5677 | 755898.3281 | 623657.8446 |
| 12-Hydroxystearic acid | 49072864.57 | 49992545.52 | 47968819.31 | 52079233.27 | 53954887.04 | 58120844.39 | 49498400.69 | 52837085.08 | 53738533.09 | 50792946.9 | 52904767.16 | 52676388.03 |
| PC 8:0_22:3 | 452683.5622 | 545026.2956 | 224892.5979 | 330533.8664 | 2533449.558 | 2273816.863 | 2069266.51 | 2321392.751 | 2389603.12 | 2452728.382 | 1729283.415 | 1356071.568 |
| 3-Methyl-2-(3-oxo-3H-benzo[d]isothiazol-2-yl)pentanoic acid | 3827095.162 | 676906.9852 | 3721661.056 | 3549490.337 | 2091838.207 | 1748464.123 | 18483065.45 | 18716968.21 | 5314845.099 | 1415755.759 | 3196804.818 | 2593964.938 |
| PC O-32:4 | 4722954.431 | 6368744.575 | 3793811.48 | 4126965.345 | 1092462.808 | 874435.5673 | 2794281.423 | 1434896.252 | 2038795.971 | 2418477.59 | 2164802.041 | 1159406.864 |
| DLCL 16:1_16:3 | 96641.13343 | 97481.4943 | 94687.86441 | 95500.63862 | 106130.4144 | 121390.8034 | 100962.4323 | 98960.29324 | 137768.196 | 105875.1523 | 109180.6468 | 90236.88644 |
| SM 14:3;2O/7:0 | 731663.499 | 343382.1625 | 160292.9172 | 568838.7039 | 372173.8589 | 495688.698 | 498304.2551 | 126206.143 | 246399.0994 | 511870.9534 | 262608.685 | 134645.3607 |
| DG 34:9 | 1159421.066 | 1161978.244 | 1069675.138 | 1089705.103 | 1358867.151 | 1104366.2 | 1343451.209 | 1167792.953 | 1143178.738 | 1507230.787 | 1145323.988 | 1220759.566 |
| 1-[(2-Hydroxyethyl)amino]-4-(methylamino)anthra-9,10-quinone | 1355032.515 | 258048.9364 | 857685.1456 | 955838.7081 | 684973.2199 | 1372097.057 | 22965668.99 | 2584789.326 | 3949852.578 | 1870100.49 | 1236048.868 | 1940259.785 |

| Metabolites | PBS_1 | PBS_2 | PBS_3 | PBS_4 | BCG_1 | BCG_2 | BCG_3 | BCG_4 | BCGΔ*pe18*_1 | BCGΔ*pe18*_2 | BCGΔ*pe18*_3 | BCGΔ*pe18*_4 |
|---|---|---|---|---|---|---|---|---|---|---|---|---|
| butabarbital | 1584615.736 | 808338.1877 | 1763537.3 | 993274.6683 | 14742846.72 | 10824807.36 | 12928113.71 | 10076730.37 | 7797600.999 | 6476304.197 | 9458514.436 | 11976626.95 |
| PC O-38:3 | 2064658.8 | 1268714.527 | 2838986.418 | 2240467.746 | 1328859.875 | 82082.20711 | 1632947.549 | 2775702.64 | 1907347.187 | 1575978.7 | 1057905.659 | 2660393.581 |
| Arg-His | 8540771.689 | 7443320.38 | 7840847.733 | 7537502.247 | 7093550.904 | 6263517.466 | 6623827.867 | 6115545.01 | 6213766.003 | 7146976.952 | 6700884.692 | 12094908.66 |
| gamma-Glutamyl-gamma-glutamyl-methionine | 0 | 0 | 0 | 5100.675654 | 99937.10201 | 169461.3624 | 233356.1314 | 261201.0632 | 260079.8493 | 199587.3666 | 309787.1753 | 478533.5292 |
| FA 24:5 | 1419862.207 | 1304433.391 | 1611055.513 | 1445049.304 | 1142267.345 | 2286220.446 | 1064380.463 | 1615771.741 | 1296141.042 | 745487.7931 | 1212703.597 | 1365425.383 |
| 2,4-Dihydroxybutanoic acid | 1646296.365 | 1578492.045 | 1446917.156 | 1505881.86 | 1506883.17 | 1523666.045 | 1497075.994 | 1572424.781 | 1556715.457 | 1552871.634 | 1517626.731 | 1433742.438 |
| Dimethocaine | 2950507.212 | 3149111.504 | 3006113.048 | 2957859.853 | 3312943.861 | 3138581.942 | 3125037.482 | 3033647.347 | 2770055.32 | 3445827.101 | 2876083.548 | 2956532.976 |
| LPE 18:2 | 2233602.104 | 2195010.18 | 3713120.305 | 2450990.926 | 6682160.228 | 6159976.325 | 6245809.602 | 6590619.324 | 6819839.882 | 6336324.464 | 7169522.685 | 8186529.386 |
| PC O-9:0_13:0 | 142294.754 | 102631.3609 | 229995.3355 | 149016.7581 | 609062.2727 | 629349.4648 | 536933.7806 | 421146.3275 | 476157.1589 | 528471.9319 | 540479.3025 | 160676.2667 |
| PC O-31:5 | 1112374.486 | 1325743.485 | 986115.0441 | 886601.4401 | 869197.1856 | 884231.8785 | 1007826.149 | 1010998.012 | 774870.3505 | 1104999.299 | 866140.8972 | 529239.5542 |
| 1-linoleoyl-sn-glycero-3-phospho-(1'-sn-glycerol) | 503996.1514 | 651643.1164 | 1174750.396 | 664066.9788 | 2619760.894 | 2453753.247 | 3252013.408 | 3114212.211 | 3207391.617 | 3193515.729 | 3689821.856 | 2514396.236 |

| Metabolites | PBS_1 | PBS_2 | PBS_3 | PBS_4 | BCG_1 | BCG_2 | BCG_3 | BCG_4 | BCGΔ*pe18*_1 | BCGΔ*pe18*_2 | BCGΔ*pe18*_3 | BCGΔ*pe18*_4 |
|---|---|---|---|---|---|---|---|---|---|---|---|---|
| LNAPE 16:0/N-18:2 | 226502.9068 | 169781.7237 | 301077.5403 | 176675.1105 | 945466.0975 | 1103265.425 | 956710.5134 | 947906.2382 | 698094.999 | 829803.0462 | 1505870.491 | 702201.1593 |
| [2.3-dihydroxypropoxy][2-[octadeca-9.12-dienoyloxy]-3-(octadecanoyloxy)propoxy]phosphinic acid | 471426.1187 | 661646.0231 | 735027.6455 | 468603.4045 | 414389.0994 | 139722.919 | 335038.1997 | 145818.16 | 725447.2944 | 633082.7586 | 795862.222 | 165980.8521 |
| 4(1H)-Quinolinone, 3-bromo-8-methoxy-2-methyl- | 1597725.022 | 1818559.587 | 1490472.876 | 1639335.852 | 1646814.369 | 1560468.087 | 1443490.612 | 1475418.556 | 1436658.688 | 1529816.833 | 1551430.056 | 1495154.319 |
| DGGA 16:4_20:4 | 747277.9498 | 774653.3555 | 738838.9234 | 760186.1519 | 793145.6847 | 834752.5925 | 937437.3154 | 753881.6093 | 740601.505 | 788081.9297 | 896163.1521 | 697051.7325 |
| Hex3Cer 17:3;2O | 732131.8916 | 916674.9802 | 253033.2833 | 526822.4162 | 442469.4875 | 539944.8006 | 390717.3507 | 391670.9203 | 474258.565 | 481748.6489 | 472934.8024 | 419407.3826 |
| 2-(Hexopyranosyloxy)-3-hydroxypropyl (7Z,10Z,13Z)-hexadeca-7,10,13-trienoate | 1036710.147 | 695080.4204 | 3513655.476 | 973953.9481 | 1354332.255 | 1111074.917 | 2090953.683 | 1301190.22 | 867659.9054 | 997934.579 | 1736036.194 | 461780.742 |
| 4-(Dimethylamino)pyridine | 5948742.029 | 3034711.904 | 18435302.77 | 18506094.4 | 20113859.49 | 17896933.67 | 1488350.847 | 4799975.616 | 3063579.098 | 6145264.067 | 2252819.957 | 18967231.44 |
| MG 20:2 | 301712. | 120926. | 351084. | 230203. | 628361. | 568113. | 272364. | 486597. | 266173. | 298461. | 316671. | 255303. |

| Metabolites | PBS_1 | PBS_2 | PBS_3 | PBS_4 | BCG_1 | BCG_2 | BCG_3 | BCG_4 | BCGΔ*pe18*_1 | BCGΔ*pe18*_2 | BCGΔ*pe18*_3 | BCGΔ*pe18*_4 |
|---|---|---|---|---|---|---|---|---|---|---|---|---|
| | 9133 | 6302 | 1537 | 6594 | 5137 | 2728 | 4307 | 9407 | 0472 | 3817 | 9744 | 8864 |
| 3-O-acetyloleanolic acid | 1183540.263 | 2221551.606 | 1224932.336 | 2548638.402 | 1119898.554 | 1339562.46 | 1256036.589 | 1617515.701 | 979129.8162 | 1783596.353 | 1185575.913 | 1225770.74 |
| PS 2:0_17:0 | 415129.8801 | 399356.5125 | 301714.215 | 345965.6214 | 382151.76 | 324774.6093 | 389516.4353 | 355385.0789 | 382219.056 | 354591.4701 | 354860.578 | 313976.6803 |
| .beta.-D-Glucopyranose, 6-[(2E)-3-phenyl-2-propenoate] 1-(3,4,5-trihydroxybenzoate) | 484164.5584 | 481256.5567 | 487264.1112 | 497620.3618 | 519479.8447 | 502638.8436 | 482959.5868 | 416268.1171 | 462082.4468 | 443523.9756 | 452701.524 | 450272.9337 |
| phosphatidylglycerol (1-palmitoyl, 2-lactobacilloyl) | 6734040.675 | 6092626.612 | 6953647.009 | 6353282.07 | 3976424.404 | 5796139.519 | 6234291.151 | 6445712.355 | 5843336.792 | 6066880.18 | 6502637.184 | 2054217.329 |
| Cer 16:0;2O/3:0 | 2058151.569 | 1900577.456 | 1824603.766 | 2032247.716 | 2522396.782 | 3757370.884 | 2479034.829 | 2542492.208 | 2474259.852 | 2169200.509 | 2683372.577 | 1451812.266 |
| LPC 22:6 | 3195257.95 | 3938976.963 | 4197945.765 | 3406993.69 | 5894468.904 | 6292100.167 | 5014484.205 | 5420342.8 | 6084845.476 | 4379530.615 | 5711400.013 | 5250777.383 |
| PI 2:0_17:0 | 168048.2155 | 231425.8372 | 157950.5342 | 186686.0984 | 228064.368 | 197596.2422 | 227093.308 | 205293.6875 | 214840.6707 | 192933.6852 | 215018.9693 | 185845.0231 |
| SM 25:1;2O/23:0 | 514034.6672 | 417132.9748 | 322173.4023 | 292029.4701 | 282970.144 | 0 | 236642.6661 | 271264.8257 | 109985.4525 | 198043.9293 | 306936.4682 | 144432.5302 |
| Cys-Tyr | 670821.467 | 651854.5704 | 996077.9248 | 600009.7455 | 1720776.094 | 983325.3559 | 1369318.421 | 1050151.681 | 1088666.033 | 963120.1393 | 835331.4684 | 736796.0763 |

| **Metabolites** | **PBS_1** | **PBS_2** | **PBS_3** | **PBS_4** | **BCG_1** | **BCG_2** | **BCG_3** | **BCG_4** | **BCGΔ*pe18*_1** | **BCGΔ*pe18*_2** | **BCGΔ*pe18*_3** | **BCGΔ*pe18*_4** |
|---|---|---|---|---|---|---|---|---|---|---|---|---|
| 6-Pentyl-1,4-dioxan-2-one | 1218325.793 | 1094796.576 | 768997.7415 | 2910645.792 | 872500.2091 | 907414.3309 | 3114364.132 | 694047.5382 | 1742815.36 | 1127727.536 | 774364.6864 | 4032656.988 |
| PG O-17:2_18:1 | 121292.2164 | 200937.9042 | 160019.1332 | 238417.1816 | 402686.8931 | 468104.344 | 450700.9807 | 392326.1398 | 466337.3059 | 522793.8353 | 110266.5146 | 427231.3705 |
| 3-Hydroxybenzoic acid | 1058422.224 | 1107447.336 | 1090578.333 | 1063243.016 | 2389073.023 | 2539454.972 | 3849955.968 | 2474850.198 | 2515515.359 | 2681712.898 | 2587966.958 | 2785235.607 |
| 2-Hydroxypyridine | 1250533.231 | 1473387.228 | 1448939.095 | 1306007.258 | 1155805.012 | 1158994.787 | 1130555.656 | 1306086.859 | 1258252.246 | 1257035.067 | 1430089.913 | 1212939.58 |
| Varenicline | 16834817.89 | 2250217.416 | 3419908.354 | 3519517.017 | 4861417.312 | 1092551.83 | 1888710.413 | 77449582.95 | 11226131.48 | 3468183.018 | 7511583.275 | 4855880.762 |
| 1-O-Hexadecyl-2-O-(4Z,7Z,10Z,13Z,16Z,19Z-docosahexaenoyl)-sn-glyceryl-3-phosphorylcholine | 122458783 | 150246116.5 | 48303225.06 | 81526995.23 | 42126999.55 | 33341334.65 | 47015765.07 | 57582538.94 | 53665860.24 | 62912898.93 | 37573401.6 | 36677236.88 |
| Benzyldimethylstearylammonium cation | 9196967.011 | 189408536.4 | 10230952.03 | 17491744.28 | 3020243.075 | 3767091.329 | 6887400.888 | 69801983.51 | 3759988.084 | 16461101.21 | 8823625.655 | 4946534.939 |
| S-Ethyl-N-[4-(trifluoromethyl)phenyl]isothiourea | 3610545.746 | 3130691.014 | 3158056.668 | 4173878.27 | 3096207.895 | 3430825.758 | 3250187.749 | 3293087.448 | 3180081.081 | 3704729.391 | 2851702.272 | 3557922.854 |
| N5-(1-Iminoethyl)-L-ornithine | 1249527.369 | 1089724.067 | 1981395.818 | 1633321.522 | 961952.6387 | 1415103.737 | 1052889.957 | 947307.5661 | 1089770.005 | 640924.0606 | 772854.4824 | 1427133.985 |

| **Metabolites** | **PBS_1** | **PBS_2** | **PBS_3** | **PBS_4** | **BCG_1** | **BCG_2** | **BCG_3** | **BCG_4** | **BCGΔ *pe18*_1** | **BCGΔ *pe18*_2** | **BCGΔ *pe18*_3** | **BCGΔ*pe18*_4** |
|---|---|---|---|---|---|---|---|---|---|---|---|---|
| (3beta)-lup-20(29)-ene-3,28-diol | 1043356.638 | 1040430.209 | 851938.3197 | 1088486.677 | 645269.0834 | 902941.6753 | 435159.9194 | 947794.4321 | 1091122.548 | 1091816.944 | 811636.5405 | 755807.5525 |
| 1-Oleoyl-2-palmitoyl-sn-glycero-3-phosphocholine | 5294133.415 | 15054268.26 | 3653069.615 | 18435372.67 | 34824115.5 | 9808756.035 | 13752712.6 | 46230737.14 | 11989847.06 | 53715982.14 | 12351361.46 | 5816972.85 |
| Macamide B | 5044629.784 | 5236282.435 | 4529319.155 | 4702946.284 | 5020197.079 | 4895136.949 | 4942930.578 | 4703992.147 | 4999235.257 | 4819682.09 | 4995530.642 | 4330235.083 |
| N-(9Z-Octadecenoyl)-4E,14Z-sphingadienine-1-phosphocholine | 1809335.043 | 1121561.585 | 1232575.123 | 707647.3568 | 3313660.397 | 1917230.133 | 4167425.623 | 3036534.861 | 3717921.715 | 4060614.576 | 3134752.238 | 1652598.526 |
| LPG O-22:1 | 1190837.001 | 1327415.391 | 1180533.834 | 1207599.138 | 1215618.368 | 1382892.219 | 1176258.252 | 1156251.148 | 1255540.744 | 1179950.965 | 1189724.811 | 1042442.867 |
| 2-Aminopurine | 3376186.606 | 1441135.2 | 2536541.14 | 1510836.882 | 3366873.197 | 3041130.619 | 5772137.852 | 3226362.601 | 2887028.465 | 2568360.88 | 3328038.748 | 2431704.92 |
| Montelukast-1,2-diol | 280982.5708 | 239958.13 | 297697.157 | 214076.5699 | 241129.2593 | 214992.497 | 102325.7242 | 214914.7695 | 121592.2636 | 276162.681 | 286405.8757 | 258694.351 |
| 22-Hydroxymifepristone | 2068731.652 | 230973.8184 | 1199623.819 | 5517588.562 | 559336.3832 | 2371824.44 | 709490.8499 | 909242.573 | 1434783.024 | 1184896.105 | 2339067.083 | 318041.5964 |
| PEtOH 22:2_20:4 | 234857.8024 | 184141.8927 | 209081.1924 | 674916.4242 | 175122.7122 | 203019.5957 | 423731.8032 | 1549836.389 | 195354.5666 | 198030.6664 | 95499.06019 | 1606965.924 |
| alpha,alpha'-Trehalose 6-phosphate | 0 | 9119.974841 | 14642.16993 | 0 | 344719.6393 | 260140.4594 | 270949.5628 | 246540.5776 | 225624.405 | 264296.2166 | 185273.2572 | 309044.6991 |

| Metabolites | PBS_1 | PBS_2 | PBS_3 | PBS_4 | BCG_1 | BCG_2 | BCG_3 | BCG_4 | BCGΔ*pe18*_1 | BCGΔ*pe18*_2 | BCGΔ*pe18*_3 | BCGΔ*pe18*_4 |
|---|---|---|---|---|---|---|---|---|---|---|---|---|
| (1R,5S)-9-Methyl-9-azabicyclo[3.3.1]nonan-3-amine | 2421933.885 | 8026540.682 | 9783590.555 | 8069053.412 | 2221772.323 | 4206125.128 | 3952073.327 | 1901754.257 | 3005809.275 | 2498702.081 | 1895306.325 | 9166080.577 |
| Cer 16:0;2O/16:0 | 485488.749 | 476440.1232 | 571169.6658 | 585208.4311 | 190626.3315 | 261226.2408 | 259532.4526 | 254891.2478 | 254015.8324 | 330246.6339 | 339150.4896 | 239859.6131 |
| (3E)-10b-(2-Methylbut-3-en-2-yl)-3-((5-(2-methylbut-3-en-2-yl)-1H-imidazol-4-yl)methylidene)-6,10b,11,11a-tetrahydro-2H-pyrazino[1',2':1,5]pyrrolo[2,3-b]indole-1,4(3H,5aH)-dione | 100636.7867 | 77540.65168 | 114706.4198 | 98818.06554 | 561259.0905 | 458536.0607 | 287741.9303 | 366544.0237 | 127960.7106 | 99212.45146 | 130234.6839 | 152748.4232 |
| Keracyanin Chloride | 1118871.337 | 871096.7097 | 1783590.413 | 925818.4623 | 976006.5652 | 1301493.626 | 1228018.428 | 1101202.012 | 845107.1809 | 1011560.355 | 948812.7852 | 981479.879 |
| Albuterol | 616707.8161 | 742226.9199 | 458121.1 | 5340273.491 | 456155.9751 | 572132.7438 | 5191152.187 | 345453.8517 | 744083.3168 | 657038.9733 | 385595.6142 | 9334019.431 |
| SHexCer 34:2;3O | 2657847.053 | 2965954.274 | 795632.6007 | 1520124.292 | 12927837.9 | 4227218.135 | 9722101.746 | 7385086.167 | 9717275.897 | 14427743.15 | 8471022.9 | 4694311.155 |
| 17.beta.-Dihydroequilenin | 1323574.03 | 1439259.758 | 1189346.704 | 1156957.982 | 1316158.909 | 1407511.747 | 1846820.252 | 1351173.437 | 1409205.098 | 1374615.561 | 1293378.291 | 1194592.834 |

| Metabolites | PBS_1 | PBS_2 | PBS_3 | PBS_4 | BCG_1 | BCG_2 | BCG_3 | BCG_4 | BCGΔ *pe18*_1 | BCGΔ *pe18*_2 | BCGΔ *pe18*_3 | BCGΔ*pe18*_4 |
|---|---|---|---|---|---|---|---|---|---|---|---|---|
| 5-Ethyl-7-hydroxy-2-methylpyrazolo[1,5-a]pyrimidine-3-carbonitrile | 237376.3439 | 288706.7773 | 338743.5615 | 294854.2975 | 7741436.56 | 5758749.015 | 5270326.455 | 4463673.783 | 1727356.173 | 1561018.279 | 1807794.987 | 2021584.752 |
| PE O-13:1_4:0 | 5738043.681 | 1461069.031 | 186977.7649 | 1571089.481 | 1200988.698 | 529337.6442 | 2062731.278 | 123778.8422 | 1081263.307 | 665156.0816 | 1625652.405 | 300823.5631 |
| Cer 17:0;3O/42:0;(2OH) | 59504.89499 | 104315.6189 | 66139.88919 | 85984.01948 | 172033.7007 | 104135.9152 | 215642.7766 | 166834.7293 | 115125.1985 | 150835.2734 | 63334.54173 | 68883.17584 |
| (5.Xi.,9.Xi.,18.Xi.)-3-Oxolup-20(29)-ene-27,28-dioic acid | 343091.5894 | 253369.3056 | 556580.6454 | 492847.5696 | 622267.3262 | 382503.4753 | 520480.5703 | 374597.7267 | 425842.9795 | 471735.9177 | 616677.4582 | 343480.7272 |
| Galeopsin | 426032.0698 | 651593.136 | 1322459.294 | 485112.7043 | 1540210.381 | 1492434.37 | 1402122.735 | 945725.7413 | 1038058.721 | 715833.6987 | 940801.8095 | 1089694.502 |
| SM 34:0;2O/6:0 | 36085.37478 | 563281.2153 | 488057.412 | 583576.0095 | 404455.6051 | 1039629.365 | 787491.6206 | 373620.2822 | 283563.7461 | 366871.9835 | 408091.9859 | 1004320.76 |
| 4-Hydroxy-6-methyl-2-oxo-1,2-dihydropyridine-3-carboxamide | 1415340.234 | 1140566.143 | 3735741.201 | 1898983.235 | 1189414.519 | 2141460.936 | 2886495.567 | 3669321.596 | 1902785.238 | 999372.0857 | 2677843.592 | 3010605.015 |
| PG 18:1_26:2 | 0 | 0 | 0 | 8653.050332 | 2578673.346 | 3108305.304 | 2251131.65 | 2543686.606 | 4631039.496 | 3142047.653 | 3458058.254 | 2882442.352 |

| Metabolites | PBS_1 | PBS_2 | PBS_3 | PBS_4 | BCG_1 | BCG_2 | BCG_3 | BCG_4 | BCGΔ*pe18*_1 | BCGΔ*pe18*_2 | BCGΔ*pe18*_3 | BCGΔ*pe18*_4 |
|---|---|---|---|---|---|---|---|---|---|---|---|---|
| 1,4a-Dimethyl-4,4a,5,6,7,8-hexahydro-2(3H)-naphthalenone | 3977762.131 | 4334996.544 | 3177266.668 | 3352870.582 | 4121648.413 | 3891729.181 | 4120059.598 | 4092102.705 | 4323805.589 | 3851578.031 | 4627646.084 | 4120347.143 |
| LPS 22:6 | 8695530.91 | 7522876.639 | 8819764.96 | 7959778.66 | 13226003.35 | 14414571.38 | 14102097.58 | 14463161.08 | 21097298.99 | 14230245.86 | 16267680.49 | 16284619.09 |
| LPE O-18:3 | 1301100.121 | 1073229.714 | 1243862.659 | 1101414.656 | 1188565.902 | 1294047.312 | 1448541.994 | 1340033.207 | 1695919.98 | 1599089.016 | 1654528.67 | 947737.6035 |
| Cer 17:0;2O/16:2 | 19830.97423 | 56166.90039 | 23779.55913 | 74820.53917 | 242806.9312 | 377249.8749 | 323187.293 | 297424.8695 | 271665.1412 | 164741.7989 | 304609.9359 | 130389.8392 |
| 3,4-Methylenedioxy-.alpha.-pyrrolidinobutyrophenone | 16687.38499 | 235321.8494 | 221075.0278 | 235321.8692 | 151667.5315 | 219772.5027 | 231975.1923 | 14564.30284 | 15490.92068 | 243965.9514 | 217725.5953 | 215644.5584 |
| NAGlySer 19:2/26:4 | 0 | 0 | 0 | 0 | 652909.9388 | 1285342.111 | 1418033.526 | 1629412.944 | 862426.8775 | 434571.1607 | 993436.3109 | 279580.413 |
| Solasodiene (not validated) | 4891264.164 | 4999870.961 | 4670453.872 | 4304637.155 | 4842421.035 | 4747501.771 | 4607316.491 | 5020110.104 | 4489768.014 | 5234508.511 | 4717567.023 | 4742688.833 |
| LPS 15:0 | 427247.8257 | 452467.9489 | 237933.0842 | 345730.1221 | 1848656.181 | 1794377.268 | 1870852.975 | 1669944.374 | 1306522.859 | 1388020.251 | 1722269.671 | 1603836.323 |
| M437T229_2 | 76054.6228 | 50487.56188 | 138846.1509 | 98417.65055 | 227296.9959 | 188486.4072 | 183286.4507 | 219908.5265 | 139327.2886 | 145983.965 | 156342.1205 | 100165.5225 |
| Cohibin_D | 5614695.803 | 4994042.481 | 4383270.908 | 3858599.233 | 9148423.967 | 9588660.142 | 13473181.97 | 10967490.3 | 8579526.38 | 11271872.1 | 11056324.59 | 5496985.345 |
| Ciclopirox | 90666.3 | 721528. | 154007 | 118650 | 280474. | 16997.6 | 186599. | 58973.4 | 123765. | 265805. | 493854. | 261954 |

| Metabolites | PBS_1 | PBS_2 | PBS_3 | PBS_4 | BCG_1 | BCG_2 | BCG_3 | BCG_4 | BCGΔ*pe18*_1 | BCGΔ*pe18*_2 | BCGΔ*pe18*_3 | BCGΔ*pe18*_4 |
|---|---|---|---|---|---|---|---|---|---|---|---|---|
| | 5759 | 7555 | 7.33 | 8.408 | 343 | 8576 | 2735 | 3081 | 4434 | 7949 | 1669 | 1.84 |
| Dethiobiotin | 183057.636 | 179376.4652 | 234256.6812 | 177575.6079 | 356712.8493 | 868804.8279 | 644708.5889 | 724180.5257 | 1188161.156 | 544378.2874 | 678779.9555 | 154337.1033 |
| Cyclo(isoleucylprolyl) | 1487363.339 | 1524311.643 | 1641411.822 | 1441998.83 | 1411149.287 | 1603303.585 | 1490345.027 | 1538171.709 | 1457560.121 | 1262620.959 | 1295659.266 | 1517734.673 |
| GDP-fucose | 654592.1303 | 634773.7171 | 743232.6171 | 588152.5607 | 950345.741 | 776592.5807 | 910004.5068 | 902283.1934 | 827795.2874 | 820448.9329 | 871821.5425 | 1021883.969 |
| 2,4-Dihydroxybenzophenone | 190506.1021 | 45938.23814 | 69769.02287 | 86808.73607 | 985730.4998 | 1333041.311 | 1201240.937 | 1229864.162 | 1097981.94 | 1244520.189 | 1336017.835 | 1670814.551 |
| 3-Dodecyloxypropanoic acid | 665579.8221 | 381643.6735 | 574799.2659 | 455502.1343 | 970310.6862 | 1019116.833 | 822691.7113 | 933110.4261 | 1158903.308 | 735161.6899 | 802963.9901 | 1089437.967 |
| 2-[2-(2-Quinolinyl)ethenyl]phenol | 33238.95841 | 372545.0943 | 815284.7511 | 660877.6056 | 21615.85109 | 0 | 217599.209 | 0 | 44717.69752 | 0 | 17024.26392 | 399016.3311 |
| DLCL 20:4_24:5 | 40879.63715 | 56674.156 | 34181.35055 | 50135.84325 | 111349.761 | 62337.04142 | 78302.78854 | 66203.13884 | 77997.81211 | 97820.84752 | 108350.8666 | 76463.01891 |
| PC 18:0_20:4 | 1435465.847 | 1101965.36 | 1335741.744 | 1800362.562 | 1368475.008 | 0 | 1564296.26 | 2866643.77 | 2234574.286 | 1840675.866 | 840637.9891 | 2539059.101 |
| PC 50:11 | 3026876.663 | 3483478.526 | 2657007.848 | 3081950.018 | 3411607.934 | 3279933.41 | 3318015.669 | 3585573.941 | 3200755.547 | 3367823.84 | 3896971.96 | 3098366.387 |
| 4-Phosphonobenzoic acid | 1395932.725 | 1290144.469 | 2360821.347 | 1690505.365 | 1988336.553 | 2038139.292 | 1640503.744 | 1804808.671 | 1466383.737 | 901531.4739 | 1419121.203 | 1495556.693 |
| PE 26:5 | 11281745.15 | 9547200.803 | 19363246 | 12337225.4 | 7830339.372 | 6183998.825 | 9619369.809 | 7105547.868 | 3404381.961 | 3667648.904 | 4538159.513 | 2845708.835 |
| PC 18:4_18:4 | 207240. | 334323. | 474350. | 185829. | 158142. | 478781. | 247341. | 308756. | 276312. | 273333. | 246461. | 299705. |

| Metabolites | PBS_1 | PBS_2 | PBS_3 | PBS_4 | BCG_1 | BCG_2 | BCG_3 | BCG_4 | BCGΔ*pe18*_1 | BCGΔ*pe18*_2 | BCGΔ*pe18*_3 | BCGΔ*pe18*_4 |
|---|---|---|---|---|---|---|---|---|---|---|---|---|
| | 7759 | 2114 | 0676 | 0713 | 4054 | 3973 | 2583 | 8623 | 5153 | 8357 | 6709 | 2842 |
| (4OH,8Z,t18:1)_sphingosine | 5231195.846 | 4158207.278 | 8672463.07 | 4406531.884 | 5289357.046 | 7669354.599 | 9286400.783 | 7604173.617 | 10427055.13 | 3369790.597 | 14780835.27 | 11786229.37 |
| Ligustrazine | 11390043.73 | 4191130.38 | 4450862.951 | 4379259.44 | 4541708.139 | 4369560.588 | 4829544.378 | 4485730.29 | 4744428.319 | 4475360.95 | 4087190.617 | 4437878.627 |
| cis-3-Hexenyl_pyruvate | 2988457.552 | 2651942.344 | 2670371.461 | 5729167.875 | 2846981.411 | 2801460.331 | 7066794.04 | 3842209.697 | 3959628.785 | 3294021.217 | 2232637.672 | 5325954.717 |
| BENZHEXOL | 374593.1017 | 358457.0807 | 398803.5534 | 359741.9907 | 377050.2717 | 420636.5396 | 411624.2768 | 378219.1473 | 421756.5167 | 436829.9269 | 423926.7321 | 375851.673 |
| N-Lactoylphenylalanine | 311550.4685 | 223918.9031 | 493899.0505 | 316029.3052 | 502239.72 | 439283.1228 | 708786.2337 | 460894.516 | 562792.3868 | 618549.4927 | 678948.6835 | 540071.7581 |
| 13-Hydroperoxyoctadeca-9,11-dienoic acid | 636694.2269 | 736608.5253 | 963098.997 | 874126.7156 | 1251599.462 | 1323413.141 | 1321985.687 | 1156870.137 | 1049249.965 | 696196.2062 | 1120725.748 | 1534501.631 |
| LPC O-15:1 | 148766.9278 | 363994.0353 | 343575.8569 | 436798.9033 | 1156952.739 | 911284.3243 | 733097.2286 | 868257.9457 | 888596.848 | 571200.3094 | 649655.4042 | 643040.9272 |
| PE-Cer 12:1;2O/16:3 | 792907.1727 | 812257.4099 | 790722.165 | 897417.6813 | 1536346.423 | 1604132.838 | 1291334.547 | 1220660.287 | 765904.9273 | 905705.5729 | 556479.3918 | 480892.7144 |
| M525T340 | 273250.8256 | 293338.3243 | 277905.9931 | 272777.5257 | 283610.2351 | 263322.7582 | 314948.1263 | 274004.1643 | 260573.8405 | 254901.3039 | 262871.2986 | 281753.8473 |
| Butanoic acid | 859276.3917 | 857163.8063 | 1077766.208 | 860577.2368 | 964797.3836 | 1038648.926 | 1029982.347 | 1244069.369 | 912735.1094 | 891114.5413 | 897050.1107 | 1092431.665 |
| LDGTS 20:3 | 579517.1396 | 1137816.164 | 430922.0912 | 1125159.957 | 470849.1816 | 746468.1006 | 1066229.709 | 1025164.924 | 371316.9898 | 195384.6254 | 258847.1358 | 416847.8793 |

| Metabolites | PBS_1 | PBS_2 | PBS_3 | PBS_4 | BCG_1 | BCG_2 | BCG_3 | BCG_4 | BCGΔ*pe18*_1 | BCGΔ*pe18*_2 | BCGΔ*pe18*_3 | BCGΔ*pe18*_4 |
|---|---|---|---|---|---|---|---|---|---|---|---|---|
| PC(20:2(11Z,14Z)/15:0) | 1860361.797 | 3314577.652 | 6341366.687 | 6431289.505 | 54568023.2 | 14489735.47 | 37865405.76 | 16903030.13 | 48064522.74 | 18049271.21 | 38737184.98 | 31858647.67 |
| PE(P-38:6) | 133041693.7 | 173265709.2 | 116096117.8 | 128227076.9 | 201194815.5 | 155277175.7 | 185950295.8 | 186387468.4 | 190333673.5 | 175346622.5 | 152008657.2 | 103777079 |
| 1-(2,4-Dihydroxyphenyl)-2-(4-nitrophenyl)ethanone | 331452.712 | 1335633.11 | 126845.2017 | 935614.4194 | 2338497.217 | 785157.7428 | 89760.68896 | 59131.21962 | 465345.9646 | 1168358.414 | 589515.6005 | 215460.843 |
| name | 23843.18514 | 15794.68356 | 31119.14607 | 25189.18805 | 148945.9709 | 139458.2038 | 113678.7683 | 120984.6011 | 76966.42975 | 122276.2583 | 114235.892 | 90935.30287 |
| GDP-3,6-dideoxy-galactose | 0 | 4055.893492 | 10788.10691 | 10200.15268 | 185289.1701 | 128121.0937 | 374089.3992 | 171908.5181 | 81501.18299 | 49991.35382 | 135457.0958 | 147743.7359 |
| M798T462 | 1107812.041 | 755327.1208 | 1069825.809 | 1229343.882 | 1164100.237 | 1104322.914 | 1285431.701 | 1201876.082 | 971208.1385 | 1069410.484 | 1169107.88 | 960941.8919 |
| Huperzine B | 0 | 528596.9929 | 1092759.563 | 1831023.669 | 0 | 36291.62566 | 240201.2355 | 0 | 71100.38483 | 24221.04584 | 0 | 1277674.765 |
| 2,2-Dimethylsuccinic acid | 2193536.492 | 1855991.881 | 1567567.034 | 2039436.693 | 12375370.89 | 6947978.579 | 8997968.947 | 9689121.685 | 5787558.206 | 4106914.525 | 5915704.994 | 5047081.081 |
| MGDG O-22:3_16:1 | 5074414.774 | 4423029.994 | 2019906.196 | 3911742.926 | 1259803.418 | 1003843.583 | 933711.4476 | 1304747.822 | 1579662.879 | 1218927.233 | 920496.1325 | 1112361.085 |
| (2S,4S)-4-Cyclohexylpyrrolidine-2-carboxylic acid | 1298349.882 | 1951286.844 | 974439.1988 | 917170.2581 | 1858190.967 | 1579437.354 | 4534216.208 | 2013032.319 | 1962133.068 | 1523655.884 | 1767008.422 | 1376012.731 |
| tetracosatetraenoate | 1916213.246 | 2028639.154 | 4645287.41 | 2150687.237 | 820100.9195 | 1883104.32 | 1453796.703 | 1521908.708 | 1409882.295 | 1323571.574 | 1685604.564 | 650805.748 |

| **Metabolites** | **PBS_1** | **PBS_2** | **PBS_3** | **PBS_4** | **BCG_1** | **BCG_2** | **BCG_3** | **BCG_4** | **BCGΔ*pe18*_1** | **BCGΔ*pe18*_2** | **BCGΔ*pe18*_3** | **BCGΔ*pe18*_4** |
|---|---|---|---|---|---|---|---|---|---|---|---|---|
| Peimine | 261941.5402 | 239738.3133 | 213477.3858 | 218579.4703 | 388106.9742 | 377190.2929 | 296328.2587 | 387426.3157 | 335617.9636 | 226869.3026 | 222792.9555 | 283806.9706 |
| PEtOH 21:0_20:3 | 201129.9823 | 233874.4878 | 220917.0198 | 156604.9645 | 329568.0128 | 277241.4141 | 302740.4296 | 279159.4327 | 265629.5592 | 254117.754 | 311607.1343 | 195678.2801 |
| N-Cyclohexanecarbonyltetradecylamine | 1288243.088 | 1348858.696 | 880349.8325 | 863202.9462 | 798740.3859 | 2704247.336 | 1399622.06 | 1191060.392 | 1479050.219 | 897381.8117 | 1326326.528 | 1004623.876 |
| 3-(4-Methyl-3-pentenyl)thiophene | 920302.9646 | 664211.4949 | 1310794.52 | 1074441.925 | 1815827.721 | 1484893.408 | 1834490.361 | 1547935.874 | 1666802.482 | 982938.3418 | 1276990.419 | 1741219.465 |
| Tazarotenic acid sulfoxide | 1102102.337 | 797592.8028 | 1176054.705 | 933755.1627 | 3085523.611 | 1631455.426 | 2519345.376 | 2140892.175 | 2435387.882 | 2076078.428 | 1978682.869 | 1625700.734 |
| SMGDG O-9:0_7:0 | 810949.2316 | 749004.8621 | 924479.1914 | 842523.6269 | 1070137.527 | 1017840.406 | 733773.6422 | 921513.8922 | 1684175.036 | 976637.0561 | 1444427.468 | 947360.8032 |
| 4'-O-Methylglucoliquiritigenin | 1090228.311 | 975369.2585 | 819645.7199 | 868655.7212 | 803140.5277 | 982523.0055 | 846105.1886 | 1066351.818 | 946493.1515 | 864028.3674 | 966911.4282 | 854430.9243 |
| PG 3:0_26:1 | 1974942.682 | 1369197.427 | 868202.2391 | 2078361.353 | 1711151.288 | 1368002.692 | 1187322.834 | 1413110.769 | 1647257.381 | 1693992.53 | 1365380.525 | 815090.917 |
| Val-Trp | 434039.7347 | 392487.4041 | 452761.0328 | 356973.78 | 365844.7652 | 255493.3542 | 353998.7933 | 332826.6812 | 341803.4361 | 334905.4602 | 412820.4228 | 254111.0073 |
| 4-(3-Bromopyrazolo[1,5-a]pyrimidin-6-yl)phenol | 648368.1061 | 557492.7988 | 778310.6392 | 571694.7405 | 1327601.655 | 1930369.616 | 1332168.743 | 1742060.235 | 1604298.297 | 1541457.487 | 1597050.005 | 1508229.535 |
| 3-(2-Bromophenyl)-1H-pyrazol-5-amin | 910118.3742 | 681625.7761 | 1303297.163 | 972677.7926 | 4464382.078 | 2992107.588 | 4582739.634 | 3483688.479 | 2863678.352 | 3680516.912 | 3712063.958 | 2755995.855 |

| **Metabolites** | **PBS_1** | **PBS_2** | **PBS_3** | **PBS_4** | **BCG_1** | **BCG_2** | **BCG_3** | **BCG_4** | **BCGΔ*pe18*_1** | **BCGΔ*pe18*_2** | **BCGΔ*pe18*_3** | **BCGΔ*pe18*_4** |
|---|---|---|---|---|---|---|---|---|---|---|---|---|
| e | | | | | | | | | | | | |
| DGDG O-16:4_4:0 | 1117226.076 | 1282223.763 | 796152.5936 | 1090647.866 | 1004451.983 | 738063.1163 | 719591.1306 | 882447.9519 | 1120071.596 | 1069280.257 | 1104175.779 | 967348.8469 |
| (+)-Cloprostenol methylamide | 1049030.669 | 594620.2787 | 969142.1523 | 662715.2696 | 654106.4225 | 1300802.712 | 601507.8008 | 668072.3753 | 908745.3194 | 579740.3196 | 960534.4984 | 672279.9695 |
| Kukoamine B | 10777.44531 | 4739.391639 | 11190.33313 | 6520.336634 | 21995.66527 | 5230.808422 | 0 | 6756.597008 | 18690.01749 | 54496.74111 | 15354.30987 | 0 |
| PG 18:2_16:3 | 44458.01981 | 69590.31382 | 34564.7918 | 31141.8807 | 541036.9938 | 246932.3124 | 431708.5312 | 510288.283 | 1068079.034 | 739074.5096 | 344130.5746 | 480377.8059 |
| Hupehenine | 106327.1987 | 111693.207 | 226665.4365 | 159901.7891 | 363148.0256 | 325505.1753 | 279187.2232 | 456376.0158 | 435918.1162 | 16351.45756 | 200727.6932 | 317175.9985 |
| 1-(1,3-Dimethyl-1H-pyrazol-4-yl)ethanone | 1893269.519 | 1476214.637 | 1621049.31 | 1319374.74 | 992935.5858 | 3010172.459 | 1104011.316 | 786507.3307 | 1138306.755 | 1133227.368 | 1581332.613 | 1506907.344 |
| Cer 13:2;2O/22:5 | 124469.4437 | 150514.3401 | 153132.4846 | 149973.0651 | 135291.6555 | 289333.6741 | 198432.3314 | 165570.6794 | 225065.3095 | 137684.1644 | 234267.8864 | 62291.97579 |
| Boldenone cypionate | 6495476.086 | 5829909.993 | 9142046.698 | 6964226.506 | 10511887.24 | 17246351.73 | 12118121.67 | 9861880.708 | 6496787.944 | 7072193.932 | 11570178.53 | 7291963.945 |
| 3-Cyano-7-methoxycoumarin | 1395654.272 | 1097718.114 | 1323897.416 | 1569489.279 | 1556144.295 | 1684170.117 | 1146462.408 | 1534355.98 | 1290263.846 | 507440.9821 | 538694.8084 | 1330622.627 |
| Caffeic acid | 1107689.386 | 1203987.668 | 1097162.794 | 1175641.078 | 1281384.394 | 1277743.538 | 1271396.412 | 1180855.525 | 1242494.665 | 1270047.622 | 1263269.109 | 1431388.312 |
| N-oleoyl-aspartate | 964879. | 112207 | 392375. | 623323. | 333806. | 347635. | 363739. | 358927. | 492848. | 552780. | 423209. | 272797. |

| Metabolites | PBS_1 | PBS_2 | PBS_3 | PBS_4 | BCG_1 | BCG_2 | BCG_3 | BCG_4 | BCGΔ*pe18*_1 | BCGΔ*pe18*_2 | BCGΔ*pe18*_3 | BCGΔ*pe18*_4 |
|---|---|---|---|---|---|---|---|---|---|---|---|---|
| | 9539 | 5.308 | 3697 | 946 | 9026 | 9928 | 8262 | 8194 | 2265 | 5412 | 266 | 9814 |
| Thiopropionic acid, S-ethyl ester | 970500.6144 | 789204.3344 | 790683.1366 | 799000.6536 | 927674.7436 | 150558.2842 | 713960.0963 | 848939.8079 | 817600.6793 | 802607.956 | 667396.3923 | 809389.6759 |
| 1-22:0-2-18:2-phosphatidylserine | 325220.088 | 233514.6303 | 193090.477 | 245108.2297 | 55353.93691 | 679961.0199 | 50908.44069 | 503590.6964 | 425398.2765 | 412507.623 | 553017.0102 | 453078.8438 |
| Vitamin K1 2,3-epoxide | 316413.7003 | 358494.2203 | 236135.5373 | 321728.6353 | 453402.3608 | 277421.3866 | 273350.4396 | 306874.4471 | 296222.5544 | 250067.6121 | 293503.7149 | 293089.5956 |
| Norbuprenorphine glucuronide | 166478.2015 | 116548.7863 | 170649.2171 | 120632.2835 | 212275.7776 | 202607.6838 | 377164.5003 | 215200.3667 | 149576.2305 | 188642.4978 | 198848.8247 | 122803.6322 |
| LDGTS 20:4 | 2041607.484 | 2264491.168 | 2090576.803 | 2064489.746 | 1857394.389 | 2558315.94 | 1716317.876 | 1800716.581 | 1782982.325 | 1990808.221 | 2087308.485 | 2122150.319 |
| 2S-Amino-4E-pentadecene-1,3R-diol | 8725153.193 | 8964672.282 | 8417802.295 | 8329242.261 | 8799270.127 | 8746398.941 | 8831831.423 | 9074993.05 | 8533712.107 | 8572323.548 | 8741327.976 | 7631135.077 |
| 2-Butyl-3-methylpyrazine | 802840.662 | 817641.3163 | 825109.6547 | 795920.5253 | 856823.6376 | 776828.9257 | 798304.2831 | 753977.0154 | 787094.4664 | 808883.751 | 793232.2119 | 938950.8132 |
| Mangiferdesmethylursanone | 1670105.678 | 1357246.712 | 844222.5608 | 1170606.111 | 1508900.294 | 1252748.459 | 1838197.157 | 1425354.422 | 1733872.087 | 1688260.771 | 1273535.816 | 800671.778 |
| Menadione (Vitamin K3) | 605823.8292 | 519634.6825 | 634341.6138 | 509691.0974 | 502545.3517 | 603108.8693 | 608032.9673 | 545828.7472 | 563858.2444 | 490054.9567 | 564894.5853 | 536522.714 |
| M309T422_1 | 1605216.594 | 1351118.82 | 3223019.093 | 1134204.41 | 1121763.496 | 1262743.274 | 3053595.242 | 1136265.76 | 1637271.513 | 3464572.912 | 3436842.247 | 1431261.667 |
| 5'-Methoxycurcumin | 157289.3281 | 240099.9162 | 148049.5754 | 119133.496 | 142585.6811 | 216114.8582 | 133475.244 | 133784.0921 | 167610.0504 | 171621.3233 | 157307.7752 | 180705.3294 |
| SM 21:3;2O/21:0 | 512248. | 434501. | 463898. | 107057 | 413265. | 731858. | 904382. | 108175 | 507531. | 300772. | 0 | 154777 |

| Metabolites | PBS_1 | PBS_2 | PBS_3 | PBS_4 | BCG_1 | BCG_2 | BCG_3 | BCG_4 | BCGΔ*pe18*_1 | BCGΔ*pe18*_2 | BCGΔ*pe18*_3 | BCGΔ*pe18*_4 |
|---|---|---|---|---|---|---|---|---|---|---|---|---|
| | 5634 | 5464 | 2174 | 1.842 | 8525 | 1068 | 5447 | 1.118 | 8441 | 0997 | | 5.671 |
| 4-(Dimethylamino) phenylalanine | 628371.2243 | 800712.5671 | 839150.1035 | 1083030.769 | 597383.8565 | 569962.6351 | 911186.0893 | 801590.6368 | 714532.7946 | 814389.6735 | 822642.743 | 1603162.191 |
| Ile-Ala | 2984514.528 | 2948183.476 | 4185084.014 | 2695938.319 | 3270266.908 | 2644208.072 | 3359708.432 | 3189560.802 | 2462470.314 | 2253228.266 | 2872243.668 | 2630836.869 |
| (S)-3,4-Dihydroxybutyric acid (lithium hydrate) | 1559496.99 | 1377184.57 | 1301274.387 | 1641271.523 | 632157.8066 | 808433.4286 | 545218.7865 | 818290.2515 | 702207.9587 | 575272.5838 | 783211.5216 | 1066908.696 |
| L-Azetidine-2-carboxylic acid | 3334192.731 | 2568398.143 | 1795757.554 | 3459403.093 | 7623146.554 | 6641046.047 | 5590585.544 | 4734566.668 | 7839002.81 | 3357518.533 | 3836437.074 | 4016868.651 |
| SL 10:0;O/12:0 | 634213.1224 | 650031.8158 | 614703.8693 | 690646.1863 | 1201825.016 | 669970.4814 | 1830409.228 | 958244.0566 | 1353190.878 | 1748316.144 | 1557170.446 | 604811.1946 |
| N-Methyl-L-serine | 3511101.478 | 2589406.876 | 10834044.96 | 3336048.228 | 2084773.078 | 8772541.683 | 2443386.652 | 2512461 | 2793797.915 | 2471193.708 | 6237464.656 | 2854574.606 |
| Heptaminol | 1306624.843 | 1155965.747 | 1456312.112 | 1510014.636 | 1009440.137 | 1144602.267 | 1533731.169 | 1689893.671 | 897813.1442 | 1259548.135 | 1610585.381 | 1449807.572 |
| DGDG O-8:0_4:0 | 608167.07 | 354620.5773 | 646604.2926 | 478937.4305 | 1222347.102 | 1810867.013 | 639817.4291 | 522928.0471 | 989783.3958 | 1027315.978 | 1533747.665 | 1279631.948 |
| Amprolium cation | 846561.7356 | 1114207.45 | 1055350.825 | 1103170.209 | 778495.7226 | 769425.8342 | 1008449.215 | 795394.4671 | 676010.6827 | 784408.4701 | 654328.085 | 1192329.161 |
| (2-{2-[(3-Methoxyphenyl)amino]-2-oxoethoxy}phenyl)pho | 801876.2958 | 847067.1061 | 1988826.009 | 1203746.21 | 1238415.32 | 1475573.527 | 1386207.584 | 1425534.398 | 1024326.601 | 808711.5267 | 1559916.907 | 1327605.222 |

| Metabolites | PBS_1 | PBS_2 | PBS_3 | PBS_4 | BCG_1 | BCG_2 | BCG_3 | BCG_4 | BCGΔ *pe18*_1 | BCGΔ *pe18*_2 | BCGΔ *pe18*_3 | BCGΔ*pe18*_4 |
|---|---|---|---|---|---|---|---|---|---|---|---|---|
| sphonic acid | | | | | | | | | | | | |
| PE O-17:1_22:5 | 445156.8551 | 314577.5612 | 477855.0253 | 249534.0893 | 477057.8209 | 349816.1084 | 563669.9689 | 381962.5552 | 351448.4371 | 433993.0466 | 550450.249 | 257227.4967 |
| 5-(Trifluoromethyl)isoindoline | 14900853.84 | 7523245.263 | 23038585.61 | 11964457.77 | 17842779.32 | 20367859.54 | 13399070.11 | 12647502.38 | 13070195.69 | 13401706.99 | 12622364.48 | 12233713.68 |
| 6-(3,5-Dimethyl-1H-pyrazol-1-yl)-3(2H)-pyridazinone | 220810.8428 | 109835.4109 | 304166.9604 | 189304.4892 | 2917996.06 | 2417748.08 | 3086896.207 | 2192399.134 | 4706643.428 | 3022115.632 | 8629289.149 | 1678716.139 |
| Succinyl-CoA | 0 | 0 | 0 | 0 | 75669.98758 | 72546.08941 | 44968.24341 | 46074.45648 | 95874.29182 | 40249.34972 | 70116.50359 | 120353.4682 |
| Diacetyl | 1659336.449 | 1598603.512 | 1488296.358 | 1597818.426 | 4806429.804 | 2009251.659 | 4350557.958 | 3247550.64 | 2684387.721 | 1837158.231 | 1978217.834 | 1867851.694 |
| 3-O-Sulfogalactosylceramide (d18:1/24:0) | 356082.9601 | 687110.0456 | 272553.4267 | 789434.9729 | 374822.868 | 318698.7574 | 322096.3444 | 416375.4668 | 280515.3388 | 502320.6829 | 384021.4699 | 303753.8313 |
| SMGDG O-26:7_12:0 | 878378.2569 | 898641.7981 | 291336.728 | 696481.2817 | 639214.2264 | 2243744.277 | 484286.5017 | 575610.4105 | 901190.7237 | 555112.8216 | 330068.2406 | 445625.551 |
| 7-Chloro-8-methyl-2-(4-pyridinyl)-4-quinolinecarboxylic acid | 2000054.734 | 1258527.779 | 2528393.897 | 1358973.243 | 2152552.471 | 1852187.222 | 1711892.131 | 1414636.063 | 1444356.986 | 1377991.675 | 1293388.018 | 999159.0788 |

| Metabolites | PBS_1 | PBS_2 | PBS_3 | PBS_4 | BCG_1 | BCG_2 | BCG_3 | BCG_4 | BCGΔ*pe18*_1 | BCGΔ*pe18*_2 | BCGΔ*pe18*_3 | BCGΔ*pe18*_4 |
|---|---|---|---|---|---|---|---|---|---|---|---|---|
| Cer 25:3;3O/28:1;(2OH) | 19909.09764 | 43842.18605 | 8599.436783 | 0 | 70386.0715 | 43738.2576 | 32230.92838 | 0 | 8580.058393 | 44288.67994 | 70139.30103 | 452240.819 |
| 4-(Difluoromethyl)-6-(furan-2-yl)-2-hydrazinylpyrimidine | 501514.2611 | 502179.3371 | 492522.6395 | 522967.1632 | 381770.9304 | 622629.4746 | 479440.0824 | 519567.9943 | 505694.5774 | 519187.9519 | 500824.1184 | 505757.7481 |
| 2-Hydroxyphenylacetic acid | 361333.3646 | 285614.0106 | 397206.7797 | 353733.6495 | 545920.8451 | 503696.8259 | 547295.6313 | 452433.4357 | 426242.1546 | 427047.2128 | 413228.5775 | 405053.9552 |
| 1-(Cyclopentylcarbonyl)-4-piperidinamine | 9412304.582 | 11161632.25 | 9486887.259 | 9127389.481 | 8342197.021 | 8893849.918 | 8648625.439 | 8581210.81 | 8417995.721 | 8342382.28 | 8453327.489 | 9188629.111 |
| Benzenemethanaminium, N-cyclopentyl-N,N-dimethyl-3,4-bis(tetradecyloxy)- | 912092.1002 | 1465630.355 | 1364865.371 | 1486283.308 | 1184878.337 | 1483237.086 | 886193.6573 | 1021701.3 | 990925.8231 | 792488.9972 | 13169.12829 | 1546185.739 |
| Squamosamide | 431830.6255 | 488997.4288 | 435693.3711 | 451881.7916 | 460801.8084 | 495949.7678 | 496764.9591 | 465320.7956 | 433399.2894 | 479711.6298 | 503891.1773 | 465873.1672 |
| (4-Acetyl-2-methoxyphenoxy)acetic acid | 516479.0261 | 422326.2341 | 540029.2885 | 912888.7186 | 3257034.752 | 2328398.748 | 2305927.853 | 1942718.887 | 718581.4288 | 714417.3463 | 781782.1857 | 727280.6912 |
| 2,4,5-Trimethoxydihydrocinnamic acid | 2635481.719 | 1887054.504 | 2314810.156 | 2236053.392 | 2020628.591 | 2394039.7 | 2162879.409 | 1837642.442 | 2117223.714 | 3121679.047 | 2006110.915 | 2558920.617 |

| **Metabolites** | **PBS_1** | **PBS_2** | **PBS_3** | **PBS_4** | **BCG_1** | **BCG_2** | **BCG_3** | **BCG_4** | **BCGΔ*pe18*_1** | **BCGΔ*pe18*_2** | **BCGΔ*pe18*_3** | **BCGΔ*pe18*_4** |
|---|---|---|---|---|---|---|---|---|---|---|---|---|
| MGDG 20:4_18:5 | 542827.1578 | 585036.3613 | 254579.8764 | 398639.1835 | 482030.9729 | 404154.6113 | 482036.9375 | 512934.9531 | 674731.1808 | 624143.4053 | 404098.6214 | 386217.2462 |
| 11-Hydroxyundecanoic acid | 934540.3246 | 1071892.604 | 1148956.339 | 839209.386 | 1287496.339 | 1480367.72 | 1308487.421 | 1231200.217 | 1361828.519 | 1413319.098 | 4248564.312 | 1581483.304 |
| Phylloporphyrin | 75050.16789 | 180963.0524 | 110882.0288 | 145949.0073 | 227942.3969 | 485879.581 | 318716.5065 | 277776.1154 | 272679.0798 | 355907.3091 | 539235.9623 | 295168.2571 |
| 5-Hydroxyindole-3-acetic acid | 577009.1924 | 444918.3002 | 542949.5218 | 540813.9465 | 744606.5409 | 677989.9439 | 662778.4342 | 782480.8615 | 646893.4556 | 342973.5424 | 388606.9536 | 601473.4175 |
| DL-Coniine | 2063038.437 | 777216.7423 | 1942247.56 | 1908290.346 | 1346677.866 | 690652.4661 | 494496.8292 | 465547.338 | 485927.517 | 1005295.8 | 894086.261 | 318925.8896 |
| MG 16:0 | 2152351.798 | 2249102.746 | 5882128.14 | 2497126.975 | 6502386.917 | 4730329.243 | 9641414.585 | 5409536.506 | 1715880.159 | 4007801.25 | 4127772.265 | 1458026.807 |
| Histamine | 3859987.14 | 2883853.458 | 1728880.091 | 3382323.769 | 3801113.705 | 2921691.116 | 2387265.803 | 3737612.522 | 4199130.457 | 5660486.843 | 1660734.27 | 3003848.499 |
| 5-Amino-2-(1,3-benzoxazol-2-yl)phenol | 709765.9467 | 546226.7333 | 507493.985 | 521667.1997 | 462182.4608 | 466982.173 | 550309.462 | 468973.3541 | 490068.493 | 687904.0536 | 713874.6941 | 512834.8682 |
| PC 16:0_20:3 | 465038.0381 | 442362.0249 | 426742.1746 | 470614.4254 | 875610.9442 | 936875.9047 | 928342.8833 | 925230.3684 | 847664.7296 | 1075780.58 | 880910.7502 | 1120456.805 |
| 3-[(Ethylanilino)methyl]benzenesulfonic acid | 229621.6527 | 208869.8512 | 355406.0737 | 344249.0181 | 299169.1951 | 322386.5226 | 320590.7831 | 284998.8537 | 258687.4172 | 183625.0134 | 179690.0307 | 266867.2146 |
| Cer 12:2;2O/4:0 | 2136102.455 | 730602.173 | 392131.4353 | 1411446.361 | 769255.0187 | 685793.5715 | 599624.3367 | 1197731.46 | 801079.2945 | 616568.2932 | 831817.2001 | 1030756.076 |

| **Metabolites** | **PBS_1** | **PBS_2** | **PBS_3** | **PBS_4** | **BCG_1** | **BCG_2** | **BCG_3** | **BCG_4** | **BCGΔ*pe18*_1** | **BCGΔ*pe18*_2** | **BCGΔ*pe18*_3** | **BCGΔ*pe18*_4** |
|---|---|---|---|---|---|---|---|---|---|---|---|---|
| DGTS 13:0_20:5 | 219859.4123 | 245382.1411 | 300183.0137 | 166498.1923 | 618677.639 | 686790.9876 | 645873.8137 | 658741.8829 | 162659.7676 | 134832.0169 | 176413.8028 | 204594.4079 |
| Quercitrin | 343266.7767 | 891998.4358 | 513572.6748 | 890352.7852 | 487106.9578 | 442242.5316 | 827000.8507 | 965684.537 | 674687.1001 | 880150.6978 | 1063592.051 | 651252.6143 |
| Betaxolol | 4401468.111 | 15408984.19 | 2443919.907 | 4806123.916 | 2931553.04 | 2364488.358 | 2354728.039 | 2327097.479 | 3007090.382 | 2785179.426 | 7675514.525 | 3055236.615 |
| Imazamethabenz-methyl | 278593.414 | 40193.34438 | 251089.6137 | 237191.074 | 174317.7708 | 371528.9581 | 5979880.356 | 665143.3166 | 973605.3199 | 442800.9873 | 325121.7199 | 446583.6973 |
| LPE 15:0 | 184118.3463 | 95945.69357 | 205042.7174 | 185048.5997 | 596012.4964 | 580132.6398 | 579503.4956 | 564952.1964 | 595694.681 | 702789.2121 | 703747.1375 | 518143.8897 |
| 2,4-Dichloro-N-(1-methyl-4-piperidinyl)benzamide | 1745004.497 | 1553482.992 | 2157252.737 | 1613259.809 | 2333660.931 | 2722239.374 | 2369800.872 | 2092499.93 | 2044515.571 | 1889507.847 | 2280136.696 | 1992934.237 |
| LPE O-18:0 | 720034.2397 | 1026729.423 | 659526.9178 | 785619.1676 | 670612.6337 | 871353.8394 | 678860.9862 | 545581.6274 | 430796.7083 | 452230.0659 | 682985.5035 | 1701576.732 |
| SL 17:1;O/17:0 | 81408.41931 | 121525.0746 | 104613.7242 | 133141.5495 | 120136.506 | 104252.4053 | 75897.37399 | 87414.60972 | 44549.98117 | 78656.68867 | 153979.6858 | 112354.8002 |
| 6-Chloro-1,3-benzoxazole-2(3H)-thione | 3582903.535 | 2701110.614 | 3920630.387 | 2784366.555 | 7049190.528 | 4336640.781 | 7919890.219 | 5338391.093 | 3651261.985 | 5552068.157 | 5362558.323 | 2602178.499 |
| 3-Hydroxycinnamic acid | 1449117.258 | 1246589.288 | 1995371.79 | 1551073.139 | 3158787.985 | 2304451.611 | 2825321.089 | 2632725.043 | 2724237.458 | 1775093.518 | 2059864.49 | 2534045.273 |
| NAE 14:1 | 131349445.8 | 138474037.5 | 25256447.99 | 26285264.92 | 120707596.5 | 109540462.2 | 128372842.4 | 125787808.3 | 40196798.06 | 37217612.63 | 122254102.1 | 28030207.65 |

| **Metabolites** | **PBS_1** | **PBS_2** | **PBS_3** | **PBS_4** | **BCG_1** | **BCG_2** | **BCG_3** | **BCG_4** | **BCGΔ*pe18*_1** | **BCGΔ*pe18*_2** | **BCGΔ*pe18*_3** | **BCGΔ*pe18*_4** |
|---|---|---|---|---|---|---|---|---|---|---|---|---|
| 2-Oxiraneoctanoic acid, .eta.-hydroxy-3-octyl- | 1605902.194 | 1609556.566 | 2389571.838 | 2621339.205 | 2728001.665 | 2815257.18 | 2865835.34 | 2477258.28 | 2099860.635 | 2134636.129 | 2847398.285 | 3100890.565 |
| 2,4,6-Tris(4-morpholinylmethyl)phenol | 309770.0341 | 305766.8075 | 326700.6097 | 363986.8497 | 357293.2899 | 443110.731 | 383617.0963 | 369477.6449 | 277441.8097 | 319340.1952 | 482411.7269 | 400808.1694 |
| Carbetapentane | 21841271.41 | 18188814.03 | 12538737.04 | 19370293.85 | 16711308.4 | 18087243.41 | 16324877.02 | 16339419.67 | 17020524.81 | 17513289.38 | 17017693.7 | 15612292.25 |
| PE O-26:5_16:0 | 787418.0704 | 773872.8843 | 854466.1445 | 667748.9874 | 769993.6814 | 965594.4744 | 787659.6877 | 717808.5595 | 759220.4363 | 702388.6188 | 857813.1382 | 731783.2455 |
| N-Cyclohexanecarbonylpentadecylamine | 6106883.663 | 5307247.279 | 3104920.833 | 21547834.5 | 2277499.501 | 2330689.973 | 2503945.706 | 2338526.558 | 3309286.795 | 2790540.051 | 2876754.842 | 2155862.993 |
| 4-Aminobutyric acid (GABA) | 655018.2627 | 305448.4116 | 300248.5031 | 312280.0717 | 3276087.499 | 2680476.45 | 3483634.129 | 2817744.093 | 5960240.97 | 4974145.687 | 5369947.851 | 4626435.537 |
| SL 21:2;O/26:5 | 2421131.848 | 1312577.159 | 991848.4264 | 1224767.78 | 1090742.291 | 2431530.296 | 1470341.631 | 1492690.579 | 1470884.665 | 1160299.846 | 764109.7283 | 1241849.672 |
| 2,4-Dihydroxy-7,8-dimethoxy-2H-1,4-benzoxazin-3(4H)-one_2-glucoside | 138360.9303 | 73987.34858 | 147839.6782 | 89898.47049 | 286486.1798 | 299357.0634 | 278343.2461 | 319290.3152 | 316683.9514 | 314535.619 | 258373.0148 | 227442.3077 |
| DL-Leucinamide | 1133402.727 | 3430636.404 | 5036938.948 | 3718680.495 | 1195469.567 | 1476339.988 | 1142514.28 | 1436581.393 | 1624518.13 | 1218254.413 | 703329.5342 | 7893913.281 |
| Canellin E | 97699.9 | 53784.2 | 36599.6 | 51632.1 | 417729. | 280430. | 486964. | 346237. | 217586. | 232696. | 383216. | 155079. |

| Metabolites | PBS_1 | PBS_2 | PBS_3 | PBS_4 | BCG_1 | BCG_2 | BCG_3 | BCG_4 | BCGΔ *pe18*_1 | BCGΔ *pe18*_2 | BCGΔ *pe18*_3 | BCGΔ*pe18*_4 |
|---|---|---|---|---|---|---|---|---|---|---|---|---|
| | 871 | 6845 | 015 | 1248 | 3984 | 2945 | 528 | 3695 | 6411 | 4313 | 9807 | 2777 |
| Deoxycholic acid | 728581.7868 | 961041.6433 | 1103004.852 | 2001379.881 | 695466.5473 | 1162736.022 | 962468.2383 | 774080.8096 | 860814.4541 | 689500.5402 | 741553.3499 | 804666.2826 |
| 5-Ethyl-N-phenyl-2-pyridinecarbothioamide | 3020882.935 | 2672061.032 | 5393052.775 | 3551031.899 | 3628408.59 | 3132284.978 | 3519803.122 | 3092426.801 | 3159040.321 | 2386798.927 | 2700778.269 | 3057404.314 |
| 2-Methylhippuric acid | 809966.8977 | 706539.9081 | 1280797.876 | 982255.7535 | 1054536.495 | 1106724.197 | 1259774.115 | 1014474.971 | 835413.9057 | 518528.1823 | 611322.8326 | 891237.1934 |
| 1H-Pyrrolo[3,4-b]quinoline-1,9(4H)-dione, 2,3-dihydro-4-methyl-3-(2-methylpropyl)- | 2453370.175 | 958370.3104 | 1395934.518 | 3193810.947 | 5828439.941 | 3799395.247 | 1009294.871 | 1262916.358 | 2346935.541 | 1336093.179 | 1781600.318 | 1009565.793 |
| Clafen_(Cyclophosphamide) | 1569536.244 | 2143463.338 | 1665709.306 | 1724061.879 | 1921740.431 | 1497959.987 | 1822767.081 | 1343337.407 | 1814282.244 | 1136021.411 | 1192929.457 | 1435296.491 |
| LPE 17:0 | 1158368.735 | 688380.7851 | 667852.3686 | 438383.6263 | 2582394.248 | 2082555.463 | 2435130.585 | 1967879.711 | 2309236.895 | 2482959.753 | 2476876.664 | 1647838.303 |
| Tetraheptylammonium cation | 1076032.279 | 1116863.194 | 299947.3845 | 786168.512 | 351654.8867 | 232579.2961 | 263799.2767 | 379241.4862 | 427763.0599 | 409169.6216 | 232788.853 | 234415.2805 |
| Ginkgolide_A | 16479.97621 | 29072.6235 | 40718.88083 | 21609.00823 | 4300400.795 | 5094886.462 | 3732006.941 | 4168079.297 | 4771223.352 | 3942470.299 | 4346910.684 | 5250518.884 |
| LPE(14:0) | 166709.7068 | 113764.6771 | 193247.0312 | 149036.1915 | 1151375.349 | 1189857.907 | 1158185.134 | 887925.4414 | 1159585.664 | 1185680.128 | 1481817.609 | 1063406.882 |

| **Metabolites** | **PBS_1** | **PBS_2** | **PBS_3** | **PBS_4** | **BCG_1** | **BCG_2** | **BCG_3** | **BCG_4** | **BCGΔ*pe18*_1** | **BCGΔ*pe18*_2** | **BCGΔ*pe18*_3** | **BCGΔ*pe18*_4** |
|---|---|---|---|---|---|---|---|---|---|---|---|---|
| SL 12:1;O/18:2;O | 1382550.692 | 1324603.252 | 1037593.142 | 1707329.524 | 4884339.687 | 3422359.786 | 2226388.791 | 3665218.202 | 2378677.088 | 947184.9382 | 880531.3017 | 2658571.445 |
| 3-(2,4-Dioxo-3,4-dihydro-1(2H)-pyrimidinyl)alanine | 2280072.318 | 3275218.713 | 2620130.322 | 2461602.071 | 3278147.08 | 3382539.668 | 2314548.041 | 2426067.865 | 2372372.931 | 2381962.159 | 2332120.024 | 3218098.047 |
| PS 14:1_26:4 | 2236029.175 | 6702521.447 | 10465889.26 | 10230811.14 | 10226867.13 | 5246937.876 | 5182200.884 | 8654569.301 | 2761435.133 | 5549483.825 | 3217822.849 | 2338163.261 |
| Glucobrassicanapin | 304157.4828 | 173857.0833 | 1351630.707 | 523040.077 | 1014064.999 | 1479428.302 | 1098554.976 | 1097923.821 | 990203.297 | 204399.1052 | 884157.0759 | 1314021.474 |
| 3,6,9,12-Tetraoxatetracosan-1-ol | 3557856.157 | 5029192.984 | 2985268.971 | 4383570.081 | 5490807.566 | 5655845.67 | 3809070.524 | 6717008.664 | 4193047.876 | 3011779.053 | 3165861.233 | 5582436.401 |
| Hinokitiol | 10724728.34 | 12959874.63 | 10312479.28 | 10411009.99 | 11935211.19 | 12694274.71 | 12337900.45 | 12297777.56 | 11455727.33 | 11325005.58 | 11725846.84 | 10193325.45 |
| 2,4-Thiazolidinedione, 5-[[4-[2-[5-(1-hydroxyethyl)-2-pyridinyl]ethoxy]phenyl]methyl]- | 851064.6002 | 774820.3262 | 812320.0687 | 715395.087 | 668733.7093 | 604166.5486 | 727307.2207 | 671509.6918 | 653799.8101 | 792793.7001 | 658729.1331 | 970734.444 |
| 13,14-dihydro-15-keto-PGA2 | 53425.45019 | 25482.91042 | 151388.7329 | 51614.94363 | 1499493.771 | 2149499.952 | 1122694.468 | 1862373.381 | 2052727.132 | 770793.9862 | 759511.2821 | 2702907.561 |
| Arachidonic sulfonic acid | 3659371.398 | 2627590.955 | 747754.9809 | 1009064.009 | 5019671.59 | 3816213.914 | 4616683.57 | 4262120.145 | 3995158.101 | 4942537.347 | 1551108.649 | 3749636.293 |
| PMeOH 16:0_22:6 | 985125. | 589268. | 680155. | 696201. | 570256. | 927491. | 622571. | 533616. | 895447. | 824641. | 866909. | 500816. |

| Metabolites | PBS_1 | PBS_2 | PBS_3 | PBS_4 | BCG_1 | BCG_2 | BCG_3 | BCG_4 | BCGΔ*pe18*_1 | BCGΔ*pe18*_2 | BCGΔ*pe18*_3 | BCGΔ*pe18*_4 |
|---|---|---|---|---|---|---|---|---|---|---|---|---|
| | 9845 | 9977 | 088 | 9206 | 0715 | 0428 | 3515 | 3183 | 1339 | 1006 | 6223 | 7681 |
| PG O-16:2_22:5 | 597473.6728 | 445848.8056 | 424187.0244 | 387917.8606 | 565379.9362 | 460351.7811 | 431093.9571 | 439692.6568 | 460276.2105 | 396145.6885 | 432787.1392 | 493747.401 |
| Pendimethalin | 2097847.258 | 2166472.122 | 2214432.981 | 2038158.776 | 2295428.878 | 2169934.676 | 2435833.794 | 2453753.518 | 2278949.502 | 2181023.424 | 2181231.069 | 1865678.783 |
| TG(22:6(4Z,7Z,10Z,13Z,16Z,19Z)/20:5(5Z,8Z,11Z,14Z,17Z)/22:6(4Z,7Z,10Z,13Z,16Z,19Z))[iso3] | 0 | 0 | 0 | 0 | 241924.768 | 394568.0937 | 723481.7638 | 904394.5189 | 511479.5431 | 1333556.808 | 455515.8421 | 352530.098 |
| MGDG O-17:2_22:6 | 1732644.319 | 2567015.13 | 1823026.525 | 1781464.243 | 1846111.582 | 2017386.516 | 1870623.841 | 1699132.773 | 1316582.691 | 1640206.645 | 1749139.956 | 1576604.518 |
| NAGlySer 26:7/22:4 | 0 | 0 | 0 | 5515.353948 | 671282.2446 | 424729.8015 | 284494.2099 | 359605.5574 | 268968.4413 | 216864.8022 | 182589.3409 | 321425.4661 |
| L-N5-(1-Imino-3-pentenyl)ornithine | 2310811.56 | 16845777.66 | 1343821.555 | 17957501.99 | 23233575.7 | 13940675.65 | 1079125.137 | 1041070.407 | 4111907.041 | 12813359.72 | 7339591.914 | 1894601.271 |
| Tomatidine | 4528339.861 | 4697682.507 | 3708256.671 | 4180243.118 | 4159617.258 | 4148753.82 | 4398495.063 | 4221115.159 | 4485637.776 | 4293872.037 | 4239604.175 | 3613950.378 |
| gamma-Glutamylalanine | 450055.027 | 544729.6703 | 757289.8116 | 452700.8618 | 5482143.263 | 3510886.17 | 4009607.098 | 3410522.812 | 7656794.241 | 7645287.916 | 6281642.999 | 7290745.865 |
| 2-Deoxyribose 5-phosphate | 156161.2022 | 356757.5739 | 180200.1513 | 182859.1326 | 3105721.843 | 435417.8159 | 31003.05554 | 1078546.72 | 792481.1709 | 1557649.583 | 1526230.454 | 249901.3815 |
| 1-decanesulfonate | 262714 | 357317 | 207569 | 309997 | 305420 | 142244 | 371166 | 336788 | 240708 | 221623 | 250662 | 329174 |

| Metabolites | PBS_1 | PBS_2 | PBS_3 | PBS_4 | BCG_1 | BCG_2 | BCG_3 | BCG_4 | BCGΔ*pe18*_1 | BCGΔ*pe18*_2 | BCGΔ*pe18*_3 | BCGΔ*pe18*_4 |
|---|---|---|---|---|---|---|---|---|---|---|---|---|
| | 9.505 | 5.689 | 4.112 | 2.535 | 2.321 | 5.176 | 1.951 | 7.245 | 0.137 | 1.926 | 9.459 | 3.013 |
| Cer 16:0;2O/2:0 | 4130865.778 | 4346243.223 | 3848780.608 | 4442075.044 | 4433850.27 | 4352461.27 | 4544505.115 | 4766177.507 | 4247855.47 | 4366568.269 | 4447626.875 | 3773256.748 |
| 4-Chloro-N-methyl-3-nitro-benzenesulfonamide | 386210.2486 | 416764.983 | 437671.6964 | 455496.4664 | 395703.095 | 419424.7569 | 414516.445 | 442046.9819 | 374735.7173 | 473808.0969 | 395626.5106 | 430774.7379 |
| N-[(5-Bromo-2-fluorophenyl)methyl]cyclopropanamine | 2283618.444 | 2253258.325 | 2221115.004 | 2234533.642 | 2200439.41 | 2184432.032 | 2096088.785 | 2244228.325 | 2152504.483 | 1975035.28 | 2075003.573 | 2046850.565 |
| PE 17:1_17:1 | 1415091.385 | 1271795.808 | 2144360.091 | 1412220.802 | 8156448.322 | 5552761.619 | 6974710.618 | 5034504.627 | 7090553.076 | 6912346.096 | 6900941.947 | 4582286.08 |
| Tridecanedioylcarnitine | 3042230.199 | 4484960.707 | 652274.5589 | 3420976.325 | 1286931.086 | 1265862.49 | 1545193.078 | 1050556.905 | 2401072.597 | 1897846.414 | 1075001.706 | 2376121.858 |
| LPS 19:1 | 945024.6783 | 485588.9464 | 1004982.551 | 1073181.9 | 1500610.496 | 1091299.116 | 1523871.794 | 930548.1471 | 820374.5016 | 1047403.175 | 1272733.497 | 545212.2756 |
| OCTYL GALLATE | 8385485.088 | 8442391.333 | 8211469.366 | 8475784.011 | 8090300.138 | 8175909.538 | 8404668.05 | 8284335.774 | 1127286.521 | 8341033.043 | 2257939.914 | 8351345.772 |
| 6-(Trifluoromethyl)-1H-pyrazolo[3,4-d]pyrimidin-4-ol | 380329.6892 | 439504.0234 | 376154.5674 | 409363.4465 | 375595.3216 | 374689.4331 | 372596.4821 | 359948.5007 | 391675.4851 | 369055.9171 | 390093.8337 | 375412.9035 |
| PS(22:4(7Z,10Z,13Z,16Z)/22:4(7Z,10Z,13Z,16Z)) | 21868270.57 | 24214374.71 | 10269346.64 | 16232688.9 | 5074244.367 | 3925049.409 | 6098601.18 | 6761140.381 | 6396617.55 | 7212410.832 | 4910016.234 | 4562457.496 |
| Nimbidiol | 185660 | 197685 | 179462 | 193261 | 188939 | 178133 | 172313 | 181202 | 177182 | 171280 | 177613 | 171087 |

| **Metabolites** | **PBS_1** | **PBS_2** | **PBS_3** | **PBS_4** | **BCG_1** | **BCG_2** | **BCG_3** | **BCG_4** | **BCGΔ*pe18*_1** | **BCGΔ*pe18*_2** | **BCGΔ*pe18*_3** | **BCGΔ*pe18*_4** |
|---|---|---|---|---|---|---|---|---|---|---|---|---|
| | 7.393 | 5.128 | 3.768 | 4.322 | 6.736 | 0.819 | 8.384 | 1.113 | 6.327 | 3.364 | 1.939 | 9.021 |
| 4'-Hydroxywarfarin | 240942.1695 | 325337.4759 | 301045.2453 | 331881.0025 | 330412.9683 | 288581.8925 | 288347.6058 | 305547.6965 | 338085.554 | 345976.9369 | 365364.6868 | 614623.0123 |
| PC(42:9) | 42383013.62 | 51097583.83 | 20687686.02 | 30804235.72 | 30110876.31 | 23007906.95 | 28752836.07 | 31159675.95 | 37362989.8 | 38610370.62 | 24928079.01 | 20843386.26 |
| Moclobemide | 894543.8805 | 813118.2805 | 1522207.742 | 1119165.481 | 1072330.22 | 1233748.633 | 1197408.221 | 1062024.243 | 1176311.646 | 507256.9893 | 672757.4552 | 1289235.998 |
| 19-O-Acetylchaetoglobosin D | 8329.066108 | 8589.221682 | 21806.36954 | 10295.91931 | 113786.633 | 85392.49911 | 78999.55015 | 86752.6526 | 101325.9977 | 67905.09321 | 96453.52324 | 108349.6473 |
| dTDP-6-deoxy-beta-talose | 0 | 0 | 0 | 0 | 3993813.154 | 3326705.248 | 2959384.851 | 2851371.009 | 4672576.851 | 3534801.377 | 4409173.256 | 5279322.264 |
| Trapidil | 1932429.049 | 4548498.912 | 4864425.469 | 5652778.38 | 4214185.664 | 4098505.325 | 6160671.082 | 3937779.848 | 4991011.32 | 4169142.335 | 5576468.605 | 5698274.315 |
| PS 18:1_24:4 | 270310.4713 | 270604.3691 | 191525.0304 | 247237.5847 | 151170.7017 | 351747.8669 | 122678.013 | 98616.20161 | 117143.6812 | 145551.7992 | 145562.9802 | 447302.8203 |
| Hematoxylin | 3848093.118 | 2373984.573 | 7310048.89 | 2035443.034 | 3083438.151 | 792794.267 | 3935786.94 | 1055715.387 | 458970.5485 | 979327.0062 | 1584551.453 | 1087348.659 |
| 4-Hydroxyproline | 3437275.862 | 3013455.817 | 3689935.084 | 3023064.353 | 14369844.36 | 7400492.296 | 11872140.56 | 10367307.18 | 9431852.828 | 10284993.02 | 9263513.612 | 7335649.108 |
| N-[[[4-[[5-(Dimethylamino)-1-oxopentyl]amino]phenyl]amino]thioxomethyl]-4-(1,1-dimethyleth | 52967.65613 | 11693.86747 | 39004.23389 | 56245.953 | 368633.7668 | 241027.1298 | 243454.3134 | 211402.6591 | 301252.2825 | 357553.5596 | 317329.3513 | 360791.858 |

| Metabolites | PBS_1 | PBS_2 | PBS_3 | PBS_4 | BCG_1 | BCG_2 | BCG_3 | BCG_4 | BCGΔ *pe18*_1 | BCGΔ *pe18*_2 | BCGΔ *pe18*_3 | BCGΔ*pe18*_4 |
|---|---|---|---|---|---|---|---|---|---|---|---|---|
| yl)benzamide | | | | | | | | | | | | |
| 2-hexyl-3-oxodecanoate | 4352050.417 | 4692505.541 | 4320847.305 | 4643694.96 | 3943268.411 | 3369089.324 | 3596264.137 | 3462793.458 | 3832949.348 | 3752001.47 | 3869048.254 | 3692574.526 |
| 4-Hydroxy-N-(4-hydroxyphenyl)benzamide | 1839507.776 | 1409806.277 | 2295261.137 | 1510479.899 | 3430119.873 | 2121547.566 | 2717584.129 | 2724684.341 | 2641408.408 | 2256009.559 | 2327017.916 | 1745460.568 |
| 3-[(2-Chloro-3-pyridinyl)amino]-2-cyclohexen-1-one | 2061421.623 | 1064214.302 | 3142088.786 | 1514518.226 | 2238681.393 | 2336010.515 | 1613681.855 | 1347290.536 | 1272409.363 | 973904.6897 | 936944.3669 | 995335.7295 |
| (6E)-8-Methoxy-3,7-dimethyl-8-oxoocta-1,6-dien-3-yl 6-O-(4-carboxy-3-hydroxy-3-methylbutanoyl)hexopyranoside | 320417.6759 | 231856.6241 | 360621.6002 | 391291.8039 | 236677.7808 | 251487.8358 | 1687441.713 | 315333.9082 | 420606.345 | 387637.2438 | 224361.9173 | 381774.6508 |
| M528T476_1 | 153004.3973 | 131419.4147 | 220547.2674 | 136847.304 | 195247.6577 | 417801.7588 | 468722.1678 | 145183.4908 | 380503.0139 | 438378.1227 | 511225.0388 | 131817.1706 |
| 2-(4-(Diethylamino)but-2-yn-1-yl)isoindoline-1,3-dione | 1014046.411 | 1049861.407 | 896068.442 | 945428.6958 | 973036.5245 | 1087094.919 | 916165.3134 | 970791.1043 | 1029966.225 | 1009434.139 | 983840.998 | 885765.3823 |

| **Metabolites** | **PBS_1** | **PBS_2** | **PBS_3** | **PBS_4** | **BCG_1** | **BCG_2** | **BCG_3** | **BCG_4** | **BCGΔ*pe18*_1** | **BCGΔ*pe18*_2** | **BCGΔ*pe18*_3** | **BCGΔ*pe18*_4** |
|---|---|---|---|---|---|---|---|---|---|---|---|---|
| PI O-16:2_18:2 | 107225.9164 | 91900.09656 | 29544.57496 | 204529.4686 | 539907.9887 | 276693.3742 | 522723.3968 | 363678.7248 | 573845.8487 | 686079.7736 | 516861.8547 | 257729.3145 |
| PE-Cer 12:1;2O/17:1 | 5576.386504 | 4459.804267 | 0 | 5954.215917 | 132780.7271 | 71357.89384 | 99906.93182 | 102797.9831 | 40494.17944 | 75777.87521 | 58785.02837 | 34114.92407 |
| 6,7,8,9,14,15-Hexahydro-7-methyl-5H-indolo[3,2-f][3]benzazecine | 1442075.162 | 1310542.792 | 1282747.463 | 1221447.629 | 614420.844 | 786595.4288 | 1058275.609 | 859708.2527 | 868121.0329 | 776928.8069 | 965945.3367 | 671535.3533 |
| D-Glucosaminic acid | 593850.9744 | 679637.7821 | 578194.6361 | 626976.1241 | 1106485.2 | 832525.6814 | 3670750.758 | 3143060.37 | 973818.4516 | 713465.6934 | 1112583.005 | 133836.4271 |
| SMGDG O-13:1_26:7 | 169762.6603 | 302588.9575 | 134273.3223 | 105383.3459 | 675172.0118 | 1099059.607 | 615988.1344 | 592985.7032 | 1042770.844 | 979508.4646 | 844184.753 | 933748.5879 |
| Sucrose | 6229.943972 | 13027.11296 | 34877.51122 | 434233.6736 | 2321743.264 | 1925094.058 | 1673306.673 | 1465547.267 | 2194829.585 | 1865984.062 | 2112761.721 | 2429479.646 |
| Pyrazoxyfen | 32627.00383 | 0 | 21171.8482 | 4193.830698 | 527343.7376 | 501515.1404 | 453231.9353 | 441323.1003 | 507794.9701 | 412191.3109 | 441173.4403 | 526970.5207 |
| M273T228 | 995920.8158 | 57742.00723 | 538981.9666 | 804650.3416 | 369403.6615 | 879997.2716 | 18245400.61 | 2079651.426 | 2923838.92 | 810199.4545 | 1032329.48 | 876010.8311 |
| SMGDG O-18:5_4:0 | 1294032.044 | 805976.3215 | 1387931.737 | 1268426.019 | 1241700.637 | 855382.7124 | 934956.6354 | 870904.4998 | 1445648.757 | 1337191.184 | 981000.5053 | 392760.1444 |
| 5-amino-2,4,5-trioxopentanoate | 100186.7252 | 118240.6547 | 114795.5728 | 112116.0022 | 341047.41 | 255724.7918 | 363209.2139 | 304906.7989 | 228384.1208 | 323154.8844 | 370997.9982 | 259745.8678 |
| 2-Hydroxymethylbenzoic acid | 361333.3646 | 285614.0106 | 397206.7797 | 353733.6495 | 545920.8451 | 503696.8259 | 547295.6313 | 452433.4357 | 426242.1546 | 427047.2128 | 413228.5775 | 405053.9552 |

| Metabolites | PBS_1 | PBS_2 | PBS_3 | PBS_4 | BCG_1 | BCG_2 | BCG_3 | BCG_4 | BCGΔ *pe18*_1 | BCGΔ *pe18*_2 | BCGΔ *pe18*_3 | BCGΔ*pe18*_4 |
|---|---|---|---|---|---|---|---|---|---|---|---|---|
| SMGDG O-8:0_13:1 | 0 | 0 | 8084.817225 | 0 | 91365.46496 | 76899.06015 | 96510.80685 | 81066.36983 | 56426.13293 | 67594.57594 | 85135.90218 | 34786.71924 |
| Phytosphingosine 1-phosphate | 959314.659 | 1000058.444 | 1003274.762 | 800135.0571 | 922536.7509 | 976369.5986 | 996071.9115 | 986684.6124 | 985512.7784 | 979547.1566 | 1017718.65 | 958458.4225 |
| 4-Methylsalicylic acid | 915165.9202 | 740263.8129 | 782693.1935 | 932905.0078 | 944553.2178 | 785908.9465 | 923158.334 | 943421.5404 | 854376.671 | 933475.4828 | 783359.2076 | 832788.861 |
| SHexCer 36:3;3O | 1197208.426 | 2756414.87 | 1251571.129 | 1684201.443 | 399254.8805 | 324164.7204 | 575516.7542 | 622990.6288 | 622632.4824 | 658316.1876 | 466514.283 | 362689.597 |
| trans-pterostilbene | 8450487.16 | 8815510.679 | 13751625.29 | 30317052.42 | 1679801.689 | 1290571.182 | 1421895.602 | 1251609.386 | 1866296.244 | 1893929.648 | 1472151.203 | 4479700.771 |
| Gangaleoidin | 500360.6994 | 443898.6803 | 801120.9928 | 641263.256 | 645787.8108 | 726401.5219 | 684570.184 | 579180.3019 | 616434.5469 | 300646.5573 | 341778.3238 | 661845.087 |
| Sterol 3-beta-glucoside | 584209.6776 | 474835.4349 | 582810.2666 | 352883.4926 | 821817.2118 | 674538.1714 | 981719.2854 | 857676.2084 | 677709.2072 | 582192.6049 | 993508.5434 | 334727.6413 |
| Gabapentin | 336927.2112 | 404418.0576 | 197629.4928 | 969878.0889 | 448497.3149 | 245912.4923 | 901008.1311 | 157570.6919 | 480554.2091 | 367679.0466 | 162767.33 | 1153492.06 |
| Leu-Val | 469623.0197 | 493328.4911 | 469500.7772 | 349399.356 | 415618.4998 | 251868.8787 | 349398.5556 | 312471.822 | 424218.2854 | 533094.5722 | 499549.307 | 233442.1776 |
| SM 23:1;2O/17:0 | 693726.0735 | 10255500.78 | 10402939.01 | 8750373.965 | 18753292.56 | 14679417.75 | 12692717.15 | 7433112.221 | 27931850.06 | 29139876.88 | 13180406.66 | 11614343.71 |
| LNAPE 20:4/N-19:1 | 312014.2079 | 215994.5883 | 337328.2429 | 297724.9047 | 135012.3401 | 63796.13585 | 482414.4279 | 155261.7993 | 331651.3368 | 209414.007 | 334215.1089 | 419976.6207 |
| Cer 9:1;2O/22:6 | 279336.352 | 240704.4053 | 280721.1167 | 429669.0681 | 254534.1878 | 269411.1333 | 320351.8718 | 254443.3532 | 234631.3936 | 220955.5528 | 197688.5236 | 468831.1101 |

| Metabolites | PBS_1 | PBS_2 | PBS_3 | PBS_4 | BCG_1 | BCG_2 | BCG_3 | BCG_4 | BCGΔ*pe18*_1 | BCGΔ*pe18*_2 | BCGΔ*pe18*_3 | BCGΔ*pe18*_4 |
|---|---|---|---|---|---|---|---|---|---|---|---|---|
| 2-Hexyl-4,5-dimethyloxazole | 1425054.117 | 1213659.505 | 1219570.999 | 2698112.374 | 1693212.001 | 2147749.06 | 4007524.058 | 1420612.231 | 2393340.132 | 1834378.576 | 1916337.182 | 2760019.794 |
| CE 16:2 | 643522.7903 | 604271.1874 | 479414.1345 | 439257.2863 | 348022.0927 | 360403.9779 | 526067.7561 | 466861.5059 | 478915.0982 | 345312.8619 | 346065.327 | 291452.1201 |
| 6.beta.-Naltrexol | 1301962.843 | 1296699.39 | 1152752.916 | 1424216.094 | 1241256.26 | 1315478.601 | 1288586.249 | 1241120.708 | 1237870.534 | 1250404.908 | 1234589.714 | 1227062.382 |
| 1H-indol-4-ol | 381350.9361 | 320350.8391 | 306797.9378 | 351250.8981 | 686287.1862 | 706508.2285 | 664265.9246 | 643513.6101 | 715014.6493 | 559171.7249 | 546803.1548 | 564735.2839 |
| IS1 | 12163325.98 | 13780518.24 | 10188499.77 | 10861368.8 | 13023054.99 | 11481493.49 | 12742180.43 | 11810031.38 | 12437414.94 | 13315515.57 | 10880299.97 | 10170339.51 |
| Methanone, (8-bromo-1-naphthalenyl)(1-pentyl-1H-indol-3-yl)- | 68892.87571 | 29639.88527 | 138353.8044 | 61960.20507 | 336145.8641 | 321556.0109 | 375813.9535 | 275605.886 | 274214.578 | 260442.474 | 265134.6249 | 166133.461 |
| PI 18:0_18:0;2O | 178385.7595 | 177140.3269 | 43195.06518 | 126246.8971 | 489762.0729 | 285505.7 | 764698.0257 | 511684.605 | 1072547.325 | 781277.2859 | 421738.4107 | 555908.0706 |
| 3-Oxo-4-cholestenoic acid | 2327617.671 | 360162.4414 | 3091771.094 | 264195.493 | 7857202.349 | 4686055.768 | 191106.8918 | 4194830.47 | 1409063.265 | 2018930.517 | 2538402.054 | 2938178.83 |
| 1-Methoxy-4-(phenylsulfonyl)benzene | 1196965.758 | 689400.9308 | 1075481.609 | 1250305.545 | 1356976.337 | 1164710.963 | 2115216.456 | 1387504.176 | 1277260.748 | 1007460.734 | 952867.3558 | 1462711.028 |
| Licorisoflavan A | 0 | 0 | 0 | 0 | 591833.9723 | 1091047.82 | 619814.3152 | 959190.5749 | 395282.9725 | 127515.0854 | 574603.5963 | 470491.2504 |
| PC O-22:5_20:4 | 0 | 0 | 0 | 0 | 1131491.685 | 849783.4258 | 791517.9795 | 945953.6886 | 407189.4379 | 305700.5253 | 290511.0349 | 276604.3204 |

| Metabolites | PBS_1 | PBS_2 | PBS_3 | PBS_4 | BCG_1 | BCG_2 | BCG_3 | BCG_4 | BCGΔ*pe18*_1 | BCGΔ*pe18*_2 | BCGΔ*pe18*_3 | BCGΔ*pe18*_4 |
|---|---|---|---|---|---|---|---|---|---|---|---|---|
| PE O-18:5_6:0 | 650996.0694 | 678788.8582 | 1033932.807 | 363121.2225 | 1018199.689 | 2135289.371 | 901371.0346 | 1021538.097 | 1155409.136 | 1081084.65 | 1249904.964 | 245320.0045 |
| Amastatin | 0 | 0 | 0 | 0 | 161655.4682 | 109773.7954 | 186724.1447 | 138534.7865 | 24562.08471 | 186598.0674 | 185340.3465 | 40216.91195 |
| PC O-31:1 | 950879.9751 | 436336.7254 | 1102821.58 | 673855.4818 | 808167.7341 | 416725.1834 | 693016.9566 | 506785.2254 | 497816.197 | 912836.3807 | 738985.652 | 611693.5819 |
| Derenofylline | 207217.6314 | 1802738.555 | 377041.6393 | 786658.0059 | 849502.123 | 123506.7672 | 830309.1787 | 1061666.891 | 198684.7114 | 1072541.26 | 196591.0067 | 663350.839 |
| Cordycepin | 7475705.384 | 8755795.751 | 342544.7164 | 7545135.332 | 8284309.279 | 6700079.309 | 4387238.841 | 4890008.777 | 8002116.805 | 4944583.601 | 2906766.459 | 8233514.289 |
| LPE 19:1 | 202692.6029 | 308738.1465 | 144873.1583 | 150312.0984 | 649591.8025 | 610953.7544 | 642739.9668 | 641439.3492 | 730993.198 | 905531.7533 | 893435.2783 | 648395.7034 |
| PG(18:1/18:1(9Z)) | 5820792.297 | 8425743.34 | 3952012.599 | 8184182.132 | 60366693.35 | 43295624.19 | 66132104.26 | 56390401.09 | 38954485.71 | 111486442.4 | 84176215.47 | 26266693.43 |
| 3H-Benzimidazol-5-yl-[4-[(5-cyclopropyl-1H-pyrazol-3-yl)amino]pyrimidin-2-yl]amine | 542503.6599 | 2493957.759 | 253699.4918 | 1558436.779 | 1975549.702 | 1385421.311 | 120712.6092 | 97160.91487 | 476138.0595 | 1271258.119 | 1075160.272 | 46648.72534 |
| trans-Crocetin | 16152213.21 | 45074900.09 | 89187764.3 | 55745485.56 | 10790922.34 | 7288423.665 | 18793415.72 | 4616528.965 | 15308842.98 | 21226034.34 | 16401434.79 | 144586160.8 |
| Tetrapentylammonium cation | 10279680.98 | 14821473.15 | 2978231.63 | 9721273.709 | 6354749.5 | 6275882.743 | 2969686.896 | 7426192.73 | 1017681.986 | 4188304.034 | 2683070.737 | 5717050.577 |
| FA 24:2 | 810794. | 653578. | 104046 | 756789. | 416463. | 493658. | 498855. | 418922. | 349391. | 333208. | 432905. | 320835. |

| **Metabolites** | **PBS_1** | **PBS_2** | **PBS_3** | **PBS_4** | **BCG_1** | **BCG_2** | **BCG_3** | **BCG_4** | **BCGΔ*pe18*_1** | **BCGΔ*pe18*_2** | **BCGΔ*pe18*_3** | **BCGΔ*pe18*_4** |
|---|---|---|---|---|---|---|---|---|---|---|---|---|
| | 6068 | 7856 | 1.596 | 6456 | 2116 | 3908 | 4831 | 9104 | 7027 | 8461 | 9623 | 7946 |
| 1-[(1R,5S)-8-Azabicyclo[3.2.1]oct-3-yl]-2-methyl-1H-benzimidazole | 103287.9595 | 1187862.37 | 3402265.033 | 1935768.816 | 49140.29521 | 46541.94396 | 86413.67058 | 38696.1128 | 308799.7502 | 387683.9717 | 59734.43518 | 6314658.451 |
| 2-(2,5-Dichlorophenyl)-5-methyl-1,2-dihydro-3H-pyrazol-3-one | 253710.5642 | 233772.2597 | 475602.4097 | 300793.0162 | 4012544.436 | 2502411.016 | 3250486.009 | 2735591.146 | 3716267.888 | 8138493.768 | 3011594.719 | 2544816.973 |
| Carbamylcholine cation | 1322264.914 | 904139.7108 | 2822078.613 | 1005439.618 | 2233867.202 | 1733351.829 | 1748086.958 | 2694334.537 | 1754005.553 | 1713664.348 | 1600771.386 | 1822481.158 |
| MGDG O-16:3_3:0 | 2397543.87 | 2221236.892 | 2142961.505 | 2385452.317 | 2551893.91 | 2160426.933 | 2429917.38 | 2744649.831 | 2388315.651 | 2648391.315 | 2480265.333 | 2273070.457 |
| (-)-trans-4-[4-(4'-Fluorophenyl)-3-piperidinylmethoxy]-2-methoxyphenol | 212465.2557 | 259108.3223 | 129629.7958 | 107039.0745 | 151161.961 | 325953.414 | 4091771.234 | 432176.6751 | 748243.3433 | 467191.6362 | 271654.1429 | 870559.2711 |
| 7-Bromopyrrolo[2,1-f][1,2,4]triazin-4-ylamine | 293608.5737 | 227509.6552 | 476580.9782 | 324360.277 | 2465018.008 | 1915317.564 | 2307232.92 | 2187793.03 | 2343335.638 | 1758301.852 | 1904844.858 | 1998955.06 |
| Kasugamycin | 4159761.528 | 4022966.318 | 3070679.916 | 5055230.38 | 3410250.541 | 4543318.245 | 3352386.87 | 5444625.86 | 4023933.586 | 5546951.395 | 3371873.222 | 3746478.642 |
| PS 41:4 | 0 | 0 | 0 | 0 | 662117.3816 | 869186.2141 | 584890.1105 | 654423.4522 | 912618.3353 | 827782.4331 | 690308.7711 | 771091.187 |

| Metabolites | PBS_1 | PBS_2 | PBS_3 | PBS_4 | BCG_1 | BCG_2 | BCG_3 | BCG_4 | BCGΔ*pe18*_1 | BCGΔ*pe18*_2 | BCGΔ*pe18*_3 | BCGΔ*pe18*_4 |
|---|---|---|---|---|---|---|---|---|---|---|---|---|
| MGDG O-14:1_28:7 | 0 | 0 | 25030.97609 | 0 | 1687745.483 | 1469917.524 | 1570892.152 | 1929795.367 | 601783.3996 | 488207.3863 | 567722.3672 | 509466.8727 |
| PC(22:2(13Z,16Z)/0:0) | 20978857.82 | 21282465.79 | 20372681.63 | 21171668.58 | 24725731.08 | 20728145.72 | 19951385.23 | 25049874.19 | 21321376.34 | 20126315.25 | 20443502.13 | 18696655.82 |
| Dorsomorphin | 493080.3849 | 433775.0855 | 327550.9611 | 434121.6151 | 515660.9573 | 453077.512 | 465418.9709 | 434373.6176 | 482790.936 | 442035.2935 | 446731.3179 | 656000.5396 |
| N-Acetyl-D-galactosaminitol | 564742.5409 | 470682.9812 | 854701.4245 | 423808.8843 | 675957.0751 | 365277.8056 | 663559.1821 | 489452.6813 | 289186.8543 | 457320.5209 | 446340.025 | 291314.0691 |
| Pentadecanedioylcarnitine | 378669.911 | 374232.8907 | 462274.9935 | 411164.3405 | 394658.6828 | 391275.0153 | 394289.8636 | 388296.6719 | 383423.0813 | 348394.1088 | 360952.7372 | 385023.3803 |
| Virolongin_B | 323318.8584 | 518707.9959 | 549413.9267 | 443706.6777 | 472618.1998 | 466192.0735 | 257313.3982 | 312351.115 | 418005.1865 | 429390.898 | 383545.7126 | 407666.6111 |
| 5-[(2-Phenylethyl)amino]-1,3,4-thiadiazole-2(3H)-thione | 566019.5689 | 345001.0763 | 770147.7223 | 402627.9412 | 1975201.046 | 974631.3836 | 1800378.724 | 1301762.61 | 1199454.587 | 1136089.221 | 1328274.517 | 1031834.228 |
| Dye X-5380-110B | 1213245.592 | 1151514.525 | 1858032.742 | 1025365.559 | 1199668.21 | 1227611.145 | 2684570.2 | 1311755.072 | 1266900.384 | 1862729.833 | 1072379.056 | 1080011.852 |
| Stearamide | 81075175.06 | 70161624.91 | 39404454.55 | 32789326.6 | 48240922.19 | 32925212.5 | 42443023.53 | 47275689.03 | 45414234.34 | 57942333.35 | 43726843.96 | 23623531.58 |
| Glucosylceramide_(d18:1/22:0) | 4816689.157 | 4962497.892 | 5229172.352 | 5436616.749 | 4791429.041 | 6436983.471 | 5240575.509 | 6130393.85 | 5672072.141 | 6601298.693 | 5564222.221 | 3744305.732 |
| SM 8:0;2O/26:2 | 1087608.152 | 371937.2076 | 1372483.515 | 764874.025 | 503368.7616 | 4629.882768 | 417990.1994 | 185896.1292 | 647815.3804 | 404590.9395 | 572680.7353 | 360337.6716 |
| M533T457 | 232865. | 265039. | 170948. | 191454. | 214245. | 245687. | 233424. | 194845. | 222465. | 212748. | 211222. | 185498. |

| Metabolites | PBS_1 | PBS_2 | PBS_3 | PBS_4 | BCG_1 | BCG_2 | BCG_3 | BCG_4 | BCGΔ*pe18*_1 | BCGΔ*pe18*_2 | BCGΔ*pe18*_3 | BCGΔ*pe18*_4 |
|---|---|---|---|---|---|---|---|---|---|---|---|---|
| | 7277 | 8183 | 0709 | 3946 | 0496 | 2298 | 1906 | 0213 | 1875 | 0885 | 5359 | 1182 |
| 4-Methylpiperidine-1-carboximidamide | 1148061.315 | 2240772.535 | 1282283.973 | 2572645.148 | 3415138.434 | 1275318.571 | 493112.6995 | 435963.9369 | 808698.1033 | 2252865.192 | 717629.3413 | 559284.9469 |
| Pipereicosalidine | 3309427.326 | 2936182.582 | 2162105.143 | 2711216.641 | 2977605.288 | 2926432.15 | 3433343.646 | 3024605.017 | 2902276.021 | 3728457.426 | 2978623.423 | 4358210.737 |
| 1-(1-Methoxy-1-methylethyl)-4-methylbenzene | 1017845.794 | 919375.5737 | 1337377.717 | 4666863.008 | 1000637.845 | 1568417.42 | 6598548.15 | 1383816.553 | 1998877.626 | 1610304.176 | 1740348.798 | 4538904.29 |
| Gliocladic acid | 2318940.485 | 3141885.9 | 2009149.11 | 2248470.716 | 2205428.171 | 2116621.114 | 2097028.575 | 2198811.252 | 2271852.802 | 2361557.378 | 2039621.243 | 2272413.35 |
| 3-Methyl-1H-pyrazole-4-carbaldehyde | 2330895.083 | 2008699.321 | 3399763.074 | 2753233.939 | 4495965.807 | 5595677.308 | 4325100.844 | 7075527.14 | 930484.0586 | 3208835.311 | 2334402.587 | 1074517.673 |
| N-(1H-Tetraazol-5-yl)decanamide | 2038764.345 | 2124914.777 | 2305067.808 | 2384513.964 | 2514484.68 | 1912072.309 | 1943512.817 | 1476490.231 | 5244734.457 | 2526110.608 | 2654809.583 | 2783843.797 |
| PC 16:0_33:0 | 258172.5491 | 154471.7135 | 187362.8355 | 168776.0534 | 179318.4047 | 0 | 246539.974 | 385220.7935 | 126311.7243 | 173449.996 | 202130.8893 | 278245.6288 |
| PC O-18:1_20:4 | 3553825.602 | 2908768.357 | 4190275.789 | 2487761.594 | 2721458.368 | 114474.1543 | 3644848.979 | 48833737.63 | 2701558.561 | 2901821.23 | 1784302.002 | 15036528.88 |
| 11-Nor-9-carboxy-.DELTA.9-tetrahydrocannabinol | 898306.7218 | 1010841.005 | 901493.0835 | 896371.4379 | 1031589.021 | 840825.8446 | 918347.7029 | 921644.474 | 843423.1834 | 968760.2024 | 987595.2729 | 919757.4171 |

| Metabolites | PBS_1 | PBS_2 | PBS_3 | PBS_4 | BCG_1 | BCG_2 | BCG_3 | BCG_4 | BCGΔ*pe18*_1 | BCGΔ*pe18*_2 | BCGΔ*pe18*_3 | BCGΔ*pe18*_4 |
|---|---|---|---|---|---|---|---|---|---|---|---|---|
| 1H-Indol-3-yl 6-O-((2R,3R,4R)-3,4-dihydroxy-4-(hydroxymethyl)tetrahydrofuran-2-yl)-.beta.-D-glucopyranoside | 511697.068 | 420014.3289 | 575562.9489 | 544287.9468 | 1462928.17 | 444515.6436 | 1029747.966 | 968892.0614 | 789576.4964 | 1063763.711 | 664379.8278 | 713350.5869 |
| 3,4-Dihydroxy-5-((6-O-((4-(2-hydroxypropan-2-yl)cyclohex-1-en-1-yl)carbonyl)-.beta.-D-glucopyranosyl)oxy)benzoic acid | 429338.5276 | 553806.4286 | 2198236.375 | 519601.1536 | 323992.0645 | 305166.3267 | 306410.0505 | 233299.2662 | 274702.8127 | 338469.9606 | 321499.6182 | 1529129.273 |
| PC O-33:7 | 983462.9594 | 1044168 | 774681.7078 | 944288.178 | 741390.7099 | 1055195.841 | 904443.6931 | 725378.6454 | 1124666.75 | 633129.6287 | 875243.2745 | 972043.7829 |
| myo-Inositol | 1028432.193 | 856049.2809 | 1242556.068 | 1007088.063 | 3056993.045 | 1928882.924 | 2975067.846 | 2363466.169 | 2374965.805 | 2083361.771 | 1838593.79 | 2417038.927 |
| PC(18:0/18:1(9Z)) | 82612091.82 | 97449947.47 | 115177412.8 | 86665696.98 | 244881056.6 | 163780386.5 | 259884226.9 | 192622678.8 | 212322847.9 | 265237400.1 | 223356010.4 | 106788762.4 |
| SPB 19:1;2O | 11319881.07 | 11367371.38 | 10205061.74 | 9538327.305 | 13770234.67 | 10610009.76 | 10472698.43 | 10411436.31 | 12860342.85 | 9872237.086 | 10327992.47 | 8821669.08 |
| N-(Quinolin-8-yl)thiophene-2-sulfonam | 1470822.164 | 1226487.249 | 1694686.078 | 1535281.03 | 1663035.916 | 1733408.778 | 1616599.951 | 1696720.296 | 1638034.479 | 1344258.565 | 1330192.31 | 1530932.727 |

| Metabolites | PBS_1 | PBS_2 | PBS_3 | PBS_4 | BCG_1 | BCG_2 | BCG_3 | BCG_4 | BCGΔ*pe18*_1 | BCGΔ*pe18*_2 | BCGΔ*pe18*_3 | BCGΔ*pe18*_4 |
|---|---|---|---|---|---|---|---|---|---|---|---|---|
| ide | | | | | | | | | | | | |
| 3-Hexanone | 6246110.916 | 3346166.556 | 776480.9917 | 3254615.434 | 3445837.07 | 2943792.625 | 3005224.013 | 3191726.209 | 3184995.434 | 3122906.356 | 3239148.118 | 3393429.322 |
| 1,2-dibutyrin | 457456.0061 | 457892.5375 | 712535.6005 | 596968.1473 | 474944.355 | 585627.089 | 477586.2727 | 454542.2704 | 485899.2767 | 446768.5784 | 590041.0454 | 1070061.019 |
| 5-[(1S,2R,4aR)-5-(Hydroxymethyl)-1,2,4a-trimethyl-2,3,4,7,8,8a-hexahydronaphthalen-1-yl]-3-methylpentanoic acid | 1568354.17 | 1284665.439 | 1606145.428 | 1232848.035 | 4688506.82 | 4963253.634 | 273262.1697 | 4282349.382 | 2916963.012 | 1850999.187 | 3833061.875 | 2628820.653 |
| DOCOSANOL | 1261420.145 | 893798.6979 | 820927.6432 | 654523.2638 | 281828.401 | 503675.1189 | 351519.4111 | 330596.1311 | 385233.5337 | 556255.6647 | 541824.2533 | 508835.6432 |
| [6]-Gingerdiol_5-O-beta-D-glucopyranoside | 462579.9592 | 470129.9773 | 470731.3432 | 414380.2936 | 1019793.287 | 777125.8394 | 342302.6574 | 383570.782 | 319096.5516 | 289718.1562 | 380515.6963 | 743629.119 |
| LPC O-15:0 | 753840.0688 | 698286.7507 | 639233.052 | 558988.6042 | 515802.2003 | 580872.4945 | 490760.0905 | 541561.8825 | 563446.0739 | 522176.1209 | 543657.8435 | 443972.9161 |
| MGDG O-13:1_2:0 | 45803.20911 | 65447.19832 | 64176.61609 | 50634.87553 | 75567.83297 | 41514.87632 | 86626.61817 | 54936.58822 | 37783.04114 | 61595.53367 | 27452.25333 | 8559.501905 |
| PE 18:1_20:4;O | 409988.7135 | 2827161.349 | 408355.4628 | 3315487.834 | 1160827.075 | 1177151.927 | 430790.1871 | 417841.9975 | 481644.1291 | 1793116.2 | 569630.7184 | 248282.7932 |
| PE 16:0_22:5 | 681537. | 579375. | 762596. | 583862. | 206936 | 192594 | 157905 | 203406 | 131356 | 155813 | 180495 | 127903 |

| **Metabolites** | **PBS_1** | **PBS_2** | **PBS_3** | **PBS_4** | **BCG_1** | **BCG_2** | **BCG_3** | **BCG_4** | **BCGΔ*pe18*_1** | **BCGΔ*pe18*_2** | **BCGΔ*pe18*_3** | **BCGΔ*pe18*_4** |
|---|---|---|---|---|---|---|---|---|---|---|---|---|
| | 5397 | 2601 | 5746 | 0614 | 7.649 | 2.073 | 6.072 | 6.749 | 5.196 | 1.14 | 8.913 | 1.359 |
| Fenugreekine | 362669.0969 | 296976.2714 | 640269.0011 | 294942.628 | 501587.8186 | 352784.0779 | 453578.3581 | 369536.6873 | 452803.5973 | 332445.8943 | 322460.8078 | 325184.5029 |
| 4-Methylumbelliferone | 651859.1789 | 602039.6814 | 446410.5194 | 494383.2485 | 716084.2541 | 729891.3272 | 593050.4424 | 674518.4045 | 600441.3094 | 578060.196 | 458954.5642 | 574514.5431 |
| PE 18:0_18:1;3O | 226482.2082 | 109848.7231 | 251119.0646 | 106789.3683 | 209822.126 | 591706.6368 | 576792.0649 | 490871.7209 | 264354.0789 | 210502.7684 | 432334.5425 | 318011.2654 |
| 3,4-Thiophenedicarbonitrile | 150824.3428 | 195359.5598 | 828108.3681 | 445221.8496 | 394352.2345 | 480788.5083 | 492150.1941 | 381826.4218 | 419399.373 | 194180.7446 | 288631.0673 | 615729.2313 |
| M509T82 | 741442.0865 | 416445.1542 | 513576.9169 | 583645.0027 | 529628.8447 | 513957.2341 | 531189.7889 | 591672.4304 | 724359.6439 | 496214.3468 | 536462.1207 | 377491.6608 |
| MGDG O-21:1_28:6 | 2625835.819 | 3175932.466 | 1680771.064 | 2448749.637 | 1638297.879 | 1316343.882 | 1256670.628 | 1297302.158 | 1245215.119 | 819142.7608 | 1398721.697 | 246908.1616 |
| 3-Hydroxyvaleric acid | 202611.8169 | 207686.1217 | 296673.571 | 337891.3375 | 456211.808 | 320267.3982 | 438971.1186 | 422567.4387 | 383709.4382 | 1216362.522 | 1100701.373 | 892113.3174 |
| Ranolazine | 0 | 0 | 42155.14722 | 0 | 573236.2686 | 923612.2339 | 456647.0734 | 665898.735 | 475603.6476 | 163042.6785 | 706520.2851 | 194954.4067 |
| [7-(2-Hydroxypropan-2-yl)-4a-methyl-1-methylidene-2,3,4,5,6,7,8,8a-octahydronaphthalen-2-yl] acetate | 732766.1461 | 322536.7201 | 502242.5086 | 383354.5263 | 322264.1401 | 431195.3782 | 408547.396 | 341787.6536 | 383134.4572 | 356405.2207 | 402733.1814 | 405714.8503 |
| Urocanic acid | 600001 | 174541 | 560245. | 228675 | 714255. | 103120 | 823393. | 171725 | 765025. | 809863. | 878894. | 213124 |

| Metabolites | PBS_1 | PBS_2 | PBS_3 | PBS_4 | BCG_1 | BCG_2 | BCG_3 | BCG_4 | BCGΔ*pe18*_1 | BCGΔ*pe18*_2 | BCGΔ*pe18*_3 | BCGΔ*pe18*_4 |
|---|---|---|---|---|---|---|---|---|---|---|---|---|
| | 6.02 | 1.803 | 0746 | 9.666 | 1935 | 1.394 | 255 | 1.059 | 1714 | 0451 | 602 | 7.448 |
| Butyrolactone I | 891120.9706 | 1372419.222 | 1344859.344 | 1434977.127 | 494060.6518 | 1071331.293 | 636587.0503 | 538469.4162 | 424224.4984 | 469320.3914 | 743119.9506 | 1931421.892 |
| CORTISONE ACETATE | 11104.42333 | 7731.842856 | 27361.86073 | 0 | 1908863.455 | 1541652.952 | 699878.3732 | 1197737.529 | 390806.9424 | 416311.6298 | 407334.9849 | 554855.8047 |
| N-Methylglycine N,N-dimethylamide | 171847.2522 | 1836609.648 | 3323681.024 | 2730636.825 | 516004.0246 | 258468.5792 | 729374.8394 | 397896.1927 | 367338.6943 | 1747064.709 | 144893.3561 | 3882913.901 |
| SMGDG O-24:2_14:0 | 0 | 0 | 0 | 0 | 474002.7165 | 473560.9388 | 556988.5893 | 827349.9031 | 190485.6991 | 127688.9423 | 165449.1229 | 165567.0099 |
| Swertisin | 1447101.892 | 1375661.446 | 1415456.953 | 1445727.78 | 1414046.58 | 1416107.99 | 1418869.627 | 1438087.05 | 1413613.387 | 1325078.724 | 1418395.667 | 1424165.903 |
| PEtOH 20:3_22:6 | 128192.0783 | 175765.2098 | 73553.71479 | 87289.73775 | 152310.597 | 107049.211 | 217106.7649 | 85897.09751 | 75205.31093 | 378532.3417 | 195546.5999 | 132140.4008 |
| D-Glucosaminide | 0 | 38438.53526 | 243206.9685 | 32627.74666 | 268187.5914 | 442606.4824 | 426644.2593 | 328448.702 | 294423.5589 | 294436.1843 | 237200.6672 | 405205.3173 |
| 3-(4-Methoxyanilino)-5,5-dimethyl-2-cyclohexen-1-one | 1516570.009 | 1447054.871 | 1456459.936 | 1456734.303 | 1708540.42 | 1487347.374 | 1512631.251 | 1516902.49 | 1497740.454 | 1488987.083 | 1559549.346 | 1332776.193 |
| 4-(Pentafluorosulfanyl)aniline | 5021222.29 | 4408646.354 | 3984646.971 | 4542550.817 | 6611490.032 | 4353680.017 | 5278277.842 | 5356222.835 | 3798944.587 | 3698079.676 | 4503944.951 | 3460291.089 |

| Metabolites | PBS_1 | PBS_2 | PBS_3 | PBS_4 | BCG_1 | BCG_2 | BCG_3 | BCG_4 | BCGΔ*pe18*_1 | BCGΔ*pe18*_2 | BCGΔ*pe18*_3 | BCGΔ*pe18*_4 |
|---|---|---|---|---|---|---|---|---|---|---|---|---|
| (5-Acetyloxy-6-hydroxy-10a-methoxy-4,4,7,11b-tetramethyl-9-oxo-1,2,3,4a,5,6,6a,7,11,11a-decahydronaphtho[2,1-f][1]benzofuran-3-yl) acetate | 1986702.065 | 1941323.937 | 1648088.266 | 1725382.827 | 1341540.696 | 1551051.496 | 1497327.516 | 1742634.795 | 1325687.974 | 1809955.566 | 1549183.873 | 1537617.045 |
| M279T449 | 11847717.41 | 11890220.92 | 9659043.394 | 11002549.66 | 11188208.34 | 9973095.707 | 10142527.64 | 9887907.89 | 10649849.61 | 12358267.23 | 9709869.543 | 10951123.06 |
| Medroxyprogesterone | 340316.2279 | 478805.6551 | 443789.7598 | 444929.8486 | 1877981.063 | 2720309.132 | 1669198.179 | 1973425.415 | 2500256.493 | 728902.6089 | 1591409.321 | 2340227.602 |
| Dibenzylamine | 157579.3694 | 580396.7787 | 27762144.3 | 824533.3289 | 182031.1343 | 1622161.125 | 199569.4751 | 184167.8816 | 100043.592 | 146325.6856 | 164979.1819 | 409327.6042 |
| 5-Chloro-2-benzimidazolone | 5812791.499 | 1406388.62 | 9164923.05 | 4226374.786 | 5365172.602 | 3000999.378 | 1164731.261 | 4342411.414 | 2047633.389 | 4601696.276 | 2141707.438 | 2755342.035 |
| 2H-1-Benzopyran-2-one, 7-mercapto-4-methyl- | 614077.1017 | 1022967.295 | 1022762.397 | 989695.0741 | 1147143.943 | 1147627.602 | 1335902.327 | 922034.7619 | 1007897.705 | 1704737.435 | 503094.2128 | 1021723.553 |
| Glycodeoxycholic acid | 188605.1032 | 192399.9266 | 442060.881 | 246555.078 | 366464.2011 | 389201.018 | 463467.1173 | 343613.5413 | 300145.94 | 258527.0588 | 214514.3566 | 300549.0886 |
| Segetalin A | 1254396.931 | 736061.9854 | 1433617.511 | 748375.009 | 479860.0907 | 323856.9949 | 942976.3217 | 778299.5066 | 547209.3928 | 668671.5819 | 376145.9668 | 257150.1186 |

| **Metabolites** | **PBS_1** | **PBS_2** | **PBS_3** | **PBS_4** | **BCG_1** | **BCG_2** | **BCG_3** | **BCG_4** | **BCGΔ*pe18*_1** | **BCGΔ*pe18*_2** | **BCGΔ*pe18*_3** | **BCGΔ*pe18*_4** |
|---|---|---|---|---|---|---|---|---|---|---|---|---|
| 6-Hydroxy-2,6,10,10-tetramethyl-1-oxaspiro[4.5]decan-8-yl 6-O-((2R,3R,4R)-3,4-dihydroxy-4-(hydroxymethyl)tetrahydrofuran-2-yl)-.beta.-D-glucopyranoside | 779175.5124 | 1210671.551 | 610978.7947 | 699184.2661 | 505940.7406 | 1232112.638 | 548142.0352 | 430000.5205 | 581186.7321 | 490324.7402 | 666418.982 | 334430.3885 |
| 5-(4-Difluoromethylsulfanyl-phenylamino)-[1,3,4]thiadiazole-2-thiol | 1170687.03 | 869071.5279 | 1668421.84 | 1132692.09 | 844445.5398 | 1208359.759 | 691918.7486 | 919319.9919 | 1053586.63 | 544864.5658 | 766371.1115 | 929974.7032 |
| LNAPS 15:0/N-27:0 | 10106.83696 | 928315.4947 | 586373.585 | 724899.4903 | 307117.6065 | 9171148.544 | 399887.0786 | 664796.7458 | 531448.6128 | 631533.089 | 620429.2602 | 460142.6253 |
| 3-dehydropresphingofungin | 2843045.297 | 1106186.955 | 2581298.929 | 1550771.351 | 2980555.368 | 540441.5683 | 2697581.96 | 1105490.859 | 1194009.749 | 784379.2236 | 2385160.802 | 883440.9587 |
| N-Oleoylglycine | 1230411.12 | 1267330.673 | 1028321.154 | 1425157.132 | 1355123.609 | 1136671.712 | 1389879.639 | 1449527.458 | 1294617.533 | 1241986.926 | 1178560.923 | 1071305.779 |
| N-Eicosapentaenoyl Serine | 486802.9052 | 486153.7945 | 489769.8881 | 510887.6949 | 436700.4278 | 487814.4577 | 494491.8772 | 426720.5715 | 505145.0317 | 500055.2205 | 496621.7861 | 571494.6147 |
| LPS 19:2 | 392741.3322 | 365867.5673 | 410244.9165 | 395361.9415 | 715038.9071 | 756056.4564 | 831269.7243 | 747288.8515 | 796034.876 | 782847.8743 | 1136217.071 | 618802.4958 |

| Metabolites | PBS_1 | PBS_2 | PBS_3 | PBS_4 | BCG_1 | BCG_2 | BCG_3 | BCG_4 | BCGΔ*pe18*_1 | BCGΔ*pe18*_2 | BCGΔ*pe18*_3 | BCGΔ*pe18*_4 |
|---|---|---|---|---|---|---|---|---|---|---|---|---|
| LPI 20:1 | 319774.6471 | 284469.4106 | 232429.2379 | 290804.2232 | 1521424.357 | 1722509.36 | 1364583.41 | 1107819.582 | 1136743.008 | 1085749.236 | 1458807.056 | 929065.0897 |
| Prolyl-Serine | 3217374.232 | 2359688.803 | 2691052.301 | 3280735.146 | 3103590.515 | 2620572.664 | 3630442.298 | 3257107.945 | 3228719.131 | 3102271.646 | 2310908.173 | 3322793.473 |
| LPG 18:3 | 24404.28395 | 7019.690651 | 48794.11866 | 11029.57043 | 986336.7636 | 1047189.29 | 732159.9299 | 1038271.961 | 812819.2848 | 707621.4275 | 758198.224 | 890067.3809 |
| Clobazam | 761451.811 | 961256.2973 | 990906.3812 | 856794.1344 | 753894.4577 | 734408.9986 | 756516.7659 | 826133.1931 | 742048.4569 | 726236.5457 | 762315.7255 | 1049147.996 |
| LNAPS 24:5/N-16:0 | 8166880.549 | 7942485.945 | 12797879.38 | 9647171.963 | 7155636.937 | 1323679.817 | 6089391.91 | 6703136.794 | 9355819.736 | 7225412.505 | 10231487.15 | 8769642.723 |
| Kojic acid | 297016.9038 | 97100.51061 | 283188.4185 | 56108.70223 | 373124.1277 | 395670.1977 | 69914.51282 | 420510.6081 | 209964.7023 | 260740.1447 | 407795.3546 | 53765.53785 |
| DIHYDROJASMONIC_ACID,_METHYL_ESTER | 3774735.921 | 3713288.514 | 3755410.09 | 3701112.853 | 3914873.986 | 4378820.379 | 4141653.43 | 4495709.517 | 4004652.526 | 3892723.76 | 3885452.791 | 4010242.448 |
| 3,4-Methyleneazelaic acid | 889648.1908 | 791779.2503 | 1140651.218 | 814546.3717 | 1716820.468 | 1204161.077 | 6291928.892 | 1004240.176 | 550897.9953 | 3644164.219 | 639502.368 | 505651.5805 |
| Glycolithocholic acid | 252854.1656 | 190061.8432 | 32483.7041 | 241486.73 | 308885.8805 | 421529.958 | 663952.0342 | 447962.2601 | 333111.3049 | 467950.2065 | 409917.2525 | 351415.035 |
| SM 19:1;2O | 622901.0017 | 686361.2715 | 704589.7765 | 476074.471 | 565607.435 | 669907.1243 | 833247.8042 | 858006.2161 | 539504.3112 | 783154.3111 | 653203.0827 | 514791.5569 |
| PE-Cer 15:3;2O/24:2;O | 811334.3439 | 1094098.699 | 1050012.406 | 954751.3176 | 1040101.356 | 0 | 1229654.783 | 1322219.808 | 1024631.944 | 657573.9446 | 1143673.867 | 1493900.708 |

| Metabolites | PBS_1 | PBS_2 | PBS_3 | PBS_4 | BCG_1 | BCG_2 | BCG_3 | BCG_4 | BCGΔ*pe18*_1 | BCGΔ*pe18*_2 | BCGΔ*pe18*_3 | BCGΔ*pe18*_4 |
|---|---|---|---|---|---|---|---|---|---|---|---|---|
| 8-Bromo-3,7-dimethyl-3,7-dihydro-1H-purine-2,6-dione | 207482.288 | 204525.3287 | 282090.5074 | 187349.1882 | 416213.7407 | 314850.9467 | 479944.2173 | 371015.0605 | 270394.5538 | 514126.945 | 435538.3348 | 273794.7571 |
| (3E,5E)-2-(4-Methoxy-4-oxobutyl)nona-3,5-dienoic acid | 422995.6407 | 364814.9649 | 411926.4664 | 372539.5139 | 570725.1199 | 587001.8694 | 658002.76 | 675186.8972 | 527387.6862 | 545135.1093 | 590666.4838 | 306829.6071 |
| PA 18:1_18:2 | 411478.4955 | 299538.4129 | 265927.3193 | 236785.9523 | 186809.6473 | 225921.8356 | 303870.9911 | 261437.0459 | 282227.3041 | 238987.1997 | 198150.794 | 149980.9677 |
| PG 16:3_22:5 | 694909.905 | 843063.9277 | 557567.1163 | 1189989.769 | 2016093.964 | 924502.6639 | 1484997.006 | 1779167.297 | 2318123.785 | 2056739.783 | 1008036.484 | 901602.3208 |
| PS 35:0 | 19297.31455 | 0 | 0 | 0 | 3214504.686 | 3064848.235 | 3158667.469 | 2512430.008 | 3972373.433 | 3486789.256 | 4184767.377 | 2221541.839 |
| SHexCer 11:1;2O/12:1;O | 0 | 0 | 0 | 0 | 136905.189 | 99952.68427 | 76425.35802 | 67199.84768 | 214585.225 | 137092.9485 | 191520.9729 | 178836.9307 |
| 2-Chloro-6-fluorophenylalanine | 500510.0785 | 339833.1318 | 746123.3938 | 994551.5455 | 3339238.53 | 1575821.562 | 2434778.257 | 2374947.102 | 2535802.247 | 1664308.824 | 1716165.498 | 1549667.708 |
| 1-Ethynylcyclohexanamine | 542994.8862 | 581082.7674 | 560174.3944 | 1852423.646 | 496859.1827 | 603526.1394 | 2368530.158 | 431044.4857 | 1219002.252 | 807230.2714 | 486385.4338 | 2307982.206 |
| .alpha.-(1-Aminoethyl)-4-hydroxybenzyl alcohol | 1730027.115 | 1110564.552 | 1026897.954 | 1062108.64 | 967632.5149 | 1038782.34 | 1046997.013 | 1078124.867 | 1040572.119 | 1018475.356 | 994996.1438 | 943752.7883 |
| PI O-18:1_22:5 | 762067.3159 | 565708.9215 | 409559.7289 | 1160718.316 | 641521.9787 | 618666.443 | 555349.921 | 612738.7264 | 813578.7674 | 984155.8785 | 731595.8861 | 1188878.557 |

| Metabolites | PBS_1 | PBS_2 | PBS_3 | PBS_4 | BCG_1 | BCG_2 | BCG_3 | BCG_4 | BCGΔ*pe18*_1 | BCGΔ*pe18*_2 | BCGΔ*pe18*_3 | BCGΔ*pe18*_4 |
|---|---|---|---|---|---|---|---|---|---|---|---|---|
| Tetracosa-13,16-dienoylcarnitine | 832511.7102 | 7797062.829 | 2764245.058 | 2744007.123 | 3375179.703 | 3799985.19 | 3546878.581 | 886409.4781 | 2243964.07 | 2613477.001 | 2491237.966 | 527832.5362 |
| Piperochromenoic acid | 4185510.498 | 21030817.95 | 2960840.892 | 3379152.647 | 5705596.096 | 7316424.689 | 6938375.247 | 5802915.67 | 5838188.348 | 6108727.149 | 7078433.38 | 12591515.29 |
| SM 13:1;2O | 298211.2663 | 241846.1258 | 125984.8186 | 220036.4814 | 201691.1085 | 227737.2359 | 120538.5206 | 68906.07081 | 154637.0824 | 156283.9879 | 380107.5923 | 84113.29862 |
| Camelledionol | 1571928.367 | 1467585.77 | 1280250.46 | 1377820.155 | 1393513.599 | 1439106.413 | 1454479.661 | 1421206.17 | 1585762.332 | 1373070.503 | 1276564.912 | 1349664.71 |
| Ile-Leu | 654679.901 | 570747.0039 | 752705.4549 | 537415.9655 | 584405.7111 | 392442.771 | 544427.277 | 401921.2885 | 642002.5541 | 731994.4778 | 859688.6794 | 451306.4991 |
| PE 16:0_18:1;3O | 85540.6266 | 38838.95061 | 92115.63882 | 93024.61245 | 254254.7739 | 293432.0166 | 220270.1198 | 223649.452 | 179324.4452 | 202184.7673 | 224426.4342 | 174287.9947 |
| PC 16:4_28:7 | 2201935.86 | 2334945.778 | 869430.1658 | 1689980.432 | 2279703.374 | 2076724.122 | 1865774.014 | 2343794.631 | 3212183.397 | 2142676.359 | 1373276.425 | 2213762.058 |
| N-Octadecanoyl-L-homoserine lactone | 4099187.429 | 4179058.216 | 3007817.617 | 3709046.829 | 3898170.668 | 4026252.26 | 3639167.934 | 3726725.305 | 3547897.9 | 3949847.883 | 3842378.829 | 3446086.019 |
| PC(20:3(8Z,11Z,14Z)/14:1(9Z)) | 1017524.522 | 772545.2255 | 897810.5887 | 596897.5921 | 2608388.761 | 2055377.658 | 12355593.52 | 2901355.648 | 2830289.191 | 8371449.281 | 1693424.76 | 1929446.686 |
| Antibiotic TAN 1446A | 1166429.074 | 1135661.426 | 1118131.147 | 1135251.995 | 1063720.528 | 1106075.02 | 1064190.205 | 1094962.534 | 1100183.757 | 1024168.718 | 1065922.921 | 1012094.696 |
| 3,3-Dimethylglutaric acid | 487744.1141 | 433515.5341 | 382584.3484 | 438819.3679 | 2759399.109 | 1475477.106 | 2931381.034 | 2062198.103 | 2045316.961 | 603121.5778 | 834279.6532 | 915359.306 |
| Xanthine | 104733.358 | 107517.9923 | 114477.4718 | 133971.7315 | 1550326.016 | 861700.1338 | 349051.3165 | 1201422.325 | 988655.4293 | 151869.1795 | 1315035.655 | 188479.5862 |

| Metabolites | PBS_1 | PBS_2 | PBS_3 | PBS_4 | BCG_1 | BCG_2 | BCG_3 | BCG_4 | BCGΔ*pe18*_1 | BCGΔ*pe18*_2 | BCGΔ*pe18*_3 | BCGΔ*pe18*_4 |
|---|---|---|---|---|---|---|---|---|---|---|---|---|
| alpha-Solanine | 86657.92082 | 50382.69309 | 419269.4663 | 127933.7262 | 115345.4541 | 169301.6497 | 157907.8988 | 178717.7947 | 38126.78697 | 66824.49313 | 97791.36964 | 45476.30199 |
| PC(15:0/20:1(11Z)) | 420907.6185 | 158940.1697 | 473486.47 | 255382.723 | 1577914.338 | 1437597.372 | 1356052.503 | 1648600.343 | 1926865.014 | 1770460.166 | 1263467.638 | 1902062.819 |
| Curine | 0 | 0 | 9411.463703 | 0 | 177202.0987 | 109390.0618 | 179439.7279 | 129278.2605 | 138592.6709 | 116892.0745 | 126081.5962 | 157141.0754 |
| SHexCer 15:3;2O/24:2;O | 0 | 0 | 0 | 0 | 739950.1776 | 1460214.306 | 919178.2552 | 1068587.734 | 1231292.791 | 743637.4092 | 845240.942 | 777597.3044 |
| Dimethylditetradecylammonium cation | 372697.171 | 450495.5117 | 148848.5755 | 290804.4674 | 55175.59774 | 25982.34847 | 62525.32746 | 124506.5009 | 128144.5414 | 86459.16229 | 22941.26734 | 49894.58239 |
| M479T160 | 912235.8894 | 963522.6353 | 704408.8051 | 923730.2152 | 869690.9868 | 891649.6288 | 1219227.351 | 924798.9389 | 660885.3313 | 902600.4383 | 677381.2195 | 822771.5446 |
| (25S)-7-Dafachronic acid | 300138.7268 | 99831.49657 | 297635.7057 | 198433.3601 | 5847185.141 | 3698963.865 | 4084864.287 | 6418002.01 | 4338574.034 | 4480918.501 | 3984998.501 | 3634301.144 |
| Azeliragon | 288753.3994 | 230159.8267 | 550476.5883 | 251370.8807 | 330841.3851 | 399354.7413 | 627928.6753 | 620307.6427 | 524116.7821 | 376647.8418 | 495856.8526 | 437400.757 |
| agrocinopine B | 0 | 0 | 0 | 0 | 784247.1146 | 525882.8817 | 363644.5855 | 483997.1987 | 445814.2642 | 448836.6404 | 719137.8951 | 659244.1325 |
| Hexadecanedioic acid | 872204.9484 | 1045329.362 | 1515474.156 | 1345912.356 | 1272113.071 | 1311719.347 | 1535068.698 | 1155883.356 | 1083387.484 | 1014976.951 | 1109900.917 | 2076268.567 |
| 2-(2-Butoxyethoxy)acetic acid | 150241.9286 | 151925.2142 | 120386.5435 | 140029.6012 | 1396494.936 | 1310045.536 | 953278.1211 | 1187021.45 | 638790.863 | 410700.2469 | 472551.7061 | 589598.5723 |
| Gly-Pro-Arg-Pro-amide | 708024.9453 | 777045.7865 | 1690632.322 | 839478.3478 | 1175456.823 | 2251259.819 | 1140126.622 | 1507327.484 | 749604.0136 | 469805.3431 | 1032641.992 | 1238569.995 |

| Metabolites | PBS_1 | PBS_2 | PBS_3 | PBS_4 | BCG_1 | BCG_2 | BCG_3 | BCG_4 | BCGΔ*pe18*_1 | BCGΔ*pe18*_2 | BCGΔ*pe18*_3 | BCGΔ*pe18*_4 |
|---|---|---|---|---|---|---|---|---|---|---|---|---|
| DGGA 14:1_26:4 | 356042.2622 | 272538.3522 | 238385.7387 | 423444.3563 | 246280.7503 | 430751.8715 | 219763.421 | 245597.5246 | 252629.6214 | 77804.71363 | 280676.1374 | 326176.6192 |
| 7-Hydroxy-3-oxo-cholest-4-en-26-oic acid | 415485.7216 | 227007.004 | 114218.8147 | 59973.9007 | 1986168.931 | 1403982.596 | 1922787.85 | 1472962.964 | 817775.6538 | 974812.1955 | 1069489.439 | 923972.8539 |
| FA 16:4 | 711209.9081 | 1074883.655 | 736580.5317 | 831294.5103 | 1018883.958 | 957675.417 | 792172.3512 | 865115.6494 | 804350.4074 | 1190677.764 | 735429.1736 | 899550.6392 |
| SMGDG O-8:0_24:2 | 115957.3048 | 47741.26391 | 160631.9207 | 164198.1438 | 160577.4087 | 149741.5722 | 110118.3816 | 106093.3389 | 159043.9285 | 133300.4744 | 100270.1986 | 72447.54709 |
| Cer 21:3;3O/29:0;(2OH) | 0 | 0 | 0 | 0 | 46494.78033 | 29242.48293 | 60408.53717 | 40044.33418 | 21689.87289 | 9590.109195 | 32524.34994 | 47588.79204 |
| SM 23:1;2O/15:0 | 2520361.849 | 2957106.109 | 2794592.01 | 3502100.792 | 2387654.886 | 11687894.83 | 1989049.581 | 2794358.286 | 2993810.875 | 2655714.672 | 2223846.124 | 3031518.176 |
| 4-Acetamido-2-aminobutanoic_acid | 79085.7968 | 186892.4975 | 227137.5352 | 86406.76686 | 3606511.541 | 3671077.823 | 2466704.264 | 2702325.545 | 2941337.399 | 2006468.632 | 2529939.326 | 3020191.274 |
| PE O-13:1_2:0 | 624439.4285 | 1077889.932 | 438064.6978 | 847886.4789 | 742743.7008 | 341238.2447 | 418555.1857 | 400594.2025 | 422770.8934 | 400769.7501 | 351421.5754 | 91658.55138 |
| 1-Stearoylglycerophosphoserine | 528969.48 | 657524.314 | 323900.6546 | 565916.2235 | 534295.5438 | 287882.9047 | 573710.396 | 386332.4408 | 715535.6257 | 496012.0128 | 678819.1761 | 336027.7319 |
| 7-Amino-4-methyl-2H-1-benzopyran-2-one arachidonoyl amide | 284738.7955 | 232332.522 | 274825.6494 | 185778.5272 | 232512.5165 | 261613.7583 | 269345.09 | 0 | 320412.3863 | 316918.3164 | 314851.7895 | 183796.8219 |

| Metabolites | PBS_1 | PBS_2 | PBS_3 | PBS_4 | BCG_1 | BCG_2 | BCG_3 | BCG_4 | BCGΔ*pe18*_1 | BCGΔ*pe18*_2 | BCGΔ*pe18*_3 | BCGΔ*pe18*_4 |
|---|---|---|---|---|---|---|---|---|---|---|---|---|
| 4-Hydroxybenzaldehyde | 4827781.538 | 4546814.131 | 4376810.209 | 5521992.087 | 6802046.662 | 6911741.092 | 4234523.184 | 6761522.72 | 4336655.422 | 4380442.328 | 4307332.475 | 4419341.884 |
| .beta.-Alanine, N-[2-(2-pyridinyl)-6-(1,2,4,5-tetrahydro-3H-3-benzazepin-3-yl)-4-pyrimidinyl]-, ethyl ester | 989531.1743 | 1817364.775 | 914166.6979 | 986977.5962 | 1087569.983 | 1126413.673 | 1020969.654 | 1009533.622 | 1051138.196 | 1164470.291 | 1024617.059 | 929118.8653 |
| isoferulic acid | 1096108.305 | 1282399.457 | 1221608 | 1422770.448 | 1557040.368 | 1434289.774 | 1461575.527 | 1230212.835 | 1238003.735 | 1544332.818 | 1194935.76 | 2663240.525 |
| N-Acetyl-S-geranylgeranyl-L-cysteine | 147222.9525 | 0 | 315919.6884 | 165148.7006 | 773421.6147 | 472378.5588 | 777175.2678 | 575349.9555 | 420984.6896 | 636394.1054 | 736143.1171 | 238515.6385 |
| 7-(4-Methoxyphenyl)-2,4-pteridinediol | 729458.1261 | 734570.5723 | 668554.69 | 674946.9661 | 685115.01 | 665217.5959 | 679285.1642 | 718244.7924 | 752045.6667 | 673966.0614 | 691261.329 | 700899.4573 |
| 1-(1-Pyrrolidinyl)-2-propanone | 2498032.473 | 2079927.771 | 1637401.344 | 3494762.594 | 2207573.549 | 1662369.455 | 3961240.199 | 1608014.656 | 2393312.837 | 3005126.141 | 2034832.582 | 2599117.307 |
| Argininosuccinic acid | 318160.6819 | 326328.0299 | 353599.4151 | 325606.0368 | 1747037.708 | 1159240.655 | 1174921.105 | 1490213.805 | 1549857.196 | 1368321.053 | 1025507.113 | 1618448.977 |
| 4-Methylacetophenone | 772911.3318 | 683304.6909 | 782368.579 | 4289784.602 | 1238246.637 | 5817203.612 | 4348232.421 | 2469569.974 | 4661977.42 | 4816509.177 | 586701.7986 | 3464226.057 |
| N-Ethyl-4-menthane-3-carboxamide | 6395214.854 | 5536893.108 | 4203599.921 | 5013953.17 | 23598528.82 | 25031644.24 | 8703914.111 | 22419262.77 | 22656408.81 | 10317165.12 | 22936191.25 | 18292812.95 |
| LPC 21:0 | 2282348.041 | 2217491.759 | 2048509.341 | 2009996.227 | 2160327.201 | 2174956.871 | 2183613.256 | 2207831.454 | 2084806.142 | 2126620.713 | 2112762.806 | 2059619.576 |

| Metabolites | PBS_1 | PBS_2 | PBS_3 | PBS_4 | BCG_1 | BCG_2 | BCG_3 | BCG_4 | BCGΔ*pe18*_1 | BCGΔ*pe18*_2 | BCGΔ*pe18*_3 | BCGΔ*pe18*_4 |
|---|---|---|---|---|---|---|---|---|---|---|---|---|
| 3-(2-Amino-4-methyl-6-oxo-1,6-dihydro-5-pyrimidinyl)propanoic acid | 3704307.013 | 3064374.184 | 3874801.891 | 4154246.841 | 4095125.649 | 4599254.054 | 3609246.638 | 4348856.935 | 3992643.54 | 1518290.613 | 1654455.544 | 3855561.627 |
| 6-Chloro-N-cyclohexyl-1-methyl-1H-pyrazolo[3,4-d]pyrimidin-4-amine | 721647.6315 | 264845.453 | 619094.6034 | 243841.3764 | 572086.0692 | 712899.2671 | 660602.1555 | 653851.6097 | 398018.1141 | 814813.8575 | 786978.2314 | 251201.0833 |
| Cer 20:0;2O/4:0 | 4065951.259 | 3966200.965 | 2659783.657 | 4755996.713 | 2750644.11 | 2839847.719 | 3085138.438 | 3467028.683 | 4310243.911 | 4481970.281 | 3177262.984 | 2743518.405 |
| Dye X-16768-260-Ba | 0 | 0 | 0 | 0 | 152828.5763 | 137802.9377 | 128108.918 | 93706.07574 | 442091.3569 | 372240.9353 | 432035.1373 | 485131.8935 |
| 3,3'-Disulfanediyldibenzoic acid | 2235682.839 | 2954811.841 | 3921112.799 | 3177035.304 | 3208355.749 | 3786669.671 | 2535690.267 | 1625362.261 | 2273115.09 | 2303736.562 | 3567131.635 | 3552869.791 |
| 4-[[2-[(E)-3-Hydroxybut-1-enyl]-5-oxooxolan-3-yl]amino]benzaldehyde | 665209.7943 | 79149.46818 | 711960.6715 | 218301.8618 | 701636.1513 | 1280431.642 | 5690063.636 | 4117511.545 | 1628746.186 | 772821.7379 | 1252425.368 | 420332.7921 |
| (Desmethyl)hydroxycerivastatin | 720243.8112 | 588352.2127 | 650352.5397 | 552001.9468 | 561113.3173 | 507241.9536 | 655412.4833 | 627934.7908 | 508868.9651 | 701647.4498 | 443842.346 | 605281.2583 |
| N-(3-Chlorophenyl)-N'-(3-fluorophenyl)urea | 653120.4486 | 693641.1542 | 515459.4886 | 412940.7196 | 348488.6257 | 704480.7312 | 565091.5893 | 1290018.407 | 640096.8233 | 739023.8675 | 391848.0975 | 305893.8391 |
| Dicaffeoylputrescin | 505319 | 488805 | 495923 | 506100 | 499402 | 515917 | 517396 | 493319 | 496621 | 514746 | 498104 | 514484 |

| Metabolites | PBS_1 | PBS_2 | PBS_3 | PBS_4 | BCG_1 | BCG_2 | BCG_3 | BCG_4 | BCGΔ*pe18*_1 | BCGΔ*pe18*_2 | BCGΔ*pe18*_3 | BCGΔ*pe18*_4 |
|---|---|---|---|---|---|---|---|---|---|---|---|---|
| e | 1.489 | 4.127 | 4.544 | 8.313 | 9.104 | 9.004 | 8.557 | 6.462 | 0.286 | 5.5 | 3.387 | 3.419 |
| Cyclo-prolylglycine | 1513948.238 | 1415128.894 | 913026.3386 | 225572.4848 | 1183553.638 | 333475.137 | 1641617.067 | 1053470.537 | 2072792.595 | 368708.4804 | 422695.4364 | 1027992.551 |
| 2-(Hydroxymethyl)-6-methylpyridin-3-ol | 664098.7047 | 1039983.719 | 827073.3059 | 988500.2008 | 968440.6265 | 717723.7775 | 729829.0699 | 618277.9287 | 696168.5765 | 722587.6614 | 699177.5418 | 857277.8914 |
| Octafluorocyclobutane | 1124822.855 | 627577.323 | 1052073.423 | 783816.4613 | 1050631.133 | 1585074.953 | 2171386.802 | 1025597.186 | 708757.6679 | 830219.4217 | 1645710.246 | 4288316.969 |
| DLCL 16:0_16:1 | 614820.9413 | 569899.2178 | 142395.5977 | 573509.3678 | 205836.8178 | 64671.99172 | 383477.7078 | 188713.2523 | 368563.5094 | 187683.1452 | 284131.7407 | 277965.387 |
| GR 89696 | 25300.63795 | 11383.68147 | 38509.40327 | 14284.67057 | 159765.1628 | 111932.7969 | 174416.7714 | 154434.8501 | 103973.9373 | 99754.66482 | 143096.438 | 129234.1891 |
| .alpha.-Pyrrolidinooctanophenone | 4601909.105 | 4614885.852 | 2710931.456 | 4468343.989 | 4890078.121 | 4084009.943 | 3792896.661 | 4255447.964 | 4300341.823 | 4926318.909 | 4319562.162 | 3383972.755 |
| Taurohyodeoxycholic acid | 3455937.586 | 3194197.459 | 3754366.847 | 3263979.532 | 5894826.896 | 5201696.435 | 6431847.391 | 3372338.399 | 6596763.581 | 4795432.523 | 2891005.96 | 2630891.293 |
| FA 13:1 | 1797119.702 | 1002424.69 | 2319623.588 | 1971170.927 | 1573757.881 | 1991738.523 | 2138781.697 | 1221039.184 | 1368811.382 | 2719515.55 | 1660035.836 | 1573066.045 |
| Diospyrin | 4311.466513 | 3204.690055 | 38013.11129 | 0 | 190802.9134 | 272271.9067 | 149612.8175 | 227039.8595 | 271523.9451 | 303994.5223 | 427714.562 | 276006.9615 |
| 1-(Cyclododec-1-en-1-yl)ethanone | 1522859.671 | 1561216.106 | 1523163.269 | 1575746.647 | 952653.8219 | 1269296.149 | 1368315.053 | 1522768.259 | 1297399.025 | 1564297.203 | 1383612.255 | 1249805.713 |
| Pepstatin A | 1191520.655 | 1800696.494 | 3540886.241 | 1121171.076 | 20807428.28 | 14381359.15 | 22521417.56 | 17207048.57 | 18278589.02 | 9107772.443 | 18452197.8 | 9815537.591 |

| **Metabolites** | **PBS_1** | **PBS_2** | **PBS_3** | **PBS_4** | **BCG_1** | **BCG_2** | **BCG_3** | **BCG_4** | **BCGΔ*pe18*_1** | **BCGΔ*pe18*_2** | **BCGΔ*pe18*_3** | **BCGΔ*pe18*_4** |
|---|---|---|---|---|---|---|---|---|---|---|---|---|
| Adrenochrome | 4625012.434 | 3610525.786 | 4396007.119 | 5281073.896 | 5071900.423 | 5488214.064 | 3621561.288 | 4988908.176 | 4427134.181 | 1699556.495 | 1880432.772 | 4333549.946 |
| Creatinine | 536099.4076 | 918341.233 | 442473.3161 | 467878.5148 | 460334.6517 | 402574.7467 | 455087.9201 | 388964.222 | 507237.2811 | 485149.4223 | 391989.5204 | 361569.7681 |
| Chlorazol black E | 758283.1132 | 178977.6175 | 38846.12566 | 0 | 0 | 71202.20321 | 98356.99724 | 203423.0938 | 239614.8084 | 81229.0463 | 2985202.276 | 161312.9948 |
| Carbocyclic thromboxane A2 | 499498.6226 | 485840.7358 | 724952.9874 | 542257.8218 | 94235.82653 | 200133.5579 | 267990.7923 | 234727.9713 | 190286.8856 | 231524.8911 | 259120.9672 | 59950.41133 |
| PC O-22:3_14:1 | 552246.8974 | 300768.6616 | 564498.8409 | 688399.225 | 450999.9053 | 0 | 552895.5307 | 1067215.706 | 341514.1913 | 329633.3149 | 376245.029 | 473980.1456 |
| Dioscoretine | 7987699.569 | 9274864.206 | 10256996.73 | 11608743 | 7931776.277 | 8339833.976 | 7653848.06 | 8405374.255 | 5890612.33 | 7654932.988 | 6172295.395 | 11424399.87 |
| 1-indanol | 422688.9814 | 473285.5629 | 366911.3286 | 436723.8267 | 453942.7715 | 442699.218 | 462564.0214 | 374519.8373 | 410340.3563 | 390586.5213 | 426829.2956 | 400059.5398 |
| (4-Bromobutyl)(triphenyl)phosphonium cation | 624974.2035 | 676526.6459 | 633313.6909 | 630143.3285 | 634558.5965 | 638500.233 | 629572.0899 | 631355.1924 | 634548.9225 | 694434.9006 | 573971.682 | 627724.7981 |
| 1-Dodecyl-2-pyrrolidinone | 7746511.499 | 8792852.91 | 4810158.094 | 6766626.858 | 6135938.227 | 5822191.033 | 5578325.325 | 6486701.002 | 4823368.839 | 3726527.32 | 3964014.148 | 3050297.222 |
| 3-Hydorxy-3-methylglutaric acid | 5794597.234 | 5074979.532 | 6202967.405 | 5466758.029 | 18807261.68 | 11705800.64 | 14832493.45 | 14441027.75 | 14085240.73 | 8651745.934 | 9438729.931 | 9091649.386 |
| 26,27-Dinorcholest-5-en-24-yne-3,20-diol, (3.beta.)- | 360667.5952 | 242867.0392 | 319333.1307 | 347250.9608 | 1156008.607 | 657276.5504 | 1189101.474 | 934662.3889 | 450457.0369 | 752531.4253 | 732865.6599 | 362535.2661 |

| Metabolites | PBS_1 | PBS_2 | PBS_3 | PBS_4 | BCG_1 | BCG_2 | BCG_3 | BCG_4 | BCGΔ*pe18*_1 | BCGΔ*pe18*_2 | BCGΔ*pe18*_3 | BCGΔ*pe18*_4 |
|---|---|---|---|---|---|---|---|---|---|---|---|---|
| cis-9-Palmitoleic acid | 80035013.65 | 79804049.04 | 73262335.81 | 64718096.93 | 87777968.53 | 115722510.8 | 101358063.9 | 100173445.1 | 71036271.6 | 71913684.15 | 88261273.18 | 71099174.85 |
| 9-Benzyl-9-azabicyclo[3.3.1]nonan-3-one | 916873.2038 | 952101.6288 | 879321.0393 | 861921.2145 | 861289.4321 | 845895.0317 | 953859.2222 | 951883.6725 | 855791.8736 | 867191.5226 | 845863.8332 | 902521.6754 |
| MGDG O-13:1_22:4 | 372839.9053 | 457393.5346 | 130873.0101 | 300887.4269 | 533707.2953 | 180133.6077 | 368510.6995 | 363624.1711 | 659583.8047 | 46467.84911 | 34074.85641 | 353758.613 |
| LPE 24:4 | 409136.3912 | 1083259.616 | 815175.8524 | 545384.2856 | 1251091.412 | 1145945.055 | 1081844.423 | 1073912.297 | 1233169.846 | 1755553.412 | 1246772.877 | 786113.2604 |
| N-(1-Allylcyclohexyl)-2,6-diethylaniline | 1785457.573 | 1956697.23 | 1592084.999 | 1639066.653 | 1796960.656 | 1825170.528 | 1794093.815 | 1701451.583 | 1880365.129 | 1784235.008 | 1609372.779 | 1685124.172 |
| 2-Pyrrolidinone, 5-[(1E,3R)-4,4-difluoro-3-hydroxy-4-phenyl-1-buten-1-yl]-1-[6-(2H-tetrazol-5-yl)hexyl]-, (5R)- | 269774.4695 | 243232.4029 | 296606.1606 | 257482.2112 | 266515.6326 | 244654.7678 | 240955.5656 | 241824.2021 | 255209.2313 | 279464.076 | 263857.7142 | 228847.8956 |
| 3-(5-methoxy-1H-indol-3-yl)propanoic acid | 1118612.252 | 1015752.508 | 1420394.573 | 1041479.091 | 3588882.939 | 2505845.487 | 3760004.26 | 2978723.202 | 2549438.724 | 2771936.25 | 2883644.558 | 1275863.159 |
| 2-Aminoisobutyric acid | 1256884.019 | 1288692.913 | 234874.1517 | 503330.9591 | 1101237.502 | 839997.8496 | 1451609.977 | 861860.94 | 882256.7403 | 864681.7414 | 957588.9364 | 849060.738 |

| Metabolites | PBS_1 | PBS_2 | PBS_3 | PBS_4 | BCG_1 | BCG_2 | BCG_3 | BCG_4 | BCGΔ*pe18*_1 | BCGΔ*pe18*_2 | BCGΔ*pe18*_3 | BCGΔ*pe18*_4 |
|---|---|---|---|---|---|---|---|---|---|---|---|---|
| 1-Linoleoyl-2-oleoyl-rac-glycerol | 77212.5365 | 93360.08657 | 79897.25181 | 59968.3146 | 65373.8351 | 66005.23188 | 67832.69645 | 71838.5676 | 80371.53111 | 62605.33655 | 90800.55561 | 60215.10101 |
| 5-(Tetradecyloxy)-2-furoic acid | 1476426.858 | 1696928.706 | 153100.9415 | 257563.1432 | 826223.1277 | 1855193.649 | 853483.7304 | 915356.0655 | 756406.3488 | 594717.849 | 935892.623 | 992767.2081 |
| arborinine | 30736046.56 | 11834778.6 | 21702485.06 | 11226920.12 | 617306.0227 | 8513849.388 | 135193828.2 | 138143227.7 | 16394919.04 | 5751323.057 | 21576252.11 | 8830130.52 |
| delta-Tocotrienol | 895934.2424 | 827006.7387 | 751458.98 | 1272078.913 | 818664.5506 | 1099652.316 | 841730.1028 | 894613.9774 | 800391.6193 | 817966.295 | 1164704.94 | 682537.7713 |
| 6a-Hydroxy-9,10a-dimethoxy-2-(prop-1-en-2-yl)-1,2,12,12a-tetrahydrochromeno[3,4-b]furo[2,3-h]chromene-6,8(6aH,10aH)-dione | 523896.9794 | 382222.5657 | 699380.1968 | 616521.2884 | 414595.8478 | 510339.7587 | 501103.0242 | 637432.4372 | 508167.2836 | 625185.879 | 364684.1841 | 605036.6369 |
| SM 37:4;3O | 12519988.61 | 11637382.03 | 8329267.528 | 9416749.189 | 1263985.947 | 1235560.648 | 2009636.375 | 1462793.972 | 3547086.189 | 1910287.154 | 1534503.189 | 2881000.979 |
| 1-Methyl-4-piperidinecarboxylic acid | 7808717.552 | 5604446.229 | 10178782.89 | 6959554.252 | 10874474.02 | 7709744.484 | 12239432.51 | 10125209.29 | 7121802.704 | 8778877.951 | 9531434.293 | 5977316.333 |
| 5,13,15-Trihydroxy-9-methyl-10-oxabicyclo[10.4.0]hexadeca-1(12),13,15-triene-3,11-dione | 568308.0009 | 418111.6308 | 851625.2681 | 578748.6002 | 1908317.193 | 1029710.678 | 1925161.409 | 1327229.64 | 1109007.629 | 1386715.752 | 1554679.522 | 916992.8139 |

| Metabolites | PBS_1 | PBS_2 | PBS_3 | PBS_4 | BCG_1 | BCG_2 | BCG_3 | BCG_4 | BCGΔ*pe18*_1 | BCGΔ*pe18*_2 | BCGΔ*pe18*_3 | BCGΔ*pe18*_4 |
|---|---|---|---|---|---|---|---|---|---|---|---|---|
| 1H-Indol-4-ol, 3-[2-(dimethylamino)ethyl]-, 4-acetate | 196309.6707 | 187408.977 | 194738.4529 | 210418.6224 | 676828.3474 | 448790.7503 | 459015.9024 | 555457.2228 | 499852.8619 | 313156.2063 | 322146.1471 | 459916.4038 |
| 2-(Acetyloxy)-3-methylbenzoic acid | 955788.091 | 1083383.433 | 934458.1925 | 982251.0669 | 1030053.87 | 1007439.18 | 899230.6741 | 936386.6843 | 931711.905 | 957854.0129 | 912969.0213 | 990310.439 |
| 3-Butynoic acid | 1741963.973 | 2166108.273 | 1978487.376 | 2038701.481 | 12550061.54 | 8477698.419 | 13643778.36 | 10021996.56 | 8784075.562 | 12896023.77 | 13053050.29 | 5468930.178 |
| 6'-O--D-Apiofuranosylsweroside | 3479.178554 | 0 | 0 | 0 | 125870.5731 | 91867.74579 | 101231.7148 | 98436.81515 | 756523.2798 | 516495.7792 | 661907.3259 | 874685.8392 |
| Ramipril | 109305.5407 | 112635.3959 | 126561.8497 | 112768.4635 | 142072.6508 | 155567.5407 | 150189.9236 | 135636.0307 | 153764.1408 | 163330.7262 | 142196.0535 | 154886.9657 |
| limonin | 455950.5573 | 663256.9739 | 304783.2345 | 493888.2837 | 1564481.485 | 567050.1397 | 985037.9659 | 757220.2302 | 689813.8261 | 737025.6243 | 467695.4172 | 534603.0669 |
| Cer 9:1;2O/24:6 | 254375.8447 | 336412.669 | 333337.3426 | 324327.8539 | 287398.1033 | 239168.0469 | 214902.8594 | 266268.5069 | 248821.7154 | 245092.8925 | 250534.2574 | 222824.9414 |
| 2',4'-Dihydroxyacetophenone | 915165.9202 | 740263.8129 | 782693.1935 | 932905.0078 | 944553.2178 | 785908.9465 | 923158.334 | 943421.5404 | 854376.671 | 933475.4828 | 783359.2076 | 832788.861 |
| 3-Furoic acid | 374652.415 | 540314.5938 | 558916.1057 | 309896.9059 | 752488.3637 | 669824.5203 | 779144.581 | 698228.6847 | 1554601.242 | 1420264.163 | 2456953.49 | 1362115.455 |
| Methylxanthoxylin | 877688.515 | 815360.2448 | 822466.3023 | 834922.4112 | 832293.0828 | 1272728.951 | 849508.5087 | 1287266.459 | 817056.3555 | 1179700.955 | 858874.6017 | 803102.7503 |
| PA 12:0_20:5 | 1390604.776 | 1318958.759 | 731095.7497 | 1140665.917 | 2442458.865 | 2040059.943 | 1589599.649 | 2482199.254 | 2320022.83 | 1438530.472 | 1516320.022 | 1466784.192 |
| TG 10:0_14:0_22:6 | 220215 | 146417 | 374056 | 215704 | 143916 | 957129. | 125357 | 107607 | 172738 | 168279 | 120552 | 795827. |

| **Metabolites** | **PBS_1** | **PBS_2** | **PBS_3** | **PBS_4** | **BCG_1** | **BCG_2** | **BCG_3** | **BCG_4** | **BCGΔ*pe18*_1** | **BCGΔ*pe18*_2** | **BCGΔ*pe18*_3** | **BCGΔ*pe18*_4** |
|---|---|---|---|---|---|---|---|---|---|---|---|---|
| | 4.766 | 3.786 | 8.671 | 8.793 | 2.05 | 2319 | 7.736 | 7.808 | 5.906 | 3.921 | 9.627 | 9555 |
| PE O-18:2_22:6 | 7668516.409 | 8519977.849 | 6449574.965 | 7499309.602 | 8847323.141 | 8944426.001 | 7298276.054 | 8938928.592 | 8707467.369 | 7016471.966 | 7701925.237 | 7278031.832 |
| LNAPS 17:2/N-26:7 | 1516725.244 | 2351781.972 | 275848.3421 | 858375.6587 | 499965.0698 | 435459.126 | 584679.1868 | 482085.351 | 420714.9394 | 535820.0074 | 340241.5103 | 295994.4702 |
| SM(d18:1/20:0) | 3215105.316 | 3259605.481 | 4817724.252 | 3087936.177 | 3358786.113 | 3023042.266 | 4611484.297 | 3547304.523 | 3571092.443 | 4550173.845 | 3954795.071 | 1727494.353 |
| Vincristine | 11377.51903 | 20588.16697 | 42292.22659 | 34746.39042 | 119615.9618 | 91397.78521 | 154787.8783 | 102881.8305 | 79389.595 | 103934.2933 | 128789.4902 | 89529.12017 |
| 1,3,5-Triazine-2-methanol, 4-amino-6-(phenylamino)-, 2-(4-nitrobenzoate) | 4605888.684 | 3895961.831 | 4045397.955 | 3975286.715 | 3876928.192 | 4160150.142 | 2955193.042 | 3528199.9 | 4058652.8 | 3343225.292 | 3358341.965 | 2947887.875 |
| Polyporenic acid C | 895674.6406 | 755663.4832 | 980267.3689 | 867727.0098 | 1044222.918 | 699393.3875 | 1452436.115 | 609746.9286 | 569174.3859 | 679064.371 | 949451.6297 | 453359.0884 |
| (+)-.gamma.-Tocopherol | 1923213.413 | 2136968.98 | 1811036.933 | 1743834.652 | 2142372.311 | 2097649.746 | 1956183.339 | 2051667.053 | 2019042.385 | 1808290.251 | 2094958.241 | 1646551.262 |
| PS 3:0_16:2 | 220330.1403 | 192993.3409 | 188133.4052 | 80234.88379 | 677110.6985 | 646395.6191 | 441313.9513 | 558544.8964 | 333192.3813 | 512844.4944 | 514820.5826 | 363690.7569 |
| Indolelactic acid | 823731.569 | 786740.6854 | 1541161.991 | 950238.0829 | 1218097.35 | 1212072.709 | 1430956.734 | 1158187.601 | 1254666.904 | 765212.6419 | 926777.2157 | 1135476.646 |
| SPB 15:0;2O | 25332602.78 | 25830670.08 | 23337158.41 | 23725652.46 | 24267801.17 | 23809051.25 | 25062606.38 | 24401088.44 | 24848847.31 | 24473528.66 | 24061987.8 | 22660420.64 |

| Metabolites | PBS_1 | PBS_2 | PBS_3 | PBS_4 | BCG_1 | BCG_2 | BCG_3 | BCG_4 | BCGΔ*pe18*_1 | BCGΔ*pe18*_2 | BCGΔ*pe18*_3 | BCGΔ*pe18*_4 |
|---|---|---|---|---|---|---|---|---|---|---|---|---|
| Hexadecasphinganine | 2951194.219 | 2712551.454 | 2565654.648 | 2672593.14 | 3038224.349 | 3415812.081 | 2902182.653 | 3692643.759 | 2649611.608 | 2743132.502 | 2824074.724 | 2219454.624 |
| Cucurbitacin B | 513038.1654 | 558353.1105 | 473235.1468 | 502460.8798 | 537771.0427 | 563040.6754 | 562597.7957 | 538396.4161 | 532692.8049 | 578037.1403 | 511166.7986 | 523764.8421 |
| .delta.-Tocotrienol | 119688.6353 | 165203.1273 | 136759.8497 | 244895.7546 | 179314.187 | 158538.2196 | 148249.7395 | 143440.9331 | 169665.2971 | 178481.5163 | 159429.4009 | 176717.236 |
| PG 18:0_20:4;O | 252918.2247 | 172455.7889 | 240479.2389 | 211948.6366 | 492401.2652 | 236356.5201 | 489974.2068 | 363644.2703 | 423832.1867 | 538693.3212 | 473502.2517 | 414549.4796 |
| Furazolidone | 4314445.949 | 3774853.574 | 5995213.736 | 4647796.136 | 5484952.288 | 5746427.244 | 5394773.102 | 5059586.263 | 5746754.441 | 3680886.703 | 4145547.794 | 5253062.469 |
| 2-Carboxy-4-dodecanolide | 374839.5574 | 283057.8959 | 358954.9747 | 418157.5915 | 403559.8957 | 393203.5608 | 424537.262 | 435231.533 | 372591.5513 | 470754.5303 | 479545.3611 | 592579.8747 |
| LPS 22:5 | 5748408.461 | 7162475.205 | 7315648.65 | 5123391.869 | 10609864.99 | 9639398.867 | 5291260.543 | 6133901.838 | 7863000.739 | 6730265.438 | 10759397.94 | 7406024.649 |
| FA 24:4 | 2609063.813 | 2751032.146 | 4812019.522 | 2702335.427 | 999567.5808 | 2209673.141 | 1631777.722 | 1733225.638 | 1324225.646 | 1360256.213 | 1953640.372 | 1096102.597 |
| PC O-30:2 | 9259494.647 | 12551295.89 | 9910773.421 | 12138816.66 | 11473037.5 | 10510641.74 | 9950770.676 | 10160324.54 | 9678896.043 | 9237163.24 | 8930414.725 | 10914608.38 |
| PI 16:4_20:5 | 533956.0859 | 329101.5245 | 642468.4505 | 352494.1942 | 214418.4343 | 0 | 164012.5784 | 241394.4183 | 371768.0548 | 253866.9228 | 156572.4895 | 124683.8629 |
| PG 16:0_16:0 | 2758421.84 | 3223434.005 | 2163685.207 | 5318845.211 | 4560213.677 | 22328.13564 | 1998144.335 | 1089942.61 | 1965121.544 | 2852345.296 | 1770240.897 | 651821.5864 |
| 2-(4-tert-Butylphenoxy)ethanamine | 589688.6937 | 681162.7877 | 554112.5977 | 658093.3306 | 573454.656 | 558337.3491 | 668959.7947 | 531601.1424 | 572133.171 | 627518.5197 | 560627.2165 | 790361.9712 |

| Metabolites | PBS_1 | PBS_2 | PBS_3 | PBS_4 | BCG_1 | BCG_2 | BCG_3 | BCG_4 | BCGΔ*pe18*_1 | BCGΔ*pe18*_2 | BCGΔ*pe18*_3 | BCGΔ*pe18*_4 |
|---|---|---|---|---|---|---|---|---|---|---|---|---|
| N-Acetylalanine | 1313705.642 | 1034633.221 | 1811294.232 | 1349218.81 | 2448412.581 | 1982515.747 | 2585157.976 | 2099831.521 | 1742750.484 | 1760564.546 | 2128147.732 | 1647878.243 |
| (E)-3-(4-Acetyloxy-2,3-dihydroxy-2,5,5,8a-tetramethyl-3,4,4a,6,7,8-hexahydro-1H-naphthalen-1-yl)prop-2-enoic acid | 1125413.309 | 859433.6653 | 959595.0918 | 1558473.429 | 696898.7897 | 897125.2565 | 737637.4038 | 729685.9716 | 747229.1371 | 850025.7606 | 934530.0574 | 749202.7901 |
| MITOMYCIN C | 998722.5006 | 795119.1692 | 1446951.204 | 792217.8388 | 1201731.424 | 1100027.639 | 1514277.916 | 1013692.355 | 969962.4447 | 1576091.232 | 1317193.54 | 798194.5027 |
| Oxopalmitoylcarnitine (Car(16:1-O)) | 151734.7184 | 24266.83368 | 213332.9226 | 121179.0709 | 674416.2602 | 450625.4382 | 965782.5026 | 579371.3737 | 387027.1459 | 766292.5474 | 358974.9825 | 196176.3411 |
| PI 12:0_25:0 | 562361.6619 | 479537.5875 | 279567.1196 | 581593.8756 | 126941.6262 | 150557.226 | 130958.8589 | 156859.006 | 202368.8953 | 197582.5774 | 126910.7173 | 142684.8382 |
| Deoxythymidine triphosphate | 921681.1857 | 402252.8409 | 998611.4298 | 2303882.209 | 1663282.377 | 1039342.766 | 778337.162 | 990617.004 | 787426.1328 | 678438.4516 | 895409.4666 | 1908912.279 |
| PI O-10:0_22:3 | 1446401.563 | 2589452.702 | 1719306.047 | 2585905.542 | 2205384.247 | 1491351.176 | 1670359.094 | 1750225.328 | 2575888.68 | 1981887.288 | 1553283.174 | 1410509.035 |
| Cholenate | 2057896.027 | 1471670.429 | 1362991.302 | 2583692.029 | 2356590.125 | 1889571.781 | 2580782.02 | 1992006.955 | 1056966.822 | 788091.9642 | 1378392.522 | 2180205.409 |
| 2.alpha.-Methyl-5.alpha.-androstan-3.alpha.-ol-17-one glucuronide | 600945.4826 | 386183.1434 | 630899.3904 | 591025.9119 | 352147.1811 | 630998.5557 | 365451.9299 | 690789.7093 | 182138.2636 | 405473.1097 | 555947.6048 | 461141.5508 |

| Metabolites | PBS_1 | PBS_2 | PBS_3 | PBS_4 | BCG_1 | BCG_2 | BCG_3 | BCG_4 | BCGΔ*pe18*_1 | BCGΔ*pe18*_2 | BCGΔ*pe18*_3 | BCGΔ*pe18*_4 |
|---|---|---|---|---|---|---|---|---|---|---|---|---|
| 2-Chloro-4-(dimethylamino)-5-fluoropyrimidine | 234948.5364 | 51733.76995 | 259113.0448 | 59256.27432 | 785284.7712 | 1851312.962 | 11318739.48 | 9339735.18 | 2808944.848 | 867134.4599 | 1721257.868 | 321490.7783 |
| PE O-18:4_3:0 | 52422.14687 | 35415.87693 | 65017.89411 | 51624.03489 | 56073.36959 | 30835.39362 | 81161.86151 | 48849.68314 | 29291.20838 | 81957.90368 | 21684.79818 | 14061.35355 |
| PG O-16:3_20:4 | 121684.9452 | 201056.5061 | 302218.0104 | 182366.9978 | 274917.9554 | 164061.6138 | 120315.3695 | 267165.5743 | 213510.1557 | 260581.6843 | 225068.0735 | 179830.6113 |
| 2-Formyl-1-hydroxy-5,5,8a-trimethyl-4-[(2E,4E,6E)-octa-2,4,6-trienoyl]oxy-4a,6,7,8-tetrahydro-4H-naphthalene-1-carboxylic acid | 145223.102 | 204283.4876 | 85237.52948 | 123440.4241 | 527253.5748 | 363359.8932 | 185569.6756 | 316609.293 | 140056.5588 | 133502.0457 | 119209.1285 | 127966.2429 |
| [6-Chloro-9-[(4-methoxy-3,5-dimethyl-2-pyridyl)methyl]purin-2-yl]amine | 141592.0191 | 7448.531465 | 86866.08291 | 100028.1311 | 58694.3209 | 143528.0195 | 2298633.345 | 279345.4601 | 398884.493 | 202682.1997 | 119556.8241 | 167304.3844 |
| M295T419 | 7604558.041 | 10849799.57 | 11515680.6 | 11739960.79 | 16034957.53 | 17115197.42 | 9909430.179 | 7059634.345 | 7319578.606 | 8438107.521 | 15686781.73 | 12117519.47 |
| 4-Nonylphenol | 2125300.913 | 2075506.075 | 2040318.037 | 2624590.627 | 1998474.273 | 2070058.078 | 2288106.043 | 2064576.996 | 2243234.562 | 2396307.914 | 2049867.783 | 2117410.82 |
| Proscillaridin A | 744468.3385 | 846738.6712 | 548890.9739 | 605610.4152 | 581223.9833 | 740362.0718 | 807168.8988 | 813832.7167 | 706671.4758 | 680981.4553 | 598549.3272 | 590920.0801 |

| Metabolites | PBS_1 | PBS_2 | PBS_3 | PBS_4 | BCG_1 | BCG_2 | BCG_3 | BCG_4 | BCGΔ *pe18*_1 | BCGΔ *pe18*_2 | BCGΔ *pe18*_3 | BCGΔ*pe18*_4 |
|---|---|---|---|---|---|---|---|---|---|---|---|---|
| (7E)-11,12-Dihydroxy-13-methoxy-4,5,8-trimethyl-3-(2-methylpropyl)-3,3a,4,6a,9,10,11,12,13,14-decahydro-1H-cycloundeca[d]isoindole-1,15(2H)-dione | 471621.3605 | 522177.3683 | 1001839.403 | 1457994.76 | 2306878.244 | 1876848.716 | 2999823.817 | 2044788.451 | 652884.8137 | 3682107.775 | 4176059.46 | 1627324.502 |
| PE O-9:0_11:0 | 1093729.31 | 1022070.505 | 764174.6672 | 808385.1419 | 716392.904 | 1185348.704 | 1189165.143 | 762789.5392 | 784249.3752 | 810724.2865 | 870073.1486 | 187263.1216 |
| Chlorophyll_b | 1980418.57 | 1779630.99 | 1029574.592 | 1697837.085 | 111148.6989 | 250333.129 | 346088.9749 | 404113.9542 | 370530.8654 | 572412.0803 | 457200.998 | 215047.4337 |
| Cholest-4-en-26-oic acid, 7alpha-hydroxy-3-oxo | 516764.1352 | 584985.3386 | 660233.7627 | 580030.5951 | 292859.8189 | 291118.4279 | 281197.7017 | 278931.3607 | 226846.6163 | 442753.1563 | 322545.693 | 357297.572 |
| 4-Hydroxyphenyl 3,6-bis-O-((2E)-3-(1-hydroxy-4-oxocyclohexa-2,5-dien-1-yl)prop-2-enoyl)-.beta.-D-glucopyranoside | 10460.51111 | 9069.533373 | 37072.01688 | 27087.93513 | 77081.74903 | 76189.44254 | 67689.36699 | 60848.66012 | 67613.76904 | 39196.03083 | 81656.51296 | 124138.4823 |
| N2-Acetyllysine | 950297. | 796683. | 221033 | 984700. | 370370 | 229455 | 335293 | 253153 | 255276 | 258055 | 263733 | 206664 |

| **Metabolites** | **PBS_1** | **PBS_2** | **PBS_3** | **PBS_4** | **BCG_1** | **BCG_2** | **BCG_3** | **BCG_4** | **BCGΔ*pe18*_1** | **BCGΔ*pe18*_2** | **BCGΔ*pe18*_3** | **BCGΔ*pe18*_4** |
|---|---|---|---|---|---|---|---|---|---|---|---|---|
| | 9687 | 8324 | 1.43 | 6127 | 7.987 | 9.512 | 0.182 | 9.095 | 5.149 | 9.246 | 6.962 | 0.595 |
| (3-Methylbutyl)(6-methylheptan-2-yl)amine | 415120.1793 | 654412.4204 | 663213.6718 | 323337.8113 | 529611.0157 | 470660.7351 | 547022.0768 | 583174.0612 | 549095.904 | 497475.7756 | 667259.3788 | 612347.3376 |
| 7.alpha.-Thiospironolactone | 96690.04447 | 58025.93422 | 151825.6185 | 89514.24048 | 2771672.365 | 1702015.692 | 3119579.647 | 2115681.279 | 1531551.862 | 2059691.539 | 2148855.523 | 1263208.123 |
| DLCL 22:3_20:4 | 335999.4433 | 284452.8475 | 421652.2077 | 305839.5891 | 232492.5752 | 220770.3381 | 374651.2109 | 180998.1557 | 245154.8976 | 396551.46 | 372166.7835 | 289962.4165 |
| PG 17:1_20:4 | 1642472.166 | 434747.9726 | 693696.3582 | 327985.2383 | 2490978.046 | 2322694.967 | 5398496.827 | 3117161.441 | 4325296.208 | 6733085.295 | 3945051.176 | 2254772.61 |
| 1-Isopropyl-1H-benzimidazole-2-thiol | 1258205.729 | 1307288.984 | 1393655.044 | 1258876.608 | 1113497.046 | 1142703.43 | 1210452.323 | 1053121.781 | 1200397.546 | 1090384.822 | 1167251.829 | 1170561.222 |
| Resazurin | 290748.8209 | 371442.4193 | 564350.6081 | 446342.8746 | 1884161.281 | 1344529.649 | 974233.0438 | 837160.9324 | 942940.7253 | 1540833.162 | 1498167.278 | 870819.759 |
| PC(P-16:0/0:0) | 4287647.935 | 4255295.33 | 2924177.241 | 3743981.226 | 4129163.935 | 3110674.601 | 3634628.04 | 3395655.44 | 3410773.075 | 3008162.598 | 3459561.936 | 2456872.815 |
| 1-Methyladenosine | 175991.9279 | 180561.3307 | 197858.411 | 201163.2437 | 2489396.6 | 520537.3961 | 1074740.227 | 1930453.265 | 1622157.884 | 674971.531 | 1933680.218 | 573267.9048 |
| Etofylline | 1045535.692 | 791990.3047 | 1251626.955 | 819253.9658 | 1705434.405 | 1075402.871 | 1431484.988 | 1465274.043 | 1335389.182 | 1293601.087 | 1391765.593 | 983988.8473 |
| SHexCer 35:2;3O | 1256294.286 | 641880.3399 | 411842.1677 | 608206.0117 | 4455817.27 | 2886863.693 | 3675342.477 | 3867850.472 | 5897300.285 | 4361121.827 | 2715155.545 | 3208331.64 |
| PC 16:0_20:5;O | 529134.3434 | 504905.5717 | 301882.1487 | 395594.4389 | 547148.6598 | 852776.4469 | 556032.7622 | 506898.4121 | 460564.5041 | 376230.4822 | 568060.2102 | 509739.1996 |

| Metabolites | PBS_1 | PBS_2 | PBS_3 | PBS_4 | BCG_1 | BCG_2 | BCG_3 | BCG_4 | BCGΔ*pe18*_1 | BCGΔ*pe18*_2 | BCGΔ*pe18*_3 | BCGΔ*pe18*_4 |
|---|---|---|---|---|---|---|---|---|---|---|---|---|
| Oxaceprol | 384338.1719 | 1861769.297 | 201938.1139 | 93062.50872 | 1961239.142 | 1180810.214 | 2142227.655 | 1777983.307 | 1069650.172 | 1430941.74 | 1546140.769 | 1027415.113 |
| Ceanothine_E | 28580071.76 | 25351930.59 | 25615440.09 | 25650319.54 | 25993203.7 | 22959714.14 | 28014389.98 | 25907940.46 | 25840081.79 | 27435890.23 | 27313799.85 | 26006471.47 |
| CAR 26:7 | 3871225.977 | 4634673.572 | 4030687.197 | 5333203.749 | 3268835.842 | 3321584.102 | 2896026.282 | 2192164.382 | 2061044.892 | 2920893.691 | 4054991.982 | 1860249.662 |
| 2-(1-Piperidyl)propan-2-ol | 613781.9078 | 1066706.075 | 1637439.038 | 1391719.049 | 522406.0604 | 330466.3921 | 662992.875 | 403461.8489 | 495057.2996 | 650661.0002 | 403952.9694 | 2667486.326 |
| Sorbitane Monostearate - Polysorbate 60 in-source fragment | 66630.57026 | 62634.95603 | 61254.56497 | 48682.35201 | 59292.80676 | 71520.97327 | 49494.36413 | 59427.98234 | 45694.68821 | 42134.53682 | 60219.77429 | 43951.09544 |
| lacty (2) diphospho-(5')adenosine | 66268.27704 | 88548.83732 | 103491.9711 | 81523.94606 | 327304.9119 | 243783.0234 | 287779.5902 | 263904.4094 | 274628.198 | 288285.9446 | 288162.4145 | 291813.3268 |
| Unoprostone | 6353986.06 | 5814330.719 | 6070350.425 | 5758459.7 | 6277627.431 | 5193863.317 | 4788302.855 | 5267984.554 | 5490957.413 | 5586762.774 | 6499795.865 | 5329661.259 |
| 2-Naphthalenethiol | 173886.1426 | 44099964.47 | 157373.4193 | 33718382.64 | 2445651.742 | 2415401.081 | 2025558.067 | 2223990.594 | 2955004.832 | 2655359.141 | 2507629.142 | 3332566.523 |
| PE O-22:6_18:2 | 324934.6268 | 446745.6479 | 657588.9103 | 542053.3624 | 1203759.456 | 536799.1678 | 1241127.2 | 754145.6373 | 1291180.677 | 1827184.856 | 1067336.338 | 913264.1484 |
| Dipentylamine | 526164.1771 | 1139475.231 | 1253933.924 | 835315.2576 | 602449.2214 | 607131.0686 | 890632.5654 | 683211.0969 | 484431.4809 | 565498.3927 | 540979.1235 | 2033697.241 |
| 1-Methylinosine | 554961. | 579710. | 131715 | 124920 | 790566 | 386561 | 638737 | 478988 | 545613 | 534945 | 688688 | 473048 |

| Metabolites | PBS_1 | PBS_2 | PBS_3 | PBS_4 | BCG_1 | BCG_2 | BCG_3 | BCG_4 | BCGΔ*pe18*_1 | BCGΔ*pe18*_2 | BCGΔ*pe18*_3 | BCGΔ*pe18*_4 |
|---|---|---|---|---|---|---|---|---|---|---|---|---|
| | 4927 | 3652 | 7.969 | 4.17 | 0.175 | 6.873 | 1.405 | 8.088 | 4.819 | 3.317 | 6.366 | 9.459 |
| Phe(Benzoyl)-Leu-Arg | 174219.4798 | 130515.1121 | 55645.88009 | 56740.27553 | 147147.6356 | 140699.7215 | 168001.4423 | 139698.0113 | 70112.84853 | 109990.6313 | 198094.1963 | 107437.6034 |
| Arecoline | 1868321.123 | 2147089.747 | 1723502.224 | 1954178.493 | 1940199.11 | 1947841.073 | 1834038.475 | 1829776.943 | 1814068.342 | 1823090.893 | 1904088.197 | 1809413.018 |
| PC O-20:4_28:7 | 168742.1765 | 118880.514 | 132951.1573 | 137041.7206 | 206448.273 | 229563.7131 | 116426.219 | 106883.67 | 157520.2649 | 155209.6353 | 134649.608 | 189274.8911 |
| Benzeneethanamine, N-[(2-fluorophenyl)methyl]-4-iodo-2,5-dimethoxy- | 829812.1559 | 543559.7877 | 292768.1044 | 686824.9387 | 1033220.564 | 347067.3671 | 977504.0981 | 769233.2509 | 1015050.056 | 839988.0798 | 394857.4882 | 521157.8991 |
| LPI 22:4 | 12600797.29 | 12777086.11 | 13233164.29 | 10791055.47 | 6059907.44 | 4582689.29 | 6906534.654 | 4986674.601 | 5964131.313 | 8141082.731 | 9107370.744 | 4567302.67 |
| Tetraoctylammonium cation | 14400.36716 | 24021.2353 | 16809.5914 | 15174.96085 | 50583.39415 | 0 | 452924.6976 | 1126173.813 | 71638.24775 | 31123.04942 | 33078.73051 | 1023459.551 |
| LPC 20:4 | 16256258.96 | 15428118.94 | 23519053.96 | 16768615.97 | 10020490.05 | 9477793.209 | 12854720.86 | 12451598.02 | 9204265.18 | 9742486.307 | 11163615.27 | 10590941.81 |
| Tetradecenoylcarnitine (Car(14:1)) | 1608614.353 | 1106111.085 | 2647145.304 | 2166422.832 | 5487841.838 | 3550474.662 | 6040036.614 | 4785732.902 | 2354411.349 | 3693174.136 | 4029378.882 | 1969342.996 |
| ST 24:1;O4;G/20:3 | 503599.0456 | 739590.1737 | 252091.3585 | 411052.2174 | 300242.8418 | 533516.0962 | 1054619.054 | 290885.8852 | 318299.4927 | 399626.7303 | 365098.9971 | 638965.1648 |
| gamma-glutamylanilide diol | 0 | 14682.32828 | 0 | 0 | 2564558.782 | 1794513.867 | 4023561.773 | 1994818.845 | 1494522.628 | 996979.3161 | 1406237.747 | 1587322.951 |

| Metabolites | PBS_1 | PBS_2 | PBS_3 | PBS_4 | BCG_1 | BCG_2 | BCG_3 | BCG_4 | BCGΔ*pe18*_1 | BCGΔ*pe18*_2 | BCGΔ*pe18*_3 | BCGΔ*pe18*_4 |
|---|---|---|---|---|---|---|---|---|---|---|---|---|
| Propyphenazone | 1046116.432 | 1553770.741 | 1251031.132 | 1485504.144 | 1644729.656 | 1481654.082 | 1706740.556 | 1297251.391 | 1624570.516 | 1518063.14 | 1453760.805 | 1228162.307 |
| 14h-Anthra[2,1,9-mna]thioxanthen-14-one | 2457174.939 | 2545312.145 | 2978759.026 | 2844925.55 | 2965502.636 | 2765383.16 | 3499026.457 | 3001520.507 | 2844091.962 | 2875763.408 | 2983571.02 | 2621572 |
| 5-Amino-2-(3,4'-dihydroxy-4,4,7,8a-tetramethyl-6'-oxospiro[2,3,4a,5,6,7-hexahydro-1H-naphthalene-8,2'-3,8-dihydrofuro[2,3-e]isoindole]-7'-yl)-5-oxopentanoic acid | 4368529.942 | 5583797.296 | 6550298.626 | 4396449.998 | 985826.8196 | 2783512.239 | 1950492.49 | 1147692.043 | 1264545.77 | 1799929.428 | 2501140.123 | 2034763.836 |
| 4(5H)-Thiazolone, 5-(6-quinolinylmethylene)-2-[(2-thienylmethyl)amino]-, (5Z)- | 868546.2262 | 249073.8812 | 735665.1625 | 417916.9978 | 16983.39561 | 255319.1275 | 2423547.893 | 2510243.678 | 354427.7022 | 84286.51762 | 491181.2851 | 18454.40575 |
| 2-(1-Adamantyl)-N-hydroxyacetamide | 780568.8351 | 860721.8003 | 568962.2966 | 942554.7881 | 593305.8072 | 631356.7895 | 862045.2724 | 545866.2885 | 761944.5164 | 689791.0697 | 610019.189 | 1059790.77 |
| 1-hexadecyl-glycero-3-phosphate | 1940844.177 | 1755064.898 | 2297186.441 | 1809179.565 | 2660680.397 | 1258150.18 | 1950539.8 | 1722997.439 | 1530529.333 | 1462683.548 | 1738154.257 | 1107766.828 |
| Guanosine | 571556. | 170113. | 176152. | 163472. | 744621. | 555061. | 707801. | 597151. | 384432. | 343641. | 416011. | 394140. |

| Metabolites | PBS_1 | PBS_2 | PBS_3 | PBS_4 | BCG_1 | BCG_2 | BCG_3 | BCG_4 | BCGΔ*pe18*_1 | BCGΔ*pe18*_2 | BCGΔ*pe18*_3 | BCGΔ*pe18*_4 |
|---|---|---|---|---|---|---|---|---|---|---|---|---|
| | 0421 | 7438 | 5666 | 3111 | 2355 | 8886 | 675 | 7138 | 9251 | 4384 | 1879 | 0019 |
| SHexCer 30:3;3O | 154867.1574 | 184994.2326 | 59935.30915 | 128995.6262 | 124690.6932 | 79802.6441 | 113722.1236 | 106252.9067 | 123125.233 | 135561.791 | 121224.0638 | 98207.42616 |
| (+)-Fluprostenol | 1561135.868 | 1532344.796 | 1419551.974 | 1401144.202 | 1208335.807 | 1246801.67 | 1486144.635 | 1237313.74 | 553056.8003 | 1398884.561 | 1306321.678 | 1240876.643 |
| 3,5-Di-tert-butyl-2-hydroxybenzaldehyde | 2161739.478 | 2679542.583 | 2244224.592 | 2027699.203 | 2397584.19 | 2545072.16 | 3436648.14 | 2539150.594 | 2270947.5 | 2283045.245 | 2656564.061 | 2416799.577 |
| cis-Aconitic acid | 1374965.607 | 1128257.439 | 1496809.641 | 1113170.91 | 6644382.804 | 3445725.342 | 7937975.216 | 4913408.893 | 3100592.921 | 5037344.921 | 4515483.281 | 2797435.092 |
| Acetylsalicylic acid | 575123.1062 | 617788.9283 | 630946.3146 | 640522.0352 | 558203.9991 | 571253.6031 | 625799.642 | 617523.0956 | 636298.004 | 673201.2034 | 622018.7633 | 750804.3437 |
| 1,2-Di-(9Z,12Z,15Z-octadecatrienoyl)-sn-glycero-3-phosphocholine | 5059973.238 | 5323840.193 | 2567451.751 | 3624873.145 | 14037123.55 | 11828676.05 | 12606556.11 | 14226525.01 | 16480789.27 | 16414155.01 | 13393620.72 | 13928721.07 |
| Phenformin | 180269.8007 | 531507.6066 | 448585.4284 | 589097.1874 | 446072.2007 | 369744.4056 | 622893.3325 | 423086.075 | 508097.9537 | 448446.2664 | 562097.9669 | 649717.5717 |
| 4-Aminopiperidine-4-carboxylic acid | 1431026.714 | 1194409.572 | 1219400.456 | 1744014.689 | 1091355.948 | 971796.2415 | 1006735.969 | 209632.3389 | 1086164.464 | 1406102.031 | 3025788.128 | 1486908.25 |
| LPG O-17:1 | 574408.3584 | 309981.8281 | 505769.517 | 349198.895 | 878450.0124 | 548117.389 | 508579.9451 | 532886.1114 | 521527.3172 | 518421.9515 | 532940.9644 | 442402.3208 |
| LPC O-13:1 | 1551762.346 | 1684174.99 | 414785.7274 | 1210357.682 | 599245.1762 | 435559.1935 | 509807.2627 | 429615.4847 | 575973.4123 | 542578.1634 | 363574.5454 | 315718.1878 |

| Metabolites | PBS_1 | PBS_2 | PBS_3 | PBS_4 | BCG_1 | BCG_2 | BCG_3 | BCG_4 | BCGΔ*pe18*_1 | BCGΔ*pe18*_2 | BCGΔ*pe18*_3 | BCGΔ*pe18*_4 |
|---|---|---|---|---|---|---|---|---|---|---|---|---|
| Gabapentin related compound E | 779228.0595 | 454515.2507 | 584318.6524 | 452975.8895 | 332525.6765 | 331805.5277 | 361572.6831 | 311608.1927 | 510614.5912 | 339320.4257 | 316300.603 | 780975.8587 |
| SM 36:2;3O | 878680.398 | 959586.2952 | 894576.2431 | 909842.214 | 712395.3276 | 652809.7852 | 565991.043 | 735053.5071 | 772643.6236 | 705058.8231 | 735279.5044 | 1045501.418 |
| N-[2-(4-Prenyloxyphenyl)ethyl]tiglamide | 233011.6363 | 249751.737 | 114392.456 | 145625.8895 | 277689.607 | 234417.5349 | 250142.5794 | 216127.6192 | 203408.9989 | 238446.578 | 258590.9184 | 130128.7887 |
| PC O-33:4 | 1431533.134 | 742604.7134 | 2131317.859 | 1177107.641 | 1233036.663 | 3503353.059 | 1204395.477 | 820653.7066 | 1696090.648 | 1798600.139 | 1088399.617 | 916483.3576 |
| 8-[(2-Hydroxyethyl)amino]-1,3,7-trimethyl-3,7-dihydro-1H-purine-2,6-dione | 1225252.362 | 5816723.442 | 582871.1598 | 3930994.939 | 4629980.311 | 4448900.515 | 31382.69685 | 101845.0467 | 1047475.375 | 2345897.263 | 3083752.716 | 722738.0765 |
| Alanylalanine (Ala-Ala) | 54131.6945 | 40789.89274 | 108334.6284 | 53074.08226 | 6038434.144 | 5191264.241 | 4314899.127 | 4324393.637 | 4371177.357 | 3140276.197 | 3741017.334 | 4156506.003 |
| .beta.-D-Glucopyranosyl 6-O-(8-methylnonanoyl)-.beta.-D-glucopyranoside | 75748.06733 | 30719.56164 | 130572.4121 | 317770.9767 | 409108.5042 | 491306.685 | 422925.3838 | 517448.1073 | 331824.8073 | 320223.1586 | 431153.4987 | 354140.5698 |
| Gardenin B | 1381391.451 | 754680.1572 | 1524523.864 | 1101977.483 | 1818448.224 | 1163827.735 | 3590260.537 | 2357419.437 | 1385348.001 | 1846078.16 | 1393883.211 | 520239.9393 |
| 1-(4-Pyridinyl)piperazine | 4397357.369 | 5048430.315 | 4514698.981 | 4022375.788 | 4349573.217 | 4444174.355 | 4283570.877 | 6457761.253 | 4368706.819 | 4236366.937 | 4140450.043 | 4014921.157 |

| **Metabolites** | **PBS_1** | **PBS_2** | **PBS_3** | **PBS_4** | **BCG_1** | **BCG_2** | **BCG_3** | **BCG_4** | **BCGΔ*pe18*_1** | **BCGΔ*pe18*_2** | **BCGΔ*pe18*_3** | **BCGΔ*pe18*_4** |
|---|---|---|---|---|---|---|---|---|---|---|---|---|
| 3-(2-Pyrimidinyl)propanoic acid | 2644643.902 | 4274796.36 | 4173118.114 | 4148399.305 | 4081378.557 | 4140412.109 | 2351448.927 | 4177532.88 | 4125076.691 | 2590648.866 | 4131860.339 | 2470658.154 |
| 2-Methyl-1-phenyl-2-propanyl butyrate | 2389460.953 | 3252231.042 | 2439674.251 | 2752321.547 | 2226899.399 | 2999533.333 | 2284450.016 | 2506687.201 | 2348570.876 | 2172450.938 | 2056206.103 | 2823461.516 |
| LPG 20:5 | 1359946.972 | 1221907.715 | 1554682.158 | 2086474.016 | 1024517.105 | 1523624.541 | 1314497.954 | 1590041.604 | 1558905.487 | 1305246.181 | 1170065.664 | 2112221.177 |
| 5,5-Dimethylimidazolidine-2,4-dione | 6395486.011 | 6364979.106 | 6615609.191 | 17569745.07 | 6428257.572 | 4994356.198 | 7150875.729 | 9570442.09 | 5981524.362 | 3724100.5 | 5949107.913 | 3014548.457 |
| PC O-13:1_22:6 | 171715.213 | 165995.986 | 171724.141 | 172349.4993 | 550267.3402 | 769003.0396 | 664067.0732 | 396030.1914 | 562496.0759 | 579193.2965 | 499611.6624 | 389230.6574 |
| Duroquinone | 883874.406 | 913783.8215 | 794527.2205 | 1225005.33 | 631017.452 | 750368.5024 | 770334.3566 | 798900.5777 | 773448.771 | 1003335.575 | 806755.7465 | 1172702.252 |
| 4-O-acetyl-N-acetyl mannosamine | 8386232.491 | 2921996.051 | 15490425.36 | 3212933.216 | 6679749.456 | 4093042.763 | 8559781.139 | 5612870.447 | 3009468.88 | 1617605.437 | 1782986.246 | 2826223.569 |
| Raffinose | 14419.36082 | 0 | 21893.57483 | 0 | 1951150.409 | 1643043.096 | 1684159.898 | 1237510.047 | 699905.6589 | 401601.026 | 504119.8559 | 695528.4678 |
| 5-Hydroxyvalproic acid | 1701432.309 | 1450287.399 | 1584972.414 | 1334001.329 | 1856708.516 | 1333907.551 | 2159146.077 | 2030308.725 | 1375930.061 | 1584777.323 | 1665216.467 | 1882777.256 |
| CAR 18:4 | 1312437.352 | 1195108.254 | 1439743.258 | 1138449.642 | 1673184.38 | 1296317.492 | 1457669.936 | 1402278.877 | 1331744.084 | 1349277.271 | 1345221.035 | 1288004.596 |
| 2-(3-Phenyl-1H-1,2,4-triazol-5-yl)pyridine | 652845.4764 | 492010.9446 | 641848.6976 | 684750.9917 | 732503.4875 | 1084131.717 | 731745.8565 | 681635.9395 | 706470.5617 | 806802.585 | 707572.1275 | 507400.2839 |
| MGDG O-22:6_7:0 | 52134.3 | 49139.5 | 85724.2 | 43584.4 | 39173.4 | 40567.7 | 57746.7 | 37133.4 | 0 | 13745.1 | 25525.5 | 0 |

| Metabolites | PBS_1 | PBS_2 | PBS_3 | PBS_4 | BCG_1 | BCG_2 | BCG_3 | BCG_4 | BCGΔ*pe18*_1 | BCGΔ*pe18*_2 | BCGΔ*pe18*_3 | BCGΔ*pe18*_4 |
|---|---|---|---|---|---|---|---|---|---|---|---|---|
| | 5891 | 3025 | 9699 | 0649 | 0383 | 8235 | 3532 | 0399 | | 9671 | 0403 | |
| ST 24:1;O4;T/18:2 | 5126491.25 | 3294356.584 | 5837527.858 | 5646242.693 | 7035650.092 | 4212267.767 | 7198903.003 | 4889160.873 | 7175831.901 | 7473820.777 | 8198880.964 | 2333944.095 |
| NAOrn 22:6/22:5 | 2062965.787 | 1924796.451 | 647255.6072 | 485087.965 | 1210398.669 | 1224680.631 | 1345773.643 | 1104212.009 | 1292924.413 | 1326039.525 | 1063382.978 | 1021519.568 |
| 2,4-Di-tert-amylphenol | 240959.4629 | 2284371.617 | 2398151.75 | 268947.5799 | 1307025.742 | 981374.2218 | 1365450.307 | 1858859.596 | 959407.3987 | 649430.0362 | 605705.0744 | 1188595.997 |
| PE O-16:1_20:2 | 785949.997 | 200840.0516 | 826058.7872 | 512758.4835 | 417227.1619 | 0 | 623864.8136 | 347731.5428 | 494176.4847 | 759670.8786 | 398556.8534 | 529747.2506 |
| 2-(4-Phenyl-1,3-thiazol-2-yl)ethylamine | 509303.7624 | 493783.813 | 516709.6601 | 506647.3166 | 433133.4048 | 219164.359 | 464736.2948 | 265230.4823 | 279666.4654 | 269223.6799 | 332068.8399 | 236737.6794 |
| Lawsone | 3099426.283 | 2218688.586 | 3345447.052 | 1777197.544 | 4161562.004 | 3046154.572 | 3253365.124 | 2562548.289 | 2976134.484 | 2250504.53 | 2203917.873 | 729656.3837 |
| 7-Chloro-4-oxo-4H-chromene-2-carboxylic acid | 1812272.492 | 566866.6641 | 1920725.407 | 582115.8535 | 2753031.762 | 1514075.819 | 1445500.691 | 2620015.439 | 1047406.995 | 1037596.245 | 1598715.713 | 1049787.992 |
| LPG(18:1(9Z)) | 3354975.279 | 5186397.267 | 4400800.722 | 3641762.871 | 23586145.25 | 18574222.62 | 22640389.07 | 18145660.42 | 17659679.55 | 19127584.31 | 24238709.51 | 17010985.73 |
| Piperidine | 3989902.442 | 3289131.591 | 4192455.113 | 3427493.411 | 6832603.526 | 4762479.549 | 6228618.993 | 5364117.839 | 4682597.82 | 5905125.068 | 5055111.242 | 4404334.746 |
| Cytidine | 445309.5894 | 468860.7278 | 728116.5403 | 506122.4069 | 707019.6721 | 501113.3285 | 793825.4468 | 558794.5909 | 557428.839 | 490340.4228 | 483312.6547 | 523262.1717 |

| Metabolites | PBS_1 | PBS_2 | PBS_3 | PBS_4 | BCG_1 | BCG_2 | BCG_3 | BCG_4 | BCGΔ *pe18*_1 | BCGΔ *pe18*_2 | BCGΔ *pe18*_3 | BCGΔ*pe18*_4 |
|---|---|---|---|---|---|---|---|---|---|---|---|---|
| 5-Heptenoic acid, 7-[(1R,2R,3S,5S)-2-[(1E,3R)-3-(2,3-dihydro-1H-inden-2-yl)-3-hydroxy-1-propen-1-yl]-3-fluoro-5-hydroxycyclopentyl]-, (5Z)- | 831801.936 | 680463.5722 | 557162.7368 | 817625.0281 | 1135485.416 | 1090137.592 | 932409.9979 | 649057.1486 | 949478.6541 | 925712.8756 | 723162.069 | 668814.1225 |
| SQDG 18:1_22:4 | 1756351.5 | 1958954.465 | 1858039.58 | 1785059.845 | 1920204.865 | 1954209.228 | 1892211.743 | 1777051.306 | 1907651.386 | 1761850.338 | 1819529.277 | 1894636.882 |
| 3-Hydroxydecanoic acid | 346721.8788 | 341913.6909 | 465114.4237 | 397484.9098 | 409300.9292 | 438911.0661 | 453713.7941 | 400329.6392 | 365230.6629 | 448883.153 | 424634.6845 | 408657.0348 |
| Clascoterone | 159370.0178 | 184368.2856 | 141737.1959 | 188197.7062 | 228794.4186 | 251431.8207 | 235663.0864 | 232609.1348 | 235362.3514 | 212393.7167 | 204880.8858 | 246690.9949 |
| 4-Amino-2-methylenebutanoic acid | 1427270.096 | 1273844.852 | 1470158.279 | 1353919.835 | 3221709.797 | 1756353.454 | 1714875.236 | 1628494.699 | 1786917.953 | 1697375.731 | 1948964.181 | 1044530.314 |
| Lepidine_D | 1469176.447 | 982715.2041 | 910987.9541 | 1034718.917 | 1054473.134 | 810129.1139 | 1150710.051 | 1366081.316 | 929525.0854 | 460442.695 | 624625.2403 | 381940.1573 |
| Diclofensine | 806017.8084 | 542091.3675 | 1167891.036 | 537073.4427 | 853704.476 | 825009.4072 | 783028.4845 | 579847.5041 | 595531.5515 | 549290.1972 | 492182.8421 | 336452.6655 |
| PI 19:0_20:4 | 871651.3314 | 538933.3478 | 870967.0636 | 862628.0828 | 446240.6468 | 0 | 342988.657 | 7639.773808 | 745279.5642 | 641349.1898 | 479408.2078 | 6961.954936 |
| 3-Deacetylsalannin | 698318.939 | 476316.1409 | 638974.0514 | 1332694.167 | 696966.8415 | 1451363.919 | 911376.1039 | 463859.0104 | 497320.7167 | 1438147.467 | 547772.0551 | 502660.1812 |

| Metabolites | PBS_1 | PBS_2 | PBS_3 | PBS_4 | BCG_1 | BCG_2 | BCG_3 | BCG_4 | BCGΔ*pe18*_1 | BCGΔ*pe18*_2 | BCGΔ*pe18*_3 | BCGΔ*pe18*_4 |
|---|---|---|---|---|---|---|---|---|---|---|---|---|
| SM 22:1;2O/20:3 | 1969972.889 | 2921226.956 | 1880259.573 | 6538179.958 | 4098609.499 | 30054.67499 | 3355261.61 | 3159144.718 | 3272506.695 | 1744247.681 | 2748400.878 | 1397885.425 |
| Propionylcarnitine (Car(3:0)) | 19499777.37 | 9820309.397 | 27479981.31 | 15920555.83 | 11794328.62 | 18195378.9 | 13436455.75 | 14658646.15 | 11916764.03 | 8679451.302 | 18559268.76 | 20503414.46 |
| Benzoic acid, 4-[4-[[5-(4,5-dimethyl-2-nitrophenyl)-2-furanyl]methylene]-4,5-dihydro-3-methyl-5-oxo-1H-pyrazol-1-yl]- | 430397.6139 | 509696.3235 | 1123179.987 | 400055.1998 | 772079.8051 | 772088.6417 | 868201.9961 | 719860.6394 | 1222953.154 | 539819.4059 | 594692.8708 | 633372.4932 |
| PC 8:0_28:5 | 1335109.392 | 1107356.271 | 1675747.08 | 1116178.284 | 1071804.826 | 0 | 792850.9267 | 650284.3592 | 647354.4889 | 931053.9607 | 1200732.877 | 321326.4022 |
| 4-[4-(2,4-Dicarboxybenzoyl)phenoxy]phthalic acid | 40933.82307 | 0 | 0 | 26225.22356 | 61594.86632 | 59693.95695 | 117221.9112 | 61520.32417 | 45614.66078 | 0 | 0 | 41697.09823 |
| PS 40:6 | 1252463.364 | 907459.737 | 1557641.438 | 1156643.62 | 1200464.92 | 382442.9914 | 1183319.885 | 1165377.276 | 1418585.706 | 1093959.076 | 1143401.305 | 932882.8292 |
| 1H-Indole-3-carboxamide, N-[(1S)-1-(aminocarbonyl)-2,2-dimethylpropyl]-1-(5-fluoropentyl)- | 1310892.879 | 1431902.375 | 1440638.994 | 1677415.614 | 1401676.971 | 1485248.186 | 1522270.729 | 1436865.886 | 1374303.684 | 1658034.386 | 1381638.63 | 1316571.038 |

| Metabolites | PBS_1 | PBS_2 | PBS_3 | PBS_4 | BCG_1 | BCG_2 | BCG_3 | BCG_4 | BCGΔ*pe18*_1 | BCGΔ*pe18*_2 | BCGΔ*pe18*_3 | BCGΔ*pe18*_4 |
|---|---|---|---|---|---|---|---|---|---|---|---|---|
| Lauramine oxide | 278554365.4 | 295382481.8 | 272552820.4 | 272289720.4 | 274480850.1 | 276122115.1 | 273186597.1 | 305668937.9 | 263084499.7 | 265842766.5 | 249593263.9 | 273716718.4 |
| 5-Amino-2,4-dichlorophenol | 6226356.879 | 3542797.955 | 3976268.576 | 5585673.37 | 5942687.867 | 4971239.929 | 2553398.629 | 4052777.786 | 3693931.388 | 5175369.322 | 4222175.887 | 4214995.457 |
| 1-[3-(Acetylsulfanyl)propanoyl]-6-methyl-2-piperidinecarboxylic acid | 787199.3543 | 743003.2459 | 833727.0254 | 859435.3601 | 1732748.508 | 1254253.111 | 1754863.393 | 1461324.051 | 1721482.898 | 1225372.791 | 954674.7824 | 1329666.335 |
| PC(38:7) | 2620787.409 | 3073801.784 | 2139765.295 | 2753357.633 | 3033798.976 | 2910000.364 | 3541743.506 | 2728490.358 | 3524893.256 | 2883418.703 | 1150726.574 | 2117489.381 |
| 4-Methylpentanoylcarnitine | 2011067.494 | 1994685.531 | 2179427.964 | 2365197.622 | 1626023.622 | 1595958.912 | 2126204.839 | 1695830.042 | 1789072.087 | 1889184.405 | 1769073.9 | 2944671.209 |
| 3-(2-Hydroxy-4-oxo-4H-pyran-3-yl)propanoic acid | 471335.3557 | 424850.0573 | 570386.7158 | 451400.3809 | 2387154.457 | 3521026.975 | 2652062.279 | 1944023.927 | 2167734.725 | 5453752.56 | 1962913.862 | 3844766.225 |
| 4-deoxy-beta-erythro-hex-4-enopyranuronosyl-(1,4)-N-sulfoglucosamine | 172725.9027 | 76229.0328 | 363545.0076 | 169825.2455 | 818195.6232 | 559254.0537 | 868758.4993 | 680691.8035 | 460600.7234 | 461162.2347 | 746538.9441 | 449300.1607 |
| Ethcathinone | 1762391.375 | 1562631.224 | 1759682.547 | 1746920.085 | 971520.4071 | 1137130.171 | 1094438.289 | 1172556.195 | 1155240.891 | 1104709.921 | 1184362.314 | 1423763.372 |
| Lucidenic Acid E2 | 97916.27521 | 137480.7032 | 145946.4417 | 121959.5428 | 222011.6272 | 117307.9842 | 167152.8312 | 90215.19809 | 129818.3436 | 101659.6009 | 192297.3537 | 165489.3982 |
| PC O-33:2 | 220639 | 205114 | 228114 | 343054 | 126685 | 544961 | 220296 | 453255 | 277980 | 188504 | 589468. | 419926 |

| Metabolites | PBS_1 | PBS_2 | PBS_3 | PBS_4 | BCG_1 | BCG_2 | BCG_3 | BCG_4 | BCGΔ*pe18*_1 | BCGΔ*pe18*_2 | BCGΔ*pe18*_3 | BCGΔ*pe18*_4 |
|---|---|---|---|---|---|---|---|---|---|---|---|---|
| | 7.596 | 2.857 | 6.06 | 5.257 | 5.661 | 9.858 | 3.175 | 9.376 | 8.825 | 6.267 | 0971 | 7.231 |
| LNAPE 18:0/N-22:5 | 11275621.74 | 6175326.296 | 9165156.37 | 8195414.756 | 7669050.499 | 53615.94558 | 6570426.942 | 3196831.337 | 9066790.656 | 7286146.576 | 7745159.793 | 7647794.865 |
| 2-(2H-Benzotriazol-2-yl)-4,6-bis(1-methyl-1-phenylethyl)phenol | 256485.0265 | 260960.206 | 207030.6707 | 234339.1139 | 277332.4462 | 289958.4078 | 282219.0494 | 327309.1551 | 311840.5089 | 314052.5525 | 275249.945 | 226416.1285 |
| PS 29:5 | 514632.6057 | 353745.4194 | 781625.5012 | 317089.416 | 237114.9904 | 175497.4388 | 379203.6514 | 287853.8722 | 162379.8108 | 175284.131 | 215722.4885 | 465972.655 |
| LDGTS 11:0 | 62739.24038 | 46402.40282 | 62838.18237 | 55018.35874 | 825391.6233 | 453603.2092 | 655442.026 | 528001.5999 | 295178.8709 | 554309.8838 | 707112.0103 | 256172.6156 |
| Droloxifene | 208843.6948 | 47730.513 | 251502.4547 | 52953.51194 | 437364.7866 | 612901.4554 | 1691480.555 | 1426515.468 | 570859.9333 | 408151.6429 | 852481.4158 | 385243.9248 |
| 6-Hydroxy-6-(3-hydroxy-2,3-dimethylcyclopentyl)-2-methylheptanoic acid | 557592.2962 | 597725.4856 | 702132.388 | 656304.2437 | 689842.9435 | 801245.7543 | 838940.3343 | 632706.4001 | 597474.0493 | 600310.6816 | 622659.7171 | 769405.0728 |
| 3-Oxostearic acid | 1601109.429 | 1843026.287 | 1801401.01 | 1506369.27 | 1902713.139 | 1737997.678 | 1529656.147 | 1519786.714 | 1378810.48 | 1293786.11 | 1348618.176 | 4234095.933 |
| PE O-26:4_10:0 | 4383888.124 | 207090.2067 | 4285731.59 | 1588439.906 | 1718308.77 | 228792.9165 | 1779429.235 | 1329272.795 | 2788658.305 | 2008383.832 | 1144264.166 | 1639068.522 |
| FAHFA 18:1/20:3 | 1364413.531 | 1867435.867 | 1594377.604 | 893597.4982 | 505381.0674 | 4155627.843 | 614503.8667 | 800081.4139 | 264237.2285 | 158599.2533 | 687847.4979 | 633880.0585 |

| Metabolites | PBS_1 | PBS_2 | PBS_3 | PBS_4 | BCG_1 | BCG_2 | BCG_3 | BCG_4 | BCGΔ*pe18*_1 | BCGΔ*pe18*_2 | BCGΔ*pe18*_3 | BCGΔ*pe18*_4 |
|---|---|---|---|---|---|---|---|---|---|---|---|---|
| 2-Bromo-N-(2-methylphenyl)benzamide | 463534.9399 | 320153.8607 | 650118.5209 | 416312.6139 | 4081462.964 | 2055869.196 | 3932914.01 | 2885819.999 | 2090622.694 | 3217423.239 | 2751084.056 | 1813539.801 |
| SL 10:0;O/10:0 | 235059.5787 | 266473.8812 | 232438.485 | 237465.2073 | 2626132.689 | 1910820.258 | 3719542.202 | 2082961.042 | 2258415.136 | 3561071.873 | 3964677.361 | 1665975.605 |
| 2-Bromoacetophenone | 1366304.797 | 911470.7999 | 1772102.961 | 1174449.155 | 2840720.434 | 2038074.532 | 2440136.143 | 2099367.02 | 2269753.614 | 1560044.908 | 2137640.827 | 1873197.495 |
| ADP-ribose | 228203.7181 | 213945.3507 | 327891.842 | 262951.0703 | 1795947.037 | 1752311.787 | 1307419.748 | 1427713.645 | 1688620.875 | 1469770.009 | 2083958.469 | 2529309.157 |
| 4-[2-(2,6-dimethoxy-4-prop-2-enylphenoxy)-1-hydroxypropyl]-2-methoxyphenol | 829399.6561 | 888087.7614 | 742982.7337 | 769543.9798 | 805949.3701 | 813230.9884 | 818120.3165 | 778296.2772 | 870758.8842 | 793532.7715 | 732828.0292 | 664940.0947 |
| 5-Chloro-3-fluoro-2-(pyrrolidin-1-yl)pyridine | 2933179.125 | 2794887.284 | 2764306.809 | 590504.5059 | 3228605.395 | 2541992.869 | 2695177.268 | 684011.8453 | 2718012.911 | 3410662.008 | 749145.4321 | 3330535.314 |
| N6-(delta2-Isopentenyl)-adenosine 5'-monophosphate | 0 | 0 | 0 | 0 | 71665.92205 | 57256.91803 | 52537.93853 | 52024.23415 | 74617.02436 | 87070.04836 | 92666.64049 | 110195.2604 |
| PC O-18:0_20:4 | 511938.7196 | 493629.3658 | 725669.2257 | 1592390.918 | 732223.6145 | 0 | 1148831.076 | 1820765.897 | 892365.3501 | 1755855.773 | 278330.8939 | 1254364.348 |
| SMGDG O-22:0_28:3 | 1133123.641 | 948899.0576 | 1036319.18 | 1072287.996 | 656581.6781 | 0 | 581732.2213 | 562500.508 | 587281.0879 | 674405.6206 | 625930.6537 | 2570839.3 |

| Metabolites | PBS_1 | PBS_2 | PBS_3 | PBS_4 | BCG_1 | BCG_2 | BCG_3 | BCG_4 | BCGΔ*pe18*_1 | BCGΔ*pe18*_2 | BCGΔ*pe18*_3 | BCGΔ*pe18*_4 |
|---|---|---|---|---|---|---|---|---|---|---|---|---|
| Glucocheirolin | 261895.1182 | 251461.1993 | 578451.5464 | 439553.148 | 676943.2017 | 1031469.401 | 486045.9749 | 471734.7815 | 478178.0027 | 153590.6081 | 266500.9571 | 1019676.934 |
| Mirtazapine | 191381.6352 | 166004.7769 | 173538.3933 | 197106.963 | 184284.3136 | 175634.9287 | 169018.1542 | 173202.3802 | 210614.164 | 190863.5504 | 157706.8736 | 207291.0778 |
| PI 8:0_25:0 | 176652.7195 | 170644.4714 | 192782.1504 | 177259.4096 | 104490.1876 | 88482.05693 | 130186.508 | 102671.951 | 79405.84804 | 57870.3319 | 69202.8657 | 35234.96162 |
| PC O-43:7 | 1872832.008 | 1628856.579 | 1359099.305 | 1900260.302 | 1377594.578 | 1388995.663 | 1630021.717 | 1498227.337 | 1594123.343 | 1506537.36 | 1376007.031 | 1610373.803 |
| LPE(16:0) | 10988896.14 | 9823887.664 | 10810685.21 | 10123160.6 | 44453656.65 | 31776559.01 | 33301374.89 | 34230280.66 | 38366263.51 | 38852382.72 | 55675630.7 | 32494336.66 |
| Xestoaminol C | 6034901.895 | 6000733.959 | 6214120.569 | 6216918.992 | 6322062.715 | 6005137.144 | 6168508.215 | 5477157.705 | 5832835.405 | 6438690.855 | 5286271.493 | 6294413.766 |
| 2-Imino-1-imidazolidineacetic acid | 1011686.143 | 721400.4388 | 748879.6227 | 881247.9064 | 925624.6115 | 930794.7717 | 876095.2791 | 659777.5574 | 717335.5698 | 762181.4134 | 867811.9518 | 600318.0527 |
| 11-Hydroxy-9-tridecenoic acid | 809785.5982 | 684002.911 | 754581.4902 | 696756.0311 | 7132373.677 | 10721493.55 | 6411875.644 | 8130820.021 | 4975582.775 | 2993298.586 | 6575020.375 | 6230663.593 |
| PI 20:1 | 118440.3447 | 93575.34627 | 372296.8384 | 152087.7612 | 106539.326 | 89549.70361 | 145862.4293 | 97830.82083 | 29586.723 | 17395.01617 | 52957.71846 | 19969.57083 |
| 4-(Methylamino)butanoic acid | 8040917.451 | 6197617.415 | 9042932.683 | 5437619.613 | 12567989.11 | 9543779.758 | 7085264.717 | 1794418.242 | 8721292.517 | 2811318.463 | 2196368.576 | 6040477.139 |
| .alpha.-Paxitriol | 1547537.716 | 1621982.24 | 1795326.276 | 364528.8832 | 2075015.829 | 1360801.719 | 2332616.041 | 1963626.219 | 1557572.051 | 1436392.632 | 1897061.89 | 1865791.768 |
| 2-Propenyl 2,4-hexadienoate | 1291436.017 | 849874.07 | 896297.5081 | 2829709.639 | 687551.7688 | 948169.3866 | 3710177.069 | 685768.154 | 1791224.336 | 1159595.407 | 706141.3494 | 3341107.116 |

| Metabolites | PBS_1 | PBS_2 | PBS_3 | PBS_4 | BCG_1 | BCG_2 | BCG_3 | BCG_4 | BCGΔ*pe18*_1 | BCGΔ*pe18*_2 | BCGΔ*pe18*_3 | BCGΔ*pe18*_4 |
|---|---|---|---|---|---|---|---|---|---|---|---|---|
| SHexCer 33:0;2O | 814963.9926 | 825733.6598 | 522490.8641 | 772861.8695 | 1268165.803 | 822179.9824 | 875146.4361 | 684832.2575 | 840560.4489 | 764329.6163 | 1312663.076 | 1165663.522 |
| 16.beta.-Hydroxyfurazabol | 2427107.798 | 3496942.238 | 2119176.956 | 2372148.055 | 4705103.549 | 3632717.984 | 6225635.613 | 4201441.804 | 2946698.43 | 4052990.395 | 4229636.553 | 2468234.01 |
| PC 16:1_20:2 | 1495215.175 | 1122889.102 | 921896.7487 | 1252779.157 | 1147581.875 | 162572.5679 | 1233965.901 | 2285690.312 | 2257141.015 | 1540282.224 | 711947.8352 | 798999.2096 |
| Cys-Gly-Cys | 4494873.135 | 3945945.387 | 6157640.739 | 4912355.549 | 11620010.4 | 10324240.76 | 24469242.41 | 10229852.76 | 7123356.092 | 6899278.633 | 8364096.734 | 6564823.101 |
| Arg-Gln | 2374819.59 | 368031.8822 | 2318165.625 | 856651.4836 | 2556048.341 | 3604029.7 | 19649181.69 | 10041155.18 | 5863185.502 | 3239111.76 | 5163308.177 | 1800220.205 |
| oxacyclohexadecan-2-one | 1370175.293 | 1395899.048 | 1456839.215 | 1278612.492 | 1323479.712 | 1384574.622 | 1508943.792 | 1352522.96 | 1315400.278 | 1327982.381 | 1319516.57 | 1356491.684 |
| PEtOH 22:6_22:6 | 1566451.966 | 1342186.282 | 596375.0076 | 1358226.15 | 1501622.492 | 0 | 947141.9203 | 1110429.601 | 1717024.476 | 1009300.858 | 980613.7851 | 725431.7061 |
| sn-Glycerol 3-phosphate | 1200206.502 | 958723.1686 | 1700152.759 | 1132852.363 | 4055139.058 | 2177756.045 | 2967153.454 | 2772054.959 | 2969181.527 | 2345501.091 | 2340676.216 | 2029166.069 |
| Syrosingopine | 92327.58165 | 91288.23921 | 87400.5779 | 99048.61469 | 90001.48413 | 101505.8424 | 99747.58405 | 105547.2963 | 80055.53719 | 114062.1938 | 71615.39239 | 92586.38513 |
| M325T332 | 584069.0458 | 645768.0655 | 570481.7479 | 629751.7702 | 1148958.758 | 956073.6418 | 855046.1064 | 898766.4388 | 843427.1123 | 770137.4415 | 790303.2748 | 822255.0899 |
| 1H-Pyrrole-3-carboxylic acid | 888098.8851 | 2465611.562 | 958650.0502 | 835570.2467 | 2929455.25 | 1639408.727 | 1803171.148 | 716263.1485 | 780729.0908 | 2460752.484 | 623977.7045 | 1044883.746 |

| Metabolites | PBS_1 | PBS_2 | PBS_3 | PBS_4 | BCG_1 | BCG_2 | BCG_3 | BCG_4 | BCGΔ *pe18*_1 | BCGΔ *pe18*_2 | BCGΔ *pe18*_3 | BCGΔ*pe18*_4 |
|---|---|---|---|---|---|---|---|---|---|---|---|---|
| Ethyl 4-(1-hydroxy-1-methylethyl)-2-propyl-1H-imidazole-5-carboxylate | 801013.1893 | 857089.2874 | 693848.2174 | 757209.469 | 844818.4444 | 773139.6412 | 726543.7263 | 761615.6451 | 720361.2233 | 762272.9983 | 639370.6912 | 913974.574 |
| 3-[4-(4-Hydroxy-2-quinazolinyl)benzyl]-2,4-imidazolidinedione | 1129892.823 | 1170509.786 | 1064586.046 | 1178081.642 | 1240982.215 | 1160342.277 | 1230633.71 | 1196060.097 | 1570531.458 | 1121316.464 | 1028837.777 | 857530.0143 |
| 5-(3,5-Dinitrophenyl)-1,3,4-thiadiazol-2-amine | 387765.9741 | 328397.719 | 447722.2851 | 361756.6777 | 1925248.031 | 1520722.17 | 1563058.151 | 1552907.062 | 1656454.882 | 1328944.228 | 1413091.824 | 1384478.702 |
| PG 16:0_22:1 | 13530.62069 | 18278.04466 | 0 | 76266.43855 | 598876.0063 | 832896.7077 | 361213.2539 | 490509.3328 | 275758.8857 | 213813.2416 | 326089.3691 | 279541.4351 |
| Isoproturon | 2050873.508 | 3115223.96 | 2259782.274 | 1878437.361 | 2199018.142 | 1911237.194 | 2637989.071 | 2590544.646 | 2552304.002 | 2516535.527 | 2285586.535 | 2615449.775 |
| 2-Naphthalenecarboxamide, N-[2-(4-oxo-1-phenyl-1,3,8-triazaspiro[4.5]dec-8-yl)ethyl]- | 82197.04644 | 106632.8407 | 109863.8825 | 103270.6546 | 202366.2833 | 26890.68671 | 190095.5441 | 164000.0541 | 192353.584 | 181221.4373 | 180220.5146 | 221286.239 |
| SM 8:0;2O/38:9 | 0 | 0 | 6206.300728 | 0 | 206660.9228 | 220768.765 | 231359.4551 | 278381.739 | 72310.38454 | 64454.1716 | 53771.55862 | 68739.76092 |

| Metabolites | PBS_1 | PBS_2 | PBS_3 | PBS_4 | BCG_1 | BCG_2 | BCG_3 | BCG_4 | BCGΔ*pe18*_1 | BCGΔ*pe18*_2 | BCGΔ*pe18*_3 | BCGΔ*pe18*_4 |
|---|---|---|---|---|---|---|---|---|---|---|---|---|
| 4,4,7a-Trimethyl-3a,5,6,7-tetrahydro-3H-indene-1-carboxylic acid | 383780.5015 | 1189272.332 | 486227.8867 | 496805.1928 | 791132.6318 | 524618.3311 | 441868.1301 | 1581205.117 | 516085.7409 | 459198.8139 | 506008.5794 | 764430.6823 |
| SMGDG O-28:4_8:0 | 5048313.549 | 5170554.358 | 473366.858 | 3555876.682 | 715317.6663 | 668794.8151 | 899101.3216 | 825664.9129 | 875574.4447 | 1016749.615 | 852064.2547 | 564396.3722 |
| N-Cyclohexylcyclopropanecarboxamide | 3001420.116 | 3555981.328 | 2131494.876 | 2454169.922 | 5294010.166 | 3397962.712 | 2927236.047 | 2715646.241 | 2506701.654 | 2983282.802 | 2297592.394 | 3229101.527 |
| PA 2:0_24:4 | 86124.34626 | 80355.25476 | 272224.3274 | 109301.6428 | 244968.7845 | 183564.3187 | 262795.4029 | 200461.3166 | 107517.842 | 132134.3604 | 191034.9247 | 75614.25488 |
| {4-[(Cyclopropylcarbonyl)amino]phenyl}acetic acid | 1230535.931 | 1230535.597 | 994609.5023 | 1136891.338 | 1092787.199 | 1101615.646 | 1042452.774 | 1142814.471 | 1087074.103 | 1084541.001 | 1053037.623 | 1073341.857 |
| PC 9:0_30:6 | 1951732.243 | 1735329.282 | 635365.1468 | 1038398.812 | 6502046.249 | 7486072.166 | 2427492.183 | 2850360.944 | 2418404.475 | 3221704.841 | 2328129.875 | 1916027.03 |
| .alpha.-Tocotrienol | 1465603.017 | 2054979.536 | 1162587.382 | 1293321.143 | 1315623.238 | 2091804.979 | 1819003.798 | 1534918.953 | 1669313.258 | 1498598.375 | 1494404.827 | 1195092.475 |
| 3-(Cyclohexylamino)-5,5-dimethylcyclohex-2-en-1-one | 1735198.349 | 1794018.446 | 1630001.703 | 1681507 | 1661603.476 | 1595790.106 | 1753187.673 | 1613321.625 | 1705359.939 | 1667439.969 | 1721676.746 | 1599493.961 |
| SM 9:0;2O/38:1 | 72051.79539 | 72123.84673 | 71959.98376 | 78567.72877 | 87349.532 | 0 | 70114.78824 | 59660.61155 | 42998.02075 | 68130.20395 | 66292.60207 | 43325.93627 |

| Metabolites | PBS_1 | PBS_2 | PBS_3 | PBS_4 | BCG_1 | BCG_2 | BCG_3 | BCG_4 | BCGΔ*pe18*_1 | BCGΔ*pe18*_2 | BCGΔ*pe18*_3 | BCGΔ*pe18*_4 |
|---|---|---|---|---|---|---|---|---|---|---|---|---|
| 1-(Aminomethyl)cyclobutan-1-amine | 2166611.634 | 1730230.576 | 230317.0438 | 2112094.152 | 4535185.408 | 3787276.791 | 2393812.287 | 151516.3817 | 1322952.129 | 3675526.875 | 2785011.251 | 2481132.146 |
| M511T474 | 280912.5284 | 324964.0466 | 545116.6143 | 279714.7302 | 247708.4499 | 258380.1456 | 245441.7501 | 222045.806 | 266296.3699 | 268017.599 | 253731.9055 | 320900.2341 |
| FA 25:0 | 70434.27093 | 78010.23782 | 91365.81831 | 26853.86852 | 166320.7734 | 60879.2249 | 124251.4894 | 305559.1942 | 29318.01874 | 96771.15042 | 336326.0609 | 194261.4933 |
| Phomalone | 3251974.759 | 4232863.686 | 3426801.406 | 3568671.833 | 3559908.171 | 4243208.322 | 4622668.251 | 3922937.878 | 3824988.53 | 3050293.698 | 2896336.823 | 6358454.689 |
| BIBR-1048 (Dabigatran) | 16715.76609 | 0 | 19989.71768 | 0 | 160014.1772 | 106763.4479 | 336427.719 | 173133.9097 | 89104.71964 | 72079.75567 | 199410.0481 | 103344.5894 |
| PC O-38:7 | 10123029.09 | 6938441.372 | 3563055.179 | 6772501.565 | 3106475.812 | 5477068.835 | 2106682.035 | 2216376.956 | 4326525.973 | 3512282.667 | 2679228.786 | 2118025.898 |
| Lycopaoctaene | 4100298.529 | 2569225.619 | 1765462.312 | 1855509.387 | 1274897.209 | 1111873.347 | 1652279.727 | 1342754.015 | 1835667.276 | 588015.4467 | 1099570.565 | 902458.5638 |
| Phenylacetylglycine | 809966.8977 | 706539.9081 | 1280797.876 | 982255.7535 | 1054536.495 | 1106724.197 | 1259774.115 | 1014474.971 | 835413.9057 | 518528.1823 | 611322.8326 | 891237.1934 |
| PC O-41:8 | 919219.3067 | 874237.2852 | 919928.832 | 1064500.866 | 898145.7611 | 1200668.394 | 903086.5169 | 981091.0842 | 817278.2362 | 763406.5638 | 1040731.944 | 997085.7767 |
| D-Lactosyl-.beta.1-1'-D-erythro-sphingosine | 161807.5476 | 123664.0337 | 81347.8051 | 105061.6382 | 142024.83 | 151228.9865 | 165810.2472 | 237872.2767 | 293553.9113 | 176777.0097 | 157547.1226 | 137528.4963 |
| Cer 12:2;2O/6:0 | 694626.6248 | 602403.0927 | 446236.2103 | 795962.5156 | 440231.3895 | 449984.2388 | 743061.9838 | 503137.8615 | 620469.5273 | 883813.5565 | 605529.8461 | 708462.5915 |
| Zolmitriptan | 772775. | 896075. | 744706. | 742810. | 719417. | 843001. | 730795. | 760196. | 786618. | 885488. | 947202. | 743790. |

| Metabolites | PBS_1 | PBS_2 | PBS_3 | PBS_4 | BCG_1 | BCG_2 | BCG_3 | BCG_4 | BCGΔ *pe18*_1 | BCGΔ *pe18*_2 | BCGΔ *pe18*_3 | BCGΔ*pe18*_4 |
|---|---|---|---|---|---|---|---|---|---|---|---|---|
| | 603 | 9947 | 009 | 1306 | 8314 | 9105 | 765 | 0949 | 3644 | 0906 | 1835 | 4141 |
| (3.beta.,8.xi.,9.xi.,14.xi.,17Z)-3-((2-O-(6-Deoxy-.alpha.-L-mannopyranosyl)-.beta.-D-glucopyranosyl)oxy)-16,22-dioxocholesta-5,17-dien-27-yl .beta.-D-glucopyranoside | 178621.9129 | 203695.7798 | 63471.11133 | 143349.2844 | 138551.8421 | 139168.995 | 122489.3028 | 134431.0364 | 150885.0326 | 167095.1939 | 139012.7122 | 107350.6885 |
| Palmitoylglycine | 455678.5043 | 404590.8657 | 249198.7619 | 347269.201 | 452571.8065 | 580212.4525 | 358578.9329 | 422466.2621 | 262715.2375 | 282563.4457 | 379001.1995 | 433593.9588 |
| Inosine | 174399.5873 | 243949.1767 | 426974.8572 | 120805.9064 | 1984092.61 | 646285.6906 | 2919484.911 | 1710512.814 | 918024.2389 | 1674392.261 | 1137276.289 | 208185.9363 |
| Diallyldimethylammonium cation | 2461595.881 | 1078054.27 | 2056722.99 | 2785212.323 | 2036401.052 | 1838854.287 | 4030638.737 | 847857.0572 | 2059459.752 | 1059876.171 | 924602.2452 | 2604873.466 |
| BMP 22:6_22:6 | 2185176.733 | 3630631.595 | 734466.9889 | 2115867.656 | 2217683.372 | 7051317.651 | 1601380.904 | 2932673.507 | 2799041.509 | 1494399.159 | 1350075.556 | 1793212.38 |
| Anisomycin | 1057382.212 | 794293.1249 | 857123.543 | 996950.7639 | 1406960.554 | 939192.9147 | 935439.6892 | 1096589.997 | 924871.8687 | 683667.2152 | 722358.6084 | 821783.6744 |
| Arachidyl_carnitine | 325370.4433 | 354503.7268 | 198477.3272 | 311620.0172 | 120500.1657 | 82686.66377 | 133533.8187 | 189311.0801 | 142174.9643 | 133716.9005 | 112864.5405 | 36938.20336 |
| Cer 16:0;2O/4:0 | 12315533.58 | 8342577.607 | 3710851.992 | 4555244.953 | 3546496.761 | 2411314.761 | 2715267.516 | 1831214.437 | 2155206.721 | 2930089.367 | 4359363.897 | 1901954.24 |

| Metabolites | PBS_1 | PBS_2 | PBS_3 | PBS_4 | BCG_1 | BCG_2 | BCG_3 | BCG_4 | BCGΔ*pe18*_1 | BCGΔ*pe18*_2 | BCGΔ*pe18*_3 | BCGΔ*pe18*_4 |
|---|---|---|---|---|---|---|---|---|---|---|---|---|
| SL 10:0;O/16:0;O | 218612.4599 | 382771.0168 | 148861.2363 | 341749.1585 | 160378.9109 | 227670.9722 | 335665.9633 | 313652.4718 | 149465.2159 | 126669.2008 | 96937.58307 | 107460.1983 |
| 5-[(E)-(4-Nitrophenyl)diazenyl]-2,4,6(1H,3H,5H)-pyrimidinetrione | 1048320.034 | 652151.3809 | 1825068.802 | 2149448.971 | 1346513.244 | 1555435.64 | 1115348.259 | 1759430.874 | 783363.5519 | 583031.3253 | 1058632.984 | 451947.1283 |
| Beauvericin | 46987.25124 | 19511.85119 | 24637.46155 | 0 | 168495.0829 | 219806.0158 | 164295.6877 | 188788.5751 | 305618.8152 | 248963.5575 | 204139.7625 | 234782.9991 |
| Indigo | 399699.1411 | 431283.4912 | 348145.8986 | 33031897.09 | 327200.2738 | 5781189.418 | 672961.4658 | 2841021.489 | 355202.0232 | 372171.3186 | 351525.028 | 1205966.117 |
| Xanthydrol | 155989.4588 | 100387.2386 | 75721.03618 | 84787.00359 | 1184272.842 | 1347528.109 | 1315057.222 | 1338936.332 | 1513683.492 | 1269206.706 | 1523503.036 | 1592609.495 |
| FA 21:1 | 0 | 11014.22515 | 46912.28588 | 10490.65817 | 457445.8652 | 969328.7151 | 348788.4656 | 380007.3227 | 262302.6859 | 156010.4369 | 512552.4212 | 332511.6645 |
| PC 22:5_22:5 | 2234382.853 | 3369853.618 | 1406665.979 | 2056084.053 | 1801668.226 | 2320385.843 | 1593391.174 | 1703564.623 | 2640862.837 | 1793447.27 | 1615888.74 | 2023547.233 |
| SM 14:3;2O/5:0 | 1244909.05 | 1262797.528 | 1159504.375 | 1245696.491 | 1296752.489 | 1090228.086 | 1306161.832 | 1128468.921 | 1248889.271 | 1221156.59 | 1004482.884 | 1208854.858 |
| Intepirdine | 72868.74146 | 42595.0256 | 110989.7076 | 43672.62089 | 2505240.594 | 1275529.725 | 2129981.487 | 1073102.329 | 1577845.692 | 1452598.867 | 1653677.989 | 1156363.446 |
| 5-Chloro-3-[(1E,3E)-3,5-dimethylhepta-1,3-dienyl]-7,8a-dihydroxy-1-methoxy | 2769853.718 | 1434450.499 | 3637389.669 | 1978156.376 | 1866662.707 | 1608440.313 | 3815480.052 | 3050340.174 | 1844370.411 | 1827716.034 | 1670302.067 | 667588.7751 |

| Metabolites | PBS_1 | PBS_2 | PBS_3 | PBS_4 | BCG_1 | BCG_2 | BCG_3 | BCG_4 | BCGΔ *pe18*_1 | BCGΔ *pe18*_2 | BCGΔ *pe18*_3 | BCGΔ*pe18*_4 |
|---|---|---|---|---|---|---|---|---|---|---|---|---|
| -7-methyl-1H-isochromene-6,8-dione | | | | | | | | | | | | |
| [1]Benzopyrano[3,4-b]furo[2,3-h][1]benzopyran-6(12H)-one, 1,2-dihydro-9-hydroxy-2-[1-(hydroxymethyl)ethenyl]-8-methoxy- | 377950.7518 | 495444.7434 | 513967.1008 | 820007.0211 | 538224.6575 | 568050.9236 | 602644.2874 | 694871.8924 | 462902.9181 | 849771.8151 | 338043.068 | 754606.4704 |
| o-Toluidine | 3170196.855 | 1754526.311 | 2436501.201 | 2225253.181 | 2274328.291 | 1442446.662 | 6386840.737 | 2366750.698 | 2579679.95 | 2417674.644 | 1481256.388 | 2214952.24 |
| Cer 9:0;2O/18:1 | 145294.4169 | 139619.3214 | 186752.8501 | 75011.28212 | 213320.4401 | 324537.7451 | 248439.3147 | 346436.3673 | 147711.9817 | 122672.1756 | 226595.1459 | 304969.7342 |
| PE O-16:1_20:5 | 4542385.312 | 4885155.704 | 3513095.742 | 3092580.061 | 5800982.148 | 5050769.57 | 3854386.473 | 5053369.603 | 4647502.515 | 4175989.192 | 4418334.809 | 3822386.759 |
| 8R-HpODE | 5388246.242 | 7031507.947 | 6253776.615 | 7942039.773 | 6838181.561 | 8600721.591 | 5716189.522 | 6835788.679 | 4801535.882 | 5014697.108 | 7130125.556 | 6343927.886 |
| PC O-34:9 | 568785.3052 | 960395.3295 | 875092.9166 | 278063.9282 | 442382.9274 | 966940.4922 | 661104.5635 | 506749.671 | 547061.6562 | 616329.3157 | 356535.4655 | 570005.8157 |
| PI 16:0_20:5 | 664364.671 | 472537.8782 | 410854.3222 | 637077.4722 | 1066856.434 | 1255801.741 | 1905015.567 | 1360351.148 | 2103112.839 | 1719276.034 | 1040376.31 | 1462534.74 |

| Metabolites | PBS_1 | PBS_2 | PBS_3 | PBS_4 | BCG_1 | BCG_2 | BCG_3 | BCG_4 | BCGΔ*pe18*_1 | BCGΔ*pe18*_2 | BCGΔ*pe18*_3 | BCGΔ*pe18*_4 |
|---|---|---|---|---|---|---|---|---|---|---|---|---|
| Trandolaprilat | 160216.7574 | 130422.0585 | 107695.9935 | 146217.0805 | 174013.3881 | 116405.8301 | 148791.5657 | 162973.5601 | 178259.7709 | 152462.6341 | 154237.0545 | 126133.7001 |
| NAOrn 11:0/10:0 | 4071121.408 | 4502594.853 | 3344489.335 | 5094787.129 | 2080824.856 | 2923728.811 | 3794291.01 | 3147897.648 | 3211169.317 | 2950485.01 | 2072019.064 | 2221614.912 |
| PI 30:5 | 5142.954297 | 14975.30497 | 62809.12256 | 8954.375249 | 35726.09265 | 55338.18686 | 87890.76496 | 72373.11253 | 39300.67589 | 62519.23618 | 40609.76062 | 22343.20861 |
| PC(34:3) | 34469892.4 | 38746631 | 36827572.54 | 37086058 | 28563584.15 | 23099026.09 | 31538531.79 | 25253321.91 | 25078269.21 | 32401326.97 | 28827043.07 | 13506717.66 |
| Histidinol | 488787.2166 | 442736.5476 | 400537.8116 | 414509.4872 | 9153398.506 | 18372503.92 | 7599559.73 | 11238200.49 | 15858118.13 | 6098095.554 | 9314882.896 | 13226699.18 |
| LPI(18:1) | 97579.25247 | 100910.8493 | 168698.1755 | 110045.2372 | 862491.1926 | 478348.8082 | 606886.2837 | 504309.0452 | 604913.6651 | 466253.6565 | 597362.3898 | 518239.7056 |
| 2-O-Methylcytosine | 339804.3707 | 403701.8137 | 560585.4358 | 659650.5024 | 577097.677 | 638586.1612 | 646794.5889 | 560632.106 | 610504.5871 | 640890.0526 | 524412.9636 | 860798.6736 |
| NAE 26:7 | 198920.832 | 188853.955 | 199877.333 | 198176.8265 | 182016.1578 | 197873.8601 | 199042.0401 | 179864.9217 | 201095.0658 | 205055.4086 | 195109.6905 | 193882.0824 |
| Car(18:0) | 21305064.44 | 19845258.5 | 16234677.07 | 18520189.35 | 15606126.94 | 10348595.84 | 17273749.45 | 15976654.76 | 12914372.26 | 51618.24586 | 10375918.11 | 5818344.156 |
| LPE 20:1 | 728874.1783 | 707772.3863 | 837298.9733 | 642154.8534 | 1833358.367 | 1200681.315 | 1863206.521 | 1529121.223 | 1463510.985 | 1372528.647 | 1605818.707 | 659436.5398 |
| PG 11:0_22:3 | 20517.37728 | 0 | 8020.66554 | 0 | 1616666.322 | 533627.5969 | 1988220.018 | 619552.3599 | 739954.1409 | 1628963.997 | 1439839.343 | 483840.1455 |
| 2,6-Xylidine | 702206.4618 | 706564.7342 | 714701.0862 | 693808.7786 | 1092954.17 | 818116.9835 | 854880.7698 | 802311.682 | 845138.1166 | 603227.8023 | 2153070.183 | 810588.0908 |

| Metabolites | PBS_1 | PBS_2 | PBS_3 | PBS_4 | BCG_1 | BCG_2 | BCG_3 | BCG_4 | BCGΔ*pe18*_1 | BCGΔ*pe18*_2 | BCGΔ*pe18*_3 | BCGΔ*pe18*_4 |
|---|---|---|---|---|---|---|---|---|---|---|---|---|
| Glyco-gamma-muricholic acid | 54939.00816 | 24803.52862 | 105287.8449 | 47773.96758 | 96299.66663 | 87161.91962 | 115579.4757 | 76552.66033 | 67234.95877 | 54039.13567 | 50219.10622 | 74054.08849 |
| Diclofenac | 170660.6128 | 195204.9115 | 165666.9083 | 177118.2728 | 188564.5378 | 183825.6423 | 2865490.738 | 41928.34023 | 185163.3804 | 2513499.095 | 2340559.104 | 145981.0325 |
| Delphinidin_3-lathyroside_5-glucoside | 8058899.728 | 3353087.276 | 9761374.306 | 11694774.16 | 10248944.59 | 8697132.758 | 9822403.211 | 8020930.757 | 7846167.367 | 9343518.101 | 10870733.79 | 4194701.585 |
| Dextrorphan O-.beta.-D-glucuronide | 147573.5494 | 144205.812 | 105470.6665 | 137913.0872 | 239584.2709 | 173325.6359 | 194343.5279 | 138580.2701 | 132128.4371 | 137499.3899 | 126629.5097 | 132624.3798 |
| PI 18:0_22:5 | 4348935.647 | 2224706.61 | 3517108.112 | 2649046.992 | 5464828.268 | 5999071.616 | 4594646.315 | 4785012.35 | 4937553.635 | 5826420.421 | 5413569.747 | 3015998.208 |
| N-Cyclohexylbenzenecarbothioamide | 769892.4389 | 701629.5826 | 799387.7429 | 836430.3257 | 1280094.181 | 1006919.471 | 1574923.042 | 1136619.238 | 1273766.709 | 805395.4115 | 883350.5772 | 909758.3948 |
| 3-Hydroxyphenylacetic acid | 361333.3646 | 285614.0106 | 397206.7797 | 353733.6495 | 545920.8451 | 503696.8259 | 547295.6313 | 452433.4357 | 426242.1546 | 427047.2128 | 413228.5775 | 405053.9552 |
| Panaxacol | 860164.6559 | 883218.9544 | 751903.0759 | 712677.4991 | 931723.8195 | 902536.0619 | 899343.1526 | 756899.0211 | 816850.8982 | 849047.9033 | 992444.3809 | 792098.1755 |
| Hexadecatrienoylcarnitine (Car(16:3)) | 925060.1414 | 905697.4735 | 1676577.652 | 1263475.337 | 263963.9004 | 212296.3596 | 282089.0913 | 303476.9261 | 71218.09289 | 257681.5773 | 201358.4629 | 170784.8716 |
| Cer 8:0;2O/26:2 | 42615.69938 | 33125.66253 | 23678.84498 | 48071.09511 | 76978.07644 | 107999.447 | 79736.09575 | 95044.03797 | 60825.18174 | 76798.19707 | 77390.34406 | 59649.11906 |
| IRIGENIN | 437629.3259 | 326119.4575 | 459220.7853 | 322018.5169 | 3564051.149 | 3285687.53 | 3004854.403 | 2997299.207 | 3382633.53 | 2464145.274 | 3195149.34 | 2902701.528 |

| **Metabolites** | **PBS_1** | **PBS_2** | **PBS_3** | **PBS_4** | **BCG_1** | **BCG_2** | **BCG_3** | **BCG_4** | **BCGΔ*pe18*_1** | **BCGΔ*pe18*_2** | **BCGΔ*pe18*_3** | **BCGΔ*pe18*_4** |
|---|---|---|---|---|---|---|---|---|---|---|---|---|
| Alanine | 921091.933 | 1726621.24 | 1850578.28 | 849914.3759 | 9985984.966 | 3622395.064 | 9873332.789 | 5661336.361 | 4246504.242 | 5472696.098 | 5088721.626 | 4503867.145 |
| 3,4-dichlorobenzoate | 988282.2263 | 633662.4705 | 991266.3697 | 1586568.443 | 272299.0711 | 972310.6522 | 400122.8593 | 2606356.591 | 215468.3119 | 2513195.458 | 930822.6914 | 1763284.16 |
| N-Oleoyl-L-serine | 119611.0224 | 104542.0564 | 80147.57148 | 69659.41356 | 598117.9042 | 665648.3385 | 404579.7861 | 468995.5103 | 339803.3696 | 207000.1352 | 449747.407 | 335263.3663 |
| Dexanabinol | 1247318.087 | 5192079.987 | 1956338.019 | 986961.3998 | 871896.2948 | 1724212.458 | 393558.3782 | 661136.263 | 1129737.449 | 450813.7911 | 473650.0623 | 572287.1019 |
| Dextrorphan | 161227.3691 | 96302.16222 | 164308.2972 | 152031.5457 | 91670.98489 | 140546.5834 | 132434.1919 | 166441.5312 | 168310.076 | 86036.96352 | 157077.0621 | 176217.1693 |
| ulvanobiouronic acid 3-sulfate A | 10071.54653 | 0 | 0 | 26081.72034 | 934455.367 | 1115016.599 | 548311.9256 | 709178.7098 | 1393796.296 | 1325251.627 | 1159373.996 | 639371.5988 |
| LPE 21:2 | 649945.8819 | 1044040.944 | 792701.164 | 853967.763 | 741472.7429 | 970685.5268 | 676795.5757 | 545423.1287 | 489494.0283 | 490143.824 | 627260.9995 | 447986.1145 |
| 2-Phenylbenzo[d]imidazo[2,1-b]thiazole | 334240.9186 | 258242.9832 | 546411.7612 | 324239.0042 | 1428561.385 | 987158.5992 | 1422739.643 | 1101781.812 | 1552489.128 | 1476779.26 | 1475754.329 | 1358986.562 |
| 3-[Hexadecyl(dimethyl)ammonio]-1-propanesulfonate | 1400582.11 | 1398685.149 | 1251174.743 | 1294263.065 | 1470654.099 | 1327372.995 | 1337610.045 | 4038979.583 | 4004670.772 | 1387908.052 | 1374543.014 | 1386008.893 |
| LPC O-20:4 | 2343875.32 | 2988075.302 | 2873100.422 | 2707688.521 | 1560764.462 | 2677824.6 | 2274931.519 | 1764954.088 | 1659120.175 | 1452468.826 | 2107652.478 | 1296153.246 |
| SMGDG O-9:0_22:3 | 809910.6227 | 351966.9426 | 1084898.007 | 611578.1643 | 712931.6203 | 262203.4678 | 631275.9111 | 396723.1577 | 524765.1903 | 632242.9391 | 482781.2486 | 200015.7026 |

| Metabolites | PBS_1 | PBS_2 | PBS_3 | PBS_4 | BCG_1 | BCG_2 | BCG_3 | BCG_4 | BCGΔ *pe18*_1 | BCGΔ *pe18*_2 | BCGΔ *pe18*_3 | BCGΔ*pe18*_4 |
|---|---|---|---|---|---|---|---|---|---|---|---|---|
| 2-Hydroxymyristic acid | 1344186.688 | 777329.5645 | 1009207.453 | 837519.6082 | 876009.9614 | 879216.4557 | 855759.1731 | 892478.7537 | 871512.8704 | 957057.2981 | 907487.4181 | 979080.2339 |
| 2-Hydroxybenzothiazole | 400755.518 | 384580.8256 | 449495.8026 | 590480.9713 | 337227.2681 | 375244.6203 | 317090.1051 | 497563.1067 | 285759.9722 | 292002.7334 | 311246.1597 | 454325.1054 |
| MGDG O-20:4_20:3 | 0 | 0 | 0 | 0 | 770496.924 | 973086.2278 | 784542.601 | 747461.6854 | 1411931.317 | 929527.3796 | 1043728.812 | 925573.8716 |
| 7-Hydroxyfluphenazine | 231344.1274 | 187165.5177 | 416092.5877 | 202915.6175 | 1553508.953 | 1180277.63 | 1621713.911 | 1325792.403 | 1320464.077 | 1341740.698 | 1276812.18 | 1099958.625 |
| Methanone, (2-iodo-5-nitrophenyl)[1-[[(2S)-1-methyl-2-piperidinyl]methyl]-1H-indol-3-yl]- | 81218.71795 | 505839.2606 | 1206969.231 | 768298.2844 | 65750.3673 | 19301.39701 | 155596.8905 | 0 | 75284.08631 | 100048.8595 | 0 | 1546761.031 |
| PI-Cer 12:1;2O/22:2;O | 114968.903 | 53278.48925 | 169680.3939 | 116860.9962 | 996234.0506 | 647395.0264 | 756064.9553 | 821632.545 | 1121006.762 | 838854.1629 | 668307.0675 | 569924.1012 |
| Ethanone, 1-[3,5-bis(1,1-dimethylethyl)-4-hydroxyphenyl]-2-[2-[(3-hydroxypropyl)amino]-5,6-dimethyl-1H-benzimidazol-1-yl]- | 694368.1048 | 557126.5896 | 753950.4874 | 780962.8595 | 626178.7764 | 747970.224 | 590281.4648 | 631054.6951 | 3267077.958 | 2082097.272 | 706325.4836 | 744822.8655 |

| Metabolites | PBS_1 | PBS_2 | PBS_3 | PBS_4 | BCG_1 | BCG_2 | BCG_3 | BCG_4 | BCGΔ*pe18*_1 | BCGΔ*pe18*_2 | BCGΔ*pe18*_3 | BCGΔ*pe18*_4 |
|---|---|---|---|---|---|---|---|---|---|---|---|---|
| 4-Imidazolidineheptanoic acid, 3-[(2-cyclohexyl-2-hydroxyethyl)amino]-2,5-dioxo-1-(phenylmethyl)- | 906410.4487 | 851834.734 | 508510.6097 | 701645.471 | 481512.5762 | 308236.6935 | 425394.3095 | 442579.4043 | 437111.8598 | 381211.6303 | 461799.6664 | 269305.6757 |
| hyperin | 2466363.941 | 2216415.556 | 2391093.706 | 2622770.314 | 2211108.537 | 2826639.511 | 2969372.798 | 2995756.724 | 2338131.463 | 2217235.829 | 1876710.087 | 1443007.254 |
| (3E,7E,11E,15E)-5,9,13,17,18-Pentahydroxy-4,6,8,10-tetramethyl-2-[(E)-2-methylbut-2-enyl]nonadeca-3,7,11,15-tetraenedioic acid | 263718.3748 | 345881.01 | 238654.7257 | 324712.6101 | 263674.35 | 220819.2655 | 360070.4501 | 285455.5971 | 294023.7468 | 263142.6726 | 292771.8511 | 269499.7795 |
| N-Phenyl-1-naphthylamine | 1947712.543 | 1643983.279 | 2050770.747 | 1789618.775 | 2280195.814 | 2197229.634 | 1938927.166 | 2193307.89 | 2224928.36 | 1949463.568 | 2231315.896 | 1868732.645 |
| 8-Bromo-2-chloro-6-fluoroquinazoline | 1414563.129 | 1713988.091 | 1627456.252 | 1597336.932 | 1901752.063 | 1891559.37 | 1836197.804 | 1881070.028 | 1771033.511 | 1821861.385 | 3684735.139 | 2186791.348 |
| all-trans-Phytoene | 1158948.864 | 1769855.696 | 649201.9337 | 762956.9294 | 414158.4446 | 2275716.585 | 775976.8813 | 940128.2595 | 695037.1122 | 567203.3574 | 746548.7344 | 641889.2971 |
| Oxamate | 2858165.639 | 1884557.99 | 2420076.621 | 2733570.902 | 618108.1209 | 2706757.296 | 2577396.959 | 3003186.001 | 1011590.901 | 1856385.361 | 2784626.564 | 2637096.494 |
| Cholic acid | 106374. | 47416.3 | 60640.5 | 47723.5 | 91354.0 | 188569. | 87493.7 | 146250. | 56373.7 | 69513.1 | 134251. | 59969.4 |

| **Metabolites** | **PBS_1** | **PBS_2** | **PBS_3** | **PBS_4** | **BCG_1** | **BCG_2** | **BCG_3** | **BCG_4** | **BCGΔ*pe18*_1** | **BCGΔ*pe18*_2** | **BCGΔ*pe18*_3** | **BCGΔ*pe18*_4** |
|---|---|---|---|---|---|---|---|---|---|---|---|---|
| 7-sulfate | 6773 | 8722 | 376 | 4767 | 9623 | 5508 | 4477 | 1427 | 6 | 1459 | 0444 | 5474 |
| 10-Oxomorphine | 697070.7348 | 606052.0049 | 1105416.014 | 922141.5974 | 1493731.982 | 1113490.086 | 1380996.847 | 1405037.498 | 953013.3872 | 791433.8504 | 1224432.64 | 957253.3391 |
| 1-Cyclobutyl-4-piperidinamine | 336777.8662 | 380797.4056 | 325235.7251 | 357204.9869 | 324595.6444 | 336909.1089 | 301124.6069 | 313772.6367 | 304618.7361 | 341152.2132 | 326560.3422 | 313586.213 |
| CerP 19:1;2O/18:1 | 6098141.618 | 6750063.324 | 5542913.908 | 7089892.662 | 18137741.19 | 15493098.63 | 7451155.523 | 18339055.37 | 9425488.008 | 4852977.93 | 4342002.635 | 11069129.65 |
| ST 24:1;O5;G;S | 49395.8099 | 64676.18491 | 34854.50279 | 100758.4093 | 235092.4126 | 243682.0931 | 266220.7141 | 184836.2413 | 215170.7661 | 242150.6499 | 77013.69706 | 170320.7668 |
| Cyclo(L-Leu-L-Pip-L-Aoe-D-Phe) | 786677.2442 | 635605.3257 | 1055060.384 | 1013800.978 | 574775.8816 | 969858.9112 | 1071910.785 | 893473.4825 | 1015096.886 | 965501.8614 | 896314.0249 | 998791.8563 |
| PE(P-18:1(9Z)/20:5(5Z,8Z,11Z,14Z,17Z)) | 1459971.777 | 4927702.059 | 4295328.755 | 3915618.075 | 7951025.109 | 3562569.045 | 3509661.814 | 2996246.738 | 5993520.22 | 1556787.565 | 3549653.216 | 2304953.677 |
| PC 11:0_11:0 | 482646.1873 | 526162.6072 | 255244.7466 | 368993.0981 | 929518.4681 | 845398.9029 | 519106.418 | 739251.7368 | 815166.1237 | 495244.4841 | 528981.9498 | 452049.7179 |
| Cholylphenylalanine | 973246.899 | 989804.3104 | 794840.4915 | 906145.4777 | 971557.7224 | 1015334.392 | 953585.6111 | 947537.1508 | 889051.4351 | 918658.5388 | 896665.7564 | 802383.6249 |
| PC 25:3 | 33946.42906 | 73781.43405 | 28474.24714 | 56505.82633 | 279267.3676 | 227871.6018 | 293602.5 | 36110.91436 | 179331.4231 | 223512.1605 | 218649.7796 | 166551.3258 |
| 3-(glyceryl)-diphospho-5'-guanosine | 535144.1626 | 375637.9052 | 854347.9481 | 860107.9936 | 545223.2721 | 415616.677 | 548639.8925 | 578864.9314 | 365540.7304 | 183473.2021 | 493486.2708 | 944391.5541 |
| SPB 14:1;2O | 2009371.613 | 2457646.724 | 2175303.033 | 2186624.025 | 3270796.195 | 3054627.886 | 3014018.695 | 2850556.91 | 2858173.817 | 3103656.158 | 2976971.314 | 3375938.429 |

| **Metabolites** | **PBS_1** | **PBS_2** | **PBS_3** | **PBS_4** | **BCG_1** | **BCG_2** | **BCG_3** | **BCG_4** | **BCGΔ*pe18*_1** | **BCGΔ*pe18*_2** | **BCGΔ*pe18*_3** | **BCGΔ*pe18*_4** |
|---|---|---|---|---|---|---|---|---|---|---|---|---|
| PG(20:4/20:4) | 2394502.064 | 3356127.468 | 1133998.679 | 2028242.416 | 448197.8433 | 620525.5435 | 542927.9893 | 561386.4842 | 634558.9068 | 527231.2789 | 354588.3599 | 442089.4588 |
| (S,S)-(+)-Tetrandrine | 147069.0941 | 147225.5177 | 136004.2945 | 155662.7782 | 57012.07087 | 182499.8566 | 265890.6565 | 126230.245 | 356148.5473 | 71066.34133 | 274735.2835 | 208603.6772 |
| 2,4-Dinitrophenol | 1709727.322 | 1564891.492 | 1698500.589 | 2245532.066 | 1058763.046 | 1894420.057 | 1223612.918 | 1507364.333 | 1186297.009 | 2039935.957 | 1428411.664 | 2314286.229 |
| Licoricesaponin G2 | 139472.5697 | 217244.6907 | 34877.28213 | 136521.8041 | 80032.97246 | 107951.0625 | 97672.01618 | 93196.61104 | 146841.4093 | 115567.2647 | 83839.71866 | 79007.9259 |
| Dihydrouracil | 1266243.5 | 1327758.488 | 1729149.259 | 1861695.752 | 1208108.418 | 1145385.598 | 1253282.953 | 1310383.924 | 1290706.348 | 1261003.646 | 1159998.778 | 1492656.391 |
| Pro-Leu | 8862143.656 | 6540662.714 | 9656053.19 | 7460452.929 | 7881579.255 | 4889214.727 | 8411087.37 | 6527023.099 | 4403696.541 | 5600029.833 | 6278854.172 | 3601650.988 |
| N-Desmethylcyclobenzaprine | 2634172.396 | 2622275.096 | 2478054.401 | 2283116.861 | 2104955.619 | 2291814.812 | 2041445.2 | 2150572.846 | 2255095.32 | 2389475.664 | 2228017.753 | 2861209.087 |
| Hex3Cer 21:3;2O | 251341.0947 | 281477.7894 | 146147.7089 | 240176.6547 | 194258.7955 | 203794.1563 | 209301.5187 | 200164.2835 | 228943.5857 | 238559.3316 | 236824.3859 | 171308.4952 |
| 3-Chloro-2-hydroxy-N,N,N-trimethyl-1-propanaminium cation | 1086212.672 | 718228.8918 | 1457168.888 | 903770.572 | 1380651.49 | 1374345.137 | 1378269.08 | 1437511.129 | 1271968.52 | 242322.1569 | 749755.4013 | 4753353.01 |
| SM 34:6;3O | 2002028.788 | 2893449.98 | 2751686.828 | 1931982.736 | 1847085.167 | 2937385.641 | 3072585.025 | 2105817.326 | 2712319.652 | 3068530.148 | 3038557.309 | 2656222.897 |
| PS 24:2_16:4 | 1884459.992 | 2075771.969 | 2831768.782 | 2157482.166 | 2187865.449 | 391286.5417 | 1848658.805 | 3097605.914 | 1923879.294 | 1832311.699 | 1984384.123 | 2352413.423 |

| **Metabolites** | **PBS_1** | **PBS_2** | **PBS_3** | **PBS_4** | **BCG_1** | **BCG_2** | **BCG_3** | **BCG_4** | **BCGΔ*pe18*_1** | **BCGΔ*pe18*_2** | **BCGΔ*pe18*_3** | **BCGΔ*pe18*_4** |
|---|---|---|---|---|---|---|---|---|---|---|---|---|
| DGDG O-14:1_28:7 | 0 | 0 | 0 | 0 | 439814.5166 | 346842.3633 | 432575.4579 | 403598.0493 | 709738.1435 | 498320.8181 | 625722.275 | 471415.6318 |
| 3-Hydroxystanozolol glucuronide | 67606.23264 | 41302.82799 | 26500.3511 | 62475.07929 | 89584.13043 | 97634.48902 | 51692.61946 | 64832.1279 | 42015.84123 | 83863.56207 | 136586.7521 | 72793.73888 |
| PE 25:4 | 200156.5263 | 207505.7615 | 248941.4157 | 261736.4701 | 165346.7413 | 127511.0948 | 181398.7262 | 182756.4132 | 201183.4111 | 178542.3148 | 215911.3924 | 130881.9699 |
| LPE O-14:1 | 601304.5785 | 413058.6679 | 575754.5551 | 478579.3849 | 1087386.24 | 979767.7781 | 907154.9705 | 941663.0042 | 950955.5238 | 858226.2437 | 913151.5956 | 663099.4925 |
| 2-Chloro-6-fluoro-1H-1,3-benzodiazole | 312284.9267 | 267218.9453 | 493930.6111 | 414893.2123 | 725719.2624 | 790056.9214 | 763285.6252 | 531470.6473 | 672546.9952 | 584686.4827 | 622605.6999 | 511277.5375 |
| DL-Norvaline | 2955215.378 | 2297966.296 | 14131757.31 | 2766872.233 | 3328072.356 | 2589234.605 | 2563012.588 | 2466930.428 | 4316302.057 | 1430005.346 | 2076617.465 | 2047702.19 |
| (S)-3,5-Dihydroxyphenylglycine | 992750.9305 | 1202119.92 | 1526876.543 | 2300343.413 | 1459704.194 | 1557948.914 | 984545.5901 | 1362590.869 | 1150840.016 | 642041.983 | 637110.9418 | 1840932.52 |
| Carbamic acid, N-10-undecyn-1-yl-, 3'-(aminocarbonyl)[1,1'-biphenyl]-3-yl ester | 97200.95263 | 104384.3749 | 192075.8376 | 97799.19624 | 174815.036 | 103425.5645 | 215365.9127 | 146141.507 | 97177.66477 | 153098.5317 | 108923.3926 | 62157.318 |
| 5-(p-Chlorobenzylidene)thiobarbituric acid | 60739.49861 | 105792.6844 | 174575.7853 | 77054.94275 | 17809.95742 | 29698.79608 | 40988.20716 | 41366.10924 | 43135.34054 | 12590.66684 | 20366.9498 | 0 |
| Hexanoylcarnitine | 127156 | 110994 | 280580 | 167665 | 272886 | 359706 | 384163 | 337734 | 187081 | 259234 | 364720 | 123258 |

| Metabolites | PBS_1 | PBS_2 | PBS_3 | PBS_4 | BCG_1 | BCG_2 | BCG_3 | BCG_4 | BCGΔ *pe18*_1 | BCGΔ *pe18*_2 | BCGΔ *pe18*_3 | BCGΔ*pe18*_4 |
|---|---|---|---|---|---|---|---|---|---|---|---|---|
| (Car(6:0)) | 3.845 | 4.195 | 2.296 | 2.287 | 1.321 | 9.141 | 3.068 | 2.752 | 0.331 | 3.213 | 1.462 | 2.071 |
| SM 40:7;3O | 2490161.373 | 2572868.954 | 1333374.017 | 1717099.66 | 835950.999 | 1031416.809 | 781285.8232 | 1091514.305 | 1177086.324 | 854421.6618 | 629329.178 | 851879.7841 |
| 2,4-Pyrimidinediamine, 5-[[3-methoxy-4-[(4-methoxyphenyl)methoxy]phenyl]methyl]- | 479467.3549 | 869634.8021 | 505845.9263 | 1420565.902 | 317837.673 | 460120.8855 | 455701.3052 | 529728.2257 | 358974.7225 | 335287.6989 | 350832.1396 | 580613.3281 |
| Ganoderenic_acid_D | 2292880.172 | 951176.1206 | 817943.8265 | 870573.2383 | 1868482.682 | 2838276.767 | 2282391.283 | 2882939.105 | 1284902.802 | 800092.1923 | 1806387.865 | 2947225.977 |
| LDGCC 10:0 | 63346.23678 | 121388.6187 | 59980.19521 | 347164.9155 | 141222.1638 | 67093.2339 | 142733.5471 | 66670.34867 | 78832.52196 | 71757.02415 | 213876.7001 | 82981.44428 |
| PG(18:2(9Z,12Z)/18:3(6Z,9Z,12Z)) | 1136474.298 | 877818.8872 | 775528.0061 | 486430.6557 | 1791629.229 | 2510089.284 | 2051197.453 | 1979731.812 | 3279620.949 | 2137764.692 | 1506783.178 | 1557254.397 |
| [DAla2] .beta.-Casomorphin (1-3), amide, bovine | 88083.54753 | 86337.6957 | 122528.1847 | 92009.55276 | 219284.3887 | 143841.4576 | 207348.1255 | 166452.4596 | 246852.7989 | 258321.1394 | 278481.3895 | 162990.5392 |
| PI 27:0_18:5 | 0 | 13366.74353 | 0 | 9074.937998 | 649859.6343 | 744356.2129 | 672713.738 | 648754.9052 | 117870.9097 | 151678.7479 | 226639.6914 | 104633.6281 |
| PC 18:0_20:3;2O | 0 | 0 | 0 | 0 | 861764.2571 | 830735.4483 | 845398.5028 | 745076.7096 | 399188.8227 | 270456.4235 | 204280.6377 | 295655.0356 |
| Neantine | 1356785.911 | 2302691.548 | 1530582.985 | 1721511.521 | 2278589.218 | 3006415.746 | 1691811.811 | 1869735.353 | 1851068.534 | 2342733.698 | 2622696.69 | 2282344.385 |

| Metabolites | PBS_1 | PBS_2 | PBS_3 | PBS_4 | BCG_1 | BCG_2 | BCG_3 | BCG_4 | BCGΔ*pe18*_1 | BCGΔ*pe18*_2 | BCGΔ*pe18*_3 | BCGΔ*pe18*_4 |
|---|---|---|---|---|---|---|---|---|---|---|---|---|
| SM 46:9;3O | 0 | 0 | 0 | 0 | 79727.60736 | 157344.8761 | 84151.6563 | 113123.9496 | 56177.38331 | 53978.11323 | 56895.6698 | 64864.59343 |
| NAE 17:2 | 3879102.438 | 3813152.263 | 3485350.136 | 3670005.393 | 3282006.912 | 3085695.068 | 3110540.719 | 3128558.767 | 3060729.36 | 3201764.375 | 3459427.452 | 2680553.562 |
| Acetylleucine | 222429.9679 | 290074.5913 | 337934.8265 | 317352.1325 | 344502.4002 | 329626.643 | 358981.1602 | 316506.563 | 299587.0461 | 269234.0207 | 334671.9071 | 290042.5941 |
| Calpain Inhibitor II | 154856.6351 | 180143.7846 | 223968.7053 | 191496.0962 | 163622.6197 | 204327.3355 | 31448.54421 | 185090.2739 | 160895.1527 | 127280.8798 | 176422.2826 | 174465.737 |
| Psychosine | 529839.8281 | 478179.1056 | 350591.4114 | 365484.9249 | 471969.9901 | 363809.855 | 501359.2686 | 450408.8268 | 503939.7628 | 429517.9687 | 517906.1629 | 300901.7714 |
| PI 19:2 | 1053423.177 | 957115.461 | 1770785.078 | 1173477.411 | 761444.7479 | 597740.5053 | 944099.5473 | 649259.8429 | 323199.5588 | 339890.315 | 412011.81 | 295174.5371 |
| 1,2-Dipalmitoyl-sn-glycero-3-phospho-(1'-rac-glycerol) | 4606744.918 | 4816408.481 | 3050406.072 | 3875598.76 | 9341582.954 | 6253553.666 | 6986571.476 | 7698724.487 | 8062870.186 | 7175144.814 | 5658787.318 | 4233489.177 |
| Thiamine | 3934164.53 | 3279816.639 | 4058815.556 | 3701221.326 | 3684428.26 | 4073420.292 | 3741439.118 | 3944361.136 | 3642509.726 | 4064285.511 | 3836968.396 | 3168115.415 |
| PC 19:1_19:1 | 6189137.838 | 2812149.193 | 5034928.072 | 2438457.973 | 4469619.845 | 311840.2394 | 3545866.698 | 2033795.275 | 1851588.267 | 1529690.426 | 1618246.252 | 1081521.423 |
| M516T443 | 641443.0236 | 524057.4635 | 479537.0638 | 576808.5001 | 498133.4978 | 447217.0009 | 493714.9371 | 507283.9272 | 489534.0914 | 558801.2561 | 456629.5075 | 496149.0076 |

| Metabolites | PBS_1 | PBS_2 | PBS_3 | PBS_4 | BCG_1 | BCG_2 | BCG_3 | BCG_4 | BCGΔ*pe18*_1 | BCGΔ*pe18*_2 | BCGΔ*pe18*_3 | BCGΔ*pe18*_4 |
|---|---|---|---|---|---|---|---|---|---|---|---|---|
| (3.beta.,5.Xi.,9.Xi.)-3-((3-O-Acetyl-.beta.-D-xylopyranosyl-(1->4)-6-deoxy-.alpha.-L-mannopyranosyl-(1->2)-.beta.-D-xylopyranosyl-(1->2)-.beta.-D-glucopyranosyl)oxy)-23-hydroxyolean-12-en-28-oic acid | 170233.7853 | 216050.3542 | 116824.6657 | 170352.5967 | 149911.6427 | 114017.7181 | 119291.6772 | 117827.9203 | 143963.3271 | 165597.1322 | 132689.8141 | 123883.5853 |
| PI 16:0_18:1;3O | 335541.1569 | 345994.8484 | 310823.3943 | 134805.6379 | 394125.7809 | 326010.9916 | 366373.7257 | 458432.2626 | 649820.3369 | 431292.8848 | 261818.3391 | 303600.0053 |
| PI O-22:4_24:5 | 47008.71466 | 41922.3366 | 78044.35907 | 13623.41785 | 414957.5414 | 328879.733 | 428611.8306 | 308162.4066 | 350160.3453 | 321465.7592 | 516808.6846 | 299364.8468 |
| HBMP 16:4_18:5_20:5 | 0 | 194067.4284 | 0 | 225212.9738 | 193381.9017 | 0 | 98588.80701 | 180910.858 | 8516.06114 | 205545.0938 | 14270.12817 | 0 |
| LPC(17:0/0:0) | 3018967.049 | 2495046.614 | 4627100.798 | 2979117.148 | 6979320.306 | 4909120.462 | 6542572.12 | 6107682.04 | 5467448.233 | 4550995.618 | 5170188.093 | 3556564.418 |
| Cer 8:0;2O/18:4 | 991029.7014 | 877418.4847 | 422332.5251 | 335983.5481 | 77453.12128 | 731260.0733 | 803962.6245 | 885531.7388 | 402191.4373 | 773831.724 | 573661.8377 | 743917.0888 |
| Neohancoside D | 271540.2275 | 276301.5325 | 246883.7195 | 250287.8815 | 290666.7661 | 240563.3007 | 231813.2576 | 268348.4755 | 278463.8035 | 269282.7997 | 283302.0556 | 249545.1563 |
| FAHFA 18:0/20:4 | 639978. | 507443. | 827025. | 101633 | 197026 | 307359. | 109950 | 277645 | 146066 | 174128 | 784923. | 116369 |

| **Metabolites** | **PBS_1** | **PBS_2** | **PBS_3** | **PBS_4** | **BCG_1** | **BCG_2** | **BCG_3** | **BCG_4** | **BCGΔ*pe18*_1** | **BCGΔ*pe18*_2** | **BCGΔ*pe18*_3** | **BCGΔ*pe18*_4** |
|---|---|---|---|---|---|---|---|---|---|---|---|---|
| | 7568 | 2387 | 461 | 1.44 | 9.996 | 2011 | 5.75 | 2.79 | 1.614 | 7.84 | 9783 | 2.57 |
| PE O-16:4_2:0 | 42193.58578 | 37135.13928 | 27267.74579 | 50397.17886 | 33560.89947 | 36301.70513 | 42486.12662 | 42575.85442 | 44027.9503 | 40485.72438 | 39538.01976 | 12300.83279 |
| Santinamide cation | 921316.6497 | 636356.9731 | 1858508.289 | 672999.2467 | 1832873.757 | 1040957.46 | 522846.0621 | 3573685.081 | 785512.2994 | 987262.9911 | 733199.3751 | 1073275.203 |
| PI 32:5 | 25002.70957 | 6898.202729 | 82405.79283 | 44299.78863 | 44485.73947 | 50209.94618 | 102710.1199 | 48101.17825 | 36686.0838 | 46413.25804 | 83960.68297 | 296718.3024 |
| PE 25:3 | 515163.5687 | 260365.0557 | 971160.2237 | 527753.7422 | 2959983.396 | 1863338.221 | 3110209.571 | 2182411.957 | 1351380.786 | 1748600.265 | 2581528.858 | 1324877.276 |
| SM 42:2;3O | 7609.889269 | 240177.5363 | 173799.0761 | 269125.6521 | 146316.768 | 221431.1573 | 100943.6067 | 166845.6222 | 175413.9614 | 166029.5752 | 168645.759 | 131991.5087 |
| Homoserine | 352588.3947 | 289158.942 | 442691.4399 | 204730.7959 | 1012157.589 | 773382.2738 | 1007640.078 | 819534.8507 | 849828.9532 | 827618.3119 | 696261.721 | 727059.3177 |
| PE O-18:1_20:4 | 4619381.08 | 1253505.003 | 3249646.186 | 2211185.234 | 2788275.333 | 0 | 3961803.155 | 5032713.563 | 2636154.452 | 747072.6702 | 2121054.034 | 1387560.765 |
| 3-Hydroxy-5-((3-hydroxy-6-methyl-5-(pentofuranosyloxy)octanoyl)oxy)-6-methyloctanoic acid | 633225.3804 | 813904.0331 | 550480.4628 | 721731.3304 | 2052259.938 | 1599127.119 | 1061434.301 | 1391549.624 | 1438593.015 | 924798.9343 | 927043.8817 | 1550110.518 |
| Salicyluric acid | 487289.9665 | 453909.1212 | 880772.0318 | 606389.2853 | 592224.4558 | 688590.1053 | 615861.0739 | 607510.5637 | 636516.8709 | 317264.3934 | 370793.8413 | 600586.8772 |

| Metabolites | PBS_1 | PBS_2 | PBS_3 | PBS_4 | BCG_1 | BCG_2 | BCG_3 | BCG_4 | BCGΔ*pe18*_1 | BCGΔ*pe18*_2 | BCGΔ*pe18*_3 | BCGΔ*pe18*_4 |
|---|---|---|---|---|---|---|---|---|---|---|---|---|
| Methyl 2-chloro-6-(trifluoromethyl)pyridine-3-carboxylate | 9105648.701 | 4365670.259 | 9681498.704 | 7520274.974 | 4205647.406 | 6789636.658 | 6443019.223 | 11212816.82 | 4479581.625 | 4872966.901 | 10774464.91 | 5725604.721 |
| 1-(10Z-Heptadecenoyl)-sn-glycero-3-phospho-(1'-rac-glycerol) | 217758.3798 | 322303.8751 | 142992.4645 | 247370.6367 | 726499.6053 | 578036.4256 | 856583.5914 | 607190.4828 | 1088621.779 | 1075047.854 | 1164270.594 | 676415.1346 |
| SHexCer 14:3;2O/26:1;O | 1053302.552 | 997520.9838 | 977709.7396 | 1076182.182 | 976642.4841 | 1071098.538 | 900909.2872 | 880953.8404 | 839843.1906 | 863110.0982 | 1057469.125 | 1018634.622 |
| Arecaidine | 1487765.252 | 1330927.148 | 1282265.47 | 1420476.023 | 2582039.387 | 2166266.339 | 2549384.287 | 2120031.446 | 1797421.561 | 1905655.82 | 2309653.938 | 1614510.864 |
| Diethylthiophosphate | 963662.4038 | 921347.9476 | 913476.3802 | 697582.1425 | 1386369.53 | 1940718.486 | 3256439.34 | 3056799.194 | 1807485.557 | 1986538.048 | 1690822.804 | 1133479.191 |
| (Cyanomethyl)(triphenyl)phosphonium cation | 494133.0793 | 542381.1164 | 347162.3559 | 781460.4496 | 12974023.59 | 11904338.26 | 5600070.565 | 6806192.783 | 19851271.35 | 14303357.47 | 15028139.99 | 8401555.964 |
| N-(3-Methylbutanoyl)valine | 156207.2618 | 140318.8666 | 125622.8425 | 2141805.621 | 24661.13605 | 160020.3049 | 1495494.078 | 26534.84482 | 606609.4632 | 299199.65 | 41925.48915 | 1213836.315 |
| Flavin adenine dinucleotide (FAD) | 34281.87921 | 29419.39556 | 44435.47886 | 27607.91271 | 276569.4186 | 147916.6056 | 251628.8584 | 191021.645 | 200845.4506 | 211984.5333 | 221567.1803 | 256629.5497 |
| PS 16:0_20:5;2O | 80179.61778 | 79309.04954 | 169702.0911 | 91863.21756 | 104671.84 | 65958.3454 | 96412.74486 | 85211.12473 | 30518.57604 | 47289.45388 | 57882.57087 | 29526.80097 |

| **Metabolites** | **PBS_1** | **PBS_2** | **PBS_3** | **PBS_4** | **BCG_1** | **BCG_2** | **BCG_3** | **BCG_4** | **BCGΔ*pe18*_1** | **BCGΔ*pe18*_2** | **BCGΔ*pe18*_3** | **BCGΔ*pe18*_4** |
|---|---|---|---|---|---|---|---|---|---|---|---|---|
| Ethyl 4-amino-1-piperidinecarboxylate | 2723538.846 | 2154316.844 | 1076522.558 | 1624662.865 | 2320198.313 | 2158438.817 | 1759946.714 | 2541909.041 | 2348341.635 | 2778072.842 | 2527778.989 | 2427239.599 |
| cis-resveratrol 2-C-glucoside | 50730.27002 | 16624.3843 | 72327.70782 | 12941.87326 | 1051410.738 | 742878.3375 | 1278863.402 | 661014.4077 | 698949.7192 | 1141705.324 | 768142.7443 | 341049.7889 |
| MGDG O-13:1_3:0 | 65253.81971 | 92927.11563 | 144103.1708 | 89188.86436 | 653914.8835 | 442674.6819 | 674623.628 | 602411.4326 | 451424.9808 | 543820.2395 | 621912.528 | 410553.5676 |
| 3,3,3-Trifluorolactic acid | 2210178.109 | 1198199.252 | 1436727.748 | 1232039.35 | 1664118.733 | 1018715.33 | 863677.236 | 1115987.533 | 853443.4058 | 912398.9093 | 1797173.187 | 791076.2605 |
| Tadalafil | 525552.0196 | 423862.21 | 261628.8139 | 324414.7428 | 589294.7219 | 486769.808 | 389685.5468 | 415121.5953 | 484239.2026 | 692873.748 | 443934.209 | 455124.4845 |
| 25-Azacholesterol | 6228626.811 | 102518563.8 | 4361083.952 | 5550631.522 | 28863445.16 | 27652033.16 | 24018721.14 | 4859005.608 | 5292970.972 | 5242207.193 | 4823209.343 | 4183548.371 |
| 2-Aminobutyric acid | 24181447.3 | 23762078.33 | 23478636.85 | 23205157.2 | 25391826.02 | 23718245.62 | 26065254.79 | 23992551.83 | 25097055.24 | 22818349.53 | 25985513.64 | 25113994.93 |
| Isoguvacine | 989401.8169 | 1958182.373 | 1989266.683 | 1217832.53 | 2934890.611 | 1220950.115 | 815610.2898 | 342872.1173 | 762465.0892 | 1794477.403 | 2560547.66 | 407329.6041 |
| 2H-Pyran-2-one, tetrahydro-4-hydroxy-6-pentyl- | 909141.7124 | 911370.443 | 757317.9722 | 760631.814 | 884809.4542 | 726787.9092 | 967991.4202 | 889512.3602 | 936293.1774 | 916712.908 | 918146.717 | 938974.733 |
| Xanthyletin | 306610.0007 | 151605.2719 | 295545.7882 | 168780.145 | 260226.3902 | 161335.0825 | 964895.0361 | 921168.6816 | 498287.5888 | 255606.525 | 435408.7643 | 181464.4359 |

| Metabolites | PBS_1 | PBS_2 | PBS_3 | PBS_4 | BCG_1 | BCG_2 | BCG_3 | BCG_4 | BCGΔ*pe18*_1 | BCGΔ*pe18*_2 | BCGΔ*pe18*_3 | BCGΔ*pe18*_4 |
|---|---|---|---|---|---|---|---|---|---|---|---|---|
| .alpha.-Hydroxyfarnesylphosphonic acid | 186089.777 | 0 | 138747.1267 | 61299.93256 | 127966.2706 | 189227.2162 | 1325532.187 | 740427.7306 | 383363.3493 | 183041.734 | 290499.178 | 75565.46791 |
| 3-Amino-4-oxobutanoic acid | 1918582.237 | 14232735.78 | 2465369.226 | 4672766.839 | 4462988.454 | 2555080.733 | 3985861.096 | 2643895.984 | 2453641.614 | 1935440.869 | 2413433.366 | 1806286.529 |
| FA 24:6 | 1648280.544 | 1237887.696 | 1612417.84 | 1633334.132 | 1416102.958 | 1640262.716 | 1255115.322 | 1314516.975 | 1317512.43 | 1006214.978 | 1091241.15 | 1143999.019 |
| LPC O-18:0 | 1139380.059 | 1565421.384 | 660253.1951 | 933653.6443 | 680399.5663 | 744271.6795 | 855171.987 | 831186.9151 | 569597.7983 | 603941.0362 | 617344.8862 | 331637.135 |
| DG 27:4 | 298536.8217 | 416915.6337 | 190377.3081 | 257998.6872 | 294917.6661 | 727658.6369 | 270294.4348 | 275864.566 | 246832.8557 | 215833.0624 | 329582.8471 | 225680.4726 |
| DGDG O-14:1_2:0 | 97872.63802 | 166776.1631 | 71696.20671 | 176164.7398 | 75425.4916 | 36394.11582 | 54455.2454 | 38134.77266 | 72013.92004 | 51265.28188 | 84046.14284 | 66591.61131 |
| Pemetrexed | 74365.59136 | 56801.14332 | 77724.03474 | 71002.38657 | 204239.3993 | 67606.69939 | 147977.1153 | 158053.343 | 125477.3973 | 153956.3627 | 101893.0481 | 78172.76353 |
| Pilocarpine | 214408.6223 | 208412.1444 | 251146.1342 | 241758.808 | 226762.362 | 230672.2062 | 216616.23 | 217044.7938 | 212430.8033 | 220421.381 | 206715.2823 | 200882.051 |
| 3-(Dimethylamino)-4-methylbenzene-1-sulfonamide | 1062287.94 | 1139159.828 | 968394.381 | 1148793.936 | 1034292.272 | 1143302.875 | 1032724.111 | 1080341.011 | 1119972.512 | 1114778.046 | 1130805.52 | 1020556.097 |
| 3'-O-Desmethyletoposide | 1098916.875 | 1037906.445 | 944153.218 | 831331.2774 | 1244623.913 | 782685.4607 | 882167.3547 | 821291.4099 | 919177.2907 | 829579.8685 | 932325.8949 | 775259.7935 |
| PC 18:0_18:4 | 1282419.484 | 2026982.411 | 981449.418 | 1007924.1 | 912239.9061 | 1081093.986 | 1003278.852 | 1464653.704 | 584177.9991 | 1164492.437 | 1041856.432 | 850998.0537 |

| **Metabolites** | **PBS_1** | **PBS_2** | **PBS_3** | **PBS_4** | **BCG_1** | **BCG_2** | **BCG_3** | **BCG_4** | **BCGΔ*pe18*_1** | **BCGΔ*pe18*_2** | **BCGΔ*pe18*_3** | **BCGΔ*pe18*_4** |
|---|---|---|---|---|---|---|---|---|---|---|---|---|
| Nonadecanoylcarnitine (Car(19:0)) | 94305.8754 | 148675.8023 | 40139.18696 | 86698.58994 | 278452.5182 | 180683.7853 | 842541.713 | 961523.147 | 682496.8353 | 645887.017 | 574992.7132 | 412357.7067 |
| 2-(2,5-Dichlorophenyl)-1,3-benzoxazol-5-amine | 1513792.168 | 2399016.109 | 956175.5128 | 798823.6729 | 2736411.382 | 600463.6651 | 1888125.75 | 1321460.559 | 1238149.819 | 2854674.31 | 814543.1055 | 272402.9439 |
| DGDG O-16:4_6:0 | 49431.13632 | 48006.12522 | 19574.79009 | 43773.52666 | 33438.08996 | 27473.71803 | 34911.88688 | 31009.82818 | 37826.12407 | 53073.56684 | 59997.6947 | 52908.56869 |
| PI 24:2 | 57059.99256 | 87225.24578 | 79117.41741 | 61858.31936 | 83040.41836 | 55184.76805 | 62042.37262 | 76020.82351 | 71491.7135 | 71645.01987 | 50780.22554 | 48384.98329 |
| 5-Bromo-1H-1,2,3-benzotriazole | 569584.3916 | 3300494.777 | 10174009.84 | 5556657.605 | 1423259.576 | 1154251.077 | 2325037.377 | 1281617.03 | 1730710.409 | 3006975.664 | 1848201.615 | 14149634.23 |
| (1-Hydroxycyclohexyl)acetic acid | 4382047.519 | 3940598.124 | 4015721.471 | 4102776.219 | 3637844.293 | 3981484.981 | 3671445.032 | 3815892.862 | 910788.2136 | 1068125.49 | 3801017.116 | 4385620.7 |
| Piperazine | 376905.3098 | 7763852.869 | 14774321.32 | 6997972.24 | 801943.2595 | 470864.1483 | 1922736.925 | 971465.8037 | 1677076.208 | 1987796.342 | 791023.7957 | 15935767.32 |
| 2-Methoxy-3-methyl-9H-carbazole | 34940.62062 | 119578.1678 | 732240.832 | 532208.6954 | 1458494.164 | 37416.20767 | 14025.21556 | 0 | 28146.64699 | 60192.12281 | 84173.74741 | 1205166.307 |
| SSR161421 | 0 | 0 | 8382.457396 | 0 | 54272.08639 | 69727.65221 | 524191.7807 | 47086.62186 | 363525.846 | 146902.7472 | 50161.91781 | 515056.0131 |
| Methyl_jasmonate | 273363.0733 | 234012.2975 | 253154.3867 | 260539.014 | 240855.7035 | 207914.1863 | 283966.6371 | 210087.1627 | 196008.6507 | 229109.8484 | 252250.5384 | 281981.327 |
| N-Desmethyltramadol | 964254.7387 | 886088.4376 | 788808.3184 | 822222.2333 | 867538.7414 | 834683.3431 | 901957.263 | 862173.9981 | 855852.2778 | 857419.3949 | 897749.1242 | 678719.4174 |

| Metabolites | PBS_1 | PBS_2 | PBS_3 | PBS_4 | BCG_1 | BCG_2 | BCG_3 | BCG_4 | BCGΔ *pe18*_1 | BCGΔ *pe18*_2 | BCGΔ *pe18*_3 | BCGΔ*pe18*_4 |
|---|---|---|---|---|---|---|---|---|---|---|---|---|
| .beta.-Alanine, N-[2-(2-pyridinyl)-6-(1,2,4,5-tetrahydro-3H-3-benzazepin-3-yl)-4-pyrimidinyl]- | 1054703.561 | 1132843.077 | 1080792.615 | 1060994.726 | 1074376.173 | 1093037.67 | 1055008.963 | 1065333.36 | 1091676.499 | 1072951.788 | 1072167.571 | 1103573.521 |
| N-acetyl-beta-glucosamine-1,6-anhydro-N-acetyl-beta-muramate | 0 | 0 | 0 | 0 | 369302.8479 | 249178.9274 | 229365.5007 | 189339.9596 | 382022.1129 | 378241.8144 | 406573.1788 | 196829.3168 |
| beta-Guanidinopropionic acid | 165264.1716 | 0 | 353100.1354 | 0 | 1930581.77 | 1066908.158 | 2802976.772 | 1775134.719 | 924365.1422 | 1361608.367 | 1948925.756 | 1075627.386 |
| SM 17:3;2O | 114871.0165 | 137308.807 | 117389.0162 | 126709.8716 | 119499.2644 | 140254.1428 | 156390.7662 | 137139.6738 | 126256.2647 | 145053.1366 | 118436.8826 | 121579.5535 |
| FAHFA 10:0/26:4 | 0 | 10680.90633 | 41051.90413 | 8141.458222 | 72765.67977 | 132342.8814 | 40691.67786 | 97230.85748 | 27213.16177 | 30108.2733 | 183399.5837 | 35704.45702 |
| 4-[4-(4-Chlorophenyl)-1,3-thiazol-2-yl]morpholine | 26880.13293 | 17441.17766 | 0 | 16113.1539 | 12663.68983 | 98248.34349 | 25745.98073 | 11155.16579 | 11035.25162 | 0 | 48482.22718 | 20007.94676 |
| LPE(18:1(9Z)/0:0) | 983253.1053 | 787531.6097 | 1327082.155 | 872105.2798 | 3963493.183 | 2788876.818 | 3794018.504 | 3120712.693 | 4009953.076 | 4228003.71 | 4499979.164 | 3111866.216 |
| Nornidulin | 295464.9329 | 227839.124 | 141345.2099 | 297279.1895 | 860420.3595 | 631598.3958 | 856384.7408 | 544211.7601 | 460597.6508 | 246810.9281 | 294924.5544 | 409527.9599 |
| DLCL 16:0_22:5 | 0 | 0 | 0 | 0 | 119094. | 188099. | 77891.2 | 134626. | 189982. | 81776.1 | 94595.5 | 137020. |

| Metabolites | PBS_1 | PBS_2 | PBS_3 | PBS_4 | BCG_1 | BCG_2 | BCG_3 | BCG_4 | BCGΔ*pe18*_1 | BCGΔ*pe18*_2 | BCGΔ*pe18*_3 | BCGΔ*pe18*_4 |
|---|---|---|---|---|---|---|---|---|---|---|---|---|
| | | | | | 1589 | 0064 | 4479 | 7918 | 188 | 6676 | 6275 | 083 |
| PC(15:0/18:0) | 4915882.161 | 5461219.119 | 11920301.29 | 5679463.88 | 9641839.853 | 8383599.033 | 10763444.39 | 9638818.777 | 20101648.34 | 11630537.85 | 8359544.807 | 18171194.49 |
| N-Acetylneuraminic acid | 1870199.234 | 1632308.36 | 1910746.304 | 1557371.503 | 1421934.62 | 943702.825 | 2524222.082 | 1427413.546 | 744180.203 | 1227448.765 | 1371338.519 | 713475.2629 |
| FA 19:2 | 392800.7906 | 1115633.099 | 1014121.22 | 447224.6471 | 1157376.757 | 1036534.308 | 847185.5982 | 1241212.275 | 1016019.815 | 655150.8705 | 871107.0408 | 1001311.905 |
| DGDG O-16:4_26:5 | 0 | 0 | 0 | 0 | 538214.9273 | 459142.1521 | 275479.687 | 570845.1583 | 74326.27745 | 52452.8636 | 95024.27236 | 0 |
| Diisodecyl phthalate | 582396.0825 | 613177.101 | 332635.2562 | 799959.6273 | 337240.1695 | 832364.6114 | 250644.8167 | 433281.667 | 282365.4559 | 325000.7516 | 350261.955 | 996866.3653 |
| N-Methylpelletierine | 2161275.541 | 1731862.402 | 2047453.085 | 2144571.274 | 2239981.467 | 2829051.359 | 2041723.932 | 2409895.777 | 2271194.889 | 1956617.899 | 2252442.795 | 1847916.491 |
| PG 5:0_18:5 | 735617.5025 | 388010.5814 | 258369.0808 | 375804.488 | 287099.0441 | 752297.0654 | 699563.9408 | 330167.4585 | 355819.1249 | 321325.7706 | 375521.6901 | 283467.7655 |
| 5-[(Z)-14-(3,5-Dihydroxyphenyl)tetradec-8-enyl]benzene-1,3-diol | 335988.9735 | 321298.777 | 399554.3396 | 375466.8247 | 221125.0543 | 138446.5199 | 515553.0943 | 308630.0516 | 206661.1957 | 275298.4914 | 207274.5443 | 242596.1004 |
| Jervine | 64971.83231 | 67354.99361 | 71406.16025 | 81404.25394 | 71618.85684 | 80873.90848 | 77802.47107 | 72401.69183 | 88273.20153 | 67525.07421 | 62708.32004 | 79972.51488 |
| Luffariellolide | 863229.3048 | 1349405.677 | 863272.103 | 820413.8181 | 2422270.04 | 2519776.303 | 1574822.338 | 1287885.774 | 745951.1221 | 632691.5365 | 1210983.112 | 787628.0608 |

| **Metabolites** | **PBS_1** | **PBS_2** | **PBS_3** | **PBS_4** | **BCG_1** | **BCG_2** | **BCG_3** | **BCG_4** | **BCGΔ*pe18*_1** | **BCGΔ*pe18*_2** | **BCGΔ*pe18*_3** | **BCGΔ*pe18*_4** |
|---|---|---|---|---|---|---|---|---|---|---|---|---|
| N-Isobutyl-3-methylbutanamide | 8911073.369 | 9192987.42 | 9095748.254 | 9440447.093 | 8924989.548 | 9178050.218 | 8584266.696 | 9131661.224 | 8933305.118 | 9237165.022 | 9436238.655 | 8831755.464 |
| Lithocholic acid 3-sulfate | 25927.67533 | 100503.412 | 111934.8338 | 87877.06325 | 828944.5358 | 641414.1963 | 793195.4416 | 593536.8508 | 1031672.785 | 742241.8882 | 861285.936 | 521552.4992 |
| MG 21:1 | 4190898.78 | 6181808.229 | 3452299.35 | 4212886.583 | 5864866.261 | 4301063.34 | 4105017.293 | 4047889.912 | 3767785.24 | 4494652.98 | 5509356.154 | 3633079.332 |
| PG(20:4(5Z,8Z,11Z,14Z)/20:4(5Z,8Z,11Z,14Z)) | 6588590.025 | 7409279.848 | 5053675.479 | 6477797.367 | 1035552.025 | 3059451.695 | 4743976.825 | 3915939.361 | 730968.181 | 5832572.438 | 4962356.092 | 3610556.777 |
| MGDG O-15:1_22:5 | 246502.2834 | 74564.66891 | 228012.5328 | 153208.6685 | 249953.2455 | 1541048.032 | 791922.7482 | 801221.1849 | 459455.7469 | 415075.2405 | 1407458.561 | 1487569.396 |
| PC O-20:5_20:4 | 752249.7491 | 362294.3863 | 632606.7148 | 651338.0741 | 536219.541 | 450093.9988 | 967293.3861 | 795570.5258 | 802612.5561 | 788903.1593 | 547945.9534 | 611195.4676 |
| Acetoin | 3646083.856 | 7903518.447 | 3592037.737 | 5005113.31 | 4606548.68 | 3504683.486 | 3706767.204 | 3509451.385 | 5759754.268 | 4062734.629 | 4037440.157 | 4621468.121 |
| 4-Aminoantipyrine | 4044130.816 | 14724188.93 | 3697711.181 | 15061797.5 | 13190362.96 | 14231249.27 | 12336536.78 | 13856116.88 | 11699732.97 | 3357977.457 | 13944863.85 | 15560138.55 |
| (4-(.beta.-D-Glucopyranosyloxy)-2,6-bis(3-methylbut-2-en-1-yl)phenyl)(hydroxy)acetic acid | 437015.5424 | 633798.1799 | 512273.6252 | 341948.0565 | 541997.176 | 534246.9575 | 464938.4028 | 555514.3157 | 465940.3844 | 426692.3824 | 508613.5911 | 509042.6293 |
| PI 18:0_22:4 | 2043548.957 | 2928488.359 | 2589969.316 | 1718079.597 | 1980240.925 | 35781.48282 | 1838977.486 | 1081106.033 | 3155087.629 | 5264127.39 | 1592376.574 | 1316686.508 |

| Metabolites | PBS_1 | PBS_2 | PBS_3 | PBS_4 | BCG_1 | BCG_2 | BCG_3 | BCG_4 | BCGΔ *pe18*_1 | BCGΔ *pe18*_2 | BCGΔ *pe18*_3 | BCGΔ*pe18*_4 |
|---|---|---|---|---|---|---|---|---|---|---|---|---|
| 2-[(2,6-Dimethylphenyl)amino]-N,N,N-triethyl-2-oxoethanaminium cation | 613322.9095 | 786676.825 | 487042.3351 | 578922.925 | 745138.9373 | 564795.9969 | 674713.4198 | 716866.8379 | 833203.1791 | 753361.8746 | 695801.052 | 719451.9112 |
| (3.beta.,9.xi.)-3-((6-Deoxyhexopyranosyl-(1->2)hexopyranosyl-(1->3)hexopyranosyl)oxy)-14-hydroxybufa-4,20,22-trienolide | 395258.4109 | 516444.9568 | 207368.386 | 415773.7859 | 374786.6577 | 362002.3712 | 372906.3109 | 337460.4672 | 391972.178 | 359124.9608 | 373068.2912 | 281716.1636 |
| 5-(4-Pyridinyl)-5-(2-butenyl)-2,7-nonadiene | 2303621.104 | 2384839.642 | 2169558.032 | 2236278.688 | 2393216.934 | 2225933.719 | 2203615.646 | 2031471.9 | 2198898.254 | 2226681.467 | 2076489.091 | 2169286.603 |
| octadeca-10,12-dienoylcarnitine | 970683.457 | 976891.1912 | 936153.8932 | 932568.4618 | 873220.7334 | 907561.153 | 1201567.103 | 929958.5751 | 917349.3651 | 1129246.872 | 1172074.7 | 811838.53 |
| Cer 16:1;3O/30:2 | 118156.0452 | 147795.7168 | 145371.1308 | 145190.8229 | 129149.5375 | 158483.7054 | 118110.9552 | 155975.1522 | 132369.9716 | 150098.4171 | 138116.6016 | 107206.7039 |
| Maprotiline (hydrochloride) | 1764213.569 | 1084052.008 | 1002516.663 | 1155403.319 | 1023974.803 | 1230686.232 | 1139924.052 | 1167371.44 | 1164646.721 | 842670.072 | 989499.0154 | 982314.4724 |
| Sinensetin | 1767184.878 | 1551406.874 | 1895919.133 | 1802786.87 | 1689930.081 | 1648761.619 | 1472103.115 | 1622029.353 | 1819097.41 | 1719018.862 | 1767071.079 | 1614712.045 |
| Halopemide | 160086.7477 | 167184.1199 | 150512.0292 | 162640.6357 | 164463.6621 | 164367.1558 | 172754.3895 | 150927.1214 | 164387.9646 | 162522.7197 | 160446.3516 | 162837.054 |

| Metabolites | PBS_1 | PBS_2 | PBS_3 | PBS_4 | BCG_1 | BCG_2 | BCG_3 | BCG_4 | BCGΔ*pe18*_1 | BCGΔ*pe18*_2 | BCGΔ*pe18*_3 | BCGΔ*pe18*_4 |
|---|---|---|---|---|---|---|---|---|---|---|---|---|
| Kaempferol-3-O-glucoside-6"-p-coumaroyl | 237615.4133 | 175252.7268 | 190722.1331 | 200224.5769 | 186661.88 | 223887.5825 | 148365.8391 | 140684.9341 | 170289.5337 | 186108.1267 | 196751.894 | 154469.5005 |
| 1H-Tetraazol-5-ylacetonitrile | 59523.13676 | 1393486.889 | 949648.1258 | 1531725.503 | 62942.40071 | 53225.00318 | 112637.8063 | 0 | 78873.32844 | 135562.0482 | 126287.5078 | 852687.0452 |
| PE-Cer 12:0;2O/22:1 | 198809.9556 | 390888.6519 | 110594.1087 | 433425.4176 | 218385.5464 | 48230.24669 | 198978.3148 | 396711.8599 | 476677.2333 | 232817.4739 | 67945.25093 | 791989.5584 |
| PC(35:3) | 4945992.178 | 15971448.94 | 19817823.75 | 15783734.63 | 13741291.24 | 6797194.045 | 6757197.366 | 6279025.351 | 10288195.44 | 4069637.827 | 8158590.374 | 3620403.14 |
| Dexamethasone phosphate | 495339.6113 | 436810.6363 | 433173.3396 | 410299.2507 | 495790.2215 | 424670.6574 | 455228.7004 | 424785.4565 | 463573.9632 | 442158.1848 | 386480.9821 | 422191.3568 |
| Aureonitol | 313938.0235 | 346296.8131 | 400141.2772 | 391477.4896 | 346101.7984 | 350572.2422 | 357506.6388 | 307563.7525 | 367678.0545 | 364600.0199 | 1462985.536 | 581669.1302 |
| 4-Quinazolinamine, 2-(hexahydro-4-methyl-1H-1,4-diazepin-1-yl)-6,7-dimethoxy-N-[1-(phenylmethyl)-4-piperidinyl]- | 714388.0327 | 744284.1711 | 729137.9077 | 674726.4211 | 625413.7436 | 763115.6231 | 760665.9346 | 760967.2464 | 761517.2097 | 680193.7298 | 775225.6793 | 717531.4287 |

| Metabolites | PBS_1 | PBS_2 | PBS_3 | PBS_4 | BCG_1 | BCG_2 | BCG_3 | BCG_4 | BCGΔ*pe18*_1 | BCGΔ*pe18*_2 | BCGΔ*pe18*_3 | BCGΔ*pe18*_4 |
|---|---|---|---|---|---|---|---|---|---|---|---|---|
| (8R,9a'S,10a'S)-1',1',4,4,12'-Pentamethyl-6',7',8',9',10',10a'-hexahydro-1'H,4H,11'H-spiro[1,4-dioxepino[2,3-g]indole-8,2'-[3a,9a](epiminomethano)cyclopenta[b]quinolizine]-9,11'(10H)-dione | 224386.979 | 238696.2167 | 166481.7983 | 232757.4531 | 237537.822 | 228977.7442 | 221727.3386 | 219036.2661 | 227777.7608 | 228390.5528 | 217963.4982 | 216977.5962 |
| 3-Hydroxyarachidonoylcarnitine | 421664.628 | 395817.0613 | 409686.4086 | 382058.1002 | 395574.3291 | 160704.7226 | 196153.593 | 348483.4497 | 589650.9874 | 425835.7542 | 408342.0933 | 307378.4981 |
| [2-(4-Bromophenyl)-5-oxo-1-cyclopenten-1-yl]acetic acid | 1631175.784 | 1434084.766 | 1793853.918 | 1562297.253 | 1630689.416 | 1334757.062 | 1862802.918 | 1814855.852 | 1551717.798 | 1686901.194 | 1783704.682 | 1564722.98 |
| MG 15:1 | 883955.4707 | 889311.4876 | 870712.0163 | 857592.2339 | 886508.6672 | 885858.4973 | 875227.0958 | 869122.1673 | 896931.2216 | 872711.2389 | 865018.5578 | 882416.5559 |
| TG O-8:0_20:5_22:6 | 718926.3397 | 1359467.416 | 811409.5925 | 1209455.515 | 738404.8051 | 1203374.408 | 834514.3344 | 1096161.706 | 884248.4595 | 926971.7355 | 808754.921 | 1119730.86 |
| 3-(3,4-Dimethoxyphenyl)-1-(2-hydroxy-4,6-dimethoxyphenyl)propan-1-one | 645691.891 | 7382796.461 | 13521183.61 | 8143189.466 | 646365.0136 | 244521.2641 | 2594811.083 | 1210189.973 | 2132075.325 | 1505855.543 | 2300918.354 | 30458234.17 |

| Metabolites | PBS_1 | PBS_2 | PBS_3 | PBS_4 | BCG_1 | BCG_2 | BCG_3 | BCG_4 | BCGΔ*pe18*_1 | BCGΔ*pe18*_2 | BCGΔ*pe18*_3 | BCGΔ*pe18*_4 |
|---|---|---|---|---|---|---|---|---|---|---|---|---|
| Tiglylcarnitine (Car(5:1)) | 620751.4155 | 512109.9482 | 854779.6384 | 560016.5707 | 1361313.72 | 680197.5539 | 1067400.279 | 1020636.005 | 748078.4957 | 676799.3726 | 588346.0243 | 519373.6486 |
| S1P(d18:1) | 2606638.42 | 2563983.403 | 2842333.978 | 2599650.514 | 2331846.652 | 1883368.284 | 2726647.55 | 2215393.743 | 1432403.502 | 2001855.446 | 1939907.581 | 1519696.75 |
| Tetrahydropersin | 150098.3207 | 102178.2683 | 67734.21167 | 104248.9195 | 59327.88776 | 192384.1606 | 47114.96164 | 103414.7678 | 37351.09198 | 42437.27098 | 109717.5309 | 175946.1677 |
| Benzenethiol, 4,4'-thiobis- | 406076.7192 | 361674.4459 | 433093.8634 | 479178.901 | 543282.412 | 601804.8743 | 309191.5652 | 534315.5739 | 555510.642 | 160282.2166 | 193913.5456 | 533969.2818 |
| 2-Amino-4-bromo-5-fluorophenol | 423917.7963 | 285898.6568 | 389285.7655 | 172330.7071 | 390215.6475 | 429650.5772 | 245567.8949 | 196464.6481 | 420451.6753 | 270603.3485 | 151823.734 | 307958.8206 |
| 2'-O-Methylinosine | 1824510.017 | 1550000.131 | 3015458.776 | 2193639.599 | 21818446.13 | 12755697.06 | 19929345.93 | 15267571.9 | 17954209.92 | 17495190.41 | 20816321.64 | 15520543.8 |
| Testosterone_enanthate | 670121.7319 | 731050.0718 | 4370836.693 | 633149.8648 | 752625.3707 | 513735.1644 | 647006.4718 | 733824.6887 | 730386.3028 | 821520.6864 | 645390.328 | 526899.0782 |
| PC O-16:0_18:2 | 489572.7122 | 595233.0601 | 497392.5253 | 552956.7939 | 351841.877 | 53717.24082 | 346140.73 | 429279.281 | 280571.7775 | 366538.1348 | 371181.4001 | 282569.593 |
| 3,3'-Disulfanediylbis(2-(2,2,2-trifluoroacetamido)propanoic acid) | 135448.7382 | 73686.05492 | 59768.03304 | 17997.6209 | 0 | 16036.88686 | 402845.5075 | 441812.3959 | 52693.66658 | 103672.0448 | 48493.70902 | 132248.8662 |
| 12(S)-HHTrE | 3508651.658 | 3981859.404 | 6979073.729 | 6342820.124 | 5692505.59 | 8773613.016 | 17750092.21 | 8590157.653 | 8329341.897 | 2681357.075 | 3787656.756 | 8441157.929 |
| 5-methylhexanoate | 328621.2875 | 300942.6644 | 375357.9398 | 1524834.342 | 474677.4053 | 377202.3683 | 2080405.252 | 377800.1731 | 910603.6661 | 958066.3741 | 401562.2586 | 1672034.116 |

| Metabolites | PBS_1 | PBS_2 | PBS_3 | PBS_4 | BCG_1 | BCG_2 | BCG_3 | BCG_4 | BCGΔ*pe18*_1 | BCGΔ*pe18*_2 | BCGΔ*pe18*_3 | BCGΔ*pe18*_4 |
|---|---|---|---|---|---|---|---|---|---|---|---|---|
| SL 16:0;O/16:0;O | 262955.2302 | 202553.7971 | 187460.5233 | 114043.2063 | 147661.5392 | 117636.3473 | 206869.413 | 156236.8714 | 165629.0889 | 130953.7327 | 146052.3804 | 109537.932 |
| Decanoylcarnitine (Car(10:0)) | 64536.9057 | 153606.9706 | 367540.2457 | 380772.6607 | 992645.8287 | 614830.7637 | 1143008.66 | 899301.5622 | 456236.7822 | 626514.0378 | 709679.5915 | 346672.6136 |
| (2Z,6E,10Z)-12-Acetyloxy-10-(acetyloxymethyl)-6-methyl-2-(4-methylpent-3-enyl)dodeca-2,6,10-trienoic acid | 1371351.185 | 1160522.404 | 1106334.284 | 1245942.459 | 920379.084 | 1233260.882 | 983897.3431 | 1618532.321 | 974911.4368 | 413457.4448 | 611291.8253 | 926923.6045 |
| 1,2-Distearoyl-sn-glycero-3-phospho-L-serine | 13702260.26 | 2946185.785 | 0 | 0 | 13645115.56 | 4036287.343 | 7047192.336 | 5812021.455 | 10767945.31 | 2638857.378 | 6745282.342 | 5784836.026 |
| Oleoylserotonin | 194918.8866 | 228832.8191 | 381486.8914 | 232864.1181 | 168737.7064 | 517729.87 | 280290.0166 | 316972.6489 | 295911.4042 | 171859.0663 | 337834.7891 | 223764.0105 |
| 1-{[3-(Trifluoromethyl)phenyl]sulfonyl}-4-piperidinecarboxylic acid | 210705.3184 | 231005.9344 | 185654.4564 | 206102.0146 | 484011.6198 | 248887.5806 | 423227.2842 | 244619.8003 | 254350.1403 | 401336.3518 | 240032.9961 | 363075.391 |
| sn-glycero-3-phospho-N-(linoleoyl)ethanolamine | 1050346.997 | 1541795.958 | 483957.0415 | 220154.3267 | 118342.8128 | 182692.0247 | 118461.3859 | 206089.0662 | 222242.8408 | 304908.3628 | 286910.221 | 135186.7687 |
| LNAPS 2:0/N-20:3 | 179120.8492 | 192250.252 | 140084.2758 | 169812.0684 | 164384.8205 | 149097.3006 | 180704.9641 | 173140.2503 | 182240.5825 | 150238.4991 | 175476.8258 | 144542.7787 |

| Metabolites | PBS_1 | PBS_2 | PBS_3 | PBS_4 | BCG_1 | BCG_2 | BCG_3 | BCG_4 | BCGΔ *pe18*_1 | BCGΔ *pe18*_2 | BCGΔ *pe18*_3 | BCGΔ*pe18*_4 |
|---|---|---|---|---|---|---|---|---|---|---|---|---|
| Piperidine, 4-(5H-dibenzo[a,d]cyclohepten-5-ylidene)-1-[4-(2H-tetrazol-5-yl)butyl]- | 2145214.699 | 1876117.837 | 1890991.673 | 2512250.742 | 2096721.082 | 2407365.555 | 2354227.579 | 1839471.755 | 2108724.883 | 3354689.868 | 2744290.226 | 2221576.967 |
| 1,2-Distearoyl-sn-glycero-3-phospho-(1'-rac-glycerol) | 6404519.951 | 7985409.168 | 6718895.537 | 8620042.033 | 11327809.97 | 6887525.231 | 13657737.76 | 9766866.218 | 13470467.36 | 7788863.673 | 11681637.96 | 4040674.843 |
| 4-Acetamidobutyric acid | 287848.4065 | 545900.1443 | 309757.7223 | 332796.9878 | 305097.3976 | 322078.4711 | 330088.6763 | 336733.4898 | 304878.5547 | 212828.2674 | 164976.4613 | 425617.2761 |
| 1,2-Dilinoleoyl-sn-glycero-3-phosphoethanolamine | 9223811.689 | 11378189.73 | 10609165.57 | 7659699.778 | 20187899.53 | 19855209.21 | 23188659.22 | 2156132 5.4 | 17043414.56 | 21336287.94 | 24437964.84 | 14949050.19 |
| 4,4-Dimethyl-1,3-cyclohexanedione | 947203.2138 | 885577.4301 | 847211.4718 | 1326235.774 | 1014223.934 | 715246.6199 | 824282.9577 | 870687.7629 | 643806.7702 | 1413467.496 | 757064.9504 | 806903.5976 |
| (1.beta.,2.alpha.,9.Xi.,11.beta.,12.alpha.,15.beta.)-1,2,11,12,14,15-Hexahydroxypicras-4-en-16-one | 108519.5588 | 142423.6947 | 133198.8555 | 93089.16166 | 262998.7424 | 220299.9819 | 226435.6757 | 314408.1216 | 282576.3876 | 236110.3491 | 259858.9828 | 288865.3343 |
| PIP(16:2(9Z,12Z)/18:0) | 41155.16582 | 3429.774018 | 42480.44799 | 8400.944658 | 44595.85768 | 0 | 112286.0811 | 40846.62516 | 16201.61754 | 27496.4521 | 9442.483803 | 0 |

| Metabolites | PBS_1 | PBS_2 | PBS_3 | PBS_4 | BCG_1 | BCG_2 | BCG_3 | BCG_4 | BCGΔ *pe18*_1 | BCGΔ *pe18*_2 | BCGΔ *pe18*_3 | BCGΔ*pe18*_4 |
|---|---|---|---|---|---|---|---|---|---|---|---|---|
| N1-citryl-N3-decanoyl-N3-hydroxy-1,3-diaminopropane | 312456.7759 | 317647.5755 | 280668.7035 | 314025.4065 | 319835.1556 | 319536.9728 | 307376.9621 | 315448.1014 | 290724.7609 | 287924.4663 | 275151.694 | 274648.0794 |
| Mupirocin | 670903.5812 | 892107.5736 | 564634.6308 | 916085.6927 | 718633.9184 | 1591693.562 | 1274740.746 | 1348284.558 | 1134842.753 | 855486.737 | 700766.4051 | 1029149.011 |
| Lupinine | 2802976.692 | 3031042.231 | 2519492.492 | 2161462.075 | 2289276.518 | 2145286.627 | 1973282.085 | 2529668.594 | 2383472.126 | 2565365.227 | 2482296.804 | 1658289.563 |
| N-Nervonoyl Glutamic acid | 647858.2967 | 699438.7832 | 815817.1012 | 485565.2297 | 411357.1595 | 645872.7346 | 421936.513 | 397098.4743 | 463268.2535 | 406859.7011 | 904041.0051 | 452544.4836 |
| PE 18:1_20:5;3O | 622241.4938 | 642440.8341 | 558321.0359 | 635816.3906 | 690161.7252 | 664566.6754 | 780243.2094 | 652938.2782 | 631405.826 | 675336.9598 | 675687.3944 | 584012.6888 |
| Isocytosine | 4618650.032 | 5259390.173 | 7063381.321 | 5171259.724 | 17381080.06 | 6110267.319 | 10606434.29 | 9129081.176 | 7851999.094 | 6726358.912 | 13402389.27 | 6613320.946 |
| LNAPE 16:0/N-20:4 | 978952.6063 | 807437.31 | 1240908.747 | 918128.1371 | 1958911.595 | 2359638.104 | 2488843.259 | 2058052.125 | 1634952.96 | 1962921.786 | 2441242.535 | 1561157.621 |
| 2-Docosahexaenoyl-1-palmitoyl-sn-glycero-3-phosphoethanolamine | 640418.5014 | 1073484.512 | 1006724.358 | 621145.3042 | 1825262.265 | 2076035.994 | 1482279.366 | 1937297.949 | 1569795.012 | 1343781.865 | 1678325.6 | 1583942.001 |
| Tyr-Leu | 790777.9298 | 696336.6474 | 853855.0092 | 647395.8585 | 900074.0946 | 659470.3939 | 789944.2487 | 678551.1967 | 799714.6252 | 814354.4259 | 942210.2556 | 680597.2431 |
| Imidazoleacetic acid | 509766.6083 | 468518.4385 | 584093.2465 | 485157.4977 | 2354486.705 | 1735322.037 | 2412526.554 | 1862278.534 | 1521912.366 | 1580876.634 | 1667837.503 | 1256640.984 |
| N-Lauroyl | 239317. | 270095. | 385710. | 317134. | 301446. | 303515. | 395749. | 328287. | 261551. | 300984. | 251331. | 268634. |

| Metabolites | PBS_1 | PBS_2 | PBS_3 | PBS_4 | BCG_1 | BCG_2 | BCG_3 | BCG_4 | BCGΔ*pe18*_1 | BCGΔ*pe18*_2 | BCGΔ*pe18*_3 | BCGΔ*pe18*_4 |
|---|---|---|---|---|---|---|---|---|---|---|---|---|
| Phenylalanine | 4074 | 4365 | 9441 | 9885 | 6538 | 8618 | 7545 | 9862 | 2543 | 9274 | 7973 | 1952 |
| PI O-17:1_18:1 | 483122.1242 | 728049.9684 | 672965.5936 | 618993.3408 | 359504.041 | 402712.7641 | 287761.0998 | 178102.0631 | 383078.4065 | 313791.4933 | 418788.9593 | 664869.714 |
| N-[3-hydroxy-1-{[3.4.5-trihydroxy-6-(hydroxymethyl)oxan-2-yl]oxy}octadec-4-en-2-yl]tetracosanamide | 252016.3872 | 334010.2574 | 306737.4003 | 372821.9733 | 433704.786 | 548323.5921 | 464766.6797 | 433986.0971 | 398230.0209 | 449345.9779 | 351993.999 | 337006.8735 |
| 3-Aminophthalimide | 47193.90594 | 594224.1034 | 382670.9461 | 372355.1513 | 50420.38179 | 54408.90826 | 85096.58048 | 30220.46187 | 84682.29944 | 81409.7187 | 82405.96329 | 1057459.12 |
| 1-Aminoanthraquinone-2-carboxylic acid | 2407474.273 | 2521817.821 | 6579450.534 | 823673.7576 | 1147449.957 | 999274.8919 | 3874022.519 | 1774824.826 | 290227.2137 | 970112.3411 | 1316006.941 | 542164.3331 |
| PI 2:0_16:0 | 254712.657 | 174520.075 | 161713.4519 | 182140.3407 | 177721.4214 | 153677.1326 | 209571.2453 | 172648.2193 | 169040.7122 | 161957.5018 | 250274.2329 | 163410.7712 |
| 7-Methoxycoumarin-4-acetic acid | 2080282.503 | 4770850.916 | 2408457.06 | 4049941.984 | 4218151.87 | 1635666.374 | 3518238.432 | 3026173.295 | 2804863.449 | 3580536.26 | 3399378.128 | 2389287.739 |
| Dye X-12948-139D | 230046.7315 | 373954.0694 | 229628.3535 | 227759.9546 | 475065.6112 | 236224.4585 | 465175.6813 | 294022.9421 | 257726.8442 | 373747.4184 | 284089.0424 | 153759.9222 |
| (p-Tolylsulfonyl)acetic acid | 853368.595 | 4547003.64 | 8457797.153 | 6268509.281 | 837938.4073 | 281146.1086 | 1429706.059 | 391946.1544 | 1370248.573 | 2055174.16 | 1998531.042 | 9555080.196 |
| M395T453 | 521259.7212 | 413705.8008 | 520859.0796 | 477948.9307 | 812286.6711 | 694805.6212 | 670675.2159 | 656340.7273 | 1050871.392 | 412547.4204 | 398842.1306 | 602122.9808 |

| Metabolites | PBS_1 | PBS_2 | PBS_3 | PBS_4 | BCG_1 | BCG_2 | BCG_3 | BCG_4 | BCGΔ*pe18*_1 | BCGΔ*pe18*_2 | BCGΔ*pe18*_3 | BCGΔ*pe18*_4 |
|---|---|---|---|---|---|---|---|---|---|---|---|---|
| PE O-18:5_2:0 | 1319573.091 | 1327046.698 | 1132300.058 | 1258021.58 | 4715680.562 | 4063641.94 | 2656391.016 | 3451235.463 | 3714154.556 | 3518582.656 | 2866834.83 | 4118386.171 |
| Cer 8:1;2O/18:2 | 727039.3328 | 829076.9326 | 314545.5389 | 401188.1635 | 436194.8325 | 474407.431 | 547853.2824 | 589589.1429 | 510651.9768 | 715280.5835 | 545190.2909 | 401247.8274 |
| 4-(2-Cyclohexylethoxy)aniline | 1047486.952 | 1212282.008 | 794798.9244 | 949410.1713 | 1087727.669 | 841690.6724 | 1072784.908 | 1113827.104 | 1025627.083 | 1089118.606 | 944517.4996 | 873673.5304 |
| 3-Chloro-N-(2,2-difluoroethyl)aniline | 893662.4826 | 670216.9294 | 2002897.169 | 1677888.91 | 2660467.386 | 2719917.744 | 3035388.507 | 2879706.2 | 2700080.587 | 67790.95702 | 2194636.08 | 2632795.561 |
| N2-Benzyl-1,3,5-triazine-2,4-diamine | 97301.45614 | 332889.8148 | 469508.1994 | 386261.0005 | 252694.5221 | 203031.2681 | 404314.2003 | 158190.8761 | 125253.8134 | 318832.1561 | 132024.1619 | 1198127.234 |
| Tagitinin C | 1196907.701 | 642538.7979 | 958397.2286 | 891528.2918 | 1246122.412 | 861683.9838 | 1085312.678 | 1398113.62 | 1171299.706 | 390679.7313 | 529149.4621 | 362001.285 |
| SM 18:1;2O/24:2 | 4829625.449 | 6092407.672 | 6164092.684 | 6970089.328 | 5701471.714 | 11927821.32 | 4854708.593 | 4821116.605 | 5031760.311 | 3232528.8 | 346704.4822 | 7650655.737 |
| Testosterone decanoate | 0 | 0 | 120596.4152 | 0 | 236369.3356 | 221621.5675 | 444217.1835 | 264652.5598 | 171307.576 | 190265.7649 | 232845.4411 | 58311.53651 |
| LPC(20:0) | 720312.4776 | 81674.89855 | 392809.1514 | 638769.4851 | 841614.4699 | 613107.6835 | 1020188.439 | 1004013.58 | 679248.1525 | 0 | 656533.0629 | 476901.3822 |
| Pseudolaric Acid A | 2256230.98 | 8826779.097 | 17592527.59 | 10851340.64 | 1791487.203 | 1165274.378 | 2795472.968 | 736974.7288 | 3152803.443 | 2915682.498 | 2943900.346 | 25506628.31 |
| Dronedarone | 420033.4622 | 226380.2298 | 153641.1565 | 257402.5057 | 200165.2564 | 179569.5818 | 154571.0079 | 168266.7708 | 211190.4305 | 191475.699 | 249086.8932 | 202619.4973 |
| LPC O-22:0 | 2810057.871 | 1163262.213 | 830449.5098 | 947288.5287 | 416965.6077 | 491455.9677 | 179082.0396 | 293241.1143 | 504247.4558 | 545703.1909 | 266288.0999 | 66215.95183 |

| Metabolites | PBS_1 | PBS_2 | PBS_3 | PBS_4 | BCG_1 | BCG_2 | BCG_3 | BCG_4 | BCGΔ *pe18*_1 | BCGΔ *pe18*_2 | BCGΔ *pe18*_3 | BCGΔ*pe18*_4 |
|---|---|---|---|---|---|---|---|---|---|---|---|---|
| 12-Methyltridecanoic acid | 160372505.1 | 131148594.7 | 92195255.14 | 108107327.1 | 109844256.1 | 151540767.3 | 107300967.2 | 126756146 | 110850148.6 | 99552336.99 | 128953929.1 | 119343003.2 |
| .beta.-D-Glucopyranosiduronic acid, 4-(7-hydroxy-4-oxo-4H-1-benzopyran-3-yl)phenyl | 1022699.655 | 934199.8316 | 899471.7348 | 649550.6746 | 977280.2146 | 1153702.893 | 916867.0297 | 920549.1249 | 1013881.145 | 501608.9072 | 520799.9555 | 890360.268 |
| Dobutamine | 530166.3924 | 515317.1756 | 449671.2254 | 658102.8787 | 536823.4135 | 710412.0417 | 493405.2744 | 520474.3741 | 524905.1686 | 495550.9111 | 502507.2043 | 460524.517 |
| 8,11-Tridecadienoic acid, 13-(3-pentyl-2-oxiranyl)-, methyl ester, (8Z,11Z)- | 368559.8192 | 233208.0185 | 537568.5514 | 380417.8202 | 1012244.365 | 1407393.505 | 1262195.404 | 931973.6579 | 1531315.596 | 1290037.441 | 1755570.974 | 994297.3177 |
| Cer 8:0;2O/38:5 | 128221.3378 | 110838.7247 | 125016.5919 | 125518.2018 | 111811.8716 | 132738.9723 | 135146.708 | 125486.237 | 138856.803 | 91339.85841 | 137719.0228 | 108357.9684 |
| 4-Chloro-N-phenylpyridine-2-carboxamide | 382400.1579 | 901216.7337 | 929276.7687 | 936140.2713 | 902908.0736 | 946536.3737 | 946142.0119 | 2020710.561 | 1006577.632 | 830649.2823 | 1411938.842 | 495910.0072 |
| PG 15:0_18:1 | 182733.5996 | 111283.3211 | 221904.2978 | 41916.5916 | 1206882.379 | 853540.2966 | 1399160.584 | 1555470.421 | 1797428.558 | 1288279.607 | 1656985.53 | 649092.5726 |
| PC O-20:0 | 2930896.613 | 3089601.067 | 1456320.601 | 1870409.602 | 3751247.153 | 5394093.428 | 4543404.244 | 3651727.908 | 3562722.334 | 2362407.941 | 3715272.679 | 2237030.4 |
| M461T509 | 298608. | 236532. | 341320. | 278756. | 310632. | 370238. | 467632. | 378026. | 410956. | 433360. | 389009. | 160609. |

| Metabolites | PBS_1 | PBS_2 | PBS_3 | PBS_4 | BCG_1 | BCG_2 | BCG_3 | BCG_4 | BCGΔ*pe18*_1 | BCGΔ*pe18*_2 | BCGΔ*pe18*_3 | BCGΔ*pe18*_4 |
|---|---|---|---|---|---|---|---|---|---|---|---|---|
| | 7985 | 4241 | 4188 | 9259 | 8668 | 9887 | 12 | 3612 | 8603 | 8407 | 2831 | 6506 |
| PC 16:1_18:1 | 1638339.115 | 662311.9013 | 1151169.916 | 2231844.242 | 931429.9875 | 9723.624459 | 1401916.404 | 397780.3339 | 1484018.262 | 1472191.156 | 1889546.288 | 448290.6861 |
| N-Acetyl-beta-alanine | 1229271.328 | 1034252.899 | 1693266.846 | 1223773.956 | 2374909.008 | 1850129.51 | 2850320.247 | 2029635.412 | 1996631.685 | 1605638.006 | 1864771.308 | 1324698.877 |
| 1-Pyrrolidinecarboxamide, N,N'-(4-methyl-1,3-phenylene)bis- | 459250.3401 | 478468.7439 | 456744.6722 | 436895.8665 | 467647.1822 | 469922.0977 | 496813.3966 | 444861.4686 | 459847.7884 | 508077.2953 | 445776.0956 | 438237.6739 |
| Garcinoic acid | 0 | 7286.187984 | 5091.230056 | 0 | 465817.977 | 394253.7979 | 310561.5991 | 344907.3309 | 37568.16768 | 43109.372 | 34995.10572 | 41920.9038 |
| Hexadecyltrimethylammonium cation | 5979895.23 | 7083406.574 | 3619744.425 | 5104803.349 | 2068840.871 | 1578212.915 | 2252023.79 | 2472198.246 | 2646520.178 | 2516442.806 | 1983191.563 | 1844084.543 |
| N-Stearoyl Aspartic acid | 185610.4023 | 171517.4613 | 359824.0807 | 222069.5103 | 1131191.409 | 877191.0136 | 1086302.242 | 986682.0835 | 618033.6425 | 934275.7984 | 1008997.886 | 539678.4469 |
| Normetanephrine | 4035829.432 | 1204670.556 | 1484702.771 | 1607594.206 | 7667663.389 | 2348400.319 | 3084290.667 | 2260861.062 | 1935246.75 | 1579763.835 | 1608057.25 | 2648602.464 |
| PS(15:0/22:6(4Z,7Z,10Z,13Z,16Z,19Z)) | 7595998.583 | 7743678.838 | 6745733.534 | 6989093.21 | 7032302.832 | 5575754.203 | 6648504.036 | 6882155.924 | 6985184.1 | 7185767.995 | 6360222.353 | 5184823.65 |
| M598T493 | 2730188.863 | 2865274.443 | 3083493.212 | 2700487.185 | 2208765.942 | 2435565.674 | 2432196.539 | 2025248.312 | 2180284.526 | 2927083.874 | 3428190.988 | 3058413.093 |
| Cyclopassifloic acid C | 1341417.539 | 1572004.086 | 1282915.468 | 1765679.008 | 1443608.049 | 1510862.197 | 1499946.17 | 1388294.98 | 1504325.593 | 1727816.786 | 1241317.744 | 1525711.62 |
| 6-Pentyl-2H-pyran- | 722573. | 615007. | 523995. | 775910. | 532515. | 507894. | 475172. | 482848. | 521593. | 538285. | 729698. | 102248 |

| Metabolites | PBS_1 | PBS_2 | PBS_3 | PBS_4 | BCG_1 | BCG_2 | BCG_3 | BCG_4 | BCGΔ*pe18*_1 | BCGΔ*pe18*_2 | BCGΔ*pe18*_3 | BCGΔ*pe18*_4 |
|---|---|---|---|---|---|---|---|---|---|---|---|---|
| 2-one | 4654 | 3295 | 9408 | 6338 | 4468 | 3064 | 5073 | 1052 | 4542 | 2484 | 485 | 9.857 |
| Fagomine | 770536.2311 | 4409411.213 | 7372890.245 | 6177571.325 | 1143435.806 | 675664.9768 | 2864342.079 | 1088443.401 | 969452.1903 | 1928532.984 | 608668.8184 | 10644027.17 |
| Omarigliptin | 418854.3334 | 194260.3986 | 196316.7325 | 192372.5127 | 350535.7442 | 293341.2511 | 225926.9621 | 287877.7551 | 927544.7946 | 264531.7162 | 861358.2442 | 399602.7061 |
| 1-Palmitoyl-2-(5-keto-8-oxo-6-octenoyl)-sn-glycero-3-phosphocholine | 937983.7242 | 990659.7406 | 1200031.368 | 1020605.979 | 1010292.916 | 990697.8386 | 887313.3858 | 936164.147 | 1095788.418 | 964800.7326 | 1075020.832 | 1013629.832 |
| 5-(4-Methylpiperazin-1-yl)-5-oxopentanoic acid | 97611.13157 | 276908.2588 | 558111.4016 | 567707.6604 | 75639.07619 | 48874.29008 | 91472.96936 | 43231.72328 | 15929.16964 | 116918.4077 | 34461.77492 | 633274.0801 |
| Isoflavone base + 3O, O-MalonylHex | 234969.3742 | 210317.481 | 233912.0269 | 229289.0612 | 225152.2943 | 234914.0428 | 211983.8213 | 236205.7087 | 221640.6998 | 219082.303 | 238529.7141 | 234677.2783 |
| 10-Undecenoic acid | 1351281.072 | 1200602.776 | 1510477.679 | 1180462.383 | 1360146.653 | 1299272.505 | 1350731.197 | 1691443.938 | 1249413.609 | 1233207.052 | 1307180.05 | 1515750.315 |
| Cer(d18:0/14:0) | 186988.5066 | 496389.2368 | 306910.8772 | 291352.0605 | 253316.0011 | 344751.2368 | 219793.1832 | 401734.7943 | 173437.7841 | 113405.6507 | 487499.5341 | 227264.6913 |
| N-(1-Allylcyclohexyl)-2-ethyl-6-methylaniline | 2526183.755 | 2584911.052 | 2258040.606 | 2345541.907 | 2555242.85 | 2465468.004 | 2560244.124 | 2459627.546 | 2531954.096 | 2451567.279 | 2505968.612 | 2278253.252 |
| 2-Pyridinylmethyl 2-(4-isobutylphenyl)propanoate | 3072523.741 | 2634770.299 | 1805504.495 | 2039843.516 | 3038756.734 | 2728586.712 | 3411463.195 | 3428874.631 | 3735927.209 | 3356560.996 | 3277346.431 | 1961736.926 |

| Metabolites | PBS_1 | PBS_2 | PBS_3 | PBS_4 | BCG_1 | BCG_2 | BCG_3 | BCG_4 | BCGΔ*pe18*_1 | BCGΔ*pe18*_2 | BCGΔ*pe18*_3 | BCGΔ*pe18*_4 |
|---|---|---|---|---|---|---|---|---|---|---|---|---|
| PC(44:8) | 3679844.065 | 4924407.288 | 3289413.462 | 3167767.151 | 794468.851 | 1272402.386 | 2588623.88 | 1287821.943 | 1891196.46 | 3592387.481 | 2191571.418 | 1379340.132 |
| LPC O-20:1 | 1245174.066 | 1394815.533 | 1015616.55 | 1162374.645 | 838059.4166 | 1079770.39 | 1019571.139 | 773420.3794 | 761993.9195 | 936929.3414 | 958910.2541 | 696657.246 |
| 4-(Dodecylamino)Phenol | 441540.3125 | 439060.088 | 445844.1572 | 397930.4056 | 438992.5554 | 514541.4509 | 499777.2345 | 415511.7239 | 436336.4654 | 505411.5199 | 360967.8981 | 506685.3176 |
| Hymexelsin | 352208.3737 | 348493.2237 | 316752.1043 | 316937.1472 | 351105.4814 | 346842.6268 | 323299.2073 | 339296.7116 | 366108.1299 | 301419.5922 | 346897.0592 | 293623.2695 |
| 4-(7-Hydroxy-3-(hydroxymethyl)-5-(3-hydroxypropyl)-2,3-dihydro-1-benzofuran-2-yl)-2-methoxyphenyl 6-deoxy-.alpha.-D-mannopyranoside | 0 | 0 | 0 | 0 | 219089.1564 | 185165.7977 | 153581.5151 | 134407.1111 | 166299.1466 | 114201.3779 | 161890.1777 | 152461.4862 |
| LNAPS 3:0/N-20:2 | 173304.9362 | 157551.5764 | 341943.9085 | 263056.2409 | 286921.1446 | 319539.4715 | 384488.9273 | 316491.655 | 332089.6538 | 184763.4105 | 214063.2505 | 269332.1685 |
| PE-Cer 12:1;2O/18:2 | 0 | 51822.54403 | 0 | 46526.25141 | 0 | 5969.564405 | 0 | 2906.976268 | 25147.03687 | 28051.90275 | 0 | 51133.92836 |
| Flumiclorac-pentyl | 252491.4 | 236382.2199 | 205454.8704 | 201319.4581 | 211000.3194 | 194677.255 | 174739.0774 | 217011.8482 | 172841.2859 | 184200.8024 | 187023.0451 | 134214.0741 |
| PG 20:4_20:5 | 40907748.16 | 46216007.11 | 17471179.03 | 19371144.47 | 10953647.43 | 9040378.402 | 11613081.34 | 9043459.198 | 16446355.75 | 16861191.22 | 9874012.382 | 8612211.854 |

| Metabolites | PBS_1 | PBS_2 | PBS_3 | PBS_4 | BCG_1 | BCG_2 | BCG_3 | BCG_4 | BCGΔ*pe18*_1 | BCGΔ*pe18*_2 | BCGΔ*pe18*_3 | BCGΔ*pe18*_4 |
|---|---|---|---|---|---|---|---|---|---|---|---|---|
| Andrastin A | 312138.3755 | 343170.8985 | 707960.405 | 315157.2202 | 763440.9912 | 503005.1057 | 945353.9875 | 886166.5925 | 683428.744 | 602168.185 | 557889.8272 | 249681.0194 |
| Ethanox 310 | 1572781.461 | 6783831.247 | 683021.2511 | 1522525.668 | 3805467.44 | 1336183.932 | 3784792.716 | 2926221.489 | 1187631.797 | 5479649.474 | 2028508.774 | 1072625.562 |
| 1H-Benzimidazole, 2-[[4-(5-ethyl-4-pyrimidinyl)-1-piperazinyl]methyl]-6-(trifluoromethyl)- | 122801.0662 | 128204.6502 | 130723.8234 | 102750.1136 | 187082.6334 | 143416.5495 | 158855.2081 | 144420.7198 | 141752.514 | 160729.1356 | 144997.9807 | 148261.9574 |
| DG 24:2 | 157123.4523 | 96418.97741 | 210503.9299 | 112927.4629 | 590255.2321 | 427141.4939 | 734767.5649 | 450669.3678 | 433314.1497 | 573580.4324 | 625252.5381 | 318215.6966 |
| LPE 16:1 | 560151.6671 | 515560.3894 | 972867.1615 | 752223.9042 | 4259643.207 | 3742733.13 | 4116451.088 | 3313494.267 | 4226819.467 | 4026946.295 | 5307187.301 | 3947238.78 |
| PE O-10:0_8:0 | 695234.7921 | 727296.4704 | 510184.8843 | 687409.8887 | 416314.7845 | 288206.6641 | 573811.5535 | 485032.1708 | 467751.0644 | 463625.4466 | 620372.5841 | 192934.7162 |
| 4-Methylquinolin-2-ol | 7539664.888 | 5696051.92 | 2756236.922 | 2494321.751 | 2355112.076 | 3654516.286 | 9438468.41 | 2362299.127 | 6886201.087 | 3082900.743 | 2996882.281 | 1773754.336 |
| .gamma.-Nonalactone | 216049.1705 | 204125.137 | 209824.6407 | 219815.1904 | 301923.465 | 838804.9244 | 4428636.413 | 212822.2235 | 2666611.803 | 983624.8658 | 346685.0507 | 3755450.68 |
| PG 16:3_20:4 | 1780576.773 | 1744743.279 | 1311355.51 | 1597701.021 | 427981.3162 | 506278.9817 | 620478.1496 | 545542.3253 | 940055.1184 | 765427.4873 | 544943.4064 | 495560.3138 |
| 2'-Deoxyguanosine 5'-diphosphate (dGDP) | 165357.1135 | 206162.3984 | 234838.4948 | 187702.6865 | 250400.5412 | 204855.4913 | 241627.9037 | 695722.77 | 382030.6711 | 363968.6293 | 201840.9126 | 2174234.18 |

| Metabolites | PBS_1 | PBS_2 | PBS_3 | PBS_4 | BCG_1 | BCG_2 | BCG_3 | BCG_4 | BCGΔ *pe18*_1 | BCGΔ *pe18*_2 | BCGΔ *pe18*_3 | BCGΔ*pe18*_4 |
|---|---|---|---|---|---|---|---|---|---|---|---|---|
| 1-20:0-2-18:1-phosphatidylserine | 1375345.048 | 1679429.698 | 490257.135 | 1017649.613 | 573130.833 | 105021.406 | 422412.1854 | 534429.4732 | 338183.7115 | 415085.3348 | 507369.1207 | 587907.5902 |
| D-myo-Inositol, 1-[2-hydroxy-3-[(1-oxo-9,12-octadecadienyl)oxy]propyl hydrogen phosphate], [S-(Z,Z)]- | 151818.2726 | 125202.7923 | 234047.6025 | 181012.2935 | 428701.585 | 261195.0775 | 358332.2631 | 286369.9034 | 339000.5931 | 265917.3949 | 349739.8627 | 308024.3144 |
| Hexadecatetraenoyl carnitine (Car(16:4)) | 249177.9122 | 386573.3846 | 182565.0333 | 160231.7998 | 247398.0306 | 113618.6629 | 288456.6848 | 229008.8168 | 251754.6054 | 321939.2748 | 56057.00852 | 121821.7624 |
| Nafcillin | 9300.390463 | 8650.149542 | 0 | 11313.92305 | 515676.7266 | 315695.9806 | 644794.4419 | 498461.4451 | 509574.1441 | 591884.8748 | 614630.4376 | 639419.0571 |
| PC 18:1_34:2 | 0 | 11652.54839 | 0 | 0 | 78351.78028 | 0 | 45040.01235 | 0 | 142526.7979 | 18796.84997 | 38780.93856 | 0 |
| M258T150 | 5011553.096 | 6637035.267 | 7654004.711 | 6259688.235 | 4028796.027 | 4115977.526 | 3760450.039 | 3819146.261 | 4421012.698 | 3805044.358 | 4555459.149 | 15276855.53 |
| Ferulic acid 4-O-glucuronide | 440969.2059 | 451202.1723 | 814212.5372 | 505317.6173 | 499867.0958 | 755700.837 | 672660.802 | 454440.3914 | 410389.6756 | 561636.8961 | 634810.0219 | 710327.6762 |
| SHexCer 32:2;3O | 118953.7108 | 193283.8631 | 209295.6215 | 130563.836 | 408743.9293 | 483777.8697 | 367336.6268 | 392885.7602 | 352608.2937 | 309322.3524 | 573933.216 | 1489679.509 |
| Terbutaline | 538830.7858 | 436632.0334 | 381295.7189 | 685341.7473 | 585023.767 | 417647.3811 | 570249.1183 | 296284.9627 | 532753.0243 | 528746.8781 | 442642.1535 | 567899.2523 |

| **Metabolites** | **PBS_1** | **PBS_2** | **PBS_3** | **PBS_4** | **BCG_1** | **BCG_2** | **BCG_3** | **BCG_4** | **BCGΔ*pe18*_1** | **BCGΔ*pe18*_2** | **BCGΔ*pe18*_3** | **BCGΔ*pe18*_4** |
|---|---|---|---|---|---|---|---|---|---|---|---|---|
| 3.alpha.,4.beta.,3.alpha.-Galactotetraose | 26461.66641 | 8605.248048 | 23113.80062 | 11604.86643 | 936054.7765 | 763601.4534 | 649662.5093 | 579205.7994 | 705828.5217 | 578133.1558 | 730421.849 | 852311.7002 |
| Asparagine | 437330.27 | 342806.3179 | 463023.4898 | 257896.7347 | 1507851.344 | 614432.0744 | 807477.9705 | 1045173.871 | 988776.2317 | 1250058.25 | 650863.0446 | 746852.3368 |
| 5-Aminolevulinic acid | 681549.3994 | 827181.6967 | 929263.0737 | 582648.447 | 3235889.381 | 1842238.429 | 3081697.055 | 2783407.787 | 2422470.391 | 2203732.023 | 2094647.448 | 1887434.117 |
| NAOrn 22:6/21:2 | 11497547.89 | 3519971.955 | 21028654.46 | 9443849.352 | 8180741.43 | 185206.7334 | 10364627.66 | 11100335.63 | 6979018.582 | 9365394.737 | 6621263.172 | 24086443.24 |
| LNAPS 19:1/N-26:6 | 0 | 0 | 0 | 0 | 416462 | 448166.6963 | 378418.634 | 386280.3562 | 152498.5635 | 125133.6879 | 134386.0818 | 112783.0681 |
| N2-Succinoylarginine | 111825.752 | 80116.82319 | 128298.9357 | 138444.5834 | 1003476.88 | 773317.5141 | 680187.6531 | 649871.3788 | 1203796.488 | 786450.8469 | 821710.5741 | 1415502.466 |
| 4-(tert-Butoxycarbonyl)benzoic acid | 3274898.324 | 3602978.08 | 4377796.793 | 3900742.399 | 4263388.672 | 4216273.5 | 4231064.9 | 3624128.007 | 3189135.658 | 2912313.947 | 3801105.516 | 3976967.387 |
| Acetophenone | 1044579.222 | 965510.4526 | 790841.701 | 878994.4998 | 821562.8924 | 854296.2221 | 794862.1079 | 824538.7704 | 832371.5526 | 824964.7882 | 1060787.419 | 778943.0703 |
| M551T211 | 0 | 0 | 6371.735837 | 0 | 513329.3359 | 469030.9699 | 393555.4052 | 365826.8163 | 459115.3973 | 342229.2526 | 404497.7704 | 456394.4475 |
| SQDG 15:0_18:1 | 219064.8028 | 207171.928 | 209441.5523 | 201501.7327 | 215776.1481 | 241299.3703 | 208764.4632 | 200773.0053 | 186583.1835 | 225457.3268 | 244334.8637 | 218828.3632 |
| Gambogic acid | 113106.8799 | 90657.40571 | 157109.5958 | 127536.506 | 119992.688 | 96280.29928 | 117791.0298 | 109728.9575 | 99941.2591 | 123865.5399 | 110247.5208 | 115339.0034 |
| PI 21:1 | 102073. | 79027.8 | 78714.1 | 71506.4 | 58345.6 | 73532.8 | 69779.0 | 93820.6 | 17455.1 | 80775.1 | 21622.8 | 51991.9 |

| Metabolites | PBS_1 | PBS_2 | PBS_3 | PBS_4 | BCG_1 | BCG_2 | BCG_3 | BCG_4 | BCGΔ*pe18*_1 | BCGΔ*pe18*_2 | BCGΔ*pe18*_3 | BCGΔ*pe18*_4 |
|---|---|---|---|---|---|---|---|---|---|---|---|---|
| | 2701 | 1956 | 074 | 5102 | 5787 | 5636 | 9343 | 49 | 7028 | 8107 | 2679 | 4203 |
| PI 20:4_20:4 | 2464698.609 | 3826447.073 | 966535.68 | 1384686.508 | 255638.5142 | 366321.8405 | 368163.7195 | 327506.5131 | 475174.5066 | 545931.5893 | 310147.2903 | 393338.3091 |
| 3-(2,5-Dimethoxyphenyl)propionic acid | 975865.2905 | 1142170.891 | 1332934.605 | 1304066.441 | 922034.9595 | 930329.664 | 2032055.703 | 818920.0687 | 1044065.439 | 951622.6093 | 965263.438 | 1815590.703 |
| LPA 24:1 | 55860.29457 | 58508.25064 | 26704.02051 | 47046.65777 | 24635.75225 | 26049.09948 | 30508.35396 | 21315.11742 | 18997.31374 | 29660.13313 | 22100.69792 | 6020.151768 |
| Hypoxanthine | 4234116.138 | 3993913.383 | 5947974.926 | 4112183.928 | 5389862.703 | 3575453.125 | 5801248.623 | 4496476.793 | 3405516.291 | 4197724.834 | 4895689.042 | 2764264.052 |
| N-Demethylmifepristone | 516027.0608 | 254170.5757 | 740773.2823 | 792963.1873 | 662397.3006 | 688291.7436 | 687865.7578 | 601184.9748 | 557315.9292 | 716960.8066 | 746846.9411 | 619750.6555 |
| Soyasaponin Ba | 678349.1896 | 852979.4577 | 1177825.342 | 1298402.758 | 1038737.25 | 915548.9534 | 901061.7108 | 866662.6238 | 1076516.892 | 969064.0689 | 288221.94 | 1184922.91 |
| LNAPE 2:0/N-18:4 | 185976.6653 | 195298.309 | 129338.892 | 150776.2748 | 153960.9917 | 186871.431 | 195465.0246 | 227500.0435 | 157115.0377 | 189340.8845 | 166654.5999 | 172193.6765 |
| Entacapone 3-.beta.-D-glucuronide | 738197.6365 | 534510.1635 | 1090703.332 | 720556.429 | 2538877.335 | 1536284.523 | 2016364.435 | 1849444.755 | 1862818.975 | 1311510.701 | 1600971.236 | 1397001.533 |
| PI 19:4 | 97591.54834 | 59632.60745 | 78008.02724 | 61904.95269 | 76824.42392 | 74733.58242 | 37373.66593 | 88185.95671 | 65689.45535 | 54900.63569 | 56308.28156 | 51927.48035 |
| LPG 16:2 | 37191.26431 | 26038.18978 | 87482.56538 | 54834.65208 | 298495.3702 | 410025.9814 | 534811.9521 | 316337.8159 | 387205.4665 | 304578.5114 | 310063.9756 | 325835.9902 |
| Aztreonam | 347702. | 266155. | 159695. | 211105. | 343135. | 234501. | 233815. | 201238. | 276060. | 380557. | 276713. | 277110. |

| Metabolites | PBS_1 | PBS_2 | PBS_3 | PBS_4 | BCG_1 | BCG_2 | BCG_3 | BCG_4 | BCGΔ*pe18*_1 | BCGΔ*pe18*_2 | BCGΔ*pe18*_3 | BCGΔ*pe18*_4 |
|---|---|---|---|---|---|---|---|---|---|---|---|---|
| | 9591 | 5723 | 3069 | 4191 | 5716 | 5781 | 1976 | 7007 | 3622 | 0155 | 3655 | 0546 |
| PE 16:0_20:3;4O | 502232.8316 | 540198.5246 | 463871.8282 | 442755.2155 | 465275.6435 | 503275.0111 | 498750.664 | 626706.1868 | 339891.961 | 468386.8022 | 526435.5054 | 454093.9666 |
| PI 22:0 | 77242.93545 | 45411.89147 | 137104.689 | 57597.66626 | 95002.98129 | 96179.54258 | 132607.956 | 131327.6633 | 111514.7608 | 96673.93598 | 124555.8503 | 82417.49364 |
| Picric acid | 142979.4186 | 245971.3382 | 531530.5888 | 299163.6461 | 126974.9307 | 97388.99872 | 131541.5729 | 105099.0862 | 131363.3759 | 133210.356 | 123905.9127 | 937039.5045 |
| 1,3-Dimethylpyrazole | 3425752.087 | 2132445.027 | 2357240.86 | 1803488.077 | 2418703.435 | 2584894.977 | 2289023.607 | 2646916.089 | 2545751.767 | 3642423.543 | 2427154.966 | 758028.289 |
| 16,16-Dimethylprostaglandin A2 | 324983.8818 | 411579.068 | 368031.8893 | 371813.9687 | 382728.859 | 344334.8939 | 352686.5234 | 374639.1953 | 382239.9002 | 388124.7869 | 342569.4408 | 324125.0913 |
| Tuberonic acid | 5735300.054 | 5609196.352 | 5332254.432 | 5533977.534 | 5525939.564 | 5487006.431 | 5432679.98 | 5484290.626 | 5462468.449 | 5440754.691 | 5467747.319 | 5664460.936 |
| (2E,4Z)-9-Oxooctadeca-2,4-dienoic acid | 3084859.192 | 3049628.066 | 3167583.109 | 1984909.006 | 4682374.235 | 4677711.229 | 4947988.433 | 4280081.192 | 3931444.855 | 3576444.222 | 5751807.195 | 4135704.057 |
| 3-Chloro-4-hydrazinophenylmethylsulfonylmethyl sulfone | 8892858.712 | 6394233.058 | 9307881.505 | 10532468.88 | 9018099.547 | 7923469.092 | 8580914.593 | 8434864.434 | 9710142.938 | 8601378.925 | 9054386.017 | 8814860.16 |
| 13(S)-HpOTrE | 714417.7519 | 936253.531 | 848370.5649 | 719153.1712 | 841219.0466 | 976462.0052 | 803841.5254 | 775964.2256 | 721093.6365 | 883038.8334 | 648871.1119 | 836734.792 |
| Vanillin | 387060.3714 | 315895.4031 | 282696.8352 | 224542.4261 | 206304.0721 | 265128.8897 | 240405.6315 | 250846.1694 | 285889.7215 | 210880.2461 | 251829.8946 | 355000.8857 |

| Metabolites | PBS_1 | PBS_2 | PBS_3 | PBS_4 | BCG_1 | BCG_2 | BCG_3 | BCG_4 | BCGΔ*pe18*_1 | BCGΔ*pe18*_2 | BCGΔ*pe18*_3 | BCGΔ*pe18*_4 |
|---|---|---|---|---|---|---|---|---|---|---|---|---|
| Tridodecylmethylammonium cation | 981922.1163 | 1018810.938 | 1055118.884 | 790310.3614 | 118298.7007 | 33663.75599 | 182672.6331 | 286956.1833 | 246759.8885 | 232955.9931 | 131166.3764 | 169619.5197 |
| 3,5-dihydroxydecanoic acid | 824270.6095 | 686192.7901 | 551335.2495 | 5667363.827 | 554353.936 | 1169521.369 | 9022294.428 | 565951.6097 | 3229804.327 | 2107675.939 | 692428.3874 | 6590494.188 |
| DGCC 9:0_12:0 | 3016719.485 | 3096489.723 | 3002644.112 | 2906238.659 | 3011011.547 | 3229546.98 | 2977450.145 | 3149387.644 | 2998761.784 | 2834838.809 | 3142137.429 | 3287443.907 |
| 6-Ethylthiopurine | 3079161.001 | 4066968.251 | 2998581.961 | 3185414.371 | 5327298.938 | 3652228.508 | 5627799.394 | 3334561.039 | 2542310.668 | 4568026.618 | 3516727.124 | 2827621.921 |
| Coumestan base + 2O, O-Hex | 817516.8501 | 667949.2875 | 974472.6915 | 820395.6971 | 2433540.583 | 1825669.623 | 2228402.522 | 1854989.37 | 2123606.223 | 1643714.013 | 1863627.321 | 1634860.92 |
| Cer 8:0;2O/16:2 | 139117.4959 | 111581.5247 | 72416.31058 | 62686.26096 | 151247.0144 | 203628.5507 | 79933.84462 | 103468.5729 | 65432.83763 | 59210.02932 | 102957.7858 | 132724.3931 |
| PC 8:0_34:8 | 13954130.71 | 16186898.33 | 8837334.076 | 10520851.52 | 2683285.607 | 3341121.37 | 3628169.234 | 2777189.267 | 4805483.65 | 5085309.989 | 4573106.19 | 2998400.322 |
| Orbifloxacin | 162139.0347 | 140558.7203 | 174736.0142 | 186652.2681 | 153587.67 | 226604.6826 | 148246.7208 | 172165.9316 | 193868.3328 | 70680.20798 | 91260.81641 | 144320.1348 |
| PS 18:0_22:6;3O | 288183.8969 | 279812.9347 | 203689.282 | 230713.9148 | 853946.1475 | 655137.4059 | 1076428.764 | 787659.8502 | 817819.8921 | 866208.8996 | 773591.3443 | 512951.8369 |
| Ac2PIM2 18:2_18:2 | 7060.899233 | 0 | 7443.646076 | 6306.135192 | 54715.8225 | 25823.07918 | 78235.94117 | 36280.1271 | 17532.99113 | 21647.04294 | 73855.29064 | 27583.99083 |
| Histidine | 5494407.081 | 8560004.51 | 6461367.419 | 6563273.839 | 16417752.52 | 6877779.025 | 12517144.58 | 8980775.798 | 11022856.96 | 4647828.504 | 8546259.426 | 5889618.265 |
| 4-O-feruloyl-quinate | 5746048.994 | 3318065.556 | 4568301.495 | 3126567.638 | 1342452.972 | 1266538.267 | 2883627.857 | 2539020.201 | 939556.9801 | 1173937.685 | 1970431.03 | 1021223.922 |

| **Metabolites** | **PBS_1** | **PBS_2** | **PBS_3** | **PBS_4** | **BCG_1** | **BCG_2** | **BCG_3** | **BCG_4** | **BCGΔ*pe18*_1** | **BCGΔ*pe18*_2** | **BCGΔ*pe18*_3** | **BCGΔ*pe18*_4** |
|---|---|---|---|---|---|---|---|---|---|---|---|---|
| 3',4'-Dihydroxyacetophenone | 915165.9202 | 740263.8129 | 782693.1935 | 932905.0078 | 944553.2178 | 785908.9465 | 923158.334 | 943421.5404 | 854376.671 | 933475.4828 | 783359.2076 | 832788.861 |
| LPS 20:1 | 234842.8035 | 287507.473 | 210880.1721 | 207582.4121 | 437566.261 | 382466.9906 | 186268.3844 | 235571.2452 | 287927.5668 | 452533.0946 | 346697.2451 | 220053.3475 |
| 2-Pyrimidinylacetic acid | 6000016.02 | 1745411.803 | 560245.0746 | 2286759.666 | 714255.1935 | 1031201.394 | 823393.255 | 1717251.059 | 765025.1714 | 809863.0451 | 878894.602 | 2131247.448 |
| PI O-22:3_26:7 | 0 | 0 | 0 | 0 | 1270373.062 | 1057039.856 | 1194912.251 | 1058288.153 | 148636.2487 | 147109.015 | 202709.8902 | 111196.5064 |
| PE-Cer 12:1;2O/16:2 | 254911.5137 | 282944.2133 | 201007.3943 | 300867.6129 | 88105.39192 | 298218.7751 | 227335.1974 | 236059.9106 | 142529.5426 | 92032.05674 | 268605.7504 | 562986.3349 |
| 1H-Indazole-3-carboxamide, 1-(5-fluoropentyl)-N-8-quinolinyl- | 236168.9631 | 931647.8413 | 1921981.3 | 1063764.633 | 183343.9913 | 350372.5501 | 211332.6703 | 332017.1209 | 180743.7538 | 309674.3754 | 182184.9306 | 3322401.759 |
| (8-Methyl-8-azabicyclo[3.2.1]octan-3-yl) 3-methoxy-2-phenylpropanoate | 349849.7666 | 365037.58 | 327526.7181 | 347392.5568 | 344092.2125 | 360431.4515 | 319701.8408 | 339100.4057 | 358552.3555 | 343063.2201 | 343415.742 | 338739.9685 |
| Prostaglandin D2 (PGD2) | 104524.1426 | 97639.58858 | 147916.1577 | 127112.4868 | 3978598.809 | 5580463.293 | 2745266.979 | 4475237.352 | 6172162.222 | 1837598.459 | 2234567.183 | 6709843.609 |

| **Metabolites** | **PBS_1** | **PBS_2** | **PBS_3** | **PBS_4** | **BCG_1** | **BCG_2** | **BCG_3** | **BCG_4** | **BCGΔ *pe18*_1** | **BCGΔ *pe18*_2** | **BCGΔ *pe18*_3** | **BCGΔ*pe18*_4** |
|---|---|---|---|---|---|---|---|---|---|---|---|---|
| 2-Propenamide, 3,3'-(1,4-phenylene) bis[N-[4-(4,5-dihydro-1H-imidazol-2-yl)phenyl]- | 335218.2322 | 281074.0755 | 398383.6925 | 571434.6972 | 274678.9094 | 332160.9517 | 1954080.494 | 381518.383 | 476766.5682 | 469020.4767 | 256671.7697 | 506475.913 |
| SL 23:3;O/18:3 | 2278189.097 | 1275465.001 | 1891857.373 | 1562064.352 | 3121674.109 | 263378.3496 | 1950159.073 | 1099387.452 | 1696795.419 | 2411390.715 | 2301426.317 | 1053018.703 |
| Propentofylline | 1199212.068 | 1152804.878 | 1216180.743 | 1201877.276 | 1098766.596 | 1093027.963 | 1192344.721 | 1126014.413 | 1320706.904 | 1159402.02 | 1170117.849 | 1175757.35 |
| Leupeptin | 293585.7465 | 229122.2628 | 386957.3769 | 360507.365 | 353237.4956 | 171272.9053 | 267793.9258 | 328809.367 | 354406.6758 | 304928.4609 | 265985.5916 | 283110.8868 |
| LPC(16:0) | 65245198.77 | 10835595.27 | 12254008.43 | 10468703.25 | 123948299.6 | 16429711.58 | 21843325.32 | 107307505.1 | 93435494.97 | 105268118.1 | 18848640.74 | 13016596.38 |
| LPE(O-20:2) | 233277.7252 | 290609.9346 | 31877.1201 | 233248.1885 | 188031.086 | 71826.05888 | 34928.67283 | 142535.673 | 154001.3181 | 180462.2842 | 121988.1792 | 28583.20257 |
| PE 36:10 | 28445.02312 | 33146.20747 | 44505.59932 | 43624.68903 | 144001.9571 | 113706.548 | 164454.2983 | 178042.0745 | 148459.6226 | 204387.4401 | 181676.7625 | 119701.08 |
| Loperamide | 165646.0238 | 191903.7099 | 221767.3646 | 184988.5626 | 175777.2507 | 175140.399 | 148902.4688 | 187795.8624 | 206776.1302 | 175327.0346 | 192913.2038 | 61541.39405 |
| N-Octadecyl-N'-propylsulfamide | 369179.6485 | 2409319.494 | 263464.8713 | 373912.7178 | 167985.9939 | 305249.1896 | 300702.7529 | 3097308.858 | 140395.4081 | 295666.5283 | 256910.9713 | 335207.1436 |
| CerP 18:1;2O/2:0 | 230747.4991 | 242871.2732 | 195326.2247 | 217527.5226 | 214066.78 | 230158.711 | 218361.1958 | 251670.4807 | 234669.1935 | 222334.2698 | 217030.5512 | 194055.3988 |
| LPE 24:6 | 383623. | 375205. | 300070. | 427304. | 101486 | 103271 | 810874. | 900378. | 111070 | 912180. | 856416. | 851363. |

| Metabolites | PBS_1 | PBS_2 | PBS_3 | PBS_4 | BCG_1 | BCG_2 | BCG_3 | BCG_4 | BCGΔ*pe18*_1 | BCGΔ*pe18*_2 | BCGΔ*pe18*_3 | BCGΔ*pe18*_4 |
|---|---|---|---|---|---|---|---|---|---|---|---|---|
| | 9783 | 7453 | 656 | 6676 | 8.369 | 8.447 | 3363 | 2218 | 1.627 | 2374 | 1521 | 4697 |
| hypericin | 76048.10706 | 110475.9123 | 139752.7176 | 116284.5976 | 64981.805 | 134690.7316 | 238814.0115 | 214036.4367 | 18908.64947 | 70127.8192 | 79811.18529 | 167702.4084 |
| 5-Methylcytosine | 712822.694 | 628298.3276 | 1282905.642 | 808392.9662 | 1201015.432 | 1099081.866 | 1214568.569 | 1048415.144 | 912433.4319 | 657542.4847 | 794292.2252 | 962091.0727 |
| 2-(4-Bromophenyl)-2,3-dihydro-1H-perimidine | 1870854.1 | 1476577.926 | 1606407.028 | 1608410.505 | 1529360.771 | 1355752.834 | 1293562.176 | 1289474.131 | 2088290.27 | 1319871.24 | 1165307.907 | 1219890.192 |
| Glucoiberin | 107636.4396 | 90717.3523 | 125654.799 | 53914.15388 | 381979.4781 | 272915.0306 | 487629.087 | 315920.3142 | 361326.9129 | 377361.8792 | 368934.0357 | 337214.8945 |
| PC 9:0_20:4 | 930610.3992 | 1811665.146 | 716193.6226 | 1186992.383 | 59543.22118 | 59442.41852 | 114971.5106 | 99399.83981 | 150904.3136 | 296631.8205 | 86105.31339 | 145548.2363 |
| Isobutyrylcarnitine (Car(4:0)) | 1498770.229 | 1415680.123 | 2266337.753 | 1842132.578 | 6270345.198 | 4135892.423 | 5959990.308 | 4654622.898 | 4491243.977 | 4185253.664 | 5537269.119 | 4838675.842 |
| Hex3Cer 17:0;2O | 113364.6741 | 90083.56998 | 120033.0296 | 109113.0903 | 164041.1302 | 186875.6589 | 96319.47111 | 201752.2094 | 114979.1515 | 86613.32699 | 126726.9024 | 109979.8501 |
| Cytidine monophosphate N-acetylneuraminic acid | 258107.9872 | 311838.32 | 377230.816 | 297209.701 | 623737.7998 | 323413.44 | 739903.1787 | 488477.8442 | 308115.2969 | 475656.1465 | 478874.395 | 373701.7772 |
| N-Docosahexaenoyl Proline | 464291.1709 | 384780.3856 | 608669.0743 | 633079.3255 | 855621.8152 | 263715.0606 | 712483.5665 | 629156.9728 | 345895.3498 | 504019.8323 | 545011.5203 | 267799.4531 |

| **Metabolites** | **PBS_1** | **PBS_2** | **PBS_3** | **PBS_4** | **BCG_1** | **BCG_2** | **BCG_3** | **BCG_4** | **BCGΔ*pe18*_1** | **BCGΔ*pe18*_2** | **BCGΔ*pe18*_3** | **BCGΔ*pe18*_4** |
|---|---|---|---|---|---|---|---|---|---|---|---|---|
| (3.beta.,5.Xi.,8.alpha.,9.beta.,10.alpha.,16.beta.)-16-Hydroxykauran-3-yl .beta.-D-glucopyranoside | 1847941.485 | 1332087.804 | 2325698.597 | 1776658.834 | 2591083.466 | 2721949.639 | 2198068.227 | 2543007.12 | 2521444.611 | 4085067.942 | 3592168.183 | 2061647.054 |
| 21-Desacetyldeflazacort | 49557.63718 | 43561.41026 | 51397.43706 | 34731.0188 | 128987.4236 | 70218.89631 | 133005.0135 | 89440.696 | 109149.7162 | 166436.0007 | 165260.9952 | 103807.7972 |
| 4-Methylumbelliferyl oleate | 133796.9397 | 116083.7797 | 267719.1007 | 185904.0619 | 142822.8292 | 191494.5404 | 161457.1982 | 150142.073 | 142108.4669 | 96163.5581 | 170059.2742 | 177492.2629 |
| Dimethylmalonic acid | 309768.2713 | 358368.0698 | 542094.4597 | 591038.5151 | 627473.7012 | 623198.3854 | 577485.0945 | 669704.7477 | 469140.4903 | 627621.8691 | 539271.4065 | 606648.5666 |
| 2-oxopenta-3,4-dienoate | 713756.8196 | 2004355.677 | 2717057.986 | 2173860.806 | 3671076.405 | 3412907.191 | 2004959.058 | 1899266.883 | 3929909.393 | 3641147.74 | 6007843.881 | 4094971.8 |
| CerP 18:0;2O/2:0 | 947565.2192 | 914581.2014 | 622418.5437 | 849141.6197 | 937260.5822 | 614574.1139 | 848487.8586 | 831676.5961 | 701617.0293 | 674282.4396 | 885802.6397 | 600630.9165 |
| 2'-O-Methyluridine | 839234.6092 | 804803.4153 | 1415395.464 | 977929.1922 | 3539406.284 | 1923019.012 | 3104847.766 | 2449665.888 | 2248990.327 | 2437620.443 | 2855088.87 | 2046152.328 |
| 5-Guanidino-2-oxopentanoate | 1512793.128 | 1100962.632 | 862968.2755 | 1324192.266 | 9854952.668 | 2983751.84 | 2175558.338 | 3519900.694 | 3817763.781 | 3638336.864 | 1006917.126 | 2825101.166 |
| DGTS 12:0_20:5 | 4741998.679 | 6026136.969 | 4238642.04 | 6143776.051 | 11192892.47 | 10634188.72 | 6037664.453 | 9228720.159 | 7779723.463 | 3308533.349 | 3056799.592 | 9362475.43 |
| 3,4-Dihydroxy-4-(4-methoxyphenyl)-1,3-dihydroquinolin-2 | 518080.9655 | 383403.4267 | 608904.8273 | 436515.0349 | 959744.4408 | 643670.4271 | 753097.6348 | 773754.6079 | 770993.807 | 634519.7977 | 617433.7804 | 497578.3855 |

| Metabolites | PBS_1 | PBS_2 | PBS_3 | PBS_4 | BCG_1 | BCG_2 | BCG_3 | BCG_4 | BCGΔ *pe18*_1 | BCGΔ *pe18*_2 | BCGΔ *pe18*_3 | BCGΔ*pe18*_4 |
|---|---|---|---|---|---|---|---|---|---|---|---|---|
| -one | | | | | | | | | | | | |
| SL 10:1;O/15:1 | 181268.3246 | 147699.1591 | 306857.6156 | 197519.1788 | 137047.3672 | 91136.36493 | 124920.6007 | 122629.2259 | 58298.592 | 66844.15971 | 77339.63438 | 45118.79608 |
| Bixin | 223701.9734 | 205823.5784 | 247886.06 | 166550.5718 | 232490.0347 | 226115.7987 | 218056.8356 | 218479.4038 | 216428.7228 | 250588.5675 | 216282.3388 | 209379.1435 |
| 7-Ketodeoxycholic acid | 18962.92623 | 5495.27438 | 44757.77749 | 13483.42152 | 19568.2336 | 15907.1735 | 34263.86729 | 32165.59302 | 14955.63136 | 28745.48385 | 23636.19676 | 6648.918377 |
| 1-Palmitoyl-2-arachidonoyl-sn-glycero-3-phosphoserine | 3499705.063 | 3020720.344 | 1598532.173 | 2343359.857 | 2702894.242 | 2075605.36 | 5458446.352 | 4498837.677 | 3167452.61 | 3478164.132 | 5345264.498 | 3393560.775 |
| SMGDG O-9:0_16:4 | 102389.817 | 100633.4526 | 121645.8142 | 96592.68898 | 103665.4212 | 112199.3325 | 97547.29406 | 105866.507 | 106215.7466 | 112347.5518 | 111780.34 | 106017.6126 |
| M573T228 | 5150.222284 | 0 | 0 | 0 | 198340.5386 | 202762.1385 | 134545.9324 | 136624.9754 | 175440.7907 | 147225.9709 | 149187.6841 | 134885.7633 |
| Uric acid | 2744249.667 | 2198853.88 | 4999039.566 | 2821845.86 | 3574551.5 | 3597200.92 | 6305117.388 | 3831755.251 | 3567383.092 | 2778863.943 | 3171245.604 | 3916071.569 |
| PG O-11:0_22:4 | 26149.71589 | 32282.59196 | 32204.65298 | 31695.86138 | 33404.19126 | 30733.96599 | 36300.55235 | 35066.59936 | 31716.33021 | 33385.94294 | 35005.68321 | 30980.78267 |
| Hydroxytyrosol Acetate | 84163.17176 | 85840.34641 | 92441.11688 | 84620.86135 | 85310.20776 | 60743.66725 | 88039.31891 | 67185.46388 | 73473.33118 | 65008.01606 | 66934.91993 | 98496.13747 |

| Metabolites | PBS_1 | PBS_2 | PBS_3 | PBS_4 | BCG_1 | BCG_2 | BCG_3 | BCG_4 | BCGΔ*pe18*_1 | BCGΔ*pe18*_2 | BCGΔ*pe18*_3 | BCGΔ*pe18*_4 |
|---|---|---|---|---|---|---|---|---|---|---|---|---|
| TG(22:5(7Z,10Z,13Z,16Z,19Z)/20:5(5Z,8Z,11Z,14Z,17Z)/22:6(4Z,7Z,10Z,13Z,16Z,19Z))[iso6] | 1652193.841 | 909094.2947 | 930233.1794 | 829958.8282 | 644815.3062 | 552905.7666 | 1052084.123 | 1136586.791 | 919960.1738 | 1467958.241 | 841577.0043 | 444488.8914 |
| PMeOH 22:4_26:4 | 372178.8925 | 313413.0226 | 366785.7822 | 435636.4904 | 290453.1871 | 101820.485 | 493928.5038 | 439000.1506 | 306042.6317 | 376275.2014 | 343545.2883 | 385777.3285 |
| CE 18:5 | 0 | 0 | 27955.87472 | 0 | 1002117.67 | 776693.2269 | 898556.8745 | 1097715.892 | 694911.4085 | 632295.5082 | 729419.1412 | 698882.4311 |
| PE O-16:1_18:2 | 3235903.836 | 1452738.275 | 2493019.906 | 2270771.592 | 1516492.14 | 153753.4618 | 1127150.374 | 1126420.89 | 1927071.2 | 2243701.069 | 1306211.996 | 1056542.453 |
| 4-Hydroxyphenylacetic acid | 361333.3646 | 285614.0106 | 397206.7797 | 353733.6495 | 545920.8451 | 503696.8259 | 547295.6313 | 452433.4357 | 426242.1546 | 427047.2128 | 413228.5775 | 405053.9552 |
| Cer 8:1;2O/22:6 | 637044.2164 | 807868.7366 | 736638.8727 | 608708.836 | 804574.4803 | 770223.0121 | 637302.0841 | 651673.7612 | 753403.9693 | 747535.6762 | 617122.1043 | 797589.7077 |
| 3-Chloro-1H-indazole | 1075673.89 | 1215762.321 | 1885959.289 | 1644750.978 | 1377504.926 | 1874511.338 | 1369330.229 | 1348226.236 | 1345287.81 | 1221561.736 | 1741715.291 | 1507843.756 |
| 5-Amino-6-methyl-1,3-dihydro-2H-benzimidazol-2-one | 1507133.397 | 2945038.104 | 2290702.111 | 1545652.382 | 4338361.533 | 2383974.341 | 2102214.063 | 2196610.506 | 2886525.506 | 2594091.372 | 2631513.042 | 1469583.705 |
| PE 18:0_22:6;4O | 65757.58092 | 30866.10336 | 58902.96365 | 53992.32458 | 322376.7981 | 295356.6808 | 254135.8379 | 237790.5078 | 267512.6606 | 297305.7542 | 274648.9417 | 221410.6946 |
| MGDG O-10:0_2:0 | 0 | 18183.55058 | 9334.49085 | 7141.381085 | 136227.0326 | 87620.26514 | 37875.47993 | 82496.70351 | 31474.20652 | 9572.590501 | 10258.57779 | 22134.09462 |

| Metabolites | PBS_1 | PBS_2 | PBS_3 | PBS_4 | BCG_1 | BCG_2 | BCG_3 | BCG_4 | BCGΔ*pe18*_1 | BCGΔ*pe18*_2 | BCGΔ*pe18*_3 | BCGΔ*pe18*_4 |
|---|---|---|---|---|---|---|---|---|---|---|---|---|
| LPC O-16:4 | 1168312.281 | 1310641.634 | 729078.0076 | 978596.9755 | 687914.5084 | 877521.8675 | 603126.7967 | 659944 | 625355.8039 | 667850.9094 | 732398.2195 | 343890.3021 |
| Myricetin, 3-Galactopyranoside | 0 | 0 | 0 | 0 | 281113.1264 | 125172.962 | 126129.3705 | 94137.17219 | 178800.3579 | 224749.2426 | 387251.1716 | 260785.9108 |
| 19(R)-Hydroxyprostaglandin B2 | 2806987.125 | 991066.2919 | 383198.7474 | 872297.9564 | 226989.2028 | 2259783.608 | 246365.596 | 810806.8479 | 239242.5702 | 503450.6672 | 833399.6391 | 1201701.523 |
| 4-Hydroxypropranolol .beta.-D-glucuronide | 867529.8869 | 886016.1478 | 818344.2602 | 710678.6458 | 108646.2972 | 95062.3979 | 132469.393 | 211354.456 | 170427.5493 | 217910.8693 | 119339.0201 | 194150.1621 |
| N-Arachidonoyl-3-hydroxy-.gamma.-aminobutyric acid | 187158.5072 | 293450.9961 | 102518.7492 | 160828.6336 | 566233.7919 | 665078.2237 | 608652.3445 | 440734.924 | 888274.6428 | 617879.2056 | 703409.9847 | 517508.9956 |
| Methyl Paraben | 915165.9202 | 740263.8129 | 782693.1935 | 932905.0078 | 944553.2178 | 785908.9465 | 923158.334 | 943421.5404 | 854376.671 | 933475.4828 | 783359.2076 | 832788.861 |
| PI 26:5 | 140951.5055 | 145746.4155 | 149839.9752 | 96382.50805 | 160231.1855 | 146054.2608 | 141677.6266 | 146159.6448 | 121499.816 | 149437.3587 | 149706.5941 | 133969.9895 |
| gamma-Glutamylvaline | 311536.272 | 427011.4564 | 356307.8174 | 446711.1244 | 1112040.636 | 1089211.835 | 1157920.12 | 1065933.739 | 783097.5318 | 911480.9245 | 771125.7686 | 1016016.757 |
| LPC O-20:0 | 421668.6311 | 603924.6167 | 119403.5694 | 308449.5407 | 84545.81543 | 87881.78307 | 103885.5452 | 131017.1148 | 175292.8763 | 159390.8556 | 59693.29744 | 109725.2453 |
| 1-(3,4-Diethoxybenzylidene)-6,7-diethoxy-1,2,3,4-tetrahy | 243804.2639 | 286840.1571 | 256756.3928 | 261544.3316 | 289030.7853 | 249297.3509 | 262429.2318 | 272984.5626 | 260116.4127 | 270747.2833 | 264132.1559 | 269705.587 |

| Metabolites | PBS_1 | PBS_2 | PBS_3 | PBS_4 | BCG_1 | BCG_2 | BCG_3 | BCG_4 | BCGΔ *pe18*_1 | BCGΔ *pe18*_2 | BCGΔ *pe18*_3 | BCGΔ*pe18*_4 |
|---|---|---|---|---|---|---|---|---|---|---|---|---|
| droisoquinoline | | | | | | | | | | | | |
| Kaurane-17,18-dioic acid | 3234435.344 | 2597910.831 | 3437494.922 | 2719925.35 | 1909394.794 | 1949469.916 | 2349618.483 | 3038010.927 | 3624804.747 | 1424405.03 | 3376453.448 | 5125137.236 |
| SQDG 15:0_20:4 | 385747.5898 | 457995.1676 | 445463.5195 | 386794.4499 | 387592.5318 | 421590.4555 | 415079.7894 | 380806.9609 | 450028.3861 | 483728.7577 | 492971.5113 | 383530.9611 |
| 2-Amino-4,4,4-trifluorobutanoic acid | 518484.3342 | 356577.7417 | 668543.5705 | 503145.3057 | 2560235.837 | 1454620.293 | 2313243.762 | 1754408.249 | 1729863.82 | 1268046.229 | 1473209.911 | 1298087.07 |
| 4-[2,5-Dimethyl-1-(tetrahydrofuran-2-ylmethyl)-1H-pyrrol-3-yl]-1,3-thiazol-2-amine | 812636.4136 | 418477.5388 | 1157316.705 | 427110.6891 | 1646100.669 | 1054587.004 | 2024958.083 | 1226280.292 | 1022068.123 | 1448496.014 | 1577703.936 | 765490.3209 |
| LNAPS 9:0/N-24:7 | 173352.7484 | 668080.6547 | 128636.5203 | 683059.0629 | 449728.4588 | 363744.4927 | 1876746.127 | 797980.1827 | 388064.1848 | 808485.9243 | 264714.6808 | 839624.324 |
| MGDG O-10:0_7:0 | 743187.5493 | 762215.4011 | 645082.8835 | 814752.4644 | 831956.1697 | 797641.3565 | 1158858.385 | 949660.8227 | 798246.6626 | 832858.8853 | 720857.0961 | 733476.4779 |
| LPC O-13:0 | 299122.2471 | 296632.6093 | 240256.0123 | 216905.85 | 250154.0297 | 258384.009 | 202868.6185 | 238567.5586 | 213128.0911 | 157685.4483 | 218785.2047 | 211613.6165 |
| 1,3-Dimethyl-8-phenyl-3,9-dihydro-1H-purine-2,6-dione | 96964.21305 | 93903.94576 | 122447.2277 | 82014.60257 | 462721.7555 | 491338.9643 | 413695.0186 | 347986.8671 | 353148.9605 | 378784.4966 | 412125.8622 | 337926.6061 |
| 1-(3-Ethyl-2,4-dihy | 58967.6 | 78922.9 | 98846.4 | 80977.3 | 247670 | 179584 | 168128 | 137830 | 176044. | 172617. | 214353. | 196665. |

| Metabolites | PBS_1 | PBS_2 | PBS_3 | PBS_4 | BCG_1 | BCG_2 | BCG_3 | BCG_4 | BCGΔ*pe18*_1 | BCGΔ*pe18*_2 | BCGΔ*pe18*_3 | BCGΔ*pe18*_4 |
|---|---|---|---|---|---|---|---|---|---|---|---|---|
| droxy-6-methoxyphenyl)-1-butanone | 0128 | 9258 | 6473 | 0773 | 3.633 | 2.072 | 5.7 | 7.791 | 8385 | 8243 | 0093 | 2855 |
| (S)-(+)-4-Amino-3-hydroxybutyric acid | 866550.7804 | 538380.5257 | 1015859.346 | 582814.4835 | 3911287.841 | 2559561.92 | 3242462.288 | 3065671.379 | 3479819.26 | 1437503.967 | 1290359.165 | 1410714.786 |
| [(5-Chloro-1H-benzimidazol-2-yl)sulfanyl]acetic acid | 1224187.53 | 1192778.146 | 897565.8726 | 1015722.871 | 1101919.272 | 1598980.912 | 1171521.816 | 1023064.454 | 1318397.18 | 1247426.985 | 1124673.157 | 1780562.641 |
| LNAPE 17:0/N-24:4 | 343071.7206 | 130544.7494 | 710339.751 | 540563.3763 | 248789.4972 | 5279.051995 | 220412.771 | 329440.6382 | 397405.8382 | 251002.3719 | 172659.0539 | 249445.2188 |
| D-erythro-Sphingosine C-20 | 1412189.342 | 1900120.313 | 1545956.583 | 1030648.549 | 1234688.039 | 1835980.493 | 1499134.872 | 1229992.538 | 1204663.903 | 1657337.11 | 1485412.201 | 1087349.097 |
| PC O-19:1 | 1611834.001 | 1735193.537 | 1671264.821 | 1307014.586 | 3517310.148 | 5378703.408 | 3919957.842 | 3395884.548 | 2742548.396 | 2753558.678 | 3994341.984 | 2626615.154 |
| PE 18:0_20:5;4O | 152867.5814 | 99624.27501 | 138058.4044 | 92253.662 | 364064.926 | 425280.9878 | 339192.3832 | 312708.0997 | 250787.686 | 309865.3245 | 427749.1098 | 250917.2351 |
| PS 25:4 | 0 | 0 | 0 | 0 | 172443.3194 | 77747.03948 | 90784.96758 | 106546.3574 | 107695.8751 | 19046.31351 | 76653.77752 | 40093.4584 |
| Methyl 3-hydroxy-4-nitrobenzoate | 145663.8732 | 123340.5634 | 30211.87161 | 153193.3214 | 153579.2472 | 98012.53745 | 99488.55007 | 91120.95319 | 90126.62886 | 87372.30138 | 82473.97928 | 65037.69783 |
| ST 24:1;O3;G/16:2 | 178271.8536 | 184028.0779 | 142357.2208 | 159667.2381 | 168204.3214 | 183729.003 | 149499.8785 | 144775.8761 | 164582.4833 | 159888.7203 | 175607.8341 | 149960.7809 |
| SM 23:3;2O/30:2 | 693570.5961 | 521002.3456 | 1300441.649 | 664446.8479 | 181018.7292 | 0 | 227343.0925 | 135865.2086 | 163683.3451 | 177489.6134 | 175974.7669 | 67039.72234 |

| Metabolites | PBS_1 | PBS_2 | PBS_3 | PBS_4 | BCG_1 | BCG_2 | BCG_3 | BCG_4 | BCGΔ*pe18*_1 | BCGΔ*pe18*_2 | BCGΔ*pe18*_3 | BCGΔ*pe18*_4 |
|---|---|---|---|---|---|---|---|---|---|---|---|---|
| DGDG O-18:4_4:0 | 256573.2879 | 247265.4702 | 242188.734 | 252390.7653 | 237049.165 | 248045.6554 | 289224.0228 | 266606.5356 | 248169.3255 | 258139.7583 | 279202.1918 | 229851.0149 |
| SL 11:1;O/22:4 | 129276.8594 | 158816.2614 | 118168.0833 | 132527.9168 | 153418.7692 | 147822.4959 | 131747.8209 | 152713.5631 | 143715.4563 | 140536.0529 | 137929.9161 | 103193.4229 |
| Bakers_yeast_extract | 499308.7148 | 584402.2968 | 808151.1508 | 777783.3057 | 836683.4054 | 1180322.505 | 1057400.499 | 1056471.596 | 1154064.17 | 814243.5259 | 755543.2753 | 1336097.411 |
| 3-Methylhistidine | 1109850.185 | 796308.2006 | 3258210.575 | 828641.8788 | 3234900.122 | 2661435.159 | 4272144.983 | 2557588.732 | 2635466.932 | 2851665.881 | 2700424.499 | 1998192.202 |
| cis-8,11,14-Eicosatrienoic acid | 8624689.183 | 8359638.294 | 21340030.61 | 13800917.89 | 7139986.308 | 8420637.158 | 10080403.91 | 8293640.302 | 7521062.23 | 6857984.623 | 9338003.2 | 7281451.562 |
| DGDG O-15:1_28:5 | 102262.7373 | 0 | 220561.6745 | 172384.495 | 939960.3383 | 487042.9507 | 918428.7588 | 762552.2007 | 923318.0381 | 1123837.663 | 1091878.448 | 379814.1063 |
| 6-[3-[(3,4-dimethoxyphenyl)methyl]-4-methoxy-2-(methoxymethyl)butyl]-4-methoxy-1,3-benzodioxole | 336528.4288 | 337643.3509 | 320703.3724 | 377513.9112 | 348696.9288 | 339997.9042 | 394640.6363 | 369882.375 | 343487.0162 | 360787.6492 | 349993.9085 | 332132.2113 |
| 5-O-Demethylnobiletin | 163637.4497 | 123851.8227 | 256384.6097 | 169602.4952 | 141335.253 | 110521.4265 | 158446.2697 | 112514.0591 | 109762.2351 | 137054.7566 | 135736.618 | 100360.9744 |
| (+)-Corydaline | 1243265.357 | 1241907.126 | 1091511.562 | 1251336.397 | 1261247.929 | 1298116.84 | 1182911.226 | 1254450.13 | 1206999.502 | 1174863.107 | 1220335.426 | 1255368.674 |
| Cer 18:0;2O/16:0 | 3541981.065 | 1953373.563 | 3191332.61 | 4167384.858 | 2743467.063 | 63901.9694 | 1719109.624 | 968044.6355 | 3533880.331 | 2404379.525 | 2063694.928 | 1204871.2 |

| Metabolites | PBS_1 | PBS_2 | PBS_3 | PBS_4 | BCG_1 | BCG_2 | BCG_3 | BCG_4 | BCGΔ*pe18*_1 | BCGΔ*pe18*_2 | BCGΔ*pe18*_3 | BCGΔ*pe18*_4 |
|---|---|---|---|---|---|---|---|---|---|---|---|---|
| Cys-Met | 2009852.302 | 1564262.695 | 2483481.542 | 1997357.76 | 2639945.076 | 2733766.597 | 31579.36968 | 2491675.818 | 2336374.987 | 1886397.317 | 2031676.569 | 2192359.425 |
| 5'-Deoxyadenosine | 335044.7114 | 208552.5023 | 805913.93 | 376122.4776 | 4079067.43 | 3713319.635 | 3492634.84 | 3210751.668 | 3199537.889 | 2392341.722 | 2789614.954 | 2990251.498 |
| Tridecanedioic acid | 7364.627146 | 19061.49343 | 26296.34048 | 15722.67087 | 2292631.245 | 10108220.36 | 3165408.005 | 2972964.655 | 1136188.487 | 488756.6459 | 1451598.123 | 2453399.965 |
| 3-(2-Oxocyclohexyl)propanoic acid | 401020.4674 | 386099.9278 | 717279.0247 | 693727.5455 | 316329.3154 | 468503.0319 | 696167.0616 | 726204.8744 | 398050.1591 | 415836.4891 | 408124.9207 | 488834.2377 |
| Oleoyl 3-carbacyclic phosphatidic acid | 59328.49876 | 29569.46367 | 15880.68797 | 40224.485 | 500625.6334 | 287610.9515 | 128114.9024 | 98876.18601 | 242264.5584 | 517093.2623 | 245100.6667 | 13437.14879 |
| IS3 | 18693650.28 | 19795330.58 | 20438117.6 | 19482479.3 | 19188580.31 | 20253370.25 | 20237794.13 | 19754265.48 | 19767491.27 | 17077092 | 19186649.55 | 17861545.24 |
| 1-[(Dimethylamino)carbonyl]-3-piperidinecarboxylic acid | 0 | 784851.4529 | 1555543.662 | 1872291.765 | 0 | 0 | 299251.0447 | 0 | 70795.23668 | 199876.226 | 57555.76838 | 2849854.826 |
| DGGA 26:4_18:5 | 1404343.78 | 2115501.425 | 1248531.123 | 1705746.599 | 1592376.534 | 1755194.235 | 989489.4009 | 1166570.279 | 1351735.014 | 1645034.345 | 1556082.321 | 879993.9432 |
| TG 8:0_9:0_16:4 | 380298.9376 | 331965.1773 | 424028.576 | 408627.7595 | 116303.4668 | 125539.5015 | 178363.0467 | 120809.4973 | 77218.77221 | 102506.1845 | 97365.48459 | 64435.2652 |
| Naftopidil | 209030.1496 | 101668.8016 | 279474.2235 | 145411.7839 | 200569.8648 | 202683.2248 | 693882.8011 | 396744.8853 | 171667.3601 | 201955.3988 | 261030.5043 | 152742.5784 |
| SL 10:1;O/12:0 | 15457.1 | 23120.7 | 13944.8 | 28534.2 | 101675 | 819926. | 111616 | 906569. | 101492 | 101695 | 903297. | 640783. |

| Metabolites | PBS_1 | PBS_2 | PBS_3 | PBS_4 | BCG_1 | BCG_2 | BCG_3 | BCG_4 | BCGΔ *pe18*_1 | BCGΔ *pe18*_2 | BCGΔ *pe18*_3 | BCGΔ*pe18*_4 |
|---|---|---|---|---|---|---|---|---|---|---|---|---|
| | 1118 | 116 | 3771 | 6281 | 0.718 | 5007 | 8.576 | 524 | 9.51 | 5.257 | 7305 | 2883 |
| (2Z)-2-Fluorobut-2-enedioic acid | 967697.4139 | 964587.1973 | 1604370.844 | 1267354.382 | 1246439.795 | 15376774.51 | 1104263.897 | 1113851.723 | 1114765.383 | 1130467.827 | 1663514.025 | 1333995.152 |
| 4'-Hydroxy-3'-methylacetophenone | 469357.3227 | 411192.0641 | 540137.8528 | 464280.5025 | 565449.2124 | 521610.515 | 447724.3909 | 518253.8203 | 518079.5871 | 312234.0585 | 393581.8531 | 486326.1621 |
| SM 8:1;2O/2:0 | 769489.2791 | 124166.3008 | 1000579.245 | 237940.0445 | 1278948.154 | 1537761.877 | 5431338.246 | 4699091.606 | 1907351.064 | 1208943.092 | 2093928.925 | 646769.269 |
| Bavachin | 1921107.847 | 2008409.409 | 1793538.378 | 1958476.471 | 1846350.346 | 1928008.612 | 1829410.661 | 1963876.127 | 1947728.4 | 1920847.449 | 2089151.446 | 1517242.751 |
| Manidipine (Manyper) | 583900.4889 | 603228.8648 | 449641.7289 | 536655.7329 | 578808.9745 | 492398.6152 | 489154.1809 | 492240.655 | 536005.7185 | 557323.3793 | 602556.7143 | 488847.6532 |
| PE(22:5(7Z,10Z,13Z,16Z,19Z)/16:1(9Z)) | 1368263.025 | 1467498.031 | 2191536.83 | 1758949.818 | 3179612.471 | 2539500.773 | 2479403.414 | 2230676.576 | 2283727.294 | 2243015.649 | 2687730.84 | 1973379.038 |
| PS 16:0_18:2 | 1797154.461 | 2411579.61 | 3434583.383 | 2221492.859 | 1323192.655 | 34032.58987 | 16269066.26 | 768954.0007 | 1881379.601 | 1415138.543 | 915485.543 | 738759.678 |
| PC 35:0_16:1 | 328151.2816 | 224863.8335 | 216532.4892 | 212127.0346 | 205684.0195 | 15814.39188 | 208152.873 | 178251.3614 | 177377.5913 | 178802.5559 | 257322.045 | 128417.6462 |
| Benzene, 1-fluoro-2-[(1E)-2-(4-methoxyphenyl)ethenyl]- | 818076.1459 | 813249.7054 | 769848.4479 | 821438.355 | 887132.6563 | 830918.1822 | 915683.7566 | 888764.233 | 796107.9167 | 803121.372 | 789411.2936 | 762128.8727 |
| PE O-19:1_20:3 | 227838.2442 | 260263.6969 | 208867.1333 | 212374.2317 | 254017.8024 | 0 | 151640.1343 | 52983.47008 | 173092.1413 | 128159.1623 | 219315.7138 | 42048.65122 |

| Metabolites | PBS_1 | PBS_2 | PBS_3 | PBS_4 | BCG_1 | BCG_2 | BCG_3 | BCG_4 | BCGΔ*pe18*_1 | BCGΔ*pe18*_2 | BCGΔ*pe18*_3 | BCGΔ*pe18*_4 |
|---|---|---|---|---|---|---|---|---|---|---|---|---|
| Tetradecadienoylcarnitine (Car(14:2)) | 277361.4511 | 269099.3212 | 604622.8822 | 320187.7983 | 488612.5284 | 289545.2104 | 538205.9022 | 481797.9929 | 203972.7992 | 373829.0598 | 285673.5303 | 137405.5701 |
| Mibefradil | 523338.5459 | 799159.9994 | 275685.6257 | 457231.6123 | 353578.4783 | 683540.9305 | 1254479.899 | 274662.1122 | 307369.5448 | 494837.5597 | 401015.4376 | 678024.8663 |
| 2-[3-(2-Quinolylmethoxy)anilino]benzoic acid | 908674.7806 | 1097753.922 | 1261455.638 | 849853.4379 | 584623.7764 | 1011840.617 | 635806.2268 | 597510.288 | 458708.9182 | 560849.1694 | 673802.2374 | 2352876.647 |
| LPC 16:2 | 215459.4394 | 205831.5332 | 369497.6296 | 222594.3977 | 474218.6316 | 562440.7841 | 552101.827 | 547413.3059 | 629315.6808 | 499145.7038 | 583525.1763 | 542464.4815 |
| Artemoin_C | 134176.634 | 252599.6396 | 113661.67 | 378168.4918 | 741981.9069 | 577065.4399 | 851185.3362 | 798469.3962 | 826857.0212 | 1112828.661 | 898166.8632 | 495757.4887 |
| Nylidrin | 596307.7083 | 382008.932 | 578607.3259 | 522150.5015 | 543567.3487 | 545519.5486 | 562042.105 | 527586.3592 | 450001.1815 | 389262.5381 | 486177.3881 | 435979.6113 |
| 2-Hexenoylcholine | 15070.12915 | 662602.219 | 1303299.486 | 1105707.03 | 216683.2677 | 25215.61206 | 183605.7609 | 30259.98326 | 160836.8293 | 464198.1022 | 45870.4298 | 2572158.476 |
| Cucurbitacin_B | 350064.1328 | 423706.4722 | 243745.6848 | 403174.8723 | 181050.3586 | 96808.63532 | 144361.8013 | 149965.2341 | 205146.9497 | 161957.6572 | 145455.4537 | 153676.957 |
| 2-[2-[19-acetamido-6-(3,4-dicarboxybutanoyloxy)-16,18-dihydroxy-5,9-dimethylicosan-7-yl]oxy-2-oxoethyl]butanedioic aci | 144147.3873 | 139251.7407 | 144558.2806 | 152087.7578 | 154882.1449 | 170442.2132 | 139027.778 | 145648.8002 | 163919.4541 | 169322.1794 | 142045.9775 | 142566.6959 |

| Metabolites | PBS_1 | PBS_2 | PBS_3 | PBS_4 | BCG_1 | BCG_2 | BCG_3 | BCG_4 | BCGΔ*pe18*_1 | BCGΔ*pe18*_2 | BCGΔ*pe18*_3 | BCGΔ*pe18*_4 |
|---|---|---|---|---|---|---|---|---|---|---|---|---|
| Dioctyl sebacate | 70083.16439 | 26691.63891 | 80033.56351 | 71159.57081 | 46010.29897 | 179025.2566 | 33482.69074 | 103568.2733 | 10051.19526 | 12692.94451 | 52973.42684 | 89255.64353 |
| Osthol | 59641.87285 | 102899.8793 | 240793.4005 | 199475.3434 | 67699.92794 | 60206.57765 | 68324.64225 | 61687.08025 | 36733.74843 | 59453.93969 | 63948.29391 | 167269.6325 |
| (R)-4-((1R,3S,5S,7R,8S,9S,10S,13R,14S,17R)-1,3,7-trihydroxy-10,13-dimethylhexadecahydro-1H-cyclopenta[a]phenanthren-17-yl)pentanoic acid | 12402.86883 | 30672.47803 | 36654.65421 | 35156.77686 | 90517.82695 | 59852.57053 | 0 | 83191.13293 | 31771.8862 | 311743.8663 | 93019.92927 | 18454.20509 |
| Cyanidin_7-arabinoside | 492782.3204 | 284496.3898 | 845192.8999 | 374369.5275 | 8295992.906 | 5045361.337 | 8012636.997 | 5920350.173 | 4798515.705 | 4984139.019 | 5698818.733 | 3916940.608 |
| PC 54:15 | 306521.1483 | 491485.0327 | 521943.8191 | 578031.369 | 15630.03298 | 12342.49911 | 14666.49111 | 12591.08348 | 0 | 15975.63687 | 7320.631508 | 22806.46472 |
| PC O-18:1_22:5 | 270209.4195 | 313850.7107 | 509444.7693 | 408271.9676 | 487807.365 | 12215.44521 | 428767.9851 | 1938879.52 | 368080.2591 | 297122.5008 | 401002.2676 | 887125.3934 |
| PA 4:0_22:6 | 279290.041 | 274881.066 | 246751.6102 | 274938.4285 | 216955.6147 | 249759.3352 | 215451.4016 | 212377.7901 | 247869.9437 | 256540.8481 | 256016.5524 | 199192.5796 |
| trans-Epoxysuccinyl-L-leucylamido(4-guanidino)butane | 437517.7334 | 526236.376 | 637277.0582 | 596724.5463 | 627371.8204 | 681095.3054 | 477660.7805 | 550367.7213 | 724513.2138 | 519005.7803 | 565613.4628 | 1151080.21 |
| SMGDG | 107237 | 113208 | 133323 | 119957 | 352611 | 320706 | 358241 | 337092 | 391586 | 411188 | 419344 | 306968 |

| Metabolites | PBS_1 | PBS_2 | PBS_3 | PBS_4 | BCG_1 | BCG_2 | BCG_3 | BCG_4 | BCGΔ*pe18*_1 | BCGΔ*pe18*_2 | BCGΔ*pe18*_3 | BCGΔ*pe18*_4 |
|---|---|---|---|---|---|---|---|---|---|---|---|---|
| O-16:3_4:0 | 16.69 | 99.12 | 20.87 | 95.82 | 3.929 | 6.19 | 9.013 | 8.703 | 3.095 | 0.18 | 5.328 | 0.288 |
| LPI 18:3 | 85485.62007 | 45090.51752 | 141548.0564 | 56845.71702 | 123018.1884 | 77229.79385 | 248462.1923 | 135822.6226 | 71680.30361 | 77619.11723 | 116828.1478 | 44820.14726 |
| Tanshinone I | 153170.0399 | 197009.8018 | 234896.2617 | 192882.3401 | 227766.2204 | 427369.8343 | 309770.3819 | 201591.3411 | 210093.3138 | 163486.0484 | 208429.5056 | 174868.6274 |
| Clofazimine | 234434.7796 | 448402.6267 | 419851.6001 | 205414.8899 | 304505.2614 | 337688.4157 | 376657.0132 | 288078.1048 | 258761.6823 | 139192.8968 | 164174.5322 | 78227.46575 |
| Cynaroside | 1440606.475 | 865740.9025 | 1608923.804 | 1159240.763 | 429112.6043 | 1102097.117 | 755256.103 | 923432.4274 | 716597.7682 | 297122.4165 | 250440.8247 | 617334.2025 |
| Verbasoside | 0 | 4298.425124 | 0 | 0 | 519961.4044 | 506757.0756 | 347992.5658 | 388208.1817 | 362048.9079 | 348711.6594 | 350363.1443 | 342443.4964 |
| Heneicosanoic acid | 157374.0526 | 53560.24038 | 176151.6556 | 152597.1395 | 193283.4391 | 474211.2854 | 173529.565 | 285080.5932 | 150655.9007 | 144161.9744 | 157431.811 | 186261.3457 |
| Methanone, [1-(5-methylhexyl)-1H-indol-3-yl](2,2,3,3-tetramethylcyclopropyl)- | 1086269.544 | 1142526.689 | 1227104.118 | 1172640.362 | 1189296.7 | 1350553.103 | 1268027.907 | 1201206.165 | 1200172.596 | 1285358.602 | 1225587.683 | 1258555.512 |
| 7-Chloro-5-methyl-2H-1,4-benzothiazin-3(4H)-one | 88831.61362 | 59338.51561 | 96297.01757 | 72900.58053 | 116870.577 | 74663.00882 | 243649.6108 | 99959.5311 | 70473.71481 | 86945.40699 | 92075.18831 | 65594.80581 |
| Thiolutin | 311732.5966 | 252036.7869 | 154575.8465 | 278758.9576 | 814124.8386 | 626970.451 | 925722.0617 | 760424.2582 | 696275.416 | 925475.8459 | 973496.4746 | 374386.5295 |
| cis-11.14-Eicosadie | 487643 | 530878 | 888722 | 550644 | 402705 | 538236 | 435838 | 424439 | 450601 | 451416 | 543632 | 398085 |

| **Metabolites** | **PBS_1** | **PBS_2** | **PBS_3** | **PBS_4** | **BCG_1** | **BCG_2** | **BCG_3** | **BCG_4** | **BCGΔ*pe18*_1** | **BCGΔ*pe18*_2** | **BCGΔ*pe18*_3** | **BCGΔ*pe18*_4** |
|---|---|---|---|---|---|---|---|---|---|---|---|---|
| noic acid | 4.847 | 8.399 | 6.269 | 5.256 | 9.006 | 3.931 | 3.71 | 6.663 | 5.638 | 7.075 | 7.954 | 0.574 |
| CAR 19:0 | 144993.6786 | 335467.3347 | 468279.0342 | 368317.7979 | 2485848.624 | 1980655.249 | 2478073.37 | 2714306.851 | 1803292.409 | 1776161.84 | 1860611.828 | 1229759.644 |
| 1,2-Di(4Z,7Z,10Z,13Z,16Z,19Z-docosahexaenoyl)-sn-glycero-3-phosphoethanolamine | 171688.5664 | 180540.3451 | 380455.5863 | 916065.7596 | 206214.8403 | 60366.39754 | 893409.3939 | 51245.71401 | 482426.1634 | 143591.7032 | 73436.86222 | 507908.1437 |
| Aspulvinone_H | 737764.4707 | 834549.8365 | 1214739.438 | 883814.818 | 750034.643 | 837973.8548 | 694930.7374 | 1013767.971 | 829363.9085 | 815782.8946 | 1037146.678 | 965212.8746 |
| GlcCer(d18:1/16:0) | 19507805.58 | 17372192.59 | 21159751.24 | 14481060.5 | 26384150.09 | 25367531.92 | 35415642.36 | 32507692.44 | 31436679.67 | 38168145.87 | 31054341.58 | 15971189.15 |
| Apigenin_7-[rhamnosyl-(1->2)-galacturonide] | 690996.2758 | 306036.6106 | 979999.3329 | 459441.0638 | 699965.3832 | 475430.8436 | 612997.736 | 406499.6223 | 388024.4484 | 337521.4285 | 294824.0956 | 310259.9095 |
| 2-Amino-6-(methylsulfanyl)-3,5-pyridinedicarbonitrile | 2967822.94 | 2357502.107 | 3999179.457 | 3332878.002 | 13874297.32 | 11989275.14 | 12718164.41 | 11311293.98 | 12424202.56 | 6950448.178 | 9066382.071 | 10186758.42 |
| SL 26:0;O/26:0;O | 56589.10979 | 38417.94779 | 253323.0697 | 27452.77983 | 151443.9255 | 94988.044 | 379336.4063 | 351156.9104 | 262748.2051 | 178840.1957 | 358784.7741 | 121075.7397 |
| SMGDG O-22:6_3:0 | 64539.95206 | 73407.52393 | 44558.39893 | 63112.07186 | 53989.62292 | 62324.99601 | 69883.56335 | 52430.6971 | 58654.33109 | 61280.68774 | 59178.24835 | 52437.62437 |
| Dimethyldioctadecylammonium cation | 633626.2386 | 2254219.352 | 857205.7349 | 881176.423 | 2665084.784 | 1221578.921 | 1116011.201 | 583682.2445 | 883789.1391 | 1385125.203 | 31074.18865 | 736810.2784 |

| **Metabolites** | **PBS_1** | **PBS_2** | **PBS_3** | **PBS_4** | **BCG_1** | **BCG_2** | **BCG_3** | **BCG_4** | **BCGΔ*pe18*_1** | **BCGΔ*pe18*_2** | **BCGΔ*pe18*_3** | **BCGΔ*pe18*_4** |
|---|---|---|---|---|---|---|---|---|---|---|---|---|
| GAMMA-CAPROLACTONE | 385222.1419 | 551247.816 | 371805.9226 | 394656.242 | 399702.4677 | 404085.8319 | 428814.0091 | 396782.6246 | 417176.7335 | 389545.7369 | 685242.9799 | 372521.1556 |
| Cinncassiol_E | 149962.7207 | 127449.2324 | 102198.78 | 101878.8735 | 157968.0853 | 145033.0176 | 77331.98211 | 106806.3827 | 160790.2403 | 136428.0375 | 133629.3191 | 148388.2879 |
| Maltotriose | 907127.9644 | 940822.2463 | 734091.1353 | 915921.9653 | 834334.1127 | 833772.1911 | 882320.7955 | 872004.4122 | 841358.964 | 871528.0934 | 843957.7805 | 866588.0327 |
| CerP 16:0;2O/22:0 | 84148.32891 | 102868.0471 | 81672.34191 | 81978.72199 | 104262.5712 | 96672.29442 | 82105.12936 | 82104.16931 | 85729.68579 | 76285.2988 | 69755.33017 | 75988.97744 |
| FA 16:3 | 2064709.62 | 1909468.201 | 4387491.818 | 2430177.773 | 188608.8972 | 439420.7967 | 296275.3974 | 501450.4777 | 508725.5275 | 195065.3673 | 369323.3586 | 521947.6932 |
| PC O-15:1_22:6 | 168442.7213 | 76429.77992 | 228516.7687 | 154310.5195 | 339083.9124 | 891441.0931 | 499413.9136 | 380039.5163 | 224498.2727 | 36023.34019 | 731591.1148 | 704694.953 |
| HexCer 9:0;2O/9:0 | 188146.4175 | 228236.2032 | 200392.5411 | 290364.8956 | 223942.3154 | 223985.4785 | 238953.5319 | 236433.6217 | 128135.3718 | 87712.67999 | 209274.2131 | 217488.3319 |
| 2-Octenoylcarnitine | 752218.8813 | 835500.0429 | 747492.2382 | 803078.7191 | 798497.2888 | 785052.0001 | 831481.8499 | 749738.7151 | 760488.9579 | 743580.8516 | 760649.05 | 767613.71 |
| 3-Methyladipic acid | 487744.1141 | 433515.5341 | 382584.3484 | 438819.3679 | 2759399.109 | 1475477.106 | 2931381.034 | 2062198.103 | 2045316.961 | 603121.5778 | 834279.6532 | 915359.306 |
| N-(6-Amino-1-butyl-2,4-dioxo-1,2,3,4-tetrahydro-5-pyrimidinyl)-N-methylbenzamide | 18353.17928 | 35648.42265 | 67056.01261 | 45495.42787 | 5887.022994 | 7213.591302 | 5381.851214 | 13332.92107 | 4449.129888 | 18646.23747 | 6434.962466 | 106864.5278 |
| MOPS | 381218. | 585486. | 169685. | 39316.0 | 282455. | 130515 | 356788. | 437176. | 759163. | 594149. | 761930. | 421618. |

| Metabolites | PBS_1 | PBS_2 | PBS_3 | PBS_4 | BCG_1 | BCG_2 | BCG_3 | BCG_4 | BCGΔ*pe18*_1 | BCGΔ*pe18*_2 | BCGΔ*pe18*_3 | BCGΔ*pe18*_4 |
|---|---|---|---|---|---|---|---|---|---|---|---|---|
| | 6 | 9229 | 2531 | 4166 | 2879 | 7.817 | 0074 | 134 | 2698 | 6949 | 2271 | 365 |
| PS 3:0_19:1 | 136697.8784 | 84788.34392 | 145152.3401 | 124903.5483 | 136718.562 | 160401.6618 | 118597.2859 | 127564.5705 | 150793.4238 | 117896.5796 | 136807.999 | 91998.89495 |
| LNAPE 2:0/N-12:0 | 312518.9786 | 274524.9305 | 225623.2467 | 274176.031 | 309773.6456 | 247531.2488 | 303944.6829 | 263421.0425 | 245297.3927 | 247079.4814 | 236560.8448 | 262725.351 |
| ST 24:1;O4;T;S | 101607.1737 | 64105.79767 | 92391.98883 | 126263.9331 | 150228.7965 | 125499.0901 | 81313.86739 | 118360.529 | 92670.65018 | 155048.1822 | 95572.27761 | 138757.4192 |
| LPC(20:5) | 211747.0862 | 173798.1256 | 172085.1944 | 198366.2822 | 474173.3993 | 350297.3488 | 344293.6198 | 374191.1338 | 407403.8835 | 285458.0122 | 353436.9355 | 341629.9815 |
| (R)-4-((3R,4R,5S,7R,8R,9S,10R,12S,13R,14S,17R)-3,4,7,12-tetrahydroxy-10,13-dimethylhexadecahydro-1H-cyclopenta[a]phenanthren-17-yl)pentanoic acid | 1670353.429 | 1835022.164 | 1494924.431 | 1433823.276 | 1671951.177 | 1954399.888 | 1327142.436 | 1659439.726 | 1520451.449 | 1614317.959 | 1640965.429 | 1563520.864 |
| Nervonic acid | 2395613.591 | 1769498.363 | 1766195.158 | 1748388.827 | 1704337.612 | 2799587.632 | 1623582.647 | 2227087.114 | 1411187.667 | 1063908.351 | 1767805.839 | 1308113.815 |
| Diglycine | 437330.27 | 342806.3179 | 463023.4898 | 257896.7347 | 1507851.344 | 614432.0744 | 807477.9705 | 1045173.871 | 988776.2317 | 1250058.25 | 650863.0446 | 746852.3368 |
| PI 16:4_22:6 | 27041.23646 | 30028.56737 | 206507.944 | 87178.01105 | 739375.4406 | 395579.2818 | 765102.8091 | 413328.1274 | 268540.9147 | 643386.3291 | 756619.0143 | 241536.8818 |

| Metabolites | PBS_1 | PBS_2 | PBS_3 | PBS_4 | BCG_1 | BCG_2 | BCG_3 | BCG_4 | BCGΔ *pe18*_1 | BCGΔ *pe18*_2 | BCGΔ *pe18*_3 | BCGΔ*pe18*_4 |
|---|---|---|---|---|---|---|---|---|---|---|---|---|
| [(2R,3S,4S,5R,6R)-3,4,5-Trihydroxy-6-[2-(3-hydroxy-5-oxooxolan-3-yl)propoxy]oxan-2-yl]methyl (E)-3-(4-hydroxyphenyl)prop-2-enoate | 0 | 0 | 0 | 0 | 1326939.546 | 1262944.16 | 913803.8433 | 1022770.531 | 1104492.858 | 651130.3705 | 818852.758 | 1088130.826 |
| trans-4-Hydroxycyclohexanecarboxylic acid | 1096410.919 | 907266.3435 | 1006713.273 | 982471.8225 | 943416.5459 | 1193394.13 | 1062547.3 | 922584.1105 | 902949.6143 | 1022042.702 | 856331.342 | 1019696.291 |
| PI O-22:6_16:0 | 1161025.975 | 1482128.972 | 359424.0875 | 1130131.045 | 434205.4562 | 330782.4441 | 394012.0724 | 476878.1674 | 652557.9057 | 524442.2151 | 412747.4658 | 128140.8249 |
| M514T459 | 977971.5294 | 960907.7652 | 813870.9043 | 1019989.868 | 1048458.138 | 932333.3468 | 1103143.807 | 988089.1347 | 1035082.634 | 1016145.388 | 906510.098 | 895814.6845 |
| PS 25:2 | 129118.5492 | 170123.0363 | 82620.14165 | 139845.7226 | 196150.0602 | 68451.96482 | 162325.9896 | 128105.5343 | 99589.35853 | 156529.5079 | 112573.9438 | 81657.31566 |
| Phenothrin | 138173.5402 | 486915.3307 | 156101.7421 | 493338.9197 | 1009595.605 | 518346.1175 | 575633.0949 | 192541.8068 | 335418.1747 | 718165.386 | 519006.936 | 132653.4075 |
| SL 10:0;O/23:0 | 402648.3637 | 358993.7193 | 469412.2152 | 675822.3576 | 313545.0913 | 222865.4604 | 317030.3231 | 408482.9147 | 233126.8145 | 329579.4845 | 246550.92 | 161462.1827 |
| M215T106 | 2297164.239 | 2100963.663 | 1919679.423 | 2622256.41 | 2041456.582 | 2233521.261 | 2229607.252 | 2016964.126 | 2331506.637 | 1797632.541 | 2146990.242 | 2047490.136 |
| Cer 18:1;2O/20:0 | 187796. | 152304. | 216063. | 263500. | 130098. | 209634. | 102237. | 118667. | 129819. | 106881. | 0 | 203404. |

| Metabolites | PBS_1 | PBS_2 | PBS_3 | PBS_4 | BCG_1 | BCG_2 | BCG_3 | BCG_4 | BCGΔ*pe18*_1 | BCGΔ*pe18*_2 | BCGΔ*pe18*_3 | BCGΔ*pe18*_4 |
|---|---|---|---|---|---|---|---|---|---|---|---|---|
| | 7209 | 9441 | 3499 | 8894 | 0914 | 4613 | 897 | 734 | 4278 | 1208 | | 9313 |
| N-[3-([1,1'-Biphenyl]-4-yloxy)-3-(4-fluorophenyl)propyl]-N-methylglycine | 1679390.675 | 1808341.399 | 2167939.558 | 1816865.298 | 1761715.27 | 1780907.039 | 1576167.584 | 1540980.311 | 1870051.541 | 1596565.893 | 1739409.906 | 3051849.161 |
| PI O-20:5_26:7 | 144990.2036 | 106508.6621 | 49366.94601 | 51826.35525 | 127686.0861 | 148375.8949 | 127413.0088 | 81330.53591 | 94637.77107 | 123374.7357 | 98827.24937 | 53312.76631 |
| PC O-16:0_18:1 | 229810.3622 | 55751.02581 | 432068.46 | 322806.3759 | 234594.2458 | 10836.34451 | 182006.8548 | 200989.5605 | 203600.8159 | 191874.8007 | 165794.5504 | 252132.4554 |
| Leukotriene B4 3-aminopropylamide | 293043.2395 | 448031.3549 | 232308.216 | 517668.7001 | 290696.828 | 441955.9651 | 448637.7908 | 402144.0398 | 428196.5106 | 489880.7241 | 404103.4835 | 446345.4218 |
| Adenosine 5'-diphosphoribose | 573093.6373 | 366532.6491 | 614405.4997 | 527606.3918 | 2240922.102 | 2314282.809 | 2475251.159 | 2701893.476 | 3237806.398 | 2546354.495 | 2663571.241 | 3993114.889 |
| Cedrin | 1244567.949 | 861414.8602 | 801940.3297 | 714296.479 | 1448939.384 | 1107286.738 | 1232929.277 | 1226010.739 | 1083412.013 | 1222208.295 | 883606.5451 | 421139.6943 |
| Hyodeoxycholic acid | 25017.90944 | 83124.85562 | 171961.3388 | 390541.7161 | 0 | 115281.5525 | 177712.2839 | 145487.2895 | 109661.1905 | 0 | 61615.55055 | 92265.08239 |
| N-Nervonoyltaurine | 463422.8881 | 312999.9059 | 360155.3598 | 361573.6564 | 140006.29 | 141086.2213 | 117919.3356 | 0 | 41246.37289 | 0 | 111878.6627 | 57387.79503 |
| 5-Methoxydimethyltryptamine | 1406733.226 | 1433750.057 | 960459.3676 | 1149775.024 | 1318314.084 | 1269128.725 | 1278862.404 | 1384026.532 | 1352553.543 | 1308397.029 | 1358047.467 | 1193010.78 |

| Metabolites | PBS_1 | PBS_2 | PBS_3 | PBS_4 | BCG_1 | BCG_2 | BCG_3 | BCG_4 | BCGΔ*pe18*_1 | BCGΔ*pe18*_2 | BCGΔ*pe18*_3 | BCGΔ*pe18*_4 |
|---|---|---|---|---|---|---|---|---|---|---|---|---|
| 7-[cis-3-(1-Azetidinylmethyl)cyclobutyl]-5-[3-(phenylmethoxy)phenyl]-7H-pyrrolo[2,3-d]pyrimidin-4-amine | 2449015.315 | 2597544.188 | 2290561.243 | 2257237.958 | 2296288.25 | 2271884.437 | 2566929.607 | 2570923.779 | 2432543.818 | 2269193.905 | 2379685.652 | 2336937.15 |
| Betaine aldehyde hydrate [M] | 16462919.91 | 16453207.82 | 17259036.06 | 17379826.57 | 18358445.46 | 16290453.45 | 16792483.77 | 17049197.92 | 17126800.88 | 17870690.81 | 15075262.61 | 15372543.03 |
| 5-(2-Methyl-4-nitrophenyl)-2-furoic acid | 1999083.441 | 1018430.017 | 2813890.04 | 1104923.985 | 1841952.962 | 1725191.228 | 1538254.284 | 1179427.677 | 1218821.967 | 1025153.598 | 956296.5073 | 691517.1628 |
| NAGlySer 16:4/22:0 | 234576.636 | 262327.055 | 231570.0197 | 252469.8778 | 228323.049 | 275107.9727 | 343325.131 | 255670.9 | 254169.0175 | 286163.4659 | 241001.0963 | 258361.8774 |
| Thromboxane B3 | 1065020.185 | 1575892.477 | 989516.3295 | 1086401.326 | 1020677.392 | 942376.7458 | 1974537.219 | 1070506.245 | 999612.7321 | 1235370.543 | 919380.5919 | 1772845.397 |
| nicotinamide guanine dinucleotide | 26941.23797 | 13796.38916 | 38963.7502 | 28067.1006 | 197970.3899 | 127143.3395 | 201869.9995 | 161176.4917 | 148703.6077 | 158181.8768 | 134795.7709 | 121735.75 |
| 1-acyl-PAF | 0 | 20050.61751 | 37445.31646 | 14156.59338 | 2304607.593 | 776666.7844 | 1779747.949 | 807272.822 | 1007835.686 | 0 | 1268252.722 | 677863.7028 |
| 9-Hydroxynonanoic acid | 667285.3809 | 592367.837 | 578041.3646 | 655501.3498 | 519629.5461 | 593754.0709 | 582870.977 | 615227.8586 | 538558.6222 | 876043.9844 | 638876.2348 | 732504.2694 |

| Metabolites | PBS_1 | PBS_2 | PBS_3 | PBS_4 | BCG_1 | BCG_2 | BCG_3 | BCG_4 | BCGΔ *pe18*_1 | BCGΔ *pe18*_2 | BCGΔ *pe18*_3 | BCGΔ*pe18*_4 |
|---|---|---|---|---|---|---|---|---|---|---|---|---|
| 3-Methylhexahydropyrrolo[1,2-a]pyrazine-1,4-dione | 611668.547 | 609146.2358 | 672236.018 | 441582.2255 | 545704.6084 | 647426.1131 | 457383.9479 | 2824644.142 | 350761.253 | 491226.5983 | 1459032.467 | 1198532.48 |
| PI 17:0_18:0 | 1555058.175 | 1761082.507 | 1680008.002 | 1984799.306 | 544912.8452 | 19370.03002 | 735222.1543 | 81675.15553 | 1762974.903 | 521625.0513 | 1120549.631 | 81800.35827 |
| FAHFA 22:6/2:0 | 1019546.784 | 1217986.546 | 1549617.836 | 1085694.508 | 275510.4794 | 934419.6219 | 399568.4307 | 344170.1434 | 268778.561 | 243887.8046 | 401059.601 | 275752.4181 |
| DLCL 16:5_22:7 | 0 | 0 | 0 | 0 | 456648.851 | 597649.7497 | 522924.7351 | 456023.2502 | 343705.7364 | 240187.8321 | 339367.027 | 222297.3333 |
| BMP 20:4_22:6 | 7089265.297 | 9449994.313 | 3504585.992 | 5625349.101 | 996384.8465 | 2396799.363 | 1361414.438 | 1491255.067 | 1769652.219 | 1606019.553 | 1381657.309 | 1333564.822 |
| [3-(hexadecanoyloxy)-2-[icosa-5.8.11-trienoyloxy]propoxy]({[2.3.4.5.6-pentahydroxycyclohexyl]oxy})phosphinic acid | 269248.4848 | 721249.6914 | 340534.0489 | 536546.3752 | 366925.6573 | 51567.95362 | 477002.4651 | 391212.4436 | 384816.9657 | 545089.1566 | 53066.87021 | 141264.696 |
| 2-Oleoyl-1-palmitoyl-sn-glycero-3-phosphoserine | 25575.8277 | 74844.442 | 132187.5715 | 42137.60928 | 2237618.976 | 346970.8513 | 2319308.526 | 602345.7477 | 468160.8879 | 924465.7797 | 534310.5827 | 262440.3064 |
| Decoquinate | 169470.096 | 176957.5798 | 169347.6042 | 162112.2093 | 191735.9378 | 182221.2845 | 192260.8096 | 191506.6153 | 185204.222 | 175622.6404 | 157001.7493 | 158324.1702 |
| LPC O-24:1 | 662291.9298 | 971693.5244 | 239611.3891 | 801759.6679 | 13138.22042 | 65095.43954 | 29151.72158 | 33288.87038 | 86732.07494 | 23416.32186 | 67935.09221 | 87608.19927 |

| Metabolites | PBS_1 | PBS_2 | PBS_3 | PBS_4 | BCG_1 | BCG_2 | BCG_3 | BCG_4 | BCGΔ*pe18*_1 | BCGΔ*pe18*_2 | BCGΔ*pe18*_3 | BCGΔ*pe18*_4 |
|---|---|---|---|---|---|---|---|---|---|---|---|---|
| Phenacetin | 2634474.458 | 353014.8686 | 2828986.29 | 945197.113 | 8905580.724 | 7692550.895 | 7892637.677 | 8162075.674 | 413225.0783 | 441542.1654 | 140858.1138 | 888511.5284 |
| Cer 17:2;2O/28:6 | 395515.4032 | 429403.9594 | 168314.4601 | 360062.1214 | 213016.4998 | 401702.8164 | 351230.2693 | 251759.9863 | 346867.9327 | 313985.9562 | 298539.8983 | 272629.779 |
| Bortezomib (Velcade) | 138282.0601 | 162586.2635 | 233732.8429 | 176413.1592 | 233630.8821 | 222373.7513 | 186542.691 | 188737.0408 | 241449.7535 | 200410.2765 | 192490.2248 | 400736.9799 |
| PC O-16:4_22:5 | 529051.9048 | 515390.7959 | 445997.9848 | 407547.4152 | 284571.911 | 410355.5137 | 322955.2244 | 322631.1608 | 220037.5439 | 227591.4438 | 348727.9831 | 326411.488 |
| 1H-Imidazole-4,5-dicarbohydrazide | 734045.6808 | 692642.3355 | 800964.841 | 1118688.659 | 963465.2855 | 914029.2341 | 971778.9789 | 711030.5689 | 810421.9725 | 413197.7515 | 717723.0502 | 911343.8751 |
| Hex2Cer 12:1;2O | 68315.61598 | 104616.6665 | 106779.5112 | 68565.58618 | 61252.06594 | 110725.8005 | 94822.9518 | 57588.30649 | 70883.4126 | 73089.54303 | 90492.94741 | 84635.9325 |
| 2-Dechloroethylifosfamide | 1325720.548 | 1068492.833 | 889023.738 | 1830434.072 | 1296527.509 | 847010.8484 | 1301036.607 | 1201566.26 | 887619.0224 | 1674766.479 | 1544048.602 | 2346249.579 |
| Betonicine | 684973.2687 | 1512797.832 | 1377631.617 | 1246593.391 | 12315115.39 | 250684.9726 | 320634.2818 | 265844.1816 | 484712.2905 | 767255.5997 | 1089867.655 | 1846542.656 |
| 1-(6Z,9Z,12Z,15Z-octadecatetraenoyl)-glycero-3-phosphate | 100382.7412 | 101664.8678 | 97857.13769 | 94862.67369 | 96181.24944 | 91817.80206 | 80477.79508 | 93240.72747 | 97438.39119 | 106007.533 | 99179.80362 | 94031.47014 |
| 5-Benzylidene-1-phenylbarbituric acid | 1808893.616 | 1686147.337 | 1618828.439 | 1987436.338 | 1569549.423 | 1238774.437 | 1516016.236 | 1511869.539 | 3765593.149 | 1621550.283 | 1781767.88 | 2231990.916 |
| Go 6983 | 992781.6153 | 884365.1076 | 761185.0356 | 814664.7323 | 762883.4215 | 701012.2502 | 822759.7461 | 982235.8567 | 755255.7645 | 842158.7408 | 719955.4216 | 731859.7238 |

| **Metabolites** | **PBS_1** | **PBS_2** | **PBS_3** | **PBS_4** | **BCG_1** | **BCG_2** | **BCG_3** | **BCG_4** | **BCGΔ*pe18*_1** | **BCGΔ*pe18*_2** | **BCGΔ*pe18*_3** | **BCGΔ*pe18*_4** |
|---|---|---|---|---|---|---|---|---|---|---|---|---|
| Platensimycin | 157287.5805 | 118592.6932 | 114514.0438 | 145285.5059 | 108558.112 | 187696.5838 | 153527.1475 | 145232.5295 | 137334.9702 | 144083.1616 | 161595.4718 | 225873.0157 |
| Isoleucyl-Alanine | 375690.6685 | 449435.9098 | 1548523.385 | 490404.4241 | 989793.267 | 606041.2506 | 1013508.341 | 722404.7705 | 558126.8144 | 719199.7434 | 771299.6625 | 918159.8515 |
| PI 4:0_18:5 | 105819.0981 | 95185.02543 | 79242.99387 | 64969.48063 | 151511.3489 | 104008.3288 | 86461.99044 | 82489.70899 | 125824.0598 | 88059.87887 | 90458.05446 | 86084.43921 |
| N-Cyclohexyl-N'-(1H-tetraazol-5-yl)urea | 2641419.564 | 2766581.088 | 2886796.881 | 2612005.847 | 2842841.009 | 2903485.687 | 2886005.3 | 2608935.334 | 2713007.026 | 2778215.428 | 3088525.963 | 2644845.094 |
| LPG O-24:6 | 49453.71149 | 82692.52691 | 58881.48095 | 41623.19291 | 261847.4839 | 285295.0728 | 201064.9618 | 229052.337 | 212558.9912 | 726969.4372 | 237224.3465 | 186549.7116 |
| (R)-4-((8S,9S,10R,13R,14S,17R)-10,13-dimethyl-3-oxo-2,3,8,9,10,11,12,13,14,15,16,17-dodecahydro-1H-cyclopenta[a]phenanthren-17-yl)pentanoic acid | 92294.25094 | 31894.37355 | 106544.1148 | 20345.76809 | 381824.0325 | 284517.6447 | 474619.835 | 319346.1683 | 123193.8913 | 94991.61839 | 62637.71656 | 126702.3674 |
| SM 8:0;2O/7:0 | 75166.36859 | 70399.85802 | 56921.7447 | 66045.65871 | 51604.42534 | 55419.20623 | 47659.40292 | 71574.79586 | 83284.34374 | 67488.46373 | 46819.54362 | 50722.47282 |
| Leu-Asp | 314877.6968 | 320255.4485 | 400861.9947 | 318159.397 | 1282962.087 | 1005681.417 | 1189065.007 | 929804.8491 | 973467.714 | 977859.5407 | 996569.0553 | 906799.154 |
| PC 9:0_28:6 | 425610. | 234743 | 459380. | 562493. | 619329 | 711030 | 446040 | 622295 | 598102 | 586270 | 441387 | 449036 |

| Metabolites | PBS_1 | PBS_2 | PBS_3 | PBS_4 | BCG_1 | BCG_2 | BCG_3 | BCG_4 | BCGΔ*pe18*_1 | BCGΔ*pe18*_2 | BCGΔ*pe18*_3 | BCGΔ*pe18*_4 |
|---|---|---|---|---|---|---|---|---|---|---|---|---|
| | 9001 | 6.017 | 4675 | 1167 | 9.474 | 6.371 | 8.841 | 2.791 | 2.863 | 0.724 | 1.737 | 4.807 |
| MLS001148580-01!6ALPHA,9ALPHA-DIFLUOROPREDNISOLONE 21-ACETATE 17-BUTYRATE | 103931.9513 | 40478.43663 | 459537.2605 | 126043.0116 | 73623.53565 | 108886.116 | 145663.4875 | 138882.0516 | 28505.10857 | 20506.24865 | 52562.81889 | 33119.07838 |
| rac-trans-4-Cotinine carboxylic acid | 560635.9959 | 291186.7864 | 375955.0845 | 300874.68 | 430992.1082 | 432085.3598 | 392442.7694 | 400252.0953 | 433448.8907 | 366015.8125 | 360743.594 | 410637.0059 |
| 5-{[2-(3,4-Dimethoxyphenyl)ethyl]amino}-1,3,4-thiadiazole-2-thiol | 2233891.942 | 1623185.664 | 4474141.375 | 1094431.013 | 2249308.731 | 393364.8967 | 3423481.516 | 1992018.478 | 292719.6561 | 1085264.757 | 1837256.89 | 529783.5094 |
| .alpha.-Naphthylthiourea | 780971.0965 | 616383.7036 | 1456626.038 | 887629.371 | 1120823.674 | 1108518.761 | 1276552.576 | 1301412.588 | 879472.5865 | 644059.1091 | 756990.5683 | 860375.7261 |
| LNAPE 5:0/N-15:0 | 359528.3781 | 534355.7685 | 153590.0818 | 428763.0581 | 162705.8536 | 218842.5727 | 144042.7367 | 179772.3831 | 182614.3525 | 185629.2903 | 132889.5994 | 143347.9862 |
| Isonicotinic acid | 3869974.369 | 7174557.91 | 6743822.849 | 8511258.185 | 6627930.801 | 4058247.505 | 2627977.764 | 4788994.375 | 7658888.17 | 1404747.68 | 8889052.335 | 7646665.261 |
| N-oleoylserotonin | 181696.8736 | 196509.7622 | 174505.4473 | 220567.5696 | 180879.1941 | 210930.1351 | 178593.8315 | 189620.2779 | 180705.1784 | 173818.0749 | 224101.2473 | 146191.8432 |
| Oxypurinol | 104733.358 | 107517.9923 | 114477.4718 | 133971.7315 | 1550326.016 | 861700.1338 | 349051.3165 | 1201422.325 | 988655.4293 | 151869.1795 | 1315035.655 | 188479.5862 |
| Eicosadienoylcarnit | 836193. | 660622. | 101107 | 698844. | 579484. | 494623. | 661475. | 706333. | 279304. | 476113. | 551577. | 189274. |

| Metabolites | PBS_1 | PBS_2 | PBS_3 | PBS_4 | BCG_1 | BCG_2 | BCG_3 | BCG_4 | BCGΔ*pe18*_1 | BCGΔ*pe18*_2 | BCGΔ*pe18*_3 | BCGΔ*pe18*_4 |
|---|---|---|---|---|---|---|---|---|---|---|---|---|
| ine | 074 | 7053 | 6.596 | 424 | 3505 | 1909 | 645 | 9288 | 0408 | 6861 | 941 | 4571 |
| Ganoderic Acid C2 | 94922.98816 | 68271.90556 | 106128.4663 | 61874.26687 | 224930.1377 | 127783.9244 | 269633.5552 | 228836.1921 | 123764.5894 | 129816.0883 | 117595.5619 | 83174.49453 |
| PC(10:0/10:0) | 563984.0791 | 497842.0475 | 791487.6962 | 388915.5344 | 846216.4317 | 740896.9417 | 829953.1312 | 858066.0894 | 816683.7139 | 906995.7765 | 971289.3032 | 670288.6333 |
| tert-Butyl 3-carbamothioylpiperidinecarboxylate | 405944.2407 | 385012.0225 | 555206.7506 | 485611.8008 | 384507.1197 | 357226.236 | 533466.4713 | 393913.8245 | 433683.3183 | 397542.411 | 385860.6138 | 400281.9672 |
| DGDG O-28:3_15:1 | 5514422.923 | 5508453.894 | 2657751.88 | 4224238.013 | 4669398.843 | 6129757.886 | 6484524.731 | 6053603.115 | 3851013.755 | 3474622.309 | 3784865.159 | 990563.2602 |
| Methanone, [1-(cyclohexylmethyl)-1H-indol-3-yl](4-methoxy-1-naphthalenyl)- | 781662.7242 | 872487.6977 | 822751.0908 | 848950.0949 | 672795.1906 | 898325.3538 | 783049.1997 | 894329.6668 | 672743.0537 | 521603.2614 | 776115.5305 | 737495.6765 |
| 6-Fluoro-4-hydroxycoumarin | 16134484.45 | 916172.6402 | 1016981.257 | 725771.6349 | 617215.4168 | 708011.3851 | 898686.8247 | 656798.3455 | 728355.769 | 626336.2822 | 658148.6843 | 907341.5717 |
| Ac2PIM1 18:0_16:1 | 38297.51963 | 93184.87527 | 20470.36802 | 97892.0974 | 124687.1148 | 70068.5227 | 327930.7306 | 96568.8749 | 0 | 115887.9302 | 0 | 32522.72561 |
| 1-18:0-2-18:1-phosphatidylserine | 3206627.489 | 3328119.548 | 3277255.111 | 4229492.997 | 5495482.575 | 4518377.883 | 5176746.122 | 2738011.375 | 2153684.767 | 984424.5726 | 955016.5474 | 2205051.97 |
| PI(18:2(9Z,12Z)/18:2(9Z,12Z)) | 383967.5589 | 426901.1225 | 182234.3436 | 283803.8672 | 779512.9144 | 417571.5127 | 538953.8 | 488148.6482 | 550851.9508 | 810790.7847 | 665436.3674 | 386877.9793 |

| Metabolites | PBS_1 | PBS_2 | PBS_3 | PBS_4 | BCG_1 | BCG_2 | BCG_3 | BCG_4 | BCGΔ*pe18*_1 | BCGΔ*pe18*_2 | BCGΔ*pe18*_3 | BCGΔ*pe18*_4 |
|---|---|---|---|---|---|---|---|---|---|---|---|---|
| AHexCer (O-22:0)18:1;2O/16:2;O | 230527.6813 | 370542.0059 | 404672.0985 | 337718.967 | 11308.80938 | 8128.31991 | 0 | 7793.365466 | 9217.895994 | 7523.040304 | 6455.231903 | 9385.974716 |
| 5-Methoxy-.alpha.-ethyltryptamine | 1985355.186 | 1596548.496 | 2063759.552 | 1523294.702 | 1562766.797 | 1660793.026 | 1556378.18 | 1583776.511 | 1695760.927 | 1668090.681 | 1664788.756 | 1602874.771 |
| SL 22:1;O/18:2 | 275038.2939 | 252311.5877 | 196587.6384 | 312698.0718 | 507036.6963 | 402489.9394 | 299750.3333 | 545752.602 | 585585.1309 | 202081.9554 | 344300.4126 | 315367.193 |
| SM 23:1;2O/10:0 | 1471936.814 | 1771378.981 | 1417376.814 | 936467.1013 | 1036699.601 | 0 | 1209754.667 | 898103.2805 | 1333346.896 | 801315.2187 | 1256338.274 | 858739.3353 |
| PC O-42:10 | 3085096.608 | 3820439.011 | 945441.6919 | 2502307.974 | 301223.3837 | 219513.7825 | 403627.9119 | 505332.9804 | 643676.1177 | 501273.9511 | 254630.8382 | 345796.2981 |
| CE 13:1 | 875845.7288 | 613756.0062 | 842984.7049 | 814278.0554 | 424894.3231 | 319606.0689 | 257180.9542 | 457892.22 | 258586.4 | 172197.2477 | 242399.8934 | 399097.0066 |
| LPI 20:5 | 1078062.18 | 823081.6061 | 695569.7632 | 924661.4188 | 578253.3046 | 170490.0479 | 355499.3469 | 397382.6783 | 593438.8063 | 465398.5246 | 400836.2415 | 92154.09943 |
| Tridecanoylcarnitine (Car(13:0)) | 315059.0507 | 230008.6635 | 360920.4518 | 301634.2129 | 2213158.868 | 1502005.001 | 2244540.778 | 1711541.905 | 1023427.784 | 0 | 1605796.459 | 767801.3586 |
| 17-Hydroxycalactin | 329662.9282 | 384921.3401 | 218810.6479 | 316726.5309 | 303547.2535 | 285960.4495 | 226882.4499 | 254038.3006 | 263157.4365 | 291858.0562 | 248183.6602 | 252301.8626 |
| 18.beta.-Glycyrrhetinic acid | 122505.4771 | 143232.6631 | 125609.898 | 108498.468 | 173582.0255 | 163997.4629 | 237087.6803 | 165832.8204 | 163135.619 | 115357.6967 | 206832.9689 | 164417.0818 |
| Glyco-beta-muricholic acid | 605272.2136 | 585046.7641 | 1186782.291 | 651428.2444 | 856531.2635 | 1000519.368 | 1122381.011 | 805869.3435 | 732857.0285 | 541718.018 | 598439.4587 | 714372.3657 |
| PA 22:4_22:4 | 208984. | 124199. | 147358. | 139246. | 105267. | 0 | 70531.3 | 100237. | 172555. | 166971. | 144217. | 105624. |

| **Metabolites** | **PBS_1** | **PBS_2** | **PBS_3** | **PBS_4** | **BCG_1** | **BCG_2** | **BCG_3** | **BCG_4** | **BCGΔ*pe18*_1** | **BCGΔ*pe18*_2** | **BCGΔ*pe18*_3** | **BCGΔ*pe18*_4** |
|---|---|---|---|---|---|---|---|---|---|---|---|---|
| | 7924 | 2216 | 2794 | 5362 | 606 | | 3938 | 2566 | 9174 | 7184 | 9504 | 7995 |
| Ala-Gly | 1681742.824 | 1688220.9 | 1981922.724 | 1739981.68 | 6489290.496 | 5483850.853 | 5875438.602 | 5765094.048 | 5931315.09 | 4731312.104 | 3871408.726 | 3999472.212 |
| M493T62 | 2087455.052 | 2201515.556 | 1029984.035 | 1454156.742 | 868196.0567 | 536498.0107 | 1451503.82 | 918174.1994 | 936051.0304 | 1430915.825 | 878545.4184 | 342492.6357 |
| NAOrn 11:0/18:0 | 11546.06544 | 65312.37616 | 20560.49567 | 36088.25468 | 0 | 12541.78343 | 32314.21763 | 41762.63019 | 32469.32033 | 49728.23156 | 0 | 43054.28282 |
| C75 trans (Drug) | 1428749.501 | 1496550.397 | 1472640.454 | 1770184.114 | 1472687.372 | 1522643.54 | 1426989.3 | 1460739.747 | 1517592.389 | 1538052.319 | 1383724.202 | 2117230.559 |
| ST 28:2;O;S | 79232.45547 | 132534.3809 | 62309.15842 | 57937.26731 | 98860.64707 | 149297.4621 | 74327.18201 | 105610.0896 | 106090.6219 | 96909.56185 | 105655.8427 | 31031.29741 |
| 1-Cyclopentyl-4-piperidinylamine | 2534148.468 | 2220216.924 | 2660560.99 | 2472426.986 | 2352503.989 | 1879846.868 | 2466193.619 | 2863367.578 | 2562286.159 | 2107869.849 | 2890589.824 | 2658832.124 |
| Thymopoietin II (32-34) | 835024.4088 | 1135445.844 | 1452765.603 | 3325853.689 | 962002.8749 | 690687.3157 | 899333.4083 | 603826.1347 | 703531.6364 | 1095536.032 | 1011264.179 | 2233646.784 |
| (+)-Norpseudoephedrine | 563257.2722 | 594772.2503 | 504943.8244 | 600171.0313 | 505098.6179 | 520915.7995 | 579544.4383 | 516777.062 | 494740.0718 | 487028.937 | 523109.0394 | 582045.0248 |
| 2-[(Dimethylamino)methyl]cyclohexanone | 1942836.715 | 1477007.563 | 1818911.202 | 1556938.727 | 1293762.715 | 1694178.618 | 1337276.092 | 1998929.084 | 1736256.78 | 1170713.895 | 1635506.061 | 1503409.092 |
| Benzophenone | 390568.4686 | 559589.8871 | 448505.5588 | 443657.7077 | 406583.6105 | 459188.1688 | 482846.9099 | 624211.937 | 485736.8615 | 409185.7377 | 353093.4955 | 381059.4323 |

| Metabolites | PBS_1 | PBS_2 | PBS_3 | PBS_4 | BCG_1 | BCG_2 | BCG_3 | BCG_4 | BCGΔ*pe18*_1 | BCGΔ*pe18*_2 | BCGΔ*pe18*_3 | BCGΔ*pe18*_4 |
|---|---|---|---|---|---|---|---|---|---|---|---|---|
| 6-Fluoro-3-methyl-1-benzothiophene-2-carboxylic acid | 1146286.282 | 1213342.636 | 1132446.052 | 1172444.73 | 1402006.992 | 1159281.465 | 1232557.006 | 1357480.908 | 1136730.522 | 1182104.776 | 1296309.423 | 1196029.456 |
| 3-Hydroxyoctanoic acid | 716486.2646 | 574860.1635 | 712320.1999 | 758454.5445 | 705392.0914 | 753633.2435 | 846331.5239 | 754515.1778 | 690746.3245 | 943313.4448 | 741314.9074 | 810168.7605 |
| DG 40:7 | 970080.4506 | 805156.1315 | 787961.5997 | 754142.2372 | 822361.6568 | 715794.9718 | 945826.4027 | 994657.646 | 846199.597 | 865891.9469 | 763860.4385 | 625311.5496 |
| PI 21:5 | 135865.3645 | 113236.4918 | 195334.6783 | 153309.3528 | 100380.9657 | 61125.32482 | 85617.28411 | 71748.13469 | 20002.33943 | 21408.7284 | 38546.14267 | 21459.51159 |
| NAE 13:1 | 3933699.166 | 4228127.328 | 5719459.265 | 4063378.191 | 3199282.679 | 3348281.471 | 3579586.378 | 3224939.79 | 3154649.209 | 3512497.616 | 3076304.227 | 5773083.902 |
| Cer 16:2;3O/13:1 | 553486.2817 | 638853.0609 | 469857.0806 | 523726.9206 | 484257.4067 | 470644.6932 | 497877.9231 | 543541.0303 | 534014.2386 | 492233.0266 | 460061.9561 | 475474.049 |
| 5-Amino-1-cyclopropyl-7-[(3R,5S)-3,5-dimethylpiperazin-1-yl]-6,8-difluoro-4-oxo-1,4-dihydroquinoline-3-carboxylic acid | 1398356.648 | 1638863.537 | 2094865.628 | 1181583.691 | 2297090.986 | 990983.7439 | 1083563.909 | 1350313.147 | 531210.8375 | 1223100.238 | 1951100.323 | 2564379.282 |
| Ginsenoyne A linoleate | 1489124.527 | 1753461.19 | 586813.6748 | 1245309.49 | 1040850.981 | 624538.1865 | 1028446.782 | 923826.3116 | 1280938.926 | 1235867.413 | 880216.5173 | 565861.0581 |
| Fexofenadine (Drug) | 132188.2573 | 102152.1128 | 110836.799 | 93396.73694 | 415998.1919 | 248305.103 | 471431.3261 | 338104.7737 | 254904.7001 | 302659.9815 | 365656.5557 | 194955.5687 |

| Metabolites | PBS_1 | PBS_2 | PBS_3 | PBS_4 | BCG_1 | BCG_2 | BCG_3 | BCG_4 | BCGΔ*pe18*_1 | BCGΔ*pe18*_2 | BCGΔ*pe18*_3 | BCGΔ*pe18*_4 |
|---|---|---|---|---|---|---|---|---|---|---|---|---|
| PI 18:1_22:5 | 744035.1782 | 778040.2655 | 347126.4472 | 783678.6404 | 1195781.346 | 784643.8368 | 1023165.201 | 1053443.771 | 1333760.446 | 1053131.98 | 130460.2819 | 968454.8673 |
| NAOrn 16:0/20:3 | 226479.7809 | 395373.2921 | 171774.8285 | 299350.6315 | 413712.1339 | 280222.729 | 343051.5562 | 307215.0381 | 344098.0152 | 301421.6863 | 342413.4014 | 304079.8813 |
| 1-Methylpseudouridine | 609048.7666 | 446653.9024 | 790216.8466 | 585455.6279 | 1566098.017 | 1125132.784 | 1605432.605 | 1299366.23 | 1110772.405 | 1231126.339 | 1398364.249 | 931945.574 |
| Cer 10:0;2O/24:6 | 607969.0173 | 770258.6985 | 684033.3424 | 944788.7966 | 2974504.612 | 2339668.593 | 1248128.592 | 2303836.072 | 1651415.012 | 593145.1446 | 519076.4345 | 1629475.041 |
| 2-16:0-lysophosphatidylglycerol | 131752.7964 | 85338.20726 | 139234.1054 | 160443.9766 | 71178.70972 | 72166.96993 | 79080.68655 | 111950.2063 | 71525.69458 | 66033.89525 | 79411.83571 | 0 |
| Procyanidin B2 | 0 | 381191.6947 | 40358.20325 | 328511.6652 | 82184.98529 | 128925.6267 | 243055.4585 | 303976.1031 | 297447.9179 | 163710.6264 | 0 | 222984.4196 |
| H-Val-Tyr-OH | 335274.6323 | 307125.2931 | 458417.2624 | 369078.7618 | 655311.8989 | 499500.3396 | 764010.8816 | 621907.0259 | 605089.5467 | 645562.9941 | 743645.5429 | 431726.1107 |
| PE-Cer 13:0;2O/12:0;O | 6669.134836 | 71008.08049 | 5513.809371 | 57057.30659 | 43244.29314 | 26254.0504 | 104907.6794 | 10611.32656 | 26001.09854 | 28440.67771 | 5590.332332 | 0 |
| PA 2:0_18:2 | 807212.0699 | 761441.4344 | 656438.7367 | 684978.4046 | 741836.7895 | 734890.4272 | 757410.6854 | 662552.2095 | 689822.4562 | 725965.9104 | 713187.535 | 661267.1371 |
| Altretamine | 1226970.846 | 578396.1659 | 716752.4192 | 502489.1075 | 744964.1577 | 644909.4842 | 673110.7419 | 446214.0797 | 849775.6041 | 654706.552 | 897378.948 | 668550.1575 |
| 7-Bromo-2,3-dihydrothieno[3,4-b][1,4]dioxine-5-carbaldehyde | 2377401.052 | 2217967.405 | 2547246.976 | 2570908.422 | 2226862.131 | 2433287.325 | 2448710.161 | 2392667.058 | 2216114.651 | 2501117.86 | 2511599.014 | 2484443.474 |

| Metabolites | PBS_1 | PBS_2 | PBS_3 | PBS_4 | BCG_1 | BCG_2 | BCG_3 | BCG_4 | BCGΔ*pe18*_1 | BCGΔ*pe18*_2 | BCGΔ*pe18*_3 | BCGΔ*pe18*_4 |
|---|---|---|---|---|---|---|---|---|---|---|---|---|
| Cer 18:2;2O/16:0 | 391570.4693 | 477627.9621 | 665740.701 | 492700.7055 | 365514.8503 | 739235.0915 | 409462.5266 | 429005.0025 | 391222.6697 | 246868.4464 | 380689.0228 | 432418.4542 |
| (indol-3-yl)acetyl-valine | 2400051.761 | 542158.6185 | 1643004.697 | 1831004.795 | 1358104.552 | 2915654.121 | 41795794.91 | 5052847.297 | 7438760.208 | 3626341.986 | 2363268.797 | 3824584.46 |
| 2-Hexadec-7-enyloctadecanedioic acid | 259582.6062 | 159424.6429 | 116988.4675 | 102644.854 | 129164.2067 | 99779.15386 | 183927.7887 | 87364.08283 | 299624.2731 | 86887.23908 | 104465.2136 | 66548.56304 |
| 2,6-Dimethyl-4-hydroxybenzaldehyde | 469357.3227 | 411192.0641 | 540137.8528 | 464280.5025 | 565449.2124 | 521610.515 | 447724.3909 | 518253.8203 | 518079.5871 | 312234.0585 | 393581.8531 | 486326.1621 |
| Tauro-beta-muricholic acid | 1096547.017 | 1077546.653 | 1824525.352 | 1223309.042 | 1558784.851 | 1885646.717 | 1765456.324 | 1393907.031 | 1372841.067 | 895441.921 | 933016.971 | 1372264.349 |
| 4-{(4E)-4-[(3-Methylphenyl)imino]-3,4-dihydro-1-phthalazinyl}benzamide | 1115250.679 | 953024.8753 | 821242.0552 | 958387.8115 | 822316.6945 | 758940.8407 | 956201.4045 | 1121615.437 | 795649.131 | 931981.854 | 827494.8481 | 1047806.344 |
| 1-Chloro-3-fluoro-5-(4-nitrophenyl)benzene | 48311.76065 | 91886.53492 | 35770.08689 | 52692.6525 | 121750.4184 | 31175.51127 | 150776.1435 | 91799.55158 | 20117.2155 | 59214.75396 | 30557.22205 | 49420.18389 |
| Platencin | 4844834.513 | 4585796.502 | 728635.9021 | 3966423.147 | 5035038.716 | 4191810.272 | 4082943.093 | 5777381.68 | 4293398.254 | 4465898.358 | 4087388.516 | 4484338.013 |
| 10-Hydroperoxy-12-[6-[pent-2-en-1-yl]-2,3-dioxabicyclo[2.2.1]heptan-5-yl]dodeca-4,7,11-trienoyl | 171472.5901 | 201839.2903 | 175316.568 | 184077.7567 | 125852.1412 | 175115.2221 | 178110.1714 | 164753.2715 | 176237.2685 | 147115.8551 | 171691.371 | 180013.4676 |

| Metabolites | PBS_1 | PBS_2 | PBS_3 | PBS_4 | BCG_1 | BCG_2 | BCG_3 | BCG_4 | BCGΔ *pe18*_1 | BCGΔ *pe18*_2 | BCGΔ *pe18*_3 | BCGΔ*pe18*_4 |
|---|---|---|---|---|---|---|---|---|---|---|---|---|
| carnitine | | | | | | | | | | | | |
| astilbin | 548586.1982 | 464054.8707 | 586805.444 | 464190.5134 | 756043.4785 | 685443.433 | 633693.5797 | 562618.7082 | 660811.6595 | 607363.9479 | 604981.8193 | 614148.0846 |
| SL 10:0;O/21:0 | 25212.39986 | 36748.14189 | 72430.28928 | 22962.09512 | 21397.96842 | 14514.02285 | 47057.35631 | 21554.37113 | 19407.48611 | 21831.71049 | 32881.45349 | 7917.102003 |
| MGDG 3:0_13:1 | 266200.2928 | 415894.9778 | 136927.6043 | 180544.0779 | 75810.1647 | 476266.7574 | 94425.34193 | 101277.8594 | 48077.69494 | 50644.30133 | 99553.79745 | 48499.87985 |
| 5-[(Benzylsulfanyl)methyl]-2-furoic acid | 8422027.233 | 6611556.288 | 9500209.219 | 5104534.11 | 10753219.69 | 8976655.321 | 8813379.197 | 7696495.463 | 9373978.267 | 7007783.85 | 6961254.269 | 3169620.93 |
| tert-Butyl 3-(aminomethyl)piperazine-1-carboxylate | 149439.6188 | 746694.2047 | 212290.8729 | 628854.6564 | 1305579.435 | 1392271.3 | 106810.8363 | 113925.484 | 653903.9353 | 526532.6313 | 558276.4296 | 433590.853 |
| LDGCC 14:0 | 351692.9376 | 309201.3174 | 288843.6715 | 361756.9609 | 342173.5899 | 386760.605 | 343842.027 | 314979.9524 | 349038.696 | 374511.8093 | 319225.344 | 222592.5669 |
| 7-[(6-hydroxy-5,5,8a-trimethyl-2-methylidene-3,4,4a,6,7,8-hexahydro-1H-naphthalen-1-yl)methox | 313528.1758 | 300423.7303 | 385601.6733 | 290834.6779 | 1469390.175 | 724238.7324 | 1352356.32 | 863038.7915 | 808989.9981 | 883466.5235 | 985754.6734 | 702135.5778 |

| Metabolites | PBS_1 | PBS_2 | PBS_3 | PBS_4 | BCG_1 | BCG_2 | BCG_3 | BCG_4 | BCGΔ *pe18*_1 | BCGΔ *pe18*_2 | BCGΔ *pe18*_3 | BCGΔ*pe18*_4 |
|---|---|---|---|---|---|---|---|---|---|---|---|---|
| y]chromen-2-one | | | | | | | | | | | | |
| 1-Phenyl-2,3,4,5-tetrahydro-1H-3-benzazepine | 888219.9376 | 913838.9734 | 849026.733 | 963310.0274 | 1001518.611 | 853544.0547 | 967186.6472 | 994435.8011 | 952918.2847 | 888135.5331 | 922907.0871 | 413918.2645 |
| LNAPS 3:0/N-22:2 | 153330.6891 | 143480.7664 | 153369.8801 | 168301.3918 | 248221.2908 | 254144.1043 | 253455.8336 | 273366.8101 | 242186.6098 | 139067.686 | 171386.0751 | 209686.27 |
| Cytidylyl-(5'-3')-5'-inosinic acid homopolymer | 4402.00888 | 0 | 3935.716964 | 0 | 70408.7991 | 23048.35687 | 46904.17221 | 29361.00164 | 46570.50242 | 24282.01564 | 86724.03816 | 4527.516731 |
| Glu-Arg | 360048.0865 | 268604.0871 | 436382.7727 | 238680.0615 | 631279.8009 | 693553.8419 | 992695.2299 | 508020.7904 | 563716.1446 | 635048.7283 | 664899.0727 | 878446.5187 |
| PC(37:2) | 1348034.467 | 2048677.851 | 1912715.79 | 1254386.719 | 20105245.61 | 14839827.3 | 19393209.41 | 16694469.07 | 19350876.25 | 20967083.61 | 16867688.88 | 9914323.052 |
| LPE 16:2 | 25821.80749 | 19580.66378 | 36103.16618 | 26144.37358 | 150631.8803 | 131226.785 | 125754.8254 | 59942.51756 | 146835.9171 | 158111.5354 | 183083.3884 | 151251.6874 |
| Tetradecyl sulfate (sodium) | 5385587.132 | 1312015.103 | 2181221.959 | 2290323.396 | 2643176.868 | 2194144.689 | 1769387.196 | 2084645.692 | 1972050.791 | 1791358.168 | 2012555.219 | 1388248.214 |
| Hygrine | 2173243.65 | 6165395.994 | 3052006.94 | 2115136.392 | 2496687.764 | 3218115.08 | 2756384.586 | 3043857.314 | 2590906.615 | 2794417.259 | 2350078.431 | 2759959.646 |
| PG(16:1(9Z)/22:6(4Z,7Z,10Z,13Z,16Z, | 141347.9707 | 259607.2508 | 0 | 85881.25356 | 1071777.508 | 810702.882 | 1148983.302 | 1049735.995 | 1100624.588 | 1141604.632 | 991063.0454 | 782184.9957 |

| Metabolites | PBS_1 | PBS_2 | PBS_3 | PBS_4 | BCG_1 | BCG_2 | BCG_3 | BCG_4 | BCGΔ *pe18*_1 | BCGΔ *pe18*_2 | BCGΔ *pe18*_3 | BCGΔ*pe18*_4 |
|---|---|---|---|---|---|---|---|---|---|---|---|---|
| 19Z)) | | | | | | | | | | | | |
| 6H-Dibenzo[b,d]pyran, 3-(1,1-dimethylbutyl)-6a,7,10,10a-tetrahydro-6,6,9-trimethyl-, (6aR,10aR)- | 691109.8984 | 825038.6676 | 288087.5256 | 2502704.782 | 317987.6906 | 652370.7663 | 3572460.416 | 193379.7715 | 2011882.203 | 873961.4658 | 413947.586 | 2918480.775 |
| Pentadecanoylcarnitine | 1479715.159 | 1330160.274 | 1027550.538 | 1254947.493 | 1388558.693 | 1371648.078 | 1375569.936 | 1314304.466 | 1268586.142 | 1358786.654 | 1307981.163 | 1075358.341 |
| LNAPS 18:0/N-18:2 | 678595.4525 | 578694.0875 | 760269.9071 | 877725.2386 | 846712.64 | 1107866.521 | 680808.8014 | 553280.2492 | 979900.8736 | 700375.9374 | 215664.6073 | 2227357.075 |
| M434T301 | 182529.4804 | 183302.0505 | 131858.9226 | 160624.9357 | 176183.9365 | 163550.4749 | 146759.7536 | 187847.9714 | 32932.34459 | 160871.4367 | 192074.3614 | 157638.9884 |
| Formylanthranilic acid | 1285785.17 | 1613948.503 | 1747147.466 | 1961066.971 | 1412330.312 | 1134860.534 | 1488548.409 | 1092076.919 | 1108850.877 | 1219576.447 | 1071991.736 | 3006286.892 |
| Tetradecenedioylcarnitine (Car(14:1-O2)) | 34129.26246 | 7363.42501 | 27309.88373 | 20497.44857 | 279547.1315 | 190045.4201 | 279490.0101 | 205358.4205 | 141316.3125 | 233206.0282 | 313001.2473 | 122598.23 |
| Cer 13:2;2O/16:4 | 387408.4219 | 359937.5622 | 335856.3633 | 379145.0598 | 334701.0002 | 359989.7518 | 377222.7664 | 325202.2184 | 356703.6694 | 378030.9554 | 331936.7576 | 363329.2528 |
| 1-Propanesulfonamide, N-[3-[(5-chloro-1H | 40270.4411 | 6214.548377 | 39660.82655 | 31404.50907 | 113300.966 | 23281.04684 | 86861.19272 | 29509.67127 | 18934.62891 | 100921.4428 | 117429.8549 | 52751.08769 |

| Metabolites | PBS_1 | PBS_2 | PBS_3 | PBS_4 | BCG_1 | BCG_2 | BCG_3 | BCG_4 | BCGΔ *pe18*_1 | BCGΔ *pe18*_2 | BCGΔ *pe18*_3 | BCGΔ*pe18*_4 |
|---|---|---|---|---|---|---|---|---|---|---|---|---|
| -pyrrolo[2,3-b]pyridin-3-yl)carbonyl]-2,4-difluorophenyl]- | | | | | | | | | | | | |
| PC O-22:6_16:0 | 177446.8442 | 166676.4129 | 133330.8224 | 159752.965 | 102398.7989 | 194942.3959 | 92296.7 | 231680.8329 | 107846.2093 | 52210.18537 | 65359.43765 | 120501.8981 |
| Obatoclax | 505650.9587 | 573654.2968 | 494878.6335 | 454508.6214 | 3672435.33 | 459937.3487 | 3678425.289 | 429245.4713 | 491418.1991 | 511640.853 | 514399.768 | 465521.2174 |
| 4-(Cyclohexyloxy)benzoic acid | 151388.408 | 212509.712 | 184076.9597 | 163510.2386 | 174016.3678 | 92661.41564 | 239472.653 | 138659.358 | 171709.3406 | 170599.0815 | 149588.4189 | 110811.1358 |
| PE 18:1_22:6;2O | 1685911.085 | 2964047.387 | 3853018.162 | 1803009.995 | 303611.3828 | 212424.6495 | 209632.9802 | 88712.26639 | 250573.2375 | 186265.9756 | 146515.7453 | 1120047.869 |
| PE(20:1(11Z)/14:1(9Z)) | 42971.19829 | 44367.94983 | 61963.12315 | 24555.7052 | 758330.1665 | 813124.7597 | 1221749.212 | 1735305.598 | 877227.9333 | 1954990.476 | 771764.1594 | 626699.8658 |
| 7-Aminoflunitrazepam | 6121773.583 | 3590095.448 | 8994516.472 | 7794577.861 | 11842438.3 | 6719083.339 | 10675876.5 | 6285054.653 | 7073971.143 | 4582483.81 | 2746765.271 | 3357842.944 |
| Artocarpin | 129301.5029 | 143106.9738 | 131659.8955 | 98350.80166 | 90077.56672 | 108628.0054 | 114928.1177 | 115377.1549 | 135288.2111 | 135290.3407 | 88906.30686 | 177988.9838 |
| N8,N8,3-Trimethyl-2,8-phenazinediamine | 1069453.935 | 1293442.364 | 1348287.171 | 1188020.953 | 1031198.652 | 980277.5157 | 1127874.457 | 1272145.48 | 1187287.192 | 1283244.585 | 983969.3368 | 1228246.069 |
| LPE O-19:0 | 233973.1167 | 273888.3104 | 187372.9233 | 208704.7882 | 198860.7827 | 171745.4579 | 221882.06 | 256693.0609 | 237368.764 | 238309.0552 | 216821.0446 | 309014.6194 |
| 2-Methyl-4,6-dinitr | 859013. | 772413. | 788903. | 134791 | 602955. | 914245. | 626998. | 638048. | 647341. | 791842. | 664106. | 931695. |

| **Metabolites** | **PBS_1** | **PBS_2** | **PBS_3** | **PBS_4** | **BCG_1** | **BCG_2** | **BCG_3** | **BCG_4** | **BCGΔ*pe18*_1** | **BCGΔ*pe18*_2** | **BCGΔ*pe18*_3** | **BCGΔ*pe18*_4** |
|---|---|---|---|---|---|---|---|---|---|---|---|---|
| ophenol | 2318 | 1935 | 4403 | 5.328 | 2592 | 0464 | 146 | 929 | 799 | 4242 | 2686 | 3472 |
| Tetrahydrocannabivarin | 977719.2416 | 1234762.211 | 1635008.181 | 1785486.968 | 789314.7588 | 1268731.565 | 992475.6929 | 797757.2391 | 779392.2415 | 884342.6851 | 778086.7117 | 1362929.125 |
| Cer 14:3;2O/18:4 | 1105248.199 | 631870.235 | 561124.313 | 1710293.085 | 1867626.911 | 1028908.284 | 922055.2875 | 678153.732 | 913693.5792 | 1194348.098 | 836639.6522 | 692547.2802 |
| Lyciumoside_IX | 2152723.479 | 2231657.546 | 2177001.877 | 2135239.911 | 2153268.645 | 2184494.884 | 2225236.622 | 2168812.88 | 2275294.782 | 2143510.865 | 2091915.314 | 2139910.723 |
| 2-Hydroxypalmitic acid | 1043677.063 | 1190293.588 | 1606037.1 | 1429955.418 | 949696.5125 | 1651472.339 | 1574368.778 | 1419741.749 | 894557.4416 | 1674299.17 | 1009250.137 | 2276932.375 |
| 2-Methyl-4-(2-thienyl)-3-butyn-2-ol | 461450.932 | 280686.96 | 690468.0188 | 495116.9365 | 517448.7504 | 516347.6349 | 408147.8903 | 264078.5793 | 260802.066 | 294644.4163 | 325027.5969 | 254092.5144 |
| PI 14:1_20:3 | 156437.1441 | 228386.8447 | 35042.27375 | 106122.0668 | 156458.3777 | 184753.4945 | 235532.5872 | 169398.0131 | 247307.6657 | 233659.1108 | 158188.3747 | 141199.3851 |
| 11-Oxatetradecanoic acid | 355048.6956 | 352083.6813 | 500255.3503 | 358736.2603 | 1309015.581 | 2501038.086 | 1445593.873 | 1658462.407 | 1222889.202 | 902664.7385 | 1441877.435 | 2275771.941 |
| Sodium lauryl sulfoacetate | 1309352.608 | 2647937.703 | 3533910.202 | 3445067.555 | 1386433.808 | 1172909.35 | 2070532.756 | 1090512.901 | 1847326.932 | 2523057.211 | 1910194.46 | 6765041.417 |
| (3.beta.)-Allopregnanolone sulfate | 198271.5116 | 244754.8748 | 404240.5711 | 363903.5632 | 538300.8147 | 467826.5781 | 836996.7273 | 537181.792 | 427219.2764 | 465205.0967 | 399438.4062 | 429453.9375 |
| Arg-Glu | 4318968.084 | 4272103.854 | 4085415.595 | 4702543.758 | 3806251.26 | 4399573.355 | 4449027.526 | 4671530.004 | 4200999.066 | 4124183.163 | 4710589.569 | 3870748.912 |
| (4E,6E)-3-(4-Formylanilino)-8-hydroxyocta-4,6-dienoic | 254884.3542 | 33306.95977 | 274191.505 | 45802.41391 | 237189.0734 | 410649.6123 | 2037335.753 | 1429984.805 | 565939.5799 | 254158.954 | 451279.9667 | 150898.5677 |

| Metabolites | PBS_1 | PBS_2 | PBS_3 | PBS_4 | BCG_1 | BCG_2 | BCG_3 | BCG_4 | BCGΔ*pe18*_1 | BCGΔ*pe18*_2 | BCGΔ*pe18*_3 | BCGΔ*pe18*_4 |
|---|---|---|---|---|---|---|---|---|---|---|---|---|
| acid | | | | | | | | | | | | |
| Man2a | 15745.31785 | 36482.21374 | 27552.64872 | 28689.48358 | 187165.9596 | 96896.03671 | 148146.374 | 116356.4794 | 125675.3303 | 109287.6538 | 172746.3979 | 146592.6939 |
| P1,P4-Bis(5'-uridyl) tetraphosphate | 0 | 0 | 15211.66725 | 11865.97379 | 87111.5876 | 171299.0616 | 126760.4585 | 182008.8494 | 160454.4283 | 75555.50167 | 138507.4031 | 193076.3533 |
| PE O-10:0_4:0 | 239507.8944 | 94452.55094 | 340991.3191 | 221267.0568 | 2086528.233 | 291893.3657 | 1985215.488 | 1634846.439 | 2312024.256 | 1998036.976 | 2681960.328 | 1759737.516 |
| Hex3Cer 19:0;2O | 17718.75994 | 69653.96609 | 58938.51269 | 42632.02468 | 45087.19283 | 4953.858849 | 53445.14469 | 38319.54461 | 11649.87086 | 34844.26086 | 76916.90709 | 200671.4146 |
| SM 33:2;3O | 865251.9513 | 999402.8499 | 884186.292 | 1108606.606 | 944033.6003 | 939385.1446 | 897456.2855 | 875114.0433 | 887438.6918 | 681460.9024 | 834010.8393 | 1126955.134 |
| N-(2-Chlorophenyl)-4-fluorobenzenesulfonamide | 1124803.018 | 1009705.364 | 1309894.365 | 1135953.813 | 1860785.185 | 1407370.443 | 1937470.354 | 1585969.927 | 1434610.459 | 1609242.319 | 1779046.766 | 1245801.368 |
| 4-(7-Hydroxy-3-(hydroxymethyl)-5-(3-hydroxypropyl)-2,3-dihydro-1-benzofuran-2-yl)-2-methoxyphenyl .beta.-D-xylopyranoside | 0 | 0 | 0 | 0 | 288409.0492 | 198959.8086 | 210889.4712 | 162618.9731 | 233167.4843 | 222780.9333 | 250630.8881 | 204258.4806 |
| 3-Hydroxyoleylcarn | 254623. | 235247. | 403571. | 239581. | 111244 | 808122. | 176507 | 38580.2 | 262762. | 0 | 682761. | 492858. |

| Metabolites | PBS_1 | PBS_2 | PBS_3 | PBS_4 | BCG_1 | BCG_2 | BCG_3 | BCG_4 | BCGΔ*pe18*_1 | BCGΔ*pe18*_2 | BCGΔ*pe18*_3 | BCGΔ*pe18*_4 |
|---|---|---|---|---|---|---|---|---|---|---|---|---|
| itine | 5181 | 7737 | 9797 | 2805 | 0.98 | 3613 | 9.668 | 6258 | 4345 | | 7672 | 2769 |
| Car(18:1) | 32015359.63 | 23606256.08 | 41490628.74 | 29645334.02 | 64271694.4 | 43276879.24 | 66974154.12 | 58065387.34 | 33548167.78 | 45575893.48 | 45189903.09 | 24762215.47 |
| Mannose | 3495504.439 | 2460028.514 | 3371648.778 | 2906362.522 | 1496898.451 | 3543713.359 | 3170554.618 | 2080247.96 | 3142979.894 | 3371011.312 | 3535382.167 | 2574971.467 |
| N-Arachidonoyldopamine | 459287.6207 | 306916.1367 | 354521.5779 | 318989.6961 | 334493.0019 | 405073.8875 | 385559.7215 | 440110.5054 | 393634.0369 | 366989.3499 | 366685.866 | 292706.4879 |
| PI O-17:0_26:5 | 54200.66614 | 64443.42198 | 66894.74171 | 55471.10816 | 88328.74011 | 104790.0393 | 61127.0119 | 60909.62856 | 56116.3186 | 87992.57743 | 34016.66623 | 68583.53935 |
| 2,4-Dodecadienamide, N-(p-hydroxyphenethyl)-, (E,E)- | 6802217.633 | 5448985.042 | 4461259.997 | 4133766.218 | 4899971.557 | 5058471.593 | 4457450.3 | 4871306.127 | 4868455.387 | 5017953.298 | 4975745.013 | 4308146.75 |
| LPE O-20:1 | 567865.1921 | 922167.7899 | 352601.3986 | 407658.4696 | 184945.5927 | 421353.1522 | 222793.6186 | 240877.7125 | 148004.2379 | 176996.1221 | 259397.6178 | 116198.2041 |
| PC(38:6) | 10729707.78 | 13241779.56 | 4478605.426 | 7229389.696 | 1174809.62 | 1022630.485 | 1418918.542 | 1401428.005 | 1490322.293 | 2263885.931 | 1404094.22 | 2274442.52 |
| Piperonylic acid | 2705278.709 | 2490302.98 | 3180872.188 | 3472120.186 | 2030782.147 | 2651225.994 | 2762232.86 | 2514550.739 | 2371968.549 | 4610884.596 | 2391723.018 | 2552119.269 |
| Kifunensine | 1908654.24 | 1656496.944 | 1360561.451 | 1548140.719 | 1903377.424 | 1031380.122 | 1298448.041 | 823062.6603 | 3420510.912 | 1435318.374 | 1628954.45 | 1707152.277 |
| Actarit | 3054082.912 | 1486133.992 | 2787402.855 | 2359268.299 | 1932863.841 | 2258858.908 | 2087312.072 | 1257946.901 | 1206958.341 | 2211616.432 | 2190713.549 | 1210980.617 |
| Ciprostene | 76376.0 | 72667.1 | 77677.2 | 67377.0 | 220903. | 700451. | 155252. | 275367. | 115254. | 106688. | 271769. | 268829. |

| **Metabolites** | **PBS_1** | **PBS_2** | **PBS_3** | **PBS_4** | **BCG_1** | **BCG_2** | **BCG_3** | **BCG_4** | **BCGΔ *pe18*_1** | **BCGΔ *pe18*_2** | **BCGΔ *pe18*_3** | **BCGΔ*pe18*_4** |
|---|---|---|---|---|---|---|---|---|---|---|---|---|
| | 9073 | 7244 | 3916 | 1463 | 8135 | 676 | 5452 | 4073 | 5651 | 8504 | 665 | 1839 |
| N1-ACETYLSPERMINE | 377701.029 | 197693.1957 | 237188.5596 | 144264.1267 | 150414.1475 | 153953.1592 | 475096.5835 | 431011.1113 | 424172.4606 | 394143.6318 | 308756.9305 | 384480.0928 |
| Annocherin_A | 385251.0962 | 361471.1174 | 681655.604 | 402645.1638 | 111967.9005 | 173529.1416 | 246391.1331 | 274715.8123 | 173527.3741 | 177605.8431 | 0 | 82151.8731 |
| Wogonoside | 370254.5101 | 488347.6411 | 396363.3634 | 427389.7265 | 246585.1534 | 295463.4746 | 437965.3513 | 330667.8944 | 1955451.986 | 375427.4239 | 358790.0155 | 342171.949 |
| PI 29:1 | 269200.1979 | 279959.3454 | 203723.5568 | 226843.7439 | 217290.2149 | 210329.5621 | 243927.2794 | 214383.9181 | 299974.6437 | 251373.2199 | 292286.7852 | 258522.1065 |
| Dye X-6012-7A | 151917.8835 | 159581.4361 | 143247.5647 | 140710.0907 | 157824.2445 | 143805.5595 | 151719.2749 | 145062.7568 | 167510.243 | 123088.4716 | 147056.7672 | 140933.9623 |
| HexCer 9:0;2O/32:9 | 1328212.619 | 1241982.22 | 2603373.386 | 2086750.521 | 2724854.922 | 2547719.943 | 2439050.348 | 3074663.855 | 2857148.834 | 3081992.324 | 2473413.947 | 1181361.353 |
| PE(20:4(5Z,8Z,11Z,14Z)/16:0) | 310972.9224 | 509628.8579 | 655889.6266 | 265413.689 | 2325308.503 | 1208251.542 | 2075441.399 | 1796284.185 | 1650634.45 | 2409261.336 | 1561634.258 | 1229036.663 |
| PC(16:1(9Z)/14:0) | 387813.9564 | 415661.952 | 282269.636 | 182336.9705 | 2660594.497 | 1817832.88 | 3080170.519 | 2853791.164 | 2564592.03 | 4532621.345 | 2492617.78 | 1599720.308 |
| Andrographiside | 43653.03206 | 50689.74736 | 48560.8843 | 51013.90866 | 48703.47612 | 49497.48694 | 52018.09275 | 51098.1362 | 76234.87596 | 46503.45694 | 56015.13681 | 43715.57498 |
| SHexCer 23:1;3O | 203791.7062 | 271624.0614 | 186917.1796 | 260527.3422 | 184910.9172 | 208689.8749 | 153992.0786 | 123622.8812 | 150579.4911 | 106334.4748 | 160578.2873 | 77369.45216 |
| Propanoyl-CoA | 0 | 0 | 0 | 0 | 101268.3445 | 81066.05431 | 49380.51563 | 58242.94099 | 69271.1649 | 62582.31099 | 51734.74918 | 77940.99905 |
| Diphenyl Phosphate | 123269 | 594467 | 121555 | 902863 | 226811 | 810248. | 285813 | 481943. | 242159 | 436960 | 254059 | 870496 |

| Metabolites | PBS_1 | PBS_2 | PBS_3 | PBS_4 | BCG_1 | BCG_2 | BCG_3 | BCG_4 | BCGΔ*pe18*_1 | BCGΔ*pe18*_2 | BCGΔ*pe18*_3 | BCGΔ*pe18*_4 |
|---|---|---|---|---|---|---|---|---|---|---|---|---|
| | 0.749 | 3.659 | 38.55 | 7.508 | 7.97 | 6095 | 7.705 | 3542 | 2.908 | 4.988 | 0.324 | 6.141 |
| SMGDG O-26:3_10:0 | 25233.7471 | 0 | 36303.29183 | 19783.50807 | 93532.21785 | 202407.4143 | 116781.1438 | 132488.2822 | 29148.81708 | 18860.14941 | 106081.4508 | 108299.5557 |
| 2-[6-(3,4-Dihydroxy-2-methylidenebutanoyl)oxy-4-ethenyl-2-hydroxy-3-(3-hydroxyprop-1-en-2-yl)-4-methylcyclohexyl]prop-2-enoic acid | 484915.8971 | 432309.9881 | 322293.5153 | 447191.017 | 477706.9773 | 381208.9602 | 539021.2825 | 391484.5641 | 470305.66 | 427330.5089 | 425440.3732 | 517495.6385 |
| 3-Hydroxyurs-12-en-23-oic acid | 940274.4701 | 2286053.56 | 1747449.448 | 2961211.811 | 1528414.662 | 2572631.552 | 2754264.629 | 1677560.847 | 2843542.477 | 1609861.045 | 1396663.908 | 1099516.435 |
| LPC(O-20:1) | 156407.9255 | 166828.6901 | 85178.86364 | 105745.1636 | 141930.6232 | 110221.3345 | 192173.755 | 65487.55383 | 107202.3664 | 191729.714 | 147717.8639 | 88169.21123 |
| 11S,15S-dihydroxy-14R-(S-glutathionyl)-5Z,8Z,12E-eicosatrienoic acid | 41179.2929 | 54020.23971 | 126750.139 | 316022.6405 | 0 | 30452.95826 | 34956.19516 | 32499.48036 | 8877.08616 | 7000.426051 | 8146.261813 | 15866.42403 |
| N,N-Dimethyltryptamine | 1014561.292 | 1425232.409 | 643880.8061 | 824691.276 | 587321.6188 | 1441334.443 | 1187829.722 | 822438.7092 | 565975.0992 | 567266.0011 | 875280.6022 | 1804589.371 |
| beta-Muricholic acid | 390744.7545 | 222693.71 | 337763.8528 | 269728.6292 | 384166.0987 | 522088.6098 | 344759.558 | 372599.0815 | 262691.4746 | 269310.8861 | 566134.897 | 295428.0458 |
| linoleyl-AMP | 45938.7 | 30415.5 | 39253.1 | 41881.1 | 173083. | 120907. | 68046.0 | 84606.0 | 103468. | 163991. | 94398.1 | 133149. |

| **Metabolites** | **PBS_1** | **PBS_2** | **PBS_3** | **PBS_4** | **BCG_1** | **BCG_2** | **BCG_3** | **BCG_4** | **BCGΔ *pe18*_1** | **BCGΔ *pe18*_2** | **BCGΔ *pe18*_3** | **BCGΔ*p e18*_4** |
|---|---|---|---|---|---|---|---|---|---|---|---|---|
| | 9205 | 5694 | 0169 | 5959 | 9903 | 366 | 5232 | 4375 | 9269 | 8629 | 5161 | 0723 |
| 5-Bromo-2-hydroxy benzamide | 115451 4.683 | 127447 5.412 | 195061 8.079 | 124363 1.417 | 113834 7.627 | 116015 5.094 | 100086 6.137 | 120321 6.153 | 114940 0.488 | 117497 0.025 | 104746 7.463 | 134712 5.442 |
| Rhodopin | 107059 5.832 | 125889 2.37 | 435055. 0981 | 882446. 851 | 560216. 5901 | 452434. 4532 | 529354. 4282 | 593390. 423 | 668341. 9123 | 759458. 2419 | 529333. 7593 | 462238. 7134 |
| 20-Hydroxy-N-arac hidonoyltaurine | 35789.2 9786 | 83539.9 9995 | 70266.1 8531 | 46280.2 7463 | 219918. 9754 | 146438. 3251 | 259309. 8483 | 173929. 1906 | 375486. 9928 | 213698. 5808 | 218715. 4207 | 146242. 3093 |
| PS 20:3 | 0 | 0 | 0 | 0 | 907198. 7416 | 534963 5.67 | 128561 0.644 | 583869. 6654 | 138113. 5645 | 104206. 9632 | 815144. 8616 | 359481. 4404 |
| N6,N6,N6-Trimeth yllysine | 112642 2.759 | 118008 1.387 | 122130 6.989 | 116362 9.76 | 875404. 2446 | 688006. 7774 | 189746 7.024 | 938337. 5093 | 767597. 1564 | 130522 0.305 | 957984. 519 | 139218 1.506 |
| 7-(Acetyloxy)-2',2', 3,6,11a-pentamethy l-8-methylidene-1,4 ,6',12-tetraoxohexa hydro-4'H,10H-spir o[3a,7a-epoxy-6,11 b-methanofuro[3,4- e][3]benzoxocine-9, 3'-pyran]-11-yl (2E)-2-methylbut-2 -enoate | 0 | 0 | 0 | 3589.93 1315 | 146032. 2041 | 214887. 5874 | 80377.7 2685 | 117825. 33 | 35337.6 8429 | 36579.9 8116 | 10517.7 0443 | 60442.4 0987 |

| Metabolites | PBS_1 | PBS_2 | PBS_3 | PBS_4 | BCG_1 | BCG_2 | BCG_3 | BCG_4 | BCGΔ*pe18*_1 | BCGΔ*pe18*_2 | BCGΔ*pe18*_3 | BCGΔ*pe18*_4 |
|---|---|---|---|---|---|---|---|---|---|---|---|---|
| 1,2-Dipalmitoylphosphatidylinositol-4,5-diphosphonate | 480089.2875 | 606199.7417 | 173518.0921 | 573667.6259 | 57911.70034 | 89187.10079 | 67682.95951 | 95678.77164 | 176654.1556 | 68484.25085 | 74225.43594 | 92785.46484 |
| Docosanoylcarnitine | 170043.5235 | 178114.3569 | 124437.8873 | 160966.5034 | 167523.3232 | 157150.3195 | 164373.0028 | 133456.565 | 174626.9921 | 182198.0452 | 154381.0955 | 109696.6713 |
| Ikarugamycin | 56425.40809 | 41041.17291 | 91968.29919 | 54034.10892 | 175535.5344 | 114875.1982 | 166300.2034 | 175723.9306 | 200741.3175 | 178148.7982 | 156380.0715 | 167234.2643 |
| 3-Oxocholic acid | 18962.92623 | 5495.27438 | 44757.77749 | 13483.42152 | 19568.2336 | 15907.1735 | 34263.86729 | 32165.59302 | 14955.63136 | 28745.48385 | 23636.19676 | 6648.918377 |
| Iromycin A | 1489692.475 | 1135648.848 | 1274976.342 | 17599944.2 | 234344.5833 | 2414471.989 | 15186608.71 | 118440.0395 | 5790266.17 | 3907496.809 | 396114.3019 | 11269356.22 |
| Trandolapril | 452209.4089 | 0 | 677132.9856 | 138607.6389 | 1201064.406 | 517891.8962 | 457619.3176 | 551657.2216 | 625409.5542 | 672151.5255 | 1197413.944 | 720278.5113 |
| all-trans-13,14-Dihydroretinol | 3442390.11 | 3344799.772 | 3300403.801 | 3064313.286 | 3459394.273 | 3671341.485 | 3462145.941 | 3400092.47 | 3628182.452 | 3363430.966 | 3187876.175 | 3396626.637 |
| M343T317 | 488696.9226 | 492433.9967 | 472662.1681 | 481410.9339 | 485433.6392 | 457878.4359 | 469294.5417 | 472930.383 | 489713.6217 | 468539.4894 | 479323.132 | 474195.6468 |
| 4-(2-Oxo-1,2,3,4-tetrahydroquinolin-6-yl)oxycilostazol | 8245222.755 | 3887309.659 | 5607348.506 | 3809284.213 | 4372727.463 | 7503934.121 | 9371144.368 | 10606791.38 | 8150605.636 | 5311711.374 | 10795017.16 | 3317334.812 |
| 11-Dehydro-2,3-dinor-txb2 | 346404.844 | 346964.5548 | 352179.7776 | 353285.7747 | 334289.7463 | 379000.7197 | 413979.8517 | 363304.1416 | 357754.8503 | 339626.5535 | 338114.4322 | 376919.8657 |
| LNAPE 3:0/N-22:4 | 460659.3406 | 506481.3444 | 307921.6935 | 508836.4259 | 75948.06323 | 72032.35958 | 111965.7973 | 64351.98793 | 50669.80474 | 77873.85722 | 79027.40245 | 57801.00709 |

| **Metabolites** | **PBS_1** | **PBS_2** | **PBS_3** | **PBS_4** | **BCG_1** | **BCG_2** | **BCG_3** | **BCG_4** | **BCGΔ*pe18*_1** | **BCGΔ*pe18*_2** | **BCGΔ*pe18*_3** | **BCGΔ*pe18*_4** |
|---|---|---|---|---|---|---|---|---|---|---|---|---|
| Uridine 5'-diphosphate (UDP) | 200091.2194 | 219820.2335 | 225194.1412 | 180048.4416 | 843235.1184 | 559563.3332 | 847297.4844 | 653487.165 | 717470.1218 | 750790.4192 | 760141.9164 | 833148.7552 |
| Octadecanedioic acid | 414171.0353 | 463605.8611 | 559747.0109 | 493804.917 | 1133427.675 | 1296994.905 | 1085689.071 | 1308070.428 | 975002.9042 | 653607.7648 | 1067414.458 | 1232208.034 |
| SL 10:0;O/22:0 | 602986.1065 | 606723.1007 | 449695.6343 | 260276.225 | 365091.2957 | 323104.2995 | 332539.0576 | 345436.4518 | 371751.9194 | 476412.2799 | 378652.6564 | 174700.5841 |
| PC(18:4(6Z,9Z,12Z,15Z)/20:3(8Z,11Z,14Z)) | 1950696.008 | 1626335.559 | 5768793.714 | 1056150.699 | 1102391.987 | 868372.9318 | 1777174.064 | 1053480.751 | 1569601.134 | 2122660.673 | 1191070.511 | 835672.3922 |
| LPC O-18:2 | 933676.5311 | 1017725.517 | 804828.5391 | 822534.4401 | 490113.6282 | 556421.6187 | 478201.4836 | 538001.8549 | 580899.0622 | 433980.11 | 462606.0887 | 421568.9351 |
| Apomorphine | 863226.1225 | 1001970.642 | 1309860.698 | 1000877.11 | 1314667.499 | 1317728.852 | 1323694.966 | 1177027.199 | 1273993.104 | 710277.5306 | 742083.4003 | 1132843.726 |
| Glycolate | 7292410.675 | 5965225.752 | 8473618.115 | 7025492.255 | 7538526.285 | 8645624.816 | 8465528.013 | 7494872.975 | 7027336.889 | 4448784.264 | 9659454.125 | 19850987.68 |
| 2-Hydroxybutyric acid | 2752904.267 | 1129923.182 | 2140696.638 | 2820984.262 | 2929405.815 | 1769671.562 | 1923798.13 | 4033504.525 | 2325692.257 | 3103243.586 | 1420908.079 | 1532185.252 |
| [(E)-4-Acetyloxy-5-(3,6-diacetyloxy-5,5,8a-trimethyl-2-methylidene-4-oxo-4a,6,7,8-tetrahydro-1H-naphthalen-1-yl)-3 | 160980.4021 | 131462.9081 | 167534.5481 | 187147.9623 | 252893.0716 | 281147.6307 | 239139.8085 | 282741.0125 | 287429.196 | 150199.4355 | 125463.0561 | 220504.2648 |

| Metabolites | PBS_1 | PBS_2 | PBS_3 | PBS_4 | BCG_1 | BCG_2 | BCG_3 | BCG_4 | BCGΔ *pe18*_1 | BCGΔ *pe18*_2 | BCGΔ *pe18*_3 | BCGΔ*pe18*_4 |
|---|---|---|---|---|---|---|---|---|---|---|---|---|
| -methylpent-2-enyl] acetate | | | | | | | | | | | | |
| M520T482 | 547001.1072 | 666814.2658 | 361009.2117 | 535392.6965 | 284699.234 | 376088.6341 | 341583.4462 | 389904.3669 | 397099.4416 | 412169.1043 | 358663.9895 | 131366.4179 |
| Jasminoside | 680977.323 | 608355.731 | 570329.1881 | 679261.7655 | 700831.5132 | 666105.3858 | 649729.6998 | 632756.9235 | 619410.278 | 575639.317 | 602461.4638 | 607348.0995 |
| Solanine D | 133943.0408 | 72141.04298 | 89709.21776 | 63032.76341 | 127381.9084 | 178387.3688 | 183961.584 | 165520.6933 | 106220.6705 | 97141.39198 | 187821.2928 | 42343.39413 |
| 3-(2-(1,3-Benzodioxol-5-yl)-7-methoxy-1-benzofuran-5-yl)-3-hydroxypropyl hexopyranoside | 931583.7332 | 900115.5263 | 772277.9385 | 806820.5896 | 805896.9159 | 855623.827 | 867859.6115 | 961260.541 | 897623.8146 | 959943.8882 | 911455.8239 | 829931.4197 |
| MGDG 2:0_18:3 | 178974.7931 | 71690.57868 | 103260.4726 | 83629.87853 | 35484.22965 | 130499.6496 | 77323.83267 | 38784.5905 | 60612.72476 | 23382.40588 | 55895.57327 | 0 |
| 2-Propyl-4-pentenoic acid | 615760.3492 | 822585.756 | 593067.1655 | 595162.5191 | 814925.7279 | 641144.0923 | 605874.4343 | 601979.5425 | 699510.3216 | 572828.9616 | 678091.8304 | 660418.8902 |
| Methylguanidine | 1328108.43 | 2285073.759 | 3192524.918 | 2492354.072 | 4260227.092 | 3394678.566 | 4659730.211 | 3716153.553 | 2162956.878 | 3217705.368 | 3943549.476 | 2548692.383 |
| Alverine | 1849078.861 | 1927615.969 | 1639095.622 | 1693493.021 | 1732685.948 | 2848104.475 | 1669050.769 | 1606333.467 | 1831845.659 | 1679240.917 | 1765774.598 | 1585004.333 |

| Metabolites | PBS_1 | PBS_2 | PBS_3 | PBS_4 | BCG_1 | BCG_2 | BCG_3 | BCG_4 | BCGΔ*pe18*_1 | BCGΔ*pe18*_2 | BCGΔ*pe18*_3 | BCGΔ*pe18*_4 |
|---|---|---|---|---|---|---|---|---|---|---|---|---|
| Diethylcarbamazine_N-Oxide | 2334055.241 | 10158630.17 | 1414563.663 | 8792533.764 | 11781276.49 | 8467184.636 | 247227.2738 | 465089.2523 | 2455958.439 | 5306692.711 | 5425103.269 | 1646179.861 |
| Tridihexethyl | 3450562.17 | 4298651.925 | 3840826.918 | 4527293.539 | 5786597.994 | 1747991.749 | 4159717.043 | 6112903.265 | 3909486.55 | 4700357.931 | 3653124.246 | 5468630.045 |
| 16-Oxopalmitate | 891329.4322 | 929427.056 | 871257.5232 | 943386.9339 | 846435.0218 | 1043610.908 | 838398.0843 | 842570.2137 | 792451.5134 | 791937.9365 | 803968.1861 | 768714.4007 |
| LPA 18:2 | 1039685.602 | 928931.6975 | 931825.4486 | 875881.8039 | 1070823.342 | 881790.9399 | 997087.2522 | 982737.6142 | 934478.4107 | 837499.6822 | 918178.6046 | 947525.2546 |
| 4′-Hydroxy-2′-methylacetophenone | 469357.3227 | 411192.0641 | 540137.8528 | 464280.5025 | 565449.2124 | 521610.515 | 447724.3909 | 518253.8203 | 518079.5871 | 312234.0585 | 393581.8531 | 486326.1621 |
| Benzoylcarnitine (Car(7:4)) | 974445.6854 | 906796.4936 | 704871.5844 | 822169.3247 | 1833451.529 | 917755.9528 | 972705.405 | 1118732.899 | 980420.2446 | 741214.3251 | 576927.4882 | 600445.6928 |
| 1-Bromo-3-chloro-4-(piperidinocarbonyl)benzene | 473699.9637 | 1168987.602 | 757165.3166 | 1013213.295 | 5957302.602 | 1477292.779 | 861004.2142 | 1807670.449 | 916708.865 | 2637465.907 | 558755.7924 | 461836.891 |
| 5-(2-Chlorophenyl)-1,3,4-oxadiazole-2(3H)-thione | 2595422.355 | 2756316.028 | 3804003.068 | 3617149.808 | 3504660.861 | 4431637.85 | 3068649.896 | 3212503.857 | 3245693.064 | 2886462.399 | 4688535.383 | 3835086.238 |
| Avenestergenin_A1 | 41698.34681 | 28840.35638 | 23312.24856 | 30768.85769 | 0 | 6858.392964 | 75471.14424 | 8969.293219 | 26949.39773 | 66717.17277 | 62090.49298 | 59022.24453 |
| MGDG O-21:2_3:0 | 1495340.878 | 1831440.961 | 1431403.209 | 3011062.154 | 2154925.667 | 2317901.068 | 1555947.996 | 1534758.239 | 1879777.652 | 2140622.786 | 1558026.754 | 1708101.051 |
| Thiomiltefosine | 24392.77779 | 24232.16234 | 4814.584407 | 0 | 20293.41104 | 18131.74898 | 20010.42486 | 16731.06601 | 15150.81028 | 0 | 22911.83266 | 20593.96463 |

| **Metabolites** | **PBS_1** | **PBS_2** | **PBS_3** | **PBS_4** | **BCG_1** | **BCG_2** | **BCG_3** | **BCG_4** | **BCGΔ*pe18*_1** | **BCGΔ*pe18*_2** | **BCGΔ*pe18*_3** | **BCGΔ*pe18*_4** |
|---|---|---|---|---|---|---|---|---|---|---|---|---|
| dTDP-3-N,N-dimethylamino-2,3,6-trideoxy-4-keto-glucose | 178642.4802 | 130989.5216 | 212595.1332 | 150405.7208 | 585080.3065 | 323756.3754 | 471050.2523 | 367433.0854 | 418340.9555 | 307853.2764 | 319299.8865 | 273486.7597 |
| Ceftibuten | 852842.4 | 466459.4261 | 2151430.488 | 918064.6813 | 548774.2196 | 642678.6159 | 862299.7331 | 762543.5149 | 443392.8972 | 205728.5988 | 542201.5178 | 706197.1874 |
| (2E,4E,12Z)-N-(2-methylpropyl)octadeca-2,4,12-trienamide | 2567433.565 | 2668837.442 | 1976800.021 | 2137689.938 | 2145824.223 | 2347763.345 | 2112516.892 | 2583302.67 | 2177244.317 | 2308178.079 | 2071913.839 | 1962756.096 |
| DLCL 14:1_16:4 | 281171.6814 | 319102.0394 | 224753.875 | 268750.3197 | 263230.8158 | 314134.6372 | 323610.657 | 288198.8142 | 263218.5785 | 279195.714 | 297091.0266 | 242182.9001 |
| Maltotetraose | 0 | 0 | 0 | 0 | 175740.1156 | 128182.6692 | 95968.739 | 95223.04902 | 164675.5358 | 120678.1885 | 108679.539 | 134701.4688 |
| CE 15:1 | 115468.9916 | 178958.3664 | 116788.7413 | 104865.1455 | 85467.94202 | 115716.104 | 110212.8407 | 130593.6849 | 161542.6719 | 130573.8235 | 131798.1667 | 80607.55643 |
| LPC 20:1 | 256265.6889 | 286463.0871 | 230883.059 | 207940.8972 | 626076.772 | 962038.48 | 547992.4441 | 627118.6611 | 598791.4529 | 541186.2892 | 567199.3078 | 324028.2396 |
| (5Z)-5-(4-Hydroxy-3-methoxybenzylidene)-1,3-thiazolidine-2,4-dione | 2833415.92 | 2783077.515 | 2482441.671 | 2828703.28 | 2898027.21 | 2836937.616 | 2834670.915 | 2811420.948 | 2753003.067 | 2962758.561 | 2809218.143 | 2598358.511 |
| Pinanethromboxane A2 | 768649.8353 | 681757.0765 | 884874.0046 | 827438.4481 | 778918.9751 | 711407.3688 | 850067.3941 | 735237.265 | 767118.6245 | 832689.5946 | 790749.0523 | 642734.954 |

| **Metabolites** | **PBS_1** | **PBS_2** | **PBS_3** | **PBS_4** | **BCG_1** | **BCG_2** | **BCG_3** | **BCG_4** | **BCGΔ*pe18*_1** | **BCGΔ*pe18*_2** | **BCGΔ*pe18*_3** | **BCGΔ*pe18*_4** |
|---|---|---|---|---|---|---|---|---|---|---|---|---|
| Methyl 3-(acetyloxy)-16-hydroxy-4,4,8,12,16-pentamethyl-15,17,19-trioxoandrost-11-ene-14-carboxylate | 757350.3964 | 819724.569 | 725654.9121 | 701974.6045 | 545236.6628 | 694830.5539 | 778918.3811 | 858371.3492 | 279110.7566 | 737538.4649 | 748343.2119 | 717245.1155 |
| DLCL 18:1_16:4 | 253075.1113 | 325379.0006 | 211377.9918 | 257112.2318 | 252769.6211 | 278321.4408 | 249642.1687 | 255516.1329 | 288337.2907 | 240409.6168 | 271665.2668 | 277618.951 |
| LDGCC 32:5 | 0 | 133295.9225 | 0 | 53498.85155 | 706789.0155 | 646229.7247 | 147227.7046 | 311347.1222 | 203302.1327 | 0 | 87455.51478 | 420400.5342 |
| Triacetonamine | 16024233.57 | 14484952.43 | 7963103.159 | 9764040.889 | 10082011.24 | 12017019.13 | 12698866.49 | 9966402.554 | 10914836.03 | 11129495.71 | 9613813.805 | 12309365.56 |
| Deoxythymidine 5'-phosphate (dTMP) | 91939.57337 | 135127.0962 | 108486.4255 | 96985.178 | 1079035.742 | 883035.0485 | 800492.3279 | 721804.1693 | 1392652.49 | 1193647.418 | 1307428.972 | 1437866.796 |
| Pindolol | 239778.3972 | 0 | 251296.7764 | 145526.9968 | 174693.4667 | 256950.3673 | 154366.1337 | 272582.2195 | 133836.5454 | 204385.0483 | 174246.7771 | 262006.3524 |
| Lurasidone | 1892473.056 | 1988145.711 | 1755679.437 | 2040826 | 1839376.106 | 1896435.923 | 1918211.896 | 1908970.08 | 1839053.652 | 1822884.378 | 1724584.462 | 1615778.574 |
| Leucine conjugated chenodeoxycholic acid | 1014397.493 | 753959.8525 | 769147.0268 | 1145443.038 | 256746.5894 | 872106.9559 | 1351287.005 | 367527.0445 | 767419.6899 | 1206017.79 | 675855.7887 | 1572907.264 |
| Mangiferin | 111586.8092 | 8979.166746 | 157319.8397 | 23504.75986 | 401789.392 | 515944.3476 | 125758.8543 | 525123.7312 | 796202.0816 | 788963.8296 | 485068.5597 | 392873.4003 |

| Metabolites | PBS_1 | PBS_2 | PBS_3 | PBS_4 | BCG_1 | BCG_2 | BCG_3 | BCG_4 | BCGΔ*pe18*_1 | BCGΔ*pe18*_2 | BCGΔ*pe18*_3 | BCGΔ*pe18*_4 |
|---|---|---|---|---|---|---|---|---|---|---|---|---|
| DG(16:0/16:0/0:0) | 56615.372 | 53963.05483 | 46028.77392 | 68540.14044 | 58171.33273 | 23457.10697 | 64014.07016 | 35027.72485 | 49949.39063 | 36689.10696 | 68374.31782 | 68507.70664 |
| PEtOH 16:0_21:0 | 381644.5444 | 138812.4422 | 537624.391 | 382303.2539 | 142445.0082 | 0 | 138756.1771 | 58553.10724 | 54670.32139 | 133061.8724 | 161485.1832 | 69057.85049 |
| Edulisin_VI | 0 | 0 | 0 | 0 | 256522.1851 | 194889.014 | 164299.0389 | 151072.1767 | 391856.6301 | 309857.8981 | 380286.5669 | 349844.2524 |
| PI(18:1(11Z)/20:4(5Z,8Z,11Z,14Z)) | 633584.9361 | 673416.2303 | 240917.7219 | 472410.8945 | 1536123.998 | 1000391.599 | 1419886.82 | 1295934.163 | 1488828.484 | 1919882.885 | 1493789.251 | 954747.5834 |
| Arachidonoylcarnitine (Car(20:4)) | 10017088.57 | 8148168.184 | 12006525.84 | 9887093.211 | 790324.6164 | 864027.6857 | 1290657.727 | 1177243.947 | 804023.908 | 1212866.109 | 1091879.079 | 623842.3169 |
| Glufosinate | 900956.9817 | 868294.593 | 6186178.692 | 1526508.005 | 1484583.786 | 2483271.408 | 1359740.675 | 1657873.759 | 807755.9539 | 305835.0935 | 2239013.298 | 3244661.153 |
| Nicotinamide riboside (NR) | 1178187.693 | 874374.1654 | 2014154.495 | 1415385.162 | 5241976.15 | 4021023.298 | 4897572.424 | 3455975.32 | 3063771.438 | 3052157.917 | 3445017.989 | 2395788.369 |
| Ursodeoxycholic acid | 25017.90944 | 83124.85562 | 171961.3388 | 390541.7161 | 0 | 115281.5525 | 177712.2839 | 145487.2895 | 109661.1905 | 0 | 61615.55055 | 92265.08239 |
| (R)-N-(1-(4-Hydroxyphenyl)-2-hydroxyethyl)oleamide | 167739.3265 | 183986.8764 | 235527.7618 | 73908.03431 | 216548.7168 | 177750.7038 | 188079.5756 | 212576.3606 | 159927.6404 | 166577.6472 | 158406.4965 | 225218.9937 |
| Creatine | 165264.1716 | 0 | 353100.1354 | 0 | 1930581.77 | 1066908.158 | 2802976.772 | 1775134.719 | 924365.1422 | 1361608.367 | 1948925.756 | 1075627.386 |

| Metabolites | PBS_1 | PBS_2 | PBS_3 | PBS_4 | BCG_1 | BCG_2 | BCG_3 | BCG_4 | BCGΔ *pe18*_1 | BCGΔ *pe18*_2 | BCGΔ *pe18*_3 | BCGΔ*pe18*_4 |
|---|---|---|---|---|---|---|---|---|---|---|---|---|
| (1S,4R)-4-[[(2S,3aS,4S)-2-Butan-2-yl-4-hydroxy-1-oxo-3,3a-dihydro-2H-imidazo[1,2-a]indol-4-yl]methyl]-1-methyl-2,4-dihydro-1H-pyrazino[2,1-b]quinazoline-3,6-dione | 296759.5745 | 408937.5198 | 375929.821 | 237060.6388 | 260304.5882 | 325267.435 | 277193.5726 | 318344.3159 | 325147.6705 | 214510.1581 | 263260.5792 | 158259.7524 |
| 2-(5-Oxovaleryl)phosphatidylcholine | 209300.9023 | 205493.1243 | 60618.48481 | 142403.6276 | 142551.329 | 72673.99351 | 92881.082 | 130279.9127 | 127867.2422 | 140851.1118 | 79949.52391 | 139820.3835 |
| PC(8:0/8:0) | 16169.12445 | 23140.19071 | 108617.411 | 113045.3026 | 400709.9208 | 330842.6926 | 392145.6574 | 343219.8076 | 382350.0382 | 263797.6215 | 379184.0293 | 373732.7729 |
| Sph(d18:0) | 13689288.95 | 13949526.93 | 8770339.204 | 10343451.26 | 10204738.45 | 7735728.758 | 10303378.45 | 10064544.61 | 9998137.828 | 11645321.82 | 11129411.1 | 4621739.653 |
| Apicidin | 88001.67992 | 115935.2293 | 79465.40927 | 94450.59802 | 94796.2652 | 86192.85669 | 102606.6868 | 70373.29695 | 84218.01376 | 90749.50165 | 85503.50075 | 88976.87604 |
| Octanoylcarnitine (Car(8:0)) | 481754.5542 | 480517.8506 | 473385.723 | 667784.3681 | 1669195.038 | 805697.6201 | 1798894.917 | 527184.4489 | 714997.4396 | 1249690.222 | 903069.4657 | 433400.4328 |
| Cer 18:1;2O/26:1 | 66758.93032 | 40822.15725 | 32823.08954 | 26659.49301 | 44664.99078 | 49875.01153 | 45614.13165 | 24575.40035 | 37224.05163 | 33071.52763 | 30711.25439 | 63634.06537 |
| N-Decyl-N,N-dimethyl-1-decanaminiu | 1384103.63 | 1421118.984 | 990598.2556 | 2002747.149 | 1603709.779 | 1330624.871 | 1168898.283 | 1506246.466 | 1478010.018 | 1207391.679 | 1246957.119 | 1447790.739 |

| Metabolites | PBS_1 | PBS_2 | PBS_3 | PBS_4 | BCG_1 | BCG_2 | BCG_3 | BCG_4 | BCGΔ*pe18*_1 | BCGΔ*pe18*_2 | BCGΔ*pe18*_3 | BCGΔ*pe18*_4 |
|---|---|---|---|---|---|---|---|---|---|---|---|---|
| m cation | | | | | | | | | | | | |
| Cer(d18:1/18:0) | 106867.8501 | 57070.15122 | 145380.2879 | 98802.8151 | 66526.19348 | 55459.65692 | 68275.34583 | 119712.4166 | 87010.48685 | 71508.16785 | 93678.58415 | 125589.5818 |
| PI 12:0_22:6 | 0 | 0 | 14891.17805 | 9247.166259 | 88986.47485 | 36460.37134 | 97478.12278 | 55544.89351 | 37381.63197 | 35967.33871 | 47920.04424 | 31189.36629 |
| FA 26:2 | 10437.56211 | 8823.562879 | 7989.96682 | 4539.280222 | 148733.4848 | 249044.82 | 149404.7392 | 264565.8139 | 109037.2586 | 57959.94149 | 124291.8731 | 143885.5981 |
| Tris(2-butoxyethyl) phosphate | 990038.8736 | 1028513.903 | 1085475.219 | 1113853.991 | 1053787.906 | 1163499 | 1140164.54 | 1076450.153 | 1185902.888 | 1071818.564 | 1203741.86 | 1184160.498 |
| 6-Benzyl-3-butan-2-yl-9-(7,8-dihydroxy-6-oxooctyl)-1,4,7,10-tetrazabicyclo[10.4.0]hexadecane-2,5,8,11-tetrone | 40403.86037 | 38747.85471 | 16843.55941 | 53278.01316 | 42518.74266 | 66075.1123 | 47931.4573 | 49769.99918 | 34225.60729 | 27960.54155 | 32315.21793 | 23823.86298 |
| Met-Ser | 169448.4633 | 224451.9596 | 193039.3253 | 207137.5644 | 212886.1114 | 415130.3548 | 178558.1288 | 234289.8492 | 188131.0766 | 181622.7753 | 225997.5141 | 214832.3224 |
| Frangulanine | 595928.9672 | 533713.0084 | 244052.5403 | 504367.9986 | 2516753.899 | 1970614.28 | 605248.5407 | 1955744.069 | 1359175.16 | 223893.1117 | 342892.1964 | 1097111.986 |
| Glycocholic acid | 54939.00816 | 24803.52862 | 105287.8449 | 47773.96758 | 96299.66663 | 87161.91962 | 115579.4757 | 76552.66033 | 67234.95877 | 54039.13567 | 50219.10622 | 74054.08849 |
| Traumatic acid | 378662.3954 | 372922.4204 | 439302.9353 | 506449.1542 | 533194.1338 | 399193.951 | 1031739.117 | 345797.4784 | 534501.4578 | 517571.2229 | 331317.0471 | 610466.5974 |

| Metabolites | PBS_1 | PBS_2 | PBS_3 | PBS_4 | BCG_1 | BCG_2 | BCG_3 | BCG_4 | BCGΔ*pe18*_1 | BCGΔ*pe18*_2 | BCGΔ*pe18*_3 | BCGΔ*pe18*_4 |
|---|---|---|---|---|---|---|---|---|---|---|---|---|
| LPC O-14:0 | 872162.8911 | 1095417.828 | 1496452.642 | 1325167.1 | 831521.8116 | 717619.1173 | 1111121.211 | 730851.9352 | 842968.6524 | 629409.6931 | 101949.0118 | 1615780.319 |
| PC 33:0_20:4 | 251176.5726 | 123016.9186 | 188032.6673 | 142860.216 | 114335.3358 | 0 | 151842.6952 | 199709.0959 | 44293.4034 | 114969.7674 | 77776.17239 | 50617.08315 |
| PC 16:0_44:10 | 126102.9103 | 69573.41242 | 82975.14162 | 43026.9476 | 71160.91855 | 0 | 92023.40978 | 48575.10108 | 56633.36497 | 111053.5265 | 105586.6961 | 0 |
| Butyllactate | 1062905.291 | 1055162.43 | 1143845.488 | 1123950.764 | 1212004.592 | 1003355.196 | 1065115.234 | 957820.4718 | 1048592.632 | 1051131.089 | 1115196.815 | 1017359.478 |
| DGDG O-28:7_18:0 | 172878.9999 | 189777.5394 | 56939.90208 | 151987.1064 | 228655.5919 | 317175.0127 | 203470.5042 | 174318.7584 | 154229.6645 | 150391.3088 | 130685.1731 | 0 |
| 2-[(2-Ethylbutanoyl)amino]-4,5-dimethoxybenzoic acid | 29430.88783 | 28189.49723 | 30603.32579 | 23260.68549 | 29019.44205 | 25929.83068 | 27266.26781 | 28631.24897 | 27363.20674 | 26089.12835 | 27601.86892 | 30199.32001 |
| FAHFA 18:0/2:0 | 591878.2532 | 779039.5988 | 505151.2708 | 595704.8772 | 495233.648 | 482897.6824 | 543720.2287 | 488749.9697 | 579105.7994 | 444628.8596 | 501358.6694 | 878778.2973 |
| PC 18:1_20:4 | 576342.6446 | 506957.6947 | 509659.2528 | 534787.9997 | 559698.6358 | 712970.7974 | 888405.7662 | 584908.3552 | 448411.5121 | 514038.7561 | 561319.4137 | 553220.1015 |
| Neolinustatin | 11379.58301 | 10463.62203 | 15340.47092 | 15461.30234 | 21589.55221 | 17338.66031 | 22871.48392 | 17891.10842 | 34459.84456 | 27641.39933 | 33492.57488 | 19393.66695 |
| Leucylphenylalanine | 134609.5722 | 127216.6649 | 176952.9407 | 118246.3326 | 176785.8097 | 130779.1647 | 152732.4959 | 114931.1105 | 154604.1666 | 159769.5575 | 164475.4289 | 99740.98468 |
| M891T521 | 125500.0262 | 127834.1053 | 153560.1333 | 111898.9537 | 206225.1029 | 0 | 278304.6778 | 1407718.269 | 212698.0784 | 111916.831 | 102122.249 | 406323.6712 |
| Cer 13:2;2O/2:0 | 992609. | 803357. | 712334. | 785084. | 768820. | 811344. | 878550. | 798783. | 811340. | 101567 | 798480. | 785776. |

| Metabolites | PBS_1 | PBS_2 | PBS_3 | PBS_4 | BCG_1 | BCG_2 | BCG_3 | BCG_4 | BCGΔ*pe18*_1 | BCGΔ*pe18*_2 | BCGΔ*pe18*_3 | BCGΔ*pe18*_4 |
|---|---|---|---|---|---|---|---|---|---|---|---|---|
| | 9242 | 686 | 5976 | 6614 | 372 | 2946 | 2442 | 2127 | 8116 | 7.926 | 3179 | 1358 |
| PG 18:1_20:6;O | 227260.1674 | 183516.4367 | 279794.5386 | 183433.2071 | 367052.0921 | 132658.5555 | 223678.7009 | 289802.3694 | 307919.4209 | 227102.8645 | 203058.2654 | 75723.21015 |
| N-(Deshydroxyethyl)dasatinib | 0 | 0 | 0 | 0 | 523708.4509 | 461594.0351 | 250761.0687 | 300174.0349 | 789267.0438 | 879935.3146 | 610798.6801 | 1101100.97 |
| 2-(1,3-Thiazol-4-yl)-1H-benzimidazol-5-amine | 7152957.587 | 7046688.201 | 7732453.828 | 7728241.313 | 8400748.921 | 8241034.758 | 7076765.346 | 7700784.882 | 8325268.131 | 6304224.249 | 6363930.869 | 7010006.589 |
| Mandelic acid | 361333.3646 | 285614.0106 | 397206.7797 | 353733.6495 | 545920.8451 | 503696.8259 | 547295.6313 | 452433.4357 | 426242.1546 | 427047.2128 | 413228.5775 | 405053.9552 |
| Glutarylcarnitine | 319418.1463 | 364969.4733 | 399609.6121 | 385882.225 | 817295.1134 | 509560.4452 | 614733.4167 | 522789.3187 | 637030.5671 | 379857.8011 | 498567.1313 | 575631.4732 |
| Hypaphorine | 101373.911 | 141063.8865 | 165666.3403 | 106078.59 | 175778.9944 | 154394.158 | 173451.3896 | 169404.7266 | 197606.1069 | 94471.57381 | 205860.3304 | 157502.3284 |
| 1,3 Cyclohexanedione | 1091336.227 | 1106158.68 | 1138746.914 | 1219434.489 | 1137180.22 | 1125484.923 | 838851.8845 | 1158664.731 | 1172275.79 | 1107485.79 | 1120830.352 | 935429.4397 |
| 2-amino-8-hydroxydecanoate | 11503.40732 | 411632.6754 | 366132.3289 | 459123.1473 | 87578.60698 | 8446.941286 | 138850.3621 | 0 | 97578.76409 | 194043.9718 | 51790.12349 | 1892405.765 |
| PS(18:1/18:1) | 1892691.114 | 1385366.721 | 1623498.473 | 1138141.022 | 2254136.25 | 1844478.918 | 1949685.469 | 2303980.762 | 2406616.567 | 1982413.734 | 1965212.011 | 2094543.492 |
| .gamma.-Octalactone | 589337.8588 | 579783.5326 | 503694.0824 | 978486.248 | 553225.5221 | 615296.0225 | 1384930.188 | 425409.1087 | 887524.1315 | 793516.5956 | 580900.7274 | 1128068.991 |
| CAR 13:0 | 774290.4936 | 1612158.714 | 1025077.33 | 837620.5334 | 3052995.896 | 1802837.611 | 2855347.263 | 2278670.779 | 1540854.409 | 2631080.911 | 2690500.433 | 1453493.502 |

| Metabolites | PBS_1 | PBS_2 | PBS_3 | PBS_4 | BCG_1 | BCG_2 | BCG_3 | BCG_4 | BCGΔ *pe18*_1 | BCGΔ *pe18*_2 | BCGΔ *pe18*_3 | BCGΔ*pe18*_4 |
|---|---|---|---|---|---|---|---|---|---|---|---|---|
| 1H-Benzimidazol-2-ylcyanamide | 142292.7001 | 416998.4665 | 313602.7318 | 397417.8616 | 533808.297 | 372752.1914 | 380046.7477 | 441161.8395 | 380022.0516 | 310247.6191 | 331816.351 | 297684.3891 |
| Ethyl (2Z)-2-cyano-3-(1H-indol-3-yl)prop-2-enoate | 2047968.112 | 2231589.846 | 1833449.247 | 1938987.825 | 1874786.441 | 1932336.789 | 2049865.428 | 1865694.217 | 1979676.955 | 2093417.503 | 2020168.217 | 1782280.275 |
| PI 20:4_22:5 | 173127.9159 | 1030992.491 | 157819.2079 | 444692.9388 | 11811.81892 | 41617.80704 | 32237.89691 | 39828.75116 | 55121.96286 | 51843.19622 | 26647.69889 | 54462.12003 |
| Saccharopine | 103796.9878 | 98187.0121 | 118434.2819 | 115691.9003 | 2139052.775 | 1377308.285 | 1340780.352 | 1196575.687 | 1708933.675 | 1498102.305 | 1600360.011 | 2096969.786 |
| Oleoyl ethylamide | 736596.2859 | 1164030.297 | 601544.5 | 656877.8591 | 294238.4306 | 375949.5632 | 287033.1495 | 329233.3312 | 330459.6358 | 230555.0487 | 305673.5185 | 239912.7207 |
| Melamine | 1231243.393 | 1385046.891 | 1971690.156 | 1875259.158 | 1309544.67 | 1297464.969 | 1309866.456 | 1329584.227 | 1355267.061 | 1035194.311 | 1266754.778 | 2149732.453 |
| Thr-Leu | 769188.9897 | 820316.7935 | 1078220.441 | 852097.6931 | 738550.396 | 547315.7233 | 886976.7395 | 730005.622 | 824385.1481 | 817434.2517 | 1029567.426 | 929061.2897 |
| N1,N8-Bis(4-coumaroyl)spermidine | 482783.2315 | 653288.1824 | 310289.2132 | 556436.5881 | 771759.4133 | 434932.0365 | 674203.7331 | 518918.0457 | 670482.7334 | 643973.371 | 447494.3599 | 773934.7079 |
| 1-Butanol, 2-[[9-(1-methylethyl)-6-[[[4-(2-pyridinyl)phenyl]methyl]amino]-9H-purin-2-yl]amino]-, (2R)- | 25362.93191 | 32459.94352 | 40106.9064 | 37175.16864 | 185392.8295 | 72744.88786 | 184001.6026 | 124973.0417 | 111581.4211 | 133440.2984 | 149974.6185 | 99268.66306 |

| **Metabolites** | **PBS_1** | **PBS_2** | **PBS_3** | **PBS_4** | **BCG_1** | **BCG_2** | **BCG_3** | **BCG_4** | **BCGΔ*pe18*_1** | **BCGΔ*pe18*_2** | **BCGΔ*pe18*_3** | **BCGΔ*pe18*_4** |
|---|---|---|---|---|---|---|---|---|---|---|---|---|
| Capsaicin | 298607.2234 | 235290.6872 | 307287.2533 | 224123.3176 | 300111.6106 | 298867.8131 | 316328.0325 | 295342.7099 | 306069.2964 | 306667.7398 | 306329.7033 | 311732.9073 |
| PI 4:0_22:6 | 18152.7958 | 18381.99572 | 25518.53292 | 25574.47448 | 21656.55546 | 21227.53321 | 21589.9001 | 19517.90288 | 21624.27848 | 20274.38279 | 32025.21398 | 21228.55215 |
| 3b,18b-3-Methoxy-11-oxo-12-oleanen-30-oic_acid | 2399274.403 | 2803644.245 | 2350535.096 | 2477716.223 | 1955633.69 | 2273029.583 | 1930125.249 | 2493095.264 | 2008930.339 | 2271996.534 | 1881519.878 | 1752105.146 |
| Hexacosanoic acid | 886329.0416 | 549436.8579 | 375871.8311 | 667829.8309 | 1193630.299 | 1820878.048 | 1263485.238 | 1174040.127 | 481771.1381 | 391734.1997 | 500779.129 | 390443.7952 |
| PC 18:0_36:5 | 0 | 22652.80824 | 0 | 0 | 109059.4496 | 0 | 139330.7015 | 458457.2301 | 97877.2873 | 41898.01871 | 73508.58929 | 191355.2628 |
| FAHFA 18:0/20:2 | 0 | 45591.99667 | 25682.3736 | 0 | 0 | 197958.3696 | 21839.57005 | 27016.90984 | 0 | 0 | 70766.37268 | 50860.07794 |
| LNAPS 24:2/N-26:7 | 0 | 58992.50884 | 61199.64296 | 67045.73126 | 38617.85555 | 39139.15681 | 0 | 0 | 5774.52598 | 3648.894161 | 0 | 63052.64362 |
| SHexCer 21:1;3O | 55054.86282 | 70513.58243 | 55946.87749 | 58393.88417 | 75039.47979 | 69983.70022 | 68278.80661 | 60455.17665 | 66232.92788 | 57699.69682 | 70249.74013 | 65467.19484 |
| (3a,5b)-24-oxo-24-[(2-sulfoethyl)amino]cholan-3-yl-b-Glucopyranosiduronic acid | 76097.73641 | 84000.20082 | 34085.51999 | 78544.76751 | 103808.6608 | 47580.77912 | 77114.68706 | 73449.1152 | 86822.88125 | 96373.31844 | 103785.0753 | 75437.49812 |
| Erucamide | 645470.9954 | 576904.7511 | 705035.5893 | 493569.4569 | 272839.6813 | 375274.7193 | 367969.3328 | 912078.8745 | 429920.1698 | 444587.8565 | 831087.4599 | 631142.8402 |

| Metabolites | PBS_1 | PBS_2 | PBS_3 | PBS_4 | BCG_1 | BCG_2 | BCG_3 | BCG_4 | BCGΔ*pe18*_1 | BCGΔ*pe18*_2 | BCGΔ*pe18*_3 | BCGΔ*pe18*_4 |
|---|---|---|---|---|---|---|---|---|---|---|---|---|
| Polyoxyethylene_dioleate | 70020.55759 | 146577.008 | 186341.9298 | 152451.0334 | 567285.3118 | 529982.8419 | 552785.6232 | 505373.4965 | 623295.0504 | 681108.8603 | 545099.2545 | 307584.1573 |
| Lignoceroylcarnitine [M] | 72140.25711 | 71093.40125 | 76720.16249 | 17765.58363 | 51822.6222 | 66346.21579 | 80924.19068 | 74464.77094 | 63175.92105 | 75646.44842 | 98176.21502 | 29635.48442 |
| (3.beta.,5.Xi.,9.Xi.,22.beta.)-22-Hydroxyolean-12-en-3-yl 6-deoxyhexopyranosyl-(1->2)hexopyranosyl-(1->2)hexopyranosiduronic acid | 56978.39937 | 0 | 9185.738275 | 26241.90555 | 256096.8089 | 672063.8426 | 508184.7699 | 329092.7937 | 314987.3792 | 311976.2473 | 518500.6027 | 303935.2073 |
| FAHFA 2:0/20:0 | 433819.4193 | 477456.6486 | 254685.553 | 345208.2694 | 224876.8693 | 454620.8363 | 185724.1561 | 318967.1664 | 148160.519 | 207537.0228 | 243595.347 | 568038.3925 |
| Isodeoxycholic acid | 728581.7868 | 961041.6433 | 1103004.852 | 2001379.881 | 695466.5473 | 1162736.022 | 962468.2383 | 774080.8096 | 860814.4541 | 689500.5402 | 741553.3499 | 804666.2826 |
| 5-Cholesten-3beta-25-diol-3-sulfate | 76687.41253 | 38503.96986 | 105409.9727 | 80090.901 | 214800.8507 | 161699.9629 | 177744.492 | 228643.0494 | 161681.8779 | 185921.0661 | 199541.4696 | 229369.4527 |
| Nicotine | 2460115.171 | 5995504.613 | 8336952.264 | 6527328.169 | 1968925.869 | 1518197.07 | 2215933.282 | 2588343.747 | 2440159.979 | 2097209.49 | 3412657.809 | 19656217.18 |
| Pantothenol | 2608610.735 | 6653264.963 | 5592147.249 | 7308126.779 | 5459796.761 | 4669501.651 | 7456623.604 | 4486736.144 | 6447681.648 | 5429203.546 | 6828384.622 | 6227701.814 |
| LPG O-11:0 | 312244.8603 | 364266.3275 | 607215.8012 | 419331.8921 | 606093.714 | 599835.1069 | 864441.2325 | 549298.0564 | 570422.0994 | 466882.1947 | 450748.0057 | 488349.5959 |
| PS 30:6 | 95424.3 | 90879.5 | 126307. | 131366. | 194596. | 195140. | 199254. | 168995. | 88510.1 | 98058.9 | 203398. | 230894. |

| Metabolites | PBS_1 | PBS_2 | PBS_3 | PBS_4 | BCG_1 | BCG_2 | BCG_3 | BCG_4 | BCGΔ*pe18*_1 | BCGΔ*pe18*_2 | BCGΔ*pe18*_3 | BCGΔ*pe18*_4 |
|---|---|---|---|---|---|---|---|---|---|---|---|---|
| | 9417 | 5445 | 3518 | 5724 | 8833 | 6428 | 2505 | 0631 | 2156 | 3964 | 2592 | 1017 |
| 1-Oleoyl-sn-glycerol 3-phosphate | 240769.4219 | 249546.6121 | 243379.6099 | 315506.541 | 443620.6368 | 515608.326 | 428147.3079 | 422023.7179 | 351542.8697 | 386767.8904 | 502445.7854 | 288738.4855 |
| Acrimarine_I | 84146.6179 | 69219.61942 | 67381.30822 | 106010.3214 | 80371.59374 | 51452.00106 | 101523.0198 | 81233.43161 | 118254.0901 | 89508.95289 | 39171.5321 | 58012.32936 |
| DGTS 7:0_26:4 | 198721.7275 | 598897.4286 | 262081.908 | 472286.4847 | 357518.6505 | 332158.0179 | 254969.8673 | 563368.8877 | 223272.9481 | 520371.2729 | 260188.3846 | 354552.8552 |
| SM 42:9;3O | 739508.5177 | 1143869.782 | 437915.9931 | 624964.7964 | 195537.6648 | 336570.5784 | 271025.4588 | 240981.3554 | 254493.7844 | 268426.0518 | 365684.7815 | 149885.8377 |
| PI 44:5 | 189283.3288 | 258041.3706 | 185022.7066 | 283960.3695 | 234429.7779 | 320720.739 | 370785.548 | 285328.6899 | 160268.5675 | 382935.8406 | 399953.9731 | 391020.5993 |
| ST 28:1;O;S | 251416.4269 | 207895.0198 | 159113.1023 | 207617.8611 | 194722.9684 | 190419.4389 | 175626.2541 | 218880.3439 | 274827.2347 | 239041.8125 | 198888.8141 | 160411.3861 |
| Ampiroxicam | 335177.8185 | 111242.5714 | 350852.268 | 176828.5026 | 280578.7057 | 357976.1003 | 371069.5444 | 42629.78124 | 169924.4992 | 114217.6833 | 48257.42517 | 94041.21339 |
| LPA 22:2 | 90480.25019 | 58877.89227 | 52454.265 | 57323.08331 | 63471.05855 | 58262.2457 | 59475.8131 | 57308.64171 | 52900.4426 | 61917.58276 | 72512.0437 | 103898.2673 |
| Gambogic_acid | 4319646.218 | 4088992.293 | 4554625.702 | 3990388.005 | 4917469.41 | 3994424.555 | 4508184.281 | 4683834.3 | 4779949.762 | 4639188.915 | 4667846.417 | 3963261.57 |
| Cinncassiol_D1_glucoside | 944023.0519 | 463395.39 | 1335583.584 | 1441530.219 | 1812562.368 | 1189329.336 | 1371002.089 | 1320775.151 | 1792496.269 | 1040845.16 | 1668110.823 | 1015070.435 |
| 3-Chloro-5,7-dihydroxy-4-methylcoumarin | 1435334.741 | 1020133.797 | 959222.954 | 1327289.026 | 1082756.917 | 922779.9311 | 1815267.084 | 1731214.572 | 1749671.782 | 1365114.172 | 1514227.531 | 1465198.518 |

| Metabolites | PBS_1 | PBS_2 | PBS_3 | PBS_4 | BCG_1 | BCG_2 | BCG_3 | BCG_4 | BCGΔ*pe18*_1 | BCGΔ*pe18*_2 | BCGΔ*pe18*_3 | BCGΔ*pe18*_4 |
|---|---|---|---|---|---|---|---|---|---|---|---|---|
| Allocholic acid | 205281.9058 | 78531.48361 | 201356.0458 | 182708.7116 | 174499.9499 | 298825.8015 | 221116.4669 | 203117.1547 | 148155.5929 | 141901.7265 | 227233.3137 | 137074.1507 |
| N-Phenyl-N'-[3-(trifluoromethyl)phenyl]urea | 55373.68313 | 52831.95712 | 53310.97097 | 54421.24751 | 57557.03807 | 57797.29491 | 57217.5545 | 48841.28733 | 55249.99453 | 50805.36243 | 48400.5267 | 59073.04563 |
| (E/Z)-4-Hydroxytamoxifen | 208843.6948 | 47730.513 | 251502.4547 | 52953.51194 | 437364.7866 | 612901.4554 | 1691480.555 | 1426515.468 | 570859.9333 | 408151.6429 | 852481.4158 | 385243.9248 |
| 3,5,9-Trioxa-4-phosphatetracosan-1-aminium, 7-(acetyloxy)-24-carboxy-4-hydroxy-N,N,N-trimethyl-, inner salt, 4-oxide, (R)- | 37243.02074 | 18529.99771 | 18569.25992 | 14341.22301 | 427400.8246 | 380577.0174 | 265014.3402 | 327491.4454 | 311491.7986 | 254792.3495 | 350914.6739 | 301903.9134 |
| SMGDG O-8:0_24:3 | 270014.5519 | 425488.9356 | 308796.7898 | 357720.3286 | 496057.0764 | 404619.774 | 421756.8451 | 403479.8193 | 554398.7717 | 374272.515 | 482162.8978 | 432537.3795 |
| CRUSTECDYSONE | 72164.62708 | 77362.00164 | 235591.2436 | 121775.7505 | 93524.83906 | 56968.4745 | 82316.7665 | 67600.92432 | 50113.08826 | 39554.98341 | 31649.05472 | 49810.80922 |
| LPC O-9:0 | 2713959.716 | 2661343.023 | 2860441.638 | 2355383.854 | 2778326.036 | 2726683.125 | 2640731.813 | 2668607.751 | 2552369.038 | 2738550.023 | 2764858.929 | 2678248.191 |
| 3-hydroxypentadecanoyl carnitine | 71883.36963 | 122728.1611 | 51022.35077 | 75384.26832 | 216658.0543 | 65617.35133 | 253482.7404 | 140998.1722 | 150232.8192 | 155532.3061 | 92891.54007 | 54860.17233 |
| Xanthone | 108794 | 224800 | 142414 | 155513 | 146987 | 146003 | 151649 | 148855 | 142781 | 144042 | 154157 | 149996 |

| **Metabolites** | **PBS_1** | **PBS_2** | **PBS_3** | **PBS_4** | **BCG_1** | **BCG_2** | **BCG_3** | **BCG_4** | **BCGΔ*pe18*_1** | **BCGΔ*pe18*_2** | **BCGΔ*pe18*_3** | **BCGΔ*pe18*_4** |
|---|---|---|---|---|---|---|---|---|---|---|---|---|
| | 9.798 | 7.727 | 2.233 | 1.043 | 4.028 | 3.846 | 0.907 | 1.828 | 6.115 | 9.831 | 6.22 | 1.23 |
| PG 18:1_18:1;O | 82188.23159 | 108173.6395 | 294239.4207 | 427152.2738 | 231710.8914 | 271527.9106 | 276342.3151 | 243785.1659 | 192002.4816 | 228168.0037 | 343821.2573 | 208974.5044 |
| M485T482_2 | 1286686.756 | 1581206.756 | 774921.2586 | 1034059.07 | 739694.9715 | 685892.884 | 815703.0172 | 842591.0309 | 930530.2812 | 924317.0109 | 824156.0022 | 593077.3877 |
| Adenosine 3',5'-diphosphate (PAP) | 378323.2436 | 503121.1916 | 379543.4121 | 411862.2418 | 454716.9533 | 372587.8604 | 172244.3132 | 244547.0491 | 300716.4026 | 671351.7392 | 350092.3163 | 1560790.557 |
| PI 40:0 | 364826.4896 | 211374.4964 | 146186.3279 | 233276.0073 | 202619.8997 | 233390.6921 | 217614.8512 | 197855.9868 | 237788.9309 | 198489.3086 | 179273.2317 | 87258.05311 |
| MGDG 2:0_6:0 | 0 | 0 | 0 | 0 | 110106.9164 | 75905.49186 | 72865.58696 | 47441.4668 | 72330.47412 | 60870.79315 | 69147.0338 | 57412.89661 |
| 3beta-Ergosta-5,23-dien-3-ol | 474769.2763 | 514127.9125 | 340557.7929 | 385712.9862 | 279786.5092 | 337146.1599 | 237745.148 | 358629.1981 | 271237.6581 | 302481.325 | 318675.6837 | 476529.1384 |
| Fulvestrant | 118702.6013 | 117011.0717 | 111627.9401 | 123094.1593 | 64550.39995 | 29220.54811 | 55466.36945 | 0 | 0 | 0 | 38816.13495 | 36156.04831 |
| PC O-22:6_20:4 | 0 | 0 | 0 | 0 | 885742.4968 | 794677.8893 | 837921.6666 | 778702.9152 | 27927.14138 | 203237.8975 | 252762.6338 | 185688.7351 |

| **Metabolites** | **PBS_1** | **PBS_2** | **PBS_3** | **PBS_4** | **BCG_1** | **BCG_2** | **BCG_3** | **BCG_4** | **BCGΔ*pe18*_1** | **BCGΔ*pe18*_2** | **BCGΔ*pe18*_3** | **BCGΔ*pe18*_4** |
|---|---|---|---|---|---|---|---|---|---|---|---|---|
| 2-[3-[(E)-6-Acetyloxy-2-hydroxy-6-methyl-3-oxohept-4-en-2-yl]-6-(carboxymethyl)-2-hydroxy-3a,5a,9b-trimethyl-5-oxo-2,3,4,6,9,9a-hexahydro-1H-cyclopenta[a]naphthalen-7-yl]-2-methylpropanoic acid | 104007.2363 | 71768.58535 | 84521.68927 | 46920.18575 | 98450.55459 | 94684.25147 | 85599.20539 | 89641.09353 | 66618.55387 | 89807.53985 | 114638.8635 | 83617.44254 |
| [(5-Chloro-1,3-benzothiazol-2-yl)sulfanyl]acetic acid | 1281259.324 | 982130.2352 | 1157206.999 | 1574681.379 | 1872272.804 | 2288291.9 | 1780931.56 | 1920000.045 | 1067844.008 | 1044132.166 | 2156651.481 | 1878436.002 |
| PS 12:0_26:4 | 208316.2271 | 410766.5207 | 310517.9647 | 260300.732 | 362775.8557 | 824919.1006 | 67722.19689 | 276900.8631 | 226763.7578 | 178212.3056 | 0 | 258446.4477 |
| TG 9:0_18:3_28:4 | 110392.8606 | 61216.50301 | 112471.862 | 110960.8161 | 103373.3694 | 200097.5301 | 111660.3694 | 104662.3381 | 85729.31899 | 70689.28644 | 125645.1644 | 59448.96539 |
| Cer 18:1;2O/12:0 | 2816598.873 | 2388750.037 | 773711.4253 | 1184960.189 | 342655.4737 | 491239.391 | 401227.5441 | 691282.7253 | 450020.6475 | 264067.4693 | 413629.5353 | 372964.6828 |
| Finasteridecarboxylic acid | 84218.74217 | 95176.89631 | 97100.45705 | 156865.3269 | 302965.364 | 279426.8296 | 146464.87 | 223035.2466 | 207888.011 | 112831.6942 | 196258.3663 | 118736.467 |
| (+)-Fluprostenol methyl ester | 665627.2234 | 550925.4268 | 996037.4488 | 958587.1106 | 662512.8054 | 1015489.004 | 782856.4907 | 891218.415 | 644586.201 | 502583.3154 | 758604.9642 | 703108.4402 |

| Metabolites | PBS_1 | PBS_2 | PBS_3 | PBS_4 | BCG_1 | BCG_2 | BCG_3 | BCG_4 | BCGΔ*pe18*_1 | BCGΔ*pe18*_2 | BCGΔ*pe18*_3 | BCGΔ*pe18*_4 |
|---|---|---|---|---|---|---|---|---|---|---|---|---|
| DL-2-Methylglutamic acid | 515713.4784 | 506964.4092 | 637400.3102 | 899120.2283 | 1103433.084 | 744398.8 | 1078410.517 | 901643.5129 | 813048.129 | 798454.1214 | 919385.7319 | 691175.6257 |
| 4-Amino-1-methyl-4-piperidinecarbonitrile | 975370.527 | 421115.698 | 824516.2596 | 801345.0531 | 6094498.678 | 1167478.061 | 676349.768 | 840450.5384 | 417000.7926 | 792672.7631 | 771736.8389 | 865618.0388 |
| 3-Phenylpropanoic acid | 469357.3227 | 411192.0641 | 540137.8528 | 464280.5025 | 565449.2124 | 521610.515 | 447724.3909 | 518253.8203 | 518079.5871 | 312234.0585 | 393581.8531 | 486326.1621 |
| 3-Amino-2-hydroxyheptanoic acid | 166673.2102 | 591611.151 | 376999.4121 | 458369.1287 | 224299.239 | 183798.8637 | 407480.5659 | 166193.319 | 177470.8227 | 727255.3866 | 185758.5546 | 844979.6833 |
| L-Alaninamide, N-[2-[2-(hydroxyamino)-2-oxoethyl]-4-methyl-1-oxopentyl]-3-methyl-L-valyl-N-(2-aminoethyl)- | 964011.9457 | 1111860.556 | 1203677.365 | 1095794.823 | 1045961.015 | 1059991.516 | 1101550.092 | 1011722.903 | 1174645.517 | 858245.2739 | 1054755.205 | 1123501.003 |
| Pyruvaldehyde | 442902.8281 | 390279.3118 | 608183.9561 | 489315.0141 | 1406110.075 | 1069163.347 | 2013414.712 | 1401647.225 | 983765.8706 | 1560223.676 | 1378318.961 | 1004064.213 |
| N-Phenylacetamide | 329118.8117 | 254207.0968 | 446044.726 | 394593.607 | 434988.4679 | 434623.2673 | 436898.5639 | 582258.5051 | 407154.3337 | 244748.6498 | 260811.6977 | 344885.9209 |

| Metabolites | PBS_1 | PBS_2 | PBS_3 | PBS_4 | BCG_1 | BCG_2 | BCG_3 | BCG_4 | BCGΔ*pe18*_1 | BCGΔ*pe18*_2 | BCGΔ*pe18*_3 | BCGΔ*pe18*_4 |
|---|---|---|---|---|---|---|---|---|---|---|---|---|
| 2,4,6-Trihydroxy-5-[1-(4-hydroxy-1,1,4,7-tetramethyl-1a,2,3,4a,5,6,7a,7b-octahydrocyclopropa[h]azulen-7-yl)-3-methylbutyl]benzene-1,3-dicarbaldehyde | 406881.7587 | 220097.951 | 297967.0321 | 340557.3877 | 214374.4622 | 281381.1012 | 303992.9191 | 101134.981 | 177639.0135 | 362799.8204 | 139503.4728 | 277621.6878 |
| Glyceric acid | 519712.452 | 425670.9168 | 555536.1391 | 504526.0253 | 1596044.241 | 1292912.4 | 1373978.559 | 1330107.958 | 1747426.907 | 1539357.589 | 1598940.785 | 1811635.44 |
| MGDG O-17:2_22:5 | 0 | 0 | 55505.54798 | 0 | 575595.0262 | 803392.9395 | 525389.5901 | 613017.8289 | 589109.458 | 533720.4125 | 660083.5198 | 683656.985 |
| Cer 23:0;3O/22:1;(2OH) | 752171.7949 | 596211.5232 | 743276.1259 | 636429.5693 | 693524.4747 | 695126.7181 | 723975.4426 | 533489.2103 | 658173.538 | 618792.2286 | 700165.091 | 654497.4569 |
| 1-Palmitoyl-2-(5-hydroxy-8-oxo-6-octenedioyl)-sn-glycero-3-phosphocholine | 132244.3926 | 162619.6218 | 143875.9283 | 163529.2646 | 156752.2059 | 139116.9689 | 138196.3048 | 122906.542 | 180997.1603 | 189666.2252 | 182496.8703 | 243335.8882 |
| LPA(18:0) | 343135.3079 | 382762.4151 | 259392.0914 | 369230.7963 | 323617.2696 | 344712.0462 | 335281.8763 | 333459.335 | 423252.6373 | 392161.4018 | 424801.7539 | 353121.5459 |
| TRICOSANOIC ACID | 366910.6493 | 125175.8891 | 163079.0574 | 117214.3674 | 119220.8834 | 410595.5443 | 103129.5779 | 172397.8033 | 34698.44142 | 37821.99315 | 140587.1382 | 99689.52503 |
| Cer | 64563.5 | 55425.7 | 15775.5 | 68026.8 | 56710.4 | 50032.5 | 61291.1 | 40888.5 | 60607.2 | 53758.0 | 59328.8 | 41640.2 |

| **Metabolites** | **PBS_1** | **PBS_2** | **PBS_3** | **PBS_4** | **BCG_1** | **BCG_2** | **BCG_3** | **BCG_4** | **BCGΔ*pe18*_1** | **BCGΔ*pe18*_2** | **BCGΔ*pe18*_3** | **BCGΔ*pe18*_4** |
|---|---|---|---|---|---|---|---|---|---|---|---|---|
| 25:3;3O/17:0;(2OH) | 769 | 1965 | 8684 | 641 | 8261 | 5903 | 719 | 8103 | 4766 | 2759 | 7959 | 3802 |
| HexCer 8:1;2O/16:4 | 1196626.243 | 1232783.378 | 1223950.999 | 1187999.264 | 1090669.176 | 1239503.356 | 1086084.547 | 1117915.016 | 1205361.58 | 1189510.31 | 1185854.851 | 1101887.929 |
| 2'-O-methylcytidine | 350981.1227 | 353055.7777 | 719480.0649 | 563822.4959 | 1887707.508 | 1188184.591 | 1795630.215 | 1324007.797 | 1263115.321 | 1179729.943 | 1252865.079 | 925475.4044 |
| LPC(12:0) | 21157.436 | 492004.7201 | 242675.0163 | 145373.5073 | 119689.0248 | 97874.2055 | 154199.1054 | 85371.57541 | 123273.237 | 0 | 95833.51637 | 71628.36915 |
| 1-(2-Hydroxyethyl) pyrazole | 301734.8329 | 894503.28 | 824498.7654 | 1311245.989 | 875553.964 | 845619.822 | 834728.7861 | 856444.6671 | 760708.2128 | 750310.9377 | 852571.2389 | 599453.5954 |
| Wilforlide A | 50255.50422 | 60196.38322 | 72723.21734 | 65910.44668 | 73655.56911 | 86344.67289 | 80582.5378 | 50582.88955 | 66775.19832 | 67867.68214 | 80130.46784 | 72741.20887 |
| 1-{[4-(Dimethylamino)benzyl]amino}-2-propanol | 900450.723 | 884792.6752 | 945227.8689 | 903527.5949 | 874774.4952 | 933223.7896 | 939449.8997 | 899704.9295 | 540586.7882 | 1385553.965 | 858657.8286 | 871471.4595 |
| Phe-Leu | 134609.5722 | 127216.6649 | 176952.9407 | 118246.3326 | 176785.8097 | 130779.1647 | 152732.4959 | 114931.1105 | 154604.1666 | 159769.5575 | 164475.4289 | 99740.98468 |
| Biliverdin | 58687.01288 | 105938.257 | 107588.7856 | 86299.8429 | 705144.8776 | 420040.6036 | 589859.3648 | 369888.8016 | 303763.8699 | 403355.7575 | 410954.4 | 323741.9371 |
| Eszopiclone | 193649.68 | 122337.3857 | 482398.7703 | 125574.4365 | 116504.2906 | 479693.7922 | 232542.7701 | 112113.7538 | 203104.2058 | 164328.755 | 158919.9563 | 112230.1564 |

| Metabolites | PBS_1 | PBS_2 | PBS_3 | PBS_4 | BCG_1 | BCG_2 | BCG_3 | BCG_4 | BCGΔ*pe18*_1 | BCGΔ*pe18*_2 | BCGΔ*pe18*_3 | BCGΔ*pe18*_4 |
|---|---|---|---|---|---|---|---|---|---|---|---|---|
| [(2R,4R)-2-(Bromomethyl)-2-(2,4-dichlorophenyl)-1,3-dioxolan-4-yl]methyl benzoate | 109772.3518 | 105415.0917 | 195920.2181 | 109897.6564 | 221737.8607 | 194003.6929 | 300523.9458 | 176474.712 | 303682.5831 | 158110.4617 | 102399.3154 | 196511.8408 |
| SM 25:1;2O/34:2 | 0 | 0 | 0 | 0 | 14191.60776 | 812697.8416 | 114917.6167 | 115441.4971 | 0 | 0 | 46266.27531 | 176594.7404 |
| L-Leucinamide, N-[(phenylmethoxy)carbonyl]-L-leucyl-N-[(1S)-1-formylbutyl]- | 159737.2664 | 173401.5087 | 173357.3439 | 163990.3604 | 136902.9496 | 128936.4588 | 110813.6968 | 112479.5139 | 146993.4819 | 152057.9297 | 165934.6042 | 170432.0481 |
| DGGA 13:0_20:4 | 38061.94935 | 24008.065 | 125506.7524 | 38292.51818 | 24331.28316 | 13864.67737 | 28977.98826 | 24670.75351 | 0 | 3979.851388 | 4836.710405 | 0 |
| SMGDG O-18:5_7:0 | 150930.5094 | 115916.8313 | 98123.13616 | 106045.7623 | 128517.2734 | 93299.1628 | 108616.0546 | 116479.2331 | 115758.2353 | 119293.6842 | 116249.0125 | 108599.4874 |
| PS 40:10 | 0 | 0 | 0 | 0 | 192270.6944 | 21918.20315 | 191221.8598 | 106642.8764 | 59558.08162 | 93657.48558 | 140243.9597 | 38201.62469 |
| (3.beta.,9.Xi.)-3-(.beta.-D-Glucopyranosyloxy)-14-hydroxycard-20(22)-enolide | 54541.33555 | 22810.76595 | 108860.3571 | 72857.35719 | 81283.01833 | 74550.04993 | 204214.3805 | 98067.03523 | 119539.0522 | 186485.1183 | 169098.5274 | 124705.8547 |
